# Drivers of dispersal and diversification in bromeliads

**DOI:** 10.1101/2022.11.04.515068

**Authors:** Igor M. Kessous, Harith Farooq, Weston Testo, María Fernanda T. Jiménez, Beatriz Neves, Alessandra R. Pinto, Fabiano Salgueiro, Andrea F. Costa, Christine D. Bacon

## Abstract

- Dispersal strategies strongly influence an array of plant traits, especially the shape and function of fruits and seeds, and can be important drivers of diversification dynamics. In this study we investigated how fruit morphology and habitat influence dispersal capacity and diversification rate in bromeliads. We hypothesize that (1) the evolution of berry fruits increased dispersal capacity and diversification rates; and (2) climatic factors contribute to increased dispersal capacity and diversification rates.
- To understand the influence of fruit and habitat traits on evolutionary dynamics, we generated a time-calibrated phylogeny including 1,268 species of bromeliads and integrated that evolutionary framework with distribution, habitat, and morphological trait data.
- We find that lineages with berry fruits have the highest rates of diversification. We also identify significant correlation between diversification rates and both elevation and forest canopy height. We demonstrate that dispersal capacity is not related to fruit morphology and covaries with forest canopy height and mean annual temperature.
- We show that factors influencing the dispersal capacity and diversification are heterogeneous among the subfamilies. These new insights into the rise and spread of bromeliads emphasize the importance of considering the plurality of morphological and ecological features to improve the understanding of the evolutionary dynamics.

## Introduction

Dispersal capacity mediates species occupancy and drives biodiversity patterns through its influence on reproductive isolation and speciation (Antonelli & Sanmartín, 2011; Smith et al., 2014; Steinbauer et al., 2016). In plants, dispersal-related traits such as adaptations in the shape of fruits and seeds are closely associated with dispersal mechanisms associated with wind, animals, amongst others (Seale & Nakayama, 2020). In addition to morphological adaptions that facilitate dispersal, different climatic conditions may also favor different dispersal strategies and thus, the evolution of different fruit and seed morphologies (Seale & Nakayama, 2020).

There are higher levels of biotic interactions (Schemske et al., 2009; Antonelli & Sanmartín, 2011; Brown, 2014; Andresen et al., 2018) and higher dispersal limitations (Janzen, 1967) in tropical compared to temperate regions. Both these patterns drive the current latitudinal diversity gradient (e.g. Mittelbach et al., 2007). Tropical rainforests are characterized by closed canopies and high temperatures and precipitation levels (Eiserhardt et al., 2017). Species richness generally peaks at low latitudes, where forests present the highest canopies (Zhang et al., 2015). Particularly for arboreal communities, such as epiphytes, higher canopies favor the presence of higher niche variation, because of the vertical stratification and microclimatic variation (de la Rosa-Manzano et al., 2014; Oliveira & Scheffers, 2019). In addition to latitude and canopy height, elevation also has an effect on dispersal and diversification, where high elevations have increased speciation rates (Madriñán et al., 2013; Lagomarsino et al., 2016; Testo et al., 2019).

Tropical America (the Neotropics) harbors unparalleled species richness and includes several biodiversity hotspots, with high levels of endemism in both animals and plants (Myers et al., 2000). Previous work has indicated that a unique combination of abiotic and biotic processes interacting over millions of years has led to a unique biodiversity and the high rates of species richness and endemism in the region (Antonelli & Sanmartín, 2011). Endemism is most common at the species level, but there are 52 plant families that are endemic (or nearly so) to the Americas, almost all of which are species-poor (fewer than 100 recognized species; Ulloa Ulloa et al., 2017; Givnish, 2017). A remarkable exception is Bromeliaceae (bromeliads), a hyperdiverse family of monocots that include air plants and pineapples. All but one of the ca. 3,700 recognized species of bromeliads are endemic to the “New World”, most of them restricted to the Neotropics (Smith & Downs, 1974; Ulloa Ulloa et al., 2017; Givnish, 2017; Gouda et al., 2020 [cont. updated]). Bromeliads are ecologically diverse and occur from sea level to ca. 4,000 m in elevation. This ecological diversity is linked to morphological, ecological, and physiological adaptations, including a tank habit, epiphytism, Crassulacean Acid Metabolism (CAM) photosynthesis, and a myriad of biotic interactions (Smith & Downs, 1974; Benzing, 2000; Givnish et al., 2014). The uniqueness in the distribution, ecology, and endemism of bromeliads provide an excellent opportunity to understand the relationship between dispersal-related traits, diversity patterns, and spatial distribution in the Neotropics.

There are two means of dispersal in bromeliads: (1) abiotic, present in capsule-fruited species (with plumose or winged seeds and its varieties) dispersed by wind or gravity (Benzing, 2000; Smith & Downs, 1974) and (2) biotic, in berry-fruited species (Givnish et al., 2011, 2014; Silva et al., 2020; Fig. 1) with “naked” seeds, often with mucilaginous and sticky appendages, dispersed by vertebrates or insects (Smith & Till, 1998; Siva et al., 2020; Leme et al., 2021). Some berry-fruited plants have higher diversification rates than other fruit types (Lagomarsino et al., 2016), since they are able to disperse at longer distances, which may increase the opportunities for lineage diversification. In bromeliads, the relationship between dispersal and morphological or habitat traits remains poorly known, and further, has not been correlated with diversification.

**Fig. 1.**
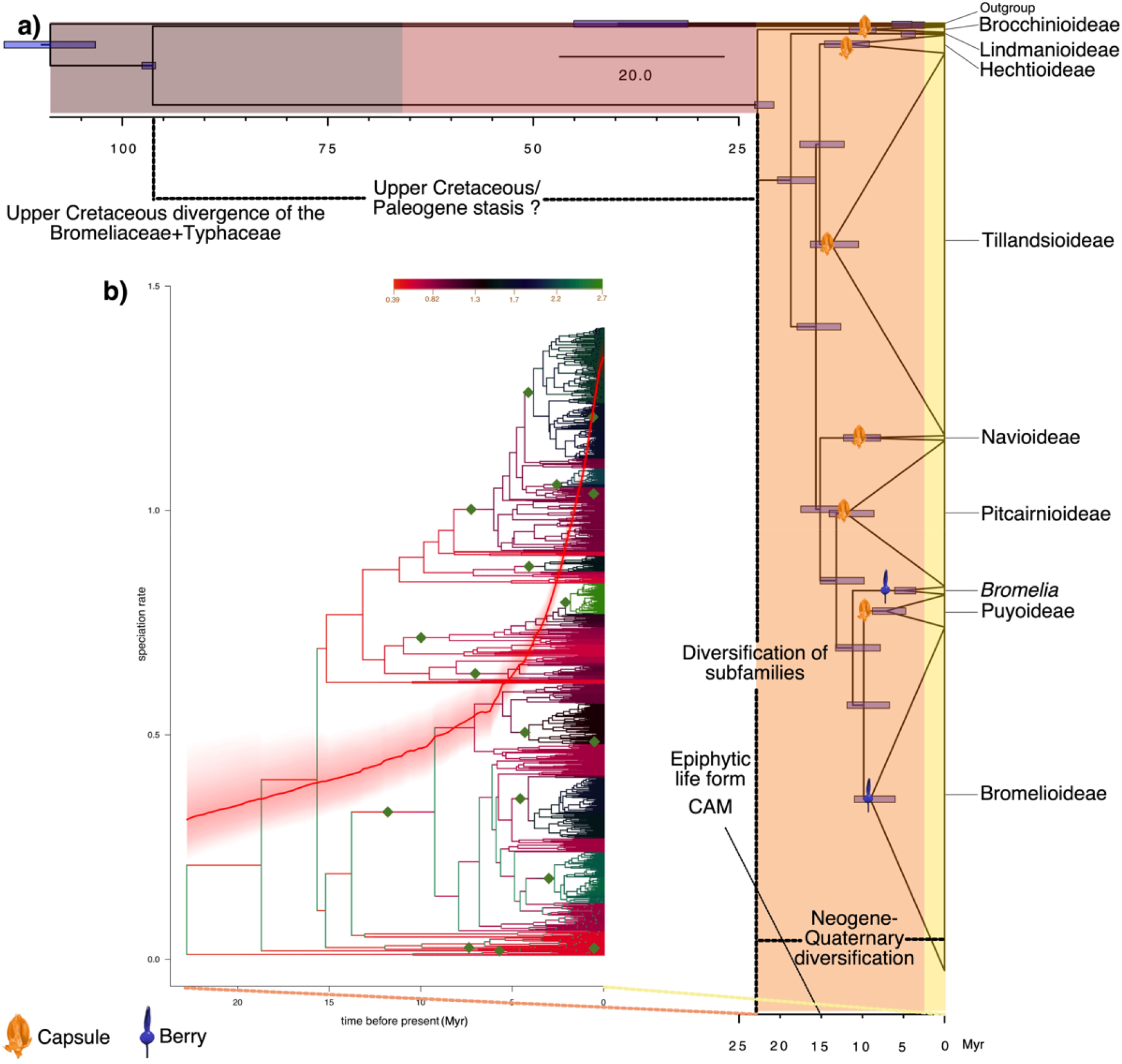
Phylogeny and diversification of bromeliads. a) Maximum clade credibility (MCC) using the combined sequence data of 1,273 taxa. Colored vertical bars refer to geological periods: Quaternary (yellow); Neogene (orange); Paleogene (brownish red); Cretaceous (gray). Horizontal bars on the nodes indicate 95% CI resulted from the treePL analysis. The topology of this time-calibrated tree suggests a Cretaceous origin, Upper Cretaceous-Paleogene stasis and a Neogene-Quaternary diversification of the group; b) BAMM output tree overlapped in the LTT (Lineage Through Time) plot. Time axes are in Myr.

In this study, we identify how fruit morphology and habitat influence dispersal capacity and diversification rate. To do so, we compiled DNA sequence data for 1,268 species of bromeliads, imputed unsampled taxa to produce a comprehensive species-level phylogeny, assembled morphological, distributional, and ecological data sets, and conducted comparative phylogenetic analyses. Specifically, we hypothesize that (1) the evolution of berry fruits increased dispersal capacity and diversification rates. We also hypothesize that (2) climatic factors inherent in certain habitat types, such as precipitation, canopy height, temperature and elevation, contribute to increased dispersal capacity and diversification rates. Determining the dynamics of dispersal and diversification is fundamental to understanding the economically-important bromeliads as well as the Neotropical environments where they are distributed.

## Materials and Methods

We used the Encyclopaedia of Bromeliads (Gouda et al., 2020 [cont. updated], https://bromeliad.nl/encyclopedia/) to standardize the taxonomic classification used in the molecular and spatial datasets (see Appendix S1).

### Taxon and sequence sampling

We obtained sequence data for 13 chloroplast (*agt1, ycf1, rps16-intron, rps16-trnK, rpl32, matK, nadH, petD, trnL-trnF, rpoB, atpB-trnC, psbA-trnH*, and *trnC-petN*) and three nuclear (*PHYC, PRK*, and *LEAFY*) loci from Genbank for a total of 1,268 species of Bromeliaceae (ca. 30% of species in the family) and five outgroups from Typhaceae and Rapateaceae (see Appendix S2). We aligned each locus with MAFFT (Katoh & Toh, 2008) using default parameters. We removed poorly aligned and divergent regions using Gblocks (Castresana, 2000), allowing gap positions in the final blocks and less strict flanking positions. Both alignment analyses were performed in R v. 3.5.3 (R Core Team, 2019), using the package *ips* 0.0.11 (Heibl, 2019). We concatenated the alignments using Geneious Prime (version 2021.0.1, Biomatters, New Zealand) and Mesquite v. 3.61 (Maddison & Maddison, 2019; see Appendix S3) under the criterion of mean pairwise identity over all pairs in the column of at least 30%.

### Phylogenetic analysis and molecular dating

To perform our dispersal and diversification analyses we used two different trees: (1) the backbone tree (BB), output from the phylogenetic analysis including 1,268 species, and; (2) the species level tree (SL) including the BB terminals and the remaining species of the family. To converge multiple chains of the BB tree, we first generated an XML file with BEAUTi v. 1.10.4 (Suchard et al., 2018) to run an exploratory analysis to identify the parameters with disproportionately high effective sample sizes (ESS) and reduce their operator weights in subsequent analyses. We then identified the parameters with disproportionately high effective sample sizes (ESS) compared to others and reduced their operator weights, under the assumption that the data was informative enough for their estimation, which allowed us to speed up data analysis. We did not have calibrations in the tree, however the large number of terminals required high performance computing resources. Thus, to facilitate Markov chain Monte Carlo (MCMC) convergence, model complexity was reduced by selecting the HKY and strict molecular clock (Yule, 1925; Gernhard, 2008). After adjusting operator weights, we used BEAST v. 1.10.4 (Suchard et al., 2018) to run three rounds of two independent chains, each with 300 million MCMC generations sampled every 30,000 generations (see Appendix S4). After every round, we selected the last tree of the analysis with the best ESS values and used it as the starting tree of a subsequent round in order to reach stationarity in both chains. To obtain the maximum clade credibility (MCC) tree, we applied a 15% burn-in to the resulting tree distribution such that the remaining ESS values were > 600. After all rounds, we reached 549,060,000 sampled generations. We performed all phylogenetic analyses at the Swedish National Infrastructure for Computing (https://www.snic.se/) and in the CIPRES Science Gateway V. 3.3 (https://www.phylo.org/).

After estimating the BB tree, we inferred a SL tree of Bromeliaceae by imputing missing taxa using the R package *V*.*Phylomaker* 0.1.0 (Jin & Qian, 2019), after matching the taxa in the BB tree with the species list in Gouda et al. (cont. updt.). We imputed unsampled species only at the genus level. In this case, we used scenario 3 (S3) where new tips of an existing genus were bound in the basal node of this genus (see Qian & Jin, 2016; Jin & Qian, 2019). For the imputation, we used the *build*.*nodes*.*2* function that uses backbone node information based on our species list and the *bind*.*relative* function to include the taxa.

Both the BB and SL trees were dated *a posteriori* using penalized likelihood performed in treePL (Smith & O’Meara, 2012). Because of the absence of reliable fossils of Bromeliaceae (Kessous et al., 2021), we assigned age constraints based on secondary calibrations from Givnish et al. (2018), with minimum and maximum age bounds set at 20% younger and older than the median ages reported for the stem (96 –) 120 (–144) Ma and crown (16–) 20 (–24) Ma of the family. We ran this analysis 100 times, used TreeAnnotator (Bouckaert et al., 2014) to generate a consensus tree and calculate the 95% highest posterior density (HPD) for each node age.

### Spatial data and dispersal capacity

We downloaded 121,978 records of bromeliads from GBIF (www.gbif.org; 03 February 2021): https://doi.org/10.15468/dl.ny8dnt). To remove duplicates and erroneous occurrence records, we performed two sequential analyses with the R package *CoordinateCleaner* 2.0.18 (Zizka et al., 2019), flagging “capitals”, “centroids”, “equal”, “institutions”, “outliers” and “zeros”. We removed species with a single occurrence point, rounded all coordinates to two decimals, and deleted duplicates. The final dataset consisted of 99,863 records of 2,720 species (75% of known species).

We derived a proxy for dispersal capacity using the mean distance between all pairwise occurrence combinations for each species with the function *earth*.*dist* of the R package *fossil 0*.*4*.*0* (Vavrek & Vavrek, 2020), which returned a list in kilometers. Our proxy alleviates issues inherent in other metrics, such as range size and maximum distance between occurrences, which result in problematic inferences for widespread species or those with disjunct ranges. Collection bias may influence the dispersal metrics, since the distance between individuals may be affected by the number of points of each species. We opted to use a different proxy because in convex hull restricted species can be more sensitive to errors (Burgman & Fox, 2003). We also observed widespread species inferred with convex hull with overestimated areas.

### Biotic and abiotic traits

Based on the literature and herbarium specimens, we scored each species for fruit type (according to Smith & Downs, 1974; Benzing, 2000; Givnish et al., 2011; Gouda et al., 2020 [cont. updated]): (1) berry and (2) capsule. Because of the heterogeneity of seeds in bromeliads, as in the case of “naked” seeds belonging to the subfamily Bromelioideae and Navioideae that have appendages with distinct anatomical origins (Silva et al., 2020, Leme et al., 2021), the great variation in winged seeds (Smith & Downs, 1974; Benzing, 2000), and non-homology of appendages in some plumose seeds (Palací et al., 2004), we only used the fruit morphology.

We transformed the fruit type trait into binary presence-absence matrices for the regression analyses. In addition, we scored four continuous variables concerning habitat: canopy height, elevation, mean annual precipitation, and mean annual temperature. We downloaded a canopy height raster with 1 km resolution (https://landscape.jpl.nasa.gov/; accessed on 22 March 2021) and an elevation raster with 30 m resolution (https://www.usgs.gov/centers/eros/science/usgs-eros-archive-digital-elevation-shuttle-radar-topography-mission-srtm?qt-science_center_objects=0#qt-science_center_objects; accessed on 22 March 2021). We downloaded the mean annual precipitation and mean annual temperature variables from CHELSA v1.2 (Karger & Zimmermann, 2019; https://envicloud.os.zhdk.cloud.switch.ch/chelsa/chelsa_V1/climatologies/bio/; accessed on 22 March 2021). We extracted the values of each geographic occurrence point using the R package *raster* 3.4.13 (Hijmans et al., 2015), and calculated the mean for each species.

### Phylogenetic regression

Using the SL tree, we determined the effect of traits on the dispersal capacity of each species by performing phylogenetic regressions using the *pgls* function of the R package *caper* 1.0.1 (Orme et al., 2013). For the phylogenetic regression, we log-transformed and scaled all variables to mean=0 and standard deviation=1. We estimated Pagel’s lambda (λ) using the phylogenetic generalized linear square (PGLS) test, which varies from 0 to 1 (towards the Brownian motion expectation). We determined the most likely model to describe our data using the likelihood ratio test implemented through the function *lrtest* in the R package *lmtest* 0.9.38 (Hothorn et al., 2015). We tested all possible combinations of traits and used the Akaike information criterion (AIC) to select the best-fit model. Finally, we performed an ANOVA test with the function *anova*.*pgls* in *caper* to investigate the effect under each trait. We tested model assumptions for all models by visual inspection of the residuals after applying the *qqnorm* function.

### Diversification analyses

We used state-dependent diversification models to identify the relationship of species traits and rates of diversification in the SL tree. We used BiSSE (binary state speciation and extinction; Maddison et al., 2007) to analyze the fruit type. We conducted all analyses in the R package *diversitree* 0.9.15 (FitzJohn, 2012) and evaluated six different models separately, from the simplest (constraining all parameters) to the most complex (allowing all parameters, in which all rates of speciation depend on the character state). We used the AIC to select the best-fit model, through an ANOVA, and then used the function *make*.*prior*.*exponential* to set exponential priors (FitzJohn, 2012). We ran a preliminary MCMC chain of 100 steps to set the control parameter (*w*) and sequentially subsampled it in the 0.05-0.95 interval. We then ran the final MCMC chain of 10,000 steps, sampling every 10 steps, applying a burn-in of 5% and calculated the net diversification rate for the fruit type trait.

The reliability of SSE methods to account for diversification has been widely discussed in the literature (Davis et al., 2013; Rabosky & Goldberg, 2015; Maddison & FitzJohn, 2015; Herrera-Alsina et al., 2019). Because state-dependent analysis is sensitive to large differences between tip ratios and small-sized trees (Davis et al., 2013; Rabosky & Goldberg, 2015), we used the fully sampled tree (SL) and the fully coded traits. It has been suggested that the state tip ratio of 3:1 has high rates of power in large sampled (>500 taxa) asymmetrical simulations and that a 10 % minimal threshold of tip ratio and 300 terminals should be used to better interpret BiSSE results (Davis et al., 2013). We obtained a ratio of 26% (berry) and 74% (capsule) and the analysis was based on 3,585 taxa (SL tree).

We then performed a time-dependent analysis to obtain the net diversification tip rates and shifts using BAMM v. 2.5.0 and analyzed the outputs using the R package BAMMTools 2.1.7 (Rabosky et al., 2014). Unlike the previous analyses, in this case, we used the BB tree, since null branch-lengths, such as some in the SL tree, can influence the measurement of diversification across clades. However, to solve the incomplete sampling and heterogeneity of the BB tree (totaling 30% for the whole family), we specified different sampling fractions for each clade, based on the proportion of recognized species present in the phylogeny: Bromelioideae + Puyoideae (0.42), Tillandsioideae (0.33), Navioideae (0.06), Brocchinioideae (0.4), Lindmanioideae (0.07), and Pitcairnioideae (0.3). We ran four reversible jump MCMC chains, each for twenty million generations, sampling each 20,000 generations. We discarded the first 20% generations as burn-in and checked the convergence in the estimated sample size with a threshold >200. We then performed Spearman’s correlation tests and plotted the extracted diversification (*lambda*) and extinction (*mu*) tip rates of each terminal, in addition to the dispersal capacity (mean distance), in order to compare the influence of each numerical variable.

BAMM analyses do not allow negative or null branch-lengths and the higher rates of diversification found in young lineages may be influenced by the “pull of present”, those that have had less time to become extinct (Eiserhardt et al., 2017; Helmstetter et al., 2022). Moore et al. (2016) presented a critique of the BAMM method, mainly regarding the likelihood function, rate-shifts, and diversification rates. However, Rabosky et al. (2017) concluded that BAMM has high accuracy and consistency to account for diversification and rate shifts, and argued that Moore et al. (2016) based their discussion on a set of low-power analyses. Following Rabosky et al. (2017), we used this method and compared the distribution of the morphological, habitat and diversification data with each tip in the trees we used the R package *ggtree* 2.99.0 (Yu et al., 2017).

## Results

### Alignment and phylogenetic trees

Our dataset resulted in an alignment of 21,271 bp from 1,273 taxa (1,268 bromeliads + 5 outgroups; see Appendices S3 and S4). Major clades within the family (subfamilies, tribes, and subtribes) were strongly supported (Fig. 1; see Appendix S5; Tab. S1). The relationships of lineages within the family correspond to those reported by Givnish et al. (2011), except for a strongly-supported, novel clade of *Bromelia* + (Bromelioideae + Puyoideae) (PP=1). Some relationships at the generic and infrageneric level were poorly-supported, especially for the taxonomically-problematic genera (see Appendix S5 for BB tree; and S6 for SL tree; 3,585 taxa).

### Divergence time estimate

The divergence between Bromeliaceae and Typhaceae occurred during the Upper Cretaceous (∼96 Ma; Fig. 1, Tab. S1) however, diversification in Bromeliaceae only started at the beginning of the Neogene (∼22 Ma). Divergence of the major bromeliad groups occurred during the Miocene (∼22 Ma), with Brocchinioideae and Lindmanioideae as the early divergent subfamilies. Despite the early divergence of these groups, the diversification of all subfamilies only started in the Mid-Upper Miocene, except Lindmainioideae, which diversified during the Pliocene (4.6 Ma).

### Phylogenetic regression

The best-fit model for dispersal capacity included annual temperature and canopy height (AIC value of 5937.534; Tab. S2, Fig. S1), which was corroborated with significant ANOVA results (Tab. 1). Although annual temperature was significant in the full model ANOVA (Tab. S3), the slope was not significant (p=0.274). For the best-fit model, we conducted a phylogenetic regression to identify the effect of temperature and canopy height on dispersal capacity. Our analyses showed that annual temperature and canopy height were strongly significant (p<0.0001). The effect of annual temperature (0.180 ± 0.028) was marginally stronger than the canopy height (0.137 ± 0.028) and moderately supported (λ=0.75, 95% CI=0.64 – 0.82). All the effects of independently tested variables on dispersal capacity had high AIC values, and all of the trait variables were non-significant.

**Tab. 1.** ANOVA of the PGLS (Phylogenetic Generalized Least Squares) best-fit dispersal capacity model according to AIC (Akaike Information Criterion) values (MD∼AT+CH). Fruit morphology is not a predictor for dispersal capacity.

| Trait | df | Sum Sq | Mean Sq | F value | p-value |
| --- | --- | --- | --- | --- | --- |
| AT | 1 | 1719 | 1718.63 | 42.612 | 8.273e-11 |
| CH | 1 | 921 | 921.30 | 22.843 | 1.875e-06 |
| Residuals | 2186 | 88166 | 40.33 | - | - |

*Notes:* Estimated value of lambda of 0.748 considering the influence of all variables under the mean distances. AT: mean annual temperature; CH: canopy height; df: degrees of freedom.

### Diversification

Among the six models tested in the trait-dependent diversification analyses, the best fit was the model where character change (*q*) was constrained for fruit type. Our analyses indicated that berry-fruited species (λ=0.20, μ=0.000034) have higher net diversification rates than capsule-fruited species (λ=0.13; Fig. 2).

**Fig. 2.**
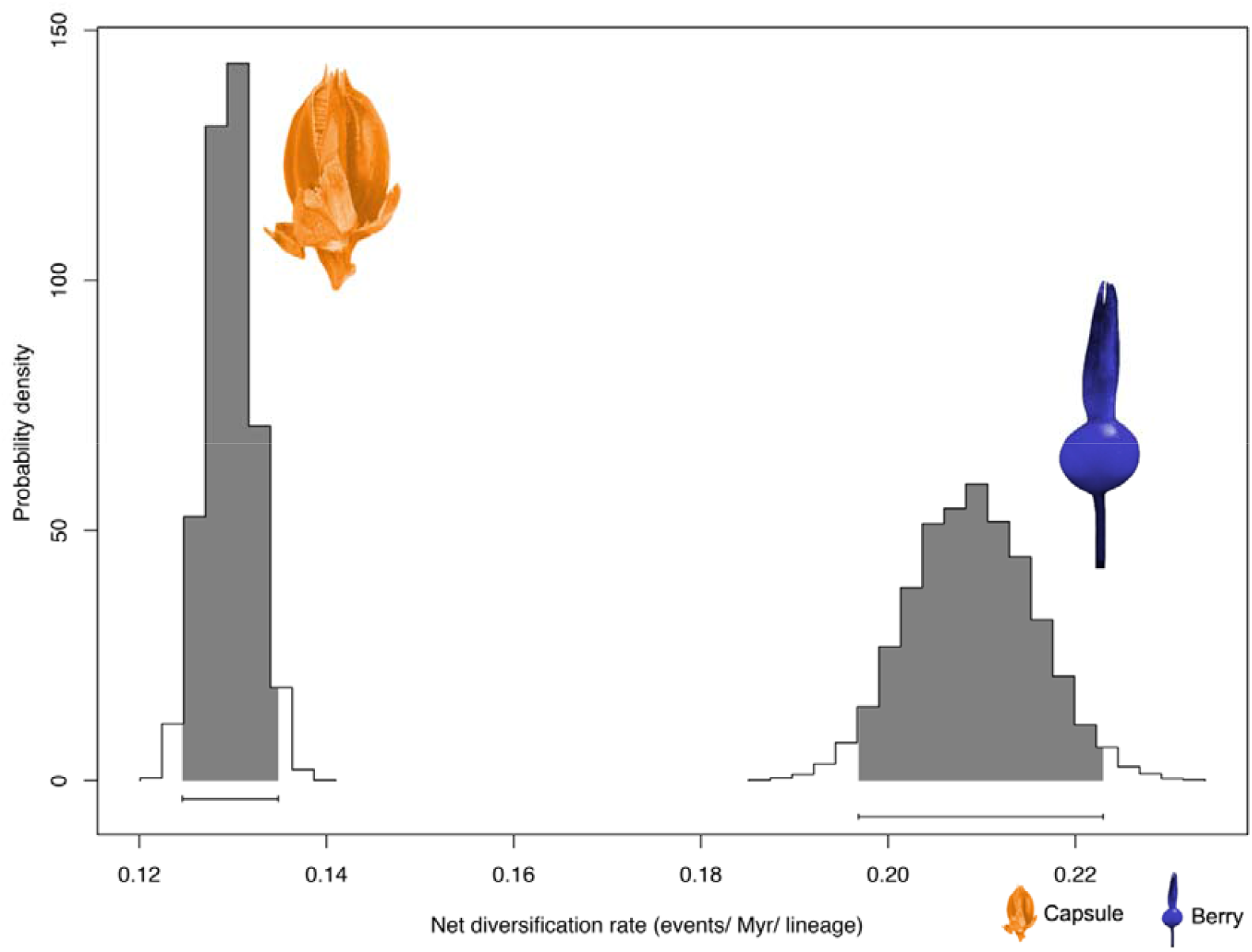
BiSSE (Binary state speciation and extinction) indicating the effect of the fruit types on net diversification rate.

The BAMM diversification analysis identified heterogeneous rates across the tree (Fig. 1, see Appendix S7). Higher rates were observed in the clades of *Wittmackia* Mez, *Dyckia* Schult.f., and a group comprising North and Central American *Tillandsia* L. (see Appendix S7). Seventeen diversification rate shifts were identified in the Upper Cenozoic, six in the Miocene, five in the Pliocene and six in the Pleistocene, coinciding with the divergence and diversification of several lineages (Fig. 1).

Considering bromeliads at the family level, our results identified a significant negative correlation between net diversification rate with elevation (R=-0.15, p=2e-06) and canopy height (R=-0.11, p=0.00068). Annual precipitation (R=-0.011, p=0.72) was not significant and annual temperature (R=0.064, p=0.039) had a significant but low positive correlation with net diversification rate (Fig. 3, S2). Considering each subfamily separately (Tab. 2), the variables had different effects on the net diversification rate, and when significant, the slopes frequently had a stronger correlation compared to those at the family level.

**Tab. 2.**
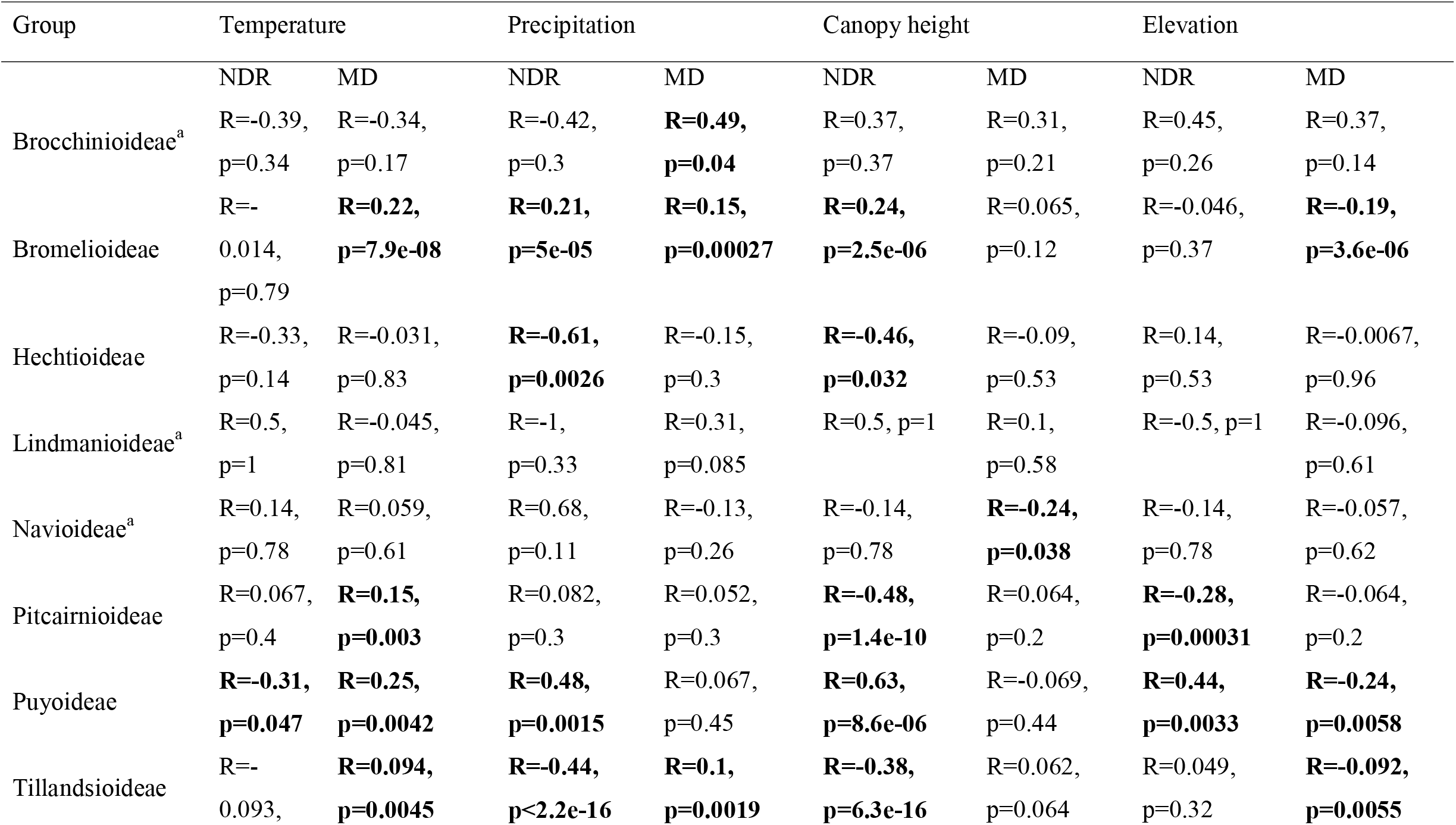
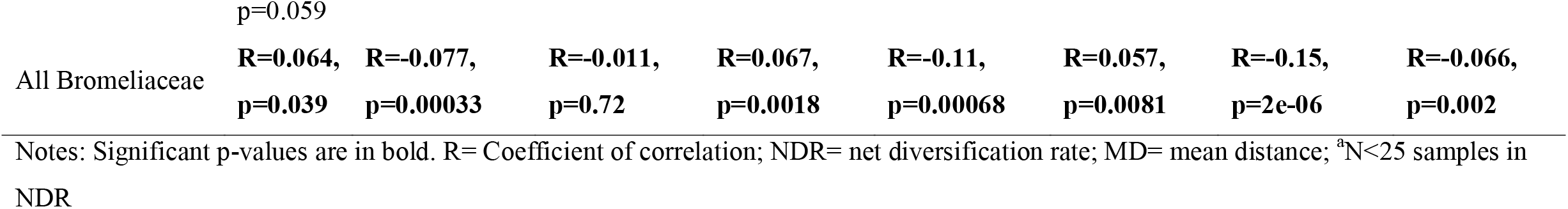
Correlation between the numeric variables and net diversification rate and mean distance for each bromeliad group.

**Fig. 3.**
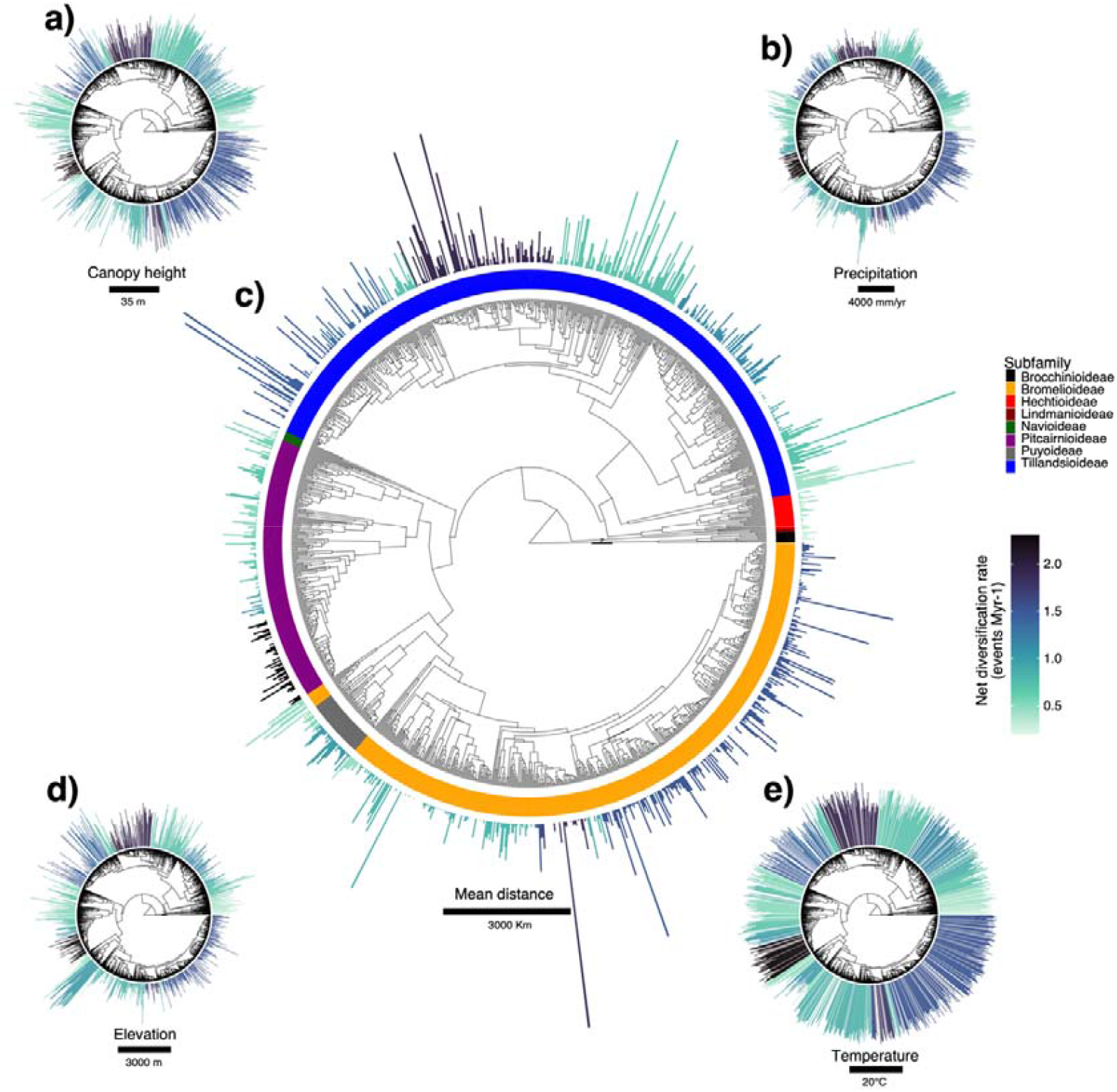
Phylogeny of Bromeliaceae, habitat and diversification. Bar plots indicate habitat type in size and net diversification rate in color: a) Canopy height values; b) Precipitation values; c) Mean distance values; d) Elevation values; e) Temperature values.

We investigated the effects of the variables in each group of fruit types. Among the variables we tested, canopy height and annual precipitation had significant correlations with net diversification rate (Fig. 4a–b), positive for the berry fruit type (R=0.24, p=2.5e-06 and R=0.21, p=5e-05, respectively) and negative for capsules (R=-0.28, p=1.4e-13 and R=-0.16, p=3.9e-05, respectively). For dispersal capacity, annual temperature and elevation had significant correlations with mean distance (Fig. 4c–d), with annual temperature positively correlated for both types of fruits (berry: R=0.22, p=7.9e-08; and capsule: R=0.056, p=0.024) and elevation negatively correlated (berry: R=-0.19, p=3.6e-08; and capsule: R=-0.054, p=0.03). In both cases, the correlation was stronger in berry-fruited species. Results on maps, data distribution and geographic information are in Fig. S3, see Appendices 8 and 9.

**Fig. 4.**
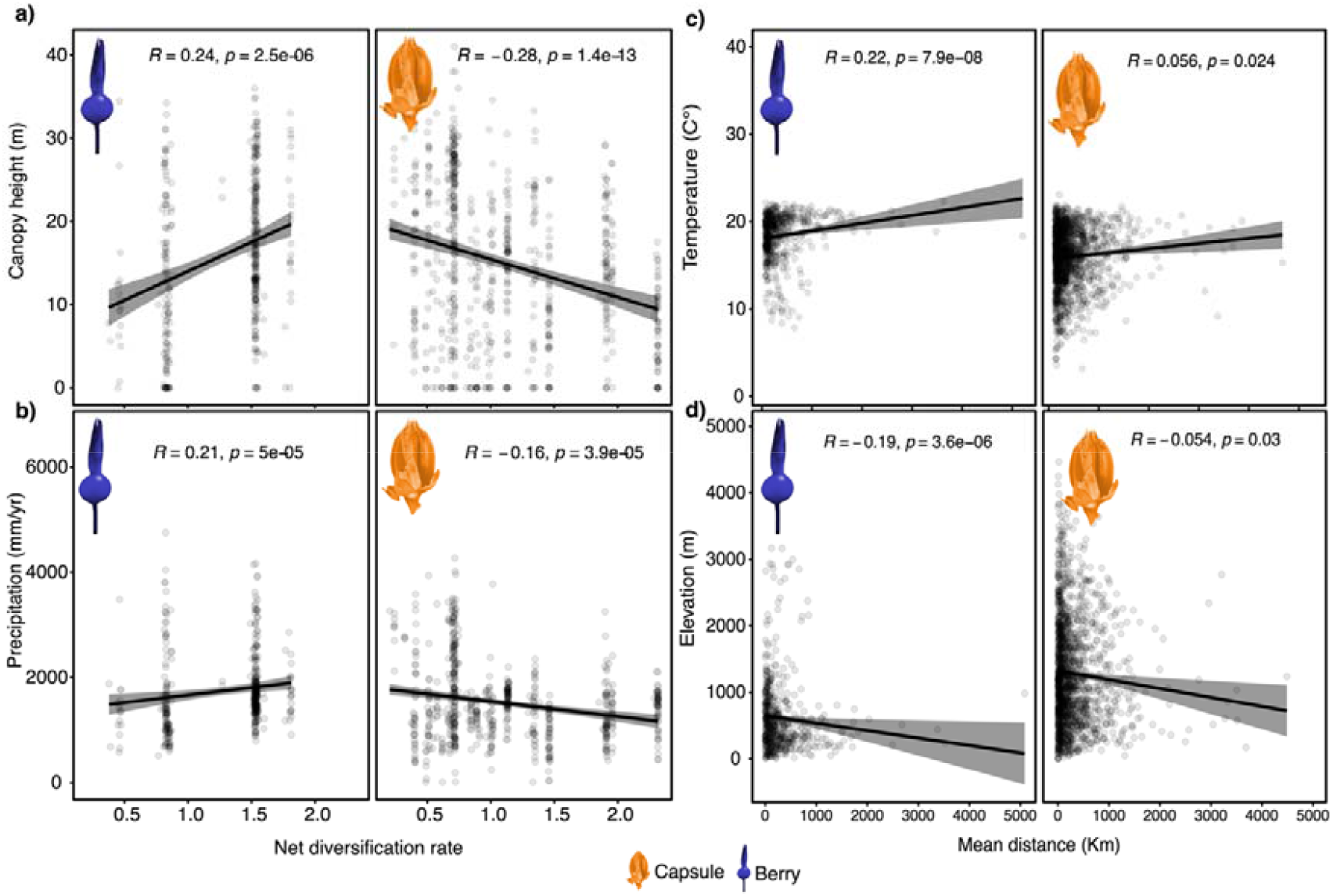
Correlation plots of the habitat types and net diversification rates (NDR)/mean distances (MD) within different types of fruits. The analysis showed canopy height (a) and precipitation (b) correlated to the net diversification rate (positively in berry and negatively in capsule, in both) and temperature (c) and elevation (d) correlated to mean distance (positively in temperature and negatively in elevation, in both).

## Discussion

The relationship between biological dispersal and distribution remains a large knowledge gap fundamental to the geographic distribution of species. It is also poorly understood how this relationship interacts with diversification, which together drive biodiversity patterns in time and space. Here, we use a large molecular phylogeny of bromeliads, fruit and climatic data to test how habitat and morphology influence dispersal capacity and lineage diversification.

### Berry-fruited species have higher diversification rates

Our results demonstrate that berry-fruited species have higher speciation rates and lower extinction rates compared to the capsule-fruited species (Fig. 2). The evolution of fleshy fruits, including berries, is considered a key innovation in bromeliads (Givnish et al., 2014). In the Neotropical bellflowers, berries were strongly correlated to closed forest canopy habitats (Lagomarsino et al., 2014, 2016), as here observed for bromeliads (Fig. 4). Berries are primarily bird-dispersed, and the Neotropics has the highest species richness of frugivorous birds in the world (Kissling et al., 2012). Moreover, fleshy-fruited plants and their associated frugivores are commonly generalists (Eriksson, 2016; Bello & Barreto, 2021), meaning that seed dispersal does not rely on a single species and can promote diversification via increasing dispersal capacity (de Queiroz, 2002).

Our BAMM results identify three bromeliad clades that had the highest diversification rates (see Appendix S7): (1) *Wittmackia* (berry fruit, Bromelioideae); (2) *Dyckia* (capsular fruit, Pitcairnioideae); (3) North and Central American *Tillandsia* (capsular fruit, Tillandsioideae) (Fig. 1). Aguirre-Santoro et al. (2020) also showed high diversification rates in *Wittmackia*, which was suggested to result from a recent long-dispersal event from the Brazilian Atlantic Forest to the Caribbean, causing a rapid radiation with a myriad of morphological adaptations. Central America (*sensu* Givnish et al., 2011) is considered an important biogeographic region for bromeliads (Givnish et al., 2011; Granados-Mendoza et al., 2017; Kessous et al., 2020), since it is one of the three species diversity centers of bromeliads (Smith & Downs, 1974; Givnish et al., 2011; Zizka et al., 2020). Also, events occurred in the Miocene-Pliocene, as the seasonality and aridity, were potential drivers of diversification of North and Central American *Tillandsia* (Granados-Mendoza et al., 2017). Our results indicate that, considering bromeliads as a whole, berry-fruited species have higher diversification, although some individual capsule-fruited clades also have high diversification rates.

### Habitat drives dispersal and diversification

Environmental conditions at high elevations contribute to diversification in mountain ecosystems (Rahbek et al., 2019), including Neotropical plants (Madriñán et al., 2013; Lagomarsino et al., 2016; Testo et al., 2019). Considering bromeliads as a whole, only temperature and canopy height influence in the dispersal capacity (Tab. 1). Fruit morphology was not found to be a predictor of dispersal capacity, probably because bromeliads have two different long-distance dispersal mechanisms (anemochory and zoochory) in their largest groups (see the distribution of the data in Fig. S4). Also, we observed an overall negative correlation between elevation and net diversification of bromeliads (Fig. S2a). In both cases, the relation among the traits, dispersal capacity and diversification are clearer considering case-by-case.

Considering subfamilies common in high elevations such as the Andean Puyoideae, we found a strong positive correlation with net diversification rate (Tab. 2). Bromeliads have great morphological diversity, favoring the occurrence across broad elevation gradients ranging from coastal Brazilian *restinga* forests (e.g. *Neoregelia cruenta* (Graham) L.B.Sm.) to Andean highlands at almost 5,000 m (e.g. *Puya raimondii* Harms; Smith & Downs, 1974).

According to our data, of the 2,718 species with an estimated mean elevation, only 15% occur higher than 2,000 m and 3% higher than 3,000 m. In contrast, among our 180 analyzed species of Puyoideae, 67% have a mean elevation higher than 2,000 m and 33% higher than 3,000 m. Thus, it is important to recognize that elevation-diversification dynamics differ considerably across clades and some patterns are driven largely by single groups.

High elevation areas have high endemism, due to the geographic isolation of the species found there (Steinbauer et al., 2016), and to the lower species richness of frugivore dispersers (Sam et al., 2019). Our analysis shows that dispersal capacity is negatively correlated with elevation in Puyoideae (Tab. 2) and berry-fruited species (Fig. 4), suggesting that high elevation species tend to be endemic to smaller regions, probably owing to the specific attributes and adaptations to these habitats.

Plant lineages often have strong niche conservatism with climatic preferences such as temperature and precipitation (Punyasena et al., 2008), and are therefore vulnerable to climate change (Liu et al., 2020). Here, we show that bromeliad subfamilies have specific climate preferences. As a whole, in bromeliads, there was no correlation between annual temperature and net diversification rate (R=0.064, p=0.039, Tab. 2), but when considering only Puyoideae we observe a negative correlation (R=-0.31, p=0.047, Tab. 2). In Puyoideae, high rates of diversification are associated with lower temperatures, potentially suggesting that a lack of competition in high elevation environments benefits the diversification in the group. In contrast, we observed that high temperatures positively influence the dispersal ability of Puyoideae (R=0.25, p=0.0042, Tab. 2), Bromelioideae (R=0.22, p=7.9e-08, Tab. 2) and Pitcairnioideae (R=0.15, p=0.003, Tab. 2), indicating that for these groups, lower temperatures lead to higher rates of endemism.

Annual precipitation is correlated with dispersal capacity and net diversification rate in some bromeliad subfamilies. Low humidity regions may positively affect dispersal in capsule-fruited species (with wind-dispersed seeds; Heydel & Tackenberg, 2017). The diversification in Tillandsioideae and Hechtioideae, both widely distributed in dry habitats, are negatively correlated with precipitation, likely because of their seed morphology that is adapted to dispersal by wind and/or gravity. On the other hand, precipitation positively influences the diversification of Bromelioideae and Puyoideae, which despite their close phylogenetic relationship (Fig. 1, Tab. 2), have differences in fruit and seed morphology (berry/naked versus capsule/winged, respectively). Precipitation is positively correlated with dispersal ability in Brocchinioideae (endemic to Amazonia), Bromelioideae, and Tillandsioideae (Tab. 2), the latter two being the most widely distributed subfamilies (Zizka et al., 2020). Co-occurrence may be reflected by similar habitat preferences among different groups (Punyasena et al., 2008) as clearly observed among these niche-conserved bromeliads (see Zizka et al., 2020).

### Canopy height influences bromeliad dispersal and diversification

The net diversification rate of the family decreases with a higher canopy height (Tab. 1, Figs. S2, 3a, 4), perhaps due to the predominance of species with capsular fruits (>70% of the species), since in berry-fruited species (Bromelioideae, mostly epiphytic) the net diversification rate increases with a higher canopy height and annual precipitation (Fig. 4a–b). A higher canopy can behave as a physical barrier for the wind-dispersed seeds present in capsular fruits and also may positively influence the diversity of vertebrate dispersers, (Walter et al., 2017), favoring the dispersal in berry-fruited species. On the other hand, open canopies can favor the wind-dispersed seeds (Correa et al., 2022).

In the Neotropics, the end of the Cretaceous (Maastrichtian: 72–66 Ma) was characterized by open canopied forests and the presence of several vascular plant groups. In contrast, early Paleocene forests (66–61 Ma) were denser, but with a lower diversity compared to the Maastrichtian (Carvalho et al., 2021). This different habitat composition is likely due to the (1) presence of large herbivores, mainly dinosaurs, that physically disturbed forests, (2) infertility of the soils in the Maastrichtian, and (3) the selective extinction of gymnosperms that occurred in the Cretaceous-Paleogene boundary (Carvalho et al., 2021). The reduction of 45% of the Paleocene plant diversity resulted from the slow recovery from the Cretaceous-Paleogene mass extinction caused by the Chicxulub impact and intense volcanic activity (Schulte et al., 2010; Hull et al., 2020), which suppressed sunlight, changed the atmosphere, and reduced global temperatures (Alvarez et al., 1980; Schulte et al., 2010; Vajda et al., 2014; Vajda et al., 2015; Hull et al., 2020). The increased complexity of Neotropical Paleocene forests provided opportunities that included vertical diversity, as in the case of the epiphytes, primarily because to changes in the water and light availability within closed canopies (Carvalho et al., 2021).

The first bromeliads were terrestrial and inhabited open environments (Givnish et al., 2014; Bouchenak-Khelladi et al., 2014), however, the evolution of other life forms and morphology, such as epiphytism, possibly behaved as a trigger for high and rapid diversification (Givnish et al., 2014; Givnish et al., 2017). The causes of the early divergence and late diversification of the bromeliads are still unknown (Kessous et al., 2021). According to our results we hypothesize that the open forests of the Upper Cretaceous favored the rise of the terrestrial/capsular bromeliads. However, after the mass extinction, during the Paleocene, the closed canopied forests negatively affected the bromeliad diversification, once the epiphytic life form of this group appeared for the first time in the Mid Miocene (Givnish et al., 2014). The Eocene-Oligocene extinction event resulted in semi-open canopies (Carvalho et al., 2021), which also may be one of the factors that triggered the diversification of bromeliads in the Miocene (Fig. 1). After the development of the epiphytic habitat (Fig. 1, 3a), the higher canopy height may have also favored diversification, as for the berry-fruited species. We hypothesize that both higher and lower canopies were triggers of bromeliad diversification, but in different time frames and for different fruit morphologies.

Taken together and according to our results, we offer two hypotheses: (1) capsular fruits have seeds that are primarily wind-dispersed, and closed canopies serve as a physical barrier to dispersal and subsequent allopatric speciation (2) the abundance of frugivores (such as large birds and bats) can be higher in dense forests (Walter et al., 2017) and benefit the dispersal of berry-fruited species. Although the first documented frugivore interactions are from the Upper Cretaceous, fleshy fruits evolved in several plant groups in the Miocene (Eriksson, 2016), as observed here in bromeliads.

## Conclusion

Here we test drivers of dispersal and diversification in a hyperdiverse plant family in a phylogenetic framework. We hypothesize that both lower and higher canopy heights influenced the dispersal and diversification of bromeliads, and that the low canopies heights of paleoforests prompted the origin and diversification of the family. We also demonstrate that berry fruits promoted diversification in bromeliads. Fruit morphology is not a predictor for dispersal capacity and factors influencing fruit types are heterogeneous among the subfamilies. Our analyses show the importance of evaluating different groups as separate entities together with different effects of climatic variables in each group. Furthermore, we show that the evolutionary success of bromeliads does not rely on a single key innovation, but rather on a complex network of strategies on different temporal scales and that are linked to dispersal, colonization, and adaptation. We highlight the importance of investigating the influence of traits and variables on dispersal and diversification, while accounting for the ecological and morphological diversity of its subgroups.

## Supporting information

Supplemental Folder

## Data availability

The data underlying this article are included in the Supporting Information.

## Acknowledgements

This work was supported by Coordenação de Aperfeiçoamento de Pessoal de Nível Superior (CAPES Foundation), CAPES-STINT (Project 88881.304776/2018-01; 8887.477452/2020-00); the Helge Ax:son Johnsons stiftelse (F21-0212); the Sven and Dagmar Saléns Foundation; the Wilhelm and Martina Lundgrens Foundation (2020-3489); the Swedish Research Council (2017-04980); and the Biodiversity and Ecosystems Services in a Changing Climate (BECC) Strategic Research Area at the University of Gothenburg. We thank Søren Faurby and the members of the Gothenburg Global Biodiversity Centre for valuable discussions; and the Swedish National Infrastructure for Computing (SNIC) for computational support (SNIC 2020/9-216; SNIC 2020/10-114).

## Conflict of interest

The authors declare no conflict of interest.

## Author contributions

IMK, AFC, FS and CDB planned and designed the study. IMK, HF, WT, MFT, BN and ARP ran the statistical and phylogenetic analyses. IMK, AFC and CDB interpreted the results.

IMK wrote the manuscript with suggestions from all remaining authors.

## Supporting Information

**App.S1** List of accepted species names of Bromeliaceae according Gouda et al., 2020 [cont. updated], <https://bromeliad.nl/encyclopedia/> (December 2020).

**App. S2** List of the sequences of the sampled taxa. Available in GenBank (ncbi.nlm.nih.gov/genbank/).

**App. S3** Concatenated matrix totaling 21,271 bp from 1,273 taxa. Sequence data for 13 chloroplast (*agt1, ycf1, rps16-intron, rps16-trnK, rpl32, matK, nadH, petD, trnL-trnF, rpoB, atpB-trnC, psbA-trnH*, and *trnC-petN*) and three nuclear (*PHYC, PRK*, and *LEAFY*) loci.

**App. S4** XML final file generated with BEAUTi v. 1.10.4 (Suchard et al., 2018).

**App. S5** Backbone tree (BB tree). Maximum clade credibility (MCC) using the combined sequence data of 1,273 taxa resulted from a 21,271 bp alignment.

**App.S6** Species-level tree (SL tree). All-species bromeliad tree obtained from the BB tree by imputing missing taxa using the R package *V*.*Phylomaker* 0.1.0 (Jin & Qian, 2019).

**App. S7** Tip rates out of net diversification obtained from BAMM v. 2.5.0 diversification analysis.

**App.S8** Records, geographical, morphological and habitat information.

**App.S9** Species and transformed data (log-transformed, scale, dummy variables), habitat and morphological information.

**Tab. S1** Main clade support and timing of crown and stem ages.

**Tab. S2** Comparison of trait-based best-fit models of dispersal represented by mean distance (MD).

**Tab. S3** ANOVA of the PGLS complete model. Estimated value of lambda of 0.736 (0.620, 0.815), kappa and delta fixed in 1, considering the influence of all variables under the mean distances. The levels of berry (FT) and winged (ST) were considered points of comparison, thus not estimated in the analysis. All variables were scale 0 modified.

**Fig. S1** Residuals of the PGLS (Phylogenetic Generalized Least Squares) best-fit model according to AIC (Akaike Information Criterion).

**Fig. S2** Correlation plots of the habitat types and net diversification rates (NDR). Elevation and Canopy height showed negative correlation with NDR, considering the whole family. a) Elevation; b) Temperature; c) Precipitation; d) Canopy height. Marginal lines represent the density distribution of the data.

**Fig. S3** Maps, tree and distribution of the data. a) Precipitation maps and density plots; b) Temperature maps and density plots; c) Canopy height maps and density plots; d) Elevation maps and density plots; e) Species-level tree (SL tree) with bar plots of the habitat data.

**Fig. S4** Violin plots representing the distribution of the mean distance (proxy for dispersal capacity) between the two fruit types.

## Notes

### Competing Interest Statement

The authors have declared no competing interest.

