## Supplemental Folder for "Drivers of dispersal and diversification in bromeliads": App S2.rtf

;agt1

			>Acanthostachys_pitcairnioides
						TCGCAGAAAGCTCTTTCCCTGCCTACTGGAATGGGCATTCTTTGCGCGAGTCCGAAAGCTCTAGAAGCTTCCAAAACGGCCAGGTCGGTCAGGGTTTTCTTCGACTGGAATGACTACTTGAAGTTYTACAAGCTCGGAACTTACTGGCCTTAACGCCTTCCATCCAACTACTTTACGGGCTGAGAGCAGCTTTGGATCTCATCTTTGAGGAGGGGCTCGACAATGTCATCGAGAGGCATCGTCGCTTAGGAAAGGCCACAAGGTTAAGTGAAAGCTATACATCT-AATAGCCGCTTTCGCCGTGTTGGATAGATATATTTACTCGAACTATCATCCTCTTGTAACAATGAGCAGACTTGCCGTCGAAG

			>Acanthostachys_strobilacea
			????????????????????????????????????????????????????????????????????????????????????????????????????????????????????????????????????????????????????????????????????????????????????????????????????????????????????????????????????????????????????????????????????????????????????????????????????????????????????????????????????????????????????????????????????????????????

			>Aechmea_abbreviata
			TCGCAGAAAGCTCTYTCCCTSCCTACTGGAATGGGCATTCTCTGCGCGAGCCCGAAAGCTCTAGAAGCATCCAAAACAGCCAAGTCGGTCAGGGTTTTCTTCGACTGGAATGACTACTTGAAGTTCTACAAGCTCGGAACTTACTGGCCTTAACGCCGTCCATCCAACTGCTTTACGGGCTGAGAGCAGCTTTGGATCTCATCTTCGAGGAGGGGCTCGACAATGTCATCGAAAGGCACCGTCGCTTAGGAAAGGCCACRAGGTTAAGTAAAAGCTGTACATCT-AATAGCCGCCTTCGCCGTGTTGGATAGATATATTACTCGGCGCTATTGTTCTCTCGTAACAATGAGCAGACTYGCCGTCGAAG

			>Aechmea_aculeatosepala
			????????????????????????????????????????????????????????????????????????????????????????????????????????????????????????????????????????????????????????????????????????????????????????????????????????????????????????????????????????????????????????????????????????????????????????????????????????????????????????????????????????????????????????????????????????????????

			>Aechmea_aiuruocensis
			TCGCAGAAGGCTCTTTCCCTGCCTACTGGAATGGGCATTCTCTGCGCGAGCCCGAAAGCTCTAGAAGCTTCCAAAACAGCCAAGTCGGTCAGGGTTTTCTTCGACTGGAATGACTACTTGAAGTTCTACAAGCTCGGAACCTACTGGCCTTAACGCCGTCCATCCAACTGCTTTACGGGCTGAGAGCAGCTTTGGATCTCATCTTTGAGGAGGGGCTCGACAATGTCATCGAAAGGCATCGTCGCTTAGGAAAGGCCACAAGGTTAAGTAAAAGCTATACATCT-AATAGCCGCCTYCGCCGTGTTGGATAGATATATTACTTGGCACTGTTGTYCTCTTGTAACAATGAGCAGACTTGCCGTCGAAG

			>Aechmea_alba
			????????????????????????????????????????????????????????????????????????????????????????????????????????????????????????????????????????????????????????????????????????????????????????????????????????????????????????????????????????????????????????????????????????????????????????????????????????????????????????????????????????????????????????????????????????????????

			>Aechmea_alopecurus
			TCACAGAAAGCTCTTTCCCTGCCTACTGGAATGGGCATTCTTTGCGCGAGCCCGAAAGCTCTAGAAGCTTCCAAAACAGCCAAGTCGGTCAGGGTTTTCTTCGACTGGAATGACTACTTGAAGTTCTACAAGCTCGGAACTTACTGGCCTTAACGCCGTCCATCCAACTGCTTTACGGGCTGAGAGCAGCTTTGGATCTCATCTTTGAGGAGGGGCTCGACAATGTCATCGAAAGGCACCGTCGCTTAGGAAAGGCCACAAGGTTAAGTAAAAGCTACACATCT-AATAGCCGCCTTCGCCGTGTCGGATAGATATATTACTTGGTACTGTTGTTCTCTTGTAACAATGAGCAGACTTGCTGTCGAAG

			>Aechmea_angustifolia
			????????????????????????????????????????????????????????????????????????????????????????????????????????????????????????????????????????????????????????????????????????????????????????????????????????????????????????????????????????????????????????????????????????????????????????????????????????????????????????????????????????????????????????????????????????????????

			>Aechmea_aquilega
			????????????????????????????????????????????????????????????????????????????????????????????????????????????????????????????????????????????????????????????????????????????????????????????????????????????????????????????????????????????????????????????????????????????????????????????????????????????????????????????????????????????????????????????????????????????????

			>Aechmea_arenaria
			????????????????????????????????????????????????????????????????????????????????????????????????????????????????????????????????????????????????????????????????????????????????????????????????????????????????????????????????????????????????????????????????????????????????????????????????????????????????????????????????????????????????????????????????????????????????

			>Aechmea_bambusoides
			????????????????????????????????????????????????????????????????????????????????????????????????????????????????????????????????????????????????????????????????????????????????????????????????????????????????????????????????????????????????????????????????????????????????????????????????????????????????????????????????????????????????????????????????????????????????

			>Aechmea_blanchetiana
			????????????????????????????????????????????????????????????????????????????????????????????????????????????????????????????????????????????????????????????????????????????????????????????????????????????????????????????????????????????????????????????????????????????????????????????????????????????????????????????????????????????????????????????????????????????????

			>Aechmea_blumenavii
			????????????????????????????????????????????????????????????????????????????????????????????????????????????????????????????????????????????????????????????????????????????????????????????????????????????????????????????????????????????????????????????????????????????????????????????????????????????????????????????????????????????????????????????????????????????????

			>Aechmea_bocainensis
			TCGCAGAARGCTCTTTCCCTGCCTACTGGAATGGGCATTCTCTGCGCGAGCCCGAAAGCTCTAGAAGCTTCCAAAACAGCCAAGTCGGTCAGGGTTTTCTTCGACTGGAATGACTACTTGAAGTTCTACAAGCTCGGAACTTACTGGCCTTAACGCCGTCCATCCAACTGCTTTAYGGGCTGAGAGCAGCTTTGGATCTCATCTTTGAGGAGGGGCTCGACAATGTCATCGAAAGGCAYCGTCGCTTAGGAAAGGCCACAAGGTTAAGTAAAAGCTATACATCTAAATAGCCGCCTTCGCYGTGTTGGATAGATATATTACTTGGCACTATTGTTCTCTCCTAACAATGAGCAGACTTGCCGTCGAAG

			>Aechmea_bracteata
			????????????????????????????????????????????????????????????????????????????????????????????????????????????????????????????????????????????????????????????????????????????????????????????????????????????????????????????????????????????????????????????????????????????????????????????????????????????????????????????????????????????????????????????????????????????????

			>Aechmea_bracteata_var._pacifica
			????????????????????????????????????????????????????????????????????????????????????????????????????????????????????????????????????????????????????????????????????????????????????????????????????????????????????????????????????????????????????????????????????????????????????????????????????????????????????????????????????????????????????????????????????????????????

			>Aechmea_brassicoides
			????????????????????????????????????????????????????????????????????????????????????????????????????????????????????????????????????????????????????????????????????????????????????????????????????????????????????????????????????????????????????????????????????????????????????????????????????????????????????????????????????????????????????????????????????????????????

			>Aechmea_brevicollis
			????????????????????????????????????????????????????????????????????????????????????????????????????????????????????????????????????????????????????????????????????????????????????????????????????????????????????????????????????????????????????????????????????????????????????????????????????????????????????????????????????????????????????????????????????????????????

			>Aechmea_bromeliifolia
			TCGCAGAAAGCTCTTTCCCTGCCTACTGGAATGGGCATTCTCTGCGCGAGCCCGAAAGCTCTAGAAGCTTCCAAAACAGCCAAGTCGGTCAGGGTTTTCTTCGACTGGAATGACTACTTGAAGTTCTACAAGCTCGGAACTTACTGGCCTTAACGCCGTCCATCCAACTGCTTTACGGGCTGAGAGCAGCTTTGGATCTCATCTTTGAGGAGGGGCTCGACAACGTCATCGAAAGGCACCGTCGCTTAGGAAAGGCCACAAGGTCAAGTAAAAGCT????????????????????????????????????????????????????????????????????????????????????????????

			>Aechmea_bromeliifolia_var._albobracteata
			????????????????????????????????????????????????????????????????????????????????????????????????????????????????????????????????????????????????????????????????????????????????????????????????????????????????????????????????????????????????????????????????????????????????????????????????????????????????????????????????????????????????????????????????????????????????

			>Aechmea_bruggeri
			????????????????????????????????????????????????????????????????????????????????????????????????????????????????????????????????????????????????????????????????????????????????????????????????????????????????????????????????????????????????????????????????????????????????????????????????????????????????????????????????????????????????????????????????????????????????

			>Aechmea_caesia
			????????????????????????????????????????????????????????????????????????????????????????????????????????????????????????????????????????????????????????????????????????????????????????????????????????????????????????????????????????????????????????????????????????????????????????????????????????????????????????????????????????????????????????????????????????????????

			>Aechmea_calyculata
			????????????????????????????????????????????????????????????????????????????????????????????????????????????????????????????????????????????????????????????????????????????????????????????????????????????????????????????????????????????????????????????????????????????????????????????????????????????????????????????????????????????????????????????????????????????????

			>Aechmea_candida
			TCGCAGAAAGCTCTTTCCCTGCCTACTGGAATGGGGATTATCTGCGCGAGCCCGAAAGCTCTAGAAGCTTCCAAAACAGCCAAGTCGGTCAGGGTTTTCTTCGACTGGAATGACTACTTGAAGTTCTACAAGCTCGGAACTTACTGGCCTTAACGCCGTCCATCCAACTGCTTTACGGGCTGAGAGCAGCTTTGGACCTCATCTTCGAGGAGGGGCTCGACAATGTCATCGAAAGGCATCGTCGCTTAGGAAAGGCCACAAGGTTAAGTAAAAGCTATACACCTAAATAGCCGCCTTCGCAGTGTTGGATAGATATATTACTTGGCACTGTTGTTCTCTTGTAACAATGGGCAGACTTGCCGTCGATG

			>Aechmea_capixabae
			????????????????????????????????????????????????????????????????????????????????????????????????????????????????????????????????????????????????????????????????????????????????????????????????????????????????????????????????????????????????????????????????????????????????????????????????????????????????????????????????????????????????????????????????????????????????

			>Aechmea_carvalhoi
			????????????????????????????????????????????????????????????????????????????????????????????????????????????????????????????????????????????????????????????????????????????????????????????????????????????????????????????????????????????????????????????????????????????????????????????????????????????????????????????????????????????????????????????????????????????????

			>Aechmea_castanea
			????????????????????????????????????????????????????????????????????????????????????????????????????????????????????????????????????????????????????????????????????????????????????????????????????????????????????????????????????????????????????????????????????????????????????????????????????????????????????????????????????????????????????????????????????????????????

			>Aechmea_castelnavi
			????????????????????????????????????????????????????????????????????????????????????????????????????????????????????????????????????????????????????????????????????????????????????????????????????????????????????????????????????????????????????????????????????????????????????????????????????????????????????????????????????????????????????????????????????????????????

			>Aechmea_castelnavii
			????????????????????????????????????????????????????????????????????????????????????????????????????????????????????????????????????????????????????????????????????????????????????????????????????????????????????????????????????????????????????????????????????????????????????????????????????????????????????????????????????????????????????????????????????????????????

			>Aechmea_caudata
			TCGCAGAAAGCTCTTTCCCTGCCTACTGGAATGGGGATTATCTGCGCGAGCCCGAAAGCTCTAGAAGCTTCCAAAACAGCCAAGTCGGTCAGGGTTTTCTTCGACTGGAATGACTACTTGAAGTTCTACAAGCTCGGAACTTACTGGCCTTAACGCCGTCCATCCAACTGCTTTACGGGCTGAGAGCAGCTTTGGACCTCATCTTYGAGGAGGGGCTCGACAATGTCATCGAAAGGCATCGTCGCTTAGGAAAGGCCACAAGGTTAAGTAAAAGCTATACATCTAAATAGCCGCCTTCRCCGTGTTGGATAGATATATTACTTGGCACTGTTGTTCTCTTGTAACAATGAGCAGACTTGCCGTCGAAG

			>Aechmea_cephaloides
			????????????????????????????????????????????????????????????????????????????????????????????????????????????????????????????????????????????????????????????????????????????????????????????????????????????????????????????????????????????????????????????????????????????????????????????????????????????????????????????????????????????????????????????????????????????????

			>Aechmea_chantinii
			????????????????????????????????????????????????????????????????????????????????????????????????????????????????????????????????????????????????????????????????????????????????????????????????????????????????????????????????????????????????????????????????????????????????????????????????????????????????????????????????????????????????????????????????????????????????

			>Aechmea_chrysocoma
			????????????????????????????????????????????????????????????????????????????????????????????????????????????????????????????????????????????????????????????????????????????????????????????????????????????????????????????????????????????????????????????????????????????????????????????????????????????????????????????????????????????????????????????????????????????????

			>Aechmea_coelestis
			TCGCAGAAAGCTCTTTCCCTGCCTACTGGAATGGGGATTATCTGCGCGAGCCCGAAAGCTCTAGAAGCTTCCAAAACAGCCAAGTCGGTCAGGGTTTTCTTCGACTGGAATGACTACTTGAAGTTCTACAAGCTCGGAACTTACTGGCCTTAACGCCGTCCATCCAACTGCTTTACGGGCTGAGAGCAGCTTTGGACCTCATCTTCGAGGAGGGGCTCGACAATGTCATCGAAAGGCATCGTCGCTTAGGAAAGGCCACAAGGTTAAGTAAAAGCTATACACCTAAATAGCCGCCTTCGCAGTGTTGGATAGATATATTACTTGGCACTGTTGTTCTCTTGTAACAATGGGCAGACTTGCCGTCGATG

			>Aechmea_colombiana
			????????????????????????????????????????????????????????????????????????????????????????????????????????????????????????????????????????????????????????????????????????????????????????????????????????????????????????????????????????????????????????????????????????????????????????????????????????????????????????????????????????????????????????????????????????????????

			>Aechmea_comata
			TCGCAGAAAGCTCTTTCCCTGCCTACTGGAATGGGGATTATCTGCGCGAGCCCGAAAGCTCTAGAAGCTTCCAAAACAGCCAAGTCGGTCAGGGTTTTCTTCGACTGGAATGACTACTTGAAGTTCTACAAGCTCGGAACTTACTGGCCTTAACGCCGTCCATCCAACTGCTTTACGGGCTGAGAGCAGCTTTGGACCTCATCTTTGAGGAGGGGCTCGACAATGTCATCGAAAGGCATCGTCGCTTAGGAAAGGCCACAAGGTTAAGTACAAGCTATACATCTAAATAGCCGCCTTCGCCGTGTTGGATAGATATATTACTTGGCACTGTTGTTATCTTGTAACAATGAGCAGACTTGCCGTCGAAG

			>Aechmea_contracta
			TCGCAGAAAGCTCTTTCCCTGCCTACTGGAATGGGCATTCTCTGCGCGAGCCCGAAAGCTCTAGAAGCATCCAAAACAGCCAAGTCGGTCAGGGTTTTCTTCGACTGGAATGACTACTTGAAGTTCTACAAGCTCGGAACTTACTGGCCTTAACGCCGTCCATCCAACTGCTTTACGGGCTGAGAGCAGCTTTGGATCTCATCTTCGAGGAGGGGCTCGACAAYGTCATCGAAAGGCACCGTCGCTTAGGAAAGGCCACAAGGTTAAGTAAAAGCTATACATCT-AATAGCCGCCTTCGCCGTGTTGGATAGATATATTACTCGGCGCTATTGTTCTCTCGTAACAATGAGCAGACTTGCCGTCGAAG

			>Aechmea_correia-araujoi
			????????????????????????????????????????????????????????????????????????????????????????????????????????????????????????????????????????????????????????????????????????????????????????????????????????????????????????????????????????????????????????????????????????????????????????????????????????????????????????????????????????????????????????????????????????????????

			>Aechmea_costantinii
			????????????????????????????????????????????????????????????????????????????????????????????????????????????????????????????????????????????????????????????????????????????????????????????????????????????????????????????????????????????????????????????????????????????????????????????????????????????????????????????????????????????????????????????????????????????????

			>Aechmea_cucullata
			TCGCAGAAAGCTCTTTCCCTGCCTACTGGAATGGGCATTCTCTGCGCGAGCCCGAAAGCTCTAGAAGCATCCAAAACAGCCAAGTCGGTCAGGGTTTTCTTCGACTGGAATGACTACTTGAAGTTCTACAAGCTCGGAACTTACTGGCCTTAACGCCGTCCATCCAACTGCTTTACGGGCTGAGAGCAGCTTTGGATCTCATCTTCGAGGAGGGGCTCGACAATGTCATCGAAAGGCACCGTCGCTTAGGAAAGGCCACAAGGTTAAGTAAAAGCTATACATCT-AATAGCCGCCTTCGCCGTGTTGGATAGATATATTACTCGGCGCTATTGTTCTCTCGTAACAATGAGCAGACTTGCCGTCGAAG

			>Aechmea_cylindrata
			????????????CTCTCCCTGCCTACTGGAATGGGGATTATCTGCGCGAGCCCGAAAGCTCTAGAAGCTTCCAAAACAGCCAAGTCGGTCAGGGTTTTCTTCGACTGGAATGACTACTTGAAGTTCTACAAGCTCGGAACTTACTGGCCTTAACGCCGTCCATCCAACTGCTTTACGGGCTGAGAGCAGCTTTGGACCTCATCTTCGAGGAGGGGCTCGACAATGTCATCGAAAGGCATCGTCGCTTAGGAAAGGCCACAAGGTTAAGTAAAAGCTATACATCTAAATAGCCACCTTCGCCGTGTTGGATAGATATATTGCTTGGCACTGTTGTTCTCTTGTAACAATGAGCAGACTTGCCGTCGAAG

			>Aechmea_dactylina
			????????????????????????????????????????????????????????????????????????????????????????????????????????????????????????????????????????????????????????????????????????????????????????????????????????????????????????????????????????????????????????????????????????????????????????????????????????????????????????????????????????????????????????????????????????????????

			>Aechmea_dealbata
			????????????????????????????????????????????????????????????????????????????????????????????????????????????????????????????????????????????????????????????????????????????????????????????????????????????????????????????????????????????????????????????????????????????????????????????????????????????????????????????????????????????????????????????????????????????????

			>Aechmea_digitata
			????????????????????????????????????????????????????????????????????????????????????????????????????????????????????????????????????????????????????????????????????????????????????????????????????????????????????????????????????????????????????????????????????????????????????????????????????????????????????????????????????????????????????????????????????????????????

			>Aechmea_discordiae
			????????????????????????????????????????????????????????????????????????????????????????????????????????????????????????????????????????????????????????????????????????????????????????????????????????????????????????????????????????????????????????????????????????????????????????????????????????????????????????????????????????????????????????????????????????????????

			>Aechmea_disjuncta
			????????????????????????????????????????????????????????????????????????????????????????????????????????????????????????????????????????????????????????????????????????????????????????????????????????????????????????????????????????????????????????????????????????????????????????????????????????????????????????????????????????????????????????????????????????????????

			>Aechmea_distichantha
			????????????????????????????????????????????????????????????????????????????????????????????????????????????????????????????????????????????????????????????????????????????????????????????????????????????????????????????????????????????????????????????????????????????????????????????????????????????????????????????????????????????????????????????????????????????????

			>Aechmea_distichantha_var._glaziovii
			????????????????????????????????????????????????????????????????????????????????????????????????????????????????????????????????????????????????????????????????????????????????????????????????????????????????????????????????????????????????????????????????????????????????????????????????????????????????????????????????????????????????????????????????????????????????

			>Aechmea_egleriana
			TCGCAGAAAGCTCTTTCCCTGCCAACCGGAATGGGCATTCTCTGCGCGAGCCCGAAAGCTCTAGAAGCATCCAAAACAGCCAAGTCGGTCAGGGTTTTCTTCGACTGGAATGACTACTTGAAGTTCTACAAGCTCGGAACTTACTGGCCTTAACGCCATCCATCCAACTGCTTTACGGGCTGAGAGCAGCTTTGGATCTCATCTTTGAGGAGGGGCTCGACAACGTCATCGAAAGGCATCGTCGCTTAGGAAAGGCCACAAGGTTAAGTAAAAGCTATACATCT-AATAGCCGCCTTCGCCGTGTTGGATAGATATATTACTTGGCACTATTGTTCTCTTGTAACGATGAGCAGACTTGCTGTCGAAG

			>Aechmea_eurycorymbus
			????????????????????????????????????????????????????????????????????????????????????????????????????????????????????????????????????????????????????????????????????????????????????????????????????????????????????????????????????????????????????????????????????????????????????????????????????????????????????????????????????????????????????????????????????????????????

			>Aechmea_farinosa
			????????????????????????????????????????????????????????????????????????????????????????????????????????????????????????????????????????????????????????????????????????????????????????????????????????????????????????????????????????????????????????????????????????????????????????????????????????????????????????????????????????????????????????????????????????????????

			>Aechmea_fasciata
			TCGCAGAAAGCTCTTTCCCTGCCTACTGGAATGGGCATTCTCTGCGCGAGCCCGAAAGCTCTAGAAGCTTCCAAAACAGCCAAGTCCGTCAGGGTTTTCTTCGACTGGAATGACTACTTGAAGTTCTACAAGCTCGGAACTTACTGGCCTTAACGCCGTCCATCCAACTGCTTTACGGGCTGAGAGCAGCTTTGGATCTCATCTTTGAGGAGGGGCTCGACAATGTCATCGAAAGGCACCGTCGCTTAGGAAAGGCCACAAGGTTAAGTAAAAGCTATACATCTAAATAGCCGCCTTCGCCGTGTCGGATAGATATTTTACATTGCACTGTTGTTCTCTTGTAACAATGAGCAGACTTGCCGTCGAAG

			>Aechmea_filicaulis
			TCGCAGAAAGCTCTTTCCCTGCCTACTGGAATGGGCATTCTCTGCGCGAGCCCGAAAGCTCTAGAAGCATCCAAAACAGCCAAGTCGGTCAGGGTTTTCTTCGACTGGAATGACTACTTGAAGTTCTACAAGCTCGGAACTTACTGGCCTTAACGCCGTCCATCCAACTGCTTTACGGGCTGAGAGCAGCTTTGGATCTCATCTTTGAGGAGGGGCTCGACAACGTCATCGAAAGGCATCGTCGCTTGGGAAAGGCCACAAGGTTAAGTAAACGCTATACGTCT-AATAGCCGCCTTCGCCGTGTCGGATAGATATTTTACTTTGCACTATTGTCCTCTTGTAACAATGAGCAGACTTGCTGTCGAAG

			>Aechmea_flavorosea
			TCGCAGAAAGCTCTTTCCCTGCCTACTGGAATGGGCATTCTCTGCGCGAGCCCGAAAGCTCTAGAAGCTTCCAAAACAGCCAAGTCGGTCAGGGTTTTCTTCGACTGGAATGACTACTTGAAGTTCTACAAGCTCGGAACTTACTGGCCTTAACGCCGTCCATCCAACTGCTTTACGGGCTGAGAGCAGCTTTGGATCTCATCTTTGAGGAGGGGCTCGACAATGTCATCGAAAGGCACCGTCGCTTAGGAAAGGCCACAAGGTTAAGTAAAAGCTATACATCTAAATAGCCGCCTTCGCCGTGTTGGATAGATATTTTACATTGCACTGTTGTTCTCTTGTAACAATGAGCAGACTTGCCGTCGAAG

			>Aechmea_floribunda
			????????????????????????????????????????????????????????????????????????????????????????????????????????????????????????????????????????????????????????????????????????????????????????????????????????????????????????????????????????????????????????????????????????????????????????????????????????????????????????????????????????????????????????????????????????????????

			>Aechmea_fosteriana
			????????????????????????????????????????????????????????????????????????????????????????????????????????????????????????????????????????????????????????????????????????????????????????????????????????????????????????????????????????????????????????????????????????????????????????????????????????????????????????????????????????????????????????????????????????????????

			>Aechmea_fulgens
			????????????????????????????????????????????????????????????????????????????????????????????????????????????????????????????????????????????????????????????????????????????????????????????????????????????????????????????????????????????????????????????????????????????????????????????????????????????????????????????????????????????????????????????????????????????????

			>Aechmea_fulgens_var._discolor
			????????????????????????????????????????????????????????????????????????????????????????????????????????????????????????????????????????????????????????????????????????????????????????????????????????????????????????????????????????????????????????????????????????????????????????????????????????????????????????????????????????????????????????????????????????????????

			>Aechmea_gamosepala
			????????????????????????????????????????????????????????????????????????????????????????????????????????????????????????????????????????????????????????????????????????????????????????????????????????????????????????????????????????????????????????????????????????????????????????????????????????????????????????????????????????????????????????????????????????????????

			>Aechmea_gracilis
			TCGCAGAAAGCTCTTTCCCTGCCTACTGGAATGGGGATTATCTGCGCGAGCCCGAAAGCTCTAGAAGCTTCCAAAACAGCCAAGTCGGTCAGGGTTTTCTTCGACTGGAATGACTACTTGAAGTTCTACAAGCTCGGAACTTACTGGCCTTAACGCCGTCCATCCAACTGCTTTACGGGCTGAGAGCAGCTTTGGACCTCATCTTCGAGGAGGGGCTCGACAATGTCATCGAAAGGCATCGTCGCTTAGGAAAGGCCACAAGGTTAAGTAAAAGCTATACATCTAAATAGCCACCTTCGCCGTGTTGGATAGATATATTGCTTGGCACTGTTGTTCTCTTGTAACAATGAGCAGACTTGCCGTCGAAG

			>Aechmea_guainumbiorum
			????????????????????????????????????????????????????????????????????????????????????????????????????????????????????????????????????????????????????????????????????????????????????????????????????????????????????????????????????????????????????????????????????????????????????????????????????????????????????????????????????????????????????????????????????????????????

			>Aechmea_haltonii
			????????????????????????????????????????????????????????????????????????????????????????????????????????????????????????????????????????????????????????????????????????????????????????????????????????????????????????????????????????????????????????????????????????????????????????????????????????????????????????????????????????????????????????????????????????????????

			>Aechmea_heterosepala
			????????????????????????????????????????????????????????????????????????????????????????????????????????????????????????????????????????????????????????????????????????????????????????????????????????????????????????????????????????????????????????????????????????????????????????????????????????????????????????????????????????????????????????????????????????????????

			>Aechmea_hoppii
			TCGCAGAAAGCTCTTTCCCTGCCTACTGGAATGGGCATTCTCTGCGCGAGCCCGAAAGCTCTAGAAGCATCCAAAACAGCCAAGTCGGTCAGGGTTTTCTTCGACTGGAATGACTACTTGAAGTTCTACAAGCTCGGAACTTACTGGCCTTAACGCCGTCCATCCAACTGCTTTACGGGCTGAGAGCAGCTTTGGATCTCATCTTTGAGGAGGGGCTCGACAATGTCATCGAAAGGCACCGTCGCTTAGGAAAGGCCACAAGGTTAAGTAAAAGCTATACATCT-AATAGCCGCCTYCGCCGTGTTGSATAGATATATTACTTGGCRCTTTTGTTCTCTTGTAACAATGRGCAGACTTGCTGTCGAAG

			>Aechmea_jungurudoensis
			????????????????????????????????????????????????????????????????????????????????????????????????????????????????????????????????????????????????????????????????????????????????????????????????????????????????????????????????????????????????????????????????????????????????????????????????????????????????????????????????????????????????????????????????????????????????

			>Aechmea_kertesziae
			????????????????????????????????????????????????????????????????????????????????????????????????????????????????????????????????????????????????????????????????????????????????????????????????????????????????????????????????????????????????????????????????????????????????????????????????????????????????????????????????????????????????????????????????????????????????

			>Aechmea_kleinii
			????????????????????????????????????????????????????????????????????????????????????????????????????????????????????????????????????????????????????????????????????????????????????????????????????????????????????????????????????????????????????????????????????????????????????????????????????????????????????????????????????????????????????????????????????????????????

			>Aechmea_lamarchei
			????????????????????????????????????????????????????????????????????????????????????????????????????????????????????????????????????????????????????????????????????????????????????????????????????????????????????????????????????????????????????????????????????????????????????????????????????????????????????????????????????????????????????????????????????????????????

			>Aechmea_leptantha
			????????????????????????????????????????????????????????????????????????????????????????????????????????????????????????????????????????????????????????????????????????????????????????????????????????????????????????????????????????????????????????????????????????????????????????????????????????????????????????????????????????????????????????????????????????????????

			>Aechmea_leucolepis
			????????????????????????????????????????????????????????????????????????????????????????????????????????????????????????????????????????????????????????????????????????????????????????????????????????????????????????????????????????????????????????????????????????????????????????????????????????????????????????????????????????????????????????????????????????????????

			>Aechmea_lilacinantha
			TCGCAGAAAGCTCTTTCCCTGCCTACCGGAATGGGCATTCTYTGCGCGAGCCCGAAAGCTCTAGAAGCTTCCAAAACAGCCAAGTCGGTCAGGGTTTTCTTCGACTGGAATGACTACTTGAAGTTCTACAAGCTCGGAACCTACTGGCCTTAACGCCGTCCATCCAACTGCTTTACGGGCTGAGAGCAGCTTTGGATCTCATCTTTGAGGAGGGGCTCGACAATGTCATCGAAAGGCACCGTCGCTTAGGAAAGGCGACAAGGTTAAGTAAAAGCTATACATCT-AATAGCCGCTTTCGCCGTGTTGCATAGATATATTACTTGGTACTGTTGTTCTCTTGTAACAATGAGCAGACTTGCCGTCGAAG

			>Aechmea_longicuspis_var._kuntzeana
			????????????????????????????????????????????????????????????????????????????????????????????????????????????????????????????????????????????????????????????????????????????????????????????????????????????????????????????????????????????????????????????????????????????????????????????????????????????????????????????????????????????????????????????????????????????????

			>Aechmea_lueddemanniana
			TCGCAGAAAGCTCTTTCCCTGCCTACTGGAATGGGCATTCTCTGCGCGAGCCCGAAAGCTCTAGAAGCTTCCAAAACAGCCAAGTCGGTCAGGGTTTTCTTCGACTGGAATGACTACTTGAAGTTCTACAAGCTCGGAACTTACTGGCCTTAACGCCTTCCATCCAACTACTTTACGGGCTGAGAGCAGCTTTGGATCTCATCTTTGAGGAGGGGCTCGACAATGTCATCGAAAGGCATCGTCGCTTAGGAAAGGCCACAAGGTTAAGTAAAAGCTATACATCT-AATAGCCGCTTTCGCCGCGTTGGATAGATATTTTACTTTGCGCTATTGTCCTCTTGTAACAATGAGCAGACTTGCCGTCGAAG

			>Aechmea_maasii
			????????????????????????????????????????????????????????????????????????????????????????????????????????????????????????????????????????????????????????????????????????????????????????????????????????????????????????????????????????????????????????????????????????????????????????????????????????????????????????????????????????????????????????????????????????????????

			>Aechmea_magdalenae
			????????????????????????????????????????????????????????????????????????????????????????????????????????????????????????????????????????????????????????????????????????????????????????????????????????????????????????????????????????????????????????????????????????????????????????????????????????????????????????????????????????????????????????????????????????????????

			>Aechmea_manzanaresiana
			????????????????????????????????????????????????????????????????????????????????????????????????????????????????????????????????????????????????????????????????????????????????????????????????????????????????????????????????????????????????????????????????????????????????????????????????????????????????????????????????????????????????????????????????????????????????

			>Aechmea_marauensis
			????????????????????????????????????????????????????????????????????????????????????????????????????????????????????????????????????????????????????????????????????????????????????????????????????????????????????????????????????????????????????????????????????????????????????????????????????????????????????????????????????????????????????????????????????????????????

			>Aechmea_marginalis
			TCGCAGAAAGCTCTTTCCCTGCCTACTGGAATGGGCATTCTCTGCGCGAGCCCGAAAGCTCTAGAGGCTTCCAAAACAGCCAAGTCGGTCAGGGTTTTCTTCGACTGGAATGACTACTTGAAGTTCTACAAGCTCGGAACTTACTGGCCTTAACGCCTTCCATCCAACTACTTTACGGGCTGAGAGCAGCTTTGGATCTCATCTTTGAGGAGGGGCTCGACAATGTCATCGAAAGGCATCGTCGCTTAGGAAAGGCCACAAGGTTAAGCAAAAGCTATACATCT-AATAGCCGCTTTCGCCGCGTTGGATAGATATTTTACTTTGCGCTATTGTCCTCTTGTAACAATGAGCAGACTTGCTGTCGAAG

			>Aechmea_mariae-reginae
			TCGCAGAAAGCTCTTTCCCTGCCTACTGGAATGGGCATTCTCTGCGCGAGCCCGAAAGCTCTAGARGCTTCCAAAACAGCCAAGTCGGTCAGGGTTTTCTTCGACTGGAATGACTACTTGAAGTTCTACAAGCTCGGAACTTACTGGCCTTAACGCCTTCCATCCAACTACTTTACGGGCTGAGAGCAGCTTTGGATCTCATCTTTGAGGAGGGGCTCGACAATGTCATCGAAAGGCATCGTCGCTTAGGAAAGGCCACAAGGTTAAGYAAAAGCTATACATCT-AATAGCCGCTTTCGCCGCGTTGGATAGATATTTTACTTTGCGCTATTGTCCTCTTGTAACAATGAGCAGACTTGCTGTCGAAG

			>Aechmea_melinonii
			????????????????????????????????????????????????????????????????????????????????????????????????????????????????????????????????????????????????????????????????????????????????????????????????????????????????????????????????????????????????????????????????????????????????????????????????????????????????????????????????????????????????????????????????????????????????

			>Aechmea_mertensii
			????????????????????????????????????????????????????????????????????????????????????????????????????????????????????????????????????????????????????????????????????????????????????????????????????????????????????????????????????????????????????????????????????????????????????????????????????????????????????????????????????????????????????????????????????????????????

			>Aechmea_mexicana
			????????????????????????????????????????????????????????????????????????????????????????????????????????????????????????????????????????????????????????????????????????????????????????????????????????????????????????????????????????????????????????????????????????????????????????????????????????????????????????????????????????????????????????????????????????????????

			>Aechmea_microcephala
			????????????????????????????????????????????????????????????????????????????????????????????????????????????????????????????????????????????????????????????????????????????????????????????????????????????????????????????????????????????????????????????????????????????????????????????????????????????????????????????????????????????????????????????????????????????????

			>Aechmea_miniata
			????????????????????????????????????????????????????????????????????????????????????????????????????????????????????????????????????????????????????????????????????????????????????????????????????????????????????????????????????????????????????????????????????????????????????????????????????????????????????????????????????????????????????????????????????????????????

			>Aechmea_miniata_var._discolor
			????????????????????????????????????????????????????????????????????????????????????????????????????????????????????????????????????????????????????????????????????????????????????????????????????????????????????????????????????????????????????????????????????????????????????????????????????????????????????????????????????????????????????????????????????????????????

			>Aechmea_mira
			TCGCAGAAAGCTCTATCCCTGCCTACTGGAATGGGCATTCTTTGCGCGAGCCCGAAAGCTCTAGAAGCTTCCAAAACAGCCAAGTCGGTCAGGGTTTTCTTCGACTGGAATGACTACTTGAAGTTCTACAAGCTCGGAACTTACTGGCCTTAACGCCTTCCATCCAACTGCTTTACGGGCTGAGAGCAGCTTTGGATCTCATCTTTGAGGAGGGGCTCGACAATGTCATCGAAAGGCACCGTCGCTTAGGAAAGGCCACAAGGTTAAGTAAAAGCTATACATCT-AATAGCCGCCTTCGCCGTGTTGGATAGATATATTACTTGGTACTGTTGTTCTCTTGTAACAATGAGCAGACTTGCTGTCGAAG

			>Aechmea_moorei
			TCGCAGAAAGCTCTTTCCCTGCCTACYGGAATGGGCATTCTCTGCGCGAGCCCGAAAGCTCTAGAAGCATCCAAAACAGCCAAGTCGGTCAGGGTTTTCTTCGACTGGAATGACTACTTGAAGTTCTACAAGCTCGGAACTTACTGGCCTTAACGCCGTCCATCCAACTGCTTTACGGGCTGAGAGCAGCTTTGGATCTCATCTTCGAGGAGGGGCTCGACAATGTCATCGAAAGGCACCGTCGCTTAGGAAAGGCCACAAGGTTAAGTAAAAGCTATACATCT-AATAGCCGCCTTCGCCGTGTTGGATAGATATATTACTCGGCGCTATTGTTCTCTCGTAACAATGAGCAGACTTGCCGTCGAAG

			>Aechmea_murcae
			????????????????????????????????????????????????????????????????????????????????????????????????????????????????????????????????????????????????????????????????????????????????????????????????????????????????????????????????????????????????????????????????????????????????????????????????????????????????????????????????????????????????????????????????????????????????

			>Aechmea_muricata
			TCGCAGAAAGCTCTTTCCCTGCCTACTGGAATGGGCATTCTTTGCGCGAGCCCGAAAGCTCTAGAAGCTTCCAAAACAGCCAAGTCGGTCAGGGTTTTCTTCGACTGGAATGACTACTTGAAGTTCTACAAGCTCGGAACTTACTGGCCTTAACGCCGTCCATCCAACTGCTTTACGGGCTGAGAGCAGCTTTGGATCTCATCTTTGAGGAGGGGCTCGACAATGTCATCGAAAGGCACCGTCGCTTAGGAAAGGCCACAAGGTTAAGTAAAAGCTATACATCT-------------------------CTGATATATTAGTTGGTACTGTTGTTCTCTTGTAACAATGAGCAGACTTGCCGTCGAAG

			>Aechmea_mutica
			????????????????????????????????????????????????????????????????????????????????????????????????????????????????????????????????????????????????????????????????????????????????????????????????????????????????????????????????????????????????????????????????????????????????????????????????????????????????????????????????????????????????????????????????????????????????

			>Aechmea_nallyi
			????????????????????????????????????????????????????????????????????????????????????????????????????????????????????????????????????????????????????????????????????????????????????????????????????????????????????????????????????????????????????????????????????????????????????????????????????????????????????????????????????????????????????????????????????????????????

			>Aechmea_napoensis
			????????????????????????????????????????????????????????????????????????????????????????????????????????????????????????????????????????????????????????????????????????????????????????????????????????????????????????????????????????????????????????????????????????????????????????????????????????????????????????????????????????????????????????????????????????????????

			>Aechmea_nidularioides
			TCGCAGAAAGCTCTTTCCCTGCCTACTGGAATGGGCATTCTCTGCGCGAGCCCGAAAGCTCTAGAAGCTTCCAAAACAGCCAAGTCGGTCAGGGTTTTCTTCGACTGGARTGACTACTTGAAGTTCTACAAGCTCGGAACTTACTGGCCTTAACGCCGTCCATCCAACTGCTTTACGGGCTGAGAGCAGCTTTGGATCTCATCTTTGAGGAGGGGCTCGACAAYGTCATCGAAAGGCACCGTCGCTTAGGAAAGGCCACAAGGTTAAGTAAAAGCTATACATCT-AATAGCCGCCTTCGCCGTGTTGGATAGATATATTACTTGGCGCYWTYGTTCTCTTGTAACAATGAGCAGACTTGCTGTCGAAG

			>Aechmea_nudicaulis
			TCGCAGAAAGCTCTTTCCCTGCCTACTGGAATGGGCATTCTCTGCGCGAGCCCGAAAGCTCTAGAAGCTTCCAAAACAGCCAAGTCGGTCAGGGTTTTCTTCGACTGGAATGACTACTTGAAGTTCTACAAGCTCGGAACTTACTGGCCTTAACGCCGTCCATCCAACTGCTTTACGGGCTGAGAGCAGCTTTGGACCTCATCTTTGAGGAGGGGCTCGACAATGTCATCGAAAGGCATCGTCGCTTAGGAAAGGCCACAAGGTTAAGTAAAAGCTATACATCTAAATAGCCGCCTTCGCCGTGTCGGATAGATATATTACTTGGCACTGTTGTTCTCTTGTAACAATGAGCAGACTTGCCGTCGAAG

			>Aechmea_organensis
			????????????????????????????????????????????????????????????????????????????????????????????????????????????????????????????????????????????????????????????????????????????????????????????????????????????????????????????????????????????????????????????????????????????????????????????????????????????????????????????????????????????????????????????????????????????????

			>Aechmea_orlandiana
			????????????????????????????????????????????????????????????????????????????????????????????????????????????????????????????????????????????????????????????????????????????????????????????????????????????????????????????????????????????????????????????????????????????????????????????????????????????????????????????????????????????????????????????????????????????????

			>Aechmea_ornata
			????????????????????????????????????????????????????????????????????????????????????????????????????????????????????????????????????????????????????????????????????????????????????????????????????????????????????????????????????????????????????????????????????????????????????????????????????????????????????????????????????????????????????????????????????????????????

			>Aechmea_paniculata
			TCGCAGAAAGCTCTTTCCCTGCCTACYGGAATGGGCATTCTCTGCGCGAGCCCGAAAGCTCTAGAAGCATCCAAAACWGCCAAGTCGGTCAGGGTTTTCTTCGACTGGAGTGACTACTTGAAGTTCTACAAGCTCGGAACTTACTGGCCTTAACGCCGTCCATCCAACTGCTTTACGGGCTGAGAGCAGCTTTGGATCTCATCTTTGAGGAGGGGCTCGACAACGTCATCGAAAGGCATCGTCGCTTAGGAAAGGCCACAAGGTTAAGTAAAAGCTAYACATCT-AATAGCCGCTTTCGCCGTGTTGGATAGATATATTACTTGCCACTRTTGTTCTCTTGTAACAATGAGCAGACTTGCTGTCGAAG

			>Aechmea_paratiensis
			????????????????????????????????????????????????????????????????????????????????????????????????????????????????????????????????????????????????????????????????????????????????????????????????????????????????????????????????????????????????????????????????????????????????????????????????????????????????????????????????????????????????????????????????????????????????

			>Aechmea_patriciae
			????????????????????????????????????????????????????????????????????????????????????????????????????????????????????????????????????????????????????????????????????????????????????????????????????????????????????????????????????????????????????????????????????????????????????????????????????????????????????????????????????????????????????????????????????????????????

			>Aechmea_pectinata
			TCGCAGAAAGCTCTTTCCCTGCCTACTGGAATGGGCATTCTCTGCGCGAGCCCGAAAGCTCTAGAAGCTTCCAAAACAGCCAAGTCGGTCAGGGTTTTCTTCGACTGGAATGACTACTTGAAGTTCTACAAGCTCGGAACTTACTGGCCTTAACGCCTTCCATCCAACTACTTTACGGGCTGAGAGCAGCTTTGGATCTCATCTTTGAGGAGGGGCTCGACAATGTCATCGAAAGGCATCGTCGCTTAGGAAAGGCCACAAGGTCAAGTAAAAGCTATACATCA-AATAGCCGCTCTCGCCGCGTTGGATAGATATATAACTTGGCACTATTGTCCTCTTGTAACAATGAGCAGACTCGCTGTCGAAG

			>Aechmea_pedicellata
			????????????????????????????????????????????????????????????????????????????????????????????????????????????????????????????????????????????????????????????????????????????????????????????????????????????????????????????????????????????????????????????????????????????????????????????????????????????????????????????????????????????????????????????????????????????????

			>Aechmea_penduliflora
			????????????????????????????????????????????????????????????????????????????????????????????????????????????????????????????????????????????????????????????????????????????????????????????????????????????????????????????????????????????????????????????????????????????????????????????????????????????????????????????????????????????????????????????????????????????????

			>Aechmea_perforata
			????????????????????????????????????????????????????????????????????????????????????????????????????????????????????????????????????????????????????????????????????????????????????????????????????????????????????????????????????????????????????????????????????????????????????????????????????????????????????????????????????????????????????????????????????????????????

			>Aechmea_phanerophlebia
			TCGCAGAAAGCTCTTTCCCTGCCTACTGGAATGGGCATTCTCTGCGCGAGCCCGAAAGCTCTAGAAGCTTCCAAAACAGCCAAGTCGGTCAGGGTTTTCTTCGACTGGAATGACTACTTGAAGTTCTACAAGCTCGGAACCTACTGGCCTTAACGCCGTCCATCCAACTGCTTTACGGGCTGAGAGCAGCTTTGGATCTCATCTTTGAGGAGGGGCTCGACAATGTCATCGAAAGGCATCGTCGCTTAGGAAAGGCCACAAGGTTAAGTAAAAGCTATACATCT-AATAGCCGCCTTCGCCGTGTTGGATAGATATATTACTCGGCACTGTTGTTCTCTCGTAACAATGAGCAGACTTGCCGTCGAAG

			>Aechmea_pimenti-velosoi
			????????????????????????????????????????????????????????????????????????????????????????????????????????????????????????????????????????????????????????????????????????????????????????????????????????????????????????????????????????????????????????????????????????????????????????????????????????????????????????????????????????????????????????????????????????????????

			>Aechmea_pineliana
			????????????????????????????????????????????????????????????????????????????????????????????????????????????????????????????????????????????????????????????????????????????????????????????????????????????????????????????????????????????????????????????????????????????????????????????????????????????????????????????????????????????????????????????????????????????????

			>Aechmea_polyantha
			TCGCAGAAAGCTCTTTCCCTGCCAACCGGAATGGGCATTCTCTGCGCGAGCCCGAAAGCTCTAGAAGCATCCAAAACAGCCAAGTCGGTCMGGGTTTTCTTCGACTGGAATGACTACTTGAAGTTCTACAAGCTCGGAACTTACTGGCCTTAACGCCGTCCATCCAACTGCTYTACGGGCTGAGAGCAGCTTTGGATCTCATCTTTGAGGAGGGGCTCGACAACGTCATCGAAAGGCATCGTCGCTTAGGAAAGGCCACAAGGTTAAGTAAAAGCTATACATCT-AATAGCCGCCTTCGCCGTGTTGGATASATATATTACTTGGCACTATTGTTCTCTTGTAACGATGAGCAGACTTGCTGTCGAAG

			>Aechmea_pubescens
			????????????????????????????????????????????????????????????????????????????????????????????????????????????????????????????????????????????????????????????????????????????????????????????????????????????????????????????????????????????????????????????????????????????????????????????????????????????????????????????????????????????????????????????????????????????????

			>Aechmea_racinae
			????????????????????????????????????????????????????????????????????????????????????????????????????????????????????????????????????????????????????????????????????????????????????????????????????????????????????????????????????????????????????????????????????????????????????????????????????????????????????????????????????????????????????????????????????????????????

			>Aechmea_ramosa
			????????????????????????????????????????????????????????????????????????????????????????????????????????????????????????????????????????????????????????????????????????????????????????????????????????????????????????????????????????????????????????????????????????????????????????????????????????????????????????????????????????????????????????????????????????????????

			>Aechmea_recurvata
			????????????????????????????????????????????????????????????????????????????????????????????????????????????????????????????????????????????????????????????????????????????????????????????????????????????????????????????????????????????????????????????????????????????????????????????????????????????????????????????????????????????????????????????????????????????????

			>Aechmea_roberto-seidelii
			????????????????????????????????????????????????????????????????????????????????????????????????????????????????????????????????????????????????????????????????????????????????????????????????????????????????????????????????????????????????????????????????????????????????????????????????????????????????????????????????????????????????????????????????????????????????

			>Aechmea_rodriguesiana
			TCGCAGAAAGCTCTTTCCCTGCCAACCGGAATGGGCATTCTCTGCGCGAGCCCGAAAGCTCTAGAAGCATCCAAAACAGCCAAGTCGGTCAGGGTTTTCTTCGACTGGAATGACTACTTGAAGTTCTACAAGCTCGGAACTTACTGGCCTTAACGCCGTCCATCCAACTGCTTTACGGGCTGAGAGCAGCTTTGGATCTCATCTTTGAGGAGGGGCTCGACAACGTCATCGAAAGGCATCGTCGCTTAGGAAAGGCCACAAGGTTAAGTAAAAGCTATACATCT-AATAGCCGCCTTCGCCGTGTTGGATAGATATATTACTTGGCACTATTGTTCTCTTGTAACGATGAGCAGACTTGCTGTCGAAG

			>Aechmea_roeseliae
			TCGCAGAAAGCTCTTTCCCTGCCTACTGGAATGGGCATTCTCTGCGCGAGCCCGAAAGCTCTAGAAGCATCCAAAACAGCCAAGTCGGTCAGGGTTTTCTTCGACTGGAATGACTACTTGAAGTTCTACAAGCTCGGAACTTACTGGCCTTAACGCCGTCCATCCAACTGCTTTACGGGCTGAGAGCAGCTTTGGATCTCATCTTCGAGGAGGGGCTCGACAATGTCATCGAAAGGCACCGTCGCTTAGGAAAGGCCACAAGGTTAAGTAAAAGCTATACATCT-AATAGCTGCCTTCGCCGTGTTGGATAGATATATTACTCGGCGCTATTGTTCTCTCGTAACAATGAGCAGACTTGCCGTCGAAG

			>Aechmea_romeroi
			????????????????????????????????????????????????????????????????????????????????????????????????????????????????????????????????????????????????????????????????????????????????????????????????????????????????????????????????????????????????????????????????????????????????????????????????????????????????????????????????????????????????????????????????????????????????

			>Aechmea_rubiginosa
			????????????????????????????????????????????????????????????????????????????????????????????????????????????????????????????????????????????????????????????????????????????????????????????????????????????????????????????????????????????????????????????????????????????????????????????????????????????????????????????????????????????????????????????????????????????????

			>Aechmea_serragrandensis
			????????????????????????????????????????????????????????????????????????????????????????????????????????????????????????????????????????????????????????????????????????????????????????????????????????????????????????????????????????????????????????????????????????????????????????????????????????????????????????????????????????????????????????????????????????????????

			>Aechmea_servitensis
			????????????????????????????????????????????????????????????????????????????????????????????????????????????????????????????????????????????????????????????????????????????????????????????????????????????????????????????????????????????????????????????????????????????????????????????????????????????????????????????????????????????????????????????????????????????????

			>Aechmea_servitensis_var._exigua
			????????????????????????????????????????????????????????????????????????????????????????????????????????????????????????????????????????????????????????????????????????????????????????????????????????????????????????????????????????????????????????????????????????????????????????????????????????????????????????????????????????????????????????????????????????????????

			>Aechmea_sphaerocephala
			TCGCAGAAAGCTCTTTCCCTGCCTACTGGAATGGGCATTCTTTGCGCGAGCCCGAAAGCTCTAGAAGCTTCCAAAACAGCCAAGTCGGTCAGGGTTTTCTTCGACTGGAATGACTACTTGAAGTTCTACAAGCTCGGAACTTACTGGCCTTAACGCCGTCCATCCAACTGCTTTACGGGCTGAGAGCAGCTTTGGATCTCATCTTTGAGGAGGGGCTCGACAATGTCATCGAAAGGCACCGTCGCTTAGGAAAGGCCACAAGGTTAAGTAAAAGCTATACATCT-AACAGCCGCTTTCGCCGTGTTGGATAGATATATTACTTGGTACTGTTGTTCTCTTGTAACAATGAGCAGACTTGCCGTCGAAG

			>Aechmea_squarrosa
			????????????????????????????????????????????????????????????????????????????????????????????????????????????????????????????????????????????????????????????????????????????????????????????????????????????????????????????????????????????????????????????????????????????????????????????????????????????????????????????????????????????????????????????????????????????????

			>Aechmea_strobilacea
			????????????????????????????????????????????????????????????????????????????????????????????????????????????????????????????????????????????????????????????????????????????????????????????????????????????????????????????????????????????????????????????????????????????????????????????????????????????????????????????????????????????????????????????????????????????????

			>Aechmea_tayoensis
			TCGCAGAAAGCTCTTTCGATGCCKACTGGAATGGGCATTCTTTGCGCGAGCCCGAAAGCTCTAGAAGCTTCCAAAACAGCCAGGTCGGTCAGGGTTTTCTTCGACTGGAATGACTACTTGAAGTTCTACAAGCTCGGAACTTACTGGCCTTAACGCCTTCCATCCAACTACTTTACGGGCTGAGAGCAGCTCTGGATCTCATCTTTGAGGAGGGGCTCGACAACGTCAACGAAAGGCATCGTCGCTTAGGAAAGGCCACAAGGTTAAGCAAGAGCTATRCGTCT-CATAGCCGCTTTCGCCGCGTTGGATAGATATATTACTTGGCGCTATTGTCCTCTCGTAACAATGAGCAGACTTGCTGTCGAAG

			>Aechmea_tessmannii
			TCGCAGAAAGCTCTTTCCCTGCCTACTGGAATGGGCATTCTCTGCGCGAGCCCGAAAGCTCTAGAAGCATCCAAAACAGCCAAGTCGGTCAGGGTTTTCTTCGACTGGAATGACTACTTGAAGTTCTACAAGCTCGGAACTTACTGGCCTTAACGCCGTCCATCCAACTGCTTTACGGGCTGAGAGCAGCTTTGGATCTCATCTTCGAGGAGGGGCTCGACAATGTCATCGAAAGGCACCGTCGCTTAGGAAAGGCCACAAGGTTAAGTAAAAGCTATACATCT-AATAGCCGCCTTCGCCGTGTTGGATAGATATATTACTCGGCGCTATTGTTCTCTCGTAACAATGAGCAGACTTGCCGTCGAAG

			>Aechmea_tillandsioides
			????????????????????????????????????????????????????????????????????????????????????????????????????????????????????????????????????????????????????????????????????????????????????????????????????????????????????????????????????????????????????????????????????????????????????????????????????????????????????????????????????????????????????????????????????????????????

			>Aechmea_triangularis
			TCGCAGAAAGCTCTTTCCCTGCCTACTGGAATGGGCATTCTTTGCGCGAGCCCGAAAGCTCTAGAAGCTTCCAAAACAGCCAAGTCGGTCAGGGTTTTCTTCGACTGGAATGACTACTTGAAGTTCTACAAGCTCGGAACTTACTGGCCTTAACGCCGTCCATCCAACTGCTTTACGGGCTGAGAGCAGCTCTGGATCTCATCTTTGAGGAGGGGCTCGACAACGTCATCGAAAGGCACCGTCGCTTAGGAAAGGCCACAAGGTCAAGTAAAAGCTATACATCT-AATAGCCGCCTTCGCCGTGTTGGATAGATATATTACTGTGTACTGTTGTTCTCTTGTAACAATGGGCAGACTTGCCGTCGAAG

			>Aechmea_triticina
			TCGCAGAAAGCTCTTTCCCTGCCTACTGGAATGGGCATTCTYTGCGCGAGCCCGAAAGCTCTAGAAGCTTCCAAAACAGCCAAGTCGGTCAGGGTTTTCTTCGACTGGAATGACTACTTGAAGTTCTACAAGCTCGGAACTTACTGGCCTTAACGCCGTCCATCCAACTGCTTTACGGGCTGAGAGCAGCTTTGGATCTCATCTTTGAGGAGGGGCTCGACAATGTCATCGAAAGGCATCGTCGCTTAGGAAAGGCCACAAGGTTAAGTAAAAGCTATACATCTAAATAGCCGCCTTCGCCGCGTTGGATAGATATATTACTCGGCACTATTGTCCTCTTGTAACAATGAGCAGACTTGCCGTCGAAG

			>Aechmea_vallerandii
			????????????????????????????????????????????????????????????????????????????????????????????????????????????????????????????????????????????????????????????????????????????????????????????????????????????????????????????????????????????????????????????????????????????????????????????????????????????????????????????????????????????????????????????????????????????????

			>Aechmea_vanhoutteana
			????????????????????????????????????????????????????????????????????????????????????????????????????????????????????????????????????????????????????????????????????????????????????????????????????????????????????????????????????????????????????????????????????????????????????????????????????????????????????????????????????????????????????????????????????????????????

			>Aechmea_victoriana
			????????????????????????????????????????????????????????????????????????????????????????????????????????????????????????????????????????????????????????????????????????????????????????????????????????????????????????????????????????????????????????????????????????????????????????????????????????????????????????????????????????????????????????????????????????????????

			>Aechmea_weilbachii
			????????????????????????????????????????????????????????????????????????????????????????????????????????????????????????????????????????????????????????????????????????????????????????????????????????????????????????????????????????????????????????????????????????????????????????????????????????????????????????????????????????????????????????????????????????????????

			>Aechmea_werdermannii
			????????????????????????????????????????????????????????????????????????????????????????????????????????????????????????????????????????????????????????????????????????????????????????????????????????????????????????????????????????????????????????????????????????????????????????????????????????????????????????????????????????????????????????????????????????????????

			>Aechmea_winkleri
			????????????????????????????????????????????????????????????????????????????????????????????????????????????????????????????????????????????????????????????????????????????????????????????????????????????????????????????????????????????????????????????????????????????????????????????????????????????????????????????????????????????????????????????????????????????????

			>Aechmea_woronowii
			????????????????????????????????????????????????????????????????????????????????????????????????????????????????????????????????????????????????????????????????????????????????????????????????????????????????????????????????????????????????????????????????????????????????????????????????????????????????????????????????????????????????????????????????????????????????

			>Aechmea_zebrina
			????????????????????????????????????????????????????????????????????????????????????????????????????????????????????????????????????????????????????????????????????????????????????????????????????????????????????????????????????????????????????????????????????????????????????????????????????????????????????????????????????????????????????????????????????????????????

			>Alcantarea_aurantiaca
			????????????????????????????????????????????????????????????????????????????????????????????????????????????????????????????????????????????????????????????????????????????????????????????????????????????????????????????????????????????????????????????????????????????????????????????????????????????????????????????????????????????????????????????????????????????????

			>Alcantarea_benzingii
			????????????????????????????????????????????????????????????????????????????????????????????????????????????????????????????????????????????????????????????????????????????????????????????????????????????????????????????????????????????????????????????????????????????????????????????????????????????????????????????????????????????????????????????????????????????????

			>Alcantarea_burle-marxii
			????????????????????????????????????????????????????????????????????????????????????????????????????????????????????????????????????????????????????????????????????????????????????????????????????????????????????????????????????????????????????????????????????????????????????????????????????????????????????????????????????????????????????????????????????????????????

			>Alcantarea_compacta
			????????????????????????????????????????????????????????????????????????????????????????????????????????????????????????????????????????????????????????????????????????????????????????????????????????????????????????????????????????????????????????????????????????????????????????????????????????????????????????????????????????????????????????????????????????????????

			>Alcantarea_duarteana
			????????????????????????????????????????????????????????????????????????????????????????????????????????????????????????????????????????????????????????????????????????????????????????????????????????????????????????????????????????????????????????????????????????????????????????????????????????????????????????????????????????????????????????????????????????????????

			>Alcantarea_farneyi
			????????????????????????????????????????????????????????????????????????????????????????????????????????????????????????????????????????????????????????????????????????????????????????????????????????????????????????????????????????????????????????????????????????????????????????????????????????????????????????????????????????????????????????????????????????????????

			>Alcantarea_geniculata
			????????????????????????????????????????????????????????????????????????????????????????????????????????????????????????????????????????????????????????????????????????????????????????????????????????????????????????????????????????????????????????????????????????????????????????????????????????????????????????????????????????????????????????????????????????????????

			>Alcantarea_glaziouana
			????????????????????????????????????????????????????????????????????????????????????????????????????????????????????????????????????????????????????????????????????????????????????????????????????????????????????????????????????????????????????????????????????????????????????????????????????????????????????????????????????????????????????????????????????????????????

			>Alcantarea_heloisae
			????????????????????????????????????????????????????????????????????????????????????????????????????????????????????????????????????????????????????????????????????????????????????????????????????????????????????????????????????????????????????????????????????????????????????????????????????????????????????????????????????????????????????????????????????????????????

			>Alcantarea_imperialis
			????????????????????????????????????????????????????????????????????????????????????????????????????????????????????????????????????????????????????????????????????????????????????????????????????????????????????????????????????????????????????????????????????????????????????????????????????????????????????????????????????????????????????????????????????????????????

			>Alcantarea_martinellii
			????????????????????????????????????????????????????????????????????????????????????????????????????????????????????????????????????????????????????????????????????????????????????????????????????????????????????????????????????????????????????????????????????????????????????????????????????????????????????????????????????????????????????????????????????????????????

			>Alcantarea_nahoumii
			????????????????????????????????????????????????????????????????????????????????????????????????????????????????????????????????????????????????????????????????????????????????????????????????????????????????????????????????????????????????????????????????????????????????????????????????????????????????????????????????????????????????????????????????????????????????

			>Alcantarea_nevaresii
			????????????????????????????????????????????????????????????????????????????????????????????????????????????????????????????????????????????????????????????????????????????????????????????????????????????????????????????????????????????????????????????????????????????????????????????????????????????????????????????????????????????????????????????????????????????????

			>Alcantarea_odorata
			????????????????????????????????????????????????????????????????????????????????????????????????????????????????????????????????????????????????????????????????????????????????????????????????????????????????????????????????????????????????????????????????????????????????????????????????????????????????????????????????????????????????????????????????????????????????

			>Alcantarea_recurvifolia
			????????????????????????????????????????????????????????????????????????????????????????????????????????????????????????????????????????????????????????????????????????????????????????????????????????????????????????????????????????????????????????????????????????????????????????????????????????????????????????????????????????????????????????????????????????????????

			>Alcantarea_regina
			????????????????????????????????????????????????????????????????????????????????????????????????????????????????????????????????????????????????????????????????????????????????????????????????????????????????????????????????????????????????????????????????????????????????????????????????????????????????????????????????????????????????????????????????????????????????

			>Alcantarea_roberto-kautskyi
			????????????????????????????????????????????????????????????????????????????????????????????????????????????????????????????????????????????????????????????????????????????????????????????????????????????????????????????????????????????????????????????????????????????????????????????????????????????????????????????????????????????????????????????????????????????????

			>Alcantarea_vasconcelosiana
			????????????????????????????????????????????????????????????????????????????????????????????????????????????????????????????????????????????????????????????????????????????????????????????????????????????????????????????????????????????????????????????????????????????????????????????????????????????????????????????????????????????????????????????????????????????????

			>Ananas_ananassoides
			TCGCAGAAAGCTCTTTCGATGCCTACTGGAATGGGCATTCTTTGCGCRAGCCCGAAAGCTCTAGAAGCTTCCAAAACAGCCAAGTCGGTCAGGGTTTTCTTCGACTGGAATGACTACTTGAAGTTCTACAAGCTCGGAACTTACTGGCCTTAACGCCTTCCATCCAACTACTTTACGGGCTAAGAGCAGCTTTGGATCTCATCTTTGAGGAGGGGCTCGACAATGTCATCGAAAGGCATCGTCGCTTAGGAAAGGCCACAAGGTTAAGCAAAAGCTATACATCT-AATAGCCGCTTTCGCCGTGTTGGATAGATATATTACTTGGCACCATTTTCCTCTTGTAACAWTGAGCAGACTTGCTGTCGAAG

			>Ananas_ananassoides_var._nanus
			????????????????????????????????????????????????????????????????????????????????????????????????????????????????????????????????????????????????????????????????????????????????????????????????????????????????????????????????????????????????????????????????????????????????????????????????????????????????????????????????????????????????????????????????????????????????

			>Ananas_bracteatus
			TCGCAGAAAGCTCTTTCGATGCCTACTGGAATGGGCATTCTTTGCGCGAGCCCGAAAGCTCTAGAAGCTTCCAAAACAGCCAAGTCGGTCAGRGTTTTCTTCGACTGGAATGACTACTTGAAGTTCTACAAGCTCGGAACTTACTGGCCTTAACGCCTTCCATCCAACTACTTTACGGGCTAAGAGCAGCTTTGGATCTCATCTTTGAGGAGGGGCTCGACAATGTCATCGAAAGGCATCGTCGCTTAGGAAAGGCCACAAGGTTAAGYAAAAGCTATACATCT-AATAGCMGCTTTCGCCGTGTTGGATAGATATATTACTTGGCACCATTTTCCTCTTGTAACAATGAGCAGACTTGCTGTCGAAG

			>Ananas_comosus
			TCGCAGAAAGCTCTTTCGATGCCTACTGGAATGGGCATTCTTTGCGCGAGCCCGAAAGCTCTAGAAGCTTCCAAAACAGCCAAGTCGGTCAGGGTTTTCTTCGACTGGAATGACTACTTGAAGTTCTACAAGCTCGGAACTTACTGGCCTTAACGCCTTCCATCCAACTACTTTACGGGCTAAGAGCAGCTTTGGATCTCATCTTTGAGGAGGGGCTCGACAATGTCATCGAAAGGCATCGTCGCTTAGGAAAGGCCACAAGGTTAAGCAAAAGCTATACATCT-AATAGCCGCTTTCGCCGTGTTGGATAGATATATTACTTGGSACCATTTTCCTCTTGYAACAATGAGCAGACTTGCTGTCGAAG

			>Ananas_comosus_var._microstachys
			TCGCAGAAAGCTCTTTCGATGCCTACTGGAATGGGCATTCTTTGCGCRAGCCCGAAAGCTCTAGAAGCTTCCAAAACAGCCAAGTCGGTCAGGGTTTTCTTCGACTGGAATGACTACTTGAAGTTCTACAAGCTCGGAACTTACTGGCCTTAACGCCTTCCATCCAACTACTTTACGGGCTAAGAGCAGCTTTGGATCTCATCTTTGAGGAGGGGCTCGACAATGTCATCGAAAGGCATCGTCGCTTAGGAAAGGCCACAAGGTTAAGCAAAAGCTATACATCT-AATAGCCGCTTTCGCCGTGTTGGATAGATATATTACTTGGCACCATTTTCCTCTTGTAACAWTGAGCAGACTTGCTGTCGAAG

			>Ananas_comosus_var._parguazensis
			TCGCAGAAAGCTCTTTCGATGCCTACTGGAATGGGCATTCTTTGCGCGAGCCCGAAAGCTCTAGAAGCTTCCAAAACAGCCAAGTCGGTCAGGGTTTTCTTCGACTGGAATGACTACTTGAAGTTCTACAAGCTCGGAACTTACTGGCCTTAACGCCTTCCATCCAACTACTTTACGGGCTAAGAGCAGCTTTGGATCTCATCTTTGAGGAGGGGCTCGACAATGTCATCGAAAGGCATCGTCGCTTAGGAAAGGCCACAAGGTTAAGCAAAAGCTATACATCT-AATAGCCGCTTTCGCCGTGTTGGATAGRTATATTACTTGGSACCATTTTCCTCTTGTAACAATGAGCAGACTTGCTGTCGAAG

			>Ananas_fritzmuelleri
			TCGCAGAAAGCTCTTTCGATGCCTACTGGAATGGGCATTCTTTGCGCGAGCCCGAAAGCTCTAGAAGCTTCCAAAACAGCCAAGTCGGTCAGGGTTTTCTTCGACTGGAATGACTACTTGAAGTTCTACAAGCTCGGAACTTACTGGCCTTAACGCCTTCCATCCAACTACTTTACGGGCTAAGAGCAGCTTTGGATCTCATCTTTGAGGAGGGGCTCGACAATGTCATCGAAAGGCATCGTCGCTTAGGAAAGGCCACAAGGTTAAGCAAAAGCTATACATCT-AATAGCCGCTTTCGCCGTGTTGGATAGATATATTACTTGGCACCATTTTCCTCTTGTAACAATGAGCAGACTTGCTGTCGAAG

			>Ananas_sagenaria
			????????????????????????????????????????????????????????????????????????????????????????????????????????????????????????????????????????????????????????????????????????????????????????????????????????????????????????????????????????????????????????????????????????????????????????????????????????????????????????????????????????????????????????????????????????????????

			>Androlepis_fragrans
			TCGCAGAAAGCTCTTTCCCTGCCTACTGGAATGGGCATTCTCTGCGCGAGCCCGAAAGCTCTAGAAGCTTCCAAAACAGCCAAGTCGGTCAGGGTTTTCTTCGACTGGAATGACTACTTGAAGTTCTACAAGCTCGGAACTTACTGGCCTTAACGCCTTCCATCCAACTACTTTACGGGCTGAGAGCAGCTTTGGATCTCATCTTTGAGGAGGGGCTCGACAATGTCATCGAAAGGCATCGTCGCTTAGGAAAGGCCACAAGGTTACGTAAAAGCTATACATCT-AATAGCCGCTTTCGCCGCGTTGGATAGATATTTTACTTTGCGCTATTGTCCTCTTGTAACAATGAGCAGACTTGCTGTCGAAG

			>Androlepis_skinneri
			????????????????????????????????????????????????????????????????????????????????????????????????????????????????????????????????????????????????????????????????????????????????????????????????????????????????????????????????????????????????????????????????????????????????????????????????????????????????????????????????????????????????????????????????????????????????

			>Araeococcus_chlorocarpus
			????????????????????????????????????????????????????????????????????????????????????????????????????????????????????????????????????????????????????????????????????????????????????????????????????????????????????????????????????????????????????????????????????????????????????????????????????????????????????????????????????????????????????????????????????????????????

			>Araeococcus_flagellifolius
			????????????????????????????????????????????????????????????????????????????????????????????????????????????????????????????????????????????????????????????????????????????????????????????????????????????????????????????????????????????????????????????????????????????????????????????????????????????????????????????????????????????????????????????????????????????????

			>Araeococcus_goeldianus
			????????????????????????????????????????????????????????????????????????????????????????????????????????????????????????????????????????????????????????????????????????????????????????????????????????????????????????????????????????????????????????????????????????????????????????????????????????????????????????????????????????????????????????????????????????????????

			>Araeococcus_micranthus
			????????????????????????????????????????????????????????????????????????????????????????????????????????????????????????????????????????????????????????????????????????????????????????????????????????????????????????????????????????????????????????????????????????????????????????????????????????????????????????????????????????????????????????????????????????????????

			>Araeococcus_pectinatus
			????????????????????????????????????????????????????????????????????????????????????????????????????????????????????????????????????????????????????????????????????????????????????????????????????????????????????????????????????????????????????????????????????????????????????????????????????????????????????????????????????????????????????????????????????????????????

			>Barfussia_laxissima
			????????????????????????????????????????????????????????????????????????????????????????????????????????????????????????????????????????????????????????????????????????????????????????????????????????????????????????????????????????????????????????????????????????????????????????????????????????????????????????????????????????????????????????????????????????????????

			>Barfussia_platyrhachis
			????????????????????????????????????????????????????????????????????????????????????????????????????????????????????????????????????????????????????????????????????????????????????????????????????????????????????????????????????????????????????????????????????????????????????????????????????????????????????????????????????????????????????????????????????????????????

			>Barfussia_wagneriana
			????????????????????????????????????????????????????????????????????????????????????????????????????????????????????????????????????????????????????????????????????????????????????????????????????????????????????????????????????????????????????????????????????????????????????????????????????????????????????????????????????????????????????????????????????????????????

			>Billbergia_amoena
			????????????????????????????????????????????????????????????????????????????????????????????????????????????????????????????????????????????????????????????????????????????????????????????????????????????????????????????????????????????????????????????????????????????????????????????????????????????????????????????????????????????????????????????????????????????????

			>Billbergia_brachysiphon
			????????????????????????????????????????????????????????????????????????????????????????????????????????????????????????????????????????????????????????????????????????????????????????????????????????????????????????????????????????????????????????????????????????????????????????????????????????????????????????????????????????????????????????????????????????????????

			>Billbergia_brasiliensis
			TCGCAGAAAGCTCTTTCCCTGCCTACTGGAATGGGCATTCTCTGCGCGAGCCCGAAAGCTCTAGAAGCTTCCAAAACAGCCAAGTCTGTCAGGGTTTTCTTCGACTGGAATGACTACTTGAAGTTCTACAAGCTCGGAACTTACTGGCCTTAACCCCGTCCATCCAACTGCTTTACGGGCTGAGAGCAGCTTTGGATCTCATCTTTGAGGAGGGGCTCGACAACGTCATCGAAAGGCATCGTCGTTTAGGAAAGGCCACAAGGTTAAGTAAAAGCTATACATCT-ATTAGCCGCCTTCGCCGTGTTGGATAGATATATTACTTGGTACTGTTGTTCTCTTGTAACAATGAGCAGGCTTGCCGTCGAAG

			>Billbergia_chlorantha
			????????????????????????????????????????????????????????????????????????????????????????????????????????????????????????????????????????????????????????????????????????????????????????????????????????????????????????????????????????????????????????????????????????????????????????????????????????????????????????????????????????????????????????????????????????????????

			>Billbergia_decora
			????????????????????????????????????????????????????????????????????????????????????????????????????????????????????????????????????????????????????????????????????????????????????????????????????????????????????????????????????????????????????????????????????????????????????????????????????????????????????????????????????????????????????????????????????????????????

			>Billbergia_distachia
			????????????????????????????????????????????????????????????????????????????????????????????????????????????????????????????????????????????????????????????????????????????????????????????????????????????????????????????????????????????????????????????????????????????????????????????????????????????????????????????????????????????????????????????????????????????????

			>Billbergia_eloiseae
			????????????????????????????????????????????????????????????????????????????????????????????????????????????????????????????????????????????????????????????????????????????????????????????????????????????????????????????????????????????????????????????????????????????????????????????????????????????????????????????????????????????????????????????????????????????????

			>Billbergia_euphemiae
			????????????????????????????????????????????????????????????????????????????????????????????????????????????????????????????????????????????????????????????????????????????????????????????????????????????????????????????????????????????????????????????????????????????????????????????????????????????????????????????????????????????????????????????????????????????????

			>Billbergia_horrida
			????????????????????????????????????????????????????????????????????????????????????????????????????????????????????????????????????????????????????????????????????????????????????????????????????????????????????????????????????????????????????????????????????????????????????????????????????????????????????????????????????????????????????????????????????????????????

			>Billbergia_kautskyana
			????????????????????????????????????????????????????????????????????????????????????????????????????????????????????????????????????????????????????????????????????????????????????????????????????????????????????????????????????????????????????????????????????????????????????????????????????????????????????????????????????????????????????????????????????????????????

			>Billbergia_leptopoda
			????????????????????????????????????????????????????????????????????????????????????????????????????????????????????????????????????????????????????????????????????????????????????????????????????????????????????????????????????????????????????????????????????????????????????????????????????????????????????????????????????????????????????????????????????????????????

			>Billbergia_magnifica
			????????????????????????????????????????????????????????????????????????????????????????????????????????????????????????????????????????????????????????????????????????????????????????????????????????????????????????????????????????????????????????????????????????????????????????????????????????????????????????????????????????????????????????????????????????????????

			>Billbergia_manarae
			????????????????????????????????????????????????????????????????????????????????????????????????????????????????????????????????????????????????????????????????????????????????????????????????????????????????????????????????????????????????????????????????????????????????????????????????????????????????????????????????????????????????????????????????????????????????

			>Billbergia_meyeri
			????????????????????????????????????????????????????????????????????????????????????????????????????????????????????????????????????????????????????????????????????????????????????????????????????????????????????????????????????????????????????????????????????????????????????????????????????????????????????????????????????????????????????????????????????????????????

			>Billbergia_morelii
			????????????????????????????????????????????????????????????????????????????????????????????????????????????????????????????????????????????????????????????????????????????????????????????????????????????????????????????????????????????????????????????????????????????????????????????????????????????????????????????????????????????????????????????????????????????????

			>Billbergia_nana
			????????????????????????????????????????????????????????????????????????????????????????????????????????????????????????????????????????????????????????????????????????????????????????????????????????????????????????????????????????????????????????????????????????????????????????????????????????????????????????????????????????????????????????????????????????????????

			>Billbergia_nutans
			????????????????????????????????????????????????????????????????????????????????????????????????????????????????????????????????????????????????????????????????????????????????????????????????????????????????????????????????????????????????????????????????????????????????????????????????????????????????????????????????????????????????????????????????????????????????

			>Billbergia_pallidiflora
			????????????????????????????????????????????????????????????????????????????????????????????????????????????????????????????????????????????????????????????????????????????????????????????????????????????????????????????????????????????????????????????????????????????????????????????????????????????????????????????????????????????????????????????????????????????????

			>Billbergia_porteana
			????????????????????????????????????????????????????????????????????????????????????????????????????????????????????????????????????????????????????????????????????????????????????????????????????????????????????????????????????????????????????????????????????????????????????????????????????????????????????????????????????????????????????????????????????????????????

			>Billbergia_pyramidalis
			????????????????????????????????????????????????????????????????????????????????????????????????????????????????????????????????????????????????????????????????????????????????????????????????????????????????????????????????????????????????????????????????????????????????????????????????????????????????????????????????????????????????????????????????????????????????

			>Billbergia_robert-readii
			????????????????????????????????????????????????????????????????????????????????????????????????????????????????????????????????????????????????????????????????????????????????????????????????????????????????????????????????????????????????????????????????????????????????????????????????????????????????????????????????????????????????????????????????????????????????

			>Billbergia_rosea
			????????????????????????????????????????????????????????????????????????????????????????????????????????????????????????????????????????????????????????????????????????????????????????????????????????????????????????????????????????????????????????????????????????????????????????????????????????????????????????????????????????????????????????????????????????????????

			>Billbergia_speciosa
			????????????????????????????????????????????????????????????????????????????????????????????????????????????????????????????????????????????????????????????????????????????????????????????????????????????????????????????????????????????????????????????????????????????????????????????????????????????????????????????????????????????????????????????????????????????????

			>Billbergia_stenopetala
			????????????????????????????????????????????????????????????????????????????????????????????????????????????????????????????????????????????????????????????????????????????????????????????????????????????????????????????????????????????????????????????????????????????????????????????????????????????????????????????????????????????????????????????????????????????????

			>Billbergia_tweedieana
			????????????????????????????????????????????????????????????????????????????????????????????????????????????????????????????????????????????????????????????????????????????????????????????????????????????????????????????????????????????????????????????????????????????????????????????????????????????????????????????????????????????????????????????????????????????????

			>Billbergia_velascana
			????????????????????????????????????????????????????????????????????????????????????????????????????????????????????????????????????????????????????????????????????????????????????????????????????????????????????????????????????????????????????????????????????????????????????????????????????????????????????????????????????????????????????????????????????????????????

			>Billbergia_violacea
			????????????????????????????????????????????????????????????????????????????????????????????????????????????????????????????????????????????????????????????????????????????????????????????????????????????????????????????????????????????????????????????????????????????????????????????????????????????????????????????????????????????????????????????????????????????????

			>Billbergia_viridiflora
			????????????????????????????????????????????????????????????????????????????????????????????????????????????????????????????????????????????????????????????????????????????????????????????????????????????????????????????????????????????????????????????????????????????????????????????????????????????????????????????????????????????????????????????????????????????????

			>Billbergia_vittata
			????????????????????????????????????????????????????????????????????????????????????????????????????????????????????????????????????????????????????????????????????????????????????????????????????????????????????????????????????????????????????????????????????????????????????????????????????????????????????????????????????????????????????????????????????????????????

			>Billbergia_zebrina
			????????????????????????????????????????????????????????????????????????????????????????????????????????????????????????????????????????????????????????????????????????????????????????????????????????????????????????????????????????????????????????????????????????????????????????????????????????????????????????????????????????????????????????????????????????????????

			>Brewcaria_reflexa
			TCGCAGAAAGCTCTTTCCCTGCCTACTGGAATGGGCATTGTTTGTGCGAGCCCGAAAGCTCTAGAAGCTTCCAAAACGGCCAAGTCGGTCAGGGTTTTCTTCGACTGGAATGACTACTTGAAGTTCTACAAGCTTGGAACTTACTGGCCTTAACGCCTTCCATCCAACTACTTTATGGGCTAAGAGCGGCCTTGGATCTCATCTTTGAGGAGGGGCTCGACAATGTCATCGAAAGGCATCGTCGTTTAGGAAAGGCCACAAGGT-AAGTAAAAGCTATACACCT-AATAGCCGATTTTACAGTGTTGGAAAGATATATTACCTGGCATTATAGTCCTCTCGTAACAATGAGCAGACTTGCTGTCGAAG

			>Brocchinia_acuminata
			????????????????????????????????????????????????????????????????????????????????????????????????????????????????????????????????????????????????????????????????????????????????????????????????????????????????????????????????????????????????????????????????????????????????????????????????????????????????????????????????????????????????????????????????????????????????

			>Brocchinia_micrantha
			????????????????????????????????????????????????????????????????????????????????????????????????????????????????????????????????????????????????????????????????????????????????????????????????????????????????????????????????????????????????????????????????????????????????????????????????????????????????????????????????????????????????????????????????????????????????

			>Brocchinia_paniculata
			????????????????????????????????????????????????????????????????????????????????????????????????????????????????????????????????????????????????????????????????????????????????????????????????????????????????????????????????????????????????????????????????????????????????????????????????????????????????????????????????????????????????????????????????????????????????

			>Brocchinia_prismatica
			????????????????????????????????????????????????????????????????????????????????????????????????????????????????????????????????????????????????????????????????????????????????????????????????????????????????????????????????????????????????????????????????????????????????????????????????????????????????????????????????????????????????????????????????????????????????

			>Brocchinia_reducta
			????????????????????????????????????????????????????????????????????????????????????????????????????????????????????????????????????????????????????????????????????????????????????????????????????????????????????????????????????????????????????????????????????????????????????????????????????????????????????????????????????????????????????????????????????????????????

			>Brocchinia_steyermarkii
			????????????????????????????????????????????????????????????????????????????????????????????????????????????????????????????????????????????????????????????????????????????????????????????????????????????????????????????????????????????????????????????????????????????????????????????????????????????????????????????????????????????????????????????????????????????????

			>Brocchinia_tatei
			????????????????????????????????????????????????????????????????????????????????????????????????????????????????????????????????????????????????????????????????????????????????????????????????????????????????????????????????????????????????????????????????????????????????????????????????????????????????????????????????????????????????????????????????????????????????

			>Brocchinia_uaipanensis
			????????????????????????????????????????????????????????????????????????????????????????????????????????????????????????????????????????????????????????????????????????????????????????????????????????????????????????????????????????????????????????????????????????????????????????????????????????????????????????????????????????????????????????????????????????????????

			>Bromelia_agavifolia
			????????????????????????????????????????????????????????????????????????????????????????????????????????????????????????????????????????????????????????????????????????????????????????????????????????????????????????????????????????????????????????????????????????????????????????????????????????????????????????????????????????????????????????????????????????????????

			>Bromelia_antiacantha
			????????????????????????????????????????????????????????????????????????????????????????????????????????????????????????????????????????????????????????????????????????????????????????????????????????????????????????????????????????????????????????????????????????????????????????????????????????????????????????????????????????????????????????????????????????????????

			>Bromelia_balansae
			????????????????????????????????????????????????????????????????????????????????????????????????????????????????????????????????????????????????????????????????????????????????????????????????????????????????????????????????????????????????????????????????????????????????????????????????????????????????????????????????????????????????????????????????????????????????

			>Bromelia_chrysantha
			????????????????????????????????????????????????????????????????????????????????????????????????????????????????????????????????????????????????????????????????????????????????????????????????????????????????????????????????????????????????????????????????????????????????????????????????????????????????????????????????????????????????????????????????????????????????

			>Bromelia_flemingii
			????????????????????????????????????????????????????????????????????????????????????????????????????????????????????????????????????????????????????????????????????????????????????????????????????????????????????????????????????????????????????????????????????????????????????????????????????????????????????????????????????????????????????????????????????????????????

			>Bromelia_hemisphaerica
			????????????????????????????????????????????????????????????????????????????????????????????????????????????????????????????????????????????????????????????????????????????????????????????????????????????????????????????????????????????????????????????????????????????????????????????????????????????????????????????????????????????????????????????????????????????????

			>Bromelia_irwinii
			????????????????????????????????????????????????????????????????????????????????????????????????????????????????????????????????????????????????????????????????????????????????????????????????????????????????????????????????????????????????????????????????????????????????????????????????????????????????????????????????????????????????????????????????????????????????

			>Bromelia_karatas
			????????????????????????????????????????????????????????????????????????????????????????????????????????????????????????????????????????????????????????????????????????????????????????????????????????????????????????????????????????????????????????????????????????????????????????????????????????????????????????????????????????????????????????????????????????????????

			>Bromelia_pinguin
			????????????????????????????????????????????????????????????????????????????????????????????????????????????????????????????????????????????????????????????????????????????????????????????????????????????????????????????????????????????????????????????????????????????????????????????????????????????????????????????????????????????????????????????????????????????????

			>Bromelia_serra
			????????????????????????????????????????????????????????????????????????????????????????????????????????????????????????????????????????????????????????????????????????????????????????????????????????????????????????????????????????????????????????????????????????????????????????????????????????????????????????????????????????????????????????????????????????????????

			>Bromelia_trianae
			????????????????????????????????????????????????????????????????????????????????????????????????????????????????????????????????????????????????????????????????????????????????????????????????????????????????????????????????????????????????????????????????????????????????????????????????????????????????????????????????????????????????????????????????????????????????

			>Canistropsis_albiflora
			????????????????????????????????????????????????????????????????????????????????????????????????????????????????????????????????????????????????????????????????????????????????????????????????????????????????????????????????????????????????????????????????????????????????????????????????????????????????????????????????????????????????????????????????????????????????

			>Canistropsis_billbergioides
			TCGCAGAAAGCTCTTTCCCTGCCTACTGGAATGGGGATTCTCTGCGCGAGCCCGAAAGCTCTAGAAGCTTCCAAAACAGCCAAGTCGGTCAGGGTTTTCTTCGACTGGAATGACTACTTGAAGTTCTACAAGCTCGGAACTTACTGGCCTTAACGCCTTCCATCCAACTACTTTACGGGCTGAGAGCAGCTTTGGACCTCATCTTTGATGAGGGGCTCGACAATGTCATCGAAAGGCATCGTCGCTTAGGAAAGGCCACAAGGTTAACTAAAAGCTATACATCTAAATAGCCGCTTTCGCCGTGTTGGATAGATATATTACTTTGCACTACTGTCCTCTTGTAACAATGGGCAGACTTGCCGTCGAAG

			>Canistropsis_burchellii
			????????????????????????????????????????????????????????????????????????????????????????????????????????????????????????????????????????????????????????????????????????????????????????????????????????????????????????????????????????????????????????????????????????????????????????????????????????????????????????????????????????????????????????????????????????????????

			>Canistropsis_correia-araujoi
			TCGCAGAAAGCTCTTTCCCTGCCTACTGGAATGGGGATTCTCTGCGCGAGCCCGAAAGCTCTAGAAGCTTCCAAAACAGCCAAGTCGGTCAGGGTTTTCTTCGACTGGAATGACTACTTGAAGTTCTACAAGCTCGGAACTTACTGGCCTTAACGCCTTCCATCCAACTACTTTACGGGCTGAGAGCAGCTTTGGACCTCATCTTTGAGGAGGGGCTCGACAATGTCATCGAAAGGCATCGTCGCTTAGGAAAGGCCACAAGGTTAASTAAAAGCTATACATCTAAATAGCCGCTTTCGCCGTGTTGGATAGATATATTACTTTGCACTRYTGTCCTCTTGTAACAATGGGCAGACTTGCCGTCGAAG

			>Canistropsis_elata
			????????????????????????????????????????????????????????????????????????????????????????????????????????????????????????????????????????????????????????????????????????????????????????????????????????????????????????????????????????????????????????????????????????????????????????????????????????????????????????????????????????????????????????????????????????????????

			>Canistropsis_exigua
			TCGCAGAAAGCTCTTTCCCTGCCTACTGGAATGGGGATTCTCTGCGCGAGCCCGAAAGCTCTAGAAGCTTCCAAAACAGCCAAGTCGGTCAGGGTTTTCTTCGACTGGAATGACTACTTGAAGTTCTACAAGCTCGGAACTTACTGGCCTTAACGCCTTCCATCCAACTACTTTACGGGCTGAGAGCAGCTTTGGACCTCATCTTTGAGGAGGGGCTCGACAATGTCATCGAAAGGCATCGTCGMTTAGGAAAGGCCACAAGGTTAAGTAAAAGCTATACATCTAAATAGCCGCTTTCGCCGTGTTGGATAGATATATTACTTTGCACTACTGTCCTCTTGTAACAATGGGCAGACTTGCCGTCGAAG

			>Canistropsis_marceloi
			TCGCAGAAAGCTCTTTCCCTGCCTACTGGAATGGGGATTCTCTGCGCGAGCCCGAAAGCTCTAGAAGCTTCSAAAACAGCCRAGTCGGTCAGGGTTTTCTTCGACTGGAATGACTACTTGAAGTTCTACAAGCTCGGAACTTACTGGCCTTAACGCCTTCCATCCAACTACTTTACGGGCTGAGAGCAGCTTTGGACCTCATCTTTGAGGAGGGGCTCGACAATGTCRTCGAAAGGCATCGTCGCTTAGGAAAGGCCACAAGGTTAAGTAAAAGCTATACATCTAAATAGCCGCTTTCGCCGTGTTGGATAGATATATTACTCTGCACTACTGTCCTCTTGTAACAATGGGCAGACTTGCCGTCGAAG

			>Canistropsis_microps
			????????????????????????????????????????????????????????????????????????????????????????????????????????????????????????????????????????????????????????????????????????????????????????????????????????????????????????????????????????????????????????????????????????????????????????????????????????????????????????????????????????????????????????????????????????????????

			>Canistropsis_pulcherrima
			????????????????????????????????????????????????????????????????????????????????????????????????????????????????????????????????????????????????????????????????????????????????????????????????????????????????????????????????????????????????????????????????????????????????????????????????????????????????????????????????????????????????????????????????????????????????

			>Canistropsis_simulans
			????????????????????????????????????????????????????????????????????????????????????????????????????????????????????????????????????????????????????????????????????????????????????????????????????????????????????????????????????????????????????????????????????????????????????????????????????????????????????????????????????????????????????????????????????????????????

			>Canistrum_alagoanum
			????????????????????????????????????????????????????????????????????????????????????????????????????????????????????????????????????????????????????????????????????????????????????????????????????????????????????????????????????????????????????????????????????????????????????????????????????????????????????????????????????????????????????????????????????????????????

			>Canistrum_aurantiacum
			????????????????????????????????????????????????????????????????????????????????????????????????????????????????????????????????????????????????????????????????????????????????????????????????????????????????????????????????????????????????????????????????????????????????????????????????????????????????????????????????????????????????????????????????????????????????

			>Canistrum_camacaense
			????????????????????????????????????????????????????????????????????????????????????????????????????????????????????????????????????????????????????????????????????????????????????????????????????????????????????????????????????????????????????????????????????????????????????????????????????????????????????????????????????????????????????????????????????????????????

			>Canistrum_fosterianum
			????????????????????????????????????????????????????????????????????????????????????????????????????????????????????????????????????????????????????????????????????????????????????????????????????????????????????????????????????????????????????????????????????????????????????????????????????????????????????????????????????????????????????????????????????????????????

			>Canistrum_pickelii
			????????????????????????????????????????????????????????????????????????????????????????????????????????????????????????????????????????????????????????????????????????????????????????????????????????????????????????????????????????????????????????????????????????????????????????????????????????????????????????????????????????????????????????????????????????????????

			>Canistrum_triangulare
			????????????????????????????????????????????????????????????????????????????????????????????????????????????????????????????????????????????????????????????????????????????????????????????????????????????????????????????????????????????????????????????????????????????????????????????????????????????????????????????????????????????????????????????????????????????????

			>Catopsis_berteroniana
			????????????????????????????????????????????????????????????????????????????????????????????????????????????????????????????????????????????????????????????????????????????????????????????????????????????????????????????????????????????????????????????????????????????????????????????????????????????????????????????????????????????????????????????????????????????????

			>Catopsis_floribunda
			????????????????????????????????????????????????????????????????????????????????????????????????????????????????????????????????????????????????????????????????????????????????????????????????????????????????????????????????????????????????????????????????????????????????????????????????????????????????????????????????????????????????????????????????????????????????

			>Catopsis_juncifolia
			????????????????????????????????????????????????????????????????????????????????????????????????????????????????????????????????????????????????????????????????????????????????????????????????????????????????????????????????????????????????????????????????????????????????????????????????????????????????????????????????????????????????????????????????????????????????

			>Catopsis_morreniana
			????????????????????????????????????????????????????????????????????????????????????????????????????????????????????????????????????????????????????????????????????????????????????????????????????????????????????????????????????????????????????????????????????????????????????????????????????????????????????????????????????????????????????????????????????????????????

			>Catopsis_nitida
			????????????????????????????????????????????????????????????????????????????????????????????????????????????????????????????????????????????????????????????????????????????????????????????????????????????????????????????????????????????????????????????????????????????????????????????????????????????????????????????????????????????????????????????????????????????????

			>Catopsis_nutans
			????????????????????????????????????????????????????????????????????????????????????????????????????????????????????????????????????????????????????????????????????????????????????????????????????????????????????????????????????????????????????????????????????????????????????????????????????????????????????????????????????????????????????????????????????????????????

			>Catopsis_nutans_var._nutans
			????????????????????????????????????????????????????????????????????????????????????????????????????????????????????????????????????????????????????????????????????????????????????????????????????????????????????????????????????????????????????????????????????????????????????????????????????????????????????????????????????????????????????????????????????????????????

			>Catopsis_paniculata
			????????????????????????????????????????????????????????????????????????????????????????????????????????????????????????????????????????????????????????????????????????????????????????????????????????????????????????????????????????????????????????????????????????????????????????????????????????????????????????????????????????????????????????????????????????????????

			>Catopsis_sessiliflora
			????????????????????????????????????????????????????????????????????????????????????????????????????????????????????????????????????????????????????????????????????????????????????????????????????????????????????????????????????????????????????????????????????????????????????????????????????????????????????????????????????????????????????????????????????????????????

			>Catopsis_subulata
			????????????????????????????????????????????????????????????????????????????????????????????????????????????????????????????????????????????????????????????????????????????????????????????????????????????????????????????????????????????????????????????????????????????????????????????????????????????????????????????????????????????????????????????????????????????????

			>Catopsis_wangerinii
			????????????????????????????????????????????????????????????????????????????????????????????????????????????????????????????????????????????????????????????????????????????????????????????????????????????????????????????????????????????????????????????????????????????????????????????????????????????????????????????????????????????????????????????????????????????????

			>Cipuropsis_amicorum
			????????????????????????????????????????????????????????????????????????????????????????????????????????????????????????????????????????????????????????????????????????????????????????????????????????????????????????????????????????????????????????????????????????????????????????????????????????????????????????????????????????????????????????????????????????????????

			>Connellia_nutans
			????????????????????????????????????????????????????????????????????????????????????????????????????????????????????????????????????????????????????????????????????????????????????????????????????????????????????????????????????????????????????????????????????????????????????????????????????????????????????????????????????????????????????????????????????????????????

			>Cottendorfia_florida
			TCGCAGAAAGCTCTTTCCCTGCCTACCGGAATGGGGATTGTTTGTGCGAGCCCGAAAGCTCTAGAAGCTTCCAAGACAGCCAAGTCGGTTAGGGTTTTCTTCGACTGGAATGACTACTTGAAGTTCTACAAGCTTGGAACTTACTGGCCTTAACGCCTTCCATCCAACTACTTTATGGGCTAAGAGCAGCTTTGGATCTCATCTTCGAGGAGGGGCTCGACAATGTCATCGAGAGGCATCGTCGTTTAGGAAAGGCCACAAGG-TAAGTA---------------CATAGCCGATTTTACTGTGTTGGACAGATATATTACTTGGCATTATAGTCCTCTCATAACAATCAGCAGACTTGCTGTTGAAG

			>Cryptanthus_acaulis
			????????????????????????????????????????????????????????????????????????????????????????????????????????????????????????????????????????????????????????????????????????????????????????????????????????????????????????????????????????????????????????????????????????????????????????????????????????????????????????????????????????????????????????????????????????????????

			>Cryptanthus_bahianus
			TCGCAGAAAGCTCTTTCGATGCCTACTGGAATGGGCATTCTTTGCGCGAGCCCGAAAGCTCTAGAAGCTTCCAAAACAGCCAAGTCGGTCAGGGTTTTCTTCGACTGGAATGACTACTTGAAGTTCTACAAGCTCGGAACTTACTGGCCTTAACGCCTTCCATCCAACTACTTTACGGGCTAAGAGCAGCTCTGGATCTCATCTTTGAGGAGGGGCTCGACAGCGTCATCGAAAGGCATCGTCGCTTAGGAAAGGCCACAAGGTTAAGTAAAAGCTGTACATCT-AATAGCCGCTTGCGCCGTGTTGGATAGATATATTACTTGGCGCTATTGTCCTCTTGTAACAATGAGCAGACTTGCTGTCGAAG

			>Cryptanthus_beuckeri
			????????????????????????????????????????????????????????????????????????????????????????????????????????????????????????????????????????????????????????????????????????????????????????????????????????????????????????????????????????????????????????????????????????????????????????????????????????????????????????????????????????????????????????????????????????????????

			>Cryptanthus_bromelioides
			????????????????????????????????????????????????????????????????????????????????????????????????????????????????????????????????????????????????????????????????????????????????????????????????????????????????????????????????????????????????????????????????????????????????????????????????????????????????????????????????????????????????????????????????????????????????

			>Cryptanthus_burle-marxii
			TCGCAGAAAGCCCTTTCGATGCCTACTGGAATGGGCATTCTTTGCGCGAGCCCGAAAGCTCTAGAAGCTTCCAAAACAGCCAAGTCGGTCAGGGTTTTCTTCGACTGGAATGACTACTTGAAGTTCTACAAGCTCGGAACTTACTGGCCTTAACGCCTTCCATCCAACTACTTTACGGGCTAAGAGCAGCTTTGGATCTCATCTTTGAGGAGGGGCTCGACAATGTCGTCGAAAGGCATCGTCGCTTAGGAAAGGCCACAAGGTTAAGCAAAAGCCATACATCT-AATAGCCGCTTTCGCCGCGTTGGATAGATATATTACTTGGCACCATTTTCCTCTTGTAACAATGAGC????????????????

			>Cryptanthus_colnagoi
			TCGCAGAAAGCCCTTTCGATGCCTACTGGAATGGGCATTCTTTGCGCGAGCCCGAAAGCTCTAGAAGCTTCCAAAACAGCCARGTCGGTCAGGGTTTTCTTCGACTGGAATGACTACTTGAAGTTCTACAAGCTCGGAACTTACTGGCCTTAACGCCTTCCATCCAACTACTTTACGGGCTAAGAGCAGCTTTGGATCTCATCTTTGAGGAGGGGCTCGACAATGTCGTCGAAAGGCATCGTCGCTTAGGAAAGGCCACAAGGTTAAGCAAAAGCCATACATCY-AATAGCCGCTTTCGCCGCGTTGGATAGATATATTACTTGGCACCATTTTCCTCTTGTAACAATGAGCAGACTTGCYGTCGAAG

			>Cryptanthus_diamantinensis
			TCGCAGAAAGCCCTTTCGATGCCTACTGGAATGGGCATTCTTTGCGCGAGCCCGAAAGCTCTAGAAGCTTCCAAAACAGCCAAGTCGGTCAGGGTTTTCTTCGACTGGAATGACTACTTGAAGTTCTACAAGCTCGGAACTTACTGGCCTTAACGCCTTCCATCCAACTACTTTACGGGCTAAGAGCAGCTTTGGATCTCATCTTTGAGGAGGGGCTCGACAATGTCGTCGAAAGGCATCGTCGCTTAGGAAAGGCCACAAGGTTAAGCAAAAGCCATACATCT-AATAGCCGCTTTCGCCGCGTTGGATAGATATATTACTTGGCACCATTTTCCTCTTGTAACAATGAGCAGACTTGCTGTCGAAG

			>Cryptanthus_dianae
			TCGCAGAAAGCCCTTTCGATGCCTACTGGAATGGGCATTCTTTGCGCGAGCCCGAAAGCTCTAGAAGCTTCCAAAACAGCCAAGTCGGTCAGGGTTTTCTTCGACTGGAATGACTACTTGAAGTTCTACAAGCTCGGAACTTACTGGCCTTAACGCCTTCCATCCAACTACTTTACGGGCTAAGAGCAGCTTTGGATCTCATCTTTGAGGAGGGGCTCGACAATGTCGTCGAAAGGCATCGTCGCTTAGGAAAGGCCACAAGGTTAAGCAAAAGCCATACATCT-AATAGCCGCTTTCGCCGCGTTGGATAGATATATTACTTGGCACCATTTTCCTCTTGTAACAATGAGCAGACTTGCTGTCGAAG

			>Cryptanthus_maritimus
			TCGCAGAAAGCCCTTTCGATGCCTACTGGAATGGGCATTCTTTGCGCGAGCCCGAAAGCTCTAGAAGCTTCCAAAACAGCCAAGTCGGTCAGGGTTTTCTTCGACTGGAATGACTACTTGAAGTTCTACAAGCTCGGAACTTACTGGCCTTAACGCCTTCCATCCAACTACTTTACGGGCTAAGAGCAGCTTTGGATCTCATCTTTGAGGAGGGGCTCGACAATGTCGTCGAAAGGCATCGTCGCTTAGGAAAGGCCACAAGGTCAAGCAAAAGCCATACATCT-AATAGCCGCTTTCGCCGCGTTGGATAGATATATTACTTGGCACCATTTTCCTCTTGTAACAATGAGCAGACTTGCTGTCGAAG

			>Cryptanthus_ruthae
			????????????????????????????????????????????????????????????????????????????????????????????????????????????????????????????????????????????????????????????????????????????????????????????????????????????????????????????????????????????????????????????????????????????????????????????????????????????????????????????????????????????????????????????????????????????????

			>Cryptanthus_teretifolius
			????????????????????????????????????????????????????????????????????????????????????????????????????????????????????????????????????????????????????????????????????????????????????????????????????????????????????????????????????????????????????????????????????????????????????????????????????????????????????????????????????????????????????????????????????????????????

			>Cryptanthus_warren-loosei
			????????????????????????????????????????????????????????????????????????????????????????????????????????????????????????????????????????????????????????????????????????????????????????????????????????????????????????????????????????????????????????????????????????????????????????????????????????????????????????????????????????????????????????????????????????????????

			>Cryptanthus_zonatus
			????????????????????????????????????????????????????????????????????????????????????????????????????????????????????????????????????????????????????????????????????????????????????????????????????????????????????????????????????????????????????????????????????????????????????????????????????????????????????????????????????????????????????????????????????????????????

			>Deinacanthon_urbanianum
			????????????????????????????????????????????????????????????????????????????????????????????????????????????????????????????????????????????????????????????????????????????????????????????????????????????????????????????????????????????????????????????????????????????????????????????????????????????????????????????????????????????????????????????????????????????????

			>Deuterocohnia_abstrusa
			TCGCAGAAAGCCCTTTCCCTGCCTACTGGAATGGGCATTTTATGCGCGAGCCCGAAAGCTCTTGAAGCTTCCAAAACAGCCAAGTCGGTCAGGGTTTTCTTCGACTGGAATGACTACTTGAAGTTCTACAAGCTCGGAACTTACTGGCCTTAACGCCTTCCATCCAACTACTGTATGGGCTAAGAGCAGCTTTGGATCTCATCTTCGAGGAGGGGCTCGACAATGTCATCGAAAGGCATCGTCGTTTAGGAAAGGCCACAAGGT-AAGTAAAAGCTATACACCT-AATAGCCGCTTTTACAGCGTCGGATAGATATATTACTTGGTATT---------TTGTAACAATGAGCAGGCTCGCTGTCGAAG

			>Deuterocohnia_brevifolia
			TCGCAGAAAGCCCTTTCCCTGCCTACTGGAATGGGCATTTTATGCGCGAGCCCGAAAGCTCTAGAAGCTTCCAAAACAGCCAAGTCGGTCAGGGTTTTCTTCGACTGGAATGACTACTTGAAGTTCTACAAGCTCGGAACTTACTGGCCTTAACGCCTTCCATCCAACTACTGTACGGGCTAAGAGCAGCTTTGGATCTCATCTTCGAGGAGGGGCTCGACAATGTCATCGAAAGGCATCGTCGTTTAGGAAAGGCCACAAGGT-AAGTAAAAGCTATACACCT-AATAGCCGCTTTTACAGTGTCGGATAGATATATTACTTGGTATT---------TTGTAACAATGAGCAGGCTCGCTGTCGAAA

			>Deuterocohnia_brevispicata
			TCGCAGAAAGCCCTTTCCCTGCCTACTGGAATGGGCATTTTATGCGCGAGCCCGAAAGCTCTAGAAGCTTCCAAAACAGCCAAGTCGGTCAGGGTTTTCTTCGACTGGAATGACTACTTGAAGTTCTACAAGCTCGGAACTTACTGGCCTTAACACCTTCCATCCAACTACTGTATGGGCTAAGAGCAGCTTTGGATCTCATCTTCGAGGAGGGGCTCGACAATGTCATCGAAAGGCATCGTCGTTTAGGAAAGGCCACAAGGT-AAGTAAAAGCTATACACCT-AATAGCCGCTTTTACAGTGTCGGATAGATATATTACTTGGTATT---------TTGTAAC-----GCAGGCTCGCTGTCGAAG

			>Deuterocohnia_chrysantha
			????????????????????????????????????????????????????????????????????????????????????????????????????????????????????????????????????????????????????????????????????????????????????????????????????????????????????????????????????????????????????????????????????????????????????????????????????????????????????????????????????????????????????????????????????????????????

			>Deuterocohnia_digitata
			??????AAAGCCCTTTCCCTGCCTACTGGAATGGGCATTTTATGCGCGAGCCCGAAAGCTCTAGAAGCTTCCAAAACAGCCAAGTCGGTCAGGGTTTTCTTCGACTGGAATGACTACTTGAAGTTCTACAAGCTCGGAACTTACTGGCCTTAACGCCTTCCATCCAACTACTGTACGGGCTAAGAGCAGCTTTGGATCTCATCTTCGAGGAGGGGCTCGACAATGTCATCGAAAGGCATCGTCGTTTAGGAAAGGCCACAAGGTAAAGTAAAAGCTGTACACCT-AATAGCCGCTTTTACAGCGTCGGATAGATATATTACTTGGTATT---------TTGTAACAATGAGCAGGCTCGCTGTCGAAG

			>Deuterocohnia_gableana
			????????????????????????????????????????????????????????????????????????????????????????????????????????????????????????????????????????????????????????????????????????????????????????????????????????????????????????????????????????????????????????????????????????????????????????????????????????????????????????????????????????????????????????????????????????????????

			>Deuterocohnia_glandulosa
			????????????????????????????????????????????????????????????????????????????????????????????????????????????????????????????????????????????????????????????????????????????????????????????????????????????????????????????????????????????????????????????????????????????????????????????????????????????????????????????????????????????????????????????????????????????????

			>Deuterocohnia_haumanii
			????????????????????????????????????????????????????????????????????????????????????????????????????????????????????????????????????????????????????????????????????????????????????????????????????????????????????????????????????????????????????????????????????????????????????????????????????????????????????????????????????????????????????????????????????????????????

			>Deuterocohnia_longipetala
			????????????????????????????????????????????????????????????????????????????????????????????????????????????????????????????????????????????????????????????????????????????????????????????????????????????????????????????????????????????????????????????????????????????????????????????????????????????????????????????????????????????????????????????????????????????????

			>Deuterocohnia_lotteae
			????????????????????????????????????????????????????????????????????????????????????????????????????????????????????????????????????????????????????????????????????????????????????????????????????????????????????????????????????????????????????????????????????????????????????????????????????????????????????????????????????????????????????????????????????????????????

			>Deuterocohnia_meziana
			????????????????????????????????????????????????????????????????????????????????????????????????????????????????????????????????????????????????????????????????????????????????????????????????????????????????????????????????????????????????????????????????????????????????????????????????????????????????????????????????????????????????????????????????????????????????

			>Deuterocohnia_meziana_subsp._carmineoviridiflora
			????????????????????????????????????????????????????????????????????????????????????????????????????????????????????????????????????????????????????????????????????????????????????????????????????????????????????????????????????????????????????????????????????????????????????????????????????????????????????????????????????????????????????????????????????????????????

			>Deuterocohnia_meziana_subsp._pedicellata
			????????????????????????????????????????????????????????????????????????????????????????????????????????????????????????????????????????????????????????????????????????????????????????????????????????????????????????????????????????????????????????????????????????????????????????????????????????????????????????????????????????????????????????????????????????????????

			>Deuterocohnia_recurvipetala
			????????????????????????????????????????????????????????????????????????????????????????????????????????????????????????????????????????????????????????????????????????????????????????????????????????????????????????????????????????????????????????????????????????????????????????????????????????????????????????????????????????????????????????????????????????????????

			>Deuterocohnia_scapigera
			????????????????????????????????????????????????????????????????????????????????????????????????????????????????????????????????????????????????????????????????????????????????????????????????????????????????????????????????????????????????????????????????????????????????????????????????????????????????????????????????????????????????????????????????????????????????

			>Deuterocohnia_schreiteri
			????????????????????????????????????????????????????????????????????????????????????????????????????????????????????????????????????????????????????????????????????????????????????????????????????????????????????????????????????????????????????????????????????????????????????????????????????????????????????????????????????????????????????????????????????????????????

			>Deuterocohnia_seramisiana
			????????????????????????????????????????????????????????????????????????????????????????????????????????????????????????????????????????????????????????????????????????????????????????????????????????????????????????????????????????????????????????????????????????????????????????????????????????????????????????????????????????????????????????????????????????????????

			>Deuterocohnia_strobilifera_var._inermis
			????????????????????????????????????????????????????????????????????????????????????????????????????????????????????????????????????????????????????????????????????????????????????????????????????????????????????????????????????????????????????????????????????????????????????????????????????????????????????????????????????????????????????????????????????????????????

			>Deuterocohnia_strobilifera_var._strobilifera
			????????????????????????????????????????????????????????????????????????????????????????????????????????????????????????????????????????????????????????????????????????????????????????????????????????????????????????????????????????????????????????????????????????????????????????????????????????????????????????????????????????????????????????????????????????????????

			>Disteganthus_basilateralis
			????????????????????????????????????????????????????????????????????????????????????????????????????????????????????????????????????????????????????????????????????????????????????????????????????????????????????????????????????????????????????????????????????????????????????????????????????????????????????????????????????????????????????????????????????????????????

			>Disteganthus_lateralis
			????????????????????????????????????????????????????????????????????????????????????????????????????????????????????????????????????????????????????????????????????????????????????????????????????????????????????????????????????????????????????????????????????????????????????????????????????????????????????????????????????????????????????????????????????????????????

			>Dyckia_acutiflora
			????????????????????????????????????????????????????????????????????????????????????????????????????????????????????????????????????????????????????????????????????????????????????????????????????????????????????????????????????????????????????????????????????????????????????????????????????????????????????????????????????????????????????????????????????????????????

			>Dyckia_areniticola
			????????????????????????????????????????????????????????????????????????????????????????????????????????????????????????????????????????????????????????????????????????????????????????????????????????????????????????????????????????????????????????????????????????????????????????????????????????????????????????????????????????????????????????????????????????????????

			>Dyckia_aurea
			????????????????????????????????????????????????????????????????????????????????????????????????????????????????????????????????????????????????????????????????????????????????????????????????????????????????????????????????????????????????????????????????????????????????????????????????????????????????????????????????????????????????????????????????????????????????

			>Dyckia_beateae
			????????????????????????????????????????????????????????????????????????????????????????????????????????????????????????????????????????????????????????????????????????????????????????????????????????????????????????????????????????????????????????????????????????????????????????????????????????????????????????????????????????????????????????????????????????????????

			>Dyckia_brachyphylla
			????????????????????????????????????????????????????????????????????????????????????????????????????????????????????????????????????????????????????????????????????????????????????????????????????????????????????????????????????????????????????????????????????????????????????????????????????????????????????????????????????????????????????????????????????????????????

			>Dyckia_bracteata
			????????????????????????????????????????????????????????????????????????????????????????????????????????????????????????????????????????????????????????????????????????????????????????????????????????????????????????????????????????????????????????????????????????????????????????????????????????????????????????????????????????????????????????????????????????????????

			>Dyckia_braunii
			????????????????????????????????????????????????????????????????????????????????????????????????????????????????????????????????????????????????????????????????????????????????????????????????????????????????????????????????????????????????????????????????????????????????????????????????????????????????????????????????????????????????????????????????????????????????

			>Dyckia_brevifolia
			????????????????????????????????????????????????????????????????????????????????????????????????????????????????????????????????????????????????????????????????????????????????????????????????????????????????????????????????????????????????????????????????????????????????????????????????????????????????????????????????????????????????????????????????????????????????

			>Dyckia_choristaminea
			????????????????????????????????????????????????????????????????????????????????????????????????????????????????????????????????????????????????????????????????????????????????????????????????????????????????????????????????????????????????????????????????????????????????????????????????????????????????????????????????????????????????????????????????????????????????

			>Dyckia_cinerea
			????????????????????????????????????????????????????????????????????????????????????????????????????????????????????????????????????????????????????????????????????????????????????????????????????????????????????????????????????????????????????????????????????????????????????????????????????????????????????????????????????????????????????????????????????????????????

			>Dyckia_dawsonii
			????????????????????????????????????????????????????????????????????????????????????????????????????????????????????????????????????????????????????????????????????????????????????????????????????????????????????????????????????????????????????????????????????????????????????????????????????????????????????????????????????????????????????????????????????????????????

			>Dyckia_delicata
			????????????????????????????????????????????????????????????????????????????????????????????????????????????????????????????????????????????????????????????????????????????????????????????????????????????????????????????????????????????????????????????????????????????????????????????????????????????????????????????????????????????????????????????????????????????????

			>Dyckia_densiflora
			????????????????????????????????????????????????????????????????????????????????????????????????????????????????????????????????????????????????????????????????????????????????????????????????????????????????????????????????????????????????????????????????????????????????????????????????????????????????????????????????????????????????????????????????????????????????

			>Dyckia_dissitiflora
			????????????????????????????????????????????????????????????????????????????????????????????????????????????????????????????????????????????????????????????????????????????????????????????????????????????????????????????????????????????????????????????????????????????????????????????????????????????????????????????????????????????????????????????????????????????????

			>Dyckia_distachya
			????????????????????????????????????????????????????????????????????????????????????????????????????????????????????????????????????????????????????????????????????????????????????????????????????????????????????????????????????????????????????????????????????????????????????????????????????????????????????????????????????????????????????????????????????????????????

			>Dyckia_duckei
			????????????????????????????????????????????????????????????????????????????????????????????????????????????????????????????????????????????????????????????????????????????????????????????????????????????????????????????????????????????????????????????????????????????????????????????????????????????????????????????????????????????????????????????????????????????????

			>Dyckia_dusenii
			????????????????????????????????????????????????????????????????????????????????????????????????????????????????????????????????????????????????????????????????????????????????????????????????????????????????????????????????????????????????????????????????????????????????????????????????????????????????????????????????????????????????????????????????????????????????

			>Dyckia_elisabethae
			????????????????????????????????????????????????????????????????????????????????????????????????????????????????????????????????????????????????????????????????????????????????????????????????????????????????????????????????????????????????????????????????????????????????????????????????????????????????????????????????????????????????????????????????????????????????

			>Dyckia_encholirioides
			????????????????????????????????????????????????????????????????????????????????????????????????????????????????????????????????????????????????????????????????????????????????????????????????????????????????????????????????????????????????????????????????????????????????????????????????????????????????????????????????????????????????????????????????????????????????

			>Dyckia_espiritosantensis
			????????????????????????????????????????????????????????????????????????????????????????????????????????????????????????????????????????????????????????????????????????????????????????????????????????????????????????????????????????????????????????????????????????????????????????????????????????????????????????????????????????????????????????????????????????????????

			>Dyckia_estevesii
			????????????????????????????????????????????????????????????????????????????????????????????????????????????????????????????????????????????????????????????????????????????????????????????????????????????????????????????????????????????????????????????????????????????????????????????????????????????????????????????????????????????????????????????????????????????????

			>Dyckia_estevesii_var._braunii
			????????????????????????????????????????????????????????????????????????????????????????????????????????????????????????????????????????????????????????????????????????????????????????????????????????????????????????????????????????????????????????????????????????????????????????????????????????????????????????????????????????????????????????????????????????????????

			>Dyckia_excelsa
			????????????????????????????????????????????????????????????????????????????????????????????????????????????????????????????????????????????????????????????????????????????????????????????????????????????????????????????????????????????????????????????????????????????????????????????????????????????????????????????????????????????????????????????????????????????????

			>Dyckia_ferox
			????????????????????????????????????????????????????????????????????????????????????????????????????????????????????????????????????????????????????????????????????????????????????????????????????????????????????????????????????????????????????????????????????????????????????????????????????????????????????????????????????????????????????????????????????????????????

			>Dyckia_ferruginea
			????????????????????????????????????????????????????????????????????????????????????????????????????????????????????????????????????????????????????????????????????????????????????????????????????????????????????????????????????????????????????????????????????????????????????????????????????????????????????????????????????????????????????????????????????????????????

			>Dyckia_floribunda
			????????????????????????????????????????????????????????????????????????????????????????????????????????????????????????????????????????????????????????????????????????????????????????????????????????????????????????????????????????????????????????????????????????????????????????????????????????????????????????????????????????????????????????????????????????????????

			>Dyckia_fosteriana
			????????????????????????????????????????????????????????????????????????????????????????????????????????????????????????????????????????????????????????????????????????????????????????????????????????????????????????????????????????????????????????????????????????????????????????????????????????????????????????????????????????????????????????????????????????????????

			>Dyckia_goehringii
			????????????????????????????????????????????????????????????????????????????????????????????????????????????????????????????????????????????????????????????????????????????????????????????????????????????????????????????????????????????????????????????????????????????????????????????????????????????????????????????????????????????????????????????????????????????????

			>Dyckia_gouveiana
			????????????????????????????????????????????????????????????????????????????????????????????????????????????????????????????????????????????????????????????????????????????????????????????????????????????????????????????????????????????????????????????????????????????????????????????????????????????????????????????????????????????????????????????????????????????????

			>Dyckia_grandidentata
			????????????????????????????????????????????????????????????????????????????????????????????????????????????????????????????????????????????????????????????????????????????????????????????????????????????????????????????????????????????????????????????????????????????????????????????????????????????????????????????????????????????????????????????????????????????????

			>Dyckia_granmogulensis
			????????????????????????????????????????????????????????????????????????????????????????????????????????????????????????????????????????????????????????????????????????????????????????????????????????????????????????????????????????????????????????????????????????????????????????????????????????????????????????????????????????????????????????????????????????????????

			>Dyckia_hebdingii
			????????????????????????????????????????????????????????????????????????????????????????????????????????????????????????????????????????????????????????????????????????????????????????????????????????????????????????????????????????????????????????????????????????????????????????????????????????????????????????????????????????????????????????????????????????????????

			>Dyckia_hohenbergioides
			????????????????????????????????????????????????????????????????????????????????????????????????????????????????????????????????????????????????????????????????????????????????????????????????????????????????????????????????????????????????????????????????????????????????????????????????????????????????????????????????????????????????????????????????????????????????

			>Dyckia_ibiramensis
			????????????????????????????????????????????????????????????????????????????????????????????????????????????????????????????????????????????????????????????????????????????????????????????????????????????????????????????????????????????????????????????????????????????????????????????????????????????????????????????????????????????????????????????????????????????????

			>Dyckia_jonesiana
			????????????????????????????????????????????????????????????????????????????????????????????????????????????????????????????????????????????????????????????????????????????????????????????????????????????????????????????????????????????????????????????????????????????????????????????????????????????????????????????????????????????????????????????????????????????????

			>Dyckia_kranziana
			????????????????????????????????????????????????????????????????????????????????????????????????????????????????????????????????????????????????????????????????????????????????????????????????????????????????????????????????????????????????????????????????????????????????????????????????????????????????????????????????????????????????????????????????????????????????

			>Dyckia_leptostachya
			????????????????????????????????????????????????????????????????????????????????????????????????????????????????????????????????????????????????????????????????????????????????????????????????????????????????????????????????????????????????????????????????????????????????????????????????????????????????????????????????????????????????????????????????????????????????

			>Dyckia_lindevaldae
			????????????????????????????????????????????????????????????????????????????????????????????????????????????????????????????????????????????????????????????????????????????????????????????????????????????????????????????????????????????????????????????????????????????????????????????????????????????????????????????????????????????????????????????????????????????????

			>Dyckia_lunaris
			????????????????????????????????????????????????????????????????????????????????????????????????????????????????????????????????????????????????????????????????????????????????????????????????????????????????????????????????????????????????????????????????????????????????????????????????????????????????????????????????????????????????????????????????????????????????

			>Dyckia_macedoi
			????????????????????????????????????????????????????????????????????????????????????????????????????????????????????????????????????????????????????????????????????????????????????????????????????????????????????????????????????????????????????????????????????????????????????????????????????????????????????????????????????????????????????????????????????????????????

			>Dyckia_machrisiana
			????????????????????????????????????????????????????????????????????????????????????????????????????????????????????????????????????????????????????????????????????????????????????????????????????????????????????????????????????????????????????????????????????????????????????????????????????????????????????????????????????????????????????????????????????????????????

			>Dyckia_maracasensis
			????????????????????????????????????????????????????????????????????????????????????????????????????????????????????????????????????????????????????????????????????????????????????????????????????????????????????????????????????????????????????????????????????????????????????????????????????????????????????????????????????????????????????????????????????????????????

			>Dyckia_maritima
			????????????????????????????????????????????????????????????????????????????????????????????????????????????????????????????????????????????????????????????????????????????????????????????????????????????????????????????????????????????????????????????????????????????????????????????????????????????????????????????????????????????????????????????????????????????????

			>Dyckia_marnier-lapostollei
			????????????????????????????????????????????????????????????????????????????????????????????????????????????????????????????????????????????????????????????????????????????????????????????????????????????????????????????????????????????????????????????????????????????????????????????????????????????????????????????????????????????????????????????????????????????????

			>Dyckia_marnier-lapostollei_var._estevesii
			????????????????????????????????????????????????????????????????????????????????????????????????????????????????????????????????????????????????????????????????????????????????????????????????????????????????????????????????????????????????????????????????????????????????????????????????????????????????????????????????????????????????????????????????????????????????

			>Dyckia_martinellii
			????????????????????????????????????????????????????????????????????????????????????????????????????????????????????????????????????????????????????????????????????????????????????????????????????????????????????????????????????????????????????????????????????????????????????????????????????????????????????????????????????????????????????????????????????????????????

			>Dyckia_mellobarretoi
			????????????????????????????????????????????????????????????????????????????????????????????????????????????????????????????????????????????????????????????????????????????????????????????????????????????????????????????????????????????????????????????????????????????????????????????????????????????????????????????????????????????????????????????????????????????????

			>Dyckia_microcalyx
			????????????????????????????????????????????????????????????????????????????????????????????????????????????????????????????????????????????????????????????????????????????????????????????????????????????????????????????????????????????????????????????????????????????????????????????????????????????????????????????????????????????????????????????????????????????????

			>Dyckia_microcalyx_var._ostenii
			????????????????????????????????????????????????????????????????????????????????????????????????????????????????????????????????????????????????????????????????????????????????????????????????????????????????????????????????????????????????????????????????????????????????????????????????????????????????????????????????????????????????????????????????????????????????

			>Dyckia_milagrensis
			????????????????????????????????????????????????????????????????????????????????????????????????????????????????????????????????????????????????????????????????????????????????????????????????????????????????????????????????????????????????????????????????????????????????????????????????????????????????????????????????????????????????????????????????????????????????

			>Dyckia_mirandiana
			????????????????????????????????????????????????????????????????????????????????????????????????????????????????????????????????????????????????????????????????????????????????????????????????????????????????????????????????????????????????????????????????????????????????????????????????????????????????????????????????????????????????????????????????????????????????

			>Dyckia_montezumensis
			????????????????????????????????????????????????????????????????????????????????????????????????????????????????????????????????????????????????????????????????????????????????????????????????????????????????????????????????????????????????????????????????????????????????????????????????????????????????????????????????????????????????????????????????????????????????

			>Dyckia_monticola
			????????????????????????????????????????????????????????????????????????????????????????????????????????????????????????????????????????????????????????????????????????????????????????????????????????????????????????????????????????????????????????????????????????????????????????????????????????????????????????????????????????????????????????????????????????????????

			>Dyckia_nana
			????????????????????????????????????????????????????????????????????????????????????????????????????????????????????????????????????????????????????????????????????????????????????????????????????????????????????????????????????????????????????????????????????????????????????????????????????????????????????????????????????????????????????????????????????????????????

			>Dyckia_niederleinii
			????????????????????????????????????????????????????????????????????????????????????????????????????????????????????????????????????????????????????????????????????????????????????????????????????????????????????????????????????????????????????????????????????????????????????????????????????????????????????????????????????????????????????????????????????????????????

			>Dyckia_oligantha
			????????????????????????????????????????????????????????????????????????????????????????????????????????????????????????????????????????????????????????????????????????????????????????????????????????????????????????????????????????????????????????????????????????????????????????????????????????????????????????????????????????????????????????????????????????????????

			>Dyckia_paraensis
			????????????????????????????????????????????????????????????????????????????????????????????????????????????????????????????????????????????????????????????????????????????????????????????????????????????????????????????????????????????????????????????????????????????????????????????????????????????????????????????????????????????????????????????????????????????????

			>Dyckia_pectinata
			????????????????????????????????????????????????????????????????????????????????????????????????????????????????????????????????????????????????????????????????????????????????????????????????????????????????????????????????????????????????????????????????????????????????????????????????????????????????????????????????????????????????????????????????????????????????

			>Dyckia_pernambucana
			????????????????????????????????????????????????????????????????????????????????????????????????????????????????????????????????????????????????????????????????????????????????????????????????????????????????????????????????????????????????????????????????????????????????????????????????????????????????????????????????????????????????????????????????????????????????

			>Dyckia_platyphylla
			????????????????????????????????????????????????????????????????????????????????????????????????????????????????????????????????????????????????????????????????????????????????????????????????????????????????????????????????????????????????????????????????????????????????????????????????????????????????????????????????????????????????????????????????????????????????

			>Dyckia_pottiorum
			????????????????????????????????????????????????????????????????????????????????????????????????????????????????????????????????????????????????????????????????????????????????????????????????????????????????????????????????????????????????????????????????????????????????????????????????????????????????????????????????????????????????????????????????????????????????

			>Dyckia_pseudococcinea
			????????????????????????????????????????????????????????????????????????????????????????????????????????????????????????????????????????????????????????????????????????????????????????????????????????????????????????????????????????????????????????????????????????????????????????????????????????????????????????????????????????????????????????????????????????????????

			>Dyckia_pulquinensis
			????????????????????????????????????????????????????????????????????????????????????????????????????????????????????????????????????????????????????????????????????????????????????????????????????????????????????????????????????????????????????????????????????????????????????????????????????????????????????????????????????????????????????????????????????????????????

			>Dyckia_pumila
			????????????????????????????????????????????????????????????????????????????????????????????????????????????????????????????????????????????????????????????????????????????????????????????????????????????????????????????????????????????????????????????????????????????????????????????????????????????????????????????????????????????????????????????????????????????????

			>Dyckia_rariflora
			????????????????????????????????????????????????????????????????????????????????????????????????????????????????????????????????????????????????????????????????????????????????????????????????????????????????????????????????????????????????????????????????????????????????????????????????????????????????????????????????????????????????????????????????????????????????

			>Dyckia_remotiflora
			????????????????????????????????????????????????????????????????????????????????????????????????????????????????????????????????????????????????????????????????????????????????????????????????????????????????????????????????????????????????????????????????????????????????????????????????????????????????????????????????????????????????????????????????????????????????

			>Dyckia_rondonopolitana
			????????????????????????????????????????????????????????????????????????????????????????????????????????????????????????????????????????????????????????????????????????????????????????????????????????????????????????????????????????????????????????????????????????????????????????????????????????????????????????????????????????????????????????????????????????????????

			>Dyckia_saxatilis
			????????????????????????????????????????????????????????????????????????????????????????????????????????????????????????????????????????????????????????????????????????????????????????????????????????????????????????????????????????????????????????????????????????????????????????????????????????????????????????????????????????????????????????????????????????????????

			>Dyckia_secunda
			????????????????????????????????????????????????????????????????????????????????????????????????????????????????????????????????????????????????????????????????????????????????????????????????????????????????????????????????????????????????????????????????????????????????????????????????????????????????????????????????????????????????????????????????????????????????

			>Dyckia_selloa
			????????????????????????????????????????????????????????????????????????????????????????????????????????????????????????????????????????????????????????????????????????????????????????????????????????????????????????????????????????????????????????????????????????????????????????????????????????????????????????????????????????????????????????????????????????????????

			>Dyckia_tenebrosa
			????????????????????????????????????????????????????????????????????????????????????????????????????????????????????????????????????????????????????????????????????????????????????????????????????????????????????????????????????????????????????????????????????????????????????????????????????????????????????????????????????????????????????????????????????????????????

			>Dyckia_tobatiensis
			????????????????????????????????????????????????????????????????????????????????????????????????????????????????????????????????????????????????????????????????????????????????????????????????????????????????????????????????????????????????????????????????????????????????????????????????????????????????????????????????????????????????????????????????????????????????

			>Dyckia_tomentella
			????????????????????????????????????????????????????????????????????????????????????????????????????????????????????????????????????????????????????????????????????????????????????????????????????????????????????????????????????????????????????????????????????????????????????????????????????????????????????????????????????????????????????????????????????????????????

			>Dyckia_ursina
			????????????????????????????????????????????????????????????????????????????????????????????????????????????????????????????????????????????????????????????????????????????????????????????????????????????????????????????????????????????????????????????????????????????????????????????????????????????????????????????????????????????????????????????????????????????????

			>Dyckia_velascana
			????????????????????????????????????????????????????????????????????????????????????????????????????????????????????????????????????????????????????????????????????????????????????????????????????????????????????????????????????????????????????????????????????????????????????????????????????????????????????????????????????????????????????????????????????????????????

			>Dyckia_vestita
			????????????????????????????????????????????????????????????????????????????????????????????????????????????????????????????????????????????????????????????????????????????????????????????????????????????????????????????????????????????????????????????????????????????????????????????????????????????????????????????????????????????????????????????????????????????????

			>Dyckia_vicentensis
			????????????????????????????????????????????????????????????????????????????????????????????????????????????????????????????????????????????????????????????????????????????????????????????????????????????????????????????????????????????????????????????????????????????????????????????????????????????????????????????????????????????????????????????????????????????????

			>Dyckia_walteriana
			????????????????????????????????????????????????????????????????????????????????????????????????????????????????????????????????????????????????????????????????????????????????????????????????????????????????????????????????????????????????????????????????????????????????????????????????????????????????????????????????????????????????????????????????????????????????

			>Edmundoa_ambigua
			TCGCAGAAAGCTCTTTCCCTGCCTACTGGAATGGGGATTCTCTGCGCGAGCCCGAAAGCTCTAGAAGCTTCCAAAACAGCCAAGTCGGTCAGGGTTTTCTTCGACTGGAATGACTACTTGAAGTTCTACAAGCTCGGAACTTACTGGCCTTAACGCCTTCCATCCAACTACTTTACGGGCTGAGAGCAGCTTTGGACCTCATCTTTGAGGAGGGGCTCGACAATGTCATCGAAAGGCATCGTCGCTTAGGAAAGGCCACAAGGTTAAGTAAAAGTTATACATCTAAATAGCCGCTTTCGCCGTGTTGGATAGATATATTACTTTGCACTATTGTCCTCTTGTAACAATGGGCAGACTTGCCGTCGAAG

			>Edmundoa_lindenii
			TCGCAGAAAGCTCTTTCCCTGCCTACTGGAATGGGGATTCTCTGCGCGAGCCCGAAAGCTCTAGAAGCTTCCAAAACAGCCAAGTCGGTCAGGGTTTTCTTCGACTGGAATGACTACTTGAAGTTCTACAAGCTCGGAACTTACTGGCCTTAACGCCTTCCATCCAACTACTTTACGGGCTGAGAGCAGCTTTGGACCTCATCTTTGAGGAGGGGCTCGACAATGTCATCGAAAGGCATCGTCGCTTAGGAAAGGCCACAAGGTTAACTAAAAGCTATACATCTAAATAGCCGCTTTCGCCGTGTTGGATAGATATATTACTTTGCACTACTGTCCTCTTGTAACAATGGGCAGACTTGCCGTCGAAG

			>Edmundoa_lindenii_var._rosaea
			????????????????????????????????????????????????????????????????????????????????????????????????????????????????????????????????????????????????????????????????????????????????????????????????????????????????????????????????????????????????????????????????????????????????????????????????????????????????????????????????????????????????????????????????????????????????

			>Edmundoa_perplexa
			TCGCAGAAAGCTCTTTCCCTGCCTACTGGAATGGGGATTCTCTGCGCGAGCCCGAAAGCTCTAGAAGCTTCCAAAACAGCCAAGTCGGTCAGGGTTTTCTTCGACTGGAATGACTACTTGAAGTTCTACAAGCTCGGAACTTACTGGCCTTAACGCCTTCCATCCAACTACTTTACGGGCTGAGAGCAGCTTTGGACCTCATCTTTGAGGAGGGGCTCGACAATGTCATCGAAAGGCATCGTCGCTTAGGAAAGGCCACAAGGTTAAGTMAAAGTTATACATCTAAATAGCCGCTTTCGCCGTGTTGGATAGATATATTACTTTGCACTATTGTCCTCTTGTAACAATGGGCAGACTTGCCGTCGAAG

			>Eduandrea_selloana
			TCGCAGAAAGCTCTTTCCCTGCCTACTGGAATGGGGATTCTCTGCGCGAGCCCGAAAGCTCTAGAAGCTTCCAAAACAGCCAAGTCGGTCAGGGTTTTCTTCGACTGGAATGACTACTTGAAGTTCTACAAGCTCGGAACTTACTGGCCTTAACGCCTTCCATCCAACTACTTTACGGGCTGAGAGCAGCTTTGGACCTCATCTTTGAGGAGGGGCTCGACAATGTCATCGAAAGGCATCGTCGCTTAGGAAAGGCCACAAGGTTAAGTAAAAGCTATACATCTAAATAGCCGCTTTCGCCGTGTTGGATAGATATATTACTTTGCACTACTGTCCTCTTGTAACAATGGGCAGACTTGCCGTCGAAG

			>Encholirium_agavoides
			????????????????????????????????????????????????????????????????????????????????????????????????????????????????????????????????????????????????????????????????????????????????????????????????????????????????????????????????????????????????????????????????????????????????????????????????????????????????????????????????????????????????????????????????????????????????

			>Encholirium_biflorum
			????????????????????????????????????????????????????????????????????????????????????????????????????????????????????????????????????????????????????????????????????????????????????????????????????????????????????????????????????????????????????????????????????????????????????????????????????????????????????????????????????????????????????????????????????????????????

			>Encholirium_brachypodum
			????????????????????????????????????????????????????????????????????????????????????????????????????????????????????????????????????????????????????????????????????????????????????????????????????????????????????????????????????????????????????????????????????????????????????????????????????????????????????????????????????????????????????????????????????????????????

			>Encholirium_ctenophyllum
			????????????????????????????????????????????????????????????????????????????????????????????????????????????????????????????????????????????????????????????????????????????????????????????????????????????????????????????????????????????????????????????????????????????????????????????????????????????????????????????????????????????????????????????????????????????????

			>Encholirium_diamantinum
			????????????????????????????????????????????????????????????????????????????????????????????????????????????????????????????????????????????????????????????????????????????????????????????????????????????????????????????????????????????????????????????????????????????????????????????????????????????????????????????????????????????????????????????????????????????????

			>Encholirium_eddie-estevesii
			????????????????????????????????????????????????????????????????????????????????????????????????????????????????????????????????????????????????????????????????????????????????????????????????????????????????????????????????????????????????????????????????????????????????????????????????????????????????????????????????????????????????????????????????????????????????

			>Encholirium_erectiflorum
			????????????????????????????????????????????????????????????????????????????????????????????????????????????????????????????????????????????????????????????????????????????????????????????????????????????????????????????????????????????????????????????????????????????????????????????????????????????????????????????????????????????????????????????????????????????????

			>Encholirium_fragae
			????????????????????????????????????????????????????????????????????????????????????????????????????????????????????????????????????????????????????????????????????????????????????????????????????????????????????????????????????????????????????????????????????????????????????????????????????????????????????????????????????????????????????????????????????????????????

			>Encholirium_gracile
			????????????????????????????????????????????????????????????????????????????????????????????????????????????????????????????????????????????????????????????????????????????????????????????????????????????????????????????????????????????????????????????????????????????????????????????????????????????????????????????????????????????????????????????????????????????????

			>Encholirium_heloisae
			????????????????????????????????????????????????????????????????????????????????????????????????????????????????????????????????????????????????????????????????????????????????????????????????????????????????????????????????????????????????????????????????????????????????????????????????????????????????????????????????????????????????????????????????????????????????

			>Encholirium_horridum
			????????????????????????????????????????????????????????????????????????????????????????????????????????????????????????????????????????????????????????????????????????????????????????????????????????????????????????????????????????????????????????????????????????????????????????????????????????????????????????????????????????????????????????????????????????????????

			>Encholirium_irwinii
			????????????????????????????????????????????????????????????????????????????????????????????????????????????????????????????????????????????????????????????????????????????????????????????????????????????????????????????????????????????????????????????????????????????????????????????????????????????????????????????????????????????????????????????????????????????????

			>Encholirium_kranzianum
			????????????????????????????????????????????????????????????????????????????????????????????????????????????????????????????????????????????????????????????????????????????????????????????????????????????????????????????????????????????????????????????????????????????????????????????????????????????????????????????????????????????????????????????????????????????????

			>Encholirium_longiflorum
			????????????????????????????????????????????????????????????????????????????????????????????????????????????????????????????????????????????????????????????????????????????????????????????????????????????????????????????????????????????????????????????????????????????????????????????????????????????????????????????????????????????????????????????????????????????????

			>Encholirium_luxor
			????????????????????????????????????????????????????????????????????????????????????????????????????????????????????????????????????????????????????????????????????????????????????????????????????????????????????????????????????????????????????????????????????????????????????????????????????????????????????????????????????????????????????????????????????????????????

			>Encholirium_lymanianum
			????????????????????????????????????????????????????????????????????????????????????????????????????????????????????????????????????????????????????????????????????????????????????????????????????????????????????????????????????????????????????????????????????????????????????????????????????????????????????????????????????????????????????????????????????????????????

			>Encholirium_magalhaesii
			????????????????????????????????????????????????????????????????????????????????????????????????????????????????????????????????????????????????????????????????????????????????????????????????????????????????????????????????????????????????????????????????????????????????????????????????????????????????????????????????????????????????????????????????????????????????

			>Encholirium_maximum
			????????????????????????????????????????????????????????????????????????????????????????????????????????????????????????????????????????????????????????????????????????????????????????????????????????????????????????????????????????????????????????????????????????????????????????????????????????????????????????????????????????????????????????????????????????????????

			>Encholirium_pedicellatum
			????????????????????????????????????????????????????????????????????????????????????????????????????????????????????????????????????????????????????????????????????????????????????????????????????????????????????????????????????????????????????????????????????????????????????????????????????????????????????????????????????????????????????????????????????????????????

			>Encholirium_pulchrum
			????????????????????????????????????????????????????????????????????????????????????????????????????????????????????????????????????????????????????????????????????????????????????????????????????????????????????????????????????????????????????????????????????????????????????????????????????????????????????????????????????????????????????????????????????????????????

			>Encholirium_scrutor
			????????????????????????????????????????????????????????????????????????????????????????????????????????????????????????????????????????????????????????????????????????????????????????????????????????????????????????????????????????????????????????????????????????????????????????????????????????????????????????????????????????????????????????????????????????????????

			>Encholirium_spectabile
			????????????????????????????????????????????????????????????????????????????????????????????????????????????????????????????????????????????????????????????????????????????????????????????????????????????????????????????????????????????????????????????????????????????????????????????????????????????????????????????????????????????????????????????????????????????????

			>Encholirium_splendidum
			????????????????????????????????????????????????????????????????????????????????????????????????????????????????????????????????????????????????????????????????????????????????????????????????????????????????????????????????????????????????????????????????????????????????????????????????????????????????????????????????????????????????????????????????????????????????

			>Encholirium_structor
			????????????????????????????????????????????????????????????????????????????????????????????????????????????????????????????????????????????????????????????????????????????????????????????????????????????????????????????????????????????????????????????????????????????????????????????????????????????????????????????????????????????????????????????????????????????????

			>Encholirium_subsecundum
			????????????????????????????????????????????????????????????????????????????????????????????????????????????????????????????????????????????????????????????????????????????????????????????????????????????????????????????????????????????????????????????????????????????????????????????????????????????????????????????????????????????????????????????????????????????????

			>Encholirium_vogelii
			????????????????????????????????????????????????????????????????????????????????????????????????????????????????????????????????????????????????????????????????????????????????????????????????????????????????????????????????????????????????????????????????????????????????????????????????????????????????????????????????????????????????????????????????????????????????

			>Epidryos_allenii
			????????????????????????????????????????????????????????????????????????????????????????????????????????????????????????????????????????????????????????????????????????????????????????????????????????????????????????????????????????????????????????????????????????????????????????????????????????????????????????????????????????????????????????????????????????????????

			>Fascicularia_bicolor
			????????????????????????????????????????????????????????????????????????????????????????????????????????????????????????????????????????????????????????????????????????????????????????????????????????????????????????????????????????????????????????????????????????????????????????????????????????????????????????????????????????????????????????????????????????????????

			>Fernseea_itatiaiae
			????????????????????????????????????????????????????????????????????????????????????????????????????????????????????????????????????????????????????????????????????????????????????????????????????????????????????????????????????????????????????????????????????????????????????????????????????????????????????????????????????????????????????????????????????????????????

			>Forzzaea_leopoldo-horstii
			TCGCAGAAGGCTCTTTCGATGCCTACTGGAATGGGCATTCTTTGCGCGAGCCCGAAAGCTCTAGAAGCTTCCAAAACAGCCAAGTCGGTCAGGGTTTTCTTCGACTGGAATGACTACTTGAAGTTCTACAAGCTCGGAACTTACTGGCCTTAACGCCTTCCATCCAACTACTTTACGGGCTAAGAGCAGCTTTGGATCTCATCTTTGAGGAGGGGCTCGACAATGTCATCGAAAGGCATCGTCGCTTAGGAAAGGCCACAAGGTCCAGTAAAAGCTATACATCTATATAGCCGCTTTCGCCGTGTTGGATAGATATATTACTCGGCGCTATTGGCCTCTTGTAACAATGAGCAGACTTGCTGTCGAAG

			>Forzzaea_warasii
			????????????????????????????????????????????????????????????????????????????????????????????????????????????????????????????????????????????????????????????????????????????????????????????????????????????????????????????????????????????????????????????????????????????????????????????????????????????????????????????????????????????????????????????????????????????????

			>Fosterella_albicans
			????????????????????????????????????????????????????????????????????????????????????????????????????????????????????????????????????????????????????????????????????????????????????????????????????????????????????????????????????????????????????????????????????????????????????????????????????????????????????????????????????????????????????????????????????????????????

			>Fosterella_batistana
			TCGCAGAAAGCACTTTCCCTGCCTACTGGAATGGGCATTGTTTGCGCGAGCCCGAAAGCTCTGGAAGCTTCCAAAAATGCCAAGTCGGTCAGAGTTTTCTTCGACTGGAATGACTACTTGAAGTTCTACAAGCTTGGAACTTACTGGCCGTAACGCCTTCCATCCAACTACTCTACGGGCTAAGAGCAGCTTTGGATCTCATCTTTGAGGAGGGGCTCGAAAATGTCATCGAAAGGCATCGTCGTTTAGGAAAGGCCACGAGG-TAAGTAAAAGCT--------AAATAGCCGATTTCACAGTATAGGAAAGATATGTAATTAGGCATTATTGTCCTCTTGTAACAATGAGCAGACTTGCCGTCGAAG

			>Fosterella_caulescens
			????????????????????????????????????????????????????????????????????????????????????????????????????????????????????????????????????????????????????????????????????????????????????????????????????????????????????????????????????????????????????????????????????????????????????????????????????????????????????????????????????????????????????????????????????????????????

			>Fosterella_christophii
			????????????????????????????????????????????????????????????????????????????????????????????????????????????????????????????????????????????????????????????????????????????????????????????????????????????????????????????????????????????????????????????????????????????????????????????????????????????????????????????????????????????????????????????????????????????????

			>Fosterella_cotacajensis
			????????????????????????????????????????????????????????????????????????????????????????????????????????????????????????????????????????????????????????????????????????????????????????????????????????????????????????????????????????????????????????????????????????????????????????????????????????????????????????????????????????????????????????????????????????????????

			>Fosterella_floridensis
			????????????????????????????????????????????????????????????????????????????????????????????????????????????????????????????????????????????????????????????????????????????????????????????????????????????????????????????????????????????????????????????????????????????????????????????????????????????????????????????????????????????????????????????????????????????????

			>Fosterella_gracilis
			????????????????????????????????????????????????????????????????????????????????????????????????????????????????????????????????????????????????????????????????????????????????????????????????????????????????????????????????????????????????????????????????????????????????????????????????????????????????????????????????????????????????????????????????????????????????

			>Fosterella_graminea
			????????????????????????????????????????????????????????????????????????????????????????????????????????????????????????????????????????????????????????????????????????????????????????????????????????????????????????????????????????????????????????????????????????????????????????????????????????????????????????????????????????????????????????????????????????????????

			>Fosterella_hatschbachii
			????????????????????????????????????????????????????????????????????????????????????????????????????????????????????????????????????????????????????????????????????????????????????????????????????????????????????????????????????????????????????????????????????????????????????????????????????????????????????????????????????????????????????????????????????????????????

			>Fosterella_heterophylla
			????????????????????????????????????????????????????????????????????????????????????????????????????????????????????????????????????????????????????????????????????????????????????????????????????????????????????????????????????????????????????????????????????????????????????????????????????????????????????????????????????????????????????????????????????????????????

			>Fosterella_kroemeri
			????????????????????????????????????????????????????????????????????????????????????????????????????????????????????????????????????????????????????????????????????????????????????????????????????????????????????????????????????????????????????????????????????????????????????????????????????????????????????????????????????????????????????????????????????????????????

			>Fosterella_micrantha
			????????????????????????????????????????????????????????????????????????????????????????????????????????????????????????????????????????????????????????????????????????????????????????????????????????????????????????????????????????????????????????????????????????????????????????????????????????????????????????????????????????????????????????????????????????????????

			>Fosterella_penduliflora
			????????????????????????????????????????????????????????????????????????????????????????????????????????????????????????????????????????????????????????????????????????????????????????????????????????????????????????????????????????????????????????????????????????????????????????????????????????????????????????????????????????????????????????????????????????????????

			>Fosterella_petiolata
			????????????????????????????????????????????????????????????????????????????????????????????????????????????????????????????????????????????????????????????????????????????????????????????????????????????????????????????????????????????????????????????????????????????????????????????????????????????????????????????????????????????????????????????????????????????????

			>Fosterella_rexiae
			????????????????????????????????????????????????????????????????????????????????????????????????????????????????????????????????????????????????????????????????????????????????????????????????????????????????????????????????????????????????????????????????????????????????????????????????????????????????????????????????????????????????????????????????????????????????

			>Fosterella_robertreadii
			????????????????????????????????????????????????????????????????????????????????????????????????????????????????????????????????????????????????????????????????????????????????????????????????????????????????????????????????????????????????????????????????????????????????????????????????????????????????????????????????????????????????????????????????????????????????

			>Fosterella_rusbyi
			TCGCAGAAAGCTCTTTCCCTGCCTACTGGAATGGGCATTGTTTGCGCGAGCCCAAAAGCGCTAGAAGCTTCCAAAACAGCCAGGTCGGTCCGGGTTTTCTTCGACTGGAATGACTACTTGAAGTTCTACAAGCTCGGAACTTACTGGCCTTAACGCCTTCCATCCAACTACTTTACGGGCTGAGGGCAGCTTTGGATCTCATCTTCGAGGAGGGGCTCGAGAATGTCATCGAAAGGCATCGTCGTCTAGGAAAGGCCACGAGGT-AAGTAACAGCT--------------CTGTTTTTACAGCATCGGATAGATATATCTGCTTGTATTATTGTCCTCTTGTAACAATGAGCAGGCTCGCTGTCGAAG

			>Fosterella_schidosperma
			????????????????????????????????????????????????????????????????????????????????????????????????????????????????????????????????????????????????????????????????????????????????????????????????????????????????????????????????????????????????????????????????????????????????????????????????????????????????????????????????????????????????????????????????????????????????

			>Fosterella_spectabilis
			????????????????????????????????????????????????????????????????????????????????????????????????????????????????????????????????????????????????????????????????????????????????????????????????????????????????????????????????????????????????????????????????????????????????????????????????????????????????????????????????????????????????????????????????????????????????

			>Fosterella_vasquezii
			????????????????????????????????????????????????????????????????????????????????????????????????????????????????????????????????????????????????????????????????????????????????????????????????????????????????????????????????????????????????????????????????????????????????????????????????????????????????????????????????????????????????????????????????????????????????

			>Fosterella_villosula
			????????????????????????????????????????????????????????????????????????????????????????????????????????????????????????????????????????????????????????????????????????????????????????????????????????????????????????????????????????????????????????????????????????????????????????????????????????????????????????????????????????????????????????????????????????????????

			>Fosterella_weberbaueri
			????????????????????????????????????????????????????????????????????????????????????????????????????????????????????????????????????????????????????????????????????????????????????????????????????????????????????????????????????????????????????????????????????????????????????????????????????????????????????????????????????????????????????????????????????????????????

			>Fosterella_weddelliana
			????????????????????????????????????????????????????????????????????????????????????????????????????????????????????????????????????????????????????????????????????????????????????????????????????????????????????????????????????????????????????????????????????????????????????????????????????????????????????????????????????????????????????????????????????????????????

			>Fosterella_windischii
			????????????????????????????????????????????????????????????????????????????????????????????????????????????????????????????????????????????????????????????????????????????????????????????????????????????????????????????????????????????????????????????????????????????????????????????????????????????????????????????????????????????????????????????????????????????????

			>Fosterella_yuvinkae
			????????????????????????????????????????????????????????????????????????????????????????????????????????????????????????????????????????????????????????????????????????????????????????????????????????????????????????????????????????????????????????????????????????????????????????????????????????????????????????????????????????????????????????????????????????????????

			>Glomeropitcairnia_erectiflora
			????????????????????????????????????????????????????????????????????????????????????????????????????????????????????????????????????????????????????????????????????????????????????????????????????????????????????????????????????????????????????????????????????????????????????????????????????????????????????????????????????????????????????????????????????????????????

			>Glomeropitcairnia_penduliflora
			????????????????????????????????????????????????????????????????????????????????????????????????????????????????????????????????????????????????????????????????????????????????????????????????????????????????????????????????????????????????????????????????????????????????????????????????????????????????????????????????????????????????????????????????????????????????

			>Goudaea_chrysostachys
			????????????????????????????????????????????????????????????????????????????????????????????????????????????????????????????????????????????????????????????????????????????????????????????????????????????????????????????????????????????????????????????????????????????????????????????????????????????????????????????????????????????????????????????????????????????????

			>Goudaea_ospinae
			TCGCAGAAAGCACTTTCCCTGCCTACTGGAATGGGCATTGTTTGCGCGAGCCCGAAAGCTCTAGAAGCTTCCAAAAATGCCAAGTCGGTCAGAGTTTTCTTCGACTGGAATGACTACTTGAAGTTCTACAAGCTTGGAACTTACTGGCCGTAACGCCTTCCATCCAACTACTCTAYGGGCTAAGAGCAGCTTTGGATCTCATCTTTGAGGAGGGGCTCGAAAATGTCATCGAAAGGCATCGCCGTTTAGGAAAGGCCACGAGG-TAAGTA-AAGCC--------ATATAGCCGATCTTAC--AATGGCAGAGACGTCTAGTTGCATATTATTTC------ATAACAATGAGCAGGCTTGCTGTGGAAG

			>Goudaea_ospinae_var._gruberi
			????????????????????????????????????????????????????????????????????????????????????????????????????????????????????????????????????????????????????????????????????????????????????????????????????????????????????????????????????????????????????????????????????????????????????????????????????????????????????????????????????????????????????????????????????????????????

			>Gregbrownia_fulgens
			????????????????????????????????????????????????????????????????????????????????????????????????????????????????????????????????????????????????????????????????????????????????????????????????????????????????????????????????????????????????????????????????????????????????????????????????????????????????????????????????????????????????????????????????????????????????

			>Gregbrownia_hutchisonii
			????????????????????????????????????????????????????????????????????????????????????????????????????????????????????????????????????????????????????????????????????????????????????????????????????????????????????????????????????????????????????????????????????????????????????????????????????????????????????????????????????????????????????????????????????????????????

			>Greigia_mulfordii
			????????????????????????????????????????????????????????????????????????????????????????????????????????????????????????????????????????????????????????????????????????????????????????????????????????????????????????????????????????????????????????????????????????????????????????????????????????????????????????????????????????????????????????????????????????????????

			>Greigia_sphacelata
			????????????????????????????????????????????????????????????????????????????????????????????????????????????????????????????????????????????????????????????????????????????????????????????????????????????????????????????????????????????????????????????????????????????????????????????????????????????????????????????????????????????????????????????????????????????????

			>Guzmania_acorifolia
			????????????????????????????????????????????????????????????????????????????????????????????????????????????????????????????????????????????????????????????????????????????????????????????????????????????????????????????????????????????????????????????????????????????????????????????????????????????????????????????????????????????????????????????????????????????????

			>Guzmania_angustifolia
			????????????????????????????????????????????????????????????????????????????????????????????????????????????????????????????????????????????????????????????????????????????????????????????????????????????????????????????????????????????????????????????????????????????????????????????????????????????????????????????????????????????????????????????????????????????????

			>Guzmania_butcheri
			????????????????????????????????????????????????????????????????????????????????????????????????????????????????????????????????????????????????????????????????????????????????????????????????????????????????????????????????????????????????????????????????????????????????????????????????????????????????????????????????????????????????????????????????????????????????

			>Guzmania_condensata
			????????????????????????????????????????????????????????????????????????????????????????????????????????????????????????????????????????????????????????????????????????????????????????????????????????????????????????????????????????????????????????????????????????????????????????????????????????????????????????????????????????????????????????????????????????????????

			>Guzmania_coriostachya
			????????????????????????????????????????????????????????????????????????????????????????????????????????????????????????????????????????????????????????????????????????????????????????????????????????????????????????????????????????????????????????????????????????????????????????????????????????????????????????????????????????????????????????????????????????????????

			>Guzmania_donnellsmithii
			????????????????????????????????????????????????????????????????????????????????????????????????????????????????????????????????????????????????????????????????????????????????????????????????????????????????????????????????????????????????????????????????????????????????????????????????????????????????????????????????????????????????????????????????????????????????

			>Guzmania_glaucophylla
			????????????????????????????????????????????????????????????????????????????????????????????????????????????????????????????????????????????????????????????????????????????????????????????????????????????????????????????????????????????????????????????????????????????????????????????????????????????????????????????????????????????????????????????????????????????????

			>Guzmania_graminifolia
			????????????????????????????????????????????????????????????????????????????????????????????????????????????????????????????????????????????????????????????????????????????????????????????????????????????????????????????????????????????????????????????????????????????????????????????????????????????????????????????????????????????????????????????????????????????????

			>Guzmania_herrerae
			????????????????????????????????????????????????????????????????????????????????????????????????????????????????????????????????????????????????????????????????????????????????????????????????????????????????????????????????????????????????????????????????????????????????????????????????????????????????????????????????????????????????????????????????????????????????

			>Guzmania_lingulata
			TCGCAGAAAGCACTTTCCCTGCCTACTGGAATGGGCATTGTTTGCGCGAGCCCGAAAGCTCTAGAAGCTTCCAAAAATGCCAAGTCGGTCAGAGTTTTCTTCGACTGGAATGACTACTTGAAGTTCTACAAGCTCGGAACTTACTGGCCGTAACGCCTTCCATCCAACTACTTTACGGGCTGAGAGCAGCTTTGGATCTCATCTTCGAGGAGGGGCTCGAAAATGTCATCGAAAGGCATCGTCGTTTAGGAAAGGCCACGAGG-TAAGTA-AAGCT--------AAATAGCCGATTTTACGG--TAGGAAAGATATGTAATTAGCCATTATTGTCCTCTTGTAACAATGAGCAGACTTGCTGTCGAAG

			>Guzmania_melinonis
			????????????????????????????????????????????????????????????????????????????????????????????????????????????????????????????????????????????????????????????????????????????????????????????????????????????????????????????????????????????????????????????????????????????????????????????????????????????????????????????????????????????????????????????????????????????????

			>Guzmania_monostachia
			????????????????????????????????????????????????????????????????????????????????????????????????????????????????????????????????????????????????????????????????????????????????????????????????????????????????????????????????????????????????????????????????????????????????????????????????????????????????????????????????????????????????????????????????????????????????

			>Guzmania_mucronata
			????????????????????????????????????????????????????????????????????????????????????????????????????????????????????????????????????????????????????????????????????????????????????????????????????????????????????????????????????????????????????????????????????????????????????????????????????????????????????????????????????????????????????????????????????????????????

			>Guzmania_multiflora
			TCGCAGAAGGCACTTTCC?TGCCTACTGG?ATGGGCATTGTTTGCGCGAGCCCGAAAGCTCTAGAAGCTTCCAAAAATGCCAAGTCGGTCAGAGTTTTCTTCGACTGGAATGACTACTTGAAGTTCTACAAGCTCGGAACTTACTGGCCGTAACSCCTTCCWTCCAACTACTTTACGGGCTGAGAGCAGCTTTGGATCTCATCTTCGAGGAGGGGCTCGAAAATGTCATCGAAAGGCATCGTCGTTTAGGAAAGGCCACGAGG-TAAGTA-AAGCT--------AAATAGCCGATTTTACGG--TAGGAAAGATATATAATTAGCCATTATTGTCCTCTTGTAACAATGAGCAGACTTGCTGTCGAAG

			>Guzmania_musaica
			TCGCAGAAAGCACTTTCCCTGCCTACTGGAATGGGCATTGTTTGCGCGAGCCCGAAAGCTCTAGAAGCTTCCAAAAATGCCAAGTCGGTCAGAGTTTTCTTCGACTGGAATGACTACTTGAAGTTCTACAAGCTTGGAACTTACTGGCCGTAACGCCTTCCATCCAACTACTTTACGGGCTAAGAGCAGCTTTGGATCTCATCTTTGAGGAGGGGCTCGAAAATGTCATCGAAAGGCATCGTCGTTTAGGAAAGGCCACCAGG-TAAGTAAAACCT--------AAATAGCCGATTTTACAGTATAGGAAAGATATATAATTAGTCATTATTGTCTCTTCGTAACAACGAGCAGACTTGCTGTCGAAG

			>Guzmania_nicaraguensis
			????????????????????????????????????????????????????????????????????????????????????????????????????????????????????????????????????????????????????????????????????????????????????????????????????????????????????????????????????????????????????????????????????????????????????????????????????????????????????????????????????????????????????????????????????????????????

			>Guzmania_osyana
			????????????????????????????????????????????????????????????????????????????????????????????????????????????????????????????????????????????????????????????????????????????????????????????????????????????????????????????????????????????????????????????????????????????????????????????????????????????????????????????????????????????????????????????????????????????????

			>Guzmania_patula
			????????????????????????????????????????????????????????????????????????????????????????????????????????????????????????????????????????????????????????????????????????????????????????????????????????????????????????????????????????????????????????????????????????????????????????????????????????????????????????????????????????????????????????????????????????????????

			>Guzmania_plicatifolia
			????????????????????????????????????????????????????????????????????????????????????????????????????????????????????????????????????????????????????????????????????????????????????????????????????????????????????????????????????????????????????????????????????????????????????????????????????????????????????????????????????????????????????????????????????????????????

			>Guzmania_polycephala
			????????????????????????????????????????????????????????????????????????????????????????????????????????????????????????????????????????????????????????????????????????????????????????????????????????????????????????????????????????????????????????????????????????????????????????????????????????????????????????????????????????????????????????????????????????????????

			>Guzmania_rhonhofiana
			TCGCAGAAAGCACTTTCCCTGCCTACTGGAATGGGCATTGTTTGCGCGAGCCCGAAAGCTCTAGAAGCTTCCAAAAATGCCAAGTCGGTCAGGGTTTTCTTCGACTGGAATGACTACTTGAAGTTCTACAAGCTCGGAACTTACTGGCCGTAACGCCTTCCATCCAACTACTCTACGGGCTGAGAGCAGCTTTGGATCTCATCTTCGAGGAGGGGCTCGAAAATGTCATCGAAAGGCATCGTCGTTTAGGAAAGGCCACGAGG-TAAGTA-AAGCT--------AAATAGCCGATTTTACGG--TAGGAAAGATATATAATTAGCCATTATTGTCCTCTTGTAACAATGAGCAGACTTGCTGTCGAAG

			>Guzmania_roezlii
			????????????????????????????????????????????????????????????????????????????????????????????????????????????????????????????????????????????????????????????????????????????????????????????????????????????????????????????????????????????????????????????????????????????????????????????????????????????????????????????????????????????????????????????????????????????????

			>Guzmania_sanguinea
			????????????????????????????????????????????????????????????????????????????????????????????????????????????????????????????????????????????????????????????????????????????????????????????????????????????????????????????????????????????????????????????????????????????????????????????????????????????????????????????????????????????????????????????????????????????????

			>Guzmania_scherzeriana
			TCGCAGAAAGCACTTTCCCTGCCTACTGGAATGGGCATTGTTTGCGCGAGCCCGAAAGCTCTAGAAGCTTCCAAAAATGCCAAGTCGGTCAGAGTTTTCTTCGACTGGAATGACTACTTGAAGTTCTACAAGCTCGGAACTTACTGGCCGTAACGCCTTCCATCCAACTACTTTACGGGCTAAGAGCAGCTTTGGATCTCATCTTCGAGGAGGGGCTCGAAAATGTCATCGAAAGGCATCGTCGTTTAGGAAAGGCCACGAGG-TAAGTA-AAGCT--------AAATAGCCGATTTTTCGG--TAGGAAAGATATATAATTAGCCATTATTG------------AATGAGCAGACTTGCTGTGGAAG

			>Guzmania_spectabilis
			????????????????????????????????????????????????????????????????????????????????????????????????????????????????????????????????????????????????????????????????????????????????????????????????????????????????????????????????????????????????????????????????????????????????????????????????????????????????????????????????????????????????????????????????????????????????

			>Guzmania_subcorymbosa
			????????????????????????????????????????????????????????????????????????????????????????????????????????????????????????????????????????????????????????????????????????????????????????????????????????????????????????????????????????????????????????????????????????????????????????????????????????????????????????????????????????????????????????????????????????????????

			>Guzmania_variegata
			????????????????????????????????????????????????????????????????????????????????????????????????????????????????????????????????????????????????????????????????????????????????????????????????????????????????????????????????????????????????????????????????????????????????????????????????????????????????????????????????????????????????????????????????????????????????

			>Guzmania_wittmackii
			????????????????????????????????????????????????????????????????????????????????????????????????????????????????????????????????????????????????????????????????????????????????????????????????????????????????????????????????????????????????????????????????????????????????????????????????????????????????????????????????????????????????????????????????????????????????

			>Guzmania_zahnii
			TCGCAGAAGGCACTTTCCCTGCCTACTGGAATGGGCATTGTTTGCGCGAGCCCGAAAGCTCTAGAAGCTTCCAAAAATGCCAAGTCGGTCAGAGTTTTCTTCGACTGGAATGACTACTTGAAGTTCTACAAGCTCGGAACTTACTGGCCGTAACGCCTTCCATCCAACTACTTTACGGGCTAAGAGCAGCTTTGGATCTCATCTTCGAGGAGGGGCTCGAAAATGTCATCGAAAGGCATCGTCGTTTAGGAAAGGCCACGAGG-TAAGTA-AAGCT--------AAAGAGCTGATTTTACGG--TAGGAAAGATATATAATTAGCCATTATTGTACTCTTGTGACAATGAGCAGACTTGCTGTCGAAG

			>Hechtia_argentea
			????????????????????????????????????????????????????????????????????????????????????????????????????????????????????????????????????????????????????????????????????????????????????????????????????????????????????????????????????????????????????????????????????????????????????????????????????????????????????????????????????????????????????????????????????????????????

			>Hechtia_caerulea
			????????????????????????????????????????????????????????????????????????????????????????????????????????????????????????????????????????????????????????????????????????????????????????????????????????????????????????????????????????????????????????????????????????????????????????????????????????????????????????????????????????????????????????????????????????????????

			>Hechtia_carlsoniae
			????????????????????????????????????????????????????????????????????????????????????????????????????????????????????????????????????????????????????????????????????????????????????????????????????????????????????????????????????????????????????????????????????????????????????????????????????????????????????????????????????????????????????????????????????????????????

			>Hechtia_deceptrix
			????????????????????????????????????????????????????????????????????????????????????????????????????????????????????????????????????????????????????????????????????????????????????????????????????????????????????????????????????????????????????????????????????????????????????????????????????????????????????????????????????????????????????????????????????????????????

			>Hechtia_dichroantha
			????????????????????????????????????????????????????????????????????????????????????????????????????????????????????????????????????????????????????????????????????????????????????????????????????????????????????????????????????????????????????????????????????????????????????????????????????????????????????????????????????????????????????????????????????????????????

			>Hechtia_epigyna
			????????????????????????????????????????????????????????????????????????????????????????????????????????????????????????????????????????????????????????????????????????????????????????????????????????????????????????????????????????????????????????????????????????????????????????????????????????????????????????????????????????????????????????????????????????????????

			>Hechtia_fosteriana
			????????????????????????????????????????????????????????????????????????????????????????????????????????????????????????????????????????????????????????????????????????????????????????????????????????????????????????????????????????????????????????????????????????????????????????????????????????????????????????????????????????????????????????????????????????????????

			>Hechtia_ghiesbreghtii
			????????????????????????????????????????????????????????????????????????????????????????????????????????????????????????????????????????????????????????????????????????????????????????????????????????????????????????????????????????????????????????????????????????????????????????????????????????????????????????????????????????????????????????????????????????????????

			>Hechtia_glomerata
			????????????????????????????????????????????????????????????????????????????????????????????????????????????????????????????????????????????????????????????????????????????????????????????????????????????????????????????????????????????????????????????????????????????????????????????????????????????????????????????????????????????????????????????????????????????????

			>Hechtia_guatemalensis
			TCGCAGAAAGCTCTTTCCCTGCCTACTGGAATGGGCATTCTTTGTGCGAGCCCGAAAGCTTTAGAAGCTTCAAAAACGGCCAAGTCGGTTAGAGTTTTCTTCGACTGGAATGACTACTTGAAGTTCTACAAGCTTGGAACTTACTGGCCTTAACGCCTTCCATCCAACTACTGTACGGGCTAAGAGCAGCTTTGGATCTCATCTTTGAGGAGGGGCTCGATAACGTCATCGAAAGGCATCATCGTTTAGGAAAGGCCACAAGGT-AAGTAAAAGCTATACGCCT-AATAGTTGATTTTTTCCTGTTGGAAAGATTTACTACTTGGCATTGCCGTCCTCTTGTAACAATGAGCAGACTTGCTGTCGAAG

			>Hechtia_hernandez-sandovalii
			????????????????????????????????????????????????????????????????????????????????????????????????????????????????????????????????????????????????????????????????????????????????????????????????????????????????????????????????????????????????????????????????????????????????????????????????????????????????????????????????????????????????????????????????????????????????

			>Hechtia_isthmusiana
			????????????????????????????????????????????????????????????????????????????????????????????????????????????????????????????????????????????????????????????????????????????????????????????????????????????????????????????????????????????????????????????????????????????????????????????????????????????????????????????????????????????????????????????????????????????????

			>Hechtia_lepidophylla
			????????????????????????????????????????????????????????????????????????????????????????????????????????????????????????????????????????????????????????????????????????????????????????????????????????????????????????????????????????????????????????????????????????????????????????????????????????????????????????????????????????????????????????????????????????????????

			>Hechtia_lundelliorum
			????????????????????????????????????????????????????????????????????????????????????????????????????????????????????????????????????????????????????????????????????????????????????????????????????????????????????????????????????????????????????????????????????????????????????????????????????????????????????????????????????????????????????????????????????????????????

			>Hechtia_malvernii
			????????????????????????????????????????????????????????????????????????????????????????????????????????????????????????????????????????????????????????????????????????????????????????????????????????????????????????????????????????????????????????????????????????????????????????????????????????????????????????????????????????????????????????????????????????????????

			>Hechtia_matudae
			????????????????????????????????????????????????????????????????????????????????????????????????????????????????????????????????????????????????????????????????????????????????????????????????????????????????????????????????????????????????????????????????????????????????????????????????????????????????????????????????????????????????????????????????????????????????

			>Hechtia_montana
			????????????????????????????????????????????????????????????????????????????????????????????????????????????????????????????????????????????????????????????????????????????????????????????????????????????????????????????????????????????????????????????????????????????????????????????????????????????????????????????????????????????????????????????????????????????????

			>Hechtia_myriantha
			????????????????????????????????????????????????????????????????????????????????????????????????????????????????????????????????????????????????????????????????????????????????????????????????????????????????????????????????????????????????????????????????????????????????????????????????????????????????????????????????????????????????????????????????????????????????

			>Hechtia_pretiosa
			????????????????????????????????????????????????????????????????????????????????????????????????????????????????????????????????????????????????????????????????????????????????????????????????????????????????????????????????????????????????????????????????????????????????????????????????????????????????????????????????????????????????????????????????????????????????

			>Hechtia_purpusii
			????????????????????????????????????????????????????????????????????????????????????????????????????????????????????????????????????????????????????????????????????????????????????????????????????????????????????????????????????????????????????????????????????????????????????????????????????????????????????????????????????????????????????????????????????????????????

			>Hechtia_schottii
			????????????????????????????????????????????????????????????????????????????????????????????????????????????????????????????????????????????????????????????????????????????????????????????????????????????????????????????????????????????????????????????????????????????????????????????????????????????????????????????????????????????????????????????????????????????????

			>Hechtia_stenopetala
			????????????????????????????????????????????????????????????????????????????????????????????????????????????????????????????????????????????????????????????????????????????????????????????????????????????????????????????????????????????????????????????????????????????????????????????????????????????????????????????????????????????????????????????????????????????????

			>Hechtia_subalata
			????????????????????????????????????????????????????????????????????????????????????????????????????????????????????????????????????????????????????????????????????????????????????????????????????????????????????????????????????????????????????????????????????????????????????????????????????????????????????????????????????????????????????????????????????????????????

			>Hechtia_texensis
			????????????????????????????????????????????????????????????????????????????????????????????????????????????????????????????????????????????????????????????????????????????????????????????????????????????????????????????????????????????????????????????????????????????????????????????????????????????????????????????????????????????????????????????????????????????????

			>Hechtia_tillandsioides
			????????????????????????????????????????????????????????????????????????????????????????????????????????????????????????????????????????????????????????????????????????????????????????????????????????????????????????????????????????????????????????????????????????????????????????????????????????????????????????????????????????????????????????????????????????????????

			>Hohenbergia_andina
			????????????????????????????????????????????????????????????????????????????????????????????????????????????????????????????????????????????????????????????????????????????????????????????????????????????????????????????????????????????????????????????????????????????????????????????????????????????????????????????????????????????????????????????????????????????????

			>Hohenbergia_belemii
			????????????????????????????????????????????????????????????????????????????????????????????????????????????????????????????????????????????????????????????????????????????????????????????????????????????????????????????????????????????????????????????????????????????????????????????????????????????????????????????????????????????????????????????????????????????????

			>Hohenbergia_brachycephala
			????????????????????????????????????????????????????????????????????????????????????????????????????????????????????????????????????????????????????????????????????????????????????????????????????????????????????????????????????????????????????????????????????????????????????????????????????????????????????????????????????????????????????????????????????????????????

			>Hohenbergia_capitata
			????????????????????????????????????????????????????????????????????????????????????????????????????????????????????????????????????????????????????????????????????????????????????????????????????????????????????????????????????????????????????????????????????????????????????????????????????????????????????????????????????????????????????????????????????????????????

			>Hohenbergia_catingae
			????????????????????????????????????????????????????????????????????????????????????????????????????????????????????????????????????????????????????????????????????????????????????????????????????????????????????????????????????????????????????????????????????????????????????????????????????????????????????????????????????????????????????????????????????????????????

			>Hohenbergia_correia-araujoi
			????????????????????????????????????????????????????????????????????????????????????????????????????????????????????????????????????????????????????????????????????????????????????????????????????????????????????????????????????????????????????????????????????????????????????????????????????????????????????????????????????????????????????????????????????????????????

			>Hohenbergia_edmundoi
			????????????????????????????????????????????????????????????????????????????????????????????????????????????????????????????????????????????????????????????????????????????????????????????????????????????????????????????????????????????????????????????????????????????????????????????????????????????????????????????????????????????????????????????????????????????????

			>Hohenbergia_humilis
			????????????????????????????????????????????????????????????????????????????????????????????????????????????????????????????????????????????????????????????????????????????????????????????????????????????????????????????????????????????????????????????????????????????????????????????????????????????????????????????????????????????????????????????????????????????????

			>Hohenbergia_loredanoana
			????????????????????????????????????????????????????????????????????????????????????????????????????????????????????????????????????????????????????????????????????????????????????????????????????????????????????????????????????????????????????????????????????????????????????????????????????????????????????????????????????????????????????????????????????????????????

			>Hohenbergia_pennae
			????????????????????????????????????????????????????????????????????????????????????????????????????????????????????????????????????????????????????????????????????????????????????????????????????????????????????????????????????????????????????????????????????????????????????????????????????????????????????????????????????????????????????????????????????????????????

			>Hohenbergia_ramageana
			????????????????????????????????????????????????????????????????????????????????????????????????????????????????????????????????????????????????????????????????????????????????????????????????????????????????????????????????????????????????????????????????????????????????????????????????????????????????????????????????????????????????????????????????????????????????

			>Hohenbergia_ridleyi
			????????????????????????????????????????????????????????????????????????????????????????????????????????????????????????????????????????????????????????????????????????????????????????????????????????????????????????????????????????????????????????????????????????????????????????????????????????????????????????????????????????????????????????????????????????????????

			>Hohenbergia_rosea
			????????????????????????????????????????????????????????????????????????????????????????????????????????????????????????????????????????????????????????????????????????????????????????????????????????????????????????????????????????????????????????????????????????????????????????????????????????????????????????????????????????????????????????????????????????????????

			>Hohenbergia_salzmannii
			????????????????????????????????????????????????????????????????????????????????????????????????????????????????????????????????????????????????????????????????????????????????????????????????????????????????????????????????????????????????????????????????????????????????????????????????????????????????????????????????????????????????????????????????????????????????

			>Hohenbergia_stellata
			????????????????????????????????????????????????????????????????????????????????????????????????????????????????????????????????????????????????????????????????????????????????????????????????????????????????????????????????????????????????????????????????????????????????????????????????????????????????????????????????????????????????????????????????????????????????

			>Hohenbergia_utriculosa
			????????????????????????????????????????????????????????????????????????????????????????????????????????????????????????????????????????????????????????????????????????????????????????????????????????????????????????????????????????????????????????????????????????????????????????????????????????????????????????????????????????????????????????????????????????????????

			>Hohenbergiopsis_guatemalensis
			TCGCAGAAAGCTCTTTCCCTGCCTACTGGAATGGGCATTCTCTGCGCGAGCCCGAAAGCTCTAGAAGCTTCCAAAACGGCCAAGTCGGTCAGGGTTTTCTTCGACTGGAATGACTACTTGAAGTTCTACAAGCTCGGAACTTACTGGCCTTAACGCCTTCCATCCAACTACTTTACGGGCTGAGAGCAGCTTTGGATCTCATCTTTGAGGAGGGGCTCGACAACGTCATCGAAAGGCATCGTCGCTTAGGAAAGGCCACAAGGTTAAGTAAAAGCTATACATCT-ARTAGCCGCTTTCGCCGTGTTGGATAGATATTTTACTTTGCACTATTGTCCTCTYGTRACAATGAGCAGACTTGCTGTCGAAG

			>Hoplocryptanthus_caracensis
			TCGCAGAAAGCTCTTTCGTTGCCTACTGGAATGGGCATTCTTTGCGCGAGCCCGAAAGCTCTAGAAGCTTCCAAAACAGCCAAGTCGGTCAGGGTTTTCTTCGACTGGAATGACTACTTGAAGTTCTACAAGCTCGGAACTTACTGGCCTTAACGCCTTCCATCCAACTACTTTATGGGCTAAGAGCAGCTCTGGATCTCATCTTTGAGGAGGGGCTCGACAGCGTCATCGAAAGGCATCGTCGCTTAGGAAAGGCCACAAGGTTAAGTAAAAGCTGTACATCT-AATAGCCGCTTGCGCCGTGTTGGATAGATATATTACTTGGCGCTATTGTCCTCTTGTAACAATGAGCAGACTTGCTGTCGAAG

			>Hoplocryptanthus_ferrarius
			????????????????????????????????????????????????????????????????????????????????????????????????????????????????????????????????????????????????????????????????????????????????????????????????????????????????????????????????????????????????????????????????????????????????????????????????????????????????????????????????????????????????????????????????????????????????

			>Hoplocryptanthus_glaziovii
			????????????????????????????????????????????????????????????????????????????????????????????????????????????????????????????????????????????????????????????????????????????????????????????????????????????????????????????????????????????????????????????????????????????????????????????????????????????????????????????????????????????????????????????????????????????????

			>Hoplocryptanthus_lavrasensis
			????????????????????????????????????????????????????????????????????????????????????????????????????????????????????????????????????????????????????????????????????????????????????????????????????????????????????????????????????????????????????????????????????????????????????????????????????????????????????????????????????????????????????????????????????????????????

			>Hoplocryptanthus_regius
			????????????????????????????????????????????????????????????????????????????????????????????????????????????????????????????????????????????????????????????????????????????????????????????????????????????????????????????????????????????????????????????????????????????????????????????????????????????????????????????????????????????????????????????????????????????????

			>Hoplocryptanthus_schwackeanus
			TCGCAGAAAGCTCTTTCGATGCCTACTGGAATGGGCATTCTTTGCGCGAGCCCGAAAGCTCTAGAAGCTTCCAAAACAGCCAAGTCGGTCAGGGTTTTCTTCGACTGGAATGACTACTTGAAGTTCTACAAGCTCGGAACTTACTGGCCTTAACGCCTTCCATCCAACTACTTTACGGGCTAAGAGCAGCTCTGGATCTCATCTTTGAGGAGGGGCTCGACAGCGTCATCGAAAGGCATCGTCGCTTAGGAAAGGCCACAAGGTTAAGTAAAAGCTGTACATCT-AATAGCCGCTTGCGCCGTGTTGGATAGATATATTACTTGGCACTATTGTCCTCTTGTAACAATGAGCAGACTTGCTGTCGAAG

			>Hoplocryptanthus_tiradentesensis
			????????????????????????????????????????????????????????????????????????????????????????????????????????????????????????????????????????????????????????????????????????????????????????????????????????????????????????????????????????????????????????????????????????????????????????????????????????????????????????????????????????????????????????????????????????????????

			>Hoplocryptanthus_vidaliorum
			TCGCAGAAAGCTCTTTCGATGCCTACTGGAATGGGCATTCTTTGCGCGAGCCCGAAAGCTCTAGAAGCTTCCAAAACAGCCAAGTCGGTCAGGGTTTTCTTCGACTGGAATGACTACTTGAAGTTCTACAAGCTCGGAACTTACTGGCCTTAACGCCTTCCATCCAACTACTTTACGGGCTAAGAGCAGCTCTGGATCTCATCTTTGAGGAGGGGCTCGACAGTGTCATCGAAAGGCATCGTCGCTTAGGAAAGGCCACAAGGTTAAGTAAAAGCTGTACATCT-AATAGCCGCTTACGCCGTGTTGGATAGATATATTACTTGGCGCTATTGTCCTCTTGTAACAATGAGCAGACTTGCTGTCGAAG

			>Jagrantia_monstrum
			????????????????????????????????????????????????????????????????????????????????????????????????????????????????????????????????????????????????????????????????????????????????????????????????????????????????????????????????????????????????????????????????????????????????????????????????????????????????????????????????????????????????????????????????????????????????

			>Josemania_asplundii
			????????????????????????????????????????????????????????????????????????????????????????????????????????????????????????????????????????????????????????????????????????????????????????????????????????????????????????????????????????????????????????????????????????????????????????????????????????????????????????????????????????????????????????????????????????????????

			>Josemania_singularis
			????????????????????????????????????????????????????????????????????????????????????????????????????????????????????????????????????????????????????????????????????????????????????????????????????????????????????????????????????????????????????????????????????????????????????????????????????????????????????????????????????????????????????????????????????????????????

			>Karawata_depressa
			TCGCAGAAAGCTCTTTCCCTGCCTACTGGAATGGGCATTCTCTGCGCGAGCCCGAAAGCTCTAGAAGCTTCCAAAACAGCCAAGTCGGTCAGGGTTTTCTTCGACTGGAATGACTACTTGAAGTTCTACAAGCTCGGAACTTACTGGCCTTAACGCCTTCCATCCAACTACTTTACGGGCTGAGAGCAGCTTTGGATCTCATCTTTGAGGAGGGGCTCGACAATGTCATYGAAAGGCATCGTCGCTTAGGAAAGGCCACAAGGTCAAGTAAAAGYTATACATCT-AATAGCCGCTTTYGCCGTGTTGGATAGATATATWACTTGGCACTATTGTYCTCTTGTAACAATGAGCAGACTYGCTGTCGAAG

			>Karawata_gustavoi
			TCGCAGAAAGCTCTTTCCCTGCCTACTGGAATGGGCATTCTCTGCGCGAGCCCGAAAGCTCTAGAAGCTTCCAAAACAGCCAAGTCGGTCAGGGTTTTCTTCGACTGGAATGACTACTTGAAGTTCTACAAGCTCGGAACTTACTGGCCTTAACGCCTTCCATCCAACTACTTTACGGGCTAAGAGCGGCTTTGGATCTCATCTTTGAGGAGGGGCTCGACAATGTCATCGAAAGGCATCGTCGCTTAGGAAAGGCCACAAGGTGAAGTAAAAGCTATACATCA-AATAGCCGCTTTCGCCGTGTTGGATAGATATATTACTTGGCACTATTGTCCTCTTGTAACAATGAGCAGACTTGCTGTCGAAG

			>Karawata_hostilis
			????????????????????????????????????????????????????????????????????????????????????????????????????????????????????????????????????????????????????????????????????????????????????????????????????????????????????????????????????????????????????????????????????????????????????????????????????????????????????????????????????????????????????????????????????????????????

			>Karawata_multiflora
			????????????????????????????????????????????????????????????????????????????????????????????????????????????????????????????????????????????????????????????????????????????????????????????????????????????????????????????????????????????????????????????????????????????????????????????????????????????????????????????????????????????????????????????????????????????????

			>Karawata_nigribracteata
			????????????????????????????????????????????????????????????????????????????????????????????????????????????????????????????????????????????????????????????????????????????????????????????????????????????????????????????????????????????????????????????????????????????????????????????????????????????????????????????????????????????????????????????????????????????????

			>Karawata_saxicola
			TCGCAGAAAGCTCTTTCCCTGCCTACTGGAATGGGCATTCTCTGCGCGAGCCCGAAAGCTCTAGAAGCTTCCAAAACAGCCAAGTCGGTCAGGGTTTTCTTCGACTGGAATGACTACTTGAAGTTCTACAAGCTCGGAACTTACTGGCCTTAACGCCTTCCATCCAACTACTTTACGGGCTGAGAGCAGCTTTGGATCTCATCTTTGAGGAGGGGCTCGACAATGTCATCGAAAGGCATCGTCGCTTAGGAAAGGCCACAAGGTKAAGTAAAAGCTATACATCA-AATAGCCGCTTTCGCCGTGTTGGATAGATATATTACTTGGCACTATTGTCCTCTTGTAACAATGAGCAGACTTGCTGTCGAAG

			>Lapanthus_duartei
			TCGCAGAAAGCTCTTTCGATGCCTACTGGAATGGGCATTCTTTGCGCGAGCCCGAAAGCTCTAGAAGCTTCCAAAACAGCCAAGTCGGTCAGGGTTTTCTTCGACTGGAATGACTACTTGAAGTTCTACAAGCTCGGAACTTACTGGCCTTAACGCCTTCCATCCAACTACTTTACGGGCTAAGAGCAGCTTTGGATCTCATCTTTGAGGAGGGGCTCGACAATGTCGTCGAAAGGCATCGTCGCTTAGGAAAGGCCACAAGGTCAAGTAAAAGCTATACATCTATATAGCCGCTTTCGCCGTGTTGGATAGATATATTACTTGGCGCTATTGGCCTCTTGTAACAATGRGCAGACTTGCTGTCGAAG

			>Lapanthus_itambensis
			TCGCAGAAAGCTCTTTCGATGCCTACTGGAATGGGCATTCTTTGCGCGAGCCCGAAAGCTCTAGAAGCTTCCAAAACAGCCAAGTCGGTCAGGGTTTTCTTCGACTGGAATRACTACTTGAAGTTCTACAAGCTCGGAACTTACTGGCCTTAACGCCTTCCATCCAACTACTTTACGGGCTAAGAGCAGCTTTGGATCTCATCTTTGAGGAGGGGCTCGACAATGTCGTCGAAAGGCATCGTCGCTTAGGAAAGGCCACAAGGTCAAGTAAAAGCTATACATCTATATAGCCGCTTTCGCCGTGTYGGATAGATATATTACTTGGCGCTATTGGCCTCTTGTAACAATGAGCAGACTTGCTGTCGAAG

			>Lemeltonia_dodsonii
			TCGCAGAAAGCACTCTCCCTGCCTACTGGAATGGGCATTGTTTGCGCGAGCCCGAAAGCTCTAGAAGCTTCCAAAAATGCCAAGTCGGTCAGAGTTTTCTTCGACTGGAATGACTACTTGAAGTTCTACAAGCTTGGAACTTACTGGCCGTAACGCCTTCCATCCAACTACTTTACGGGCTAAGAGCAGCTTTGGATCTCATCTTTGAGGAGGGGCTCGAAAATGTCATCGAAAGGCATCGTCGTTTAGGAAAGGCCACGAGG-TAAGTA-AAGCT--------AAATAGCCGATTTTACAGTATAGGAAAGATATATAATTAGGCATTATTGTCCTCCTGTAACAATGAGCAGACTTGCTGTCGAAG

			>Lemeltonia_monadelpha
			????????????????????????????????????????????????????????????????????????????????????????????????????????????????????????????????????????????????????????????????????????????????????????????????????????????????????????????????????????????????????????????????????????????????????????????????????????????????????????????????????????????????????????????????????????????????

			>Lemeltonia_narthecioides
			????????????????????????????????????????????????????????????????????????????????????????????????????????????????????????????????????????????????????????????????????????????????????????????????????????????????????????????????????????????????????????????????????????????????????????????????????????????????????????????????????????????????????????????????????????????????

			>Lemeltonia_triglochinoides
			????????????????????????????????????????????????????????????????????????????????????????????????????????????????????????????????????????????????????????????????????????????????????????????????????????????????????????????????????????????????????????????????????????????????????????????????????????????????????????????????????????????????????????????????????????????????

			>Lindmania_guianensis
			????????????????????????????????????????????????????????????????????????????????????????????????????????????????????????????????????????????????????????????????????????????????????????????????????????????????????????????????????????????????????????????????????????????????????????????????????????????????????????????????????????????????????????????????????????????????

			>Lindmania_longipes
			????????????????????????????????????????????????????????????????????????????????????????????????????????????????????????????????????????????????????????????????????????????????????????????????????????????????????????????????????????????????????????????????????????????????????????????????????????????????????????????????????????????????????????????????????????????????

			>Lutheria_glutinosa
			????????????????????????????????????????????????????????????????????????????????????????????????????????????????????????????????????????????????????????????????????????????????????????????????????????????????????????????????????????????????????????????????????????????????????????????????????????????????????????????????????????????????????????????????????????????????

			>Lutheria_splendens
			TCGCAGAAAGCACTTTCCCTGCCTACTGGAATGGGCATTGTTTGCGCGAGCCCGAAAGCTCTAGAAGCTTCCAAAAATGCCAAGTCGGTCAGAGTTTTCTTCGACTGGAATGACTACTTGAAGTTCTACAAGCTTGGAACTTACTGGCCGTAACGCCTTCCATCCAACTACTTTACGGGCTAAGAGCAGCTTTGGATCTCATCTTTGAGGAGGGGCTCGAAAATGTCATCGAAAGGCATCGCCGTTTAGGAAAGGCCACGAGG-TAAGTAAAAGCT--------AAATAGCCGATTTTAGCA--TAGGAAAGATATATAATTACTCATTATTGTCCTCTTGTAACAATGAGCAGACTTGCTGTCGAAG

			>Lymania_alvimii
			????????????????????????????????????????????????????????????????????????????????????????????????????????????????????????????????????????????????????????????????????????????????????????????????????????????????????????????????????????????????????????????????????????????????????????????????????????????????????????????????????????????????????????????????????????????????

			>Lymania_azurea
			????????????????????????????????????????????????????????????????????????????????????????????????????????????????????????????????????????????????????????????????????????????????????????????????????????????????????????????????????????????????????????????????????????????????????????????????????????????????????????????????????????????????????????????????????????????????

			>Lymania_brachycaulis
			????????????????????????????????????????????????????????????????????????????????????????????????????????????????????????????????????????????????????????????????????????????????????????????????????????????????????????????????????????????????????????????????????????????????????????????????????????????????????????????????????????????????????????????????????????????????

			>Lymania_corallina
			????????????????????????????????????????????????????????????????????????????????????????????????????????????????????????????????????????????????????????????????????????????????????????????????????????????????????????????????????????????????????????????????????????????????????????????????????????????????????????????????????????????????????????????????????????????????

			>Lymania_globosa
			????????????????????????????????????????????????????????????????????????????????????????????????????????????????????????????????????????????????????????????????????????????????????????????????????????????????????????????????????????????????????????????????????????????????????????????????????????????????????????????????????????????????????????????????????????????????

			>Lymania_languida
			????????????????????????????????????????????????????????????????????????????????????????????????????????????????????????????????????????????????????????????????????????????????????????????????????????????????????????????????????????????????????????????????????????????????????????????????????????????????????????????????????????????????????????????????????????????????

			>Lymania_smithii
			????????????????????????????????????????????????????????????????????????????????????????????????????????????????????????????????????????????????????????????????????????????????????????????????????????????????????????????????????????????????????????????????????????????????????????????????????????????????????????????????????????????????????????????????????????????????

			>Lymania_spiculata
			????????????????????????????????????????????????????????????????????????????????????????????????????????????????????????????????????????????????????????????????????????????????????????????????????????????????????????????????????????????????????????????????????????????????????????????????????????????????????????????????????????????????????????????????????????????????

			>Mezobromelia_bicolor
			????????????????????????????????????????????????????????????????????????????????????????????????????????????????????????????????????????????????????????????????????????????????????????????????????????????????????????????????????????????????????????????????????????????????????????????????????????????????????????????????????????????????????????????????????????????????

			>Mezobromelia_capituligera
			????????????????????????????????????????????????????????????????????????????????????????????????????????????????????????????????????????????????????????????????????????????????????????????????????????????????????????????????????????????????????????????????????????????????????????????????????????????????????????????????????????????????????????????????????????????????

			>Mezobromelia_pleiosticha
			????????????????????????????????????????????????????????????????????????????????????????????????????????????????????????????????????????????????????????????????????????????????????????????????????????????????????????????????????????????????????????????????????????????????????????????????????????????????????????????????????????????????????????????????????????????????

			>Navia_barbellata
			????????????????????????????????????????????????????????????????????????????????????????????????????????????????????????????????????????????????????????????????????????????????????????????????????????????????????????????????????????????????????????????????????????????????????????????????????????????????????????????????????????????????????????????????????????????????

			>Navia_igneosicola
			????????????????????????????????????????????????????????????????????????????????????????????????????????????????????????????????????????????????????????????????????????????????????????????????????????????????????????????????????????????????????????????????????????????????????????????????????????????????????????????????????????????????????????????????????????????????

			>Navia_phelpsiae
			????????????????????????????????????????????????????????????????????????????????????????????????????????????????????????????????????????????????????????????????????????????????????????????????????????????????????????????????????????????????????????????????????????????????????????????????????????????????????????????????????????????????????????????????????????????????

			>Navia_saxicola
			TCGCAGAAAGCTCTTTCCCTGCCTACTGGAATGGGCATTGTTTGTGCGAGCCCGAAAGCTCTAGAAGCTTCCAAGACGGCCAAGTCGGTCAGGGTTTTCTTCGACTGGAATGACTACTTGAAGTTCTACAAGCTTGGAACTTACTGGCCTTAACGCCTTCCATCCAACTACTTTATGGGCTAAGAGCAGCTTTGGATCTCATCTTTGAGGAGGGGCTCGACAATGTCATCGAAAGGCATCGTCGTTTAGGAAAGGCCACAAGGT-AAGTAAAAGCTTTACASCT-ARTAGCCGATTTTACAGTGTTGGAAAGATATATTACTTGGCATTATAGTCCTCTCGTAACAATGAGCAGACTTGCTGTCGAAG

			>Neoglaziovia_burle-marxii
			TCGCAGAAAGCTCTTTCGATGCCTACTGGAATGGGCATTCTTTGCGCGAGCCCGAAAGCTCTAGAAGCTTCCAAAACAGCCAAGTCGGTCAGGGTTTTCTTCGACTGGAAYGACTACTTGAAGTTCTACAAGCTCGGAACTTACTGGCCTTAACGCCTTCCATCCAACTACTTTACGGGCTGAGAGCAGCTTTGGATCTCATCTTTGAGGAGGGGCTCGACAATGTCATCGAAAGGCATCGTCGCTTAGGAAAGGCCACAAGGTTAAGYAAAAGCTATACATCT-AATAGCCGCTTTCGCCGTGTCGGATAGATATATTACTTGGCACTATTGTCCTCTTGTAACAATGAGCAGACTTGCTGTCGAAG

			>Neoglaziovia_variegata
			TCGCAGAAAGCTCTTTCGATGCCTACTGGAATGGGCATTCTTTGCGCGAGCCCGAAAGCTCTAGAAGCTTCCAAAACAGCCAAGTCGGTCAGGGTTTTCTTCGACTGGAAYGACTACTTGAAGTTCTACAAGCTCGGAACTTAYTGGCCTTAACGCCTTCCATCCAACTACTTTACGGGCTGAGAGCAGCTTTGGATCTCATCTTYGAGGAGGGGCTCGACAATGTCATCGAAAGGCATCGTCGCTTAGGAAAGGCCACAAGGTTAAGTAAAAGCTATACATCT-AATAGCCGCTTTCGCCGTGTYGGATAGATATATTACTTGGCACTATTGTCCTCTTGTAACAATGAGCAGACTTGCTGTCGAAG

			>Neoregelia_binotii
			????????????????????????????????????????????????????????????????????????????????????????????????????????????????????????????????????????????????????????????????????????????????????????????????????????????????????????????????????????????????????????????????????????????????????????????????????????????????????????????????????????????????????????????????????????????????

			>Neoregelia_bragarum
			TCGCAGAAAGCTCTTTCCCTGCCTACTGGAATGGGGATTCTCTGCGCGAGCCCGAAAGCTCTAGAAGCTTCCAAAACAGCCAAGTCGGTCAGGGTTTTCTTCGACTGGAATGACTACTTGAAGTTCTACAAGCTCGGAACTTACTGGCCTTAACGCCTTCCATCCAACTACTTTACGGGCTGAGAGCAGCTTTGGACCTCATCTTTGAGGAGGGGCTCGACAATGTCATCGAAAGGCATCGTCGCTTAGGAAAGGCCACAAGGTTAAGTAAAAGCTATACATCTAAATAGCCGCTTTTGCCGTGTTGGATAGATATATTACTTTGCACTGTTGTCCTCTTGTAACAATGGGCAGACTTGCCGTMGAAG

			>Neoregelia_capixaba
			????????????????????????????????????????????????????????????????????????????????????????????????????????????????????????????????????????????????????????????????????????????????????????????????????????????????????????????????????????????????????????????????????????????????????????????????????????????????????????????????????????????????????????????????????????????????

			>Neoregelia_carolinae
			????????????????????????????????????????????????????????????????????????????????????????????????????????????????????????????????????????????????????????????????????????????????????????????????????????????????????????????????????????????????????????????????????????????????????????????????????????????????????????????????????????????????????????????????????????????????

			>Neoregelia_compacta
			TCGCAGAAAGCTCTTTCCCTGCCTACTGGAATGGGGATTCTCTGCGCGAGCCCGAAAGCTCTAGAAGCTTCCAAAACAGCCAAGTCGGTCAGGGTTTTCTTCGACTGGAATGACTACTTGAAGTTCTACAAGCTCGGAACTTACTGGCCTTAACGCCTTCCATCCAACTACTTTACGGGCTGAGAGCAGCTTTGGACCTCATCTTTGAGGAGGGGCTCGATAATGTCATCGAAAGGCACCGTCGCTTAGGAAAGGCCACAAGGTTAAATAAAAGCTATACATTTAAATAGCCGCTTTCGCCGTGTTGGATAGATATATTACTTTGCACTACTGTCCTCTTGTAACAATGGGCAGACTTGCCGTCGAAG

			>Neoregelia_concentrica
			TCGCAGAAAGCTCTTTCCCTGCCTACTGGAATGGGGATTCTCTGCGCGAGCCCGAAAGCTCTAGAAGCTTCCAAAACAGCCAAGTCGGTCAGGGTTTTCTTCGACTGGAATGACTACTTGAAGTTCTACAAGCTCGGAACTTACTGGCCTTAACGCCTTCCATCCAACTACTTTACGGGCTGAGAGCAGCTTTGGACCTCATCTTTGAGGAGGGGCTCGACAATGTCATCGAAAGGCATCGTCGCTTAGGAAAGGCCACAAGGTTAAGTAAAAGCCATACATCTAAATASCCGCTTTCGCCGTGTTGGATAGATATATTACTTTGCACTRYTGTCCTCTTGTAACAATGGGCAGACTTGCCGTCGAAG

			>Neoregelia_cruenta
			TCGCAGAAAGCTCTTTCCCTGCCTACTGGAATGGGGATTCTCTGCGCGAGCCCGAAAGCTCTAGAAGCTTCCAAAACAGCCAAGTCGGTCAGGGTTTTCTTCGACTGGAATGACTACTTGAAGTTCTACAAGCTCGGAACTTACTGGCCTTAACGCCTTCCATCCAACTACTTTACGGGCTGAGAGCAGCTTTGGACCTCATCTTTGAGGAGGGGCTCGACAATGTCATCGAAAGGCATCGTCGCTTAGGAAAGGCCACAAGGTTAAGYAAAAGCTATACATCTAAAWWMCCGCTTTCRCCGTGTTGGATAGATATATTACTTTGCACTGTTGTCCTCTTGTAACAATGGGCAGGCTTGCCGTCGAAG

			>Neoregelia_dungsiana
			TCGCAGAAAGCTCTTTCCCTGCCTACTGGAATGGGGATTCTCTGCGCGAGCCCGAAAGCTCTAGAAGCTTCCAAAACAGCCAAGTCGGTCAGGGTTTTCTTCGACTGGAATGACTACTTGAAGTTCTACAARCTCGGAACTTACTGGCCTTAACGCCTTCCATCCAACTACTTTACGGGCTGAGAGCAGCTTTGGACCTCATCTTTGAGGAGGGGCTCGACAATGTCATCGAAAGGCATCGTCGCTTAGGAAAGGCCACAAGGTTAAGTAAAAGCCACACATCTAAATASSCGCTTTCRCCGTGTTGGATAGATATATTACTTKGCACTGYTGTCMTCTTGTAACAATGGGCAGACTTGCCGTCGAAG

			>Neoregelia_eleutheropetala
			????????????????????????????????????????????????????????????????????????????????????????????????????????????????????????????????????????????????????????????????????????????????????????????????????????????????????????????????????????????????????????????????????????????????????????????????????????????????????????????????????????????????????????????????????????????????

			>Neoregelia_lactea
			TCGCAGAAAGCTCTTTCCCTGCCTACTGGAATGGGGATTCTCTGCGCGAGCCCGAAAGCTCTAGAAGCTTCCAAAACAGCCGAGTCGGTCAGGGTTTTCTTCGACTGGAATGACTACTTGAAGTTCTACAAGCTCGGAACTTACTGGCCTTAACGCCTTCCATCCAACTACTTTACGGGCTGAGAGCAGCTTTGGACCTCATCTTTGAGGAGGGGCTCGACAATGTCATCGAAAGGCATCGTCGCTTAGGAAAGGCCACAAGGTTGACTAAAAGCTATACATCTAAATAGCCGCTTTCGCCGTGTTGGATAGTTATATTACTCTGCACTACTGTCCTCTTGTAACAATGGGCAGACTTGCCGTCGAAG

			>Neoregelia_laevis
			????????????????????????????????????????????????????????????????????????????????????????????????????????????????????????????????????????????????????????????????????????????????????????????????????????????????????????????????????????????????????????????????????????????????????????????????????????????????????????????????????????????????????????????????????????????????

			>Neoregelia_leucophoea
			TCGCAGAAAGCTCTTTCCCTGCCTACTGGAATGGGGATTCTCTGCGCGAGCCCGAAAGCTCTAGAAGCTTCCAAAACWGCCAAGTCGGTCAGGGTTTTCTTCGACTGGAATGACTACTTGAAGTTCTACAAGCTCGGAACTTACTGGCCTTAACGCCTTCCATCCAACTACTTTACGGGCTGAGAGCAGCTTTGGACCTCATCTTYGAGGAGGGGCTCGACAATGTCATCGAAAGGCATCGTCGCTTAGGAAAGGCCACAAGGTTAAGTAAAAGCTATACATCTAAATRGCCGCTTTCGCCGTGTTGGATAGATATATTACTTTGCACTGTTGTCCTCTTGTAACAATGGGCAGACTTGCCGTCGAAG

			>Neoregelia_longisepala
			TCACAGAAAGCTCTTTCCCTGCCTACTGGAATGGGCATTCTCTGCGCGAGCCCGAAAGCTCTAGAAGCTTCCAAAACAGCCAAGTCGGTCAGGGTTTTCTTCGACTGGAATGACTACTTGAAGTTCTACAAGCTCGGAACTTACTGGCCTTAACGCCKTCCATCCAACTGCTTTACGGGCTGAGAGCAGCTTTGGATCTCATCTTTGAGGAGGGGCTCGACAATGTCATCGAAAGGCACCGYCGCTTAGGAAAGGCCACAAGGTTAAGTAAAAGCTATACATYT-AATAGCCGCCTTCGCCGTGTYGGATAGATATATTACTTGGCACTGTTGTTCTCTTGTAACAATGRGCAGACTTGCTGTCGAAG

			>Neoregelia_lymaniana
			TCGCAGAAAGCTCTTTCCCTGCCTACTGGAATGGGGATTCTCTGCGCGAGCCCGAAAGCTCTAGAAGCTTCCAAAACAGCCGAGTCGGTCAGGGTTTTCTTCGACTGGAATGACTACTTGAAGTTCTACAAGCTCGGAACTTACTGGCCTTAACGCCTTCCATCCAACTACTTTACGGGCTGAGAGCAGCTTTGGACCTCATCTTTGAGGAGGGGCTCGACAATGTCATCGAAAGGCATCGTCGCTTAGGAAAGGCCACAAGGTTGACTAAAAGCTATACATCTAAATAGCCGCTTTCGCCGTGTTGGATAGTTATATTACTCTGCACTACTGTCCTCTTGTAACAATGGGCAGACTTGCCGTCGAAG

			>Neoregelia_mucugensis
			????????????????????????????????????????????????????????????????????????????????????????????????????????????????????????????????????????????????????????????????????????????????????????????????????????????????????????????????????????????????????????????????????????????????????????????????????????????????????????????????????????????????????????????????????????????????

			>Neoregelia_myrmecophila
			????????????????????????????????????????????????????????????????????????????????????????????????????????????????????????????????????????????????????????????????????????????????????????????????????????????????????????????????????????????????????????????????????????????????????????????????????????????????????????????????????????????????????????????????????????????????

			>Neoregelia_nevaresii
			TCGCAGAAAGCTCTTTCCCTGCCTACTGGAATGGGGATTCTCTGCGCGAGCCCGAAAGCTCTAGAAGCTTCCAAAACAGCCAAGTCGGTCAGGGTTTTCTTCGACTGGAATGACTACTTGAAGTTCTACAAGCTCGGAACTTACTGGCCTTAACGCCTTCCATCCAACTACTTTACGGGCTGAGAGCAGCTTTGGACCTCATCTTTGAGGAGGGGCTCGACAACGTCATCGAAAGGCATCGTCGCTTAGGCAAGGCCACAAGGTTAAGTAAAAGCTATACATCTAAATAGCCGTTTTCGCCGTGTTGGATAGATATATTACTTTGCACTACTGTCCTCTTGTAACAATGGGCAGGCTTGCCGTCGAAG

			>Neoregelia_nivea
			TCGCAGAAAGCTCTTTCCCTGCCTACTGGAATGGGGATTCTCTGCGCGAGCCCGAAAGCTCTAGAAGCTTCCAAAACAGCCAAGTCGGTCAGGGTTTTCTTCSACTGGAATGACTACTTGAAGTTCTACAAGCTCGGAACTTACTGGCCTTAACGCCTTCCATCCAACTACTTTACGGGCTGAGAGCAGCTTTGGACCTCATCTTTGAGGAGGGGCTCGACAATGTCATCGAAAGGCATCGTCGCTTAGGAAAGGCCACAAGGTTAAGTAAAAGCTATACATCTAAATAGCCGCTTTCGCCGTGTTGGATAGATATACTACTTTGCACTACTGTCCTCTTGTAACAATGGGCAGACTTGCCGTCGAAG

			>Neoregelia_pascoaliana
			TCGCAGAAAGCTCTTTCCCTGCCTACTGGAATGGGGATTCTCTGCGCGAGCCCGAAAGCTCTAGAAGCTTCCAAAACAGCCAAGTCGGTCAGGGTTTTCTTCGACTGGAATGACTACTTGAAGTTCTACAAGCTCGGAACTTACTGGCCTTAACGCCTTCCATCCAACTACTTTACGGGCTGAGAGCAGCTTTGGACCTCATCTTTGAGGAGGGGCTCGACAATGTCATCGAAAGGCATCGTCGCTTAGGAAAGGCCACAAGGTTAACTAAAAGCTATACATCTAAATAGCCGTTTTCGCCATGTTGGATAGATATATTACTTTGCACTACTGTCCTCTTGTAACAATGGGCAGACTTGCCGTCGAAG

			>Neoregelia_pauciflora
			????????????????????????????????????????????????????????????????????????????????????????????????????????????????????????????????????????????????????????????????????????????????????????????????????????????????????????????????????????????????????????????????????????????????????????????????????????????????????????????????????????????????????????????????????????????????

			>Neoregelia_pineliana
			TCGCAGAAAGCTCTTTCCCTGCCTACTGGAATGGGGATTCTCTGCGCGAGCCCGAAAGCTCTAGAAGCTTCCAAAACAGCCAAGTCGGTCAGGGTTTTCTTCGACTGGAATGACTACTTGAAGTTCTACAAGCTCGGAACTTACTGGCCTTAACGCCTTCCATCCAACTACTTTACGGGCTGAGAGCAGCTTTGGACCTCATCTTTGAGGAGGGGCTCGACAATGTCATCGAAAGGCATCGTCGCTTAGGAAAGGCCACAAGGTTAAGTAAAAGCTATACATCTAAATAGCCGCTTTCGCCGTGTTGGATAGATATATTACTTKGCACTACTGTCCTCTTGTAACAATGGGCAGRCTTGCCGTCGAAG

			>Neoregelia_retrorsa
			TCGCAGAAAGCTCTTTCCCTGCCTACTGGAATGGGGATTCTCTGCGCGAGCCCGAAAGCTCTAGAAGCTTCCAAAACAGCCGAGTCGGTCAGGGTTTTCTTCGACTGGAATGACTACTTGAAGTTCTACAAGCTCGGAACTTACTGGCCTTAACGCCTTCCATCCAACTACTTTACGGGCTGAGAGCAGCTTTGGACCTCATCTTTGAGGAGGGGCTCGACAATGTCATCGAAAGGCATCGTCGCTTAGGAAAGGCCACAAGGTTGAGTAAAAGCTATACATCTAAATACCCGCTTTCGCCGTGTTGGATAGATATATTACTCTGCACTACTGTCCTCTTGTAACAATGGGCAGACTTGCCGTCGAAG

			>Neoregelia_richteri
			????????????????????????????????????????????????????????????????????????????????????????????????????????????????????????????????????????????????????????????????????????????????????????????????????????????????????????????????????????????????????????????????????????????????????????????????????????????????????????????????????????????????????????????????????????????????

			>Neoregelia_rothinessa
			????????????????????????????????????????????????????????????????????????????????????????????????????????????????????????????????????????????????????????????????????????????????????????????????????????????????????????????????????????????????????????????????????????????????????????????????????????????????????????????????????????????????????????????????????????????????

			>Neoregelia_rubrovittata
			TCGCAGAAAGCTCTTTCCCTGCCTACTGGAATGGGGATTCTCTGCGCGAGCCCGAAAGCTCTAGAAGCTTCCAAAACAGCCAAGTCGGTCAGGGTTTTCTTCGACTGGAATGACTACTTGAAGTTCTACAAGCTCGGAACTTACTGGCCTTAACGCCTTCCATCCAACTACTTTACGGGCTGAGAGCAGCTTTGGACCTCATCTTTGAGGAGGGGCTCGACAACGTCATCGAAAGGCATCGTCGCTTAGGAAAGGCCACAAGGTTAAGCAAAAGCTATACATCTAAATAGCCGCTTTCGCCGTGYTGGATAGATATATTACTYKRCACTATTGTCCTCTTGTAACAATGGGCAGGCTTGCCGTCGAAG

			>Neoregelia_tenebrosa
			TCGCAGAAAGCTCTTTCCCTGCCTACTGGAATGGGGATTCTCTGCGCGAGCCCGAAAGCTCTAGAAGCTTCCAAAACAGCCAAGTCGGTCAGGGTTTTCTTCGACTGGAATGACTACTTGAAGTTCTACAAGCTCGGAACTTACTGGCCTTAACGCCTTCCATCCAACTACTTTACGGGCTGAGAGCAGCTTTGGACCTCATCTTTGAGGAGGGGCTCGACAATGTCATCGAAAGGCATCGTCGCTTAGGAAAGGCCACAAGGTTAAGTAAAAGCTATACATCTAAATAGCCGCTTTCGCCGTGTTGGATAGATATATTACTTGGCACTACTGTCCTCTTGTAACAATGGGCAGGCTTGCCGTCGAAG

			>Neoregelia_viridovinosa
			TCGCAGAAAGCTCTTTCCCTGCCTACTGGAATGGGGATTCTCTGCGCGAGCCCGAAAGCTCTAGAAGCTTCCAAAACAGCCGAGTCGGTCAGGGTTTTCTTCGACTGGAATGACTACTTGAAGTTCTACAAGCTCGGAACTTACTGGCCTTAACGCCTTCCATCCAACTACTTTACGGGCTGAGAGCAGCTTTGGACCTCATCTTTGAGGAGGGGCTCGACAATGTCATCGAAAGGCATCGTCGCTTAGGAAAGGCCACAAGGTTGAGCAAAAGCTATACATCTAAATAGCCGCTTTCGCCGTGTTGGATAGATATATTACTCTGCACTACTGTCCTCTTGTAACAATGGGCAGACTTGCCGTCGAAG

			>Neoregelia_wilsoniana
			TCGCAGAAAGCTCTTTCCCTGCCTACYGGAATGGGGATTCTCTGCGCGAGCCCGAAAGCTCTAGAAGCTTCCAAAACAGCCAAGTCGGTCAGGGTTTTCTTCGACTGGAATGACTACTTGAAGTTCTACAAGCTCGGAACTTACTGGCCTTAACGCCTTCCATCCAACTACTTTACGGGCTGAGAGCAGCTTTGGACCTCATCTTTGAGGAGGGGCTCGACAATGTCATCGAAAGGCATCGTCGCTTAGGAAAGGCCACAAGGTTAAGTAAAAGCYATACATCTAAATASCCGCYTTCGYCGTGTTGGATAGATATATTWCTTKGCACTACTGTCCTCTTGTAACAATGGGCAGACTTGCCGTCGAAG

			>Neoregelia_zonata
			????????????????????????????????????????????????????????????????????????????????????????????????????????????????????????????????????????????????????????????????????????????????????????????????????????????????????????????????????????????????????????????????????????????????????????????????????????????????????????????????????????????????????????????????????????????????

			>Nidularium_altimontanum
			????????????????????????????????????????????????????????????????????????????????????????????????????????????????????????????????????????????????????????????????????????????????????????????????????????????????????????????????????????????????????????????????????????????????????????????????????????????????????????????????????????????????????????????????????????????????

			>Nidularium_amazonicum
			TCGCAGAAAGCTCTTTCCCTGCCTACTGGAATGGGGATTCTCTGCGCGAGCCCGAAAGCTCTAGAAGCTTCCAAAACAGCCAAGTCGGTCAGGGTTTTCTTCGACTGGAATGACTACTTGAAGTTCTACAAGCTCGGAACTTACTGGCCTTAACGCCTTCCATCCAACTACTTTACGGGCTGAGAGCAGCTTTGGACCTCATCTTTGAGGAGGGGCTCGACAATGTCATCGAAAGGCATCGTCGCTTAGGAAAGGCCACAAGGTTAAGTAAAAGCTATACATCTAAATAGCCGCTTTCGCCGTGTTGGATAGATATATTACTTTGCACTACTGTCCTCTTGTAACAATGGGCAGACTTGCCGTCGAAG

			>Nidularium_angustibracteatum
			????????????????????????????????????????????????????????????????????????????????????????????????????????????????????????????????????????????????????????????????????????????????????????????????????????????????????????????????????????????????????????????????????????????????????????????????????????????????????????????????????????????????????????????????????????????????

			>Nidularium_campoalegrense
			TCGCAGAAAGCTCTTTCCCTGCCTACTGGAATGGGGATTCTCTGCGCGAGCCCGAAAGCTCTAGAAGCTTCCAAAACAGCCAAGTCGGTCAGGGTTTTCTTCGACTGGAATGACTACTTGAAGTTCTACAAGCTCGGAACTTACTGGCCTTAACGCCTTCCATCCAACTACTTTACGGGCTGAGAGCAGCTTTGGACCTCATCTTTGAGGAGGGGCTCGACAATGTCATCGAAAGGCACCGTCGCTTAGGAAAGGCCACAAGGTTAAGTAAAAGCTATACATCTAAATAGCCGCTTTCGCCGTGTTGGATAGATATATTACTTTGCACTACTGTCCTCTTGTAACAATGGGCAGACTTGCCGTCGAAG

			>Nidularium_camposportoi
			????????????????????????????????????????????????????????????????????????????????????????????????????????????????????????????????????????????????????????????????????????????????????????????????????????????????????????????????????????????????????????????????????????????????????????????????????????????????????????????????????????????????????????????????????????????????

			>Nidularium_fulgens
			TCGCAGAAAGCTCTTTCCCTGCCTACTGGAATGGGGATTCTCTGCGCGAGCCCGAAAGCTCTAGAAGCTTCCAAAACAGCCAAGTCGGTCAGGGTTTTCTTCGACTGGAATGACTACTTGAAGTTCTACAAGCTCGGAACTTACTGGCCTTAACGCCTTCCATCCAACTACTTTACGGGCTGAGAGCAGCTTTGGACCTCATCTTTGAGGAGGGGCTCGACAATGTCATCGAAAGGCATCGTCGCTTAGGAAAGGCCACAAGGTTAACTAAAAGCTATACATCTAAATAGCCGCTTTCGCCGTGTTGGATAGATATATTACTCTGCACTAYTGTCCTCTTGTAACAATGGGCAGACTTGCCGTCGAAG

			>Nidularium_innocentii
			TCGCAGAARGCTCTTTCCCTGCCTACTGGAATGGGGATTCTCTGCGCGAGCCCGAAAGCTCTAGAAGCTTCCAAAACAGCCAAGTCGGTCAGGGTTTTCTTCGACTGGAATGACTACTTGAAGTTCTACAAGCTCGGAACTTACTGGCCTTAACGCCTTCCATCCAACTACTTTACGGGCTGAGAGCAGCTTTGGACCTCATCTTTGAGGAGGGGCTCGACAATGTCATCGAAAGGCATCGTCGCTTAGGAAAGGCCACAAGGTTAAGTAAAAGCTATACATCTAAATAGCCGCTTTCGCCGTGTTGGATAGATATWTTACTTGGCACTAYTGTCCTCTTGTAACAATGGGCAGACTTGCCGTCGAAG

			>Nidularium_krisgreeniae
			????????????????????????????????????????????????????????????????????????????????????????????????????????????????????????????????????????????????????????????????????????????????????????????????????????????????????????????????????????????????????????????????????????????????????????????????????????????????????????????????????????????????????????????????????????????????

			>Nidularium_marigoi
			TCGCAGAAAGCTCTTTCCCTGCCGACTGGAATGGGGATTCTCTGCGCGAGCCCGAAAGCTCTAGAAGCTTCCAAAACAGCCAAGTCGGTCAGGGTTTTCTTCGACTGGAACGACTACTTGAAGTTCTACAAGCTCGGAACTTACTGGCCTTAACGCCTTCCATCCAACTACTTTACGGGCTGAGAGCAGCTTTGGACCTCATCTTTGAGGAGGGGCTCGACAATGTCATCGAAAGGCATCATCGCTTCGGAAAGGCCACAAGGTTAAGTAAAAGCTATACATCTAAATAGCCGCTTTCGCCGTGTTGGATAGATATATTACTTTGCACTACTGTCCTCTTGTAACAATGGGCAGACTTGCCGTCGAAG

			>Nidularium_picinguabense
			????????????????????????????????????????????????????????????????????????????????????????????????????????????????????????????????????????????????????????????????????????????????????????????????????????????????????????????????????????????????????????????????????????????????????????????????????????????????????????????????????????????????????????????????????????????????

			>Nidularium_procerum
			????????????????????????????????????????????????????????????????????????????????????????????????????????????????????????????????????????????????????????????????????????????????????????????????????????????????????????????????????????????????????????????????????????????????????????????????????????????????????????????????????????????????????????????????????????????????

			>Nidularium_purpureum
			TCGCAGAAAGCTCTCTCCCTGCCTACTGGAATGGGGATTCTCTGCGCGAGCCCGAAAGCTCTAGATGCTTCCAAAACAGCCAAGTCGGTCAGGGTTTTCTTCGACTGGAATGACTACTTGAAGTTCTACAAGCTCGGAACTTACTGGCCTTAACGCCTTCCATCCAACTACTTTACGGGCTGAGAGCAGCTTTGGACCTCATCTTTGAGGAGGGGCTCGACAATGTCATCGAAAGGCATCGTCGCTTAGGAAAGGCCACAAGGTTAAGTAAAAGCTATACATCTAAATAGCCGCTTTCGCCGTGTTGGATAGATATATTACTCTGCACTATTGTCCTCTTGTAACAATGGGCAGACTTGCCGTCGAAG

			>Ochagavia_andina
			????????????????????????????????????????????????????????????????????????????????????????????????????????????????????????????????????????????????????????????????????????????????????????????????????????????????????????????????????????????????????????????????????????????????????????????????????????????????????????????????????????????????????????????????????????????????

			>Ochagavia_carnea
			????????????????????????????????????????????????????????????????????????????????????????????????????????????????????????????????????????????????????????????????????????????????????????????????????????????????????????????????????????????????????????????????????????????????????????????????????????????????????????????????????????????????????????????????????????????????

			>Ochagavia_elegans
			????????????????????????????????????????????????????????????????????????????????????????????????????????????????????????????????????????????????????????????????????????????????????????????????????????????????????????????????????????????????????????????????????????????????????????????????????????????????????????????????????????????????????????????????????????????????

			>Ochagavia_litoralis
			????????????????????????????????????????????????????????????????????????????????????????????????????????????????????????????????????????????????????????????????????????????????????????????????????????????????????????????????????????????????????????????????????????????????????????????????????????????????????????????????????????????????????????????????????????????????

			>Orthophytum_alvimii
			????????????????????????????????????????????????????????????????????????????????????????????????????????????????????????????????????????????????????????????????????????????????????????????????????????????????????????????????????????????????????????????????????????????????????????????????????????????????????????????????????????????????????????????????????????????????

			>Orthophytum_argentum
			????????????????????????????????????????????????????????????????????????????????????????????????????????????????????????????????????????????????????????????????????????????????????????????????????????????????????????????????????????????????????????????????????????????????????????????????????????????????????????????????????????????????????????????????????????????????

			>Orthophytum_benzingii
			????????????????????????????????????????????????????????????????????????????????????????????????????????????????????????????????????????????????????????????????????????????????????????????????????????????????????????????????????????????????????????????????????????????????????????????????????????????????????????????????????????????????????????????????????????????????

			>Orthophytum_boudetianum
			????????????????????????????????????????????????????????????????????????????????????????????????????????????????????????????????????????????????????????????????????????????????????????????????????????????????????????????????????????????????????????????????????????????????????????????????????????????????????????????????????????????????????????????????????????????????

			>Orthophytum_braunii
			????????????????????????????????????????????????????????????????????????????????????????????????????????????????????????????????????????????????????????????????????????????????????????????????????????????????????????????????????????????????????????????????????????????????????????????????????????????????????????????????????????????????????????????????????????????????

			>Orthophytum_compactum
			????????????????????????????????????????????????????????????????????????????????????????????????????????????????????????????????????????????????????????????????????????????????????????????????????????????????????????????????????????????????????????????????????????????????????????????????????????????????????????????????????????????????????????????????????????????????

			>Orthophytum_conquistense
			????????????????????????????????????????????????????????????????????????????????????????????????????????????????????????????????????????????????????????????????????????????????????????????????????????????????????????????????????????????????????????????????????????????????????????????????????????????????????????????????????????????????????????????????????????????????

			>Orthophytum_diamantinense
			????????????????????????????????????????????????????????????????????????????????????????????????????????????????????????????????????????????????????????????????????????????????????????????????????????????????????????????????????????????????????????????????????????????????????????????????????????????????????????????????????????????????????????????????????????????????

			>Orthophytum_disjunctum
			????????????????????????????????????????????????????????????????????????????????????????????????????????????????????????????????????????????????????????????????????????????????????????????????????????????????????????????????????????????????????????????????????????????????????????????????????????????????????????????????????????????????????????????????????????????????

			>Orthophytum_duartei
			????????????????????????????????????????????????????????????????????????????????????????????????????????????????????????????????????????????????????????????????????????????????????????????????????????????????????????????????????????????????????????????????????????????????????????????????????????????????????????????????????????????????????????????????????????????????

			>Orthophytum_estevesii
			????????????????????????????????????????????????????????????????????????????????????????????????????????????????????????????????????????????????????????????????????????????????????????????????????????????????????????????????????????????????????????????????????????????????????????????????????????????????????????????????????????????????????????????????????????????????

			>Orthophytum_falconii
			????????????????????????????????????????????????????????????????????????????????????????????????????????????????????????????????????????????????????????????????????????????????????????????????????????????????????????????????????????????????????????????????????????????????????????????????????????????????????????????????????????????????????????????????????????????????

			>Orthophytum_foliosum
			????????????????????????????????????????????????????????????????????????????????????????????????????????????????????????????????????????????????????????????????????????????????????????????????????????????????????????????????????????????????????????????????????????????????????????????????????????????????????????????????????????????????????????????????????????????????

			>Orthophytum_fosterianum
			????????????????????????????????????????????????????????????????????????????????????????????????????????????????????????????????????????????????????????????????????????????????????????????????????????????????????????????????????????????????????????????????????????????????????????????????????????????????????????????????????????????????????????????????????????????????

			>Orthophytum_glabrum
			????????????????????????????????????????????????????????????????????????????????????????????????????????????????????????????????????????????????????????????????????????????????????????????????????????????????????????????????????????????????????????????????????????????????????????????????????????????????????????????????????????????????????????????????????????????????

			>Orthophytum_graomogolense
			????????????????????????????????????????????????????????????????????????????????????????????????????????????????????????????????????????????????????????????????????????????????????????????????????????????????????????????????????????????????????????????????????????????????????????????????????????????????????????????????????????????????????????????????????????????????

			>Orthophytum_grossiorum
			????????????????????????????????????????????????????????????????????????????????????????????????????????????????????????????????????????????????????????????????????????????????????????????????????????????????????????????????????????????????????????????????????????????????????????????????????????????????????????????????????????????????????????????????????????????????

			>Orthophytum_gurkenii
			????????????????????????????????????????????????????????????????????????????????????????????????????????????????????????????????????????????????????????????????????????????????????????????????????????????????????????????????????????????????????????????????????????????????????????????????????????????????????????????????????????????????????????????????????????????????

			>Orthophytum_harleyi
			????????????????????????????????????????????????????????????????????????????????????????????????????????????????????????????????????????????????????????????????????????????????????????????????????????????????????????????????????????????????????????????????????????????????????????????????????????????????????????????????????????????????????????????????????????????????

			>Orthophytum_horridum
			????????????????????????????????????????????????????????????????????????????????????????????????????????????????????????????????????????????????????????????????????????????????????????????????????????????????????????????????????????????????????????????????????????????????????????????????????????????????????????????????????????????????????????????????????????????????

			>Orthophytum_lanuginosum
			????????????????????????????????????????????????????????????????????????????????????????????????????????????????????????????????????????????????????????????????????????????????????????????????????????????????????????????????????????????????????????????????????????????????????????????????????????????????????????????????????????????????????????????????????????????????

			>Orthophytum_lemei
			????????????????????????????????????????????????????????????????????????????????????????????????????????????????????????????????????????????????????????????????????????????????????????????????????????????????????????????????????????????????????????????????????????????????????????????????????????????????????????????????????????????????????????????????????????????????

			>Orthophytum_leprosum
			????????????????????????????????????????????????????????????????????????????????????????????????????????????????????????????????????????????????????????????????????????????????????????????????????????????????????????????????????????????????????????????????????????????????????????????????????????????????????????????????????????????????????????????????????????????????

			>Orthophytum_lucidum
			????????????????????????????????????????????????????????????????????????????????????????????????????????????????????????????????????????????????????????????????????????????????????????????????????????????????????????????????????????????????????????????????????????????????????????????????????????????????????????????????????????????????????????????????????????????????

			>Orthophytum_macroflorum
			????????????????????????????????????????????????????????????????????????????????????????????????????????????????????????????????????????????????????????????????????????????????????????????????????????????????????????????????????????????????????????????????????????????????????????????????????????????????????????????????????????????????????????????????????????????????

			>Orthophytum_magalhaesii
			????????????????????????????????????????????????????????????????????????????????????????????????????????????????????????????????????????????????????????????????????????????????????????????????????????????????????????????????????????????????????????????????????????????????????????????????????????????????????????????????????????????????????????????????????????????????

			>Orthophytum_maracasense
			????????????????????????????????????????????????????????????????????????????????????????????????????????????????????????????????????????????????????????????????????????????????????????????????????????????????????????????????????????????????????????????????????????????????????????????????????????????????????????????????????????????????????????????????????????????????

			>Orthophytum_mello-barretoi
			????????????????????????????????????????????????????????????????????????????????????????????????????????????????????????????????????????????????????????????????????????????????????????????????????????????????????????????????????????????????????????????????????????????????????????????????????????????????????????????????????????????????????????????????????????????????

			>Orthophytum_pseudostoloniferum
			????????????????????????????????????????????????????????????????????????????????????????????????????????????????????????????????????????????????????????????????????????????????????????????????????????????????????????????????????????????????????????????????????????????????????????????????????????????????????????????????????????????????????????????????????????????????

			>Orthophytum_pseudovagans
			????????????????????????????????????????????????????????????????????????????????????????????????????????????????????????????????????????????????????????????????????????????????????????????????????????????????????????????????????????????????????????????????????????????????????????????????????????????????????????????????????????????????????????????????????????????????

			>Orthophytum_riocontense
			????????????????????????????????????????????????????????????????????????????????????????????????????????????????????????????????????????????????????????????????????????????????????????????????????????????????????????????????????????????????????????????????????????????????????????????????????????????????????????????????????????????????????????????????????????????????

			>Orthophytum_roseolilacinum
			TCGCAGAAAGCTCTTTCGATGCCTACCGGAATGGGCATTCTTTGCGCGAGCCCGAAAGCTCTAGAAGCTTCCAAAACAGCAAGGTCGGTCAGGGTTTTCTTCGACTGGAATGACTACTTGAAGTTCTACAAGCTCGGAACTTACTGGCCTTAACGCCGTCCATCCAACTACTTTACGGGCTGAGAGCAGCTTTGGATCTCATCTTTGAGGAGGGGCTCGACAACGTCATCGAAAGGCATCGCCGCTTAGGAAAGGCCACAAGGT-AACCAAAAGCTATACATCTATATAGCCGCTTTCGCCGTGTCGGATAGATATGTTACTCGGCGCTATATTCCTCTTGTAACAATGAGCAGACTTGCTGTCGAAG

			>Orthophytum_rubiginosum
			????????????????????????????????????????????????????????????????????????????????????????????????????????????????????????????????????????????????????????????????????????????????????????????????????????????????????????????????????????????????????????????????????????????????????????????????????????????????????????????????????????????????????????????????????????????????

			>Orthophytum_santaritense
			????????????????????????????????????????????????????????????????????????????????????????????????????????????????????????????????????????????????????????????????????????????????????????????????????????????????????????????????????????????????????????????????????????????????????????????????????????????????????????????????????????????????????????????????????????????????

			>Orthophytum_saxicola
			TCGCAGAAAGCTCTCTCGATGCCTACGGGAATGGGCATTCTTTGCGCGAGCCCGAAAGCTCTAGAAGCCTCCAAAACAGCGAGGTCGGTCAGGGTTTTCTTCGACTGGAATGACTACTTGAAGTTCTACAAGCTCGGAACTTACTGGCCTTAACGCCGTCCATCCAACTGCTTTACGGGCTGAGAGCAGCTCTGGATCTCATCTTCGAGGAGGGGCTCGACAACGTCATCGAAAGGCATCGCCGCTTAGGAAAGGCCACAAGGTCAACTGGAAATTGAACCTCC-AACAACCGTTGTTAGAGTATCGAGAAGCTATTTTCATATCTACCACTATTCTCTCGCCGCAACGAGCAGACTCGCCGTCGAAG

			>Orthophytum_schulzianum
			????????????????????????????????????????????????????????????????????????????????????????????????????????????????????????????????????????????????????????????????????????????????????????????????????????????????????????????????????????????????????????????????????????????????????????????????????????????????????????????????????????????????????????????????????????????????

			>Orthophytum_sucrei
			????????????????????????????????????????????????????????????????????????????????????????????????????????????????????????????????????????????????????????????????????????????????????????????????????????????????????????????????????????????????????????????????????????????????????????????????????????????????????????????????????????????????????????????????????????????????

			>Orthophytum_toscanoi
			????????????????????????????????????????????????????????????????????????????????????????????????????????????????????????????????????????????????????????????????????????????????????????????????????????????????????????????????????????????????????????????????????????????????????????????????????????????????????????????????????????????????????????????????????????????????

			>Orthophytum_vagans
			????????????????????????????????????????????????????????????????????????????????????????????????????????????????????????????????????????????????????????????????????????????????????????????????????????????????????????????????????????????????????????????????????????????????????????????????????????????????????????????????????????????????????????????????????????????????

			>Orthophytum_vasconcelosianum
			????????????????????????????????????????????????????????????????????????????????????????????????????????????????????????????????????????????????????????????????????????????????????????????????????????????????????????????????????????????????????????????????????????????????????????????????????????????????????????????????????????????????????????????????????????????????

			>Orthophytum_zanonii
			????????????????????????????????????????????????????????????????????????????????????????????????????????????????????????????????????????????????????????????????????????????????????????????????????????????????????????????????????????????????????????????????????????????????????????????????????????????????????????????????????????????????????????????????????????????????

			>Pitcairnia_abundans
			????????????????????????????????????????????????????????????????????????????????????????????????????????????????????????????????????????????????????????????????????????????????????????????????????????????????????????????????????????????????????????????????????????????????????????????????????????????????????????????????????????????????????????????????????????????????

			>Pitcairnia_albiflos
			????????????????????????????????????????????????????????????????????????????????????????????????????????????????????????????????????????????????????????????????????????????????????????????????????????????????????????????????????????????????????????????????????????????????????????????????????????????????????????????????????????????????????????????????????????????????

			>Pitcairnia_atrorubens
			??????????CTCTTTCCCTGCCTACTGGAATGGGCATTGTTTGCGCGAGCCCGAAAGCTCTAGAAGCTTCCAAAACAGCCAAGTCGGTCAGGGTTTTCTTCGACTGGAATGACTACTTGAAGTTCTACAAGATCGGAACTTACTGGCCTTAACGCCGTCCATCCAACTACTTTACGGGCTAAGAGCAGCTTTGGATCTCATCTTTGAGGAGGGGCTCGACAATGTCATCGAAAGGCACCGTCGTTTAGGAAAGGCCACAAGGT-AAGTAAAAGCTATACATCT-AATAGTTTTTTTTACAGTGTCGGATAGATATATTACTTGGTATTATTGACCTCTTCTAACAATGAGCAGGCT???????????

			>Pitcairnia_beachiae
			????????????????????????????????????????????????????????????????????????????????????????????????????????????????????????????????????????????????????????????????????????????????????????????????????????????????????????????????????????????????????????????????????????????????????????????????????????????????????????????????????????????????????????????????????????????????

			>Pitcairnia_breedlovei
			????????????????????????????????????????????????????????????????????????????????????????????????????????????????????????????????????????????????????????????????????????????????????????????????????????????????????????????????????????????????????????????????????????????????????????????????????????????????????????????????????????????????????????????????????????????????

			>Pitcairnia_burle-marxii
			????????????????????????????????????????????????????????????????????????????????????????????????????????????????????????????????????????????????????????????????????????????????????????????????????????????????????????????????????????????????????????????????????????????????????????????????????????????????????????????????????????????????????????????????????????????????

			>Pitcairnia_carinata
			????????????????????????????????????????????????????????????????????????????????????????????????????????????????????????????????????????????????????????????????????????????????????????????????????????????????????????????????????????????????????????????????????????????????????????????????????????????????????????????????????????????????????????????????????????????????

			>Pitcairnia_chiapensis
			????????????????????????????????????????????????????????????????????????????????????????????????????????????????????????????????????????????????????????????????????????????????????????????????????????????????????????????????????????????????????????????????????????????????????????????????????????????????????????????????????????????????????????????????????????????????

			>Pitcairnia_corallina
			TCGCAGAAAGCTCTTTCCCTGCCTACTGGAATGGGCATTGTTTGCGCGAGCCCGAAAGCTCTAGAGGCTTCCAAAACAGCCAAGTCGGTCAGGGTTTTCTTCGACTGGAATGACTACTTGAAGTTCTACAAGCTCGGAACTTACTGGCCTTAACGCCTTCCATCCAACTACTTTATGGGCTGAGAGCAGCTTTGGATCTCATCTTTGAGGAGGGGCTCGACAATGTCATCGAAAGGCACCGTCGTTTAGGAAAGGCCACAAGGT-GAGTAAAAGCTATACATCT-AATAGCTGTTTTTACAGTGTCGGATATATATATTACTTTGTATTATTGACCTCTTCTAACAATGAGCAGGCTTGCTGTCGAAG

			>Pitcairnia_elvirae
			TCGCAGAAAGCTCTTTCCCTGCCTACTGGAATGGGCATTGTTTGCGCGAGCCCGAAAGCTCTAGAAGCTTCCAAAACAGCCAAGTCGGTCAGGGTTTTCTTCGACTGGAATGACTACTTGAAGTTCTACAAGCTCGGAACTTACTGGCCTTAACTCCTTCCATCCAACTACTTTATGGGCTAAGAGCAGCTTTGGATCTCATCTTTGAGGAGGGGCTCGACAATGTCATCGAAAGGCATCGTCGTTTAGGAAAGGCCACAAGGT-AAGTAAAAGCTATACATCT-AATAGCTGTTTTTACAGTGTCGGATATATATATTACTTGGTATTATTGACCTCTTCTAACAATGA??????????????????

			>Pitcairnia_feliciana
			????????????????????????????????????????????????????????????????????????????????????????????????????????????????????????????????????????????????????????????????????????????????????????????????????????????????????????????????????????????????????????????????????????????????????????????????????????????????????????????????????????????????????????????????????????????????

			>Pitcairnia_flammea
			????????????????????????????????????????????????????????????????????????????????????????????????????????????????????????????????????????????????????????????????????????????????????????????????????????????????????????????????????????????????????????????????????????????????????????????????????????????????????????????????????????????????????????????????????????????????

			>Pitcairnia_flammea_var._glabrior
			????????????????????????????????????????????????????????????????????????????????????????????????????????????????????????????????????????????????????????????????????????????????????????????????????????????????????????????????????????????????????????????????????????????????????????????????????????????????????????????????????????????????????????????????????????????????

			>Pitcairnia_fuertesii
			????????????????????????????????????????????????????????????????????????????????????????????????????????????????????????????????????????????????????????????????????????????????????????????????????????????????????????????????????????????????????????????????????????????????????????????????????????????????????????????????????????????????????????????????????????????????

			>Pitcairnia_heterophylla
			????????????????????????????????????????????????????????????????????????????????????????????????????????????????????????????????????????????????????????????????????????????????????????????????????????????????????????????????????????????????????????????????????????????????????????????????????????????????????????????????????????????????????????????????????????????????

			>Pitcairnia_hirtzii
			????????????????????????????????????????????????????????????????????????????????????????????????????????????????????????????????????????????????????????????????????????????????????????????????????????????????????????????????????????????????????????????????????????????????????????????????????????????????????????????????????????????????????????????????????????????????

			>Pitcairnia_integrifolia
			????????????????????????????????????????????????????????????????????????????????????????????????????????????????????????????????????????????????????????????????????????????????????????????????????????????????????????????????????????????????????????????????????????????????????????????????????????????????????????????????????????????????????????????????????????????????

			>Pitcairnia_jimenezii
			????????????????????????????????????????????????????????????????????????????????????????????????????????????????????????????????????????????????????????????????????????????????????????????????????????????????????????????????????????????????????????????????????????????????????????????????????????????????????????????????????????????????????????????????????????????????

			>Pitcairnia_lignosa
			????????????????????????????????????????????????????????????????????????????????????????????????????????????????????????????????????????????????????????????????????????????????????????????????????????????????????????????????????????????????????????????????????????????????????????????????????????????????????????????????????????????????????????????????????????????????

			>Pitcairnia_loki-schmidtiae
			????????????????????????????????????????????????????????????????????????????????????????????????????????????????????????????????????????????????????????????????????????????????????????????????????????????????????????????????????????????????????????????????????????????????????????????????????????????????????????????????????????????????????????????????????????????????

			>Pitcairnia_macrochlamys
			????????????????????????????????????????????????????????????????????????????????????????????????????????????????????????????????????????????????????????????????????????????????????????????????????????????????????????????????????????????????????????????????????????????????????????????????????????????????????????????????????????????????????????????????????????????????

			>Pitcairnia_maidifolia
			????????????????????????????????????????????????????????????????????????????????????????????????????????????????????????????????????????????????????????????????????????????????????????????????????????????????????????????????????????????????????????????????????????????????????????????????????????????????????????????????????????????????????????????????????????????????

			>Pitcairnia_nigra
			TCGCAGAAAGCTCTTTCCCTGCCTACTGGAATGGGCATTGTTTGCGCGAGCCCGAAAGCTCTAGAAGCTTCCAAAACAGCCAAGTCGGTCAGGGTTTTCTTCGACTGGAATGACTACTTGAAGTTCTACAAGCTCGGAACTTACTGGCCTTAACGCCGTCCATCCAACTACTTTATGGGCTAAGAGCAGCTTTGGACCTCATCTTTGAGGAGGGGCTCGACAATGTCATCGAAAGGCACCATCGTTTAGGAAAGGCCACAAGGT-AAGTAAAAGCTATACATCT-AATAGTTTTTTTTACAGCGTCAGATAGATATATTACTTGGTATTATTGACCTCTTCTAACAATGA??????????????????

			>Pitcairnia_orchidifolia
			????????????????????????????????????????????????????????????????????????????????????????????????????????????????????????????????????????????????????????????????????????????????????????????????????????????????????????????????????????????????????????????????????????????????????????????????????????????????????????????????????????????????????????????????????????????????

			>Pitcairnia_piepenbringii
			????????????????????????????????????????????????????????????????????????????????????????????????????????????????????????????????????????????????????????????????????????????????????????????????????????????????????????????????????????????????????????????????????????????????????????????????????????????????????????????????????????????????????????????????????????????????

			>Pitcairnia_poortmanii
			????????????????????????????????????????????????????????????????????????????????????????????????????????????????????????????????????????????????????????????????????????????????????????????????????????????????????????????????????????????????????????????????????????????????????????????????????????????????????????????????????????????????????????????????????????????????

			>Pitcairnia_prolifera
			TCGCAGAAAGCTCTTTCCCTGCCTACTGGAATGGGCATTGTTTGCGCGAGCCCGAAAGCTCTAGAAGCTTCCAAAACAGCCAAGTCGGTCAGGGTTTTCTTCGACTGGAATGACTACTTGAAGTTCTACAAGCTCGGAACTTACTGGCCTTAACGCCTTCCATCCAACTACTTTATGGGCTAAGAGCAGCTTTGGATCTCATCTTTGAGGAGGGGCTCGACAATGTCATCGAAAGGCATCATCGTTTAGGAAAGGCCACAAGGT-AAGTA----CTATTCATCT-AATAGCCGGTTTTACAGTGTCGGATATATATATTACTTGGTATTATTGTCCTCTCGTAACAATGAGCAGGCTTGCTGTCGAAG

			>Pitcairnia_pseudoundulata
			????????????????????????????????????????????????????????????????????????????????????????????????????????????????????????????????????????????????????????????????????????????????????????????????????????????????????????????????????????????????????????????????????????????????????????????????????????????????????????????????????????????????????????????????????????????????

			>Pitcairnia_pungens
			????????????????????????????????????????????????????????????????????????????????????????????????????????????????????????????????????????????????????????????????????????????????????????????????????????????????????????????????????????????????????????????????????????????????????????????????????????????????????????????????????????????????????????????????????????????????

			>Pitcairnia_punicea
			????????????????????????????????????????????????????????????????????????????????????????????????????????????????????????????????????????????????????????????????????????????????????????????????????????????????????????????????????????????????????????????????????????????????????????????????????????????????????????????????????????????????????????????????????????????????

			>Pitcairnia_rectiflora
			????????????????????????????????????????????????????????????????????????????????????????????????????????????????????????????????????????????????????????????????????????????????????????????????????????????????????????????????????????????????????????????????????????????????????????????????????????????????????????????????????????????????????????????????????????????????

			>Pitcairnia_recurvata
			???CAGAAAGCTCTTTCCCTGCCTACTGGAATGGGCATTGTTTGCGCGAGCCCGAAAGCTCTAGAAGCTTCCAAAACAGCCAAGTCGGTCAGGGTTTTCTTCGACTGGAATGACTACTTGAAGTTCTACAAGCTCGGAACTTACTGGCCTTAACTCCTTCCATCCAACTACTTTATGGGCTAAGAGCAGCTTTGGATCTCATCTTTGAGGAGGGGCTCGACAATGTCATCGAAAGGCATCGTCGTTTAGGAAAGGCCACAAGGT-AAGTAAAAGCTATACATCT-AATAGCTGTTTTTACAGTGTCGGATATATATATAACTTTGTATTATTGACCTCTTCTAACAATG???????????????????

			>Pitcairnia_reflexiflora
			TCGCAGAAAGCTCTTTCCCTGCCTACTGGAATGGGCATTGTTTGCGCGAGCCCGAAAGCTCTAGAGGCTTCCAAAACAGCCAAGTCGGTCAGGGTTTTCTTCGACTGGAATGACTACTTGAAGTTCTACAAGCTCGGAACTTACTGGCCTTAACGCCGTCCATCCAACTACTTTATGGGCTAAGAGCAGCTTTGGATCTCATCTTTGAGGAGGGGCTCGACAATGTCATCGAAAGGCATCGTCGTTTAGGAAAGGCCACAAGGT-AAGTAAAAGATATATATCT-AATAGCTGTTTTTACAGTGTCGGATATATATATTACTTGGTATTATTGACCTCTTCTAACAATGAGCAGGCTTGCTGTCG???

			>Pitcairnia_riparia
			?????????????????????????????????????TTGTTTGCGCGAGCCCGAAAGCTCTAGAGGCTTCCAAAACAGCCAAGTCGGTCAGGGTTTTCTTCGACTGGAATGACTACTTGAAGTTCTACAAGCTCGGAACTTACTGGCCTTAACGCCTTCCATCCAACTTCTTTATGGGCTGAGAGCAGCTTTGGATCTCATCTTTGAGGAGGGGCTCGACAATGTCATCGAAAGGCATCATCGTTTAGGAAAGGCCACAAGGT-AAGTAAAAGCTATTCAGCT-AATAGCCGGTTTTACAGTGTCGGATAGATATATTACTTGGTATTATTGTCCTCTCGTAAC???????????????????????

			>Pitcairnia_roseoalba
			???CAGAAAGCTCTTTCCCTGCCTACTGGAATGGGCATTGTTTGCGCGAGCCCGAAAGCTCTAGAAGCTTCCAAAACAGCCAAGTCGGTCAGGGTTTTCTTCGACTGGAATGACTACTTGAAGTTCTACAAGCTCGGAACTTACTGGCCTTAACGCCTTCCATCCAACTTCTTTATGGGCTAAGAGCAGCTTTGGATCTCATCTTCGAGGAGGGGCTCGACAATGTCATCGAAAGGCATCATCGTTTAGGAAAGGCCACAAGGT-AAGTAAAAGCTATACATCT-AATAGTTGTTTTTACAGCGTCAGATAGATATATTACTTGGTATTATTGACCTCTTGTAACAATGAGCAGGCTTGCTGTCGAAG

			>Pitcairnia_rubronigriflora
			????????????????????????????????????????????????????????????????????????????????????????????????????????????????????????????????????????????????????????????????????????????????????????????????????????????????????????????????????????????????????????????????????????????????????????????????????????????????????????????????????????????????????????????????????????????????

			>Pitcairnia_sanguinea
			????????????????????????????????????????????????????????????????????????????????????????????????????????????????????????????????????????????????????????????????????????????????????????????????????????????????????????????????????????????????????????????????????????????????????????????????????????????????????????????????????????????????????????????????????????????????

			>Pitcairnia_smithiorum
			????????????????????????????????????????????????????????????????????????????????????????????????????????????????????????????????????????????????????????????????????????????????????????????????????????????????????????????????????????????????????????????????????????????????????????????????????????????????????????????????????????????????????????????????????????????????

			>Pitcairnia_spicata
			????????????????????????????????????????????????????????????????????????????????????????????????????????????????????????????????????????????????????????????????????????????????????????????????????????????????????????????????????????????????????????????????????????????????????????????????????????????????????????????????????????????????????????????????????????????????

			>Pitcairnia_sprucei
			????????????????????????????????????????????????????????????????????????????????????????????????????????????????????????????????????????????????????????????????????????????????????????????????????????????????????????????????????????????????????????????????????????????????????????????????????????????????????????????????????????????????????????????????????????????????

			>Pitcairnia_squarrosa
			????????????????????????????????????????????????????????????????????????????????????????????????????????????????????????????????????????????????????????????????????????????????????????????????????????????????????????????????????????????????????????????????????????????????????????????????????????????????????????????????????????????????????????????????????????????????

			>Pitcairnia_staminea
			???????????TCTTTCCCTGCCTACTGGAATGGGCATTGTTTGCGCGAGCCCGAAAGCTCTAGAAGCTTCCAAAACAGCCAAGTCGGTCAGGGTTTTCTTCGACTGGAATGACTACTTGAAGTTCTACAAGCTCGGAACTTACTGGCCTTAACGCCTTCCATCCAACTTCTTTATGGGCTAAGAGCAGCTTTGGATCTCATCTTTGAGGAGGGGCTCGACAATGTCATCGAAAGGCATCATCGTTTAGGAAAGGCCACAAGGT-AAGTAAAAGCTATTCAGCT-AATAGCCGGTTTTACAGTGTCGGATAGATATGTTACTTGGTATTATTTTCCTCTTGTAACAATGA??????????????????

			>Pitcairnia_sulphurea
			????????????????????????????????????????????????????????????????????????????????????????????????????????????????????????????????????????????????????????????????????????????????????????????????????????????????????????????????????????????????????????????????????????????????????????????????????????????????????????????????????????????????????????????????????????????????

			>Pitcairnia_utcubambensis
			????????????????????????????????????????????????????????????????????????????????????????????????????????????????????????????????????????????????????????????????????????????????????????????????????????????????????????????????????????????????????????????????????????????????????????????????????????????????????????????????????????????????????????????????????????????????

			>Pitcairnia_villetaensis
			????????????????????????????????????????????????????????????????????????????????????????????????????????????????????????????????????????????????????????????????????????????????????????????????????????????????????????????????????????????????????????????????????????????????????????????????????????????????????????????????????????????????????????????????????????????????

			>Pitcairnia_wendlandii
			TCGCAGAAAGCTCTTTCCCTGCCTACTGGAATGGGCATTGTTTGCGCGAGCCCGAAAGCTCTAGAAGCTTCCAAAACAGCCAAGTCGGTCAGGGTTTTCTTCGACTGGAATGACTACTTGAAGTTCTACAAGATCGGAACTTACTGGCCTTAACGCCGTCCATCCAACTACTTTACGGGCTAAGAGCAGCTTTGGATCTCATCTTTGAGGAGGGGCTCGACAATGTCATCGAAAGGCACCGTCGTTTAGGAAAGGCCACAAGGT-AAGTAAAAGCTATACATCT-AATAGTTTTTTTTACAGTGTCGGATAGATATATTACTTGGTATTATTGACCTCTTCTAACAATGA??????????????????

			>Pitcairnia_xanthocalyx
			????????????????????????????????????????????????????????????????????????????????????????????????????????????????????????????????????????????????????????????????????????????????????????????????????????????????????????????????????????????????????????????????????????????????????????????????????????????????????????????????????????????????????????????????????????????????

			>Pitcairnia_yaupibajaensis
			????????????????????????????????????????????????????????????????????????????????????????????????????????????????????????????????????????????????????????????????????????????????????????????????????????????????????????????????????????????????????????????????????????????????????????????????????????????????????????????????????????????????????????????????????????????????

			>Portea_alatisepala
			????????????????????????????????????????????????????????????????????????????????????????????????????????????????????????????????????????????????????????????????????????????????????????????????????????????????????????????????????????????????????????????????????????????????????????????????????????????????????????????????????????????????????????????????????????????????

			>Portea_filifera
			????????????????????????????????????????????????????????????????????????????????????????????????????????????????????????????????????????????????????????????????????????????????????????????????????????????????????????????????????????????????????????????????????????????????????????????????????????????????????????????????????????????????????????????????????????????????

			>Portea_grandiflora
			????????????????????????????????????????????????????????????????????????????????????????????????????????????????????????????????????????????????????????????????????????????????????????????????????????????????????????????????????????????????????????????????????????????????????????????????????????????????????????????????????????????????????????????????????????????????

			>Portea_kermesina
			????????????????????????????????????????????????????????????????????????????????????????????????????????????????????????????????????????????????????????????????????????????????????????????????????????????????????????????????????????????????????????????????????????????????????????????????????????????????????????????????????????????????????????????????????????????????

			>Portea_nana
			????????????????????????????????????????????????????????????????????????????????????????????????????????????????????????????????????????????????????????????????????????????????????????????????????????????????????????????????????????????????????????????????????????????????????????????????????????????????????????????????????????????????????????????????????????????????

			>Portea_petropolitana
			????????????????????????????????????????????????????????????????????????????????????????????????????????????????????????????????????????????????????????????????????????????????????????????????????????????????????????????????????????????????????????????????????????????????????????????????????????????????????????????????????????????????????????????????????????????????

			>Portea_petropolitana_var._extensa
			????????????????????????????????????????????????????????????????????????????????????????????????????????????????????????????????????????????????????????????????????????????????????????????????????????????????????????????????????????????????????????????????????????????????????????????????????????????????????????????????????????????????????????????????????????????????

			>Portea_petropolitana_var._noettigii
			????????????????????????????????????????????????????????????????????????????????????????????????????????????????????????????????????????????????????????????????????????????????????????????????????????????????????????????????????????????????????????????????????????????????????????????????????????????????????????????????????????????????????????????????????????????????

			>Pseudalcantarea_grandis
			????????????????????????????????????????????????????????????????????????????????????????????????????????????????????????????????????????????????????????????????????????????????????????????????????????????????????????????????????????????????????????????????????????????????????????????????????????????????????????????????????????????????????????????????????????????????

			>Pseudalcantarea_macropetala
			????????????????????????????????????????????????????????????????????????????????????????????????????????????????????????????????????????????????????????????????????????????????????????????????????????????????????????????????????????????????????????????????????????????????????????????????????????????????????????????????????????????????????????????????????????????????

			>Pseudalcantarea_viridiflora
			????????????????????????????????????????????????????????????????????????????????????????????????????????????????????????????????????????????????????????????????????????????????????????????????????????????????????????????????????????????????????????????????????????????????????????????????????????????????????????????????????????????????????????????????????????????????

			>Pseudaraeococcus_parviflorus
			????????????????????????????????????????????????????????????????????????????????????????????????????????????????????????????????????????????????????????????????????????????????????????????????????????????????????????????????????????????????????????????????????????????????????????????????????????????????????????????????????????????????????????????????????????????????

			>Puya_aequatorialis
			????????????????????????????????????????????????????????????????????????????????????????????????????????????????????????????????????????????????????????????????????????????????????????????????????????????????????????????????????????????????????????????????????????????????????????????????????????????????????????????????????????????????????????????????????????????????

			>Puya_alpestris
			??GCAGAAAGCTCTTTCCCTGCCTACTGGAATGGGCATTGTTTGCGCGAGCCCGAAAGCTCTAGAAGCTTCCAAAACAGCCAAGTCGGTCAGGGTTTTCTTCGACTGGAATGACTACTTGAAGTTCTACAAGCTTGGAACTTACTGGCCTTAACGCCTTCCATCCAACTACTTTACGGGCTAAGAGCAGCTTTGGATCTCATCTTTGAGGAGGGGCTCGACAATGTCATCGAAAGGCATCGTCGTTTAGGAAAGGCCACAAGGT-AAGTAAAAGCTATACATCT-GACAGCCGCTWTTACAGTGTCGGATAGATATATTACTTGGTATTATTGTCCTCTTGTAACAATGAGCAGACTTGCTGTCGAAG

			>Puya_angusta
			????????????????????????????????????????????????????????????????????????????????????????????????????????????????????????????????????????????????????????????????????????????????????????????????????????????????????????????????????????????????????????????????????????????????????????????????????????????????????????????????????????????????????????????????????????????????

			>Puya_bicolor
			????????????????????????????????????????????????????????????????????????????????????????????????????????????????????????????????????????????????????????????????????????????????????????????????????????????????????????????????????????????????????????????????????????????????????????????????????????????????????????????????????????????????????????????????????????????????

			>Puya_boliviensis
			??GCAGAAAGCTCTTTCCCTGCCTACTGGAATGGGCATTGTTTGCGCGAGCCCGAAAGCTCTAGAAGCTTCCAAAACAGCCAAGTCGGTCAGGGTTTTYTTSGACKGGAATGACTACTTGAAGTTCTACAAGCTTGGAACTTACTGGCCTTAACGCCTTCCATCCAACTACTTTACGGGCTAAGAGCAGCTTTGGATCTCATCTTTGAGGARGGGCTCGACAATGTCATCGAAAGGCATCGTCGTTTAGGAAAGGCCACAAGGT-AAGTAAAAGCTATACATCT-GACAGCCGCTWTTACAGTGTCGGATAGATATATTACTTGGTATTATTGTCCTCTTGTAACAATGAGCAGACTTGCTGTCGAAG

			>Puya_castellanosii
			????????????????????????????????????????????????????????????????????????????????????????????????????????????????????????????????????????????????????????????????????????????????????????????????????????????????????????????????????????????????????????????????????????????????????????????????????????????????????????????????????????????????????????????????????????????????

			>Puya_chilensis
			TCGCAGAAAGCTCTTTCCCTGCCTACTGGAATGGGCATTGTTTGYGCGAGCCCGAAAGCTCTAGAAGCTTCCAAAACAGCCAAGTCGGTCAGGGTTTTCTTCGACTGGAATGACTACTTGAAGTTCTACAAGCTTGGAACTTACTGGCCTTAACGCCTTCCATCCAACTACTTTACGGGCTAAGAGCAGCTTTGGATCTCATCTTTGAGGAGGGGCTCGACAATGTCATCGAAAGGCATCGTCGTTTAGGAAAGGCCACAAGGT-AAGTAAAAGCTATACATCT-GACAGCCGCTATTACAGTGTCGGATAGATATATTACTTGGTATTATTGTCCTCTTGTAACAATGAGCAGACTTGCTGTCGAAG

			>Puya_coerulea
			TCGCAGAAAGCTCTTTCCCTGCCTACTGGAATGGGCATTGTTTGCGCGAGCCCGAAAGCTCTAGAAGCTTCCAAAACAGCCAAGTCGGTCAGGGTTTTCTTCGACTGGAATGACTACTTGAAGTTCTACAAGCTTGGAACTTACTGGCCTTAACGCCTTCCATCCAACTACTTTATGGGCTAAGAGCAGCTTTGGATCTCATCTTTGAGGAGGGGCTCGACAATGTCATCGAAAGGCATCGTCGTTTAGGAAAGGCCACAAGGT-AAGTAAAAGCTATACATCT-GACAGCCGCTTTTACAGTGTCGGATAGATATATTACTTGGTATTATTGTCCTCTTGTAACAATGAGCAGACTTGCTGTCGAAG

			>Puya_coerulea_var._violacea
			????????????????????????????????????????????????????????????????????????????????????????????????????????????????????????????????????????????????????????????????????????????????????????????????????????????????????????????????????????????????????????????????????????????????????????????????????????????????????????????????????????????????????????????????????????????????

			>Puya_compacta
			????????????????????????????????????????????????????????????????????????????????????????????????????????????????????????????????????????????????????????????????????????????????????????????????????????????????????????????????????????????????????????????????????????????????????????????????????????????????????????????????????????????????????????????????????????????????

			>Puya_dasylirioides
			????????????????????????????????????????????????????????????????????????????????????????????????????????????????????????????????????????????????????????????????????????????????????????????????????????????????????????????????????????????????????????????????????????????????????????????????????????????????????????????????????????????????????????????????????????????????

			>Puya_densiflora
			????????????????????????????????????????????????????????????????????????????????????????????????????????????????????????????????????????????????????????????????????????????????????????????????????????????????????????????????????????????????????????????????????????????????????????????????????????????????????????????????????????????????????????????????????????????????

			>Puya_dyckioides
			????????????????????????????????????????????????????????????????????????????????????????????????????????????????????????????????????????????????????????????????????????????????????????????????????????????????????????????????????????????????????????????????????????????????????????????????????????????????????????????????????????????????????????????????????????????????

			>Puya_ferreyrae
			????????????????????????????????????????????????????????????????????????????????????????????????????????????????????????????????????????????????????????????????????????????????????????????????????????????????????????????????????????????????????????????????????????????????????????????????????????????????????????????????????????????????????????????????????????????????

			>Puya_ferruginea
			????????????????????????????????????????????????????????????????????????????????????????????????????????????????????????????????????????????????????????????????????????????????????????????????????????????????????????????????????????????????????????????????????????????????????????????????????????????????????????????????????????????????????????????????????????????????

			>Puya_floccosa
			????????????????????????????????????????????????????????????????????????????????????????????????????????????????????????????????????????????????????????????????????????????????????????????????????????????????????????????????????????????????????????????????????????????????????????????????????????????????????????????????????????????????????????????????????????????????

			>Puya_gilmartiniae
			TCGCAGAAAGCTCTTTCCCTGCCTACTGGAATGGGCATTGTTTGCGCGAGCCCGAAAGCTCTAGAAGCTTCCAAAACAGCCAAGTCGGTCAGGGTTTTCTTCGACTGGAATGACTACTTGAAGTTCTACAAGCTTGGAACTTACTGGCCTTAACGCCTTCCATCCAACTACTTTACGGGCTAAGAGCAGCTTTGGATCTCATCTTTGAGGAAGGGCTCGACAATGTCATCGAAAGGCATCGTCGTTTAGGAAAGGCCACAAGGT-AAGTAAAAGCTATACATCT-GACAGCCGCTTTTACAGTGTCGGATAGATATATTACTTGGTATTATTGTCCTCTTGTAACAATGAGCAGACTTGCTGTCGAAG

			>Puya_goudotiana
			????????????????????????????????????????????????????????????????????????????????????????????????????????????????????????????????????????????????????????????????????????????????????????????????????????????????????????????????????????????????????????????????????????????????????????????????????????????????????????????????????????????????????????????????????????????????

			>Puya_hamata
			????????????????????????????????????????????????????????????????????????????????????????????????????????????????????????????????????????????????????????????????????????????????????????????????????????????????????????????????????????????????????????????????????????????????????????????????????????????????????????????????????????????????????????????????????????????????

			>Puya_harmsii
			????????????????????????????????????????????????????????????????????????????????????????????????????????????????????????????????????????????????????????????????????????????????????????????????????????????????????????????????????????????????????????????????????????????????????????????????????????????????????????????????????????????????????????????????????????????????

			>Puya_herrerae
			????????????????????????????????????????????????????????????????????????????????????????????????????????????????????????????????????????????????????????????????????????????????????????????????????????????????????????????????????????????????????????????????????????????????????????????????????????????????????????????????????????????????????????????????????????????????

			>Puya_herzogii
			????????????????????????????????????????????????????????????????????????????????????????????????????????????????????????????????????????????????????????????????????????????????????????????????????????????????????????????????????????????????????????????????????????????????????????????????????????????????????????????????????????????????????????????????????????????????

			>Puya_humilis
			????????????????????????????????????????????????????????????????????????????????????????????????????????????????????????????????????????????????????????????????????????????????????????????????????????????????????????????????????????????????????????????????????????????????????????????????????????????????????????????????????????????????????????????????????????????????

			>Puya_lanata
			????????????????????????????????????????????????????????????????????????????????????????????????????????????????????????????????????????????????????????????????????????????????????????????????????????????????????????????????????????????????????????????????????????????????????????????????????????????????????????????????????????????????????????????????????????????????

			>Puya_laxa
			????????????????????????????????????????????????????????????????????????????????????????????????????????????????????????????????????????????????????????????????????????????????????????????????????????????????????????????????????????????????????????????????????????????????????????????????????????????????????????????????????????????????????????????????????????????????

			>Puya_lineata
			????????????????????????????????????????????????????????????????????????????????????????????????????????????????????????????????????????????????????????????????????????????????????????????????????????????????????????????????????????????????????????????????????????????????????????????????????????????????????????????????????????????????????????????????????????????????

			>Puya_macrura
			????????????????????????????????????????????????????????????????????????????????????????????????????????????????????????????????????????????????????????????????????????????????????????????????????????????????????????????????????????????????????????????????????????????????????????????????????????????????????????????????????????????????????????????????????????????????

			>Puya_mima
			????????????????????????????????????????????????????????????????????????????????????????????????????????????????????????????????????????????????????????????????????????????????????????????????????????????????????????????????????????????????????????????????????????????????????????????????????????????????????????????????????????????????????????????????????????????????

			>Puya_mirabilis
			????????????????????????????????????????????????????????????????????????????????????????????????????????????????????????????????????????????????????????????????????????????????????????????????????????????????????????????????????????????????????????????????????????????????????????????????????????????????????????????????????????????????????????????????????????????????

			>Puya_nana
			????????????????????????????????????????????????????????????????????????????????????????????????????????????????????????????????????????????????????????????????????????????????????????????????????????????????????????????????????????????????????????????????????????????????????????????????????????????????????????????????????????????????????????????????????????????????

			>Puya_nitida
			????????????????????????????????????????????????????????????????????????????????????????????????????????????????????????????????????????????????????????????????????????????????????????????????????????????????????????????????????????????????????????????????????????????????????????????????????????????????????????????????????????????????????????????????????????????????

			>Puya_nutans
			????????????????????????????????????????????????????????????????????????????????????????????????????????????????????????????????????????????????????????????????????????????????????????????????????????????????????????????????????????????????????????????????????????????????????????????????????????????????????????????????????????????????????????????????????????????????

			>Puya_obconica
			????????????????????????????????????????????????????????????????????????????????????????????????????????????????????????????????????????????????????????????????????????????????????????????????????????????????????????????????????????????????????????????????????????????????????????????????????????????????????????????????????????????????????????????????????????????????

			>Puya_pygmaea
			????????????????????????????????????????????????????????????????????????????????????????????????????????????????????????????????????????????????????????????????????????????????????????????????????????????????????????????????????????????????????????????????????????????????????????????????????????????????????????????????????????????????????????????????????????????????

			>Puya_raimondii
			????????????????????????????????????????????????????????????????????????????????????????????????????????????????????????????????????????????????????????????????????????????????????????????????????????????????????????????????????????????????????????????????????????????????????????????????????????????????????????????????????????????????????????????????????????????????

			>Puya_santosii
			????????????????????????????????????????????????????????????????????????????????????????????????????????????????????????????????????????????????????????????????????????????????????????????????????????????????????????????????????????????????????????????????????????????????????????????????????????????????????????????????????????????????????????????????????????????????

			>Puya_spathacea
			????????????????????????????????????????????????????????????????????????????????????????????????????????????????????????????????????????????????????????????????????????????????????????????????????????????????????????????????????????????????????????????????????????????????????????????????????????????????????????????????????????????????????????????????????????????????

			>Puya_stenothyrsa
			????????????????????????????????????????????????????????????????????????????????????????????????????????????????????????????????????????????????????????????????????????????????????????????????????????????????????????????????????????????????????????????????????????????????????????????????????????????????????????????????????????????????????????????????????????????????

			>Puya_trianae
			????????????????????????????????????????????????????????????????????????????????????????????????????????????????????????????????????????????????????????????????????????????????????????????????????????????????????????????????????????????????????????????????????????????????????????????????????????????????????????????????????????????????????????????????????????????????

			>Puya_tuberosa
			????????????????????????????????????????????????????????????????????????????????????????????????????????????????????????????????????????????????????????????????????????????????????????????????????????????????????????????????????????????????????????????????????????????????????????????????????????????????????????????????????????????????????????????????????????????????

			>Puya_ultima
			????????????????????????????????????????????????????????????????????????????????????????????????????????????????????????????????????????????????????????????????????????????????????????????????????????????????????????????????????????????????????????????????????????????????????????????????????????????????????????????????????????????????????????????????????????????????

			>Puya_venusta
			????????????????????????????????????????????????????????????????????????????????????????????????????????????????????????????????????????????????????????????????????????????????????????????????????????????????????????????????????????????????????????????????????????????????????????????????????????????????????????????????????????????????????????????????????????????????

			>Puya_wrightii
			????????????????????????????????????????????????????????????????????????????????????????????????????????????????????????????????????????????????????????????????????????????????????????????????????????????????????????????????????????????????????????????????????????????????????????????????????????????????????????????????????????????????????????????????????????????????

			>Puya_yakespala
			????????????????????????????????????????????????????????????????????????????????????????????????????????????????????????????????????????????????????????????????????????????????????????????????????????????????????????????????????????????????????????????????????????????????????????????????????????????????????????????????????????????????????????????????????????????????

			>Quesnelia_arvensis
			????????????????????????????????????????????????????????????????????????????????????????????????????????????????????????????????????????????????????????????????????????????????????????????????????????????????????????????????????????????????????????????????????????????????????????????????????????????????????????????????????????????????????????????????????????????????

			>Quesnelia_conquistensis
			????????????????????????????????????????????????????????????????????????????????????????????????????????????????????????????????????????????????????????????????????????????????????????????????????????????????????????????????????????????????????????????????????????????????????????????????????????????????????????????????????????????????????????????????????????????????

			>Quesnelia_edmundoi
			????????????????????????????????????????????????????????????????????????????????????????????????????????????????????????????????????????????????????????????????????????????????????????????????????????????????????????????????????????????????????????????????????????????????????????????????????????????????????????????????????????????????????????????????????????????????

			>Quesnelia_edmundoi_var._rubrobracteata
			????????????????????????????????????????????????????????????????????????????????????????????????????????????????????????????????????????????????????????????????????????????????????????????????????????????????????????????????????????????????????????????????????????????????????????????????????????????????????????????????????????????????????????????????????????????????

			>Quesnelia_humilis
			TCGCAGAAAGCTCTTTCCCTGCCTACTGGAATGGGCATTCTCTGCGCGAGCCCGAAAGCTCTAGAAGCTTCCAAAACAGCCAAGTCGGTCAGGGTTTTYTTYGACTGGAATGACTACTTGAAGTTCTACAAGCTCGGAACTTACTGGCCTTAACGCCGTCCATCCAACTGCTTTACGGCCTGAGAGCAGCTTTGGACCTCATCTTTGAGGAGGGGTTCGACAATGTCATCGAAAGGCATCGTCGCTTAGGAAAGGCCACAAGGTTAAGTAAAAGCTATACATCTAAATAGCCGCCTTCGCCGTGTTGGATAGATATATTACTTGGCACTGTTGTTCTCTTGTAACAATGGGCAGACTTGCCGTCGAAG

			>Quesnelia_kautskyi
			TCGCAGAAAGCTCTTTCCCTGCCTACTGGAATGGGCATTCTCTGCGCGAGCCCGAAAGCTCTAGAAGCTTCCAAAACAGCCAAGTCGGTCAGGGTTTTCTTCGACTGGAATGACTACTTGAAGTTCTACAAGCTCGGAACTTACTGGCCTTAACGCCGTCCATCCAACTGCTTTATGGGCTGAGAGCAGCTTTGGATCTCATCTTCGAGGAGGGGCTCGACAATGTCATCGAAAGGCATCGTCGCTTAGGAAAGGCCACAAGGTTAAGTAAAAGCTATACATCTAAATAGCCGCCTTCGCCGTGTTGGATAGATATATTACTTGGCACTGTTGTTCTCTTGTAACAATGAGCAGACTTGCCGTCGAAG

			>Quesnelia_koltesii
			????????????????????????????????????????????????????????????????????????????????????????????????????????????????????????????????????????????????????????????????????????????????????????????????????????????????????????????????????????????????????????????????????????????????????????????????????????????????????????????????????????????????????????????????????????????????

			>Quesnelia_lateralis
			????????????????????????????????????????????????????????????????????????????????????????????????????????????????????????????????????????????????????????????????????????????????????????????????????????????????????????????????????????????????????????????????????????????????????????????????????????????????????????????????????????????????????????????????????????????????

			>Quesnelia_liboniana
			TCGCAGAAAGCTCTTTCCCTGCCTACTGGAATGGGCATTCTCTGCGCGAGCCCGAAAGCTCTAGAAGCTTCCRAAACAGCCAAGTCGGTCAGGGTTTTCTTCGACTGGAATGACTACTTGAAGTTCTACAAGCTCGGAACTTACTGGCCTTAACGCCGTCCATCCAACTGCTTTACGGGCTGAGAGCAGCTTTGGATCTCATCTTTGAGGAGGGGCTCGACAATGTCATCGAAAGGCATCGTCGCTTAGGAAAGGCCACAAGGTTAAGTAAAAGCTCTACATCTAAATAGCCGCCTTCGCCGTGTTGGATATATATATTACTTGGCACTGTTGTTCTCTTGTAACAATGAGCAGACTTGCCGTCGAAG

			>Quesnelia_marmorata
			????????????????????????????????????????????????????????????????????????????????????????????????????????????????????????????????????????????????????????????????????????????????????????????????????????????????????????????????????????????????????????????????????????????????????????????????????????????????????????????????????????????????????????????????????????????????

			>Quesnelia_quesneliana
			????????????????????????????????????????????????????????????????????????????????????????????????????????????????????????????????????????????????????????????????????????????????????????????????????????????????????????????????????????????????????????????????????????????????????????????????????????????????????????????????????????????????????????????????????????????????

			>Quesnelia_strobilispica
			????????????????????????????????????????????????????????????????????????????????????????????????????????????????????????????????????????????????????????????????????????????????????????????????????????????????????????????????????????????????????????????????????????????????????????????????????????????????????????????????????????????????????????????????????????????????

			>Quesnelia_testudo
			????????????????????????????????????????????????????????????????????????????????????????????????????????????????????????????????????????????????????????????????????????????????????????????????????????????????????????????????????????????????????????????????????????????????????????????????????????????????????????????????????????????????????????????????????????????????

			>Quesnelia_tubifolia
			????????????????????????????????????????????????????????????????????????????????????????????????????????????????????????????????????????????????????????????????????????????????????????????????????????????????????????????????????????????????????????????????????????????????????????????????????????????????????????????????????????????????????????????????????????????????

			>Racinaea_cuspidata
			????????????????????????????????????????????????????????????????????????????????????????????????????????????????????????????????????????????????????????????????????????????????????????????????????????????????????????????????????????????????????????????????????????????????????????????????????????????????????????????????????????????????????????????????????????????????

			>Racinaea_dyeriana
			????????????????????????????????????????????????????????????????????????????????????????????????????????????????????????????????????????????????????????????????????????????????????????????????????????????????????????????????????????????????????????????????????????????????????????????????????????????????????????????????????????????????????????????????????????????????

			>Racinaea_elegans
			????????????????????????????????????????????????????????????????????????????????????????????????????????????????????????????????????????????????????????????????????????????????????????????????????????????????????????????????????????????????????????????????????????????????????????????????????????????????????????????????????????????????????????????????????????????????

			>Racinaea_fraseri
			????????????????????????????????????????????????????????????????????????????????????????????????????????????????????????????????????????????????????????????????????????????????????????????????????????????????????????????????????????????????????????????????????????????????????????????????????????????????????????????????????????????????????????????????????????????????

			>Racinaea_hamaleana
			????????????????????????????????????????????????????????????????????????????????????????????????????????????????????????????????????????????????????????????????????????????????????????????????????????????????????????????????????????????????????????????????????????????????????????????????????????????????????????????????????????????????????????????????????????????????

			>Racinaea_multiflora
			????????????????????????????????????????????????????????????????????????????????????????????????????????????????????????????????????????????????????????????????????????????????????????????????????????????????????????????????????????????????????????????????????????????????????????????????????????????????????????????????????????????????????????????????????????????????

			>Racinaea_parviflora
			????????????????????????????????????????????????????????????????????????????????????????????????????????????????????????????????????????????????????????????????????????????????????????????????????????????????????????????????????????????????????????????????????????????????????????????????????????????????????????????????????????????????????????????????????????????????

			>Racinaea_pugiformis
			TCGCAGAAAGCACTTTCCCTGCCTACTGGAATGGGCATTGTTTGCGCGAGCCCGAAAGCTCTAGAAGCTTCCAAAAATGCCAAGTCGGTYAGAGTTTTCTTCGACTGGAATGACTACTTGAAGTTCTACAAGCTTGGAACTTACTGGCCGTAACGCCTTCCATCCAACTACTTTACGGGCTAAGAGCAGCTTTGGATCTCATCTTTGAGGAGGGGCTCGAAAATGTCATCGAAAGGCATCGTCGTTTAGGAAAGGCCACGAGG-TAAGTA-AAGCT--------AAATAGCCGATTTTACAGTATAGGAAAGATATATAATTAGTCATTATTGTCCTCTTGTAACAACGAGCAGACTTGCTGTCGAAG

			>Racinaea_ropalocarpa
			????????????????????????????????????????????????????????????????????????????????????????????????????????????????????????????????????????????????????????????????????????????????????????????????????????????????????????????????????????????????????????????????????????????????????????????????????????????????????????????????????????????????????????????????????????????????

			>Racinaea_rothschuhiana
			????????????????????????????????????????????????????????????????????????????????????????????????????????????????????????????????????????????????????????????????????????????????????????????????????????????????????????????????????????????????????????????????????????????????????????????????????????????????????????????????????????????????????????????????????????????????

			>Racinaea_schumanniana
			????????????????????????????????????????????????????????????????????????????????????????????????????????????????????????????????????????????????????????????????????????????????????????????????????????????????????????????????????????????????????????????????????????????????????????????????????????????????????????????????????????????????????????????????????????????????

			>Racinaea_seemannii
			????????????????????????????????????????????????????????????????????????????????????????????????????????????????????????????????????????????????????????????????????????????????????????????????????????????????????????????????????????????????????????????????????????????????????????????????????????????????????????????????????????????????????????????????????????????????

			>Racinaea_sinuosa
			?????????????????????????CTGGAATGGGCATTGTTTGCGCGAGCCCGAAAGCTCTAGAAGCTTCCAAAAATGCCAAGTCGGTCAGAGTTTTCTTCGACTGGAATGACTACTTGAAGTTCTACAAGCTTGGAACTTACTGGCCGTAACGCCTTCCATCCAACTACTTTACGGGCTAAGAGCAGCTTTGGATCTCATCTTTGAGGAGGGGCTCGAAAATGTCATCGAAAGGCATCGTCGTTTAGGAAAGGCCACGAGG-TAAGTA-AAGCT--------AAATAGCCGATTTTACAGTATAGGAAAGATATATAATTAGTCATTATTGTCCTCTTGTAACAACAAGCAGGCTTGCTGTCGAAG

			>Racinaea_spiculosa
			TCGCAGAAAGCACTTTCCCTGCCTACTGGAATGGGCATTGTTTGCGCGAGCCCGAAAGCTCTAGAAGCTTCCAAAAATGCCAAGTCGGTCAGAGTTTTCTTCGACTGGAATGAYTACTTGAAGTTCTACAAGCTTGGAACTTATTGGCCGTAACGCCTTCCATCCAACTACTTTACGGGCTAAGAGCAGCTTTGGATCTCATCTTTGAGGAGGGGCTCGAAAATGTCATCGAAAGGCATCGTCGTTTAGGAAAGGCCACGAGG-TAAGTA-AAGCT--------AAATAGCCGATTTTACAGTATAGGAAAGATATATAATTAGTCATTATTGTCCTCTTGTAAC???????????????????????

			>Racinaea_spiculosa_var._micrantha
			????????????????????????????????????????????????????????????????????????????????????????????????????????????????????????????????????????????????????????????????????????????????????????????????????????????????????????????????????????????????????????????????????????????????????????????????????????????????????????????????????????????????????????????????????????????????

			>Racinaea_venusta
			????????????????????????????????????????????????????????????????????????????????????????????????????????????????????????????????????????????????????????????????????????????????????????????????????????????????????????????????????????????????????????????????????????????????????????????????????????????????????????????????????????????????????????????????????????????????

			>Rapatea_paludosa
			????????????????????????????????????????????????????????????????????????????????????????????????????????????????????????????????????????????????????????????????????????????????????????????????????????????????????????????????????????????????????????????????????????????????????????????????????????????????????????????????????????????????????????????????????????????????

			>Rokautskyia_exaltata
			????????????????????????????????????????????????????????????????????????????????????????????????????????????????????????????????????????????????????????????????????????????????????????????????????????????????????????????????????????????????????????????????????????????????????????????????????????????????????????????????????????????????????????????????????????????????

			>Rokautskyia_fernseeoides
			????????????????????????????????????????????????????????????????????????????????????????????????????????????????????????????????????????????????????????????????????????????????????????????????????????????????????????????????????????????????????????????????????????????????????????????????????????????????????????????????????????????????????????????????????????????????

			>Rokautskyia_latifolia
			????????????????????????????????????????????????????????????????????????????????????????????????????????????????????????????????????????????????????????????????????????????????????????????????????????????????????????????????????????????????????????????????????????????????????????????????????????????????????????????????????????????????????????????????????????????????

			>Rokautskyia_microglazioui
			????????????????????????????????????????????????????????????????????????????????????????????????????????????????????????????????????????????????????????????????????????????????????????????????????????????????????????????????????????????????????????????????????????????????????????????????????????????????????????????????????????????????????????????????????????????????

			>Rokautskyia_odoratissima
			????????????????????????????????????????????????????????????????????????????????????????????????????????????????????????????????????????????????????????????????????????????????????????????????????????????????????????????????????????????????????????????????????????????????????????????????????????????????????????????????????????????????????????????????????????????????

			>Rokautskyia_roberto-kautskyi
			????????????????????????????????????????????????????????????????????????????????????????????????????????????????????????????????????????????????????????????????????????????????????????????????????????????????????????????????????????????????????????????????????????????????????????????????????????????????????????????????????????????????????????????????????????????????

			>Rokautskyia_sanctaluciae
			????????????????????????????????????????????????????????????????????????????????????????????????????????????????????????????????????????????????????????????????????????????????????????????????????????????????????????????????????????????????????????????????????????????????????????????????????????????????????????????????????????????????????????????????????????????????

			>Rokautskyia_scaposa
			????????????????????????????????????????????????????????????????????????????????????????????????????????????????????????????????????????????????????????????????????????????????????????????????????????????????????????????????????????????????????????????????????????????????????????????????????????????????????????????????????????????????????????????????????????????????

			>Ronnbergia_aciculosa
			????????????????????????????????????????????????????????????????????????????????????????????????????????????????????????????????????????????????????????????????????????????????????????????????????????????????????????????????????????????????????????????????????????????????????????????????????????????????????????????????????????????????????????????????????????????????

			>Ronnbergia_allenii
			TCGCAGAAAGCTCTTTCCCTGCCTACTGGAATGGGCATTCTCTGCGCGAGCCCGAAAGCTCTAGAAGCTTCCAAAACAGCCAAGTCGGTCAGGGTTTTCTTCGACTGGAATGACTACTTGAAGTTCTACAAGCTCGGAACTTACTGGCCTTAACGCCTTCCATCCAACTACTTTACGGGCTGAGAGCAGCTTTGGATCTCATCTTTGAGGAGGGGCTYGACAATGTCATYGAAAGGCACCGTCGCTTAGGAAAGGCCACAAGGTTAAGTAAAAGCTATACATCT-AATAGCCGCTTTCGCCGTGTYGGATAGATATATTACATGGCACTATTGTTCTCTTGTAACAATGAGCAGACTTGCTGTCGAAG

			>Ronnbergia_columbiana
			????????????????????????????????????????????????????????????????????????????????????????????????????????????????????????????????????????????????????????????????????????????????????????????????????????????????????????????????????????????????????????????????????????????????????????????????????????????????????????????????????????????????????????????????????????????????

			>Ronnbergia_deleonii
			????????????????????????????????????????????????????????????????????????????????????????????????????????????????????????????????????????????????????????????????????????????????????????????????????????????????????????????????????????????????????????????????????????????????????????????????????????????????????????????????????????????????????????????????????????????????

			>Ronnbergia_drakeana
			????????????????????????????????????????????????????????????????????????????????????????????????????????????????????????????????????????????????????????????????????????????????????????????????????????????????????????????????????????????????????????????????????????????????????????????????????????????????????????????????????????????????????????????????????????????????

			>Ronnbergia_explodens
			????????????????????????????????????????????????????????????????????????????????????????????????????????????????????????????????????????????????????????????????????????????????????????????????????????????????????????????????????????????????????????????????????????????????????????????????????????????????????????????????????????????????????????????????????????????????

			>Ronnbergia_fraseri
			TCGCAGAAAGCTCTTTCCCTGCCTACTGGAATGGGGATTCTCTGCGCGAGCCCGAAAGCTCTAGAAGCTTCCAAAACAGCCAAGTCGGTCAGGGTTTTCTTCGACTGGAATGACTACTTGAAGTTCTACAAGCTCGGAACTTACTGGCCTTAACGCCTTCCATCCAACTACTTTACGGGCTGAGAGCAGCTCTGGATCTCATCTTTGAGGAGGGGCTYGACAATGTCATCGAAAGGCATCGTCGCTTAGGAAAGGCCACAAGGTTAAGTAAAAGCTATACATCT-AATACCCGCTTTCGCCGTGTTGGATAGATATATTACTTGGSACCATTGTTCTMTTGTAACAATGAGCAGGCTTGCTGTCGAAG

			>Ronnbergia_germinyana
			????????????????????????????????????????????????????????????????????????????????????????????????????????????????????????????????????????????????????????????????????????????????????????????????????????????????????????????????????????????????????????????????????????????????????????????????????????????????????????????????????????????????????????????????????????????????

			>Ronnbergia_hathewayi
			????????????????????????????????????????????????????????????????????????????????????????????????????????????????????????????????????????????????????????????????????????????????????????????????????????????????????????????????????????????????????????????????????????????????????????????????????????????????????????????????????????????????????????????????????????????????

			>Ronnbergia_involucrata
			????????????????????????????????????????????????????????????????????????????????????????????????????????????????????????????????????????????????????????????????????????????????????????????????????????????????????????????????????????????????????????????????????????????????????????????????????????????????????????????????????????????????????????????????????????????????

			>Ronnbergia_killipiana
			????????????????????????????????????????????????????????????????????????????????????????????????????????????????????????????????????????????????????????????????????????????????????????????????????????????????????????????????????????????????????????????????????????????????????????????????????????????????????????????????????????????????????????????????????????????????

			>Ronnbergia_maidifolia
			????????????????????????????????????????????????????????????????????????????????????????????????????????????????????????????????????????????????????????????????????????????????????????????????????????????????????????????????????????????????????????????????????????????????????????????????????????????????????????????????????????????????????????????????????????????????

			>Ronnbergia_morreniana
			????????????????????????????????????????????????????????????????????????????????????????????????????????????????????????????????????????????????????????????????????????????????????????????????????????????????????????????????????????????????????????????????????????????????????????????????????????????????????????????????????????????????????????????????????????????????

			>Ronnbergia_nidularioides
			????????????????????????????????????????????????????????????????????????????????????????????????????????????????????????????????????????????????????????????????????????????????????????????????????????????????????????????????????????????????????????????????????????????????????????????????????????????????????????????????????????????????????????????????????????????????

			>Ronnbergia_petersii
			????????????????????????????????????????????????????????????????????????????????????????????????????????????????????????????????????????????????????????????????????????????????????????????????????????????????????????????????????????????????????????????????????????????????????????????????????????????????????????????????????????????????????????????????????????????????

			>Ronnbergia_tonduzii
			TCGCAGAAAGCTCTTTCCCTGCCTACTGGGATGGGCATTCTTTGCGCCAGCCCGAAAGCTCTAGAAGCTTCCAAAACAGCCAAGTCGGTCAGGGTTTTCTTCGACTGGAATGACTACTTGAAGTTCTACAAGCTCGGAACTTACTGGCCTTAACGCCGTCCATCCAACTACTTTACGGGCTGAGAGCAGCTTTGGATCTCATCTTTGAGGAGGGGCTCGACAACGTCATCGAAAGGCACCGTCGCTTAGGAAAGGCCACAAGGTTAAGTAAAAGCTATACATCT-AATAGCCGCTTTCGCCGTGTTGGATAGATATATAACTTGGCGCTATTGTTCTCTTGTAACAATGAGCAGACTTGCTGTCGAAG

			>Ronnbergia_veitchii
			????????????????????????????????????????????????????????????????????????????????????????????????????????????????????????????????????????????????????????????????????????????????????????????????????????????????????????????????????????????????????????????????????????????????????????????????????????????????????????????????????????????????????????????????????????????????

			>Ronnbergia_weberbaueri
			????????????????????????????????????????????????????????????????????????????????????????????????????????????????????????????????????????????????????????????????????????????????????????????????????????????????????????????????????????????????????????????????????????????????????????????????????????????????????????????????????????????????????????????????????????????????

			>Ronnbergia_wuelfinghoffii
			????????????????????????????????????????????????????????????????????????????????????????????????????????????????????????????????????????????????????????????????????????????????????????????????????????????????????????????????????????????????????????????????????????????????????????????????????????????????????????????????????????????????????????????????????????????????

			>Sequencia_serrata
			TCGCAGAAAGCTCTTTCCATGCCCACTGGAATGGGCATTGTTTGTGCGAGCCCGAAAGCTCTGGAAGTGTCCAAAACGGCCAAGTCGGTCAGGGTTTTCTTCGACTGGAATGACTACTTGAAGTTCTACAAGCTTGGAACTTACTGGCCTTAACACCTTCCATCCAACTACTGTATGGGCTAAGAGCAGCTTTGGATCTCATCTTTGAGGAGGGGCTCGACAATGTCATTGAAAGGCATCGTCGTTTAGGAAAGGCCACAAGGT-AAGTAAAAACTTTACACCT-AATAGCAGATTTTACAGTATTGGAAAGATATATTACTTGGCATTGTAGTCCTCTCGTACCAATGAGCAGACTAGCTGTCGAAG

			>Sincoraea_albopicta
			????????????????????????????????????????????????????????????????????????????????????????????????????????????????????????????????????????????????????????????????????????????????????????????????????????????????????????????????????????????????????????????????????????????????????????????????????????????????????????????????????????????????????????????????????????????????

			>Sincoraea_burle-marxii
			????????????????????????????????????????????????????????????????????????????????????????????????????????????????????????????????????????????????????????????????????????????????????????????????????????????????????????????????????????????????????????????????????????????????????????????????????????????????????????????????????????????????????????????????????????????????

			>Sincoraea_hatschbachii
			????????????????????????????????????????????????????????????????????????????????????????????????????????????????????????????????????????????????????????????????????????????????????????????????????????????????????????????????????????????????????????????????????????????????????????????????????????????????????????????????????????????????????????????????????????????????

			>Sincoraea_heleniceae
			????????????????????????????????????????????????????????????????????????????????????????????????????????????????????????????????????????????????????????????????????????????????????????????????????????????????????????????????????????????????????????????????????????????????????????????????????????????????????????????????????????????????????????????????????????????????

			>Sincoraea_humilis
			????????????????????????????????????????????????????????????????????????????????????????????????????????????????????????????????????????????????????????????????????????????????????????????????????????????????????????????????????????????????????????????????????????????????????????????????????????????????????????????????????????????????????????????????????????????????

			>Sincoraea_mucugensis
			????????????????????????????????????????????????????????????????????????????????????????????????????????????????????????????????????????????????????????????????????????????????????????????????????????????????????????????????????????????????????????????????????????????????????????????????????????????????????????????????????????????????????????????????????????????????

			>Sincoraea_navioides
			????????????????????????????????????????????????????????????????????????????????????????????????????????????????????????????????????????????????????????????????????????????????????????????????????????????????????????????????????????????????????????????????????????????????????????????????????????????????????????????????????????????????????????????????????????????????

			>Sincoraea_ophiuroides
			????????????????????????????????????????????????????????????????????????????????????????????????????????????????????????????????????????????????????????????????????????????????????????????????????????????????????????????????????????????????????????????????????????????????????????????????????????????????????????????????????????????????????????????????????????????????

			>Sincoraea_ulei
			????????????????????????????????????????????????????????????????????????????????????????????????????????????????????????????????????????????????????????????????????????????????????????????????????????????????????????????????????????????????????????????????????????????????????????????????????????????????????????????????????????????????????????????????????????????????

			>Stegolepis_ligulata
			????????????????????????????????????????????????????????????????????????????????????????????????????????????????????????????????????????????????????????????????????????????????????????????????????????????????????????????????????????????????????????????????????????????????????????????????????????????????????????????????????????????????????????????????????????????????

			>Stegolepis_parvipetala
			????????????????????????????????????????????????????????????????????????????????????????????????????????????????????????????????????????????????????????????????????????????????????????????????????????????????????????????????????????????????????????????????????????????????????????????????????????????????????????????????????????????????????????????????????????????????

			>Stigmatodon_amadoi
			????????????????????????????????????????????????????????????????????????????????????????????????????????????????????????????????????????????????????????????????????????????????????????????????????????????????????????????????????????????????????????????????????????????????????????????????????????????????????????????????????????????????????????????????????????????????

			>Stigmatodon_apparicianus
			????????????????????????????????????????????????????????????????????????????????????????????????????????????????????????????????????????????????????????????????????????????????????????????????????????????????????????????????????????????????????????????????????????????????????????????????????????????????????????????????????????????????????????????????????????????????

			>Stigmatodon_belloi
			????????????????????????????????????????????????????????????????????????????????????????????????????????????????????????????????????????????????????????????????????????????????????????????????????????????????????????????????????????????????????????????????????????????????????????????????????????????????????????????????????????????????????????????????????????????????

			>Stigmatodon_bifidus
			????????????????????????????????????????????????????????????????????????????????????????????????????????????????????????????????????????????????????????????????????????????????????????????????????????????????????????????????????????????????????????????????????????????????????????????????????????????????????????????????????????????????????????????????????????????????

			>Stigmatodon_brassicoides
			????????????????????????????????????????????????????????????????????????????????????????????????????????????????????????????????????????????????????????????????????????????????????????????????????????????????????????????????????????????????????????????????????????????????????????????????????????????????????????????????????????????????????????????????????????????????

			>Stigmatodon_costae
			????????????????????????????????????????????????????????????????????????????????????????????????????????????????????????????????????????????????????????????????????????????????????????????????????????????????????????????????????????????????????????????????????????????????????????????????????????????????????????????????????????????????????????????????????????????????

			>Stigmatodon_croceanus
			????????????????????????????????????????????????????????????????????????????????????????????????????????????????????????????????????????????????????????????????????????????????????????????????????????????????????????????????????????????????????????????????????????????????????????????????????????????????????????????????????????????????????????????????????????????????

			>Stigmatodon_euclidianus
			????????????????????????????????????????????????????????????????????????????????????????????????????????????????????????????????????????????????????????????????????????????????????????????????????????????????????????????????????????????????????????????????????????????????????????????????????????????????????????????????????????????????????????????????????????????????

			>Stigmatodon_funebris
			????????????????????????????????????????????????????????????????????????????????????????????????????????????????????????????????????????????????????????????????????????????????????????????????????????????????????????????????????????????????????????????????????????????????????????????????????????????????????????????????????????????????????????????????????????????????

			>Stigmatodon_gastinianus
			????????????????????????????????????????????????????????????????????????????????????????????????????????????????????????????????????????????????????????????????????????????????????????????????????????????????????????????????????????????????????????????????????????????????????????????????????????????????????????????????????????????????????????????????????????????????

			>Stigmatodon_goniorachis
			????????????????????????????????????????????????????????????????????????????????????????????????????????????????????????????????????????????????????????????????????????????????????????????????????????????????????????????????????????????????????????????????????????????????????????????????????????????????????????????????????????????????????????????????????????????????

			>Stigmatodon_harrylutheri
			????????????????????????????????????????????????????????????????????????????????????????????????????????????????????????????????????????????????????????????????????????????????????????????????????????????????????????????????????????????????????????????????????????????????????????????????????????????????????????????????????????????????????????????????????????????????

			>Stigmatodon_multifoliatus
			????????????????????????????????????????????????????????????????????????????????????????????????????????????????????????????????????????????????????????????????????????????????????????????????????????????????????????????????????????????????????????????????????????????????????????????????????????????????????????????????????????????????????????????????????????????????

			>Stigmatodon_plurifolius
			????????????????????????????????????????????????????????????????????????????????????????????????????????????????????????????????????????????????????????????????????????????????????????????????????????????????????????????????????????????????????????????????????????????????????????????????????????????????????????????????????????????????????????????????????????????????

			>Stigmatodon_rosulatulus
			????????????????????????????????????????????????????????????????????????????????????????????????????????????????????????????????????????????????????????????????????????????????????????????????????????????????????????????????????????????????????????????????????????????????????????????????????????????????????????????????????????????????????????????????????????????????

			>Stigmatodon_sanctateresensis
			????????????????????????????????????????????????????????????????????????????????????????????????????????????????????????????????????????????????????????????????????????????????????????????????????????????????????????????????????????????????????????????????????????????????????????????????????????????????????????????????????????????????????????????????????????????????

			>Tillandsia_achyrostachys
			????????????????????????????????????????????????????????????????????????????????????????????????????????????????????????????????????????????????????????????????????????????????????????????????????????????????????????????????????????????????????????????????????????????????????????????????????????????????????????????????????????????????????????????????????????????????

			>Tillandsia_adpressiflora
			????????????????????????????????????????????????????????????????????????????????????????????????????????????????????????????????????????????????????????????????????????????????????????????????????????????????????????????????????????????????????????????????????????????????????????????????????????????????????????????????????????????????????????????????????????????????

			>Tillandsia_aeranthos
			????????????????????????????????????????????????????????????????????????????????????????????????????????????????????????????????????????????????????????????????????????????????????????????????????????????????????????????????????????????????????????????????????????????????????????????????????????????????????????????????????????????????????????????????????????????????

			>Tillandsia_afonsoana
			????????????????????????????????????????????????????????????????????????????????????????????????????????????????????????????????????????????????????????????????????????????????????????????????????????????????????????????????????????????????????????????????????????????????????????????????????????????????????????????????????????????????????????????????????????????????

			>Tillandsia_aizoides
			TCGCAGAAAGCACTTTCCCTGCCTACTGGAATGGGCATTGTTTGCGCGAGCCCGAAAGCTCTGGAAGCTTCCAAAAATGCCAAGTCGGTTAGAGTTTTCTTCGACTGGAATGACTACTTGAAGTTCTACAAGCTTGGAACTTACTGGCCGTAACGCCTTCCATCCAACTACTTTACGGGCTAAGAGCAGCTTTGGATCTCATCTTTGAGGAGGGCCTCGAAAATGTCATCGAAAGGCATCGTCGTTTAGGAAAGGCCACGAGG-TAAGTATAAGCT--------AAATAGCCGATTTTACAGTATAGGAAAGATATGTAATTAGCCATTATTGTCCTCTTGTAACAATGAGCAGACTTGCTGTCGAAG

			>Tillandsia_albertiana
			????????????????????????????????????????????????????????????????????????????????????????????????????????????????????????????????????????????????????????????????????????????????????????????????????????????????????????????????????????????????????????????????????????????????????????????????????????????????????????????????????????????????????????????????????????????????

			>Tillandsia_albida
			????????????????????????????????????????????????????????????????????????????????????????????????????????????????????????????????????????????????????????????????????????????????????????????????????????????????????????????????????????????????????????????????????????????????????????????????????????????????????????????????????????????????????????????????????????????????

			>Tillandsia_andicola
			????????????????????????????????????????????????????????????????????????????????????????????????????????????????????????????????????????????????????????????????????????????????????????????????????????????????????????????????????????????????????????????????????????????????????????????????????????????????????????????????????????????????????????????????????????????????

			>Tillandsia_andrieuxii
			????????????????????????????????????????????????????????????????????????????????????????????????????????????????????????????????????????????????????????????????????????????????????????????????????????????????????????????????????????????????????????????????????????????????????????????????????????????????????????????????????????????????????????????????????????????????

			>Tillandsia_appenii
			????????????????????????????????????????????????????????????????????????????????????????????????????????????????????????????????????????????????????????????????????????????????????????????????????????????????????????????????????????????????????????????????????????????????????????????????????????????????????????????????????????????????????????????????????????????????

			>Tillandsia_araujei
			????????????????????????????????????????????????????????????????????????????????????????????????????????????????????????????????????????????????????????????????????????????????????????????????????????????????????????????????????????????????????????????????????????????????????????????????????????????????????????????????????????????????????????????????????????????????

			>Tillandsia_argentea
			????????????????????????????????????????????????????????????????????????????????????????????????????????????????????????????????????????????????????????????????????????????????????????????????????????????????????????????????????????????????????????????????????????????????????????????????????????????????????????????????????????????????????????????????????????????????

			>Tillandsia_argentina
			????????????????????????????????????????????????????????????????????????????????????????????????????????????????????????????????????????????????????????????????????????????????????????????????????????????????????????????????????????????????????????????????????????????????????????????????????????????????????????????????????????????????????????????????????????????????

			>Tillandsia_arroyoensis
			????????????????????????????????????????????????????????????????????????????????????????????????????????????????????????????????????????????????????????????????????????????????????????????????????????????????????????????????????????????????????????????????????????????????????????????????????????????????????????????????????????????????????????????????????????????????

			>Tillandsia_atenangoensis
			TCGCAGAAAGCACTTTCCCTGCCTACTGGAATGGGCATTGTTTGCGCGAGCCCGAAAGCTCTGGAAGCTTCCAAAAATGCCAAGTCGGTCAGAGTTTTCTTCGACTGGAATGACTACTTGAAGTTCTACAAGCTTGGAACTTACTGGCCGTAACGCCTTCCATCCAACTACTCTACGGCCTAAGAGCAGCTTTGGATCTCATCTTCGAGGAGGGGCTCGAAAATGTCATCGAAAGGCATCGTCGTTTAGGAAAGGCCACGAGG-TAAGTAAAAGCT--------AAATAGCCGATTTTACAGTATAGGAAAGATATGTAATTAGGCATTATTGTCCTCTTGTAACAATGAGCAGACTTGCTGTCGAAG

			>Tillandsia_atroviolacea
			TCGCAGAAAGCACTTTCCCTGCCTACGGGAATGGGCATTGTTTGCGCGAGCCCGAAAGCTCTGGAAGCTTCCAAAAATGCCAAGTCGGTCAGAGTTTTCTTCGACTGGAATGACTACTTGAAGTTCTACAAGCTTGGAACTTACTGGCCGTAACGCCTTCCATCCAACTACTCTACGGGCTAAGAGCAGCTTTGGATCTCATCTTTGAGGAGGGGCTCGAAAATGTCATCGAAAGGCATCGTCGTTTAGGAAAGGCCACGAGG-TAAGTAAAAGCT--------------------TTACAGTATAGGAAAGATATATAATTAGGCATTATTGTCCTCTTGTAACAATGAGCAGACTTGCTGTCGAAG

			>Tillandsia_atroviridipetala
			TCGCAGAAAGCACTTTCCCTGCCTACTGGAATGGGCATTGTTTGCGCGAGCCCGAAAGCTCTAGAAGCTTCCAAAAATGCCAAGTCGGTCAGAGTTTTCTTCGACTGGAATGACTACTTGAAGTTCTACAAGCTTGGAACTTACTGGCCGTAACGCCTTCCATCCAACTACTTTACGGGCTAAGAGCAGCTTTGGATCTCATCTTTGAGGAGGGGCTCGAAAATGTCATCGAAAGACATCGTCGTTTAGGAAAGGCCACGAGG-TAAGTA-AAGCT--------AAATAGCCGATTTTACAGTATAGGAAAGATATATAATTAGGCATTATTGT-----------AATGAGCAGACTTGCTGTCGAAG

			>Tillandsia_aurea
			????????????????????????????????????????????????????????????????????????????????????????????????????????????????????????????????????????????????????????????????????????????????????????????????????????????????????????????????????????????????????????????????????????????????????????????????????????????????????????????????????????????????????????????????????????????????

			>Tillandsia_australis
			????????????????????????????????????????????????????????????????????????????????????????????????????????????????????????????????????????????????????????????????????????????????????????????????????????????????????????????????????????????????????????????????????????????????????????????????????????????????????????????????????????????????????????????????????????????????

			>Tillandsia_baguagrandensis
			TCGCAGAAAGCACTTTCCCTGCCTACTGGAATGGGCATTGTTTGCGCGAGCCCGAAAGCTCTGGAAGCTTCCAAAAATGCCAAGTCGGTCAGAGTTTTCTTCGACTGGAATGACTACTTGAAGTTCTACAAGCTTGGAACTTACTGGCCGTAACGCCTTCCATCCAACTACTTTACGGGCTAAGAGCAGCTTTGGATCTCATCTTTGAGGAGGGGCTCSAAAATGTCATCGAAAGGCATCGTCGTTTAGGAAAGGCCACGAGG-TAAGTATAAGCT--------AAATAGCCGATTTTACAGTATAGGAAAGATATGTAATTAGCCATTCTTGT-----------AATGAGCAGACTTGCTGTCGAAG

			>Tillandsia_baileyi
			TCGCAGAAAGCACTTTCCCTGCCTACTGG?ATGGGCATTGTTTGCGCGAGCCCGAAAGCTCTGGAAGCTTCCAAAAATGCCAAGTCGGTCAGAGTTTTCTTCGACTGGAATGACTACTTGAAGTTCTACAAGCTTGGAACTTACTGGCCGTAACGCCTTCCATCCAACTACTCTACGGGCTAAGAGCAGCTTTGGATCTCATCTTCGAGGAGGGGCTCGAAAATGTCATCGAAAGGCATCGTCGTTTAGGAAAGGCCACGAGG-TAAGTAAAACCT--------AAATAGCCGATTTTACAGTACAGGAAAGATATATAATTAGGCATTATTGTCCTCTTGTAACAATGAGCAGAC????????????

			>Tillandsia_balbisiana
			???????????????????????????????TGGGCATTGTTTGCGCGAGCCCGAAAGCTCTGGAAGCTTCCAAAAATGCCAAGTCGGTCAGAGTTTTCTTCGACTGGAATGACTACTTGAAGTTCTACAAGCTTGGAACTTACTGGCCGTAACGCCTTCCATCCAACTACTCTACGGCCTAAGAGCAGCTTTGGATCTCATCTTTGAGGAGGGGCTCGAAAATGTCATCGAAAGGCATCGTCGTTTAGGAAAGGCCACGAGG-TAAGTAAAACCT--------AAATAGCCGATTTTACAGCATAGGAAAGATATAAAATTAGGCATTATTGTCCTCTTGTAACAATGAGCAGACTTGCTGTCG???

			>Tillandsia_baliophylla
			????????????????????????????????????????????????????????????????????????????????????????????????????????????????????????????????????????????????????????????????????????????????????????????????????????????????????????????????????????????????????????????????????????????????????????????????????????????????????????????????????????????????????????????????????????????????

			>Tillandsia_balsasensis
			TCGCAGAAAGCACTTTCCCTGCCTACTGGAATGGGCATTGTTTGCGCGAGCCCGAAAGCTCTAGAAGCTTCCAAAAATGCCAAGTCGGTCAGAGTTTTCTTCGACTGGAATGACTACTTGAAGTTCTACAAGCTCGGAACTTACTGGCCGTAACGCCTTCCATCCAACTACTTTACGGGCTAAGAGCAGCTTTGGATCTCATCTTTGAGGAGGGGCTCGAAAATGTCATCGAAAGGCATCGTCGTTTAGGAAAGGCCACGAGG-TAAGTA-AAGCT--------AAATAGCCGATTTTACAGTATAGGAAAGATATATAATTAGGCATTATTGT-----------AATGAGCAGACTTGCTGTCGAAG

			>Tillandsia_barclayana
			????????????????????????????????????????????????????????????????????????????????????????????????????????????????????????????????????????????????????????????????????????????????????????????????????????????????????????????????????????????????????????????????????????????????????????????????????????????????????????????????????????????????????????????????????????????????

			>Tillandsia_barfussii
			????????????????????????????????????????????????????????????????????????????????????????????????????????????????????????????????????????????????????????????????????????????????????????????????????????????????????????????????????????????????????????????????????????????????????????????????????????????????????????????????????????????????????????????????????????????????

			>Tillandsia_barthlottii
			????????????????????????????????????????????????????????????????????????????????????????????????????????????????????????????????????????????????????????????????????????????????????????????????????????????????????????????????????????????????????????????????????????????????????????????????????????????????????????????????????????????????????????????????????????????????

			>Tillandsia_bartramii
			????????????????????????????????????????????????????????????????????????????????????????????????????????????????????????????????????????????????????????????????????????????????????????????????????????????????????????????????????????????????????????????????????????????????????????????????????????????????????????????????????????????????????????????????????????????????

			>Tillandsia_bella
			????????????????????????????????????????????????????????????????????????????????????????????????????????????????????????????????????????????????????????????????????????????????????????????????????????????????????????????????????????????????????????????????????????????????????????????????????????????????????????????????????????????????????????????????????????????????

			>Tillandsia_belloensis
			????????????????????????????????????????????????????????????????????????????????????????????????????????????????????????????????????????????????????????????????????????????????????????????????????????????????????????????????????????????????????????????????????????????????????????????????????????????????????????????????????????????????????????????????????????????????

			>Tillandsia_bergeri
			????????????????????????????????????????????????????????????????????????????????????????????????????????????????????????????????????????????????????????????????????????????????????????????????????????????????????????????????????????????????????????????????????????????????????????????????????????????????????????????????????????????????????????????????????????????????

			>Tillandsia_bermejoensis
			TCGCAGAAAGCACTTTCCCTGCCTACTGGAATGGGCATTGTTTGTGCGAGCCCGAAAGCTCTGGAAGCTTCCAAAAATGCCAAGTCGGTCAGAGTTTTCTTCGACTGGAATGACTACTTGAAGTTCTACAAGCTTGGAACTTACTGGCCGTAACGCCTTCCATCCAACTACTTTACGGGCTAAGAGCAGCTTTGGATCTCATCTTTGAGGAGGGSCTCGAAAATGTCATCGAAAGGCATCGTCGTTTAGGAAAGGCCACGAGG-TAAGTATAAGCT--------AAATAGCCGATTTTACAGTATAGGAAAGATATGTAATTAGCCATTATTGT-----------AATGAGCAGACTTGCTGTCGAAG

			>Tillandsia_biflora
			????????????????????????????????????????????????????????????????????????????????????????????????????????????????????????????????????????????????????????????????????????????????????????????????????????????????????????????????????????????????????????????????????????????????????????????????????????????????????????????????????????????????????????????????????????????????

			>Tillandsia_bonita
			????????????????????????????????????????????????????????????????????????????????????????????????????????????????????????????????????????????????????????????????????????????????????????????????????????????????????????????????????????????????????????????????????????????????????????????????????????????????????????????????????????????????????????????????????????????????

			>Tillandsia_botterii
			????????????????????????????????????????????????????????????????????????????????????????????????????????????????????????????????????????????????????????????????????????????????????????????????????????????????????????????????????????????????????????????????????????????????????????????????????????????????????????????????????????????????????????????????????????????????

			>Tillandsia_bourgaei
			????????????????????????????????????????????????????????????????????????????????????????????????????????????????????????????????????????????????????????????????????????????????????????????????????????????????????????????????????????????????????????????????????????????????????????????????????????????????????????????????????????????????????????????????????????????????

			>Tillandsia_brachycaulos
			????????????????????????????????????????????????????????????????????????????????????????????????????????????????????????????????????????????????????????????????????????????????????????????????????????????????????????????????????????????????????????????????????????????????????????????????????????????????????????????????????????????????????????????????????????????????

			>Tillandsia_brachyphylla
			????????????????????????????????????????????????????????????????????????????????????????????????????????????????????????????????????????????????????????????????????????????????????????????????????????????????????????????????????????????????????????????????????????????????????????????????????????????????????????????????????????????????????????????????????????????????

			>Tillandsia_brevilingua
			????????????????????????????????????????????????????????????????????????????????????????????????????????????????????????????????????????????????????????????????????????????????????????????????????????????????????????????????????????????????????????????????????????????????????????????????????????????????????????????????????????????????????????????????????????????????

			>Tillandsia_bulbosa
			????????????????????????????????????????????????????????????????????????????????????????????????????????????????????????????????????????????????????????????????????????????????????????????????????????????????????????????????????????????????????????????????????????????????????????????????????????????????????????????????????????????????????????????????????????????????

			>Tillandsia_burle-marxii
			????????????????????????????????????????????????????????????????????????????????????????????????????????????????????????????????????????????????????????????????????????????????????????????????????????????????????????????????????????????????????????????????????????????????????????????????????????????????????????????????????????????????????????????????????????????????

			>Tillandsia_butzii
			TCGCAGAAAGCACTTTCCCTGCCTACTGGAATGGGCATTGTTTGCGCGAGCCCGAAAGCTCTGGAAGCTTCCAAAAATGCCAAGTCGGTCAGAGTTTTCTTCGACTGGAATGACTACTTGAAGTTCTACAAGCTTGGAACTTACTGGCCGTAACGCCTTCCATCCAACTACTCTACGGGCTAAGAGCAGCTTTGGATCTCATCTTCGAGGAGGGGCTCGAAAATGTCATCGAAAGGCATCGTCGTTTAGGAAAGGCCACGAGG-TAAGTAAAAGCT---------AATAGCCGATTTTACAGTATAGGAAAGATATATAATTAGACATTATTGTCCTCTTGTAACAATGAGCAGACTTGCTGTCGAAG

			>Tillandsia_cacticola
			????????????????????????????????????????????????????????????????????????????????????????????????????????????????????????????????????????????????????????????????????????????????????????????????????????????????????????????????????????????????????????????????????????????????????????????????????????????????????????????????????????????????????????????????????????????????

			>Tillandsia_calcicola
			????????????????????????????????????????????????????????????????????????????????????????????????????????????????????????????????????????????????????????????????????????????????????????????????????????????????????????????????????????????????????????????????????????????????????????????????????????????????????????????????????????????????????????????????????????????????

			>Tillandsia_califanii
			????????????????????????????????????????????????????????????????????????????????????????????????????????????????????????????????????????????????????????????????????????????????????????????????????????????????????????????????????????????????????????????????????????????????????????????????????????????????????????????????????????????????????????????????????????????????

			>Tillandsia_caliginosa
			TCGCAGAAAGCACTTTCCCTGCCTACTGGAATGGGCATTGTTTGCGCGAGCCCGAAAGCTCTGGAAGCTTCCAAAAATGCCAAGTCGGTCAGAGTTTTCTTCGACTGGAATGACTACTTGAAGTTCTACAAGCTTGGAACTTACTGGCCGTAACGCCTTCCATCCAACTACTTTACGGGCTAAGAGCAGCTTTGGATCTCATCTTTGAGGAGGGGCTCGAAAATGTCATCGAAAGGCATCGTCGTTTAGGAAAGGCCACGAGG-TAAGTATAAGCT--------AAATAGCCAATTTTACAGTATAGAAAAGATATATAATTAGCCTTTATTGT-----------AATGAGCAGACTTGCTGTCGAAG

			>Tillandsia_calothyrsus
			????????????????????????????????????????????????????????????????????????????????????????????????????????????????????????????????????????????????????????????????????????????????????????????????????????????????????????????????????????????????????????????????????????????????????????????????????????????????????????????????????????????????????????????????????????????????

			>Tillandsia_caloura
			????????????????????????????????????????????????????????????????????????????????????????????????????????????????????????????????????????????????????????????????????????????????????????????????????????????????????????????????????????????????????????????????????????????????????????????????????????????????????????????????????????????????????????????????????????????????

			>Tillandsia_capillaris
			TCGCAGAAAGCACTTTCCCTGCCTACTGGAATGGGCATTGTTTGCGCGAGCCCGAAAGCTCTGGAAGCTTCCAAAAATGCCAAGTCGGTCAGAGTTTTCTTCGACTGGAATGACTACTTGAAGTTCTACAAGCTTGGAACTTACTGGCCGTAACGCCTTCCATCCAACTACTTTACGGGCTAAGAGCAGCTTTGGATCTCATCTTTGAGGAGGGCCTCGAAAATGTCATCGAAAGGCATCGTCGTTTAGGAAAGGCCACGAGG-TAAGTATAAGCT--------AAATAGCCAATTTYACAGTATAGGAAAGATATGTAATTAGCCATTATTGTCCTCTTGTAACAATGAGCAGACTTGCTGTCGAAG

			>Tillandsia_capillaris_f._virescens
			TCGCAGAAAGCACTTTCCCTGCCTACTGGAATGGGCATTGTTTGCGCGAGCCCGAAAGCTCTGGAAGCTTCCAAAAATGCCAAGTCGGTCAGAGTTTTCTTCGACTGGAATGACTACTTGAAGTTCTACAAGCTTGGAACTTACTGGCCGTAACGCCTTCCATCCAACTACTTTACGGGCTAAGAGCAGCTTTGGATCTCATCTTTGAGGAGGGCCTCGAAAATGTCATCGAAAGGCATCGTCGTTTAGGAAAGGCCACGAGG-TAAGTATAAGCT--------AAATAGCCGATTTCACAGTATAGGAAAGATATGTAATTAGCCATTATTGTCCTCTTGTAACAATGAGCA???????????????

			>Tillandsia_capitata
			TCGCAGAAAGCACTTTCCCTGCCTACTGGAATGGGCATTGTTTGCGCGAGCCCGAAAGCTCTGGAAGCTTCCAAAAATGCCAAGTCGGTCAGAGTTTTCTTCGACTGGAATGACTACTTGAAGTTCTACAAGCTTGGAACTTACTGGCCRTAACGCCTTCCATCCAACTACTCTACGGGCTAAGAGCAGCTTTGGATCTCATCTTTGAGGAGGGGCTCGAAAATGTCATCGAAAGGCATCGTCGTTTAGGAAAGGCCACGAGG-TAAGTAAAAGCT--------AAATAGCCGATTTTACAGTACAGGAAAGATATATAATTAGGCATTATTTTCCTCTTGTAACAATGAGCAGACTTGCTGTCGAAG

			>Tillandsia_caput-medusae
			????????????????????????????????????????????????????????????????????????????????????????????????????????????????????????????????????????????????????????????????????????????????????????????????????????????????????????????????????????????????????????????????????????????????????????????????????????????????????????????????????????????????????????????????????????????????

			>Tillandsia_cardenasii
			TCGCAGAAAGCACTTTCCCTGCCTACTGGAATGGGCATTGTTTGYGCGAGCCCGAAAGCTCTGGAAGCTTCCAAAAATGCCAAGTCGGTTAGAGTTTTCTTCGACTGGAATGACTACTTGAAGTTCTACAAGCTTGGAACTTACTGGCCGTAACRCCTTCCATCCAACTACTTTACGGGCTAAGAGCAGCTTTGGATCTCATCTTTGAGGAGGGGCTCGAAAATGTCATCGAAAGGCATCGTCGTTTAGGAAAGGCCACGAGG-TAAGTATAAGCT--------AAATAGCCGATTTTACAGTATAGGAAAGATATGTAATTAGCCATTATTGT-----------AATGAGCAGACTTGCTGTCGAAG

			>Tillandsia_carlos-hankii
			????????????????????????????????????????????????????????????????????????????????????????????????????????????????????????????????????????????????????????????????????????????????????????????????????????????????????????????????????????????????????????????????????????????????????????????????????????????????????????????????????????????????????????????????????????????????

			>Tillandsia_carminea
			????????????????????????????????????????????????????????????????????????????????????????????????????????????????????????????????????????????????????????????????????????????????????????????????????????????????????????????????????????????????????????????????????????????????????????????????????????????????????????????????????????????????????????????????????????????????

			>Tillandsia_carnosa
			????????????????????????????????????????????????????????????????????????????????????????????????????????????????????????????????????????????????????????????????????????????????????????????????????????????????????????????????????????????????????????????????????????????????????????????????????????????????????????????????????????????????????????????????????????????????

			>Tillandsia_caulescens
			????????????????????????????????????????????????????????????????????????????????????????????????????????????????????????????????????????????????????????????????????????????????????????????????????????????????????????????????????????????????????????????????????????????????????????????????????????????????????????????????????????????????????????????????????????????????

			>Tillandsia_cereicola
			????????????????????????????????????????????????????????????????????????????????????????????????????????????????????????????????????????????????????????????????????????????????????????????????????????????????????????????????????????????????????????????????????????????????????????????????????????????????????????????????????????????????????????????????????????????????

			>Tillandsia_chapeuensis
			????????????????????????????????????????????????????????????????????????????????????????????????????????????????????????????????????????????????????????????????????????????????????????????????????????????????????????????????????????????????????????????????????????????????????????????????????????????????????????????????????????????????????????????????????????????????

			>Tillandsia_chiletensis
			????????????????????????????????????????????????????????????????????????????????????????????????????????????????????????????????????????????????????????????????????????????????????????????????????????????????????????????????????????????????????????????????????????????????????????????????????????????????????????????????????????????????????????????????????????????????

			>Tillandsia_chlorophylla
			????????????????????????????????????????????????????????????????????????????????????????????????????????????????????????????????????????????????????????????????????????????????????????????????????????????????????????????????????????????????????????????????????????????????????????????????????????????????????????????????????????????????????????????????????????????????

			>Tillandsia_circinnatoides
			????????????????????????????????????????????????????????????????????????????????????????????????????????????????????????????????????????????????????????????????????????????????????????????????????????????????????????????????????????????????????????????????????????????????????????????????????????????????????????????????????????????????????????????????????????????????

			>Tillandsia_cochabambae
			????????????????????????????????????????????????????????????????????????????????????????????????????????????????????????????????????????????????????????????????????????????????????????????????????????????????????????????????????????????????????????????????????????????????????????????????????????????????????????????????????????????????????????????????????????????????

			>Tillandsia_coinaensis
			????????????????????????????????????????????????????????????????????????????????????????????????????????????????????????????????????????????????????????????????????????????????????????????????????????????????????????????????????????????????????????????????????????????????????????????????????????????????????????????????????????????????????????????????????????????????

			>Tillandsia_comarapaensis
			????????????????????????????????????????????????????????????????????????????????????????????????????????????????????????????????????????????????????????????????????????????????????????????????????????????????????????????????????????????????????????????????????????????????????????????????????????????????????????????????????????????????????????????????????????????????

			>Tillandsia_comitanensis
			????????????????????????????????????????????????????????????????????????????????????????????????????????????????????????????????????????????????????????????????????????????????????????????????????????????????????????????????????????????????????????????????????????????????????????????????????????????????????????????????????????????????????????????????????????????????

			>Tillandsia_complanata
			????????????????????????????????????????????????????????????????????????????????????????????????????????????????????????????????????????????????????????????????????????????????????????????????????????????????????????????????????????????????????????????????????????????????????????????????????????????????????????????????????????????????????????????????????????????????

			>Tillandsia_comulcoensis
			TCGCAGAAAGCACTTTCCCTGCCTACTGGAATGGGCATTGTTTGCGCGAGCCCGAAAGCGCTGGAAGCTTCCAAAAATGCCAAGTCGGTCAGAGTTTTCTTCGACTGGAATGACTACTTGAAGTTCTACAAGCTTGGAACTTACTGGCCGTAACGCCTTCCATCCAACTACTCTACGGGCTAAGAGCAGCTTTGGATCTCATCTTCGAGGAGGGGCTCGAAAATGTCATCGAAAGGCATCGTCGTTTAGGAAAGGCCACGAGG-TAAGTAAAAGCT--------AAATAGCCGATTTTACAGTATAGGAAAGATATATAATTAGGCATTATTGTCCTCTTGTAACAATGAGCAGACTTGCTGTCGAAG

			>Tillandsia_concolor
			TCGCAGAAAGCACTTTCCCTGCCTACTGGAATGGGCATTGTTTGCGCGAGCCCGAAAGCTCTGGAAGCTTCCAAAAATGCCAAGTCGGTCAGAGTTTTCTTCGACTGGAATGACTACTTGAAGTTCTACAAGCTTGGAACTTACTGGCCGTAACGCCTTCCATCCAACTACTCTACGGCCTAAGAGCAGCTTTGGATCTCATCTTTGAGGAGGGGCTCGAAAATGTCATCGAAAGGCATCGTCGTTTAGGAAAGGCCACGAGGTTAAGTAAAACCT--------AAATAGCCGATTTTACAGTATAGGAAAGATATGTAATTAGGCATTATTGTCCTCTTGTAACAATGAGCAGACTTGCTGTCGAAG

			>Tillandsia_cossonii
			????????????????????????????????????????????????????????????????????????????????????????????????????????????????????????????????????????????????????????????????????????????????????????????????????????????????????????????????????????????????????????????????????????????????????????????????????????????????????????????????????????????????????????????????????????????????

			>Tillandsia_cryptantha
			????????????????????????????????????????????????????????????????????????????????????????????????????????????????????????????????????????????????????????????????????????????????????????????????????????????????????????????????????????????????????????????????????????????????????????????????????????????????????????????????????????????????????????????????????????????????

			>Tillandsia_cucaensis
			????????????????????????????????????????????????????????????????????????????????????????????????????????????????????????????????????????????????????????????????????????????????????????????????????????????????????????????????????????????????????????????????????????????????????????????????????????????????????????????????????????????????????????????????????????????????

			>Tillandsia_dasyliriifolia
			????????????????????????????????????????????????????????????????????????????????????????????????????????????????????????????????????????????????????????????????????????????????????????????????????????????????????????????????????????????????????????????????????????????????????????????????????????????????????????????????????????????????????????????????????????????????

			>Tillandsia_delicata
			TCGCAGAAAGCACTTTCCCTGCCTACTGGAATGGGCATTGTTTGCGCGAGCCCGAAAGCTCTGGAAGCTTCCAAAAATGCCAAGTCGGTCAGAGTTTTCTTCGACTGGAATGACTACTTGAAGTTCTACAAGCTTGGAACTTACTGGCCGTAACGCCTTCCATCCAACTACTCTACGGGCTAAGAGCAGCTTTGGATCTCATCTTTGAGGAGGGGCTCGAAAATGTCATCGAAAGGCATCGTCGTTTAGGAAAGGCCACGAGG-TAAGTAAAAGCT--------AAATAGCCGATTTTACAGTACAGGAAAGATATATAATTAGGCATTATTTTCCTCTTGTAACAATGAGCAGACTTGCTGTGGAAG

			>Tillandsia_demissa
			????????????????????????????????????????????????????????????????????????????????????????????????????????????????????????????????????????????????????????????????????????????????????????????????????????????????????????????????????????????????????????????????????????????????????????????????????????????????????????????????????????????????????????????????????????????????

			>Tillandsia_denudata
			????????????????????????????????????????????????????????????????????????????????????????????????????????????????????????????????????????????????????????????????????????????????????????????????????????????????????????????????????????????????????????????????????????????????????????????????????????????????????????????????????????????????????????????????????????????????

			>Tillandsia_denudata_var._vivipara
			????????????????????????????????????????????????????????????????????????????????????????????????????????????????????????????????????????????????????????????????????????????????????????????????????????????????????????????????????????????????????????????????????????????????????????????????????????????????????????????????????????????????????????????????????????????????

			>Tillandsia_deppeana
			????????????????????????????????????????????????????????????????????????????????????????????????????????????????????????????????????????????????????????????????????????????????????????????????????????????????????????????????????????????????????????????????????????????????????????????????????????????????????????????????????????????????????????????????????????????????

			>Tillandsia_didisticha
			TCGCAGAAAGCACTTTCCCTGCCTACTGGAATGGGCATTGTTTGCGCGAGCCCGAAAGCTCTGGAAGCTTCCAAAAATGCCAAGTCGGTCAGAGTTTTCTTCGACTGGAATGACTACTTGAAGTTCTACAAGCTTGGAACTTACTGGCCGTAACGCCTTCCATCCAACTACTTTACGGGCTAAGAGCAGCTTTGGATCTCATCTTTGAGGAGGGGCTCGAAAATGTCATCGAAAGGCATCGTCGTTTAGGAAAGGCCACGAGG-TAAGTATAAGCT--------AAATAGCCGATTTTACAGTATAGGAAAGATATGTAATTAGCCATTCTTGT-----------AATGAGCAGACTTGCTGTC????

			>Tillandsia_diguetii
			????????????????????????????????????????????????????????????????????????????????????????????????????????????????????????????????????????????????????????????????????????????????????????????????????????????????????????????????????????????????????????????????????????????????????????????????????????????????????????????????????????????????????????????????????????????????

			>Tillandsia_disticha
			TCGCAGAAAGCACTTTCACTGCCTACTGGAATGGGCATTGTTTGCGCGAGCCCGAAAGCTCTGGAAGCTTCCAAAAATGCCAAGTCGGTCAGAGTTTTCTTCGACTGGAATGACTACTTGAAGTTCTACAAGCTTGGAACTTACTGGCCGTAACGCCTTCCATCCAACTACTTTACGGGCTAAGAGCAGCTTTGGATCTCATCTTTGAGGAGGGGCTCGAAAATGTCATCGAAAGGCATCGTCGTTTAGGAAAGGCCACGAGG-TAAGTA-AAGCT--------AAATAGCCGATTTTGCGGTATAGGAAAGATATATAATTAGTCATTATTACCCTCTTGTAACAATGAGCAGACTTGCTGTCGAAG

			>Tillandsia_divaricata
			????????????????????????????????????????????????????????????????????????????????????????????????????????????????????????????????????????????????????????????????????????????????????????????????????????????????????????????????????????????????????????????????????????????????????????????????????????????????????????????????????????????????????????????????????????????????

			>Tillandsia_dugesii
			????????????????????????????????????????????????????????????????????????????????????????????????????????????????????????????????????????????????????????????????????????????????????????????????????????????????????????????????????????????????????????????????????????????????????????????????????????????????????????????????????????????????????????????????????????????????

			>Tillandsia_dura
			????????????????????????????????????????????????????????????????????????????????????????????????????????????????????????????????????????????????????????????????????????????????????????????????????????????????????????????????????????????????????????????????????????????????????????????????????????????????????????????????????????????????????????????????????????????????

			>Tillandsia_duratii
			????????????????????????????????????????????????????????????????????????????????????????????????????????????????????????????????????????????????????????????????????????????????????????????????????????????????????????????????????????????????????????????????????????????????????????????????????????????????????????????????????????????????????????????????????????????????

			>Tillandsia_ecarinata
			????????????????????????????????????????????????????????????????????????????????????????????????????????????????????????????????????????????????????????????????????????????????????????????????????????????????????????????????????????????????????????????????????????????????????????????????????????????????????????????????????????????????????????????????????????????????

			>Tillandsia_edithae
			TCGCAGAAAGCACTTTCCCTGCCTACTGGAATGGGCATTGTTTGCGCGAGCCCGAAAGCTCTAGAAGCTTCCAAAAATGCCAAGTCGGTCAGAGTTTTCTTCGACTGGAATGACTACTTGAAGTTCTACAAGCTTGGAACTTACTGGCCGTAACGCCTTCCATCCAACTACTTTACGGGCTAAGAGCAGCTTTGGATCTCATCTTTGAGGAAGGGCTCGAAAATGTCATCGATAGGCATCGTCGTTTAGGAAAGGCCACGAGG-TAAGTATAAGCT--------AAATAGTCGATTTTACAGTATAGGAAAGATATATAATTAGCCATTATTGT-----------AATGAGCA???????????????

			>Tillandsia_ehlersiana
			TCGCAGAAAGCACTTTCCCTGCCTACTGGAATGGGCATTGTTTGCGCGAGCCCGAAAGCGCTGGAAGCTTCCAAAAATGCCAAGTCGGTCAGAGTTTTCTTCGACTGGAATGACTACTTGAAGTTCTACAAGCTTGGAACTTACTGGCCGTAACGCCTTCCATCCAACTACTCTACGGGCTAAGAGCAGCTTTGGATCTCATCTTTGAGGAGGGGCTCGAAAATGTCATCGAAAGGCATCGTCGTTTAGGAAAGGCCACGAGG-TAAGTAAAAGCT---------AATAGCCGATTTTACAGTACAGGAAAGATATATAATTAGGCATTATTTTCCTCTTGTAACAATGAGCAGACTTGCTGTCGAAG

			>Tillandsia_eizii
			????????????????????????????????????????????????????????????????????????????????????????????????????????????????????????????????????????????????????????????????????????????????????????????????????????????????????????????????????????????????????????????????????????????????????????????????????????????????????????????????????????????????????????????????????????????????

			>Tillandsia_elongata
			????????????????????????????????????????????????????????????????????????????????????????????????????????????????????????????????????????????????????????????????????????????????????????????????????????????????????????????????????????????????????????????????????????????????????????????????????????????????????????????????????????????????????????????????????????????????

			>Tillandsia_elusiva
			????????????????????????????????????????????????????????????????????????????????????????????????????????????????????????????????????????????????????????????????????????????????????????????????????????????????????????????????????????????????????????????????????????????????????????????????????????????????????????????????????????????????????????????????????????????????

			>Tillandsia_erubescens
			????????????????????????????????????????????????????????????????????????????????????????????????????????????????????????????????????????????????????????????????????????????????????????????????????????????????????????????????????????????????????????????????????????????????????????????????????????????????????????????????????????????????????????????????????????????????

			>Tillandsia_espinosae
			TCGCAGAAAGCACTTTCCCTGCCTACTGGAATGGGCATTGTTTGCGCGAGCCCGAAAGCTCTAGAAGCTTCCAAAAATGCCAAGTCGGTCAGAGTTTTCTTCGACTGGAATGACTACTTGAAGTTCTACAAGCTTGGAACTTACTGGCCGTAACGCCTTCCATCCAACTACTCTATGGGCTAAGAGCAGCTTTGGATCTCATCTTTGAGGAGGGGCTCGAAAATGTCATCGAAAGGCATCGCCGTTTAGGAAAGGCCACGAGG-TAAGTA-AAGCT--------ATATAGCCGATCTTAC--AATGGCAGAGACGTCTAGTTGCATATTATTTC------ATAACAA?????????????????????

			>Tillandsia_esseriana
			TCGCAGAAAGCACTTTCCCTGCCTACTGGAATGGGCATTGTTTGCGCGAGCCCGAAAGCTCTGGAAGCTTCCAAAAATGCCAAGTCGGTCAGAGTTTTCTTCGACTGGAATGACTACTTGAAGTTCTACAAGCTTGGAACTTACTGGCCGTAACGCCTTCCATCCAACTACTTTACGGGCTAAGAGCAGCTTTGGATCTCATCTTTGAGGAGGGCCTCGAAAATGTCATCGAAAGACATCGTCGTTTAGGAAAGGCCACGAGG-TAAGTA-AAGCT--------AAATAGCCGATTTTACAGTATAGGAAAGATATGTAATTAGCCATTATTGTCCTCTTGTAACAATGAGCAGACTTGCTGTCGAAG

			>Tillandsia_exserta
			????????????????????????????????????????????????????????????????????????????????????????????????????????????????????????????????????????????????????????????????????????????????????????????????????????????????????????????????????????????????????????????????????????????????????????????????????????????????????????????????????????????????????????????????????????????????

			>Tillandsia_extensa
			????????????????????????????????????????????????????????????????????????????????????????????????????????????????????????????????????????????????????????????????????????????????????????????????????????????????????????????????????????????????????????????????????????????????????????????????????????????????????????????????????????????????????????????????????????????????

			>Tillandsia_fasciculata
			????????????????????????????????????????????????????????????????????????????????????????????????????????????????????????????????????????????????????????????????????????????????????????????????????????????????????????????????????????????????????????????????????????????????????????????????????????????????????????????????????????????????????????????????????????????????

			>Tillandsia_fasciculata_var._densispica
			????????????????????????????????????????????????????????????????????????????????????????????????????????????????????????????????????????????????????????????????????????????????????????????????????????????????????????????????????????????????????????????????????????????????????????????????????????????????????????????????????????????????????????????????????????????????

			>Tillandsia_fendleri
			TCGCAGAAAGCACTTTCCCTGCCTACTGGAATGGGCATTGTTTGCGCGAGCCCGAAAGCTCTAGAAGCTTCCAAAAATGCCAAGTCGGTCAGAGTTTTCTTCGACTGGAATGACTACTTGAAGTTCTACAAGCTCGGAACTTACTGGCCGTAACGCCTTCCATCCAACTACTTTACGGGCTAAGAGCAGCTTTGGATCTCATCTTTGAGGAGGGGCTCGAAAATGTCATCGAAAGGCATCGTCGTTTAGGAAAGGCCACGAGG-TAAGTA-AAGCT--------AAATAGCCGATTTTACAGTATAGGAAAGATATATAATTAGGCATTATTGTCCTSTTGTAACAATGAGCAGACTTGCTGTCGAAG

			>Tillandsia_ferreyrae
			????????????????????????????????????????????????????????????????????????????????????????????????????????????????????????????????????????????????????????????????????????????????????????????????????????????????????????????????????????????????????????????????????????????????????????????????????????????????????????????????????????????????????????????????????????????????

			>Tillandsia_ferrisiana
			TCGCAGAAAGCACTTTCCCTGCCTACTGGAATGGGCATTGTTTGCGCGAGCCCGAAAGCTCTGGAAGCTTCCAAAAATGCCAAGTCGGTCAGAGTTTTCTTCGACTGGAATGACTACTTGAAGTTCTACAAGCTTGGAACTTACTGGCCGTAACGCCTTCCATCCAACTACTCTACGGGCTAAGAGCAGCTTTGGATCTCATCTTCGAGGAGGGGCTCGAAAATGTCATCGAAAGGCATCGTCGTTTAGGAAAGGCCACGAGG-TAAGTAAAAGCT--------AAATAGCCGATTTTACAGTATAGGAAAGATATAAAATTAGGCATTATTGTCCTCTTGTAACAATGAGCAGACTTGCTGTCGAAG

			>Tillandsia_festucoides
			????????????????????????????????????????????????????????????????????????????????????????????????????????????????????????????????????????????????????????????????????????????????????????????????????????????????????????????????????????????????????????????????????????????????????????????????????????????????????????????????????????????????????????????????????????????????

			>Tillandsia_filifolia
			????????????????????????????????????????????????????????????????????????????????????????????????????????????????????????????????????????????????????????????????????????????????????????????????????????????????????????????????????????????????????????????????????????????????????????????????????????????????????????????????????????????????????????????????????????????????

			>Tillandsia_flabellata
			????????????????????????????????????????????????????????????????????????????????????????????????????????????????????????????????????????????????????????????????????????????????????????????????????????????????????????????????????????????????????????????????????????????????????????????????????????????????????????????????????????????????????????????????????????????????

			>Tillandsia_flavobracteata
			????????????????????????????????????????????????????????????????????????????????????????????????????????????????????????????????????????????????????????????????????????????????????????????????????????????????????????????????????????????????????????????????????????????????????????????????????????????????????????????????????????????????????????????????????????????????

			>Tillandsia_flexuosa
			????????????????????????????????????????????????????????????????????????????????????????????????????????????????????????????????????????????????????????????????????????????????????????????????????????????????????????????????????????????????????????????????????????????????????????????????????????????????????????????????????????????????????????????????????????????????

			>Tillandsia_floribunda
			????????????????????????????????????????????????????????????????????????????????????????????????????????????????????????????????????????????????????????????????????????????????????????????????????????????????????????????????????????????????????????????????????????????????????????????????????????????????????????????????????????????????????????????????????????????????

			>Tillandsia_foliosa
			????????????????????????????????????????????????????????????????????????????????????????????????????????????????????????????????????????????????????????????????????????????????????????????????????????????????????????????????????????????????????????????????????????????????????????????????????????????????????????????????????????????????????????????????????????????????

			>Tillandsia_fresnilloensis
			????????????????????????????????????????????????????????????????????????????????????????????????????????????????????????????????????????????????????????????????????????????????????????????????????????????????????????????????????????????????????????????????????????????????????????????????????????????????????????????????????????????????????????????????????????????????

			>Tillandsia_friesii
			TCGCAGAAAGCACTTTCCCTGCCTACTGGAATGGGCATTGTTTGCGCGAGCCCGAAAGCTCTGGAAGCTTCCAAAAATGCCAAGTCGGTCAGAGTTTTCTTCGACTGGAATGACTACTTGAAGTTCTACAAGCTTGGAACTTACTGGCCGTAACGCCTTCCATCCAACTACTTTACGGGCTAAGAGCAGCTTTGGATCTCATCTTTGAGGAGGGCCTCGAAAATGTCATCGAAAGGCATCGTCGTTTAGGAAAGGCCACGAGG-TAAGTATAAGCT--------AAATAGCCGATTTTACAGTATAGGAAAGATATGTAATTAGCCATTATTGT-----------AATGAGCAGACTTGCTGTCGAAG

			>Tillandsia_fuchsii
			????????????????????????????????????????????????????????????????????????????????????????????????????????????????????????????????????????????????????????????????????????????????????????????????????????????????????????????????????????????????????????????????????????????????????????????????????????????????????????????????????????????????????????????????????????????????

			>Tillandsia_funckiana
			????????????????????????????????????????????????????????????????????????????????????????????????????????????????????????????????????????????????????????????????????????????????????????????????????????????????????????????????????????????????????????????????????????????????????????????????????????????????????????????????????????????????????????????????????????????????

			>Tillandsia_funckiana_var._recurvifolia
			????????????????????????????????????????????????????????????????????????????????????????????????????????????????????????????????????????????????????????????????????????????????????????????????????????????????????????????????????????????????????????????????????????????????????????????????????????????????????????????????????????????????????????????????????????????????

			>Tillandsia_funebris
			TCGCAGAAAGCACTTTCCCTGCCTACTGGAATGGGCATTGTTTGCGCGAGCCCGAAAGCTCTGGAAGCTTCCAAAAATGCCAAGTCGGTTAGAGTTTTCTTCGACTGGAATGACTACTTGAAGTTCTACAAGCTTGGAACTTACTGGCCGTAACGCCTTCCATCCAACTACTTTACGGGCTAAGAGCAGCTTTGGATCTCATCTTTGAGGAGGGCCTCGAAAATGTCATCGAAAGGCATCGTCGTTTAGGAAAGGCCACGAGG-TAAGTATAAGCT--------AAATAGCCGATTTTACAGTATAGGAAAGATATGTAATTAGCCATTATTGTCCTCTTGTAACAATGAGCAGACTTGCTGTCGAAG

			>Tillandsia_gardneri
			????????????????????????????????????????????????????????????????????????????????????????????????????????????????????????????????????????????????????????????????????????????????????????????????????????????????????????????????????????????????????????????????????????????????????????????????????????????????????????????????????????????????????????????????????????????????

			>Tillandsia_gardneri_var._rupicola
			????????????????????????????????????????????????????????????????????????????????????????????????????????????????????????????????????????????????????????????????????????????????????????????????????????????????????????????????????????????????????????????????????????????????????????????????????????????????????????????????????????????????????????????????????????????????

			>Tillandsia_geissei
			TCGCAGAAAGCACTTTCCCTGCCTACTGGAATGGGCATTGTTTGCGCGAGCCCGAAAGCTCTAGAAGCTTCCAAAAATGCCAAGTCGGTCAGAGTTTTCTTCGACTGGAATGACTACTTGAAGTTCTACAAGCTTGGAACTTACTGGCCGTAACGCCTTCCATCCAACTACTTTACGGGCTAAGAGCAGCTTTGGATCTCATCTTTGAGGAGGGGCTCGAAAATGTCATCGAAAGGCATCGTCGTTTAGGAAAGGCCACGAGG-TAAGTA-AAGCT--------AAATAGCCGATTTTACAGTATAGGAAAGATATATGATTAGCCATTATTGT-----------AATGAGCA???????????????

			>Tillandsia_geminiflora
			????????????????????????????????????????????????????????????????????????????????????????????????????????????????????????????????????????????????????????????????????????????????????????????????????????????????????????????????????????????????????????????????????????????????????????????????????????????????????????????????????????????????????????????????????????????????

			>Tillandsia_grandispica
			TCGCAGAAAGCACTTTCCCTGCCTACTGGAATGGGCATTGTTTGCGCGAGCCCGAAAGCTCTAGAAGCTTCCAAAAATGCCAAGTCGGTCAGAGTTTTCTTCGACTGGAATGACTACTTGAAGTTCTACAAGCTTGGAACTTACTGGCCGTAACGCCTTCCATCCAACTACTTTACGGGCTAAGAGCAGCTTTGGATCTCATCTTTGAGGAGGGGCTCGAAAATGTCATCGAAAGACATCGTCGTTTAGGAAAGGCCACGAGG-TAAGTA-AAGCT--------AAATAGCCGATTTTACAGTATAGGAAAGATATATAATTAGGCATTATTGT-----------AATGAGCAGACTTGCTGTCGAAG

			>Tillandsia_grao-mogolensis
			????????????????????????????????????????????????????????????????????????????????????????????????????????????????????????????????????????????????????????????????????????????????????????????????????????????????????????????????????????????????????????????????????????????????????????????????????????????????????????????????????????????????????????????????????????????????

			>Tillandsia_guatemalensis
			TCGCAGAAAGCACTTTCCCTGCCTACTGGAATGGGCATTGTTTGCGCGAGCCCGAAAGCTCTGGAAGCTTCCAAAAATGCCAAGTCGGTCAGAGTTTTCTTCGACTGGAATGACTACTTGAAGTTCTACAAGCTTGGAACTTACTGGCCGTAACGCCTTCCATCCAACTACTCTACGGGCTAAGAGCAGCTTTGGATCTCATCTTTGAGGAGGGGCTCGAAAATGTCATCGAAAGGCATCATCGTTTAGGAAAGGCCACGAGG-TAAGTAAAAGCT---------AATAGCCGATTTTACAGTATAGGAAAGATATATAATTAGGCATTATTGTCCTCTTGTAACAATGAGCAGACTTGCTGTCGAAG

			>Tillandsia_guenther-nolleri
			TCGCAGAAGGCACTTTCCCTGCCTACTGGAATGGGCATTGTTTGCGCGAGCCCGAAAGCTCTGGAAGCTTCCAAAAATGCTAAGTCGGTCAGAGTTTTCTTCGACTGGAATGACTACTTGAAGTTCTACAAGCTTGGAACTTACTGGCCGTAACGCCTTCCATCCAACTACTCTACGGCCTAAGAGCAGCTTTGGATCTCATCTTTGAGGAGGGGCTCGAAAATGTCATCGAAAGGCATCGTCGTTTAGGAAAGGCCACGAGG-TAAGTAAAAGCT--------AAATAGCCGATTTTACAGTATAGGAAAGATATGTAATTAGGCATTATTGTCCTCTTGTAACAATGAGCAGACT???????????

			>Tillandsia_guerreroensis
			TCGCAGAAAGCACTTTCCCTGCCTACTGGAATGGGCATTGTTTGCGCGAGCCCGAAAGCGCTGGAAGCTTCGAAAAATGCCAAGTCGGTCAGAGTTTTCTTCGACTGGAATGACTACTTGAAGTTCTACAAGCTTGGAACTTACTGGCCGTAACGCCTTCCATCCAACTACTCTACGGGCTAAGAGCAGCTTTGGATCTCATCTTTGAGGAGGGGCTTGAAAATGTCATCGAAAGGCATCGTCGTTTAGGAAAGGCCACGAGG-TAAGTAAAAGCT--------AAATAGCCGATTTTACAGTACAGGAAAGATATATAATTAGGCGTTATTTTCCTCTTGTAACAATGAGCAGACTTGCTGTCGAAG

			>Tillandsia_gymnobotrya
			????????????????????????????????????????????????????????????????????????????????????????????????????????????????????????????????????????????????????????????????????????????????????????????????????????????????????????????????????????????????????????????????????????????????????????????????????????????????????????????????????????????????????????????????????????????????

			>Tillandsia_harrisii
			TCGCAGAAAGCACTTTCCCTGCCTACTGGAATGGGCATTGTTTGCGCGAGCCCGAAAGCGCTGGAAGCTTCCAAAAATGCCAAGTCGGTCAGAGTTTTCTTCGACTGGAATGACTACTTGAAGTTCTACAAGCTTGGAACTTACTGGCCGTAACGCCTTCCATCCAACTACTCTACGGGCTAAGAGCAGCTTTGGATCTCATCTTYGAGGAGGGGCTCGAAAATGTCATCGAAAGGCATCGTCGTTTAGGAAAGGCCACGAGG-TAAGTAAAAGCT---------AATAGCCGATTTTACAGTATAGGAAAGATATRTAATTAGGCATTATTGTCCTCTTGTAACAATGAGCAGACTTGCTGTGGAAG

			>Tillandsia_hegeri
			TCGCAGAAAGCACTTTCCCTGCCTACTGGAATGGGCATTGTTTGYGCGAGCCCGAAAGCTCTGGAAGCTTCCAAAAATGCCAAGTCGGTAAGAGTTTTCTTCGACTGGAATGACTACTTGAAGTTCTACAAGCTTGGAACTTACTGGCCGTAACGCCTTCCATCCAACTACTTTACGGGCTAAGAGCAGCTTTGGATCTCATCTTTGAGGAGGGGCTCGAAAATGTCATCGAAAGGCATCGTCGTTTAGGAAAGGCCACGAGG-TAAGTATAAGCT--------AAATAGCCGATTTTACAGTATAGGAAAGATATGTAATTAGCCATTATTGT-----------AATGAGCAGACTTGCTGTCGAAG

			>Tillandsia_heliconioides
			TCGCAGAAAGCACTTTCCCTGCCTACTGGAATGGGCATTGTTTGCGCGAGCCCGAAAGCTCTGGAAGCTTCCAAAAACGCCAAGTCGGTCAGAGTTTTCTTCGACTGGAATGACTACTTGAAGTTCTACAAGCTTGGAACTTACTGGCCGTAACGCCTTCCATCCAACTACTCTACGGGCTAAGAGCAGCTTTGGATCTCATCTTTGAGGAGGGGCTCGAAAATGTCATCGAAAGGCATCGTCGTTTAGGAAAGGCCACGAGG-TAAGTAAAAGCT--------AAATAGCCGATTTCACAGTATAGGAAAGATATGTAATTAGGCATTATTGTCCTCTTGTAACAATGAGCAGACTTGCCGTCGAAG

			>Tillandsia_heteromorpha
			????????????????????????????????????????????????????????????????????????????????????????????????????????????????????????????????????????????????????????????????????????????????????????????????????????????????????????????????????????????????????????????????????????????????????????????????????????????????????????????????????????????????????????????????????????????????

			>Tillandsia_heterophylla
			????????????????????????????????????????????????????????????????????????????????????????????????????????????????????????????????????????????????????????????????????????????????????????????????????????????????????????????????????????????????????????????????????????????????????????????????????????????????????????????????????????????????????????????????????????????????

			>Tillandsia_heubergeri
			????????????????????????????????????????????????????????????????????????????????????????????????????????????????????????????????????????????????????????????????????????????????????????????????????????????????????????????????????????????????????????????????????????????????????????????????????????????????????????????????????????????????????????????????????????????????

			>Tillandsia_hildae
			????????????????????????????????????????????????????????????????????????????????????????????????????????????????????????????????????????????????????????????????????????????????????????????????????????????????????????????????????????????????????????????????????????????????????????????????????????????????????????????????????????????????????????????????????????????????

			>Tillandsia_hofackeri
			????????????????????????????????????????????????????????????????????????????????????????????????????????????????????????????????????????????????????????????????????????????????????????????????????????????????????????????????????????????????????????????????????????????????????????????????????????????????????????????????????????????????????????????????????????????????

			>Tillandsia_hondurensis
			TCGCAGAAAGCACTTTCCCTGCCTACTGGAATGGGCATTGTTTGCGCGAGCCCGAAAGCTCTGGAAGCTTCCAAAAATGCCAAGTCGGTCAGAGTTTTCTTCGACTGGAATGACTACTTGAAATTCTACAAGCTTGGAACTTACTGGCCGTAACGCCTTCCATCCAACTACTCTACGGGCTAAGAGCAGCTTTGGATCTCATCTTTGAGGAGGGGCTCGAAAATGTCATCGAAAGGCATCGTCGTTTAGGAAAGGCCACGAGG-TAAGTAAAAGCT---------AATAGCCGATTTTACAGTATAGGAAAGATATATAATTAGACATTATTGTCCTCTTGTAACAATGAGCAGACTTGCTGTGGAAG

			>Tillandsia_horstii
			????????????????????????????????????????????????????????????????????????????????????????????????????????????????????????????????????????????????????????????????????????????????????????????????????????????????????????????????????????????????????????????????????????????????????????????????????????????????????????????????????????????????????????????????????????????????

			>Tillandsia_hromadnikiana
			????????????????????????????????????????????????????????????????????????????????????????????????????????????????????????????????????????????????????????????????????????????????????????????????????????????????????????????????????????????????????????????????????????????????????????????????????????????????????????????????????????????????????????????????????????????????

			>Tillandsia_huamelulaensis
			????????????????????????????????????????????????????????????????????????????????????????????????????????????????????????????????????????????????????????????????????????????????????????????????????????????????????????????????????????????????????????????????????????????????????????????????????????????????????????????????????????????????????????????????????????????????

			>Tillandsia_huarazensis
			TCGCAGAAAGCACTTTCCCTGCCTACTGGAATGGGCATTGTTTGCGCGAGCCCGAAAGCTCTAGAAGCTTCCAAAAATGCCAAGTCGGTCAGAGTTTTCTTCGACTGGAATGACTACTTGAAGTTCTACAAGCTCGGAACTTACTGGCCGTAACGCCTTCCATCCAACTACTTTACGGGCTAAGAGCAGCTTTGGATCTCATCTTTGAGGAGGGGCTCGAAAATGTCATCGAAAGGCATCGTCGTTTAGGAAAGGCCACGAGG-TAAGTA-AAGCT--------AAATAGCCGATTTTACAGTATAGGAAAGATATATAATTAGGCATTATTGTCCTCTTGTAACAATGAGCAGACTTGCTGTCG???

			>Tillandsia_humilis
			TCGCAGAAAGCACTTTCCCTGCCTACTGGAATGGGCATTGTTTGCGCGAGCCCGAAAGCTCTAGAAGCTTCCAAAAATGCCAAGTCGGTCAGAGTTTTCTTCGACTGGAATGACTACTTGAAGTTCTACAAGCTTGGAACTTACTGGCCGTAACGCCTTCCATCCAACTACTTTACGGGCTAAGAGCAGCTTTGGATCTCATCTTTGAGGAGGGGCTCGAAAATGTCATCGAAAGGCATCATCGTTTAGGAAAGGCCACGAGG-TAAGTA---------------AATAGCCGATTTTACAGTATAGGAAAGATATATAATTAGTTTTTATTAG-CATTTATAATAATGAGCAGACTTGCTGTCGAAG

			>Tillandsia_imperialis
			????????????????????????????????????????????????????????????????????????????????????????????????????????????????????????????????????????????????????????????????????????????????????????????????????????????????????????????????????????????????????????????????????????????????????????????????????????????????????????????????????????????????????????????????????????????????

			>Tillandsia_inopinata
			????????????????????????????????????????????????????????????????????????????????????????????????????????????????????????????????????????????????????????????????????????????????????????????????????????????????????????????????????????????????????????????????????????????????????????????????????????????????????????????????????????????????????????????????????????????????

			>Tillandsia_ionantha
			TCGCAGAAAGCACTTTCCCTGCCTACTGGAATGGGCATTGTTTGCGCGAGCCCGAAAGCTCTGGAAGCTTCCAAAAATGCCAAGTCGGTCAGAGTTTTCTTCGACTGGAATGACTACTTGAAGTTCTACAAGCTTGGAACTTACTGGCCGTAACGCCTTCCATCCAACTACTCTACGGGCTAAGAGCAGCTTTGGATCTCATCTTCGAGGAGGGGCTCGAAAATGTCATCGAAAGGCATCGTCGTTTAGGAAAGGCCACGAGG-TAAGTAAAAGCT--------AAATAGCCGATTTCACAGTACAGGAAAGACATATAATTAGGCATTATTTTCCTCTTGTAACAATGAGCAGACTTGCTGTGGAAG

			>Tillandsia_ionochroma
			TCGCAGAAAGCACTTTCCCTGCCTACTGGAATGGGCATTGTTTGCGCGAGCCCGAAAGCTCTAGAAGCTTCCAAAAATGCCAAGTCGGTCAGAGTTTTCTTCGACTGGAATGACTACTTGAAGTTCTACAAGCTTGGAACTTACTGGCCGTAACGCCTTCCATCCAACTACTTTACGGGCTAAGAGCAGCTTTGGATCTCATCTTTGAGGAGGGGCTCGAAAATGTCATCGAAAGGCATCGTCGTTTAGGAAAGGCCACGAGG-TAAGTA-AAGCT------------------TTTTACAGTATAGGAAAGATATATAATTAGGCATTATTGT-----------AATGAGCAGACTTGCTGTCGAAG

			>Tillandsia_ixioides
			????????????????????????????????????????????????????????????????????????????????????????????????????????????????????????????????????????????????????????????????????????????????????????????????????????????????????????????????????????????????????????????????????????????????????????????????????????????????????????????????????????????????????????????????????????????????

			>Tillandsia_izabalensis
			????????????????????????????????????????????????????????????????????????????????????????????????????????????????????????????????????????????????????????????????????????????????????????????????????????????????????????????????????????????????????????????????????????????????????????????????????????????????????????????????????????????????????????????????????????????????

			>Tillandsia_jaliscomonticola
			TCGCAGAAAGCACTTTCCCTGCCTACTGGAATGGGCATTGTTTGCGCGAGCCCGAAAGCTCTGGAAGCTTCCAAAAATGCCAAGTCGGTCAGAGTTTTCTTCGACTGGAATGACTACTTGAAGTTCTACAAGCTTGGAACTTACTGGCCGTAACRCCTTCCATCCAACTACTCTACGGCCTAAGAGCAGCTTTGGATCTCATCTTTGAGGAGGGGCTCGAAAATGTCATCGAAAGGCATCGTCGTTTAGGAAAGGCCACGAGG-TAAGTAAAAGCT--------AAATAGCCGATTTTACAGTATAGGAAAGATATGTAATTAGGCATTATTGTCCTCTTGTAACAATGAGCAGACTTGCTGTC????

			>Tillandsia_jucunda
			????????????????????????????????????????????????????????????????????????????????????????????????????????????????????????????????????????????????????????????????????????????????????????????????????????????????????????????????????????????????????????????????????????????????????????????????????????????????????????????????????????????????????????????????????????????????

			>Tillandsia_jucunda_var._vivida
			????????????????????????????????????????????????????????????????????????????????????????????????????????????????????????????????????????????????????????????????????????????????????????????????????????????????????????????????????????????????????????????????????????????????????????????????????????????????????????????????????????????????????????????????????????????????

			>Tillandsia_juerg-rutschmannii
			????????????????????????????????????????????????????????????????????????????????????????????????????????????????????????????????????????????????????????????????????????????????????????????????????????????????????????????????????????????????????????????????????????????????????????????????????????????????????????????????????????????????????????????????????????????????

			>Tillandsia_juncea
			????????????????????????????????????????????????????????????????????????????????????????????????????????????????????????????????????????????????????????????????????????????????????????????????????????????????????????????????????????????????????????????????????????????????????????????????????????????????????????????????????????????????????????????????????????????????

			>Tillandsia_kammii
			TCGCAGAAAGCACTTTCCCTGCCTACTGGAATGGGCATTGTTTGCGCGAGCCCGAAAGCGCTGGAAGCTTCCAAAAATGCCAAGTCGGTCAGAGTTTTCTTCGACTGGAATGACTACTTGAAGTTCTACAAGCTTGGAACTTACTGGCCGTAACGCCTTCCATCCAACTACTCTACGGGCTAAGAGCAGCTTTGGATCTCATCTTCGAGGAGGGGCTCGAAAATGTCATCGAAAGGCATCGTCGTTTAGGAAAGGCCACGAGG-TAAGTAAAAGCT---------AATAGCCGATTTTACAGTATAGGAAAGATATATAATTAGGCATTATTGTCCTCTTGTAACAATGAGCAGACTTGCTGTGGAAG

			>Tillandsia_karwinskyana
			????????????????????????????????????????????????????????????????????????????????????????????????????????????????????????????????????????????????????????????????????????????????????????????????????????????????????????????????????????????????????????????????????????????????????????????????????????????????????????????????????????????????????????????????????????????????

			>Tillandsia_kauffmannii
			????????????????????????????????????????????????????????????????????????????????????????????????????????????????????????????????????????????????????????????????????????????????????????????????????????????????????????????????????????????????????????????????????????????????????????????????????????????????????????????????????????????????????????????????????????????????

			>Tillandsia_kautskyi
			????????????????????????????????????????????????????????????????????????????????????????????????????????????????????????????????????????????????????????????????????????????????????????????????????????????????????????????????????????????????????????????????????????????????????????????????????????????????????????????????????????????????????????????????????????????????

			>Tillandsia_kegeliana
			TCGCAGAAAGCACTTTCACTGCCTACTGGAATGGGGATTGTTTGCGCGAGCCCGAAGGCTCTGGAAGCTTCTAAAAATGCCAAGTCGGTCAGAGTTTTCTTCGACTGGAATGACTACTTGAAGTTCTACAAGCTTGGAACTTACTGGCCGTAACGCCTTCAATCCAACTACTCTACGGGCTAAGAGCAGCTTTGGATCTCATCTTTGAGGAGGGGCTCGAAAATGTCATCGAAAGGCATCGTCGTTTAGGAAAGGCCACGAGG-TAACTAAAAGC----------------CGATTTTACAGTATAGGAAAGATATATAATTAGTCATTATTGCCCTCTTGTGACAATGAGCAGACTCGCTGTCGAAG

			>Tillandsia_kirchhoffiana
			????????????????????????????????????????????????????????????????????????????????????????????????????????????????????????????????????????????????????????????????????????????????????????????????????????????????????????????????????????????????????????????????????????????????????????????????????????????????????????????????????????????????????????????????????????????????

			>Tillandsia_kirschnekii
			????????????????????????????????????????????????????????????????????????????????????????????????????????????????????????????????????????????????????????????????????????????????????????????????????????????????????????????????????????????????????????????????????????????????????????????????????????????????????????????????????????????????????????????????????????????????

			>Tillandsia_klausii
			????????????????????????????????????????????????????????????????????????????????????????????????????????????????????????????????????????????????????????????????????????????????????????????????????????????????????????????????????????????????????????????????????????????????????????????????????????????????????????????????????????????????????????????????????????????????

			>Tillandsia_koehresiana
			????????????????????????????????????????????????????????????????????????????????????????????????????????????????????????????????????????????????????????????????????????????????????????????????????????????????????????????????????????????????????????????????????????????????????????????????????????????????????????????????????????????????????????????????????????????????

			>Tillandsia_kuehhasii
			????????????????????????????????????????????????????????????????????????????????????????????????????????????????????????????????????????????????????????????????????????????????????????????????????????????????????????????????????????????????????????????????????????????????????????????????????????????????????????????????????????????????????????????????????????????????

			>Tillandsia_lajensis
			????????????????????????????????????????????????????????????????????????????????????????????????????????????????????????????????????????????????????????????????????????????????????????????????????????????????????????????????????????????????????????????????????????????????????????????????????????????????????????????????????????????????????????????????????????????????

			>Tillandsia_lampropoda
			????????????????????????????????????????????????????????????????????????????????????????????????????????????????????????????????????????????????????????????????????????????????????????????????????????????????????????????????????????????????????????????????????????????????????????????????????????????????????????????????????????????????????????????????????????????????

			>Tillandsia_landbeckii
			TCGCAGAAAGCACTTTCCCTGCCTACTGGAATGGGCATTGTTTGCGCGAGCCCGAAAGCTCTGGAAGCTTCCAAAAATGCCAAGTCGGTCAGAGTTTTCTTCGACTGGAATGACTACTTGAAGTTCTACAAGCTTGGAACTTACTGGCCGTAACGCCTTCCATCCAACTACTTTACGGGCTAAGAGCAGCTTTGGATCTCATCTTTGAGGAGGGCCTCGAAAATGTCATCGAAAGGCATCGTCGTTTAGGAAAGGCCACGAGG-TAAGTATAAGCT--------AAATAGCCGATTTTACAGTATAGGAAAGATATGTAATTAGCCATTATTGTCCTCTTGTAACAATGAGCAGACTTGCTGTCGAAG

			>Tillandsia_landbeckii_subsp._andina
			TCGCAGAAAGCACTTTCCCTGCCTACTGGAATGGGCATTGTTTGCGCGAGCCCGAAAGCTCTGGAAGCTTCCAAAAATGCCAAGTCGGTCAGAGTTTTCTTCGACTGGAATGACTACTTGAAGTTCTACAAGCTTGGAACTTACTGGCCGTAACGCCTTCCATCCAACTACTTTACGGGCTAAGAGCAGCTTTGGATCTCATCTTTGAGGAGGGCCTCGAAAATGTCATCGAAAGGCATCGTCGTTTAGGAAAGGCCACGAGG-TAAGTATAAGCT--------AAATAGCCGATTTTACAGTATAGGAAAGATATGTAATTAGCCATTATTGTCCTCTTGTAACAATGAGCA???????????????

			>Tillandsia_landbeckii_var._rigidor
			TCGCAGAAAGCACTTTCCCTGCCTACTGGAATGGGCATTGTTTGCGCGAGCCCGAAAGCTCTGGAAGCTTCCAAAAATGCCAAGTCGGTCAGAGTTTTCTTCGACTGGAATGACTACTTGAAGTTCTACAAGCTTGGAACTTACTGGCCGTAACGCCTTCCATCCAACTACTTTACGGGCTAAGAGCAGCTTTGGATCTCATCTTTGAGGAGGGCCTCGAAAATGTCATCGAAAGGCATCGTCGTTTAGGAAAGGCCACGAGG-TAAGTATAAGCT--------AAATAGCCGATTTTACAGTATAGGAAAGATATGTAATTAGCCATTATTGTCCTCTTGTAACAATGAGCA???????????????

			>Tillandsia_langlasseana
			????????????????????????????????????????????????????????????????????????????????????????????????????????????????????????????????????????????????????????????????????????????????????????????????????????????????????????????????????????????????????????????????????????????????????????????????????????????????????????????????????????????????????????????????????????????????

			>Tillandsia_latifolia_var._divaricata
			????????????????????????????????????????????????????????????????????????????????????????????????????????????????????????????????????????????????????????????????????????????????????????????????????????????????????????????????????????????????????????????????????????????????????????????????????????????????????????????????????????????????????????????????????????????????

			>Tillandsia_laui
			????????????????????????????????????????????????????????????????????????????????????????????????????????????????????????????????????????????????????????????????????????????????????????????????????????????????????????????????????????????????????????????????????????????????????????????????????????????????????????????????????????????????????????????????????????????????

			>Tillandsia_lautneri
			????????????????????????????????????????????????????????????????????????????????????????????????????????????????????????????????????????????????????????????????????????????????????????????????????????????????????????????????????????????????????????????????????????????????????????????????????????????????????????????????????????????????????????????????????????????????

			>Tillandsia_lechneri
			????????????????????????????????????????????????????????????????????????????????????????????????????????????????????????????????????????????????????????????????????????????????????????????????????????????????????????????????????????????????????????????????????????????????????????????????????????????????????????????????????????????????????????????????????????????????

			>Tillandsia_leiboldiana
			????????????????????????????????????????????????????????????????????????????????????????????????????????????????????????????????????????????????????????????????????????????????????????????????????????????????????????????????????????????????????????????????????????????????????????????????????????????????????????????????????????????????????????????????????????????????

			>Tillandsia_lepidosepala
			????????????????????????????????????????????????????????????????????????????????????????????????????????????????????????????????????????????????????????????????????????????????????????????????????????????????????????????????????????????????????????????????????????????????????????????????????????????????????????????????????????????????????????????????????????????????

			>Tillandsia_leucopetala
			????????????????????????????????????????????????????????????????????????????????????????????????????????????????????????????????????????????????????????????????????????????????????????????????????????????????????????????????????????????????????????????????????????????????????????????????????????????????????????????????????????????????????????????????????????????????

			>Tillandsia_limbata
			????????????????????????????????????????????????????????????????????????????????????????????????????????????????????????????????????????????????????????????????????????????????????????????????????????????????????????????????????????????????????????????????????????????????????????????????????????????????????????????????????????????????????????????????????????????????

			>Tillandsia_lorentziana
			TCGCAGAAAGCACTTTCCCTGCCTACTGGAATGGGCATTGTTTGYGCGAGCCCGAAAGCTCTGGAAGCTTCCAAAAATGCCAAGTCGGTCAGAGTTTTCTTCGACTGGAATGACTACTTGAAGTTCTACAAGCTTGGAACTTACTGGCCGTAACGCCTTCCATCCAACTACTTTACGGGCTAAGAGCAGCTTTGGATCTCATCTTTGAGGAGGGGCTCGAAAATGTCATCGAAAGGCATCGTCGTTTAGGAAAGGCCACGAGG-TAAGTATAAGCT--------AAATAGCCGATTTTACAGTATAGGAAAGATATGTAATTAGCCATTATTGT-----------AATGAGCAGACTTGCTGTCGAAG

			>Tillandsia_lotteae
			????????????????????????????????????????????????????????????????????????????????????????????????????????????????????????????????????????????????????????????????????????????????????????????????????????????????????????????????????????????????????????????????????????????????????????????????????????????????????????????????????????????????????????????????????????????????

			>Tillandsia_lucida
			????????????????????????????????????????????????????????????????????????????????????????????????????????????????????????????????????????????????????????????????????????????????????????????????????????????????????????????????????????????????????????????????????????????????????????????????????????????????????????????????????????????????????????????????????????????????

			>Tillandsia_lymanii
			TCGCAGAAAGCACTTTCCCTGCCTACTGGAATGGGCATTGTTTGCGCGAGCCCGAAAGCTCTGGAAGCTTCCAAGAATGCCAAGTCGGTCAGAGTTTTCTTCGACTGGAATGACTACTTGAAGTTCTACAAGCTTGGAACTTACTGGCCGTAACGCCTTCCATACAACTACTCTACGGGCTAAGAGCAGCTTTGGATCTCATCTTTGAGGAGGGGCTCGAAAATGTCATCGAAAGGCATCGTCGTTTAGGAAAGGCCACGAGG-TAAGTAAAAGCT--------AAATAGCCGATTTTACAGTATAGGAAAGATATATAATTAGTCATTATTGTCCTCTTTTAACAATGAG?????????????????

			>Tillandsia_macbrideana
			????????????????????????????????????????????????????????????????????????????????????????????????????????????????????????????????????????????????????????????????????????????????????????????????????????????????????????????????????????????????????????????????????????????????????????????????????????????????????????????????????????????????????????????????????????????????

			>Tillandsia_macdougallii
			????????????????????????????????????????????????????????????????????????????????????????????????????????????????????????????????????????????????????????????????????????????????????????????????????????????????????????????????????????????????????????????????????????????????????????????????????????????????????????????????????????????????????????????????????????????????

			>Tillandsia_maculata
			????????????????????????????????????????????????????????????????????????????????????????????????????????????????????????????????????????????????????????????????????????????????????????????????????????????????????????????????????????????????????????????????????????????????????????????????????????????????????????????????????????????????????????????????????????????????

			>Tillandsia_magnusiana
			TCGCAGAAAGCACTTTCCCTGCCTACTGGAATGGGCATTGTTTGCGCGAGCCCGAAAGCGCTGGAAGCTTCCAAAAATGCCAAGTCGGTCAGAGTTTTCTTCGACTGGAATGACTACTTGAAGTTCTACAAGCTTGGAACTTACTGGCCGTAACGCCTTCCATCCAACTACTCTACGGGCTAAGAGCAGCTTTGGATCTCATCTTCGAGGAGGGGCTCGAAAATGTCATCGAAAGGCATCGTCGTTTAGGAAAGGCCACGAGG-TAAGTAAAAGCT--------AAATAGCCGATTTTACAGCATAGGAAAGATATATAATTAGGCATTATTGTCATCTTGTAACAATGAGCAGACTTGCTGTGGAAG

			>Tillandsia_makoyana
			????????????????????????????????????????????????????????????????????????????????????????????????????????????????????????????????????????????????????????????????????????????????????????????????????????????????????????????????????????????????????????????????????????????????????????????????????????????????????????????????????????????????????????????????????????????????

			>Tillandsia_malzinei
			????????????????????????????????????????????????????????????????????????????????????????????????????????????????????????????????????????????????????????????????????????????????????????????????????????????????????????????????????????????????????????????????????????????????????????????????????????????????????????????????????????????????????????????????????????????????

			>Tillandsia_marconae
			TCGCAGAAAGCACTTTCCCTGCCTACTGGAATGGGCATTGTTTGCGCGAGCCCGAAAGCTCTGGAAGCTTCCAAAAATGCCAAGTCGGTCAGAGTTTTCTTCGACTGGAATGACTACTTGAAGTTCTACAAGCTTGGAACTTACTGGCCGTAACGCCTTCCATCCAACTACTTTACGGGCTAAGAGCAGCTTTGGATCTCATCTTTGAGGAGGGCCTCGAAAATGTCATCGAAAGGCATCGTCGTTTAGGAAAGGCCACGAGG-TAAGTATAAGCT--------AAATAGCCGATTTTACAGTATAGGAAAGATATGTAATTAGCCATTATTGTCCTCTTGTAACAATGAGCA???????????????

			>Tillandsia_marnier-lapostollei
			????????????????????????????????????????????????????????????????????????????????????????????????????????????????????????????????????????????????????????????????????????????????????????????????????????????????????????????????????????????????????????????????????????????????????????????????????????????????????????????????????????????????????????????????????????????????

			>Tillandsia_matudae
			????????????????????????????????????????????????????????????????????????????????????????????????????????????????????????????????????????????????????????????????????????????????????????????????????????????????????????????????????????????????????????????????????????????????????????????????????????????????????????????????????????????????????????????????????????????????

			>Tillandsia_may-patii
			????????????????????????????????????????????????????????????????????????????????????????????????????????????????????????????????????????????????????????????????????????????????????????????????????????????????????????????????????????????????????????????????????????????????????????????????????????????????????????????????????????????????????????????????????????????????

			>Tillandsia_mazatlanensis
			TCGCAGAAGGCACTTTCCCTGCCTACTGGAATGGGCATTGTTTGCGCGAGCCCGAAAGCTCTGGAAGCTTCCAAAAATGCCAAGTCGGTCAGAGTTTTCTTCGACTGGAATGACTACTTGAAGTTCTACAAGCTTGGAACTTACTGGCCGTAACGCCTTCCATCCAACTACTCTACGGCCTAAGAGCAGCTTTGGATCTCATCTTTGAGGAGGGGCTCGAAAATGTCATCGAAAGGCATCGTCGTTTAGGAAAGGCCACGAGG-TAAGTAAAAGCT--------AAATAGCCGATTTTACAGCATAGGAAAGATATATAATTAGGCATTATTGTCCTCTTGTAACAATGAGC????????????????

			>Tillandsia_milagrensis
			????????????????????????????????????????????????????????????????????????????????????????????????????????????????????????????????????????????????????????????????????????????????????????????????????????????????????????????????????????????????????????????????????????????????????????????????????????????????????????????????????????????????????????????????????????????????

			>Tillandsia_mima
			TCGCAGAAAGCACTTTCCCTGCCTACTGGAATGGGCATTGTTTGCGCGAGCCCGAAAGCTCTGGAAGCTTCCAAGAATGCCAAGTCGGTCAGAGTTTTCTTCGACTGGAATGACTACTTGAAGTTCTACAAGCTTGGAACTTACTGGCCGTAACGCCTTCCATACAACTACTCTACGGGCTAAGAGCAGCTTTGGATCTCATCTTTGAGGAGGGGCTCGAAAATGTCATCGAAAGGCATCGTCGTTTAGGAAAGGCCACGAGG-TAAGTAAAAGCT--------AAATAGCCGATTTTACAGTATAGGAAAGATATATAATTAGTCATTATTGTCCTCTTTTAACAATGAGCAGACTCGCTGTCGAAG

			>Tillandsia_mirabilis
			TCGCAGAAAGCACTTTCCCTGCCTACTGGAATGGGCATTGTTTGCGCGAGCCCGAAAGCTCTGGAAGCTTCCAAAAATGCCAAGTCGGTCAGAGTTTTCTTCGACTGGAATGACTACTTGAAGTTCTACAAGCTTGGAACTTACTGGCCGTAACGCCTTCCATCCAACTACTCTACGGGCTAAGAGCAGCTTTGGATCTCATCTTTGAGGAGGGGCTCGAAAATGTCATCGAAAGGCATCGTCGTTTAGGAAAGGCCACGAGG-TAAGTAAAAGCT--------------------TTACAGTATAGGAAAGATATATAATTAGGCATTATTGTCCTCTTGTAACAATGAGCAGACTTGCTGTCGAAG

			>Tillandsia_mollis
			TCGCAGAAAGCACTTTCCCTGCCTACTGGAATGGGCATTGTTTGCGCGAGCCCGAAAGCTCTGGAAGCTTCCAAAAATGCCAAGTCGGTCAGAGTTTTCTTCGACTGGAATGACTACTTGAAGTTCTACAAGCTTGGAACTTACTGGCCGTAACGCCTTCCATCCAACTACTTTACGGGCTAAGAGCAGCTTTGGATCTCATCTTTGAGGAGGGCCTCGAAAATGTCATCGAAAGGCATCGTCGTTTAGGAAAGGCCACGAGG-TAAGTATAAGCT--------AAATAGCCGATTTTACAGTATAGGAAAGATATGTAATTAGCCATTATTGTCCTCTTGTAACAATGAGCAGACTTGCTGTCGAAG

			>Tillandsia_mooreana
			TCGCAGAAAGCACTTTCCCTGCCTACTGGAATGGGCATTGTTTGCGCGAGCCCGAAAGCGCTGGAAGCTTCCAAAAACGCCAAGTCGGTCAGAGTTTTCTTCGACTGGAATGACTACTTGAAGTTCTACAAGCTTGGAACTTACTGGCCGTAACGCCTTCCATCCAACTACTCTACGGGCTAAGAGCAGCTTTGGATCTCATCTTCGAGGAGGGGCTCGAAAATGTCATCGAAAGGCATCGTCGTTTAGGAAAGGCCACGAGG-TAAGTAAAAGTT---------AATAGCCGATTTTACAGTATAGGAAAGATATATAATTAGGCATTATTGTCCTCTTGTAACAATGAGCAGACTTGCTGTCGAAG

			>Tillandsia_moronesensis
			TCGCAGAAAGCACTTTCCCTGCCTACTGGAATGGGCATTGTTTGCGCGAGCCCGAAAGCTCTGGAAGCTTCCAAAAATGCCAAGTCGGTCAGAGTTTTCTTCGACTGGAATGACTACTTGAAGTTCTACAAGCTTGGAACTTACTGGCCGTAACGCCTTCCATCCAACTACTCTACGGGCTAAGAGCAGCTTTGGATCTCATCTTTGAGGAGGGGCTCGAAAATGTCATCGAAAGGCATCGTCGTTTAGGAAAGGCCACGAGG-TAAGTAAAAGCT--------------------TTACAGTATAGGAAAGATATATAATTAGGCATTATTGTCCTCTTGTAACAATGAGCAGACTTGCTGTCGAAG

			>Tillandsia_multicaulis
			????????????????????????????????????????????????????????????????????????????????????????????????????????????????????????????????????????????????????????????????????????????????????????????????????????????????????????????????????????????????????????????????????????????????????????????????????????????????????????????????????????????????????????????????????????????????

			>Tillandsia_myriantha
			????????????????????????????????????????????????????????????????????????????????????????????????????????????????????????????????????????????????????????????????????????????????????????????????????????????????????????????????????????????????????????????????????????????????????????????????????????????????????????????????????????????????????????????????????????????????

			>Tillandsia_nana
			TCGCAGAAAGCACTTTCCCTGCCTACTGGAATGGGCATTGTTTGCGCGAGCCCGAAAGCTCTGGAAGCTTCCAAAAATGCCAAGTCGGTCAGAGTTTTCTTCGACTGGAATGACTACTTGAAGTTCTACAAGCTTGGAACTTACTGGCCGTAACGCCTTCCATCCAACTACTTTACGGGCTAAGAGCAGCTTTGGATCTCATCTTTGAGGAGGGGCTCGAAAATGTCATCGAAAGGCATCGTCGTTTAGGAAAGGCCACGAGG-TAAGTA-ATGCT------------------TTTTACAGTATAGGAAAGATATATAATTAGGCATTATTGT-----------AATGAGCAGACTTGCTGTCGAAG

			>Tillandsia_neglecta
			????????????????????????????????????????????????????????????????????????????????????????????????????????????????????????????????????????????????????????????????????????????????????????????????????????????????????????????????????????????????????????????????????????????????????????????????????????????????????????????????????????????????????????????????????????????????

			>Tillandsia_nicolasensis
			????????????????????????????????????????????????????????????????????????????????????????????????????????????????????????????????????????????????????????????????????????????????????????????????????????????????????????????????????????????????????????????????????????????????????????????????????????????????????????????????????????????????????????????????????????????????

			>Tillandsia_nizandaensis
			TCGCAGAAAGCACTTTCCCTGCCTACTGGAATGGGCATTGTTTGCGCGAGCCCGAAAGCTCTGGAAGCTTCCAAAAATGCCAAGTCTGTCAGAGTTTTCTTCGACTGGAATGACTACTTGAAGTTCTACAAGCTTGGAACTTACTGGCCGTAACGCCTTCCATCCAACTACTCTACGGGCTAAGAGCAGCTTTGGATCTCATCTTCGAGGAGGGGCTCGAAAATGTCATCGAAAGGCATCGTCGTTTAGGAAAGGCCACGAGG-TAAGTAAAAGCT--------AAATAGCCGATTTTACAGTATAGGAAAGAGATATAATTAGGCATTATTGTCCTCTTGTAACAATGAGCAGACTTGCTGTGGAAG

			>Tillandsia_novakii
			????????????????????????????????????????????????????????????????????????????????????????????????????????????????????????????????????????????????????????????????????????????????????????????????????????????????????????????????????????????????????????????????????????????????????????????????????????????????????????????????????????????????????????????????????????????????

			>Tillandsia_nuptialis
			????????????????????????????????????????????????????????????????????????????????????????????????????????????????????????????????????????????????????????????????????????????????????????????????????????????????????????????????????????????????????????????????????????????????????????????????????????????????????????????????????????????????????????????????????????????????

			>Tillandsia_oaxacana
			????????????????????????????????????????????????????????????????????????????????????????????????????????????????????????????????????????????????????????????????????????????????????????????????????????????????????????????????????????????????????????????????????????????????????????????????????????????????????????????????????????????????????????????????????????????????

			>Tillandsia_oblivata
			TCGCAGAAAGCACTTTCCCTGCCTACTGGAATGGGCATTGTTTGCGCGAGCCCGAAAGCTCTAGAAGCTTCCAAAAATGCCAAGTCGGTCAGAGTTTTCTTCGACTGGAATGACTACTTGAAGTTCTACAAGCTCGGAACTTACTGGCCGTAACGCCTTCCATCCAACTACTTTACGGGCTAAGAGCAGCTTTGGATCTCATCTTTGAGGAGGGGCTCGAAAATGTCATCGAAAGGCATCGTCGTTTAGGAAAGGCCACGAGG-TAAGTA-AAGCT--------AAATAGCCGATTTTACAGTATAGGAAAGATATATAATTAGGCATTATTGT-----------AATGAGCAGACTTGCTGTGGAAG

			>Tillandsia_oerstediana
			????????????????????????????????????????????????????????????????????????????????????????????????????????????????????????????????????????????????????????????????????????????????????????????????????????????????????????????????????????????????????????????????????????????????????????????????????????????????????????????????????????????????????????????????????????????????

			>Tillandsia_paleacea
			TCGCAGAAAGCACTTTCCCTGCCTACTGGAATGGGCATTGTTTGCSCGAGCCCGAAAGCTCTGGAAGCTTCCAAAAATGCCAAGTCGGTCAGAGTTTTCTTCGACTGGAATGACTACTTGAAGTTCTACAAGCTTGGAACTTACTGGCCGTAACGCCTTCCATCCAACTACTTTACGGGCTAAGAGCAGCTTTGGATCTCATCTTTGAGGAGGGCCTCGAAAATGTCATCGAAAGGCATCGTCGTTTAGGAAAGGCCACGAGG-TAAGTATAAGCT--------AAAYAGCCGATTTTACAG--YAGGAAAGAGATGTAATTAGCCATTATTGTCCTCTTGTAACAATGAGCAGACTTGCTGTCGAAG

			>Tillandsia_pamelae
			TCGCAGAAAGCACTTTCCCTGCCTACTGGAATGGGCATTGTTTGCGCGAGCCCGAAAGCTCTGGAAGCTTCCAAAAATGCCAAGTCGGTCAGAGTTTTCTTCGACTGGAATGACTAYTTGAAGTTCTACAAGCTYGGAACTTACTGGCCGTAACGCCTTCCATCCAACTACTCTACGGGCTAAGAGCAGCTTTGGATCTCATCTTTGAGGAGGGGCTCGAAAATGTCATCGAAAGGCATCGTCGTTTAGGAAAGGCCACGAGG-TAAGTAAAAGCT--------------------TTACAGTATAGGAAAGATATATAATTAGGCATTATTGTCCTCTTGTAACAATGAACAGACTTGCTGTCGAAG

			>Tillandsia_paniculata
			????????????????????????????????????????????????????????????????????????????????????????????????????????????????????????????????????????????????????????????????????????????????????????????????????????????????????????????????????????????????????????????????????????????????????????????????????????????????????????????????????????????????????????????????????????????????

			>Tillandsia_paraensis
			????????????????????????????????????????????????????????????????????????????????????????????????????????????????????????????????????????????????????????????????????????????????????????????????????????????????????????????????????????????????????????????????????????????????????????????????????????????????????????????????????????????????????????????????????????????????

			>Tillandsia_paraibensis
			????????????????????????????????????????????????????????????????????????????????????????????????????????????????????????????????????????????????????????????????????????????????????????????????????????????????????????????????????????????????????????????????????????????????????????????????????????????????????????????????????????????????????????????????????????????????

			>Tillandsia_paraisoensis
			????????????????????????????????????????????????????????????????????????????????????????????????????????????????????????????????????????????????????????????????????????????????????????????????????????????????????????????????????????????????????????????????????????????????????????????????????????????????????????????????????????????????????????????????????????????????

			>Tillandsia_parryi
			????????????????????????????????????????????????????????????????????????????????????????????????????????????????????????????????????????????????????????????????????????????????????????????????????????????????????????????????????????????????????????????????????????????????????????????????????????????????????????????????????????????????????????????????????????????????

			>Tillandsia_paucifolia
			????????????????????????????????????????????????????????????????????????????????????????????????????????????????????????????????????????????????????????????????????????????????????????????????????????????????????????????????????????????????????????????????????????????????????????????????????????????????????????????????????????????????????????????????????????????????

			>Tillandsia_pinicola
			????????????????????????????????????????????????????????????????????????????????????????????????????????????????????????????????????????????????????????????????????????????????????????????????????????????????????????????????????????????????????????????????????????????????????????????????????????????????????????????????????????????????????????????????????????????????

			>Tillandsia_plagiotropica
			TCGCAGAAAGCACTTTCCCTGCCTACTGGAATGGGCATTGTTTGCGCGAGCCCGAAAGCGCTGGAAGCTTCCAAAAATGCCAAGTCGGTCAGAGTTTTCTTCGACTGGAATGACTACTTGAAGTTCTACAAGCTTGGAACTTACTGGCCGTAACGCCTTCCATCCAACTACTCTACGGGCTAAGAGCAGCTTTGGATCTCATCTTCGAGGAGGGGCTCGAAAATGTCATCGAAAGACATCGTCGTTTAGGAAAGGCCACGAGG-TAAGTAAAAGCT---------AATAGCCGATTTTACAGTATTGGAAAGATATATAATTAGGCATTATTGTCCTCTTGTAACAATGAGCAGACTTGCTGTGGAAG

			>Tillandsia_plumosa
			????????????????????????????????????????????????????????????????????????????????????????????????????????????????????????????????????????????????????????????????????????????????????????????????????????????????????????????????????????????????????????????????????????????????????????????????????????????????????????????????????????????????????????????????????????????????

			>Tillandsia_pohliana
			????????????????????????????????????????????????????????????????????????????????????????????????????????????????????????????????????????????????????????????????????????????????????????????????????????????????????????????????????????????????????????????????????????????????????????????????????????????????????????????????????????????????????????????????????????????????

			>Tillandsia_polita
			TCGCAGAAAGCACTTTCCCTGCCTACTGGAATGGGCATTGTTTGCGCGAGCCCGAAAGCGCTGGAAGCTTCCAAAAATGCCAAGTCGGTCAGAGTTTTCTTCGACTGGAATGACTACTTGAAGTTCTACAAGCTTGGAACTTACTGGCCGTAACACCTTCCATCCAACTACTCTACGGGCTAAGAGCAGCTTTGGATCTCATCTTTGAGGAGGGGCTCGAAAATGTCATCGAAAGGCATCGTCGTTTAGGAAAGGCCACGAGG-TAAGTAAAAGCT---------AATAGCCGATTTTACAGTATAGGAAAGATATATAATTAGGCATTATTGTCCTCTTGTAACAATGAGCAGACTTGCTGTCGAAG

			>Tillandsia_polystachia
			????????????????????????????????????????????????????????????????????????????????????????????????????????????????????????????????????????????????????????????????????????????????????????????????????????????????????????????????????????????????????????????????????????????????????????????????????????????????????????????????????????????????????????????????????????????????

			>Tillandsia_polzii
			????????????????????????????????????????????????????????????????????????????????????????????????????????????????????????????????????????????????????????????????????????????????????????????????????????????????????????????????????????????????????????????????????????????????????????????????????????????????????????????????????????????????????????????????????????????????

			>Tillandsia_ponderosa
			????????????????????????????????????????????????????????????????????????????????????????????????????????????????????????????????????????????????????????????????????????????????????????????????????????????????????????????????????????????????????????????????????????????????????????????????????????????????????????????????????????????????????????????????????????????????

			>Tillandsia_prodigiosa
			????????????????????????????????????????????????????????????????????????????????????????????????????????????????????????????????????????????????????????????????????????????????????????????????????????????????????????????????????????????????????????????????????????????????????????????????????????????????????????????????????????????????????????????????????????????????

			>Tillandsia_prolata
			????????????????????????????????????????????????????????????????????????????????????????????????????????????????????????????????????????????????????????????????????????????????????????????????????????????????????????????????????????????????????????????????????????????????????????????????????????????????????????????????????????????????????????????????????????????????

			>Tillandsia_propagulifera
			TCGCAGAAAGCACTTTCCCTGCCTACTGGAATGGGGATTGTTTGCGCGAGCCCGAAAGCTCTGGAAGCTTCCAAAAATGCCAAGTCGGTCAGAGTTTTCTTCGACTGGAATGACTACTTGAAGTTCTACAAGCTTGGAACTTACTGGCCGTAACGCCTTCCATCCAACTACTCTACGGGCTAAGAGCAGCTTTGGATCTTATCTTTGAGGAGGGGCTCGAAAATGTCATCGAAAGGCATCGTCGTTTAGGAAAGGCCACGAGG-TAAGTA----CT---------AATAGCCGATTTTACAGTATAGGAAAGATATATAATTAGGCATTATTGTCCTCTTGTAACAATGAGCAGACTCGCTGTCGAAG

			>Tillandsia_pseudocardenasii
			????????????????????????????????????????????????????????????????????????????????????????????????????????????????????????????????????????????????????????????????????????????????????????????????????????????????????????????????????????????????????????????????????????????????????????????????????????????????????????????????????????????????????????????????????????????????

			>Tillandsia_pseudomacbrideana
			????????????????????????????????????????????????????????????????????????????????????????????????????????????????????????????????????????????????????????????????????????????????????????????????????????????????????????????????????????????????????????????????????????????????????????????????????????????????????????????????????????????????????????????????????????????????

			>Tillandsia_pseudomicans
			TCGCAGAAAGCACTTTCCCTGCCTACTGGAATGGGCATTGTTTGCGCGAGCCCGAAAGCTCTAGAAMCTTCCAAAAATGCCAAGTCGGTCAGAGTTTTCTTCGACTGGAATGACTACTTGAAGTTCTACAAGCTTGGAACTTACTGGCCGTAACGCCTTCCATCCAACTACTTTACGGGCTAAGAGCAGCTTTGGATCTCATCTTTGAGGAGGGGCTCGAAAATGTCATCGATAGGCATCGTCGTTTAGGAAAGGCCACGAGG-TAAGTA-AAGCT--------AAATAGCCGATTTTACAGTATAGGAAAGATATATAATTAGGCATTATTGT-----------AATGAGCAGACTTGCTGTCGAAG

			>Tillandsia_pueblensis
			????????????????????????????????????????????????????????????????????????????????????????????????????????????????????????????????????????????????????????????????????????????????????????????????????????????????????????????????????????????????????????????????????????????????????????????????????????????????????????????????????????????????????????????????????????????????

			>Tillandsia_punctulata
			????????????????????????????????????????????????????????????????????????????????????????????????????????????????????????????????????????????????????????????????????????????????????????????????????????????????????????????????????????????????????????????????????????????????????????????????????????????????????????????????????????????????????????????????????????????????

			>Tillandsia_purpurea
			TCGCAGAAAGCACTTTCCCTGCCTACTGGAATGGGCATTGTTTGCGCGAGCCCGAAAGCTCTAGAAGCTTCCAAAAATGCCAAGTCGGTCAGAGTTTTCTTCGACTGGAATGACTACTTGAAGTTCTACAAGCTCGGAACTTACTGGCCGTAACGCCTTCCATCCAACTACTTTACGGGCTAAGAGCAGCTTTGGATCTCATCTTTGAGGAGGGGCTCGAAAATGTCATCGAAAGGCATCATCGTTTAGGAAAGGCCACGAGG-TAAGTA-ACGCT--------AAATAGCCGATTTTACAGTATAGGAAAGATATATAATTAGGCATTATTGT-----------AATGAGCAGACTTGCTGTCGAAG

			>Tillandsia_quaquaflorifera
			????????????????????????????????????????????????????????????????????????????????????????????????????????????????????????????????????????????????????????????????????????????????????????????????????????????????????????????????????????????????????????????????????????????????????????????????????????????????????????????????????????????????????????????????????????????????

			>Tillandsia_rauhii
			TCGCAGAAAGCACTTTCCCTGCCTACTGGAATGGGCATTGTTTGCGCGAGCCCGAAAGCTCTGGAAGCTTCCAAAAATGCCAAGTCGGTCAGAGTTTTCTTCGACTGGAATGACTACTTGAAGTTCTACAAGCTTGGAACTTACTGGCCGTAACGCCTTCCATCCAACTACTCTACGGGCTAAGAGCAGCTTTGGATCTCATCTTTGAGGAGGGGCTCGAAAATGTCATCGAAAGGCATCGTCGTTTAGGAAAGGCCACGAGG-TAAGTAAAAGC----------------CGATTTTACAGTATAGGAAAGATATATAATTAGGCATTATTGTCCTCTTGTAACAATGAGCAGACTTGCTGTCGAAG

			>Tillandsia_rectangula
			TCGCAGAAAGCACTTTCCCTGCCTACTGGAATGGGCATTGTTTGCGCGAGCCCGAAAGCTCTGGAAGCTTCCAAAAATGCCAAGTCGGTTAGAGTTTTCTTCGACTGGAATGACTACTTGAAGTTCTACAAGCTTGGAACTTACTGGCCGTAACGCCTTCCATCCAACTACTTTACGGGCTAAGAGCAGCTTTGGATCTCATCTTTGAGGAGGGCCTCGAAAATGTCATCGAAAGGCATCGTCGTTTAGGAAAGGCCACGAGG-TAAGTATAAGCT--------AAATAGCCGATTTTACAGTAT----AAGGTATGTAATTAACCATTATTGTCCTCTTGTAAC???????????????????????

			>Tillandsia_recurvata
			TCGCAGAAAGCACTTTCCCTGCCTACTGGAATGGGCATTGTTTGCGCGAGCCCAAAAGCTCTGGAAGCTTCCAAAAATGCCAAGTCGGTCCGAGTTTTCTTCGACTGGAATGACTACTTGAAGTTCTACAAGCTTGGAACTTACTGGCCGTAACGCCTTCCATCCAACTACTTTACGGGCTAAGAGCAGCTTTGGATCTCATCTTTGAGGAGGGCCTCGAAAATGTCATCGAAAGGCATCGTCGTTTAGGAAAGGCCACGAGG-TAAGTATAAGCT--------AAATAGTCAATTTTACAGTATAGGAAAGATATGTAATTAGCCATTATTGTCCTCTTGTAACAATGAGCA???????????????

			>Tillandsia_recurvifolia
			????????????????????????????????????????????????????????????????????????????????????????????????????????????????????????????????????????????????????????????????????????????????????????????????????????????????????????????????????????????????????????????????????????????????????????????????????????????????????????????????????????????????????????????????????????????????

			>Tillandsia_recurvifolia_var._subsecundifolia
			????????????????????????????????????????????????????????????????????????????????????????????????????????????????????????????????????????????????????????????????????????????????????????????????????????????????????????????????????????????????????????????????????????????????????????????????????????????????????????????????????????????????????????????????????????????????

			>Tillandsia_remota
			TCGCAGAAAGCACTTTCCCTGCCTACTGGAATGGGCATTGTTTGCGCGAGCCCGAAAGCGCTGGAAGCTTCCAAAAATGCCAAGTCGGTCAGAGTTTTCTTCGACTGGAATGACTACTTGAAGTTCTACAAGCTTGGAACTTACTGGCCGTAACGCCTTCCATCCAACTACTCTACGGGCTAAGAGCAGCTTTGGATCTCATCTTCGAGGAGGGGCTCGAAAATGTCATCGAAAGGCATCGTCGTTTAGGAAAGGCCACGAGG-TAAGTAAAAGCT--------AAATAGCCGATTTTACAGCATAGGAAAGATATATAATTAGGCATTATTGTCCTCTTGTAACAATGAGCA???????????????

			>Tillandsia_rosarioae
			????????????????????????????????????????????????????????????????????????????????????????????????????????????????????????????????????????????????????????????????????????????????????????????????????????????????????????????????????????????????????????????????????????????????????????????????????????????????????????????????????????????????????????????????????????????????

			>Tillandsia_roseoscapa
			????????????????????????????????????????????????????????????????????????????????????????????????????????????????????????????????????????????????????????????????????????????????????????????????????????????????????????????????????????????????????????????????????????????????????????????????????????????????????????????????????????????????????????????????????????????????

			>Tillandsia_rupicola
			????????????????????????????????????????????????????????????????????????????????????????????????????????????????????????????????????????????????????????????????????????????????????????????????????????????????????????????????????????????????????????????????????????????????????????????????????????????????????????????????????????????????????????????????????????????????

			>Tillandsia_sceptriformis
			TCGCAGAAAGCACTTTCCCTGCCTACTGGAATGGGCATTGTTTGCGCGAGCCCGAAAGCTCTAGAAGCTTCCAAAAATGCCAAGTCGGTCAGAGTTTTCTTCGACTGGAATGACTACTTGAAGTTCTACAAGCTTGGAACTTACTGGCCGTAACGCCTTCCATCCAACTACTTTACGGGCTAAGAGCAGCTTTGGATCTCATCTTTGAGGAGGGGCTCGAAAATGTCATCGAAAGGCATCATCGTTTAGGAAAGGCCACGAGG-TAAGTA-AAGCT--------AAATAGCCGATTTTACAG-------AAGATATATAATTAGGCATTATTGTACTCTTGTAACAATGAGCAGACTTGCTGTCGAAG

			>Tillandsia_schiedeana
			TCGCAGAAAGCACTTTCCCTGCCTACTGGAATGGGCATTGTTTGTGCGAGCCCGAAAGCTCTGGAAGCTTCCAAAAATGCCAAGTCGGTCAGAGTTTTCTTCGACTGGAATGACTACTTGAAGTTCTACAAGCTTGGAACTTACTGGCCGTAACGCCTTCCATCCAACTACTCTACGGGCTAAGAGCAGCTTTGGATCTCATCTTTGAGGAGGGGCTCGAAAATGTCATCGAAAGGCATCGTCGTTTAGGAAAGGCCACGAGG-TAAGTAAAAGCT--------AAATAGCCGATTTTACAGCATAGGAAAGATATATAATTAGGCATTATTGTCCTGTTGTAAC???????????????????????

			>Tillandsia_schimperiana
			????????????????????????????????????????????????????????????????????????????????????????????????????????????????????????????????????????????????????????????????????????????????????????????????????????????????????????????????????????????????????????????????????????????????????????????????????????????????????????????????????????????????????????????????????????????????

			>Tillandsia_secunda
			????????????????????????????????????????????????????????????????????????????????????????????????????????????????????????????????????????????????????????????????????????????????????????????????????????????????????????????????????????????????????????????????????????????????????????????????????????????????????????????????????????????????????????????????????????????????

			>Tillandsia_seleriana
			TCGCAGAAAGCACTTTCCCTGCCTACTGGAATGGGCATTGTTTGCGCGAGCCCGAAAGCGCTGGAAGCTTCCAAAAATGCCAAGTCGGTCAGAGTTTTCTTCGACTGGAATGACTACTTGAAGTTCTACAAGCTTGGAACTTACTGGCCGTAACGCCTTCCATCCAACTACTCTACGGGCTAAGAGCAGCTTTGGATCTCATCTTTGAGGAGGGGCTCGAAAATGTCATCGAAAGGCATCGTCGTTTAGGAAAGGCCACTAGG-TAAGTAAAAGCT--------AAATAGCCGATTTTACAGTAAAGGAAAGATATATAATTAGGCATTATTGTCCTCTTGTAAC???????????????????????

			>Tillandsia_sessemocinoi
			????????????????????????????????????????????????????????????????????????????????????????????????????????????????????????????????????????????????????????????????????????????????????????????????????????????????????????????????????????????????????????????????????????????????????????????????????????????????????????????????????????????????????????????????????????????????

			>Tillandsia_setacea
			????????????????????????????????????????????????????????????????????????????????????????????????????????????????????????????????????????????????????????????????????????????????????????????????????????????????????????????????????????????????????????????????????????????????????????????????????????????????????????????????????????????????????????????????????????????????

			>Tillandsia_sierrajuarezensis
			????????????????????????????????????????????????????????????????????????????????????????????????????????????????????????????????????????????????????????????????????????????????????????????????????????????????????????????????????????????????????????????????????????????????????????????????????????????????????????????????????????????????????????????????????????????????

			>Tillandsia_socialis
			????????????????????????????????????????????????????????????????????????????????????????????????????????????????????????????????????????????????????????????????????????????????????????????????????????????????????????????????????????????????????????????????????????????????????????????????????????????????????????????????????????????????????????????????????????????????

			>Tillandsia_somnians
			TCGCAGAAAGCACTTTCCCTGCCTACTGGAATGGGCATTGTTTGCGCGAGCCCGAAAGCTCTAGAAGCTTCCAAAAATGCCAAGTCGGTCAGAGTTTTCTTCGACTGGAATGACTACTTGAAGTTCTACAAGCTTGGAACTTACTGGCCGTAACGCCTTCCATCCAACTACTTTACGGGCTAAGAGCAGCTTTGGATCTCATCTTTGAGGAGGGGCTCGAAAATGTCATCGAAAGGCATCGTCGTTTAGGAAAGGCCACGAGG-TAAGTA-AAGCT--------AA----------TTACAGTATAGGAAAGATATATAATTAGGCATTATTKTMCTGTTGTAACAATGAGCAGACTTGCTGTCGAAG

			>Tillandsia_spathacea
			????????????????????????????????????????????????????????????????????????????????????????????????????????????????????????????????????????????????????????????????????????????????????????????????????????????????????????????????????????????????????????????????????????????????????????????????????????????????????????????????????????????????????????????????????????????????

			>Tillandsia_sphaerocephala
			????????????????????????????????????????????????????????????????????????????????????????????????????????????????????????????????????????????????????????????????????????????????????????????????????????????????????????????????????????????????????????????????????????????????????????????????????????????????????????????????????????????????????????????????????????????????

			>Tillandsia_spiraliflora
			TCGCAGAAAGCACTTTCCCTGCCTACTGGAATGGGGATTGTTTGCGCGAGCCCGAAAGCTCTGGAAGCTTCCAAGAATGCCAAGTCGGTCAGAGTTTTCTTCGACTGGAATGACTACTTGAAGTTCTACAAGCTTGGAACTTACTGGCCGTAACGCCTTCCATCCAACTACTCTACGGGCTAAGAGCAGCTTTGGATCTCATCTTTGAGGAGGGGCTCGAAAATGTCATCGAAAGGCATCGTCGTTTAGGAAAGGCCACGAGG-TAAGTAAAAGCT---------AATAGCCGATTTTACAGTATAGGAAAGATATATAATTAGGCATTATTGTCCTCTTGTAACAATGAGCAGACTCGCTGTCGAAG

			>Tillandsia_sprengeliana
			????????????????????????????????????????????????????????????????????????????????????????????????????????????????????????????????????????????????????????????????????????????????????????????????????????????????????????????????????????????????????????????????????????????????????????????????????????????????????????????????????????????????????????????????????????????????

			>Tillandsia_stenoura
			????????????????????????????????????????????????????????????????????????????????????????????????????????????????????????????????????????????????????????????????????????????????????????????????????????????????????????????????????????????????????????????????????????????????????????????????????????????????????????????????????????????????????????????????????????????????

			>Tillandsia_streptocarpa
			TCGCAGAAAGCACTTTCCCTGCCTACTGGAATGGGCATTGTTTGCGCGAGCCCGAAAGCTCTGGAAGCTTCCAAAAATGCCAAGTCGGTCAGAGTTTTCTTCGACTGGAATGACTACTTGAAGTTCTACAAGCTTGGAACTTACTGGCCGTAACGCCTTCCATCCAACTACTTTACGGGCTAAGAGCAGCTTTGGATCTCATCTTTGAGGAGGGCCTCGAAAATGTCATCGAAAGGCATCGTCGTTTAGGAAAGGCCACGAGG-TAAGTATAAGCT--------AAATAGCCGATTTTACAG--TAGGAAAGAGATGTAATTAGCCATTATTGTCCTCTTGTAACAATGAGCAGACTTGCTGTCGAAG

			>Tillandsia_streptophylla
			TCGCAGAAAGCACTTTCC?TGCCTACTGGAATGGGCATTGTTTGCGCGAGCCCGAAAGCGCTGGAAGCTTCCAAAAATGCCAAGTCGGTCAGAGTTTTCTTCGACTGGAATGACTACTTGAAGTTCTACAAGCTTGGAACTTACTGGCCGTAACGCCTTCCATCCAACTACTCTACGGGCTAAGAGCAGCTTTGGATCTCATCTTCGAGGAGGGGCTCGAAAATGTCATCGAAAGGCATCGTCGTTTAGGAAAGGCCACGAGGTTAAGTAAAAGCT--------AAATAGCCGATTTTACAGCATAGGAAAGATATATAATTAGGCATTATTGTCCTCTTGTAACAA?????????????????????

			>Tillandsia_stricta
			????????????????????????????????????????????????????????????????????????????????????????????????????????????????????????????????????????????????????????????????????????????????????????????????????????????????????????????????????????????????????????????????????????????????????????????????????????????????????????????????????????????????????????????????????????????????

			>Tillandsia_stricta_var._albifolia
			????????????????????????????????????????????????????????????????????????????????????????????????????????????????????????????????????????????????????????????????????????????????????????????????????????????????????????????????????????????????????????????????????????????????????????????????????????????????????????????????????????????????????????????????????????????????

			>Tillandsia_subulifera
			????????????????????????????????????????????????????????????????????????????????????????????????????????????????????????????????????????????????????????????????????????????????????????????????????????????????????????????????????????????????????????????????????????????????????????????????????????????????????????????????????????????????????????????????????????????????

			>Tillandsia_suesilliae
			????????????????????????????????????????????????????????????????????????????????????????????????????????????????????????????????????????????????????????????????????????????????????????????????????????????????????????????????????????????????????????????????????????????????????????????????????????????????????????????????????????????????????????????????????????????????

			>Tillandsia_supermexicana
			????????????????????????????????????????????????????????????????????????????????????????????????????????????????????????????????????????????????????????????????????????????????????????????????????????????????????????????????????????????????????????????????????????????????????????????????????????????????????????????????????????????????????????????????????????????????

			>Tillandsia_tecolometl
			????????????????????????????????????????????????????????????????????????????????????????????????????????????????????????????????????????????????????????????????????????????????????????????????????????????????????????????????????????????????????????????????????????????????????????????????????????????????????????????????????????????????????????????????????????????????

			>Tillandsia_tectorum
			????????????????????????????????????????????????????????????????????????????????????????????????????????????????????????????????????????????????????????????????????????????????????????????????????????????????????????????????????????????????????????????????????????????????????????????????????????????????????????????????????????????????????????????????????????????????

			>Tillandsia_tehuacana
			????????????????????????????????????????????????????????????????????????????????????????????????????????????????????????????????????????????????????????????????????????????????????????????????????????????????????????????????????????????????????????????????????????????????????????????????????????????????????????????????????????????????????????????????????????????????

			>Tillandsia_tenuifolia
			????????????????????????????????????????????????????????????????????????????????????????????????????????????????????????????????????????????????????????????????????????????????????????????????????????????????????????????????????????????????????????????????????????????????????????????????????????????????????????????????????????????????????????????????????????????????

			>Tillandsia_tenuifolia_var._albifolia
			????????????????????????????????????????????????????????????????????????????????????????????????????????????????????????????????????????????????????????????????????????????????????????????????????????????????????????????????????????????????????????????????????????????????????????????????????????????????????????????????????????????????????????????????????????????????

			>Tillandsia_tequendamae
			????????????????????????????????????????????????????????????????????????????????????????????????????????????????????????????????????????????????????????????????????????????????????????????????????????????????????????????????????????????????????????????????????????????????????????????????????????????????????????????????????????????????????????????????????????????????

			>Tillandsia_teres
			TCGCAGAAAGCACTTTCCCTGCCTACTGGAATGGGCATTGTTTGCGCGAGCCCGAAAGCTCTGGAAGCTTCYAAAAATGCCAAGTCGGTCAGAGTTTTCTTCGACTGGAATGACTACTTGAAGTTCTACAAGCTTGGAACTTACTGGCCGTAACGCCTTCCATCCAACTACTCTACGGGCTAAGAGCAGCTTTGGATCTCATCTTTGAGGAGGGGCTCGAAAATGTCATCGAAAGGCATCGTCGTTTAGGAAAGGCCACGAGG-TAAGTAAAAGC----------------CGATTTTACAGTATAGGAAAGATATATAATTAGGCATTATTGTCCTCTTGTAACAATGAGCAGACTTGCTGTCGAAG

			>Tillandsia_toropiensis
			????????????????????????????????????????????????????????????????????????????????????????????????????????????????????????????????????????????????????????????????????????????????????????????????????????????????????????????????????????????????????????????????????????????????????????????????????????????????????????????????????????????????????????????????????????????????

			>Tillandsia_tortilis
			????????????????????????????????????????????????????????????????????????????????????????????????????????????????????????????????????????????????????????????????????????????????????????????????????????????????????????????????????????????????????????????????????????????????????????????????????????????????????????????????????????????????????????????????????????????????

			>Tillandsia_tricholepis
			????????????????????????????????????????????????????????????????????????????????????????????????????????????????????????????????????????????????????????????????????????????????????????????????????????????????????????????????????????????????????????????????????????????????????????????????????????????????????????????????????????????????????????????????????????????????

			>Tillandsia_tricolor
			TCGCAGAAAGCACTTTCCCTGCCTACTGGAATGGGCATTGTTTGCGCTAGCCCGAAAGCTCTGGAAGCTTCCAAAAATGCCAAGTCGGTCAGAGTTTTCTTCGACTGGAATGACTACTTGAAGTTCTACAAGCTTGGAACTTACTGGCCGTAACGCCTTCCATCCAACTACTCTACGGGCTAAGAGCAGCTTTGGATCTCATCTTTGAGGAGGGGCTCGAAAATGTCATCGAAAGGCATCGTCGTTTAGGAAAGGCCACGAGG-TAAGTAAAAGCT--------AAATAGCCGATTTTACAGTATAGGAAAGATATATAATTAGGCATTATTGTCCTCTTGTAACAATGAGCAGACTTGCTGTCGAAG

			>Tillandsia_turneri
			????????????????????????????????????????????????????????????????????????????????????????????????????????????????????????????????????????????????????????????????????????????????????????????????????????????????????????????????????????????????????????????????????????????????????????????????????????????????????????????????????????????????????????????????????????????????

			>Tillandsia_turneri_var._orientalis
			????????????????????????????????????????????????????????????????????????????????????????????????????????????????????????????????????????????????????????????????????????????????????????????????????????????????????????????????????????????????????????????????????????????????????????????????????????????????????????????????????????????????????????????????????????????????

			>Tillandsia_usneoides
			TCGCAGAAAGCACTTTCCCTGCCTACTGGAATGGGCATTGTTTGCGCGAGCCCGAAAGCTCTGGAAGCTTCCAAAAATGCCAAGTCTGTCAGAGTTTTCTTCGACTGGAATGACTACTTGAAGTTCTACAAGCTTGGAACTTACTGGCCGTAACGCCTTCCATCCAACTACTTTACGGGCTAAGAGCAGCTTTGGATCTCATCTTTGAGGAGGGCCTCGAAAATGTCATCGAAAGGCATCGTCGTTTAGGAAAGGCCACGAGG-TAAGTATAAGCT--------AAATAGCCGATTTTACAGTATAGGAAAGATATGTAATTAGTCATTATTGTCCTCTTGTAACAATGAGCAGACTTGCTGTCGAAG

			>Tillandsia_utriculata
			TCGCAGAAGGCACTTTCTCTGCCTACTGGAATGGGCATTGTTTGCGCGAGCCCGAAAGCTCTGGAAGCTTCCAAAAATGCCAAGTCGGTCAGAGTTTTCTTCGACTGGAATGACTACTTGAAGTTCTACAAGCTTGGAACTTACTGGCCGTAACGCCTTCCATCCAACTACTCTACGGGCTAAGAGCAGCTTTGGATCTCATCTTTGAGGAGGGGCTCGAAAATGTCATCGAAAGGCATCGTCGTTTAGGAAAGGCCACGAGG-TAAGTAAAAGCT--------------------TTACAGTATAGGAAAGATATATAATTAGGCATTATTGTCCTCTTGTAACAATGAGCAGACTCGCTGTCGAAG

			>Tillandsia_utriculata_subsp._pringlei
			????????????????????????????????????????????????????????????????????????????????????????????????????????????????????????????????????????????????????????????????????????????????????????????????????????????????????????????????????????????????????????????????????????????????????????????????????????????????????????????????????????????????????????????????????????????????

			>Tillandsia_variabilis
			????????????????????????????????????????????????????????????????????????????????????????????????????????????????????????????????????????????????????????????????????????????????????????????????????????????????????????????????????????????????????????????????????????????????????????????????????????????????????????????????????????????????????????????????????????????????

			>Tillandsia_venusta
			????????????????????????????????????????????????????????????????????????????????????????????????????????????????????????????????????????????????????????????????????????????????????????????????????????????????????????????????????????????????????????????????????????????????????????????????????????????????????????????????????????????????????????????????????????????????

			>Tillandsia_violacea
			????????????????????????????????????????????????????????????????????????????????????????????????????????????????????????????????????????????????????????????????????????????????????????????????????????????????????????????????????????????????????????????????????????????????????????????????????????????????????????????????????????????????????????????????????????????????

			>Tillandsia_werdermannii
			TCGCAGAAAGCACTTTCCCTGCCTACTGGAATGGGCATTGTTTGCGCTAGCCCGAAAGCTCTGCAAGCTTCCAAAAATGCCAAGTCGGTCAGAGTTTTCTTCGACTGGAATGACTACTTGAAGTTCTACAAGCTTGGAACTTACTGGCCGTAACGCCTTCCATCCAACTACTTTACGGGCTAAGAGCAGCTTTGGATCTCATCTTTGAGGAGGGCCTCGAAAATGTCATCGAAAGGCATCGTCGTTTAGGAAAGGCCACGAGG-TAAGTA-AAGCT--------AAATAGCCGATTTTACAGTATAGGAAAGATATGTAATTAGCCATTATTGT-----------AATGAGCA???????????????

			>Tillandsia_werneriana
			????????????????????????????????????????????????????????????????????????????????????????????????????????????????????????????????????????????????????????????????????????????????????????????????????????????????????????????????????????????????????????????????????????????????????????????????????????????????????????????????????????????????????????????????????????????????

			>Tillandsia_winkleri
			????????????????????????????????????????????????????????????????????????????????????????????????????????????????????????????????????????????????????????????????????????????????????????????????????????????????????????????????????????????????????????????????????????????????????????????????????????????????????????????????????????????????????????????????????????????????

			>Tillandsia_witeckii
			????????????????????????????????????????????????????????????????????????????????????????????????????????????????????????????????????????????????????????????????????????????????????????????????????????????????????????????????????????????????????????????????????????????????????????????????????????????????????????????????????????????????????????????????????????????????

			>Tillandsia_xerographica
			????????????????????????????????????????????????????????????????????????????????????????????????????????????????????????????????????????????????????????????????????????????????????????????????????????????????????????????????????????????????????????????????????????????????????????????????????????????????????????????????????????????????????????????????????????????????

			>Tillandsia_xiphioides
			TCGCAGAAAGCACTTTCCCTGCCTACTGGAATGGGCATTGTTTGCGCGAGCCCGAAAGCTCTGGAAGCTTCCAAAAATGCCAAGTCGGTTAGAGTTTTCTTCGACTGGAATGACTACTTGAAGTTCTACAAGCTTGGAACTTACTGGCCGTAACGCCTTCCATCCAACTACTTTACGGGCTAAGAGCAGCTTTGGATCTCATCTTTGAGGAGGGGCTCGAAAATGTCATCGAAAGGCATCGTCGTTTAGGAAAGGCCACGAGG-TAAGTATAAGCT--------AAATAGCCGATTTTACAGTATAGGAAAGATATGTAATTAGCCATTATTGT-----------AATGAGCAGACTTGCTGTCGAAG

			>Typha_latifolia
			????????????????????????????????????????????????????????????????????????????????????????????????????????????????????????????????????????????????????????????????????????????????????????????????????????????????????????????????????????????????????????????????????????????????????????????????????????????????????????????????????????????????????????????????????????????????

			>Ursulaea_macvaughii
			TCGCAGAAAGCTCTTTCCCTGCCTACTGGAATGGGCATTCTCTGCGCGAGCCCGAAAGCTCTAGAAGCTTCCAAAACAGCCAAGTCGGTCAGGGTTTTCTTCGACTGGAATGACTACTTGAAGTTCTACAAGCTCGGAACTTACTGGCCTTAACGCCTTCCATCCAACTACTTTACGGGCTGAGAGCAGCTTTGGATCTCATCTTTGAGGAGGGGCTCGACAATGTCATCGAAAGGCATCGTCGTTTAGGAAAGGCCACAAGGTTAAGTAAAAGCTATACATCT-AATAGCCGCTTTCGCCGCGTTGGATAGATATTTTACTTTGCGCTATTGTCCTCTTGGTACAATGAGCAGACTTGCTGTCGAAG

			>Ursulaea_tuitensis
			TCGCAGAAAGCTCTTTCCCTGCCTACTGGAATGGGCATTCTCTGCGCGAGCCCGAAAGCTCTAGAAGCTTCCAAAACAGCCAAGTCGGTCAGGGTTTTCTTCGACTGGAATGACTACTTGAAGTTCTACAAGCTCGGAACTTACTGGCCTTAACGCCTTCCATCCAACTACTTTACGGGCTGAGAGCAGCTTTGGATCTCATCTTTGAGGAGGGGCTCGACAATGTCATCGAAAGGCATCGTCGCTTAGGAAAGGCCACAAGGTTAAGTAAAAGCTATACATCT-AATAGCCGCTTTCGCCGCGTTGGATAGATATTTTACTTTGCGCTATTGTCCTCTTGTAACAATGAGCAGACTTGCTGTCGAAG

			>Vriesea_agostiniana
			????????????????????????????????????????????????????????????????????????????????????????????????????????????????????????????????????????????????????????????????????????????????????????????????????????????????????????????????????????????????????????????????????????????????????????????????????????????????????????????????????????????????????????????????????????????????

			>Vriesea_amethystina
			????????????????????????????????????????????????????????????????????????????????????????????????????????????????????????????????????????????????????????????????????????????????????????????????????????????????????????????????????????????????????????????????????????????????????????????????????????????????????????????????????????????????????????????????????????????????

			>Vriesea_andaraiensis
			????????????????????????????????????????????????????????????????????????????????????????????????????????????????????????????????????????????????????????????????????????????????????????????????????????????????????????????????????????????????????????????????????????????????????????????????????????????????????????????????????????????????????????????????????????????????

			>Vriesea_arachnoidea
			????????????????????????????????????????????????????????????????????????????????????????????????????????????????????????????????????????????????????????????????????????????????????????????????????????????????????????????????????????????????????????????????????????????????????????????????????????????????????????????????????????????????????????????????????????????????

			>Vriesea_atra
			????????????????????????????????????????????????????????????????????????????????????????????????????????????????????????????????????????????????????????????????????????????????????????????????????????????????????????????????????????????????????????????????????????????????????????????????????????????????????????????????????????????????????????????????????????????????

			>Vriesea_bahiana
			????????????????????????????????????????????????????????????????????????????????????????????????????????????????????????????????????????????????????????????????????????????????????????????????????????????????????????????????????????????????????????????????????????????????????????????????????????????????????????????????????????????????????????????????????????????????

			>Vriesea_billbergioides
			????????????????????????????????????????????????????????????????????????????????????????????????????????????????????????????????????????????????????????????????????????????????????????????????????????????????????????????????????????????????????????????????????????????????????????????????????????????????????????????????????????????????????????????????????????????????

			>Vriesea_bituminosa
			????????????????????????????????????????????????????????????????????????????????????????????????????????????????????????????????????????????????????????????????????????????????????????????????????????????????????????????????????????????????????????????????????????????????????????????????????????????????????????????????????????????????????????????????????????????????

			>Vriesea_bleheri
			????????????????????????????????????????????????????????????????????????????????????????????????????????????????????????????????????????????????????????????????????????????????????????????????????????????????????????????????????????????????????????????????????????????????????????????????????????????????????????????????????????????????????????????????????????????????

			>Vriesea_botafogensis
			????????????????????????????????????????????????????????????????????????????????????????????????????????????????????????????????????????????????????????????????????????????????????????????????????????????????????????????????????????????????????????????????????????????????????????????????????????????????????????????????????????????????????????????????????????????????

			>Vriesea_cacuminis
			????????????????????????????????????????????????????????????????????????????????????????????????????????????????????????????????????????????????????????????????????????????????????????????????????????????????????????????????????????????????????????????????????????????????????????????????????????????????????????????????????????????????????????????????????????????????

			>Vriesea_calimaniana
			????????????????????????????????????????????????????????????????????????????????????????????????????????????????????????????????????????????????????????????????????????????????????????????????????????????????????????????????????????????????????????????????????????????????????????????????????????????????????????????????????????????????????????????????????????????????

			>Vriesea_capixabae
			????????????????????????????????????????????????????????????????????????????????????????????????????????????????????????????????????????????????????????????????????????????????????????????????????????????????????????????????????????????????????????????????????????????????????????????????????????????????????????????????????????????????????????????????????????????????

			>Vriesea_carinata
			????????????????????????????????????????????????????????????????????????????????????????????????????????????????????????????????????????????????????????????????????????????????????????????????????????????????????????????????????????????????????????????????????????????????????????????????????????????????????????????????????????????????????????????????????????????????

			>Vriesea_carinata_var._flavominiata
			????????????????????????????????????????????????????????????????????????????????????????????????????????????????????????????????????????????????????????????????????????????????????????????????????????????????????????????????????????????????????????????????????????????????????????????????????????????????????????????????????????????????????????????????????????????????

			>Vriesea_carinata_var._mangaratibensis
			????????????????????????????????????????????????????????????????????????????????????????????????????????????????????????????????????????????????????????????????????????????????????????????????????????????????????????????????????????????????????????????????????????????????????????????????????????????????????????????????????????????????????????????????????????????????

			>Vriesea_colnagoi
			????????????????????????????????????????????????????????????????????????????????????????????????????????????????????????????????????????????????????????????????????????????????????????????????????????????????????????????????????????????????????????????????????????????????????????????????????????????????????????????????????????????????????????????????????????????????

			>Vriesea_correia-araujoi
			????????????????????????????????????????????????????????????????????????????????????????????????????????????????????????????????????????????????????????????????????????????????????????????????????????????????????????????????????????????????????????????????????????????????????????????????????????????????????????????????????????????????????????????????????????????????

			>Vriesea_crassa
			????????????????????????????????????????????????????????????????????????????????????????????????????????????????????????????????????????????????????????????????????????????????????????????????????????????????????????????????????????????????????????????????????????????????????????????????????????????????????????????????????????????????????????????????????????????????

			>Vriesea_dubia
			????????????????????????????????????????????????????????????????????????????????????????????????????????????????????????????????????????????????????????????????????????????????????????????????????????????????????????????????????????????????????????????????????????????????????????????????????????????????????????????????????????????????????????????????????????????????

			>Vriesea_duvaliana
			????????????????????????????????????????????????????????????????????????????????????????????????????????????????????????????????????????????????????????????????????????????????????????????????????????????????????????????????????????????????????????????????????????????????????????????????????????????????????????????????????????????????????????????????????????????????

			>Vriesea_elata
			????????????????????????????????????????????????????????????????????????????????????????????????????????????????????????????????????????????????????????????????????????????????????????????????????????????????????????????????????????????????????????????????????????????????????????????????????????????????????????????????????????????????????????????????????????????????

			>Vriesea_eltoniana
			????????????????????????????????????????????????????????????????????????????????????????????????????????????????????????????????????????????????????????????????????????????????????????????????????????????????????????????????????????????????????????????????????????????????????????????????????????????????????????????????????????????????????????????????????????????????

			>Vriesea_ensiformis
			????????????????????????????????????????????????????????????????????????????????????????????????????????????????????????????????????????????????????????????????????????????????????????????????????????????????????????????????????????????????????????????????????????????????????????????????????????????????????????????????????????????????????????????????????????????????

			>Vriesea_ensiformis_var._bicolor
			????????????????????????????????????????????????????????????????????????????????????????????????????????????????????????????????????????????????????????????????????????????????????????????????????????????????????????????????????????????????????????????????????????????????????????????????????????????????????????????????????????????????????????????????????????????????

			>Vriesea_erythrodactylon
			????????????????????????????????????????????????????????????????????????????????????????????????????????????????????????????????????????????????????????????????????????????????????????????????????????????????????????????????????????????????????????????????????????????????????????????????????????????????????????????????????????????????????????????????????????????????

			>Vriesea_fenestralis
			????????????????????????????????????????????????????????????????????????????????????????????????????????????????????????????????????????????????????????????????????????????????????????????????????????????????????????????????????????????????????????????????????????????????????????????????????????????????????????????????????????????????????????????????????????????????

			>Vriesea_flammea
			????????????????????????????????????????????????????????????????????????????????????????????????????????????????????????????????????????????????????????????????????????????????????????????????????????????????????????????????????????????????????????????????????????????????????????????????????????????????????????????????????????????????????????????????????????????????

			>Vriesea_flava
			????????????????????????????????????????????????????????????????????????????????????????????????????????????????????????????????????????????????????????????????????????????????????????????????????????????????????????????????????????????????????????????????????????????????????????????????????????????????????????????????????????????????????????????????????????????????

			>Vriesea_fluviatilis
			????????????????????????????????????????????????????????????????????????????????????????????????????????????????????????????????????????????????????????????????????????????????????????????????????????????????????????????????????????????????????????????????????????????????????????????????????????????????????????????????????????????????????????????????????????????????

			>Vriesea_fosteriana
			????????????????????????????????????????????????????????????????????????????????????????????????????????????????????????????????????????????????????????????????????????????????????????????????????????????????????????????????????????????????????????????????????????????????????????????????????????????????????????????????????????????????????????????????????????????????

			>Vriesea_friburgensis
			????????????????????????????????????????????????????????????????????????????????????????????????????????????????????????????????????????????????????????????????????????????????????????????????????????????????????????????????????????????????????????????????????????????????????????????????????????????????????????????????????????????????????????????????????????????????

			>Vriesea_gigantea
			????????????????????????????????????????????????????????????????????????????????????????????????????????????????????????????????????????????????????????????????????????????????????????????????????????????????????????????????????????????????????????????????????????????????????????????????????????????????????????????????????????????????????????????????????????????????

			>Vriesea_gracilior
			????????????????????????????????????????????????????????????????????????????????????????????????????????????????????????????????????????????????????????????????????????????????????????????????????????????????????????????????????????????????????????????????????????????????????????????????????????????????????????????????????????????????????????????????????????????????

			>Vriesea_gradata
			????????????????????????????????????????????????????????????????????????????????????????????????????????????????????????????????????????????????????????????????????????????????????????????????????????????????????????????????????????????????????????????????????????????????????????????????????????????????????????????????????????????????????????????????????????????????

			>Vriesea_grandiflora
			????????????????????????????????????????????????????????????????????????????????????????????????????????????????????????????????????????????????????????????????????????????????????????????????????????????????????????????????????????????????????????????????????????????????????????????????????????????????????????????????????????????????????????????????????????????????

			>Vriesea_guttata
			????????????????????????????????????????????????????????????????????????????????????????????????????????????????????????????????????????????????????????????????????????????????????????????????????????????????????????????????????????????????????????????????????????????????????????????????????????????????????????????????????????????????????????????????????????????????

			>Vriesea_heterostachys
			????????????????????????????????????????????????????????????????????????????????????????????????????????????????????????????????????????????????????????????????????????????????????????????????????????????????????????????????????????????????????????????????????????????????????????????????????????????????????????????????????????????????????????????????????????????????

			>Vriesea_hydrophora
			????????????????????????????????????????????????????????????????????????????????????????????????????????????????????????????????????????????????????????????????????????????????????????????????????????????????????????????????????????????????????????????????????????????????????????????????????????????????????????????????????????????????????????????????????????????????

			>Vriesea_incurvata
			????????????????????????????????????????????????????????????????????????????????????????????????????????????????????????????????????????????????????????????????????????????????????????????????????????????????????????????????????????????????????????????????????????????????????????????????????????????????????????????????????????????????????????????????????????????????

			>Vriesea_inflata
			????????????????????????????????????????????????????????????????????????????????????????????????????????????????????????????????????????????????????????????????????????????????????????????????????????????????????????????????????????????????????????????????????????????????????????????????????????????????????????????????????????????????????????????????????????????????

			>Vriesea_interrogatoria
			????????????????????????????????????????????????????????????????????????????????????????????????????????????????????????????????????????????????????????????????????????????????????????????????????????????????????????????????????????????????????????????????????????????????????????????????????????????????????????????????????????????????????????????????????????????????

			>Vriesea_jonghei
			????????????????????????????????????????????????????????????????????????????????????????????????????????????????????????????????????????????????????????????????????????????????????????????????????????????????????????????????????????????????????????????????????????????????????????????????????????????????????????????????????????????????????????????????????????????????

			>Vriesea_limae
			????????????????????????????????????????????????????????????????????????????????????????????????????????????????????????????????????????????????????????????????????????????????????????????????????????????????????????????????????????????????????????????????????????????????????????????????????????????????????????????????????????????????????????????????????????????????

			>Vriesea_longicaulis
			????????????????????????????????????????????????????????????????????????????????????????????????????????????????????????????????????????????????????????????????????????????????????????????????????????????????????????????????????????????????????????????????????????????????????????????????????????????????????????????????????????????????????????????????????????????????

			>Vriesea_longistaminea
			????????????????????????????????????????????????????????????????????????????????????????????????????????????????????????????????????????????????????????????????????????????????????????????????????????????????????????????????????????????????????????????????????????????????????????????????????????????????????????????????????????????????????????????????????????????????

			>Vriesea_lubbersii
			????????????????????????????????????????????????????????????????????????????????????????????????????????????????????????????????????????????????????????????????????????????????????????????????????????????????????????????????????????????????????????????????????????????????????????????????????????????????????????????????????????????????????????????????????????????????

			>Vriesea_maxoniana
			????????????????????????????????????????????????????????????????????????????????????????????????????????????????????????????????????????????????????????????????????????????????????????????????????????????????????????????????????????????????????????????????????????????????????????????????????????????????????????????????????????????????????????????????????????????????

			>Vriesea_minuta
			????????????????????????????????????????????????????????????????????????????????????????????????????????????????????????????????????????????????????????????????????????????????????????????????????????????????????????????????????????????????????????????????????????????????????????????????????????????????????????????????????????????????????????????????????????????????

			>Vriesea_modesta
			????????????????????????????????????????????????????????????????????????????????????????????????????????????????????????????????????????????????????????????????????????????????????????????????????????????????????????????????????????????????????????????????????????????????????????????????????????????????????????????????????????????????????????????????????????????????

			>Vriesea_neoglutinosa
			TCGCAGAAAGCACTTTCCCTGCCTACTGGAATGGGCATTGTTTGCGCGAGCCCGAAAGCTCTAGAAGCTTCCAAAAATGCCAAGTCGGTCAGAGTTTTCTTCGACTGGAATGACTACTTGAAGTTCTACAAGCTTGGAACTTACTGGCCGTAACGCCTTCCATCCAACTACTTTACGGGCTAAGAGCAGCTTTGGATCTCATCTTTGAGGAGGGGCTCGAAAATGTCATCGAAAGGCATCGCCGTTTAGGAAAGGCCACGAGG-TAAGTAAAAGCT--------AAATAGCCGATTTTAGCA--TAGGAAAGATATATAATTACTCATTATTGTCCTCTTGTAACAATGAGCAGACTTGCTGTCGAAG

			>Vriesea_noblickii
			????????????????????????????????????????????????????????????????????????????????????????????????????????????????????????????????????????????????????????????????????????????????????????????????????????????????????????????????????????????????????????????????????????????????????????????????????????????????????????????????????????????????????????????????????????????????

			>Vriesea_ochracea
			????????????????????????????????????????????????????????????????????????????????????????????????????????????????????????????????????????????????????????????????????????????????????????????????????????????????????????????????????????????????????????????????????????????????????????????????????????????????????????????????????????????????????????????????????????????????

			>Vriesea_oligantha
			????????????????????????????????????????????????????????????????????????????????????????????????????????????????????????????????????????????????????????????????????????????????????????????????????????????????????????????????????????????????????????????????????????????????????????????????????????????????????????????????????????????????????????????????????????????????

			>Vriesea_pabstii
			????????????????????????????????????????????????????????????????????????????????????????????????????????????????????????????????????????????????????????????????????????????????????????????????????????????????????????????????????????????????????????????????????????????????????????????????????????????????????????????????????????????????????????????????????????????????

			>Vriesea_paraibica
			????????????????????????????????????????????????????????????????????????????????????????????????????????????????????????????????????????????????????????????????????????????????????????????????????????????????????????????????????????????????????????????????????????????????????????????????????????????????????????????????????????????????????????????????????????????????

			>Vriesea_pauperrima
			????????????????????????????????????????????????????????????????????????????????????????????????????????????????????????????????????????????????????????????????????????????????????????????????????????????????????????????????????????????????????????????????????????????????????????????????????????????????????????????????????????????????????????????????????????????????

			>Vriesea_platynema
			????????????????????????????????????????????????????????????????????????????????????????????????????????????????????????????????????????????????????????????????????????????????????????????????????????????????????????????????????????????????????????????????????????????????????????????????????????????????????????????????????????????????????????????????????????????????

			>Vriesea_platynema_var._rosea
			????????????????????????????????????????????????????????????????????????????????????????????????????????????????????????????????????????????????????????????????????????????????????????????????????????????????????????????????????????????????????????????????????????????????????????????????????????????????????????????????????????????????????????????????????????????????

			>Vriesea_platynema_var._variegata
			????????????????????????????????????????????????????????????????????????????????????????????????????????????????????????????????????????????????????????????????????????????????????????????????????????????????????????????????????????????????????????????????????????????????????????????????????????????????????????????????????????????????????????????????????????????????

			>Vriesea_procera
			????????????????????????????????????????????????????????????????????????????????????????????????????????????????????????????????????????????????????????????????????????????????????????????????????????????????????????????????????????????????????????????????????????????????????????????????????????????????????????????????????????????????????????????????????????????????

			>Vriesea_procera_var._tenuis
			????????????????????????????????????????????????????????????????????????????????????????????????????????????????????????????????????????????????????????????????????????????????????????????????????????????????????????????????????????????????????????????????????????????????????????????????????????????????????????????????????????????????????????????????????????????????

			>Vriesea_psittacina
			????????????????????????????????????????????????????????????????????????????????????????????????????????????????????????????????????????????????????????????????????????????????????????????????????????????????????????????????????????????????????????????????????????????????????????????????????????????????????????????????????????????????????????????????????????????????

			>Vriesea_recurvata
			????????????????????????????????????????????????????????????????????????????????????????????????????????????????????????????????????????????????????????????????????????????????????????????????????????????????????????????????????????????????????????????????????????????????????????????????????????????????????????????????????????????????????????????????????????????????

			>Vriesea_repandostachys
			????????????????????????????????????????????????????????????????????????????????????????????????????????????????????????????????????????????????????????????????????????????????????????????????????????????????????????????????????????????????????????????????????????????????????????????????????????????????????????????????????????????????????????????????????????????????

			>Vriesea_rhodostachys
			????????????????????????????????????????????????????????????????????????????????????????????????????????????????????????????????????????????????????????????????????????????????????????????????????????????????????????????????????????????????????????????????????????????????????????????????????????????????????????????????????????????????????????????????????????????????

			>Vriesea_rodigasiana
			????????????????????????????????????????????????????????????????????????????????????????????????????????????????????????????????????????????????????????????????????????????????????????????????????????????????????????????????????????????????????????????????????????????????????????????????????????????????????????????????????????????????????????????????????????????????

			>Vriesea_roethii
			????????????????????????????????????????????????????????????????????????????????????????????????????????????????????????????????????????????????????????????????????????????????????????????????????????????????????????????????????????????????????????????????????????????????????????????????????????????????????????????????????????????????????????????????????????????????

			>Vriesea_rubra
			????????????????????????????????????????????????????????????????????????????????????????????????????????????????????????????????????????????????????????????????????????????????????????????????????????????????????????????????????????????????????????????????????????????????????????????????????????????????????????????????????????????????????????????????????????????????

			>Vriesea_rubyae
			????????????????????????????????????????????????????????????????????????????????????????????????????????????????????????????????????????????????????????????????????????????????????????????????????????????????????????????????????????????????????????????????????????????????????????????????????????????????????????????????????????????????????????????????????????????????

			>Vriesea_ruschii_subsp._leonii
			????????????????????????????????????????????????????????????????????????????????????????????????????????????????????????????????????????????????????????????????????????????????????????????????????????????????????????????????????????????????????????????????????????????????????????????????????????????????????????????????????????????????????????????????????????????????

			>Vriesea_sanfranciscana
			????????????????????????????????????????????????????????????????????????????????????????????????????????????????????????????????????????????????????????????????????????????????????????????????????????????????????????????????????????????????????????????????????????????????????????????????????????????????????????????????????????????????????????????????????????????????

			>Vriesea_saundersii
			????????????????????????????????????????????????????????????????????????????????????????????????????????????????????????????????????????????????????????????????????????????????????????????????????????????????????????????????????????????????????????????????????????????????????????????????????????????????????????????????????????????????????????????????????????????????

			>Vriesea_sazimae
			????????????????????????????????????????????????????????????????????????????????????????????????????????????????????????????????????????????????????????????????????????????????????????????????????????????????????????????????????????????????????????????????????????????????????????????????????????????????????????????????????????????????????????????????????????????????

			>Vriesea_scalaris
			????????????????????????????????????????????????????????????????????????????????????????????????????????????????????????????????????????????????????????????????????????????????????????????????????????????????????????????????????????????????????????????????????????????????????????????????????????????????????????????????????????????????????????????????????????????????

			>Vriesea_scalaris_var._viridis
			????????????????????????????????????????????????????????????????????????????????????????????????????????????????????????????????????????????????????????????????????????????????????????????????????????????????????????????????????????????????????????????????????????????????????????????????????????????????????????????????????????????????????????????????????????????????

			>Vriesea_segadas-viannae
			????????????????????????????????????????????????????????????????????????????????????????????????????????????????????????????????????????????????????????????????????????????????????????????????????????????????????????????????????????????????????????????????????????????????????????????????????????????????????????????????????????????????????????????????????????????????

			>Vriesea_seideliana
			????????????????????????????????????????????????????????????????????????????????????????????????????????????????????????????????????????????????????????????????????????????????????????????????????????????????????????????????????????????????????????????????????????????????????????????????????????????????????????????????????????????????????????????????????????????????

			>Vriesea_simplex
			????????????????????????????????????????????????????????????????????????????????????????????????????????????????????????????????????????????????????????????????????????????????????????????????????????????????????????????????????????????????????????????????????????????????????????????????????????????????????????????????????????????????????????????????????????????????

			>Vriesea_sincorana
			????????????????????????????????????????????????????????????????????????????????????????????????????????????????????????????????????????????????????????????????????????????????????????????????????????????????????????????????????????????????????????????????????????????????????????????????????????????????????????????????????????????????????????????????????????????????

			>Vriesea_sparsiflora
			????????????????????????????????????????????????????????????????????????????????????????????????????????????????????????????????????????????????????????????????????????????????????????????????????????????????????????????????????????????????????????????????????????????????????????????????????????????????????????????????????????????????????????????????????????????????

			>Vriesea_sucrei
			????????????????????????????????????????????????????????????????????????????????????????????????????????????????????????????????????????????????????????????????????????????????????????????????????????????????????????????????????????????????????????????????????????????????????????????????????????????????????????????????????????????????????????????????????????????????

			>Vriesea_taritubensis
			????????????????????????????????????????????????????????????????????????????????????????????????????????????????????????????????????????????????????????????????????????????????????????????????????????????????????????????????????????????????????????????????????????????????????????????????????????????????????????????????????????????????????????????????????????????????

			>Vriesea_taritubensis_var._brevisepala
			????????????????????????????????????????????????????????????????????????????????????????????????????????????????????????????????????????????????????????????????????????????????????????????????????????????????????????????????????????????????????????????????????????????????????????????????????????????????????????????????????????????????????????????????????????????????

			>Vriesea_taritubensis_var._patens
			????????????????????????????????????????????????????????????????????????????????????????????????????????????????????????????????????????????????????????????????????????????????????????????????????????????????????????????????????????????????????????????????????????????????????????????????????????????????????????????????????????????????????????????????????????????????

			>Vriesea_teresopolitana
			????????????????????????????????????????????????????????????????????????????????????????????????????????????????????????????????????????????????????????????????????????????????????????????????????????????????????????????????????????????????????????????????????????????????????????????????????????????????????????????????????????????????????????????????????????????????

			>Vriesea_tijucana
			????????????????????????????????????????????????????????????????????????????????????????????????????????????????????????????????????????????????????????????????????????????????????????????????????????????????????????????????????????????????????????????????????????????????????????????????????????????????????????????????????????????????????????????????????????????????

			>Vriesea_unilateralis
			????????????????????????????????????????????????????????????????????????????????????????????????????????????????????????????????????????????????????????????????????????????????????????????????????????????????????????????????????????????????????????????????????????????????????????????????????????????????????????????????????????????????????????????????????????????????

			>Vriesea_vagans
			????????????????????????????????????????????????????????????????????????????????????????????????????????????????????????????????????????????????????????????????????????????????????????????????????????????????????????????????????????????????????????????????????????????????????????????????????????????????????????????????????????????????????????????????????????????????

			>Vriesea_zamorensis
			????????????????????????????????????????????????????????????????????????????????????????????????????????????????????????????????????????????????????????????????????????????????????????????????????????????????????????????????????????????????????????????????????????????????????????????????????????????????????????????????????????????????????????????????????????????????

			>Vriesea_zonata
			????????????????????????????????????????????????????????????????????????????????????????????????????????????????????????????????????????????????????????????????????????????????????????????????????????????????????????????????????????????????????????????????????????????????????????????????????????????????????????????????????????????????????????????????????????????????

			>Wallisia_anceps
			TCGCAGAAAGCACTTTCCCTGCCTACTGGAATGGGCATTGTTTGCGCGAGCCCGAAAGCTCTAGAAGCTTCCAAAAATGCCAAGTCGGTCAGAGTTTTCTTCGACTGGAATGACTACTTGAAGTTCTACAAGCTTGGAACTTACTGGCCGTAACGCCTTCCATCCAACTACTTTACGGGCTAAGAGCAGCTTTGGATCTCATCTTTGAGGAGGGGCTCGAAAATGTCATCGAAAGGCATCGTCGTTTAGGAAAGGCCACGAGG-TAAGTA-AAGCT--------AAATAGCCGATTTTACAGTATAGAAAAGATATATAATTACGCATTATTGTCCTGTTGTAACAATGAGCAGACTTGCTGTCGAAG

			>Wallisia_cyanea
			????????????????????GCCTACTGGAATGGGCATTGTTTGCGCGAGCCCGAAAGCTCTAGAAGCTTCCAAAAATGCCAAGTCGGTCAGAGTTTTCTTCGACTGGAATGACTACTTGAAGTTCTACAAGCTTGGAACTTACTGGCCGTAACCCCTTCCATCCAACTACTTTACGGGCTAAGAGCAGCTTTGGATCTCATCTTTGAGGAGGGGCTCGAAAATGTCATCGAAAGGCATCATCGTTTAKGAAAGGCCACGAGG-TAAGTA-AAGCT--------AAATAGCCGATTTTACAGTATAGGAAAGATATATAATTAGGCATTATTGTCCTCTTGTAACAATGAACAGACT???????????

			>Wallisia_lindeniana
			????????????????????????????????????????????????????????????????????????????????????????????????????????????????????????????????????????????????????????????????????????????????????????????????????????????????????????????????????????????????????????????????????????????????????????????????????????????????????????????????????????????????????????????????????????????????

			>Waltillia_hatschbachii
			????????????????????????????????????????????????????????????????????????????????????????????????????????????????????????????????????????????????????????????????????????????????????????????????????????????????????????????????????????????????????????????????????????????????????????????????????????????????????????????????????????????????????????????????????????????????

			>Werauhia_barii
			????????????????????????????????????????????????????????????????????????????????????????????????????????????????????????????????????????????????????????????????????????????????????????????????????????????????????????????????????????????????????????????????????????????????????????????????????????????????????????????????????????????????????????????????????????????????

			>Werauhia_bicolor
			????????????????????????????????????????????????????????????????????????????????????????????????????????????????????????????????????????????????????????????????????????????????????????????????????????????????????????????????????????????????????????????????????????????????????????????????????????????????????????????????????????????????????????????????????????????????

			>Werauhia_burgeri
			????????????????????????????????????????????????????????????????????????????????????????????????????????????????????????????????????????????????????????????????????????????????????????????????????????????????????????????????????????????????????????????????????????????????????????????????????????????????????????????????????????????????????????????????????????????????

			>Werauhia_gladioliflora
			????????????????????????????????????????????????????????????????????????????????????????????????????????????????????????????????????????????????????????????????????????????????????????????????????????????????????????????????????????????????????????????????????????????????????????????????????????????????????????????????????????????????????????????????????????????????

			>Werauhia_insignis
			????????????????????????????????????????????????????????????????????????????????????????????????????????????????????????????????????????????????????????????????????????????????????????????????????????????????????????????????????????????????????????????????????????????????????????????????????????????????????????????????????????????????????????????????????????????????

			>Werauhia_ororiensis
			????????????????????????????????????????????????????????????????????????????????????????????????????????????????????????????????????????????????????????????????????????????????????????????????????????????????????????????????????????????????????????????????????????????????????????????????????????????????????????????????????????????????????????????????????????????????

			>Werauhia_patzeltii
			????????????????????????????????????????????????????????????????????????????????????????????????????????????????????????????????????????????????????????????????????????????????????????????????????????????????????????????????????????????????????????????????????????????????????????????????????????????????????????????????????????????????????????????????????????????????

			>Werauhia_pectinata
			TCGCAGAAAGCACTTTCCCTGCCTACTGGAATGGGCATTGTTTGCGCGAGCCCGAAAGCTCTAGAAGCTTCCAAAAATGCCAAGTCGGTCAGAGTTTTCTTCGACTGGAATGACTACTTGAAGTTCTACAAGCTTGGAACTTACTGGCCGTAACGCCTTCCATCCAACTACTTTACGGGCTAAGAGCAGCTTTGGATCTCATCTTTGAGGAGGGGCTCGAAAATGTCATCGAAAGGCATCGTCGTTTAGGAAAGGCCACGAGG-TAAGTAAAAGCT--------AAATAGCCGATTTTAGCA--TAGGAAAGATATATAATTASTCATTATTATCCTCTTGTAACAATGAGCAGACTTGCTGTCGAAG

			>Werauhia_pedicellata
			????????????????????????????????????????????????????????????????????????????????????????????????????????????????????????????????????????????????????????????????????????????????????????????????????????????????????????????????????????????????????????????????????????????????????????????????????????????????????????????????????????????????????????????????????????????????

			>Werauhia_ringens
			????????????????????????????????????????????????????????????????????????????????????????????????????????????????????????????????????????????????????????????????????????????????????????????????????????????????????????????????????????????????????????????????????????????????????????????????????????????????????????????????????????????????????????????????????????????????

			>Werauhia_sanguinolenta
			????????????????????????????????????????????????????????????????????????????????????????????????????????????????????????????????????????????????????????????????????????????????????????????????????????????????????????????????????????????????????????????????????????????????????????????????????????????????????????????????????????????????????????????????????????????????

			>Werauhia_tarmaensis
			????????????????????????????????????????????????????????????????????????????????????????????????????????????????????????????????????????????????????????????????????????????????????????????????????????????????????????????????????????????????????????????????????????????????????????????????????????????????????????????????????????????????????????????????????????????????

			>Werauhia_viridiflora
			TCGCAGAAAGCACTTTCCCTGCCTACTGGAATGGGCATTGTTTGCGCTAGCCCGAAAGCTCTAGAAGCTTCTAAAAATGCCAAGTCGGTCAGAGTTTTCTTCGACTGGAATGACTACTTGAAGTTCTACAAGCTTGGAACTTACTGGCCGTAACGCCTTCCATCCAACTACTTTACGGGCTAAGAGCAGCTTTGGATCTCATCTTTGAGGAGGGGCTCGAAAATGTCATCGAAAGGCATCGTCGTCTAGGAAAGGCCACGAGG-TAAGTA-AAGCT--------AAATAGCCGATTTTAGCATATAGGAAAGATATATAATTAGTCATTATTATCCTCTTGTAACAATGAGCAGACTTGCTGTCGAAG

			>Werauhia_vittata
			????????????????????????????????????????????????????????????????????????????????????????????????????????????????????????????????????????????????????????????????????????????????????????????????????????????????????????????????????????????????????????????????????????????????????????????????????????????????????????????????????????????????????????????????????????????????

			>Wittmackia_abbreviata
			????????????????????????????????????????????????????????????????????????????????????????????????????????????????????????????????????????????????????????????????????????????????????????????????????????????????????????????????????????????????????????????????????????????????????????????????????????????????????????????????????????????????????????????????????????????????

			>Wittmackia_altocaririensis
			????????????????????????????????????????????????????????????????????????????????????????????????????????????????????????????????????????????????????????????????????????????????????????????????????????????????????????????????????????????????????????????????????????????????????????????????????????????????????????????????????????????????????????????????????????????????

			>Wittmackia_amorimii
			TCGCAGAAAGCTCTTTCCCTGCCTACTGGAATGGGGATTCTCTGCGCGAGCCCGAAAGCTCTAGAAGCTTCCAAAACAGCCAAGTCGGTCAGGGTTTTCTTCGACTGGAATGACTACTTGAAGTTCTACAAGCTCGGAACTTACTGGCCTTAACGCCTTCCATCCAACTACTTTACGGGCTGAGAGCAGCTCTGGATCTCATCTTTGAGGAGGGGCTCGACAATGTCATCGAAAGGCATCGTCGCTTAGGAAAGGCCACAAGGTTAAGTAAAAGCTATACATCT-AATACCCGCTTTCGCCGTGTTGGATAGATATATTACTTGGCACTATTGTTCTCTTGTAACAATGAGCAGACTTGCTGTCGAAG

			>Wittmackia_andersoniana
			????????????????????????????????????????????????????????????????????????????????????????????????????????????????????????????????????????????????????????????????????????????????????????????????????????????????????????????????????????????????????????????????????????????????????????????????????????????????????????????????????????????????????????????????????????????????

			>Wittmackia_antillana
			????????????????????????????????????????????????????????????????????????????????????????????????????????????????????????????????????????????????????????????????????????????????????????????????????????????????????????????????????????????????????????????????????????????????????????????????????????????????????????????????????????????????????????????????????????????????

			>Wittmackia_bicolor
			????????????????????????????????????????????????????????????????????????????????????????????????????????????????????????????????????????????????????????????????????????????????????????????????????????????????????????????????????????????????????????????????????????????????????????????????????????????????????????????????????????????????????????????????????????????????

			>Wittmackia_brasiliensis
			????????????????????????????????????????????????????????????????????????????????????????????????????????????????????????????????????????????????????????????????????????????????????????????????????????????????????????????????????????????????????????????????????????????????????????????????????????????????????????????????????????????????????????????????????????????????

			>Wittmackia_burle-marxii
			????????????????????????????????????????????????????????????????????????????????????????????????????????????????????????????????????????????????????????????????????????????????????????????????????????????????????????????????????????????????????????????????????????????????????????????????????????????????????????????????????????????????????????????????????????????????

			>Wittmackia_canaliculata
			????????????????????????????????????????????????????????????????????????????????????????????????????????????????????????????????????????????????????????????????????????????????????????????????????????????????????????????????????????????????????????????????????????????????????????????????????????????????????????????????????????????????????????????????????????????????

			>Wittmackia_caymanensis
			????????????????????????????????????????????????????????????????????????????????????????????????????????????????????????????????????????????????????????????????????????????????????????????????????????????????????????????????????????????????????????????????????????????????????????????????????????????????????????????????????????????????????????????????????????????????

			>Wittmackia_distans
			TCGCAGAAAGCTCTTTCCCTGCCTACTGGAATGGGCATTCTTTGCGCCAGCCCGAAAGCTCTAGAAGCTTCCAAAACAGCCAAGTCGGTCAGGGTTTTCTTCGACTGGAATGACTACTTGAAGTTCTACAAGCTCGGAACTTACTGGCCTTAACGCCTTCCATCCAACTACTTTACGGGCTGAGAGCAGCTCTGGATCTCATCTTTGAGGAGGGGCTCGACAATGTCATCGAAAGGCATCGTCGCTTAGGAAAGGCCACAAGGTTAAGTAAAAGCTATACATCT-AATACCCGCTTTCGCCGTGTTGGATAGATATATTACTTGGGACCATTGTTCTCTTGTAACAATGAGCAGGCTTGCTGTCGAAG

			>Wittmackia_eriostachya
			????????????????????????????????????????????????????????????????????????????????????????????????????????????????????????????????????????????????????????????????????????????????????????????????????????????????????????????????????????????????????????????????????????????????????????????????????????????????????????????????????????????????????????????????????????????????

			>Wittmackia_fawcettii
			????????????????????????????????????????????????????????????????????????????????????????????????????????????????????????????????????????????????????????????????????????????????????????????????????????????????????????????????????????????????????????????????????????????????????????????????????????????????????????????????????????????????????????????????????????????????

			>Wittmackia_froesii
			TCGCAGAAAGCTCTTTCCCTGCCTACTGGAATGGGGATTCTCTGCGCGAGCCCGAAAGCTCTAGAAGCTTCCAAAACAGCCAAGTCGGTCAGGGTTTTCTTCGACTGGAATGACTACTTGAAGTTCTACAAGCTCGGAACTTACTGGCCTTAACGCCTTCCATCCAACTACTTTATGGGCTGAGAGCAGCTCTGGATCTCATCTTTGAGGAGGGGCTCGACAATGTCATCGAAAGGCATCGTCGCTTAGGAAAGGCCACAAGGTTAAGTAAAAGCTATACATCT-AATACCCGCTTTCGCCGTGTTGGATAGATATATTACTTGGGACCATTGTTCTCTTGTAACAATGAGCAGGCTTGCTGTCGAAG

			>Wittmackia_incompta
			TCGCAGAAAGCTCTTTCCCTGCCTACTGGAATGGGGATTCTCTGCGCGAGCCCGAAAGCTCTAGAAGCTTCCAAAACAGCCAAGTCGGTCAGGGTTTTCTTCGACTGGAATGACTACTTGAAGTTCTACAAGCTCGGAACTTACTGGCCTTAACGCCTTCCATCCAACTACTTTACGGGCTGAGAGCAGCTCTGGATCTCATCTTTGAGGAGGGGCTCGACAATGTCATCGAAAGGCATCGTCGCTTAGGAAAGGCCACAAGGTTAAGTAAAAGCTATACATCT-AATACCCGCTTTCGCCGTGTTGGATAGATATATTACTTGGCACCATTGTTCTCTTGTAACAATGAGCAGGCTTGCTGTCGAAG

			>Wittmackia_inermis
			????????????????????????????????????????????????????????????????????????????????????????????????????????????????????????????????????????????????????????????????????????????????????????????????????????????????????????????????????????????????????????????????????????????????????????????????????????????????????????????????????????????????????????????????????????????????

			>Wittmackia_ituberaensis
			TCGCAGAAAGCTCTTTCCCTGCCTACTGGAATGGGGATTCTCTGCGCGAGCCCGAAAGCTCTAGAAGCTTCCAAAACAGCCAAGTCGGTCAGGGTTTTCTTCGACTGGAATGACTACTTGAAGTTCTACAAGCTCGGAACTTACTGGCCTTAACGCCTTCCATCCAACTACTTTACGGGCTGAGAGCAGCTCTGGATCTCATCTTTGAGGAGGGGCTCGACAACGTCATCGAAAGGCACCGTCGCTTAGGAAAGGCCACAAGGTTAAGTAAAAGCTATACATCT-AATACCCGCTTTCGCCGTGTTGGATAGATATATTACTTGGGACCATTGTTCTCTTGTAACAATGAGCAGGCTTGCTGTCGAAG

			>Wittmackia_jamaicana
			????????????????????????????????????????????????????????????????????????????????????????????????????????????????????????????????????????????????????????????????????????????????????????????????????????????????????????????????????????????????????????????????????????????????????????????????????????????????????????????????????????????????????????????????????????????????

			>Wittmackia_laesslei
			????????????????????????????????????????????????????????????????????????????????????????????????????????????????????????????????????????????????????????????????????????????????????????????????????????????????????????????????????????????????????????????????????????????????????????????????????????????????????????????????????????????????????????????????????????????????

			>Wittmackia_laevigata
			????????????????????????????????????????????????????????????????????????????????????????????????????????????????????????????????????????????????????????????????????????????????????????????????????????????????????????????????????????????????????????????????????????????????????????????????????????????????????????????????????????????????????????????????????????????????

			>Wittmackia_limae
			????????????????????????????????????????????????????????????????????????????????????????????????????????????????????????????????????????????????????????????????????????????????????????????????????????????????????????????????????????????????????????????????????????????????????????????????????????????????????????????????????????????????????????????????????????????????

			>Wittmackia_lingulata
			TCGCAGAAAGCTCTTTCCCTGCCTACTGGAATGGGGATTCTCTGCGCGAGCCCGAAAGCTCTAGAAGCTTCCAAAACAGCCAAGTCGGTCAGGGTTTTCTTCGACTGGAATGACTACTTGAAGTTCTACAAGCTCGGAACTTACTGGCCTTAACGCCTTCCATCCAACTACTTTACGGGCTGAGAGCAGCTCTGGATCTCATCTTTGAGGAGGGGCTCGACAATGTCATCGAAAGGCATCGTCGCTTAGGAAAGGCCACAAGGTTAAGTAAAAGCTATACATCT-AATACCCGCTTTCGCCGTGTTGGATAGATATATTACTTGGCACCATTGTTCTCTTGTAACAATGAGCAGGCTTGCTGTCGAAG

			>Wittmackia_lingulatoides
			TCGCAGAAAGCTCTTTCCCTGCCTACTGGAATGGGGATTCTCTGCGCGAGCCCGAAAGCTCTAGAAGCTTCCAAAACAGCCAAGTCGGTCAGGGTTTTCTTCGACTGGAATGACTACTTGAAGTTCTACAAGCTCGGAACTTACTGGCCTTAACGCCTTCCATCCAACTACTTTACGGGCTGAGAGCAGCTCTGGATCTCATCTTTGAGGAGGGGCTCGACAATGTCATCGAAAGGCATCGTCGCTTAGGAAAGGCCACAAGGTTAAGTAAAAGCTATACATCT-AATACCCGCTTTCGCCGTGTTGGATAGATATATTGTTTGGCACCATTGTTCTCTTGTAACAATGAGCAGGCTTGCTGTCGAAG

			>Wittmackia_linharesiorum
			????????????????????????????????????????????????????????????????????????????????????????????????????????????????????????????????????????????????????????????????????????????????????????????????????????????????????????????????????????????????????????????????????????????????????????????????????????????????????????????????????????????????????????????????????????????????

			>Wittmackia_maranguapensis
			????????????????????????????????????????????????????????????????????????????????????????????????????????????????????????????????????????????????????????????????????????????????????????????????????????????????????????????????????????????????????????????????????????????????????????????????????????????????????????????????????????????????????????????????????????????????

			>Wittmackia_mesoamericana
			????????????????????????????????????????????????????????????????????????????????????????????????????????????????????????????????????????????????????????????????????????????????????????????????????????????????????????????????????????????????????????????????????????????????????????????????????????????????????????????????????????????????????????????????????????????????

			>Wittmackia_negrilensis
			????????????????????????????????????????????????????????????????????????????????????????????????????????????????????????????????????????????????????????????????????????????????????????????????????????????????????????????????????????????????????????????????????????????????????????????????????????????????????????????????????????????????????????????????????????????????

			>Wittmackia_neoregelioides
			????????????????????????????????????????????????????????????????????????????????????????????????????????????????????????????????????????????????????????????????????????????????????????????????????????????????????????????????????????????????????????????????????????????????????????????????????????????????????????????????????????????????????????????????????????????????

			>Wittmackia_patentissima
			TCGCAGAAAGCTCTTTCCCTGCCTACTGGAATGGGGATTCTCTGCGCGAGCCCGAAAGCTCTAGAAGCTTCCAAAACAGCCAAGTCGGTCAGGGTTTTCTTCGACTGGAATGACTACTTGAAGTTCTACAAGCTCGGAACTTACTGGCCTTAACGCCTTCCATCCAACTACTTTATGGGCTGAGAGCAGCTCTGGATCTCATCTTTGAGGAGGGGCTCGACAATGTCATCGAAAGGCATCGTCGCTTAGGAAAGGCCACAAGGTTAAGTAAAAGCTATACATCT-AATACCCGCTTTCGCCATGTTGGATAGATATATTACTTGGGACCATTGTTCTCTTGTAACAATGAGCAGGCTTGCTGTCGAAG

			>Wittmackia_penduliflora
			????????????????????????????????????????????????????????????????????????????????????????????????????????????????????????????????????????????????????????????????????????????????????????????????????????????????????????????????????????????????????????????????????????????????????????????????????????????????????????????????????????????????????????????????????????????????

			>Wittmackia_pernambucentris
			TCGCAGAAAGCTCTTTCCCTGCCTACTGGAATGGGGATTCTCTGCGCGAGCCCGAAAGCTCTAGAAGCTTCCAAAACAGCCAAGTCGGTCAGGGTTTTCTTCGACTGGAATGACTACTTGAAGTTCTACAAGCTCGGAACTTACTGGCCTTAACGCCTTCCATCCAACTACTTTACGGGCTGAGAGCAGCTCTGGATCTCATCTTTGAGGAGGGGCTCGACAATGTCATCGAAAGGCATCGTCGCTTAGGAAAGGCTACAAGGTTAAGTATAAGCTATACATCT-AATAGCAGCTTTCGCCGTGTTGGATAGATATATTACTTGGGACCATTGTTCTCTTGTAACAATGAGCAGGCTTGCTGTCGAAG

			>Wittmackia_polycephala
			TCGCAGAAAGCTCTTTCCCTGCCTACTGGAATGGGSATTCTYTGCGCSAGCCCGAAAGCTCTAGAAGCTTCCAAAACAGCCAAGTCGGTCAGGGTTTTCTTCGACTGGAATGACTACTTGAAGTTCTACAAGCTCGGAACTTACTGGCCTTAACGCCTTCCATCCAACTACTTTACGGGCTGAGAGCAGCTCTGGATCTCATCTTTGAGGAGGGGCTCGACAATGTCATCGAAAGGCATCGTCGCTTAGGAAAGGCCACAAGGTTAAGTAAAAGCTATRCATCT-AATACCCGCTTTCGCCGTGTTGGATAGATATATTACTTGGGACCATTGTTCTCTTGTAACAATGAGCAGGCTTGCTGTCGAAG

			>Wittmackia_portoricensis
			????????????????????????????????????????????????????????????????????????????????????????????????????????????????????????????????????????????????????????????????????????????????????????????????????????????????????????????????????????????????????????????????????????????????????????????????????????????????????????????????????????????????????????????????????????????????

			>Wittmackia_silvana
			TCGCAGAAAGCTCTTTCCCTGCCTACTGGAATGGGGATTCTCTGCGCGAGCCCGAAAGCTCTAGAAGCTTCCAAAACAGCCAAGTCGGTCAGGGTTTTCTTCGACTGGAATGACTACTTGAAGTTCTACAAGCTCGGAACTTACTGGCCTTAACGCCTTCCATCCAACTACTTTACGGGCTGAGAGCAGCTCTGGATCTCATCTTTGAGGAGGGGCTCGACAACGTCATCGAAAGGCACCGTCGCTTAGGAAAGGCCACAAGGTTAAGTAAAAGCTATACGTAC-AATACCCGCTTTCGCCGTGTTGGATAGATATATTACTTGGCACTATTGTTCTCTTGTAACAATGAGCAGACTTGCTGTCGAAG

			>Wittmackia_spinulosa
			????????????????????????????????????????????????????????????????????????????????????????????????????????????????????????????????????????????????????????????????????????????????????????????????????????????????????????????????????????????????????????????????????????????????????????????????????????????????????????????????????????????????????????????????????????????????

			>Wittmackia_sulbahianensis
			TCGCAGAAAGCTCTTTCCCTGCCTACTGGAATGGGGATTCTCTGCGCGAGCCCGAAAGCTCTAGAAGCTTCCAAAACAGCCAAGTCGGTCAGGGTTTTCTTCGACTGGAATGACTACTTGAAGTTCTACAAGCTCGGAACTTACTGGCCTTAACGCCTTCCATCCAACTACTTTACGGGCTGAGAGCAGCTCTGGATCTCATCTTTGAGGAGGGGCTCGACAATGTCATCGAAAGGCATCGTCGCTTAGGAAAGGCCACAAGGTTAAGTAAAAGCTATACATMT-AATACCCGCTTTCGCCGTGTTGGATAGATATATTACTTGGCACCATTGTTCTCTTGTAACAATGAGCAGGCTTGCTGTCGAAG

			>Wittmackia_tentaculifera
			TCGCAGAAAGCTCTTTCCCTGCCTACTGGAATGGGGATTCTCTGCGCGAGCCCGAAAGCTCTAGAAGCTTCCAAAACAGCCAAGTCGGTCAGGGTTTTCTTCGACTGGAATGACTACTTGAAGTTCTACAAGCTCGGAACTTACTGGCCTTAACGCCTTCCATCCAACTACTTTACGGGCTGAGAGCAGCTCTGGATCTCATCTTTGAGGAGGGGCTCGACAACGTCATCGAAAGGCACCGTCGCTTAGGAAAGGCCACAAGGTTAAGTAAAAGCTATACATCT-AATACCCGCTTTCGCCGTGTTGGATAKATATATTACTTGGCACCATTGTTCTCTTGTAACAATGAGCAGGCTTGCTGTCGAAG

			>Wittmackia_turbinocalyx
			TCGCAGAAAGCTCTTTCCCTGCCTACTGGAATGGGGATTCTCTGCGCGAGCCCGAAAGCTCTAGAAGCTTCCAAAACAGCCAAGTCGGTCAGGGTTTTCTTCGACTGGAATGACTACTTGAAGTTCTACAAGCTCGGAACTTACTGGCCTTAACGCCTTCCATCCAACTACTTTACGGGCTGAGAGCAGCTCTGGATCTCATCTTTGAGGAGGGGCTCGACAATGTCATCGAAAGGCATCGTCGCTTAGGAAAGGCCACAAGGTTAAGTAAAAGCTATACATCT-AATACCCGCTTTCGCCGTGTTGGATAGATATATTACTTGGGACCATTGTTCTCTTGTAACAATGAGCAGGCTTGCTGTCGAAG

			>Wittmackia_urbaniana
			????????????????????????????????????????????????????????????????????????????????????????????????????????????????????????????????????????????????????????????????????????????????????????????????????????????????????????????????????????????????????????????????????????????????????????????????????????????????????????????????????????????????????????????????????????????????

			>Wittmackia_viridostigma
			????????????????????????????????????????????????????????????????????????????????????????????????????????????????????????????????????????????????????????????????????????????????????????????????????????????????????????????????????????????????????????????????????????????????????????????????????????????????????????????????????????????????????????????????????????????????

			>Wittrockia_abbreviata
			????????????????????????????????????????????????????????????????????????????????????????????????????????????????????????????????????????????????????????????????????????????????????????????????????????????????????????????????????????????????????????????????????????????????????????????????????????????????????????????????????????????????????????????????????????????????

			>Wittrockia_antillana
			????????????????????????????????????????????????????????????????????????????????????????????????????????????????????????????????????????????????????????????????????????????????????????????????????????????????????????????????????????????????????????????????????????????????????????????????????????????????????????????????????????????????????????????????????????????????

			>Wittrockia_cyathiformis
			TCGCAGAAAGCTCTTTCCCTGCCTACCGGAATGGGGATTCTCTGCGCGAGCCCGAAAGCTCTAGAAGCTTCCAAAACAGCCAAGTCGGTCAGGGTTTTCTTCGACTGGAATGACTACTTGAAGTTCTACAAGCTCGGAACTTACTGGCCTTAACGCCTTCCATCCAACTACTTTACGGGCTGAGAGCAGCTTTGGACCTCATCTTTGAGGAGGGGCTCGACAATGTCATCGAAAGGCATCGTCGCTTAGGAAAGGCCACAAGGTTAAGTAAAAGCTATACWTCTAAATRGCCGCTTTCGCCGTGTTGGATAGATATATTACTTKGCACTACTGTCCTCTYGTAACAATGGGCAGACTTGCCGTCGAAG

			>Wittrockia_gigantea
			TCGCAGAAAGCTCTTTCCCTGCCTACTGGAATGGGGATTCTCTGCGCGAGCCCGAAAGCTCTAGAAGCTTCCAAAACAGCCAAGTCGGTCAGGGTTTTCTTCGACTGGAATGACTACTTGAAGTTCTACAAGCTCGGAACTTACTGGCCTTAACGCCTTCCATCCAACTACTTTACGGGCTGAGAGCAGCTTTGGACCTCATCTTTGAGGAGGGGCTCGACAATGTCATCGAAAGGCATCGTCGCTTAGGAAAGGCCACAAGGTTAAGTAAAAGCTATACATCTAAATAGCCGCTTTCGCCGTGTTGGATAGATATATTACTTGGCACTACTGTCCTCTTGTAACAATGGGCAGACTTGCCGTCGAAG

			>Wittrockia_paulistana
			????????????????????????????????????????????????????????????????????????????????????????????????????????????????????????????????????????????????????????????????????????????????????????????????????????????????????????????????????????????????????????????????????????????????????????????????????????????????????????????????????????????????????????????????????????????????

			>Wittrockia_spiralipetala
			????????????????????????????????????????????????????????????????????????????????????????????????????????????????????????????????????????????????????????????????????????????????????????????????????????????????????????????????????????????????????????????????????????????????????????????????????????????????????????????????????????????????????????????????????????????????

			>Wittrockia_superba
			TCGCAGAAAGCTCTTTCCCTGCCTACTGGAATGGGGATTCTCTGCGCGAGCCCGAAAGCTCTAGAAGCTTCCAAAACAGCCAAGTCGGTCAGGGTTTTCTTCGACTGGAATGACTACTTGAAGTTCTACAAGCTCGGAACTTACTGGCCTTAACGCCTTCCATCCAACTACTTTACGGGCTGAGAGCAGCTTTGGACCTCATCTTTGAGGAGGGGCTTGACAATGTCATCGAAAGGCATCGTCGCTTAGGAAAGGCCACAAGGTTAAGTAAAAGCTATACATCTAAATAGCCGCTTTCGCCGTGTTGGATAGATATATTACTTTGCACTGTTGTCCTCTTGTAACAATGGGCAGACTCGCCGTCGAAG

			>Zizkaea_tuerckheimii
			????????????????????????????????????????????????????????????????????????????????????????????????????????????????????????????????????????????????????????????????????????????????????????????????????????????????????????????????????????????????????????????????????????????????????????????????????????????????????????????????????????????????????????????????????????????????

;atpb
	

			>Acanthostachys_pitcairnioides
			???????????????????????????????????TTTGGAATAATACCAAATCGAAAATAAATGTCCGATAGCAAGTTGATCAGTTAATTCAATAAGAGATAAATGGGAGATAGCACTCGATTTAGTTGGTACCACCCAACCGAATTCGATTCACTCGTTTACTCATTGAATGAGTAAATTTTCAAGTTCAGCCAATCTTTTTTTTTT-AAAAAAAAAATTGCAAGTAGATGAAATCGTGAATAAATGTGTTTCATTTCTATATCATTATATAAAAAAGGATCCATATTATATTACTTATGGAATTCGAACCCGAACTCGATTTATGATTTATTATTTGGATCTTATTGGCCTTTGTTCTTTATTTTTTATTTTTCATATAGATTTCCGTCTTTATATTATACCCTTTCGTGTATGAATTATGCCTATTTTCACATCTAGGATTTACATATACAACATATATTACTGTCAAGAGGGAATTTTTCTCAGTATTTAGATATTTAGATTCAAAATAAGAAAGGGATCAATTCAATTATAAACCTGCAAAACAAAGATTAGGATTGGGTTGGGTTGCGCTATATATATCAAAGAGTATACAATAATGATGTATTTGGTGAATCAAATACATGGTCTAATAACAAACCATTTTAACATAAAATTTTAATGAGTTGATAATATTAATTGAATATTGAATATTTTTTGAAAGATTTTTGTTAAAGGTTTCATTCATGCCTAATCCATATCGAGTAGACCTTGTTGTTGTG??

			>Acanthostachys_strobilacea
			ATAATTATAATAAAGTGAAATATGTCGAAATCCTTTTTGGAATAATACCAAATCGAAAATAAATGTCCGATAGCAAGTTGATCAGTTAATTCAATAAGAGATAAATGGGAGATAGCACTCGATTTAGTTGGTACCACCCAACCGAATCCGATTCACTCGTTTACTCATTCAATGAGTAAATTTTCAAGTTCAGCCAATCTTTTTTTTTTTAAAAAAAAAATTGCAAGTAGATGAAATCGTGAATAAATGTGTTTCATTTCTATATCATTATATAAAAAAAGATCCATATTATATTACTTATGGAATTCGAACCCGAACTCGATTTATGATTTATTATTTGGATCTTATTGGCCTTTGTTCTTTATTTTTGATTTTTCATATAGATTTCCGTCTTTATATTAT---------TGGATGAATTATGCCTATTTTCACATCTAGGATTTACATATACAACATATATTACTGTCAAGAGGGAATTTTTCTCAGTATTTAGATATTTAGATTCAAAATAAGAAAGGGATCAATTCAATTATAAACCTGCAAAACAAAGATTAGGATTGGGTTGGGTTGCGCTATATATATCAAAGAGTATACAATAATTATGTATTTGGTGAATCAAATACATGGTCTAATAACAAACCATTTTAACATAACATTTTAATGAGTTGATAATATTAATTGAATATTGAATATTTTTTGAAAGATTTTTGTTAAAGGTTTCATTCATGCCTAATCCATATCGAGTAGACCTTGTTGTTGTG??

			>Aechmea_abbreviata
			??????????????????????????????????????????????????????????????????????????????????????????????????????????????????????????????????????????????????????????????????????????????????????????????????????????????????????????????????????????????????????????????????????????????????????????????????????????????????????????????????????????????????????????????????????????????????????????????????????????????????????????????????????????????????????????????????????????????????????????????????????????????????????????????????????????????????????????????????????????????????????????????????????????????????????????????????????????????????????????????????????????????????????????????????????????????????????????????????????????????????????????????????????????????????????????????

			>Aechmea_aculeatosepala
			??????????????????????????????????????????????????????????????????????????????????????????????????????????????????????????????????????????????????????????????????????????????????????????????????????????????????????????????????????????????????????????????????????????????????????????????????????????????????????????????????????????????????????????????????????????????????????????????????????????????????????????????????????????????????????????????????????????????????????????????????????????????????????????????????????????????????????????????????????????????????????????????????????????????????????????????????????????????????????????????????????????????????????????????????????????????????????????????????????????????????????????????????????????????????????????????

			>Aechmea_aiuruocensis
			??????????????????????????????????????????????????????????????????????????????????????????????????????????????????????????????????????????????????????????????????????????????????????????????????????????????????????????????????????????????????????????????????????????????????????????????????????????????????????????????????????????????????????????????????????????????????????????????????????????????????????????????????????????????????????????????????????????????????????????????????????????????????????????????????????????????????????????????????????????????????????????????????????????????????????????????????????????????????????????????????????????????????????????????????????????????????????????????????????????????????????????????????????????????????????????????

			>Aechmea_alba
			?????????????????????????????????????TTGGATAATACCAAATCGAAAATAAATGTCCAATAGCAAGTTGATCAGTTAATTCAATAAGAGATAAATGGGAGATAGCACTCGATTTAGTTGGTACCACCCAACCGAATCCGATTCACTCGTTTACTCATTCAATGAGTAAATTTTCAAGTTCAGCCAATCTTTTTTTTTT-AAAAAAAAAATTGCAAGTAGATGAAATCGTGAATAAATGTGTTTCATTTCTATATCATTATATAAAAAAAGATCCATATTATATTACTTATGGAATTCGAACCCGAACTCGATTTATGATTTATTATTTGGATCTTATTGGCCTTTGTTCTTTATTTTTGATTTTTAATATAGATTTCCGTCTTTATATTATACCCTTTCGTGGATGAATTATGCCTATTTTCACATCTAGGATTTACATATACAACATATATTACTGTCAAGAGGGAATTTTTCTCAGTATTTAGATATTTAGATTCAAAATAAGAAAGGGATCAATTCAATTATAAACCTGCAAAACAAAGATTAGGATTGGGTTGGGTTGCGCTATATATATCAAAGAGTATACAATAATGATGTATTTGGTGAATCAAATACATGGTCTAATAACAAACCATTTTAACATAACATTTTAATGAGTTGATAATATTAATTGAATATTGAATATTTTTTGAAAGATTTTTGTTAAAGGTTTCATTCATGCCTAATCCATATCGAGTAGACCTTGTTGTTGTG??

			>Aechmea_alopecurus
			??????????????????????????????????????????????????????????????????????????????????????????????????????????????????????????????????????????????????????????????????????????????????????????????????????????????????????????????????????????????????????????????????????????????????????????????????????????????????????????????????????????????????????????????????????????????????????????????????????????????????????????????????????????????????????????????????????????????????????????????????????????????????????????????????????????????????????????????????????????????????????????????????????????????????????????????????????????????????????????????????????????????????????????????????????????????????????????????????????????????????????????????????????????????????????????????

			>Aechmea_angustifolia
			??????????????????????????????????????????????????????????????????????????????????????????????????????????????????????????????????????????????????????????????????????????????????????????????????????????????????????????????????????????????????????????????????????????????????????????????????????????????????????????????????????????????????????????????????????????????????????????????????????????????????????????????????????????????????????????????????????????????????????????????????????????????????????????????????????????????????????????????????????????????????????????????????????????????????????????????????????????????????????????????????????????????????????????????????????????????????????????????????????????????????????????????????????????????????????????????

			>Aechmea_aquilega
			??????????????????????????????????????????????????????????????????????????????????????????????????????????????????????????????????????????????????????????????????????????????????????????????????????????????????????????????????????????????????????????????????????????????????????????????????????????????????????????????????????????????????????????????????????????????????????????????????????????????????????????????????????????????????????????????????????????????????????????????????????????????????????????????????????????????????????????????????????????????????????????????????????????????????????????????????????????????????????????????????????????????????????????????????????????????????????????????????????????????????????????????????????????????????????????????

			>Aechmea_arenaria
			??????????????????????????????????????????????????????????????????????????????????????????????????????????????????????????????????????????????????????????????????????????????????????????????????????????????????????????????????????????????????????????????????????????????????????????????????????????????????????????????????????????????????????????????????????????????????????????????????????????????????????????????????????????????????????????????????????????????????????????????????????????????????????????????????????????????????????????????????????????????????????????????????????????????????????????????????????????????????????????????????????????????????????????????????????????????????????????????????????????????????????????????????????????????????????????????

			>Aechmea_bambusoides
			????????????????????????????????????TTTGGATAATACCAAATCGAAAATAAATGTCCGATAGCAAGTTGATCAGTTAATTCAATAAGAGATAAATGGGAGATAGCACTCGATTTAGTTGGTACCACCCAACCGAATCCGATTCACTCGTTTACTCATTCAATGAGTAAATTTTCAAGTTCAGCCAATCTTTTTTTTTT-AAAAAAAAGATTGCAAGTAGATGAAATCGTGAATAAATGTGTTTCATTTCTATATCATTATATAAAAAAAGATCCATATTATATTACTTATGGAATTCGAACCCGAACTCGATTTATGATTTATTATTTGGATCTTATTGGCCTTTGCTCTTTATTTTTGATTTTTCATATAGATTTCCATCTTTATATTATACCCTTTCGTGTATGAATTATGCCTATTTTCACATCTAGGATTTACATATACAACATATATTACTGTCAAGAGGGAATTTTTCTCAGTATTTAGATATTTAGATTCAAAATAAGAAAGGGATCAATTCAATTATAAACCTGCAAAACAAAGATTAGGATTGGGTTGGGTTGCGCTATATATATCAAAGAGTATACAATAATGATGTATTTGGTGAATCAAATACATGGTCTAATAACAAACCATTTTAACATAACATTTTAATGAGTTGATAATATTAATTGAATATTGAATATTTTTTGAAAGATTTTTGTTAAAGGTTTCATTCATGCCTAATCCATATCGAGTAGACCTTGTTGTTGTG??

			>Aechmea_blanchetiana
			??????????????????????????????????????????????????????????????????????????????????????????????????????????????????????????????????????????????????????????????????????????????????????????????????????????????????????????????????????????????????????????????????????????????????????????????????????????????????????????????????????????????????????????????????????????????????????????????????????????????????????????????????????????????????????????????????????????????????????????????????????????????????????????????????????????????????????????????????????????????????????????????????????????????????????????????????????????????????????????????????????????????????????????????????????????????????????????????????????????????????????????????????????????????????????????????

			>Aechmea_blumenavii
			??????????????????????????????????????????????????????????????????????????????????????????????????????????????????????????????????????????????????????????????????????????????????????????????????????????????????????????????????????????????????????????????????????????????????????????????????????????????????????????????????????????????????????????????????????????????????????????????????????????????????????????????????????????????????????????????????????????????????????????????????????????????????????????????????????????????????????????????????????????????????????????????????????????????????????????????????????????????????????????????????????????????????????????????????????????????????????????????????????????????????????????????????????????????????????????????

			>Aechmea_bocainensis
			??????????????????????????????????????????????????????????????????????????????????????????????????????????????????????????????????????????????????????????????????????????????????????????????????????????????????????????????????????????????????????????????????????????????????????????????????????????????????????????????????????????????????????????????????????????????????????????????????????????????????????????????????????????????????????????????????????????????????????????????????????????????????????????????????????????????????????????????????????????????????????????????????????????????????????????????????????????????????????????????????????????????????????????????????????????????????????????????????????????????????????????????????????????????????????????????

			>Aechmea_bracteata
			??????????????????????????????????????????????????????????????????????????????????????????????????????????????????????????????????????????????????????????????????????????????????????????????????????????????????????????????????????????????????????????????????????????????????????????????????????????????????????????????????????????????????????????????????????????????????????????????????????????????????????????????????????????????????????????????????????????????????????????????????????????????????????????????????????????????????????????????????????????????????????????????????????????????????????????????????????????????????????????????????????????????????????????????????????????????????????????????????????????????????????????????????????????????????????????????

			>Aechmea_bracteata_var._pacifica
			??????????????????????????????????????????????????????????????????????????????????????????????????????????????????????????????????????????????????????????????????????????????????????????????????????????????????????????????????????????????????????????????????????????????????????????????????????????????????????????????????????????????????????????????????????????????????????????????????????????????????????????????????????????????????????????????????????????????????????????????????????????????????????????????????????????????????????????????????????????????????????????????????????????????????????????????????????????????????????????????????????????????????????????????????????????????????????????????????????????????????????????????????????????????????????????????

			>Aechmea_brassicoides
			??????????????????????????????????????????????????????????????????????????????????????????????????????????????????????????????????????????????????????????????????????????????????????????????????????????????????????????????????????????????????????????????????????????????????????????????????????????????????????????????????????????????????????????????????????????????????????????????????????????????????????????????????????????????????????????????????????????????????????????????????????????????????????????????????????????????????????????????????????????????????????????????????????????????????????????????????????????????????????????????????????????????????????????????????????????????????????????????????????????????????????????????????????????????????????????????

			>Aechmea_brevicollis
			??????????????????????????????????????????????????????????????????????????????????????????????????????????????????????????????????????????????????????????????????????????????????????????????????????????????????????????????????????????????????????????????????????????????????????????????????????????????????????????????????????????????????????????????????????????????????????????????????????????????????????????????????????????????????????????????????????????????????????????????????????????????????????????????????????????????????????????????????????????????????????????????????????????????????????????????????????????????????????????????????????????????????????????????????????????????????????????????????????????????????????????????????????????????????????????????

			>Aechmea_bromeliifolia
			ATAATTATAATAAAGTGAAATATGTCGAAATCCTTTTTGGAATAATACCAAATCGAAAATAAATGTCCGATAGCAAGTTGATCAGTTAATTCAATAAGAGATAAATGGGAGATAGCACTCGATTTAGTTGGTACCACCCAACCGAATCCGATTCACTCGTTTACTCATTGAATGAGTCAATTTTCAAGTTCAGCCAATC--TTTTTTTTTGAAAAAAAAATTGCAAGTAGATTAAATCATGAATAAATGTGTTTCATTTCTATATCATTATAGAAAAAAAGATCCATATTATATTACTTATGGAATTCGAACCCGAACTCGATTTATGATTTATTATTTGGATCTTATTGGCCTTTGTTCTTTATTTTTGATTTTTCATATAGATTTCCGTCTTTATATTATACCCTTTCGTGGATGAATTATGCCTATTTTCACATCTAGGATTTACATATACAACATATATTACTGTCAAGAGGGAATTTTTCTCAGTATTTAGATATTTAGATTCAAAATAAGAAAGGGATCAATTCAATTATAAACCTGCAAAACAAAGATTAGGATTGGGTTGGGTTGCGCTATATATATCAAAGAGTATACAATAATGATGTATTTGGTGAATCAAATACATGGTCTAATAACAAACCATTTTAACATAACATTTTAATGAGTTGATAATATTAATTGAATATTGAATATTTTTTGAAAGATTTTTGTTAAAGGTTTCATTCATGCCTAATCCATATCGAGTAGACCTTGTTGTTGTGAG

			>Aechmea_bromeliifolia_var._albobracteata
			??????????????????????????????????????????????????????????????????????????????????????????????????????????????????????????????????????????????????????????????????????????????????????????????????????????????????????????????????????????????????????????????????????????????????????????????????????????????????????????????????????????????????????????????????????????????????????????????????????????????????????????????????????????????????????????????????????????????????????????????????????????????????????????????????????????????????????????????????????????????????????????????????????????????????????????????????????????????????????????????????????????????????????????????????????????????????????????????????????????????????????????????????????????????????????????????

			>Aechmea_bruggeri
			??????????????????????????????????????????????????????????????????????????????????????????????????????????????????????????????????????????????????????????????????????????????????????????????????????????????????????????????????????????????????????????????????????????????????????????????????????????????????????????????????????????????????????????????????????????????????????????????????????????????????????????????????????????????????????????????????????????????????????????????????????????????????????????????????????????????????????????????????????????????????????????????????????????????????????????????????????????????????????????????????????????????????????????????????????????????????????????????????????????????????????????????????????????????????????????????

			>Aechmea_caesia
			??????????????????????????????????????????????????????????????????????????????????????????????????????????????????????????????????????????????????????????????????????????????????????????????????????????????????????????????????????????????????????????????????????????????????????????????????????????????????????????????????????????????????????????????????????????????????????????????????????????????????????????????????????????????????????????????????????????????????????????????????????????????????????????????????????????????????????????????????????????????????????????????????????????????????????????????????????????????????????????????????????????????????????????????????????????????????????????????????????????????????????????????????????????????????????????????

			>Aechmea_calyculata
			ATAATTATAATAAAGTGAAATATGTCGAAATCCTTTTTGGAATAATACCAAATCGAAAATAAATGTCCGATAGCAAGTTGATCAGTTAATTCAATAAGAGATAAATGGGAGATAGCACTCGATTTAGTTGGTACCACCCAACCGAATCCGATTCACTCGTTTACTCATTGAATGAGTAAATTTTCAAGTTCAGCCAATC-TTTTTTTTTGAAAAAAAAATTGGCAAGTAGATTAAATCGTGAATAAATGTGTTTCATTTCTATATCATTATATAAAAAAAGATCCATATTATATTACTTATGGAATTCGAACCCGAACTCGATTTATGATTTATTGTTTGGATCTTATTGGCCTTTGTTCTTTATTTTTGATTTTTCATATAGATTTCCGTCTTTATATTATACCCTTTCGTGGATGAATTATGCCTATTTTCACATCTAGGATTTACATATACAACATATATTACTGTCAAGAGGGAATTTTTCTCAGTATTTAGATATTTAGATTCAAAATAAGAAAGGGATCAATTCAATTATAAACCTGCAAAACAAAGATTAGGATTGGGTTGGGTTGCGCTATATATATCAAAGAGTATACAATAATGATGTATTTGGTGAATCAAATACATGGTCTAATAACAAACCATTTTAACATAACATTTTAATGAGTTGATAATATTAATTGAATATTGAATATTTTTTGAAAGATTTTTGTTAAAGGTTTCATTCATGCCTAATCCATATCGAGTAGACCTTGTTGTTGTG??

			>Aechmea_candida
			??????????????????????????????????????????????????????????????????????????????????????????????????????????????????????????????????????????????????????????????????????????????????????????????????????????????????????????????????????????????????????????????????????????????????????????????????????????????????????????????????????????????????????????????????????????????????????????????????????????????????????????????????????????????????????????????????????????????????????????????????????????????????????????????????????????????????????????????????????????????????????????????????????????????????????????????????????????????????????????????????????????????????????????????????????????????????????????????????????????????????????????????????????????????????????????????

			>Aechmea_capixabae
			??????????????????????????????????????????????????????????????????????????????????????????????????????????????????????????????????????????????????????????????????????????????????????????????????????????????????????????????????????????????????????????????????????????????????????????????????????????????????????????????????????????????????????????????????????????????????????????????????????????????????????????????????????????????????????????????????????????????????????????????????????????????????????????????????????????????????????????????????????????????????????????????????????????????????????????????????????????????????????????????????????????????????????????????????????????????????????????????????????????????????????????????????????????????????????????????

			>Aechmea_carvalhoi
			??????????????????????????????????????????????????????????????????????????????????????????????????????????????????????????????????????????????????????????????????????????????????????????????????????????????????????????????????????????????????????????????????????????????????????????????????????????????????????????????????????????????????????????????????????????????????????????????????????????????????????????????????????????????????????????????????????????????????????????????????????????????????????????????????????????????????????????????????????????????????????????????????????????????????????????????????????????????????????????????????????????????????????????????????????????????????????????????????????????????????????????????????????????????????????????????

			>Aechmea_castanea
			??????????????????????????????????????????????????????????????????????????????????????????????????????????????????????????????????????????????????????????????????????????????????????????????????????????????????????????????????????????????????????????????????????????????????????????????????????????????????????????????????????????????????????????????????????????????????????????????????????????????????????????????????????????????????????????????????????????????????????????????????????????????????????????????????????????????????????????????????????????????????????????????????????????????????????????????????????????????????????????????????????????????????????????????????????????????????????????????????????????????????????????????????????????????????????????????

			>Aechmea_castelnavi
			??????????????????????????????????????????????????????????????????????????????????????????????????????????????????????????????????????????????????????????????????????????????????????????????????????????????????????????????????????????????????????????????????????????????????????????????????????????????????????????????????????????????????????????????????????????????????????????????????????????????????????????????????????????????????????????????????????????????????????????????????????????????????????????????????????????????????????????????????????????????????????????????????????????????????????????????????????????????????????????????????????????????????????????????????????????????????????????????????????????????????????????????????????????????????????????????

			>Aechmea_castelnavii
			??????????????????????????????????????????????????????????????????????????????????????????????????????????????????????????????????????????????????????????????????????????????????????????????????????????????????????????????????????????????????????????????????????????????????????????????????????????????????????????????????????????????????????????????????????????????????????????????????????????????????????????????????????????????????????????????????????????????????????????????????????????????????????????????????????????????????????????????????????????????????????????????????????????????????????????????????????????????????????????????????????????????????????????????????????????????????????????????????????????????????????????????????????????????????????????????

			>Aechmea_caudata
			??????????????????????????????????????????????????????????????????????????????????????????????????????????????????????????????????????????????????????????????????????????????????????????????????????????????????????????????????????????????????????????????????????????????????????????????????????????????????????????????????????????????????????????????????????????????????????????????????????????????????????????????????????????????????????????????????????????????????????????????????????????????????????????????????????????????????????????????????????????????????????????????????????????????????????????????????????????????????????????????????????????????????????????????????????????????????????????????????????????????????????????????????????????????????????????????

			>Aechmea_cephaloides
			??????????????????????????????????????????????????????????????????????????????????????????????????????????????????????????????????????????????????????????????????????????????????????????????????????????????????????????????????????????????????????????????????????????????????????????????????????????????????????????????????????????????????????????????????????????????????????????????????????????????????????????????????????????????????????????????????????????????????????????????????????????????????????????????????????????????????????????????????????????????????????????????????????????????????????????????????????????????????????????????????????????????????????????????????????????????????????????????????????????????????????????????????????????????????????????????

			>Aechmea_chantinii
			ATAATTATAATAAAGTGAAATATGTCGAAATCCTTTTTGGAATAATACCAAATTGAAAATAAATGTCCGATAGCAAGTTGATCAGTTAATTCAATAAGAGATAAATGGGAGATAGCACTCGATTTAGTTGGTACCACCCAACCGAATCCGATTCACCCGTTTACTCATTCAATGAGTAAATTTTCAAGTTCAGCCAATCTTTTTTTTTTTAAAAAAAAAATTGCAAGTAGATGAAATCGTGAATAAATGTGTTTCATTTCTATATCATTATATAAAAAAAGATCCATATTATATTACTTATGGAATTCGAACCCGAACTCGATTTATGATTTATTATTTGGATCTTATTGGCCTTTGTTCTTTATTTTTGATTTTTCATATAGATTTCCGTCTTTATATTATACCCTTTCGTGGATGAATTATGCCTATTTTCACATCTAGGATTTACATATACAACATATATTACTGTCAAGAGGGAATTTTTCTCAGTATTTAGATATTTAGATTCAAAATAAGAAAGGGATCAATTCAATTATAAACCTGCAAAACAAAGATTAGGATTGGGTTGGGTTGCGCTATATATATCAAAGAGTATACAATAATGATGTATTTGGTGAATTAAATACATGGTCTAATAACAAACCATTTTAACATAACATTTTAATGAGTTGATAATATTAATTGAATATTGAATATTTTTTGAAAGATTTTTGTTAAAGGTTTCATTCATGCCTAATCCATATCGAGTAGACCTTGTTGTTGTG??

			>Aechmea_chrysocoma
			??????????????????????????????????????????????????????????????????????????????????????????????????????????????????????????????????????????????????????????????????????????????????????????????????????????????????????????????????????????????????????????????????????????????????????????????????????????????????????????????????????????????????????????????????????????????????????????????????????????????????????????????????????????????????????????????????????????????????????????????????????????????????????????????????????????????????????????????????????????????????????????????????????????????????????????????????????????????????????????????????????????????????????????????????????????????????????????????????????????????????????????????????????????????????????????????

			>Aechmea_coelestis
			??????????????????????????????????????????????????????????????????????????????????????????????????????????????????????????????????????????????????????????????????????????????????????????????????????????????????????????????????????????????????????????????????????????????????????????????????????????????????????????????????????????????????????????????????????????????????????????????????????????????????????????????????????????????????????????????????????????????????????????????????????????????????????????????????????????????????????????????????????????????????????????????????????????????????????????????????????????????????????????????????????????????????????????????????????????????????????????????????????????????????????????????????????????????????????????????

			>Aechmea_colombiana
			????????????????????????????????????TTGGAATAATACCAAATCGAAAATAAATGTCCGATAGCAAGTTGATCAGTTAATTCAATAAGAGATAAATGGGAGATAGCACTCGATTTAGTTGGTACCACCCAACCGAATCCGATTCACTCGTTTACTCATTCAATGAGTAAATTTTCAAGTTCAGCCAATC-TTTTTTTTTTAAAAAAAAAATTGCAAGTAGATGAAATCGTGAATAAATGTGTTTCATTTCTATATCATTATATAAAAAAAGATCCATATTATATTACTTATGGAATTCGAACCCGAACTCGATTTATGATTTATTATTTGGATCTTATTGGCCTTTGTTCTTTATTTTTGATTTTTCATATAGATTTCCGTCTTTATATTATACCCTTTCGTGGATGAATTATGCCTATTTTCACATCTAGGATTTACATATACAACATATATTACTGTCAAGAGGGAATTTTTCTCAGTATTTAGATATTTAGATTCAAAATAAGAAAGGGATCAATTCAATTATAAACCTGCAAAACAAAGATTAGGATTGGGTTGGGTTGCGCTATATATATCAAAGAGTATACAATAATGATGTATTTGGTGAATCAAATACATGGTCTAATAACAAACCATTTTAACATAACATTTTAATGAGTTGATAATATTAATTGAATATTGAATATTTTTTGAAAGATTTTTGTTAAAGGTTTCATTCATGCCTAATCCATATCGAGTAGACCTTGTTGTTGTG??

			>Aechmea_comata
			??????????????????????????????????????????????????????????????????????????????????????????????????????????????????????????????????????????????????????????????????????????????????????????????????????????????????????????????????????????????????????????????????????????????????????????????????????????????????????????????????????????????????????????????????????????????????????????????????????????????????????????????????????????????????????????????????????????????????????????????????????????????????????????????????????????????????????????????????????????????????????????????????????????????????????????????????????????????????????????????????????????????????????????????????????????????????????????????????????????????????????????????????????????????????????????????

			>Aechmea_contracta
			??????????????????????????????????????????????????????????????????????????????????????????????????????????????????????????????????????????????????????????????????????????????????????????????????????????????????????????????????????????????????????????????????????????????????????????????????????????????????????????????????????????????????????????????????????????????????????????????????????????????????????????????????????????????????????????????????????????????????????????????????????????????????????????????????????????????????????????????????????????????????????????????????????????????????????????????????????????????????????????????????????????????????????????????????????????????????????????????????????????????????????????????????????????????????????????????

			>Aechmea_correia-araujoi
			??????????????????????????????????????????????????????????????????????????????????????????????????????????????????????????????????????????????????????????????????????????????????????????????????????????????????????????????????????????????????????????????????????????????????????????????????????????????????????????????????????????????????????????????????????????????????????????????????????????????????????????????????????????????????????????????????????????????????????????????????????????????????????????????????????????????????????????????????????????????????????????????????????????????????????????????????????????????????????????????????????????????????????????????????????????????????????????????????????????????????????????????????????????????????????????????

			>Aechmea_costantinii
			??????????????????????????????????????????????????????????????????????????????????????????????????????????????????????????????????????????????????????????????????????????????????????????????????????????????????????????????????????????????????????????????????????????????????????????????????????????????????????????????????????????????????????????????????????????????????????????????????????????????????????????????????????????????????????????????????????????????????????????????????????????????????????????????????????????????????????????????????????????????????????????????????????????????????????????????????????????????????????????????????????????????????????????????????????????????????????????????????????????????????????????????????????????????????????????????

			>Aechmea_cucullata
			??????????????????????????????????????????????????????????????????????????????????????????????????????????????????????????????????????????????????????????????????????????????????????????????????????????????????????????????????????????????????????????????????????????????????????????????????????????????????????????????????????????????????????????????????????????????????????????????????????????????????????????????????????????????????????????????????????????????????????????????????????????????????????????????????????????????????????????????????????????????????????????????????????????????????????????????????????????????????????????????????????????????????????????????????????????????????????????????????????????????????????????????????????????????????????????????

			>Aechmea_cylindrata
			??????????????????????????????????????????????????????????????????????????????????????????????????????????????????????????????????????????????????????????????????????????????????????????????????????????????????????????????????????????????????????????????????????????????????????????????????????????????????????????????????????????????????????????????????????????????????????????????????????????????????????????????????????????????????????????????????????????????????????????????????????????????????????????????????????????????????????????????????????????????????????????????????????????????????????????????????????????????????????????????????????????????????????????????????????????????????????????????????????????????????????????????????????????????????????????????

			>Aechmea_dactylina
			??????????????????????????????????????????????????????????????????????????????????????????????????????????????????????????????????????????????????????????????????????????????????????????????????????????????????????????????????????????????????????????????????????????????????????????????????????????????????????????????????????????????????????????????????????????????????????????????????????????????????????????????????????????????????????????????????????????????????????????????????????????????????????????????????????????????????????????????????????????????????????????????????????????????????????????????????????????????????????????????????????????????????????????????????????????????????????????????????????????????????????????????????????????????????????????????

			>Aechmea_dealbata
			??????????????????????????????????????????????????????????????????????????????????????????????????????????????????????????????????????????????????????????????????????????????????????????????????????????????????????????????????????????????????????????????????????????????????????????????????????????????????????????????????????????????????????????????????????????????????????????????????????????????????????????????????????????????????????????????????????????????????????????????????????????????????????????????????????????????????????????????????????????????????????????????????????????????????????????????????????????????????????????????????????????????????????????????????????????????????????????????????????????????????????????????????????????????????????????????

			>Aechmea_digitata
			??????????????????????????????????????????????????????????????????????????????????????????????????????????????????????????????????????????????????????????????????????????????????????????????????????????????????????????????????????????????????????????????????????????????????????????????????????????????????????????????????????????????????????????????????????????????????????????????????????????????????????????????????????????????????????????????????????????????????????????????????????????????????????????????????????????????????????????????????????????????????????????????????????????????????????????????????????????????????????????????????????????????????????????????????????????????????????????????????????????????????????????????????????????????????????????????

			>Aechmea_discordiae
			??????????????????????????????????????????????????????????????????????????????????????????????????????????????????????????????????????????????????????????????????????????????????????????????????????????????????????????????????????????????????????????????????????????????????????????????????????????????????????????????????????????????????????????????????????????????????????????????????????????????????????????????????????????????????????????????????????????????????????????????????????????????????????????????????????????????????????????????????????????????????????????????????????????????????????????????????????????????????????????????????????????????????????????????????????????????????????????????????????????????????????????????????????????????????????????????

			>Aechmea_disjuncta
			??????????????????????????????????????????????????????????????????????????????????????????????????????????????????????????????????????????????????????????????????????????????????????????????????????????????????????????????????????????????????????????????????????????????????????????????????????????????????????????????????????????????????????????????????????????????????????????????????????????????????????????????????????????????????????????????????????????????????????????????????????????????????????????????????????????????????????????????????????????????????????????????????????????????????????????????????????????????????????????????????????????????????????????????????????????????????????????????????????????????????????????????????????????????????????????????

			>Aechmea_distichantha
			??????????????????????????????????????????????????????????????????????????????????????????????????????????????????????????????????????????????????????????????????????????????????????????????????????????????????????????????????????????????????????????????????????????????????????????????????????????????????????????????????????????????????????????????????????????????????????????????????????????????????????????????????????????????????????????????????????????????????????????????????????????????????????????????????????????????????????????????????????????????????????????????????????????????????????????????????????????????????????????????????????????????????????????????????????????????????????????????????????????????????????????????????????????????????????????????

			>Aechmea_distichantha_var._glaziovii
			ATCATTATAATAAAGTGAAATATGTCGAAATCCTTTTTGGAATAATACCAAATC-AAAATAAATGTCCGATAGCAAGTTGATCAGTTAATTCAATAAGAGATAAATGGGAGATAGCACTCGATTTAGTTGGTACCACCCAACCGAATCCGATTCACTCGTTTACTCATTCAATGAGTAAATTTTCAAGTTCAGCCAATCTTTTTTTTTTTAAAAAAAAAATTGCAAGTAGATGAAATCGTGAATAAATGTGTTTCATTTCTATATCATTATATAAAAAAAGATCCATATTATATTACTTATGGAATTCGAACCCGAACTCGATTTATGATTTATTATTTGGATCTTATTGGCCTTTGTTCTTTATTTTTGATTTTTAATATAGATTTCCGTCTTTATATTATACCCTTTCGTGGATGAATTATGCCTATTTTCACATCTAGGATTTACATATACAACATATATTACTGTCAAGAGGGAATTTTTCTCAGTATTTAGATATTTAGATTCAAAATAAGAAAGGGATCAATTCAATTATAAACCTGCAAAACAAAGATTAGGATTGGGTTGGGTTGCGCTATATATATCAAAGAGTATACAATAATGATGTATTTGGTGAATCAAATACATGGTCTAATAACAAACCATTTTAACATAACATTTTAATGAGTTGATAATATTAATTGAATATTGAATATTTTTTGAAAGATTTTTGTTAAAGGTTTCATTCATGCCTAATCCATATCGAGTAGACCTTGTTGTTGTG??

			>Aechmea_egleriana
			??????????????????????????????????????????????????????????????????????????????????????????????????????????????????????????????????????????????????????????????????????????????????????????????????????????????????????????????????????????????????????????????????????????????????????????????????????????????????????????????????????????????????????????????????????????????????????????????????????????????????????????????????????????????????????????????????????????????????????????????????????????????????????????????????????????????????????????????????????????????????????????????????????????????????????????????????????????????????????????????????????????????????????????????????????????????????????????????????????????????????????????????????????????????????????????????

			>Aechmea_eurycorymbus
			??????????????????????????????????????????????????????????????????????????????????????????????????????????????????????????????????????????????????????????????????????????????????????????????????????????????????????????????????????????????????????????????????????????????????????????????????????????????????????????????????????????????????????????????????????????????????????????????????????????????????????????????????????????????????????????????????????????????????????????????????????????????????????????????????????????????????????????????????????????????????????????????????????????????????????????????????????????????????????????????????????????????????????????????????????????????????????????????????????????????????????????????????????????????????????????????

			>Aechmea_farinosa
			ATAATTATAATAAAGTGAAATATGTCGAAATCCTTTTTGGAATAATACCAAATCGAAAATAAATGTCCGATAGCAAGTTGATCAGTTAATTCAATAAGAGATAAATGGGAGATAGCACTCGATTTAGTTGGTACCACCCAACCGAATCCGATTCACTCGTTTACTCATTCAATGAGTAAATTTTCAAGTTCAGCCAATC-TTTTTTTTTGAAAAAAAAAATTGCAAGTAGATGAAATCGTGAATAAATGTGTTTCATTTCTATATCATTATATAAAAAAAGATCCATATTATATTACTTATGGAATTCGAACCCGAACTCGATTTATGATTTATTATTTGGATCTTATTGGCCTTTGTTCTTTATTTTTGATTTTTCATATGGATTTCCGTCTTTATATTATACCCTTTCGTGGATGAATTATGCCTATTTTCACATCTAGGATTTACATATACAACATATATTACTGTCAAGAGGGAATTTTTCTCAGTATTTAGATATTTAGATTCAAAATAAGAAAGGGATCAATTCAATTATAAACCTGCAAAACAAAGATTAGGATTGGGTTGGGTTGCGCTATATATATCAAAGAGTATACAATAATGATGTATTTGGTGAATCAAATACATGGTCTAATAACAAACCATTTTAACATAACATTTTAATGAGTTGATAATATTAATTGAATATTGAATATTTTTTGAAAGATTTTTGTTAAAGGTTTCATTCATGCCTAATCCATATCGAGTAGACCTTGTTGTTGTG??

			>Aechmea_fasciata
			ATAATTATAATAAAGTGAAATATGTCGAAATCCTTTTTGGAATAATACCAAATCGAAAATAAATGTCCGATAGCAAGTTGATCAGTTAATTCAATAAGAGATAAATGGGAGATAGCACTCGATTTAGTTGGTACCACCCAACCGAATCCGATTCACTCGTTTACTCATTCAATGAGTAAATTTTCAAGTTCAGCCAATCTTTTTTTTTTTAAAAAAGAAATTGCAAGTAGATGAAATCGTGAATAAATGTGTTTCATTTCTATATCATTATATAAAAAAAGATCCATATTATATTACTTATGGAATTCGAACCCGAACTCGATTTATGATTTATTATTTGGATCTTATTGGCCTTTGTTCTTTATTTTTGATTTTTCATATAGATTTCCGTCTTTATATTATACCCTTTCGTGGATGAATTATGCCTATTTTCACATCTAGGATTTACATATACAACATATATTACTGTCAAGAGGGAATTTTTCTCAGTATTTAGATATTTAGATTCAAAATAAGAAAGGG-----TTCAATTATAAACCTGCAAAACAAAGATTAGGATTGGGTTGGGTTGCGCTATATATATCAAAGAGTATACAATAATGATGTATTTGGTGAATCAAATACATGGTCTAATAACAAACCATTTTAACATAACATTTTAATGAGTTGATAATATTAATTGAATATTGAATATTTTTTGAAAGATTTTTGTTAAAGGTTTCATTCATGCCTAATCCATATCGAGTAGACCTTGTTGTTGTG??

			>Aechmea_filicaulis
			ATAATTATAATAAAGTGAAATATGTCGAAATCCTTTTTGGAATAATACCAAATTGAAAATAAATGTCCGATAGCAAGTTGATCAGTTAATTCAATAAGAGATAAATGGGAGATAGCACTCGATTTAGTTGGTACCACCCAACCGAATCCGATTCACTCGTTTACTCATTGAATGAGTAAATTTTCAAGTTCAGCCAATC---TTTTTTTTAAAAAAAAAATTGCAAGTAGATGAAATCGTGAATAAATGTGTTTCATTTCTATATCATTATATAAAAAAAGATCCATATTATATTACTTATGGAATTCGAACCCGAACTCGATTTATGATTTATTATTTGGATCTTATTGGCCTTTGTTCTTTATTTTTGATTTTTCATATAGATTTCCGTCTTTATATTATACCCTTTCGTGGATGAATTATGCCTATTTTCACATCTAGGATTTACATATACAACATATATTACTGTCAAGAGGGAATTTTTCTCAGTATTTAGATATTTAGATTCAAAATAAGAAAGGGATCAATTCAATTATAAACCTGCAAAACAAAGATTAGGATTGGGTTGGGTTGCGCTATATATATCAAAGAGTATACAATAATGATGTATTTGGTGAATCAAATACATGGTCTAATAACAAACCATTTTAACATAACATTTTAATGAGTTGATAATATTAATTGAATATTGAATATTTTTTGAAAGATTTTTGTTAAAGGTTTCATTCATGCCTAATCCATATCGAGTAGACCTTGTTGTTGTG??

			>Aechmea_flavorosea
			??????????????????????????????????????????????????????????????????????????????????????????????????????????????????????????????????????????????????????????????????????????????????????????????????????????????????????????????????????????????????????????????????????????????????????????????????????????????????????????????????????????????????????????????????????????????????????????????????????????????????????????????????????????????????????????????????????????????????????????????????????????????????????????????????????????????????????????????????????????????????????????????????????????????????????????????????????????????????????????????????????????????????????????????????????????????????????????????????????????????????????????????????????????????????????????????

			>Aechmea_floribunda
			??????????????????????????????????????????????????????????????????????????????????????????????????????????????????????????????????????????????????????????????????????????????????????????????????????????????????????????????????????????????????????????????????????????????????????????????????????????????????????????????????????????????????????????????????????????????????????????????????????????????????????????????????????????????????????????????????????????????????????????????????????????????????????????????????????????????????????????????????????????????????????????????????????????????????????????????????????????????????????????????????????????????????????????????????????????????????????????????????????????????????????????????????????????????????????????????

			>Aechmea_fosteriana
			??????????????????????????????????????????????????????????????????????????????????????????????????????????????????????????????????????????????????????????????????????????????????????????????????????????????????????????????????????????????????????????????????????????????????????????????????????????????????????????????????????????????????????????????????????????????????????????????????????????????????????????????????????????????????????????????????????????????????????????????????????????????????????????????????????????????????????????????????????????????????????????????????????????????????????????????????????????????????????????????????????????????????????????????????????????????????????????????????????????????????????????????????????????????????????????????

			>Aechmea_fulgens
			ATAATTATAATAAAGTGAAATATGTCGAAATCCTTTTTGGAATAATACCAAATCGAAAATAAATGTCCGATAGCAAGTTGATCAGTTAATTCAATAAGAGATAAATGGGAGATAGCACTCGATTTAGTTGGTACCACCCAACCGAATCCGATTCACTCGTTTACTCATTCAATGAGTAAATTTTCAAGTTCAGCCAATC-TTTTTTTTTGAAAAAAAAAATTGCAAGTAGATGAAATCGTGAATAAATGTGTTTCATTTCTATATCATTATATAAAAAAAGATCCATATTATATTACTTATGGAATTCGAACCCGAACTCGATTTATGATTTATTATTTGGATCTTATTGGCCTTTGTTCTTTATTTTTGATTTTTCATATAGATTTCCGTCTTTATATTATACCCTTTCGTGGATGAATTATGCCTATTTTCACATCTAGGATTTACATATACAACATATATTACTGTCAAGAGGGAATTTTTCTCAGTATTTAGATATTTAGATTCAAAATAAGAAAGGGATCAATTCAATTATAAACCTGCAAAACAAAGATTAGGATTGGGTTGGGTTGCGCTATATATATCAAAGAGTATACAATAATGATGTATTTGGTGAATCAAATACATGGTCTAATAACAAACCATTTTAACATAACATTTTAATGAGTTGATAATATTAATTGAATATTGAATATTTTTTGAAAGATTTTTGTTAAAGGTTTCATTCATGCCTAATCCATATCGAGTAGACCTTGTTGTTGTG??

			>Aechmea_fulgens_var._discolor
			??????????????????????????????????????????????????????????????????????????????????????????????????????????????????????????????????????????????????????????????????????????????????????????????????????????????????????????????????????????????????????????????????????????????????????????????????????????????????????????????????????????????????????????????????????????????????????????????????????????????????????????????????????????????????????????????????????????????????????????????????????????????????????????????????????????????????????????????????????????????????????????????????????????????????????????????????????????????????????????????????????????????????????????????????????????????????????????????????????????????????????????????????????????????????????????????

			>Aechmea_gamosepala
			??????????????????????????????????????????????????????????????????????????????????CAGTTAATTCAATAAGAGATAAATGGGAGATAGCACTCGATTTAGTTGGTACCACCCAACCGAATCCGATTCACTCGTTTACTCATTCAATGAGTAAATTTTCAAGTTCAGCCAATCTTTTTTTTTTGAAAAAAAAAATTGCAAGTAGATTAAATCGTGAATAAATGTGTTTCATTTCTATATCATTATATAAAAAAAGATCCATATTATATTACTTATGGAATTCGAACCCGAACTCGATTTATGATTTATTGTTTGGATCTTATTGGCCTTTGTTCTTTATTTTTGATTTTTCATATAGATTTCCGTCTTTATATTATACCCTTTCGTGGATGAATTATGCCTATTTTCACATCTAGGATTTACATATACAACATATATTACTGTCAAGAGGGAATTTTTCTCAGTATTTAGATATTTAGATTCAAAATAAGAAAGGGATCAATTCAATTATAAACCCGCAAAACAAAGATTAGGATTGGGTTGGGTTGCGCTATATATATCAAAGAGTATACAATAATGATGTATTTGGTGAATCAAATACATGGTCTAATAACAAACCATTTTAACATAACATTTTAATGAGTTGATAATATTAATTGAATATTGAATATTTTTTGAAAGATTTTTGTTAAAGGTTTCATTCATGCCTAATCCATATCGAGTAGACCTTGTTGTTGTGAG

			>Aechmea_gracilis
			ATAATTATAATAAAGTGAAATATGTCGAAATCCTTTTTGGAATAATACCAAATCGAAAATAAATGTCCGATAGCAAGTTGATCAGTTAATTCAATAAGAGATAAATGGGAGATAGCACTCGATTTAGTTGGTACCACCCAACCGAATCCGATTCACTCGTTTACTCATTCAATGAGTAAATTTTCAAGTTCAGCCAATCTTTTTTTTTTGAAAAAAAAAATTGCAAGTAGATTAAATCGTGAATAAATGTGTTTCATTTCTATATCATTATATAAAAAAAGATCCATATTATATTACTTATGGAATTCGAACCCGAACTCGATTTATGATTTATTGTTTGGATCTTATTGGCCTTTGTTCTTTATTTTGGATTTTTCATATAGATTTCCGTCTTTATATTATACCCTTTCGTGGATGAATTATGCCTATTTTCACATCTAGGATTTACATATACAACATATATTACTGTCAAGAGGGAATTTTTCTCAGTATTTAGATATTTAGATTCAAAATAAGAAAGGGATCAATTCAATTATAAACCTGCAAAACAAAGATTAGGATTGGGTTGGGTTGCGCTATATATATCAAAGAGTATACAATAATGATGTATTTGGTGAATTAAATACATGGTCTAATAACAAACCATTTTAACATAACATTTTAATGAGTTGATAATATTAATTGAATATTGAATATTTTTTGAAAGATTTTTGTTAAAGGTTTCATTCATGCCTAATCCATATCGAGTAGACCTTGTTGTTGTG??

			>Aechmea_guainumbiorum
			???????????????????????????????????TTTGGAATAATACCAAATCGAAAATAAATGTCCGATAGCAAGTTGATCAGTTAATTCAATAAGAGATAAATGGGAGATAGCACTCGATTTAGTTGGTACCACCCAACCGAATCCGATTCACTCGTTTACTCATTCAATGAGTAAATTTTCAAGTTCAGCCAATC--TTTTTTTTTAAAAAAAAAATTGCAAGTAGATGAAATCGTGAATAAATGTGTTTCATTTCTATATCATTATATAAAAAAAGATCCATATTATATTACTTATGGAATTCGAACCCGAACTCGATTTATGATTTATTATTTGGATCTTATTGGCCTTTGTTCTTTATTTTTGATTTTTCATATAGATTTCCGTCTTTATATTATACCCTTTCGTGGATGAATTATGCCTATTTTCACATCTAGGATTTACATATACAACATATATTACTGTCAAGAGGGAATTTTTCTCAGTATTTAGATATTTAGATTCAAAATAAGAAAGGGATCAATTCAATTATAAACCTGCAAAACAAAGATTAGGATTGGGTTGGGTTGCGCTATATATATCAAAGAGTATACAATAATGATGTATTTGGTGAATCAAATACATGGTCTAATAACAAACCATTTTAACATAACATTTTAATGAGTTGATAATATTAATTGAATATTGAATATTTTTTGAAAGATTTTTGTTAAAGGTTTCATTCATGCCTAATCCATATCGAGTAGACCTTGTTGTTGTG??

			>Aechmea_haltonii
			ATAATTATAATAAAGTGAAATATGTCGAAATCCTTTTTGGAATAATACCAAATCGAAAATAAATGTCCGATAGCAAGTTGATCAGTTAATTCAATAAGAGATAAATGGGAGATAGCACTCGATTTAGTTGGTACCACCCAACCGAATCCGATTCACTCGTTTACTCATTCAATGAGTAAATTTTCAAGTTCAGCCAATC-TTTTTTTTTGAAAAAAAAAATTGCAAGTAGATGAAATCGTGAATAAATGTGTTTCATTTCTATATCATTATATAAAAAAAGATCCATATTATATTACTTATGGAATTCGAACCCGAACTCGATTTATGATTTATTATTTGGATCTTATTGGCCTTTGTTCTTTATTTTTGATTTTTCATATAGATTTCCGTCTTTATATTATACCCTTTCGTGGATGAATTATGCCTATTTTCACATCTAGGATTTACATATACAACATATATTACTGTCAAGAGGGAATTTTTCTCAGTATTTAGATATTTAGATTCAAAATAAGAAAGGGATCAATTCAATTATAAACCTGCAAAACAAAGATTAGGATTGGGTTGGGTTGCGCTATATATATCAAAGAGTATACAATAATGATGTATTTGGTGAATCAAATACATGGTCTAATAACAAACCATTTTAACATAACATTTTAATGAGTTGATAATATTAATTGAATATTGAATATTTTTTGAAAGATTTTTGTTAAAGGTTTCATTCATGCCTAATCCATATCGAGTAGACCTTGTTGTTGTGAG

			>Aechmea_heterosepala
			??????????????????????????????????????????????????????????????????????????????????????????????????????????????????????????????????????????????????????????????????????????????????????????????????????????????????????????????????????????????????????????????????????????????????????????????????????????????????????????????????????????????????????????????????????????????????????????????????????????????????????????????????????????????????????????????????????????????????????????????????????????????????????????????????????????????????????????????????????????????????????????????????????????????????????????????????????????????????????????????????????????????????????????????????????????????????????????????????????????????????????????????????????????????????????????????

			>Aechmea_hoppii
			??????????????????????????????????????????????????????????????????????????????????????????????????????????????????????????????????????????????????????????????????????????????????????????????????????????????????????????????????????????????????????????????????????????????????????????????????????????????????????????????????????????????????????????????????????????????????????????????????????????????????????????????????????????????????????????????????????????????????????????????????????????????????????????????????????????????????????????????????????????????????????????????????????????????????????????????????????????????????????????????????????????????????????????????????????????????????????????????????????????????????????????????????????????????????????????????

			>Aechmea_jungurudoensis
			??????????????????????????????????????????????????????????????????????????????????????????????????????????????????????????????????????????????????????????????????????????????????????????????????????????????????????????????????????????????????????????????????????????????????????????????????????????????????????????????????????????????????????????????????????????????????????????????????????????????????????????????????????????????????????????????????????????????????????????????????????????????????????????????????????????????????????????????????????????????????????????????????????????????????????????????????????????????????????????????????????????????????????????????????????????????????????????????????????????????????????????????????????????????????????????????

			>Aechmea_kertesziae
			??????????????????????????????????????????????????????????????????????????????????CAGTTAATTCCATCAGAGATAAATGGGAGATAGCACTCGATTTAGTTGGTACCACCCAACCGAATCCGATTCAATCGTTTACTCATTCAATGAGTCAATTTTCAAGTTCAGTCAATCCTTTTTTTTTTCAAAAAAAAATTGCAAGTAGATGAAATCGTGAATAAATGTGTTTTATTTCTCTATCATTATAGAAAATCAGATACATATTATATTACTTATGGAATTCGAACCCGAACTCGATTTATGATTCATTATTTCGATCTTATTGGCCTTTGTTCTTTATTTTTGATTTTTCATATAGATTTCCGTCTTTATATTATACCCTTTCGTGGATGAATTATGCTTATTTTCACATCTAGGATTTACATATACAACATATATGACTGTCAAGAGGGAATTTTTCTCAGTATTTAGATATTTAGATTCAAAATAAGAAAGGGATCAATTCAATTATAAACCTGCAAAGCAAAGATTAGGATTGGGTTGGGTTGCGCTATATATATCAAAGAGTATACAATAATGATGTATTTGGTGAATCAAATACATGGTCTAATAACAAACCATTTTAACATAACATTTTAATGAGTTGATAATATTAATTGAATATTGAATATTTTTTGAAAGATTTTTGTTAAAGGTTTGATTCATGCTTAATCCATATCGAGTAGACCTTGTTGTTGTGAG

			>Aechmea_kleinii
			??????????????????????????????????????????????????????????????????????????????????????????????????????????????????????????????????????????????????????????????????????????????????????????????????????????????????????????????????????????????????????????????????????????????????????????????????????????????????????????????????????????????????????????????????????????????????????????????????????????????????????????????????????????????????????????????????????????????????????????????????????????????????????????????????????????????????????????????????????????????????????????????????????????????????????????????????????????????????????????????????????????????????????????????????????????????????????????????????????????????????????????????????????????????????????????????

			>Aechmea_lamarchei
			ATAATTATAATAAAGTGAAATATGTCGAAATCCTTTTTGGAATAATACCAAATCGAAAATAAATGTCCGATAGCAAGTTGATCAGTTAATTCAATAAGAGATAAATGGGAGATAGCACTCGATTTAGTTGGTACCACCCAACCGAATCCGATTCACTCGTTTACTCATTGAATGAGTAAATTTTCAAGTTCAGCCAATC--TTTTTTTTTAAAAAAAAAATTGCAAGTAGATTAAATCATGAATAAATGTGTTTCATTTCTATATCATTATATAAAAAAAGATCCATATTATATTACTTATGGAATTCGAACCCGAACTCGATTTATGATTTATTATTTGGATCTTATTGGCCTTTGTTCTTTATTTTTGATTTTTCATATAGATTTCCGTCTTTATATTATACCCTTTCGTGGATGAATTATGCCTATTTTCACATCTAGGATTTACATATACAACATATATTACTGTCAAGAGGGAATTTTTCTCAGTATTTAGATATTTAGATTCAAAATAAGAAAGGGATCAATTCAATTATAAACCTGCAAAACAAAGATTAGGATTGGGTTGGGTTGCGCTATATATATCAAAGAGTATACAATAATGATGTATTTGGTGAATCAAATACATGGTCTAATAACAAACCATTTTAACATAACATTTTAATGAGTTGATAATATTAATTGAATATTGAATATTTTTTGAAAGATTTTTGTTAAAGGTTTCATTCATGCCTAATCCATATCGAGTAGACCTTGTTGTTGTG??

			>Aechmea_leptantha
			ATAATTATAATAAAGTGAAATATGTCGAAATCCTTTTTGGAATAATACCAAATCGAAAATAAATGTCCGATAGCAAGTTGATCAGTTAATTCAATAAGAGATAAATGGGAGATAGCACTCGATTTAGTTGGTACCACCCAACCGAATCCGATTCACTCGTTTACTCATTCAATGAGTAAATTTTCAAGTTCAGCCAATC-TTTTTTTTTTAAAAAAAAAATTGCAAGTAGATGAAATCGTGAATAAATGTGTTTCATTTCTATATCATTATATAAAAAAAGATCCATATTTTATTACTTATGGAATTCGAACCCGAACTCGATTTATGATTTATTATTTGGATCTTATTGGCCTTTGTTCTTTATTTTTGATTTTTCATATAGATTTCCGTCTTTATATTATACCCTTTCGTGGATGAATTATGCCTATTTTCACATCTAGGATTTACATATACAACATATATTACTGTCAAGAGGGAATTTTTCTCAGTATTTAGATATTTAGATTCAAAATAAGAAAGGGATCAATTCAATTATAAACCTGCAAAACAAAGATTAGGATTGGGTTGGGTTGCGCTATATATATCAAAGAGTATACAATAATGATGTATTTGGTGAATCAAATACATGGTCTAATAACAAACCATTTTAACATAACATTTTAACGAGTTGATAATATTAATTGAATATTGAATATTTTTTGAAAGATTTTTGTTAAAGGTTTCATTCATGCCTAATCCATATCGAGTAGACCTTGTTATTGTG??

			>Aechmea_leucolepis
			??????????????????????????????????????????????????????????????????????????????????????????????????????????????????????????????????????????????????????????????????????????????????????????????????????????????????????????????????????????????????????????????????????????????????????????????????????????????????????????????????????????????????????????????????????????????????????????????????????????????????????????????????????????????????????????????????????????????????????????????????????????????????????????????????????????????????????????????????????????????????????????????????????????????????????????????????????????????????????????????????????????????????????????????????????????????????????????????????????????????????????????????????????????????????????????????

			>Aechmea_lilacinantha
			??????????????????????????????????????????????????????????????????????????????????????????????????????????????????????????????????????????????????????????????????????????????????????????????????????????????????????????????????????????????????????????????????????????????????????????????????????????????????????????????????????????????????????????????????????????????????????????????????????????????????????????????????????????????????????????????????????????????????????????????????????????????????????????????????????????????????????????????????????????????????????????????????????????????????????????????????????????????????????????????????????????????????????????????????????????????????????????????????????????????????????????????????????????????????????????????

			>Aechmea_longicuspis_var._kuntzeana
			??????????????????????????????????????????????????????????????????????????????????????????????????????????????????????????????????????????????????????????????????????????????????????????????????????????????????????????????????????????????????????????????????????????????????????????????????????????????????????????????????????????????????????????????????????????????????????????????????????????????????????????????????????????????????????????????????????????????????????????????????????????????????????????????????????????????????????????????????????????????????????????????????????????????????????????????????????????????????????????????????????????????????????????????????????????????????????????????????????????????????????????????????????????????????????????????

			>Aechmea_lueddemanniana
			ATAATTATAATAAAGTGAAATATGTCGAAATCCTTTTTGGAATAATACCAAATCGAAAATAAATGTCCGATAGCAAGTTGATCAGTTAATTCAATAAGAGATAAATGGGAGATAGCACTCGATTTAGTTGGTACCACCCAACCGAATCCGATTCACTCGTTTACTCATTGAATGAGTAAATTTTCAAGTTCAGCCAATCTTTTTTTTTGAAAAAAAAAAATTGCAAGTAGATGAAATCGTGAATAAATGTGTTTCATTTCTATATCATTATATAAAAAAAGATCCATATTATATTACTTATGGAATTCGAACCCGAACTCGATTTATGATTTATTATTTGGATCTTATTGGCCTTTGTTCTTTATTTTTGATTTTTCATATAGATTTCCGTCTTTATATTATACCCTTTCGTGGATGAATTATGCCTATTTTCACATCTAGGATTTACATATACAACATATATTACTGTCAAGAGGGAATTTTTCTCAGTATTTAGATATTTAGATTCAAAATAAGAAAGGGATCAATTCAATTATAAACCTGCAAAACAAAGATTAGGATTGGGTTGGGTTGCGCTATATATATCAAAGAGTATACAATAATGATGTATTTGGTGAATCAAATACATGGTCTAATAACAAACCATTTTAACATAACATTTTAATGAGTTGATAATATTAATTGAATATTGAATATTTTTTGAAAGATTTTTGTTAAAGGTTTCATTCATGCCTAATCCATATCGAGTAGACCTTGTTGTTGTG??

			>Aechmea_maasii
			??????????????????????????????????????????????????????????????????????????????????????????????????????????????????????????????????????????????????????????????????????????????????????????????????????????????????????????????????????????????????????????????????????????????????????????????????????????????????????????????????????????????????????????????????????????????????????????????????????????????????????????????????????????????????????????????????????????????????????????????????????????????????????????????????????????????????????????????????????????????????????????????????????????????????????????????????????????????????????????????????????????????????????????????????????????????????????????????????????????????????????????????????????????????????????????????

			>Aechmea_magdalenae
			??????????????????????????????????????????????????????????????????????????????????????????????????????????????????????????????????????????????????????????????????????????????????????????????????????????????????????????????????????????????????????????????????????????????????????????????????????????????????????????????????????????????????????????????????????????????????????????????????????????????????????????????????????????????????????????????????????????????????????????????????????????????????????????????????????????????????????????????????????????????????????????????????????????????????????????????????????????????????????????????????????????????????????????????????????????????????????????????????????????????????????????????????????????????????????????????

			>Aechmea_manzanaresiana
			??????????????????????????????????????????????????????????????????????????????????????????????????????????????????????????????????????????????????????????????????????????????????????????????????????????????????????????????????????????????????????????????????????????????????????????????????????????????????????????????????????????????????????????????????????????????????????????????????????????????????????????????????????????????????????????????????????????????????????????????????????????????????????????????????????????????????????????????????????????????????????????????????????????????????????????????????????????????????????????????????????????????????????????????????????????????????????????????????????????????????????????????????????????????????????????????

			>Aechmea_marauensis
			??????????????????????????????????????????????????????????????????????????????????????????????????????????????????????????????????????????????????????????????????????????????????????????????????????????????????????????????????????????????????????????????????????????????????????????????????????????????????????????????????????????????????????????????????????????????????????????????????????????????????????????????????????????????????????????????????????????????????????????????????????????????????????????????????????????????????????????????????????????????????????????????????????????????????????????????????????????????????????????????????????????????????????????????????????????????????????????????????????????????????????????????????????????????????????????????

			>Aechmea_marginalis
			??????????????????????????????????????????????????????????????????????????????????????????????????????????????????????????????????????????????????????????????????????????????????????????????????????????????????????????????????????????????????????????????????????????????????????????????????????????????????????????????????????????????????????????????????????????????????????????????????????????????????????????????????????????????????????????????????????????????????????????????????????????????????????????????????????????????????????????????????????????????????????????????????????????????????????????????????????????????????????????????????????????????????????????????????????????????????????????????????????????????????????????????????????????????????????????????

			>Aechmea_mariae-reginae
			??????????????????????????????????????????????????????????????????????????????????????????????????????????????????????????????????????????????????????????????????????????????????????????????????????????????????????????????????????????????????????????????????????????????????????????????????????????????????????????????????????????????????????????????????????????????????????????????????????????????????????????????????????????????????????????????????????????????????????????????????????????????????????????????????????????????????????????????????????????????????????????????????????????????????????????????????????????????????????????????????????????????????????????????????????????????????????????????????????????????????????????????????????????????????????????????

			>Aechmea_melinonii
			??????????????????????????????????????????????????????????????????????????????????????????????????????????????????????????????????????????????????????????????????????????????????????????????????????????????????????????????????????????????????????????????????????????????????????????????????????????????????????????????????????????????????????????????????????????????????????????????????????????????????????????????????????????????????????????????????????????????????????????????????????????????????????????????????????????????????????????????????????????????????????????????????????????????????????????????????????????????????????????????????????????????????????????????????????????????????????????????????????????????????????????????????????????????????????????????

			>Aechmea_mertensii
			ATAATTATAATAAAGTGAAATATGTCGAAATCCTTTTTGGAATAATACCAAATC-AAAATAAATGTCCGATAGCAAGTTGATCAGTTAATTCAATAAGAGATAAATGGGAGATAGCACTCGATTTAGTTGGTACCACCCAACCGAATCCGATTCACCCGTTTACTCATTCAATGAGTAAATTTTCAAGTTCAGCCAATCTTTTTTTTTTTAAAAAAAAGATTTCAAGTAGATGAAATCGTGAATAAATGTGTTTCATTTCTATATCATTATATAAAAAAAGATCCATATTATATTACTTATGGAATTCAAACCCGAACTCGATTTATGATTTATTATTTGGATCTTATTGGCCTTTGTTCTTTATTTTTTATTTTTCATATAGATTTCCGTCTTTATATTATACCCTTTCGTGGATGAATTATGCCTATTTTCACATCTAGGATTTACATATACAACATATATTACTGTCAAGAGGGAATTGTTCTCAGTATTTAGATATTTAGATTCAAAATAAGAAAGGGATCAATTCAATTATAAACCTGCAAAACAAAGATTAGGATTGGGTTGGGTTGCGCTATATATATCAAAGAGTATACAATAATGATGTATTTGGTGAATCAAATACATGGTCTAATAACAAACCATTTTAACATAACATTTTAATGAGTTGATAATATTAATTGAATATTGAATATTTTTTGAAAGATTTTTGTTAAAGGTTTCATTCATGCCTAATCCATATCGAGTAGACCTTGTTGTTGTG??

			>Aechmea_mexicana
			ATAATTATAATAAAGTGAAATATGTCGAAATCCTTTTTGGAATAATACCAAATCGAAAATAAATGTCCGATAGCAAGTTGATCAGTTAATTCAATAAGAGATAAATGGGAGATAGCACTCGATTTAGTTGGTACCACCCAACCGAATCCGATTCACTCGTTTACTCATTGAATGAGTAAATTTTCAAGTTCAGCCAATCTTTTTTTTTTCAAAAAAAAAATTGCAAGTAGATGAAATCGTGAATAAATGTGTTTCATTTCTATATCATTATATAAAAAAAGATCCATATTATATTACTTATGGAATTCGAACCCGAACTCGATTTATGATTTATTATTTGGATCTTATTGGCCTTTGTTCTTTATTTTTGATTTTTCATATAGATTTCCGTCTTTATATTATACCCTTTCGTGGATGAATTATGCCTATTTTCACATCTAGGATTTACATATACAACATATATTACTGTCAAGAGGGAATTTTTCTCAGTATTTAGATATTTAGATTCAAAATAAGAAAGGGATCAATTCAATTATAAACCTGCAAAACAAAGATTAGGATTGGGTTGGGTTGCGCTATATATATCAAAGAGTATACAATAATGATGTATTTGGTGAATCAAATACATGGTCTAATAACAAACCATTTTAACATAACATTTTAATGAGTTGATAATATTAATTGAATATTGAATATTTTTTGAAAGATTTTTGTTAAAGGTTTCATTCATGCCTAATCCATATCGAGTAGACCTTGTTGTTGTG??

			>Aechmea_microcephala
			??????????????????????????????????????????????????????????????????????????????????????????????????????????????????????????????????????????????????????????????????????????????????????????????????????????????????????????????????????????????????????????????????????????????????????????????????????????????????????????????????????????????????????????????????????????????????????????????????????????????????????????????????????????????????????????????????????????????????????????????????????????????????????????????????????????????????????????????????????????????????????????????????????????????????????????????????????????????????????????????????????????????????????????????????????????????????????????????????????????????????????????????????????????????????????????????

			>Aechmea_miniata
			??????????????????????????????????????????????????????????????????????????????????????????????????????????????????????????????????????????????????????????????????????????????????????????????????????????????????????????????????????????????????????????????????????????????????????????????????????????????????????????????????????????????????????????????????????????????????????????????????????????????????????????????????????????????????????????????????????????????????????????????????????????????????????????????????????????????????????????????????????????????????????????????????????????????????????????????????????????????????????????????????????????????????????????????????????????????????????????????????????????????????????????????????????????????????????????????

			>Aechmea_miniata_var._discolor
			??????????????????????????????????????????????????????????????????????????????????????????????????????????????????????????????????????????????????????????????????????????????????????????????????????????????????????????????????????????????????????????????????????????????????????????????????????????????????????????????????????????????????????????????????????????????????????????????????????????????????????????????????????????????????????????????????????????????????????????????????????????????????????????????????????????????????????????????????????????????????????????????????????????????????????????????????????????????????????????????????????????????????????????????????????????????????????????????????????????????????????????????????????????????????????????????

			>Aechmea_mira
			??????????????????????????????????????????????????????????????????????????????????????????????????????????????????????????????????????????????????????????????????????????????????????????????????????????????????????????????????????????????????????????????????????????????????????????????????????????????????????????????????????????????????????????????????????????????????????????????????????????????????????????????????????????????????????????????????????????????????????????????????????????????????????????????????????????????????????????????????????????????????????????????????????????????????????????????????????????????????????????????????????????????????????????????????????????????????????????????????????????????????????????????????????????????????????????????

			>Aechmea_moorei
			??????????????????????????????????????????????????????????????????????????????????????????????????????????????????????????????????????????????????????????????????????????????????????????????????????????????????????????????????????????????????????????????????????????????????????????????????????????????????????????????????????????????????????????????????????????????????????????????????????????????????????????????????????????????????????????????????????????????????????????????????????????????????????????????????????????????????????????????????????????????????????????????????????????????????????????????????????????????????????????????????????????????????????????????????????????????????????????????????????????????????????????????????????????????????????????????

			>Aechmea_murcae
			??????????????????????????????????????????????????????????????????????????????????????????????????????????????????????????????????????????????????????????????????????????????????????????????????????????????????????????????????????????????????????????????????????????????????????????????????????????????????????????????????????????????????????????????????????????????????????????????????????????????????????????????????????????????????????????????????????????????????????????????????????????????????????????????????????????????????????????????????????????????????????????????????????????????????????????????????????????????????????????????????????????????????????????????????????????????????????????????????????????????????????????????????????????????????????????????

			>Aechmea_muricata
			??????????????????????????????????????????????????????????????????????????????????????????????????????????????????????????????????????????????????????????????????????????????????????????????????????????????????????????????????????????????????????????????????????????????????????????????????????????????????????????????????????????????????????????????????????????????????????????????????????????????????????????????????????????????????????????????????????????????????????????????????????????????????????????????????????????????????????????????????????????????????????????????????????????????????????????????????????????????????????????????????????????????????????????????????????????????????????????????????????????????????????????????????????????????????????????????

			>Aechmea_mutica
			??????????????????????????????????????????????????????????????????????????????????????????????????????????????????????????????????????????????????????????????????????????????????????????????????????????????????????????????????????????????????????????????????????????????????????????????????????????????????????????????????????????????????????????????????????????????????????????????????????????????????????????????????????????????????????????????????????????????????????????????????????????????????????????????????????????????????????????????????????????????????????????????????????????????????????????????????????????????????????????????????????????????????????????????????????????????????????????????????????????????????????????????????????????????????????????????

			>Aechmea_nallyi
			??????????????????????????????????????????????????????????????????????????????????????????????????????????????????????????????????????????????????????????????????????????????????????????????????????????????????????????????????????????????????????????????????????????????????????????????????????????????????????????????????????????????????????????????????????????????????????????????????????????????????????????????????????????????????????????????????????????????????????????????????????????????????????????????????????????????????????????????????????????????????????????????????????????????????????????????????????????????????????????????????????????????????????????????????????????????????????????????????????????????????????????????????????????????????????????????

			>Aechmea_napoensis
			??????????????????????????????????????????????????????????????????????????????????????????????????????????????????????????????????????????????????????????????????????????????????????????????????????????????????????????????????????????????????????????????????????????????????????????????????????????????????????????????????????????????????????????????????????????????????????????????????????????????????????????????????????????????????????????????????????????????????????????????????????????????????????????????????????????????????????????????????????????????????????????????????????????????????????????????????????????????????????????????????????????????????????????????????????????????????????????????????????????????????????????????????????????????????????????????

			>Aechmea_nidularioides
			??????????????????????????????????????????????????????????????????????????????????????????????????????????????????????????????????????????????????????????????????????????????????????????????????????????????????????????????????????????????????????????????????????????????????????????????????????????????????????????????????????????????????????????????????????????????????????????????????????????????????????????????????????????????????????????????????????????????????????????????????????????????????????????????????????????????????????????????????????????????????????????????????????????????????????????????????????????????????????????????????????????????????????????????????????????????????????????????????????????????????????????????????????????????????????????????

			>Aechmea_nudicaulis
			ATAATTATAATAAAGTGAAATATGTCGAAATCCTTTTTGGAATAATACCAAATC-AAAATAAATGTCCGATAGCAAGTTGATCAGTTAATTCAATAAGAGATAAATGGGAGATAGCACTCGATTTAGTTGGTACCACCCAACCAAATCCGATTCACTCGTTTACTCATTGAATGAGTAAATTTTCAAGTTCAGCCAATCTTTTTTTTTTTAAAAAAAAAATTGCAAGTAGATGAAATCGTGAATAAATGTGTTTCATTTCTATATCATTATATAAAAAAGGATCCATATTATATTACTTATGGAATTCGAACCCGAACTCGATTTATGATTTATTATTTGGATCTTATTGGCCTTTGTTCTTTATTTTTGATTTTTCATATAGATTTCCGTCTTTATATTATACCCTTTCGTGGATGAATTATGCCTATTTTCACATCTAGGATTTACATATACAACATATATTACTGTCAAGAGGGAATTTTTCTCAGTATTTAGATATTTAGATTCAAAATAAGAAAGGGATCAATTCAATTATAAACCTGCAAAACAAAGATTAGGATTGGGTTGGGTTGCGCTATATATATCAAAGAGTATACAATAATGATGTATTTGGTGAATCAAATACATGGTCTAATAACAAACCATTTTAACATAACATTTTAATGAGTTGATAATATTAATTGAATATTTAATATTTTTTGAAAGATTTTTGTTAAAGGTTTCATTCATGCCTAATCCATATCGAGTAGACCTTGTTGTTGTG??

			>Aechmea_organensis
			ATAATTATAATAAAGTGAAATATGTCGAAATCCTTTTTGGAATAATACCAAATCGAAAATAAATGTCCGATAGCAAGTTGATCAGTTAATTCAATAAGAGATAAATGGGAGATAGCACTCGATTTAGTTGGTACCACCCAACCGAATCCGATTCACTCGTTTACTCATTCAATGAGTAAATTTTCAAGTTCAGCCAATC-TTTTTTTTTGAAAAAAAAAATTGCAAGTAGATTAAATCGTGAATAAATGTGTTTCATTTCTATATCATTATATAAAAAAAGATCCATATTATATTACTTATGGAATTCGAACCCGAACTCGATTTATGATTTATTGTTTGGATCTTATTGGCCTTTGTTCTTTATTTTGGATTTTTCATATAGATTTCCGTCTTTATATTATACCCTTTCGTGGATGAATTATGCCTATTTTCACATCTAGGATTTACATATACAACATATATTACTGTCAAGAGGGAATTTCTCTCAGTATTTAGATATTTAGATTCAAAATAAGAAAGGGATCAATTCAATTATAAACCTGCAAAACAAAGATTAGGATTGGGTTGGGTTGCGCTATATATATCAAAGAGTATACAATAATGATGTATTTGGTGAATTAAATACATGGTCTAATAACAAACCATTTTAACATAACATTTTAATGAGTTGATAATATTAATTGAATATTGAATATTTTTTGAAAGATTTTTGTTAAAGGTTTCATTCATGCCTAATCCATATCGAGTAGACCTTGTTGTTGTGAG

			>Aechmea_orlandiana
			??????????????????????????????????????????????????????????????????????????????????CAGTTAATTCAATAAGAGATAAATGGGAGATAGCACTCGATTTAGTTGGTACCACCCAACCGAATCCGATTCACTCGTTTACTCATTCAATGAGTAAATTTTCAAGTTCAGCCAATCTTTTTTTTTTTAAAAAAAAGATTGCAAGTAGATGAAATCGTGAATAAATGTGTTTCATTTCTATATCATTATATAAAAAAAGATCCATATTATATTACTTATGGAATTCGAACCCGAACTCGATTTATGATTTATTATTTGGATCTTATTGGCCTTTGCTCTTTATTTTTGATTTTTCATATAGATTTCCATCTTTATATTATACCCTTTCGTGTATGAATTATGCCTATTTTCACATCTAGGATTTACATATACAACATATATTACTGTCAAGAGGGAATTTTTCTCAGTATTTAGATATTTAGATTCAAAATAAGAAAGGGATCAATTCAATTATAAACCTGCAAAACAAAGATTAGGATTGGGTTGGGTTGCGCTATATATATCAAAGAGTATACAATAATGATGTATTTGGTGAATCAAATACATGGTCTAATAACAAACCATTTTAACATAACATTTTAATGAGTTGATAATATTAATTGAATATTGAATATTTTTTGAAAGATTTTTGTTAAAGGTTTCATTCATGCCTAATCCATATCGAGTAGACCTTGTTGTTGTGAG

			>Aechmea_ornata
			??????????????????????????????????????????????????????????????????????????????????????????????????????????????????????????????????????????????????????????????????????????????????????????????????????????????????????????????????????????????????????????????????????????????????????????????????????????????????????????????????????????????????????????????????????????????????????????????????????????????????????????????????????????????????????????????????????????????????????????????????????????????????????????????????????????????????????????????????????????????????????????????????????????????????????????????????????????????????????????????????????????????????????????????????????????????????????????????????????????????????????????????????????????????????????????????

			>Aechmea_paniculata
			???????????????????????????????????TTTGGAATAATACCAAATC-AAAATAAATGTCCGATAGCAAGTTGATCAGTTAATTCAATAAGAGATAAATGGGAGATAGCACTCGATTTAGTTGGTACCACCCAACCGA------------------------CAATGAGTAAATTTTCAAGTTCAGCCAATC--TTTTTTTTTTAAAAAAAAATTGCAAGTAGATGAAATCGTGAATAAATGTGTTTCATTTCTATATCATTATATAAAAAAAGATCCATATTATATTACTTATGGAATTCGAACCCGAACTCGATTTATGATTTATTATTTGGATCTTATTGGCCTTTGTTCTTTATTTTTGATTTTTCATATAGATTTCCGTCTTTATATTATACCCTTTCGTGGATGAATTATGCCTATTTTCACATCTAGGATTTACATATACAACATATATTACTGTCAAGAGGGAATTTTTCTCAGTATTTAGATATTTAGATTCAAAATAAGAAAGGGATCAATTCAATTATAAACCTGCAAAACAAAGATTAGGATTGGGTTGGGTTGCGCTATATATATCAAAGAGTATACAATAATGATGTATTTGGTGAATCAAATACATGGTCTAATAACAAACCATTTTAACATAACATTTTAATGAGTTGATAATATTAATTGAATATTGAATATTTTTTGAAAGATTTTTGTTAAAGGTTTCATTCATGCCTAATCCATATCGAGTAGACCTTGTTGTTGTG??

			>Aechmea_paratiensis
			??????????????????????????????????????????????????????????????????????????????????????????????????????????????????????????????????????????????????????????????????????????????????????????????????????????????????????????????????????????????????????????????????????????????????????????????????????????????????????????????????????????????????????????????????????????????????????????????????????????????????????????????????????????????????????????????????????????????????????????????????????????????????????????????????????????????????????????????????????????????????????????????????????????????????????????????????????????????????????????????????????????????????????????????????????????????????????????????????????????????????????????????????????????????????????????????

			>Aechmea_patriciae
			??????????????????????????????????????????????????????????????????????????????????????????????????????????????????????????????????????????????????????????????????????????????????????????????????????????????????????????????????????????????????????????????????????????????????????????????????????????????????????????????????????????????????????????????????????????????????????????????????????????????????????????????????????????????????????????????????????????????????????????????????????????????????????????????????????????????????????????????????????????????????????????????????????????????????????????????????????????????????????????????????????????????????????????????????????????????????????????????????????????????????????????????????????????????????????????????

			>Aechmea_pectinata
			??????????????????????????????????????????????????????????????????????????????????????????????????????????????????????????????????????????????????????????????????????????????????????????????????????????????????????????????????????????????????????????????????????????????????????????????????????????????????????????????????????????????????????????????????????????????????????????????????????????????????????????????????????????????????????????????????????????????????????????????????????????????????????????????????????????????????????????????????????????????????????????????????????????????????????????????????????????????????????????????????????????????????????????????????????????????????????????????????????????????????????????????????????????????????????????????

			>Aechmea_pedicellata
			??????????????????????????????????????????????????????????????????????????????????????????????????????????????????????????????????????????????????????????????????????????????????????????????????????????????????????????????????????????????????????????????????????????????????????????????????????????????????????????????????????????????????????????????????????????????????????????????????????????????????????????????????????????????????????????????????????????????????????????????????????????????????????????????????????????????????????????????????????????????????????????????????????????????????????????????????????????????????????????????????????????????????????????????????????????????????????????????????????????????????????????????????????????????????????????????

			>Aechmea_penduliflora
			??????????????????????????????????????????????????????????????????????????????????????????????????????????????????????????????????????????????????????????????????????????????????????????????????????????????????????????????????????????????????????????????????????????????????????????????????????????????????????????????????????????????????????????????????????????????????????????????????????????????????????????????????????????????????????????????????????????????????????????????????????????????????????????????????????????????????????????????????????????????????????????????????????????????????????????????????????????????????????????????????????????????????????????????????????????????????????????????????????????????????????????????????????????????????????????????

			>Aechmea_perforata
			??????????????????????????????????????????????????????????????????????????????????????????????????????????????????????????????????????????????????????????????????????????????????????????????????????????????????????????????????????????????????????????????????????????????????????????????????????????????????????????????????????????????????????????????????????????????????????????????????????????????????????????????????????????????????????????????????????????????????????????????????????????????????????????????????????????????????????????????????????????????????????????????????????????????????????????????????????????????????????????????????????????????????????????????????????????????????????????????????????????????????????????????????????????????????????????????

			>Aechmea_phanerophlebia
			??????????????????????????????????????????????????????????????????????????????????????????????????????????????????????????????????????????????????????????????????????????????????????????????????????????????????????????????????????????????????????????????????????????????????????????????????????????????????????????????????????????????????????????????????????????????????????????????????????????????????????????????????????????????????????????????????????????????????????????????????????????????????????????????????????????????????????????????????????????????????????????????????????????????????????????????????????????????????????????????????????????????????????????????????????????????????????????????????????????????????????????????????????????????????????????????

			>Aechmea_pimenti-velosoi
			ATAATTATAATAAAGTGAAATATGTCGAAATCCTTTTTGGAATAATACCAAATCGAAAATAAATGTCCGATAGCAAGTTGATCAGTTAATTCAATAAGAGATAAATGGGAGATAGCACTCGATTTAGTTGGTACCACCCAACCGAATCCGATTCACTCGTTTACTCATTCAATGAGTAAATTTTCAAGTTCAGCCAATCTTTTTTTTT--AAAAAAAAAATTGCAAGTAGATGAAATCGTGAATAAATGTGTTTCATTTCTATATCATTATATAAAAAAAGATCCATATTATATTACTTATGGAATTCGAACCCGAACTCGATTTATGATTTATTATTTGGATCTTATTGGCCTTTATTCTTTATTTTTGATTTTTCATATGGATTTCCGTCTTTATATTATACCCTTTCGTGGATGAATTATGCCTATTTTCACATCTAGGATTTACATATACAACATATATTACCCTCAAGAGGGAATTTTTCTCAGTATTTAGATATTTAGATTCAAAATAAGAAAGGGATCAATTCAATTATAAACCTGCAAAACAAAGATTAGGATTGGGTTGGGTTGCGCTATATATATCAAAGAGTATACAATAATGATGTATTTGGTGAATCAAATACATGGTCTAATAACAAACCATTTTAACATAACATTTTAATGAGTTGATAATATTAATTGAATATTGAATATTTTTTGAAAGATTTTTGTTAAAGGTTTCATTCATGCCTAATCCATATCGAGTAGACCTTGTTGTTGTG??

			>Aechmea_pineliana
			??????????????????????????????????????????????????????????????????????????????????????????????????????????????????????????????????????????????????????????????????????????????????????????????????????????????????????????????????????????????????????????????????????????????????????????????????????????????????????????????????????????????????????????????????????????????????????????????????????????????????????????????????????????????????????????????????????????????????????????????????????????????????????????????????????????????????????????????????????????????????????????????????????????????????????????????????????????????????????????????????????????????????????????????????????????????????????????????????????????????????????????????????????????????????????????????

			>Aechmea_polyantha
			??????????????????????????????????????????????????????????????????????????????????????????????????????????????????????????????????????????????????????????????????????????????????????????????????????????????????????????????????????????????????????????????????????????????????????????????????????????????????????????????????????????????????????????????????????????????????????????????????????????????????????????????????????????????????????????????????????????????????????????????????????????????????????????????????????????????????????????????????????????????????????????????????????????????????????????????????????????????????????????????????????????????????????????????????????????????????????????????????????????????????????????????????????????????????????????????

			>Aechmea_pubescens
			??????????????????????????????????????????????????????????????????????????????????????????????????????????????????????????????????????????????????????????????????????????????????????????????????????????????????????????????????????????????????????????????????????????????????????????????????????????????????????????????????????????????????????????????????????????????????????????????????????????????????????????????????????????????????????????????????????????????????????????????????????????????????????????????????????????????????????????????????????????????????????????????????????????????????????????????????????????????????????????????????????????????????????????????????????????????????????????????????????????????????????????????????????????????????????????????

			>Aechmea_racinae
			ATAATTATAATAAAGTGAAATATGTCGAAATCCTTTTTGGAATAATACCAAATCGAAAATAAATGTCCGATAGCAAGTTGATCAGTTAATTCAATAAGAGATAAATGGGAGATAGCACTCGATTTAGTTGGTACCACCCAACCGAATCCGATTCACTCGTTTACTCATTGAATGAGTAAATTTTCAAGTTCAGCCAATC-TTTTTTTTTGAAAAAAAAATTGGCAAGTAGATKAAATCGTGAATAAATGTGTTTCATTTCTATATCATTATATAAAAAAAGATCCATATTATATTACTTATGGAATTCGAACCCGAACTCGATTTATGATTTATTGTTTGGATCTTATTGGCCTTTGTTCTTTATTTTTGATTTTTCATATAGATTTCCGTCTTTATATTATACCCTTTCGTGGATGAATTATGCCTATTTTCACATCTAGGATTTACATATACAACATATATTACTGTCAAGAGGGAATTTTTCTCAGTATTTAGATATTTAGATTCAAAATAAGAAAGGGATCAATTCAATTATAAACCTGCAAAACAAAGATTAGGATTGGGTTGGGTTGCGCTATATATATCAAAGAGTATACAATAATGATGTATTTGGTGAATCAAATACATGGTCTAATAACAAACCATTTTAACATAACATTTTAATGAGTTGATAATATTAATTGAATATTGAATATTTTTTGAAAGATTTTTGTTAAAGGTTTCATTCATGCCTAATCCATATCGAGTAGACCTTGTTGTTGTG??

			>Aechmea_ramosa
			??????????????????????????????????????????????????????????????????????????????????????????????????????????????????????????????????????????????????????????????????????????????????????????????????????????????????????????????????????????????????????????????????????????????????????????????????????????????????????????????????????????????????????????????????????????????????????????????????????????????????????????????????????????????????????????????????????????????????????????????????????????????????????????????????????????????????????????????????????????????????????????????????????????????????????????????????????????????????????????????????????????????????????????????????????????????????????????????????????????????????????????????????????????????????????????????

			>Aechmea_recurvata
			???????????????????????????????????TTTGGAATAATACCAAATCGAAAATAAATGTCCGATAGCAAGTTGATCAGTTAATTCAATAAGAGATAAATGGGAGATAGCACTCTATTTAGTTGGTACCACCCAACCGAATCCGATTCACTCGTTTACTCATTCAATGAGTAAATTTTCAAGTTCAGCCAATC-TTTTTTTTTTAAAAAAAAAATTGCAAGTAGATGAAATCGTGAATAAATGTGTTTCATTTCTATATCATTATATAAAAAAAGATCCATATTATATTACTTATGGAATTCGAACCCGAACTCGATTTATGATTTATTGTTTGGATCTTATTGGCCTTTGTTCTTTATTTTTGATTTTTCATATGGATTTCCGTCTTTATATTATACCCTTTCGTGGATGAATTATGCCTATTTTCACATCTAGGATTTACATATACAACATATATTACTGTCAAGAGGGAATTTTTCTCAGTATTTAGATATTTAGATTCAAAATAAGAAAGGGATCAATTCAATTATAAACCTGCAAAACAAAGATTAGGATTGGGTTGGGTTGCGCTATATATATCAAAGAGTATACAATAATGATGTATTTGGTGAATCAAATACATGGTCTAATAACAAACCATTTTAACATAACATTTTAATGAGTTGATAATATTAATTGAATA-------TTTTTTGAAAGATTTTTGTTAAAGGTTTCATTCATGCCTAATCCACATCGAGTAGACCTTGTTGTTGTG??

			>Aechmea_roberto-seidelii
			??????????????????????????????????????????????????????????????????????????????????????????????????????????????????????????????????????????????????????????????????????????????????????????????????????????????????????????????????????????????????????????????????????????????????????????????????????????????????????????????????????????????????????????????????????????????????????????????????????????????????????????????????????????????????????????????????????????????????????????????????????????????????????????????????????????????????????????????????????????????????????????????????????????????????????????????????????????????????????????????????????????????????????????????????????????????????????????????????????????????????????????????????????????????????????????????

			>Aechmea_rodriguesiana
			??????????????????????????????????????????????????????????????????????????????????????????????????????????????????????????????????????????????????????????????????????????????????????????????????????????????????????????????????????????????????????????????????????????????????????????????????????????????????????????????????????????????????????????????????????????????????????????????????????????????????????????????????????????????????????????????????????????????????????????????????????????????????????????????????????????????????????????????????????????????????????????????????????????????????????????????????????????????????????????????????????????????????????????????????????????????????????????????????????????????????????????????????????????????????????????????

			>Aechmea_roeseliae
			??????????????????????????????????????????????????????????????????????????????????????????????????????????????????????????????????????????????????????????????????????????????????????????????????????????????????????????????????????????????????????????????????????????????????????????????????????????????????????????????????????????????????????????????????????????????????????????????????????????????????????????????????????????????????????????????????????????????????????????????????????????????????????????????????????????????????????????????????????????????????????????????????????????????????????????????????????????????????????????????????????????????????????????????????????????????????????????????????????????????????????????????????????????????????????????????

			>Aechmea_romeroi
			????????????????????????????????????TTGGAATAATACCAAATTGAAAATAAATGTCCGATAGCAAGTTGATCAGTTAATTCAATAAGAGATAAATGGGAGATAGCACTCGATTTAGTTGGTACCACCCAACCGAATCCGATTCACCCGTTTACTCATTCAATGAGTAAATTTTCAAGTTCAGCCAATCTTTTTTTTTT-AAAAAAAAAATTGCAAGTAGATGAAATCGTGAATAAATGTGTTTCATTTCTATATCATTATATAAAAAAAGATCCATATTATATTACTTATGGAATTCGAACCCGAACTCGATTTATGATTTATTATTTGGATCTTATTGGCCTTTGTTCTTTATTTTTGATTTTTCATATAGATTTCCGTCTTTATATTATA--CTTTCGTGGATGAATTATGCCTATTTTCACATCTAGGATTTACATATACAACATATATTACTGTCAAGAGGGAATTTTTCTCAGTATTTAGATATTTAGATTCAAAATAAGAAAGGGATCAATTCAATTATAAACCTGCAAAACAAAGATTAGGATTGGGTTGGGTTGCGCTATATATATCAAAGAGTATACAATAATGATGTATTTGGTGAATTAAATACATGGTCTAATAACAAACCATTTTAACATAACATTTTAATGAGTTGATAATATTAATTGAATATTGAATATTTTTTGAAAGATTTTTGTTAAAGGTTTCATTCATGCCTAATCCATATCGAGTAGACCTTGTTGTTGTG??

			>Aechmea_rubiginosa
			??????????????????????????????????????????????????????????????????????????????????????????????????????????????????????????????????????????????????????????????????????????????????????????????????????????????????????????????????????????????????????????????????????????????????????????????????????????????????????????????????????????????????????????????????????????????????????????????????????????????????????????????????????????????????????????????????????????????????????????????????????????????????????????????????????????????????????????????????????????????????????????????????????????????????????????????????????????????????????????????????????????????????????????????????????????????????????????????????????????????????????????????????????????????????????????????

			>Aechmea_serragrandensis
			??????????????????????????????????????????????????????????????????????????????????????????????????????????????????????????????????????????????????????????????????????????????????????????????????????????????????????????????????????????????????????????????????????????????????????????????????????????????????????????????????????????????????????????????????????????????????????????????????????????????????????????????????????????????????????????????????????????????????????????????????????????????????????????????????????????????????????????????????????????????????????????????????????????????????????????????????????????????????????????????????????????????????????????????????????????????????????????????????????????????????????????????????????????????????????????????

			>Aechmea_servitensis
			??????????????????????????????????????????????????????????????????????????????????????????????????????????????????????????????????????????????????????????????????????????????????????????????????????????????????????????????????????????????????????????????????????????????????????????????????????????????????????????????????????????????????????????????????????????????????????????????????????????????????????????????????????????????????????????????????????????????????????????????????????????????????????????????????????????????????????????????????????????????????????????????????????????????????????????????????????????????????????????????????????????????????????????????????????????????????????????????????????????????????????????????????????????????????????????????

			>Aechmea_servitensis_var._exigua
			??????????????????????????????????????????????????????????????????????????????????????????????????????????????????????????????????????????????????????????????????????????????????????????????????????????????????????????????????????????????????????????????????????????????????????????????????????????????????????????????????????????????????????????????????????????????????????????????????????????????????????????????????????????????????????????????????????????????????????????????????????????????????????????????????????????????????????????????????????????????????????????????????????????????????????????????????????????????????????????????????????????????????????????????????????????????????????????????????????????????????????????????????????????????????????????????

			>Aechmea_sphaerocephala
			ATAATTATAATAAAGTGAAATATGTCGAAATCCTTTTTGGAATAATACCAAATCGAAAATAAATGTCCGATAGCAAGTTGATCAGTTAATTCAATAAGAGATAAATGGGAGATAGCACTCGATTTAGTTGGTACCACCCAACCGAATCCGATTCACTCGTTTACTCATTGAATGAGTAAATTTTCAAGTTCAGCCAATCTTTTTTTTTTTAAAAAAAAAATTGCAAGTAGATGAAATCGTGAATAAATGTGTTTCATTTCTATATCATTATATAAAAAAAGATCCATATTATATTACTTATGGAATTCGAACCCGAACTCGATTTATGATTTATTATTTGGATCTTATTGGCCTTTGTTCTTTATTTTTGATTTTTCATATAGATTTCCGTCTTTATATTATACCCTTTCGTGGATGAATTATGCCTATTTTCACATCTAGGATTTACATATACAACATATATTACTGTCAAGAGGGAATTTTTCTCAGTATTTAGATATTTAGATTCAAAATAAGAAAAGGATCAATTCAATTATAAACCTGCAAAACAAAGATTAGGATTGGGTTGGGTTGCGCTATATATATCAAAGAGTATACAATAATGATGTATTTGGTGAATCAAATACATGGTCTAATAACAAACCATTTTAACATAACATTTTAATGAGTTGATAATATTAATTGAATATTGAATATTTTTTGAAAGATTTTTGTTAAAGGTTTCATTCATGCCTAATCCATATCGAGTAGACCTTGTTGTTGTG??

			>Aechmea_squarrosa
			????????AATAAAGTGAAATATGTCGAAATCCTTTTTGGAATAATACCAAATCGAAAAAAAATGTCCGATAGCAAGTTGATCAGTTAATTCAATAAGAGATAAATGGGAGATAGCACTCGATTTAGTTGGTACCACCCAACCGAATCCGATTCACTCGTTTACTCATTGAATGAGTAAATTTTCAAGTTCAGCCAATCTTTTTTTTTTCAAAAAAAAAATTGCAAGTAGATTAAATCATGAATAAATGTGTTTCATTTCTATATCATTATAGAAAAAAAGATCCATATTATATTACTTATGGAATTCGAACCCGAACTCGATTTATGATTTATTATTTGGATCTTATTGGCCTTTGTTCTTTATTTTTGATTTTTCATATAGATTTCCGTCTTTATATTATACCCTTTCGTGGATGAATTATGCCTATTTTCACATCTAGGATTTACATATACAACATATATTACTGTCAAGAGGGAATTTTTCTCAGTATTTAGATATTTAGATTCAAAATAAGAAAGGGATCAATTCAATTATAAACCTGCAAAACAAAGATTAGGATTGGGTTGGGTTGCGCTATATATATCAAAGAGTATACAATAATGATGTATTTGGTGAATCAAATACATGGTCTAATAACAAACCATTTTAACATAACATTTTAATGAGTTGATAATATTAATTGAATATTGAATATTTTTTGAAAGATTTTTGTTAAAGGTTTCATTCATGCCTAATCCAT?????????????????????????

			>Aechmea_strobilacea
			??????????????????????????????????????????????????????????????????????????????????????????????????????????????????????????????????????????????????????????????????????????????????????????????????????????????????????????????????????????????????????????????????????????????????????????????????????????????????????????????????????????????????????????????????????????????????????????????????????????????????????????????????????????????????????????????????????????????????????????????????????????????????????????????????????????????????????????????????????????????????????????????????????????????????????????????????????????????????????????????????????????????????????????????????????????????????????????????????????????????????????????????????????????????????????????????

			>Aechmea_tayoensis
			??????????????????????????????????????????????????????????????????????????????????????????????????????????????????????????????????????????????????????????????????????????????????????????????????????????????????????????????????????????????????????????????????????????????????????????????????????????????????????????????????????????????????????????????????????????????????????????????????????????????????????????????????????????????????????????????????????????????????????????????????????????????????????????????????????????????????????????????????????????????????????????????????????????????????????????????????????????????????????????????????????????????????????????????????????????????????????????????????????????????????????????????????????????????????????????????

			>Aechmea_tessmannii
			??????????????????????????????????????????????????????????????????????????????????????????????????????????????????????????????????????????????????????????????????????????????????????????????????????????????????????????????????????????????????????????????????????????????????????????????????????????????????????????????????????????????????????????????????????????????????????????????????????????????????????????????????????????????????????????????????????????????????????????????????????????????????????????????????????????????????????????????????????????????????????????????????????????????????????????????????????????????????????????????????????????????????????????????????????????????????????????????????????????????????????????????????????????????????????????????

			>Aechmea_tillandsioides
			??????????????????????????????????????????????????????????????????????????????????????????????????????????????????????????????????????????????????????????????????????????????????????????????????????????????????????????????????????????????????????????????????????????????????????????????????????????????????????????????????????????????????????????????????????????????????????????????????????????????????????????????????????????????????????????????????????????????????????????????????????????????????????????????????????????????????????????????????????????????????????????????????????????????????????????????????????????????????????????????????????????????????????????????????????????????????????????????????????????????????????????????????????????????????????????????

			>Aechmea_triangularis
			??????????????????????????????????????????????????????????????????????????????????????????????????????????????????????????????????????????????????????????????????????????????????????????????????????????????????????????????????????????????????????????????????????????????????????????????????????????????????????????????????????????????????????????????????????????????????????????????????????????????????????????????????????????????????????????????????????????????????????????????????????????????????????????????????????????????????????????????????????????????????????????????????????????????????????????????????????????????????????????????????????????????????????????????????????????????????????????????????????????????????????????????????????????????????????????????

			>Aechmea_triticina
			??????????????????????????????????????????????????????????????????????????????????????????????????????????????????????????????????????????????????????????????????????????????????????????????????????????????????????????????????????????????????????????????????????????????????????????????????????????????????????????????????????????????????????????????????????????????????????????????????????????????????????????????????????????????????????????????????????????????????????????????????????????????????????????????????????????????????????????????????????????????????????????????????????????????????????????????????????????????????????????????????????????????????????????????????????????????????????????????????????????????????????????????????????????????????????????????

			>Aechmea_vallerandii
			???????????????????????????????????TTTGGAATAATACCAAATCGAAAATAAATGTCCGATAGCAAGTTGATCAGTTAATTCAATAAGAGATAAATGGGAGATAGCACTCGATTTAGTTGGTACCACCCAACTGAATCCGATTCACTCGTTTACTCATTGAATGAGTAAATTTTCAAGTTCAGCCAATC-TTTTTTTTTTAAAAAAAAAATTGCAAGTAGATGAAATCGTGAATAAATGTGTTTCATTTCTATATCATTATATAAAAAAAGATCCATATTATATTACTTATGGAATTCGAACCCGAACTCGATTTATGATTTATTATTTGGATCTTATTGGCCTTTGTTCTTTATTTTTGATTTTTCATATAGATTTCCGTCTTTATATTATACCCTTTCGTGGATGAATTATGCCTATTTTCACATCTAGGATTTACATATACAACATATATTACTGTCAAGAGGGAATTTTTCTCAGTATTTAGATATTTAGATTCAAAATAAGAAAGGGATCAATTCAATTATAAACCTGCAAAACAAAGATTAGGATTGGGTTGGGTTGCGCTATATATATCAAAGAGTATACAATAATGATGTATTTGGTGAATCAAATACATGGTCTAATAACAAACCATTTTAACATAACATTTTAATGAGTTGATAATATTAATTGAATATTGAATATTTTTTGAAAGATTTTTGTTAAAGGTTTCATTCATGCCTAATCCATATCGAGTAGACCTTGTTGTTGTG??

			>Aechmea_vanhoutteana
			??????????????????????????????????????????????????????????????????????????????????????????????????????????????????????????????????????????????????????????????????????????????????????????????????????????????????????????????????????????????????????????????????????????????????????????????????????????????????????????????????????????????????????????????????????????????????????????????????????????????????????????????????????????????????????????????????????????????????????????????????????????????????????????????????????????????????????????????????????????????????????????????????????????????????????????????????????????????????????????????????????????????????????????????????????????????????????????????????????????????????????????????????????????????????????????????

			>Aechmea_victoriana
			??????????????????????????????????????????????????????????????????????????????????????????????????????????????????????????????????????????????????????????????????????????????????????????????????????????????????????????????????????????????????????????????????????????????????????????????????????????????????????????????????????????????????????????????????????????????????????????????????????????????????????????????????????????????????????????????????????????????????????????????????????????????????????????????????????????????????????????????????????????????????????????????????????????????????????????????????????????????????????????????????????????????????????????????????????????????????????????????????????????????????????????????????????????????????????????????

			>Aechmea_weilbachii
			??????????????????????????????????????????????????????????????????????????????????????????????????????????????????????????????????????????????????????????????????????????????????????????????????????????????????????????????????????????????????????????????????????????????????????????????????????????????????????????????????????????????????????????????????????????????????????????????????????????????????????????????????????????????????????????????????????????????????????????????????????????????????????????????????????????????????????????????????????????????????????????????????????????????????????????????????????????????????????????????????????????????????????????????????????????????????????????????????????????????????????????????????????????????????????????????

			>Aechmea_werdermannii
			??????????????????????????????????????????????????????????????????????????????????????????????????????????????????????????????????????????????????????????????????????????????????????????????????????????????????????????????????????????????????????????????????????????????????????????????????????????????????????????????????????????????????????????????????????????????????????????????????????????????????????????????????????????????????????????????????????????????????????????????????????????????????????????????????????????????????????????????????????????????????????????????????????????????????????????????????????????????????????????????????????????????????????????????????????????????????????????????????????????????????????????????????????????????????????????????

			>Aechmea_winkleri
			??????????????????????????????????????????????????????????????????????????????????????????????????????????????????????????????????????????????????????????????????????????????????????????????????????????????????????????????????????????????????????????????????????????????????????????????????????????????????????????????????????????????????????????????????????????????????????????????????????????????????????????????????????????????????????????????????????????????????????????????????????????????????????????????????????????????????????????????????????????????????????????????????????????????????????????????????????????????????????????????????????????????????????????????????????????????????????????????????????????????????????????????????????????????????????????????

			>Aechmea_woronowii
			??????????????????????????????????????????????????????????????????????????????????????????????????????????????????????????????????????????????????????????????????????????????????????????????????????????????????????????????????????????????????????????????????????????????????????????????????????????????????????????????????????????????????????????????????????????????????????????????????????????????????????????????????????????????????????????????????????????????????????????????????????????????????????????????????????????????????????????????????????????????????????????????????????????????????????????????????????????????????????????????????????????????????????????????????????????????????????????????????????????????????????????????????????????????????????????????

			>Aechmea_zebrina
			??????????????????????????????????????????????????????????????????????????????????????????????????????????????????????????????????????????????????????????????????????????????????????????????????????????????????????????????????????????????????????????????????????????????????????????????????????????????????????????????????????????????????????????????????????????????????????????????????????????????????????????????????????????????????????????????????????????????????????????????????????????????????????????????????????????????????????????????????????????????????????????????????????????????????????????????????????????????????????????????????????????????????????????????????????????????????????????????????????????????????????????????????????????????????????????????

			>Alcantarea_aurantiaca
			??????????????????????????????????????????????????????????????????????????????????????????????????????????????????????????????????????????????????????????????????????????????????????????????????????????????????????????????????????????????????????????????????????????????????????????????????????????????????????????????????????????????????????????????????????????????????????????????????????????????????????????????????????????????????????????????????????????????????????????????????????????????????????????????????????????????????????????????????????????????????????????????????????????????????????????????????????????????????????????????????????????????????????????????????????????????????????????????????????????????????????????????????????????????????????????????

			>Alcantarea_benzingii
			??????????????????????????????????????????????????????????????????????????????????????????????????????????????????????????????????????????????????????????????????????????????????????????????????????????????????????????????????????????????????????????????????????????????????????????????????????????????????????????????????????????????????????????????????????????????????????????????????????????????????????????????????????????????????????????????????????????????????????????????????????????????????????????????????????????????????????????????????????????????????????????????????????????????????????????????????????????????????????????????????????????????????????????????????????????????????????????????????????????????????????????????????????????????????????????????

			>Alcantarea_burle-marxii
			??????????????????????????????????????????????????????????????????????????????????????????????????????????????????????????????????????????????????????????????????????????????????????????????????????????????????????????????????????????????????????????????????????????????????????????????????????????????????????????????????????????????????????????????????????????????????????????????????????????????????????????????????????????????????????????????????????????????????????????????????????????????????????????????????????????????????????????????????????????????????????????????????????????????????????????????????????????????????????????????????????????????????????????????????????????????????????????????????????????????????????????????????????????????????????????????

			>Alcantarea_compacta
			??????????????????????????????????????????????????????????????????????????????????????????????????????????????????????????????????????????????????????????????????????????????????????????????????????????????????????????????????????????????????????????????????????????????????????????????????????????????????????????????????????????????????????????????????????????????????????????????????????????????????????????????????????????????????????????????????????????????????????????????????????????????????????????????????????????????????????????????????????????????????????????????????????????????????????????????????????????????????????????????????????????????????????????????????????????????????????????????????????????????????????????????????????????????????????????????

			>Alcantarea_duarteana
			ATAATTATAATAAAGTGAAATATGTCGAAATCCTTTTTGGAATAATACCAAATCGAAAATAAATGTCCGATAGCAAGTTGATCAGTTAATTCAATAAGAGATAAATGGGAGATAGCACTCGATTTAGTTGGTACCACCCAACCGAATCCGATTCAATCGTTTACTCATTCAATGAGTAAATTTTCAAGTTCAGCCAATCTTTTTTTTTTCAAAAAAAAAATTGCAAGTAAATGAAATCGTGAATAAATGTGTTTCATTTCTCTATCATTATAGAAAAAAAGATCCATATTATATTACTTATGGAATTCGAGCCCGAACTCGATTTATGATTCATTATTTCGATCTTATTGGCCTTTGTTCTTTATTTTTTATTTTTCATATAGATTTCCGTCTTTATATTATACCCTTTCGTAGATGAATTATGCTTATTTTCACATCTAGGATTTACATATACAACATATATTACTGTCAAGAGGGAATTTTTCTCAGTATTTAGATATTTAGATTCAAAATAAGAAAGGGATCAATTCAATTATAAACCTGCAAAACAAAGATTAGGATTGGGTTGGGTTGCGCTATATATATCAAAGAGTATACAATAATGATGTATTTGGTGAATCAAATACATGGTCTAATAACAAACCATTTTAACATAACATTTTAATGAGTTGATAATATTAATTGAATATTGAATATTTTTTGAAAGATTTTTGTTAAAGGTTTCATTCATGCCTAATCCATATCGAGTAGACCTTGTTGTTGTGAG

			>Alcantarea_farneyi
			??????????????????????????????????????????????????????????????????????????????????????????????????????????????????????????????????????????????????????????????????????????????????????????????????????????????????????????????????????????????????????????????????????????????????????????????????????????????????????????????????????????????????????????????????????????????????????????????????????????????????????????????????????????????????????????????????????????????????????????????????????????????????????????????????????????????????????????????????????????????????????????????????????????????????????????????????????????????????????????????????????????????????????????????????????????????????????????????????????????????????????????????????????????????????????????????

			>Alcantarea_geniculata
			??????????????????????????????????????????????????????????????????????????????????????????????????????????????????????????????????????????????????????????????????????????????????????????????????????????????????????????????????????????????????????????????????????????????????????????????????????????????????????????????????????????????????????????????????????????????????????????????????????????????????????????????????????????????????????????????????????????????????????????????????????????????????????????????????????????????????????????????????????????????????????????????????????????????????????????????????????????????????????????????????????????????????????????????????????????????????????????????????????????????????????????????????????????????????????????????

			>Alcantarea_glaziouana
			??????????????????????????????????????????????????????????????????????????????????????????????????????????????????????????????????????????????????????????????????????????????????????????????????????????????????????????????????????????????????????????????????????????????????????????????????????????????????????????????????????????????????????????????????????????????????????????????????????????????????????????????????????????????????????????????????????????????????????????????????????????????????????????????????????????????????????????????????????????????????????????????????????????????????????????????????????????????????????????????????????????????????????????????????????????????????????????????????????????????????????????????????????????????????????????????

			>Alcantarea_heloisae
			??????????????????????????????????????????????????????????????????????????????????????????????????????????????????????????????????????????????????????????????????????????????????????????????????????????????????????????????????????????????????????????????????????????????????????????????????????????????????????????????????????????????????????????????????????????????????????????????????????????????????????????????????????????????????????????????????????????????????????????????????????????????????????????????????????????????????????????????????????????????????????????????????????????????????????????????????????????????????????????????????????????????????????????????????????????????????????????????????????????????????????????????????????????????????????????????

			>Alcantarea_imperialis
			ATAATTATAATAAAGTGAAATATGTCGAAATCCTTTTTGGAATAATACCAAATCGAAAATAAATGTCCGATAGCAAGTTGATCAGTTAATTCAATAAGAGATAAATGGGAGATAGCACTCGATTTAGTTGGTACCACCCAACCGAATCCTATTCAATCGTTTACTCATTCAATGAGTAAATTTGCAAGTTCAGCCAATCTTTTTTTTT--AAAAAAAAAATTGCAAGTAAATGAAATCGTGAATAAATGTGTTTCATTTCTCTATCATTATAGAAAAAAAGATCCATATTATATTACTTATGAAATTCGAGCCCGAACTCGATTTATGATTCATTATTTCGATCTTATTGGCCTTTGTTCTTTATTTTTTATTTTTCATATAGATTTCCGTCTTTATATTATACCCTTTCGTAGATGAATTATGCTTATTTTCACATCTAGGATTTACATATACAACATATATTACTGTCAAGAGGGAATTTTTCTCAGTATTTAGATATTTAGATTCAAAATAAGAAAGGG-----TTCAATTATAAACCTGCAAAACAAAGATTAGGATTGGGTTGGGTTGCGCTATATATATCAAAGAGTATACAATAATGATGTATTTGGTGAATCAAATACATGGTCTAATAACAAACCATTTTAACATAACATTTTAATGAGTTGATAATATTAATTGAATATTGAATATTTTTTGAAAGATTTTTGTTAAAGGTTTCATTCATGCCCAATCCATATCGAGTAGACCTTGTTGTTGTGAG

			>Alcantarea_martinellii
			??????????????????????????????????????????????????????????????????????????????????????????????????????????????????????????????????????????????????????????????????????????????????????????????????????????????????????????????????????????????????????????????????????????????????????????????????????????????????????????????????????????????????????????????????????????????????????????????????????????????????????????????????????????????????????????????????????????????????????????????????????????????????????????????????????????????????????????????????????????????????????????????????????????????????????????????????????????????????????????????????????????????????????????????????????????????????????????????????????????????????????????????????????????????????????????????

			>Alcantarea_nahoumii
			??????????????????????????????????????????????????????????????????????????????????????????????????????????????????????????????????????????????????????????????????????????????????????????????????????????????????????????????????????????????????????????????????????????????????????????????????????????????????????????????????????????????????????????????????????????????????????????????????????????????????????????????????????????????????????????????????????????????????????????????????????????????????????????????????????????????????????????????????????????????????????????????????????????????????????????????????????????????????????????????????????????????????????????????????????????????????????????????????????????????????????????????????????????????????????????????

			>Alcantarea_nevaresii
			??????????????????????????????????????????????????????????????????????????????????????????????????????????????????????????????????????????????????????????????????????????????????????????????????????????????????????????????????????????????????????????????????????????????????????????????????????????????????????????????????????????????????????????????????????????????????????????????????????????????????????????????????????????????????????????????????????????????????????????????????????????????????????????????????????????????????????????????????????????????????????????????????????????????????????????????????????????????????????????????????????????????????????????????????????????????????????????????????????????????????????????????????????????????????????????????

			>Alcantarea_odorata
			??????????????????????????????????????????????????????????????????????????????????????????????????????????????????????????????????????????????????????????????????????????????????????????????????????????????????????????????????????????????????????????????????????????????????????????????????????????????????????????????????????????????????????????????????????????????????????????????????????????????????????????????????????????????????????????????????????????????????????????????????????????????????????????????????????????????????????????????????????????????????????????????????????????????????????????????????????????????????????????????????????????????????????????????????????????????????????????????????????????????????????????????????????????????????????????????

			>Alcantarea_recurvifolia
			??????????????????????????????????????????????????????????????????????????????????????????????????????????????????????????????????????????????????????????????????????????????????????????????????????????????????????????????????????????????????????????????????????????????????????????????????????????????????????????????????????????????????????????????????????????????????????????????????????????????????????????????????????????????????????????????????????????????????????????????????????????????????????????????????????????????????????????????????????????????????????????????????????????????????????????????????????????????????????????????????????????????????????????????????????????????????????????????????????????????????????????????????????????????????????????????

			>Alcantarea_regina
			??????????????????????????????????????????????????????????????????????????????????????????????????????????????????????????????????????????????????????????????????????????????????????????????????????????????????????????????????????????????????????????????????????????????????????????????????????????????????????????????????????????????????????????????????????????????????????????????????????????????????????????????????????????????????????????????????????????????????????????????????????????????????????????????????????????????????????????????????????????????????????????????????????????????????????????????????????????????????????????????????????????????????????????????????????????????????????????????????????????????????????????????????????????????????????????????

			>Alcantarea_roberto-kautskyi
			??????????????????????????????????????????????????????????????????????????????????????????????????????????????????????????????????????????????????????????????????????????????????????????????????????????????????????????????????????????????????????????????????????????????????????????????????????????????????????????????????????????????????????????????????????????????????????????????????????????????????????????????????????????????????????????????????????????????????????????????????????????????????????????????????????????????????????????????????????????????????????????????????????????????????????????????????????????????????????????????????????????????????????????????????????????????????????????????????????????????????????????????????????????????????????????????

			>Alcantarea_vasconcelosiana
			??????????????????????????????????????????????????????????????????????????????????????????????????????????????????????????????????????????????????????????????????????????????????????????????????????????????????????????????????????????????????????????????????????????????????????????????????????????????????????????????????????????????????????????????????????????????????????????????????????????????????????????????????????????????????????????????????????????????????????????????????????????????????????????????????????????????????????????????????????????????????????????????????????????????????????????????????????????????????????????????????????????????????????????????????????????????????????????????????????????????????????????????????????????????????????????????

			>Ananas_ananassoides
			ATAATTATAATAAAGTGAAATATGTCGAAATCCTTTTTGGAATAATACCAAATCGAAAATAAATGTCCGATAGCAAGTTGATCAGTTAATTCAATAAGAGATAAATGGGAGATAGCACTCGATTTAGTTGGTACCACCCAACCGAATCCGATTCACTCGTTTACTCATTCAATGAGTAAATTTTCAAGTTCAGCCAATCTTTTTTTTTTT-AAAAAAAAATTGCAAGTAGATGAAATCGTGAATAAATGTGTTTCATTTCTCTATCATTATATAAAAAAAGATCCATATTATATTACTTATGGAATTCGAACCCGAACTCGATTTATGATTTATTATTTGGATCTTATTGGCCTTTGTTCTTTATTTTGGATTTTTCATATAGATTTCCGTCTTTATATTATACCCTTTCGTGGATGAATTATGCCTATTTTCACATCTAGGATTTACATATACAACATATATTACTGTCAAGGGGGAATTTTTCTCAGTATTTAGATATTTAGATTCAAAATAAGAAAGGGATCAATTCAATTATAAACCTGCAAAACAAAGATTAGGATTGGGTTGGGTTGCGCTATATATATCAAAGAGTATACAATAATGATGTATTTGGTGAATCAAATACATGGTCTAATAACAAACCGTTTTAACATAACATTTTAATGAGTTGATAATATTAATTGAATATTGAATATTTTTTGAAAGATTTTTGTTAAAGGTTTCATTCATGCCTAATCCATATCGAGTAGACCTTGTTGTTGTGAG

			>Ananas_ananassoides_var._nanus
			ATAATTATAATAAAGTGAAATATGTCGAAATCCTTTTTGGAATAATACCAAATCGAAAATAAATGTCCGATAGCAAGTTGATCAGTTAATTCAATAAGAGATAAATGGGAGATAGCACTCGATTTAGTTGGTACCACCCAACCGAATCCGATTCACTCGTTTACTCATTCAATGAGTAAATTTTCAAGTTCAGCCAATCTTTTTTTTTTTAAAAAAAAAATTGCAAGTAGATGAAATCGTGAATAAATGTGTTTCATTTCTCTATCATTATATAAAAAAAGATCCATATTATATTACTTATGGAATTCGAACCCGAACTCGATTTATGATTTATTATTTGGATCTTATTGGCCTTTGTTCTTTATTTTGGATTTTTCATATAGATTTCCGTCTTTATATTATACCCTTTCGTGGATGAATTATGCCTATTTTCACATCTAGGATTTACATATACAACATATATTACTGTCAAGGGGGAATTTTTCTCAGTATTTAGATATTTAGATTCAAAATAAGAAAGGGATCAATTCAATTATAAACCTGCAAAACAAAGATTAGGATTGGGTTGGGTTGCGCTATATATATCAAAGAGTATACAATAATGATGTATTTGGTGAATCAAATACATGGTCTAATAACAAACCGTTTTAACATAACATTTTAATGAGTTGATAATATTAATTGAATATTGAATATTTTTTGAAAGATTTTTGTTAAAGGTTTCATTCATGCCTAATCCATATCGAGTAGACCTTGTTGTTGTG??

			>Ananas_bracteatus
			??????????????????????????????ATCCTTTTTGGATAATACCAAATCGAAAATAAATGTCCGATAGCAAGTTGATCAGTTAATTCAATAAGAGATAAATGGGAGATAGCACTCGATTTAGTTGGTACCACCCAACCGAATCCGATTCACTCGTTTACTCATTCAATGAGTAAATTTTCAAGTTCAGCCAATCTTTTTTTTTTTAAAAAAAA??TTGCAAGTAGATGAAATCGTGAATAAATGTGTTTCATTTCTCTATCATTATATAAAAAAAGATCCATATTATATTACTTATGGAATTCGAACCCGAACTCGATTTATGATTTATTATTTGGATCTTATTGGCCTTTGTTCTTTATTTTGGATTTTTCATATAGATTTCCGTCTTTATATTATACCCTTTCGTGGATGAATTATGCCTA-TTTCACATCTAGGATTTACATATACAACATATATTACTGTCAAGGGGGAATTTTTCTCAGTATTTAGATATTTAGATTCAAAATAAGAAAGGGATCAATTCAATTATAAACCTGCAAAACAAAGATTAGGATTGGGTTGGGTTGCGCTATATATATCAAAGAGTATACAATAATGATGTATTTGGTGAATCAAATACATGGTCTAATAACAAACCGTTTTAACATAACATTTTAATGAGTTGATAATATTAATTGAATATTGAATATTTTTTGAAAGATTTTTGTTAAAGGTTTCATTCATGCCTAATCCATATCGAGTAGACCTTGTTGTTGTG??

			>Ananas_comosus
			??AATTATAATAAAGTGAAATATGTCGAAATCCTTTTTGGAATAATACCAAATCGAAAATAAATGTCCGATAGCAAGTTGATCAGTTAATTCAATAAGAGATAAATGGGAGATAGCACTCGATTTAGTTGGTACCACCCAACCGAATCCGATTCACTCGTTTACTCATTCAATGAGTAAATTTTCAAGTTCAGCCAATCTTTTTTTTTTT-AAAAAAAAATTGCAAGTAGATGAAATCGTGAATAAATGTGTTTCATTTCTCTATCATTATATAAAAAAAGATCCATATTATATTACTTATGGAATTCGAACCCGAACTCGATTTATGATTTATTATTTGGATCTTATTGGCCTTTGTTCTTTATTTTGGATTTTTCATATAGATTTCCGTCTTTATATTATACCCTTTCGTGGATGAATTATGCCTATTTTCACATCTAGGATTTACATATAC?ACATATATTACTGTCAAGGGGGAATTTTTCTCAGTATTTAGATATTTAGATTCAAAATAAGAAAGGGATCAATTCAATTATAAACCTGCAAAACAAAGATTAGGATTGGGTTGGGTTGCGCTATATATATCAAAGAGTATACAATAATGATGTATTTGGTGAATCAAATACATGGTCTAATAACAAACCGTTTTAACATAACATTTTAATGAGTTGATAATATTAATTGAATATTGAATATTTTTTGAAAGATTTTTGTTAAAGGTTTCATTCATGCCTAATCCATATCGAGTAGACCTTGTTGTTGTGAG

			>Ananas_comosus_var._microstachys
			??????????????????????????????????????????????????????????????????????????????????????????????????????????????????????????????????????????????????????????????????????????????????????????????????????????????????????????????????????????????????????????????????????????????????????????????????????????????????????????????????????????????????????????????????????????????????????????????????????????????????????????????????????????????????????????????????????????????????????????????????????????????????????????????????????????????????????????????????????????????????????????????????????????????????????????????????????????????????????????????????????????????????????????????????????????????????????????????????????????????????????????????????????????????????????????????

			>Ananas_comosus_var._parguazensis
			??????????????????????????????????????????????????????????????????????????????????????????????????????????????????????????????????????????????????????????????????????????????????????????????????????????????????????????????????????????????????????????????????????????????????????????????????????????????????????????????????????????????????????????????????????????????????????????????????????????????????????????????????????????????????????????????????????????????????????????????????????????????????????????????????????????????????????????????????????????????????????????????????????????????????????????????????????????????????????????????????????????????????????????????????????????????????????????????????????????????????????????????????????????????????????????????

			>Ananas_fritzmuelleri
			ATAATTATAATAAAGTGAAATATGTCGAAATCCTTTTTGGAATAATACCAAATCGAAAATAAATGTCCGATAGCAAGTTGATCAGTTAATTCAATAAGAGATAAATGGGAGATAGCACTCGATTTAGTTGGTACCACCCAACCGAATCCGATTCACTCGTTTACTCATTCAATGAGTAAATTTTCAAGTTCAGCCAATCTTTTTTTTTTTAAAAAAAAAATTGCAAGTAGATGAAATCGTGAATAAATGTGTTTCATTTCTCTATCATTATATAAAAAAAGATCCATATTATATTACTTATGGAATTCGAACCCGAACTCGATTTATGATTTATTATTTGGATCTTATTGGCCTTTGTTCTTTATTTTGGATTTTTCATATAGATTTCCGTCTTTATATTATACCCTTTCGTGGATGAATTATGCCTATTTTCACATCTAGGATTTACATATACAACATATATTACTGTCAAGGGGGAATTTTTCTCAGTATTTAGATATTTAGATTCAAAATAAGAAAGGGATCAATTCAATTATAAACCTGCAAAACAAAGATTAGGATTGGGTTGGGTTGCGCTATATATATCAAAGAGTATACAATAATGATGTATTTGGTGAATCAAATACATGGTCTAATAACAAACCGTTTTAACATAACATTTTAATGAGTTGATAATATTAATTG??????????????????????????????????????????????????????????????????????????????????

			>Ananas_sagenaria
			ATAATTATAATAAAGTGAAATATGTCGAAATCCTTTTTGGAATAATACCAAATCGAAAATAAATGTCCGATAGCAAGTTGATCAGTTAATTCAATAAGAGATAAATGGGAGATAGCACTCGATTTAGTTGGTACCACCCAACCGAATCCGATTCACTCGTTTACTCATTCAATGAGTAAATTTTCAAGTTCAGCCAATCTTTTTTTTTTTCAAAAAAAAATTGCAAGTAGATGAAATCGTGAATAAATGTGTTTCATTTCTCTATCATTATATAAAAAAAGATCCATATTATATTACTTATGGAATTCGAACCCGAACTCGATTTATGATTTATTATTTGGATCTTATTGGCCTTTGTTCTTTATTTTGGATTTTTCATATAGATTTCCGTCTTTATATTATACCCTTTCGTGGATGAATTATGCCTATTTTCACATCTAGGATTTACATATACAACATATATTACTGTCAAGGGGGAATTTTTCTCAGTATTTAGATATTTAGATTCAAAATAAGAAAGGGATCAATTCAATTATAAACCTGCAAAACAAAGATTAGGATTGGGTTGGGTTGCGCTATATATATCAAAGAGTATACAATAATGATGTATTTGGTGAATCAAATACATGGTCTAATAACAAACCGTTTTAACATAACATTTTAATGAGTTGATAATATTAATTGAATATTGAATATTTTTTGAAAGATTTTTGTTAAAGGTTTCATTCATGCCTAATCCATATCGAGTAGACCTTGTTGTTGTGAG

			>Androlepis_fragrans
			??????????????????????????????????????????????????????????????????????????????????????????????????????????????????????????????????????????????????????????????????????????????????????????????????????????????????????????????????????????????????????????????????????????????????????????????????????????????????????????????????????????????????????????????????????????????????????????????????????????????????????????????????????????????????????????????????????????????????????????????????????????????????????????????????????????????????????????????????????????????????????????????????????????????????????????????????????????????????????????????????????????????????????????????????????????????????????????????????????????????????????????????????????????????????????????????

			>Androlepis_skinneri
			ATAATTATAATAAAGTGAAATATGTCGAAATCCTTTTTGGAATAATACCAAATCGAAAATAAATGTCCGATAGCAAGTTGATCAGTTAATTCAATAAGAGATAAATGGGAGATAGCACTCGATTTAGTTGGTACCACCCAACCGAATCCGATTCACTCGTTTACTCATTGAATGAGTAAATTTTCAAGTTCAGCCAATC-TTTTTTTTTTAAAAAAAAAATTGCAAGTAGATGAAATCGTGAATAAATGTGTTTCATTTCTATATCATTATATAAAAAAAGATCCATATTATATTACTTATGGAATTCGAACCCGAACTCGATTTATGATTTATTATTTGGATCTTATTGGCCTTTGTTCTTTATTTTTGATTTTTCATATAGATTTCCGTCTTTATATTATACCCTTTCGTGGATGAATTATGCCTATTTTCACATCTAGGATTTACATATACAACATATATTACTGTCAAGAGGGAATTTTTCTCAGTATTTAGATATTTAGATTCAAAATAAGAAAGGGATCAATTCAATTATAAACCTGCAAAACAAAGATTAGGATTGGGTTGGGTTGCGCTATATATATCAAAGAGTATACAATAATGATGTATTTGGTGAATCAAATACATGGTCTAATAACAAACCATTTTAACATAACATTTTAATGAGTTGATAATATTAATTGAATATTGAATATTTTTTGAAAGATTTTTGTTAAAGGTTTCATTCATGCCTAATCCATATCGAGTAGACCTTGTTGTTGTG??

			>Araeococcus_chlorocarpus
			??????????????????????????????????????????????????????????????????????????????????????????????????????????????????????????????????????????????????????????????????????????????????????????????????????????????????????????????????????????????????????????????????????????????????????????????????????????????????????????????????????????????????????????????????????????????????????????????????????????????????????????????????????????????????????????????????????????????????????????????????????????????????????????????????????????????????????????????????????????????????????????????????????????????????????????????????????????????????????????????????????????????????????????????????????????????????????????????????????????????????????????????????????????????????????????????

			>Araeococcus_flagellifolius
			ATAATTATAATAAAGTGAAATATGTCGAAATCCTTTTTGGAATAATACCAAATCGAAAATAAATGTCCGATAGCAAGTTGATCAGTTAATTCAATAAGAGATAAATGGGAGATAGCACTCGATTTAGTTGGTACCACCCAACCGAATCCGATTCACTCGTTTACTCATTCAATGAGTAAATTTTCAAGTTCAGCCAATCTTTTTTTTTTTTGAAAAAAAATTGCAAGTAGATGAAATCGTGAATAAATGTGTTTCATTTCTATATCATTATATAAAAAAGGATCCATATTATATTACTTATGGAATTCGAACCCGAACTCGATTTATGATTTATTATTTGGATCTTATTGGCCTTTGTTCTTTATTTTTGATTTTTCATATAGATTTCCGTCTTTATATTATACCCTTTCGTGGATGAATTATGCCTATTTTCACATCTAGGATTTACATATACAACATATATTACTGTCAAGAGGGAATTTTTCTCAGTATTTAGATATTTAGATTCAAAATAAGAAAGGGATCAATTCAATTATAAACCTGCAAAACAAAGATTAGGATTGGGTTGGGTTGCGCTATATATATCAAAGAGTATACAATAATGATGTATTTGGTGAATCAAATACATGGTCTAATAACAAACCATTTTAACATAACATTTTAATGAGTTGATAATATTAATTGAATATTGAATATTTTTTGAAAGATTTTTGTTAAAGGTTTCATTCATGCCTAATCCATATCGAGTAGACCTTGTTGTTGTG??

			>Araeococcus_goeldianus
			ATAATTATAATAAAGTGAAATATGTCGAAATCCTTTTTGGAATAATACCAAATCGAAAATAAATGTCCGATAGCAAGTTGATCAGTTAATTCAATAAGAGATAAATGGGAGATAGCACTCGATTTAGTTGGTACCACCCAACCGAATCCGATTCACTCGTTTACTCATTCAATGAGTAAATTTTCAAGTTCAGCCAATC-TTTTTTTTTTGAAAAAAAAATTGCAAGTAGATGAAATCGTGAATAAATGTGTTTCATTTCTATATCATTATATAAAAAAGGATCCATATTATATTACTTATGGAATTCGAACCCGAACTCGATTTATGATTTATTATTTGGATCTTATTGGCCTTTGTTCTTTATTTTTGATTTTTCATATAGATTTCCGTCTTTATATTATACCCTTTCGTGGATGAATTATGCCTATTTTCACATCTAGGATTTACATATACAACATATATTACTGTCAAGAGGGAATTTTTCTCAGTATTTAGATATTTAGATTCAAAATAAGAAAGGGATCAATTCAATTATAAACCTGCAAAACAAAGATTAGGATTGGGTTGGGTTGCGCTATATATATCAAAGAGTATACAATAATGATGTATTTGGTGAATCAAATACATGGTCTAATAACAAACCATTTTAACATAACATTTTAATGAGTTGATAATATTAATTGAATATTGAATATTTTTTGAAAGATTTTTGTTAAAGGTTTCATTCATGCCTAATCCATATCGAGTAGACCTTGTTGTTGTG??

			>Araeococcus_micranthus
			??????????????????????????????????????????????????????????????????????????????????????????????????????????????????????????????????????????????????????????????????????????????????????????????????????????????????????????????????????????????????????????????????????????????????????????????????????????????????????????????????????????????????????????????????????????????????????????????????????????????????????????????????????????????????????????????????????????????????????????????????????????????????????????????????????????????????????????????????????????????????????????????????????????????????????????????????????????????????????????????????????????????????????????????????????????????????????????????????????????????????????????????????????????????????????????????

			>Araeococcus_pectinatus
			ATAATTATAATAAAGTGAAATATGTCGAAATCCTTTTTGGAATAATACCAAATC-AAAATAAATGTCCGATAGCAAGTTGATCAGTTAATTCAATAAGAGATAAATGGGAGATAGCACTCGATTTAGTTGGTACCACCCAACCGAATCCGATTCACTCGTTTACTCATTCAATGAGTAAATTTTCAAGTTCAGCCAATCTTTTTTTTTTTCAAAAAAGAATTGCAAGTAGATGAAATCGTGAATAAATGTGTTTCATTTCTATATCATTATATAAAAAAAGATCCATATTATATTACTTATGGAATTCGAACCCGAACTCGATTTATGATTTATTATTTGGATCTTATTGGCCTTTGTTCTTTATTTTTGATTTTTCATATAGATTTCCGTCTTTATATTATACCCTTTCGTGGATGAATTATGCCTATTTTCACATCTAGGATTTACATATACAACATATATTACTGTCAAGAGGGAATTTTTCTCAGTATTTAGATATTTAGATTCAAAATAAGAAAGGGATCAATTCAATTATAAACCTGCAAAACAAAGATTAGGATTGGGTTGGGTTGCGCTATATATATCAAAGAGTATACAATAATGATGTATTTGGTGAATCAAATACATGGTCTAATAACAAACCATTTTAACATAACATTTTAATGAGTTGATAATATTAATTGAATATTGAATATTTTTTGAAAGATTTTTGTTAAAGGTTTCATTCATGCCTAATCCATATCGAGTAGACCTTGTTGTTGTGAG

			>Barfussia_laxissima
			??????????????????????????????????????????????????????????????????????????????????????????????????????????????????????????????????????????????????????????????????????????????????????????????????????????????????????????????????????????????????????????????????????????????????????????????????????????????????????????????????????????????????????????????????????????????????????????????????????????????????????????????????????????????????????????????????????????????????????????????????????????????????????????????????????????????????????????????????????????????????????????????????????????????????????????????????????????????????????????????????????????????????????????????????????????????????????????????????????????????????????????????????????????????????????????????

			>Barfussia_platyrhachis
			??????????????????????????????????????????????????????????????????????????????????????????????????????????????????????????????????????????????????????????????????????????????????????????????????????????????????????????????????????????????????????????????????????????????????????????????????????????????????????????????????????????????????????????????????????????????????????????????????????????????????????????????????????????????????????????????????????????????????????????????????????????????????????????????????????????????????????????????????????????????????????????????????????????????????????????????????????????????????????????????????????????????????????????????????????????????????????????????????????????????????????????????????????????????????????????????

			>Barfussia_wagneriana
			ATAATTATAATAAAGTTAAATATGTCGAAATCTTTTTTGGAATAATACCAAATCGAAAATAAATGTCCGATAGCAAGTTGATCAGTTAATTCAATAAGAGATAAATGGGAGATAGCACTCGATTTAGTTGGTACCACCCAACCGAATCCGATTCAATCGTTTACTCATTCAATGAGTCAATTTTCAAGTTCAGCCAATCTTTTTTTTTTTCAAAAAAAAATTGAAAGTAAATGAAATCGTGAATAAATGTGTTTCATTTCTCTATCATTATAGAAAAAAAGATCCATATTATATTACTTATGGAATTCGAACCCGAACTCGATTTATGATTCATTATTTCGATCTTATTGGCCTTTGTTCTTTATTTTTTATTTTTCATATAGATTTCCGTCTTTATATTATACCCTTTCGTAGATGAATTATGCTTATTTTCACATCTAGGATTTACATATACAACATATATTACTGTCAAGAGGGAATTTTTCTCAGTATTTAGATATTTAGATTCAAAATAAGAAAGGG-----TTCAATTATAAACCCGCAAAACAAAGATTAGGATTGGGTTGGGTTGCGCTATATATATCAAAGAGTATACAATAATGATGTATTTGGTGAATCAAATACATGGTCTAATAACAAACCATTTTAACATAACATTTTAATGAGTTGATAATATTAATTGAATATTGAATATTTTTTGAAAGATTTTTGTTAAGGGTTTCATTCATGCCTAATCCATATCGAGTAGACCTTGTTGTTGTGAG

			>Billbergia_amoena
			???????????????????????????????????TTTGGAATAATACCAAATCGAAAATAAATGTCCGATAGCAAGTTGATCAGTTAATTCAATAAGAGATAAATGGGAGATAGCACTCGATTTAGTTGGTACCACCCAACCGAATCCGATTCACTCGTTTACTCATTCAATGAGTAAATTTTCAAGTTCAGCCAATC-TTTTTTTTTTAAAAAAGAAATTGCAAGTAGATGAAATCGTGAATAAATGTGTTTCATTTCTATATCATTATATAAAAACAGATCCATATTATATTACTTATGGAATTCGAACCCGAACTCGATTTATGATTTATTATTTGGATCTTATTGGCCTTTGTTCTTTATTTTTGATTTTTCATATATATTTCCGTCTTTATATTATACCCTTTCGTGGATGAATTATGCCTATTTTCACATCTAGGATTTACATATACAACATATATTACTGTCAAGAGGGAATTTTTCTCAGTATTTAGATATTTAGATTCAAAATAAGAAAGGGATCAATTCAATTATAAACCTGCAAAACAAAGATTAGGATTGGGTTGGGTTGCGCTATATATATCAAAGAGTATACAATAATGATGTATTTGGTGAATCAAATACATGGTCTAATAACAAACCATTTTAACATAACATTTTAATGAGTTGATAATATTAATTGAATATTGAATATTTTTTGAAAGATTTTTGTTAAAGGTTTCATTCATGCCTAATCCATATCGAGTAGACCTTGTTGTTGTG??

			>Billbergia_brachysiphon
			??????????????????????????????????????????????????????????????????????????????????????????????????????????????????????????????????????????????????????????????????????????????????????????????????????????????????????????????????????????????????????????????????????????????????????????????????????????????????????????????????????????????????????????????????????????????????????????????????????????????????????????????????????????????????????????????????????????????????????????????????????????????????????????????????????????????????????????????????????????????????????????????????????????????????????????????????????????????????????????????????????????????????????????????????????????????????????????????????????????????????????????????????????????????????????????????

			>Billbergia_brasiliensis
			??????????????????????????????????????????????????????????????????????????????????????????????????????????????????????????????????????????????????????????????????????????????????????????????????????????????????????????????????????????????????????????????????????????????????????????????????????????????????????????????????????????????????????????????????????????????????????????????????????????????????????????????????????????????????????????????????????????????????????????????????????????????????????????????????????????????????????????????????????????????????????????????????????????????????????????????????????????????????????????????????????????????????????????????????????????????????????????????????????????????????????????????????????????????????????????????

			>Billbergia_chlorantha
			??????????????????????????????????????????????????????????????????????????????????????????????????????????????????????????????????????????????????????????????????????????????????????????????????????????????????????????????????????????????????????????????????????????????????????????????????????????????????????????????????????????????????????????????????????????????????????????????????????????????????????????????????????????????????????????????????????????????????????????????????????????????????????????????????????????????????????????????????????????????????????????????????????????????????????????????????????????????????????????????????????????????????????????????????????????????????????????????????????????????????????????????????????????????????????????????

			>Billbergia_decora
			ATAATTATAATAAAGTGAAATATGTCGAAATCCTTTTTGAAATAATACCAAATCGAAAATAAATGTCCGATAGCAAGTTGATCAGTTAATTCAATAAGAGATAAATGGGAGATAGCACTCGATTTAGTTGGTACCACCCAACCGAATCCGATTCACTCGTTTACTCATTCAATGAGTAAATTTTCAAGTTCAGCCAATCTTTTTTTTTTTAAAAAAGAAATTGCAAGTAGATGAAATCGTGAATAAATGTGTTTCATTTCTATATCATTATATAAAAAAAGATCCATATTATATTACTTATGGAATTCGAACCCGAACTCGATTTATGATTTATTATTTGGATCTTATGGGCCTTTGTTCTTTATTTTTGATTTTTCATATAGATTTCCGTCTTTATATTATACCCTTTCGTGGATGAATTATGCCTATTTTCACATCTAGGATTTACATATACAACATATATTACTGTCAAGAGGGAATTTTTCTCAGTATTTAGATATTTAGATTCAAAATAAGAAAGGGATCAATTCAATTATAAACCTGCAAAACAAAGATTAGGATTGGGTTGGGTTGCGCTATATATATCAAAGAGTATACAATAATGATGTATTTGGTGAATCAAATACATGGTCTAATAACAAACCATTTTAACATAACATTTTAATGAGTTGATAATATTAATTGAATATTGAATATTTTTTGAAAGATTTTTGTTAAAGGTTTCATTCATGCCTAATCCATATCGAGTAGACCTTGTTGTTGTG??

			>Billbergia_distachia
			??????????????????????????????????????????????????????????????????????????????????????????????????????????????????????????????????????????????????????????????????????????????????????????????????????????????????????????????????????????????????????????????????????????????????????????????????????????????????????????????????????????????????????????????????????????????????????????????????????????????????????????????????????????????????????????????????????????????????????????????????????????????????????????????????????????????????????????????????????????????????????????????????????????????????????????????????????????????????????????????????????????????????????????????????????????????????????????????????????????????????????????????????????????????????????????????

			>Billbergia_eloiseae
			??????????????????????????????????????????????????????????????????????????????????????????????????????????????????????????????????????????????????????????????????????????????????????????????????????????????????????????????????????????????????????????????????????????????????????????????????????????????????????????????????????????????????????????????????????????????????????????????????????????????????????????????????????????????????????????????????????????????????????????????????????????????????????????????????????????????????????????????????????????????????????????????????????????????????????????????????????????????????????????????????????????????????????????????????????????????????????????????????????????????????????????????????????????????????????????????

			>Billbergia_euphemiae
			??????????????????????????????????????????????????????????????????????????????????????????????????????????????????????????????????????????????????????????????????????????????????????????????????????????????????????????????????????????????????????????????????????????????????????????????????????????????????????????????????????????????????????????????????????????????????????????????????????????????????????????????????????????????????????????????????????????????????????????????????????????????????????????????????????????????????????????????????????????????????????????????????????????????????????????????????????????????????????????????????????????????????????????????????????????????????????????????????????????????????????????????????????????????????????????????

			>Billbergia_horrida
			??????????????????????????????????????????????????????????????????????????????????????????????????????????????????????????????????????????????????????????????????????????????????????????????????????????????????????????????????????????????????????????????????????????????????????????????????????????????????????????????????????????????????????????????????????????????????????????????????????????????????????????????????????????????????????????????????????????????????????????????????????????????????????????????????????????????????????????????????????????????????????????????????????????????????????????????????????????????????????????????????????????????????????????????????????????????????????????????????????????????????????????????????????????????????????????????

			>Billbergia_kautskyana
			??????????????????????????????????????????????????????????????????????????????????????????????????????????????????????????????????????????????????????????????????????????????????????????????????????????????????????????????????????????????????????????????????????????????????????????????????????????????????????????????????????????????????????????????????????????????????????????????????????????????????????????????????????????????????????????????????????????????????????????????????????????????????????????????????????????????????????????????????????????????????????????????????????????????????????????????????????????????????????????????????????????????????????????????????????????????????????????????????????????????????????????????????????????????????????????????

			>Billbergia_leptopoda
			??????????????????????????????????????????????????????????????????????????????????????????????????????????????????????????????????????????????????????????????????????????????????????????????????????????????????????????????????????????????????????????????????????????????????????????????????????????????????????????????????????????????????????????????????????????????????????????????????????????????????????????????????????????????????????????????????????????????????????????????????????????????????????????????????????????????????????????????????????????????????????????????????????????????????????????????????????????????????????????????????????????????????????????????????????????????????????????????????????????????????????????????????????????????????????????????

			>Billbergia_magnifica
			??????????????????????????????????????????????????????????????????????????????????????????????????????????????????????????????????????????????????????????????????????????????????????????????????????????????????????????????????????????????????????????????????????????????????????????????????????????????????????????????????????????????????????????????????????????????????????????????????????????????????????????????????????????????????????????????????????????????????????????????????????????????????????????????????????????????????????????????????????????????????????????????????????????????????????????????????????????????????????????????????????????????????????????????????????????????????????????????????????????????????????????????????????????????????????????????

			>Billbergia_manarae
			??????????????????????????????????????????????????????????????????????????????????????????????????????????????????????????????????????????????????????????????????????????????????????????????????????????????????????????????????????????????????????????????????????????????????????????????????????????????????????????????????????????????????????????????????????????????????????????????????????????????????????????????????????????????????????????????????????????????????????????????????????????????????????????????????????????????????????????????????????????????????????????????????????????????????????????????????????????????????????????????????????????????????????????????????????????????????????????????????????????????????????????????????????????????????????????????

			>Billbergia_meyeri
			??????????????????????????????????????????????????????????????????????????????????????????????????????????????????????????????????????????????????????????????????????????????????????????????????????????????????????????????????????????????????????????????????????????????????????????????????????????????????????????????????????????????????????????????????????????????????????????????????????????????????????????????????????????????????????????????????????????????????????????????????????????????????????????????????????????????????????????????????????????????????????????????????????????????????????????????????????????????????????????????????????????????????????????????????????????????????????????????????????????????????????????????????????????????????????????????

			>Billbergia_morelii
			??????????????????????????????????????????????????????????????????????????????????????????????????????????????????????????????????????????????????????????????????????????????????????????????????????????????????????????????????????????????????????????????????????????????????????????????????????????????????????????????????????????????????????????????????????????????????????????????????????????????????????????????????????????????????????????????????????????????????????????????????????????????????????????????????????????????????????????????????????????????????????????????????????????????????????????????????????????????????????????????????????????????????????????????????????????????????????????????????????????????????????????????????????????????????????????????

			>Billbergia_nana
			??????????????????????????????????????????????????????????????????????????????????????????????????????????????????????????????????????????????????????????????????????????????????????????????????????????????????????????????????????????????????????????????????????????????????????????????????????????????????????????????????????????????????????????????????????????????????????????????????????????????????????????????????????????????????????????????????????????????????????????????????????????????????????????????????????????????????????????????????????????????????????????????????????????????????????????????????????????????????????????????????????????????????????????????????????????????????????????????????????????????????????????????????????????????????????????????

			>Billbergia_nutans
			ATAATTATAATAAAGTGAAATATGTCGAAATCCTTTTTGGAATAATACCAAATCGAAAATAAATGTCCGATAGCAAGTTGATCAGTTAATTCAATAAGAGATAAATGGGAGATAGCACTCGATTTAGTTGGTACCACCCAACCGAATCCGATTCACTCGTTTACTCATTGAATGAGTAAATTTTCAAGTTCAGCCAATC--TTTTTTTTTAAAAAAGAAATTGCAAGTAGATGAAATCGTGAATAAATGTGTTTCATTTCTATATCATTATATAAAAAAAGATCCATATTATATTACTTATGGAATTCGAACCCGAACTCGATTTATGATTTATTATTTGGATCTTATTGGCCTTTGTTCTTTATTTTTGATTTTTCATATAGATTTCCGTCTTTATATTATACCCTTTCGTGGATGAATTATGCCTATTTTCACATCTAGGATTTACATATACAACATATATTACTGTCAAGAGGGAATTTTTCTCAGTATTTAGATATTTAGATTCAAAATAAGAAAGGGATCAATTCAATTATAAACCTGCAAAACAAAGATTAGGATTGGGTTGGGTTGCGCTATATATATCAAAGAGTATACAATAATGATGTATTTGGTGAATCAAATACATGGTCTAATAACAAACCATTTTAACATAACATTTTAATGAGTTGATAATATTAATTGAATATTGAATATTTTTTGAAAGATTTTTGTTAAAGGTTTCATTCATGCCTAATCCATATCGAGTAGACCTTGTTGTTGTG??

			>Billbergia_pallidiflora
			??????????????????????????????????????????????????????????????????????????????????????????????????????????????????????????????????????????????????????????????????????????????????????????????????????????????????????????????????????????????????????????????????????????????????????????????????????????????????????????????????????????????????????????????????????????????????????????????????????????????????????????????????????????????????????????????????????????????????????????????????????????????????????????????????????????????????????????????????????????????????????????????????????????????????????????????????????????????????????????????????????????????????????????????????????????????????????????????????????????????????????????????????????????????????????????????

			>Billbergia_porteana
			??????????????????????????????????????????????????????????????????????????????????????????????????????????????????????????????????????????????????????????????????????????????????????????????????????????????????????????????????????????????????????????????????????????????????????????????????????????????????????????????????????????????????????????????????????????????????????????????????????????????????????????????????????????????????????????????????????????????????????????????????????????????????????????????????????????????????????????????????????????????????????????????????????????????????????????????????????????????????????????????????????????????????????????????????????????????????????????????????????????????????????????????????????????????????????????????

			>Billbergia_pyramidalis
			??????????????????????????????????????????????????????????????????????????????????????????????????????????????????????????????????????????????????????????????????????????????????????????????????????????????????????????????????????????????????????????????????????????????????????????????????????????????????????????????????????????????????????????????????????????????????????????????????????????????????????????????????????????????????????????????????????????????????????????????????????????????????????????????????????????????????????????????????????????????????????????????????????????????????????????????????????????????????????????????????????????????????????????????????????????????????????????????????????????????????????????????????????????????????????????????

			>Billbergia_robert-readii
			??????????????????????????????????????????????????????????????????????????????????????????????????????????????????????????????????????????????????????????????????????????????????????????????????????????????????????????????????????????????????????????????????????????????????????????????????????????????????????????????????????????????????????????????????????????????????????????????????????????????????????????????????????????????????????????????????????????????????????????????????????????????????????????????????????????????????????????????????????????????????????????????????????????????????????????????????????????????????????????????????????????????????????????????????????????????????????????????????????????????????????????????????????????????????????????????

			>Billbergia_rosea
			??????????????????????????????????????????????????????????????????????????????????????????????????????????????????????????????????????????????????????????????????????????????????????????????????????????????????????????????????????????????????????????????????????????????????????????????????????????????????????????????????????????????????????????????????????????????????????????????????????????????????????????????????????????????????????????????????????????????????????????????????????????????????????????????????????????????????????????????????????????????????????????????????????????????????????????????????????????????????????????????????????????????????????????????????????????????????????????????????????????????????????????????????????????????????????????????

			>Billbergia_speciosa
			??????????????????????????????????????????????????????????????????????????????????????????????????????????????????????????????????????????????????????????????????????????????????????????????????????????????????????????????????????????????????????????????????????????????????????????????????????????????????????????????????????????????????????????????????????????????????????????????????????????????????????????????????????????????????????????????????????????????????????????????????????????????????????????????????????????????????????????????????????????????????????????????????????????????????????????????????????????????????????????????????????????????????????????????????????????????????????????????????????????????????????????????????????????????????????????????

			>Billbergia_stenopetala
			??????????????????????????????????????????????????????????????????????????????????????????????????????????????????????????????????????????????????????????????????????????????????????????????????????????????????????????????????????????????????????????????????????????????????????????????????????????????????????????????????????????????????????????????????????????????????????????????????????????????????????????????????????????????????????????????????????????????????????????????????????????????????????????????????????????????????????????????????????????????????????????????????????????????????????????????????????????????????????????????????????????????????????????????????????????????????????????????????????????????????????????????????????????????????????????????

			>Billbergia_tweedieana
			???????????????????????????????????TTTTGGATAATACCAAATCGAAAATAAATGTCCAATAGCAAGTTGATCAGTTAATTCAATAAGAGATAAATGGGAGATAGCACTCGATTTAGTTGGTACCACCCAACCGAATCCGATTCACTCGTTTACTCATTCAATGAGTAAATTTTCAAGTTCAGCCAATCTTTTTTTTTT--AAAAAAAAATTGCAAGTAGATGAAATCGTGAATAAATGTGTTTCATTTCTATATCATTATATAAAAAAAGATCCATATTATATTACTTATGGAATTCGAACCCGAACTCGATTTATGATTTATTATTTGGATCTTATTGGCCTTTGTTCTTTATTTTTGATTTTTAATATAGATTTCCGTCTTTATATTATACCCTTTCGGGGATGAATTATGCCTATTTTCACATCTAGGATTTACATATACAACATATATTACTGTCAAGAGGGAATTTTTCTCAGTATTTAGATATTTAGATTCAAAATAAGAAAGGGATCAATTCAATTATAAACCTGCAAAACAAAGATTAGGATTGGGTTGGGTTGCGCTATATATATCAAAGAGTATACAATAATGATGTATTTGGTGAATCAAATACATGGTCTAATAACAAACCATTTTAACATAACATTTTAATGAGTTGATAATATTAATTGAATATTGAATATTTTTTGAAAGATTTTTGTTAAAGGTTTCATTCATGCCTAATCCATATCGAGTAGACCTTGTTGTTGTG??

			>Billbergia_velascana
			??????????????????????????????????????????????????????????????????????????????????????????????????????????????????????????????????????????????????????????????????????????????????????????????????????????????????????????????????????????????????????????????????????????????????????????????????????????????????????????????????????????????????????????????????????????????????????????????????????????????????????????????????????????????????????????????????????????????????????????????????????????????????????????????????????????????????????????????????????????????????????????????????????????????????????????????????????????????????????????????????????????????????????????????????????????????????????????????????????????????????????????????????????????????????????????????

			>Billbergia_violacea
			??????????????????????????????????????????????????????????????????????????????????????????????????????????????????????????????????????????????????????????????????????????????????????????????????????????????????????????????????????????????????????????????????????????????????????????????????????????????????????????????????????????????????????????????????????????????????????????????????????????????????????????????????????????????????????????????????????????????????????????????????????????????????????????????????????????????????????????????????????????????????????????????????????????????????????????????????????????????????????????????????????????????????????????????????????????????????????????????????????????????????????????????????????????????????????????????

			>Billbergia_viridiflora
			??????????????????????????????????????????????????????????????????????????????????????????????????????????????????????????????????????????????????????????????????????????????????????????????????????????????????????????????????????????????????????????????????????????????????????????????????????????????????????????????????????????????????????????????????????????????????????????????????????????????????????????????????????????????????????????????????????????????????????????????????????????????????????????????????????????????????????????????????????????????????????????????????????????????????????????????????????????????????????????????????????????????????????????????????????????????????????????????????????????????????????????????????????????????????????????????

			>Billbergia_vittata
			??????????????????????????????????????????????????????????????????????????????????????????????????????????????????????????????????????????????????????????????????????????????????????????????????????????????????????????????????????????????????????????????????????????????????????????????????????????????????????????????????????????????????????????????????????????????????????????????????????????????????????????????????????????????????????????????????????????????????????????????????????????????????????????????????????????????????????????????????????????????????????????????????????????????????????????????????????????????????????????????????????????????????????????????????????????????????????????????????????????????????????????????????????????????????????????????

			>Billbergia_zebrina
			??????????????????????????????????????????????????????????????????????????????????????????????????????????????????????????????????????????????????????????????????????????????????????????????????????????????????????????????????????????????????????????????????????????????????????????????????????????????????????????????????????????????????????????????????????????????????????????????????????????????????????????????????????????????????????????????????????????????????????????????????????????????????????????????????????????????????????????????????????????????????????????????????????????????????????????????????????????????????????????????????????????????????????????????????????????????????????????????????????????????????????????????????????????????????????????????

			>Brewcaria_reflexa
			ATAATTATAATAAAGTGAAATATGTCGAAATCCTTTTTGGAATAATACCAAATCGAAAATAAATGTCCGATAGCAAGTTGATCAGTTAATTCAATAAGAGATAAATGGGAGATAGCACTCGATTTAGTTGGTACCACCCAACCGAATCCGATTCAATCGTTTACTCATTCAATGAGTCAATTTGCAAGTTCAGCCAATCTTTTTTTTTTCAAAAAAAAAATTGCAAGTAGATGAAATCGTGAATAAATGTGTTTCATTTCTCTATCATTATAGAAAATAAGATCCATATTATATTACTTATGGAATTCGAACCCGAACTCGATTTATGATTCATTATTTCGATCTTATTGGCCTTTGTTCGGTATTTTTGATTTTTCATATAGATTTCCGTCTTTATATTATACCCTTTCGTGGATGAATTATGCCTATTTTCACATCTAGGATTTACATATGCAACATATATTACTGTCAAGAGGGAATTTTCCTCAGTATTTAGATATTTAGATTCAAAATAAGAAAGGGATCAATTCAATTATAAACCTGCAAAACAAAGATTAGGA-----TTGGGTTGCGCTATATATATCAAAGAGTATACAATAATGATGTATTTGGTGAATCAAATACATGGTCTAATAACAAACCATTTTAACATAACATTTTAATGAGTTGATAATATTAATTGAATATTGAATATTTTTTGAAAGATTTTTGTTAAAGGTTTCATTCATGCCTAATCCATATCGAGTATACCTTGTTGTTGTGAG

			>Brocchinia_acuminata
			??????????????????????????????????????????????????????????????????????????????????CAGTTAATTCAATAAGAGATAAATGGGGGATAGCACTCGATTTAGTTGGTACCACCCAGCCGAATCCGATTCAATCGTTTACTCATTCAATCAGTCAATTTTCAAGTTCAGCCAATCTTTTTTTTTTCAAAAAAAAAATTGCAAGTAGATGAAATCGTGAATAAATGTGTTTCATTTCTCTATCATTATAGAAAATAAGATCCCTATTATATTACCTATGGAATTCCAACCCGAACTCGATTTATGATTCATTATTTCGATCTTATCGGCCTTTGTTCTTTATTTTTTATTTTTCATATAGATTTCCGTCTTTATATTATACCCCTTCGTGGATGAATTATGCCTATTTTCACATCTAGGATTTACATATACAACATATATTACTGTCAAGAGGGAATTTTTCTCAGTATTTAGATATTTAGATTCAAAATAAGAAAGGGATCAATTCAATTATAAACCCGCAAAACAAAGATTAGGATTGGATTGGGTTGCGCTATATATATCAAAGAGTATACAATAATGATGTATTTGGTGAATCAAATACATGGTCTAATAACAAACCATTTTAACATAACATTTTAATGAGTTGATAATATTAATTGAATATTGAATATCTTTTGAAAGATTTTTGTTAAAGGTTCCATTCATGCCTAATCCATATCGAGTAGACCTTGTTGTTGTGAG

			>Brocchinia_micrantha
			ATAATTATAATAAAGTGAAATATGTCGAAAT----ATTGGAATAATACCAAATCAAAAATAAATGTCCGATAGCAAGTTGATCAGTTAATTCAATAAGAGATAAATGGGGGATAGCACTCGATTTAGTTGGTACCACCCAGCCGAATCCGATTCAATCGTTTACTCATTCAATCAGTCAATTTTCAAGTTCAGCCAATCTTTTTTTTTTTAAAAAAAAAATTGCAAGTAGATGAAATCGTGAATAAATGTGTTTCATTTCTCTATCATTATAGAAAATAAGATCCCTATTATATTACCTATGGAATTCCAACCCGCACTCGATTTATGATTCATTATTTCGATCTTATCGGCCTTTGTTCTTTATTTTTGATTTTTCATATAGATTTCCGTCTTTATATTATACCCCTTCGTGGATGAATTATGCCTATTTTCACATCTAGGATTTACATATACAACATATATTACTGTTAAGAGGGAATTTTTCTCAGTATTTAGATATTTAGATTCAAAATAAGAAAGGGATCAATTCAATTATAAACCCGCAAAACAAAGATTAGGATTGGATTGGGTTGCGCTATATATATCAAAGAGTATACAATAATGATGTATTTGGTGAATCAAATACATGGTCTAATAACAAACCATTTTAACATAACATTTTAATGAGTTGATAATATTAATTGAATATTGAATATCTTTTGAAAGATTTTTGTTAAAGGTTCCATTCATGCCTAATCCATATCGAGTAGACCTTGTTGTTGTGAG

			>Brocchinia_paniculata
			??????????????????????????????????????????????????????????????????????????????????????????????????????????????????????????????????????????????????????????????????????????????????????????????????????????????????????????????????????????????????????????????????????????????????????????????????????????????????????????????????????????????????????????????????????????????????????????????????????????????????????????????????????????????????????????????????????????????????????????????????????????????????????????????????????????????????????????????????????????????????????????????????????????????????????????????????????????????????????????????????????????????????????????????????????????????????????????????????????????????????????????????????????????????????????????????

			>Brocchinia_prismatica
			ATAATTATAATAAAGTGAAATATGTCGAAAT----TTTGGAATAATACCAAATCAAAAATAAAGGTCCGATAGCAAGTTGATCAGTTAATTCAATAAGAGATAAATGGGAGATAGCACTCGATTTAGTTGGTACCACCCAGCCGAATTCGATTCAATGGTTTATTCATTCAATCAGTCAATTTTCAAGTTCAGCCAATC--TTTTTTTTTAAAAAAAAAATTGCAAGTAGATGAAATCGTGAATAAATGTGTTTCATTTCTCTATCATTATAGAAAATAAGATCCCTATTATATTACCTATGGAATTCGAACCCGAACTCGATTTATGATTCATTATTTCGATCTTATTGGCCTTTGTTCTTTATTTTTTATTTTTCATATAGATTTCCGTCTTTATATTATACCCCTTCGTGGATGAATTATGCCTATTTTCACATCTAGGATTTACATATACAACATATATTACTGTCAAGAGGGAATTTTTCTCAGTATTTAGATATTTAGATTCAAAATAAGAAAGGGATCAATTCAATTATAAACCTGCAAAACAAAGATTAGGATTGGATTGGGTTGCGCTATATATATCAAAGAGTATACAATAATGATGTATTTGGTGAATCAAATACATGGTCTAATAACAAACCATTTTAACATAACATTTTAATGAGTTGATAATATTAATTGAATATTGAATATCTTTTGAAAGATTTTTGTTAAAGGTTTCATTCATGCCTAATCCGTATCGAGTAGACCTTGTTGTTGTGAG

			>Brocchinia_reducta
			ATAATTATAATAAAGTGAAATATGTCGAAAT----TTTGGAATAATACCAAATCAAAAATAAATGTCCGATAGCAAGTTGATCAGTTAATTCAATAAGAGATAAATGGGAGATAGCACTCGATTTAGTTGGTACCACCCAGCCGAATCCGATTCAATCGTTTACTCATTCAATCAGTCAATTTTCAAGTTCAGCTAATCTTTTTTCTTTTAAAAAAA---TTGCAAGTAGATGAAATCGTGAATAAATGTGTTTCATTTCTCTATCATTATAGAAAAAAGGATCCCTATTATATTACCTATGGAATTCCAACCCGAACTCGATTTATGATTCATTATTTCGATCTTATTGGCCTTTGTTCTTTATTTTTTATTTTTCATATAGATTTCCGTCTTTATATTATACCCCTTCGTAGGTGAATTATGCCTATTTTCACATCTAGGATTTACATATACAACATATATTACTGTCAAGAGGGAATTTTTCTCAGTATTTAGATATTTAGATTCAAAATAAGAAAGGGATCAATTCAATTATAAACCCGCAAAACAAAGATTAGGATTGGATTGGGTTGCGCTATATATATCAAAGAGTACACAATAATGATGTATTTGGTGAATCAAATACATGGTCTAATAACAAACCATTTTAACATAACATTTTAATGAGTTGATAATATTAATTGAATATTGAATATCTTTTGAAAGATTTTTGTTAAAGGTTTCATTCATGCCTAATCCATATCGAGTAGACCTTGTTGTTGTGAG

			>Brocchinia_steyermarkii
			ATAATTATAATAAAGTGAAATATGTCGAAAT----ATTGGAATAATACCAAATCAAAAATAAATGTCCGATAGCAAGTTGATCAGTTAATTCAATAAGAGATAAATGGGGGATAGCACTCGATTTAGTTGGTACCACCCAGCCGAATCCGATTCAATCGTTTACTCATTCAATCAGTCAATTTTCAAGTTCAGCCAATCTTTTTTTTTTTAAAAAAAAAATTGCAAGTAGATGAAATCGTGAATAAATGTGTTTCATTTCTCTATCATTATAGAAAATAAGATCCCTATTATATTACCTATGGAATTCCAACCCGAACTCGATTTATGATTCATTATTTCGATCTTATCGGCCTTTGTTCTTTATTTTTGATTTTTCATATAGATTTCCGTCTTTATATTATACCCCTTCGTGGATGAATTATGCCTATTTTCACATCTAGGATTTACATATACAACATATATTACTGTTAAGAGGGAATTTTTCTCAGTATTTAGATATTTAGATTCAAAATAAGAAAGGGATCAATTCAATTATAAACCCGCAAAACAAAGATTAGGATTGGATTGGGTTGCGCTATATATATCAAAGAGTATACAATAATGATGTATTTGGTGAATCAAATACATGGTCTAATAACAAACCATTTTAACATAACATTTTAATGAGTTGATAATATTAATTGAATATTGAATATCTTTTGAAAGATTTTTGTTAAAGGTTCCATTCATGCCTAATCCATATCGAGTAGACCTTGTTGTTGTGAG

			>Brocchinia_tatei
			ATAATTATAATAAAGTGAAATATGTCGAAAT----TTTGGAATAATACCAAATCAAAAATAAATGTCCGATAGCAAGTTGATCAGTTAATTCAATAAGAGATAAATGGGAGATAGCACTCGATTTAGTTGGTACCACCCAGCCGAATCCGATTCAATCGTTTACTCATTCAATCAGTCAATTTTCAAGTTCAGCTAATCTTTTTTTTTTCAAAAAA----TTGCAAGTAGATGAAATCGTGAATAAATGTGTTTCATTTCTCTATCATTATAGAAAAAAAGATCCCTATTATATTACCTATGGAATTCCAACCCGAACTCGATTTATGATTCATTATTTCGATCTTATTGGCCTTTGTTCTTTATTTTTTATTTTTCATATAGATTTCCGTCTTTATATTATACCCCTTCGTGGATGAATTATGCCTATTTTCACATCTAGGATTTACATATACAACATATATCACTGTCAAGAGAGAATTTTTCTCAGTATTTAGATATTTAGATTCAAAATAAGAAAGGGATCAATTCAATTATAAACCCACAAAACAAAGATTAGGATTGGATTGGGTTGCGCTATATATATCAAAGAGTATACAATAATGATGTATTTGGTGAATCAAATACATGGTCTAATAACAAACCATTTTAACATAACATTTTAATGAGTTGATAATATTAATTGAATATTGAATATCTTTTGAAAGATTTTTGTTAAAGGTTTCATTCATGCCTAATCCATATCGAGTAGACCTTGTTGTTGTGAG

			>Brocchinia_uaipanensis
			??????????????????????????????????????????????????????????????????????????????????CAGTTAATTCAATAAGAGATAAATGGGAAATAGCACTCGATTTAGTTGGTACCACCCAGCCGAATCCGATTCAATCGTTTACTCATTCAATCAGTCAATTTTAAAGTTCAGCCAATCTTTTTTTTTTT------TTAATTGCAAGTAGATGAAATCGTGAATAAATGTGTTTCATTTCTCTATCATTATAGAAAATAAGATCCCTATTATATTACCTATGGAATTCGAACCCGAACTCGATTTATGATTCATTATTTCGATCTTATTGGCCTTTGTTCTTTATTTTTTATTTTTCATATAGATTTCCGTCTTTATATTATACCCCCTCGTGGATGAATTATGCCTATTTTCACATCTAGGATTTACATATACAACATATATTACTGTCAAGAGGGAATTTTTCTCAGTATTTAGATATTTAGATTCAAAATAAGAAAGGGATCAATTCAATTATAAACCTGCAAAACAAAGATTAGGATTGGATTGGGTTGCGCTATATATATCAAAGAGTATACAATAATGATGTATTTGGTGAATCAAATACATGGTCTAATAACAAACCATTTTAACATAACATTTTAATGAGTTGATAATATTAATTGAATATTGAATATCCTTTGAAAGATTTTTGTTAAAGGTTTCATTCATGCCTAATCCATATCGAGTAGACCTTGTTGTTGTGAG

			>Bromelia_agavifolia
			??????????????????????????????????????????????????????????????????????????????????????????????????????????????????????????????????????????????????????????????????????????????????????????????????????????????????????????????????????????????????????????????????????????????????????????????????????????????????????????????????????????????????????????????????????????????????????????????????????????????????????????????????????????????????????????????????????????????????????????????????????????????????????????????????????????????????????????????????????????????????????????????????????????????????????????????????????????????????????????????????????????????????????????????????????????????????????????????????????????????????????????????????????????????????????????????

			>Bromelia_antiacantha
			???????????????????????????????????TTTGGAATAATACCAAATC-AAAATAAATGTCCGATAGCAAGTTGATCAGTTAATTCAATAAGAGATAAATGGGAGATAGCACTCGATTTAGTTGGTACCACCCAACCGAATCCGATTCAATCGTTTACTCATTGAATGAGTAAATTTTCAAGTTCAGCCAATCTTTTTTTTTT-AAAAAAAAAATTGCAAGTAGATGAAATCGTGAATAAATGTGTTTCATTTCTCTATCATTATATAAAATAAGATCCATATTATATTACTTATGGAATTCGAACCCGAACTCGATTTATGATTTATTATTTCGATCTTATTGGCCTTTGTTCTTTATTTTTGATTTTTCATATAGATTTCCGTCTTTATATTATACCCTTTCGTGGATGAATTATGCCTATTTTCACATCTAGGATTTACATATACAACATATATTACTGTCAAGAGGGAATTTTTCTCAGTATTTATATATTTAGATTCAAAATAAGAAAGGGATCAATTCAATTATAAACCTGCAAAATAAAGATTAGGA-----TTGGGTTGCGCTATATATATCAAAGAGTATACAATAATGATGTATTTGGTGAATCAAATACATGGTCTAATAACAAACCATTTTAACATAACATTTTAATGAGTTGATAATATTAATTGAATATTGAATATTTTTTGAAAGATTTTTGTTAAAGGTTTCATTCATGCCTAATCCATATCGAGTAGACCTTGTTCTTGTG??

			>Bromelia_balansae
			??????????????????????????????????????????????????????????????????????????????????????????????????????????????????????????????????????????????????????????????????????????????????????????????????????????????????????????????????????????????????????????????????????????????????????????????????????????????????????????????????????????????????????????????????????????????????????????????????????????????????????????????????????????????????????????????????????????????????????????????????????????????????????????????????????????????????????????????????????????????????????????????????????????????????????????????????????????????????????????????????????????????????????????????????????????????????????????????????????????????????????????????????????????????????????????????

			>Bromelia_chrysantha
			??????????????????????????????????????????????????????????????????????????????????????????????????????????????????????????????????????????????????????????????????????????????????????????????????????????????????????????????????????????????????????????????????????????????????????????????????????????????????????????????????????????????????????????????????????????????????????????????????????????????????????????????????????????????????????????????????????????????????????????????????????????????????????????????????????????????????????????????????????????????????????????????????????????????????????????????????????????????????????????????????????????????????????????????????????????????????????????????????????????????????????????????????????????????????????????????

			>Bromelia_flemingii
			ATAATTATAATAAAGTGAAATACGTCGAAATCCTTTTTGGAATAATACCAAATC-AAAATAAATGTCCGATAGCAAGTTGATCAGTTAATTCAATAAGAGATAAATGGGAGATAGCACTCGATTTAGTTGGTACCACCCAACCGAATCCGATTCAATCGTTTACTCATTGAATGAGTAAATTTTCAAGTTCAGCCAATCTTTTTTTTTTTAAAAAAAAAATTGCAAGTAGATGAAATCGTGAATAAATGTGTTTCATTTCTCTATCATTATATAAAATAAGATCCATATTATATTACTTATGGAATTCGAACCCGAACTCGATTTATGATTTATTATTTCGATCTTATTGGCCTTTGTTCTTTATTTTTGATTTTTCATATAGATTTCCGTCTTTATATTATACCCTTTCGTGGATGAATTATGCCTATTTTCACATCTAGGATTTACATATACAACATATATTACTGTCAAGAGGGAATTTTTCTCAGTATTTATATATTTAGATTCAAAATAAGAAAGGGATCAATTCAATTATAAACCTGCAAAATAAAGATTAGGA-----TTGGGTTGCGCTATATATATCAAAGAGTATACAATAATGATGTATTTGGTGAATCAAATACATGGTCTAATAACAAACCATTTTAACATAACATTTTAATGAGTTGATAATATTAATTGAATATTGAATATTTTTTGAAAGATTTTTGTTAAAGGTTTCATTCATGCCTAATCCATATCGAGTAGACCTTGTTCTTGTGAG

			>Bromelia_hemisphaerica
			??????????????????????????????????????????????????????????????????????????????????????????????????????????????????????????????????????????????????????????????????????????????????????????????????????????????????????????????????????????????????????????????????????????????????????????????????????????????????????????????????????????????????????????????????????????????????????????????????????????????????????????????????????????????????????????????????????????????????????????????????????????????????????????????????????????????????????????????????????????????????????????????????????????????????????????????????????????????????????????????????????????????????????????????????????????????????????????????????????????????????????????????????????????????????????????????

			>Bromelia_irwinii
			???????????????????????????????????TTTGGAATAATACCAAATC-AAAATAAATGTCCGATAGCAAGTTGATCAGTTAATTCAATAAGAGATAAATGGGAGATAGCACTCGATTTAGTTGGTACCACCCAACCGAATCCGATTCAATCGTTTACTCATTGAATGAGTAAATTTTCAAGTTCAGCCAATCTTTTTTTTTT-AAAAAAAAAATTGCAAGTAGATGAAATCGTGAATAAATGTGTTTCATTTCTCTATCATTATATAAAATAAGATCCATATTATATTACTTATGGAATTCGAACCCGAACTCGATTTATGATTTATTATTTCGATCTTATTGGCCTTTGTTCTTTATTTTTGATTTTTCATATAGATTTCCGTCTTTATATTATACCCTTTCGTGGATGAATTATGCCTATTTTCACATCTAGGATTTACATATACAACATATATTACTGTCAAGAGGGAATTTTTCTCAGTATTTATATATTTAGATTCAAAATAAGAAAGGGATCAATTCAATTATAAACCTGCAAAATAAAGATTAGGA-----TTGGGTTGCGCTATATATATCAAAGAGTATACAATAATGATGTATTTGGTGAATCAAATACATGGTCTAATAACAAACCATTTTAACATAACATTTTAATGAGTTGATAATATTAATTGAATATTGAATATTTTTTGAAAGATTTTTGTTAAAGGTTTCATTCATGCCTAATCCATATCGAGTAGACCTTGTTCTTGTG??

			>Bromelia_karatas
			ATAATTATAATAAAGTGAAATACGTCGAAATCCTTTTTGGAATAATACCAAATC-AAAATAAATGTCCGATAGCAAGTTGATCAGTTAATTCAATAAGAGATAAATGGGAGATAGCACTCGATTTAGTTGGTACCACCCAACCGAATCCGATTCAATCGTTTACTCATTGAATGAGTAAATTTTCAAGTTCAGCCAATCTTTTTTTTTTTAAAAAAAAAATTGCAAGTAGATGAAATCGTGAATAAATGTGTTTCATTTCTCTATCATTATATAAAATAAGATCCATATTATATTACTTATGGAATTCGAACCCGAACTCGATTTATGATTTATTATTTCGATCTTATTGGCCTTTGTTCTTTATTTTTGATTTTTCATATAGATTTCCGTCTTTATATTATACCCTTTCGTGGATGAATTATGCCTATTTTCACATCTAGGATTTACATATACAACATATATTACTGTCAAGAGGGAATTTTTCTCAGTATTTATATATTTAGATTCAAAATAAGAAAGGGATCAATTCAATTATAAACCTGCAAAATAAAGATTAGGA-----TTGGGTTGCGCTATATATATCAAAGAGTATACAATAATGATGTATTTGGTGAATCAAATACATGGTCTAATAACAAACCATTTTAACATAACATTTTAATGAGTTGATAATATTAATTGAATATTGAATATTTTTTGAAAGATTTTTGTTAAAGGTTTCATTCATGCCTAATCCATATCGAGTAGACCTTGTTCTTGTGAG

			>Bromelia_pinguin
			??????????????????????????????????????????????????????????????????????????????????CAGTTAATTCAATAAGAGATAAATGGGAGATAGCACTCGATTTAGTTGGTACCACCCAACCGAATCCGATTCAATCGTTTACTCATTGAATGAGTAAATTTTCAAGTTCAGCCAATCTTTTTTTTTTTAAAAAAAAAATTGCAAGTAGATGAAATCGTGAATAAATGTGTTTCATTTCTCTATCATTATATAAAATAAGATCCATATTATATTACTTATGGAATTCGAACCCGAACTCGATTTATGATTTATTATTTCGATCTTATTGGCCTTTGTTCTTTATTTTTGATTTTTCATATAGATTTCCGTCTTTATATTATACCCTTTCGTGGATGAATTATGCCTATTTTCACATCTAGGATTTACATATACAACATATATTACTGTCAAGAGGGAATTTTTCTCAGTATTTATATATTTAGATTCAAAATAAGAAAGGGATCAATTCAATTATAAACCTGCAAAATAAAGATTAGGA-----TTGGGTTGCGCTATATATATCAAAGAGTATACAATAATGATGTATTTGGTGAATCAAATACATGGTCTAATAACAAACCATTTTAACATAACATTTTAATGAGTTGATAATATTAATTGAATATTGAATATTTTTTGAAAGATTTTTGTTAAAGGTTTCATTCATGCCTAATCCATATCGAGTAGACCTTGTTCTTGTGAG

			>Bromelia_serra
			?????????????????????????????????????????????????????????????????????????????????????????TTCAATAAGAGATAAATGGGAGATAGCACTCGATTTAGTTGGTACCACCCAACCGAATCCGATTCAATCGTTTACTCATTGAATGAGTAAATTTTCAAGTTCAGCCAATCTTTTTTTTTTTAAAAAAAAAATTGCAAGTAGATGAAATCGTGAATAAATGTGTTTCATTTCTCTATCATTATATAAAATAAGATCCATATTATATTACTTATGGAATTCGAACCCGAACTCGATTTATGATTTATTATTTCGATCTTATTGGCCTTTGTTCTTTATTTTTGATTTTTCATATAGATTTCCGTCTTTATATTATACCCTTTCGTGGATGAATTATGCCTATTTTCACATCTAGGATTTACATATACAACATATATTACTGTCAAGAGGGAATTTTTCTCAGTATTTATATATTTAGATTCAAAATAAGAAAGGGATCAATTCAATTATAAACCTGCAAAATAAAGATTAGGA-----TTGGGTTGCGCTATATATATCAAAGAGTATACAATAATGATGTATTTGGTGAATCAAATACATGGTCTAATAACAAACCATTTTAACATAACATTTTAATGAGTTGATAATATTAATTGAATATTGAATATTTTTTGAAAGATTTTTGTTAAAGGTTTCATTCATGCCTAATCCATATCGAGTAGACCTTGTTCTTGTGAG

			>Bromelia_trianae
			??????????????????????????????????????????????????????????????????????????????????????????????????????????????????????????????????????????????????????????????????????????????????????????????????????????????????????????????????????????????????????????????????????????????????????????????????????????????????????????????????????????????????????????????????????????????????????????????????????????????????????????????????????????????????????????????????????????????????????????????????????????????????????????????????????????????????????????????????????????????????????????????????????????????????????????????????????????????????????????????????????????????????????????????????????????????????????????????????????????????????????????????????????????????????????????????

			>Canistropsis_albiflora
			??????????????????????????????????????????????????????????????????????????????????????????????????????????????????????????????????????????????????????????????????????????????????????????????????????????????????????????????????????????????????????????????????????????????????????????????????????????????????????????????????????????????????????????????????????????????????????????????????????????????????????????????????????????????????????????????????????????????????????????????????????????????????????????????????????????????????????????????????????????????????????????????????????????????????????????????????????????????????????????????????????????????????????????????????????????????????????????????????????????????????????????????????????????????????????????????

			>Canistropsis_billbergioides
			ATAATTATAATAAAGTGAAATATGTCGAAATCCTTTTTGGAATAATACCAAATCGAAAATAAATGTCCGATAGCAAGTTGATCAGTTAATTCAATAAGAGATAAATGGGAGATAGCACTCGATTTAGTTGGTACCACCCAACCGAATCCGATTCACTCGTTTACTCATTCAATGAGTAAATTTTCAAGTTCAGCCAATCTTTTTTTTTTTCAAAAAAAAATTGCAAGTAGATGAAATCGCGAATAAATGTGTTTCATTTC--TA----TATATAAAAAAAGATCCATATTATATTACTTATGGAATTCGAACCCGAACTCGATTTATGATTTATTATTTGGATCTTATTGGCCTTTGTTCTTTATTTTTGATTTTTCATATAGATTTCCGTCTTTATATTATACCCTTTCGTGGATGAATTATGCCTATTTTCACATCTAGGATTTACATATACAACATATATTACTGTCAAGAGGGAATTTTTCTCAGTATTTAGATATTTTGATTCAAAATAAGAAAGGGATCAATTCAATTATAAACCTGCAAAACAAAGATTAGGATTGGGTTGGGTTGCGCTATATATATCAAAGAGTATACAATAATGATGTATTTGGTGAATCAAATACATGGTCTAATAACAAACCATTTTAACATAACATTTTAATGAGTTGATAATATTAATTGAATATTGAATATTTTTTGAAAGATTTTTGTTAAAGGTTTCATTCATGCCTAATCCATATCGAGTAGACCTTGTTGTTGTGAG

			>Canistropsis_burchellii
			??????????????????????????????????????????????????????????????????????????????????????????????????????????????????????????????????????????????????????????????????????????????????????????????????????????????????????????????????????????????????????????????????????????????????????????????????????????????????????????????????????????????????????????????????????????????????????????????????????????????????????????????????????????????????????????????????????????????????????????????????????????????????????????????????????????????????????????????????????????????????????????????????????????????????????????????????????????????????????????????????????????????????????????????????????????????????????????????????????????????????????????????????????????????????????????????

			>Canistropsis_correia-araujoi
			??????????????????????????????????????????????????????????????????????????????????????????????????????????????????????????????????????????????????????????????????????????????????????????????????????????????????????????????????????????????????????????????????????????????????????????????????????????????????????????????????????????????????????????????????????????????????????????????????????????????????????????????????????????????????????????????????????????????????????????????????????????????????????????????????????????????????????????????????????????????????????????????????????????????????????????????????????????????????????????????????????????????????????????????????????????????????????????????????????????????????????????????????????????????????????????????

			>Canistropsis_elata
			??????????????????????????????????????????????????????????????????????????????????????????????????????????????????????????????????????????????????????????????????????????????????????????????????????????????????????????????????????????????????????????????????????????????????????????????????????????????????????????????????????????????????????????????????????????????????????????????????????????????????????????????????????????????????????????????????????????????????????????????????????????????????????????????????????????????????????????????????????????????????????????????????????????????????????????????????????????????????????????????????????????????????????????????????????????????????????????????????????????????????????????????????????????????????????????????

			>Canistropsis_exigua
			??????????????????????????????????????????????????????????????????????????????????????????????????????????????????????????????????????????????????????????????????????????????????????????????????????????????????????????????????????????????????????????????????????????????????????????????????????????????????????????????????????????????????????????????????????????????????????????????????????????????????????????????????????????????????????????????????????????????????????????????????????????????????????????????????????????????????????????????????????????????????????????????????????????????????????????????????????????????????????????????????????????????????????????????????????????????????????????????????????????????????????????????????????????????????????????????

			>Canistropsis_marceloi
			??????????????????????????????????????????????????????????????????????????????????????????????????????????????????????????????????????????????????????????????????????????????????????????????????????????????????????????????????????????????????????????????????????????????????????????????????????????????????????????????????????????????????????????????????????????????????????????????????????????????????????????????????????????????????????????????????????????????????????????????????????????????????????????????????????????????????????????????????????????????????????????????????????????????????????????????????????????????????????????????????????????????????????????????????????????????????????????????????????????????????????????????????????????????????????????????

			>Canistropsis_microps
			???????????????????????????????????TTTGGAATAATACCAAATCGAAAATAAATGTCCGATAGCAAGTTGATCAGTTAATTCAATAAGAGATAAATGGGAGATAGCACTCGATTTAGTTGGTACCACCCAACCGAATCCGATTCACTCGTTTACTCATTCAATGAGTAAATTTTCAAGTTCAGCCAATC-TTTTTTTTTTACAAAAAAAATTGCAAGTAGATGAAATCGCGAATAAATGTGTTTCATTTCTATATCATTATATAAAAAAAGATCCATATTATATTACTTATGGAATTCGAACCCGAACTCGATTTATGATTTATTATTTGGATCTTATTGGCCTTTGTTCTTTATTTTTGATTTTTCATATAGATTTCCGTCTTTATATTATACCCTTTCGTGGATGAATTATGCCTATTTTCACATCTAGGATTTACATATACAACATATATTACTGTCAAGAGGGAATTTTTCTCAGTATTTAGATATTTAGATTCAAAATAAGAAAGGGATCAATTCAATTATAAACCTGCAAAACAAAGATTAGGATTGGGTTGGGTTGCGCTATATATATCAAAGAGTATACAATAATGATGTATTTGGTGAATCAAATACATGGTCTAATAACAAACCATTTTAACATAACATTTTAATGAGTTGATAATATTAATTGAATATTGAATATTTTTTGAAAGATTTTTGTTAAAGGTTTCATTCATGCCTAATCCATATCGAGTAGACCTTGTTGTTGTG??

			>Canistropsis_pulcherrima
			??????????????????????????????????????????????????????????????????????????????????????????????????????????????????????????????????????????????????????????????????????????????????????????????????????????????????????????????????????????????????????????????????????????????????????????????????????????????????????????????????????????????????????????????????????????????????????????????????????????????????????????????????????????????????????????????????????????????????????????????????????????????????????????????????????????????????????????????????????????????????????????????????????????????????????????????????????????????????????????????????????????????????????????????????????????????????????????????????????????????????????????????????????????????????????????????

			>Canistropsis_simulans
			??????????????????????????????????????????????????????????????????????????????????????????????????????????????????????????????????????????????????????????????????????????????????????????????????????????????????????????????????????????????????????????????????????????????????????????????????????????????????????????????????????????????????????????????????????????????????????????????????????????????????????????????????????????????????????????????????????????????????????????????????????????????????????????????????????????????????????????????????????????????????????????????????????????????????????????????????????????????????????????????????????????????????????????????????????????????????????????????????????????????????????????????????????????????????????????????

			>Canistrum_alagoanum
			??????????????????????????????????????????????????????????????????????????????????????????????????????????????????????????????????????????????????????????????????????????????????????????????????????????????????????????????????????????????????????????????????????????????????????????????????????????????????????????????????????????????????????????????????????????????????????????????????????????????????????????????????????????????????????????????????????????????????????????????????????????????????????????????????????????????????????????????????????????????????????????????????????????????????????????????????????????????????????????????????????????????????????????????????????????????????????????????????????????????????????????????????????????????????????????????

			>Canistrum_aurantiacum
			ATAATTATAATAAAGTGAAATATGTCGAAATCCTTTTTGGAATAATACCAAATCGAAAATAAATGTCCGATAGCAAGTTGATCAGTTAATTCAATAAGAGATAAATGGGAGATAGCACTCGATTTAGTTGGTACCACCCAACCGAATCCGATTCACTCGTTTACTCATTCAATGAGTAAATTTTCAAGTTCAGCCAATC--TTTTTTTTTAAAAAAAAAATTGCAAGTAGATGAAATCGTGAATAAATGTGTTTCATTTCTATATCATTATATAAAAAAAGATCCATATTTTATTACTTATGGAATTCGAACCCGAACTCGATTTATGATTTATTATTTGGATCTTATTGGCCTTTGTTCTTTATTTTTGATTTTTCATATAGATTTCCGTCTTTATATTATACCCTTTCGTGGATGAATTATGCCTATTTTCACATCTAGGATTTACATATACAACATATATTACTGTCAAGAGGGAATTTTTCTCAGTATTTAGATATTTAGATTCAAAATAAGAAAGGGATCAATTCAATTATAAACCTGCAAAACAAAGATTAGGATTGGGTTGGGTTGCGCTATATATATCAAAGAGTATACAATAATGATGTATTTGGTGAATCAAATACATGGTCTAATAACAAACCATTTTAACATAACATTTTAACGAGTTGATAATATTAATTGAATATTGAATATTTTTTGAAAGATTTTTGTTAAAGGTTTCATTCATGCCTAATCCATATCGAGTAGACCTTGTTATTGTGAG

			>Canistrum_camacaense
			??????????????????????TGTCGA---TCCTTTTTGGATAATACCAAATCGAAAATAAATGTCCGATAGCAAGTTGATCAGTTAATTCAATAAGAGATAAATGGGAGATAGCACTCGATTTAGTTGGTACCACCCAACCGAATCCGATTCACTCATTTACTCATTGAATGAGTAAATTTTCAAGTTCAGCCAATCTTTTTTTTTTTAAAAAAAAAATTGCAAGTAGATGAAATCGTGAATAAATGTGTTTCATTTCTATATCATTATATAAAAAAAGATCCATATTATATTACTTATGGAATTCGAACCCGAACTCGATTTATGATTTATTATTTGGATCTTATTGGCCTTTGTTCTTTATTTTTGATTTTTCATATAGATTTCCGTCTTTATATTATACCCTTTCGTGGATGAATTATGCCTATTTTCACATCTAGGATTTACATATACAACATATATTACTGTCAAGAGGGAATTTTTCTCAGTATTTAGATATTTAGATTCAAAATAAGAAAGGGATCAATTCAATTATAAACCTGCAAAACAAAGATTAGGATTGGGTTGGGTTGCGCTATATATATCAAAGAGTATACAATAATGATGTATTTGGTGAATCAAATACATGGTCTAATAACAAACCATTTTAACATAACATTTTAATGAGTTGATAATATTAATTGAATATTGAATATTTTTTGAAAGATTTTTGTTAAAGGTTTCATTCATGCCTAATCCATATCGAGTAGACCTTGTTGTTGTG??

			>Canistrum_fosterianum
			ATAATTATAATAAAGTGAAATATGTCGAAATCCTTTTTGGAATAATACCAAATCGAAAATAAATGTCCGATAGCAAGTTGATCAGTTAATTCAATAAGAGATAAATGGGAGATAGCACTCGATTTAGTTGGTACCACCCAACCGAATCCGATTCACTCGTTTACTCATTCAATGAGTAAATTTTCAAGTTCAGCCAATCTTTTTTTTTTTAAAAAAAAGATTGCAAGTAGATGAAATCGTGAATAAATGTGTTTCATTTCTATATCATTATATAAAAAAAGATCCATATTATATTACTTATGGAATTCGAACCCGAACTCGATTTATGATTTATTATTTGGATCTTATTGGCCTTTGCTCTTTATTTTTTATTTTTCATATAGATTTCCATCTTTATATTATACCCTTTCGTGGATGAATTATGCCTATTTTCACATCTAGGATTTACATATACAACATATATTACTGTCAAGAGGGAATTTTTCTCAGTATTTAGATATTTAGATTCAAAATAAGAAAGGGATCAATTCAATTATAAACCTGCAAAACAAAGATTAGGATTGGGTTGGGTTGCGCTATATATATCAAAGAGTATACAATAATGATGTATTTGGTGAATCAAATACATGGTCTAATAACAAACCATTTTAACATAACATTTTAATGAGTTGATAATATTAATTGAATATTGAATATTTTTTGAAAGATTTTTGTTAAAGGTTTCATTCATGCCTAATCCATATCGAGTAGACCTTGTTGTTGTG??

			>Canistrum_pickelii
			??????????????????????????????????????????????????????????????????????????????????????????????????????????????????????????????????????????????????????????????????????????????????????????????????????????????????????????????????????????????????????????????????????????????????????????????????????????????????????????????????????????????????????????????????????????????????????????????????????????????????????????????????????????????????????????????????????????????????????????????????????????????????????????????????????????????????????????????????????????????????????????????????????????????????????????????????????????????????????????????????????????????????????????????????????????????????????????????????????????????????????????????????????????????????????????????

			>Canistrum_triangulare
			??????????????????????????????????????????????????????????????????????????????????????????????????????????????????????????????????????????????????????????????????????????????????????????????????????????????????????????????????????????????????????????????????????????????????????????????????????????????????????????????????????????????????????????????????????????????????????????????????????????????????????????????????????????????????????????????????????????????????????????????????????????????????????????????????????????????????????????????????????????????????????????????????????????????????????????????????????????????????????????????????????????????????????????????????????????????????????????????????????????????????????????????????????????????????????????????

			>Catopsis_berteroniana
			??????????????????????????????????????????????????????????????????????????????????????????????????????????????????????????????????????????????????????????????????????????????????????????????????????????????????????????????????????????????????????????????????????????????????????????????????????????????????????????????????????????????????????????????????????????????????????????????????????????????????????????????????????????????????????????????????????????????????????????????????????????????????????????????????????????????????????????????????????????????????????????????????????????????????????????????????????????????????????????????????????????????????????????????????????????????????????????????????????????????????????????????????????????????????????????????

			>Catopsis_floribunda
			??????????????????????????????????????????????????????????????????????????????????CAGTTAATTCAATAAGAGATAAATGGGAGATAGCACTCGATTTAGTTGGTACCACCCAACCGAATCCGATTCAATCGTTTACTCATTCAATGAGTCAATTTTCAAGTTCAGTCAATCCTTTTTTTTTTCAAAAAAAAATTGCAAGTAGATGAAATCGTGAATAAATGTGTTTTATTTCTCTATCATTATAGAAAATAAGATACATATTATATTACTTATGGAATTCGAACCCGAACTCGATTTATGATTCATTATTTCGATCTTATTGGCCTTTGTTCTTTATTTTTGATTTTTCATATAGATTTCCGTCTTTATATTATACCCTTTCGTGGATGAATTATGCTTATTTTCACATCTAGGATTTACATATACAACATATATTACTGTCAAGAGGGAATTTTTCTCAGTATTTAGATATTTAGATTCAAAATAAGAAAGGGATCAATTCAATTATAAACCTGCAAAGCAAAGATTAGGATTGGGTTGGGTTGCGCTATATATATCAAAGAGTATACAATAATGATGTATTTGGTGAATCAAATACATGGTCTAATAACAAACCATTTTAACATAACATTTTAATGAGTTGATAATATTAATTGAATATTGAATATTTTTTGAAAGATTTTTGTTAAAGGTTTCATTCATGCTTAATCCATATCGAGTAGACCTTGTTGTTGTGAG

			>Catopsis_juncifolia
			ATAATTATAATAAAGTGAAATATGTCGAAATCCTTTTTGGAATAATACCAAATCAAAAATAAATGTCCGATAGCAAGTTGATCAGTTAATTCAATAAGAGATAAATGGGAGATAGCACTCGATTTAGTTGGTACCACCCAACCGAATCCGATTCAATCGTTTACTCATTCAATGAGTCAATTTTCAAGTTCAGTCAACCCTTTTTTTTTGAAAAAAAAAATTGCAAGTAGATGAAATCGTGAATAAATGTGTTTCATTTCTCTATCATTATAGAAAATAAGATACATATTATATTACTTATGGAATTCGAACCCGAACTCGATTTATGATTCATTATTTCGATCTTATTGGCCTTTGTTCTTTATTTTTTATTTTTCATATAGATTTCCGTCTTTATATTATACCCTTTCGTGGATGAATTATGCTTATTTTCACATCTAGGATTTACATATACAACATATATTACTGTCAAGAGGGAATTTTTCTCAGTATTTAGATATTTAGATTCAAAATAAGAAAGGGATCAATTCAATTATAAACCTGCAAAGCAAAGATTAGGATTGGGTTGGGTTGCGCTATATATATCAAAGAGTATACAATAATGATGTATTTGGTGAATCAAATACATGGTCTAATAACAAACCATTTTAACATAACATTTTAATGAGTTGATAATATTAATCGAATATTGAATATTTTTTGAAAGATTTTTGTTAAAGGTTTCATTCATGCTTAATCTATATCGAGTAGACCTTGTTGTTGTGAG

			>Catopsis_morreniana
			ATAATTATAATAAAGTGAAATATGTCGAAATCCTTTTTGGAATAATACCAAATCAAAAATAAATGTCCGATAGCAAGTTGATCAGTTAATTCAATAAGAGATAAATGGGAGATAGCACTCGATTTAGTTGGTACCACCCAACCGAATCCGATTCAATCGTTTACTCATTCAATGAGTCAATTTTCAAGTTCAGTCAATCCTTTTTTTTT-CAAAAAAAAATTGCAAGTAGATGAAATCGTGAATAAATGTGTTTTATTTCTCTATCATTATAGAAAATAAGATACATATTATATTACTTATGGAATTCGAACCCGAACTCGATTTATGATTCATTATTTCGATCTTATTGGCCTTTGTTCTTTATTTTTGATTTTTCATATAGATTTCCGTCTTTATATTATACCCTTTCGTGGATGAATTATGCTTATTTTCACATCTAGGATTTACATATACAACATATATTACTGTCAAGAGGGAATTTTTCTCAGTATTTAGATATTTAGATTCAAAATAAGAAAGGGATCAATTCAATTATAAACCTGCAAAGCAAAGATTAGGATTGGGTTGGGTTGCGCTATATATATCAAAGAGTATACAATAATGATGTATTTGGTGAATCAAATACATGGTCTAATAACAAACCATTTTAACATAACATTTTAATGAGTTGATAATATTAATTGAATATTGAATATTTTTTGAAAGATTTTTGTTAAAGGTTTCATTCATGCTTAATCCATATCGAGTAGACCTTGTTGTTGTGAG

			>Catopsis_nitida
			??????????????????????????????????????????????????????????????????????????????????????????????????????????????????????????????????????????????????????????????????????????????????????????????????????????????????????????????????????????????????????????????????????????????????????????????????????????????????????????????????????????????????????????????????????????????????????????????????????????????????????????????????????????????????????????????????????????????????????????????????????????????????????????????????????????????????????????????????????????????????????????????????????????????????????????????????????????????????????????????????????????????????????????????????????????????????????????????????????????????????????????????????????????????????????????????

			>Catopsis_nutans
			ATAATTATAATAAAGTGAAATATGTCGAAATCCTTTTTGGAATAATACCAAATCAAAAATAAATGTCCGATAGCAAGTTGATCAGTTAATTCAATAAGAGATAAATGGGAGATAGCACTCGATTTAGTTGGTACCACCCAACCGAATCCGATTCAATCGTTTACTCATTCAATGAGTCAATTTTCAAGTTCAGTCAATCCTTTTTTTTTTAAAAAAAAAATTGCAAGTAGATGAAATCGTGAATAAATGTGTTTCATTTCTCTATCATTATAGAAAATAAGATACATATTATATTACTTATGGAATTCGAACCCGAACTCGATTTATGATTCATTATTTCGATCTTATTGGCCTTTGTTCTTTATTTTTGATTTTTCATATAGATTTCCGTCTTTATATTATACCCTTTCGTGGATGAATTATGCTTATTTTCACATCTAGGATTTACATATACAACATATATTACTGTCAAGAGGGAATTTTTCTCAG--------TATTTAGATTCAAAATAAGAAAGGGATCAATTCAATTATAAACCTGCAAAGCAAAGATTAGGATTGGGTTGGGTTGCGCTATATATATCAAAGAGTATACAATAATGATGTATTTGGTGAATCAAATACATGGTCTAATAACAAACCATTTTAACATAACATTTTAATGAGTTGATAATATTAATTGAATATTGAATATTTTTTGAAAGATTTTTGTTAAAGGTTTCATTCATGCTTAATCTATATCGAGTAGACCTTGTTGTTGTGAG

			>Catopsis_nutans_var._nutans
			??????????????????????????????????????????????????????????????????????????????????????????????????????????????????????????????????????????????????????????????????????????????????????????????????????????????????????????????????????????????????????????????????????????????????????????????????????????????????????????????????????????????????????????????????????????????????????????????????????????????????????????????????????????????????????????????????????????????????????????????????????????????????????????????????????????????????????????????????????????????????????????????????????????????????????????????????????????????????????????????????????????????????????????????????????????????????????????????????????????????????????????????????????????????????????????????

			>Catopsis_paniculata
			??????????????????????????????????????????????????????????????????????????????????????????????????????????????????????????????????????????????????????????????????????????????????????????????????????????????????????????????????????????????????????????????????????????????????????????????????????????????????????????????????????????????????????????????????????????????????????????????????????????????????????????????????????????????????????????????????????????????????????????????????????????????????????????????????????????????????????????????????????????????????????????????????????????????????????????????????????????????????????????????????????????????????????????????????????????????????????????????????????????????????????????????????????????????????????????????

			>Catopsis_sessiliflora
			??????????????????????????????????????????????????????????????????????????????????????????????????????????????????????????????????????????????????????????????????????????????????????????????????????????????????????????????????????????????????????????????????????????????????????????????????????????????????????????????????????????????????????????????????????????????????????????????????????????????????????????????????????????????????????????????????????????????????????????????????????????????????????????????????????????????????????????????????????????????????????????????????????????????????????????????????????????????????????????????????????????????????????????????????????????????????????????????????????????????????????????????????????????????????????????????

			>Catopsis_subulata
			ATAATTATAATAAAGTGAAATATGTCGAAATCCTTTTTGGAATAATACCAAATCAAAAATAAATGTCCGATAGCAAGTTGATCAGTTAATTCAATAAGAGATAAATGGGAGATAGCACTCGATTTAGTTGGTACCACCCAACCGAATCCGATTCAATCGTTTACTCATTCAATGAGTCAATTTTCAAGTTCAGTCAACCCCCTTTTTTTTCAAAAAAAAATTGCAAGTAGATGAAATCGTGAATAAATGTGTTTCATTTCTCTATCATTATAGAAAATAAGATACATATTATATTACTTATGGAATTCGAACCCGAACTCGATTTATGATTCATTATTTCGATCTTATTGGCCTTTGCTCTTTATTTTTTATTTTTCATATAGATTTCCGTCTTTATATTATACCCTTTCGTGGATGAATTATGCTTATTTTCACATCTAGGATTTACATATACAACATATATTACTGTCAAGAGGGAATTTTTCTCAGTATTTAGATATTTAGATTCAAAATAAGAAAGGGATCAATTCAATTATAAACCTGCAAAGCAAAGATTAGGATTGGGTTGGGTTGCGCTATATATATCAAAGAGTATACAATAATGATGTATTTGGTGAATCAAATACATGGTCTAATAACAAACCATTTTAACATAACATTTTAATGAGTTGATAATATTAATTGAATATTGAATATTTTTTGAAAGATTTTTGTTAAAGGTTTCATTCATGCTTAATCTATATCGAGTAGACCTTGTTGTTGTGAG

			>Catopsis_wangerinii
			??????????????????????????????????????????????????????????????????????????????????????????????????????????????????????????????????????????????????????????????????????????????????????????????????????????????????????????????????????????????????????????????????????????????????????????????????????????????????????????????????????????????????????????????????????????????????????????????????????????????????????????????????????????????????????????????????????????????????????????????????????????????????????????????????????????????????????????????????????????????????????????????????????????????????????????????????????????????????????????????????????????????????????????????????????????????????????????????????????????????????????????????????????????????????????????????

			>Cipuropsis_amicorum
			??????????????????????????????????????????????????????????????????????????????????????????????????????????????????????????????????????????????????????????????????????????????????????????????????????????????????????????????????????????????????????????????????????????????????????????????????????????????????????????????????????????????????????????????????????????????????????????????????????????????????????????????????????????????????????????????????????????????????????????????????????????????????????????????????????????????????????????????????????????????????????????????????????????????????????????????????????????????????????????????????????????????????????????????????????????????????????????????????????????????????????????????????????????????????????????????

			>Connellia_nutans
			??????????????????????????????????????????????????????????????????????????????????????????????????????????????????????????????????????????????????????????????????????????????????????????????????????????????????????????????????????????????????????????????????????????????????????????????????????????????????????????????????????????????????????????????????????????????????????????????????????????????????????????????????????????????????????????????????????????????????????????????????????????????????????????????????????????????????????????????????????????????????????????????????????????????????????????????????????????????????????????????????????????????????????????????????????????????????????????????????????????????????????????????????????????????????????????????

			>Cottendorfia_florida
			ATAATTATAATAAAGTGAAATATGTCGAAATCCTTTTTGGAATAATAACAAATCGAAAATAAATGTCCGATAGCAAGTTGATCAGTTAATTCAATAAGAGATAAATGGGAGATAGCACTCGATTTAGTTGGTACCACCCAACCGAATCCGATTCAATCGTTTACTCATTCAATGAGTCAATTTTCAAGTTCAGCCAATC--TTTTTTTTTAAAAAAAAAATTGTAAGTAGATGAAATCGTGAATAAATGTGATTCATTTCTCTATCATTATAGAAAATAAGATACGTATTATATTACTTATGGAATTCTAACCCGAACTCGATTTATGATTCATTATTTCGATCTTATTGGCCTTTGTTCTTTATTTTTTATTTTTCATATAGATTTCCGTCTTTATATTATACCCTTTCGTGGATGAATTATGCCTATTTTCACATCTAGGATTTACATATACAACATATATTACTGTCAAAAGGGAATTTTCCTCAGTATTTAGATATTTAGATTCAAAATAAGAAAGGGATCAATTCAATTATAAACCTGCAAAACAAAGATTAGGA-----TTGGGTTGCGCTATATATATCAAAGAGTATACAATAATGATGTATTTGGTGAATAAAATACATGGTCTAATAACAAACCATTTTAACATAACATTTTAATGAGTTGATAATATTAATTGAATATTGAATATTTTTTGAAAGATTTTTGTTAAAGGTTTCATTCATGCCTAATCCATATCGAGTAGACCTTGTTGTTGTGAG

			>Cryptanthus_acaulis
			??????????????????????????????????????????????????????????????????????????????????????????????????????????????????????????????????????????????????????????????????????????????????????????????????????????????????????????????????????????????????????????????????????????????????????????????????????????????????????????????????????????????????????????????????????????????????????????????????????????????????????????????????????????????????????????????????????????????????????????????????????????????????????????????????????????????????????????????????????????????????????????????????????????????????????????????????????????????????????????????????????????????????????????????????????????????????????????????????????????????????????????????????????????????????????????????

			>Cryptanthus_bahianus
			ATAATTTTAATAAAGTGAAATATGTCGAAATCCTTTTTGGAATAATACCAAATCGAAAATAAATGTCCGATAGCAAGTTGATCAGTTAATTCAATAAGAGATAAATGGGAGATAGCACTCGATTTAGTTGGTACCACCCAACCGAATCCGATTCACTCGTTTACTCATTGAATGAGTCAATTTTCGAGTTCAGCCAATC-CTTTT----------------------------------------------------------------------------------------------------------------------------------------------------------------------TTTTTCATATAGATTTCCGTCTTTATATTATACCCTTTCGTGGATGAATTATGCCTATTTTCACATCTAGGATTTACATATACAACATATATTACTGTCAAGAGGGAATTTTTCTCAG--------TATTTAGATTCAAAATAAGAAAGGGATCAATTCAATTATAAACCTGCAAAACAAAGATTAGGATTGGGTTGGGTTGCGCTATATATATCAAAGAGTATACAATAATGATGTATTTGGTGAATCAAATACATGGTCTAATAACAAACCATTTTAACATAACATTTTAATGAGTTGATAATATTAATTGAATATTGAATATTTTTTGAAAGATTTTTGTTAAAGGTTTCATTCATGCCTAATCCATATCGAGTAGACCTTGTTGTTGTG??

			>Cryptanthus_beuckeri
			ATAATTTTAATAAAGCGAAATATGTCGAAATCCTTTTTGGAATAATACCAAATCGAAAATAAATGTCCGATAGCAGGTTGATCAGTTAATTCAATAAGAGATAAATGGG-GATAGCACTCGATTTAGTTGGTACCACCCAACCGAATCCGATTCACTCGTTTACTCATTCAATGAGTAAATTTTCAA---------------------------------TT---------------AATGAGTAAATTTT---------------------------------------------------------AAGTTCAGACA----------------------ATCTT---------------------T---TTTTTCATATAGATTTCCGTCTTTATATTATACCCTTTCGTGGATGAATTATGCCTATTTTCACATCTAGGATTTACATATACAACATATATTACTGTCAAGAGGGAATTTTTCTCAG--------TATTTAGATTCAAAATAAGAAAGGGATCAATTCAATTATAAACCTGCAAAACAAAGATTAGGATTGGGTTGGGTTGCGCTATATATATCAAAGAGTATACAATAATGATGTATTTGGTGAATCAAATACATGGTCTAATAACAAACCATTTTAACATAACATTTTAATGAGTTGATAATATTAATTGAATATTGAATATTTTTTGAAAGATTTTTGTTAAAGGTTTCATTCATGCCTAATCCATATCGAGTAGACCTTGTTGTTGTGAG

			>Cryptanthus_bromelioides
			??????????????????????????????????????????????????????????????????????????????????????????????????????????????????????????????????????????????????????????????????????????????????????????????????????????????????????????????????????????????????????????????????????????????????????????????????????????????????????????????????????????????????????????????????????????????????????????????????????????????????????????????????????????????????????????????????????????????????????????????????????????????????????????????????????????????????????????????????????????????????????????????????????????????????????????????????????????????????????????????????????????????????????????????????????????????????????????????????????????????????????????????????????????????????????????????

			>Cryptanthus_burle-marxii
			??????????????????????????????????????????????????????????????????????????????????????????????????????????????????????????????????????????????????????????????????????????????????????????????????????????????????????????????????????????????????????????????????????????????????????????????????????????????????????????????????????????????????????????????????????????????????????????????????????????????????????????????????????????????????????????????????????????????????????????????????????????????????????????????????????????????????????????????????????????????????????????????????????????????????????????????????????????????????????????????????????????????????????????????????????????????????????????????????????????????????????????????????????????????????????????????

			>Cryptanthus_colnagoi
			???????????????????????????????????TTTGGAATAATACCAAATCGAAAATAAATGTCCGATAGCAAGTTGATCAGTTAATTCAATAAGAGATAAATGGG-GATAGCACTCGATTTAGTTGGTACCACCCAACCGAATCCGATTCACTCGTTTACTCATTCAATGAGTAAATTTTCAAGTTCAGCCCATCTTTTTT----------------------------------------------------------------------------------------------------------------------------------------------------------------------TTTTTCATATAGATTTCCGTCTTTATATTATACCCTTTCGTGGATGAATTATGCCTATTTTCACATCTAGGATTTACATATACAACATATATTACTGTCAAGAGGGAATTTTTCTCAG--------TATTTAGATTCAAAATAAGAAAGGGATCAATTCAATTATAAAC?TGCAAAACAAAGATTAGGATTGGGTTGGGTTGCGCTATATATATCAAAGAGTATACAATAATGATGTATTTGGTGAATCAAATACATGGTCTAATAACAAACCATTTTAACATAACATTTTAATGAGTTGATAATATTAATTGAATATTGAATATTTTTTGAAAGATTTTTGTTAAAGGTTTCATTCATGC?TAATCCATATCGAGTAGAC?TTGTTGTTGTG??

			>Cryptanthus_diamantinensis
			???????????????????????????????????TTTGGAATAATACCAAATCGAAAATAAATGTCCGATAGCAAGTTGATCAGTTAATTCAATAAGAGATAAATGGG-GATAGCACTCGATTTAGTTGGTACCACCCAACCGAATCCGATTCACTCGTTTACTCATTCAATGAGTAAATTTTCAAGTTCAGCCAATC--TTTT----------------------------------------------------------------------------------------------------------------------------------------------------------------------TTTTTCATATAGATTTCCGTCTTTATATTATACCCTTTCGTGGATGAATTATGCCTATTTTCACATCTAGGATTTACATATACAACATATATTACTGTCAAGAGGGAATTTTTCTCAG--------TATTTAGATTCAAAATAAGAAAGGGATCAATTCAATTATAAACCTGCAAAACAAAGATTAGGATTGGGTTGGGTTGCGCTATATATATCAAAGAGTATACAATAATGATGTATTTGGTGAATCAAATACATGGTCTAATAACAAACCATTTTAACATAACATTTTAATGAGTTGATAATATTAATTGAATATTGAATATTTTTTGAAAGATTTTTGTTAAAGGTTTCATTCATGCCTAATCCATATCGAGTAGACCTTGTTGTTGTG??

			>Cryptanthus_dianae
			??????????????????????????????????????????????????????????????????????????????????????????????????????????????????????????????????????????????????????????????????????????????????????????????????????????????????????????????????????????????????????????????????????????????????????????????????????????????????????????????????????????????????????????????????????????????????????????????????????????????????????????????????????????????????????????????????????????????????????????????????????????????????????????????????????????????????????????????????????????????????????????????????????????????????????????????????????????????????????????????????????????????????????????????????????????????????????????????????????????????????????????????????????????????????????????????

			>Cryptanthus_maritimus
			??????????????????????????????????????????????????????????????????????????????????????????????????????????????????????????????????????????????????????????????????????????????????????????????????????????????????????????????????????????????????????????????????????????????????????????????????????????????????????????????????????????????????????????????????????????????????????????????????????????????????????????????????????????????????????????????????????????????????????????????????????????????????????????????????????????????????????????????????????????????????????????????????????????????????????????????????????????????????????????????????????????????????????????????????????????????????????????????????????????????????????????????????????????????????????????????

			>Cryptanthus_ruthae
			??????????????????????????????????????????????????????????????????????????????????????????????????????????????????????????????????????????????????????????????????????????????????????????????????????????????????????????????????????????????????????????????????????????????????????????????????????????????????????????????????????????????????????????????????????????????????????????????????????????????????????????????????????????????????????????????????????????????????????????????????????????????????????????????????????????????????????????????????????????????????????????????????????????????????????????????????????????????????????????????????????????????????????????????????????????????????????????????????????????????????????????????????????????????????????????????

			>Cryptanthus_teretifolius
			??????????????????????????????????????????????????????????????????????????????????????????????????????????????????????????????????????????????????????????????????????????????????????????????????????????????????????????????????????????????????????????????????????????????????????????????????????????????????????????????????????????????????????????????????????????????????????????????????????????????????????????????????????????????????????????????????????????????????????????????????????????????????????????????????????????????????????????????????????????????????????????????????????????????????????????????????????????????????????????????????????????????????????????????????????????????????????????????????????????????????????????????????????????????????????????????

			>Cryptanthus_warren-loosei
			????????????????????????????AATCCT-TTTTGGATAATACCAAATCGAAAATAAATGTCCRATAGCAAGTTGATCAGTTAATTCAATAAGAGATAAATGGGAGATAGCACTCGATTTAGTTGGTACCACCCAACCGAATCCGATTCACTCGTTTACTCATTCAATGAGTAAATTTTCAAGTTCAGCCAATCTTTTTTTTTTTAAAAAAAAAATTGCAAGTAGATGAAATCGTGAATAAATGTGTTTCATTTTTATATCATTATATAAAAAAAGATCCATATTATATTACTTATGGAATTCGAACCCGAACTCGATTTATGATTTATTATTTGGATCTTATTGGCCTTTGTTCTTTATTTTTGATTTTTAATATAGATTTCCGTCTTTATATTATACCCTTTCGTGGATGAATTATGCCTATTTTCACATCTAGGATTTACATATACAACATATATTACTGTCAAGAGGGAATTTTTCTCAGTATTTAGATATTTAGATTCAAAATAAGAAAGGGATCAATTCAATTATAAACCTGCAAAACAAAGATTAGGATTGGGTTGGGTTGCGCTATATATATCAAAGAGTATACAATAATGATGTATTTGGTGAATCAAATACATGGTCTAATAACAAACCATTTTAACATAACATTTTAATGAGTTGATAATATTAATTGAATATTGAATATTTTTTGAAAGATTTTTGTTAAAGGTTTCATTCATGCCTAATCCATATCGAGTAGACCTTGTTGTTGTG??

			>Cryptanthus_zonatus
			??????????????????????????????????????????????????????????????????????????????????????????????????????????????????????????????????????????????????????????????????????????????????????????????????????????????????????????????????????????????????????????????????????????????????????????????????????????????????????????????????????????????????????????????????????????????????????????????????????????????????????????????????????????????????????????????????????????????????????????????????????????????????????????????????????????????????????????????????????????????????????????????????????????????????????????????????????????????????????????????????????????????????????????????????????????????????????????????????????????????????????????????????????????????????????????????

			>Deinacanthon_urbanianum
			??????????????????????????????????????????????????????????????????????????????????CAGTTAATTCAATAAGAGATAAATGGGAGATAGCACTCGATTTAGTTGGTACCACCCAACCGAATCCGATTCAATCGTTTACTCATTCAATGAGTAAATTTTTAAGTTCAGCCAATCTTTTTTTTTTTAAAAAAAAAATTGCAAGTAGATGAAATCGTGAATAAATGTGTTTCATTTCTCTATCATTATATAAAATAAGATCCATATTATATTACTTAGGGAATTCGAACCCGAACTCGATTTATGATTTATTATTTCGATCTTATTGGCCTTTGTTCTTTATTTTTGATTTTTCATATGGATTCCCGTCTTTATATTATACCCTTTCGTGGATGAATTATGCCTATTTTCACATCTAGGATTTACATATACAACATATATTACTGTCAAGAGGGAATTTTTCTCAGTATTTAGATATTTAGATTCAAAATAAGAAAGGGATCAATTCAATTATAAACCCGCAAAACAAAGATTAGGATTGGGTTGGGTTGCGCTATATATATCAAAGAGTATACAATAATGATGTATTTGGTGAATCAAATACATGGTCTAATAACAAACCATTTTAACATAACATTTTAATGAGTTGATAATATTAATTGAATATTGAATATTTTTTGAAAGATTTTTGTTAAAGGTTTCATTCATGCCTAATCCATATCGAGTAGACCTTGTTGTTGTGAG

			>Deuterocohnia_abstrusa
			??????????????????????????????????????????????????????????????????????????????????????????????????????????????????????????????????????????????????????????????????????????????????????????????????????????????????????????????????????????????????????????????????????????????????????????????????????????????????????????????????????????????????????????????????????????????????????????????????????????????????????????????????????????????????????????????????????????????????????????????????????????????????????????????????????????????????????????????????????????????????????????????????????????????????????????????????????????????????????????????????????????????????????????????????????????????????????????????????????????????????????????????????????????????????????????????

			>Deuterocohnia_brevifolia
			??????????????????????????????????????????????????????????????????????????????????CAGTTAATTCAATAAGAGATAAATGGGAGATAGCACTCGGTTTAGTTGGTACCACCCAACCGAATCCGATTCAATCGTTTACTCATTGAATGAGTAAATTTTCAAGTTCAACCAATCTTTTTTTTTTTAAAAAAAAAATTGCAAGTAGATGAAATCGTGAATAAATGTGTTTCATTTCTCTATCATTATAGAAAATAAGATCCATATTATATTACTTATGGAATTCGAACCCGAACTCGATTTATGATTCATTATTTCGATCTTATTGGCCTTTGTTCTTTATTTTTTATTTTTCATATAGATTTCCGTCTTTATATTATACCCTTTCGTGGATGAATTATGCCTCTTTTCACATCTAGGATTTACATATACAACATATATTACTGTCAAGAGGGAATTTTTCTCAGTATTTAGATATTTAGATTCAAAATAAGAAAGGGATCAATTCAATTATAAACCTGTAAAACAAAGATTAGGATTGGGTTGGGTTGCGCTATATATATCAAAGAGTATACAATAATGATGTATTTGGTGAATCAAATACATGGTCTAATAACAAACCATTTTAACATAACATTTTAATGAGTTGATAATATTAATTGAATATTGAATATTTTTTGAAAGATTTTTGTTAAAGGTTTCATTCATGCCTAATCCATATCGAGTAGACCTTGTTGTTGTGAG

			>Deuterocohnia_brevispicata
			??????????????????????????????????????????????????????????????????????????????????CGGTTAATTCAATAAGAGATAAATGGAAGATAGCACTCGGTTTAGTTGGTACCACCCAACCGAATCCGATTCAATCGTTTACTCATTGAATGAGTCAATTTTCAAGTTCAGCCAATCTTTTTTTTTG-AAAAAATAAATTGCAAGTAGATGAAATCGTGAATAAATGTGTTTCATTTCTCTATCATTATAGAAAATAAGATCCATATTATATTACTTATGGAATTCGAACCCGAACTCGATTTATGATTCATTATTTCGATCTTATTGGCCTTTGTTCTTTATTTTTGATTTTTCATATGGATTTCCGTCTTTATATTATACCCTTTCGTGGATGAATTATGCCTCTTTTCACATCTAGGATTTACATATACAACATATATTACTGTCAAGAGGGAATTTTTCTCAGTATTTAGATATTTAGATTCAAAATAAGAAAGGGATCAATTCAATTATAAACCTGTAAAACAAAGATTAGGATTGGGTTGGGTTGCGCTATATATATCAAAGAGTATACAATAATGATGTATTTGGTGAATCAAATACATGGTCTAATAACAAACCATTTTAACATAACATTTTAATGAGTTGATAATATTAATTGAATATTGAATATTTTTTGAAAGATTTTTGTTAAAGGTTTCATTCATGCCTAATCCATATCGAGTAGACCTTGTTGTTGTGAG

			>Deuterocohnia_chrysantha
			??????????????????????????????????????????????????????????????????????????????????????????????????????????????????????????????????????????????????????????????????????????????????????????????????????????????????????????????????????????????????????????????????????????????????????????????????????????????????????????????????????????????????????????????????????????????????????????????????????????????????????????????????????????????????????????????????????????????????????????????????????????????????????????????????????????????????????????????????????????????????????????????????????????????????????????????????????????????????????????????????????????????????????????????????????????????????????????????????????????????????????????????????????????????????????????????

			>Deuterocohnia_digitata
			??????????????????????????????????????????????????????????????????????????????????????????????????????????????????????????????????????????????????????????????????????????????????????????????????????????????????????????????????????????????????????????????????????????????????????????????????????????????????????????????????????????????????????????????????????????????????????????????????????????????????????????????????????????????????????????????????????????????????????????????????????????????????????????????????????????????????????????????????????????????????????????????????????????????????????????????????????????????????????????????????????????????????????????????????????????????????????????????????????????????????????????????????????????????????????????????

			>Deuterocohnia_gableana
			??????????????????????????????????????????????????????????????????????????????????????????????????????????????????????????????????????????????????????????????????????????????????????????????????????????????????????????????????????????????????????????????????????????????????????????????????????????????????????????????????????????????????????????????????????????????????????????????????????????????????????????????????????????????????????????????????????????????????????????????????????????????????????????????????????????????????????????????????????????????????????????????????????????????????????????????????????????????????????????????????????????????????????????????????????????????????????????????????????????????????????????????????????????????????????????????

			>Deuterocohnia_glandulosa
			??????????????????????????????????????????????????????????????????????????????????CAGTTAATTCAATAAGAGATAAATGGGAGATAGCACTCGGTTTAGTTGGTACCACCCAACCGAATCCGATTCAATCGTTTACTCATTGAATGAGTCAATTTTCAAGTTCAGCCAATCTTTTTTTTTTGAAAAAAAAAATTGCAAGTAGATGAAATCGTGAATAAATGTGTTTCATTTCTCTATCATTATAGAAAATAAGATCCATATTATATTACTTATGGAATTCGAACCCGAACTCGATTTATGATTCATTATTTCGATCTTCTTGGCCTTTGTTCTTTATTTTTGATTTTTCATATAGATTTCCGTCTTTATATTATACCCTTTCGTGGATGAATTATGCCTCTTTTCACATCTAGGATTTACATATACAACATATATTACTGTCAAGAGGGAATTTTTCTCAGTATTTAGATATTTAGATTCAAAATAAGAAAGGGATCAATTCAATTATAAACCTGTAAAACAAGGATTAGGA-----TTGGGTTGCGCTATATATATCAAAGAGTATACAATAATGATGTATTTGGTGAATCAAATACATGGTCTAATAACAAACCATTTTAACATAACATTTTAATGAGTTGATAATATTAATTGAATATTGAATATTTTTTGAAAGATTTTTGTTAAAGGTTTCATTCATGCCTAATCCATATCGAGTAGACCTTGTTGTTGTGAG

			>Deuterocohnia_haumanii
			??????????????????????????????????????????????????????????????????????????????????????????????????????????????????????????????????????????????????????????????????????????????????????????????????????????????????????????????????????????????????????????????????????????????????????????????????????????????????????????????????????????????????????????????????????????????????????????????????????????????????????????????????????????????????????????????????????????????????????????????????????????????????????????????????????????????????????????????????????????????????????????????????????????????????????????????????????????????????????????????????????????????????????????????????????????????????????????????????????????????????????????????????????????????????????????????

			>Deuterocohnia_longipetala
			ATAATTATAATAAAGTTAAATATGTCGAAATCCTTTTTGGAATAATACCAAATCTAAAATAAATGTCCGATAGCAAGTTGATCAGTTAATTCAATAAGAGATAAATGGGAGATAGCACTCGGTTTAGTTGGTACCACCCAACCGAATCCGATTCAATCGTTTACTCATTGAATGAGTCAATTTTCAAGTTCAGCCAATCTTTTTTTTTTTAAA----AAATTGCAAGTAGATGAAATCGTGAATAAATGTGTTTCATTTCTCTATCATTATAGAAAATAAGATCCATATTATATTACTTATGGAATTCGAACCCGAACTCGATTTATGATTCATTATTTCGATCTTATTGGCCTTTGTTCTTTATTTTTGATTTTTCATATAGATTTCCGTCTTTATATTATACCCTTTCGTGGATGAATTATGCCTCTTTTCACATCTAGGATTTACATATACAACATATATTACTGTCAAGAGGGAATTTTTCTCAGTATTTAGATATTTAGATTCAAAATAAGAAAGGGATCAATTCAATTATAAACCTGTAAAACAAAGATTAGGATTGGGTTGGGTTGCGCTATATATATCAAAGAGTATACAATAATGATGTATTTGGTGAATCAAATACATGGTCTAATAACAAACCATTTTAACATAACATTTTAATGAGTTGATAATATTAATTGAATATTGAATATTTTTTGAAAGATTTTTGTTAAAGGTTTCATTCATGCCTAATCCATATCGAGTAGACCTTGTTGTTGTGAG

			>Deuterocohnia_lotteae
			??????????????????????????????????????????????????????????????????????????????????CAGTTAATTCAATAAGAGATAAATGGGAGATAGCACTCGGTTTAGTTGGTACCACCCAACCGAATCCGATTCAATCGTTTACTCATTGAATGAGTCAATTTTCAAGTTCAGCCAATCTTTTTTTTTTGAAAAAAAAAATTGCAAGTAGATGAAATCGTGAATAAATGTGTTTCATTTCTCTATCATTATAGAAAATAAGATCCATATTATATTACTTATGGAATTCGAACCCGAACTCGATTTATGATTCATTATTTCGATCTTCTTGGCCTTTGTTCTTTATTTTTGATTTTTCATATAGATTTCCGTCTTTATATTATACCCTTTCGTGGATGAATTATGCCTCTTTTCACATCTAGGATTTACATATACAACATATATTACTGTCAAGAGGGAATTTTTCTCAGTATTTAGATATTTAGATTCAAAATAAGAAAGGGATCAATTCAATTATAAACCTGTAAAACAAAGATTAGGA-----TTGGGTTGCGCTATATATATCAAAGAGTATACAATAATGATGTATTTGGTGAATCAAATACATGGTCTAATAACAAACCATTTTAACATAACATTTTAATGAGTTGATAATATTAATTGAATATTGAATATTTTTTGAAAGATTTTTGTTAAAGGTTTCATTCATGCCTAATCCATAACGAGTAGACCTTGTTGTTGTGAG

			>Deuterocohnia_meziana
			??????????????????????????????????????????????????????????????????????????????????????????????????????????????????????????????????????????????????????????????????????????????????????????????????????????????????????????????????????????????????????????????????????????????????????????????????????????????????????????????????????????????????????????????????????????????????????????????????????????????????????????????????????????????????????????????????????????????????????????????????????????????????????????????????????????????????????????????????????????????????????????????????????????????????????????????????????????????????????????????????????????????????????????????????????????????????????????????????????????????????????????????????????????????????????????????

			>Deuterocohnia_meziana_subsp._carmineoviridiflora
			??????????????????????????????????????????????????????????????????????????????????????????????????????????????????????????????????????????????????????????????????????????????????????????????????????????????????????????????????????????????????????????????????????????????????????????????????????????????????????????????????????????????????????????????????????????????????????????????????????????????????????????????????????????????????????????????????????????????????????????????????????????????????????????????????????????????????????????????????????????????????????????????????????????????????????????????????????????????????????????????????????????????????????????????????????????????????????????????????????????????????????????????????????????????????????????????

			>Deuterocohnia_meziana_subsp._pedicellata
			??????????????????????????????????????????????????????????????????????????????????????????????????????????????????????????????????????????????????????????????????????????????????????????????????????????????????????????????????????????????????????????????????????????????????????????????????????????????????????????????????????????????????????????????????????????????????????????????????????????????????????????????????????????????????????????????????????????????????????????????????????????????????????????????????????????????????????????????????????????????????????????????????????????????????????????????????????????????????????????????????????????????????????????????????????????????????????????????????????????????????????????????????????????????????????????????

			>Deuterocohnia_recurvipetala
			??????????????????????????????????????????????????????????????????????????????????????????????????????????????????????????????????????????????????????????????????????????????????????????????????????????????????????????????????????????????????????????????????????????????????????????????????????????????????????????????????????????????????????????????????????????????????????????????????????????????????????????????????????????????????????????????????????????????????????????????????????????????????????????????????????????????????????????????????????????????????????????????????????????????????????????????????????????????????????????????????????????????????????????????????????????????????????????????????????????????????????????????????????????????????????????????

			>Deuterocohnia_scapigera
			??????????????????????????????????????????????????????????????????????????????????CGGTTAATTCAATAAGAGATAAATGGAAGATAGCACTCGGTTTAGTTGGTACCACCCAACCGAATCCGATTCAATCGTTTACTCATTGAATGAGTCAATTTTCAAGTTCAGCCAATCTTTTTTTTTG-AAAAAATAAATTGCAAGTAGATGAAATCGTGAATAAATGTGTTTCATTTCTCTATCATTATAGAAAATAAGATCCATATTATATTACTTATGGAATTCGAACCCGAACTCGATTTATGATTCATTATTTCGATCTTATTGGCCTTTGTTCTTTATTTTTGATTTTTCATATGGATTTCCGTCTTTATATTATACCCTTTCGTGGATGAATTATGCCTCTTTTCACATCTAGGATTTACATATACAACATATATTACTGTCAAGAGGGAATTTTTCTCAGTATTTAGATATTTAGATTCAAAATAAGAAAGGGATCAATTCAATTATAAACCTGTAAAACAAAGATTAGGATTGGGTTGGGTTGCGCTATATATATCAAAGAGTATACAATAATGATGTATTTGGTGAATCAAATACATGGTCTAATAACAAACCATTTTAACATAACATTTTAATGAGTTGATAATATTAATTGAATATTGAATATTTTTTGAAAGATTTTTGTTAAAGGTTTCATTCATGCCTAATCCATATCGAGTAGACCTTGTTGTTGTGAG

			>Deuterocohnia_schreiteri
			??????????????????????????????????????????????????????????????????????????????????????????????????????????????????????????????????????????????????????????????????????????????????????????????????????????????????????????????????????????????????????????????????????????????????????????????????????????????????????????????????????????????????????????????????????????????????????????????????????????????????????????????????????????????????????????????????????????????????????????????????????????????????????????????????????????????????????????????????????????????????????????????????????????????????????????????????????????????????????????????????????????????????????????????????????????????????????????????????????????????????????????????????????????????????????????????

			>Deuterocohnia_seramisiana
			??????????????????????????????????????????????????????????????????????????????????????????????????????????????????????????????????????????????????????????????????????????????????????????????????????????????????????????????????????????????????????????????????????????????????????????????????????????????????????????????????????????????????????????????????????????????????????????????????????????????????????????????????????????????????????????????????????????????????????????????????????????????????????????????????????????????????????????????????????????????????????????????????????????????????????????????????????????????????????????????????????????????????????????????????????????????????????????????????????????????????????????????????????????????????????????????

			>Deuterocohnia_strobilifera_var._inermis
			??????????????????????????????????????????????????????????????????????????????????????????????????????????????????????????????????????????????????????????????????????????????????????????????????????????????????????????????????????????????????????????????????????????????????????????????????????????????????????????????????????????????????????????????????????????????????????????????????????????????????????????????????????????????????????????????????????????????????????????????????????????????????????????????????????????????????????????????????????????????????????????????????????????????????????????????????????????????????????????????????????????????????????????????????????????????????????????????????????????????????????????????????????????????????????????????

			>Deuterocohnia_strobilifera_var._strobilifera
			??????????????????????????????????????????????????????????????????????????????????????????????????????????????????????????????????????????????????????????????????????????????????????????????????????????????????????????????????????????????????????????????????????????????????????????????????????????????????????????????????????????????????????????????????????????????????????????????????????????????????????????????????????????????????????????????????????????????????????????????????????????????????????????????????????????????????????????????????????????????????????????????????????????????????????????????????????????????????????????????????????????????????????????????????????????????????????????????????????????????????????????????????????????????????????????????

			>Disteganthus_basilateralis
			????????????????????????????????????TTTGGATAATACCAAATCGAAAATAAATGTCCGATAGCAAGTTGATCAGTTAATTCAATAAGAGATAAATGGGAGATAGCACTCGATTTAGTTGGTACCACCCAACCGAATCCGATTCACTCGTTTACTCATTCAATGAGTAAATTTTCAAGTTCAGCCAATCTTTTTTTTTTC-AAAAAAAAATTGCAAGTAGATGAAATCGTGAATAAATGTGTTTCATTTCTCTATCATTATATAAAAAAAGATCCATATTATATTACTTATGGAATTCGAACCCGAACTCGATTTATGATTTATTATTTGGATCTTATTGGCCTTTGTTCTTTATTTTGGATTTTTCATATAGATTTCCGTCTTTATATTATACCCTTTCGTGGATGAATTATGCCTATTTTCACATCTAGGATTTACATATACAACATATATTACTGTCAAGGGGGAATTTTTCTCAGTATTTAGATATTTAGATTCCAAATAAGAAAGGGATCAATTCAATTATAAACCTGCAAAACAAAGATTAGGATTGGGTTGGGTTGCGCTATATATATCAAAGAGTATACAATAATGATGTATTTGGTGAATCAAATACATGGTCTAATAACAAACCGTTTTAACATAACATTTTAATGAGTTGATAATATTAATTGAATATTGAATATTTTTTGAAAGATTTTTGTTAAAGGTTTCATTCATGCCTAATCCATATCGAGTAGACCTTGTTGTTGTG??

			>Disteganthus_lateralis
			??????????????????????????????????????????????????????????????????????????????????????????????????????????????????????????????????????????????????????????????????????????????????????????????????????????????????????????????????????????????????????????????????????????????????????????????????????????????????????????????????????????????????????????????????????????????????????????????????????????????????????????????????????????????????????????????????????????????????????????????????????????????????????????????????????????????????????????????????????????????????????????????????????????????????????????????????????????????????????????????????????????????????????????????????????????????????????????????????????????????????????????????????????????????????????????????

			>Dyckia_acutiflora
			??????????????????????????????????????????????????????????????????????????????????????????????????????????????????????????????????????????????????????????????????????????????????????????????????????????????????????????????????????????????????????????????????????????????????????????????????????????????????????????????????????????????????????????????????????????????????????????????????????????????????????????????????????????????????????????????????????????????????????????????????????????????????????????????????????????????????????????????????????????????????????????????????????????????????????????????????????????????????????????????????????????????????????????????????????????????????????????????????????????????????????????????????????????????????????????????

			>Dyckia_areniticola
			??????????????????????????????????????????????????????????????????????????????????????????????????????????????????????????????????????????????????????????????????????????????????????????????????????????????????????????????????????????????????????????????????????????????????????????????????????????????????????????????????????????????????????????????????????????????????????????????????????????????????????????????????????????????????????????????????????????????????????????????????????????????????????????????????????????????????????????????????????????????????????????????????????????????????????????????????????????????????????????????????????????????????????????????????????????????????????????????????????????????????????????????????????????????????????????????

			>Dyckia_aurea
			??????????????????????????????????????????????????????????????????????????????????????????????????????????????????????????????????????????????????????????????????????????????????????????????????????????????????????????????????????????????????????????????????????????????????????????????????????????????????????????????????????????????????????????????????????????????????????????????????????????????????????????????????????????????????????????????????????????????????????????????????????????????????????????????????????????????????????????????????????????????????????????????????????????????????????????????????????????????????????????????????????????????????????????????????????????????????????????????????????????????????????????????????????????????????????????????

			>Dyckia_beateae
			??????????????????????????????????????????????????????????????????????????????????????????????????????????????????????????????????????????????????????????????????????????????????????????????????????????????????????????????????????????????????????????????????????????????????????????????????????????????????????????????????????????????????????????????????????????????????????????????????????????????????????????????????????????????????????????????????????????????????????????????????????????????????????????????????????????????????????????????????????????????????????????????????????????????????????????????????????????????????????????????????????????????????????????????????????????????????????????????????????????????????????????????????????????????????????????????

			>Dyckia_brachyphylla
			??????????????????????????????????????????????????????????????????????????????????????????????????????????????????????????????????????????????????????????????????????????????????????????????????????????????????????????????????????????????????????????????????????????????????????????????????????????????????????????????????????????????????????????????????????????????????????????????????????????????????????????????????????????????????????????????????????????????????????????????????????????????????????????????????????????????????????????????????????????????????????????????????????????????????????????????????????????????????????????????????????????????????????????????????????????????????????????????????????????????????????????????????????????????????????????????

			>Dyckia_bracteata
			??????????????????????????????????????????????????????????????????????????????????????????????????????????????????????????????????????????????????????????????????????????????????????????????????????????????????????????????????????????????????????????????????????????????????????????????????????????????????????????????????????????????????????????????????????????????????????????????????????????????????????????????????????????????????????????????????????????????????????????????????????????????????????????????????????????????????????????????????????????????????????????????????????????????????????????????????????????????????????????????????????????????????????????????????????????????????????????????????????????????????????????????????????????????????????????????

			>Dyckia_braunii
			??????????????????????????????????????????????????????????????????????????????????????????????????????????????????????????????????????????????????????????????????????????????????????????????????????????????????????????????????????????????????????????????????????????????????????????????????????????????????????????????????????????????????????????????????????????????????????????????????????????????????????????????????????????????????????????????????????????????????????????????????????????????????????????????????????????????????????????????????????????????????????????????????????????????????????????????????????????????????????????????????????????????????????????????????????????????????????????????????????????????????????????????????????????????????????????????

			>Dyckia_brevifolia
			??????????????????????????????????????????????????????????????????????????????????????????????????????????????????????????????????????????????????????????????????????????????????????????????????????????????????????????????????????????????????????????????????????????????????????????????????????????????????????????????????????????????????????????????????????????????????????????????????????????????????????????????????????????????????????????????????????????????????????????????????????????????????????????????????????????????????????????????????????????????????????????????????????????????????????????????????????????????????????????????????????????????????????????????????????????????????????????????????????????????????????????????????????????????????????????????

			>Dyckia_choristaminea
			??????????????????????????????????????????????????????????????????????????????????????????????????????????????????????????????????????????????????????????????????????????????????????????????????????????????????????????????????????????????????????????????????????????????????????????????????????????????????????????????????????????????????????????????????????????????????????????????????????????????????????????????????????????????????????????????????????????????????????????????????????????????????????????????????????????????????????????????????????????????????????????????????????????????????????????????????????????????????????????????????????????????????????????????????????????????????????????????????????????????????????????????????????????????????????????????

			>Dyckia_cinerea
			??????????????????????????????????????????????????????????????????????????????????????????????????????????????????????????????????????????????????????????????????????????????????????????????????????????????????????????????????????????????????????????????????????????????????????????????????????????????????????????????????????????????????????????????????????????????????????????????????????????????????????????????????????????????????????????????????????????????????????????????????????????????????????????????????????????????????????????????????????????????????????????????????????????????????????????????????????????????????????????????????????????????????????????????????????????????????????????????????????????????????????????????????????????????????????????????

			>Dyckia_dawsonii
			ATAATTATAATAAAGTTAAATATGTCGAAATCCTTTTTGGAATAATACCAAATCTAAAATAAATGTCCGATAGCAAGTTGATCAGTTAATTCAATAAGAGATAAATGGAGGATAGCACTCGGTTTAGTTGGTACCACCCAACCGAATCCGATTCAATCGTTTACTCATTGAATGAGTCAATTTTCAAGTTCAGCCAATCTTTTTTTTTT-AAAAAAGAAATTGCAAGTAGATGAAATCGTGAATAAATGTGTTTCATTTCTCTATCCTTATAGAAAATAAGATCCATATTATATTACTTATGGAATTCGAACCCGAACTCGATTTATGATTCATTATTTCGATCTTATTGGCCTTTGTTCTTTTTTTTTTATTTTTCATATGGATTTCCGTCTTTATATTATACCCTTTCGTGGATGAATTATGCCTCTTTTCACATCTAGGATTTACATATACAACATATATTACTGTCAAGAGGGAATTTTTCTCAGTATTTAGATATTTAGATTCAAAATAAGAAAGGGATCAATTCAATTATAAACCTGTAAAACAAAGATTAGGATTGGGTTGGGTTGCGCTATATATATCAAAGAGTATACAATAATGATGTATTTGGTGAATCAAATACATGGTCTAATAACAAACCATTTTAACATAACATTTTAATGAGTTGATAATATTAATTGAATATTGAATATTTTTTGAAAGATTTTTGTTAAAGGTTTCATTCATGCCTAATCCATATCGAGTAGACCTTGTTGTTGTGAG

			>Dyckia_delicata
			??????????????????????????????????????????????????????????????????????????????????????????????????????????????????????????????????????????????????????????????????????????????????????????????????????????????????????????????????????????????????????????????????????????????????????????????????????????????????????????????????????????????????????????????????????????????????????????????????????????????????????????????????????????????????????????????????????????????????????????????????????????????????????????????????????????????????????????????????????????????????????????????????????????????????????????????????????????????????????????????????????????????????????????????????????????????????????????????????????????????????????????????????????????????????????????????

			>Dyckia_densiflora
			??????????????????????????????????????????????????????????????????????????????????????????????????????????????????????????????????????????????????????????????????????????????????????????????????????????????????????????????????????????????????????????????????????????????????????????????????????????????????????????????????????????????????????????????????????????????????????????????????????????????????????????????????????????????????????????????????????????????????????????????????????????????????????????????????????????????????????????????????????????????????????????????????????????????????????????????????????????????????????????????????????????????????????????????????????????????????????????????????????????????????????????????????????????????????????????????

			>Dyckia_dissitiflora
			??????????????????????????????????????????????????????????????????????????????????????????????????????????????????????????????????????????????????????????????????????????????????????????????????????????????????????????????????????????????????????????????????????????????????????????????????????????????????????????????????????????????????????????????????????????????????????????????????????????????????????????????????????????????????????????????????????????????????????????????????????????????????????????????????????????????????????????????????????????????????????????????????????????????????????????????????????????????????????????????????????????????????????????????????????????????????????????????????????????????????????????????????????????????????????????????

			>Dyckia_distachya
			??????????????????????????????????????????????????????????????????????????????????????????????????????????????????????????????????????????????????????????????????????????????????????????????????????????????????????????????????????????????????????????????????????????????????????????????????????????????????????????????????????????????????????????????????????????????????????????????????????????????????????????????????????????????????????????????????????????????????????????????????????????????????????????????????????????????????????????????????????????????????????????????????????????????????????????????????????????????????????????????????????????????????????????????????????????????????????????????????????????????????????????????????????????????????????????????

			>Dyckia_duckei
			??????????????????????????????????????????????????????????????????????????????????????????????????????????????????????????????????????????????????????????????????????????????????????????????????????????????????????????????????????????????????????????????????????????????????????????????????????????????????????????????????????????????????????????????????????????????????????????????????????????????????????????????????????????????????????????????????????????????????????????????????????????????????????????????????????????????????????????????????????????????????????????????????????????????????????????????????????????????????????????????????????????????????????????????????????????????????????????????????????????????????????????????????????????????????????????????

			>Dyckia_dusenii
			??????????????????????????????????????????????????????????????????????????????????????????????????????????????????????????????????????????????????????????????????????????????????????????????????????????????????????????????????????????????????????????????????????????????????????????????????????????????????????????????????????????????????????????????????????????????????????????????????????????????????????????????????????????????????????????????????????????????????????????????????????????????????????????????????????????????????????????????????????????????????????????????????????????????????????????????????????????????????????????????????????????????????????????????????????????????????????????????????????????????????????????????????????????????????????????????

			>Dyckia_elisabethae
			??????????????????????????????????????????????????????????????????????????????????????????????????????????????????????????????????????????????????????????????????????????????????????????????????????????????????????????????????????????????????????????????????????????????????????????????????????????????????????????????????????????????????????????????????????????????????????????????????????????????????????????????????????????????????????????????????????????????????????????????????????????????????????????????????????????????????????????????????????????????????????????????????????????????????????????????????????????????????????????????????????????????????????????????????????????????????????????????????????????????????????????????????????????????????????????????

			>Dyckia_encholirioides
			??????????????????????????????????????????????????????????????????????????????????CAGTTAATTCAATAAGAGATAAATGGGAGATAGCACTCGGTTTAGTTGGTACCACCCAACCGAATCCGATTCAATCGTTTACTCATTGAATGAGTCAATTTTCAAGTTCAGCCAATC-TTTTTTTTTTAAAAAATAAATTGCAAGTAGATGAAATCGTGAATAAATGTGTTTCATTTCTCTATCATTATAGAAAATAAGATCCATATTATATTACTTATGGAATTCGAACCCGAACTCGATTTATGATTCATTATTTCGATCTTATTGGCCTTTGTTCTTTTTTTTTTATTTTTCATATGGATTTCCGTCTTTATATTATACCCTTTCGTGGATGAATTATGCCTCTTTTCACATCTAGGATTTACATATACAACATATATTACTGTCAAGAGGGAATTTTTCTCAGTATTTAGATATTTAGATTCAAAATAAGAAAGGGATCAATTCAATTATAAACCTGTAAAACAAAGATTAGGATTGGGTTGGGTTGCGCTATATATATCAAAGAGTATACAATAATGATGTATTTGGTGAATCAAATACATGGTCTAATAACAAACCATTTTAACATAACATTTTAATGAGTTGATAATATTAATTGAATATTGAATATTTTTTGAAAGATTTTTGTTAAAGGTTTCATTCATGCCTAATCCATATCGAGTAGACCTTGTTGTTGTGAG

			>Dyckia_espiritosantensis
			??????????????????????????????????????????????????????????????????????????????????????????????????????????????????????????????????????????????????????????????????????????????????????????????????????????????????????????????????????????????????????????????????????????????????????????????????????????????????????????????????????????????????????????????????????????????????????????????????????????????????????????????????????????????????????????????????????????????????????????????????????????????????????????????????????????????????????????????????????????????????????????????????????????????????????????????????????????????????????????????????????????????????????????????????????????????????????????????????????????????????????????????????????????????????????????????

			>Dyckia_estevesii
			??????????????????????????????????????????????????????????????????????????????????CAGTTAATTCAATAAGAGATAAATGGAAGATAGCACTCGGTTTAGTTGGTACCACCCAACCGAATCCGATTCAATCGTTTACTCATTGAATGAGTCAATTTTCAAGTTCAGCCAATC-TTTTTTTTTTAAAAAATAAATTGCAAGTAGATGAAATCGTGAATAAATGTGTTTCATTTCTCTATCATTATAGAAAATAAGATCCATATTATATTACTTATGGAATTCGAACCCGAACTCGATTTATGATTCATTATTTCGATCTTATTGGCCTTTGTTCTTTTTTTTTTATTTTTCATATGGATTTCCGTCTTTATATTATACCCTTTCGTGGATGAATTATGCCTCTTTTCACATCTAGGATTTACATATACAACATATATTACTGTCAAGAGGGAATTTTTCTCAGTATTTAGATATTTAGATTCAAAATAAGAAAGGGATCAATTCAATTATAAACCTGTAAAACAAAGATTAGGATTGGGTTGGGTTGCGCTATATATATCAAAGAGTATACAATAATGATGTATTTGGTGAATCAAATACATGGTCTAATAACAAACCATTTTAACATAACATTTTAATGAGTTGATAATATTAATTGAATATTGAATATTTTTTGAAAGATTTTTGTTAAAGGTTTCATTCATGCCTAATCCATATCGAGTAGACCTTGTTGTTGTGAG

			>Dyckia_estevesii_var._braunii
			??????????????????????????????????????????????????????????????????????????????????????????????????????????????????????????????????????????????????????????????????????????????????????????????????????????????????????????????????????????????????????????????????????????????????????????????????????????????????????????????????????????????????????????????????????????????????????????????????????????????????????????????????????????????????????????????????????????????????????????????????????????????????????????????????????????????????????????????????????????????????????????????????????????????????????????????????????????????????????????????????????????????????????????????????????????????????????????????????????????????????????????????????????????????????????????????

			>Dyckia_excelsa
			??????????????????????????????????????????????????????????????????????????????????????????????????????????????????????????????????????????????????????????????????????????????????????????????????????????????????????????????????????????????????????????????????????????????????????????????????????????????????????????????????????????????????????????????????????????????????????????????????????????????????????????????????????????????????????????????????????????????????????????????????????????????????????????????????????????????????????????????????????????????????????????????????????????????????????????????????????????????????????????????????????????????????????????????????????????????????????????????????????????????????????????????????????????????????????????????

			>Dyckia_ferox
			ATAATTATAATAAAGTTAAATATGTCGAAATCCTTTTTGGAATAATACCAAATCTAAAATAAATGTCCGATAGCAAGTTGATCAGTTAATTCAATAAGAGATAAATGGAAGATAGCACTCGGTTTAGTTGGTACCACCCAACCGAATCCGATTCAATCGTTTACTCATTGAATGAGTCAATTTTCAAGTTCAGCCAATCTTTTTTTTTTTAAAAAATAAATTGCAAGTAGATGAAATCGTGAATAAATGTGTTTCATTTCTCTATCATTATAGAAAATAAGATCCATATTATATTACTTATGGAATTCGAACCCGAACTCGATTTATGATTCATTATTTCGATCTTATTGGCCTTTGTTCTTTTTTTTTTATTTTTCATATGGATTTCCGTCTTTATATTATACCCTTTCGTGGATGAATTATGCCTCTTTTCACATCTAGGATTTACATATACAACATATATTACTGTCAAGAGGGAATTTTTCTCAGTATTTAGATATTTAGATTCAAAATAAGAAAGGGATCAATTCAATTATAAACCTGTAAAACAAAGATTAGGATTGGGTTGGGTTGCGCTATATATATCAAAGAGTATACAATAATGATGTATTTGGTGAATCAAATACATGGTCTAATAACAAACCATTTTAACATAACATTTTAATGAGTTGATAATATTAATTGAATATTGAATATTTTTTGAAAGATTTTTGTTAAAGGTTTCATTCATGCCTAATCCATATCGAGTAGACCTTGTTGTTGTGAG

			>Dyckia_ferruginea
			??????????????????????????????????????????????????????????????????????????????????????????????????????????????????????????????????????????????????????????????????????????????????????????????????????????????????????????????????????????????????????????????????????????????????????????????????????????????????????????????????????????????????????????????????????????????????????????????????????????????????????????????????????????????????????????????????????????????????????????????????????????????????????????????????????????????????????????????????????????????????????????????????????????????????????????????????????????????????????????????????????????????????????????????????????????????????????????????????????????????????????????????????????????????????????????????

			>Dyckia_floribunda
			??????????????????????????????????????????????????????????????????????????????????????????????????????????????????????????????????????????????????????????????????????????????????????????????????????????????????????????????????????????????????????????????????????????????????????????????????????????????????????????????????????????????????????????????????????????????????????????????????????????????????????????????????????????????????????????????????????????????????????????????????????????????????????????????????????????????????????????????????????????????????????????????????????????????????????????????????????????????????????????????????????????????????????????????????????????????????????????????????????????????????????????????????????????????????????????????

			>Dyckia_fosteriana
			??????????????????????????????????????????????????????????????????????????????????????????????????????????????????????????????????????????????????????????????????????????????????????????????????????????????????????????????????????????????????????????????????????????????????????????????????????????????????????????????????????????????????????????????????????????????????????????????????????????????????????????????????????????????????????????????????????????????????????????????????????????????????????????????????????????????????????????????????????????????????????????????????????????????????????????????????????????????????????????????????????????????????????????????????????????????????????????????????????????????????????????????????????????????????????????????

			>Dyckia_goehringii
			??????????????????????????????????????????????????????????????????????????????????CAGTTAATTCAATAAGAGATAAATGGAAGATAGCACTCGGTTTAGTTGGTACCACCCAACCGAATCCGATTCAATCGTTTACTCATTGAATGAGTCAATTTTCAAGTTCAGCCAATCTTTTTTTTTTTAAAAAATAAATTGCAAGTAGATGAAATCGTGAATAAATGTGTTTCATTTCTCTATCATTATAGAAAATAAGATCCATATTATATTACTTATGGAATTCGAACCCGAACTCGATTTATGATTCATTATTTCGATCTTATTGGCCTTTGTTCTTTTTTTTTTATTTTTCATATGGATTTCCGTCTTTATATTATACCCTTTCGTGGATGAATTATGCCTCTTTTCACATCTAGGATTTACATATACAACATATATTACTGTCAAGAGGGAATTTTTCTCAGTATTTAGATATTTAGATTCAAAATAAGAAAGGGATCAATTCAATTATAAACCTGTAAAACAAAGATTAGGATTGGGTTGGGTTGCGCTATATATATCAAAGAGTATACAATAATGATGTATTTGGTGAATCAAATACATGGTCTAATAACAAACCATTTTAACATAACATTTTAATGAGTTGATAATATTAATTGAATATTGAATATTTTTTGAAAGATTTTTGTTAAAGGTTTCATTCATGCCTAATCCATATCGAGTAGACCTTGTTGTTGTGAG

			>Dyckia_gouveiana
			??????????????????????????????????????????????????????????????????????????????????????????????????????????????????????????????????????????????????????????????????????????????????????????????????????????????????????????????????????????????????????????????????????????????????????????????????????????????????????????????????????????????????????????????????????????????????????????????????????????????????????????????????????????????????????????????????????????????????????????????????????????????????????????????????????????????????????????????????????????????????????????????????????????????????????????????????????????????????????????????????????????????????????????????????????????????????????????????????????????????????????????????????????????????????????????????

			>Dyckia_grandidentata
			??????????????????????????????????????????????????????????????????????????????????????????????????????????????????????????????????????????????????????????????????????????????????????????????????????????????????????????????????????????????????????????????????????????????????????????????????????????????????????????????????????????????????????????????????????????????????????????????????????????????????????????????????????????????????????????????????????????????????????????????????????????????????????????????????????????????????????????????????????????????????????????????????????????????????????????????????????????????????????????????????????????????????????????????????????????????????????????????????????????????????????????????????????????????????????????????

			>Dyckia_granmogulensis
			??????????????????????????????????????????????????????????????????????????????????????????????????????????????????????????????????????????????????????????????????????????????????????????????????????????????????????????????????????????????????????????????????????????????????????????????????????????????????????????????????????????????????????????????????????????????????????????????????????????????????????????????????????????????????????????????????????????????????????????????????????????????????????????????????????????????????????????????????????????????????????????????????????????????????????????????????????????????????????????????????????????????????????????????????????????????????????????????????????????????????????????????????????????????????????????????

			>Dyckia_hebdingii
			??????????????????????????????????????????????????????????????????????????????????????????????????????????????????????????????????????????????????????????????????????????????????????????????????????????????????????????????????????????????????????????????????????????????????????????????????????????????????????????????????????????????????????????????????????????????????????????????????????????????????????????????????????????????????????????????????????????????????????????????????????????????????????????????????????????????????????????????????????????????????????????????????????????????????????????????????????????????????????????????????????????????????????????????????????????????????????????????????????????????????????????????????????????????????????????????

			>Dyckia_hohenbergioides
			??????????????????????????????????????????????????????????????????????????????????????????????????????????????????????????????????????????????????????????????????????????????????????????????????????????????????????????????????????????????????????????????????????????????????????????????????????????????????????????????????????????????????????????????????????????????????????????????????????????????????????????????????????????????????????????????????????????????????????????????????????????????????????????????????????????????????????????????????????????????????????????????????????????????????????????????????????????????????????????????????????????????????????????????????????????????????????????????????????????????????????????????????????????????????????????????

			>Dyckia_ibiramensis
			??????????????????????????????????????????????????????????????????????????????????????????????????????????????????????????????????????????????????????????????????????????????????????????????????????????????????????????????????????????????????????????????????????????????????????????????????????????????????????????????????????????????????????????????????????????????????????????????????????????????????????????????????????????????????????????????????????????????????????????????????????????????????????????????????????????????????????????????????????????????????????????????????????????????????????????????????????????????????????????????????????????????????????????????????????????????????????????????????????????????????????????????????????????????????????????????

			>Dyckia_jonesiana
			??????????????????????????????????????????????????????????????????????????????????????????????????????????????????????????????????????????????????????????????????????????????????????????????????????????????????????????????????????????????????????????????????????????????????????????????????????????????????????????????????????????????????????????????????????????????????????????????????????????????????????????????????????????????????????????????????????????????????????????????????????????????????????????????????????????????????????????????????????????????????????????????????????????????????????????????????????????????????????????????????????????????????????????????????????????????????????????????????????????????????????????????????????????????????????????????

			>Dyckia_kranziana
			??????????????????????????????????????????????????????????????????????????????????????????????????????????????????????????????????????????????????????????????????????????????????????????????????????????????????????????????????????????????????????????????????????????????????????????????????????????????????????????????????????????????????????????????????????????????????????????????????????????????????????????????????????????????????????????????????????????????????????????????????????????????????????????????????????????????????????????????????????????????????????????????????????????????????????????????????????????????????????????????????????????????????????????????????????????????????????????????????????????????????????????????????????????????????????????????

			>Dyckia_leptostachya
			??????????????????????????????????????????????????????????????????????????????????????????????????????????????????????????????????????????????????????????????????????????????????????????????????????????????????????????????????????????????????????????????????????????????????????????????????????????????????????????????????????????????????????????????????????????????????????????????????????????????????????????????????????????????????????????????????????????????????????????????????????????????????????????????????????????????????????????????????????????????????????????????????????????????????????????????????????????????????????????????????????????????????????????????????????????????????????????????????????????????????????????????????????????????????????????????

			>Dyckia_lindevaldae
			??????????????????????????????????????????????????????????????????????????????????????????????????????????????????????????????????????????????????????????????????????????????????????????????????????????????????????????????????????????????????????????????????????????????????????????????????????????????????????????????????????????????????????????????????????????????????????????????????????????????????????????????????????????????????????????????????????????????????????????????????????????????????????????????????????????????????????????????????????????????????????????????????????????????????????????????????????????????????????????????????????????????????????????????????????????????????????????????????????????????????????????????????????????????????????????????

			>Dyckia_lunaris
			??????????????????????????????????????????????????????????????????????????????????????????????????????????????????????????????????????????????????????????????????????????????????????????????????????????????????????????????????????????????????????????????????????????????????????????????????????????????????????????????????????????????????????????????????????????????????????????????????????????????????????????????????????????????????????????????????????????????????????????????????????????????????????????????????????????????????????????????????????????????????????????????????????????????????????????????????????????????????????????????????????????????????????????????????????????????????????????????????????????????????????????????????????????????????????????????

			>Dyckia_macedoi
			??????????????????????????????????????????????????????????????????????????????????????????????????????????????????????????????????????????????????????????????????????????????????????????????????????????????????????????????????????????????????????????????????????????????????????????????????????????????????????????????????????????????????????????????????????????????????????????????????????????????????????????????????????????????????????????????????????????????????????????????????????????????????????????????????????????????????????????????????????????????????????????????????????????????????????????????????????????????????????????????????????????????????????????????????????????????????????????????????????????????????????????????????????????????????????????????

			>Dyckia_machrisiana
			??????????????????????????????????????????????????????????????????????????????????????????????????????????????????????????????????????????????????????????????????????????????????????????????????????????????????????????????????????????????????????????????????????????????????????????????????????????????????????????????????????????????????????????????????????????????????????????????????????????????????????????????????????????????????????????????????????????????????????????????????????????????????????????????????????????????????????????????????????????????????????????????????????????????????????????????????????????????????????????????????????????????????????????????????????????????????????????????????????????????????????????????????????????????????????????????

			>Dyckia_maracasensis
			??????????????????????????????????????????????????????????????????????????????????????????????????????????????????????????????????????????????????????????????????????????????????????????????????????????????????????????????????????????????????????????????????????????????????????????????????????????????????????????????????????????????????????????????????????????????????????????????????????????????????????????????????????????????????????????????????????????????????????????????????????????????????????????????????????????????????????????????????????????????????????????????????????????????????????????????????????????????????????????????????????????????????????????????????????????????????????????????????????????????????????????????????????????????????????????????

			>Dyckia_maritima
			??????????????????????????????????????????????????????????????????????????????????????????????????????????????????????????????????????????????????????????????????????????????????????????????????????????????????????????????????????????????????????????????????????????????????????????????????????????????????????????????????????????????????????????????????????????????????????????????????????????????????????????????????????????????????????????????????????????????????????????????????????????????????????????????????????????????????????????????????????????????????????????????????????????????????????????????????????????????????????????????????????????????????????????????????????????????????????????????????????????????????????????????????????????????????????????????

			>Dyckia_marnier-lapostollei
			??????????????????????????????????????????????????????????????????????????????????????????????????????????????????????????????????????????????????????????????????????????????????????????????????????????????????????????????????????????????????????????????????????????????????????????????????????????????????????????????????????????????????????????????????????????????????????????????????????????????????????????????????????????????????????????????????????????????????????????????????????????????????????????????????????????????????????????????????????????????????????????????????????????????????????????????????????????????????????????????????????????????????????????????????????????????????????????????????????????????????????????????????????????????????????????????

			>Dyckia_marnier-lapostollei_var._estevesii
			??????????????????????????????????????????????????????????????????????????????????????????????????????????????????????????????????????????????????????????????????????????????????????????????????????????????????????????????????????????????????????????????????????????????????????????????????????????????????????????????????????????????????????????????????????????????????????????????????????????????????????????????????????????????????????????????????????????????????????????????????????????????????????????????????????????????????????????????????????????????????????????????????????????????????????????????????????????????????????????????????????????????????????????????????????????????????????????????????????????????????????????????????????????????????????????????

			>Dyckia_martinellii
			??????????????????????????????????????????????????????????????????????????????????????????????????????????????????????????????????????????????????????????????????????????????????????????????????????????????????????????????????????????????????????????????????????????????????????????????????????????????????????????????????????????????????????????????????????????????????????????????????????????????????????????????????????????????????????????????????????????????????????????????????????????????????????????????????????????????????????????????????????????????????????????????????????????????????????????????????????????????????????????????????????????????????????????????????????????????????????????????????????????????????????????????????????????????????????????????

			>Dyckia_mellobarretoi
			??????????????????????????????????????????????????????????????????????????????????????????????????????????????????????????????????????????????????????????????????????????????????????????????????????????????????????????????????????????????????????????????????????????????????????????????????????????????????????????????????????????????????????????????????????????????????????????????????????????????????????????????????????????????????????????????????????????????????????????????????????????????????????????????????????????????????????????????????????????????????????????????????????????????????????????????????????????????????????????????????????????????????????????????????????????????????????????????????????????????????????????????????????????????????????????????

			>Dyckia_microcalyx
			??????????????????????????????????????????????????????????????????????????????????????????????????????????????????????????????????????????????????????????????????????????????????????????????????????????????????????????????????????????????????????????????????????????????????????????????????????????????????????????????????????????????????????????????????????????????????????????????????????????????????????????????????????????????????????????????????????????????????????????????????????????????????????????????????????????????????????????????????????????????????????????????????????????????????????????????????????????????????????????????????????????????????????????????????????????????????????????????????????????????????????????????????????????????????????????????

			>Dyckia_microcalyx_var._ostenii
			??????????????????????????????????????????????????????????????????????????????????????????????????????????????????????????????????????????????????????????????????????????????????????????????????????????????????????????????????????????????????????????????????????????????????????????????????????????????????????????????????????????????????????????????????????????????????????????????????????????????????????????????????????????????????????????????????????????????????????????????????????????????????????????????????????????????????????????????????????????????????????????????????????????????????????????????????????????????????????????????????????????????????????????????????????????????????????????????????????????????????????????????????????????????????????????????

			>Dyckia_milagrensis
			??????????????????????????????????????????????????????????????????????????????????????????????????????????????????????????????????????????????????????????????????????????????????????????????????????????????????????????????????????????????????????????????????????????????????????????????????????????????????????????????????????????????????????????????????????????????????????????????????????????????????????????????????????????????????????????????????????????????????????????????????????????????????????????????????????????????????????????????????????????????????????????????????????????????????????????????????????????????????????????????????????????????????????????????????????????????????????????????????????????????????????????????????????????????????????????????

			>Dyckia_mirandiana
			??????????????????????????????????????????????????????????????????????????????????????????????????????????????????????????????????????????????????????????????????????????????????????????????????????????????????????????????????????????????????????????????????????????????????????????????????????????????????????????????????????????????????????????????????????????????????????????????????????????????????????????????????????????????????????????????????????????????????????????????????????????????????????????????????????????????????????????????????????????????????????????????????????????????????????????????????????????????????????????????????????????????????????????????????????????????????????????????????????????????????????????????????????????????????????????????

			>Dyckia_montezumensis
			??????????????????????????????????????????????????????????????????????????????????????????????????????????????????????????????????????????????????????????????????????????????????????????????????????????????????????????????????????????????????????????????????????????????????????????????????????????????????????????????????????????????????????????????????????????????????????????????????????????????????????????????????????????????????????????????????????????????????????????????????????????????????????????????????????????????????????????????????????????????????????????????????????????????????????????????????????????????????????????????????????????????????????????????????????????????????????????????????????????????????????????????????????????????????????????????

			>Dyckia_monticola
			??????????????????????????????????????????????????????????????????????????????????????????????????????????????????????????????????????????????????????????????????????????????????????????????????????????????????????????????????????????????????????????????????????????????????????????????????????????????????????????????????????????????????????????????????????????????????????????????????????????????????????????????????????????????????????????????????????????????????????????????????????????????????????????????????????????????????????????????????????????????????????????????????????????????????????????????????????????????????????????????????????????????????????????????????????????????????????????????????????????????????????????????????????????????????????????????

			>Dyckia_nana
			??????????????????????????????????????????????????????????????????????????????????????????????????????????????????????????????????????????????????????????????????????????????????????????????????????????????????????????????????????????????????????????????????????????????????????????????????????????????????????????????????????????????????????????????????????????????????????????????????????????????????????????????????????????????????????????????????????????????????????????????????????????????????????????????????????????????????????????????????????????????????????????????????????????????????????????????????????????????????????????????????????????????????????????????????????????????????????????????????????????????????????????????????????????????????????????????

			>Dyckia_niederleinii
			??????????????????????????????????????????????????????????????????????????????????????????????????????????????????????????????????????????????????????????????????????????????????????????????????????????????????????????????????????????????????????????????????????????????????????????????????????????????????????????????????????????????????????????????????????????????????????????????????????????????????????????????????????????????????????????????????????????????????????????????????????????????????????????????????????????????????????????????????????????????????????????????????????????????????????????????????????????????????????????????????????????????????????????????????????????????????????????????????????????????????????????????????????????????????????????????

			>Dyckia_oligantha
			??????????????????????????????????????????????????????????????????????????????????????????????????????????????????????????????????????????????????????????????????????????????????????????????????????????????????????????????????????????????????????????????????????????????????????????????????????????????????????????????????????????????????????????????????????????????????????????????????????????????????????????????????????????????????????????????????????????????????????????????????????????????????????????????????????????????????????????????????????????????????????????????????????????????????????????????????????????????????????????????????????????????????????????????????????????????????????????????????????????????????????????????????????????????????????????????

			>Dyckia_paraensis
			??????????????????????????????????????????????????????????????????????????????????????????????????????????????????????????????????????????????????????????????????????????????????????????????????????????????????????????????????????????????????????????????????????????????????????????????????????????????????????????????????????????????????????????????????????????????????????????????????????????????????????????????????????????????????????????????????????????????????????????????????????????????????????????????????????????????????????????????????????????????????????????????????????????????????????????????????????????????????????????????????????????????????????????????????????????????????????????????????????????????????????????????????????????????????????????????

			>Dyckia_pectinata
			??????????????????????????????????????????????????????????????????????????????????????????????????????????????????????????????????????????????????????????????????????????????????????????????????????????????????????????????????????????????????????????????????????????????????????????????????????????????????????????????????????????????????????????????????????????????????????????????????????????????????????????????????????????????????????????????????????????????????????????????????????????????????????????????????????????????????????????????????????????????????????????????????????????????????????????????????????????????????????????????????????????????????????????????????????????????????????????????????????????????????????????????????????????????????????????????

			>Dyckia_pernambucana
			??????????????????????????????????????????????????????????????????????????????????????????????????????????????????????????????????????????????????????????????????????????????????????????????????????????????????????????????????????????????????????????????????????????????????????????????????????????????????????????????????????????????????????????????????????????????????????????????????????????????????????????????????????????????????????????????????????????????????????????????????????????????????????????????????????????????????????????????????????????????????????????????????????????????????????????????????????????????????????????????????????????????????????????????????????????????????????????????????????????????????????????????????????????????????????????????

			>Dyckia_platyphylla
			??????????????????????????????????????????????????????????????????????????????????????????????????????????????????????????????????????????????????????????????????????????????????????????????????????????????????????????????????????????????????????????????????????????????????????????????????????????????????????????????????????????????????????????????????????????????????????????????????????????????????????????????????????????????????????????????????????????????????????????????????????????????????????????????????????????????????????????????????????????????????????????????????????????????????????????????????????????????????????????????????????????????????????????????????????????????????????????????????????????????????????????????????????????????????????????????

			>Dyckia_pottiorum
			??????????????????????????????????????????????????????????????????????????????????????????????????????????????????????????????????????????????????????????????????????????????????????????????????????????????????????????????????????????????????????????????????????????????????????????????????????????????????????????????????????????????????????????????????????????????????????????????????????????????????????????????????????????????????????????????????????????????????????????????????????????????????????????????????????????????????????????????????????????????????????????????????????????????????????????????????????????????????????????????????????????????????????????????????????????????????????????????????????????????????????????????????????????????????????????????

			>Dyckia_pseudococcinea
			??????????????????????????????????????????????????????????????????????????????????????????????????????????????????????????????????????????????????????????????????????????????????????????????????????????????????????????????????????????????????????????????????????????????????????????????????????????????????????????????????????????????????????????????????????????????????????????????????????????????????????????????????????????????????????????????????????????????????????????????????????????????????????????????????????????????????????????????????????????????????????????????????????????????????????????????????????????????????????????????????????????????????????????????????????????????????????????????????????????????????????????????????????????????????????????????

			>Dyckia_pulquinensis
			??????????????????????????????????????????????????????????????????????????????????????????????????????????????????????????????????????????????????????????????????????????????????????????????????????????????????????????????????????????????????????????????????????????????????????????????????????????????????????????????????????????????????????????????????????????????????????????????????????????????????????????????????????????????????????????????????????????????????????????????????????????????????????????????????????????????????????????????????????????????????????????????????????????????????????????????????????????????????????????????????????????????????????????????????????????????????????????????????????????????????????????????????????????????????????????????

			>Dyckia_pumila
			??????????????????????????????????????????????????????????????????????????????????????????????????????????????????????????????????????????????????????????????????????????????????????????????????????????????????????????????????????????????????????????????????????????????????????????????????????????????????????????????????????????????????????????????????????????????????????????????????????????????????????????????????????????????????????????????????????????????????????????????????????????????????????????????????????????????????????????????????????????????????????????????????????????????????????????????????????????????????????????????????????????????????????????????????????????????????????????????????????????????????????????????????????????????????????????????

			>Dyckia_rariflora
			??????????????????????????????????????????????????????????????????????????????????????????????????????????????????????????????????????????????????????????????????????????????????????????????????????????????????????????????????????????????????????????????????????????????????????????????????????????????????????????????????????????????????????????????????????????????????????????????????????????????????????????????????????????????????????????????????????????????????????????????????????????????????????????????????????????????????????????????????????????????????????????????????????????????????????????????????????????????????????????????????????????????????????????????????????????????????????????????????????????????????????????????????????????????????????????????

			>Dyckia_remotiflora
			??????????????????????????????????????????????????????????????????????????????????????????????????????????????????????????????????????????????????????????????????????????????????????????????????????????????????????????????????????????????????????????????????????????????????????????????????????????????????????????????????????????????????????????????????????????????????????????????????????????????????????????????????????????????????????????????????????????????????????????????????????????????????????????????????????????????????????????????????????????????????????????????????????????????????????????????????????????????????????????????????????????????????????????????????????????????????????????????????????????????????????????????????????????????????????????????

			>Dyckia_rondonopolitana
			??????????????????????????????????????????????????????????????????????????????????????????????????????????????????????????????????????????????????????????????????????????????????????????????????????????????????????????????????????????????????????????????????????????????????????????????????????????????????????????????????????????????????????????????????????????????????????????????????????????????????????????????????????????????????????????????????????????????????????????????????????????????????????????????????????????????????????????????????????????????????????????????????????????????????????????????????????????????????????????????????????????????????????????????????????????????????????????????????????????????????????????????????????????????????????????????

			>Dyckia_saxatilis
			??????????????????????????????????????????????????????????????????????????????????????????????????????????????????????????????????????????????????????????????????????????????????????????????????????????????????????????????????????????????????????????????????????????????????????????????????????????????????????????????????????????????????????????????????????????????????????????????????????????????????????????????????????????????????????????????????????????????????????????????????????????????????????????????????????????????????????????????????????????????????????????????????????????????????????????????????????????????????????????????????????????????????????????????????????????????????????????????????????????????????????????????????????????????????????????????

			>Dyckia_secunda
			??????????????????????????????????????????????????????????????????????????????????????????????????????????????????????????????????????????????????????????????????????????????????????????????????????????????????????????????????????????????????????????????????????????????????????????????????????????????????????????????????????????????????????????????????????????????????????????????????????????????????????????????????????????????????????????????????????????????????????????????????????????????????????????????????????????????????????????????????????????????????????????????????????????????????????????????????????????????????????????????????????????????????????????????????????????????????????????????????????????????????????????????????????????????????????????????

			>Dyckia_selloa
			??????????????????????????????????????????????????????????????????????????????????????????????????????????????????????????????????????????????????????????????????????????????????????????????????????????????????????????????????????????????????????????????????????????????????????????????????????????????????????????????????????????????????????????????????????????????????????????????????????????????????????????????????????????????????????????????????????????????????????????????????????????????????????????????????????????????????????????????????????????????????????????????????????????????????????????????????????????????????????????????????????????????????????????????????????????????????????????????????????????????????????????????????????????????????????????????

			>Dyckia_tenebrosa
			??????????????????????????????????????????????????????????????????????????????????????????????????????????????????????????????????????????????????????????????????????????????????????????????????????????????????????????????????????????????????????????????????????????????????????????????????????????????????????????????????????????????????????????????????????????????????????????????????????????????????????????????????????????????????????????????????????????????????????????????????????????????????????????????????????????????????????????????????????????????????????????????????????????????????????????????????????????????????????????????????????????????????????????????????????????????????????????????????????????????????????????????????????????????????????????????

			>Dyckia_tobatiensis
			??????????????????????????????????????????????????????????????????????????????????????????????????????????????????????????????????????????????????????????????????????????????????????????????????????????????????????????????????????????????????????????????????????????????????????????????????????????????????????????????????????????????????????????????????????????????????????????????????????????????????????????????????????????????????????????????????????????????????????????????????????????????????????????????????????????????????????????????????????????????????????????????????????????????????????????????????????????????????????????????????????????????????????????????????????????????????????????????????????????????????????????????????????????????????????????????

			>Dyckia_tomentella
			??????????????????????????????????????????????????????????????????????????????????????????????????????????????????????????????????????????????????????????????????????????????????????????????????????????????????????????????????????????????????????????????????????????????????????????????????????????????????????????????????????????????????????????????????????????????????????????????????????????????????????????????????????????????????????????????????????????????????????????????????????????????????????????????????????????????????????????????????????????????????????????????????????????????????????????????????????????????????????????????????????????????????????????????????????????????????????????????????????????????????????????????????????????????????????????????

			>Dyckia_ursina
			??????????????????????????????????????????????????????????????????????????????????????????????????????????????????????????????????????????????????????????????????????????????????????????????????????????????????????????????????????????????????????????????????????????????????????????????????????????????????????????????????????????????????????????????????????????????????????????????????????????????????????????????????????????????????????????????????????????????????????????????????????????????????????????????????????????????????????????????????????????????????????????????????????????????????????????????????????????????????????????????????????????????????????????????????????????????????????????????????????????????????????????????????????????????????????????????

			>Dyckia_velascana
			??????????????????????????????????????????????????????????????????????????????????????????????????????????????????????????????????????????????????????????????????????????????????????????????????????????????????????????????????????????????????????????????????????????????????????????????????????????????????????????????????????????????????????????????????????????????????????????????????????????????????????????????????????????????????????????????????????????????????????????????????????????????????????????????????????????????????????????????????????????????????????????????????????????????????????????????????????????????????????????????????????????????????????????????????????????????????????????????????????????????????????????????????????????????????????????????

			>Dyckia_vestita
			??????????????????????????????????????????????????????????????????????????????????????????????????????????????????????????????????????????????????????????????????????????????????????????????????????????????????????????????????????????????????????????????????????????????????????????????????????????????????????????????????????????????????????????????????????????????????????????????????????????????????????????????????????????????????????????????????????????????????????????????????????????????????????????????????????????????????????????????????????????????????????????????????????????????????????????????????????????????????????????????????????????????????????????????????????????????????????????????????????????????????????????????????????????????????????????????

			>Dyckia_vicentensis
			??????????????????????????????????????????????????????????????????????????????????????????????????????????????????????????????????????????????????????????????????????????????????????????????????????????????????????????????????????????????????????????????????????????????????????????????????????????????????????????????????????????????????????????????????????????????????????????????????????????????????????????????????????????????????????????????????????????????????????????????????????????????????????????????????????????????????????????????????????????????????????????????????????????????????????????????????????????????????????????????????????????????????????????????????????????????????????????????????????????????????????????????????????????????????????????????

			>Dyckia_walteriana
			??????????????????????????????????????????????????????????????????????????????????????????????????????????????????????????????????????????????????????????????????????????????????????????????????????????????????????????????????????????????????????????????????????????????????????????????????????????????????????????????????????????????????????????????????????????????????????????????????????????????????????????????????????????????????????????????????????????????????????????????????????????????????????????????????????????????????????????????????????????????????????????????????????????????????????????????????????????????????????????????????????????????????????????????????????????????????????????????????????????????????????????????????????????????????????????????

			>Edmundoa_ambigua
			???????????????????????????????????TTTTGGATA-TACCAAATCGAAAATAAATGTCCAATAGCAAGTTGATCAGTTAATTCAATAAGAGATAAATGGGAGATAGCACTCGATTTAGTTGGTACCACCCAACCGAATCCGATTCACTCGTTTACTCATTCAATGAGTAAATTTTCAAGTTCAGCCAATCTTTTTTTTTT-AAAAAAAAAATTGCAAGTAGATGAAATCGTGAATAAATGTGTTTCATTTCTATATCATTATATAAAAAAAGATCCATATTATATTACTTATGGAATTCGAACCCGAACTCGATTTA?GATTTATTATT?GGATCTTATTGGCCTTTGTTCTTTATTTTTGATTTTTAATATAGATTTCCGTCTTTATATTATACCCTTTCGGGGATGAATTATGCCTATTTTCACATCTAGGATTTACATATACAACATATATTACTGTCAAGAGGGAATTTTTCTCAGTATTTAGATATTTAGATTCAAAATAAGAAAGGGATCAATTCAATTATAAACCTGCAAAACAAAGATTAGGATTGGGTTGGGTTGCGCTATATATATCAAAGAGTATACAATAATGATGTATTTGGTGAATCAAATACATGGTCTAATAACAAACCATTTTAACATAACATTTTAATGAGTTGATAATATTAATTGAATATTGAATATTTTTTGAAAGATTTTTGTTAAAGGTTTCATTCATGCCTAATCCATATCGAGTAGACCTTGTTGTTGTG??

			>Edmundoa_lindenii
			??????????????????????????????????????????????????????????????????????????????????CAGTTAATTCAATAAGAGATAAATGGGAGATAGCACTCGATTTAGTTGGTACCACCCAACCGAATCCGATTCACTCGTTTACTCATTGAATGAGTAAATTTTCAAGTTCAGCCAATCTTTTTTTTTT-CAAAAAAAAATTGCAAGTAGATGAAATCGCGAATAAATGTGTTTCATTTCTATATCATTATATAAAAAAAGATCCATATTATATTACTTATGGAATTCGAACCCGAACTCGATTTATGATTTATTATTTGGATCTTATTGGCCTTTGTTCTTTATTTTTGATTTTTCATATAGATTTCCGTCTTTATATTATACCCTTTCGTGGATGAATTATGCCTATTTTCACATCTAGGATTTACATATACAACATATATTACTGTCAAGAGGGAATTTTTCTCAGTATTTAGATATTTAGATTCAAAATAAGAAAGGGATCAATTCAATTATAAACCTGCAAAACAAAGATTAGGATTGGGTTGGGTTGCGCTATATATATCAAAGAGTATACAATAATGATGTATTTGGTGAATCAAATACATGGTCTAATAACAAACCATTTTAACATAACATTTTAATGAGTTGATAATATTAATTGAATATTGAATATTTTTTGAAAGATTTTTGTTAAAGGTTTCATTCATGCCTAATCCATATCGAGTAGACCTTGTTGTTGTGAG

			>Edmundoa_lindenii_var._rosaea
			??????????????????????????????????????????????????????????????????????????????????????????????????????????????????????????????????????????????????????????????????????????????????????????????????????????????????????????????????????????????????????????????????????????????????????????????????????????????????????????????????????????????????????????????????????????????????????????????????????????????????????????????????????????????????????????????????????????????????????????????????????????????????????????????????????????????????????????????????????????????????????????????????????????????????????????????????????????????????????????????????????????????????????????????????????????????????????????????????????????????????????????????????????????????????????????????

			>Edmundoa_perplexa
			ATAATTATAATAAAGTGAAATATGTCGAAATCCTTTTTGGAATAATACCAAATC-AAAATAAATGTCCGATAGCAAGTTGATCAGTTAATTCAATAAGAGATAAATGGGGGATAGCACTCGATTTAGTTGGTACCACCCAACCGAATCCGATTCACTCGTTTACTCATTGAATGAGTAAATTTTCAAGTTCAGCCAATCTTTTTTTTTTCAAAAAAAAAATTGCAAGTAGATGAAATCGCGAATAAATGTGTTTCATTTCTATATCATTATATAAAAAAAGATCCATATTATATTACTTATGGAATTCGAACCCGAACTCGATTTATGATTTATTATTTGGATCTTATTGGCCTTTGTTCTTTATTTTTGATTTTTCATATAGATTTCCGTCTTTATATTATACCCTTTCGTGGATGAATTATGCCTATTTTCACATCTAGGATTTACATATACAACATATATTACTGTCAAGAGGGAATTTTTCTCAGTATTTAGATATTTAGATTCAAAATAAGAAAGGGATCAATTCAATTATAAACCTGCAAAACAAAGATTAGGATTGGGTTGGGTTGCGCTATATATATCAAAGAGTATACAATAATGATGTATTTGGTGAATCAAATACATGGTCTAATAACAAACCATTTTAACATAACATTTTAATGAGTTGATAATATTAATTGAATATTGAATATTTTTTGAAAGATTTTTGTTAAAGGTTTCATTCATGCCTAATCCATATCGAGTAGACCTTGTTGTTGTGAG

			>Eduandrea_selloana
			ATAATTATAATAAAGTGAAATATGTCGAAATCCTTTTTGGAATAATACCAAATCGAAAATAAATGTCCGATAGCAAGTTGATCAGTTAATTCAATAAGAGATAAATGGGAGATAGCACTCGATTTAGTTGGTACCACCCAACCGAATCCGATTCACTCGTTTACTCATTCAATGAGTAAATTTTCAAGTTCAGCCAATCTTTTTTTTT--AAAAAAAAAATTGCAAGTAGATGAAATCGTGAATAAATGTGTTTCATTTCTATATCATTATATAAAAAAAGATCCATATTATATTACTTATGGAATTCGAACCCGAACTCGATTTATGATTTATTATTTGGATCTTATTGGCCTTTGTTCTTTATTTTTGATTTTTCATATGGATTTCCGTCTTTATATTATACCCTTTCGTGGATGAATTATGCCTATTTTCACATCTAGGATTTACATATACAACATATATTACCCTCAAGAGGGAATTTTTCTCAGTATTTAGATATTTAGATTCAAAATAAGAAAGGGATCAATTCAATTATAAACCTGCAAAACAAAGATTAGGATTGGGTTGGGTTGCGCTATATATATCAAAGAGTATACAATAATGATGTATTTGGTGAATCAAATACATGGTCTAATAACAAACCATTTTAACATAACATTTTAATGAGTTGATAATATTAATTGAATATTGAATATTTTTTGAAAGATTTTTGTTAAAGGTTTCATTCATGCCTAATCCATATCGAGTAGACCTTGTTGTTGTGAG

			>Encholirium_agavoides
			??????????????????????????????????????????????????????????????????????????????????????????????????????????????????????????????????????????????????????????????????????????????????????????????????????????????????????????????????????????????????????????????????????????????????????????????????????????????????????????????????????????????????????????????????????????????????????????????????????????????????????????????????????????????????????????????????????????????????????????????????????????????????????????????????????????????????????????????????????????????????????????????????????????????????????????????????????????????????????????????????????????????????????????????????????????????????????????????????????????????????????????????????????????????????????????????

			>Encholirium_biflorum
			??????????????????????????????????????????????????????????????????????????????????????????????????????????????????????????????????????????????????????????????????????????????????????????????????????????????????????????????????????????????????????????????????????????????????????????????????????????????????????????????????????????????????????????????????????????????????????????????????????????????????????????????????????????????????????????????????????????????????????????????????????????????????????????????????????????????????????????????????????????????????????????????????????????????????????????????????????????????????????????????????????????????????????????????????????????????????????????????????????????????????????????????????????????????????????????????

			>Encholirium_brachypodum
			??????????????????????????????????????????????????????????????????????????????????????????????????????????????????????????????????????????????????????????????????????????????????????????????????????????????????????????????????????????????????????????????????????????????????????????????????????????????????????????????????????????????????????????????????????????????????????????????????????????????????????????????????????????????????????????????????????????????????????????????????????????????????????????????????????????????????????????????????????????????????????????????????????????????????????????????????????????????????????????????????????????????????????????????????????????????????????????????????????????????????????????????????????????????????????????????

			>Encholirium_ctenophyllum
			??????????????????????????????????????????????????????????????????????????????????????????????????????????????????????????????????????????????????????????????????????????????????????????????????????????????????????????????????????????????????????????????????????????????????????????????????????????????????????????????????????????????????????????????????????????????????????????????????????????????????????????????????????????????????????????????????????????????????????????????????????????????????????????????????????????????????????????????????????????????????????????????????????????????????????????????????????????????????????????????????????????????????????????????????????????????????????????????????????????????????????????????????????????????????????????????

			>Encholirium_diamantinum
			??????????????????????????????????????????????????????????????????????????????????????????????????????????????????????????????????????????????????????????????????????????????????????????????????????????????????????????????????????????????????????????????????????????????????????????????????????????????????????????????????????????????????????????????????????????????????????????????????????????????????????????????????????????????????????????????????????????????????????????????????????????????????????????????????????????????????????????????????????????????????????????????????????????????????????????????????????????????????????????????????????????????????????????????????????????????????????????????????????????????????????????????????????????????????????????????

			>Encholirium_eddie-estevesii
			??????????????????????????????????????????????????????????????????????????????????????????????????????????????????????????????????????????????????????????????????????????????????????????????????????????????????????????????????????????????????????????????????????????????????????????????????????????????????????????????????????????????????????????????????????????????????????????????????????????????????????????????????????????????????????????????????????????????????????????????????????????????????????????????????????????????????????????????????????????????????????????????????????????????????????????????????????????????????????????????????????????????????????????????????????????????????????????????????????????????????????????????????????????????????????????????

			>Encholirium_erectiflorum
			??????????????????????????????????????????????????????????????????????????????????????????????????????????????????????????????????????????????????????????????????????????????????????????????????????????????????????????????????????????????????????????????????????????????????????????????????????????????????????????????????????????????????????????????????????????????????????????????????????????????????????????????????????????????????????????????????????????????????????????????????????????????????????????????????????????????????????????????????????????????????????????????????????????????????????????????????????????????????????????????????????????????????????????????????????????????????????????????????????????????????????????????????????????????????????????????

			>Encholirium_fragae
			??????????????????????????????????????????????????????????????????????????????????????????????????????????????????????????????????????????????????????????????????????????????????????????????????????????????????????????????????????????????????????????????????????????????????????????????????????????????????????????????????????????????????????????????????????????????????????????????????????????????????????????????????????????????????????????????????????????????????????????????????????????????????????????????????????????????????????????????????????????????????????????????????????????????????????????????????????????????????????????????????????????????????????????????????????????????????????????????????????????????????????????????????????????????????????????????

			>Encholirium_gracile
			??????????????????????????????????????????????????????????????????????????????????????????????????????????????????????????????????????????????????????????????????????????????????????????????????????????????????????????????????????????????????????????????????????????????????????????????????????????????????????????????????????????????????????????????????????????????????????????????????????????????????????????????????????????????????????????????????????????????????????????????????????????????????????????????????????????????????????????????????????????????????????????????????????????????????????????????????????????????????????????????????????????????????????????????????????????????????????????????????????????????????????????????????????????????????????????????

			>Encholirium_heloisae
			??????????????????????????????????????????????????????????????????????????????????????????????????????????????????????????????????????????????????????????????????????????????????????????????????????????????????????????????????????????????????????????????????????????????????????????????????????????????????????????????????????????????????????????????????????????????????????????????????????????????????????????????????????????????????????????????????????????????????????????????????????????????????????????????????????????????????????????????????????????????????????????????????????????????????????????????????????????????????????????????????????????????????????????????????????????????????????????????????????????????????????????????????????????????????????????????

			>Encholirium_horridum
			??????????????????????????????????????????????????????????????????????????????????CAGTTAATTCAATAAGAGATAAATGGAAGATAGCACTCGGTTTAGTTGGTATCACCCAACCGAATCCGATTCAATCGTTTACTCATTGAATGAGTCAATTTTCAAGTTCAGCCAATCTTTTTTTTTTTAAAAAATAAATTGCAAGTAGATGAAATCGTGAATAAATGTGTTTCATTTCTCTATCATTATAGAAAATAAGATCCATATTATATTACTTATGGAATTCGAACCCGAACTCGATTTATGATTCATTATTTCGATCTTATTGGCCTTTGTTCTTTATTTTTGATTTTTCATATGGATTTCCGTCTTTATATTATACCCTTTCGTGGATGAATTATGCCTCTTTTCACATCTAGGATTTACATATACAACATATATTACTGTCAAGAGGGAATTTTTCTCAGTATTTAGATATTTAGATTCAAAATAAGAAAGGGATCAATTCTTTTATAAACCTGTAAAACAAAGATTAGGATTGGGTTGGGTTGCGCTATATATATCAAAGAGTATACAATAATGATGTATTTGGTGAATCAAATACATGGTCTAATAACAAACCATTTTAACATAACATTTTAATGAGTTGATAATATTAATTGAATATTGAATATTTTTTGAAAGATTTTTGTTAAAGGTTTCATTCATGCCTAATCCATATCGAGTAGACCTTGTTGTTGTGAG

			>Encholirium_irwinii
			ATAATTATAATAAAGTTAAATATGTCGAAATCCTTTTTGGAATAATACCAAATCTAAAATAAATGTCCGATAGCAAGTTGATCAGTTAATTCAATAAGAGATAAATGGAAGATAGCACTCGGTTTAGTTGGTACCACCCAACCGAATCCGATTCAATCGTTTACTCATTGAATGAGTCAATTTTCAAGTTCAGCCAATCTTTTTTTTTTTAAAAAATAAATTGCAAGTAGATGAAATCGTGAATAAATGTGTTTCATTTCTCTATCATTATAGAAAATAAGATCCATATTATATTACTTATGGAATTCGAACCCGAACTCGATTTATGATTCATTATTTCGATCTTATTGGCCTTTGTTCTTTATTTTTGATTTTTCATATGGATTTCCGTCTTTATATTATACCCTTTCGTGGATGAATTATGCCTCTTTTCACATCTAGGATTTACATATACAACATATATTACTGTCAAGAGGGAATTTTTCTCAGTATTTAGATATTTAGATTCAAAATAAGAAAGGGATCAATTCAATTATAAACCTGTAAAACAAAGATTAGGATTGGGTTGGGTTGCGCTATATATATCAAAGAGTATACAATAATGATGTATTTGGTGAATCAAATACATGGTCTAATAACAAACCATTTTAACATAACATTTTAATGAGTTGATAATATTAATTGAATATTGAATATTTTTTGAAAGATTTTTGTTAAAGGTTTCATTCATGCCTAATCCATATCGAGTAGACCTTGTTGTTGTGAG

			>Encholirium_kranzianum
			??????????????????????????????????????????????????????????????????????????????????????????????????????????????????????????????????????????????????????????????????????????????????????????????????????????????????????????????????????????????????????????????????????????????????????????????????????????????????????????????????????????????????????????????????????????????????????????????????????????????????????????????????????????????????????????????????????????????????????????????????????????????????????????????????????????????????????????????????????????????????????????????????????????????????????????????????????????????????????????????????????????????????????????????????????????????????????????????????????????????????????????????????????????????????????????????

			>Encholirium_longiflorum
			??????????????????????????????????????????????????????????????????????????????????????????????????????????????????????????????????????????????????????????????????????????????????????????????????????????????????????????????????????????????????????????????????????????????????????????????????????????????????????????????????????????????????????????????????????????????????????????????????????????????????????????????????????????????????????????????????????????????????????????????????????????????????????????????????????????????????????????????????????????????????????????????????????????????????????????????????????????????????????????????????????????????????????????????????????????????????????????????????????????????????????????????????????????????????????????????

			>Encholirium_luxor
			??????????????????????????????????????????????????????????????????????????????????????????????????????????????????????????????????????????????????????????????????????????????????????????????????????????????????????????????????????????????????????????????????????????????????????????????????????????????????????????????????????????????????????????????????????????????????????????????????????????????????????????????????????????????????????????????????????????????????????????????????????????????????????????????????????????????????????????????????????????????????????????????????????????????????????????????????????????????????????????????????????????????????????????????????????????????????????????????????????????????????????????????????????????????????????????????

			>Encholirium_lymanianum
			??????????????????????????????????????????????????????????????????????????????????????????????????????????????????????????????????????????????????????????????????????????????????????????????????????????????????????????????????????????????????????????????????????????????????????????????????????????????????????????????????????????????????????????????????????????????????????????????????????????????????????????????????????????????????????????????????????????????????????????????????????????????????????????????????????????????????????????????????????????????????????????????????????????????????????????????????????????????????????????????????????????????????????????????????????????????????????????????????????????????????????????????????????????????????????????????

			>Encholirium_magalhaesii
			??????????????????????????????????????????????????????????????????????????????????????????????????????????????????????????????????????????????????????????????????????????????????????????????????????????????????????????????????????????????????????????????????????????????????????????????????????????????????????????????????????????????????????????????????????????????????????????????????????????????????????????????????????????????????????????????????????????????????????????????????????????????????????????????????????????????????????????????????????????????????????????????????????????????????????????????????????????????????????????????????????????????????????????????????????????????????????????????????????????????????????????????????????????????????????????????

			>Encholirium_maximum
			??????????????????????????????????????????????????????????????????????????????????????????????????????????????????????????????????????????????????????????????????????????????????????????????????????????????????????????????????????????????????????????????????????????????????????????????????????????????????????????????????????????????????????????????????????????????????????????????????????????????????????????????????????????????????????????????????????????????????????????????????????????????????????????????????????????????????????????????????????????????????????????????????????????????????????????????????????????????????????????????????????????????????????????????????????????????????????????????????????????????????????????????????????????????????????????????

			>Encholirium_pedicellatum
			??????????????????????????????????????????????????????????????????????????????????????????????????????????????????????????????????????????????????????????????????????????????????????????????????????????????????????????????????????????????????????????????????????????????????????????????????????????????????????????????????????????????????????????????????????????????????????????????????????????????????????????????????????????????????????????????????????????????????????????????????????????????????????????????????????????????????????????????????????????????????????????????????????????????????????????????????????????????????????????????????????????????????????????????????????????????????????????????????????????????????????????????????????????????????????????????

			>Encholirium_pulchrum
			??????????????????????????????????????????????????????????????????????????????????????????????????????????????????????????????????????????????????????????????????????????????????????????????????????????????????????????????????????????????????????????????????????????????????????????????????????????????????????????????????????????????????????????????????????????????????????????????????????????????????????????????????????????????????????????????????????????????????????????????????????????????????????????????????????????????????????????????????????????????????????????????????????????????????????????????????????????????????????????????????????????????????????????????????????????????????????????????????????????????????????????????????????????????????????????????

			>Encholirium_scrutor
			ATAATTATAATAAAGTTAAATATGTCGAAAT----CTTGGAATAATACCAAATCTAAAATAAATGTCCGATAGCAAGTTGATCAGTTAATTCAATAAGAGATAAATGGAAGATAGCACTCGGTTTAGTTGGTACCACCCAACCGAATCCGATTCAATCGTTTACTCATTGAATGAGTCAATTTTCAAGTTCAGCCAATC--TTTTTTT--------GAAATTGCAAGTAGATGAAATCGTGAATAAATGTGTTTCATTTCTCTATCATTATAGAAAATAAGATCC-----ATATTATTTATGGAATTCGAACCCGAACTCGATTTATGATTCATTATTTCGATCTTATTGGCCTTTGTTCTTTATTTT-------C-ATATGGATTTCCGTCTTTATATTATACCCTTTC-TGGATGAATTATGCCTCTTTTCACATCTAGGATTTACATATACAACATATATTACTGTCAAGAGGGAATTTTTCTCAG--------TATTTAGATTCAAAATAAGAAAGGGATCAATTCAATTATAAACCTGTAAAACAAAGA-----------TTGGGTTGCGCTATATATATCAAAGAGTATACAATAATGATGTATTTGGTGAATCAAATACATGGTC-----ACAAACCATTTTA-----ACATTTTAATGAGTTGATAATATTAATTGAATATTGGAAA----------------GATT-----TTTCATTCATGCCTAATCCATATCGAGTAGACCTTGTTGTTGTGAG

			>Encholirium_spectabile
			??????????????????????????????????????????????????????????????????????????????????????????????????????????????????????????????????????????????????????????????????????????????????????????????????????????????????????????????????????????????????????????????????????????????????????????????????????????????????????????????????????????????????????????????????????????????????????????????????????????????????????????????????????????????????????????????????????????????????????????????????????????????????????????????????????????????????????????????????????????????????????????????????????????????????????????????????????????????????????????????????????????????????????????????????????????????????????????????????????????????????????????????????????????????????????????????

			>Encholirium_splendidum
			??????????????????????????????????????????????????????????????????????????????????????????????????????????????????????????????????????????????????????????????????????????????????????????????????????????????????????????????????????????????????????????????????????????????????????????????????????????????????????????????????????????????????????????????????????????????????????????????????????????????????????????????????????????????????????????????????????????????????????????????????????????????????????????????????????????????????????????????????????????????????????????????????????????????????????????????????????????????????????????????????????????????????????????????????????????????????????????????????????????????????????????????????????????????????????????????

			>Encholirium_structor
			??????????????????????????????????????????????????????????????????????????????????????????????????????????????????????????????????????????????????????????????????????????????????????????????????????????????????????????????????????????????????????????????????????????????????????????????????????????????????????????????????????????????????????????????????????????????????????????????????????????????????????????????????????????????????????????????????????????????????????????????????????????????????????????????????????????????????????????????????????????????????????????????????????????????????????????????????????????????????????????????????????????????????????????????????????????????????????????????????????????????????????????????????????????????????????????????

			>Encholirium_subsecundum
			??????????????????????????????????????????????????????????????????????????????????????????????????????????????????????????????????????????????????????????????????????????????????????????????????????????????????????????????????????????????????????????????????????????????????????????????????????????????????????????????????????????????????????????????????????????????????????????????????????????????????????????????????????????????????????????????????????????????????????????????????????????????????????????????????????????????????????????????????????????????????????????????????????????????????????????????????????????????????????????????????????????????????????????????????????????????????????????????????????????????????????????????????????????????????????????????

			>Encholirium_vogelii
			??????????????????????????????????????????????????????????????????????????????????????????????????????????????????????????????????????????????????????????????????????????????????????????????????????????????????????????????????????????????????????????????????????????????????????????????????????????????????????????????????????????????????????????????????????????????????????????????????????????????????????????????????????????????????????????????????????????????????????????????????????????????????????????????????????????????????????????????????????????????????????????????????????????????????????????????????????????????????????????????????????????????????????????????????????????????????????????????????????????????????????????????????????????????????????????????

			>Epidryos_allenii
			??????????????????????????????????????????????????????????????????????????????????????????????????????????????????????????????????????????????????????????????????????????????????????????????????????????????????????????????????????????????????????????????????????????????????????????????????????????????????????????????????????????????????????????????????????????????????????????????????????????????????????????????????????????????????????????????????????????????????????????????????????????????????????????????????????????????????????????????????????????????????????????????????????????????????????????????????????????????????????????????????????????????????????????????????????????????????????????????????????????????????????????????????????????????????????????????

			>Fascicularia_bicolor
			ATAATTATAATAAAGTAAAATATGTCGAAATCCTTTTTGGAATAATACCAAATCGAAAATAAATGTCCGATAGCAAGTTGATCAGTTAATTCAATAAGAGATAAATGGGAGATAGCACTCGATTTAGTTGGTACCACCCAACCGAATCCGATTCAATCGTTTACTCATTCAATGAGTAAATTTGCAAGTTCAGCCAATCTTTTTTTTTTTAAAAAAAAAATTGCAAGTAGATGAAATCGTGAATAAATGTGTTTCATTTCTCTATCATTATATAAAATAAGATCCATATTATATTACTTATGGAATTCGAACCCGAACTCGATTTATGATTTATTATTTCGATCTTATTGGCCTTTGTTCTTTATTTTTGATTTTGCATATAGATTTCCGTCTTTATATTATACCCTCTCGTGGATGAATTATGCCTATTTTCACATCTAGGATTTACATATACAACATATATTACTGTCAAGAGGGAATTTTTCTCAGTATTTAGATATTTAGATTCAAAATAAGAAAGGGATCAATTCAATTATAAACCTGCAAAACAAAGATTAGGATTGGGTTGGGTTGCGCTATATATATCAAAGAGTATACAATAATGATGTATTTGGTGAATCAAATACATGGTCTAATAACAAACCATTTTAACATAACATTTTAATGAGTTGATAATATTAATTGAATATTGAATATTTTTTGAAAGATTTTTGTTAAAGGTTTCATTCATGCCTAATCCATATCGAGTAGACCTTGTTGTTGTG??

			>Fernseea_itatiaiae
			ATAATTATAATAAAGTGAAATATGTCGAAATCCTTTTTGGAATAATACCAAATC-AAAATAAATGTCCGATAGCAAGTTGATCAGTTAATTCAATAAGAGATAAATGGGAGATAGCACTCGATTTAGTTGGTACCACCCAACCGAATCCGATTCAATCGTTTACTCATTCAATGAGTAAATTTTAAAGTTCAGCCAATCTTTTTTTTTTGAAAAAAAAAATTGCAAGTAGATGAAATCGTGAATAAATGTGTTTCATTTCTCTATCATTATATAAAAAAAGATCCATATTATATTACTTATGGAATTCGAACCCGAACTCGATTTATGATTTATTATTTCGATCTTATTGGCCTTTGTTCTTTATTTTGGATTTTTCATATAGATTTCCGTCTTTATATTATACCCTTTCGTGGATGAATTATGCCTATTTTCACATCTAGGATTTACATATACAACATATATTACTGTCAAGAGGGAATTTTTCTCAGTATTTAGATATTTAGATTCAAAATAAGAAAGGGATCAATTCAATTATAAACCTGCAAAACAAAGATTAGGATTGGGTTGGGTTGCGCTATATATATCAAAGAGTATACAATAATGATGTATTTGGTGAATCAAATACATGGTCTAATAACAAACCATTTTAACATAACATTTTAATGAGTTGATAATATTAATTGAATA-------TTTTTTGAAAGATTTTTGTTAAAGGTTTCATTCATGCCTAATCCATATCGAGTAGACCTTGTTGTTGTG??

			>Forzzaea_leopoldo-horstii
			??????????????????????????????????????????????????????????????????????????????????????????????????????????????????????????????????????????????????????????????????????????????????????????????????????????????????????????????????????????????????????????????????????????????????????????????????????????????????????????????????????????????????????????????????????????????????????????????????????????????????????????????????????????????????????????????????????????????????????????????????????????????????????????????????????????????????????????????????????????????????????????????????????????????????????????????????????????????????????????????????????????????????????????????????????????????????????????????????????????????????????????????????????????????????????????????

			>Forzzaea_warasii
			??????????????????????????????????????????????????????????????????????????????????????????????????????????????????????????????????????????????????????????????????????????????????????????????????????????????????????????????????????????????????????????????????????????????????????????????????????????????????????????????????????????????????????????????????????????????????????????????????????????????????????????????????????????????????????????????????????????????????????????????????????????????????????????????????????????????????????????????????????????????????????????????????????????????????????????????????????????????????????????????????????????????????????????????????????????????????????????????????????????????????????????????????????????????????????????????

			>Fosterella_albicans
			??????????????????????????????????????????????????????????????????????????????????CAGTTAATTCAATAAGAGATAAATGGGCGATAGCACTCGGTTTAGTTGGTATCACCCAACCGAATCCGATTCAATCGTTTACTCATTCAATGAGTCAATTTTCAAGTTCAGCCAATC--TTTTTTTTTAAAAAAAAAATTGCAAGTAGATGAAATCGTGAATAAATGTGTTTCATTTCTCTATCATTATAGAAAATAGGATCC-----ATATTACTTATGGAATTCGAACCCGAACTCGATTTATGATTCATTATTTCGATCTTATTGGCCTTTGTTCTTTATTTTTGATTTTTCATATTGATTTCCGTCTTTATATTATACCCTTTCGTGGATGAATTATGCCTATTTTCACATCTAGGATTTACATATACAACATATATTACTGTCAAGAGGGAATTTTTCTCAGTATTTAGATATTTAGATTCAAAATAAGAAAGGGATCAATTCAATTATAAACCTGTAAAACAAGGATTAGGA-----TTGGGTTGCGCTATATATATCAAAGAGTATACAATAATGATGTATTTGGTGAATCAAATACATGGTCTAATAACAAACCATTTTA-----ACATTTTAATGAGTTGATAATATTAATTGAATATTGAATATTTTTTGAAAGATTTTTGTTAAAGGTTTCATTCATGCCTAATCCATATCGAGTAGACCTTGTTGTTGTGAG

			>Fosterella_batistana
			??????????????????????????????????????????????????????????????????????????????????????????????????????????????????????????????????????????????????????????????????????????????????????????????????????????????????????????????????????????????????????????????????????????????????????????????????????????????????????????????????????????????????????????????????????????????????????????????????????????????????????????????????????????????????????????????????????????????????????????????????????????????????????????????????????????????????????????????????????????????????????????????????????????????????????????????????????????????????????????????????????????????????????????????????????????????????????????????????????????????????????????????????????????????????????????????

			>Fosterella_caulescens
			??????????????????????????????????????????????????????????????????????????????????CAGTTAATTCAATAAGAGATAAATGGGCGATAGCACTCGGTTTAGTTGGTATCACCCAACCGAATCCGATTCAATCGTTTACTCATTCAATGAGTCAATTTTCAAGTTCAGCCAATC--TTTTTTTTTAAAAAAAAAATTGCAAGTAGATGAAATCGTGAATAAATGTGTTTCATTTCTCTATCATTATAGAAAATAGGATCC-----ATATTACTTATGGAATTCGAACCCGAACTCGATTTATGATTCATTATTTCGATCTTATTGGCCTTTGTTCTTTATTTTTGATTTTTCATATTGATTTCCGTCTTTATATTATACCCTTTCGTGGATGAATTATGCCTATTTTCACATCTAGGATTTACATATACAACATATATTACTGTCAAGAGGGAATTTTTCTCAGTATTTAGATATTTAGATTCAAAATAAGAAAGGGATCAATTCAATTATAAACCTGTAAAACAAGGATTAGGA-----TTGGGTTGCGCTATATATATCAAAGAGTATACAATAATGATGTATTTGGTGAATCAAATACATGGTCTAATAACAAACCATTTTA-----ACATTTTAATGAGTTGATAATATTAATTGAATATTGAATATTTTTTGAAAGATTTTTGTTAAAGGTTTCATTCATGCCTAATCCATATCGAGTAGACCTTGTTGTTGTGAG

			>Fosterella_christophii
			??????????????????????????????????????????????????????????????????????????????????CAGTTAATTCAATAAGAGATAAATGGGCGATAGCACTCGATTTAGTTGGTATCACCCAACCGAATCCGATTCAATCGTTTACTCATTCAATGAGTCAATTTGCAAGTTCAGCCAATCCTTTTTTTTTTAAAAAAAAAATTGCAAGTAGATGAAATCGTGAAAAAATGTGTTTCATTTCTCTATCATTATAGAAAA-----------------------------------------------------------------------------------------TTTGATTTTTCATATTGATTTCCGTCTTTATATTATACCCTTTCGTGGATGAATTATGCCTATTTTCACATCTAGGATTTACATATACAACATATATTACTGTCAAGAGGGAATTTTTCTCAGTATTTAGATATTTAGATTCAAAATAAGAAAGGGATCAATTCAATTATAAACCTGTAAAACAAGGATTAGGA-----TTGGGTTGCGCTATATATATCAAAGAGTATACAATAATGATGTATTTGGTGAATCAAATACATGGTCTAATAACAAACCATTTTA-----ACATTTTAATGAGTTGATAATATTAATTGAATATTGAATATCTTTTGAAAGATTTTTGTTAAAGGTTTCATTCATGCCTAATCCATATCGAGTAGACCTTGTTGTTGTGAG

			>Fosterella_cotacajensis
			??????????????????????????????????????????????????????????????????????????????????CAGTTAATTCAATAAGAGATAAATGGGCGATAGCACTCGGTTTAGTTGGTATCACCCAACCGAATCCGATTCAATCGTTTACTCATTCAATGAGTCAATTTTCAAGTTCAGCCAATC--TTTTTTTTT--------AATTTCAAGTAGATGAAATCGTGAATAAATGTGTTTCATTTCTCTATCATTATAGAAAATAGGATCC-----ATATTACTTATGGAATTCGAACCCGAACTCGATTTATGATTCATTATTTCGATCTTATTGGCCTTTGCTCTTTATTTTTGATTTTTCATATTGATTTCCGTCTTTATATTATACCCTTTCGTGGATGAATTATGCCTATTTTCACATCTAGGATTTACATATACAACATATATTACTGTCAAGAGGGAATTTTTCTCAGTATTTAGATATTTAGATTCAAAATAAGAAAGGGATCAATTCAATTATAAACCTATAAAACAAGGATTAGGA-----TTGGGTTGCGCTATATATATCAAAGAGTATACAATAATGATGTATTTGGTGAATCAAATACATGGTCTAATAACAAACCATTTTA-----ACATTTTAATGAGTTGATAATATTAATTGAATATTGAATATTTTTTGAAAGATTTTTGTTAAAGGTTTCATTCATGCCTAATCCATATCGAGTAGACCTTGTTGTTGTGAG

			>Fosterella_floridensis
			??????????????????????????????????????????????????????????????????????????????????CAATTAATTCAATAAGAGATAAATGGGCGATAGCACTCGGTTTAGTTGGTATCACCCAACCGAATCCGATTCAATCGTTTACTCATTCAATGAGTCAATTTTCAAGTTCAGCCAATC--TTTTTTTTGAAAAAAAAAATTGCAAGTAGATGAAATCGTGAATAAATGTGTTTCATTTCTCTATCATTATAGAAAATGGGATCC-----ATATTACTTATGGAATTCGAACCCGAACTCGATTTATGATTCATTATTTCGATCTTATTGGCCTTTGTTCTTTATTTTTGATTTTTCATATTGATTTCCGTCTTTATATTATACCCTTTCGTGGATGAATTATGCCTATTTTCACATCTAGGATTTACATATACAACATATATTACTGTCAAGAGGGAATTTTTCTCAGTATTTAGATATTTAGATTCAAAATAAGAAAGGGATCAATTCAATTATAAACCTGTAAAACAAGGATTAGGA-----TTGGGTTGCGCTATATATATCAAAGAGTATACAATAATGATGTATTTGGTGAATCAAATACATGGTCTAATAACAAACCATTTTA-----ACATTTTAATGAGTTGATAATATTAATTGAATATTCAATATTTTTTGAAAGATTTTTGTTAAAGGTTTCATTCATGCCTAATCCATATCGAGTAGACCTTGTTGTTGTGAG

			>Fosterella_gracilis
			??????????????????????????????????????????????????????????????????????????????????CAGTTAATTCAATAAGAGATAAATGGGCGATAGCACTCGGTTTAGTTGGTATCACCCAACCGAATCCGATTCAATCGTTTACTCATTCAATGAGTCAATTTTCAAGTTCAGCCAATC--TTTTTTTTT------TTAATTGCAAATAGATGAAATCGTGAATAAATGTGTTTCATTTCTCTATCATTATAGAAAATAGGATCC-----ATATTACTTATGGAATTCGAACCCGAACTCGATTTATGATTCATTATTTCGATCTTATTGGCCTTTGTTCTTTATTTTTTATTTTTCATATTGATTTCCGTCTTTATATTATACCCTTTCGTGGATGAATTATGCCTATTTTCACATCTAGGATTTACATATACAACATATATTACTGTCAAGAGGGAATTTTTCTCAGTATTTAGATATTTAGATTCAAAAAAAGAAAGGGATCAATTCAATTATAAACTTGTAAAACAAGGATTAGGA-----TTGGGTTGCGCTATATATATCAAAGAGTATACAATAATGATGTATTTGGTGAATCAAATACATGGTCTAATAACAAACCATTTTA-----ACATTTTAATGAGTTGATAATATTAATTGAATATTGAATATTTTTTGAAATATTTTTGTTAAAGGTTTCATTCATGCCTAATCCATATCGGGTAGACCTTGTTGTTGTGAG

			>Fosterella_graminea
			??????????????????????????????????????????????????????????????????????????????????CAGTTAATTCAATAAGAGATAAATGGGCGATAGCACTCGGTTTAGTTGGTATCACCCAACCGAATCCGATTCAATCGTTTACTCATTCAATGAGTCAATTTTCAAGTTCAGCCAATC--TTTTTTTTTAAAAAAAAAATTGCAAGTAGATGAAATCGTGAATAAATGTGTTTCATTTCTCTATCATTATAGAAAATAGGATCC-----ATATTACTTATGGAATTCGAACCCGAACTCGATTTTTGATTCATTATTTCGATCTTATTGGCCTTTGTTCTTTATTTTTGATTTTTCATATTGATTTCCGTCTTTATATTATACCCTTTCGTGGATGAATTATGCCTATTTTCACATCTAGGATTTACATATACAACATATATTACTGTCAAGAGGGAATTTTTCTCAGTATTTAGATATTTAGATTCAAAATAAGAAAGGGATCAATTCAATTATAAACCTGTAAAACAAGGATTAGGA-----TTGGGTTGCGCTATATATATCAAAGAGTATACAATAATGATGTATTTGGTGAATCAAATACATGGTCTAATAACAAACCATTTTA-----ACATTTTAATGAGTTGATAATATTAATTGAATATTGAATATTTTTTGAAAGATTTTTGGTAAAGGTTTCATTCATGCCTAATCCATATCGAGTAGACCTTGTTGTTGTGAG

			>Fosterella_hatschbachii
			??????????????????????????????????????????????????????????????????????????????????CAATTAATTCAATAAGAGATAAATGGGCGATAGCACTCGGTTTAGTTGGTATCACCCAACCGAATCCGATTCAATCGTTTACTCATTCAATGAGTAAATTTTCAAGTTCAGCCAATC--TTTTTTTTT---------------------------------------------------------------------------------------------------------------------------------------------------------------------TCATATTGATTTCCGTCTTTATATTATACCCTTTCGTGGATGAATTATGCCTATTTTCACATCTAGGATTTACATATACAACATATATTACTGTCAAGAGGGAATTTTTCTCAGTATTTAGATATTTAGATTCAAAATAAGAAAGGGATCAATTCAATTATAAACCTGTAAAACAAGGATTAGGA-----TTGGGTTGCGCTATATATATCAAAGAGTATACAATAATGATGTATTTGGTGAATCAAATACATGGTCTAATAACAAACCATTTTA-----ACATTTTAATGAGTTGATAATATTAATTGAATATTCAATATTTTTTGAAAGATTTTTGTTAAAGGTTTCATTCATGCCTAATCCATATCGAGTAGACCTTGTTGTTGTGAG

			>Fosterella_heterophylla
			??????????????????????????????????????????????????????????????????????????????????CAGTTAATTCAATAAGAGATAAATGGGCGATAGCACTCGGTTTAGTTGGTATCACCCAACCGAATCCGATTCAATCGTTTACTCATTCAATGAGTCAATTTTCAAGTTCAGCCAATC--TTTTTTTTTAAAAAAAAAATTGCAAGTAGATGAAATCGTGAATAAATGTGTTTCATTTCTCTATCATTATAGAAAATAGGATCC-----ATATTACTTATGGAATTCGAACCCGAACTCGATTTATGATTCATTATTTCGATCATATTGGCCTTTGTTCTTTATTTTTGATTTTTCATATTGATTTCCGTCTTTATATTATACCCTCTCGTGGATGAATTATGCCTATTTTCACATCTAGGATTTACATATACAACATATATTACTGTCAAGAGGGAATTTTTCTCAGTATTTAGATATTTAGATTCAAAATAAGAAAGGGATCAATTCAATTATAAACCTGTAAAACAAGGATTAGGA-----TTGGGTTGCGCTATATATATCAAAGAGTATACAATAATGATGTATTTGGTGAATCAAATACATGGTCTAATAACAAACCATTTTA-----ACATTTTAATGAGTTGATAATATTAATTGAATATTGAATATTTTTTGAAAGATTTTTGTTAAAGGTTTCATTCATGCCTAATCCATATCGAGTAGACCTTGTTGTTGTGAG

			>Fosterella_kroemeri
			??????????????????????????????????????????????????????????????????????????????????????????????????????????????????????????????????????????????????????????????????????????????????????????????????????????????????????????????????????????????????????????????????????????????????????????????????????????????????????????????????????????????????????????????????????????????????????????????????????????????????????????????????????????????????????????????????????????????????????????????????????????????????????????????????????????????????????????????????????????????????????????????????????????????????????????????????????????????????????????????????????????????????????????????????????????????????????????????????????????????????????????????????????????????????????????????

			>Fosterella_micrantha
			??????????????????????????????????????????????????????????????????????????????????CAGTTAATTCAATAAGAGATAAATGGGCGATAGCACTCGATTTAGTTGGTATCACCCAACCGAATCCGATTCAATCGTTTACTCATTCAATGAGTCAATTTGCAAGTTCAGCCAATCCTTTTTTTTTTAAAAAAAAAATTGCAAGTAGATGAAATCGTGAAAAAATGTGTTTCATTTCTCTATCATTATAGAAAA-----------------------------------------------------------------------------------------TTTGATTTTTCATATTGATTTCCGTCTTTATATTATATCCTTTCGTGGATGAATTATGCCTATTTTCACATCTAGGATTTACATATACAACATATATTACTGTCAAGAGGGAATTTTTCTCAGTATTTAGATATTTAGATTCAAAATAAGAAAGGGATCAATTCAATTATAAACCTGTAAAACAAGGATTAGGA-----TTGGGTTGCGCTATATATATCAAAGAGTATACAATAATGATGTATTTGGTGAATCAAATACATGGTCTAATAACAAACCATTTTA-----ACATTTTAATGAGTTGATAATATTAATTGAATATTGAATATCTTTTGAAAGATTTTTGTTAAAGGTTTCATTCATGCCTAATCCATATCGAGTAGACCTTGTTGTTGTGAG

			>Fosterella_penduliflora
			??????????????????????????????????????????????????????????????????????????????????CAGTTAATTCAATAAGAGATAAATGGGCGATAGCACTCGGTTTAGTTGGTATCACCCAACCGAATCCGATTCAATCGTTTACTCATTCAATGAGTCAATTTTCAAGTTCAGCCAATC--TTTTTTTTT-------TAATTGCAAATAGATGAAATCGTGAATAAATGTGTTTCATTTCTCTATCATTATAGAAAATAGGATCC-----ATATTACTTATGGAATTCGAACCCGAACTCGATTTATGATTCATTATTTCGATCTTATTGGCCTTTGTTCTTTATTTTTGATTTTTCATATTGATTTCCGTCTTTATATTATACCCTTTCGTGGATGAATTATGCCTATTTTCACATCTAGGATTTACATATACAACATATATTACTGTCAAGAGGGAATTTTTCTCAGTATTTAGATATTTAGATTCAAAAAAAGAAAGGGATCAATTCAATTATAAACTTGTAAAACAAGGATTAGGA-----TTGGGTTGCGCTATATATATCAAAGAGTATACAATAATGATGTATTTGGTGAATCAAATACATGGTCTAATAACAAACCATTTTA-----ACATTTTAATGAGTTGATAATATTAATTGAATATTGAATATTTTTTGAAATATTTTTGTTAAAGGTTTCATTCATGCCTAATCCATATCGAGTAGACCTTGTTGTTGTCAG

			>Fosterella_petiolata
			??????????????????????????????????????????????????????????????????????????????????CAGTTAATTCAATAAGAGATAAATGGGCGATAGCACTCGGTTTAGTTGGTATCACCCAACCGAATCCGATTCAATCGTTTACTCATTCAATGAGTCAATTTTCAAGTTCAGCCAATC--TTTTTTTTTAAAAAAAAAATTGCAAGTAGATGAAATCGTGAATAAATGTGTTTCATTTCTCTATCATTATAGAAAATAGGATCC-----ATATTACTTATGGAATTCGAACCCGAACTCGATTTATGATTCATTATTTCGATCATATTGGCCTTTGTTCTTTATTTTTGATTTTTCATATTGATTTCCGTCTTTATATTATACCCTCTCGTGGATGAATTATGCCTATTTTCACATCTAGGATTTACATATACAACATATATTACTGTCAAGAGGGAATTTTTCTCAGTATTTAGATATTTAGATTCAAAATAAGAAAGGGATCAATTCAATTATAAACCTGTAAAACAAGGATTAGGA-----TTGGGTTGCGCTATATATATCAAAGAGTATACAATAATGATGTATTTGGTGAATCAAATACATGGTCTAATAACAAACCATTTTA-----ACATTTTAATGAGTTGATAATATTAATTGAATATTGAATATTTTTTGAAAGATTTTTGTTAAAGGTTTCATTCATGCCTAATCCATATCGAGTAGACCTTGTTGTTGTGAG

			>Fosterella_rexiae
			??????????????????????????????????????????????????????????????????????????????????CAGTTAATTCAATAAGAGATAAATGGGCGATAGCACTCGGTTTAGTTGGTATCACCCAACCGAATCCGATTCAATCGTTTACTCATTCAATGAGTCAATTTTCAAGTTCAGCCAATC--TTTTTTTTTAAAAAAAAAATTGCAAGTAGATGAAATCGTGAATAAATGTGTTTCATTTCTCTATCATTATAGAAAATAGGATCC-----ATATTACTTATGGAATTCGAACCCGAACTCGATTTATGATTCATTATTTCGATCTTATTGGCCTTTGTTCTTTATTTTTGATTTTTCATATTGATTTCCGTCTTTATATTATACCCTTTCGTGGATGAATTATGCCTATTTTCACATCTAGGATTTACATATACAACATATATTACTGTCAAGAGGGAATTTTTCTCAGTATTTAGATATTTAGATTCAAAATAAGAAAGGGATCAATTCAATTATAAACCTGTAAAACAAGGATTAGGA-----TTGGGTTGCGCTATATATATCAAAGAGTATACAATAATGATGTATTTGGTGAATCAAATACATGGTCTAATAACAAACCATTTTA-----ACATTTTAATGAGTTGATAATATTAATTGAATATTGAATATTTTTTGAAAGATTTTTGTTAAAGGTTTCATTCATGCCTAATCCATATCGAGTAGACCTTGTTGTTGTGAG

			>Fosterella_robertreadii
			??????????????????????????????????????????????????????????????????????????????????CAGTTAATTCAATAAGAGATAAATGGGCGATAGCACTCGGTTTAGTTGGTATCACCCAACCGAATCCGATTCAATCGTTTACTCATTCAATGAGTCAATTTTCAAGTTCAGCCAATC--TTTTTTTTTAAAAAAAAAATTGCAAGTAGATGAAATCGTGAATAAATGTGTTTCATTTCTCTATCATTATAGAAAATAGGATCC-----ATATTACTTATGGAATTCGAACCCGAACTCGATTTTTGATTCATTATTTCGATCTTATTGGCCTTTGTTCTTTATTTTTGATTTTTCATATTGATTTCCGTCTTTATATTATACCCTTTCGTGGATGAATTATGCCTATTTTCACATCTAGGATTTACATATACAACATATATTACTGTCAAGAGGGAATTTTTCTCAGTATTTAGATATTTAGATTCAAAATAAGAAAGGGATCAATTCAATTATAAACCTGTAAAACAAGGATTAGGA-----TTGGGTTGCGCTATATATATCAAAGAGTATACAATAATGATGTATTTGGTGAATCAAATACATGGTCTAATAACAAACCATTTTA-----ACATTTTAATGAGTTGATAATATTAATTGAATATTGAATATTTTTTGAAAGATTTTTGTTAAAGGTTTCATTCATGCCTAATCCATATCGAGTAGACCTTGTTGTTGTGAG

			>Fosterella_rusbyi
			??????????????????????????????????????????????????????????????????????????????????CAGTTAATTCAATAAGAGATAAATGGGCGATAGCACTCGGTTTAGTTGGTATCACCCAACCGAATCCGATTCAATCGTTTACTCATTCAATGAGTCAATTTTCAAGTTCAGCCAATC--TTTTTTTTTAAAAAAAAAATTGCAAGTAGATGAAATCGTGAATAAATGTGTTTCATTTCTCTATCATTATAGAAAATAGGATCC-----ATATTACTTATGGAATTCGAACCCGAACTCGATTTTTGATTCATTATTTCGATCTTATTGGCCTTTGTTCTTTATTTTTGATTTTTCATATTGATTTCCGTCTTTATATTATACCCTTTCGTGGATGAATTATGCCTATTTTCACATCTAGGATTTACATATACAACATATATTACTGTCAAGAGGGAATTTTTCTCAGTATTTAGATATTTAGATTCAAAATAAGAAAGGGATCAATTCAATTATAAACCTGTAAAACAAGGATTAGGA-----TTGGGTTGCGCTATATATATCAAAGAGTATACAATAATGATGTATTTGGTGAATCAAATACATGGTCTAATAACAAACCATTTTA-----ACATTTTAATGAGTTGATAATATTAATTGAATATTGAATATTTTTTGAAAGATTTTTGTTAAAGGTTTCATTCATGCCTAATCCATATCGAGTAGACCTTGTTGTTGTGAG

			>Fosterella_schidosperma
			??????????????????????????????????????????????????????????????????????????????????CAGTTAATTCAATAAGAGATAAATGGGCGATAGCACTCGGTTTAGTTGGTATCACCCAACCGAATCCGATTCAATCGTTTACTCATTCAATGAGTCAATTTTCAAGTTCAGCCAATC--TTTTTTTTTAAAAAAAAAATTGCAAGTAGATGAAATCGTGAATAAATGTGTTTCATTTCTCTATCATTATAGAAAATAGGATCC-----ATATTACTTATGGAATTCGAACCCGAACTCGATTTTTGATTCATTATTTCGATCTTATTGGCCTTTGTTCTTTATTTTTGATTTTTCATATTGATTTCCGTCTTTATATTATACCCTTTCGTGGATGAATTATGCCTATTTTCACATCTAGGATTTACATATACAACATATATTACTGTCAAGAGGGAATTTTTCTCAGTATTTAGATATTTAGATTCAAAATAAGAAAGGGATCAATTCAATTATAAACCTGTAAAACAAGGATTAGGA-----TTGGGTTGCGCTATATATATCAAAGAGTATACAATAATGATGTATTTGGTGAATCAAATACATGGTCTAATAACAAACCATTTTA-----ACATTTTAATGAGTTGATAATATTAATTGAATATTGAATATTTTTTGAAAGATTTTTGTTAAAGGTTTCATTCATGCCTAATCCATATCGAGTAGACCTTGTTGTTGTGAG

			>Fosterella_spectabilis
			??????????????????????????????????????????????????????????????????????????????????CAATTAATTCAATAAGAGATAAATGGGCGATAGCACTCGGTTTAGTTGGTATCACCCAACCGAATCCGATTCAATCGTTTACTCATTCAATGAGTCAATTTTCAAGTTCAGCCAATC--TTTTTTTTTAAAAAAAAAATTGCAAGTAGATGAAATCGTGAATAAATGTGTTTCATTTCTCTATCATTATAGAAAATGGGATCC-----ATATTACTTATGGAATTCGAACCCGAACTCGATTTATGATTCATTATTTCGATCTTATTGGCCTTTGTTCTTTATTTTTGATTTTTCATATTGATTTCCGTCTTTATATTATACCCTTTCGTGGATGAATTATGCCTATTTTCACATCTAGGATTTACATATACAACATATATTACTGTCAAGAGGGAATTTTTCTCAGTATTTAGATATTTAGATTCAAAATAAGAAAGGGATCAATTCAATTATAAACCTGTAAAACAAGGATTAGGA-----TTGGGTTGCGCTATATATATCAAAGAGTATACAATAATGATGTATTTGGTGAATCAAATACATGGTCTAATAACAAACCATTTTA-----ACATTTTAATGAGTTGATAATATTAATTGAATATTCAATATTTTTTGAAAGATTTTTGTTAAAGGTTTCATTCATGCCTAATCCATATCGAGTAGACCTTGTTGTTGTGAG

			>Fosterella_vasquezii
			??????????????????????????????????????????????????????????????????????????????????CAATTAATTCAATAAGAGATAAATGGGCGATAGCACTCGGTTTAGTTGGTATCACCCAACCGAATCCGATTCAATCGTTTACTCATTCAATGAGTAAATTTTCAAGTTCAGCCAATC--TTTTTTTTGAAAAAAAAAATTGCAAGTAGATGAAATCGTGAATAAATGTGTTTCATTTCTCTATCATTATAGAAAATGGGATCC-----ATATTACTTATGGAATTCGAACCCGAACTCGATTTATGATTCATTATTTCGATCTTATTGGCCTTTGTTCTTTATTTTTGATTTTTCATATTGATTTCCGTCTTTATATTATACCCTTTCGTGGATGAATTATGCCTATTTTCACATCTAGGATTTACATATACAACATATATTACTGTCAAGAGGGAATTTTTCTCAGTATTTAGATATTTAGATTCAAAATAAGAAAGGGATCAATTCAATTATAAACCTGTAAAACAAGGATTAGGA-----TTGGGTTGCGCTATATATATCAAAGAGTATACAATAATGATGTATTTGGTGAATCAAATACATGGTCTAATAACAAACCATTTTA-----ACATTTTAATGAGTTGATAATATTAATTGAATATTCAATATTTTTTGAAAGATTTTTGTTAAAGGTTTCATTCATGCCTAATCCATATCGAGTAGACCTTGTTGTTGTGAG

			>Fosterella_villosula
			??????????????????????????????????????????????????????????????????????????????????CAGTTAATTCAATAAGAGATAAATGGGCGATAGCACTCGATTTAGTTGGTATCACCCAACCGAATCCGATTCAATCGTTTACTCATTCAATGAGTCAATTTGCAAGTTCAGCCAATCTTTTTTTTTTTAAAAAAAAAATTGCAAGTAGATGAAATCGTGAAAAAATGTGTTTCATTTCTCTATCATTATAGAAAA-----------------------------------------------------------------------------------------TTTGATTTTTCATATTGATTTCCGTCTTTATATTATATCCTTTCGTGGATGAATTATGCCTATTTTCACATCTAGGATTTACATATACAACATATATTACTGTCAAGAGGGAATTTTTCTCAGTATTTAGATATTTAGATTCAAAATAAGAAAGGGATCAATTCAATTATAAACCTGTAAAACAAGGATTAGGA-----TTGGGTTGCGCTATATATATCAAAGAGTATACAATAATGATGTATTTGGTGAATCAAATACATGGTCTAATAACAAACCATTTTA-----ACATTTTAATGAGTTGATAATATTAATTGAATATTGAATATCTTTTGAAAGATTTTTGTTAAAGGTTTCATTCATGCCTAATCCATATCGAGTAGACCTTGTTGTTGTGAG

			>Fosterella_weberbaueri
			??????????????????????????????????????????????????????????????????????????????????CAGTTAATTCAATAAGAGATAAATGGGCGATAGCACTCGATTTAGTTGGTATCACCCAACCGAATCCGATTCAATCGTTTACTCATTCAATGAGTCAATTTGCAAGTTCAGCCAATCTTTTTTTTTTTAAAAAAAAAATTGCAAGTAGATGAAATCGTGAATAAATGTGTTTCATTTCTCTATCATTATAGAAAATAGGATCC-----ATATTACTTATGGAATTCGAACCCGAACTCGATTTATGATTCATTATTTCGATCTTATTGGCCTTTGTTCTTTATTTTTGATTTTTCATATTGATTTCCGTCTTTATATTATACCCTTTCGTGGATGAATTATGCCTATTTTCACATCTAGGATTTACATATACAACATATATTACTGTCAAGAGGGAATTTTTCTCAGTATTTAGATATTTAGATTCAAAATAAGAAAGGGATCAATTCAATTATAAACCTGTAAAACAAGGATTAGGA-----TTGGGTTGCGCTATATATATCATAGAGTATACAATAATGATGTATTTGGTGAATCAAATACATGGTCTAATAACAAACCATTTTA-----ACATTTAAATAAGTTGATAATATTAATTGAATATTGAATATTTTTTGAAAGATTTTTGTTAAAGGTTTCATTCATGCCTAATCCATATCGAGTAGACCTTGTTGTTGTGAG

			>Fosterella_weddelliana
			??????????????????????????????????????????????????????????????????????????????????CAGTTAATTCAATAAGAGATAAATGGGCGATAGCACTCGGTTTAGTTGGTATCACCCAACCGAATCCGATTCAATCGTTTACTCATTCAATGAGTCAATTTTCAAGTTCAGCCAATC--TTTTTTTTT--------AATTTCAAGTAGATGAAATCGTGAATAAATGTGTTTCATTTCTCTATCATTATAGAAAATAGGATCC-----ATATTACTTATGGAATTCGAACCCGAACTCGATTTATGATTCATTATTTCGATCTTATTGGCCTTTGCTCTTTATTTTTGATTTTTCATATTGATTTCCGTCTTTATATTATACCCTTTCGTGGATGAATTATGCCTATTTTCACATCTAGGATTTACATATACAACATATATTACTGTCAAGAGGGAATTTTTCTCAGTATTTAGATATTTAGATTCAAAATAAGAAAGGGATCAATTCAATTATAAACCTATAAAACAAGGATTAGGA-----TTGGGTTGCGCTATATATATCAAAGAGTATACAATAATGATGTATTTGGTGAATCAAATACATGGTCTAATAACAAACCATTTTA-----ACATTTTAATGAGTTGATAATATTAATTGAATATTGAATATTTTTTGAAAGATTTTTGTTAAAGGTTTCATTCATGCCTAATCCATATCGAGTAGACCTTGTTGTTGTGAG

			>Fosterella_windischii
			??????????????????????????????????????????????????????????????????????????????????CAATTAATTCAATAAGAGATAAATGGGCGATAGCACTCGGTTTAGTTGGTATCACCCAACCGAATCCGATTCAATCGTTTACTCATTCAATGAGTAAATTTTCAAGTTCAGCCAATC--TTTTTTTTGAAAAAAAAAATTGCAAGTAGATGAAATCGTGAATAAATGTGTTTCATTTCTCTATCATTATAGAAAATGGGATCC-----ATATTACTTATGGAATTCGAACCCGAACTCGATTTATGATTCATTATTTCGATCTTATTGGCCTTTGTTCTTTATTTTTGATTTTTCATATTGATTTCCGTCTTTATATTATACCCTTTCGTGGATGAATTATGCCTATTTTCACATCTAGGATTTACATATACAACATATATTACTGTCAAGAGGGAATTTTTCTCAGTATTTAGATATTTAGATTCAAAATAAGAAAGGGATCAATTCAATTATAAACCTGTAAAACAAGGATTAGGA-----TTGGGTTGCGCTATATATATCAAAGAGTATACAATAATGATGTATTTGGTGAATCAAATACATGGTCTAATAACAAACCATTTTA-----ACATTTTAATGAGTTGATAATATTAATTGAATATTCAATATTTTTTGAAAGATTTTTGTTAAAGGTTTCATTCATGCCTAATCCATATCGAGTAGACCTTGTTGTTGTGAG

			>Fosterella_yuvinkae
			??????????????????????????????????????????????????????????????????????????????????CAATTAATTCAATAAGAGATAAATGGGCGATAGCACTCGGTTTAGTTGGTATCACCCAACCGAATCCGATTCAATCGTTTACTCATTCAATGAGTAAATTTTCAAGTTCAGCCAATC--TTTTTTTTGAAAAAAAAAATTGCAAGTAGATGAAATCGTGAATAAATGTGTTTCATTTCTCTATCATTATAGAAAATGGGATCC-----ATATTACTTATGGAATTCGAACCCGAACTCGATTTATGATTCATTATTTCGATCTTATTGGCCTTTGTTCTTTATTTTTGATTTTTCATATTGATTTCCGTCTTTATATTATACCCTTTCGTGGATGAATTATGCCTATTTTCACATCTAGGATTTACATATACAACATATATTACTGTCAAGAGGGAATTTTTCTCAGTATTTAGATATTTAGATTCAAAATAAGAAAGGGATCAATTCAATTATAAACCTGTAAAACAAGGATTAGGA-----TTGGGTTGCGCTATATATATCAAAGAGTATACAATAATGATGTATTTGGTGAATCAAATACATGGTCTAATAACAAACCATTTTA-----ACATTTTAATGAGTTGATAATATTAATTGAATATTCAATATTTTTTGAAAGATTTTTGTTAAAGGTTTCATTCATGCCTAATCCATATCGAGTAGACCTTGTTGTTGTGAG

			>Glomeropitcairnia_erectiflora
			ATAATTATAATAAAGTGAAATATGTCGAAATCCTTTTTGGAATAATACCAAATC-AAAATAAATGTCCGATAGCAAGTTGATCAGTTAATTCAATAAGAGATAAATGGGAGATAGCACTCGATTTAGTTGGTACCACCCAACCGAATCCGATTCAATCGTTTACTCATTCAATGAGTAAATTTGCAAGTTCAGCCAATCTTCTTTTTTTTGAAAAAGAAATTGCAAGTAGATGAAATCGTGAATAAATGTGTTTCATTTCTCTATCATTATAGAAAATAAGATCCATATTATATTACTTATGGAATTCGAACCCGAACTCGATTTATGATTCATTATTTCGATCTTATTGGCCTTTGTTCTTTATTTTTTATTTTTCATATTGATTTCCGTCTTTATATTATACCCTTTCGTGGATGAATTATGCTTATTTTCACATCTAGGATTTACATATACAACATATATTACTGTCAAGAGGGAATTTTTCTCAGTATTTAGATATTTAGATTCAAAATAAGAAAGGGATCAATTCAATTATAAACCTGCAAAGCAAAGATTAGGA-----TTGGGTTGCGCTATATATATCAAAGAGTATACAATAATGATGTATTTGGTGAATCAAATACATGGTCTAATAACAAACCATTTTAACATAACATTTTAATGAGTTGATAATATTAATTGAATATTGAATATCTTTTGAAAGATTTTTGTTAAAGGTTTCATTCATGCCTAATCCATATCGAGTAGACCTTGTTGTTGTGAG

			>Glomeropitcairnia_penduliflora
			ATAATTATAATAAAGTGAAATATGTCGAAATCCTTTTTGGAATAATACCAAATC-AAAATGAATGTCCGATAGCAAGTTGATCAGTTAATTCAATAAGAGATAAATGGGAGATAGCACTCGATTTAGTTGGTACCACCCAACCGAATCCGATTCAATCGTTTACTCATTCAATGAGTCAATTTTCAAGTTCAGCCAATC--TTTTTTTTTGAAAAAGAAATTGCAAGTAGATGAAATCGTGAATAAATGTGTTTCATTTCTCTATCATTATAGAAAATAAGATCCATATTATATTACTTATGGAATTCGAACCCGAACTCGATTTATGATTCATTATTTCGATCTTATTGGCCTTTGTTCTTTATTTTTTATTTTTCATATTGATTTCCGTCTTTATATTATACCCTTTCGTGGATGAATTATGCTTATTTTCACATCTAGGATTTACATATACAACATATATTACTGTCAAGAGGGAATTTTTCTCAGTATTTAGATATTTAGATTCAAAATAAGAAAGGGATCAATTCAATTATAAACCTGCAAAGCAAAGATTAGGA-----TTGGGTTGCGCTATATATATCAAAGAGTATACAATAATGATGTATTTGGTGAATCAAATACATGGTCTAATAACAAACCATTTTAACATAACATTTTAATGAGTTGATAATATTAATTGAATATTGAATATCTTTTGAAAGATTTTTGTTAAAGGTTTCATTCATGCCTAATCCATATCGAGTAGACCTTGTTGTTGTGAG

			>Goudaea_chrysostachys
			ATAATTATAATAAAGTGAAATATGTCGAAATCCTTTTTGGAATAATACCAAATC-AAAATAAATGTCCGATAGCAAGTTGATCAGTTAATTCAATAAGAGATAAATGGGAGATAGCACTCGATTTAGTTGGTACCACCCAACCGAATCCGATTCAATCGTTTACTCATTCAATGAGTCAATTTTCAAGTTCAGCCAATCTTTTTTTTTTCAAAAAAAAAATTGCAAGTAAATGAAATCGTGAATAAATGTGTTTCATTTCTCTATCATTATAGAAAAAAAGATCCATATTATATTACTTATGGAATTCGAACCCGAACTCGATTTATGATTCATTATTTCGATCTTATTGGCCTTTGTTCTTTATTTTTGATTTTTCATATAGATTTCCGTCTTTATATTATACCCTTTCGTAGATGAATTATGCTTATTTTCACATCTAGGATTTACATATACAACATATATTACTGTCAAGAGGGAATTTTTCTCAGTATTTAGATATTTAGATTCAAAATAAGAAAGGG-----TTCAATTATAAACCTGCAAAACAAAGATTAGGATTGGGTTGGGTTGCGCTATATATATCAAAGAGTATACAATAATGATGTATTTGGTGAATCAAATACATGGTCTAATAACAAACCATTTTAACATAACATTTTAATGAGTTGATAATATTAATTGAATATTGAATATTTTTTGAAAGATTTTTGTTAAAGGTTTCATTCATGCCTAATCCATATCGAGTAGACCTTGTTGTTGTGAG

			>Goudaea_ospinae
			ATAATTATAATAAAGTGAAATATGTCGAAATCCTTTTTGGATTAATACCAAATCGAAAATAAATGTCCGATAGCAAGTTGATCAGTTAATTCAATAAGAGATAAATGGGAGATAGCACTCGATTTAGTTGGTACCACCCAACCGAATCCGATTCAATCGTTTACTCATTCAATGAGTCAATTTTCAAGTTCAGCCAATCTTTTTTTTTTCAAAAAAAAAATTGCAAGTAAATGAAATCGTGAATAAATGTGTTTCATTTCTCTATCATTATAGAAAAAAAGATCCATATTATATTACTTATGGAATTCGAACCCGAACTCGATTTATGATTCATTATTTTGATCTTATTGGCCTTTGTTCTTTATTTTTTATTTTTCATATAGATTTCCGTCTTTATATTATACCCTTTCGTAGATGAATTATGCTTATTTTCACATCTAGGATTTACATATACAACATATATTACTGTCAAGAGGGAATTTTTCTCAGTATTTAGATATTTAGATTCAAAATAAGAAAGGGATTAATTCAATTATAAACCTGCAAAACAAAGATTAGGATTGGGTTGGGTTGCGCTATATATATCAAAGAATATACAATAATGATGTATTTGGTGAATCAAATACATGGTCTAATAACAAACCATTTTAACATAACATTTTAATGAGTTGATAATATTAATTGAATATTGAATATTTTTTGAAAGATTTTTGTTAAAGGTTTCATTCATGCCTAATCCATATCGAGTAGACCTTGTTGTTGTGAG

			>Goudaea_ospinae_var._gruberi
			??????????????????????????????????????????????????????????????????????????????????????????????????????????????????????????????????????????????????????????????????????????????????????????????????????????????????????????????????????????????????????????????????????????????????????????????????????????????????????????????????????????????????????????????????????????????????????????????????????????????????????????????????????????????????????????????????????????????????????????????????????????????????????????????????????????????????????????????????????????????????????????????????????????????????????????????????????????????????????????????????????????????????????????????????????????????????????????????????????????????????????????????????????????????????????????????

			>Gregbrownia_fulgens
			??????????????????????????????????????????????????????????????????????????????????????????????????????????????????????????????????????????????????????????????????????????????????????????????????????????????????????????????????????????????????????????????????????????????????????????????????????????????????????????????????????????????????????????????????????????????????????????????????????????????????????????????????????????????????????????????????????????????????????????????????????????????????????????????????????????????????????????????????????????????????????????????????????????????????????????????????????????????????????????????????????????????????????????????????????????????????????????????????????????????????????????????????????????????????????????????

			>Gregbrownia_hutchisonii
			ATAATTATAATAAAGTGAAATATGTCGAAATCCTTTTTGGAATAATACCAAATCAAAAATAAATGTCCGATAGCAAGTTGATCAGTTAATTCAATAAGAGATAAATGGGAGATAGCACTCGATTTAGTTGGTACCACCCAACCGAATCCGATTCAATCGTTTACTCATTCAATGAGTAAATTTTCAAGTTCAGCCAATCTTTTTTTT---AAAAAAAAAATTGCAAGTAAATGAAATCGTGAATAAATGTGTTTCATTTCTCTATCATTATAGAAAAAAAGATCCATATTATATTACTTATGGAATTCGAACCCGAACTCGATTTATGATTCATTATTTCGATCTTATTGGCCTTTGTTCTTTATTTTTTATTTTTCATATAGATTTTCGTCTTTATATTATACCCTTTCGTAGATGAATTATGCTTATTTTCACATCTAGGATTTACATATACAACATATATTACTGTCAAGAGGGAATTTTTCTCAGTATTTAGATATTTAGATTCAAAATAAGAAAGGGATCAATTCAATTATAAACCTGCAAAACAAAGATTAGGATTGGGTTGGGTTGCGCTATATATATCAAAGAGTATACAATAATGATGTATTTGGTGAATCAAATACATGGTCTAATAACAAACCATTTTAACATAACATTTTAATGAGTTGATAATATTAATTGAATATTGAATATTTTTTGAAAGATTTTTGTTAAAGGTTTCATTCATGCCTAATCCATATCGAGTAGACCTTGTTGTTGTGAG

			>Greigia_mulfordii
			ATAATTATAATAAAGTGAAATATGTCGAAATCCTTTTTGGAATAATACCAAATC-AAAATAAATGTCCGATAGCAAGTTGATCAGTTAATTCAATAAGAG----------------------------------------------------------------------------------------------------------------------------------------------ATAAATGTGTTTCATTTCTCTATCATTATATAAAATAAGATCCATATTATATTACTTATGGAATTCGAACCCGAACTCGATTTATGATTTATTATTTCGATCTTATTGGCCTTTGTTCTTTATTTTGGATTTTTCATATAGATTTCCGTCTTTATATTATACCCTTTCGTGGATGAATTATGCCTATTTTAACATCTAGGATTTACATATACAACATATATTACTGTCAAGAGGGAATTTTTCTCAGTATTTAGATATTTAGATTCAAAATAAGAAAGGGATCAATTCAATTATAAACCTGCAAAACAAAGATTAGGATTGGGTTGGGTTGCGCTATATATATCAAAGAGTATACAATAATGATGTATTTGGTGAATCAAATACATGGTCTAATAACAAACCATTTTAACATAACATTTTAATGAGTTGATAATATTAATTGAATA-------TTTTTTGAAAGATTTTTGTTAAAGGTTTCATTCATGCCTAATCCATATCGAGTAGACCTTGTTGTTGTG??

			>Greigia_sphacelata
			ATAATTATAATAAAGTGAAATATGTCGAAATCCTTTTTGGAATAATACCAAATC-AAAATAAATGTCCGATAGCAAGTTGATCAGTTAATTCAATAAGAG----------------------------------------------------------------------------------------------------------------------------------------------ATAAATGTGTTTCATTTCTCTATCATTATATAAAATAAGATCCATATTATATTACTTATGGAATTCGAACCCGAACTCGATTTATGATTTATTATTTAGATCTTATTGGCCTTTGTTCTTTATTTTTTATTTTTCATATGGATTTCCGTCTTTATATTATACCCTTTCGTGGATGAATTATGCCTATTTTCACATCTAGGATTTACATATACAACATATATTACTGTCAAGAGGGAATTTTTCTCAGTATTTAGATATTTAGATTCAAAATAAGAAAGGGATCAATTCAATTATAAACCTGCAAAACAAAGATTTGGATTGGGTTGGGTTGCGCTATATATATCAAAGAGTATACAATAATGATGTATTTGGTGAATCAAATACATGGTCTAATAACAAACCATTTTAACATAACATTTTAATGAGTTGATAATATTAATTGAATATTGAATATTTTTTGAAAGATTTTTGTTAAAGGTTTCATTCATGCCTAATCCATATCGAGTAGACCTTGTTGTTGTG??

			>Guzmania_acorifolia
			ATAATTATAATAAAGTGAAATATGTCGAAATCCTTTTTGGAATAATACCAAATCGAAAATAAATGTCCGATAGCAAGTTGATCAGTTAATTCAATAAGAGATAAATGGGAAATAGCACTCGATTTAGTTGGTACCACCCAACCGAATCCGATTCAATCGTTTACTCATTCAATGAGTAAATTTTCAAGTTCAGCCAATC-TTTTTTTTTT--------AATTGCAAGTAAATGAAATCGTGAATAAATGTGTTTCATTTCTCTATCATTATAGAAAAAAAGATCCATATTATATTACTTATGGAATTCGAACCCGAACTCAATTTATGATTCATTATTTCGATCTTATTGGCCTTTGTTCTTTATTTTTTATTTTTCATATAGATTTCCATCTTTATATTATACCCTTTCGTAGATTAATTATGCTTATTTTCACATCTAGGATTTACATATACAACATATATTACTGTCAAGAGGGAATTTTTCTCAGTATTTAGATATTTAGATTCAAAATAAGAAAGGGATCAATTCAATTATAAACCTGCAAAACAAAGATTAGAATTGGGTTGGGTTGCGCTATATATATCAAAGAGTATACAATAATGATGTATTTGGTGAATCAAATACATGGTCTAATAACAAACCATTTTAACATAACATTTTAATGAGTTGATAATATTAATTGAATATTGAATATTTTTTGAAAGATTTTTGTTAAAGGTTTCATTCATGCCTAATCCATATCGAGTAGACCTTGTTGTTGTGAG

			>Guzmania_angustifolia
			ATAATTATAATAAAGTGAAATATGTCGAAATCCTTTTTGGAATAATACCAAATCGAAAATAAATGTCCGATAGCAAGTTGATCAGTTAATTCAATAAGAGATAAATGGGAAATAGCACTCAATTTAGTTGGTACCACCCAACCGAACCCGATTCAATCGTTTACTCATTGAATGAGTAAATTTTCAAGTTCAGCCAATCCTTTTTTTTTT------TTAATTGCAAGTAAATGAAATCGTGAATAAATGTGTTTCATTTCTCTATCATTATAGAAAAAAAGATCCATATTATATTACTTATGGAATTCGAACCCGAACTCAATTTATGATTCATTATTTCGATCTTATTGGCCTTTGTTCTTTA-------TTTTTCATATAAATTTCGGTCTTTATATTATACCCTTTCGTAGATGAATTATGCTTATTTTCACATCTAGGATTTACATATACAACATATATTACTGTCAAGAGGGAATTTTTCTCAGTATTTAGATATTTAGATTCAAAATAAGAAAGGGATCAATTCAATTATAAACCTGCAAAACAAAGATTAGGATTGGGTTGGGTTGCGCTATATATATCAAAGAGTATACAATAATGATGTATTTGGTGAATCAAATACATGGTCTAATAACAAACCATTTTAACATAACATTTTAATGAGTTGATAATATTAATTGAATATTGAATATTTTTTGAAAGATTTTTGTTAAAGGTTTCATTCATGCCTAATCCATATCGAGTAGACCTTGTTGTTGTGAG

			>Guzmania_butcheri
			??????????????????????????????????????????????????????????????????????????????????????????????????????????????????????????????????????????????????????????????????????????????????????????????????????????????????????????????????????????????????????????????????????????????????????????????????????????????????????????????????????????????????????????????????????????????????????????????????????????????????????????????????????????????????????????????????????????????????????????????????????????????????????????????????????????????????????????????????????????????????????????????????????????????????????????????????????????????????????????????????????????????????????????????????????????????????????????????????????????????????????????????????????????????????????????????

			>Guzmania_condensata
			ATAATTATAATAAAGTGAAATATGTCGAAATCCTTTTTGGAATAATACCAAATCGAAAATAAATGTCCGATAGCAAGTTGATCAGTTAATTCAATAAGAGATAAATGGGAAATAGCACTCGATTTAGTTGGTACCACCCAACCGAATCCGATTCAATCGTTTACTCATTCAATGAGTAAATTTGCAAGTTCAGCCAATCTTTTTTTTTTT--------AATTGCAAGTAAATGAAATCGTGAATAAATGTGTTTCATTTCTCTATCATTATAGAAAAAAAGATCCATATTATATTACTTATGGAATTCGAACCCGAACTCAATTTATGATTCATTATTTCGATCTTATTGGCCTTTGTTCTTTATTTTTTATTTTTCATATAGATTTCCATCTTTATATTATACCCTTTCGTAGATGAATTATGCTTATTTTCACATCTAGGATTTACATATACAACATATATTACTGTCAAGAGGGAATTTTTCTCAGTATTTAGATATTTAGATTCAAAATAAGAAAGGGATCAATTCAATTATAAACCTGCAAAACAAAGATTAGGATTGGGTTGGGTTGCGCTATATATATCAAAGAGTATACAATAATGATGTATTTGGTGAATCAAATACATGGTCTAATAACAAACCATTTTAACATAACATTTTAATGAGTTGATAATATTAATTGAATATTGAATATTTTTTGAAAGATTTTTGTTAAAGGTTTCATTCATGCCTAATCCATATCGAGTAGACCTTGTTGTTGTGAG

			>Guzmania_coriostachya
			??????????????????????????????????????????????????????????????????????????????????????????????????????????????????????????????????????????????????????????????????????????????????????????????????????????????????????????????????????????????????????????????????????????????????????????????????????????????????????????????????????????????????????????????????????????????????????????????????????????????????????????????????????????????????????????????????????????????????????????????????????????????????????????????????????????????????????????????????????????????????????????????????????????????????????????????????????????????????????????????????????????????????????????????????????????????????????????????????????????????????????????????????????????????????????????????

			>Guzmania_donnellsmithii
			ATAATTATAATAAAGTGAAATATGTCGAAATCCTTTTTGGAATAATACCAAATCGAAAATAAATGTCCGATAGCAAGTTGATCAGTTAATTCAATAAGAGATAAATGGGAAATAGCACTCGATTTAGTTGGTACCACCCAACCGAATCCGATTCAATCGTTTACTCATTCAATGAGTAAATTTTCAAGTTCAGCCAATCCTTTTTTTTTT--------AATTGCAAGTAAATGAAATCGTGAATAAATGTGTTTCATTTCTCTATCATTATAGAGAAAAAGATCCATATTATATTATTTATGGAATTCGAACCCGAACTCAATTTATGATTCATTATTTCGATCTTATTGGCCTTTGTTCTTTATTTTTTATTTTTCATATAGATTTCCATCTTTATATTATACCCTTTCGTAGATGAATTATGCTTATTTTCACATCTAGGATTTACATATACAACATATATTACTGTCAAGAGGGAATTTTTCTCAGTATTTAGATATTTAGATTCAAAATAAGAAAGGGATCAATTCAATTATAAACCTGCAAAACAAAGATTAGGATTGGGTTGGGTTGCGCTATATATATCAAAGAGTATACAATAATGATGTATTTGGTGAATCAAATACATGGTCTAATAACAAACCATTTTAACATAACATTTTAATGAGTTGATAATATTAATTGAATATTGAATATTTTTTGAAAGATTTTTGTTAAAGGTTTCATTCATGCCTAATCCATATCGAGTAGACCTTGTTGTTGTGAG

			>Guzmania_glaucophylla
			??????????????????????????????????????????????????????????????????????????????????????????????????AGATAAATGGGAAATAGCACTCGATTTAGTTGGTACCACCCAACCGAATCCGATTCAATCGTTTACTCATTCAATGAGTAAATTTTCAAGTTCAGCCAATCCTTTTTTTTTT------TTAATTGCAAGTAAATGAAATCGTGAATAAATGTGTTTCATTTCTCTATCATTATAGAAAAAAAGATCCATATTATATTACTTATGGAATTCGAACCCGAACTCAATTTATGATTCATTATTTCGATCTTATTGGCCTTTGTTCTTTATTTTTTATTTTTCATATAGATTTCCATCTTTATATTATACCCTTTCGTAGATGAATTATGCTTATTTTCACATCTAGGATTTACATATACAACATATATTACTGTCAAGAGAGAATTTTTCTCAGTATTTAGATATTTAGATTCAAAATAAGAAAGGGATCAATTCAATTATAAACCTGCAAAACAAAGATTAGGATTGGGTTGGGTTGCGCTATATATATCAAAGAGTATACAATAATGATGTATTTGGTGAATCAAATACATGGTCTAATAACAAACCATTTTAACATAACATTTTAATGAGTTGATAATATTAATTGAATATTGAATATTTTTTGAAAGATTTTTGTTAAAGGTTTCATTCATGCCTAATCCATATCGAGTAGAC-TTGTTGTTGTGAG

			>Guzmania_graminifolia
			ATAATTATAATAAAGTGAAATATGTCGAAATCCTTTTTGGAATAATACCAAATCGAAAATAAATGTCCGATAGCAAGTTGATCAGTTAATTCAATAAGAGATAAATGGGAAATAGCACTCGATTTAGTTGGTACCACCCAACCGAATCCGATTCAATCGTTTACTCATTCAATGAGTAAATTTTCAAGTTCAGCCAATCTTTTTTTTTTT--------AATTGCAAGTAAATGAAATCGTGAATAAATGTGTTTCACTTCTCTATCATTATAGAAAAAAAGATCCATATTATATTACTTATGGAATTCGAACCCGAACTCAATTTATGATTCATTATTTCGATCTTATTGGCCTTTGTTCTTTATTTTTTATTTTTCATATAGATTTCCATCTTTATATTATACCCTTTCGTAGATGAATTATGCTTATTTTCACATCTAGGATTTACATATACAACATATATTACTGTCAAGAGGGAATTTTTCTCAGTATTTAGATATTTAGATTCAAAATAAGAAAGGGATCAATTCAATTATAAACCTGCAAAACAAAGATTAGGATTGGGTTGGGTTGCGCTATATATATCAAAGAGTATACAATAATGATGTATTTGGTGAATCAAATACATGGTCTAATAACAAACCATTTTAACATAACATTTTAATGAGTTGATAATATTAATTGAATATTGAATATTTTTTGAAAGATTTTTGTTAAAGGTTTCATTCATGCCTAATCCATATCGAGTAGACCTTGTTGTTGTGAG

			>Guzmania_herrerae
			ATAATTATAATAAAGTGAAATATGTCGAAATCCTTTTTGGAATAATACCAAATCGAAAATAAATGTCCGATAGCAAGTTGATCAGTTAATTCAATAAGAGATAAATGGGAAATAGCACTCGATTTAGTTGGTACCACCCAACCGAATCCGATTCAATCGTTTACTCATTCAATGAGTAAATTTTCAAGTTCAGCCAATC--TTTTTTTTT--------AATTGCAAGTAAATGAAATCGTGAATAAATGTGTTTCATTTCTCTATCATTATAGAAAAAAAGATCCATATTATATTACTTATGGAATTCGAACCCGAACTCAATTTATGATTCATTATTTCGATCTTATTGGCCTTTGTTCTTTATTTTTTATTTTTCATATAGATTTCCATCTTTATATTATACCCTTTCGTAGATGAATTATGCTTATTTTCACATCTAGGATTTACATATACAACATATATTACTGTCAAGAGGGAATTTTTCTCAGTATTTAGATATTTAGATTCAAAATAAGAAAGGGATCAATTCAATTATAAACCTGCAAAACAAAGATTAGGATTGGGTTGGGTTGCGCTATATATATCAAAGAGTATACAATAATGATGTATTTGGTGAATCAAATACATGGTCTAATAACAAACCATTTTAACATAACATTTTAATGAGTTGATAATATTAATTGAATATTGAATATTTTTTGAAAGATTTTTGTTAAAGGTTTCATTCATGCCTAATCCATATCGAGTAGACCTTGTTGTTGTGAG

			>Guzmania_lingulata
			??????????????????????????????????????????????????????????????????????????????????????????????????????????????????????????????????????????????????????????????????????????????????????????????????????????????????????????????????????????????????????????????????????????????????????????????????????????????????????????????????????????????????????????????????????????????????????????????????????????????????????????????????????????????????????????????????????????????????????????????????????????????????????????????????????????????????????????????????????????????????????????????????????????????????????????????????????????????????????????????????????????????????????????????????????????????????????????????????????????????????????????????????????????????????????????????

			>Guzmania_melinonis
			ATAATTATAATAAAGTGAAATATGTTGAAATCCTTTTTGGAATAATACCAAATCGAAAATAAATGTCCGATAGCAAGTTGATCAGTTAATTCAATAAGAGATAAATGGGAAATAGCACTCAATTTAGTTGGTACCACCCAACCGAATCCGATTCAATCGTTTACTCATTGAATGAGTAAATTTTCAAGTTCAGCCAATCTTTTTTTTTTT--------TATTGCAAGTAAATGAAATCGTGAATAAATGTGTTTCATTTCTCTATCATTATAGAAAAAAAGATCCATATTATATTACTTATGGAATTCGAACCCGAACTCAATTTATGATTCATTATTTCGATCTTATTGGCCTTTGTTCTTTATTTTTTATTTTTCATATAGATTTCCGTCTTTATATTATACCCTTTCGTAGATGAATTATGCTTATTTTCACATCTAGGATTTACATATACAACATATATTACTGTCAAGAGGGAATTTTTCTCAGTATTTAGATATTTAGATTCAAAATAAGAAAGGGATCAATTCAATTATAAACCTGCAAAACAAAGATTAGGATTGGGTTGGGTTGCGCTATATATATCAAAGAGTATACAATAATGATGTATTTGGTGAATCAAATACATGGTCTAATAACAAACCATTTTAACATAACATTTTAATGAGTTGATAATATTAATTGAATATTGAATATTTTTTGAAAGATTTTTGTTAAAGGTTTCATTCATGCCTAATCCATATCGAGTAGACCTTGTTGTTGTGAG

			>Guzmania_monostachia
			ATAATTATAATAAAGTGAAATATGTCGAAATCCTTTTTGGAATAATACCAAATCGAAAATAAATGTCCGATAGCAAGTTGATCAGTTAATTCAATAAGAGATAAATGGGAAATAGCACTCAATTTAGTTGGTACCACCCAACCGAATCCGATTCAATCGTTTACTCATTCAATGAGTAAATTTTCAAGTTCAGCCAATCCTTTTTTTC--AAAAAAAAAATTGCAAGTAAATGAAATCGTGAATAAATGTGTTTCATTTCTCTATCATTATAGAAAAAAAGATCCATATTATATTACTTATGGAATTCGAACCCGAACTCAATTTATGATTCATTATTTCGATCTTATTGGCCTTTGTTCTTCATTTTTTATTTTTCATATAAATTTCCGTCTTTATATTATACCCTTTCGTAGATGAATTATGCTTATTTTCACATCTAGGATTTACATATACAACATATATTACTGTCAAGAGGGAATTTTTCTCAGTATTTAGATATTTAGATTCAAAATAAGAAAGGGATCAATTCAATTATAAACCTGCAAAACAAAGATTAGGATTGGGTTGGGTTGCGCTATATATATCAAAGAGTATACAATAATGATGTATTTGGTGAATCAAATACATGGTCTAATAACAAACCATTTTAACATAACATTTTAATGAGTTGATAATATTAATTGAATATTGAATATTTTTTGAAAGATTTTTGTTAAAGGTTTCATTCATGCCTAATCCATATMGAGTAGACCTTGTTGTTGTG??

			>Guzmania_mucronata
			??????????????????????????????????????????????????????????????????????????????????????????????????????????????????????????????????????????????????????????????????????????????????????????????????????????????????????????????????????????????????????????????????????????????????????????????????????????????????????????????????????????????????????????????????????????????????????????????????????????????????????????????????????????????????????????????????????????????????????????????????????????????????????????????????????????????????????????????????????????????????????????????????????????????????????????????????????????????????????????????????????????????????????????????????????????????????????????????????????????????????????????????????????????????????????????????

			>Guzmania_multiflora
			ATAATTATAATAAAGTGAAATATGTCGAAATCCTTTTTGGAATAATACCAAATCGAAAATAAATGTCCGATAGCAAGTTGATCAGTTAATTCAATAAGAGATAAATGGGAAATAGCACTCGATTTAGTTGGTACCACCCAACCGAATCCGATTCAATCGTTTACTCATTCAATGAGTAAATTTGCAAGTTCAGCCAATCTTTTTTTTTTT--------AATTGCAAGTAAATGAAATCGTGAATAAATGTGTTTCATTTCTCTATCATTATAGAAAAAAAGATCCATATTATATTACTTATGGAATTCGAACCCGAACTCAATTTATGATTCATTATTTCGATCTTATTGGCCTTTGTTCTTTATTTTTTATTTTTCATATAGATTTCCATCTTTATATTATACCCTTTCGTAGATGAATTATGCTTATTTTCACATCTAGGATTTACATATACAACATATATTACTGTCAAGAGGGAATTTTTCTCAGTATTTAGATATTTAGATTCAAAATAAGAAAGGGATCAATTCAATTATAAACCTGCAAAACAAAGATTAGGATTGGGTTGGGTTGCGCTATATATATCAAAGAGTATACAATAATGATGTATTTGGTGAATCAAATACATGGTCTAATAACAAACCATTTTAACATAACATTTTAATGAGTTGATAATATTAATTGAATATTGAATATTTTTTGAAAGATTTTTGTTAAAGGTTTCATTCATGCCTAATCCATATCGAGTAGACCTTGTTGTTGTGAG

			>Guzmania_musaica
			??????????????????????????????????????????????????????????????????????????????????CAGTTAATTCAATAAGAGATAAATGGGAGATAGCACTCGATTTAGTTGGTACCACCCAACCGAATCCGATTCAATCGTTTACTCATTCAATGAGTAAATTTTCAAGTTCAGCCAATCCTTTTTTTTTTAAAAAAAAAATTGCAAGTAAATGAAATCGTGAATAAATGTGTTTCATTTCTCTATCATTATAGAAAAAAGGATCCATATTATATTACTTATGGAATTCGAGCCCGAACTCGATTTATGATTCATTATTTCGATCTTATTGGCCTTTGTTCTTTATTTTTTATTTTTCATATAGATTTCCGTCTTTATATTATACCCTTTCGTAGATGAATTATGCTTATTTTCACATCTAGGATTTACATATACAACATATATTACTGTCAAGAGGGAATTTTTCTCAGTATTTAGATATTTAGATTCAAAATAAGAAAGGGATCAATTCAATTATAAACCTGCAAAACAAAGATTAGGATTGGGTTGGGTTGCGCTATATATATCAAAGAGTATACAATAATGATGTATTTGGTGAATCAAATACATGGTCTAATAACAAACCATTTTAACATAACATTTTAATGAGTTGATAATATTAATTGAATATTGAATATTTTTTGAAAGATTTTTGTTAAAGGTTTCATTCATGCCT-ATCCATATCGAGTAGACCTTGTTGTTGTGAG

			>Guzmania_nicaraguensis
			??????????????????????????????????????????????????????????????????????????????????????????????????????????????????????????????????????????????????????????????????????????????????????????????????????????????????????????????????????????????????????????????????????????????????????????????????????????????????????????????????????????????????????????????????????????????????????????????????????????????????????????????????????????????????????????????????????????????????????????????????????????????????????????????????????????????????????????????????????????????????????????????????????????????????????????????????????????????????????????????????????????????????????????????????????????????????????????????????????????????????????????????????????????????????????????????

			>Guzmania_osyana
			??????????????????????????????????????????????????????????????????????????????????????????????????????????????????????????????????????????????????????????????????????????????????????????????????????????????????????????????????????????????????????????????????????????????????????????????????????????????????????????????????????????????????????????????????????????????????????????????????????????????????????????????????????????????????????????????????????????????????????????????????????????????????????????????????????????????????????????????????????????????????????????????????????????????????????????????????????????????????????????????????????????????????????????????????????????????????????????????????????????????????????????????????????????????????????????????

			>Guzmania_patula
			ATAATTATAATAAAGTGAAATATGTCGAAATCCTTTTTGGAATAATACCAAATCGAAAATAAATGTCCGATAGCAAGTTGATCAGTTAATTCAATAAGAGATAAATGGGAAATAGCACTCGATTTAGTTGGTACCACCCAACCGAATCCGATTCAATCGTTTACTCATTCAATGAGTAAATTTTCAAGTTCAGCCAATCCTTTTTTTTTT--------AATTGCAAGTAAATGAAATCGTGAATAAATGTGTTTCATTTCTCTATCATTATAGAAAAAAAGATCCATATTATATTACTTATGGAATTCGAACCCGAACTCAATTTATGATTCATTATTTCGATCTTATTGGCCTTTGTTCTTTATTTTTTATTTTTCATATAGATTTCCATCTTTATATTATACCCTTTCGTAGATGAATTATGCTTATTTTCACATCTAGGATTTACATATACAACATATATTACTGTCAAGAGGGAATTTTTCTCAGTATTTAGATATTTAGATTCAAAATAAGAAAGGGATCAATTCAATTATAAACCTGCAAAACAAAGATTAGGATTGGGTTGGGTTGCGCTATATATATCAAAGAGTATACAATAATGATGTATTTGGTGAATCAAATACATGGTCTAATAACAAACCATTTTAACATAACATTTTAATGAGTTGATAATATTAATTGAATATTGAATATTTTTTGAAAGATTTTTGTTAAAGGTTTCATTCATGCCTAATCCATATCGAGTAGACCTTGTTGTTGTGAG

			>Guzmania_plicatifolia
			??????????????????????????????????????????????????????????????????????????????????????????????????????????????????????????????????????????????????????????????????????????????????????????????????????????????????????????????????????????????????????????????????????????????????????????????????????????????????????????????????????????????????????????????????????????????????????????????????????????????????????????????????????????????????????????????????????????????????????????????????????????????????????????????????????????????????????????????????????????????????????????????????????????????????????????????????????????????????????????????????????????????????????????????????????????????????????????????????????????????????????????????????????????????????????????????

			>Guzmania_polycephala
			??????????????????????????????????????????????????????????????????????????????????????????????????????????????????????????????????????????????????????????????????????????????????????????????????????????????????????????????????????????????????????????????????????????????????????????????????????????????????????????????????????????????????????????????????????????????????????????????????????????????????????????????????????????????????????????????????????????????????????????????????????????????????????????????????????????????????????????????????????????????????????????????????????????????????????????????????????????????????????????????????????????????????????????????????????????????????????????????????????????????????????????????????????????????????????????????

			>Guzmania_rhonhofiana
			ATAATTATAATAAAGTGAAATATGTCGAAATCCTTTTTGGAATAATACCAAATCGAAAATAAATGTCCGATAGCAAGTTGATCAGTTAATTCAATAAGAGATAAATGGGAAATAGCACTCGATTTAGTTGGTACCACCCAACCGAATCCGATTCAATCGTTTACTCATTCAATGAGTAAATTTTCAAGTTCAGCCAATCCTTTTTTTTTT--------AATTGCAAGTAAATGAAATCGTGAATAAATGTGTTTCATTTCTCTATCATTATAGAAAAAAAGATCCATATTATATTACTTATGGAATTCGAACCCGAACTCAATTTATGATTCATTATTTCGATCTTATTGGCCTTTGTTCTTTATTTTTTATTTTTCATATAGATTTCCATCTTTATATTATACCCTTTCGTAGATGAATTATGCTTATTTTCACATCTAGGATTTACATATACAACATATATTACTGTCAAGAGGGAATTTTTCTCAGTATTTAGATATTTAGATTCAAAATAAGAAAGGGATCAATTCAATTATAAACCTGCAAAACAAAGATTAGGGTTGGGTTGGGTTGCGCTATATATATCAAAGAGTATACAATAATGATGTATTTGGTGAATCAAATACATGGTCTAATAACAAACCATTTTAACATAACATTTTAATGAGTTGATAATATTAATTGAATATTGAATATTTTTTGAAAGATTTTTGTTAAAGGTTTCATTCATGCCTAATCCATATCGAGTAGACCTTGTTGTTGTGAG

			>Guzmania_roezlii
			ATAATTATAATAAAGTGAAATATGTCGAAATCCTTTTTGGAATAATACCAAATCGAAAATAAATGTCCGATAGCAAGTTGATCAGTTAATTCAATAAGAGATAAATGGGAAATAGCACTCGATTTAGTTGGTACCACCCAACCGAATCCGATTCAATCGTTTACTCATTCAATGAGTAAATTTTCAAGTTCAGCCAATCCTTTATTTTTT------TTAATTGCAAGTAAATGAAATCGTGAATAAATGTGTTTCATTTCTCTATCATTATAGAAAAAAAGATCCATATTATATTACTTATGGAATTCGAACCCGAACTCAATTTATGATTCATTATTTCGATCTTATTGGCCTTTGCTCTTTATTTTTTATTTTTCATATAGATTTCCATCTTTATATTATACCCTTTCGTAGATGAATTATGCTTATTTTCACATCTAGGATTTACATATACAACATATATTACTGTCAAGAGAGAATTTTTCTCAGTATTTAGATATTTAGATTCAAAATAAGAAAGGGATCAATTCAATTATAAACCTGCAAAACAAAGATTAGGATTGGGTTGGGTTGCGCTATATATATCAAAGAGTATACAATAATGATGTATTTGGTGAATCAAATACATGGTCTAATAACAAACCATTTTAACATAACATTTTAATGAGTTGATAATATTAATTGAATATTGAATATTTTTTGAAAGATTTTTGTTAAAGGTTTCATTCATGCCTAATCCATATCGAGTAGACCTTGTTGTTGTGAG

			>Guzmania_sanguinea
			??????????????????????????????????????????????????????????????????????????????????????????????????????????????????????????????????????????????????????????????????????????????????????????????????????????????????????????????????????????????????????????????????????????????????????????????????????????????????????????????????????????????????????????????????????????????????????????????????????????????????????????????????????????????????????????????????????????????????????????????????????????????????????????????????????????????????????????????????????????????????????????????????????????????????????????????????????????????????????????????????????????????????????????????????????????????????????????????????????????????????????????????????????????????????????????????

			>Guzmania_scherzeriana
			??????????????????????????????????????????????????????????????????????????????????????????????????????????????????????????????????????????????????????????????????????????????????????????????????????????????????????????????????????????????????????????????????????????????????????????????????????????????????????????????????????????????????????????????????????????????????????????????????????????????????????????????????????????????????????????????????????????????????????????????????????????????????????????????????????????????????????????????????????????????????????????????????????????????????????????????????????????????????????????????????????????????????????????????????????????????????????????????????????????????????????????????????????????????????????????????

			>Guzmania_spectabilis
			??????????????????????????????????????????????????????????????????????????????????????????????????????????????????????????????????????????????????????????????????????????????????????????????????????????????????????????????????????????????????????????????????????????????????????????????????????????????????????????????????????????????????????????????????????????????????????????????????????????????????????????????????????????????????????????????????????????????????????????????????????????????????????????????????????????????????????????????????????????????????????????????????????????????????????????????????????????????????????????????????????????????????????????????????????????????????????????????????????????????????????????????????????????????????????????????

			>Guzmania_subcorymbosa
			??????????????????????????????????????????????????????????????????????????????????????????????????????????????????????????????????????????????????????????????????????????????????????????????????????????????????????????????????????????????????????????????????????????????????????????????????????????????????????????????????????????????????????????????????????????????????????????????????????????????????????????????????????????????????????????????????????????????????????????????????????????????????????????????????????????????????????????????????????????????????????????????????????????????????????????????????????????????????????????????????????????????????????????????????????????????????????????????????????????????????????????????????????????????????????????????

			>Guzmania_variegata
			ATAATTATAATAAAGTGAAATATGTCGAAATCCTTTTTGGAATAATACCAAATTGAAAATAAATGTCCGATAGCAAGTTGATCAGTTAATTCAATAAGAGATAAATGGGAAATAGCACTCGATTTAGTTGGTACCACCCAACCGAATCCGATTCAATCGTTTACTCATTCAATGAGTCAATTTTCAAGTTCAGCCAATC--TTTTTTTTGAAAAAAAAAATTGCAAGTAAATGAAATCGTGAATAAATGTGTTTCATTTCTCTATCATTATAGAAAAAAAGATCCATATTATATTACTTATGGAATTCGAACCCGAACTCGATTTATGATTCATTATTTCGATCTTATTGGCCTTTGTTCTTTATTTTTTATTTTGCATATAGATTTCCGTCTTTATATTATACCCTTTCGTAGATGAATTATGCTTATTTTCACATCTAGGATTTACATATACAACATATATTACTGTCAAGAGGGAATTTTTCTCAGTATTTAGATATTTAGATTCAAAATAAGAAAGGGATCAATTCAATTATAAACCTGCAAAACAAAGATTAGGATTGGGTTGGGTTGCGCTATATATATCAAAGAGTATACAATAATGATGTATTTGGTGAATCAAATACATGGTCTAATAACAAACCATTTTAACATAACATTTTAATGAGTTGATAATATTAATTGAATATTGAATATTTTTTGAAAGATTTTTGTTAAAGGTTTCATTCATGCCTAATCCATATCGAGTAGACCTTGTTGTTGTGAG

			>Guzmania_wittmackii
			ATAATTATAATAAAGTGAAATATGTCGAAATCCTTTTTGGAATAATACCAAATTGAAAATAAATGTCCGATAGCAAGTTGATCAGTTAATTCAATAAGAGATAAATGGGAAATAGCACTCGATTTAGTTGGTACCACCCAACCGAATCCGATTCAATCGTTTACTCATTCAATGAGTAAATTTTCAAGTTCAGCCAATCTTTTTTTTTTT------TTAATTGCAAGTAAATGAAATCGTGAATAAATGTGTTTCATTTCTCTATCATTATAGAAAAAAAGATCCATATTATATTACTTATGGAATTCGAACCCGAACTCGATTTATGATTCATTATTTTGATCTTATTGGCCTTTGTTCTTTATTTTTTATTTTTCATATAGATTTCCGTCTTTATATTATACCCTTTCGTAGATGAATTATGCTTATTTTCACATCTAGGATTTACATATACAACATATATTACTGTCAAGAGGGAATTTTTCTCAGTATTTAGATATTTAGATTCAAAATAAGAAAGGGATCAATTCAATTATAAACCTGCAAAACAAAGATTAGGATTGGGTTGGGTTGCGCTATATATATCAAAGAGTATACAATAATGATGTATTTGGTGAATCAAATACATGGTCTAATAACAAACCATTTTAACATAACATTTTAATGAGTTGATAATATTAATTGAATATTGAATATTTTTTGAAAGATTTTTGTTAAAGGTTTCATTCATGCCTAATCCATATCGAGTAGACCTTGTTGTTGTGAG

			>Guzmania_zahnii
			??????????????????????????????????????????????????????????????????????????????????????????????????????????????????????????????????????????????????????????????????????????????????????????????????????????????????????????????????????????????????????????????????????????????????????????????????????????????????????????????????????????????????????????????????????????????????????????????????????????????????????????????????????????????????????????????????????????????????????????????????????????????????????????????????????????????????????????????????????????????????????????????????????????????????????????????????????????????????????????????????????????????????????????????????????????????????????????????????????????????????????????????????????????????????????????????

			>Hechtia_argentea
			??????????????????????????????????????????????????????????????????????????????????CAGTTAATTCAATAAGAGATAAATGGGAGATAGCACTCGATTTAGTTGGTACCAACCAACCGAATCCAATTCAATCGTTTACTCA--------------TTGCAAGTTCAGCCAATC-TTTTTTTTTTCAAAAAAAAATTGCAAGTAGATGAAATCGTAAATAAATGTGTTTCATTTCTCTATCATTATAGAAAATAAGATCCATATTATATTACTTATGGAATTCGAACCCGAACTCGATTTATGATTCATTATTTCGATCTTATTGGCCTTTGTTCTTTATTTTTGATTTTTCATATGGATTTCCGTCTTTATATTATACCCTTTCGTGGATGAATGATGCCTATTTTCACATCTAGGATTTACATATACAACATATATTACTGTCAAGAGGGAATTTTTCTCAGTATTTAGATATTTAGATTCAAAATAAGAAAGGGATCAATTCAATTATAAACCTGCAAAACAAAGATTAGGATTGGGTTGGGTTGCGCTATATATATCAAAGAGTATACAATAATGATGTATTTGGTGAATTAAATACATGGTCTAATAACAAACCATTTTAACATAACATTTTAATGAGTTGATAATATTAATTGAATATTGAATATTTTTTGAAAGATTTTTGTTAAAGGTTTCATTCATGCCTAATTCATATCGAGTAGACCTTGTTGTTGTGAG

			>Hechtia_caerulea
			??????????????????????????????????????????????????????????????????????????????????????????????????????????????????????????????????????????????????????????????????????????????????????????????????????????????????????????????????????????????????????????????????????????????????????????????????????????????????????????????????????????????????????????????????????????????????????????????????????????????????????????????????????????????????????????????????????????????????????????????????????????????????????????????????????????????????????????????????????????????????????????????????????????????????????????????????????????????????????????????????????????????????????????????????????????????????????????????????????????????????????????????????????????????????????????????

			>Hechtia_carlsoniae
			ATAATTATAATAAAGTGAAATATGTCGAAATCCTTTTTGGAATAATACCAAATCGAAAATAAATGTCCGATAGCAAGTTGATCAGTTAATTCAATAAGAGATAAATGGGAGATAGCACTCGATTTAGTTGGTACCAACCAACCGAATCCAATTCAATCGTTTACTCATTGAATGAGTAAATTTTCAAGTTCAGCCAATCTTTTTTTTTTTT-------AATTGCAAGTAGATGAAATCGTAAATAAATGTGTTTCATTTCTCTATCATTATAGAAAAAAAGATCCATATTATATTACTTATGGAATTCGAACCCGAACTCGATTTATGATTCATTATTTCGATCTTATTGGCCTTTGTTCTTTATTTTTGATTTTTCATATGGATTTCCGTCTTTATATTATACCCTTTCGTGGATGAATGATGCCTATTTTCACATCTAGGATTTACATATACAACATATATTACTGTCAAGAGGGAATTTTTCTCAGTATTTAGATATTTAGATTCAAAATAAGAAAGGGATCAATTCAATTATAAACCTGCAAAACAAAGATTAGGATTGGGTTGGGTTGCGCTATATATATCAAAGAGTATACAATAATGATGTATTTGGTGAATTAAATACATGGTCTAATAACAAACCATTTTAACATAACATTTTAATGAGTTGATAATATTAATTGAATATTGAATATTTTTTGAAAGATTTTTGTTAAAGGTTTCATTCATGCCTAATTCATATCGAGTAGACCTTGTTGTTGTGAG

			>Hechtia_deceptrix
			??????????????????????????????????????????????????????????????????????????????????????????????????????????????????????????????????????????????????????????????????????????????????????????????????????????????????????????????????????????????????????????????????????????????????????????????????????????????????????????????????????????????????????????????????????????????????????????????????????????????????????????????????????????????????????????????????????????????????????????????????????????????????????????????????????????????????????????????????????????????????????????????????????????????????????????????????????????????????????????????????????????????????????????????????????????????????????????????????????????????????????????????????????????????????????????????

			>Hechtia_dichroantha
			??????????????????????????????????????????????????????????????????????????????????????????????????????????????????????????????????????????????????????????????????????????????????????????????????????????????????????????????????????????????????????????????????????????????????????????????????????????????????????????????????????????????????????????????????????????????????????????????????????????????????????????????????????????????????????????????????????????????????????????????????????????????????????????????????????????????????????????????????????????????????????????????????????????????????????????????????????????????????????????????????????????????????????????????????????????????????????????????????????????????????????????????????????????????????????????????

			>Hechtia_epigyna
			??????????????????????????????????????????????????????????????????????????????????????????????????????????????????????????????????????????????????????????????????????????????????????????????????????????????????????????????????????????????????????????????????????????????????????????????????????????????????????????????????????????????????????????????????????????????????????????????????????????????????????????????????????????????????????????????????????????????????????????????????????????????????????????????????????????????????????????????????????????????????????????????????????????????????????????????????????????????????????????????????????????????????????????????????????????????????????????????????????????????????????????????????????????????????????????????

			>Hechtia_fosteriana
			??????????????????????????????????????????????????????????????????????????????????????????????????????????????????????????????????????????????????????????????????????????????????????????????????????????????????????????????????????????????????????????????????????????????????????????????????????????????????????????????????????????????????????????????????????????????????????????????????????????????????????????????????????????????????????????????????????????????????????????????????????????????????????????????????????????????????????????????????????????????????????????????????????????????????????????????????????????????????????????????????????????????????????????????????????????????????????????????????????????????????????????????????????????????????????????????

			>Hechtia_ghiesbreghtii
			??????????????????????????????????????????????????????????????????????????????????????????????????????????????????????????????????????????????????????????????????????????????????????????????????????????????????????????????????????????????????????????????????????????????????????????????????????????????????????????????????????????????????????????????????????????????????????????????????????????????????????????????????????????????????????????????????????????????????????????????????????????????????????????????????????????????????????????????????????????????????????????????????????????????????????????????????????????????????????????????????????????????????????????????????????????????????????????????????????????????????????????????????????????????????????????????

			>Hechtia_glomerata
			ATAATTATAATAAAGTGAAATATGTCGAAATCCTTTTTGGAATAATACCAAATCGAAAATAAATGTCCGATAGCAAGTTGATCAGTTAATTCAATAAGAGATAAATGGGAGATAGCACTCGATTTAGTTGGTACCAACCAACCGAATCCAATTCAATCGTTTACTCATTGAATGAGTAAATTTTCAAGTTCAGCCAATC-TTTTTTTTTTCAAAAAAAAATTGCAAGTAGATGAAATCGTAAATAAATGTGTTTCATTTCTCTATCATTATAGAAAATAAGATCCATATTATATTACTTATGGAATTCGAACCCGAACTCGATTTATGATTCATTATTTCGATCTTATTGGCCTTTGTTCTTTATTTTTGATTTTTCATATGGATTTCCGTCTTTATATTATACCCTTTCGTGGATGAATGATGCCTATTTTCACATCTAGGATTTACATATACAACATATATTACTGTCAAGAGGGAATTTTTCTCAGTATTTAGATATTTAGATTCAAAATAAGAAAGGGATCAATTCAATTATAAACCTGCAAAACAAAGATTAGGATTGGGTTGGGTTGCGCTATATATATCAAAGAGTATACAATAATGATGTATTTGGTGAATTAAATACATGGTCTAATAACAAACCATTTTAACATAACATTTTAATGAGTTGATAATATTAATTGAATATTGAATATTTTTTGAAAGATTTTTGTTAAAGGTTTCATTCATGCCTAATTCATATCGAGTAGACCTTGTTGTTGTGAG

			>Hechtia_guatemalensis
			ATAATTATAATAAAGTGAAATATGTCGAAATCCTTTTTGGAATAATACCAAATCGAAAATAAATGTCCGATAGCAAGTTGATCAGTTAATTCAATAAGAGATAAATGGGAGATAGCACTCGATTTAGTTGGTACCAACCAACCGAATCCGATTCAATCGTTTACTCATTCAATGAGTCAATTTTCAAGTTCAGCCAATCTTTTTTTTTTTCAAAAAAAAATTGCAAGTAGATGAAATCGTAAATAAATGTGTTTCGTTTCTCTATCATTATAGAAAATAAGATCCATATTATATTACTTATGGAATTCGAACCCGAACTCGATTTATGATTCATTATTTCGATCTTATTGGCCTTTGTTCTTTATTTTTGATTTTTCATATAGATTTCCGTCTTTATATTATACCCTTTCGTGGATGAAAGATGCCTATTTTCACATCTAGGATTTACATATACAACATATATTACTGTCAAGAGGGAATTTTTCTCAGTATTTTGATATTTAGATTCAAAATAAGAAAAGGATCAATTCAATTATAAACCTGCAAAACAAAGATTAGGATTGGGTTGGGTTGCGCTATATATATCAAAGAGTATACAATAATGATGTATTTGGTGAATCAAATACATGGTCTAATAACAAACCATTTTAACATAACATTTTAATGAGTTGATAATATTAATTGAATATTGAATATTTTTTGAAAGATTTTTGTTAAAGGTTTCATTCATGCCTAATCCATATCGAGTAGACCTTGTTGTTGTGAG

			>Hechtia_hernandez-sandovalii
			??????????????????????????????????????????????????????????????????????????????????????????????????????????????????????????????????????????????????????????????????????????????????????????????????????????????????????????????????????????????????????????????????????????????????????????????????????????????????????????????????????????????????????????????????????????????????????????????????????????????????????????????????????????????????????????????????????????????????????????????????????????????????????????????????????????????????????????????????????????????????????????????????????????????????????????????????????????????????????????????????????????????????????????????????????????????????????????????????????????????????????????????????????????????????????????????

			>Hechtia_isthmusiana
			??????????????????????????????????????????????????????????????????????????????????????????????????????????????????????????????????????????????????????????????????????????????????????????????????????????????????????????????????????????????????????????????????????????????????????????????????????????????????????????????????????????????????????????????????????????????????????????????????????????????????????????????????????????????????????????????????????????????????????????????????????????????????????????????????????????????????????????????????????????????????????????????????????????????????????????????????????????????????????????????????????????????????????????????????????????????????????????????????????????????????????????????????????????????????????????????

			>Hechtia_lepidophylla
			??????????????????????????????????????????????????????????????????????????????????????????????????????????????????????????????????????????????????????????????????????????????????????????????????????????????????????????????????????????????????????????????????????????????????????????????????????????????????????????????????????????????????????????????????????????????????????????????????????????????????????????????????????????????????????????????????????????????????????????????????????????????????????????????????????????????????????????????????????????????????????????????????????????????????????????????????????????????????????????????????????????????????????????????????????????????????????????????????????????????????????????????????????????????????????????????

			>Hechtia_lundelliorum
			??????????????????????????????????????????????????????????????????????????????????????????????????????????????????????????????????????????????????????????????????????????????????????????????????????????????????????????????????????????????????????????????????????????????????????????????????????????????????????????????????????????????????????????????????????????????????????????????????????????????????????????????????????????????????????????????????????????????????????????????????????????????????????????????????????????????????????????????????????????????????????????????????????????????????????????????????????????????????????????????????????????????????????????????????????????????????????????????????????????????????????????????????????????????????????????????

			>Hechtia_malvernii
			??????????????????????????????????????????????????????????????????????????????????????????????????????????????????????????????????????????????????????????????????????????????????????????????????????????????????????????????????????????????????????????????????????????????????????????????????????????????????????????????????????????????????????????????????????????????????????????????????????????????????????????????????????????????????????????????????????????????????????????????????????????????????????????????????????????????????????????????????????????????????????????????????????????????????????????????????????????????????????????????????????????????????????????????????????????????????????????????????????????????????????????????????????????????????????????????

			>Hechtia_matudae
			??????????????????????????????????????????????????????????????????????????????????????????????????????????????????????????????????????????????????????????????????????????????????????????????????????????????????????????????????????????????????????????????????????????????????????????????????????????????????????????????????????????????????????????????????????????????????????????????????????????????????????????????????????????????????????????????????????????????????????????????????????????????????????????????????????????????????????????????????????????????????????????????????????????????????????????????????????????????????????????????????????????????????????????????????????????????????????????????????????????????????????????????????????????????????????????????

			>Hechtia_montana
			??????????????????????????????????????????????????????????????????????????????????????????????????????????????????????????????????????????????????????????????????????????????????????????????????????????????????????????????????????????????????????????????????????????????????????????????????????????????????????????????????????????????????????????????????????????????????????????????????????????????????????????????????????????????????????????????????????????????????????????????????????????????????????????????????????????????????????????????????????????????????????????????????????????????????????????????????????????????????????????????????????????????????????????????????????????????????????????????????????????????????????????????????????????????????????????????

			>Hechtia_myriantha
			??????????????????????????????????????????????????????????????????????????????????????????????????????????????????????????????????????????????????????????????????????????????????????????????????????????????????????????????????????????????????????????????????????????????????????????????????????????????????????????????????????????????????????????????????????????????????????????????????????????????????????????????????????????????????????????????????????????????????????????????????????????????????????????????????????????????????????????????????????????????????????????????????????????????????????????????????????????????????????????????????????????????????????????????????????????????????????????????????????????????????????????????????????????????????????????????

			>Hechtia_pretiosa
			??????????????????????????????????????????????????????????????????????????????????????????????????????????????????????????????????????????????????????????????????????????????????????????????????????????????????????????????????????????????????????????????????????????????????????????????????????????????????????????????????????????????????????????????????????????????????????????????????????????????????????????????????????????????????????????????????????????????????????????????????????????????????????????????????????????????????????????????????????????????????????????????????????????????????????????????????????????????????????????????????????????????????????????????????????????????????????????????????????????????????????????????????????????????????????????????

			>Hechtia_purpusii
			ATAATTATAATAAAGTGAAATATGTCGAAATCCTTTTTGGAATAATACCAAATCGAAAATAAATGTCCGATAGCAAGTTGATCAGTTAATTCAATAAGAGATAAATGGGAGATAGCACTCGATTTAGTTGGTACCAACCAACCGAATCCAATTCAATCGTTTACTCATTCAATGAGTAAATTTTCAAGTTCAGCCAATCTTTTTTTTTTTCAAAAAAAAATTGCAAGTAGATGAAATCGTAAATAAATGTGTTTCATTTCTCTATCATTATAGAAAATAAGATCCATATTATATTACTTATGGAATTCGAACCCGAACTCGATTTATGATTCATTATTTCGATCTTATTGGCCTTTGTTCTTTATTTTGGATTTTTCATGTGGATTTCCGTCTTTATATTATACCCTTTCGTGGATGAATGATGCCTATTTTCACATCTAGGATTTACATATACAACATATATTACTGTCAAGAGGGAATTTTTCTCAGTATTTAGATATTTAGATTCAAAATAAGAAAGGGATCAATTCAATTATAAACCTGCAAAACAAAGATTAGGATTGGATTGGGTTGCGCTATATATATCAAAGAGTATACAATAATGATGTATTTGGTGAATTAAATACATGGTCTAATAACAAACCATTTTAACATAACATTTTAATGAGTTGATAATATTAATTGAATATTGAATATTTTTTGAAAGATTTTTGTTAAAGGTTTCATTCATGCCTAATTCATATCGAGTAGACCTTGTTGTTGTGAG

			>Hechtia_schottii
			??????????????????????????????????????????????????????????????????????????????????????????????????????????????????????????????????????????????????????????????????????????????????????????????????????????????????????????????????????????????????????????????????????????????????????????????????????????????????????????????????????????????????????????????????????????????????????????????????????????????????????????????????????????????????????????????????????????????????????????????????????????????????????????????????????????????????????????????????????????????????????????????????????????????????????????????????????????????????????????????????????????????????????????????????????????????????????????????????????????????????????????????????????????????????????????????

			>Hechtia_stenopetala
			??????????????????????????????????????????????????????????????????????????????????CAGTTAATTCAATAAGAGATAAATGGGAGATAGCACTCGATTTAGTTGGTACCAACCAACCGAATCCAATTCAATCGTTTACTCATTCAATGAGTAAATTTTCAAGTTCAGCCAATC-TTTTTTTTTTCAAAAAAAAATTGCAAGTAGATGAAATCGTAAATAAATGTGTTTCATTTCTCTACCATTATAGAAAATAAGATCCATATTATATTACTTATGGAATTCGAACCCGAACTCGATTTATGATTCATTATTTCGATCTTATTGGCCTTTGTTCTTTATTTTGGATTTTTCATATGGATTTCCGTCTTTATATTATACCCTTTCGTGGATGAATGATGCCTATTTTCACATCTAGGATTTACATATACAACATATATTACTGTCAAGAGGGAATTTTTCTCAGTATTTAGATATTTAGATTCAAAATAAGAAAGGGATCAATTCAATTATAAACCTGCAAAACAAAGATTAGGATTGGATTGGGTTGCGCTATATATATCAAAGAGTATACAATAATGATGTATTTGGTGAATTAAATACATGGTCTAATAACAAACCATTTTAACATAACATTTTAATGAGTTGATAATATTAATTGAATATTGAATATTTTTTGAAAGATTTTTGTTAAAGGTTTCATTCATGCCTAATTCATATCGAGTAGACCTTGTTGTTGTGAG

			>Hechtia_subalata
			??????????????????????????????????????????????????????????????????????????????????????????????????????????????????????????????????????????????????????????????????????????????????????????????????????????????????????????????????????????????????????????????????????????????????????????????????????????????????????????????????????????????????????????????????????????????????????????????????????????????????????????????????????????????????????????????????????????????????????????????????????????????????????????????????????????????????????????????????????????????????????????????????????????????????????????????????????????????????????????????????????????????????????????????????????????????????????????????????????????????????????????????????????????????????????????????

			>Hechtia_texensis
			??????????????????????????????????????????????????????????????????????????????????????????????????????????????????????????????????????????????????????????????????????????????????????????????????????????????????????????????????????????????????????????????????????????????????????????????????????????????????????????????????????????????????????????????????????????????????????????????????????????????????????????????????????????????????????????????????????????????????????????????????????????????????????????????????????????????????????????????????????????????????????????????????????????????????????????????????????????????????????????????????????????????????????????????????????????????????????????????????????????????????????????????????????????????????????????????

			>Hechtia_tillandsioides
			??????????????????????????????????????????????????????????????????????????????????????????????????????????????????????????????????????????????????????????????????????????????????????????????????????????????????????????????????????????????????????????????????????????????????????????????????????????????????????????????????????????????????????????????????????????????????????????????????????????????????????????????????????????????????????????????????????????????????????????????????????????????????????????????????????????????????????????????????????????????????????????????????????????????????????????????????????????????????????????????????????????????????????????????????????????????????????????????????????????????????????????????????????????????????????????????

			>Hohenbergia_andina
			??????????????????????????????????????????????????????????????????????????????????????????????????????????????????????????????????????????????????????????????????????????????????????????????????????????????????????????????????????????????????????????????????????????????????????????????????????????????????????????????????????????????????????????????????????????????????????????????????????????????????????????????????????????????????????????????????????????????????????????????????????????????????????????????????????????????????????????????????????????????????????????????????????????????????????????????????????????????????????????????????????????????????????????????????????????????????????????????????????????????????????????????????????????????????????????????

			>Hohenbergia_belemii
			??????????????????????????????????????????????????????????????????????????????????????????????????????????????????????????????????????????????????????????????????????????????????????????????????????????????????????????????????????????????????????????????????????????????????????????????????????????????????????????????????????????????????????????????????????????????????????????????????????????????????????????????????????????????????????????????????????????????????????????????????????????????????????????????????????????????????????????????????????????????????????????????????????????????????????????????????????????????????????????????????????????????????????????????????????????????????????????????????????????????????????????????????????????????????????????????

			>Hohenbergia_brachycephala
			??????????????????????????????????????????????????????????????????????????????????????????????????????????????????????????????????????????????????????????????????????????????????????????????????????????????????????????????????????????????????????????????????????????????????????????????????????????????????????????????????????????????????????????????????????????????????????????????????????????????????????????????????????????????????????????????????????????????????????????????????????????????????????????????????????????????????????????????????????????????????????????????????????????????????????????????????????????????????????????????????????????????????????????????????????????????????????????????????????????????????????????????????????????????????????????????

			>Hohenbergia_capitata
			??????????????????????????????????????????????????????????????????????????????????????????????????????????????????????????????????????????????????????????????????????????????????????????????????????????????????????????????????????????????????????????????????????????????????????????????????????????????????????????????????????????????????????????????????????????????????????????????????????????????????????????????????????????????????????????????????????????????????????????????????????????????????????????????????????????????????????????????????????????????????????????????????????????????????????????????????????????????????????????????????????????????????????????????????????????????????????????????????????????????????????????????????????????????????????????????

			>Hohenbergia_catingae
			??????????????????????????????????????????????????????????????????????????????????????????????????????????????????????????????????????????????????????????????????????????????????????????????????????????????????????????????????????????????????????????????????????????????????????????????????????????????????????????????????????????????????????????????????????????????????????????????????????????????????????????????????????????????????????????????????????????????????????????????????????????????????????????????????????????????????????????????????????????????????????????????????????????????????????????????????????????????????????????????????????????????????????????????????????????????????????????????????????????????????????????????????????????????????????????????

			>Hohenbergia_correia-araujoi
			??????????????????????????????????????????????????????????????????????????????????????????????????????????????????????????????????????????????????????????????????????????????????????????????????????????????????????????????????????????????????????????????????????????????????????????????????????????????????????????????????????????????????????????????????????????????????????????????????????????????????????????????????????????????????????????????????????????????????????????????????????????????????????????????????????????????????????????????????????????????????????????????????????????????????????????????????????????????????????????????????????????????????????????????????????????????????????????????????????????????????????????????????????????????????????????????

			>Hohenbergia_edmundoi
			??????????????????????????????????????????????????????????????????????????????????????????????????????????????????????????????????????????????????????????????????????????????????????????????????????????????????????????????????????????????????????????????????????????????????????????????????????????????????????????????????????????????????????????????????????????????????????????????????????????????????????????????????????????????????????????????????????????????????????????????????????????????????????????????????????????????????????????????????????????????????????????????????????????????????????????????????????????????????????????????????????????????????????????????????????????????????????????????????????????????????????????????????????????????????????????????

			>Hohenbergia_humilis
			??????????????????????????????????????????????????????????????????????????????????????????????????????????????????????????????????????????????????????????????????????????????????????????????????????????????????????????????????????????????????????????????????????????????????????????????????????????????????????????????????????????????????????????????????????????????????????????????????????????????????????????????????????????????????????????????????????????????????????????????????????????????????????????????????????????????????????????????????????????????????????????????????????????????????????????????????????????????????????????????????????????????????????????????????????????????????????????????????????????????????????????????????????????????????????????????

			>Hohenbergia_loredanoana
			??????????????????????????????????????????????????????????????????????????????????????????????????????????????????????????????????????????????????????????????????????????????????????????????????????????????????????????????????????????????????????????????????????????????????????????????????????????????????????????????????????????????????????????????????????????????????????????????????????????????????????????????????????????????????????????????????????????????????????????????????????????????????????????????????????????????????????????????????????????????????????????????????????????????????????????????????????????????????????????????????????????????????????????????????????????????????????????????????????????????????????????????????????????????????????????????

			>Hohenbergia_pennae
			??????????????????????????????????????????????????????????????????????????????????????????????????????????????????????????????????????????????????????????????????????????????????????????????????????????????????????????????????????????????????????????????????????????????????????????????????????????????????????????????????????????????????????????????????????????????????????????????????????????????????????????????????????????????????????????????????????????????????????????????????????????????????????????????????????????????????????????????????????????????????????????????????????????????????????????????????????????????????????????????????????????????????????????????????????????????????????????????????????????????????????????????????????????????????????????????

			>Hohenbergia_ramageana
			??????????????????????????????????????????????????????????????????????????????????????????????????????????????????????????????????????????????????????????????????????????????????????????????????????????????????????????????????????????????????????????????????????????????????????????????????????????????????????????????????????????????????????????????????????????????????????????????????????????????????????????????????????????????????????????????????????????????????????????????????????????????????????????????????????????????????????????????????????????????????????????????????????????????????????????????????????????????????????????????????????????????????????????????????????????????????????????????????????????????????????????????????????????????????????????????

			>Hohenbergia_ridleyi
			??????????????????????????????????????????????????????????????????????????????????????????????????????????????????????????????????????????????????????????????????????????????????????????????????????????????????????????????????????????????????????????????????????????????????????????????????????????????????????????????????????????????????????????????????????????????????????????????????????????????????????????????????????????????????????????????????????????????????????????????????????????????????????????????????????????????????????????????????????????????????????????????????????????????????????????????????????????????????????????????????????????????????????????????????????????????????????????????????????????????????????????????????????????????????????????????

			>Hohenbergia_rosea
			??????????????????????????????????????????????????????????????????????????????????????????????????????????????????????????????????????????????????????????????????????????????????????????????????????????????????????????????????????????????????????????????????????????????????????????????????????????????????????????????????????????????????????????????????????????????????????????????????????????????????????????????????????????????????????????????????????????????????????????????????????????????????????????????????????????????????????????????????????????????????????????????????????????????????????????????????????????????????????????????????????????????????????????????????????????????????????????????????????????????????????????????????????????????????????????????

			>Hohenbergia_salzmannii
			??????????????????????????????????????????????????????????????????????????????????????????????????????????????????????????????????????????????????????????????????????????????????????????????????????????????????????????????????????????????????????????????????????????????????????????????????????????????????????????????????????????????????????????????????????????????????????????????????????????????????????????????????????????????????????????????????????????????????????????????????????????????????????????????????????????????????????????????????????????????????????????????????????????????????????????????????????????????????????????????????????????????????????????????????????????????????????????????????????????????????????????????????????????????????????????????

			>Hohenbergia_stellata
			ATAATTATAATAAAGTGAAATATGTCGAAATCCTTTTTGGAATAATACCAAATCGAAAATAAATGTCCGATAGCAAGTTGATCAGTTAATTCAATAAGAGATAAATGGGAGATAGCACTCGATTTAGTTGGTACCACCCAACCGAATCCGATTCACTCATTTACTCATTGAATGAGTAAATTTTCAAGTTCAGCCAATCTTTTTTTTTTGAAAAAAAAAATTGCAAGTAGATGAAATCGTGAATAAATGTGTTTCATTTCTATATCATTATATAAAAAAAGATCCATATTATATTACTTATGGAATTCGAACCCGAACTCGATTTATGATTTATTATTTGGATCTTATTGGCCTTTGTTCTTTATTTTTGATTTTTCATATAGATTTCCGTCTTTATATTATACCCTTTCGTGGATGAATTATGCCTATTTTCACATCTAGGATTTACATATACAACATATATTACTGTCAAGAGGGAATTTTTCTCAGTATTTAGATATTTAGATTCAAAATAAGAAAGGGATCAATTCAATTATAAACCTGCAAAACAAAGATTAGGATTGGGTTGGGTTGCGCTATATATATCAAAGAGTATACAATAATGATGTATTTGGTGAATCAAATACATGGTCTAATAACAAACCATTTTAACATAACATTTTAATGAGTTGATAATATTAATTGAATATTGAATATTTTTTGAAAGATTTTTGTTAAAGGTTTCATTCATGCCTAATCCATATCGAGTAGACCTTGTTGTTGTG??

			>Hohenbergia_utriculosa
			??????????????????????????????????????????????????????????????????????????????????????????????????????????????????????????????????????????????????????????????????????????????????????????????????????????????????????????????????????????????????????????????????????????????????????????????????????????????????????????????????????????????????????????????????????????????????????????????????????????????????????????????????????????????????????????????????????????????????????????????????????????????????????????????????????????????????????????????????????????????????????????????????????????????????????????????????????????????????????????????????????????????????????????????????????????????????????????????????????????????????????????????????????????????????????????????

			>Hohenbergiopsis_guatemalensis
			ATAATTATAATAAAGTGAAATATGTCGAAATCCTTTTTGGAATAATACCAAATCGAAAATAAATGTCCGATAGCAAGTTGATCAGTTAATTCAATAAGAGATAAATGGGAGATAGCACTCGATTTAGTTGGTACCACCCAACCGAATCCGATTCACTCGTTTACTCATTCAATGAGTAAATTTTCAAGTTCAGCCAATCTTTTTTTTTGAAAAAAAAAAATTGCAAGTAGATGAAATCGTGAATAAATGTGTTTCATTTCTATATCATTATATAAAAAAAGATCCATATTATATTACTTATGGAATTCGAACCCGAACTCGATTTATGATTTATTATTTGGATCTTATTGGCCTTTGTTCTTTATTTTTGATTTTTCATATAGATTTCCGTCTTTATATTATACCCTTTCGTGGATGAATTATGCCTATTTTCACATCTAGGATTTACATATACAACATATATTACTGTCAAGAGGGAATTTTTCTCAGTATTTAGATATTTAGATTCAAAATAAGAAAGGGATCAATTCAATTATAAACCTGCAAAACAAAGATTAGGATTGGGTTGGGTTGCGCTATATATATCAAAGAGTATACAATAATGATGTATTTGGTGAATCAAATACATGGTCTAATAACAAACCATTTTAACATAACATTTTAATGAGTTGATAATATTAATTGAATATTGAATATTTTTTGAAAGATTTTTGTTAAAGGTTTCGTTCATGCCTAATCCATATCGAGTAGACCTTGTTGTTGTG??

			>Hoplocryptanthus_caracensis
			??????????????????????????????????????????????????????????????????????????????????????????????????????????????????????????????????????????????????????????????????????????????????????????????????????????????????????????????????????????????????????????????????????????????????????????????????????????????????????????????????????????????????????????????????????????????????????????????????????????????????????????????????????????????????????????????????????????????????????????????????????????????????????????????????????????????????????????????????????????????????????????????????????????????????????????????????????????????????????????????????????????????????????????????????????????????????????????????????????????????????????????????????????????????????????????????

			>Hoplocryptanthus_ferrarius
			??????????????????????????????????????????????????????????????????????????????????????????????????????????????????????????????????????????????????????????????????????????????????????????????????????????????????????????????????????????????????????????????????????????????????????????????????????????????????????????????????????????????????????????????????????????????????????????????????????????????????????????????????????????????????????????????????????????????????????????????????????????????????????????????????????????????????????????????????????????????????????????????????????????????????????????????????????????????????????????????????????????????????????????????????????????????????????????????????????????????????????????????????????????????????????????????

			>Hoplocryptanthus_glaziovii
			ATAATTATAATAAAGTAAAATATGTCGAAATCCTTTTTGGAATAATACCAAATCGAAAATAAATGTCCGATAGCAAGTTGATCAGTTAATTCAATAAGAGATAAATGGGAGATAGCACTCGATTTAGTTGGTACCACCCAACCGAATCCGATTCACTCGTTTACTCATTGAATGAGTAAATTTTCAAGTTCAGCCAATCTTTATTTATTTGAAAAAAAAATTGCAAGTAGATGAAATCGTGAATAAATGTGTTTCATTTCTATATCATTATAGAAAAAAAGATCCATATTATATTACTTATGGAATTCGAACCCGAACTCGATTTATGATTTATTATTTGGATCTTATTGGCCTTTGTTCTTTATTTTTGATTTTTCATATAGATTTCCGTCTTTATATTAT---------TGGATGAATAATGCCTATTTTCACATCTAGGATTTACATATACAACATATATTACTGTCAAGAGGGAATTTTTCTCAGTATTTAGATATTTAGATTCAAAATAAGAAAGGGATCAATTCAATTATAAACCTGCAAAACAAAGATTAGGATTGGGTTGGGTTGCGCTATATATATCAAAGAGTATACAATAATGATGTATTTGGTGAATCAAATACATGGTCTAATAACAAACCATTTTAACATACCATTTTAATGAGTTGATAATATTAATTGAATATTGAATATTTTTTGAAAGATTTTTGTTAAAGGTTTCATTCATGCCTAATCCATATCGAGTAGACCTTGTTGTTGTG??

			>Hoplocryptanthus_lavrasensis
			??????????????????????????????????????????????????????????????????????????????????????????????????????????????????????????????????????????????????????????????????????????????????????????????????????????????????????????????????????????????????????????????????????????????????????????????????????????????????????????????????????????????????????????????????????????????????????????????????????????????????????????????????????????????????????????????????????????????????????????????????????????????????????????????????????????????????????????????????????????????????????????????????????????????????????????????????????????????????????????????????????????????????????????????????????????????????????????????????????????????????????????????????????????????????????????????

			>Hoplocryptanthus_regius
			??????????????????????????????????????????????????????????????????????????????????????????????????????????????????????????????????????????????????????????????????????????????????????????????????????????????????????????????????????????????????????????????????????????????????????????????????????????????????????????????????????????????????????????????????????????????????????????????????????????????????????????????????????????????????????????????????????????????????????????????????????????????????????????????????????????????????????????????????????????????????????????????????????????????????????????????????????????????????????????????????????????????????????????????????????????????????????????????????????????????????????????????????????????????????????????????

			>Hoplocryptanthus_schwackeanus
			??????????????????????????????????????????????????????????????????????????????????????????????????????????????????????????????????????????????????????????????????????????????????????????????????????????????????????????????????????????????????????????????????????????????????????????????????????????????????????????????????????????????????????????????????????????????????????????????????????????????????????????????????????????????????????????????????????????????????????????????????????????????????????????????????????????????????????????????????????????????????????????????????????????????????????????????????????????????????????????????????????????????????????????????????????????????????????????????????????????????????????????????????????????????????????????????

			>Hoplocryptanthus_tiradentesensis
			??????????????????????????????????????????????????????????????????????????????????????????????????????????????????????????????????????????????????????????????????????????????????????????????????????????????????????????????????????????????????????????????????????????????????????????????????????????????????????????????????????????????????????????????????????????????????????????????????????????????????????????????????????????????????????????????????????????????????????????????????????????????????????????????????????????????????????????????????????????????????????????????????????????????????????????????????????????????????????????????????????????????????????????????????????????????????????????????????????????????????????????????????????????????????????????????

			>Hoplocryptanthus_vidaliorum
			??????????????????????????????????????????????????????????????????????????????????????????????????????????????????????????????????????????????????????????????????????????????????????????????????????????????????????????????????????????????????????????????????????????????????????????????????????????????????????????????????????????????????????????????????????????????????????????????????????????????????????????????????????????????????????????????????????????????????????????????????????????????????????????????????????????????????????????????????????????????????????????????????????????????????????????????????????????????????????????????????????????????????????????????????????????????????????????????????????????????????????????????????????????????????????????????

			>Jagrantia_monstrum
			ATAATTATAATAAAGTGAAATATGTCGAAATCCTTTTTGGAATAATACCAAATCGAAAATAAATGTCCGATAGCAAGTTGATCAGTTAATTCAATAAGAGATAAATGGGAGATAGCACTCGATTTAGTTGGTACCACCCAACCGAATCCGATTCAATCGTTTACTCATTCAATGAGTCAATTTTCAAGTTCAGCCAATC--TTTTTTTTCAAAAAAAAAATTGCAAGTAAATGAAATCGTGAATAAATGTGTTTCATTTCTCTATCATTATAGAAAAAAAGATCCATATTATATTACTTATGAAATTTGAACCCGAACTCGATTTATGATTCATTATTTCGATCTTATTGGCCTTTGTTCTTTATTTTTGATTTTTCATATAGATTTCCGTCTTTATATTATACCCTTTCGTAGATGAATTATGCTTATTTTCACATCTAGGATTTACATATACAACATATATTACTGTCAAGAGGGAATTTTTCTCAGTATTTAGATATTTAGATTCAAAATAAGAAAGGGATCAATTCAATTATAAACCTGCAAAACAAAGATTAGGATTGGGTTGGGTTGCGCTATATATATCAAAGAGTATACAATAATGATGTATTTGGTGAATCAAATACATGGTCTAATAACAAACCATTTTAACATAACATTTTAATGAGTTGATAATATTAATTGAATATTGAATATTTTTTGAAAGATTTTTGTTAAAGGTTTCATTCATGCCTAATCCATATCGAGTAGACCTTGTTGTTGTGAG

			>Josemania_asplundii
			??????????????????????????????????????????????????????????????????????????????????????????????????????????????????????????????????????????????????????????????????????????????????????????????????????????????????????????????????????????????????????????????????????????????????????????????????????????????????????????????????????????????????????????????????????????????????????????????????????????????????????????????????????????????????????????????????????????????????????????????????????????????????????????????????????????????????????????????????????????????????????????????????????????????????????????????????????????????????????????????????????????????????????????????????????????????????????????????????????????????????????????????????????????????????????????????

			>Josemania_singularis
			ATAATTATAATAAAGTGAAATATGTCGAAATCCTTTTTGGATTAATACCAAATCGAAAATAAATGTCCGATAGCAAGTTGATCAGTTAATTCAATAAGAGATAAATGGGAGATAGCACTCGATTTAGTTGGTACCACCCAACCGAATCCGATTCAATCGTTTACTCATTCAATGAGTCAATTTTCAAGTTCAGCCAATC--TTTTTTTTGAAAAAAAAAATTGCAAGTAAATGAAATCGTGAATAAATGTGTTTCATTTCTCTATCATTATAGAAAAAAGGATCCATATTATATTACTTATGGAATTCGAACCCGAACTCGATTTATGATTCATTATTTCGATCTTATTGGCCTTTGTTCTTTATTTTTGATTTTTCATATAGATTTCCGTCTTTATATTATACCCTTTCGTAGATGAATTATGCTTATTTTCACATCTAGGATTTACATATACAACATATATTACTGTCAAGAGGGAATTTTTCTCAGTATTTAGATATTTAGATTCAAAATAAGAAAGGGATTAATTCAATTATAAACCTGCAAAACAAAGATTAGGATTGGGTTGGGTTGCGCTATATATATCAAAGAGTATACAATAATGATGTATTTGGTGAATCAAATACATGGTCTAATAACAAACCATTTTAACATAACATTTTAATGAGTTGATAATATTAATTGAATATTGAATATTTTTTGAAAGATTTTTGTTAAAGGTTTCATTCATGCCTAATCCATATCGAGTAGACCTTGTTGTTGTGAG

			>Karawata_depressa
			??????????????????????????????????????????????????????????????????????????????????????????????????????????????????????????????????????????????????????????????????????????????????????????????????????????????????????????????????????????????????????????????????????????????????????????????????????????????????????????????????????????????????????????????????????????????????????????????????????????????????????????????????????????????????????????????????????????????????????????????????????????????????????????????????????????????????????????????????????????????????????????????????????????????????????????????????????????????????????????????????????????????????????????????????????????????????????????????????????????????????????????????????????????????????????????????

			>Karawata_gustavoi
			??????????????????????????????????????????????????????????????????????????????????????????????????????????????????????????????????????????????????????????????????????????????????????????????????????????????????????????????????????????????????????????????????????????????????????????????????????????????????????????????????????????????????????????????????????????????????????????????????????????????????????????????????????????????????????????????????????????????????????????????????????????????????????????????????????????????????????????????????????????????????????????????????????????????????????????????????????????????????????????????????????????????????????????????????????????????????????????????????????????????????????????????????????????????????????????????

			>Karawata_hostilis
			??????????????????????????????????????????????????????????????????????????????????????????????????????????????????????????????????????????????????????????????????????????????????????????????????????????????????????????????????????????????????????????????????????????????????????????????????????????????????????????????????????????????????????????????????????????????????????????????????????????????????????????????????????????????????????????????????????????????????????????????????????????????????????????????????????????????????????????????????????????????????????????????????????????????????????????????????????????????????????????????????????????????????????????????????????????????????????????????????????????????????????????????????????????????????????????????

			>Karawata_multiflora
			??????????????????????????????????????????????????????????????????????????????????????????????????????????????????????????????????????????????????????????????????????????????????????????????????????????????????????????????????????????????????????????????????????????????????????????????????????????????????????????????????????????????????????????????????????????????????????????????????????????????????????????????????????????????????????????????????????????????????????????????????????????????????????????????????????????????????????????????????????????????????????????????????????????????????????????????????????????????????????????????????????????????????????????????????????????????????????????????????????????????????????????????????????????????????????????????

			>Karawata_nigribracteata
			??????????????????????????????????????????????????????????????????????????????????????????????????????????????????????????????????????????????????????????????????????????????????????????????????????????????????????????????????????????????????????????????????????????????????????????????????????????????????????????????????????????????????????????????????????????????????????????????????????????????????????????????????????????????????????????????????????????????????????????????????????????????????????????????????????????????????????????????????????????????????????????????????????????????????????????????????????????????????????????????????????????????????????????????????????????????????????????????????????????????????????????????????????????????????????????????

			>Karawata_saxicola
			??????????????????????????????????????????????????????????????????????????????????????????????????????????????????????????????????????????????????????????????????????????????????????????????????????????????????????????????????????????????????????????????????????????????????????????????????????????????????????????????????????????????????????????????????????????????????????????????????????????????????????????????????????????????????????????????????????????????????????????????????????????????????????????????????????????????????????????????????????????????????????????????????????????????????????????????????????????????????????????????????????????????????????????????????????????????????????????????????????????????????????????????????????????????????????????????

			>Lapanthus_duartei
			ATAATTATAATAAAGTGAAATATGTCGAAATCCTTTTTGGAATAATACCAAATCGAAAATAAATGTCCGATAGCAAGTTGATCAGTTAATTCAATAAGAGATAAATGGGAGATAGCACTCGATTTAGTTGGTACCACCCAACCGAATCCGATTCACTCGTTTACTCATTGAATGAGTAAATTTTCAAGTTCAGCCAATCTTTTTTTTTTTAAAAAAAAAATTGCAAGTAGATGAAATCGTGAATAAATGTGTTTCATTTCTATATCATTATAGAAAAATAGATCCATATTATATTACTTATGGAATTCGAACCCGAACTCGATTTATGATTTATTATTTGGATCTTATTGGCCTTTGTTCTTTATTTTTGATTTTTCATATAGATTTCCGTCTTTATATTATACCCTTTCGTGGATGAATTATGCCTATTTTCACATCTAGGATTTACATATACAACATATATTACTGTCAAGAGGGAATTTTTCTCAGTATTTAGATATTTAGATTCAAAATAAGAAAGGGATCAATTCAATTATAAACCTGCAAAACAAGGATTAGGATTGGGTTGGGTTGCGCTATATATATCAAAGAGTATACAATAATGATGTATTTGGTGAATCAAATACATGGTCTAATAACAAACCATTTTAACATAACATTTTAATGAGTTGATAATATTAATTGAATATTGAATATTTTTTGAAAGATTTTTGTTAAAGGTTTCATTCATGCCTAATCCATATCGAGTAGACCTTGTTGTTGTG??

			>Lapanthus_itambensis
			??????????????????????????????????????????????????????????????????????????????????????????????????????????????????????????????????????????????????????????????????????????????????????????????????????????????????????????????????????????????????????????????????????????????????????????????????????????????????????????????????????????????????????????????????????????????????????????????????????????????????????????????????????????????????????????????????????????????????????????????????????????????????????????????????????????????????????????????????????????????????????????????????????????????????????????????????????????????????????????????????????????????????????????????????????????????????????????????????????????????????????????????????????????????????????????????

			>Lemeltonia_dodsonii
			ATAATTATAATAAAGTGAAATATGTCGAAATCCTTTTTGGAATAATACCAAATCGAAAATAAATGTCCGATAGCAAGTTGATCAGTTAATTCAATAAGAGATAAATGGGAGATAGCACTCGATTTAGTTGGTACCACCCAACCGAATCCGATTCAATCGTTTACTCATTCAATGAGTAAATTTTCAAGTTCAGCCAATC--TTTTTTTTCAAAAAAAAAATTGCAAGTAAATGAAATCGTGAATAAATGTGTTTCATTTCTCTATCATTATAGAAAAAAAGATCCATATTATATTACTTATGGAATTCGAACCCGAACTCGATTTATGATTCATTATTTCGATCTTATTGACCTTTGTTCTTTATTTTTTTTTTTTCATATAGATTTCCGTCTTTATATTATACCCTTTCGTAGATAAATTATGCTTATTTTCACATCTAGGATTTACATATACAACATATATTACTGTCAAGAGGGAATTTTTCTCAGTATTTAGATATTTAGATTCAAAATAAGAAAGGGATCAATTCAATTATAAACCCGCAAAACAAAGATTAGGATTGG-----GTTGCGCTATATATATCAAAGAGTATACAATAATGATGTATTTGGTGAATCAAATACATGGTCTAATAACAAACCATTTTAACATAACATTTTAATGAGTTGATAATATTAATTGAATATTGAATATTTTTTGAAAGATTTTTGTTAAAGGTTTCATTCATGCCTAATCCATATCGAGTAGACCTTGTTGTTGTGAG

			>Lemeltonia_monadelpha
			??????????????????????????????????????????????????????????????????????????????????????????????????????????????????????????????????????????????????????????????????????????????????????????????????????????????????????????????????????????????????????????????????????????????????????????????????????????????????????????????????????????????????????????????????????????????????????????????????????????????????????????????????????????????????????????????????????????????????????????????????????????????????????????????????????????????????????????????????????????????????????????????????????????????????????????????????????????????????????????????????????????????????????????????????????????????????????????????????????????????????????????????????????????????????????????????

			>Lemeltonia_narthecioides
			ATAATTATAATAAAGTGAAATATGTCGAAATCCTTTTTGGAATAATACCAAATCGAAAATAAATGTCCGATAGCAAGTTGATCAGTTAATTCAATAAGAGATAAATGGGAGATAGCACTCGATTTAGTTGGTACCACCCAACCGAATCCGATTCAATCGTTTACTCATTCAATGAGTAAATTTTCAAGTTCAGCCAATCTTTTTTTTC--AAAAAAAAAATTGCAAGTAAATGAAATCGTGAATAAATGTGTTTCATTTCTCTATCATTATAGAAAAAAAGATCCATATTATATTACTTATGGAATTCGAACCCGAACTCAATTTATGATTCATTATTTCGATCTTATTGGCCTTTGTTCTTTATTTTTTATTTTTCATATGGATTTCCGTCTTTATATTATACCCTTTCGTAGATAAATTATGCTTATTTTCACATCTAGGATTTACATATACAACATATATTACTGTCAAGAGGGAATTTTTCTCAGTATTTAGATATTTAGATTCAAAATAAGAAAGGG-----TTCAATTATAAACCCGCA----AAAGATTAGGA-----TTGGGTTGCGCTATATATATCAAAGAGTATACAATAATGATGTATTTGGTGAATCAAATACATGGTCTAATAACAAACCATTTTAACATAACATTTTAATGAGTTGATAATATTAATTGAATATTGAATATTTTTTGAAAGATTTTTGTTAAAGGTTTCATTCATGCCTAATCCATATCGAGTAGACCTTGTTGTTGTGAG

			>Lemeltonia_triglochinoides
			??????????????????????????????????????????????????????????????????????????????????????????????????????????????????????????????????????????????????????????????????????????????????????????????????????????????????????????????????????????????????????????????????????????????????????????????????????????????????????????????????????????????????????????????????????????????????????????????????????????????????????????????????????????????????????????????????????????????????????????????????????????????????????????????????????????????????????????????????????????????????????????????????????????????????????????????????????????????????????????????????????????????????????????????????????????????????????????????????????????????????????????????????????????????????????????????

			>Lindmania_guianensis
			ATAATTATAATAAAGTGAAATATGTCGAAATCCTTTTTGGAATAATACCAAATCGAAAATAAATGTCCGATAGCAAGTTGATCAGTTAATTCAATAAGAGATAAATGGGAGATAGCACTCTATTTAGTTGGTACCACCCAACCGAATCCGATTCAATCGTTTACTCATTCAATGAGTCAATTTTCAAGTTCAGCCAATC--TTTTTTTTTAAAAAAAAAATTGCAAGTAGATGAAATCGTGACTAAATGTGTTTCATTTCTCTATCATTATAGAAAATAAGATCCATATTATATTACTTATGGAATTCGAACCCGAACTCGATTTATGATTCATTATTTCGATCTTATTGGCCTTTGTTCTTTATTTTTTATTTTTCATATAGATTTCCGTCTTTATATTATACCCTTTCGTGGATGAATTATGTCTATTTTCACATCTAGGATTTACATATACAACATATATTACTGTCAAGAGGGAATTTTTCTCAGTATTTAGATATTTAGATTCAAAATAAGAAAGGGATCAATTCAATTATAAACCTGCAAAACAAAGATTAGGATTGGATTGGGTTGCGCTATATATATCAAAGAGTATACAATAATGATGTATTTGGTGAATCAAATACATGGTCTAATAACAAACCATTTTAACATAACATTTTAATGAGTTGATAATATTAATTGAATATTGAATATTTTTTGAAAGATTTTTGTTAAAGGTTTCAGTCATGCCTAATCCATATCGAGTAGACCTTGTTGTTGTGAG

			>Lindmania_longipes
			ATAATTATAATAAAGTGAAATATGTCGAAATCCTTTTTGGAATAATACCAAATC-AAAATAAATGTCCGATAGCAAGTTGATCAGTTAATTCAATAAGAGATAAATGGGAGATAGCACTCTATTTAGTTGGTACCACCCAACCGAATCCGATTCAATCGTTTACTCATTCAATGAGTCAATTTTCAAGTTCAGCCAATCTTTTTTTTTTCAAAAAAAAAATTGCAAGTAGATGAAATCGTGAATAAATGTGTTTCATTTCTCTATCATTATAGAAAATAAGATCCATATTATATTACTTATGGAATTCGAACCCGAACTCGATTTATGATTCATTATTTCGATCTTATTGGCCTTTGTTCTTTATTTTTGATTTTTCATATAGATTTCCGTCTTTATATTATACCCTTTCGTGGATGAATTATGCCTATTTTCACATCTAGGATTTACATATACAACATATATTACTGTCAAGAGGGAATTTTTCTCAGTATTTAGATATTTAGATTCAAAATAAGAAAGGGATCAATTCAATTATAAACCTGCAAAACAAAGATTAGGATTGGATTGGGTTGCGCTATATATATCAAAGAGTATACAATAATGATGTATTTGGTGAATCAAATACATGGTCTAATAACAAACCATTTTAACATAACATTTTAATGAGTTGATAATATTAATTGAATATTGAATATTTTTGGAAAGATTTTTGTTAAAGGTTTCAGTCATGCCTAATCCATATCGAGTAGACCTCGTTGTTGTGAG

			>Lutheria_glutinosa
			ATAATTATAATAAAGTGAAATATGTCGAAATCCTTTTTGGAATAATACCAAATCGAAAATAAATGTCCGATAGCAAGTTGATCAGTTAATTCAATAAGAGATAAATGGGAGATAGCACTCGATTTAGTTGGTACCACCCAACCGAATCCGATTCAATCGTTTACTCATTCAATGAGTCAATTTTCAAGTTCAGCCAATCCTTTTTTTTTTAAAAAAAAAATTGCAAGTAAATGAAATCGTGAATAAATGTGTTTCATTTCTCTATCATTATAGAAAAAAAGATCCATATTATATTACTTATGGAATTCGAACCCGAACTCGATTTATGATTCATTATTTTGATCTTATTGGCTTTTGTTCTTTATTTTTTATTTTTCATATAGATTTCCGTCTTTATATTATACCCTTTCGTAGATGAATTATGCTTATTTTCACATCTAGGATTTACATATACAACATATATTACTGTCAAGAGGGAATTTTTCTCAGTATTTAGATATTTAGATTCAAAATAAGAAAGGGATCAATTCAATTATAAACCTGCAAAACAAAGATTAGGATTGGGTTGGGTTGCGCTATATATATCAAAGAGTATACAATAATGATGTATTTGGTGAATCAAATACATGGTCTAATAACAAACCATTTTAACATAACATTTTAATGAGTTGATAATATTAATTGAATATTGAATATTTTTTGAAAGATTTTTGTTAAAGGTTTCATTCATGCCTAATCCATATCGAGTAGACCTTGTTGTTGTGAG

			>Lutheria_splendens
			ATAATTATAATAAAGTGAAATATGTCGAAATCCTTTTTGGAATAATACCAAATCGAAAATAAATGTCCGATAGCAAGTTGATCAGTTAATTCAATAAGAGATAAATGGGAGATAGCACTCGATTTAGTTGGTACCACCCAACCGAATCCGATTCAATCGTTTACTCATTCAATGAGTCAATTTTCAAGTTCAGCCAATCTTTTTTTTTTC-AAAAAAAAATTGCAAGTAAATGAAATCGTGAATAAATGTGTTTCATTTCTCTATCATTATAGAAAAAAAGATCCATATTATATTACTTATGGAATTCGAACCCGAACTCGATTTATGATTCATTATTTCGATCTTATTGGCCTTTGTTCTTTATTTTTTATTTTTCATATAGATTTCCGTCTTTATATTATACCCTTTCGTAGATGAATTATGCTTATTTTCACATCTAGGATTTACATATACAACATATATTACTGTCAAGAGGGAATTTTTCTCAGTATTTAGATATTTAGATTCAAAATAAGAAAGGGATCAATTCAATTATAAACCTGCAAAACAAAGATTAGGATTGGGTTGGGTTGCGCTATATATATCAAAGAGTATACAATAATGATGTATTTGGTGAATCAAATACATGGTCTAATAACAAACCATTTTAACATAACATTTTAATGAGTTGATAATATTAATTGAATATTGAATATTTTTTGAAAGATTTTTGTTAAAGGTTTCATTCATGCCTAATCCATATCGAGTAGACCTTGTTGTTGTGAG

			>Lymania_alvimii
			ATAATTATAATAAAGTGAAATATGTCGAAATCCTTTTTGGAATAATACCAAATCGAAAATAAATGTCCGATAGCAAGTTGATCAGTTAATTCAATAAGAGATAAATGGGAGATAGCACTCGATTTAGTTGGTACCACCCAACCGAATCCGATTCACTCGTTTACTCATTGAATGAGTAAATTTTCAAGTTCAGCCAATC-TTTTTTTTTTAAAAAAAAAATTGCAAGTAGATGAAATCGTGAATAAATGTGTTTCATTTCTATATCATTATATAAAAAAAGATCCATATTATATTACTTATGGAATTCGAACCCGAACTCGATTTATGATTTATTATTTGGATCTTATTGGCCTTTGTTCTTTATTTTTTATTTTTCATATAGATTTCCGTCTTTATATTATACCCTTTCGTGGATGAATTATGCCTATTTTCACATCTAGGATTTACATATACAACATATATTACTGTCAAGAGGGAATTTTTCTCAGTATTTAGATATTTAGATTCAAAATAAGAAAGGGATCAATTCAATTATAAACCTGCAAAACAAAGATTAGGATTGGGTTGGGTTGCGCTATATATATCAAAGAGTATACAATAATGATGTATTTGGTGAATCAAATACATGGTCTAATAACAAACCATTTTAACATAACATTTTAATGAGTTGATAATATTAATTGAATATTGAATATTTTTTGAAAGATTTTTGTTAAAGGTTTCATTCATGCCTAATCCATATCGAGTAGACCTTGTTGTTGTG??

			>Lymania_azurea
			??????????????????????????????????????????????????????????????????????????????????????????????????????????????????????????????????????????????????????????????????????????????????????????????????????????????????????????????????????????????????????????????????????????????????????????????????????????????????????????????????????????????????????????????????????????????????????????????????????????????????????????????????????????????????????????????????????????????????????????????????????????????????????????????????????????????????????????????????????????????????????????????????????????????????????????????????????????????????????????????????????????????????????????????????????????????????????????????????????????????????????????????????????????????????????????????

			>Lymania_brachycaulis
			??????????????????????????????????????????????????????????????????????????????????????????????????????????????????????????????????????????????????????????????????????????????????????????????????????????????????????????????????????????????????????????????????????????????????????????????????????????????????????????????????????????????????????????????????????????????????????????????????????????????????????????????????????????????????????????????????????????????????????????????????????????????????????????????????????????????????????????????????????????????????????????????????????????????????????????????????????????????????????????????????????????????????????????????????????????????????????????????????????????????????????????????????????????????????????????????

			>Lymania_corallina
			???????????????????????????????????TTTGGAATAATACCAAATCGAAAATAAATGTCCGATAGCAAGTTGATCAGTTAATTCAATAAGAGATAAATGGGAGATAGCACTCGATTTAGTTGGTACCACCCAACCGAATCCGATTCACTCGTTTACTCATTCAATGAGTAAATTTTCAAGTTCAGCCAATC-TTTTTTTTTTAAAAAAAAAATTGCAAGTAGATGAAATCGTGAATAAATGTGTTTCATTTCTATATCATTATATAAAAAAAGATCCATATTATATTACTTATGGAATTCGAACCCGAACTCGATTTATGATTTATTATTTGGATCTTATTGGCCTTTGTTCTTTATTTTTGATTTTGCATATAGATTTCCGTCTTTATATTATACCCTTTCGTGGATGAATTATGCCTATTTTCACATCTAGGATTTACATATACAACATATATTACTGTCAAGAGGGAATTTTTCTCAGTATTTAGATATTTAGATTCAAAATAAGAAAGGGATCAATTCAATTATAAA-CTGCAAAACAAAGATTAGGATTGGGTTGGGTTGCGCTATATATATCAAAGAGTATACAATAATGATGTATTTGGTGAATCAAATACATGGTCTAATAACAAACCATTTTAACATAACATTTTAATGAGTTGATAATATTAATTGAATATTGAATATTTTTTGAAAGATTTTTGTTAAAGGTTTCATTCATGCCTAATCCATATCGAGTAGACCTTGTTGTTGTG??

			>Lymania_globosa
			??????????????????????????????????????????????????????????????????????????????????????????????????????????????????????????????????????????????????????????????????????????????????????????????????????????????????????????????????????????????????????????????????????????????????????????????????????????????????????????????????????????????????????????????????????????????????????????????????????????????????????????????????????????????????????????????????????????????????????????????????????????????????????????????????????????????????????????????????????????????????????????????????????????????????????????????????????????????????????????????????????????????????????????????????????????????????????????????????????????????????????????????????????????????????????????????

			>Lymania_languida
			??????????????????????????????????????????????????????????????????????????????????????????????????????????????????????????????????????????????????????????????????????????????????????????????????????????????????????????????????????????????????????????????????????????????????????????????????????????????????????????????????????????????????????????????????????????????????????????????????????????????????????????????????????????????????????????????????????????????????????????????????????????????????????????????????????????????????????????????????????????????????????????????????????????????????????????????????????????????????????????????????????????????????????????????????????????????????????????????????????????????????????????????????????????????????????????????

			>Lymania_smithii
			??????????????????????????????????????????????????????????????????????????????????????????????????????????????????????????????????????????????????????????????????????????????????????????????????????????????????????????????????????????????????????????????????????????????????????????????????????????????????????????????????????????????????????????????????????????????????????????????????????????????????????????????????????????????????????????????????????????????????????????????????????????????????????????????????????????????????????????????????????????????????????????????????????????????????????????????????????????????????????????????????????????????????????????????????????????????????????????????????????????????????????????????????????????????????????????????

			>Lymania_spiculata
			??????????????????????????????????????????????????????????????????????????????????????????????????????????????????????????????????????????????????????????????????????????????????????????????????????????????????????????????????????????????????????????????????????????????????????????????????????????????????????????????????????????????????????????????????????????????????????????????????????????????????????????????????????????????????????????????????????????????????????????????????????????????????????????????????????????????????????????????????????????????????????????????????????????????????????????????????????????????????????????????????????????????????????????????????????????????????????????????????????????????????????????????????????????????????????????????

			>Mezobromelia_bicolor
			??????????????????????????????????????????????????????????????????????????????????????????????????????????????????????????????????????????????????????????????????????????????????????????????????????????????????????????????????????????????????????????????????????????????????????????????????????????????????????????????????????????????????????????????????????????????????????????????????????????????????????????????????????????????????????????????????????????????????????????????????????????????????????????????????????????????????????????????????????????????????????????????????????????????????????????????????????????????????????????????????????????????????????????????????????????????????????????????????????????????????????????????????????????????????????????????

			>Mezobromelia_capituligera
			??????????????????????????????????????????????????????????????????????????????????????????????????????????????????????????????????????????????????????????????????????????????????????????????????????????????????????????????????????????????????????????????????????????????????????????????????????????????????????????????????????????????????????????????????????????????????????????????????????????????????????????????????????????????????????????????????????????????????????????????????????????????????????????????????????????????????????????????????????????????????????????????????????????????????????????????????????????????????????????????????????????????????????????????????????????????????????????????????????????????????????????????????????????????????????????????

			>Mezobromelia_pleiosticha
			ATAATTATAATAAAGTGAAATATGTCGAAATCCTTTTTGGAATAATACCAAATCGAAAATAAATGTCCGATAGCAAGTTGATCAGTTAATTCAATAAGAGATAAATGGGAGATAGCACTCGATTTAGTTGGTACCACCCAACCGAATCCGATTCAATCGTTTACTCATTCAATGAGTCAATTTTCAAGTTCAGCCAATCTTTTTTTTTGAAAAAAAAAAATTGCAAGTAAATGAAATCGTGAATAAATGTGTTTCATTTCTCTATCATTATAGAAAAAAAGATCCATATTATATTACTTATGGAATTCGAACCCGAACTCGATTTATGATTCATTATTTCGATCTTATTGGCCTTTGTTCTTTATTTTTGATTTTTCATATGGATTTCCGTCTTTATATTATACCCTTTCGTAGATGAATTATGCTTATTTTCACATCTAGGATTTACATATACAACATATATTACTGTCAAGAGGGAATTTTTCTCAGTATTTAGATATTTAGATTCAAAATAAGAAAGGGATCAATTCAATTATAAACCTGCAAAACAAAGATTAGGATTGGGTTGGGTTGCGCTATATATATCAAAGAGTATACAATAATGATGTATTTGGTGAATCAAATACATGGTCTAATAACAAACCATTTTAACATAACATTTTAATGAGTTGATAATATTAATTGAATATTGAATATTTTTTGAAAGATTTTTGTTAAAGGTTTCATTCATGCCTAATCCATATCGAGTAGACCTTGTTGTTGTGAG

			>Navia_barbellata
			??????????????????????????????????????????????????????????????????????????????????????????????????????????????????????????????????????????????????????????????????????????????????????????????????????????????????????????????????????????????????????????????????????????????????????????????????????????????????????????????????????????????????????????????????????????????????????????????????????????????????????????????????????????????????????????????????????????????????????????????????????????????????????????????????????????????????????????????????????????????????????????????????????????????????????????????????????????????????????????????????????????????????????????????????????????????????????????????????????????????????????????????????????????????????????????????

			>Navia_igneosicola
			ATAATTATAATAAAGTGAAATATGTCGAAATCCTTTTTGGAATAATACCAAATCGAAAATAAATGTCCGATAGCAAGTTGATCAGTTAATTCAATAAGAGATAAATGGGAGATAGCACTCGATTTAGTTGGTACCACCCAACCGAATCCGATTCAATCGTTTACTCATTCAATGAGTCAATTTTCAAGTTCAGCCAATCTTTTTTTTTTCAAAAAAAAAATTAAAAGTAGATGAAATCGTGAATAAATGTGTTTCATTTTTCTATCATTATAGAAAATAAGATCCATATTATATTACTTATGGAATTCGAACCCGAACTCGATTTATGATTCATTATTTCGATCTTATTGGCCTTTGTTCTTTATTTTTGATTTTTCATATAGATTTCCGTCTTTATATTATACCCTTTCGTGGATGAATTATGCCTATTTGCACATCTAGGATTTACATATACAACATATATTACTGTCAAGAGGGAATTTTCCTCAGTATTTAGATATTTAGATTCAAAATAAGAAAGGGATCAATTCAATTATAAACCTGCAAAACAAAGATTAGGA-----TTGGGTTGCGCTATATATATCAAAGAGTATACAATAATGATGTATTTGGTGAATCAAATACATGGTCTAATAACAAACCATTTTAACATAACATTTTAATGAGTTGATAATATTAATTGAATATTGAATATTTTTTGAAAGATTTTTGTTAAAGGTTTCATTCATGCCTAATCCATATCGAGTAGACCTTGTTGTTGTGAG

			>Navia_phelpsiae
			ATAATTATAATAAAGTGAAATATGTCGAAATCCTTTTTGGAATAATACCAAATCGAAAATAAATGTCCGATAGCAAGTTGATCAGTTAATTCAATAAGAGATAAATGGGAGATAGCACTCGATTTAGTTGGTACCACCCAACCGAATCCGATTCAATCGTTTACTCATTCAATGAGTCAATTTTCAAGTTCAGCCAATCTTTTTTTTTTTCAAAAAAAAATTGCAAGTAGATGAAATCGTGAATAAATGTGTTTCATTTCTCTATCATTATAGAAAATAAGATCCATATTATATTACTTATGGAATTCGAACCCGAACTCGATTTATGATTCATTATTTCGATCTTATTGGCCTTTGTTCTTTATTTTTGATTTTTCATATAGATTTCCGTCTTTATATTATACCCTTTCGTGGATGAATTATGCCTATTTTCACATCTAGGATTTACATATACAACATATATTACTGTCAAGAGGGAATTTTCCTCAGTATTTAGATATTTAGATTCAAAATAAGAAAGGGATCAATTCAATTATAAACCTGCAAAACAAAGATTAGGA-----TTGGGTTGCGCTATATATATCAAAGAGTATACAATAATGATGTATTTGGTGAATCAAATACATGGTCTAATAACAAACCATTTTAACATAACATTTTAATGAGTTGATAATATTAATTGAATATTGAATATTTTTTGAAAGATTTTTGTTAAAGGTTTCATTCATGCCTAATCCATATCGAGTAGACCTTGTTGTTGTGAG

			>Navia_saxicola
			ATAATTATAATAAAGTGAAATATGTCGAAATCCTTTTTGGAATAATACCAAATCGAAAATAAATGTCCGATAGCAAGTTGATCAGTTAATTCAATAAGAGATAAATGGGAGATAGCACTCAATTTAGTTGGTACCACCCAACCGAATCCGATTCAATCGTTTACTCATTCAATGAGTCAATTTTCAAGTTCAGCCAATCTTTTTTTTTTTCAAAAAAAAATTGCAAGTAGATGAAATCGTGAATAAATGTGTTTCATTTCTCTATCATTATAGAAAATAAGATCCATATTATATTACTTATGGAATTCGAACCCGAACTCGATTTATGATTCATTATTTCGATCTTATTGGCCTTTGTTCTTTATTTTGGATTTTTCATATAGATTTCCGTCTTTATATTATACCCTTTCGTGGATGAATTATGCCTATTTTCACATCTAGGATTTACATATACAACATATATTACTGTCAAGAGGGAATTTTCCTCAGTATTTAGATATTTAGATTCAAAATAAGAAAGGGATCAATTCAATTATAAACCTGCAAAACAAAGATTAGGA-----TTGGGTTGCGCTATATATATCAAAGAGTATACAATAATGATGTATTTGGTGAATCAAATACATGGTCTAATAACAAACCATTTTAACATAACATTTTAATGAGTTGATAATATTAATTGAATATTGAATATTTTTTGAAAGATTTTTGTTAAAGGTTTCATTCATGCCTAATCCATATCGAGTAGACCTTGTTGTTGTGAG

			>Neoglaziovia_burle-marxii
			??????????????????????????????????????????????????????????????????????????????????????????????????????????????????????????????????????????????????????????????????????????????????????????????????????????????????????????????????????????????????????????????????????????????????????????????????????????????????????????????????????????????????????????????????????????????????????????????????????????????????????????????????????????????????????????????????????????????????????????????????????????????????????????????????????????????????????????????????????????????????????????????????????????????????????????????????????????????????????????????????????????????????????????????????????????????????????????????????????????????????????????????????????????????????????????????

			>Neoglaziovia_variegata
			ATAATTATAATAAAGTGAAATATGTCGAAATCCTTTTTGGAATAATACCAAATCGAAAATAAATGTCCGATAGCAAGTTGATCAGTTAATTCAATAAGAGATAAATGGGAGATAGCACTCGATTTAGTTGGTACCACCCAACCGAATCCGATTCACTCGTTTACTCATTCAATGAGTAAATTTTCAAGTTCAGCCAATCTTTTTTTTTTTCAAAAAAAAATTGCAAGTAGATGAAATCGTGAATAAATGTGTTTCATTTCTATATCATTATATAAAAAAAGATCCATATTATATTACTTATGGAATTCGAACCCGAACTCGATTTATGATTTATTATTTGGATCTTATTGGCCTTTGTTCTTTATTTTTGATTTTTCATATAGATTTCCGTCTTTATATTATACCCTTTCGTGGATGAATTATGCCTATTTTCACATCTAGGATTTACATATACAACATATATTACTGTCAAGAGGGAATTTTTCTCAGTATTTAGATATTTAGATTCAAAATAAGAAAGGGATCAATTCAATTATAAACCTGCAAAACAAAGATTAGGATTGGGTTGGGTTGCGCTATATATATCAAAGAGTATACAATAATGATGTATTTGGTGAATCAAATACATGGTCTAATAACAAACCATTTTAACATAACATTTTAATGAGTTGATAATATTAATTGAATATTGAATATTTTTTGAAAGATTTTTGTTAAAGGTTTCATTCATGCCTAATCCATATCGAGTAGACCTTGTTGTTGTG??

			>Neoregelia_binotii
			ATAATTATAATAAAGTGAAATATGTCGAAATCCTTTTTGGAATAATACCAAATCGAAAATAAATGTCCGATAGCAAGTTGATCAGTTAATTCAATAAGAGATAAATGGGAGATAGCACTCGATTTAGTTGGTACCACCCAACCGAATCCGATTCACTCGTTTACTCATTCAATGAGTAAATTTTCAAGTTCAGCCAATCAATTTTTTT--AAAAAAAAAATTGCAAGTAGATGAAATCGTGAATAAATGTGTTTCATTTCTATATCATTATATAAAAAAAGATCCATATTATATTACTTATGGAATTCGAACCCGAACTCGATTTATGATTTATTATTTGGATCTTATTGGCCTTTGTTCTTTATTTTTGATTTTTCATATGGATTTCCGTCTTTATATTATACCCTTTCGTGGATGAATTATGCCTATTTTCACATCTAGGATTTACATATACAACATATATTACCCTCAAGAGGGAATTTTTCTCAGTATTTAGATATTTAGATTCAAAATAAGAAAGGGATCAATTCAATTATAAACCTGCAAAACAAAGATTAGGATTGGGTTGGGTTGCGCTATATATATCAAAGAGTATACAATAATGATGTATTTGGTGAATCAAATACATGGTCTAATAACAAACCATTTTAACATAACATTTTAATGAGTTGATAATATTAATTGAATATTGAATATTTTTTGAAAGATTTTTGTTAAAGGTTTCATTCATGCCTAATCCATATCGAGTAGACCTTGTTGTTGTG??

			>Neoregelia_bragarum
			??????????????????????????????????????????????????????????????????????????????????????????????????????????????????????????????????????????????????????????????????????????????????????????????????????????????????????????????????????????????????????????????????????????????????????????????????????????????????????????????????????????????????????????????????????????????????????????????????????????????????????????????????????????????????????????????????????????????????????????????????????????????????????????????????????????????????????????????????????????????????????????????????????????????????????????????????????????????????????????????????????????????????????????????????????????????????????????????????????????????????????????????????????????????????????????????

			>Neoregelia_capixaba
			??????????????????????????????????????????????????????????????????????????????????????????????????????????????????????????????????????????????????????????????????????????????????????????????????????????????????????????????????????????????????????????????????????????????????????????????????????????????????????????????????????????????????????????????????????????????????????????????????????????????????????????????????????????????????????????????????????????????????????????????????????????????????????????????????????????????????????????????????????????????????????????????????????????????????????????????????????????????????????????????????????????????????????????????????????????????????????????????????????????????????????????????????????????????????????????????

			>Neoregelia_carolinae
			??????????????????????????????????????????????????????????????????????????????????????????????????????????????????????????????????????????????????????????????????????????????????????????????????????????????????????????????????????????????????????????????????????????????????????????????????????????????????????????????????????????????????????????????????????????????????????????????????????????????????????????????????????????????????????????????????????????????????????????????????????????????????????????????????????????????????????????????????????????????????????????????????????????????????????????????????????????????????????????????????????????????????????????????????????????????????????????????????????????????????????????????????????????????????????????????

			>Neoregelia_compacta
			??????????????????????????????????????????????????????????????????????????????????????????????????????????????????????????????????????????????????????????????????????????????????????????????????????????????????????????????????????????????????????????????????????????????????????????????????????????????????????????????????????????????????????????????????????????????????????????????????????????????????????????????????????????????????????????????????????????????????????????????????????????????????????????????????????????????????????????????????????????????????????????????????????????????????????????????????????????????????????????????????????????????????????????????????????????????????????????????????????????????????????????????????????????????????????????????

			>Neoregelia_concentrica
			??????????????????????????????????????????????????????????????????????????????????????????????????????????????????????????????????????????????????????????????????????????????????????????????????????????????????????????????????????????????????????????????????????????????????????????????????????????????????????????????????????????????????????????????????????????????????????????????????????????????????????????????????????????????????????????????????????????????????????????????????????????????????????????????????????????????????????????????????????????????????????????????????????????????????????????????????????????????????????????????????????????????????????????????????????????????????????????????????????????????????????????????????????????????????????????????

			>Neoregelia_cruenta
			??????????????????????????????????????????????????????????????????????????????????????????????????????????????????????????????????????????????????????????????????????????????????????????????????????????????????????????????????????????????????????????????????????????????????????????????????????????????????????????????????????????????????????????????????????????????????????????????????????????????????????????????????????????????????????????????????????????????????????????????????????????????????????????????????????????????????????????????????????????????????????????????????????????????????????????????????????????????????????????????????????????????????????????????????????????????????????????????????????????????????????????????????????????????????????????????

			>Neoregelia_dungsiana
			??????????????????????????????????????????????????????????????????????????????????????????????????????????????????????????????????????????????????????????????????????????????????????????????????????????????????????????????????????????????????????????????????????????????????????????????????????????????????????????????????????????????????????????????????????????????????????????????????????????????????????????????????????????????????????????????????????????????????????????????????????????????????????????????????????????????????????????????????????????????????????????????????????????????????????????????????????????????????????????????????????????????????????????????????????????????????????????????????????????????????????????????????????????????????????????????

			>Neoregelia_eleutheropetala
			??????????????????????????????????????????????????????????????????????????????????????????????????????????????????????????????????????????????????????????????????????????????????????????????????????????????????????????????????????????????????????????????????????????????????????????????????????????????????????????????????????????????????????????????????????????????????????????????????????????????????????????????????????????????????????????????????????????????????????????????????????????????????????????????????????????????????????????????????????????????????????????????????????????????????????????????????????????????????????????????????????????????????????????????????????????????????????????????????????????????????????????????????????????????????????????????

			>Neoregelia_lactea
			??????????????????????????????????????????????????????????????????????????????????????????????????????????????????????????????????????????????????????????????????????????????????????????????????????????????????????????????????????????????????????????????????????????????????????????????????????????????????????????????????????????????????????????????????????????????????????????????????????????????????????????????????????????????????????????????????????????????????????????????????????????????????????????????????????????????????????????????????????????????????????????????????????????????????????????????????????????????????????????????????????????????????????????????????????????????????????????????????????????????????????????????????????????????????????????????

			>Neoregelia_laevis
			??????????????????????????????????????????????????????????????????????????????????CAGTTAATTCAATAAGAGATAAATGGGAGATAGCACTCGATTTAGTTGGTACCACCCAACCGAATCCGATTCACTCGTTTACTCATTCAATGAGTAAATTTTCAAGTTCAGCCAATCTTTTTTTTTTTAAAAAAAAAATTGCAAGTAGATGAAATCGTGAATAAATGTGTTTCATTTCTATATCATTATATAAAAAAAGATCCATATTATATTACTTATGGAATTCGAACCCGAACTCGATTTATGATTTATTATTTGGATCTTATTGGCCTTTGTTCTTTATTTTTGATTTTTCATATGGATTTCCGTCTTTATATTATACCCTGTCGTGGATGAATTATGCCTATTTTCACATCTAGGATTTACATATACAACATATATTACCCTCAAGAGGGAATTTTTCTCAGTATTTAGATATTTAGATTCAAAATAAGAAAGGGATCAATTCAATTATAAACCTGCAAAGCAAAGATTAGGATTGGGTTGGGTTGCGCTATATATATCAAAGAGTATACAATAATGATGTATTTGGTGAATCAAATACATGGTCTAATAACAAACCATTTTAACATAACATTTTAATGAGTTGATAATATTAATTGAATATTGAATATTTTTTGAAAGATTTTTGTTAAGGGTTTCATTCATGCCTAATCCATATCGAGTAGACCTTGTTGTTGTGAG

			>Neoregelia_leucophoea
			??????????????????????????????????????????????????????????????????????????????????????????????????????????????????????????????????????????????????????????????????????????????????????????????????????????????????????????????????????????????????????????????????????????????????????????????????????????????????????????????????????????????????????????????????????????????????????????????????????????????????????????????????????????????????????????????????????????????????????????????????????????????????????????????????????????????????????????????????????????????????????????????????????????????????????????????????????????????????????????????????????????????????????????????????????????????????????????????????????????????????????????????????????????????????????????????

			>Neoregelia_longisepala
			??????????????????????????????????????????????????????????????????????????????????????????????????????????????????????????????????????????????????????????????????????????????????????????????????????????????????????????????????????????????????????????????????????????????????????????????????????????????????????????????????????????????????????????????????????????????????????????????????????????????????????????????????????????????????????????????????????????????????????????????????????????????????????????????????????????????????????????????????????????????????????????????????????????????????????????????????????????????????????????????????????????????????????????????????????????????????????????????????????????????????????????????????????????????????????????????

			>Neoregelia_lymaniana
			??????????????????????????????????????????????????????????????????????????????????????????????????????????????????????????????????????????????????????????????????????????????????????????????????????????????????????????????????????????????????????????????????????????????????????????????????????????????????????????????????????????????????????????????????????????????????????????????????????????????????????????????????????????????????????????????????????????????????????????????????????????????????????????????????????????????????????????????????????????????????????????????????????????????????????????????????????????????????????????????????????????????????????????????????????????????????????????????????????????????????????????????????????????????????????????????

			>Neoregelia_mucugensis
			??????????????????????????????????????????????????????????????????????????????????????????????????????????????????????????????????????????????????????????????????????????????????????????????????????????????????????????????????????????????????????????????????????????????????????????????????????????????????????????????????????????????????????????????????????????????????????????????????????????????????????????????????????????????????????????????????????????????????????????????????????????????????????????????????????????????????????????????????????????????????????????????????????????????????????????????????????????????????????????????????????????????????????????????????????????????????????????????????????????????????????????????????????????????????????????????

			>Neoregelia_myrmecophila
			??????????????????????????????????????????????????????????????????????????????????????????????????????????????????????????????????????????????????????????????????????????????????????????????????????????????????????????????????????????????????????????????????????????????????????????????????????????????????????????????????????????????????????????????????????????????????????????????????????????????????????????????????????????????????????????????????????????????????????????????????????????????????????????????????????????????????????????????????????????????????????????????????????????????????????????????????????????????????????????????????????????????????????????????????????????????????????????????????????????????????????????????????????????????????????????????

			>Neoregelia_nevaresii
			??????????????????????????????????????????????????????????????????????????????????????????????????????????????????????????????????????????????????????????????????????????????????????????????????????????????????????????????????????????????????????????????????????????????????????????????????????????????????????????????????????????????????????????????????????????????????????????????????????????????????????????????????????????????????????????????????????????????????????????????????????????????????????????????????????????????????????????????????????????????????????????????????????????????????????????????????????????????????????????????????????????????????????????????????????????????????????????????????????????????????????????????????????????????????????????????

			>Neoregelia_nivea
			??????????????????????????????????????????????????????????????????????????????????????????????????????????????????????????????????????????????????????????????????????????????????????????????????????????????????????????????????????????????????????????????????????????????????????????????????????????????????????????????????????????????????????????????????????????????????????????????????????????????????????????????????????????????????????????????????????????????????????????????????????????????????????????????????????????????????????????????????????????????????????????????????????????????????????????????????????????????????????????????????????????????????????????????????????????????????????????????????????????????????????????????????????????????????????????????

			>Neoregelia_pascoaliana
			??????????????????????????????????????????????????????????????????????????????????????????????????????????????????????????????????????????????????????????????????????????????????????????????????????????????????????????????????????????????????????????????????????????????????????????????????????????????????????????????????????????????????????????????????????????????????????????????????????????????????????????????????????????????????????????????????????????????????????????????????????????????????????????????????????????????????????????????????????????????????????????????????????????????????????????????????????????????????????????????????????????????????????????????????????????????????????????????????????????????????????????????????????????????????????????????

			>Neoregelia_pauciflora
			??????????????????????????????????????????????????????????????????????????????????????????????????????????????????????????????????????????????????????????????????????????????????????????????????????????????????????????????????????????????????????????????????????????????????????????????????????????????????????????????????????????????????????????????????????????????????????????????????????????????????????????????????????????????????????????????????????????????????????????????????????????????????????????????????????????????????????????????????????????????????????????????????????????????????????????????????????????????????????????????????????????????????????????????????????????????????????????????????????????????????????????????????????????????????????????????

			>Neoregelia_pineliana
			ATAATTATAATAAAGTGAAATATGTCGAAATCCTTTTTGGAATAATACCAAATCGAAAATAAATGTCCGATAGCAAGTTGATCAGTTAATTCAATAAGAGATAAATGGGAGATAGCACTCGATTTAGTTGGTACCACCCAACCGAATCCGATTCATTCGTTTACTCATTGAATGAGTAAATTTTCAAGTTCAGCCAATCTTTTTTTTT--AAAAAAAAAATTGCAAGTAGATGAAATCGTGAATAAATGTGTTTCATTTCTATATCATTATATAAAAAAAGATCCATATTATATTACTTATGGAATTCGAACCCGAACTCGATTTATGATTTATTATTTGGATCTTATTGGCCTTTGTTCTTTATTTTTGATTTTTCATATGGATTTCCGTCTTTATATTATACCCTTTCGTGGATGAATTATGCCTATTTTCACATCTAGGATTTACATATACAACATATATTACCCTCAAGAGGGAATTTTTCTCAGTATTTAGATATTTAGATTCAAAATAAGAAAGGGATCAATTCAATTATAAACCTGCAAAACAAAGATTAGGATTGGGTTGGGTTGCGCTATATATATCAAAGAGTATACAATAATGATGTATTTGGTGAATCAAATACATGGTCTAATAACAAACCGTTTTAACATAACATTTTAATGAGTTGATAATATTAATTGAATATTGAATATTTTTTGAAAGATTTTTGTTAAAGGTTTCATTCATGCCTAATCCATATCGAGTAGACCTTGTTGTTGTGAG

			>Neoregelia_retrorsa
			??????????????????????????????????????????????????????????????????????????????????????????????????????????????????????????????????????????????????????????????????????????????????????????????????????????????????????????????????????????????????????????????????????????????????????????????????????????????????????????????????????????????????????????????????????????????????????????????????????????????????????????????????????????????????????????????????????????????????????????????????????????????????????????????????????????????????????????????????????????????????????????????????????????????????????????????????????????????????????????????????????????????????????????????????????????????????????????????????????????????????????????????????????????????????????????????

			>Neoregelia_richteri
			??????????????????????????????????????????????????????????????????????????????????????????????????????????????????????????????????????????????????????????????????????????????????????????????????????????????????????????????????????????????????????????????????????????????????????????????????????????????????????????????????????????????????????????????????????????????????????????????????????????????????????????????????????????????????????????????????????????????????????????????????????????????????????????????????????????????????????????????????????????????????????????????????????????????????????????????????????????????????????????????????????????????????????????????????????????????????????????????????????????????????????????????????????????????????????????????

			>Neoregelia_rothinessa
			??????????????????????????????????????????????????????????????????????????????????????????????????????????????????????????????????????????????????????????????????????????????????????????????????????????????????????????????????????????????????????????????????????????????????????????????????????????????????????????????????????????????????????????????????????????????????????????????????????????????????????????????????????????????????????????????????????????????????????????????????????????????????????????????????????????????????????????????????????????????????????????????????????????????????????????????????????????????????????????????????????????????????????????????????????????????????????????????????????????????????????????????????????????????????????????????

			>Neoregelia_rubrovittata
			??????????????????????????????????????????????????????????????????????????????????????????????????????????????????????????????????????????????????????????????????????????????????????????????????????????????????????????????????????????????????????????????????????????????????????????????????????????????????????????????????????????????????????????????????????????????????????????????????????????????????????????????????????????????????????????????????????????????????????????????????????????????????????????????????????????????????????????????????????????????????????????????????????????????????????????????????????????????????????????????????????????????????????????????????????????????????????????????????????????????????????????????????????????????????????????????

			>Neoregelia_tenebrosa
			??????????????????????????????????????????????????????????????????????????????????????????????????????????????????????????????????????????????????????????????????????????????????????????????????????????????????????????????????????????????????????????????????????????????????????????????????????????????????????????????????????????????????????????????????????????????????????????????????????????????????????????????????????????????????????????????????????????????????????????????????????????????????????????????????????????????????????????????????????????????????????????????????????????????????????????????????????????????????????????????????????????????????????????????????????????????????????????????????????????????????????????????????????????????????????????????

			>Neoregelia_viridovinosa
			??????????????????????????????????????????????????????????????????????????????????????????????????????????????????????????????????????????????????????????????????????????????????????????????????????????????????????????????????????????????????????????????????????????????????????????????????????????????????????????????????????????????????????????????????????????????????????????????????????????????????????????????????????????????????????????????????????????????????????????????????????????????????????????????????????????????????????????????????????????????????????????????????????????????????????????????????????????????????????????????????????????????????????????????????????????????????????????????????????????????????????????????????????????????????????????????

			>Neoregelia_wilsoniana
			??????????????????????????????????????????????????????????????????????????????????????????????????????????????????????????????????????????????????????????????????????????????????????????????????????????????????????????????????????????????????????????????????????????????????????????????????????????????????????????????????????????????????????????????????????????????????????????????????????????????????????????????????????????????????????????????????????????????????????????????????????????????????????????????????????????????????????????????????????????????????????????????????????????????????????????????????????????????????????????????????????????????????????????????????????????????????????????????????????????????????????????????????????????????????????????????

			>Neoregelia_zonata
			??????????????????????????????????????????????????????????????????????????????????????????????????????????????????????????????????????????????????????????????????????????????????????????????????????????????????????????????????????????????????????????????????????????????????????????????????????????????????????????????????????????????????????????????????????????????????????????????????????????????????????????????????????????????????????????????????????????????????????????????????????????????????????????????????????????????????????????????????????????????????????????????????????????????????????????????????????????????????????????????????????????????????????????????????????????????????????????????????????????????????????????????????????????????????????????????

			>Nidularium_altimontanum
			??????????????????????????????????????????????????????????????????????????????????????????????????????????????????????????????????????????????????????????????????????????????????????????????????????????????????????????????????????????????????????????????????????????????????????????????????????????????????????????????????????????????????????????????????????????????????????????????????????????????????????????????????????????????????????????????????????????????????????????????????????????????????????????????????????????????????????????????????????????????????????????????????????????????????????????????????????????????????????????????????????????????????????????????????????????????????????????????????????????????????????????????????????????????????????????????

			>Nidularium_amazonicum
			??????????????????????????????????????????????????????????????????????????????????????????????????????????????????????????????????????????????????????????????????????????????????????????????????????????????????????????????????????????????????????????????????????????????????????????????????????????????????????????????????????????????????????????????????????????????????????????????????????????????????????????????????????????????????????????????????????????????????????????????????????????????????????????????????????????????????????????????????????????????????????????????????????????????????????????????????????????????????????????????????????????????????????????????????????????????????????????????????????????????????????????????????????????????????????????????

			>Nidularium_angustibracteatum
			??????????????????????????????????????????????????????????????????????????????????????????????????????????????????????????????????????????????????????????????????????????????????????????????????????????????????????????????????????????????????????????????????????????????????????????????????????????????????????????????????????????????????????????????????????????????????????????????????????????????????????????????????????????????????????????????????????????????????????????????????????????????????????????????????????????????????????????????????????????????????????????????????????????????????????????????????????????????????????????????????????????????????????????????????????????????????????????????????????????????????????????????????????????????????????????????

			>Nidularium_campoalegrense
			??????????????????????????????????????????????????????????????????????????????????????????????????????????????????????????????????????????????????????????????????????????????????????????????????????????????????????????????????????????????????????????????????????????????????????????????????????????????????????????????????????????????????????????????????????????????????????????????????????????????????????????????????????????????????????????????????????????????????????????????????????????????????????????????????????????????????????????????????????????????????????????????????????????????????????????????????????????????????????????????????????????????????????????????????????????????????????????????????????????????????????????????????????????????????????????????

			>Nidularium_camposportoi
			??????????????????????????????????????????????????????????????????????????????????????????????????????????????????????????????????????????????????????????????????????????????????????????????????????????????????????????????????????????????????????????????????????????????????????????????????????????????????????????????????????????????????????????????????????????????????????????????????????????????????????????????????????????????????????????????????????????????????????????????????????????????????????????????????????????????????????????????????????????????????????????????????????????????????????????????????????????????????????????????????????????????????????????????????????????????????????????????????????????????????????????????????????????????????????????????

			>Nidularium_fulgens
			??????????????????????????????????????????????????????????????????????????????????????????????????????????????????????????????????????????????????????????????????????????????????????????????????????????????????????????????????????????????????????????????????????????????????????????????????????????????????????????????????????????????????????????????????????????????????????????????????????????????????????????????????????????????????????????????????????????????????????????????????????????????????????????????????????????????????????????????????????????????????????????????????????????????????????????????????????????????????????????????????????????????????????????????????????????????????????????????????????????????????????????????????????????????????????????????

			>Nidularium_innocentii
			??????????????????????????????????????????????????????????????????????????????????????????????????????????????????????????????????????????????????????????????????????????????????????????????????????????????????????????????????????????????????????????????????????????????????????????????????????????????????????????????????????????????????????????????????????????????????????????????????????????????????????????????????????????????????????????????????????????????????????????????????????????????????????????????????????????????????????????????????????????????????????????????????????????????????????????????????????????????????????????????????????????????????????????????????????????????????????????????????????????????????????????????????????????????????????????????

			>Nidularium_krisgreeniae
			??????????????????????????????????????????????????????????????????????????????????????????????????????????????????????????????????????????????????????????????????????????????????????????????????????????????????????????????????????????????????????????????????????????????????????????????????????????????????????????????????????????????????????????????????????????????????????????????????????????????????????????????????????????????????????????????????????????????????????????????????????????????????????????????????????????????????????????????????????????????????????????????????????????????????????????????????????????????????????????????????????????????????????????????????????????????????????????????????????????????????????????????????????????????????????????????

			>Nidularium_marigoi
			??????????????????????????????????????????????????????????????????????????????????????????????????????????????????????????????????????????????????????????????????????????????????????????????????????????????????????????????????????????????????????????????????????????????????????????????????????????????????????????????????????????????????????????????????????????????????????????????????????????????????????????????????????????????????????????????????????????????????????????????????????????????????????????????????????????????????????????????????????????????????????????????????????????????????????????????????????????????????????????????????????????????????????????????????????????????????????????????????????????????????????????????????????????????????????????????

			>Nidularium_picinguabense
			??????????????????????????????????????????????????????????????????????????????????????????????????????????????????????????????????????????????????????????????????????????????????????????????????????????????????????????????????????????????????????????????????????????????????????????????????????????????????????????????????????????????????????????????????????????????????????????????????????????????????????????????????????????????????????????????????????????????????????????????????????????????????????????????????????????????????????????????????????????????????????????????????????????????????????????????????????????????????????????????????????????????????????????????????????????????????????????????????????????????????????????????????????????????????????????????

			>Nidularium_procerum
			ATAATTATAATAAAGTGAAATATGTCGAAATCCTTTTTGGAATAATACCAAATC-AAAATAAATGTCCGATAGCAAGTTGATCAGTTAATTCAATAAGAGATAAATGGGAGATAGCACTCGATTTAGTTGGTACCACCCAACCGAATCCGATTCACTCGTTTACTCATTGAATGAGTAAATTTTCAAGTTCAGCCAATCTTTTTTTTTTGAAAAAAAAAATTGCAAGTAGATGAAATCGTGAATAAATGTGTTTCATTTCTATATCATTATATAAAAAAAGATCCATATTATATTACTTATGGAATTCGAACCCGAACTCGATTTATGATTTATTATTTGGATCTTATTGGCCTTTGTTCTTTATTTTTGATTTTTCATATGGATTTCCGTCTTTATATTATACCCTTTCGTGGATGAATTATGCCTATTTTCACATCTAGGATTTACATATACAACATATATTACCCTCAAGAGGGAATTTTTCTCAGTATTTAGATATTTAGATTCAAAATAAGAAAGGGATCAATTCAATTATAAACCTGCAAAACAAAGATTAGGATTGGGTTGGGTTGCGCTATATATATCAAAGAGTATACAATAATGATGTATTTGGTGAATCAAATACATGGTCTAATAACAAACCATTTTAACATAACATTTTAATGAGTTGATAATATTAATTGAATATTGAATATTTTTTGAAAGATTTTTGTTAAAGGTTTCATTCATGCCTAATCCATATCGAGTAGACCTTGTTGTTGTG??

			>Nidularium_purpureum
			??????????????????????????????????????????????????????????????????????????????????????????????????????????????????????????????????????????????????????????????????????????????????????????????????????????????????????????????????????????????????????????????????????????????????????????????????????????????????????????????????????????????????????????????????????????????????????????????????????????????????????????????????????????????????????????????????????????????????????????????????????????????????????????????????????????????????????????????????????????????????????????????????????????????????????????????????????????????????????????????????????????????????????????????????????????????????????????????????????????????????????????????????????????????????????????????

			>Ochagavia_andina
			???????????????????????????????????TTTGGAATAATACCAAATCGAAAATAAATGTCCGATAGCAAGTTGATCAGTTAATTCAATAAGAGATAAATGGGAGATAGCACTCGATTTAGTTGGTACCACCCAACCGAATCCGATTCAATCGTTTACTCATTCAATGAGTAAATTTGCAAGTTCAGCCAATCTTTTTTTTTT--AAAAAAAAATTGCAAGTAGATGAAATCGTGAATAAATGTGTTTCATTTCTCTATCATTATATAAAATAAGATCCATATTATATTACTTATGGAATTCGAACCCGAACTCGATTTATGATTTATTATTTCGATCTTATTGGCCTTTGTTCTTTATTTTTGATTTTTCATATAGATTTCCGTCTTTATATTATACCCTCTCGTGGATGAATTATGCCTATTTTCACATCTAGGATTTACATATACAACATATATTACTGTCAAGAGGGAATTTTTCTCAGTATTTAGATATTTAGATTCAAAATAAGAAAGGGATCAATTCAATTATAAACCTGCAAAACAAAGATTAGGATTGGGTTGGGTTGCGCTATATATATCAAAGAGTATACAATAATGATGTATTTGGTGAATCAAATACATGGTCTAATAACAAACCATTTTAACATAACATTTTAATGAGTTGATAATATTAATTGAATATTGAATATTTTTTGAAAGATTTTTGTTAAAGGTTTCATTCATGCCTAATCCATATCGAGTAGACCTTGTTGTTGTG??

			>Ochagavia_carnea
			??????????????????????????????????????????????????????????????????????????????????CAGTTAATTCAATAAGAGATAAATGGGAGATAGCACTCGATTTAGTTGGTACCACCCAACCGAATCCGATTCAATCGTTTACTCATTCAATGAGTAAATTTGCAAGTTCAGCCAATCTTTTTTTTTTGAAAAAAAAAATTGCAAGTAGATGAAATCGTGAATAAATGTGTTTCATTTCTCTATCATTATATAAAATAAGATCCATATTATATTACTTATGGAATTCGAACCCGAACTCGATTTATGATTTATTATTTCGATCTTATTGGCCTTTGTTCTTTATTTTTGATTTTTCATATAGATTTCCGTCTTTATATTATACCCTCTCGTGGATGAATTATGCCTATTTTCACATCTAGGATTTACATATACAACATATATTACTGTCAAGAGGGAATTTTTCTCAGTATTTAGATATTTAGATTCAAAATAAGAAAGGGATCAATTCAATTATAAACCTGCAAAACAAAGATTAGGATTGGGTTGGGTTGCGCTATATATATCAAAGAGTATACAATAATGATGTATTTGGTGAATCAAATACATGGTCTAATAACAAACCATTTTAACATAACATTTTAATGAGTTGATAATATTAATTGAATATTGAATATTTTTTGAAAGATTTTTGTTAAAGGTTTCATTCATGCCTAATCCATATCGAGTAGACCTTGTTGTTGTGAG

			>Ochagavia_elegans
			ATAATTATAATAAAGTGAAATATGTCGAAATCCTTTTTGGAATAATACCAAATCGAAAATAAATGTCCGATAGCAAGTTGATCAGTTAATTCAATAAGAGATAAATGGGAGATAGCACTCGATTTAGTTGGTACCACCCAACCGAATCCGATTCAATCGTTTACTCATTCAATGAGTAAATTTGCAAGTTCAGCCAATCTTTTTTTTTTGAAATAAAAAATTGCAAGTAGATGAAATCGTGAATAAATGTGTTTCAGTTCTCTATCATTATATAAAATAAGATCCATATTATATTACTTATGGAATTCGAACCCGAACTCGATTTATGATTTATTATTTCGATCTTATTGGCCTTTGTTCTTTATTTTTTATTTTTCATATAGATTTCCGTCTTTATATTATACCCTCTCGTGGATGAATTATGCCTATTTTCACATCTAGGATTTACATATACAACATATATTACTGTCAAGAGGAAATTTTTCTCAGTATTTAGATATTTAGATTCAAAATAAGAAAGGGATCAATTCAATTATAAACCTGCAAAACAAAGATTAGGATTGGGTTGGGTTGCGCTATATATATCAAAGAGTATACAATAATGATGTATTTGGTGAATCAAATACATGGTCTAATAACAAACCATTTTAACATAACATTTTAATGAGTTGATAATATTAATTGAATATTGAATATTTTTTGAAAGATTTTTGTTAAAGGTTTCATTCATGCCTAATCCATATCGAGTAGACCTTGTTGTTGTG??

			>Ochagavia_litoralis
			ATAATTATAATAAAGTAAAATATGTCGAAATCCTTTTTGGAATAATACCAAATCGAAAATAAATGTCCGATAGCAAGTTGATCAGTTAATTCAATAAGAGATAAATGGGAGATAGCACTCGATTTAGTTGGTACCACCCAACCGAATCCGATTCAATCGTTTACTCATTCAATGAGTAAATTTGCAAGTTCAGCCAATCTTTTTTTTTTGAAAAAAAAAATTGCAAGTAGATGAAATCGTGAATAAATGTGTTTCATTTCTCTATCATTATATAAAATAAGATCCATATTATATTACTTATGGAATTCGAACCCGAACTCGATTTATGATTTATTATTTCGATCTTATTGGCCTTTGTTCTTTATTTTTGATTTTTCATATAGATTTCCGTCTTTATATTATACCCTCTCGTGGATGAATTATGCCTATTTTCACATCTAGGATTTACATATACAACATATATTACTGTCAAGAGGGAATTTTTCTCAGTATTTAGATATTTAGATTCAAAATAAGAAAGGGATCAATTCAATTATAAACCTGCAAAACAAAGATTAGGATTGGGTTGGGTTGCGCTATATATATCAAAGAGTATACAATAATGATGTATTTGGTGAATCAAATACATGGTCTAATAACAAACCATTTTAACATAACATTTTAATGAGTTGATAATATTAATTGAATATTGAATATTTTTTGAAAGATTTTTGTTAAAGGTTTCATTCATGCCTAATCCATATCGAGTAGACCTTGTTGTTGTG??

			>Orthophytum_alvimii
			??????????????????????????????????????????????????????????????????????????????????????????????????????????????????????????????????????????????????????????????????????????????????????????????????????????????????????????????????????????????????????????????????????????????????????????????????????????????????????????????????????????????????????????????????????????????????????????????????????????????????????????????????????????????????????????????????????????????????????????????????????????????????????????????????????????????????????????????????????????????????????????????????????????????????????????????????????????????????????????????????????????????????????????????????????????????????????????????????????????????????????????????????????????????????????????????

			>Orthophytum_argentum
			??????????????????????????????????????????????????????????????????????????????????????????????????????????????????????????????????????????????????????????????????????????????????????????????????????????????????????????????????????????????????????????????????????????????????????????????????????????????????????????????????????????????????????????????????????????????????????????????????????????????????????????????????????????????????????????????????????????????????????????????????????????????????????????????????????????????????????????????????????????????????????????????????????????????????????????????????????????????????????????????????????????????????????????????????????????????????????????????????????????????????????????????????????????????????????????????

			>Orthophytum_benzingii
			??????????????????????????????????????????????????????????????????????????????????????????????????????????????????????????????????????????????????????????????????????????????????????????????????????????????????????????????????????????????????????????????????????????????????????????????????????????????????????????????????????????????????????????????????????????????????????????????????????????????????????????????????????????????????????????????????????????????????????????????????????????????????????????????????????????????????????????????????????????????????????????????????????????????????????????????????????????????????????????????????????????????????????????????????????????????????????????????????????????????????????????????????????????????????????????????

			>Orthophytum_boudetianum
			??????????????????????????????????????????????????????????????????????????????????????????????????????????????????????????????????????????????????????????????????????????????????????????????????????????????????????????????????????????????????????????????????????????????????????????????????????????????????????????????????????????????????????????????????????????????????????????????????????????????????????????????????????????????????????????????????????????????????????????????????????????????????????????????????????????????????????????????????????????????????????????????????????????????????????????????????????????????????????????????????????????????????????????????????????????????????????????????????????????????????????????????????????????????????????????????

			>Orthophytum_braunii
			??????????????????????????????????????????????????????????????????????????????????????????????????????????????????????????????????????????????????????????????????????????????????????????????????????????????????????????????????????????????????????????????????????????????????????????????????????????????????????????????????????????????????????????????????????????????????????????????????????????????????????????????????????????????????????????????????????????????????????????????????????????????????????????????????????????????????????????????????????????????????????????????????????????????????????????????????????????????????????????????????????????????????????????????????????????????????????????????????????????????????????????????????????????????????????????????

			>Orthophytum_compactum
			?????????????????????????????????????TTGGATAATACCAAATCGAAAATAAATGTCCAATAGCAAGTTGATCAGTTAATTCAATAAGAGATAAATGGGAGATAGCACTCGATTTAGTTGGTACCACCCAACCGAATCCGATTCACTCGTTTACTCATTCAATGAGTAAATTTTCAAGTTCAGCCAATCTTTTTTTTTT-AAAAAAAAAATTGCAAGTAGATGAAATCGTGAATAAATGTGTTTCATTTCTATATCATTATATAAAAAAAGATCCATATTATATTACTTATGGAATTCGAACCCGAACTCGATTTATGATTTATTATTTGGATCTTATTGGCCTTTGTTCTTTATTTTTGATTTTTAATATAGATTTCCGTCTTTATATTATACCCTTTCGGGGATGAATTATGCCTATTTTCACATCTAGGATTTACATATACAACATATATTACTGTCAAGAGGGAATTTTTCTCAGTATTTAGATATTTAGATTCAAAATAAGAAAGGGATCAATTCAATTATAAACCTGCAAAACAAAGATTAGGATTGGGTTGGGTTGCGCTATATATATCAAAGAGTATACAATAATGATGTATTTGGTGAATCAAATACATGGTCTAATAACAAACCATTTTAACATAACATTTTAATGAGTTGATAATATTAATTGAATATTGAATATTTTTTGAAAGATTTTTGTTAAAGGTTTCATTCATGCCTAATCCATATCGAGTAGACCTTGTTGTTGTG??

			>Orthophytum_conquistense
			??????????????????????????????????????????????????????????????????????????????????????????????????????????????????????????????????????????????????????????????????????????????????????????????????????????????????????????????????????????????????????????????????????????????????????????????????????????????????????????????????????????????????????????????????????????????????????????????????????????????????????????????????????????????????????????????????????????????????????????????????????????????????????????????????????????????????????????????????????????????????????????????????????????????????????????????????????????????????????????????????????????????????????????????????????????????????????????????????????????????????????????????????????????????????????????????

			>Orthophytum_diamantinense
			??????????????????????TGTCGAA--TCCTTTTTGGATAATACCAAATCGAAAATAAATGTCCGATAGCAAGTTGATCAGTTAATTCAATAAGAGATAAATGGGAGATAGCACTCGATTTAGTTGGTACCACCCAACCGAATCCGATTCACTCGTTTACTCATTCAATGAGTAAATTTTCAAGTTCAGCCAATCTTTTTTTTTTTAAAAAAAAGATTGCAAGTAGATGAAATCGTGAATAAATGTGTTTCATTTCTATATCATTATATAAAAAAAGATCCATATTATATTACTTATGGAATTCGAACCCGAACTCGATTTATGATTTATTATTTGGATCTTATTGGCCTTTGTTCTTTATTTTTTATTTTTCATATAGATTTCCGTCTTTATATTAT---------TGGATGAATTATGCCTATTTTCACATCTAGGATTTACATATACAACATATATTACTGTCAAGAGGGAATTTTTCTCAGTATTTAGATATTTAGATTCAAAATAAGAAAGGGATCAATTCAATTATAAACCTGCAAAACAAAGATTAGGATTGGGTTGGGTTGCGCTATATATATCAAAGAGTATACAATAATGATGTATTTGGTGAATCAAATACATGGTCTAATAACAAACCATTTTAACATAACATTTTAATGAGTTGATAATATTAATTGAATATTGAATATTTTTTGAAAGATTTTTGTTAAAGGTTTCATTCATGCCTAATCCATATCGAGTAGACCTTGTTGTTGTG??

			>Orthophytum_disjunctum
			ATAATTATAATAAAGTGAAATATGTCGAAATCCTTTTTGGAATAATACCAAATCGAAAATAAATGTCCGATAGCAAGTTGATCAGTTAATTCAATAAGAGATAAATGGGAGATAGCACTCGATTTAGTTGGTACCACCCAACCGAATCCGATTCACTCGTTTACTCATTGAATGAGTAAATTTTCAAGTTCAGCCAATCTTTTTTTTTTTAAAAAAAAAATTGCAAGTAGATGAAATCGTGAATAAATGTGTTTCATTTCTATATCATTATATAAAAAAAGATCCATATTATATTACTTATGGAATTCGAACCCGAACTCGATTTATGATTTATTATTTGGATCTTATTGGCCTTTGTTCTTTATTTTTTATTTTTCATATAGATTTCCGTCTTTATATTAT---------TGGATGAATTATGCCTATTTTCACATCTAGGATTTACATATACAACATATATTACTGTCAAGAGGGAATTTTTCTCAGTATTTAGATATTTAGATTCAAAATAAGAAAGGGATCAATTCAATTATAAACCTGCAAAACAAAGATTAGGATTGGGTTGGGTTGCGCTATATATATCAAAGAGTATACAATAATGATGTATTTGGTGAATCAAATACATGGTCTAATAACAAACCATTTTAACATAACATTTTAATGAGTTGATAATATTAATTGAATATTGAATATTTTTTGAAAGATTTTTGTTAAAGGTTTCATTCATGCCTAATCCATATCGAGTAGACCTTGTTGTTGTG??

			>Orthophytum_duartei
			??????????????????????????????????????????????????????????????????????????????????????????????????????????????????????????????????????????????????????????????????????????????????????????????????????????????????????????????????????????????????????????????????????????????????????????????????????????????????????????????????????????????????????????????????????????????????????????????????????????????????????????????????????????????????????????????????????????????????????????????????????????????????????????????????????????????????????????????????????????????????????????????????????????????????????????????????????????????????????????????????????????????????????????????????????????????????????????????????????????????????????????????????????????????????????????????

			>Orthophytum_estevesii
			??????????????????????????????????????????????????????????????????????????????????????????????????????????????????????????????????????????????????????????????????????????????????????????????????????????????????????????????????????????????????????????????????????????????????????????????????????????????????????????????????????????????????????????????????????????????????????????????????????????????????????????????????????????????????????????????????????????????????????????????????????????????????????????????????????????????????????????????????????????????????????????????????????????????????????????????????????????????????????????????????????????????????????????????????????????????????????????????????????????????????????????????????????????????????????????????

			>Orthophytum_falconii
			??????????????????????????????????????????????????????????????????????????????????????????????????????????????????????????????????????????????????????????????????????????????????????????????????????????????????????????????????????????????????????????????????????????????????????????????????????????????????????????????????????????????????????????????????????????????????????????????????????????????????????????????????????????????????????????????????????????????????????????????????????????????????????????????????????????????????????????????????????????????????????????????????????????????????????????????????????????????????????????????????????????????????????????????????????????????????????????????????????????????????????????????????????????????????????????????

			>Orthophytum_foliosum
			??????????????????????????????????????????????????????????????????????????????????????????????????????????????????????????????????????????????????????????????????????????????????????????????????????????????????????????????????????????????????????????????????????????????????????????????????????????????????????????????????????????????????????????????????????????????????????????????????????????????????????????????????????????????????????????????????????????????????????????????????????????????????????????????????????????????????????????????????????????????????????????????????????????????????????????????????????????????????????????????????????????????????????????????????????????????????????????????????????????????????????????????????????????????????????????????

			>Orthophytum_fosterianum
			??????????????????????????????????????????????????????????????????????????????????????????????????????????????????????????????????????????????????????????????????????????????????????????????????????????????????????????????????????????????????????????????????????????????????????????????????????????????????????????????????????????????????????????????????????????????????????????????????????????????????????????????????????????????????????????????????????????????????????????????????????????????????????????????????????????????????????????????????????????????????????????????????????????????????????????????????????????????????????????????????????????????????????????????????????????????????????????????????????????????????????????????????????????????????????????????

			>Orthophytum_glabrum
			??????????????????????????????????????????????????????????????????????????????????????????????????????????????????????????????????????????????????????????????????????????????????????????????????????????????????????????????????????????????????????????????????????????????????????????????????????????????????????????????????????????????????????????????????????????????????????????????????????????????????????????????????????????????????????????????????????????????????????????????????????????????????????????????????????????????????????????????????????????????????????????????????????????????????????????????????????????????????????????????????????????????????????????????????????????????????????????????????????????????????????????????????????????????????????????????

			>Orthophytum_graomogolense
			??????????????????????????????????????????????????????????????????????????????????????????????????????????????????????????????????????????????????????????????????????????????????????????????????????????????????????????????????????????????????????????????????????????????????????????????????????????????????????????????????????????????????????????????????????????????????????????????????????????????????????????????????????????????????????????????????????????????????????????????????????????????????????????????????????????????????????????????????????????????????????????????????????????????????????????????????????????????????????????????????????????????????????????????????????????????????????????????????????????????????????????????????????????????????????????????

			>Orthophytum_grossiorum
			??????????????????????????????????????????????????????????????????????????????????????????????????????????????????????????????????????????????????????????????????????????????????????????????????????????????????????????????????????????????????????????????????????????????????????????????????????????????????????????????????????????????????????????????????????????????????????????????????????????????????????????????????????????????????????????????????????????????????????????????????????????????????????????????????????????????????????????????????????????????????????????????????????????????????????????????????????????????????????????????????????????????????????????????????????????????????????????????????????????????????????????????????????????????????????????????

			>Orthophytum_gurkenii
			??????????????????????????????????????????????????????????????????????????????????????????????????????????????????????????????????????????????????????????????????????????????????????????????????????????????????????????????????????????????????????????????????????????????????????????????????????????????????????????????????????????????????????????????????????????????????????????????????????????????????????????????????????????????????????????????????????????????????????????????????????????????????????????????????????????????????????????????????????????????????????????????????????????????????????????????????????????????????????????????????????????????????????????????????????????????????????????????????????????????????????????????????????????????????????????????

			>Orthophytum_harleyi
			???????????????????ATATGTCGA---TCCTTTTTGGATAATACCAAATCGAAAATAAATGTCCGAKAGCAAGTTGATCAGTTAATTCAATAAGAGATAAATGGGAGATAGCACTCGATTTAGTTGGTACCACCCAACCGAATCCGATTCACTCGTTTACTCATTCAATGAGTAAATTTTCAAGTTCAGCCAATCTTTTTTTTTTTAAAAAAAAAATTGCAAGTAGATGAAATCGTGAATAAATGTGTTTCATTTCTATATCATTATATAAAAAAAGATCCATATTATATTACTTATGGAATTCGAACCCGAACTCGATTTATGATTTATTATTTGGATCTTATTGGCCTTTGTTCTTTATTTTTTATTTTTCATATAGATTTCCGTCTTTATATTAT---------TGGATGAATTATGCCTATTTTCACATCTAGGATTTACATATACAACATATATTACYGTCAAGAGGGAATTTTTCTCAGTATTTAGATATTTAGATTCAAAATAAGAAAGGGATCAATTCAATTATAAACCCGCAAAACAAAGATTAGGATTGGGTTGGGTTGCGCTATATATATCAAAGAGTATACAATAATGATGTATTTGGTGAATCAAATACATGGTCTAATAACAAACCATTTTAACATAACATTTTAATGAGTTGATAATATTAATTGAATATTGAATATTTTTTGAAAGATTTTTGTTAAAGGTTTCATTCATGCATAATCCATATCGAGTAGACCTTGTTGTTGTG??

			>Orthophytum_horridum
			??????????????????????????????????????????????????????????????????????????????????????????????????????????????????????????????????????????????????????????????????????????????????????????????????????????????????????????????????????????????????????????????????????????????????????????????????????????????????????????????????????????????????????????????????????????????????????????????????????????????????????????????????????????????????????????????????????????????????????????????????????????????????????????????????????????????????????????????????????????????????????????????????????????????????????????????????????????????????????????????????????????????????????????????????????????????????????????????????????????????????????????????????????????????????????????????

			>Orthophytum_lanuginosum
			??????????????????????????????????????????????????????????????????????????????????????????????????????????????????????????????????????????????????????????????????????????????????????????????????????????????????????????????????????????????????????????????????????????????????????????????????????????????????????????????????????????????????????????????????????????????????????????????????????????????????????????????????????????????????????????????????????????????????????????????????????????????????????????????????????????????????????????????????????????????????????????????????????????????????????????????????????????????????????????????????????????????????????????????????????????????????????????????????????????????????????????????????????????????????????????????

			>Orthophytum_lemei
			??????????????????????????????????????????????????????????????????????????????????????????????????????????????????????????????????????????????????????????????????????????????????????????????????????????????????????????????????????????????????????????????????????????????????????????????????????????????????????????????????????????????????????????????????????????????????????????????????????????????????????????????????????????????????????????????????????????????????????????????????????????????????????????????????????????????????????????????????????????????????????????????????????????????????????????????????????????????????????????????????????????????????????????????????????????????????????????????????????????????????????????????????????????????????????????????

			>Orthophytum_leprosum
			??????????????????????????????????????????????????????????????????????????????????????????????????????????????????????????????????????????????????????????????????????????????????????????????????????????????????????????????????????????????????????????????????????????????????????????????????????????????????????????????????????????????????????????????????????????????????????????????????????????????????????????????????????????????????????????????????????????????????????????????????????????????????????????????????????????????????????????????????????????????????????????????????????????????????????????????????????????????????????????????????????????????????????????????????????????????????????????????????????????????????????????????????????????????????????????????

			>Orthophytum_lucidum
			??????????????????????????????????????????????????????????????????????????????????????????????????????????????????????????????????????????????????????????????????????????????????????????????????????????????????????????????????????????????????????????????????????????????????????????????????????????????????????????????????????????????????????????????????????????????????????????????????????????????????????????????????????????????????????????????????????????????????????????????????????????????????????????????????????????????????????????????????????????????????????????????????????????????????????????????????????????????????????????????????????????????????????????????????????????????????????????????????????????????????????????????????????????????????????????????

			>Orthophytum_macroflorum
			??????????????????????????????????????????????????????????????????????????????????????????????????????????????????????????????????????????????????????????????????????????????????????????????????????????????????????????????????????????????????????????????????????????????????????????????????????????????????????????????????????????????????????????????????????????????????????????????????????????????????????????????????????????????????????????????????????????????????????????????????????????????????????????????????????????????????????????????????????????????????????????????????????????????????????????????????????????????????????????????????????????????????????????????????????????????????????????????????????????????????????????????????????????????????????????????

			>Orthophytum_magalhaesii
			??????????????????????????????????????????????????????????????????????????????????????????????????????????????????????????????????????????????????????????????????????????????????????????????????????????????????????????????????????????????????????????????????????????????????????????????????????????????????????????????????????????????????????????????????????????????????????????????????????????????????????????????????????????????????????????????????????????????????????????????????????????????????????????????????????????????????????????????????????????????????????????????????????????????????????????????????????????????????????????????????????????????????????????????????????????????????????????????????????????????????????????????????????????????????????????????

			>Orthophytum_maracasense
			?????????ATAAAGTGAAATATGTCGAAATCCTTTTTGGAATAATACCAAATCGAAAATAAATGTCCGATAGCAAGTTGATCAGTTAATTCAATAAGAGATAAATGGGAGATAGCACTCGATTTAGTTGGTACCACCCAACCGAATCCGATTCACTCGTTTACTCATTCAATGAGTAAATTTTCAAGTTCAGCCAATC--TTTTTTTTGAAAAAAAAAATTGCAAGTAGATGAAATCGTGAATAAATGTGTTTCATTTCTATATCATTATATAAAAAAAGATCCATATTATATTACTTATGGAATTCGAACCCGAACTCGATTTATGATTTATTATTTGGATCTTATTGGCCTTTGTTCTTTATTTTTGATTTTTCATATGGATTTCCGTCTTTATATTATACCCTTTCGTGGATGAATTATGCCTATTTTCACATCTAGGATTTACATATACAACATATATTACTGTCAAGAGGGAATTTTTCTCAGTATTTAGATATTTAGATTCAAAATAAGAAAGGGATCAATTCAATTATAAACCTGCAAAACAAAGATTAGGATTGGGTTGGGTTGCGCTATATATATCAAAGAGTATACAATAATGATGTATTTGGTGAATCAAATACATGGTCTAATAACAAACCATTTTAACATAACATTTTAATGAGTTGATAATATTAATTGAATATTGAATATTTTTTGAAAGATTTTTGTTAAAGGTTTCATTCATGCCTAATCCATATCGAGTAGRCCTTGTTGTTGTG??

			>Orthophytum_mello-barretoi
			??????????????????????????????????????????????????????????????????????????????????????????????????????????????????????????????????????????????????????????????????????????????????????????????????????????????????????????????????????????????????????????????????????????????????????????????????????????????????????????????????????????????????????????????????????????????????????????????????????????????????????????????????????????????????????????????????????????????????????????????????????????????????????????????????????????????????????????????????????????????????????????????????????????????????????????????????????????????????????????????????????????????????????????????????????????????????????????????????????????????????????????????????????????????????????????????

			>Orthophytum_pseudostoloniferum
			??????????????????????????????????????????????????????????????????????????????????????????????????????????????????????????????????????????????????????????????????????????????????????????????????????????????????????????????????????????????????????????????????????????????????????????????????????????????????????????????????????????????????????????????????????????????????????????????????????????????????????????????????????????????????????????????????????????????????????????????????????????????????????????????????????????????????????????????????????????????????????????????????????????????????????????????????????????????????????????????????????????????????????????????????????????????????????????????????????????????????????????????????????????????????????????????

			>Orthophytum_pseudovagans
			??????????????????????????????????????????????????????????????????????????????????????????????????????????????????????????????????????????????????????????????????????????????????????????????????????????????????????????????????????????????????????????????????????????????????????????????????????????????????????????????????????????????????????????????????????????????????????????????????????????????????????????????????????????????????????????????????????????????????????????????????????????????????????????????????????????????????????????????????????????????????????????????????????????????????????????????????????????????????????????????????????????????????????????????????????????????????????????????????????????????????????????????????????????????????????????????

			>Orthophytum_riocontense
			??????????????????????????????????????????????????????????????????????????????????????????????????????????????????????????????????????????????????????????????????????????????????????????????????????????????????????????????????????????????????????????????????????????????????????????????????????????????????????????????????????????????????????????????????????????????????????????????????????????????????????????????????????????????????????????????????????????????????????????????????????????????????????????????????????????????????????????????????????????????????????????????????????????????????????????????????????????????????????????????????????????????????????????????????????????????????????????????????????????????????????????????????????????????????????????????

			>Orthophytum_roseolilacinum
			??????????????????????????????????????????????????????????????????????????????????????????????????????????????????????????????????????????????????????????????????????????????????????????????????????????????????????????????????????????????????????????????????????????????????????????????????????????????????????????????????????????????????????????????????????????????????????????????????????????????????????????????????????????????????????????????????????????????????????????????????????????????????????????????????????????????????????????????????????????????????????????????????????????????????????????????????????????????????????????????????????????????????????????????????????????????????????????????????????????????????????????????????????????????????????????????

			>Orthophytum_rubiginosum
			??????????????????????????????????????????????????????????????????????????????????????????????????????????????????????????????????????????????????????????????????????????????????????????????????????????????????????????????????????????????????????????????????????????????????????????????????????????????????????????????????????????????????????????????????????????????????????????????????????????????????????????????????????????????????????????????????????????????????????????????????????????????????????????????????????????????????????????????????????????????????????????????????????????????????????????????????????????????????????????????????????????????????????????????????????????????????????????????????????????????????????????????????????????????????????????????

			>Orthophytum_santaritense
			??????????????????????????????????????????????????????????????????????????????????????????????????????????????????????????????????????????????????????????????????????????????????????????????????????????????????????????????????????????????????????????????????????????????????????????????????????????????????????????????????????????????????????????????????????????????????????????????????????????????????????????????????????????????????????????????????????????????????????????????????????????????????????????????????????????????????????????????????????????????????????????????????????????????????????????????????????????????????????????????????????????????????????????????????????????????????????????????????????????????????????????????????????????????????????????????

			>Orthophytum_saxicola
			???????????????????????????????????TTTGGAATAATACCAAATCGAAAATAAATGTCCGATAGCAAGTTGATCAGTTAATTCAATAAGAGATAAATGGGAGATAGCACTCGATTTAGTTGGTACCACCCAACCGAATCCGATTCACTCGTTTACTCATTCAATGAGTAAATTTTCAAGTTCAGCCAATCTTTTTTTTTT-AAAAAAAAAATTGCAAGTAGATGAAATCGTGAATAAATGTGTTTCATTTCTATATCATTATATAAAAAAAGATCCATATTATATTACTTATGGAATTCGAACCCGAACTCGATTTATGATTTATTATTTGGATCTTATTGGCCTTTGTTCTTTATTTTTTATTTTTCATATAGATTTCCGTCTTTATATTAT---------TGGATGAATTATGCCTATTTTCACATCTAGGATTTACATATACAACATATATTACTGTCAAGAGGGAATTTTTCTCAGTATTTAGATATTTAGATTCAAAATAAGAAAGGGATCAATTCAATTATAAACCCGCAAAACAAAGATTAGGATTGGGTTGGGTTGCGCTATATATATCAAAGAGTATACAATAATGATGTATTTGGTGAATCAAATACATGGTCTAATAACAAACCATTTTAACATAACATTTTAATGAGTTGATAATATTAATTGAATATTGAATATTTTTTGAAAGATTTTTGTTAAAGGTTTCATTCATGCATAATCCATATCGAGTAGACCTTGTTGTTGTG??

			>Orthophytum_schulzianum
			??????????????????????????????????????????????????????????????????????????????????????????????????????????????????????????????????????????????????????????????????????????????????????????????????????????????????????????????????????????????????????????????????????????????????????????????????????????????????????????????????????????????????????????????????????????????????????????????????????????????????????????????????????????????????????????????????????????????????????????????????????????????????????????????????????????????????????????????????????????????????????????????????????????????????????????????????????????????????????????????????????????????????????????????????????????????????????????????????????????????????????????????????????????????????????????????

			>Orthophytum_sucrei
			??????????????????????????????????????????????????????????????????????????????????????????????????????????????????????????????????????????????????????????????????????????????????????????????????????????????????????????????????????????????????????????????????????????????????????????????????????????????????????????????????????????????????????????????????????????????????????????????????????????????????????????????????????????????????????????????????????????????????????????????????????????????????????????????????????????????????????????????????????????????????????????????????????????????????????????????????????????????????????????????????????????????????????????????????????????????????????????????????????????????????????????????????????????????????????????????

			>Orthophytum_toscanoi
			??????????????????????????????????????????????????????????????????????????????????????????????????????????????????????????????????????????????????????????????????????????????????????????????????????????????????????????????????????????????????????????????????????????????????????????????????????????????????????????????????????????????????????????????????????????????????????????????????????????????????????????????????????????????????????????????????????????????????????????????????????????????????????????????????????????????????????????????????????????????????????????????????????????????????????????????????????????????????????????????????????????????????????????????????????????????????????????????????????????????????????????????????????????????????????????????

			>Orthophytum_vagans
			???????????????????????????????????TTTGGAATAATACCAAATCGAAAATAAATGTCCGATAGCAAGTTGATCAGTTAATTCAATAAGAGATAAATGGGAGATAGCACTCTATTTAGTTGGTAGCACCCAACCGAATCCGATTCACTCGTTTACTCATTGAATGAGTAAATTTTCAAGT------------------------------------------------CGTGAATAAATGTGTTTCATTTCTATATCATTATATAAAAAAAGATCCATATTATATTACTTATGGAATTCGAACCCGAACTCGATTTATGATTTATTATTTGGATCTTATTGGCCTTTGTTCTTTCTTTTTGATTTTTCATATAGATTTCCGTCTTTATATTAT---------TGGATGAATTATGCCTATTTTCACATCTAGGATTTACATATACAACATATATTACTGTCAAGAGGGAATTTTTCTCAGTATTTAGATATTTAGATTCAAAATAAGAAAGGGATCAATTCAATTATAAACCTGCAAAACAAAGATTAGGATTGGGTTGGGTTGCGCTATATATATCAAAGAGTATACAATAATGATGTATTTGGTGAATCAAATACATGGTCTAATAACAAACCATTTTAACATAACATTTTAATGAGTTGATAATATTAATTGAATATTGAATATTTTTTGAAAGATTTTTGTTAAAGGTTTCATTCATGCCTAATCCATATCGAGTAGACCTTGTTGTTGTG??

			>Orthophytum_vasconcelosianum
			??????????????????????????????????????????????????????????????????????????????????????????????????????????????????????????????????????????????????????????????????????????????????????????????????????????????????????????????????????????????????????????????????????????????????????????????????????????????????????????????????????????????????????????????????????????????????????????????????????????????????????????????????????????????????????????????????????????????????????????????????????????????????????????????????????????????????????????????????????????????????????????????????????????????????????????????????????????????????????????????????????????????????????????????????????????????????????????????????????????????????????????????????????????????????????????????

			>Orthophytum_zanonii
			??????????????????????????????????????????????????????????????????????????????????????????????????????????????????????????????????????????????????????????????????????????????????????????????????????????????????????????????????????????????????????????????????????????????????????????????????????????????????????????????????????????????????????????????????????????????????????????????????????????????????????????????????????????????????????????????????????????????????????????????????????????????????????????????????????????????????????????????????????????????????????????????????????????????????????????????????????????????????????????????????????????????????????????????????????????????????????????????????????????????????????????????????????????????????????????????

			>Pitcairnia_abundans
			??????????????????????????????????????????????????????????????????????????????????????????????????????????????????????????????????????????????????????????????????????????????????????????????????????????????????????????????????????????????????????????????????????????????????????????????????????????????????????????????????????????????????????????????????????????????????????????????????????????????????????????????????????????????????????????????????????????????????????????????????????????????????????????????????????????????????????????????????????????????????????????????????????????????????????????????????????????????????????????????????????????????????????????????????????????????????????????????????????????????????????????????????????????????????????????????

			>Pitcairnia_albiflos
			??????????????????????????????????????????????????????????????????????????????????CAGTTAATTTAATAAGAGATAAATGGGAGATAGCACTCGATTTAGTTGGTACCACCCAACCGAATCCGATTCAATCGTTTACTCATTCAATGAGTAAATTTTAAAGTTCAGCCAATT--TTTTTTTTC-AAAAAAAAATTGCAAGTAGATAAAATCGTGAATAAATGTGTTTCATTTCTCTATCATTATAGAAAATAAGATCC-----ATATTACTTATGGAATTCGAACCCGAACTCGATTTATGATTCATTATTTCGATCTTATTGGCCTTTGTTCTTTATTTTTGATTTTTCATATAGATTTCCGTCTTTATATTATACCCTTTCGTGGATGAATTATGCCTATTTTCACATCTAGGATTTACATATACAACATATATTACTGTCAAGAG-----TTTTCTCAGTATTTAGATATTTAGATTCCAAATAAGAAAGGGATCAATTCAATTATAAACCTGCAAAACAAAGATTAGGA-----TTGGGTTGCGCTATATATATCAAAGAGTATACAATAATGATGTATTTGGTGAATCAAATACATGGTCTAATAACAAACCATTTTAACATAACATTTTAATGAGTTGATAATATTAATTGAATATTGAATATTTTTTGAAAGATTTTTGTTAAAGGTTTCATTCATGCCTAATCCATATCGAGTAGACCTTGTTGTTGTGAG

			>Pitcairnia_atrorubens
			??????????????????????????????????????????????????????????????????????????????????CAGTTAATTCAATAAGAGATAAATGGGAGATAGCACTCGATTTAGTTGGTACCACCCAACCGAATCCGATTCAATCGTTTACTCATTCAATGAGTCAATTTTCAAGTTCAGCCAATC-TTTTTTTTTGAAAAAAAAAATTGCAAGTAGATGAAATCGTGAATAAAAGTGTTTCATTTCTCTATCATTATAGAAAATAAGATCCATATTATATTACTTATGGAATTCGAACCCGAACTCGATTTATGATTCATTATTTCGATCTTATCGGCCTTTGTTCTTTATTTTTGATTTTTCATATAGATTTCCGTCTTTATATTATACCCTTTCGTGGATGAATTATGCCTATTTTCATATCTAGGATTTACATATACAACATATATTACTGTCAAGAGGGAATTTTTCTCAGTATTTAGATATTTAGATTCAAAATAAGAAAGGGATCAATTCAATTATAAACCTGCAAAACAAAGATTAGGATTGGGTTGGGTTGCGCTATATATATCAAAGAGTATACAATAATGATGTATTTGGTGAATCAAATACATGGTCTAATAACAAACCATTTTAACATAACATTTTAATGAGTTGATAATATTAATTGAATATTGAATATTTTTTGAAAGATTTTTGTTAAAGGTTTCATTCATGCCTAATCCATATCGAGTAGACCTTGTTGTTGTGAG

			>Pitcairnia_beachiae
			??????????????????????????????????????????????????????????????????????????????????????????????????????????????????????????????????????????????????????????????????????????????????????????????????????????????????????????????????????????????????????????????????????????????????????????????????????????????????????????????????????????????????????????????????????????????????????????????????????????????????????????????????????????????????????????????????????????????????????????????????????????????????????????????????????????????????????????????????????????????????????????????????????????????????????????????????????????????????????????????????????????????????????????????????????????????????????????????????????????????????????????????????????????????????????????????

			>Pitcairnia_breedlovei
			??????????????????????????????????????????????????????????????????????????????????CAGTTAATTCAATAAGAGATAAATGGGAGATAGCACTCGATTTAGTTGGTACCACCCAACCGAATCCGATTCAATCGTTTACTCATTCAATGAGTCAATTTTCAAGTTCAGCCAATC-TTTTTTTTTTGAAAAAAAAATTGCAAGTAGATGAAATCGTGAATAAATGTGTTTCATTTCTCTATCATTATAGAAAATAAGATCCATATTATATTACTTATGGAATTCGAACCCGAACTCGATTTATGATTCATTATTTCGATCTTATTGGCCTTTGTTCTTTATTTTTGATTTTTCATATAGATTTCCGTCTTTATATTATACCCTTTCGTGGATGAATTATGCCTATTTTCATATCTAGGATTTACATATACAACATATATGACTGTCAAGAGGGAATTTTTCTCAGTATTTAGATATTTAGATTCAAAATAAGAAAGGGATCAATTCAATTATAAACCTGCAAAACAAAGATTAGGATTGGGTTGGGTTGCGCTATATATATCAAAGAGTATACAATAATGATGTATTTGGTGAATCAAATACATGGTCTAATAACAAACCATTTTAACATAACATTTTAATGAGTTGATAATATTAATTGAATATTGAATATTTTTTGAAAGATTTTTGTTAAAGGTTTCATTCATGCCTAATCCATATCGAGTAGACCTTGTTGTTGTGAG

			>Pitcairnia_burle-marxii
			??????????????????????????????????????????????????????????????????????????????????????????????????????????????????????????????????????????????????????????????????????????????????????????????????????????????????????????????????????????????????????????????????????????????????????????????????????????????????????????????????????????????????????????????????????????????????????????????????????????????????????????????????????????????????????????????????????????????????????????????????????????????????????????????????????????????????????????????????????????????????????????????????????????????????????????????????????????????????????????????????????????????????????????????????????????????????????????????????????????????????????????????????????????????????????????????

			>Pitcairnia_carinata
			ATAATTATAATAAAGTTAAATATGTCGAAATCCTTTTTGGAATAATACCAAATCGAAAATAAATGTCCGATAGCAAGTTGATCAGTTAATTTAATAAGGGATAAATGGGAGATAGCACTCGATTTAGTTGGTACCACCCAACCGAATCCGATTCAATCGTTTACTCATTCAATGAGTAAATTTTAAAGTTCAGCCAATT--TTTTTTTTG-AAAAAAAAATTGCAAGTAGATAAAATCGTGAATAAATGTGTTTCATTTCTCTATCATTATAGAAAATAAGATCC-----ATATTACTTATGGAATTCGAACCCGAACTCGATTTATGATTCATTATTTCGATCTTATTGGCCTTTGTTCTTTATTTTTGATTTTTCATATAGATTTCCGTCTTTATATTATACCCTTTCGTGGATGAATTATGCCTATTTTCACATCTAGGATTTACATATACAACATATATTACTGTCAAGAG-----TTTTCTCAGTATTTAGATATTTAGATTCCAAATAAGAAAGGGATCAATTCAATTATAAACCTGCAAAACAAAGATTAGGA-----TTGGGTTGCGCTATATATATCAAAGAGTATACAATAATGATGTATTTGGTGAATCAAATACATGGTCTAATAACAAACCATTTTAACATAACATTTTAATGAGTTGATAATATTAATTGAATATTGAATATTTTTTGAAAGATTTTTGTTAAAGGTTTCATTCATGCCTAATCCATATCGAGTAGACCTTGTTGTTGTGAG

			>Pitcairnia_chiapensis
			??????????????????????????????????????????????????????????????????????????????????CAGTTAATTCAATAAGAGATAAATGGGAGATAGCACTCGATTTAGTTGGTACCACCCAACCGAATCCGATTCAATCGTTTACTCATTCAATGAGTCAATTTTCAAGTTCAGCCAATCTTTTTTTTTGAAAAAAAAAAATTGCAAGTAGATGAAATCGTGAATAAATGTGTTTCATTTCTCTATCATTATAGAAAATAAGATCCATATTATATTACTTATGGAATTCGAACCCGAACTCGATTTATGATTCATTATTTCGATCTTATTGGCCTTTGTTCTTTATTTTTGATTTTTCATATAGATTTCCGTCTTTATATTATACCCTTTCGTGGATGAATTATGCCTATTTTCATATCTAGGATTTACATATACAACATATATTACTGTCAAGAGGGAATTTTTCTCAGTATTTAGATATTTAGATTCAAAATAAGAAAGGGATCAATTCAATTATAAACCTGCAAAACAAAGATTAGGA-----TTGGGTTGCGCTATATATATCAAAGAGTATACAATAATGATGTATTTGGTGAATCAAATACATGGTCTAATAACAAACCATTTTAACATAACATTTTAATGAGTTGATAATATTAATTGAATATTGAATATTTTTTGAAAGATTTTTGTTAAAGGTTTCATTCATGCCTAATCCATATCGAGTAGACCTTGTTGTTGTGAG

			>Pitcairnia_corallina
			ATAATTATAATAAAGTTAAATATGTCGAAATCCTTTTTGGAATAATACCAAATCGAAAATAAATGTCCGATAGCAAGTTGATCAGTTAATTCAATAAGAGATAAATGGGAGATAGCACTCGATTTAGTTGGTACCACCCAACCGAATCCGATTCAATCGTTTACTCATTCAATGAGTCAATTTTCAAGTTCAGCCAATC--TTTTTTTTTGAAAAAAAAATTGCAAGTAGATGAAATCGTGAATAAATGTGTTTCATTTCTCTATCATTATAGAAAATAAGATCCATATTATATTACTTATGGAATTCGAACCCGAACTCGATTTATGATTCATTATTTCGATCTTATTGGCCTTTGTTCTTTATTTTTGATTTTTCATATAGATTTCCGTCTTTATATTATAACCTTTCGTGGATGAATTATGCCTATTTTCATATCTAGGATTTACATATACAACATATATGACTGTCAAGAGGGAATTTTTCTCAGTATTTAGATATTTAGATTCAAAATAAGAAAGGGATCAATTCAATTATAAACCTGCAAAACAAAGATTAGGATTGGGTTGGGTTGCGCTATATATATCAAAGAGTATACAATAATGATGTATTTGGTGAATCAAATACATGGTCTAATAACAAACCATTTTAACATAACATTTTAATGAGTTGATAATATTAATTGAATATTGAATATTTTTTGAAAGATTTTTGTTAAAGGTTTCATTCATGCCTAATCCATATCGAGTAGACCTTGTTGTTGTGAG

			>Pitcairnia_elvirae
			??????????????????????????????????????????????????????????????????????????????????????????????????????????????????????????????????????????????????????????????????????????????????????????????????????????????????????????????????????????????????????????????????????????????????????????????????????????????????????????????????????????????????????????????????????????????????????????????????????????????????????????????????????????????????????????????????????????????????????????????????????????????????????????????????????????????????????????????????????????????????????????????????????????????????????????????????????????????????????????????????????????????????????????????????????????????????????????????????????????????????????????????????????????????????????????????

			>Pitcairnia_feliciana
			ATAATTATAATAAAGTTAAATATGTCGAAATCCTTTTTGGAATAATACCAAATCGAAAATAAATGTCCGATAGCAAGTTGATCAGTTAATTCAATAAGAGATAAATGGGAGATAGCACTCGATTTAGTTGGTACCACCCAACCGAATCCGATTCAATCGTTTACTCATTCAATGAGTCAATTTTCAAGTTCAGCCAATC--TTTTTTTTGAAAAAAAAAATTGCAAGTAGATGAAATCGTGAATAAATGTGTTTCATTTCTCTATCATTATAGAAAATAAGATCCATATTATATTACTTATGGAATTCGAACCCGAACTCGATTTATGATTCATTATTTCGATCTTATTGGCCTTTGTTCTTTATTTTTGATTTTTCATATAGATTTCCGTCTTTATATTATA-CCTTTCGTGGATGAATTATGCCTATTTTCATATCTGGGATTTACATATACAACATATATTACTGTCAAGAGGGAATTTTTCTCAGTATTTAGATATTTAGATTCAAAATAAGAAAGGGATCAATTCAATTAGAAACCTGCAAAACAAAGATTAGGA-----TTGGGTTGCGCTATATATATCAAAGAGTATACAATAATGATGTATTTGGTGAATCAAATACATGGTCTAATAACAAACCATTTTAACATAACATTTTAATGAGTTGATAATATTAATTGAATATTGAATATTTTTTGAAAGATTTTTGTTAAAGGTTTCATTCATGCCTAATCCATATCGAGTAGACCTTGTTGTTGTG??

			>Pitcairnia_flammea
			??????????????????????????????????????????????????????????????????????????????????????????????????????????????????????????????????????????????????????????????????????????????????????????????????????????????????????????????????????????????????????????????????????????????????????????????????????????????????????????????????????????????????????????????????????????????????????????????????????????????????????????????????????????????????????????????????????????????????????????????????????????????????????????????????????????????????????????????????????????????????????????????????????????????????????????????????????????????????????????????????????????????????????????????????????????????????????????????????????????????????????????????????????????????????????????????

			>Pitcairnia_flammea_var._glabrior
			??????????????????????????????????????????????????????????????????????????????????????????????????????????????????????????????????????????????????????????????????????????????????????????????????????????????????????????????????????????????????????????????????????????????????????????????????????????????????????????????????????????????????????????????????????????????????????????????????????????????????????????????????????????????????????????????????????????????????????????????????????????????????????????????????????????????????????????????????????????????????????????????????????????????????????????????????????????????????????????????????????????????????????????????????????????????????????????????????????????????????????????????????????????????????????????????

			>Pitcairnia_fuertesii
			??????????????????????????????????????????????????????????????????????????????????????????????????????????????????????????????????????????????????????????????????????????????????????????????????????????????????????????????????????????????????????????????????????????????????????????????????????????????????????????????????????????????????????????????????????????????????????????????????????????????????????????????????????????????????????????????????????????????????????????????????????????????????????????????????????????????????????????????????????????????????????????????????????????????????????????????????????????????????????????????????????????????????????????????????????????????????????????????????????????????????????????????????????????????????????????????

			>Pitcairnia_heterophylla
			ATAATTATAATAAAGTTAAATATGTCGAAATCCTTTTTGGAATAATACCAAATCGAAAATAAATGTCCGATAGCAAGTTGATCAGTTAATTTAATAAGAGATAAATGGGAGATAGCACTCGATTTAGTTGGTACCACCCAACCGAATCCGATTCAATCGTTTACTCATTCAATGAGTAAATTTAAAAGTTCAGCCAATT--TTTTTTTTGAAAAAAAAAATTGCAAGTAGATAAAATCGTGAATAAATGTGTTTTATTTCTCTATCATTATAGAAAAAAAGATCC-----ATATTACTTATGGAATTCGAACCCGAACTCGATTTATGATTCATTATTTCGATCTTATTGGCCTGTGTTCTTTATTTTTGATTTTTCATATAGATTTCCGTCTTTATATTATACCCTTTCGTGGATGAATTATGCCTATTTTCACATCTAGGATTTACATATACAACATATATTACTGTCAAGAG-----TTTTCTCAGTATTTAGATATTTAGATTCCAAATAAGAAAGGGATCAATTCAATTATAAACCTGCAAAACAAAGATTAGGA-----TTGGGTTGCGCTATATATATCAAAGAGTATACAATAATGATGTATTTGGTGAATCAAATACATGGTCTAATAACAAACCATTTTAACATAACATTTTAATGAGTTGATAATATTAATTGAATATTGAATATTTTTTGAAAGATTTTTGTTAAAGGTTTCATTCATGCCTAATCCATATCGAGTAGACCTTGTTGTTGTGAG

			>Pitcairnia_hirtzii
			ATAATTATAATAAAGTTAAATATGTCGAAATCCTTTTTGGAATAATACCAAATCGAAAATAAATGTCCGATAGCAAGTTGATCAGTTAATTCAATAAGAGATAAATGGGAGATAGCACTCGATTTAGTTGGTACCACCCAACCGAATCCGATTCAATCGTTTACTCATTCAATGAGTCAATTTTCAAGTTCAGCCAATC-TTTTTTTTTTTCAAAAAAAATTGCAAGTAGATGAAATCGTGAATAAATGTGTTTCATTTCTCTATCATTATAGAAAATAAGATCCATATTATATTACTTATGGAATTCGAACCCGAACTCGATTTATGATTCATTATTTCGATCTTATTGGCCTTTGTTCTTTATTTTTGATTTTTCATATAGATTTCCGTCTTTATATTATACCCTTTCGTGGATGAATTATGCCTATTTTCATATCTAGGATTTACATATACAACATATATGACTGTCAAGAGGGAATTTTTCTCAGTATTTAGATATTTAGATTCAAAATAAGAAAGGGATCAATTCAATTATAAACCTGCAAAACAAAGATTAGGATTGGGTTGGGTTGCGCTATATATATCAAAGAGTATACAATAATGATGTATTTGGTGAATCAAATACATGGTCTAATAACAAACCATTTTAACATAACATTTTAATGAGTTGATAATATTAATTGAATATTGAATATTTTTTGAAAGATTTTTGTTAAAGGTTTCATTCATGCCTAATCCATATCGAGTAGACCTTGTTGTTGTGAG

			>Pitcairnia_integrifolia
			??????????????????????????????????????????????????????????????????????????????????????????????????????????????????????????????????????????????????????????????????????????????????????????????????????????????????????????????????????????????????????????????????????????????????????????????????????????????????????????????????????????????????????????????????????????????????????????????????????????????????????????????????????????????????????????????????????????????????????????????????????????????????????????????????????????????????????????????????????????????????????????????????????????????????????????????????????????????????????????????????????????????????????????????????????????????????????????????????????????????????????????????????????????????????????????????

			>Pitcairnia_jimenezii
			??????????????????????????????????????????????????????????????????????????????????????????????????????????????????????????????????????????????????????????????????????????????????????????????????????????????????????????????????????????????????????????????????????????????????????????????????????????????????????????????????????????????????????????????????????????????????????????????????????????????????????????????????????????????????????????????????????????????????????????????????????????????????????????????????????????????????????????????????????????????????????????????????????????????????????????????????????????????????????????????????????????????????????????????????????????????????????????????????????????????????????????????????????????????????????????????

			>Pitcairnia_lignosa
			??????????????????????????????????????????????????????????????????????????????????????????????????????????????????????????????????????????????????????????????????????????????????????????????????????????????????????????????????????????????????????????????????????????????????????????????????????????????????????????????????????????????????????????????????????????????????????????????????????????????????????????????????????????????????????????????????????????????????????????????????????????????????????????????????????????????????????????????????????????????????????????????????????????????????????????????????????????????????????????????????????????????????????????????????????????????????????????????????????????????????????????????????????????????????????????????

			>Pitcairnia_loki-schmidtiae
			??????????????????????????????????????????????????????????????????????????????????CAGTTAATTTAATAAGAGATAAATGGGAGATAGCACTCGATTTAGTTGGTACCACCCAACCGAATCCGATTCAATCGTTTACTCATTCAATGAGTAAATTTTAAAGTTCAGCCAATT--TTTTTTTTCAAAAAAAAAATTGCAAGTAGATAAAATCGTGAATAAATGTGTTTCATTTCTCTATCATTATAGAAAATAAGATCC-----ATATTACTTATGGAATTCGAACCCGAACTCGATTTATGATTCATTATTTCGATCTTATTGGCCTTTGTTCTTTATTTTTGATTTTTCATATAGATTTCCGTCTTTATATTATACCCTTTCGTGGATGAATTATGCCTATTTTCACATCTAGGATTTACATATACAACATATATTACTGTCAAGAG-----TTTTCTCAGTATTTAGATATTTAGATTCCAAATAAGAAAGGGATCAATTCAATTATAAACCTGCAAAACAAAGATTAGGA-----TTGGGTTGCGCTATATATATCAAAGAGTATACAATAATGATGTATTTGGTGAATCAAATACATGGTCTAATAACAAACCATTTTAACATAACATTTTAATGAGTTGATAATATTAATTGAATATTGAATATTTTTTGAAAGATTTTTGTTAAAGGTTTCATTCATGCCTAATCCATATCGAGTAGACCTTGTTGTTGTGAG

			>Pitcairnia_macrochlamys
			??????????????????????????????????????????????????????????????????????????????????????????????????????????????????????????????????????????????????????????????????????????????????????????????????????????????????????????????????????????????????????????????????????????????????????????????????????????????????????????????????????????????????????????????????????????????????????????????????????????????????????????????????????????????????????????????????????????????????????????????????????????????????????????????????????????????????????????????????????????????????????????????????????????????????????????????????????????????????????????????????????????????????????????????????????????????????????????????????????????????????????????????????????????????????????????????

			>Pitcairnia_maidifolia
			??????????????????????????????????????????????????????????????????????????????????????????????????????????????????????????????????????????????????????????????????????????????????????????????????????????????????????????????????????????????????????????????????????????????????????????????????????????????????????????????????????????????????????????????????????????????????????????????????????????????????????????????????????????????????????????????????????????????????????????????????????????????????????????????????????????????????????????????????????????????????????????????????????????????????????????????????????????????????????????????????????????????????????????????????????????????????????????????????????????????????????????????????????????????????????????????

			>Pitcairnia_nigra
			??????????????????????????????????????????????????????????????????????????????????????????????????????????????????????????????????????????????????????????????????????????????????????????????????????????????????????????????????????????????????????????????????????????????????????????????????????????????????????????????????????????????????????????????????????????????????????????????????????????????????????????????????????????????????????????????????????????????????????????????????????????????????????????????????????????????????????????????????????????????????????????????????????????????????????????????????????????????????????????????????????????????????????????????????????????????????????????????????????????????????????????????????????????????????????????????

			>Pitcairnia_orchidifolia
			ATAATTATAATAAAGTTAAATATGTCGAAATCCTTTTTGGAATAATACCAAATCGAAAATAAATGTCCGATAGCAAGTTGATCAGTTAATTCAATAAGAGATAAATGGGAGATAGCACTCGATTTAGTTGGTACCACCCAACCGAATCCGATTCAATCGTTTACTCATTCAATGAGTCAATTTTCAAGTTCAGCCAATCTTTTTTTTTTGAAAAAAAAAATTGCAAGTAGATGAAATCGTGAATAAATGTGTTTCATTTCTCTATCATTATAGAAAATAAGATCCATATTATATTACTTATGGAATTCGAACCCGAACTCGATTTATGATTCATTATTTCGATCTTATTGGCCTTTGTTCTTTATTTTTGATTTTTCATATAGATTTCCGTCTTTATATTATACCCTTTCGTGGATGAATTATGCCTATTTTCATATCTAGGATTTACATATACAACATATATTACTGTCAAGAGGGAATTTTTCTCAGTATTTAGATATTTAGATTCAAAATAAGAAAGGGATCAATTCAATTATAAACCTGCAAAACAAAGATTAGGA-----TTGGGTTGCGCTATATATATCAAAGAGTATACAATAATGATGTATTTGGTGAATCAAATACATGGTCTAATAACAAACCATTTTAACATAACATTTTAATGAGTTGATAATATTAATTGAATATTGAATATTTTTTGAAAGATTTTTGTTAAAGGTTTCATTCATGCCTAATCCATATCGAGTAGACCTTGTTGTTGTGAG

			>Pitcairnia_piepenbringii
			??????????????????????????????????????????????????????????????????????????????????????????????????????????????????????????????????????????????????????????????????????????????????????????????????????????????????????????????????????????????????????????????????????????????????????????????????????????????????????????????????????????????????????????????????????????????????????????????????????????????????????????????????????????????????????????????????????????????????????????????????????????????????????????????????????????????????????????????????????????????????????????????????????????????????????????????????????????????????????????????????????????????????????????????????????????????????????????????????????????????????????????????????????????????????????????????

			>Pitcairnia_poortmanii
			ATAATTATAATAAAGTTAAATATGTCGAAATCCTTTTTGGAATAATACCAAATCGAAAATAAATGTCCGATAGCAAGTTGATCAGTTAATTCAATAAGAGATAAATGGGAGATAGCACTCGATTTAGTTGGTACCACCCAACCGAATCCGATTCAATCGTTTACTCATTCAATGAGTCAATTTTCAAGTTCAGCCAATC-TTTTTTTTTTGAAAAAAAAATTGCAAGTAGATGAAATCGTGAATAAATGTGTTTCATTTCTCTATCATTATAGAAAATAAGATCCATATTATATTACTTATGGAATTCGAACCCGAACTCGATTTATGATTCATTATTTCGATCTTATTGGCCTTTGTTCTTTATTTTTTATTTTTCATATAGATTTCCGTCTTTATATTATACCCTTTCGTGGATGAATTATGCCTATTTTCATATCTAGGATTTACATATACAACATATATGACTGTCAAGAGGGAATTTTTCTCAGTATTTAGATATTTAGATTCAAAATAAGAAAGGGATCAATTCAATTATAAACCTGCAAAACAAAGATTAGGATTGGGTTGGGTTGCGCTATATATATCAAAGAGTATACAATAATGATGTATTTGGTGAATCAAATACATGGTCTAATAACAAACCATTTTAACATAACATTTTAATGAGTTGATAATATTAATTGAATATTGAATATTTTTTGAAAGATTTTTGTTAAAGGTTTCATTCATGCCTAATCCATATCGAGTAGACCTTGTTGTTGTGAG

			>Pitcairnia_prolifera
			??????????????????????????????????????????????????????????????????????????????????????????????????????????????????????????????????????????????????????????????????????????????????????????????????????????????????????????????????????????????????????????????????????????????????????????????????????????????????????????????????????????????????????????????????????????????????????????????????????????????????????????????????????????????????????????????????????????????????????????????????????????????????????????????????????????????????????????????????????????????????????????????????????????????????????????????????????????????????????????????????????????????????????????????????????????????????????????????????????????????????????????????????????????????????????????????

			>Pitcairnia_pseudoundulata
			??????????????????????????????????????????????????????????????????????????????????????????????????????????????????????????????????????????????????????????????????????????????????????????????????????????????????????????????????????????????????????????????????????????????????????????????????????????????????????????????????????????????????????????????????????????????????????????????????????????????????????????????????????????????????????????????????????????????????????????????????????????????????????????????????????????????????????????????????????????????????????????????????????????????????????????????????????????????????????????????????????????????????????????????????????????????????????????????????????????????????????????????????????????????????????????????

			>Pitcairnia_pungens
			??????????????????????????????????????????????????????????????????????????????????????????????????????????????????????????????????????????????????????????????????????????????????????????????????????????????????????????????????????????????????????????????????????????????????????????????????????????????????????????????????????????????????????????????????????????????????????????????????????????????????????????????????????????????????????????????????????????????????????????????????????????????????????????????????????????????????????????????????????????????????????????????????????????????????????????????????????????????????????????????????????????????????????????????????????????????????????????????????????????????????????????????????????????????????????????????

			>Pitcairnia_punicea
			ATAATTATAATAAAGTTAAATATGTCGAAATCCTTTTTGGAATAATACCAAATCGAAAATAAATGTCCGATAGCAAGTTGATCAGTTAATTCAATAAGAGATAAATGGGAGATAGCACTCGATTTAGTTGGTACCACCCAACCGAATCCGATTCAATCGTTTACTCATTCAATGAGTCAATTTTCAAGTTCAGCCAATC--TTTTTTTTTGAAAAAAAAATTGCAAGTAGATGAAATCGTGAATAAATGTGTTTCATTTCTCTATCATTATAGAAAATAAGATCCATATTATATTACTTATGGAATTCGAACCCGAACTCGATTTATGATTCATAATTTCGATCTTATTGGCCTTTGTTCTTTATTTTTGATTTTTCATATAGATTTCCGTCTTTATATTATACCCTTTCGTGGATGAATTATGCCTATTTTCATATCTAGGATTTACATATACAACATATATGACTGTCAAGAGGGAATTTTTCTCAGTATTTAGATATTTAGATTCAAAATAAGAAAGGGATCAATTCAATTATAAACCTGCAAAACAAAGATTAGGATTGGGTTGGGTTGCGCTATATATATCAAAGAGTATACAATAATGATGTATTTGGTGAATCAAATACATGGTCTAATAACAAACCATTTTAACATAACATTTTAATGAGTTGATAATATTAATTGAATATTGAATATTTTTTGAAAGATTTTTGTTAAAGGTTTCATTCATGCCTAATCCATATCGAGTAGACCTTGTTGTTGTGAG

			>Pitcairnia_rectiflora
			??????????????????????????????????????????????????????????????????????????????????????????????????????????????????????????????????????????????????????????????????????????????????????????????????????????????????????????????????????????????????????????????????????????????????????????????????????????????????????????????????????????????????????????????????????????????????????????????????????????????????????????????????????????????????????????????????????????????????????????????????????????????????????????????????????????????????????????????????????????????????????????????????????????????????????????????????????????????????????????????????????????????????????????????????????????????????????????????????????????????????????????????????????????????????????????????

			>Pitcairnia_recurvata
			??????????????????????????????????????????????????????????????????????????????????CAGTTAATTCAATAAGAGATAAATGGGAGATAGCACTCGATTTAGTTGGTACCACCCAACCGAATCCGATTCAATCGTTTACTCATTCAATGAGTCAATTTTCAAGTTCAGCCAATC--TTTTTTTTTGAAAAAAAAATTGCAAGTAGATGAAATCGTGAATAAATGTGTTTCATTTCTCTATCATTATAGAAAATAAGATCCCTATTATATTACTTATGGAATTCGAACCCGAACTCGATTTATGATTCATTATTTCGATCTTATTGGCCTTTGTTCTTTATTTTTGATTTTTCATATAGATTTCCGTCTTTATATTATACCCTTTCGTGGATGAATTATGCCTATTTTCATATCTAGGATTTACATATACAACATATATGACTGTCAAGAGGGAATTTTTCTCAGTATTTAGATATTTAGATTCAAAATAAGAAAGGGATCAATTCAATTATAAACCTGCAAAACAAAGATTAGGATTGGGTTGGGTTGCGCTATATATATCAAAGAGTATACAATAATGATGTATTTGGTGAATCAAATACATGGTCTAATAACAAACCATTTTAACATAACATTTTAATGAGTTGATAATATTAATTGAATATTGAATATTTTTTGAAAGATTTTTGTTAAAGGTTTCATTCATGCCTAATCCATATCGAGTAGACCTTGTTGTTGTGAG

			>Pitcairnia_reflexiflora
			??????????????????????????????????????????????????????????????????????????????????????????????????????????????????????????????????????????????????????????????????????????????????????????????????????????????????????????????????????????????????????????????????????????????????????????????????????????????????????????????????????????????????????????????????????????????????????????????????????????????????????????????????????????????????????????????????????????????????????????????????????????????????????????????????????????????????????????????????????????????????????????????????????????????????????????????????????????????????????????????????????????????????????????????????????????????????????????????????????????????????????????????????????????????????????????????

			>Pitcairnia_riparia
			??????????????????????????????????????????????????????????????????????????????????????????????????????????????????????????????????????????????????????????????????????????????????????????????????????????????????????????????????????????????????????????????????????????????????????????????????????????????????????????????????????????????????????????????????????????????????????????????????????????????????????????????????????????????????????????????????????????????????????????????????????????????????????????????????????????????????????????????????????????????????????????????????????????????????????????????????????????????????????????????????????????????????????????????????????????????????????????????????????????????????????????????????????????????????????????????

			>Pitcairnia_roseoalba
			??????????????????????????????????????????????????????????????????????????????????????????????????????????????????????????????????????????????????????????????????????????????????????????????????????????????????????????????????????????????????????????????????????????????????????????????????????????????????????????????????????????????????????????????????????????????????????????????????????????????????????????????????????????????????????????????????????????????????????????????????????????????????????????????????????????????????????????????????????????????????????????????????????????????????????????????????????????????????????????????????????????????????????????????????????????????????????????????????????????????????????????????????????????????????????????????

			>Pitcairnia_rubronigriflora
			??????????????????????????????????????????????????????????????????????????????????CAGTTAATTCAATAAGAGATAAATGGGAGATAGCACTCGATTTAGTTGGTACCACCCAACCGAATCCGATTCAATCGTTTACTCATTCAATGAGTCAATTTTCAAGTTCAGCCAATC--TTTTTTTTTGAAAAAAAAATTGCAAGTAGATGAAATCGTGAATAAATGTGTTTCATTTCTCTATCATTATAGAAAATAAGATCCATATTATATTACTTATGGAATTCGAACCCGAACTCGATTTATGATTCATTATTTCGATCTTATTGGCCTTTGTTCTTTATTTTTGATTTTTCATATAGATTTCCGTCTTTATATTATACCCTTTCGTGGATGAATTATGCCTATTTTCATATCTAGGATTTACATATACAACATATATGACTGTCAAGAGGGAATTTTTCTCAGTATTTAGATATTTAGATTCAAAATAAGAAAGGGATCAATTCAATTATAAACCTGCAAAACAAGGATTAGGATTGGGTTGGGTTGCGCTATATATATCAAAGAGTATACAATAATGATGTATTTGGTGAATCAAATACATGGTCTAATAACAAACCATTTTAACATAACATTTTAATGAGTTGATAATATTAATTGAATATTGAATATTTTTTGAAAGATTTTTGTTAAAGGTTTCATTCATGCCTAATCCATATCGAGTAGACCTTGTTGTTGTGAG

			>Pitcairnia_sanguinea
			??????????????????????????????????????????????????????????????????????????????????????????????????????????????????????????????????????????????????????????????????????????????????????????????????????????????????????????????????????????????????????????????????????????????????????????????????????????????????????????????????????????????????????????????????????????????????????????????????????????????????????????????????????????????????????????????????????????????????????????????????????????????????????????????????????????????????????????????????????????????????????????????????????????????????????????????????????????????????????????????????????????????????????????????????????????????????????????????????????????????????????????????????????????????????????????????

			>Pitcairnia_smithiorum
			??????????????????????????????????????????????????????????????????????????????????????????????????????????????????????????????????????????????????????????????????????????????????????????????????????????????????????????????????????????????????????????????????????????????????????????????????????????????????????????????????????????????????????????????????????????????????????????????????????????????????????????????????????????????????????????????????????????????????????????????????????????????????????????????????????????????????????????????????????????????????????????????????????????????????????????????????????????????????????????????????????????????????????????????????????????????????????????????????????????????????????????????????????????????????????????????

			>Pitcairnia_spicata
			??????????????????????????????????????????????????????????????????????????????????????????????????????????????????????????????????????????????????????????????????????????????????????????????????????????????????????????????????????????????????????????????????????????????????????????????????????????????????????????????????????????????????????????????????????????????????????????????????????????????????????????????????????????????????????????????????????????????????????????????????????????????????????????????????????????????????????????????????????????????????????????????????????????????????????????????????????????????????????????????????????????????????????????????????????????????????????????????????????????????????????????????????????????????????????????????

			>Pitcairnia_sprucei
			??????????????????????????????????????????????????????????????????????????????????????????????????????????????????????????????????????????????????????????????????????????????????????????????????????????????????????????????????????????????????????????????????????????????????????????????????????????????????????????????????????????????????????????????????????????????????????????????????????????????????????????????????????????????????????????????????????????????????????????????????????????????????????????????????????????????????????????????????????????????????????????????????????????????????????????????????????????????????????????????????????????????????????????????????????????????????????????????????????????????????????????????????????????????????????????????

			>Pitcairnia_squarrosa
			??????????????????????????????????????????????????????????????????????????????????????????????????????????????????????????????????????????????????????????????????????????????????????????????????????????????????????????????????????????????????????????????????????????????????????????????????????????????????????????????????????????????????????????????????????????????????????????????????????????????????????????????????????????????????????????????????????????????????????????????????????????????????????????????????????????????????????????????????????????????????????????????????????????????????????????????????????????????????????????????????????????????????????????????????????????????????????????????????????????????????????????????????????????????????????????????

			>Pitcairnia_staminea
			??????????????????????????????????????????????????????????????????????????????????????????????????????????????????????????????????????????????????????????????????????????????????????????????????????????????????????????????????????????????????????????????????????????????????????????????????????????????????????????????????????????????????????????????????????????????????????????????????????????????????????????????????????????????????????????????????????????????????????????????????????????????????????????????????????????????????????????????????????????????????????????????????????????????????????????????????????????????????????????????????????????????????????????????????????????????????????????????????????????????????????????????????????????????????????????????

			>Pitcairnia_sulphurea
			??????????????????????????????????????????????????????????????????????????????????????????????????????????????????????????????????????????????????????????????????????????????????????????????????????????????????????????????????????????????????????????????????????????????????????????????????????????????????????????????????????????????????????????????????????????????????????????????????????????????????????????????????????????????????????????????????????????????????????????????????????????????????????????????????????????????????????????????????????????????????????????????????????????????????????????????????????????????????????????????????????????????????????????????????????????????????????????????????????????????????????????????????????????????????????????????

			>Pitcairnia_utcubambensis
			??????????????????????????????????????????????????????????????????????????????????????????????????????????????????????????????????????????????????????????????????????????????????????????????????????????????????????????????????????????????????????????????????????????????????????????????????????????????????????????????????????????????????????????????????????????????????????????????????????????????????????????????????????????????????????????????????????????????????????????????????????????????????????????????????????????????????????????????????????????????????????????????????????????????????????????????????????????????????????????????????????????????????????????????????????????????????????????????????????????????????????????????????????????????????????????????

			>Pitcairnia_villetaensis
			??????????????????????????????????????????????????????????????????????????????????????????????????????????????????????????????????????????????????????????????????????????????????????????????????????????????????????????????????????????????????????????????????????????????????????????????????????????????????????????????????????????????????????????????????????????????????????????????????????????????????????????????????????????????????????????????????????????????????????????????????????????????????????????????????????????????????????????????????????????????????????????????????????????????????????????????????????????????????????????????????????????????????????????????????????????????????????????????????????????????????????????????????????????????????????????????

			>Pitcairnia_wendlandii
			ATAATTATAATAAAGTTAAATATGTCGAAATCCTTTTTGGAATAATACCAAATCGAAAATAAATGTCCGATAGCAAGTTGATCAGTTAATTCAATAAGAGATAAATGGGAGATAGCACTCGATTTAGTTGGTACCACCCAACCGAATCCGATTCAATCGTTTACTCATTCAATGAGTCAATTTTCAAGTTCAGCCAATCTTTTTTTTTTGAAAAAAAAAATTGCAAGTAGATGAAATCGTGAATAAAAGTGTTTCATTTCTCTATCATTATAGAAAATAAGATCCATATTATATTACTTATGGAATTCGAACCCGAACTCGATTTATGATTCATTATTTCGATCTTATTGGCCTTTGTTCTTTATTTTTGATTTTTCATATAGATTTCCGTCTTTATATTATACCCTTTCGTGGATGAATTATGCCTATTTTCATATCTAGGATTTACATATACAACATATATTACTGTCAAGAGGGAATTTTTCTCAGTATTTAGATATTTAGATTCAAAATAAGAAAGGGATCAATTCAATTATAAACCTGCAAAACAAAGATTAGGATTGGGTTGGGTTGCGCTATATATATCAAAGAGTATACAATAATGATGTATTTGGTGAATCAAATACATGGTCTAATAACAAACCATTTTAACATAACATTTTAATGAGTTGATAATATTAATTGAATATTGAATATTTTTTGAAAGATTTTTGTTAAAGGTTTCATTCATGCCTAATCCATATCGAGTAGACCTTGTTGTTGTGAG

			>Pitcairnia_xanthocalyx
			??????????????????????????????????????????????????????????????????????????????????????????????????????????????????????????????????????????????????????????????????????????????????????????????????????????????????????????????????????????????????????????????????????????????????????????????????????????????????????????????????????????????????????????????????????????????????????????????????????????????????????????????????????????????????????????????????????????????????????????????????????????????????????????????????????????????????????????????????????????????????????????????????????????????????????????????????????????????????????????????????????????????????????????????????????????????????????????????????????????????????????????????????????????????????????????????

			>Pitcairnia_yaupibajaensis
			??????????????????????????????????????????????????????????????????????????????????????????????????????????????????????????????????????????????????????????????????????????????????????????????????????????????????????????????????????????????????????????????????????????????????????????????????????????????????????????????????????????????????????????????????????????????????????????????????????????????????????????????????????????????????????????????????????????????????????????????????????????????????????????????????????????????????????????????????????????????????????????????????????????????????????????????????????????????????????????????????????????????????????????????????????????????????????????????????????????????????????????????????????????????????????????????

			>Portea_alatisepala
			??????????????????????????????????????????????????????????????????????????????????????????????????????????????????????????????????????????????????????????????????????????????????????????????????????????????????????????????????????????????????????????????????????????????????????????????????????????????????????????????????????????????????????????????????????????????????????????????????????????????????????????????????????????????????????????????????????????????????????????????????????????????????????????????????????????????????????????????????????????????????????????????????????????????????????????????????????????????????????????????????????????????????????????????????????????????????????????????????????????????????????????????????????????????????????????????

			>Portea_filifera
			??????????????????????????????????????????????????????????????????????????????????????????????????????????????????????????????????????????????????????????????????????????????????????????????????????????????????????????????????????????????????????????????????????????????????????????????????????????????????????????????????????????????????????????????????????????????????????????????????????????????????????????????????????????????????????????????????????????????????????????????????????????????????????????????????????????????????????????????????????????????????????????????????????????????????????????????????????????????????????????????????????????????????????????????????????????????????????????????????????????????????????????????????????????????????????????????

			>Portea_grandiflora
			ATAATTATAATAAAGTGAAATATGTCGAAATCCTTTTTGGAATAATACCAAATCGAAAATAAATGTCCGATAGCAAGTTGATCAGTTAATTCAATAAGAGATAAATGGGAGATAGCACTCGATTTAGTTGGTACCACCCAACCGAATCCGATTCACTCGTTTACTCATTCAATGAGTAAATTTTCAAGTTCAGCCAATCTTTTTTTTTTTAAAAAAAAAATTGCAAGTAGATGAAATCGTGAATAAATGTGTTTCATTTCTATATCATTATATAAAAAAAGATCCATATTATATTACTTATGGAATTCGAACCCGAACTCGATTTATGATTTATTATTTGGATCTTATTGGCCTTTGTTCTTTATTTTTGATTTTTCATATAGATTTCCGTCTTTATATTATACCCTTTCGTGGATGAATTATGCCTATTTTCACATCTAGGATTTACATATACAACATATATTACTGTCAAGAGGGAATTTTTCTCAGTATTTAGATATTTAGATTCAAAATAAGAAAGGGATCAATTCAATTATAAACCTGCAAAACAAAGATTAGGATTGGGTTGGGTTGCGCTATATATATCAAAGAGTATACAATAATGATGTATTTGGTGAATCAAATACATGGTCTAATAACAAACCATTTTAACATAACATTTTAATGAGTTGATAATATTAATTGAATATTGAATATTTTTTGAAAGATTTTTGTTAAAGGTTTCATTCATGCCTAATCCATATCGAGTAG????????????????

			>Portea_kermesina
			???????????????????????????????????TTTGGAATAATACCAAATCGAAAATAAATGTCCGATAGCAAGTTGATCAGTTAATTCAATAAGAGATAAATGGGAGATAGCACTCGATTTAGTTGGTACCACCCAACCGAATCCGATTCACTCGTTTACTCATTGAATGAGTAAATTTTCAAGTTCAGCCAATC-TTTTTTTTTTAAAAAAAAAATTGCAAGTAGATGAAATCGTGAATAAATGTGTTTCATTTCTATATCATTATATAAAAAAAGATCCATATTATATTACTTATGGAATTCGAACCCGAACTCGATTTATGATTTATTATTTGGATCTTATTGGCCTTTGTTCTTTATTTTTGATTTTTCATATAGATTTCCGTCTTTATATTATACCCTTTCGTGGATGAATTATGCCTATTTTCACATCTAGGATTTACATATACAACATATATTACTGTCAAGAGGGAATTTTTCTCAGTATTTAGATATTTAGATTCAAAATAAGAAAGGGATCAATTCAATTATAAACCTGCAAAACAAAGATTAGGATTGGGTTGGGTTGCGCTATATATATCAAAGAGTATACAATAATGATGTATTTGGTGAATCAAATACATGGTCTAATAACAAACCATTTTAACATAACATTTTAATGAGTTGATAATATTAATTGAATATTGACTATTTTTTGAAAGATTTTTGTTAAAGGTTTCATTCATGCCTAATCCATATCGAGTAGACCTTGTTGTTGTG??

			>Portea_nana
			??????????????????????????????????????????????????????????????????????????????????????????????????????????????????????????????????????????????????????????????????????????????????????????????????????????????????????????????????????????????????????????????????????????????????????????????????????????????????????????????????????????????????????????????????????????????????????????????????????????????????????????????????????????????????????????????????????????????????????????????????????????????????????????????????????????????????????????????????????????????????????????????????????????????????????????????????????????????????????????????????????????????????????????????????????????????????????????????????????????????????????????????????????????????????????????????

			>Portea_petropolitana
			??????????????????????????????????????????????????????????????????????????????????????????????????????????????????????????????????????????????????????????????????????????????????????????????????????????????????????????????????????????????????????????????????????????????????????????????????????????????????????????????????????????????????????????????????????????????????????????????????????????????????????????????????????????????????????????????????????????????????????????????????????????????????????????????????????????????????????????????????????????????????????????????????????????????????????????????????????????????????????????????????????????????????????????????????????????????????????????????????????????????????????????????????????????????????????????????

			>Portea_petropolitana_var._extensa
			ATAATTATAATAAAGTGAAATATGTCGAAATCCTTTTTGGAATAATACCAAATCGAAAATAAATGTCCGATAGCAAGTTGATCAGTTAATTCAATAAGAGATAAATGGGAGATAGCACTCGATTTAGTTGGTACCACCCAACCGAATCCGATTCACTCGTTTACTCATTCAATGAGTAAATTTTCAAGTTCAGCCAATCTTTTTTTTTTGAAAAAAAAAATTGCAAGTAGATGAAATCGTGAATAAATGTGTTTCATTTCTATATCATTATATAAAAAAAGATCCATATTATATTACTTATGGAATTCGAACCCGAACTCGATTTATGATTTATTATTTGGATCTTATTGGCCTTTGTTCTTTATTTTTGATTTTTCATATAGATTTCCGTCTTTATATTATACCCTTTCGTGGATGAATTATGCCTATTTTCACATCTAGGATTTACATATACAACATATATTACTGTCAAGAGGGAATTTTTCTCAGTATTTAGATATTTAGATTCAAAATAAGAAAGGGATCAATTCAATTATAAACCTGCAAAACAAAGATTAGGATTGGGTTGGGTTGCGCTATATATATCAAAGAGTATACAATAATGATGTATTTGGTGAATCAAATACATGGTCTAATAACAAACCATTTTAACATAACATTTTAATGAGTTGATAATATTAATTGAATATTGAATATTTTTTGAAAGATTTTTGTTAAAGGTTTCATTCATGCCTAATCCATATCGAGTAGACCTTGTTGTTGTG??

			>Portea_petropolitana_var._noettigii
			??????????????????????????????????????????????????????????????????????????????????????????????????????????????????????????????????????????????????????????????????????????????????????????????????????????????????????????????????????????????????????????????????????????????????????????????????????????????????????????????????????????????????????????????????????????????????????????????????????????????????????????????????????????????????????????????????????????????????????????????????????????????????????????????????????????????????????????????????????????????????????????????????????????????????????????????????????????????????????????????????????????????????????????????????????????????????????????????????????????????????????????????????????????????????????????????

			>Pseudalcantarea_grandis
			??????????????????????????????????????????????????????????????????????????????????????????????????????????????????????????????????????????????????????????????????????????????????????????????????????????????????????????????????????????????????????????????????????????????????????????????????????????????????????????????????????????????????????????????????????????????????????????????????????????????????????????????????????????????????????????????????????????????????????????????????????????????????????????????????????????????????????????????????????????????????????????????????????????????????????????????????????????????????????????????????????????????????????????????????????????????????????????????????????????????????????????????????????????????????????????????

			>Pseudalcantarea_macropetala
			??????????????????????????????????????????????????????????????????????????????????????????????????????????????????????????????????????????????????????????????????????????????????????????????????????????????????????????????????????????????????????????????????????????????????????????????????????????????????????????????????????????????????????????????????????????????????????????????????????????????????????????????????????????????????????????????????????????????????????????????????????????????????????????????????????????????????????????????????????????????????????????????????????????????????????????????????????????????????????????????????????????????????????????????????????????????????????????????????????????????????????????????????????????????????????????????

			>Pseudalcantarea_viridiflora
			ATAATTATAATAAAGTGAAATATGTCGAAATCCTTTTTGGAATAATACCAAATCGAAAATAAATGTCCGATAGCAAGTTGATCAGTTAATTCAATAAGAGATAAATGGGAGATAGCACTCGATTTAGTTGGTACCACCCAACCGAATCCGATTCAATCGTTTACTCATTCAATGAGTAAATTTTCAAGTTCAGCCAATCTTTTTTTTTTGAAAAAAAAAATTGCAAGTAAATGAAATCGTGAATAAATGTGTTTCATTTCTCTATCATTATAGAAAAAAAGATCCATATTATATTACTTATGGAATTCGAACCCGAACTCGATTTATGATTCATTATTTCGATCTTATTGGTCTTTGTTCTTTA-------TTTTTTATATAGATTTCCGTCTTTATATTATACCCTTTCGTAGATGAATTATGCTTATTTTCACATCTAGGATTTACATATACAACATATATTACTGTCAAGAGGGAATTTTTCTCAGTATTTAGATATTTAGATTCAAAATAAGAAAGGGATCAATTCAATTATAAACCCACAAAACAAAAATTAGGATTGGGTTGGGTTGCGCTATATATATCAAAGAGTATACAATAATGATGTATTTGGTGAATCAAATACATGGTCTAATAACAAACCATTTTAACATAACATTTTAATGAGTTGATAATATTAATTGAATATTGAATATTTTTTGAAAGATTTTTGTTAAAGGTTTCATTCATGCCTAATCCATATCGAGTAGACCTTGTTGTTGTGAG

			>Pseudaraeococcus_parviflorus
			??????????????????????????????????????????????????????????????????????????????????????????????????????????????????????????????????????????????????????????????????????????????????????????????????????????????????????????????????????????????????????????????????????????????????????????????????????????????????????????????????????????????????????????????????????????????????????????????????????????????????????????????????????????????????????????????????????????????????????????????????????????????????????????????????????????????????????????????????????????????????????????????????????????????????????????????????????????????????????????????????????????????????????????????????????????????????????????????????????????????????????????????????????????????????????????????

			>Puya_aequatorialis
			ATAATTATAATAAAGTGAAATATGTCGAAATCCTTTTTGGAATAATACCAAATCGAAAATAAATGTCCGATAGCAAGTTGATCAGTTAATTCAATAAGAGATAAATGGGAGATAGCACTCGATTTAGTTGGTACCACCCAACCGAATCCGATTCAATCGTTTACTCATTGAATGAGTAAATTTGCAAGTTCAGCCAATCTTTTTTTTTTGAAAAAAAAAATTGCAAGTAGATGAAATCGTGAATAAATGTGTTTCATTTCTCTATCATTATATAAAATAAGATCCATATTATATTACTTATGGAATTCGAACCCGAACTCGATTTATGATTTATTATTTCGATCTTATTGGCCTTTGTTCTTTATTTTTGATTTTTCATATAGATTTCCGTCTTTATATTATACCCTTTCGTGGATGAATTATGCCTATTTTCACATCTAGGATTTACATATACAACATATATTACTGTCAAGAGAGAATTTTTCTCAGTATTTAGATATTTAGATTCAAAATAAGAAAGGGATCAATTCAATTATAAACCTGCAAAACAAAGATTAGGATTGGGTTGGGTTGCGCTATATATATCAAAGAGTATACAATAATGATGTATTTGGTGAATCAAATACATGGTCTAATAACAAACCATTTTAACATAACATTTTAATGAGTTGATAATATTAATTGAATATTGAATATTTTTTGAAAGATTTTTGTTAAAGGTTTCATTCATGCCTAATCCATATCGAGTAGACCTTGTTGTTGTGAG

			>Puya_alpestris
			ATAATTATAATAAAGTGAAATATGTCGAAATCCTTTTTGGAATAATACCAAATCGAAAATAAATGTCCGATAGCAAGTTGATCAGTTAATTCAATAAGAGATAAATGGGAGATAGCACTCGATTTAGTTGGTACCACCCAACCGAATCCGATTCAATCGTTTACTCATTGAATGAGTAAATTTTCAAGTTCAGCCAATCTTTTTTTTTTTAAAAAAAAAATTGCAAGTAGATGAAATCGTGAATAAATGTGTTTCATTTCTCTATCATTATATAAAATAAGATCCATATTATATTACTTATGGAATTCGAACCCGAACTCGATTTATGATTTATTATTTCGATCTTATTGGCCTTTGTTCTTTATTTTTGATTTTTCATATAGATTTCCGTCTTTATATTATACCCTTTCGTGGATGAATTATGCCTATTTTCACATCTAGGATTTACATATACAACATATATTACTGTCAAGAGAGAATTTTTATCAGTATTTAGATATTTAGATTCAAAATAAGAAAGGGATCAATTCAATCATAAACCTGCAAAACAAAGATTAGGATTGGGTTGGGTTGCGCTATATATATCAAAGAGTATACAATAATGATGTATTTGGTGAATCAAATACATGGTCTAATAACAAACCATTTTAACATAACATTTTAATGAGTTGATAATATTAATTGAATATTGAATATTTTTTGAAAGATTTTTGTTAAAGGTTTCATTCATGCCTAATCCATATCGAGTAGACCTTGTTGTTGTG??

			>Puya_angusta
			??????????????????????????????????????????????????????????????????????????????????????????????????????????????????????????????????????????????????????????????????????????????????????????????????????????????????????????????????????????????????????????????????????????????????????????????????????????????????????????????????????????????????????????????????????????????????????????????????????????????????????????????????????????????????????????????????????????????????????????????????????????????????????????????????????????????????????????????????????????????????????????????????????????????????????????????????????????????????????????????????????????????????????????????????????????????????????????????????????????????????????????????????????????????????????????????

			>Puya_bicolor
			??????????????????????????????????????????????????????????????????????????????????????????????????????????????????????????????????????????????????????????????????????????????????????????????????????????????????????????????????????????????????????????????????????????????????????????????????????????????????????????????????????????????????????????????????????????????????????????????????????????????????????????????????????????????????????????????????????????????????????????????????????????????????????????????????????????????????????????????????????????????????????????????????????????????????????????????????????????????????????????????????????????????????????????????????????????????????????????????????????????????????????????????????????????????????????????????

			>Puya_boliviensis
			??????????????????????????????????????????????????????????????????????????????????????????????????????????????????????????????????????????????????????????????????????????????????????????????????????????????????????????????????????????????????????????????????????????????????????????????????????????????????????????????????????????????????????????????????????????????????????????????????????????????????????????????????????????????????????????????????????????????????????????????????????????????????????????????????????????????????????????????????????????????????????????????????????????????????????????????????????????????????????????????????????????????????????????????????????????????????????????????????????????????????????????????????????????????????????????????

			>Puya_castellanosii
			??????????????????????????????????????????????????????????????????????????????????????????????????????????????????????????????????????????????????????????????????????????????????????????????????????????????????????????????????????????????????????????????????????????????????????????????????????????????????????????????????????????????????????????????????????????????????????????????????????????????????????????????????????????????????????????????????????????????????????????????????????????????????????????????????????????????????????????????????????????????????????????????????????????????????????????????????????????????????????????????????????????????????????????????????????????????????????????????????????????????????????????????????????????????????????????????

			>Puya_chilensis
			ATAATTATAATAAAGTGAAATATGTCGAAATCCTTTTTGGAATAATACCAAATYGAAAATAAATGTCTGATAGCAAGTTGATCAGTTAATTCAATAAGAGATAAATGGGAGATAGCACTCGATTTAGTTGGTACCACCCAACCGAATCTGATTCAATCGTTTACTCATTCAATGAGTAAATTTTCAAGTTCAGCCAATCTTTTTTTTTTTCAAAAAAGAATTGCAAGTARATGAAATCGTGAATAAATGTGTTTCATTTCTCTATCATTATAGAAAATAAGAT-CATATTATATTACTTATGGAATTCGAACCCGAACTCGATTTATGATTTATTATTTCGATCTTATTGGCCTTTGTTCTTTATTTTTGATTTTTCATATAGATTTCCGTCTTTATATTATACCCTTTCGTGGATGAATTATGCCTATTTTCACATCTAGGATTTACATATACAACATATATTACTGTCAAGAGGGAATTTTTCTCAGTATTTAGATATTTAGATTCAAAATAAGAAAGGGATCAATTCAATTATAAACCTGCAAAACAAAGATTAGGATTGGGTTGGGTTGCGCTATATATATCAAAGAGTATACAATAATGATGTATTTGGTGAATCAAATACATGGTCTAATAACAAACCATTTTAACATAACATTTTAATGAGTTGATAATATTAATTGAATATTGAATATTTTTTGAAAGATTTTTGTTAAAGGTTTCATTCATGCCTAATCCATATCGAGTAGACCTTGTTGTTGTG??

			>Puya_coerulea
			??????????????????????????????????????????????????????????????????????????????????????????????????????????????????????????????????????????????????????????????????????????????????????????????????????????????????????????????????????????????????????????????????????????????????????????????????????????????????????????????????????????????????????????????????????????????????????????????????????????????????????????????????????????????????????????????????????????????????????????????????????????????????????????????????????????????????????????????????????????????????????????????????????????????????????????????????????????????????????????????????????????????????????????????????????????????????????????????????????????????????????????????????????????????????????????????

			>Puya_coerulea_var._violacea
			??????????????????????????????????????????????????????????????????????????????????CAGTTAATTCAATAAGAGATAAATGGGAGATAGCACTCGATTTAGTTGGTACCACCCAACCGAATCCGATTCAATCGTTTACTCATTCAATGAGTAAATTTTCAAGTTCAGCCAATCTTTTTTTTTTTCAAAAAAGAATTGCAAGTAGATGAAATCGTGAATAAATGTGTTTCATTTCTCTATCATTATAGAAAATAAGAT-CATATTATATTACTTATGGAATTCGAACCCGAACTCGATTTATGATTTATTATTTCGATCTTATTGGCCTTTGTTCTTTATTTTTGATTTTTCATATAGATTTCCGTCTTTATATTATACCCTTTCGTGGATGAATTATGCCTATTTTCACATCTAGGATTTACATATACAACATATATTACTGTCAAGAGGGAATTTTTCTCAGTATTTAGATATTTAGATTCAAAATAAGAAAGGGATCAATTCAATTATAAACCTGCAAAACAAAGATTAGGATTGGGTTGGGTTGCGCTATATATATCAAAGAGTATACAATAATGATGTATTTGGTGAATCAAATACATGGTCTAATAACAAACCATTTTAACATAACATTTTAATGAGTTGATAATATTAATTGAATATTGAATATTTTTTGAAAGATTTTTGTTAAAGGTTTCATTCATGCCTAATCCATATCGAGTAGACCTTGTTGTTGTGAG

			>Puya_compacta
			??????????????????????????????????????????????????????????????????????????????????????????????????????????????????????????????????????????????????????????????????????????????????????????????????????????????????????????????????????????????????????????????????????????????????????????????????????????????????????????????????????????????????????????????????????????????????????????????????????????????????????????????????????????????????????????????????????????????????????????????????????????????????????????????????????????????????????????????????????????????????????????????????????????????????????????????????????????????????????????????????????????????????????????????????????????????????????????????????????????????????????????????????????????????????????????????

			>Puya_dasylirioides
			??????????????????????????????????????????????????????????????????????????????????????????????????????????????????????????????????????????????????????????????????????????????????????????????????????????????????????????????????????????????????????????????????????????????????????????????????????????????????????????????????????????????????????????????????????????????????????????????????????????????????????????????????????????????????????????????????????????????????????????????????????????????????????????????????????????????????????????????????????????????????????????????????????????????????????????????????????????????????????????????????????????????????????????????????????????????????????????????????????????????????????????????????????????????????????????????

			>Puya_densiflora
			ATAATTATAATAAAGTGAAATATGTCGAAATCCTTTTTGGAATAATACCAAATCGAAAATAAATGTCCGATAGCAAGTTGATCAGTTAATTCAATAAGAGATAAATGGGAGATAGCACTCGATTTTGTTGGTACCACCCAACCGAATCCGATTCAATCGTTTACTCATTGAATGAGTAAATTTTCAAGTTCAGCCAATCTTTTTTTTTTTAAAAAAAAAATTGCAAGTAGATGAAATCGTGAATAAATGTGTTTCATTTCTCTATCATTATATAAAATAAGATCCATATTATATTACTTATGGAATTCGAACCCGAACTCGATTTATGATTTATTATTTCGATCTTATTGGCCTTTGTTCTTTATTTTTGATTTTTCATATAGATTTCCGTCTTTATATTATACCCTTTCGTGGATGAATTATGCCTATTTTCACATCTAGGATTTACATATACAACATATATTACTGTCAAGAGAGAATTTTTCTCAGTATTTAGATATTTAGATTCAAAATAAGAAAGGGATCAATTCAATTATAAACCTGCAAAACAAAGATTAGGATTGGGTTGGGTTGCGCTATATATATCAAAGAGTATACAATAATGATGTATTTGGTGAATCAAATACATGGTCTAATAACAAACCATTTTAACATAACATTTTAATGAGTTGATAATATTAATTGAATATTGAATATTTTTTGAAAGATTTTTGTTAAAGGTTTCATTCATGCCTAATCCATATCGAGTAGACCTTGTTGTTGTG??

			>Puya_dyckioides
			??????????????????????????????????????????????????????????????????????????????????????????????????????????????????????????????????????????????????????????????????????????????????????????????????????????????????????????????????????????????????????????????????????????????????????????????????????????????????????????????????????????????????????????????????????????????????????????????????????????????????????????????????????????????????????????????????????????????????????????????????????????????????????????????????????????????????????????????????????????????????????????????????????????????????????????????????????????????????????????????????????????????????????????????????????????????????????????????????????????????????????????????????????????????????????????????

			>Puya_ferreyrae
			??????????????????????????????????????????????????????????????????????????????????????????????????????????????????????????????????????????????????????????????????????????????????????????????????????????????????????????????????????????????????????????????????????????????????????????????????????????????????????????????????????????????????????????????????????????????????????????????????????????????????????????????????????????????????????????????????????????????????????????????????????????????????????????????????????????????????????????????????????????????????????????????????????????????????????????????????????????????????????????????????????????????????????????????????????????????????????????????????????????????????????????????????????????????????????????????

			>Puya_ferruginea
			ATAATTATAATAAAGTGAAATATGTCGAAATCCTTTTTGGAATAATACCAAATCGAAAATAAATGTCCGATAGCAAGTTGATCAGTTAATTCAATAAGAGATAAATGGGAGATAGCACTCGATTTAGTTGGTACCACCCAACCGAATCCGATTCAATCGTTTACTCATTGAATGAGTAAATTTTCAAGTTCAGCCAATCTTTTTTTTC--AAAAAAAAAATTGCAAGTAGATGAAATCGTGAATAAATGTGTTTCATTTCTCTATCATTATATAAAATAAGATCCATATTATATTACTTATGGAATTCGAACCCGAACTCGATTTATGATTTATTATTTCGATCTTATTGGCCTTTGTTCTTTATTTTTGATTTTTCATATAGATTTCCGTCTTTATATTATACCCTTTCGTGGATGAATTATGCCTATTTTCACATCTAGGATTTACATATACAACATATATTACTGTCAAGAGAGAATTTTTCTCAGTATTTAGATATTTAGATTCAAAATAAGAAAGGGATCAATTCAATTATAAACCTGCAAAACAAAGATTAGGATTGGGTTGGGTTGCGCTATATATATCAAAGAGTATACAATAATGATGTATTTGGTGAATCAAATACATGGTCTAATAACAAACCATTTTAACATAACATTTTAATGAGTTGATAATATTAATTGAATATTGAATATTTTTTGAAAGATTTTTGTTAAAGGTTTCATTCATGCCTAATCCATATCGAGTAGACCTTGTTGTTGTG??

			>Puya_floccosa
			??????????????????????????????????????????????????????????????????????????????????????????????????????????????????????????????????????????????????????????????????????????????????????????????????????????????????????????????????????????????????????????????????????????????????????????????????????????????????????????????????????????????????????????????????????????????????????????????????????????????????????????????????????????????????????????????????????????????????????????????????????????????????????????????????????????????????????????????????????????????????????????????????????????????????????????????????????????????????????????????????????????????????????????????????????????????????????????????????????????????????????????????????????????????????????????????

			>Puya_gilmartiniae
			??????????????????????????????????????????????????????????????????????????????????????????????????????????????????????????????????????????????????????????????????????????????????????????????????????????????????????????????????????????????????????????????????????????????????????????????????????????????????????????????????????????????????????????????????????????????????????????????????????????????????????????????????????????????????????????????????????????????????????????????????????????????????????????????????????????????????????????????????????????????????????????????????????????????????????????????????????????????????????????????????????????????????????????????????????????????????????????????????????????????????????????????????????????????????????????????

			>Puya_goudotiana
			??????????????????????????????????????????????????????????????????????????????????????????????????????????????????????????????????????????????????????????????????????????????????????????????????????????????????????????????????????????????????????????????????????????????????????????????????????????????????????????????????????????????????????????????????????????????????????????????????????????????????????????????????????????????????????????????????????????????????????????????????????????????????????????????????????????????????????????????????????????????????????????????????????????????????????????????????????????????????????????????????????????????????????????????????????????????????????????????????????????????????????????????????????????????????????????????

			>Puya_hamata
			??????????????????????????????????????????????????????????????????????????????????????????????????????????????????????????????????????????????????????????????????????????????????????????????????????????????????????????????????????????????????????????????????????????????????????????????????????????????????????????????????????????????????????????????????????????????????????????????????????????????????????????????????????????????????????????????????????????????????????????????????????????????????????????????????????????????????????????????????????????????????????????????????????????????????????????????????????????????????????????????????????????????????????????????????????????????????????????????????????????????????????????????????????????????????????????????

			>Puya_harmsii
			??????????????????????????????????????????????????????????????????????????????????????????????????????????????????????????????????????????????????????????????????????????????????????????????????????????????????????????????????????????????????????????????????????????????????????????????????????????????????????????????????????????????????????????????????????????????????????????????????????????????????????????????????????????????????????????????????????????????????????????????????????????????????????????????????????????????????????????????????????????????????????????????????????????????????????????????????????????????????????????????????????????????????????????????????????????????????????????????????????????????????????????????????????????????????????????????

			>Puya_herrerae
			??????????????????????????????????????????????????????????????????????????????????????????????????????????????????????????????????????????????????????????????????????????????????????????????????????????????????????????????????????????????????????????????????????????????????????????????????????????????????????????????????????????????????????????????????????????????????????????????????????????????????????????????????????????????????????????????????????????????????????????????????????????????????????????????????????????????????????????????????????????????????????????????????????????????????????????????????????????????????????????????????????????????????????????????????????????????????????????????????????????????????????????????????????????????????????????????

			>Puya_herzogii
			??????????????????????????????????????????????????????????????????????????????????CAGTTAATTCAATAAGAGATAAATGGGAGATAGCACTCGATTTAGTTGGTACCACCCAACCGAATCCGATTCAATCGTTTACTCATTGAATGAGTAAATTTTCAAGTTCMGCCAATCTTTTTTTTTTTAAAAAAAAAATTGCAAGTAGATGAAATCGTGAATAAATGTGTTTCATTTCTCTATCATTATATAAAATAAGATCCATATTATATTACTTATGGAATTCGAACCCGAACTCGATTTATGATTTATTATTTCGATCTTATTGGCCTTTGTTCTTTATTTTTGATTTTTCATATAGATTTCCGTCTTTATATTATACCCTTTCGTGGATGAATTATGCCTATTTTCACATCTAGGATTTACATATACAACATATATTACTGTCAAGAGAGAATTTTTCTCAGTATTTAGATATTTAGATTCAAAATAAGAAAGGGATCAATTCAATTATAAACCTGCAAAACAAAGATTAGGATTGGGTTGGGTTGCGCTATATATATCAAAGAGTATACAATAATGATGTATTTGGTGAATCAAATACATGGTCTAATAACAAACCATTTTAACATAACATTTTAATGAGTTGATAATATTAATTGAATATTGAATATTTTTTGAAAGATTTTTGTTAAAGGTTTCATTCATGCCTAATCCATATCGAGTAGACCTTGTTGTTGTGAG

			>Puya_humilis
			??????????????????????????????????????????????????????????????????????????????????????????????????????????????????????????????????????????????????????????????????????????????????????????????????????????????????????????????????????????????????????????????????????????????????????????????????????????????????????????????????????????????????????????????????????????????????????????????????????????????????????????????????????????????????????????????????????????????????????????????????????????????????????????????????????????????????????????????????????????????????????????????????????????????????????????????????????????????????????????????????????????????????????????????????????????????????????????????????????????????????????????????????????????????????????????????

			>Puya_lanata
			??????????????????????????????????????????????????????????????????????????????????????????????????????????????????????????????????????????????????????????????????????????????????????????????????????????????????????????????????????????????????????????????????????????????????????????????????????????????????????????????????????????????????????????????????????????????????????????????????????????????????????????????????????????????????????????????????????????????????????????????????????????????????????????????????????????????????????????????????????????????????????????????????????????????????????????????????????????????????????????????????????????????????????????????????????????????????????????????????????????????????????????????????????????????????????????????

			>Puya_laxa
			ATAATTATAATAAAGTGAAATATGTCGAAATCCTTTTTGGAATAATACCAAATCGAAAATAAATGTCCGATAGCAAGTTGATCAGTTAATTCAATAAGAGATAAATGGGAGATAGCACTCGATTTAGTTGGTACCACCCAACCGAATCCGATTCAATCGTTTACTCATTGAATGAGTAAATTTTCAAGTTCAGCCAATCTTTTTTTTTTTAAAAAAAAAATTGCAAGTAGATGAAATCGTGAATAAATGTGTTTCATTTCTCTATCATTATATAAAATAAGATCCATATTATATTACTTATGGAATTCGAACCCGAACTCGATTTATGATTTATTATTTCGATCTTATTGGCCTTTGTTCTTTATTTTTGATTTTTCATATAGATTTCCGTCTTTATATTATACCCTTTCGTGGATGAATTATGCCTATTTTCACATCTAGGATTTACATATACAACATATATTACTGTCAAGAGAGAATTTTTCTCAGTATTTAGATATTTAGATTCAAAATAAGAAAGGGATCAATTCAATTATAAACCTGCAAAACAAAGATTAGGATTGGGTTGGGTTGCGCTATATATATCAAAGAGTATACAATAATGATGTATTTGGTGAATCAAATACATGGTCTAATAACAAACCATTTTAACATAACATTTTAATGAGTTGATAATATTAATTGAATATTGAATATTTTTTGAAAGATTTTTGTTAAAGGTTTCATTCATGCCTAATCCATATCGAGTAGACCTTGTTGTTGTGAG

			>Puya_lineata
			??????????????????????????????????????????????????????????????????????????????????????????????????????????????????????????????????????????????????????????????????????????????????????????????????????????????????????????????????????????????????????????????????????????????????????????????????????????????????????????????????????????????????????????????????????????????????????????????????????????????????????????????????????????????????????????????????????????????????????????????????????????????????????????????????????????????????????????????????????????????????????????????????????????????????????????????????????????????????????????????????????????????????????????????????????????????????????????????????????????????????????????????????????????????????????????????

			>Puya_macrura
			??????????????????????????????????????????????????????????????????????????????????????????????????????????????????????????????????????????????????????????????????????????????????????????????????????????????????????????????????????????????????????????????????????????????????????????????????????????????????????????????????????????????????????????????????????????????????????????????????????????????????????????????????????????????????????????????????????????????????????????????????????????????????????????????????????????????????????????????????????????????????????????????????????????????????????????????????????????????????????????????????????????????????????????????????????????????????????????????????????????????????????????????????????????????????????????????

			>Puya_mima
			?????????????????????????????????????????????????????????????????????????????????????????????????????????????????????????????????????????????????????????????????????????????????????????????TCAGCCAATCTTTTTTTTTTCAAAAAAAAGATTGCAAGTAGATGAAATCGTGAATAAATGTGTTTCATTTCTCTATCATTATATAAAATAAGATCCATATTATATTACTTATGGAATTCGAACCCGAACTC?ATTTATGATTTATTATTTCGATCTTATTGGCCTTTGTTCTTTATTTTTGATTTTTCATATAGATTTCCGTCTTTATATTATACCCTTTCGTGGATGAATTATGCCTATTTTCACATCTAGGATTTACATATACAACATATATTACTGTCAAGAGAGAATTTTTCTCAGTATTTAGATATTTAGATTCAAAATAAGAAAGGGATCAATTCAATTATAAACCTGCAAAACAAAGATTAGGATTGGGTTGGGTTGCGCTATATATATCAAAGAGTATACAATAATGATGTATTTGGTGAATCAAATACATGGTCTAATAACAAACCATTTTAACATAACATTTTAATGAGTTGATAATATT------------------------------CC???????????????????????????????????????????????????????

			>Puya_mirabilis
			??????????????????????????????????????????????????????????????????????????????????CAGTTAATTCAATAAGAGATAAATGGGAGATAGCACTCGATTTAGTTGGTACCACCCAACCGAATCCGATTCAATCGTTTACTCATTGAATGAGTAAATTTTCAAGTTCAGCCAATCTTTTTTTTTTTAAAAAAAAAATTGCAAGTAGATGAAATCGTGAATAAATGTGTTTCATTTCTCTATCATTATATAAAATAAGATCCATATTATATTACTTATGGAATTCGAACCCGAACTCGATTTATGATTTATTATTTCGATCTTATTGGCCTTTGTTCTTTATTTTTGATTTTTCATATAGATTTCCGTCTTTATATTATACCCTTTCGTGGATGAATTATGCCTATTTTCACATCTAGGATTTACATATACAACATATATTACTGTCAAGAGAGAATTTTTATCAGTATTTAGATATTTAGATTCAAAATAAGAAAGGGATCAATTCAATCATAAACCTGCAAAACAAAGATTAGGATTGGGTTGGGTTGCGCTATATATATCAAAGAGTATACAATAATGATGTATTTGGTGAATCAAATACATGGTCTAATAACAAACCATTTTAACATAACATTTTAATGAGTTGATAATATTAATTGAATATTGAATATTTTTTGAAAGATTTTTGTTAAAGGTTTCATTCATGCCTAATCCATATCGAGTAGACCTTGTTGTTGTGAG

			>Puya_nana
			??????????????????????????????????????????????????????????????????????????????????????????????????????????????????????????????????????????????????????????????????????????????????????????????????????????????????????????????????????????????????????????????????????????????????????????????????????????????????????????????????????????????????????????????????????????????????????????????????????????????????????????????????????????????????????????????????????????????????????????????????????????????????????????????????????????????????????????????????????????????????????????????????????????????????????????????????????????????????????????????????????????????????????????????????????????????????????????????????????????????????????????????????????????????????????????????

			>Puya_nitida
			??????????????????????????????????????????????????????????????????????????????????????????????????????????????????????????????????????????????????????????????????????????????????????????????????????????????????????????????????????????????????????????????????????????????????????????????????????????????????????????????????????????????????????????????????????????????????????????????????????????????????????????????????????????????????????????????????????????????????????????????????????????????????????????????????????????????????????????????????????????????????????????????????????????????????????????????????????????????????????????????????????????????????????????????????????????????????????????????????????????????????????????????????????????????????????????????

			>Puya_nutans
			??????????????????????????????????????????????????????????????????????????????????????????????????????????????????????????????????????????????????????????????????????????????????????????????????????????????????????????????????????????????????????????????????????????????????????????????????????????????????????????????????????????????????????????????????????????????????????????????????????????????????????????????????????????????????????????????????????????????????????????????????????????????????????????????????????????????????????????????????????????????????????????????????????????????????????????????????????????????????????????????????????????????????????????????????????????????????????????????????????????????????????????????????????????????????????????????

			>Puya_obconica
			??????????????????????????????????????????????????????????????????????????????????????????????????????????????????????????????????????????????????????????????????????????????????????????????????????????????????????????????????????????????????????????????????????????????????????????????????????????????????????????????????????????????????????????????????????????????????????????????????????????????????????????????????????????????????????????????????????????????????????????????????????????????????????????????????????????????????????????????????????????????????????????????????????????????????????????????????????????????????????????????????????????????????????????????????????????????????????????????????????????????????????????????????????????????????????????????

			>Puya_pygmaea
			??????????????????????????????????????????????????????????????????????????????????????????????????????????????????????????????????????????????????????????????????????????????????????????????????????????????????????????????????????????????????????????????????????????????????????????????????????????????????????????????????????????????????????????????????????????????????????????????????????????????????????????????????????????????????????????????????????????????????????????????????????????????????????????????????????????????????????????????????????????????????????????????????????????????????????????????????????????????????????????????????????????????????????????????????????????????????????????????????????????????????????????????????????????????????????????????

			>Puya_raimondii
			ATAATTATAATAAAGTGAAATATGTCGAAATCCTTTTTGGAATAATACCAAATCGAAAATAAATGTCCGATAGCAAGTTGATCAGTTAATTCAATAAGAGATAAATGGGAGATAGCACTCGATTTAGTTGGTACCACCCAACCGAATCCGATTCAATCGTTTACTCATTGAATGAGTCAATTTTCAAGTTCAGCCAATCTTTTTTTTTTTAAAAAAAAAATTGCAAGTAGATGAAATCGTGAATAAATGTGTTTCATTTCTCTATCATTATATAAAAWAAGATCCATATTATATTACTTATGGAATTCGAACCCGAACTCGATTTATGATTTATTATTTCGATCTTATTGGCCTTTGTTCTTTATTTTTGATTTTTCATATAGATTTCCGTCTTTATATTATACCCTTTCGTGGATGAATTATGCCTATTTTCACATCTAGGATTTACATATACAACATATATTACTGTCAAGAGAGAATTTTTCTCAGTATTTAGATATTTAGATTCAAAATAAGAAAGGGATCAATTCAATTATAAACCTGCAAAACAAAGATTAGGATTGGGTTGGGTTGCGCTATATATATCAAAGAGTATACAATAATGATGTATTTGGTGAATCAAATACATGGTCTAATAACAAACCATTTTAACATAACATTTTAATGAGTTGATAATATTAATTGAATATTGAATATTTTTTGAAAGATTTTTGTTAAAGGTTTCATTCATGCCTAATCCATATCGAGTAGACCTTGTTGTTGTG??

			>Puya_santosii
			??????????????????????????????????????????????????????????????????????????????????????????????????????????????????????????????????????????????????????????????????????????????????????????????????????????????????????????????????????????????????????????????????????????????????????????????????????????????????????????????????????????????????????????????????????????????????????????????????????????????????????????????????????????????????????????????????????????????????????????????????????????????????????????????????????????????????????????????????????????????????????????????????????????????????????????????????????????????????????????????????????????????????????????????????????????????????????????????????????????????????????????????????????????????????????????????

			>Puya_spathacea
			??????????????????????????????????????????????????????????????????????????????????CAGTTAATTCAATAAGAGATAAATGGGAGATAGCACTCGATTTAGTTGGTACCACCCAACCGAATCCGATTCAATCGTTTACTCATTGAATGAGTAAATTTTCAAGTTCAGCCAATCTTTTTTTTTTTAAAAAAAAAATTGCAAGTAGATGAAATCGTGAATAAATGTGTTTCATTTCTCTATCATTATATAAAATAAGATCCATATTATATTACTTATGGAATTCGAACCCGAACTCGATTTATTATTTATTATTTCGATCTTATTGGCCTTTGTTCTTTATTTTTGATTTTTCATATAGATTTCCGTCTTTATATTATACCCTTTCGTGGATGAATTATGCCTATTTTCACATCTAGGATTTACATATACAACATATATTACTGTCAAGAGAGAATTTTTCTCAGTATTTAGATATTTAGATTCAAAATAAGAAAGGGATCAATTCAATTATAAACCTGCAAAACAAAGATTAGGATTGGGTTGGGTTGCGCTATATATATCAAAGAGTATACAATAATGATGTATTTGGTGAATCAAATACATGGTCTAATAACAAACCATTTTAACATAACATTTTAATGAGTTGATAATATTAATTGAATATTGAATATTTTTTGAAAGATTTTTGTTAAAGGTTTCATTCATGCCTAATCCATATCGAGTAGACCTTGTTGTTGTGAG

			>Puya_stenothyrsa
			??????????????????????????????????????????????????????????????????????????????????????????????????????????????????????????????????????????????????????????????????????????????????????????????????????????????????????????????????????????????????????????????????????????????????????????????????????????????????????????????????????????????????????????????????????????????????????????????????????????????????????????????????????????????????????????????????????????????????????????????????????????????????????????????????????????????????????????????????????????????????????????????????????????????????????????????????????????????????????????????????????????????????????????????????????????????????????????????????????????????????????????????????????????????????????????????

			>Puya_trianae
			??????????????????????????????????????????????????????????????????????????????????????????????????????????????????????????????????????????????????????????????????????????????????????????????????????????????????????????????????????????????????????????????????????????????????????????????????????????????????????????????????????????????????????????????????????????????????????????????????????????????????????????????????????????????????????????????????????????????????????????????????????????????????????????????????????????????????????????????????????????????????????????????????????????????????????????????????????????????????????????????????????????????????????????????????????????????????????????????????????????????????????????????????????????????????????????????

			>Puya_tuberosa
			??????????????????????????????????????????????????????????????????????????????????????????????????????????????????????????????????????????????????????????????????????????????????????????????????????????????????????????????????????????????????????????????????????????????????????????????????????????????????????????????????????????????????????????????????????????????????????????????????????????????????????????????????????????????????????????????????????????????????????????????????????????????????????????????????????????????????????????????????????????????????????????????????????????????????????????????????????????????????????????????????????????????????????????????????????????????????????????????????????????????????????????????????????????????????????????????

			>Puya_ultima
			??????????????????????????????????????????????????????????????????????????????????????????????????????????????????????????????????????????????????????????????????????????????????????????????????????????????????????????????????????????????????????????????????????????????????????????????????????????????????????????????????????????????????????????????????????????????????????????????????????????????????????????????????????????????????????????????????????????????????????????????????????????????????????????????????????????????????????????????????????????????????????????????????????????????????????????????????????????????????????????????????????????????????????????????????????????????????????????????????????????????????????????????????????????????????????????????

			>Puya_venusta
			??????????????????????????????????????????????????????????????????????????????????CAGTTAATTCAATAAGAGATAAATGGGAGATAGCACTCGATTTAGTTGGTACCACCCAACCGAATCTGATTCAATCGTTTACTCATTCAATGAGTAAATTTTCAAGTTCAGCCAATCTTTTTTTTTTTCAAAAAAGAATTGCAAGTAGATGAAATCGTGAATAAATGTGTTTCATTTCTCTATCATTATAGAAAATAAGAT-CATATTATATTACTTATGGAATTCGAACCCGAACTCGATTTATGATTTATTATTTCGATCTTATTGGCCTTTGTTCTTTATTTTTGATTTTTCATATAGATTTCCGTCTTTATATTATACCCTTTCGTGGATGAATTATGCCTATTTTCACATCTAGGATTTACATATACAACATATATTACTGTCAAGAGGGAATTTTTCTCAGTATTTAGATATTTAGATTCAAAATAAGAAAGGGATCAATTCAATTATAAACCTGCAAAACAAAGATTAGGATTGGGTTGGGTTGCGCTATATATATCAAAGAGTATACAATAATGATGTATTTGGTGAATCAAATACATGGTCTAATAACAAACCATTTTAACATAACATTTTAATGAGTTGATAATATTAATTGAATATTGAATATTTTTTGAAAGATTTTTGTTAAAGGTTTCATTCATGCCTAATCCATATCGAGTAGACCTTGTTGTTGTGAG

			>Puya_wrightii
			??????????????????????????????????????????????????????????????????????????????????????????????????????????????????????????????????????????????????????????????????????????????????????????????????????????????????????????????????????????????????????????????????????????????????????????????????????????????????????????????????????????????????????????????????????????????????????????????????????????????????????????????????????????????????????????????????????????????????????????????????????????????????????????????????????????????????????????????????????????????????????????????????????????????????????????????????????????????????????????????????????????????????????????????????????????????????????????????????????????????????????????????????????????????????????????????

			>Puya_yakespala
			??????????????????????????????????????????????????????????????????????????????????????????????????????????????????????????????????????????????????????????????????????????????????????????????????????????????????????????????????????????????????????????????????????????????????????????????????????????????????????????????????????????????????????????????????????????????????????????????????????????????????????????????????????????????????????????????????????????????????????????????????????????????????????????????????????????????????????????????????????????????????????????????????????????????????????????????????????????????????????????????????????????????????????????????????????????????????????????????????????????????????????????????????????????????????????????????

			>Quesnelia_arvensis
			???????????????????????GTCRAAATCCTTTTTGGAATAATACCAAATCGAAAATAAATGTCCGATAGCAAGTTGATCAGTTAATTCAATAAGAGATAAATGGGAGATAGCACTCGATTTAGTTGGTACCACCCAACCGAATCCGATTCACTCGTTTACTCATTCAATGAGTAAATTTTCAAGTTCAGCCAATCTTTTTTTTTT-GAAAAAAAAATTGCAAGTAGATGAAATCGTGAATAAATGTGTTTCATTTCTATATCATTATATAAAAAAAGATCCATATTATATTACTTATGGAATTCGAACCCGAACTCGATTTATGATTTATTATTTGGATCTTATTGGCCTTTGTTCTTTATTTTTGATTTTTCATATGGATTTCCGTCTTTATATTATACCCTTTCGTGGATGAATTATGCCTATTTTCACATCTAGGATTTACATATACAACATATATTACTGTCAAGAGGGAATTTTTCTCAGTATTTAGATATTTAGATTCAAAATAAGAAAGGGATCAATTCAATTATAAACCTGCAAAACAAAGATTAGGATTGGGTTGGGTTGCGCTATATATATCAAAGAGTATACAATAATGATGTATTTGGTGAATCAAATACATGGTCTAATAACAAACCATTTTAACATAACATTTTAATGAGTTGATAATATTAATTGAATATTGAATATTTTTTGAAAGATTTTTGTTAAAGGTTTCATTCATGCC?????????????????????????????????

			>Quesnelia_conquistensis
			??????????????????????????????????????????????????????????????????????????????????????????????????????????????????????????????????????????????????????????????????????????????????????????????????????????????????????????????????????????????????????????????????????????????????????????????????????????????????????????????????????????????????????????????????????????????????????????????????????????????????????????????????????????????????????????????????????????????????????????????????????????????????????????????????????????????????????????????????????????????????????????????????????????????????????????????????????????????????????????????????????????????????????????????????????????????????????????????????????????????????????????????????????????????????????????????

			>Quesnelia_edmundoi
			ATAATTATAATAAAGTGAAATATGTCGAAATCCTTTTTGGAATAATACCAAATCGAAAATAAATGTCCGATAGCAAGTTGATCAGTTAATTCAATAAGAGATAAATGGGAGATAGCACTCGATTTAGTTGGTACCACCCAACCGAATCCGATTCACTCGTTTACTCATTCAATGAGTAAATTTTCAAGTTCAGCCAATC-TTTTTTTTTTAAAAAAAAGATTGCAAGTAGATGAAATCGTGAATAAATGTGTTTCATTTCTATATCATTATATAAAAAAAGATCCATATTATATTACTTATGGAATTCGAACCCGAACTCGATTTATGATTTATTATTTGGATCTTATTGGCCTTTGCTCTTTATTTTTGATTTTTCATATAGATTTCCATCTTTATATTATACCCTTTCGTGGATGAATTATGCCTATTTTCACATCTAGGATTTACATATACAACATATATTACTGTCAAGAGGGAATTTTTCTCAGTATTTAGATATTTAGATTCAAAATAAGAAAGGGATCAATTCAATTATAAACCTGCAAAACAAAGATTAGGATTGGGTTGGGTTGCGCTATATATATCAAAGAGTATACAATAATGATGTATTTGGTGAATCAAATACATGGTCTAATAACAAACCATTTTAACATAACATTTTAATGAGTTGATAATATTAATTGAATATTGAATATTTTTTGAAAGATTTTTGTTAAAGGTTTCATTCATGCCTAATCCATATCGAGTAGACCTTGTTGTTGTG??

			>Quesnelia_edmundoi_var._rubrobracteata
			??????????????????????????????????????????????????????????????????????????????????????????????????????????????????????????????????????????????????????????????????????????????????????????????????????????????????????????????????????????????????????????????????????????????????????????????????????????????????????????????????????????????????????????????????????????????????????????????????????????????????????????????????????????????????????????????????????????????????????????????????????????????????????????????????????????????????????????????????????????????????????????????????????????????????????????????????????????????????????????????????????????????????????????????????????????????????????????????????????????????????????????????????????????????????????????????

			>Quesnelia_humilis
			??????????????????????????????????????????????????????????????????????????????????????????????????????????????????????????????????????????????????????????????????????????????????????????????????????????????????????????????????????????????????????????????????????????????????????????????????????????????????????????????????????????????????????????????????????????????????????????????????????????????????????????????????????????????????????????????????????????????????????????????????????????????????????????????????????????????????????????????????????????????????????????????????????????????????????????????????????????????????????????????????????????????????????????????????????????????????????????????????????????????????????????????????????????????????????????????

			>Quesnelia_kautskyi
			??????????????????????????????????????????????????????????????????????????????????????????????????????????????????????????????????????????????????????????????????????????????????????????????????????????????????????????????????????????????????????????????????????????????????????????????????????????????????????????????????????????????????????????????????????????????????????????????????????????????????????????????????????????????????????????????????????????????????????????????????????????????????????????????????????????????????????????????????????????????????????????????????????????????????????????????????????????????????????????????????????????????????????????????????????????????????????????????????????????????????????????????????????????????????????????????

			>Quesnelia_koltesii
			??????????????????????????????????????????????????????????????????????????????????????????????????????????????????????????????????????????????????????????????????????????????????????????????????????????????????????????????????????????????????????????????????????????????????????????????????????????????????????????????????????????????????????????????????????????????????????????????????????????????????????????????????????????????????????????????????????????????????????????????????????????????????????????????????????????????????????????????????????????????????????????????????????????????????????????????????????????????????????????????????????????????????????????????????????????????????????????????????????????????????????????????????????????????????????????????

			>Quesnelia_lateralis
			ATAATTATAATAAAGTTAAATATGTCGAAATCCTTTTTGGAATAATACCAAATCGAAAATAAATGTCCGATAGCAAGTTGATCAGTTAATTCAATAAGAGATAAATGGGAGATAGCACTCGATTTAGTTGGTACCACCCAACCGAATCCGATTCACTCGTTTACTCATTGAATGAGTAAATTTTCAAGTTCAGCCAATC-TTTTTTTTTGAAAAAATAAATTGCAAGTAGATGAAATCGTGAATAAATGTGTTTCATTTCTATATCATTATATAAAAAAAGATCCATATTATATTACTTATGGAATTCGAACCCGAACTCGATTTATGATTTATTATTTGGATCTTATTGGCCTTTGTTCTTTATTTTTGATTTTTCATATAGATTTCCGTCTTTATATTATACCCTTTCGTGGATGAATTATGCCTATTTTCACATCTAGGATTTACATATACAACATATATTACTGTCAAGAGGGAATTTTTCTCAGTATTTAGATATTTAGATTCAAAATAAGAAAGGGATCAATTCAATTATAAACCTGCAAAACAAAGATTAGGATTGGGTTGGGTTGCGCTATATATATCAAAGAGTATACAATAATGATGTATTTGGTGAATCAAATACATGGTCTAATAACAAACCATTTTAACATAACATTTTAATGAGTTGATAATATTAATTGAATATTGAATATTTTATGAAAGATTTTTGTTAAAGGTTTCATTCATGCCTAATCCATATCGAGTAGACCTTGTTGTTGTG??

			>Quesnelia_liboniana
			ATAATTATAATAAAGTGAAATATGTCGAAATCCTTTTTGGAATAATACCAAATCGAAAATAAATGTCCGATAGCAAGTTGATCAGTTAATTCAATAAGAGATAAATGGGAGATAGCACTCGATTTAGTTGGTACCACCCAGCCGAATCCGATTCACTCGTTTACTCATTGAATGAGTAAATTTTCAAGTTCAGCCAATC---TTTTTTTTAAAAAAGAAATTGCAAGTAGATGAAATCGTGAATAAATGTGTTTCATTTCTATATCATTATATAAAAAAAGATCCATATTATATTACTTATGGAATTCGAACCCGAACTCGATTTATGATTTATTATTTGGATCTTATTGGCCTTTGTTCTTTATTTTTGATTTTTCATATAGATTTCCGTCTTTATATTATACCCTTTCGTGGATGAATTATGCCTATTTTCACATCTAGGATTTACATATACAACATATATTACTGTCAAGAGGGAATTTTTCTCAGTATTTAGATATTTAGATTCAAAATAAGAAAGGGATCAATTCAATTATAAACCTGCAAAACAAAGATTAGGATTGGGTTGGGTTGCGCTATATATATCAAAGAGTATACAATAATGATGTATTTGGTGAATCAAATACATGGTCTAATAACAAACCATTTTAACATAACATTTTAATGAGTTGATAATATTAATTGAATATTGAATATTTTTTGAAAGATTTTTGTTAAAGGTTTCATTCATGCCTAATCCATATCGAGTAGACCTTGTTGTTGTG??

			>Quesnelia_marmorata
			??????????????????????????????????????????????????????????????????????????????????????????????????????????????????????????????????????????????????????????????????????????????????????????????????????????????????????????????????????????????????????????????????????????????????????????????????????????????????????????????????????????????????????????????????????????????????????????????????????????????????????????????????????????????????????????????????????????????????????????????????????????????????????????????????????????????????????????????????????????????????????????????????????????????????????????????????????????????????????????????????????????????????????????????????????????????????????????????????????????????????????????????????????????????????????????????

			>Quesnelia_quesneliana
			ATAATTATAATAAAGTGAAATATGTCGAAATCCTTTTTGGAATAATACCAAATCGAAAATAAATGTCCGATAGCAAGTTGATCAGTTAATTCAATAAGAGATAAATGGGAGATAGCACTCGATTTAGTTGGTACCACCCAACCGAATCCGATTCACTCGTTTACTCATTCAATGAGTAAATTTTCAAGTTCAGCCAATC-TTTTTTTTTGAAAAAAAAAATTGCAAGTAGATGAAATCGTGAATAAATGTGTTTCATTTCTATATCATTATATAAAAAAAGATCCATATTATATTACTTATGGAATTCGAACCCGAACTCGATTTATGATTTATTATTTGGATCTTATTGGCCTTTGTTCTTTATTTTTGATTTTTCATATGGATTTCCGTCTTTATATTATACCCTTTCGTGGATGAATTATGCCTATTTTCACATCTAGGATTTACATATACAACATATATTACTGTCAAGAGGGAATTTTTCTCAGTATTTAGATATTTAGATTCAAAATAAGAAAGGGATCAATTCAATTATAAACCTGCAAAACAAAGATTAGGATTGGGTTGGGTTGCGCTATATATATCAAAGAGTATACAATAATGATGTATTTGGTGAATCAAATACATGGTCTAATAACAAACCATTTTAACATAACATTTTAATGAGTTGATAATATTAATTGAATATTGAATATTTTTTGAAAGATTTTTGTTAAAGGTTTCATTCATGCCTAATCCATATCGAGTAGACCTTGTTGTTGTGAG

			>Quesnelia_strobilispica
			??????????????????????????????????????????????????????????????????????????????????????????????????????????????????????????????????????????????????????????????????????????????????????????????????????????????????????????????????????????????????????????????????????????????????????????????????????????????????????????????????????????????????????????????????????????????????????????????????????????????????????????????????????????????????????????????????????????????????????????????????????????????????????????????????????????????????????????????????????????????????????????????????????????????????????????????????????????????????????????????????????????????????????????????????????????????????????????????????????????????????????????????????????????????????????????????

			>Quesnelia_testudo
			??????????????????????????????????????????????????????????????????????????????????????????????????????????????????????????????????????????????????????????????????????????????????????????????????????????????????????????????????????????????????????????????????????????????????????????????????????????????????????????????????????????????????????????????????????????????????????????????????????????????????????????????????????????????????????????????????????????????????????????????????????????????????????????????????????????????????????????????????????????????????????????????????????????????????????????????????????????????????????????????????????????????????????????????????????????????????????????????????????????????????????????????????????????????????????????????

			>Quesnelia_tubifolia
			??????????????????????????????????????????????????????????????????????????????????????????????????????????????????????????????????????????????????????????????????????????????????????????????????????????????????????????????????????????????????????????????????????????????????????????????????????????????????????????????????????????????????????????????????????????????????????????????????????????????????????????????????????????????????????????????????????????????????????????????????????????????????????????????????????????????????????????????????????????????????????????????????????????????????????????????????????????????????????????????????????????????????????????????????????????????????????????????????????????????????????????????????????????????????????????????

			>Racinaea_cuspidata
			??????????????????????????????????????????????????????????????????????????????????????????????????????????????????????????????????????????????????????????????????????????????????????????????????????????????????????????????????????????????????????????????????????????????????????????????????????????????????????????????????????????????????????????????????????????????????????????????????????????????????????????????????????????????????????????????????????????????????????????????????????????????????????????????????????????????????????????????????????????????????????????????????????????????????????????????????????????????????????????????????????????????????????????????????????????????????????????????????????????????????????????????????????????????????????????????

			>Racinaea_dyeriana
			??????????????????????????????????????????????????????????????????????????????????????????????????????????????????????????????????????????????????????????????????????????????????????????????????????????????????????????????????????????????????????????????????????????????????????????????????????????????????????????????????????????????????????????????????????????????????????????????????????????????????????????????????????????????????????????????????????????????????????????????????????????????????????????????????????????????????????????????????????????????????????????????????????????????????????????????????????????????????????????????????????????????????????????????????????????????????????????????????????????????????????????????????????????????????????????????

			>Racinaea_elegans
			ATAATTATAATAAAGTGAAATATGTCGAAATCCTTTTTGGAATAATACCAAATCGAAAATAAATGTCCGATAGCAAGTTGATCAGTTAATTCAATAAGAGATAAATGGGAGATAGCACTCGATTTAGTTGGTACCACCCAACCGAATCCGATTCAATCGTTTACTCA------GAGTAAATTTTCAAGTTCAGCCAATCTTTTTTTTT------------------------------------------------------------------------------------------------------------------------------------------------------------------------CATATGGATTTCCGTCTTTATATTATACCCTTTTGTAGATGAATTATGCTTATTTGCACATCTAGGATTTACATATACAACATATATTACTGTCAAGAGGTAATTTTTCTCAGTATTTAGATATTTAGATTCAAAATAAGAAAGGG-----TTCAATTATAAACCCGCAAAACAAAGATTAGGATTGGGTTGGGTTGCGCTATATATATCAAAGAGTATACAATAATGATGTATTTGGTGAATCAAATACATGGTCTAATAACAAACCATTTTAACATAACATTTTAATGAGTTGATAATATTAATTGAATATTGAATATTTTTTGAAAGATTCTTGTTAAAGGTTTCATTCATGCCTAATCCATATCGAGTAGACCTTGTTGTTGTGAG

			>Racinaea_fraseri
			??????????????????????????????????????????????????????????????????????????????????CAGTTAATTCAATAAGAGATAAATGGGAGATAGCACTCGATTTAGTTGGTACCACCCAACCGAATCCGATTCAATCGTTTACTCA------GAGTAAATTTTCAAGTTCAGCCAATC-TTTTTTTTT------------------------------------------------------------------------------------------------------------------------------------------------------------------------ATATGGATTTCCGTCTTTATATTATACCCTTTCGTAGATGAATTATGCTTATTTTCATATCTAGGATTTACATATACAACATATATTACTGTCAAGAGGGAATTTTTCTCAGTATTTAGATATTTGGATTCAAAATAAGAAAGGG-----TTCAATTATAAACCCGCAAAACAAAGATTAGGATTGGGTTGGGTTGCGCTATATATATCAAAGAGTATACAATAATGATGTATTTGGTGAATCAAATACATGGTCTAATAACAAACCATTTTAACATAACATTTTAATGAGTTGATAATATTAATTGAATATTGAATATTTTTTGAAAGATTTTTGTTAAAGGTTTCATTCATGCCTAATCCATATCGAGTAGACCTTGTTGTTGTGAG

			>Racinaea_hamaleana
			??????????????????????????????????????????????????????????????????????????????????????????????????????????????????????????????????????????????????????????????????????????????????????????????????????????????????????????????????????????????????????????????????????????????????????????????????????????????????????????????????????????????????????????????????????????????????????????????????????????????????????????????????????????????????????????????????????????????????????????????????????????????????????????????????????????????????????????????????????????????????????????????????????????????????????????????????????????????????????????????????????????????????????????????????????????????????????????????????????????????????????????????????????????????????????????????

			>Racinaea_multiflora
			??????????????????????????????????????????????????????????????????????????????????????????????????????????????????????????????????????????????????????????????????????????????????????????????????????????????????????????????????????????????????????????????????????????????????????????????????????????????????????????????????????????????????????????????????????????????????????????????????????????????????????????????????????????????????????????????????????????????????????????????????????????????????????????????????????????????????????????????????????????????????????????????????????????????????????????????????????????????????????????????????????????????????????????????????????????????????????????????????????????????????????????????????????????????????????????????

			>Racinaea_parviflora
			??????????????????????????????????????????????????????????????????????????????????????????????????????????????????????????????????????????????????????????????????????????????????????????????????????????????????????????????????????????????????????????????????????????????????????????????????????????????????????????????????????????????????????????????????????????????????????????????????????????????????????????????????????????????????????????????????????????????????????????????????????????????????????????????????????????????????????????????????????????????????????????????????????????????????????????????????????????????????????????????????????????????????????????????????????????????????????????????????????????????????????????????????????????????????????????????

			>Racinaea_pugiformis
			??????????????????????????????????????????????????????????????????????????????????CAGTTAATTCAATAAGAGATAAATGGGAGATAGCACTCGATTTAGTTGGTACCACCCAACCGAATCCGATTCAATCGTTTACTCAT------AGTAAATTTTCAAGTTCAGCCAATC-TTTTTTTTT-----------------------------------------------------------------------------------------------------------------------------------------------------------------------CGTATGGATTTCCGTCTTTATATTATACCCTTTCGTAGATGAATTATGCTTATTTTCACATCTAGGATTTACATATACAACATATATTACTGTCAAGAGGTAATTTTTCTCAGTATTTAGATATTTAGATTCAAAATAAGAAAGGG-----TTCAATTATAAACCCGCAAAACAAAGATTAGGATTGGGTTGGGTTGCGCTATATATATCAAAGAGTATACAATAATGATGTATTTGGTGAATCAAATACATGGTCTAATAACAAACCATTTTAACATAACATTTTAATGAGTTGATAATATTAATTGAATATTGAATATTTTTTGAAAGATTCTTGTTAAAGGTTTCATTCATGCCTAATCCATATCGAGTAGACCTTGTTGTTGTGAG

			>Racinaea_ropalocarpa
			ATAATTATAATAAAGTGAAATATGTCGAAATCCTTTTTGGAATAATACCAAATCAAAAATAAATGTCCGATAGCAAGTTGATCAGTTAATTCAATAAGAGATAAATGGGAGATAGCACTCGATTTAGTTGGTACCACCCAACCGAATCCGATTCAATCGTTTACTCA------GAGTAAATTTTCAAGTTCAGCCAATCTT-------------------TT-----------------------------------------------------------------------------------------------------------------------------------------------------TTTTTCATATGGATTTCCGTCTTTATATTATACTCTTTCGTAGATGAATTATGCTTATTTTCACATCTAGGATTTACATATACAATATATATTACTGTCAAGAGGTAATTTTTCTCAGTATTTAGATATTTAGATTCAAAATAAGAAAGGG-----TTCAATTATAAACCCGCAAAACAAAGATTAGGATTGGGTTGGGTTGCGCTATATATATCAAAGAGTATACAATAATGATGTATTTGGTGAATCAAATACATGGTCTAATAACAAACCATTTTAACATAACATTTTAATGGGTTGATAATATTAATTGAATATTGAATATTTTTTGAAAGATTCTTGTTAAAGGTTTCATTCATGCCTAATCCATATCGAGTAGACCTTGTTGTTGTGAG

			>Racinaea_rothschuhiana
			??????????????????????????????????????????????????????????????????????????????????????????????????????????????????????????????????????????????????????????????????????????????????????????????????????????????????????????????????????????????????????????????????????????????????????????????????????????????????????????????????????????????????????????????????????????????????????????????????????????????????????????????????????????????????????????????????????????????????????????????????????????????????????????????????????????????????????????????????????????????????????????????????????????????????????????????????????????????????????????????????????????????????????????????????????????????????????????????????????????????????????????????????????????????????????????????

			>Racinaea_schumanniana
			??????????????????????????????????????????????????????????????????????????????????????????????????????????????????????????????????????????????????????????????????????????????????????????????????????????????????????????????????????????????????????????????????????????????????????????????????????????????????????????????????????????????????????????????????????????????????????????????????????????????????????????????????????????????????????????????????????????????????????????????????????????????????????????????????????????????????????????????????????????????????????????????????????????????????????????????????????????????????????????????????????????????????????????????????????????????????????????????????????????????????????????????????????????????????????????????

			>Racinaea_seemannii
			ATAATTATAATAAAGTGAAATATGTCGAAATCCTTTTTGGAATAATACCAAATCGAAAATAAATGTCCGATAGCAAGTTGATCAGTTAATTCAATAAGAGATAAATGGGAGATAGCACTCGATTTAGTTGGTACCACCCAACCGAATCCGATTCAATCGTTTACTCA------GAGTAAATTTTCAAGTTCAGCCAATCTTTTTTTTT------------------------------------------------------------------------------------------------------------------------------------------------------------------------CATATGGATTTCCGTCTTTATATTATACCCTTTCGTAGATGAATTATGCTTATTTTCACATCTAGGATTTACATATACAACATATATTACTGTCAAGAGGTAATTTTTCTCAGTATTTAGATATTTAGATTCAAAATAAGAAAGGG-----TTCAATTATAAACCCGCAAAACAAAGATTAGGATTGGGTTGGGTTGCGCTATATATATCAAAGAGTATACAATAATGATGTATTTGGTGAATCAAATACATGGTCTAATAACAAACCATTTTAACATAACATTTTAATGAGTTGATAATATTAATTGAATATTGAATATTTTTTGAAAGATTCTTGTTAAAGGTTTCATTCATGCCTAATCCATATCGAGTAGACCTTGTTGTTGTGAG

			>Racinaea_sinuosa
			??????????????????????????????????????????????????????????????????????????????????????????????????????????????????????????????????????????????????????????????????????????????????????????????????????????????????????????????????????????????????????????????????????????????????????????????????????????????????????????????????????????????????????????????????????????????????????????????????????????????????????????????????????????????????????????????????????????????????????????????????????????????????????????????????????????????????????????????????????????????????????????????????????????????????????????????????????????????????????????????????????????????????????????????????????????????????????????????????????????????????????????????????????????????????????????????

			>Racinaea_spiculosa
			ATAATTATAATAAAGTGAAATATGTCGAAATCCTTTTTGGAATAATACCAAATCGAAAATAAATGTCCGATAGCAAGTTGATCAGTTAATTCAATAAGAGATAAATGGGAGATAGCACTTGATTTAGTTGGTACCACCCAACCGAATCCGATTCAATCGTTTACTCA------GAGTAAATTTTCAAGTTCAGCCAATCTTTTTTTTT------------------------------------------------------------------------------------------------------------------------------------------------------------------------CATATGGATTTCCGTCTTTATATTATACCCTTTCGTAGATGAATTATGCTTATTTTCACATCTAGGATTTACATATACAACATATATTACTGTCAAGAGGTAATTTTTCTCAGTATTTAGATATTTAGATTCAAAATAAGAAAGGG-----TTCAATTATAAACCCGCAAAACAAAGATTAGGATTGGGTTGGGTTGCGCTATATATATCAAAGAGTATACAATAATGATGTATTTGGTGAATCAAATACATGGTCTAATAACAAACCATTTTAACATAACATTTTAATGAGTTGATAATATTAATTGAATATTGAATATTTTTTGAAAGATTCTTGTTAAAGGTTTCATTCATGCCTAATCCATATCGAGTAGACCTTGTTGTTGTGAG

			>Racinaea_spiculosa_var._micrantha
			??????????????????????????????????????????????????????????????????????????????????????????????????????????????????????????????????????????????????????????????????????????????????????????????????????????????????????????????????????????????????????????????????????????????????????????????????????????????????????????????????????????????????????????????????????????????????????????????????????????????????????????????????????????????????????????????????????????????????????????????????????????????????????????????????????????????????????????????????????????????????????????????????????????????????????????????????????????????????????????????????????????????????????????????????????????????????????????????????????????????????????????????????????????????????????????????

			>Racinaea_venusta
			ATAATTATAATAAAGTGAAATATGTCGAAATCCTTTTTGGAATAATACCAAATCGAAAATAAATGTCCGATAGCAAGTTGATCAGTTAATTCAATAAGAGATAAATGGGAGATAGCACTCGATTTAGTTGGTACCACCCAACCGAATCCGATTCAATCGTTTACTCA------GAGTAAATTTTCAAGTTCAGCCAATCTTTTTTTTT------------------------------------------------------------------------------------------------------------------------------------------------------------------------CATATGGATTTCCGTCTTTATATTATACCCTTTCGTAGATGAATTATGCTTATTTTCACATCTAGGATTTACATATACAACATATATTACTGTCAAGAGGGAATTTTTCTCAGTATTTAGATATTTAGATTCAAAATAAGAAAGGG-----TTCAATTATAAACCCGCAAAACAAAGATTAGGATTGGGTTGGGTTGCGCTATATATATCAAAGAGTATACAATAATGATGTATTTGGTGAATCAAATACATGGTCTAATAACAAACCATTTTAACATAACATTTTAATGAGTTGATAATATTAATTGAATATTGAATATTTTTTGAAAGATTTTTGTTAAAGGTTTCATTCATGCCTAATCCATATCGAGTAGACCTTGTTGTTGTGAG

			>Rapatea_paludosa
			ATAATTATAATAAAGTGAAATATGTCGAAATTTTATTTTGAATAGTGCCACATCAAAAATAAATGTCCGATAACAAGTTGATCAGTTAATTCAATAAGAGATAAATGGGAAATCGCACTTGATTTGGTTGGTAACGCCCAACCAAATCCGACTCTATCGTTTACTTA----------GAATCTTCAAGTTAAGCCAATCTTTTTTTTTTTAAAAAAGGGATTTCAAGTAGATGAAATCTTGAGTAAATGTGTTTCA--------TTAT-----AGAAAAATTTTTCTATTAGATTACCTTTTGAGTTCGAACCAGAACTCGATTTATGTTTGATTATTTCGATCTCATTTCCCATTGTTCTTTA--TTTATTTTTTCATATTGATTT---TTTCCATCTTATACCTTTTCGTGGATGAGTTATGCCCATTTTCACATCTAGGATTTACATATACAACATATATTACTGTCAAGAGGGAA-TTTTCTTAGTATTTATATACTTAGATTTTAAATAAATAGGAAATAGATTCAAATATAAACTTGCAAAACAAAGATTAGGATTGGATTGGGTTGCGCTATATATATCAAAGAGTATACAATAATGATGTATTTGGTGAATCAAATACATGGTCTAATAAGGAACCATTTTAACATAACATTTTAATGAGTCGATAATATTAATTGAATA-------TTTTTAGAAAGATTTTTGTTAAAGGTTTCATTCACGCCTAATCCATATCGAGTAGACCTTGTCGTTGTGAC

			>Rokautskyia_exaltata
			??????????????????????????????????????????????????????????????????????????????????????????????????????????????????????????????????????????????????????????????????????????????????????????????????????????????????????????????????????????????????????????????????????????????????????????????????????????????????????????????????????????????????????????????????????????????????????????????????????????????????????????????????????????????????????????????????????????????????????????????????????????????????????????????????????????????????????????????????????????????????????????????????????????????????????????????????????????????????????????????????????????????????????????????????????????????????????????????????????????????????????????????????????????????????????????????

			>Rokautskyia_fernseeoides
			??????????????????????????????????????????????????????????????????????????????????????????????????????????????????????????????????????????????????????????????????????????????????????????????????????????????????????????????????????????????????????????????????????????????????????????????????????????????????????????????????????????????????????????????????????????????????????????????????????????????????????????????????????????????????????????????????????????????????????????????????????????????????????????????????????????????????????????????????????????????????????????????????????????????????????????????????????????????????????????????????????????????????????????????????????????????????????????????????????????????????????????????????????????????????????????????

			>Rokautskyia_latifolia
			??????????????????????????????????????????????????????????????????????????????????????????????????????????????????????????????????????????????????????????????????????????????????????????????????????????????????????????????????????????????????????????????????????????????????????????????????????????????????????????????????????????????????????????????????????????????????????????????????????????????????????????????????????????????????????????????????????????????????????????????????????????????????????????????????????????????????????????????????????????????????????????????????????????????????????????????????????????????????????????????????????????????????????????????????????????????????????????????????????????????????????????????????????????????????????????????

			>Rokautskyia_microglazioui
			??????????????????????????????????????????????????????????????????????????????????????????????????????????????????????????????????????????????????????????????????????????????????????????????????????????????????????????????????????????????????????????????????????????????????????????????????????????????????????????????????????????????????????????????????????????????????????????????????????????????????????????????????????????????????????????????????????????????????????????????????????????????????????????????????????????????????????????????????????????????????????????????????????????????????????????????????????????????????????????????????????????????????????????????????????????????????????????????????????????????????????????????????????????????????????????????

			>Rokautskyia_odoratissima
			????????????????????????????AATCCT-TTTTGGATAATACCAAATCGAAAATAAATGTCCAATAGCAAGTTGATCAGTTAATTCAATAAGAGATAAATGGGAGATAGCACTCGATTTAGTTGGTACCACCCAACCGAATCCGATTCACTCGTTTACTCATTCAATGAGTAAATTTTCAAGTTCAGCCAATCTTTTTTTTTTTAAAAAAAAAATTGCAAGTAGATGAAATCGTGAATAAATGTGTTTCATTT?TATATCATTATATAAAAAAAGATCCATATTATATTACTTATGGAATTCGAACCCGAACTCGATTTATGATTTATTATTTGGATCTTATTGGCCTTTGTTCTTTATTTTTGATTTTTAATATAGATTTCCGTCTTTATATTATACCCTTTCGTGGATGAATTATGCCTATTTTCACATCTAGGATTTACATATACAACATATATTACTGTCAAGAGGGAATTTTTCTCAGTATTTAGATATTTAGATTCAAAATAAGAAAGGGATCAATTCAATTATAAACCTGCAAAACAAAGATTAGGATTGGGTTGGGTTGCGCTATATATATCAAAGAGTATACAATAATGATGTATTTGGTGAATCAAATACATGGTCTAATAACAAACCATTTTAACATAACATTTTAATGAGTTGATAATATTAATTGAATATTGAATATTTTTTGAAAGATTTTTGTTAAAGGTTTCATTCATGCCTAATCCATATCGAGTAGACCTTGTTGTTGTG??

			>Rokautskyia_roberto-kautskyi
			??????????????????????????????????????????????????????????????????????????????????????????????????????????????????????????????????????????????????????????????????????????????????????????????????????????????????????????????????????????????????????????????????????????????????????????????????????????????????????????????????????????????????????????????????????????????????????????????????????????????????????????????????????????????????????????????????????????????????????????????????????????????????????????????????????????????????????????????????????????????????????????????????????????????????????????????????????????????????????????????????????????????????????????????????????????????????????????????????????????????????????????????????????????????????????????????

			>Rokautskyia_sanctaluciae
			??????????????????????????????????????????????????????????????????????????????????????????????????????????????????????????????????????????????????????????????????????????????????????????????????????????????????????????????????????????????????????????????????????????????????????????????????????????????????????????????????????????????????????????????????????????????????????????????????????????????????????????????????????????????????????????????????????????????????????????????????????????????????????????????????????????????????????????????????????????????????????????????????????????????????????????????????????????????????????????????????????????????????????????????????????????????????????????????????????????????????????????????????????????????????????????????

			>Rokautskyia_scaposa
			??????????????????????????????????????????????????????????????????????????????????????????????????????????????????????????????????????????????????????????????????????????????????????????????????????????????????????????????????????????????????????????????????????????????????????????????????????????????????????????????????????????????????????????????????????????????????????????????????????????????????????????????????????????????????????????????????????????????????????????????????????????????????????????????????????????????????????????????????????????????????????????????????????????????????????????????????????????????????????????????????????????????????????????????????????????????????????????????????????????????????????????????????????????????????????????????

			>Ronnbergia_aciculosa
			??????????????????????????????????????????????????????????????????????????????????????????????????????????????????????????????????????????????????????????????????????????????????????????????????????????????????????????????????????????????????????????????????????????????????????????????????????????????????????????????????????????????????????????????????????????????????????????????????????????????????????????????????????????????????????????????????????????????????????????????????????????????????????????????????????????????????????????????????????????????????????????????????????????????????????????????????????????????????????????????????????????????????????????????????????????????????????????????????????????????????????????????????????????????????????????????

			>Ronnbergia_allenii
			ATAATTATAATAAAGTGAAATATGTCGAAATCCTTTTTGGAATAATACCAAATCGAAAA-AAATGTCCGATAGCAAGTTGATCAGTTAATTCAATAAGAGATAAATGGGAGATAGCACTCGATTTAGTTGGTACCACCCAACCGAATCCGATTCACTCGTTTACTCATTGAATGAGTAAATTTTCAAGTTCAGCCAATCTTTTTTTTTTTAAAAAAAAAATTGCAAGTAGATGAAATCGTGAATAAATGTGTTTCATTTCTATATCATTATATAAAAAAAGATCCATATTATATTACTTATGGAATTCGAACCCGAACTCGATTTATGATTTATTATTTGGATCTTATTGGCCTTTGTTCTTTATTTTTTATTTTTCATATAGATTTCCGTCTTTATATTATACCCTTTCGTGGATGAATTATGCCTATTTTCACATCTAGGATTTACATATACAACATATATTACTGTCAAGAGGGAATTTTTCTCAGTATTTAGATATTTAGATTCAAAATAAGAAAGGGATCAATTCAATTATAAACCTGCAAAACAAAGATTAGGATTGGGTTGGGTTGCGCTATATATATCAAAGAGTATACAATAATGATGTATTTGGTGAATCAAATACATGGTCTAATAACAAACCATTTTAACATAACATTTTAATGAGTTGATAATATTAATTGAATATTGAATATTTTTTGAAAGATTTTTGTTAAAGGTTTCATTCATGCCTAATCCATATCGAGTAGACCTTGTTGTTGTG??

			>Ronnbergia_columbiana
			??????????????????????????????????????????????????????????????????????????????????????????????????????????????????????????????????????????????????????????????????????????????????????????????????????????????????????????????????????????????????????????????????????????????????????????????????????????????????????????????????????????????????????????????????????????????????????????????????????????????????????????????????????????????????????????????????????????????????????????????????????????????????????????????????????????????????????????????????????????????????????????????????????????????????????????????????????????????????????????????????????????????????????????????????????????????????????????????????????????????????????????????????????????????????????????????

			>Ronnbergia_deleonii
			??????????????????????????????????????????????????????????????????????????????????????????????????????????????????????????????????????????????????????????????????????????????????????????????????????????????????????????????????????????????????????????????????????????????????????????????????????????????????????????????????????????????????????????????????????????????????????????????????????????????????????????????????????????????????????????????????????????????????????????????????????????????????????????????????????????????????????????????????????????????????????????????????????????????????????????????????????????????????????????????????????????????????????????????????????????????????????????????????????????????????????????????????????????????????????????????

			>Ronnbergia_drakeana
			ATAATTATAATAAAGTGAAATATGTCGAAATCCTTTTTGGAATAATACCAAATCGAAAATAAATGTCCGATAGCAAGTTGATCAGTTAATTCAATAAGAGATAAATGGGAGATAGCACTCGATTTAGTTGGTACCACCCAACCGAATCCGATTCACTCGTTTACTCATTCAATGAGTAAATTTTCAAGTTCAGCCAATCTTTTTTTTTTTCAAAAAAGAATTGCAAGTAGATGAAATCGTGAATAAATGTGTTTCATTTCTATATCATTATATAAAAAAAGATCCATATTATATTACTTATGGAATTCGAACCCGAACTCGATTTATGATTTATTATTTGGATCTTATTGGCCTTTGTTCTTTATTTTTTATTTTTCATATAGATTTCCGTCTTTATATTATACCCTTTCGTGGATGAATTATGCCTATTTTCACATCTAGGATTTACATATACAACATATATTACTGTCAAGAGGGAATTTTTCTCAGTATTTAGATATTTAGATTCAAAATAAGAAAGGGATCAATTCAATTATAAACCTGCAAAACAAAGATTAGGATTGGGTTGGGTTGCGCTATATATATCAAAGAGTATACAATAATGATGTATTTGGTGAATCAAATACATGGTCTAATAACAAACCATTTTAACATAACATTTTAATGAGTTGATAATATTAATTGAATATTGAATATTTTTTGAAAGATTTTTGTTAAAGGTTTCATTCATGCCTAATCCATATCGAGTAGACCTTGTTGTTGTG??

			>Ronnbergia_explodens
			???????????????????????????????????TTTGGAATAATACCAAATCGAAAATAAATGTCCGATAGCAAGTTGATCAGTTAATTCAATAAGAGATAAATGGGAGATAGCACTCGATTTAGTTGGTACCACCCAACCGAATCCGATTCACTCGTTTACTCATTGAATGAGTAAATTTTCAAGTTCAGCCAATCTTTTTTTTTT-CAAAAAAAAATTGCAAGTAGATGAAATCGTGAATAAATGTGTTTCATTTCTATATCATTATATAAAAAAGGATCCATATTATATTACTTATGGAATTCGAACCCGAACTCGATTTATGATTTATTATTTGGATCTTATTGGCCTTTGTTCTTTATTTTTTATTTTTCATATAGATTTCCGTCTTTATATTATACCCTTTCGTGGATGAATTATGCCTATTTTCACATCTAGGATTTACATATACAACATATATTACTGTCAAGAGGGAATTTTTCTCAGTATTTAGATATTTAGATTCAAAATAAGAAAGGGATCAATTCAATTATAAACCTGCAAAACAAAGATTAGGATTGGGTTGGGTTGCGCTATATATATCAAAGAGTATACAATAATGATGTATTTGGTGAATCAAATACATGGTCTAATAACAAACCATTTTAACATAACATTTTAATGAGTTGATAATATTAATTGAATATTGAATATTTTTTGAAAGATTTTTGTTAAAGGTTTCATTCATGCCTAATCCATATCGAGTAGACCTTGTTGTTGTG??

			>Ronnbergia_fraseri
			??????????????????????????????????????????????????????????????????????????????????????????????????????????????????????????????????????????????????????????????????????????????????????????????????????????????????????????????????????????????????????????????????????????????????????????????????????????????????????????????????????????????????????????????????????????????????????????????????????????????????????????????????????????????????????????????????????????????????????????????????????????????????????????????????????????????????????????????????????????????????????????????????????????????????????????????????????????????????????????????????????????????????????????????????????????????????????????????????????????????????????????????????????????????????????????????

			>Ronnbergia_germinyana
			??????????????????????????????????????????????????????????????????????????????????????????????????????????????????????????????????????????????????????????????????????????????????????????????????????????????????????????????????????????????????????????????????????????????????????????????????????????????????????????????????????????????????????????????????????????????????????????????????????????????????????????????????????????????????????????????????????????????????????????????????????????????????????????????????????????????????????????????????????????????????????????????????????????????????????????????????????????????????????????????????????????????????????????????????????????????????????????????????????????????????????????????????????????????????????????????

			>Ronnbergia_hathewayi
			??????????????????????????????????????????????????????????????????????????????????????????????????????????????????????????????????????????????????????????????????????????????????????????????????????????????????????????????????????????????????????????????????????????????????????????????????????????????????????????????????????????????????????????????????????????????????????????????????????????????????????????????????????????????????????????????????????????????????????????????????????????????????????????????????????????????????????????????????????????????????????????????????????????????????????????????????????????????????????????????????????????????????????????????????????????????????????????????????????????????????????????????????????????????????????????????

			>Ronnbergia_involucrata
			??????????????????????????????????????????????????????????????????????????????????????????????????????????????????????????????????????????????????????????????????????????????????????????????????????????????????????????????????????????????????????????????????????????????????????????????????????????????????????????????????????????????????????????????????????????????????????????????????????????????????????????????????????????????????????????????????????????????????????????????????????????????????????????????????????????????????????????????????????????????????????????????????????????????????????????????????????????????????????????????????????????????????????????????????????????????????????????????????????????????????????????????????????????????????????????????

			>Ronnbergia_killipiana
			??????????????????????????????????????????????????????????????????????????????????????????????????????????????????????????????????????????????????????????????????????????????????????????????????????????????????????????????????????????????????????????????????????????????????????????????????????????????????????????????????????????????????????????????????????????????????????????????????????????????????????????????????????????????????????????????????????????????????????????????????????????????????????????????????????????????????????????????????????????????????????????????????????????????????????????????????????????????????????????????????????????????????????????????????????????????????????????????????????????????????????????????????????????????????????????????

			>Ronnbergia_maidifolia
			??????????????????????????????????????????????????????????????????????????????????????????????????????????????????????????????????????????????????????????????????????????????????????????????????????????????????????????????????????????????????????????????????????????????????????????????????????????????????????????????????????????????????????????????????????????????????????????????????????????????????????????????????????????????????????????????????????????????????????????????????????????????????????????????????????????????????????????????????????????????????????????????????????????????????????????????????????????????????????????????????????????????????????????????????????????????????????????????????????????????????????????????????????????????????????????????

			>Ronnbergia_morreniana
			??????????????????????????????????????????????????????????????????????????????????????????????????????????????????????????????????????????????????????????????????????????????????????????????????????????????????????????????????????????????????????????????????????????????????????????????????????????????????????????????????????????????????????????????????????????????????????????????????????????????????????????????????????????????????????????????????????????????????????????????????????????????????????????????????????????????????????????????????????????????????????????????????????????????????????????????????????????????????????????????????????????????????????????????????????????????????????????????????????????????????????????????????????????????????????????????

			>Ronnbergia_nidularioides
			??????????????????????????????????????????????????????????????????????????????????????????????????????????????????????????????????????????????????????????????????????????????????????????????????????????????????????????????????????????????????????????????????????????????????????????????????????????????????????????????????????????????????????????????????????????????????????????????????????????????????????????????????????????????????????????????????????????????????????????????????????????????????????????????????????????????????????????????????????????????????????????????????????????????????????????????????????????????????????????????????????????????????????????????????????????????????????????????????????????????????????????????????????????????????????????????

			>Ronnbergia_petersii
			??????????????????????????????????????????????????????????????????????????????????????????????????????????????????????????????????????????????????????????????????????????????????????????????????????????????????????????????????????????????????????????????????????????????????????????????????????????????????????????????????????????????????????????????????????????????????????????????????????????????????????????????????????????????????????????????????????????????????????????????????????????????????????????????????????????????????????????????????????????????????????????????????????????????????????????????????????????????????????????????????????????????????????????????????????????????????????????????????????????????????????????????????????????????????????????????

			>Ronnbergia_tonduzii
			??????????????????????????????????????????????????????????????????????????????????????????????????????????????????????????????????????????????????????????????????????????????????????????????????????????????????????????????????????????????????????????????????????????????????????????????????????????????????????????????????????????????????????????????????????????????????????????????????????????????????????????????????????????????????????????????????????????????????????????????????????????????????????????????????????????????????????????????????????????????????????????????????????????????????????????????????????????????????????????????????????????????????????????????????????????????????????????????????????????????????????????????????????????????????????????????

			>Ronnbergia_veitchii
			??????????????????????????????????????????????????????????????????????????????????????????????????????????????????????????????????????????????????????????????????????????????????????????????????????????????????????????????????????????????????????????????????????????????????????????????????????????????????????????????????????????????????????????????????????????????????????????????????????????????????????????????????????????????????????????????????????????????????????????????????????????????????????????????????????????????????????????????????????????????????????????????????????????????????????????????????????????????????????????????????????????????????????????????????????????????????????????????????????????????????????????????????????????????????????????????

			>Ronnbergia_weberbaueri
			??????????????????????????????????????????????????????????????????????????????????????????????????????????????????????????????????????????????????????????????????????????????????????????????????????????????????????????????????????????????????????????????????????????????????????????????????????????????????????????????????????????????????????????????????????????????????????????????????????????????????????????????????????????????????????????????????????????????????????????????????????????????????????????????????????????????????????????????????????????????????????????????????????????????????????????????????????????????????????????????????????????????????????????????????????????????????????????????????????????????????????????????????????????????????????????????

			>Ronnbergia_wuelfinghoffii
			??????????????????????????????????????????????????????????????????????????????????????????????????????????????????????????????????????????????????????????????????????????????????????????????????????????????????????????????????????????????????????????????????????????????????????????????????????????????????????????????????????????????????????????????????????????????????????????????????????????????????????????????????????????????????????????????????????????????????????????????????????????????????????????????????????????????????????????????????????????????????????????????????????????????????????????????????????????????????????????????????????????????????????????????????????????????????????????????????????????????????????????????????????????????????????????????

			>Sequencia_serrata
			ATAATTATAATAAAGTGAAATATGTCGAAATCCTTTTTGGAATAATACCAAATCGAAAATAAATGTCCGATAGCAAGTTGATCAGTTAATTCAATAAGAGATAAATGGGAGATAGCACTCGATTTAGTTGGTACCACCCAACCGAATCCGATTCAATTGTTTACTCATTCAATGAGTCAATTTTCAAGTTCAGCCAATCCTTTTTTTTTTAAAAAAAAAATTGCAAGTAGATGAAATCGTGAATAAATGTGTTTCATTTCTCTATCATTATAGAAAATAAGATCCATATTATATTACTTATGGAATTCGAACCCGAACTCGATTTATGATTCATTATTTCGATCTTATTGGCCTTTGTTCTTTATTTTTGATTTTGCATATAGATTTCCGTCTTTATATTATACCCTTTCGTGGATGAATTATGCCTATTTTCACATCTAGGATTTACATATACAACATATATTACTGTCAAGAGGGAATTTTCCTCAGTATTTAGATATTTAGATTCAAAATAAGAAAGAGATCAATTCTCTTATAAACCTGCAAAACAAAGATTAGGA-----TTGGGTTGCGCTATATATATCAAAGAGTATACAATAATGATGTATTTGGTGAATCAAATACATGGTCTAATAACAAACCATTTTAACATAACATTTTAATGAGTTGATAATATTAATTGAATATTAAATATTTTTTGAAAGATTTTTGTTAAAGGTTTCATTCATGCTTAATCCATATCGAGTAGACCTTGTTGTTGTGAG

			>Sincoraea_albopicta
			????????????????????????????????????TTTGGATAATACCAAATCG-AAATAAATGTCCGATAGCAAGTTGATCAGTTAATTCAATAAGAGATAAATGGGAGATAGCACTCGATTTAGTTGGTACCACCCAACCGAATCCGATTCACTCGTTTACTCATTCAATGAGTAAATTTTCAAGTTCAGCCAATCTTTTTTTTTT-AAAAAAAAAATTGCAAGTAGATGAAATCGTGAATAAATGTGTTTCATTTCTATATCATTATATAAAAAAAGATCCATATTATATTACTTATGGAATTCGAACCCGAACTCGATTTATGATTTATTATTTGGATCTTATTGGCCTTTGTTCTTTATTTTTGATTTTTCATATAGATTTCCGTCTTTATATTATACCCTTTCGTGGATGAATTATGCCTATTTTCACATCTAGGATTTACATATACAACATATATTACTGTCAAGAGGGAATTTTTCTCAGTATTTAGATATTTAGATTCAAAATAAGAAAGGGATCAATTCAATTATAAACCTGCAAAACAAAGATTAGGATTGGGTTGGGTTGCGCTATATATATCAAAGAGTATACAATAATGATGTATTTGGTGAATCAAATACATGGTCTAATAACAAACCATTTTAACATAACATTTTAATGAGTTGATAATATTAATTGAATATTGAATATTTTTTGAAAGATTTTTGTTAAAGGTTTCATTCATGCCTAATCCATATCGAGTAGACCTTGTTGTTGTG??

			>Sincoraea_burle-marxii
			??????????????????????????????????????????????????????????????????????????????????????????????????????????????????????????????????????????????????????????????????????????????????????????????????????????????????????????????????????????????????????????????????????????????????????????????????????????????????????????????????????????????????????????????????????????????????????????????????????????????????????????????????????????????????????????????????????????????????????????????????????????????????????????????????????????????????????????????????????????????????????????????????????????????????????????????????????????????????????????????????????????????????????????????????????????????????????????????????????????????????????????????????????????????????????????????

			>Sincoraea_hatschbachii
			??????????????????????????????????????????????????????????????????????????????????????????????????????????????????????????????????????????????????????????????????????????????????????????????????????????????????????????????????????????????????????????????????????????????????????????????????????????????????????????????????????????????????????????????????????????????????????????????????????????????????????????????????????????????????????????????????????????????????????????????????????????????????????????????????????????????????????????????????????????????????????????????????????????????????????????????????????????????????????????????????????????????????????????????????????????????????????????????????????????????????????????????????????????????????????????????

			>Sincoraea_heleniceae
			??????????????????????????????????????????????????????????????????????????????????????????????????????????????????????????????????????????????????????????????????????????????????????????????????????????????????????????????????????????????????????????????????????????????????????????????????????????????????????????????????????????????????????????????????????????????????????????????????????????????????????????????????????????????????????????????????????????????????????????????????????????????????????????????????????????????????????????????????????????????????????????????????????????????????????????????????????????????????????????????????????????????????????????????????????????????????????????????????????????????????????????????????????????????????????????????

			>Sincoraea_humilis
			??????????????????????????????????????????????????????????????????????????????????????????????????????????????????????????????????????????????????????????????????????????????????????????????????????????????????????????????????????????????????????????????????????????????????????????????????????????????????????????????????????????????????????????????????????????????????????????????????????????????????????????????????????????????????????????????????????????????????????????????????????????????????????????????????????????????????????????????????????????????????????????????????????????????????????????????????????????????????????????????????????????????????????????????????????????????????????????????????????????????????????????????????????????????????????????????

			>Sincoraea_mucugensis
			??????????????????????????????????????????????????????????????????????????????????????????????????????????????????????????????????????????????????????????????????????????????????????????????????????????????????????????????????????????????????????????????????????????????????????????????????????????????????????????????????????????????????????????????????????????????????????????????????????????????????????????????????????????????????????????????????????????????????????????????????????????????????????????????????????????????????????????????????????????????????????????????????????????????????????????????????????????????????????????????????????????????????????????????????????????????????????????????????????????????????????????????????????????????????????????????

			>Sincoraea_navioides
			??????????????????????????????????????????????????????????????????????????????????????????????????????????????????????????????????????????????????????????????????????????????????????????????????????????????????????????????????????????????????????????????????????????????????????????????????????????????????????????????????????????????????????????????????????????????????????????????????????????????????????????????????????????????????????????????????????????????????????????????????????????????????????????????????????????????????????????????????????????????????????????????????????????????????????????????????????????????????????????????????????????????????????????????????????????????????????????????????????????????????????????????????????????????????????????????

			>Sincoraea_ophiuroides
			??????????????????????????????????????????????????????????????????????????????????????????????????????????????????????????????????????????????????????????????????????????????????????????????????????????????????????????????????????????????????????????????????????????????????????????????????????????????????????????????????????????????????????????????????????????????????????????????????????????????????????????????????????????????????????????????????????????????????????????????????????????????????????????????????????????????????????????????????????????????????????????????????????????????????????????????????????????????????????????????????????????????????????????????????????????????????????????????????????????????????????????????????????????????????????????????

			>Sincoraea_ulei
			???????????????????????????????????TTTGGAATAATACCAAATCGAAAATAAATGTCCGATAGCAAGTTGATCAGTTAATTCAATAAGAGATAAATGGGAGATAGCACTCGATTTAGTTGGTACCACCCAACCGAATCCGATTCACTCGTTTACGCATTCAATGAGTAAATTTTCAAGTTCAGCCAATCTTTTTTTTTT-AAAAAAAAAATTGCAAGTAGATGAAATCGTGAATAAATGTGTTTCATTTCTATATCATTATATAAAAAAAGATCCATATTATATTACTTATGGAATTCGAACCCGAACTCGATTTATGATTTATTATTTGGATCTTATTGGCCTTTGTTCTTTATTTTTGATTTTTCATATAGATTTCCGTCTTTATATTATACCCTTTCGTGGATGAATTATGCCTATTTTCACATCTAGGATTTACATATACAACATATATTACTGTCAAGAGGGAATTTTTCTCAGTATTTAGATATTTAGATTCAAAATAAGAAAGGGATCAATTCAATTATAAACCTGCAAAACAAAGATTAGGATTGGGTTGGGTTGCGCTATATATATCAAAGAGTATACAATAATGATGTATTTGGTGAATCAAATACATGGTCTAATAACAAACCATTTTAACATAACATTTTAATGAGTTGATAATATTAATTGAATATTGAATATTTTTTGAAAGATTTTTGTTAAAGGTTTCATTCATGCCTAATCCATATCGAGTAGACCTTGTTGTTGTG??

			>Stegolepis_ligulata
			ATAATTATAATAAAGTGAAATATGGCGAAATTTTTTTTTGAATAGTGCCACATCAAAAAAAAATGTCCGATAGCAAGTTGATCAGTTAATTCAATAAGAGATAAATGGGAGATAGCACTTGATTTAGTTGGTACCGCCCAACCAAATCCGATTCAATCGTTTATTTATTCAA----TGAGTTTTCAAGTTCAGCCAATC--TTTTT----------------------------------------------------------------------------------------------------------------------------------------------------------------TTTTTTTTTTCATATTGATTT---TTTCCGTCTTATACCCTTTCGTGGATGAATTATGTCCATTTTCACATCTAGGATTTACATATACAATATATATTACTGTCAAGAGGGAA-TTTTCTTAGTATTTATATACTTAGATTTTAAATAAATAAGAAATGGATTCAAATATAAACTTGCAAAACAAAAATTAGGATTGGATTGGGTTGCGCTATATATATGAAAGAGTATACAATAATGATGTATTTGGTGAATCAAATACATGGCCTAATAAGGAACCATGTTAACATAACATTTTAATGAGTTGATAATATTAATTGAATA-------TTTTTTGAAAGATTTTTGTCAAAGGTTTCATTCACGCCTAATCCATATCGAGTAGACCTTGTCGTTGTGAC

			>Stegolepis_parvipetala
			ATAATTATAATAAAGTGAAATATGGCGAAATTTTTTTTTGAATAGTGCCATATCAAAAAAAAATGTCCGATAGCAAGTTGATCAGTTAATTCAATAAGAGATAAATGGGGGATAGCACTTGATTTAGTTGGTACCGCCCAACCAAATCCGATTCAATCGTTTATTTATTCAA----TGAGTTTTCAAGTTCAGCCAATC--TTTTTTTTTAAAAAAAAAATTTCAAGTAGATGAAATCTTGAGTAAATGTGTTTCATTTCTCTATCCTTATGGAAAATCCTATTTATATTAGATTACCTTTGGAGTTCGAACCAGAACTCGATTTATGTTTGATTATTTCGATTTCATTGTCCATTGTTCCTTATTTTTTTTTTTTCATATTGATTT---TTTCCGTCTTATACCCTTTCGTGGATGAATTATGTCCATTTTCACATCTAGGATTTACATATACAATATATATTACTGTCAAGAGGGAA-TTTTCTTAGTATTTATATACTTAGATTTAAAATAAATAAGAAATGGATTCAAATATAAACTTGCAAAACAAAAATTAGGATTGGATTGGGTTGCGCTATATATATGAAAGAGTATACAATAATGATGTATTTGGTGAATCAAATACATGGCCTAATAAGGAACCATGTTAACATAACATTTTAATGAGTTGATAATATTAATTGAATA-------TTTTTTGAAAGATTTTTGTCAAAGGTTTCATTCACGCCTAATCCATATCGAGTAGACCTTGTCGTTGTGAC

			>Stigmatodon_amadoi
			??????????????????????????????????????????????????????????????????????????????????????????????????????????????????????????????????????????????????????????????????????????????????????????????????????????????????????????????????????????????????????????????????????????????????????????????????????????????????????????????????????????????????????????????????????????????????????????????????????????????????????????????????????????????????????????????????????????????????????????????????????????????????????????????????????????????????????????????????????????????????????????????????????????????????????????????????????????????????????????????????????????????????????????????????????????????????????????????????????????????????????????????????????????????????????????????

			>Stigmatodon_apparicianus
			??????????????????????????????????????????????????????????????????????????????????????????????????????????????????????????????????????????????????????????????????????????????????????????????????????????????????????????????????????????????????????????????????????????????????????????????????????????????????????????????????????????????????????????????????????????????????????????????????????????????????????????????????????????????????????????????????????????????????????????????????????????????????????????????????????????????????????????????????????????????????????????????????????????????????????????????????????????????????????????????????????????????????????????????????????????????????????????????????????????????????????????????????????????????????????????????

			>Stigmatodon_belloi
			??????????????????????????????????????????????????????????????????????????????????????????????????????????????????????????????????????????????????????????????????????????????????????????????????????????????????????????????????????????????????????????????????????????????????????????????????????????????????????????????????????????????????????????????????????????????????????????????????????????????????????????????????????????????????????????????????????????????????????????????????????????????????????????????????????????????????????????????????????????????????????????????????????????????????????????????????????????????????????????????????????????????????????????????????????????????????????????????????????????????????????????????????????????????????????????????

			>Stigmatodon_bifidus
			??????????????????????????????????????????????????????????????????????????????????????????????????????????????????????????????????????????????????????????????????????????????????????????????????????????????????????????????????????????????????????????????????????????????????????????????????????????????????????????????????????????????????????????????????????????????????????????????????????????????????????????????????????????????????????????????????????????????????????????????????????????????????????????????????????????????????????????????????????????????????????????????????????????????????????????????????????????????????????????????????????????????????????????????????????????????????????????????????????????????????????????????????????????????????????????????

			>Stigmatodon_brassicoides
			??????????????????????????????????????????????????????????????????????????????????????????????????????????????????????????????????????????????????????????????????????????????????????????????????????????????????????????????????????????????????????????????????????????????????????????????????????????????????????????????????????????????????????????????????????????????????????????????????????????????????????????????????????????????????????????????????????????????????????????????????????????????????????????????????????????????????????????????????????????????????????????????????????????????????????????????????????????????????????????????????????????????????????????????????????????????????????????????????????????????????????????????????????????????????????????????

			>Stigmatodon_costae
			??????????????????????????????????????????????????????????????????????????????????????????????????????????????????????????????????????????????????????????????????????????????????????????????????????????????????????????????????????????????????????????????????????????????????????????????????????????????????????????????????????????????????????????????????????????????????????????????????????????????????????????????????????????????????????????????????????????????????????????????????????????????????????????????????????????????????????????????????????????????????????????????????????????????????????????????????????????????????????????????????????????????????????????????????????????????????????????????????????????????????????????????????????????????????????????????

			>Stigmatodon_croceanus
			??????????????????????????????????????????????????????????????????????????????????????????????????????????????????????????????????????????????????????????????????????????????????????????????????????????????????????????????????????????????????????????????????????????????????????????????????????????????????????????????????????????????????????????????????????????????????????????????????????????????????????????????????????????????????????????????????????????????????????????????????????????????????????????????????????????????????????????????????????????????????????????????????????????????????????????????????????????????????????????????????????????????????????????????????????????????????????????????????????????????????????????????????????????????????????????????

			>Stigmatodon_euclidianus
			??????????????????????????????????????????????????????????????????????????????????????????????????????????????????????????????????????????????????????????????????????????????????????????????????????????????????????????????????????????????????????????????????????????????????????????????????????????????????????????????????????????????????????????????????????????????????????????????????????????????????????????????????????????????????????????????????????????????????????????????????????????????????????????????????????????????????????????????????????????????????????????????????????????????????????????????????????????????????????????????????????????????????????????????????????????????????????????????????????????????????????????????????????????????????????????????

			>Stigmatodon_funebris
			??????????????????????????????????????????????????????????????????????????????????????????????????????????????????????????????????????????????????????????????????????????????????????????????????????????????????????????????????????????????????????????????????????????????????????????????????????????????????????????????????????????????????????????????????????????????????????????????????????????????????????????????????????????????????????????????????????????????????????????????????????????????????????????????????????????????????????????????????????????????????????????????????????????????????????????????????????????????????????????????????????????????????????????????????????????????????????????????????????????????????????????????????????????????????????????????

			>Stigmatodon_gastinianus
			??????????????????????????????????????????????????????????????????????????????????????????????????????????????????????????????????????????????????????????????????????????????????????????????????????????????????????????????????????????????????????????????????????????????????????????????????????????????????????????????????????????????????????????????????????????????????????????????????????????????????????????????????????????????????????????????????????????????????????????????????????????????????????????????????????????????????????????????????????????????????????????????????????????????????????????????????????????????????????????????????????????????????????????????????????????????????????????????????????????????????????????????????????????????????????????????

			>Stigmatodon_goniorachis
			??????????????????????????????????????????????????????????????????????????????????????????????????????????????????????????????????????????????????????????????????????????????????????????????????????????????????????????????????????????????????????????????????????????????????????????????????????????????????????????????????????????????????????????????????????????????????????????????????????????????????????????????????????????????????????????????????????????????????????????????????????????????????????????????????????????????????????????????????????????????????????????????????????????????????????????????????????????????????????????????????????????????????????????????????????????????????????????????????????????????????????????????????????????????????????????????

			>Stigmatodon_harrylutheri
			??????????????????????????????????????????????????????????????????????????????????????????????????????????????????????????????????????????????????????????????????????????????????????????????????????????????????????????????????????????????????????????????????????????????????????????????????????????????????????????????????????????????????????????????????????????????????????????????????????????????????????????????????????????????????????????????????????????????????????????????????????????????????????????????????????????????????????????????????????????????????????????????????????????????????????????????????????????????????????????????????????????????????????????????????????????????????????????????????????????????????????????????????????????????????????????????

			>Stigmatodon_multifoliatus
			??????????????????????????????????????????????????????????????????????????????????????????????????????????????????????????????????????????????????????????????????????????????????????????????????????????????????????????????????????????????????????????????????????????????????????????????????????????????????????????????????????????????????????????????????????????????????????????????????????????????????????????????????????????????????????????????????????????????????????????????????????????????????????????????????????????????????????????????????????????????????????????????????????????????????????????????????????????????????????????????????????????????????????????????????????????????????????????????????????????????????????????????????????????????????????????????

			>Stigmatodon_plurifolius
			??????????????????????????????????????????????????????????????????????????????????????????????????????????????????????????????????????????????????????????????????????????????????????????????????????????????????????????????????????????????????????????????????????????????????????????????????????????????????????????????????????????????????????????????????????????????????????????????????????????????????????????????????????????????????????????????????????????????????????????????????????????????????????????????????????????????????????????????????????????????????????????????????????????????????????????????????????????????????????????????????????????????????????????????????????????????????????????????????????????????????????????????????????????????????????????????

			>Stigmatodon_rosulatulus
			??????????????????????????????????????????????????????????????????????????????????????????????????????????????????????????????????????????????????????????????????????????????????????????????????????????????????????????????????????????????????????????????????????????????????????????????????????????????????????????????????????????????????????????????????????????????????????????????????????????????????????????????????????????????????????????????????????????????????????????????????????????????????????????????????????????????????????????????????????????????????????????????????????????????????????????????????????????????????????????????????????????????????????????????????????????????????????????????????????????????????????????????????????????????????????????????

			>Stigmatodon_sanctateresensis
			??????????????????????????????????????????????????????????????????????????????????????????????????????????????????????????????????????????????????????????????????????????????????????????????????????????????????????????????????????????????????????????????????????????????????????????????????????????????????????????????????????????????????????????????????????????????????????????????????????????????????????????????????????????????????????????????????????????????????????????????????????????????????????????????????????????????????????????????????????????????????????????????????????????????????????????????????????????????????????????????????????????????????????????????????????????????????????????????????????????????????????????????????????????????????????????????

			>Tillandsia_achyrostachys
			ATAATTATAATAAAGTGAAATATGTCGAAATCCTTTTTGGAATAATACCAAATCGAAAATAAATGTCCGATAGCAAGTTGATCAGTTAATTCAATAAGAGATAAATGGGAGATAGCACTCGATTTAGTTGGTACCGCCCAACCGAATCCGATTCAATCGTTTACTCATTCAATGAGTAAATTTTCAAGTTCAGCCAATCCTTTTTTTTTTCAAAAAAAAATTGCAAGTAAATGAAATCGTGAATAAATGTGTTTCATTTCTCTATCATTATAGAAAAAAAGATCCATATTATATTACTTATGGAATTCGAACCCGAACTCGATTTATGATTCATTATTTCGATCTTATTGGCCTTTGTTCTTTATTTTTTATTTTTCATATAGATTTCCGTCTTTATATTATACCCTTTCGTAGATGAATTATGCTTATTTTCACATCTAGGATTTACATATACAACATATATTACTGTCAAGAGGGAATTTTTCTCAGTATTTAGATATTTAGATTCAAAATAAGAAAGGGATCAATTCAATTATAAACCCGCAAAACAAAGATTAGGATTGGGTTGGGTTGCGCTATATATATCAAAGAGTATACAATAATGATGTATTTGGTGAATCAAATACATGGTCTAATAACAAACCATTTTAACATAACATTTTAATGAGTTGATAATATTAATTGAATATTGAATATTTTTTGAAAGATTTTTGTTAAAGGTTTCATTCATGCCTAATCCATATCGAGTAGACCTTGTTGTTGTGAG

			>Tillandsia_adpressiflora
			??????????????????????????????????????????????????????????????????????????????????????????????????????????????????????????????????????????????????????????????????????????????????????????????????????????????????????????????????????????????????????????????????????????????????????????????????????????????????????????????????????????????????????????????????????????????????????????????????????????????????????????????????????????????????????????????????????????????????????????????????????????????????????????????????????????????????????????????????????????????????????????????????????????????????????????????????????????????????????????????????????????????????????????????????????????????????????????????????????????????????????????????????????????????????????????????

			>Tillandsia_aeranthos
			ATAATTATAATAAAGTGAAATATGTCGAAATCCTTTTTGGAATAATACCAAATCGAAAATAAATGTCCGATAGCAAGTTGATCAGTTAATTCAATAAGAGATAAATGGGAGATAGCACTCGATTTAGTTGGTACCACCCAACCGAATCCGATTCAATCGTTTACTCATTCAATGAGTAAATTTTCAAGTTCAGCCAATCTTTTTTTTTTT------TA-ATTGCAAGTAAATGAAATCGTGAATAAATGTGTTTCATTTCTCTATCATTATAGAAAAAAAGATTCATATTATATTACTTATGGAATTCGAACCCGAACTCTATTTATGATTCATTATTTCGATCTTATTGGCCTTTGTTCTTTATTTTTTATTTTTCATATAGATTTCCGTCTTTATATTATACCCTTTCGTAGATGAATTATGCTTACTTTCACATCTAGGATTTACATATACAACATATATTGCTGTCAAGAGGGAATTTTTCTCAGTATTTAGATATTTAGATTCAAAATAAGAAAGGGATCAATTCAATTATAAACCCGCAAAACAAAGATTAGGATTGGGTTGGGTTGCGCTATATATATCAAAGAGTATACAATAATGATGTATTTGGTGAATCAAATACATGGTCTAATAACAAACCATTTTAACATAACATTTTAATGAGTTGATAATATTAATTGACTATTGAATATTTTTTGAAAGATTTTTGTTAAAGGTTTCATTCATGCCTAATCCATATCGAGTAGACCTTGTTGTTGTGAG

			>Tillandsia_afonsoana
			??????????????????????????????????????????????????????????????????????????????????????????????????????????????????????????????????????????????????????????????????????????????????????????????????????????????????????????????????????????????????????????????????????????????????????????????????????????????????????????????????????????????????????????????????????????????????????????????????????????????????????????????????????????????????????????????????????????????????????????????????????????????????????????????????????????????????????????????????????????????????????????????????????????????????????????????????????????????????????????????????????????????????????????????????????????????????????????????????????????????????????????????????????????????????????????????

			>Tillandsia_aizoides
			??????????????????????????????????????????????????????????????????????????????????????????????????????????????????????????????????????????????????????????????????????????????????????????????????????????????????????????????????????????????????????????????????????????????????????????????????????????????????????????????????????????????????????????????????????????????????????????????????????????????????????????????????????????????????????????????????????????????????????????????????????????????????????????????????????????????????????????????????????????????????????????????????????????????????????????????????????????????????????????????????????????????????????????????????????????????????????????????????????????????????????????????????????????????????????????????

			>Tillandsia_albertiana
			ATAATTATAATAAAGTGAAATATGTCGAAATCCTTTTTGGAATAATACCAAATCGAAAATAAATGTCCGATAGCAAGTTGATCAGTTAATTCAATAAGAGATAAATGGGAGATAGCACTCGATTTAGTTGGTACCACCCAACCGAATCCGATTCAATCGTTTACTCATTCAATGAGTAAATTTTCAAGTTCAGCCAATCTTTATTT---------------------------------------------------------------------------------------------------------------------------------------------------------------------TTTTTCATATAGATTTCCGTCTTTATATTATACCCTTTCGTAGATGAATTATGCTTATTTTCACATCTAGGATTTACATATACAACATATATTACTGTCAAGAGGGAATTTTTCTCAGTATTTAGATATTTAGATTCAAAATAAGAAAGGGATCAATTCAATTATAAACCCGCAAAACAAAGATTAGGATTGGGTTGGGTTGCGCTATATATATCAAAGAGTATACAATAATGATGTATTTGGTGAATCAAATACATGGTCTAATAACAAACCATTTTAACATAACATTTTAATGAGTTGATAATATTAATTGACTATTGAATATTTTTTGAAAGATTTTTGTTAAAGGTTTCATTCATGCCTAATCCATATCGAGTAGACCTTGTTGTTGTGAG

			>Tillandsia_albida
			??????????????????????????????????????????????????????????????????????????????????????????????????????????????????????????????????????????????????????????????????????????????????????????????????????????????????????????????????????????????????????????????????????????????????????????????????????????????????????????????????????????????????????????????????????????????????????????????????????????????????????????????????????????????????????????????????????????????????????????????????????????????????????????????????????????????????????????????????????????????????????????????????????????????????????????????????????????????????????????????????????????????????????????????????????????????????????????????????????????????????????????????????????????????????????????????

			>Tillandsia_andicola
			??????????????????????????????????????????????????????????????????????????????????????????????????????????????????????????????????????????????????????????????????????????????????????????????????????????????????????????????????????????????????????????????????????????????????????????????????????????????????????????????????????????????????????????????????????????????????????????????????????????????????????????????????????????????????????????????????????????????????????????????????????????????????????????????????????????????????????????????????????????????????????????????????????????????????????????????????????????????????????????????????????????????????????????????????????????????????????????????????????????????????????????????????????????????????????????????

			>Tillandsia_andrieuxii
			ATAATTATAATAAAGTGAAATATGTCGAAATCCTTTTTGGAATAATACCAAATCGAAAATAAATGTCCGATAGCAAGTTGATCAGTTAATTCAATAAGAGATAAATGGGAGATAGCACTCGATTTAGTTGGTACCGCCCAACCGAATCCGATTCAATCATTTACTCATTCAATGAGTAAATTTTCAAGTTCAGCCAATCTTTTTTTTTTCAAAAAAAAAATTGCAAGTAAATGAAATCGTGAATAAATGTGTTTCATTTCTCTATCATTATAGAAAAAAAGATCCATATTATATTACTTATGGAATTCGAACCCGAACTCGATTTATGATTCATTATTTCGATCTTATTGGCCTTTGTTCTTTATTTTTTATTTTTCATATAGATTTCCGTCTTTATATTATACCCTTTCGTAGATGAATTATGCTTATTTTCACATCTAGGATTTACATATACAACATATATTACTGTCAAGAGGGGATTTTTCTCAATATTTAGATATTTAGATTCAAAATAAGAAAGGGATCAATTCAATTATAAACCCGCAAAACAAAGATTAGGATTGGGTTGGGTTGCGCTATATATATCAAAGAGTATACAATAATGATGTATTTGGTGAATCAAATACATGGTCTAATAACAAACCATTTTAACATAACATTTTAATGAGTTGATAATATTAATTGAATATTGAATATTTTTTGAAAGATTTTTGTTAAAGGTTTCATTCATGCCTAATCCATATCGAGTAGACCTTGTTGTTGTGAG

			>Tillandsia_appenii
			ATAATTATAATAAAGTGAAATATGTCGAAATCCTTTTTGGAATAATACCAAATCGAAAATAAATGTCCGATAGCAAGTTGATCAGTTAATTCAATAAGAGATAAATGGGAGATAGCACTCGATTTAGTTGGTACCACCCAACCGAATCCGATTCAATCGTTTACTCATTCAATGAGTCAATTTTCAAGTTCAGCCAATCTTTTTTTTTTC-AAAAAAAAATTGCAAGTAAATGAAATCGTGAATAAATGTGTTTCATTTCTCTATCATTATAGAAAAAAAGATCCATATTATATTACTTATGGAATTCGAACCCGAACTCGATTTATGATTCATTATTTCGATCTTATTGGCCTTTGTTCTTTATTTTTTATTTTTCATATAGATTTCCGTCTTTATATTATACCCTTTCGTAGATGAATTATGCTTATTTTCACATCTAGGATTTACATATACAACATATATTACTGTCAAGAGGGAATTTTTCTCAGTATTTAGATATTTAGATTCAAAATAAGAAAGGGATCAATTCAATTATAAACCCGCAAAACAAAGATTAGGGTTGGGTTGGGTTGCGCTATATATATCAAAGAGTATACAATAATGATGTATTTGGTGAATCAAATACATGGTCTAATAACAAACCATTTTAACATAACATTTTAATGAGTTGATAATATTAATTGAATATTGAATATTTTTTGAAAGATTTTTGTTAAAGGTTTCATTCATGCCTAATCCATATCGAGTAGACCTTGTTGTTGTGAG

			>Tillandsia_araujei
			??????????????????????????????????????????????????????????????????????????????????????????????????????????????????????????????????????????????????????????????????????????????????????????????????????????????????????????????????????????????????????????????????????????????????????????????????????????????????????????????????????????????????????????????????????????????????????????????????????????????????????????????????????????????????????????????????????????????????????????????????????????????????????????????????????????????????????????????????????????????????????????????????????????????????????????????????????????????????????????????????????????????????????????????????????????????????????????????????????????????????????????????????????????????????????????????

			>Tillandsia_argentea
			??????????????????????????????????????????????????????????????????????????????????????????????????????????????????????????????????????????????????????????????????????????????????????????????????????????????????????????????????????????????????????????????????????????????????????????????????????????????????????????????????????????????????????????????????????????????????????????????????????????????????????????????????????????????????????????????????????????????????????????????????????????????????????????????????????????????????????????????????????????????????????????????????????????????????????????????????????????????????????????????????????????????????????????????????????????????????????????????????????????????????????????????????????????????????????????????

			>Tillandsia_argentina
			ATAATTATAATAAAGTGAAATATGTCGAAATCCTTTTTGGAATAATACCAAATCGAAAATAAATGTCCGATAGCAAGTTGATCAGTTAATTCAATAAGAGATAAATGGGAGATAGCACTCGATTTAGTTGGTACCACCCAACCGAATCCGATTCAATCGTTTACTCATTCAATGAGTAAATTTGCAAGTTCAGACAATCTTTATTTT--------------------------------------------------------------------------------------------------------------------------------------------------------------------TTTTTCATATAGATTTCCGTCTTTATATTATACCCTTTCATAGATGAATTATGCTTATTTTCACATCTAGGATTTACATATACAACATATATTACTGTCAAGAGGGAATTTTTCTCAGTATTTAGATATTTAGATTCAAAATAAGAAAGGGATCAATTCAATTATAAACCCGCAAAACAAAGATTAGGATTGGGTTGGGTTGCGCTATATATATCAAAGAGTATACAATAATGATGTATTTGGTGAATCAAATACATGGTCTAATAACAAACCATTTTAACATAACATTTTAATGAGTTGATAATATTAATTGACTATTGAATATTTTTTGAAAGATTTTTGTTAAAGGTTTCATTCATGCCTAATCCATATCGAGTAGACCTTGTTGTTGTGAG

			>Tillandsia_arroyoensis
			??????????????????????????????????????????????????????????????????????????????????????????????????????????????????????????????????????????????????????????????????????????????????????????????????????????????????????????????????????????????????????????????????????????????????????????????????????????????????????????????????????????????????????????????????????????????????????????????????????????????????????????????????????????????????????????????????????????????????????????????????????????????????????????????????????????????????????????????????????????????????????????????????????????????????????????????????????????????????????????????????????????????????????????????????????????????????????????????????????????????????????????????????????????????????????????????

			>Tillandsia_atenangoensis
			??????????????????????????????????????????????????????????????????????????????????????????????????????????????????????????????????????????????????????????????????????????????????????????????????????????????????????????????????????????????????????????????????????????????????????????????????????????????????????????????????????????????????????????????????????????????????????????????????????????????????????????????????????????????????????????????????????????????????????????????????????????????????????????????????????????????????????????????????????????????????????????????????????????????????????????????????????????????????????????????????????????????????????????????????????????????????????????????????????????????????????????????????????????????????????????????

			>Tillandsia_atroviolacea
			??????????????????????????????????????????????????????????????????????????????????????????????????????????????????????????????????????????????????????????????????????????????????????????????????????????????????????????????????????????????????????????????????????????????????????????????????????????????????????????????????????????????????????????????????????????????????????????????????????????????????????????????????????????????????????????????????????????????????????????????????????????????????????????????????????????????????????????????????????????????????????????????????????????????????????????????????????????????????????????????????????????????????????????????????????????????????????????????????????????????????????????????????????????????????????????????

			>Tillandsia_atroviridipetala
			??????????????????????????????????????????????????????????????????????????????????????????????????????????????????????????????????????????????????????????????????????????????????????????????????????????????????????????????????????????????????????????????????????????????????????????????????????????????????????????????????????????????????????????????????????????????????????????????????????????????????????????????????????????????????????????????????????????????????????????????????????????????????????????????????????????????????????????????????????????????????????????????????????????????????????????????????????????????????????????????????????????????????????????????????????????????????????????????????????????????????????????????????????????????????????????????

			>Tillandsia_aurea
			??????????????????????????????????????????????????????????????????????????????????????????????????????????????????????????????????????????????????????????????????????????????????????????????????????????????????????????????????????????????????????????????????????????????????????????????????????????????????????????????????????????????????????????????????????????????????????????????????????????????????????????????????????????????????????????????????????????????????????????????????????????????????????????????????????????????????????????????????????????????????????????????????????????????????????????????????????????????????????????????????????????????????????????????????????????????????????????????????????????????????????????????????????????????????????????????

			>Tillandsia_australis
			??????????????????????????????????????????????????????????????????????????????????????????????????????????????????????????????????????????????????????????????????????????????????????????????????????????????????????????????????????????????????????????????????????????????????????????????????????????????????????????????????????????????????????????????????????????????????????????????????????????????????????????????????????????????????????????????????????????????????????????????????????????????????????????????????????????????????????????????????????????????????????????????????????????????????????????????????????????????????????????????????????????????????????????????????????????????????????????????????????????????????????????????????????????????????????????????

			>Tillandsia_baguagrandensis
			??????????????????????????????????????????????????????????????????????????????????????????????????????????????????????????????????????????????????????????????????????????????????????????????????????????????????????????????????????????????????????????????????????????????????????????????????????????????????????????????????????????????????????????????????????????????????????????????????????????????????????????????????????????????????????????????????????????????????????????????????????????????????????????????????????????????????????????????????????????????????????????????????????????????????????????????????????????????????????????????????????????????????????????????????????????????????????????????????????????????????????????????????????????????????????????????

			>Tillandsia_baileyi
			??????????????????????????????????????????????????????????????????????????????????????????????????????????????????????????????????????????????????????????????????????????????????????????????????????????????????????????????????????????????????????????????????????????????????????????????????????????????????????????????????????????????????????????????????????????????????????????????????????????????????????????????????????????????????????????????????????????????????????????????????????????????????????????????????????????????????????????????????????????????????????????????????????????????????????????????????????????????????????????????????????????????????????????????????????????????????????????????????????????????????????????????????????????????????????????????

			>Tillandsia_balbisiana
			??????????????????????????????????????????????????????????????????????????????????????????????????????????????????????????????????????????????????????????????????????????????????????????????????????????????????????????????????????????????????????????????????????????????????????????????????????????????????????????????????????????????????????????????????????????????????????????????????????????????????????????????????????????????????????????????????????????????????????????????????????????????????????????????????????????????????????????????????????????????????????????????????????????????????????????????????????????????????????????????????????????????????????????????????????????????????????????????????????????????????????????????????????????????????????????????

			>Tillandsia_baliophylla
			ATAATTATAATAAAGTGAAATATGTCGAAATCCTTTTTGGAATAATACCAAATCGAAAATAAATGTCCGATAGCAAGTTGATCAGTTAATTCAATAAGAGATAAATGGGAGATAGCACTCGATTTAGTTGGTACCACCCAACCGAATCTGATTCAATCGTTTACTCATTCAATGAGTAAATTTTCAAGTTCAGCCAATCTTTTTTTTTT---------AATTGCAAGTAAATGAAATCGTGAATAAATGTGTTTCATTTCTCTATCATTATAGAAAAAAAGATCCATATTATATTACTTATGGAATTCGAACCCGAACTCGATTTATGATTCATTATTTCGATCTTATTGGCCTTTGTTCTTTATTTTTTATTTTTCATATAGATTTCCGTCTTTATATTATACCCTTTCGTAGATGAATTATGCTTATTTTCACATCTAGGATTTACATATACAACATATATTACTGTCAAGAGGGAATTTTTCTCAGTATTTAGATATTTAGATTCAAA---------GGATCAATTCAATTATAAACCCGCAAAACAAAGATTAGGATTGGGTTGGGTTGCGCTATATATATCAAAGAGTATACAATAATGATGTATTTGGTGAATCAAATACATGGTCTAATAACAAACCATTTTAACATAACATTTTAATGAGTTGATAATATTAATTGAATATTGAATATTTTTTGAAAGATTTTTGTTAAAGGTTTCATTCATGCCTAATCCATATCGAGTAGACCTTGTTGTTGTGAG

			>Tillandsia_balsasensis
			??????????????????????????????????????????????????????????????????????????????????????????????????????????????????????????????????????????????????????????????????????????????????????????????????????????????????????????????????????????????????????????????????????????????????????????????????????????????????????????????????????????????????????????????????????????????????????????????????????????????????????????????????????????????????????????????????????????????????????????????????????????????????????????????????????????????????????????????????????????????????????????????????????????????????????????????????????????????????????????????????????????????????????????????????????????????????????????????????????????????????????????????????????????????????????????????

			>Tillandsia_barclayana
			ATAATTATAATAAAGTGAAATATGTCGAAATCCTTTTTGGAATAATACCAAATCGAAAATAAATGTCCGATAGCAAGTTGATCAGTTAATTCAATAAGAGATAAATGGGAGATAGCACTCGATTTAGTTGGTACCACCCAACCGAATCCGATTCAATCGTTTACTCATTCAATGAGTCAATTTTCAAGTTCAGCCAATC-TTTTTTTTTC-AAAAAAAAATTGCAAGTAAATGAAATCGTGAATAAATGTGTTTCATTTCTCTATCATTATAGAAAAAAAGATCCATATTATATTACTTATGGAATTCGAACCCGAACTCGATTTATGATTCATTATTTCGATCTTATTGGCCTTTGTTCTTTATTTTTTATTTTTCATATAGATTTCCGTCTTTATATTATACCCTTTCGTAGATGAATTATGCTTATTTTCACATCTAGGATTTACATATACAACATATATTACTGTCAAGAGGGAATTTTTCTCAGTATTTAGATATTTAGATTCAAAATAAGAAAGGGATCAATTCAATTATAAACCCGCAAAACAAAGATTAGGGTTGGGTTGGGTTGCGCTATATATATCAAAGAGTATACAATAATGATGTATTTGGTGAATCAAATACATGGTCTAATAACAAACCATTTTAACATAACATTTTAATGAGTTGATAATATTAATTGAATATTGAATATTTTTTGAAAGATTTTTGTTAAAGGTTTCATTCATGCCTAATCCATATCGAGTAGACCTTGTTGTTGTGAG

			>Tillandsia_barfussii
			??????????????????????????????????????????????????????????????????????????????????????????????????????????????????????????????????????????????????????????????????????????????????????????????????????????????????????????????????????????????????????????????????????????????????????????????????????????????????????????????????????????????????????????????????????????????????????????????????????????????????????????????????????????????????????????????????????????????????????????????????????????????????????????????????????????????????????????????????????????????????????????????????????????????????????????????????????????????????????????????????????????????????????????????????????????????????????????????????????????????????????????????????????????????????????????????

			>Tillandsia_barthlottii
			ATAATTATAATAAAGTGAAATATGTCGAAATCCTTTTTGGAATAATACCAAATCGAAAATAAATGTCCGATAGCAAGTTGATCAGTTAATTCAATAAGAGATAAATGGGAGATAGCACTCGATTTAGTTGGTACCACCCAACCGAATCCGATTCAATCGTTTACTCATTCAATGAGTAAATTTTCAAGTTCAGCCAATCTTTTTTTTATT------TAAATTGCAAGTAAATGAAATCGTGAATAAATGTTTTTCATTTCTCTATCATTATAGAAAAAAAGATCCATATTATATTACTTATGGAATTCGAACCCGAACTCGATTTATGATTCATTATTTCGATCTTATTGGCCTTTGTTCTTTATTTTTTATTTTTCATATAGATTTCCGTCTTTATATTATACCCTTTCGTAGATGAATTATGCTTATTTTCACATCTAGGATTTACATATACAACATATATTACTGTCAAGAGGGAATTTTTCTCAGTATTTAGATATTTAGATTCAAAATAAGAAAGGGATCAATTCAATTATAAACCCGCAAAACAAAGATTAGGGTTGGGTTGGGTTGCGCTATATATATCAAAGAGTATACAATAATGATGTATTTGGTGAATCAAATACATGGTCTAATAACAAACCATTTTAACATAACATTTTAATGAGTTGATAATATTAATTGAATATTGAATATTTTTTGAAAGATTTTTGTTAAAGGTTTCATTCATGCCTAATCCATATCGAGTAGACCTTGTTGTTGTGAG

			>Tillandsia_bartramii
			??????????????????????????????????????????????????????????????????????????????????????????????????????????????????????????????????????????????????????????????????????????????????????????????????????????????????????????????????????????????????????????????????????????????????????????????????????????????????????????????????????????????????????????????????????????????????????????????????????????????????????????????????????????????????????????????????????????????????????????????????????????????????????????????????????????????????????????????????????????????????????????????????????????????????????????????????????????????????????????????????????????????????????????????????????????????????????????????????????????????????????????????????????????????????????????????

			>Tillandsia_bella
			??????????????????????????????????????????????????????????????????????????????????????????????????????????????????????????????????????????????????????????????????????????????????????????????????????????????????????????????????????????????????????????????????????????????????????????????????????????????????????????????????????????????????????????????????????????????????????????????????????????????????????????????????????????????????????????????????????????????????????????????????????????????????????????????????????????????????????????????????????????????????????????????????????????????????????????????????????????????????????????????????????????????????????????????????????????????????????????????????????????????????????????????????????????????????????????????

			>Tillandsia_belloensis
			??????????????????????????????????????????????????????????????????????????????????????????????????????????????????????????????????????????????????????????????????????????????????????????????????????????????????????????????????????????????????????????????????????????????????????????????????????????????????????????????????????????????????????????????????????????????????????????????????????????????????????????????????????????????????????????????????????????????????????????????????????????????????????????????????????????????????????????????????????????????????????????????????????????????????????????????????????????????????????????????????????????????????????????????????????????????????????????????????????????????????????????????????????????????????????????????

			>Tillandsia_bergeri
			ATAATTATAATAAAGTGAAATATGTCGAAATCCTTTTTGGAATAATACCAAATCGAAAATAAATGTCCGATAGCAAGTTGATCAGTTAATTCAATAAGAGATAAATGGGAGATAGCACTCGATTTAGTTGGTACCACCCAACCGAATCCGATTCAATCGTTTACTCATTGAATGAGTAAATTTTCAAGTTCAGCCAATCTTTTTTTTTTT------TT-ATTGCAAGTAAATGAAATCGTGAATAAATGTGTTTCATTTCTCTATCATTATAGAAAAAAAGATTCATATTATATTACTTATGGAATTCGAACCCGAACTCGATTTATGATTCATTATTTCGATCTTATTGGCCTTTGTTCTTTATTTTTTATTTTTCATATAGATTTCCGTCTTTATATTATACCCTTTCGTAGATGAATTATGCTTACTTTCACATCTAGGATTTACATATACAACATATATTACTGTCAAGAGGGAATTTTTCTCAGTATTTAGATATTTAGATTCAAAATAAGAAAGGGATCAATTCAATTATAAACCCGCAAAACAAAGATTAGGATTGGGTTGGGTTGCGCTATATATATCAAAGAGTATACAATAATGATGTATTTGGTGAATCAAATACATGGTCTAATAACAAACCATTTTAACATAACATTTTAATGAGTTGATAATATTAATTGACTTTTGAATATTTTTTGAAAGATTTTTGTTAAAGGTTTCATTCATGCCTAATCCATATCGAGTAGACCTTGTTGTTGTGAG

			>Tillandsia_bermejoensis
			ATAATTATAATAAAGTGAAATATGTCGAAATCCTTTTTGGAATAATACCAAATCGAAAATAAATGTCCGATAGCAAGTTGATCAGTTAATTCAATAAGAGATAAATGGGAGATAGCACTCGATTTAGTTGGTACCACCCAACCGAATCCGATTCAATCGTTTACTCATTCAATGAGTAAATTTTCAAGTTCAGCCAATCTTTATTTT--------------------------------------------------------------------------------------------------------------------------------------------------------------------TTTTTCATATATATTTCCGTCTTTATATTATACCCTTTCATAGATGAATTATGCTTATTTTCACATCTAGGATTTACATATACAACATATATTACTGTCAAGAGGGAATTTTTCTCAGTATTTAGATATTTAGATTCAAAATAAGAAAGGGATCAATTCAATTATAAACCCGCAAAACAAGGATTAGGATTGGGTTGGGTTGCGCTATATATATCAAAGAGTATACAATAATGATGTATTTGGTGAATCAAATACATGGTCTAATAACAAACCATTTTAACATAACATTTTAATGAGTTGATAATATTAATTGACTATTGAATATTTTTTGAAAGATTTTTGTTAAAGGTTTCATTCATGCCTAATCCATATCGAGTAGACCTTGTTGTTGTGAG

			>Tillandsia_biflora
			ATAATTATAATAAAGTGAAATATGTCGAAATCCTTTTTGGAATAATACCAAATCGAAAATAAATGTCCGATAGCAAGTTGATCAGTTAATTCAATAAGAGATAAATGGGAGATAGCACTCGATTTAGTTGGTACCACCCAACCGAATCCGATTCAATCGTTTACTCATTCAATGAGTAAATTTTCAAGTTCAGCCAATCTTTTTT----------------------------------------------------------------------------------------------------------------------------------------------------------------------TTTTTCATATAGATTTCCGTCTTTATATTATACCCTTTCGTAGATGAATTATGCTTATTTTCACATCTAGGATTTACATATACAACATATATTACTGTCAAGAGGGAATTTTTCTCAGTATTTAGATATTTAGATTCAAAATAAGAAAGGGATCAATTCAATTATAAACCCGCAAAACAAAGATTAGGATTGGGTTGGGTTGCGCTATATATATCAAAGAGTATACAATAATGATGTATTTGGTGAATCAAATACATGGTCTAATAACAAACCATTTTAACATAACATTTTAATGAGTTGATAATATTA-------ATTGAATATTTTTTGAAAGATTTTTGTTAAAGGTTTCATTCATGCCTAATCCATATCGAGTAGACCTTGTTGTTGTGAG

			>Tillandsia_bonita
			??????????????????????????????????????????????????????????????????????????????????????????????????????????????????????????????????????????????????????????????????????????????????????????????????????????????????????????????????????????????????????????????????????????????????????????????????????????????????????????????????????????????????????????????????????????????????????????????????????????????????????????????????????????????????????????????????????????????????????????????????????????????????????????????????????????????????????????????????????????????????????????????????????????????????????????????????????????????????????????????????????????????????????????????????????????????????????????????????????????????????????????????????????????????????????????????

			>Tillandsia_botterii
			??????????????????????????????????????????????????????????????????????????????????????????????????????????????????????????????????????????????????????????????????????????????????????????????????????????????????????????????????????????????????????????????????????????????????????????????????????????????????????????????????????????????????????????????????????????????????????????????????????????????????????????????????????????????????????????????????????????????????????????????????????????????????????????????????????????????????????????????????????????????????????????????????????????????????????????????????????????????????????????????????????????????????????????????????????????????????????????????????????????????????????????????????????????????????????????????

			>Tillandsia_bourgaei
			??????????????????????????????????????????????????????????????????????????????????????????????????????????????????????????????????????????????????????????????????????????????????????????????????????????????????????????????????????????????????????????????????????????????????????????????????????????????????????????????????????????????????????????????????????????????????????????????????????????????????????????????????????????????????????????????????????????????????????????????????????????????????????????????????????????????????????????????????????????????????????????????????????????????????????????????????????????????????????????????????????????????????????????????????????????????????????????????????????????????????????????????????????????????????????????????

			>Tillandsia_brachycaulos
			??????????????????????????????????????????????????????????????????????????????????????????????????????????????????????????????????????????????????????????????????????????????????????????????????????????????????????????????????????????????????????????????????????????????????????????????????????????????????????????????????????????????????????????????????????????????????????????????????????????????????????????????????????????????????????????????????????????????????????????????????????????????????????????????????????????????????????????????????????????????????????????????????????????????????????????????????????????????????????????????????????????????????????????????????????????????????????????????????????????????????????????????????????????????????????????????

			>Tillandsia_brachyphylla
			ATAATTATAATAAAGTGAAATATGTCGAAATCCTTTTTGGAATAATACCAAATCGAAAATAAATGTCCGATAGCAAGTTGATCAGTTAATTCAATAAGAGATAAATGGGAGATAGCACTCGATTTAGTTGGTACCACCCAACCGAATCCGATTCAATCGTTTACTCATTGAATGAGTAAATTTTCAAGTTCAGTCAATCTTTTTTTTTTT--------AATTGCAAGTAAATGAAATCGTGAATAAATGTGTTTCATTTCTCTATCATTATAGAAAAAAAGATCCATATTATATTACTTATGGAATTCGAACCCGAACTCGATTTATGATTCATTATTTCGATCTTATTGGCCTTTGTTCTTTATTTTTTATTTTTCATATAGATTTCCGTCTTTATATTATACCCTTTCGTAGATGAATTATGCTTATTTTCACATCTAGGATTTACATATACAACATATATTACTGTCAAGAGGGAATTTTTCTCAGTATTTAGATATTTAGATTCAAAATAAGAAAGGGATCAATTCAATTATAAACCCGCAAAACAAAGATTAGGATTGGGTTGGGTTGCGCTATATATATCAAAGAGTATACAATAATGATGTATTTGGTGAATCAAATACATGGTCTAATAACAAACCATTTTAACATAACATTTTAATGAGTTGATAATATTAATTGAATATTGAATATTTTTTGAAAGATTTTTGTTAAAGGTTTCATTCATGCCTAATCCATATCGAGTAGACCTTGTTGTTGTGAG

			>Tillandsia_brevilingua
			ATAATTATAATAAAGTGAAATATGTCGAAATCCTTTTTGGAATAATACCAAATCAAAAATAAATGTCCGATAGCAAGTTGATCAGTTAATTCAATAAGAGATAAATGGGAGATAGCACTCGATTTAGTTGGTACCACCCAACCGAATCCGATTCAATCGTTTACTCATTCAATGAGTAAATTTTCAAGTTCAGCCAATCTTTTTTTTTTT--------AATTGCAAGTAAATGAAATCGTGAATAAATGTGTTTCATTTCTCTATCATTATAGAAAAAAAGATCCATATTATATTACTTATGGAATTAGAACCCGAACTCGATTTATGATTCATTATTTAGATCTTATTGGCCTTTGTTCTTTATTTTTTATTTTTCATATAGATTTTCGTCTTTATATTATACCCTTTCGTAGATGAATTATGCTTATTTTCACATCTAGGATTTACATATACAACATATATTACTGTCAAGAGGGAATTTTTCTCAGTATTTAGATATTTAGATTCAAAATAAGAAAGGGATCAATTCAATTATAAACCCGCAAAACAAAGATTAGGATTGGGTTGGGTTGCGCTATATATATCAAAGAGTATACAATAATGATGTATTTGGTGAATCAAATACATGGTCTAATAACAAACCATTTTAACATAACATTTTAATGAGTTGATAATATTAATTGAATATTGAATATTTTTTGAAAGATTTTTGTTAAAGGTTTCATTCATGCCTAATCCATATCGAGTAGACCTTGTTGTTGTGAG

			>Tillandsia_bulbosa
			??????????????????????????????????????????????????????????????????????????????????????????????????????????????????????????????????????????????????????????????????????????????????????????????????????????????????????????????????????????????????????????????????????????????????????????????????????????????????????????????????????????????????????????????????????????????????????????????????????????????????????????????????????????????????????????????????????????????????????????????????????????????????????????????????????????????????????????????????????????????????????????????????????????????????????????????????????????????????????????????????????????????????????????????????????????????????????????????????????????????????????????????????????????????????????????????

			>Tillandsia_burle-marxii
			??????????????????????????????????????????????????????????????????????????????????????????????????????????????????????????????????????????????????????????????????????????????????????????????????????????????????????????????????????????????????????????????????????????????????????????????????????????????????????????????????????????????????????????????????????????????????????????????????????????????????????????????????????????????????????????????????????????????????????????????????????????????????????????????????????????????????????????????????????????????????????????????????????????????????????????????????????????????????????????????????????????????????????????????????????????????????????????????????????????????????????????????????????????????????????????????

			>Tillandsia_butzii
			??????????????????????????????????????????????????????????????????????????????????????????????????????????????????????????????????????????????????????????????????????????????????????????????????????????????????????????????????????????????????????????????????????????????????????????????????????????????????????????????????????????????????????????????????????????????????????????????????????????????????????????????????????????????????????????????????????????????????????????????????????????????????????????????????????????????????????????????????????????????????????????????????????????????????????????????????????????????????????????????????????????????????????????????????????????????????????????????????????????????????????????????????????????????????????????????

			>Tillandsia_cacticola
			ATAATTATAATAAAGTGAAATATGTCGAAATCCTTTTTGGAATAATACCAAATCGAAAATAAATGTCCGATAGCAAGTTGATCAGTTAATTCAATAAGAGATAAATGGGAGATAGCACTCGATTTAGTTGGTACCACCCAACCGAATCCGATTCAATCGTTTACTCATTCAATGAGTAAATTTGCAAGTTCAGCCAATCTTTTTTTTCAA-AAAAAAAAATTGCAAGTAAATGAAATCGTGAATAAATGTGTTTCATTTCTCTATCATTATAGAAAAAAAGATCCATATTATATTACTTATGGAATTCGAACCCGAACTCGATTTATGATTCATTATTTCGATCTTATTGGCCTTTGTTCTTTATTTTTTATTTTTCATATAGATTTCCGTCTTTATATTATACCCTTTCGTAGATGAATTATGCTTATTTTCACATCTAGGATTTACATATACAACATATATTACTGTCAAGAGGGAATTTTTCTCAGTATTTAGATATTTAGATTCAAAATAAGAAAGGG-----TTCAATTATAAACCCGCAAAACAAAGATTAGGATTGGGTTGGGTTGCGCTATATATATCAAAGAGTATACAATAATGATGTATTTGGTGAATCAAATACATGGTCTAATAACAAACCATTTTAACATAACATTTTAATGAGTTGATAATATTAATTGAATATTGAATATTTTTTGAAAGATTTTTGTTAAAGGTTTCATTCATGCCTAATCCATATCGAGTAGACCTTGTTGTTGTGAG

			>Tillandsia_calcicola
			??????????????????????????????????????????????????????????????????????????????????????????????????????????????????????????????????????????????????????????????????????????????????????????????????????????????????????????????????????????????????????????????????????????????????????????????????????????????????????????????????????????????????????????????????????????????????????????????????????????????????????????????????????????????????????????????????????????????????????????????????????????????????????????????????????????????????????????????????????????????????????????????????????????????????????????????????????????????????????????????????????????????????????????????????????????????????????????????????????????????????????????????????????????????????????????????

			>Tillandsia_califanii
			ATAATTATAATAAAGTGAAATATGTCGAAATCCTTTTTGGAATAATACCAAATCGAAAATAAATGTCCGATAGCAAGTTGATCAGTTAATTCAATAAGAGATAAATGGGAGATAGCACTCGATTTAGTTGGTACCGCCCAACCGAATCCGATTCAATCGTTTACTCATTCAATGAGTAAATTTTCAAGTTCAGCCAATC--TTTTTTTTCAAAAAAAAAATTGCAAGTAAATGAAATCGTGAATAAATGTGTTTCATTTCTCTATCATTATAGAAAAAAAGATCCATATTATATTACTTATGGAATTCGAACCCGAACTCGATTTATGATTCATTATTTCGATCTTATTGGCCTTTGTTCTTTATTTTTTATTTTTCATATAGATTTCCGTCTTTATATTATACCCTTTCGTAGATGAATTATGCTTATTTTCACATCTAGGATTTACATATACAACATATATTACTGTCAAGAGGGAATTTTTCTCAGTATTTAGATATTTAGATTCAAAATAAGAAAGGGATCAATTCAATTATAAACCCGCAAAACAAAGATTAGGATTGGGTTGGGTTGCGCTATATATATCAAAGAGTATACAATAATGATGTATTTGGTGAATCAAATACATGGTCTAATAACAAACCATTTTAACATAACATTTTAATGAGTTGATAATATTAATTGAATATTGAATATTTTTTGAAAGATTTTTGTTAAAGGTTTCATTCATGCCTAATCCATATCGAGTAGACCTTGTTGTTGTGAG

			>Tillandsia_caliginosa
			??????????????????????????????????????????????????????????????????????????????????????????????????????????????????????????????????????????????????????????????????????????????????????????????????????????????????????????????????????????????????????????????????????????????????????????????????????????????????????????????????????????????????????????????????????????????????????????????????????????????????????????????????????????????????????????????????????????????????????????????????????????????????????????????????????????????????????????????????????????????????????????????????????????????????????????????????????????????????????????????????????????????????????????????????????????????????????????????????????????????????????????????????????????????????????????????

			>Tillandsia_calothyrsus
			??????????????????????????????????????????????????????????????????????????????????????????????????????????????????????????????????????????????????????????????????????????????????????????????????????????????????????????????????????????????????????????????????????????????????????????????????????????????????????????????????????????????????????????????????????????????????????????????????????????????????????????????????????????????????????????????????????????????????????????????????????????????????????????????????????????????????????????????????????????????????????????????????????????????????????????????????????????????????????????????????????????????????????????????????????????????????????????????????????????????????????????????????????????????????????????????

			>Tillandsia_caloura
			??????????????????????????????????????????????????????????????????????????????????????????????????????????????????????????????????????????????????????????????????????????????????????????????????????????????????????????????????????????????????????????????????????????????????????????????????????????????????????????????????????????????????????????????????????????????????????????????????????????????????????????????????????????????????????????????????????????????????????????????????????????????????????????????????????????????????????????????????????????????????????????????????????????????????????????????????????????????????????????????????????????????????????????????????????????????????????????????????????????????????????????????????????????????????????????????

			>Tillandsia_capillaris
			??????????????????????????????????????????????????????????????????????????????????????????????????????????????????????????????????????????????????????????????????????????????????????????????????????????????????????????????????????????????????????????????????????????????????????????????????????????????????????????????????????????????????????????????????????????????????????????????????????????????????????????????????????????????????????????????????????????????????????????????????????????????????????????????????????????????????????????????????????????????????????????????????????????????????????????????????????????????????????????????????????????????????????????????????????????????????????????????????????????????????????????????????????????????????????????????

			>Tillandsia_capillaris_f._virescens
			??????????????????????????????????????????????????????????????????????????????????????????????????????????????????????????????????????????????????????????????????????????????????????????????????????????????????????????????????????????????????????????????????????????????????????????????????????????????????????????????????????????????????????????????????????????????????????????????????????????????????????????????????????????????????????????????????????????????????????????????????????????????????????????????????????????????????????????????????????????????????????????????????????????????????????????????????????????????????????????????????????????????????????????????????????????????????????????????????????????????????????????????????????????????????????????????

			>Tillandsia_capitata
			??????????????????????????????????????????????????????????????????????????????????????????????????????????????????????????????????????????????????????????????????????????????????????????????????????????????????????????????????????????????????????????????????????????????????????????????????????????????????????????????????????????????????????????????????????????????????????????????????????????????????????????????????????????????????????????????????????????????????????????????????????????????????????????????????????????????????????????????????????????????????????????????????????????????????????????????????????????????????????????????????????????????????????????????????????????????????????????????????????????????????????????????????????????????????????????????

			>Tillandsia_caput-medusae
			ATAATTATAATAAAGTGAAATATGTCGAAATCCTTTTTGGAATAATACCAAATCGAAAATAAATGTCCGATAGCAAGTTGATCAGTTAATTCAATAAGAGATAAATGGGAGATAGCACTCGATTTAGTTGGTACCGCCCAACCGAATCCGATTCAATCGTTTACTCATTCAATGAGTAAATTTTCAAGTTCAGCCAATCTTTTTTTC---AAAAAAAAAATTGCAAGTAAATGAAATCGTGAATAAATGTGTTTCATTTCTCTATCATTATAGAAAAAAAGATCCATATTATATTACTTATGGAATTCGAACCCGAACTCGATTTATGATTCATTATTTCGATCTTATCGGCCTTTGTTCTTTATTTTTTATTTTTCATATAGATTTCCGTCTTTATATTATACCCTTTCGTAGATGAATTATGCTTATTTTCACATCTAGGATTTACATATACAACATATATTACTGTCAAGAGGGAATTTTTCTCAGTATTTAGATATTTAGATTCAAAATAAGAAAGGGATCAATTCAATTATAAACCCGCAAAACAAAGATTAGGATTGGGTTGGGTTGCGCTATATATATCAAAGAGTATACAATAATGATGTATTTGGTGAATCAAATACATGGTCTAATAACAAACCATTTTAACATAACATTTTAATGAGTTGATAATATTAATTGAATATTGAATATTTTTTGAAAGATTTTTGTTAAAGGTTTCATTCATGCCTAATCCATATCGAGTAGACCTTGTTGTTGTGAG

			>Tillandsia_cardenasii
			??????????????????????????????????????????????????????????????????????????????????????????????????????????????????????????????????????????????????????????????????????????????????????????????????????????????????????????????????????????????????????????????????????????????????????????????????????????????????????????????????????????????????????????????????????????????????????????????????????????????????????????????????????????????????????????????????????????????????????????????????????????????????????????????????????????????????????????????????????????????????????????????????????????????????????????????????????????????????????????????????????????????????????????????????????????????????????????????????????????????????????????????????????????????????????????????

			>Tillandsia_carlos-hankii
			ATAATTATAATAAAGTGAAATATGTCGAAATCCTTTTTGGAATAATACCAAATCGAAAATAAATGTCCGATAGCAAGTTGATCAGTTAATTCAATAAGAGATAAATGGGAGATAGCACTCGATTTAGTTGGTACCGCCCAACCGAATCCGATTCAATCGTTTACTCATTCAATGAGTAAATTTTCAAGTTCAGCCAATCTTTTTTTTTTCAAAAAAAAAATTGCAAGTAAATGAAATCGTGAATAAATGTGTTTCATTTCTCTATCATTATAGAAAAAAAGATCCATATTATATTACTTATGGAATTCGAACCCGAACTCGATTTATGATTCATTATTTCGATCTTATTGGCCTTTGTTCTTTATTTTTTATTTTTCATATAGATTTCCGTCTTTATATTATACCCTTTCGTAGATGAATTATGCTTATTTTCACATCTAGGATTTACATATACAACATATATTACTGTCAAGAGGGAATTTTTCTCAGTATTTAGATATTTAGATTCAAAATAAGAAAGGGATCAATTCAATTATAAACCCGCAAAACAAAGATTAGGATTGGGTTGGGTTGCGCTATATATATCAAAGAGTATACAATAATGATGTATTTGGTGAATCAAATACATGGTCTAATAACAAACCATTTTAACATAACATTTTAATGAGTTGATAATATTAATTGAATATTGAATATTTTTTGAAAGATTTTTGTTAAAGGTTTCATTCATGCCTAATCCATATCGAGTAGACCTTGTTGTTGTGAG

			>Tillandsia_carminea
			??????????????????????????????????????????????????????????????????????????????????????????????????????????????????????????????????????????????????????????????????????????????????????????????????????????????????????????????????????????????????????????????????????????????????????????????????????????????????????????????????????????????????????????????????????????????????????????????????????????????????????????????????????????????????????????????????????????????????????????????????????????????????????????????????????????????????????????????????????????????????????????????????????????????????????????????????????????????????????????????????????????????????????????????????????????????????????????????????????????????????????????????????????????????????????????????

			>Tillandsia_carnosa
			??????????????????????????????????????????????????????????????????????????????????????????????????????????????????????????????????????????????????????????????????????????????????????????????????????????????????????????????????????????????????????????????????????????????????????????????????????????????????????????????????????????????????????????????????????????????????????????????????????????????????????????????????????????????????????????????????????????????????????????????????????????????????????????????????????????????????????????????????????????????????????????????????????????????????????????????????????????????????????????????????????????????????????????????????????????????????????????????????????????????????????????????????????????????????????????????

			>Tillandsia_caulescens
			ATAATTATAATAAAGTGAAATATGTCGAAATCCTTTTTGGAATAATACCAAATCGAAAATAAATGTCCGATAGCAAGTTGATCAGTTAATTCAATAAGAGATAAATGGGAGATAGCACTCGATTTAGTTGGTACCACCCAACCGAATCCGATTCAATCGTTTACTCATTCAATGAGTAAATTTTCAAGTTCAGCCAATCTTTATTTT--------------------------------------------------------------------------------------------------------------------------------------------------------------------TTTTTCATATAGATTTCCGTCTTTATATTATACCCTTTCATAGATGAATTATGCTTATTTTCACATCTAGGATTTACATATACAACATATATTACTGTCAAGAGGGAATTTTTCTCAGTATTTAGATATTTAGATTCAAAATAAGAAAGGGATCAATTCAATTATAAACCCGCAAAACAAAGATTAGGATTGGGTTGGGTTGCGCTATATATATCAAAGAGTATACAATAATGATGTATTTGGTGAATCAAATACATGGTCTAATAACAAACCATTTTAACATAACATTTTAATGAGTTGATAATATTAATTGACTATTGAATATTTTTTGAAAGATTTTTGTTAAAGGTTTCATTCATGCCTAATCCATATCGAGTAGACCTTGTTGTTGTGAG

			>Tillandsia_cereicola
			??????????????????????????????????????????????????????????????????????????????????????????????????????????????????????????????????????????????????????????????????????????????????????????????????????????????????????????????????????????????????????????????????????????????????????????????????????????????????????????????????????????????????????????????????????????????????????????????????????????????????????????????????????????????????????????????????????????????????????????????????????????????????????????????????????????????????????????????????????????????????????????????????????????????????????????????????????????????????????????????????????????????????????????????????????????????????????????????????????????????????????????????????????????????????????????????

			>Tillandsia_chapeuensis
			??????????????????????????????????????????????????????????????????????????????????????????????????????????????????????????????????????????????????????????????????????????????????????????????????????????????????????????????????????????????????????????????????????????????????????????????????????????????????????????????????????????????????????????????????????????????????????????????????????????????????????????????????????????????????????????????????????????????????????????????????????????????????????????????????????????????????????????????????????????????????????????????????????????????????????????????????????????????????????????????????????????????????????????????????????????????????????????????????????????????????????????????????????????????????????????????

			>Tillandsia_chiletensis
			??????????????????????????????????????????????????????????????????????????????????????????????????????????????????????????????????????????????????????????????????????????????????????????????????????????????????????????????????????????????????????????????????????????????????????????????????????????????????????????????????????????????????????????????????????????????????????????????????????????????????????????????????????????????????????????????????????????????????????????????????????????????????????????????????????????????????????????????????????????????????????????????????????????????????????????????????????????????????????????????????????????????????????????????????????????????????????????????????????????????????????????????????????????????????????????????

			>Tillandsia_chlorophylla
			??????????????????????????????????????????????????????????????????????????????????????????????????????????????????????????????????????????????????????????????????????????????????????????????????????????????????????????????????????????????????????????????????????????????????????????????????????????????????????????????????????????????????????????????????????????????????????????????????????????????????????????????????????????????????????????????????????????????????????????????????????????????????????????????????????????????????????????????????????????????????????????????????????????????????????????????????????????????????????????????????????????????????????????????????????????????????????????????????????????????????????????????????????????????????????????????

			>Tillandsia_circinnatoides
			??????????????????????????????????????????????????????????????????????????????????????????????????????????????????????????????????????????????????????????????????????????????????????????????????????????????????????????????????????????????????????????????????????????????????????????????????????????????????????????????????????????????????????????????????????????????????????????????????????????????????????????????????????????????????????????????????????????????????????????????????????????????????????????????????????????????????????????????????????????????????????????????????????????????????????????????????????????????????????????????????????????????????????????????????????????????????????????????????????????????????????????????????????????????????????????????

			>Tillandsia_cochabambae
			??????????????????????????????????????????????????????????????????????????????????????????????????????????????????????????????????????????????????????????????????????????????????????????????????????????????????????????????????????????????????????????????????????????????????????????????????????????????????????????????????????????????????????????????????????????????????????????????????????????????????????????????????????????????????????????????????????????????????????????????????????????????????????????????????????????????????????????????????????????????????????????????????????????????????????????????????????????????????????????????????????????????????????????????????????????????????????????????????????????????????????????????????????????????????????????????

			>Tillandsia_coinaensis
			ATAATTATAATAAAGTGAAATATGTCGAAATCCTTTTTGGAATAATACCAAATCGAAAATAAATGTCCGATAGCAAGTTGATCAGTTAATTCAATAAGAGATAAATGGGAGATAGCACTCGATTTAGTTGGTACCACCCAACCGAATCCGATTCAATCGTTTACTCATTCAATGAGTAAATTTTAAAGTTCAGCCAATCTTTATTTT--------------------------------------------------------------------------------------------------------------------------------------------------------------------TTTTTCATATAGATTTCCGTCTTTATATTATACCCTTTCGTAGATGAATTATGCTTATTTTCACATCTAGGATTTACATATACAACATATATTACTGTCAAGAGGGAATTTTTCTCAGTATTTAGATATTTAGATTCAAAATAAGAAAGGGATCAATTCAATTATAAACCCGCAAAACAAAGATTAGGATTGGGTTGGGTTGCGCTATATATATCAAAGAGTATACAATAATGATGTATTTGGTGAATCAAATACATGGTCTAATAACAAACCATTTTAACATAACATTTTAATGAGTTGATAATATTAATTGAATATTGAATATTTTTTGAAAGATTTTTGTTAAAGGTTTCATTCATGCCTAATCCATATCGAGTAGACCTTGTTGTTGTGAG

			>Tillandsia_comarapaensis
			??????????????????????????????????????????????????????????????????????????????????????????????????????????????????????????????????????????????????????????????????????????????????????????????????????????????????????????????????????????????????????????????????????????????????????????????????????????????????????????????????????????????????????????????????????????????????????????????????????????????????????????????????????????????????????????????????????????????????????????????????????????????????????????????????????????????????????????????????????????????????????????????????????????????????????????????????????????????????????????????????????????????????????????????????????????????????????????????????????????????????????????????????????????????????????????????

			>Tillandsia_comitanensis
			??????????????????????????????????????????????????????????????????????????????????????????????????????????????????????????????????????????????????????????????????????????????????????????????????????????????????????????????????????????????????????????????????????????????????????????????????????????????????????????????????????????????????????????????????????????????????????????????????????????????????????????????????????????????????????????????????????????????????????????????????????????????????????????????????????????????????????????????????????????????????????????????????????????????????????????????????????????????????????????????????????????????????????????????????????????????????????????????????????????????????????????????????????????????????????????????

			>Tillandsia_complanata
			ATAATTATAATAAAGTGAAATATGTCGAAATCCTTTTTGGAATAATACCAAATC-AAAATAAATGTCCGATAGCAAGTTGATCAGTTAATTCAATAAGAGATAAATGGGAGATAGCACTCGATTTAGTTGGTACCACCCAACCGAATCCGATTCAATCGTTTACTCATTCAATGAGTAAATTTTCAAGTTCAGCCAATCCTTTATTTTT------------------------------------------------------------------------------------------------------------------------------------------------------------------TTTTTCATATAGATTTCCGTCTTTATATTATACCCTTTCGTAGATGAATTATGCTTATTTTCACATCTAGGATTTACATATACAACATATATTACTGTCAAGAGGGAATTTTTCTCAGTATTTAGATATTTAGATTCAAAATAAGAAAGGGATCAATTCAATTATAAACCCGCAAAACAAAGATTAGGATTGGGTTGGGTTGCGCTATATATATCAAAGAGTATACAATAATGATGTATTTGGTGAATCAAATACATGGTCTAATAACAAACCATTTTAACATAACATTTTAATGAGTTGATAATATTAATTGAATATTGAATATTTTTTGAAAGATTTTTGTTAAAGGTTTCATTCATGCCTAATCCATATCGAGTAGACCTTGTTGTTGTGAG

			>Tillandsia_comulcoensis
			??????????????????????????????????????????????????????????????????????????????????????????????????????????????????????????????????????????????????????????????????????????????????????????????????????????????????????????????????????????????????????????????????????????????????????????????????????????????????????????????????????????????????????????????????????????????????????????????????????????????????????????????????????????????????????????????????????????????????????????????????????????????????????????????????????????????????????????????????????????????????????????????????????????????????????????????????????????????????????????????????????????????????????????????????????????????????????????????????????????????????????????????????????????????????????????????

			>Tillandsia_concolor
			??????????????????????????????????????????????????????????????????????????????????????????????????????????????????????????????????????????????????????????????????????????????????????????????????????????????????????????????????????????????????????????????????????????????????????????????????????????????????????????????????????????????????????????????????????????????????????????????????????????????????????????????????????????????????????????????????????????????????????????????????????????????????????????????????????????????????????????????????????????????????????????????????????????????????????????????????????????????????????????????????????????????????????????????????????????????????????????????????????????????????????????????????????????????????????????????

			>Tillandsia_cossonii
			??????????????????????????????????????????????????????????????????????????????????????????????????????????????????????????????????????????????????????????????????????????????????????????????????????????????????????????????????????????????????????????????????????????????????????????????????????????????????????????????????????????????????????????????????????????????????????????????????????????????????????????????????????????????????????????????????????????????????????????????????????????????????????????????????????????????????????????????????????????????????????????????????????????????????????????????????????????????????????????????????????????????????????????????????????????????????????????????????????????????????????????????????????????????????????????????

			>Tillandsia_cryptantha
			??????????????????????????????????????????????????????????????????????????????????????????????????????????????????????????????????????????????????????????????????????????????????????????????????????????????????????????????????????????????????????????????????????????????????????????????????????????????????????????????????????????????????????????????????????????????????????????????????????????????????????????????????????????????????????????????????????????????????????????????????????????????????????????????????????????????????????????????????????????????????????????????????????????????????????????????????????????????????????????????????????????????????????????????????????????????????????????????????????????????????????????????????????????????????????????????

			>Tillandsia_cucaensis
			??????????????????????????????????????????????????????????????????????????????????????????????????????????????????????????????????????????????????????????????????????????????????????????????????????????????????????????????????????????????????????????????????????????????????????????????????????????????????????????????????????????????????????????????????????????????????????????????????????????????????????????????????????????????????????????????????????????????????????????????????????????????????????????????????????????????????????????????????????????????????????????????????????????????????????????????????????????????????????????????????????????????????????????????????????????????????????????????????????????????????????????????????????????????????????????????

			>Tillandsia_dasyliriifolia
			??????????????????????????????????????????????????????????????????????????????????????????????????????????????????????????????????????????????????????????????????????????????????????????????????????????????????????????????????????????????????????????????????????????????????????????????????????????????????????????????????????????????????????????????????????????????????????????????????????????????????????????????????????????????????????????????????????????????????????????????????????????????????????????????????????????????????????????????????????????????????????????????????????????????????????????????????????????????????????????????????????????????????????????????????????????????????????????????????????????????????????????????????????????????????????????????

			>Tillandsia_delicata
			??????????????????????????????????????????????????????????????????????????????????????????????????????????????????????????????????????????????????????????????????????????????????????????????????????????????????????????????????????????????????????????????????????????????????????????????????????????????????????????????????????????????????????????????????????????????????????????????????????????????????????????????????????????????????????????????????????????????????????????????????????????????????????????????????????????????????????????????????????????????????????????????????????????????????????????????????????????????????????????????????????????????????????????????????????????????????????????????????????????????????????????????????????????????????????????????

			>Tillandsia_demissa
			ATAATTATAATAAAGTGAAATATGTCGAAATCCTTTTTGGAATAATACCAAATCGAAAATAAATGTCCGATAGCAAGTTGATCAGTTAATTCAATAAGAGATAAATGGGAGATAGCACTCGATTTAGTTGGTACCACCCAACCGAATCCGATTCAATCGTTTACTCATTCAATGAGTAAATTTTCAAGTTCAGCCAATCTTTTTTTTTTT--------CATTGCAAGTAAATGAAATCGTGAATAAATGTGTTTCATTTCTCTATCATTATAGAAAAAAAGATCCATATTATATTACTTATGGAATTCGAACCCGAACTCGATTTATGATTCATTATTTCGATCTTATTGGCCTTTGTTCTTTATTTTTTATTTTTCATATAGATTTCCGTCTTTATATTATACCCTTTCGTAGATGAATTATGCTTATTTTCACATCTAGGATTTACATATACAACATATATTACTGTCAAGAGGGAATTTTTCTCAGTATTTAGATATTTAGATTCAAAATAAGAAAGGGATCAATTCAATTATAAACCCGCAAAACAAAGATTAGGATTGGGTTGGGTTGCGCTATATATATCAAAGAGTATACAATAATGATGTATTTGGTGAATCAAATACATGGTCTAATAACAAACCATTTTAACATAACATTTTAATGAGTTGATAATATTAATTGAATATTGAATATTTTTTGAAAGATTTTTGTTAAAGGTTTCATTCATGCCTAATCCATATCGAGTAGACCTTGTTGTTGTGAG

			>Tillandsia_denudata
			??????????????????????????????????????????????????????????????????????????????????????????????????????????????????????????????????????????????????????????????????????????????????????????????????????????????????????????????????????????????????????????????????????????????????????????????????????????????????????????????????????????????????????????????????????????????????????????????????????????????????????????????????????????????????????????????????????????????????????????????????????????????????????????????????????????????????????????????????????????????????????????????????????????????????????????????????????????????????????????????????????????????????????????????????????????????????????????????????????????????????????????????????????????????????????????????

			>Tillandsia_denudata_var._vivipara
			??????????????????????????????????????????????????????????????????????????????????????????????????????????????????????????????????????????????????????????????????????????????????????????????????????????????????????????????????????????????????????????????????????????????????????????????????????????????????????????????????????????????????????????????????????????????????????????????????????????????????????????????????????????????????????????????????????????????????????????????????????????????????????????????????????????????????????????????????????????????????????????????????????????????????????????????????????????????????????????????????????????????????????????????????????????????????????????????????????????????????????????????????????????????????????????????

			>Tillandsia_deppeana
			??????????????????????????????????????????????????????????????????????????????????????????????????????????????????????????????????????????????????????????????????????????????????????????????????????????????????????????????????????????????????????????????????????????????????????????????????????????????????????????????????????????????????????????????????????????????????????????????????????????????????????????????????????????????????????????????????????????????????????????????????????????????????????????????????????????????????????????????????????????????????????????????????????????????????????????????????????????????????????????????????????????????????????????????????????????????????????????????????????????????????????????????????????????????????????????????

			>Tillandsia_didisticha
			ATAATTATAATAAAGTGAAATATGTCGAAATCCTTTTTGGAATAATACCAAATCGAAAATAAATGTCCGATAGCAAGTTGATCAGTTAATTCAATAAGAGATAAATGGGAGATAGCACTCGATTTAGTTGGTACCACCCAACCGAATCCGATTCAATCGTTTACTCATTCAATGAGTAAATTTTCAAGTTCAGCCAATCTTTATTTTTT------------------------------------------------------------------------------------------------------------------------------------------------------------------TTTTTCATATAGATTTTCGTCTTTATATTATACCCTTTCATAGATGAATTATGCTTATTTTCACATCTAGGATTTACATATACAACATATATTACTGTCAAGAGGGAATTTTTCTCAGTATTTAGATATTTAGATTCAAAATAAGAAAGGGATCAATTCAATTATAAACCCGCAAAACAAAGATTAGGATTGGGTTGGGTTGCGCTATATATATCAAAGAGTATACAATAATGATGTATTTGGTGAATCAAATACATGGTCTAATAACAAACCATTTTAACATAACATTTTAATGAGTTGATAATATTAATTGACTATTGAATATTTTTTGAAAGATTTTTGTTAAAGGTTTCATTCATGCCTAATCCATATCGAGTAGACCTTGTTGTTGTGAG

			>Tillandsia_diguetii
			??????????????????????????????????????????????????????????????????????????????????????????????????????????????????????????????????????????????????????????????????????????????????????????????????????????????????????????????????????????????????????????????????????????????????????????????????????????????????????????????????????????????????????????????????????????????????????????????????????????????????????????????????????????????????????????????????????????????????????????????????????????????????????????????????????????????????????????????????????????????????????????????????????????????????????????????????????????????????????????????????????????????????????????????????????????????????????????????????????????????????????????????????????????????????????????????

			>Tillandsia_disticha
			ATAATTATAATAAAGTGAAATATGTCGAAATCCTTTTTGGAATAATACCAAATC-AAAATAAATGTCCGATAGCAAGTTGATCAGTTAATTCAATAAGAGATAAATGGGGGATAGCACTCGATTTAGTTGGTACCACCCAACCGAATCCGATTCAATCGTTTACTCATTCAATGAGTAAATTTTCAAGTTCAGCCAATCTTTTTTTTTTT--------AATGGCAAGTAAATGAAATCGTGAATAAATGTGTTTCATTTCTCTATCATTATAGAAAAAAGGATCCATATTATATTACTTATGGAATTCGAACCCGAACTCAATTTATGATTCATTATTTCGATCTTATTGGCCTTTGTTCTTTATTTTTTATTTTTCATATGGATTTCCGTCTTTATATTATACCCTTTCGTAGATGAATTATGCTTATTTTCACATCTAGGATTTACATATACAACATATATTACTGTCAAGAGGGAATTTTTCTCAGTATTTAGATATTTAGATTCAAAATAAGAAAGGGATCAATTCAATTATAAACCCGCAAAACAAAGATTAGGATTGGGATGGGTTGCGCTATATATATCAAAGAGTATACAATAATGATGTATTTGGTGAATCAAATACATGGTCTAATAACAAACCATTTTAACATAACATTTTAATGAGTTGAGAATATTAATTGAATATTGAATATTTTTTGAAAGATTTTTGTTAAAGGTTTCATTCATGCCTAATCCATATCGAGTAGACCTTGTTGTTGTGAG

			>Tillandsia_divaricata
			??????????????????????????????????????????????????????????????????????????????????????????????????????????????????????????????????????????????????????????????????????????????????????????????????????????????????????????????????????????????????????????????????????????????????????????????????????????????????????????????????????????????????????????????????????????????????????????????????????????????????????????????????????????????????????????????????????????????????????????????????????????????????????????????????????????????????????????????????????????????????????????????????????????????????????????????????????????????????????????????????????????????????????????????????????????????????????????????????????????????????????????????????????????????????????????????

			>Tillandsia_dugesii
			??????????????????????????????????????????????????????????????????????????????????????????????????????????????????????????????????????????????????????????????????????????????????????????????????????????????????????????????????????????????????????????????????????????????????????????????????????????????????????????????????????????????????????????????????????????????????????????????????????????????????????????????????????????????????????????????????????????????????????????????????????????????????????????????????????????????????????????????????????????????????????????????????????????????????????????????????????????????????????????????????????????????????????????????????????????????????????????????????????????????????????????????????????????????????????????????

			>Tillandsia_dura
			??????????????????????????????????????????????????????????????????????????????????????????????????????????????????????????????????????????????????????????????????????????????????????????????????????????????????????????????????????????????????????????????????????????????????????????????????????????????????????????????????????????????????????????????????????????????????????????????????????????????????????????????????????????????????????????????????????????????????????????????????????????????????????????????????????????????????????????????????????????????????????????????????????????????????????????????????????????????????????????????????????????????????????????????????????????????????????????????????????????????????????????????????????????????????????????????

			>Tillandsia_duratii
			ATAATTATAATAAAGTGAAATATGTCGAAATCCTTTTTGGAATAATACCAAATCGAAAATAAATGTCCGATAGCAAGTTGATCAGTTAATTCAATAAGAGATAAATGGGAGATAGCACTCGATTTAGTTGGTACCACCCAACCGAATCCGATTCAATCGTTTACTCATTCAATGAGTAAATTTTCAAGTTCAGCCAATCTTTATTTTTTT--------AATTGCAAGTAAATGAAATCGTGAATAAATGTGTTTCATTTCTCTATCATTATAGAAAAAAGGATCCATATTATATTACTTATGGAATTCGAACCCGAACTCGATTTATGATTCATTATTTCGATCTTATTGGCCTTTGTTCTTTATTTTTTATTTTTCATATAGATTTCCGTCTTTATATTATACCCTTTCGTAGATGAATTATGCTTATTTTCACATCTAGGATTTACATATACAACATATATTACTGTCAAGAGGGAATTTTTCTCAGTATTTAGATATTTAGATTCAAAATAAGAAAGGGATCAATTCAATTATAAACCCGCAAAACAAAGATTAGGATTGGGTTGGGTTGCGCTATATATATCAAAGAGTATACAATAATGATGTATTTGGTGAATCAAATACATGGTCTAATAACAAACCATTTTAACATAACATTTTAATGAGTTGATAATATTAATTGACTATTGAATATTTTTTGAAAGATTTTTGTTAAAGGTTTCATTCATGCCTAATCCATATCGAGTAGACCTTGTTGTTGTGAG

			>Tillandsia_ecarinata
			??????????????????????????????????????????????????????????????????????????????????????????????????????????????????????????????????????????????????????????????????????????????????????????????????????????????????????????????????????????????????????????????????????????????????????????????????????????????????????????????????????????????????????????????????????????????????????????????????????????????????????????????????????????????????????????????????????????????????????????????????????????????????????????????????????????????????????????????????????????????????????????????????????????????????????????????????????????????????????????????????????????????????????????????????????????????????????????????????????????????????????????????????????????????????????????????

			>Tillandsia_edithae
			??????????????????????????????????????????????????????????????????????????????????????????????????????????????????????????????????????????????????????????????????????????????????????????????????????????????????????????????????????????????????????????????????????????????????????????????????????????????????????????????????????????????????????????????????????????????????????????????????????????????????????????????????????????????????????????????????????????????????????????????????????????????????????????????????????????????????????????????????????????????????????????????????????????????????????????????????????????????????????????????????????????????????????????????????????????????????????????????????????????????????????????????????????????????????????????????

			>Tillandsia_ehlersiana
			??????????????????????????????????????????????????????????????????????????????????????????????????????????????????????????????????????????????????????????????????????????????????????????????????????????????????????????????????????????????????????????????????????????????????????????????????????????????????????????????????????????????????????????????????????????????????????????????????????????????????????????????????????????????????????????????????????????????????????????????????????????????????????????????????????????????????????????????????????????????????????????????????????????????????????????????????????????????????????????????????????????????????????????????????????????????????????????????????????????????????????????????????????????????????????????????

			>Tillandsia_eizii
			??????????????????????????????????????????????????????????????????????????????????????????????????????????????????????????????????????????????????????????????????????????????????????????????????????????????????????????????????????????????????????????????????????????????????????????????????????????????????????????????????????????????????????????????????????????????????????????????????????????????????????????????????????????????????????????????????????????????????????????????????????????????????????????????????????????????????????????????????????????????????????????????????????????????????????????????????????????????????????????????????????????????????????????????????????????????????????????????????????????????????????????????????????????????????????????????

			>Tillandsia_elongata
			??????????????????????????????????????????????????????????????????????????????????????????????????????????????????????????????????????????????????????????????????????????????????????????????????????????????????????????????????????????????????????????????????????????????????????????????????????????????????????????????????????????????????????????????????????????????????????????????????????????????????????????????????????????????????????????????????????????????????????????????????????????????????????????????????????????????????????????????????????????????????????????????????????????????????????????????????????????????????????????????????????????????????????????????????????????????????????????????????????????????????????????????????????????????????????????????

			>Tillandsia_elusiva
			??????????????????????????????????????????????????????????????????????????????????????????????????????????????????????????????????????????????????????????????????????????????????????????????????????????????????????????????????????????????????????????????????????????????????????????????????????????????????????????????????????????????????????????????????????????????????????????????????????????????????????????????????????????????????????????????????????????????????????????????????????????????????????????????????????????????????????????????????????????????????????????????????????????????????????????????????????????????????????????????????????????????????????????????????????????????????????????????????????????????????????????????????????????????????????????????

			>Tillandsia_erubescens
			??????????????????????????????????????????????????????????????????????????????????????????????????????????????????????????????????????????????????????????????????????????????????????????????????????????????????????????????????????????????????????????????????????????????????????????????????????????????????????????????????????????????????????????????????????????????????????????????????????????????????????????????????????????????????????????????????????????????????????????????????????????????????????????????????????????????????????????????????????????????????????????????????????????????????????????????????????????????????????????????????????????????????????????????????????????????????????????????????????????????????????????????????????????????????????????????

			>Tillandsia_espinosae
			ATAATTATAATAAAGTGAAATATGTCGAAATCCTTTTTGGAATAATACCAAATCGAAAATAAATGTCCGATAGCAAGTTGATCAGTTAATTCAATAAGAGATAAATGGGAGATAGCACTCGATTTAGTTGGTACCGCCCAACCGAATCCGATTCAATCGTTTACTCATTCAATGAGTAAATTTTCAAGTTCAGCCAATCTTTTTTTC---AAAAAAAAAATTGCAAGTAAATGAAATCGTGAATAAATGTGTTTCATTTCTCTATCATTATAGAAAAAAAGATCCATATTATATTACTTATGGAATTCGAACCCGAACTCGATTTATGATTCATTATTTCGATCTTATTGGCCTTTGTTCTTTATTTTTTATTTTTCATATAGATTTCCGTCTTTATATTATACCCTTTCGTAGATGAATTATGCTTATTTTCACATCTAGGATTTACATATACAACATATATTACTGTCAAGAGGGAATTTTTCTCAGTATTTAGATATTTAGATTCAAAATAAGAAAGGGATCAATTCAATTATAAACCCGCAAAACAAAGATTAGGATTGGGTTGGGTTGCGCTATATATATCAAAGAGTATACAATAATGATGTATTTGGTGAATCAAATACATGGTCTAATAACAAACCATTTTAACATAACATTTTAATGAGTTGATAATATTAATTGAATATTGAATATTTTTTGAAAGATTTTTGTTAAAGGTTTCATTCATGCCTAATCCATATCGAGTAGACCTTGTTGTTGTGAG

			>Tillandsia_esseriana
			ATAATTATAATAAAGTGAAATATGTCGAAATCCTTTTTGGAATAATACCAAATCGAAAATAAATGTCCGATAGCAAGTTGATCAGTTAATTCAATAAGAGATAAATGGGAGATAGCACTCGATTTAGTTGGTACCACCCAACCGAATCCGATTCAATCGTTTACTCATTCAATGAGTAAATTTTCAAGTTCAGCCAATCTTTTTTTTT----------AATTGCAAGTAAATGAAATCGTGAATAAATGTGTTTCATTTCTCTATCATTATAGAAAAAAAGATCCATATTATATTACTTATGGAATTCGACCCCGAACTCGATTTATGATTCATTATTTCGATCTTATTGGCTTTTGTTCTTTATTTTTTATTTTTCATATAGATTTCCGTCTTTATATTATACCCTTTCGTAGATGAATTATGCTTATTTTCACATCTAGGATTTACATATACAACATATATTACTGTCAAGAGGGAATTTTTCTCAGTATTTAGATATTTAGATTCAAAATAAGAAAGGG-----TTCAATTATAAACCCGCAAAACAAAGATTAGGATTGGGTTGGGTTGCGCTATATATATCAAAGAGTATACAATAATGATGTATTTGGTGAATCAAATACATGGTCTAATAACAAACCATTTTAACATAACATTTTAATGAGTTGATAATATTAATTGACTATTGAATATTTTTTGAAAGATTTTTGTTAAAGGTTTCATTCATGCCTAATCCATATCGAGTAGACCTTGTTGTTGTGAG

			>Tillandsia_exserta
			ATAATTATAATAAAGTGAAATATGTCGAAATCCTTTTTGGAATAATACCAAATCGAAAATAAATGTCCGATAGCAAGTTGATCAGTTAATTCAATAAGAGATAAATGGGAGATAGCACTCGATTTAGTTGGTACCGCCCAACCGAATCCGATTCAATCGTTTACTCATTCAATGAGTAAATTTTCAAGTTCAGCCAATCTTTTTTTC----AAAAAAAAATTGCAAGTAAATGAAATCGTGAATAAATGTGTTTCATTTCTCTATCATTATAGAAAAAAAGATCCATATTATATTACTTATGGAATTCGAACCCGAACTCGATTTATGATTCATTATTTCGATCTTATTGGCCTTTGTTCTTTATTTTTTATTTTTCATATAGATTTCCGTCTTTATATTATACCCTTTCGTAGATGAATTATGCTTATTTTCACATCTAGGATTTACATATACAACATATATTACTGTCAAGAGGGAATTTTTCTCAGTATTTAGATATTTAGATTCAAAATAAGAAAGGGATCAATTCAATTATAAACCCGCAAAACAAAGATTAGGATTGGGTTGGGTTGCGCTATATATATCAAAGAGTATACAATAATGATGTATTTGGTGAATCAAATACATGGTCTAATAACAAACCATTTTAACATAACATTTTAATGAGTTGATAATATTAATTGAATATTGAATATTTTTTGAAAGATTTTTGTTAAAGGTTTCATTCATGCCTAATCCATATCGAGTAGACCTTGTTGTTGTGAG

			>Tillandsia_extensa
			??????????????????????????????????????????????????????????????????????????????????????????????????????????????????????????????????????????????????????????????????????????????????????????????????????????????????????????????????????????????????????????????????????????????????????????????????????????????????????????????????????????????????????????????????????????????????????????????????????????????????????????????????????????????????????????????????????????????????????????????????????????????????????????????????????????????????????????????????????????????????????????????????????????????????????????????????????????????????????????????????????????????????????????????????????????????????????????????????????????????????????????????????????????????????????????????

			>Tillandsia_fasciculata
			ATAATTATAATAAAGTGAAATATGTCGAAATCCTTTTTGGAATAATACCAAATCGAAAATAAATGTCCGATAGCAAGTTGATCAGTTAATTCAATAAGAGATAAATGGGAGATAGCACTCGATTTAGTTGGTACCGCCCAACCGAATCCGATTCAATCGTTTACTCATTCAATGAGTAAATTTTCAAGTTCAGCCAATCTTTTTTTC---AAAAAAAAAATTGCAAGTAAATGAAATCGTGAATAAATGTGTTTCATTTCTCTATCATTATAGAAAAAAAGATCCATATTATATTACTTATGGAATTCGAACCCGAACTCGATTTATGATTCATTATTTCGATCTTATTGGCCTTTGTTCTTTATTTTTTATTTTTCATATAGATTTCCGTCTTTATATTATACCCTTTCGTAGATGAATTATGCTTATTTTCACATCTAGGATTTACATATACAACATATATTACTGTCAAGAGGGAATTTTTCTCAGTATTTAGATATTTAGATTCAAAATAAGAAAGGGATCAATTCAATTATAAACCCGCAAAACAAAGATTAGGATTGGGTTGGGTTGCGCTATATATATCAAAGAGTATACAATAATGATGTATTTGGTGAATCAAATACATGGTCTAATAACAAACCATTTTAACATAACATTTTAATGAGTTGATAATATTAATTGAATATTGAATATTTTTTGAAAGATTTTTGTTAAAGGTTTCATTCATGCCTAATCCATATCGAGTAGACCTTGTTGTTGTGAG

			>Tillandsia_fasciculata_var._densispica
			??????????????????????????????????????????????????????????????????????????????????????????????????????????????????????????????????????????????????????????????????????????????????????????????????????????????????????????????????????????????????????????????????????????????????????????????????????????????????????????????????????????????????????????????????????????????????????????????????????????????????????????????????????????????????????????????????????????????????????????????????????????????????????????????????????????????????????????????????????????????????????????????????????????????????????????????????????????????????????????????????????????????????????????????????????????????????????????????????????????????????????????????????????????????????????????????

			>Tillandsia_fendleri
			ATAATTATAATAAAGTGAAATATGTCGAAATCCTTTTTGGAATAATACCAAATTGAAAATAAATGTCCGATAGCAAGTTGATCAGTTAATTCAATAAGAGATAAATGGGAGATAGCACTCGATTTAGTTGGTACCACCCAACCGAATCCGATTCAATCGTTTACTCATTCAATGAGTAAATTTTCAAGTTCAGCCAATCTTTTTTTTT----------AATTGCAAGTAAATGAAATCGTGAATAAATGTGTTTCATTTCTCTATCATTATAGAAAAAAAGATCCATATTATATTACTTATGGAATTCGAACCCGAACTCGATTTATGATTCATTATTTCGATCTTATTGGACTTTGTTCTTTATTTTTTATTTTTCATATAGATTTCCGTCTTTATATTATACCCTTTCGTAGATGAATTATGCTTATTTTCACATCTAGGATTTACATATACAACATATATTACTGTCAAGAGGGAATTTTTCTCAGTATTTAGATATTTAGATTCAAAATAAGAAAGGGATCAATTCAATTATAAACCCGCAAAACAAAGATTAGGATTGGGTTGGGTTGCGCTATATATATCAAAGAGTATACAATAATGATGTATTTGGTGAATCAAATACATGGTCTAATAACAAACCATTTTAACATAACATTTTAATGAGTTGATAATATTAATTGAATATTGAATATTTTTTGAAAGATTTTTGTTAAAGGTTTCATTCATGCCTAATCCATATCGAGTAGACCTTGTTGTTGTGAG

			>Tillandsia_ferreyrae
			??????????????????????????????????????????????????????????????????????????????????????????????????????????????????????????????????????????????????????????????????????????????????????????????????????????????????????????????????????????????????????????????????????????????????????????????????????????????????????????????????????????????????????????????????????????????????????????????????????????????????????????????????????????????????????????????????????????????????????????????????????????????????????????????????????????????????????????????????????????????????????????????????????????????????????????????????????????????????????????????????????????????????????????????????????????????????????????????????????????????????????????????????????????????????????????????

			>Tillandsia_ferrisiana
			??????????????????????????????????????????????????????????????????????????????????????????????????????????????????????????????????????????????????????????????????????????????????????????????????????????????????????????????????????????????????????????????????????????????????????????????????????????????????????????????????????????????????????????????????????????????????????????????????????????????????????????????????????????????????????????????????????????????????????????????????????????????????????????????????????????????????????????????????????????????????????????????????????????????????????????????????????????????????????????????????????????????????????????????????????????????????????????????????????????????????????????????????????????????????????????????

			>Tillandsia_festucoides
			??????????????????????????????????????????????????????????????????????????????????????????????????????????????????????????????????????????????????????????????????????????????????????????????????????????????????????????????????????????????????????????????????????????????????????????????????????????????????????????????????????????????????????????????????????????????????????????????????????????????????????????????????????????????????????????????????????????????????????????????????????????????????????????????????????????????????????????????????????????????????????????????????????????????????????????????????????????????????????????????????????????????????????????????????????????????????????????????????????????????????????????????????????????????????????????????

			>Tillandsia_filifolia
			??????????????????????????????????????????????????????????????????????????????????????????????????????????????????????????????????????????????????????????????????????????????????????????????????????????????????????????????????????????????????????????????????????????????????????????????????????????????????????????????????????????????????????????????????????????????????????????????????????????????????????????????????????????????????????????????????????????????????????????????????????????????????????????????????????????????????????????????????????????????????????????????????????????????????????????????????????????????????????????????????????????????????????????????????????????????????????????????????????????????????????????????????????????????????????????????

			>Tillandsia_flabellata
			??????????????????????????????????????????????????????????????????????????????????????????????????????????????????????????????????????????????????????????????????????????????????????????????????????????????????????????????????????????????????????????????????????????????????????????????????????????????????????????????????????????????????????????????????????????????????????????????????????????????????????????????????????????????????????????????????????????????????????????????????????????????????????????????????????????????????????????????????????????????????????????????????????????????????????????????????????????????????????????????????????????????????????????????????????????????????????????????????????????????????????????????????????????????????????????????

			>Tillandsia_flavobracteata
			??????????????????????????????????????????????????????????????????????????????????????????????????????????????????????????????????????????????????????????????????????????????????????????????????????????????????????????????????????????????????????????????????????????????????????????????????????????????????????????????????????????????????????????????????????????????????????????????????????????????????????????????????????????????????????????????????????????????????????????????????????????????????????????????????????????????????????????????????????????????????????????????????????????????????????????????????????????????????????????????????????????????????????????????????????????????????????????????????????????????????????????????????????????????????????????????

			>Tillandsia_flexuosa
			??????????????????????????????????????????????????????????????????????????????????????????????????????????????????????????????????????????????????????????????????????????????????????????????????????????????????????????????????????????????????????????????????????????????????????????????????????????????????????????????????????????????????????????????????????????????????????????????????????????????????????????????????????????????????????????????????????????????????????????????????????????????????????????????????????????????????????????????????????????????????????????????????????????????????????????????????????????????????????????????????????????????????????????????????????????????????????????????????????????????????????????????????????????????????????????????

			>Tillandsia_floribunda
			??????????????????????????????????????????????????????????????????????????????????????????????????????????????????????????????????????????????????????????????????????????????????????????????????????????????????????????????????????????????????????????????????????????????????????????????????????????????????????????????????????????????????????????????????????????????????????????????????????????????????????????????????????????????????????????????????????????????????????????????????????????????????????????????????????????????????????????????????????????????????????????????????????????????????????????????????????????????????????????????????????????????????????????????????????????????????????????????????????????????????????????????????????????????????????????????

			>Tillandsia_foliosa
			??????????????????????????????????????????????????????????????????????????????????????????????????????????????????????????????????????????????????????????????????????????????????????????????????????????????????????????????????????????????????????????????????????????????????????????????????????????????????????????????????????????????????????????????????????????????????????????????????????????????????????????????????????????????????????????????????????????????????????????????????????????????????????????????????????????????????????????????????????????????????????????????????????????????????????????????????????????????????????????????????????????????????????????????????????????????????????????????????????????????????????????????????????????????????????????????

			>Tillandsia_fresnilloensis
			??????????????????????????????????????????????????????????????????????????????????????????????????????????????????????????????????????????????????????????????????????????????????????????????????????????????????????????????????????????????????????????????????????????????????????????????????????????????????????????????????????????????????????????????????????????????????????????????????????????????????????????????????????????????????????????????????????????????????????????????????????????????????????????????????????????????????????????????????????????????????????????????????????????????????????????????????????????????????????????????????????????????????????????????????????????????????????????????????????????????????????????????????????????????????????????????

			>Tillandsia_friesii
			??????????????????????????????????????????????????????????????????????????????????????????????????????????????????????????????????????????????????????????????????????????????????????????????????????????????????????????????????????????????????????????????????????????????????????????????????????????????????????????????????????????????????????????????????????????????????????????????????????????????????????????????????????????????????????????????????????????????????????????????????????????????????????????????????????????????????????????????????????????????????????????????????????????????????????????????????????????????????????????????????????????????????????????????????????????????????????????????????????????????????????????????????????????????????????????????

			>Tillandsia_fuchsii
			??????????????????????????????????????????????????????????????????????????????????????????????????????????????????????????????????????????????????????????????????????????????????????????????????????????????????????????????????????????????????????????????????????????????????????????????????????????????????????????????????????????????????????????????????????????????????????????????????????????????????????????????????????????????????????????????????????????????????????????????????????????????????????????????????????????????????????????????????????????????????????????????????????????????????????????????????????????????????????????????????????????????????????????????????????????????????????????????????????????????????????????????????????????????????????????????

			>Tillandsia_funckiana
			??????????????????????????????????????????????????????????????????????????????????????????????????????????????????????????????????????????????????????????????????????????????????????????????????????????????????????????????????????????????????????????????????????????????????????????????????????????????????????????????????????????????????????????????????????????????????????????????????????????????????????????????????????????????????????????????????????????????????????????????????????????????????????????????????????????????????????????????????????????????????????????????????????????????????????????????????????????????????????????????????????????????????????????????????????????????????????????????????????????????????????????????????????????????????????????????

			>Tillandsia_funckiana_var._recurvifolia
			??????????????????????????????????????????????????????????????????????????????????????????????????????????????????????????????????????????????????????????????????????????????????????????????????????????????????????????????????????????????????????????????????????????????????????????????????????????????????????????????????????????????????????????????????????????????????????????????????????????????????????????????????????????????????????????????????????????????????????????????????????????????????????????????????????????????????????????????????????????????????????????????????????????????????????????????????????????????????????????????????????????????????????????????????????????????????????????????????????????????????????????????????????????????????????????????

			>Tillandsia_funebris
			ATAATTATAATAAAGTGAAATATGTCAAAATCCTTTTTGGAATAATACCAAATCGAAAATAAATGTCCGATAGCAAGTTGATCAGTTAATTCAATAAGAGATAAATGGGAGATAGCACTCGATTTAGTTGGTACCACCCAACCGAATCCGATTCAATCGCTTACTCATTCAATGAGTAAATTTTAAAGTTCAGCCAATCTTTATTTTTTT----------TTGCAAGTAAATGAAATCGTGAATAAATGTGTTTCATTTCTCTATCATTATAGAAAAAAAGATCCATATTATATTACTTATGGAATTCGAACCCGAACTCGATTTATGATTCATTATTTCGATCTTATTGGCCTTTGTTCTTTATTTTTTATTTTTCATATAGATTCCCGTCTTTATATTATACCCTTTCGTAGATGAATTATGCTTATTTTCACATCTAGGATTTACATATACAACATATATTACTGTCAAGAGGGAATTTTTCCCAGTATTTAGATATTTAGATTCAAAATAAGAAAGGGATCAATTCAATTATAAACCCGCAAAACAAAGATTAGGATTGGGTTGGGTTGCGCTATATATATCAAAGAGTATACAATAATGATGTATTTGGTGAATCAAATACATGGTCTAATAACAAACCATTTTAACATAACATTTTAATGAGTTGATAATATTAATTGACTATTGAATATTTTTTGAAAGATTTTTGTTAAAGGTTTCATTCATGCCTAATCCATATCGAGTAGACCTTGTTGTTGTGAG

			>Tillandsia_gardneri
			ATAATTATAATAAAGTGAAATATGTCGAAATCCTTTTTGGAATAATACCAAATCGAAAATAAATGTCCGATAGCAAGTTGATCAGTTAATTCAATAAGAGATAAATGGGAGATAGCACTCGATTTAGTTGGTACCACCCAACCGAATCCGATTCAATCGTTTACTCATTCAATGAGTAAATTTTCAAGTTCAGCCAATCTTTATTTTTTT------A-TTGTGCAAGTAAATGAAATCGTGAATAAATGTGTTTCATTTCTCTATCATTATAGAAAAAAAGATCCATATTATATTACTTATGGAATTCGAACCCGAACTCGATTTATGATTCATTATTTCGATCTTATTGGCCTTTGTTCTTTATTTTTTATTTTTCATATAGATTTCCGTCTTTATATTATACCCTTTCGTAGATGAATTATGCTTATTTTCACATCTAGGATTTACATATACAACATATATTACTGTCAAGAGGGAATTTTTCTCAGTATTTAGATATTTAGATTCAAAATAAGAAAGGGATCAATTCAATTATAAACCCGCAAAACAAAGATTAGGATTGGGTTGGGTTGCGCTATATATATCAAAGAGTATACAATAATGATGTATTTGGTGAATCAAATACATGGTCTAATAACAAACCATTTTAACATAACATTTTAATGAGTTGATAATATTAATTGAATATTGAATATTTTTTGAAAGATTTTTGTTAAAGGTTTCATTCATGCCTAATCCATATCGAGTAGACCTTGTTGTTGTGAG

			>Tillandsia_gardneri_var._rupicola
			??????????????????????????????????????????????????????????????????????????????????????????????????????????????????????????????????????????????????????????????????????????????????????????????????????????????????????????????????????????????????????????????????????????????????????????????????????????????????????????????????????????????????????????????????????????????????????????????????????????????????????????????????????????????????????????????????????????????????????????????????????????????????????????????????????????????????????????????????????????????????????????????????????????????????????????????????????????????????????????????????????????????????????????????????????????????????????????????????????????????????????????????????????????????????????????????

			>Tillandsia_geissei
			??????????????????????????????????????????????????????????????????????????????????????????????????????????????????????????????????????????????????????????????????????????????????????????????????????????????????????????????????????????????????????????????????????????????????????????????????????????????????????????????????????????????????????????????????????????????????????????????????????????????????????????????????????????????????????????????????????????????????????????????????????????????????????????????????????????????????????????????????????????????????????????????????????????????????????????????????????????????????????????????????????????????????????????????????????????????????????????????????????????????????????????????????????????????????????????????

			>Tillandsia_geminiflora
			??????????????????????????????????????????????????????????????????????????????????????????????????????????????????????????????????????????????????????????????????????????????????????????????????????????????????????????????????????????????????????????????????????????????????????????????????????????????????????????????????????????????????????????????????????????????????????????????????????????????????????????????????????????????????????????????????????????????????????????????????????????????????????????????????????????????????????????????????????????????????????????????????????????????????????????????????????????????????????????????????????????????????????????????????????????????????????????????????????????????????????????????????????????????????????????????

			>Tillandsia_grandispica
			??????????????????????????????????????????????????????????????????????????????????????????????????????????????????????????????????????????????????????????????????????????????????????????????????????????????????????????????????????????????????????????????????????????????????????????????????????????????????????????????????????????????????????????????????????????????????????????????????????????????????????????????????????????????????????????????????????????????????????????????????????????????????????????????????????????????????????????????????????????????????????????????????????????????????????????????????????????????????????????????????????????????????????????????????????????????????????????????????????????????????????????????????????????????????????????????

			>Tillandsia_grao-mogolensis
			??????????????????????????????????????????????????????????????????????????????????????????????????????????????????????????????????????????????????????????????????????????????????????????????????????????????????????????????????????????????????????????????????????????????????????????????????????????????????????????????????????????????????????????????????????????????????????????????????????????????????????????????????????????????????????????????????????????????????????????????????????????????????????????????????????????????????????????????????????????????????????????????????????????????????????????????????????????????????????????????????????????????????????????????????????????????????????????????????????????????????????????????????????????????????????????????

			>Tillandsia_guatemalensis
			ATAATTATAATAAAGTGAAATATGTCGAAATCCTTTTTGGAATAATACCAAATCGAAAATAAATGTCCGATAGCAAGTTGATCAGTTAATTCAATAAGAGATAAATGGGAGATAGCACTCGATTTAGTTGGTACCGCCCAACCGAATCCGATTCAATCGTTTACTCATTCAATGAGTAAATTTTCAAGTTCAGCCAATCTTTTTTTTCA-AAAAAAAAAATTGCAAGTAAATGAAATCGTGAATAAATGTGTTTCATTTCTCTATCATTATAGAAAAAAAGATCCATATTATATTACTTATGGAATTCGAACCCGAACTCGATTTATGATTCATTATTTCGATCTTATTGGCCTTTGTTCTTTATTTTTTATTTTTCATATAGATTTCCGTCTTTATATTATACCCTTTCGTAGATGAATTATGCTTATTTTCACATCTAGGATTTACATATACAACATATATTACTGTCAAGAGGGAATTTTTCTCAGTATTTAGATATTTAGATTCAAAATAAGAAAGGGATCAATTCAATTATAAACCCGCAAAACAAAGATTAGGATTGGGTTGGGTTGCGCTATATATATCAAAGAGTATACAATAATGATGTATTTGGTGAATCAAATACATGGTCTAATAACAAACCATTTTAACATAACATTTTAATGAGTTGATAATATTAATTGAATATTGAATATTTTTTGAAAGATTTTTGTTAAAGGTTTCATTCATGCCTAATCCATATCGAGTAGACCTTGTTGTTGTGAG

			>Tillandsia_guenther-nolleri
			??????????????????????????????????????????????????????????????????????????????????????????????????????????????????????????????????????????????????????????????????????????????????????????????????????????????????????????????????????????????????????????????????????????????????????????????????????????????????????????????????????????????????????????????????????????????????????????????????????????????????????????????????????????????????????????????????????????????????????????????????????????????????????????????????????????????????????????????????????????????????????????????????????????????????????????????????????????????????????????????????????????????????????????????????????????????????????????????????????????????????????????????????????????????????????????????

			>Tillandsia_guerreroensis
			??????????????????????????????????????????????????????????????????????????????????????????????????????????????????????????????????????????????????????????????????????????????????????????????????????????????????????????????????????????????????????????????????????????????????????????????????????????????????????????????????????????????????????????????????????????????????????????????????????????????????????????????????????????????????????????????????????????????????????????????????????????????????????????????????????????????????????????????????????????????????????????????????????????????????????????????????????????????????????????????????????????????????????????????????????????????????????????????????????????????????????????????????????????????????????????????

			>Tillandsia_gymnobotrya
			??????????????????????????????????????????????????????????????????????????????????????????????????????????????????????????????????????????????????????????????????????????????????????????????????????????????????????????????????????????????????????????????????????????????????????????????????????????????????????????????????????????????????????????????????????????????????????????????????????????????????????????????????????????????????????????????????????????????????????????????????????????????????????????????????????????????????????????????????????????????????????????????????????????????????????????????????????????????????????????????????????????????????????????????????????????????????????????????????????????????????????????????????????????????????????????????

			>Tillandsia_harrisii
			??????????????????????????????????????????????????????????????????????????????????????????????????????????????????????????????????????????????????????????????????????????????????????????????????????????????????????????????????????????????????????????????????????????????????????????????????????????????????????????????????????????????????????????????????????????????????????????????????????????????????????????????????????????????????????????????????????????????????????????????????????????????????????????????????????????????????????????????????????????????????????????????????????????????????????????????????????????????????????????????????????????????????????????????????????????????????????????????????????????????????????????????????????????????????????????????

			>Tillandsia_hegeri
			??????????????????????????????????????????????????????????????????????????????????????????????????????????????????????????????????????????????????????????????????????????????????????????????????????????????????????????????????????????????????????????????????????????????????????????????????????????????????????????????????????????????????????????????????????????????????????????????????????????????????????????????????????????????????????????????????????????????????????????????????????????????????????????????????????????????????????????????????????????????????????????????????????????????????????????????????????????????????????????????????????????????????????????????????????????????????????????????????????????????????????????????????????????????????????????????

			>Tillandsia_heliconioides
			??????????????????????????????????????????????????????????????????????????????????????????????????????????????????????????????????????????????????????????????????????????????????????????????????????????????????????????????????????????????????????????????????????????????????????????????????????????????????????????????????????????????????????????????????????????????????????????????????????????????????????????????????????????????????????????????????????????????????????????????????????????????????????????????????????????????????????????????????????????????????????????????????????????????????????????????????????????????????????????????????????????????????????????????????????????????????????????????????????????????????????????????????????????????????????????????

			>Tillandsia_heteromorpha
			??????????????????????????????????????????????????????????????????????????????????????????????????????????????????????????????????????????????????????????????????????????????????????????????????????????????????????????????????????????????????????????????????????????????????????????????????????????????????????????????????????????????????????????????????????????????????????????????????????????????????????????????????????????????????????????????????????????????????????????????????????????????????????????????????????????????????????????????????????????????????????????????????????????????????????????????????????????????????????????????????????????????????????????????????????????????????????????????????????????????????????????????????????????????????????????????

			>Tillandsia_heterophylla
			ATAATTATAATAAAGTGAAATATGTCGAAATCCTTTTTGGAATAATACCAAATCGAAAATAAATGTCCGATAGCAAGTTGATCAGTTAATTCAATAAGAGATAAATGGGAGATAGCACTCGATTTAGTTGGTACCACCCAACCGAATCCGATTCAATCGTTTACTCATTCAATGAGTCAATTTTCAAGTTCAGCCAATCTTTTTTTTT----------AATTGCAAGTAAATGAAATCGTGAATAAATGTGTTTCATTTCTCTATCATTATAGAAAAAAAGATCCATATTATATTACTTATGGAATTCGAACCCGAACTCGATTTATGATTCATTATTTCGATCTTATTGGCCTTTGTTCTTTATTTTTTATTTTTCATATAGATTTCCGTCTTTATATTATACCCTTTCGTAGATGAATTATGCTTATTTTCACATCTAGGATTTACATATACAACATATATTACTGTCAAGAGGGAATTTTTCTCAGTATTTAGATATTTAGATTCAAAATAAGAAAGGG-----TTCAATTATAAACCCGCAAAACAAAGATTAGGATTGGGTTGGGTTGCGCTATATATATCAAAGAGTATACAATAATGATGTATTTGGTGAATCAAATACATGGTCTAATAACAAACCATTTTAACATAACATTTTAATGAGTTGATAATATTAATTGAATATTGAATATTTTTTGAAAGATTTTTGTTAAAGGTTTCATTCATGCCTAATCCATATCGAGTAGACCTTGTTGTTGTGAG

			>Tillandsia_heubergeri
			ATAATTATAATAAAGTGAAATATGTCGAAATCCTTTTTGGAATAATACCAAATCGAAAATAAATGTCCGATAGCAAGTTGATCAGTTAATTCAATAAGAGATAAATGGGAGATAGCACTCGATTTAGTTGGTACCACCCAACCGAATCCGATTCAATCGTTTACTCATTGAATGAGTAAATTTTCAAGTTCAGTCAATCTTTTTTTTTTT--------AATTGCAAGTAAATGAAATCGTGAATAAATGTGTTTCATTTCTCTATCATTATAGAAAAAAAGATCCATATTATATTACTTATGGAATTCGAACCCGAACTCGATTTATGATTCATTATTTCGATCTTATTGGCCTTTGTTCTTTATTTTTTATTTTTCATATAGATTTCCGTCTTTATATTATACCCTTTCGTAGATGAATTATGCTTATTTTCACATCTAGGATTTACATATACAACATATATTACTGTCAAGAGGGAATTTTTCTCAGTATTTAGATATTTAGATTCAAAATAAGAAAGGGATCAATTCAATTATAAACCCGCAAAACAAAGATTAGGATTGGGTTGGGTTGCGCTATATATATCAAAGAGTATACAATAATGATGTATTTGGTGAATCAAATACATGGTCTAATAACAAACCATTTTAACATAACATTTTAATGAGTTGATAATATTAATTGAATATTGAATATTTTTTGAAAGATTTTTGTTAAAGGTTTCATTCATGCCTAATCCATATCGAGTAGACCTTGTTGTTGTGAG

			>Tillandsia_hildae
			??????????????????????????????????????????????????????????????????????????????????????????????????????????????????????????????????????????????????????????????????????????????????????????????????????????????????????????????????????????????????????????????????????????????????????????????????????????????????????????????????????????????????????????????????????????????????????????????????????????????????????????????????????????????????????????????????????????????????????????????????????????????????????????????????????????????????????????????????????????????????????????????????????????????????????????????????????????????????????????????????????????????????????????????????????????????????????????????????????????????????????????????????????????????????????????????

			>Tillandsia_hofackeri
			??????????????????????????????????????????????????????????????????????????????????????????????????????????????????????????????????????????????????????????????????????????????????????????????????????????????????????????????????????????????????????????????????????????????????????????????????????????????????????????????????????????????????????????????????????????????????????????????????????????????????????????????????????????????????????????????????????????????????????????????????????????????????????????????????????????????????????????????????????????????????????????????????????????????????????????????????????????????????????????????????????????????????????????????????????????????????????????????????????????????????????????????????????????????????????????????

			>Tillandsia_hondurensis
			??????????????????????????????????????????????????????????????????????????????????????????????????????????????????????????????????????????????????????????????????????????????????????????????????????????????????????????????????????????????????????????????????????????????????????????????????????????????????????????????????????????????????????????????????????????????????????????????????????????????????????????????????????????????????????????????????????????????????????????????????????????????????????????????????????????????????????????????????????????????????????????????????????????????????????????????????????????????????????????????????????????????????????????????????????????????????????????????????????????????????????????????????????????????????????????????

			>Tillandsia_horstii
			??????????????????????????????????????????????????????????????????????????????????????????????????????????????????????????????????????????????????????????????????????????????????????????????????????????????????????????????????????????????????????????????????????????????????????????????????????????????????????????????????????????????????????????????????????????????????????????????????????????????????????????????????????????????????????????????????????????????????????????????????????????????????????????????????????????????????????????????????????????????????????????????????????????????????????????????????????????????????????????????????????????????????????????????????????????????????????????????????????????????????????????????????????????????????????????????

			>Tillandsia_hromadnikiana
			??????????????????????????????????????????????????????????????????????????????????????????????????????????????????????????????????????????????????????????????????????????????????????????????????????????????????????????????????????????????????????????????????????????????????????????????????????????????????????????????????????????????????????????????????????????????????????????????????????????????????????????????????????????????????????????????????????????????????????????????????????????????????????????????????????????????????????????????????????????????????????????????????????????????????????????????????????????????????????????????????????????????????????????????????????????????????????????????????????????????????????????????????????????????????????????????

			>Tillandsia_huamelulaensis
			??????????????????????????????????????????????????????????????????????????????????????????????????????????????????????????????????????????????????????????????????????????????????????????????????????????????????????????????????????????????????????????????????????????????????????????????????????????????????????????????????????????????????????????????????????????????????????????????????????????????????????????????????????????????????????????????????????????????????????????????????????????????????????????????????????????????????????????????????????????????????????????????????????????????????????????????????????????????????????????????????????????????????????????????????????????????????????????????????????????????????????????????????????????????????????????????

			>Tillandsia_huarazensis
			??????????????????????????????????????????????????????????????????????????????????????????????????????????????????????????????????????????????????????????????????????????????????????????????????????????????????????????????????????????????????????????????????????????????????????????????????????????????????????????????????????????????????????????????????????????????????????????????????????????????????????????????????????????????????????????????????????????????????????????????????????????????????????????????????????????????????????????????????????????????????????????????????????????????????????????????????????????????????????????????????????????????????????????????????????????????????????????????????????????????????????????????????????????????????????????????

			>Tillandsia_humilis
			??????????????????????????????????????????????????????????????????????????????????????????????????????????????????????????????????????????????????????????????????????????????????????????????????????????????????????????????????????????????????????????????????????????????????????????????????????????????????????????????????????????????????????????????????????????????????????????????????????????????????????????????????????????????????????????????????????????????????????????????????????????????????????????????????????????????????????????????????????????????????????????????????????????????????????????????????????????????????????????????????????????????????????????????????????????????????????????????????????????????????????????????????????????????????????????????

			>Tillandsia_imperialis
			??????????????????????????????????????????????????????????????????????????????????????????????????????????????????????????????????????????????????????????????????????????????????????????????????????????????????????????????????????????????????????????????????????????????????????????????????????????????????????????????????????????????????????????????????????????????????????????????????????????????????????????????????????????????????????????????????????????????????????????????????????????????????????????????????????????????????????????????????????????????????????????????????????????????????????????????????????????????????????????????????????????????????????????????????????????????????????????????????????????????????????????????????????????????????????????????

			>Tillandsia_inopinata
			??????????????????????????????????????????????????????????????????????????????????????????????????????????????????????????????????????????????????????????????????????????????????????????????????????????????????????????????????????????????????????????????????????????????????????????????????????????????????????????????????????????????????????????????????????????????????????????????????????????????????????????????????????????????????????????????????????????????????????????????????????????????????????????????????????????????????????????????????????????????????????????????????????????????????????????????????????????????????????????????????????????????????????????????????????????????????????????????????????????????????????????????????????????????????????????????

			>Tillandsia_ionantha
			ATAATTATAATAAAGTGAAATATGTCGAAATCCTTTTTGGAATAATACCAAATCGAAAATAAATGTCCGATAGCAAGTTGATCAGTTAATTCAATAAGAGATAAATGGGAGATAGCACTCGATTTAGTTGGTACCGCCCAACCGAATCCGATTCAATCGTTTACTCATTCAATGAGTAAATTTTCAAGTTCAGCCAATCTTTTTTTC---AAAAAAAAAATTGCAAGTAAATGAAATCGTGAATAAATGTGTTTCATTTCTCTATCATTATAGAAAAAAAGATCCATATTATATTACTTATGGAATTCGAACCCGAACTCGATTTATGATTCATTATTTCGATCTTATTGGCCTTTGTTCTTTATTTTTTATTTTTCATATAGATTTCCGTCTTTATATTATACCCTTTCGTAGATGAATTATGCTTATTTTCACATCTAGGATTTACATATACAACATATATTACTGTCAAGAGGGAATTTTTCTCAGTATTTAGATATTTAGATTCAAAATAAGAAAGGGATCAATTCAATTATAAACCCGCAAAACAAAGATTAGGATTGGGTTGGGTTGCGCTATATATATCAAAGAGTATACAATAATGATGTATTTGGTGAATCAAATACATGGTCTAATAACAAACCATTTTAACATAACATTTTAATGAGTTGATAATATTAATTGAATATTGAATATTTTTTGAAAGATTTTTGTTAAAGGTTTCATTCATGCCTAATCCATATCGAGTAGACCTTGTTGTTGTGAG

			>Tillandsia_ionochroma
			??????????????????????????????????????????????????????????????????????????????????????????????????????????????????????????????????????????????????????????????????????????????????????????????????????????????????????????????????????????????????????????????????????????????????????????????????????????????????????????????????????????????????????????????????????????????????????????????????????????????????????????????????????????????????????????????????????????????????????????????????????????????????????????????????????????????????????????????????????????????????????????????????????????????????????????????????????????????????????????????????????????????????????????????????????????????????????????????????????????????????????????????????????????????????????????????

			>Tillandsia_ixioides
			ATAATTATAATAAAGTGAAATATGTCGAAATCCTTTTTGGAATAATACCAAATCGAAAATAAATGTCCGATAGCAAGTTGATCAGTTAATTCAATAAGAGATAAATGGGAGATAGCACTCGATTTAGTTGGTACCACCCAACCGAATCCGATTCAATCGTTTACTCATTCAATGAGTAAATTTTCAAGTTCAGCCAATCTTTATTTTTTT------TTAATTGCAAGTAAATGAAATCGTGAATAAATGTGTTTCATTTCTCTATCATTATAGAAAAAAAGATACATATTATATTACTTATGGAATTAGAACCCGAACTCGATTTATGATTCATTATTTCGATCTTATTGGCCTTTGTTCTTTATTTTTTATTTTTCATATAGATTTCCGTCTTTATATTATACCCTTTCGTAGATGAATTATGCTTACTTTCACATCTAGGATTTACATATACAACATATATTACTGTCAAGAGGGAATTTTTCTCAGTATTTAGATATTTAGATTCAAAATAAGAAAGGGATCAATTCAATTATAAACCCGCAAAACAAAGATTAGGATTGGGTTGGGTTGCGCTATATATATCAAAGAGTATACAATAATGATGTATTTGGTGAATCAAATACATGGTCTAATAACAAACCATTTTAACATAACATTTTAATGAGTTGATAATATTAATTGACTATTGAATATTTTTTGAAAGATTTTTGTTAAAGGTTTCATTCATGCCTAATCCATATCGAGTAGACCTTGTTGTTGTGAG

			>Tillandsia_izabalensis
			??????????????????????????????????????????????????????????????????????????????????????????????????????????????????????????????????????????????????????????????????????????????????????????????????????????????????????????????????????????????????????????????????????????????????????????????????????????????????????????????????????????????????????????????????????????????????????????????????????????????????????????????????????????????????????????????????????????????????????????????????????????????????????????????????????????????????????????????????????????????????????????????????????????????????????????????????????????????????????????????????????????????????????????????????????????????????????????????????????????????????????????????????????????????????????????????

			>Tillandsia_jaliscomonticola
			??????????????????????????????????????????????????????????????????????????????????????????????????????????????????????????????????????????????????????????????????????????????????????????????????????????????????????????????????????????????????????????????????????????????????????????????????????????????????????????????????????????????????????????????????????????????????????????????????????????????????????????????????????????????????????????????????????????????????????????????????????????????????????????????????????????????????????????????????????????????????????????????????????????????????????????????????????????????????????????????????????????????????????????????????????????????????????????????????????????????????????????????????????????????????????????????

			>Tillandsia_jucunda
			??????????????????????????????????????????????????????????????????????????????????????????????????????????????????????????????????????????????????????????????????????????????????????????????????????????????????????????????????????????????????????????????????????????????????????????????????????????????????????????????????????????????????????????????????????????????????????????????????????????????????????????????????????????????????????????????????????????????????????????????????????????????????????????????????????????????????????????????????????????????????????????????????????????????????????????????????????????????????????????????????????????????????????????????????????????????????????????????????????????????????????????????????????????????????????????????

			>Tillandsia_jucunda_var._vivida
			??????????????????????????????????????????????????????????????????????????????????????????????????????????????????????????????????????????????????????????????????????????????????????????????????????????????????????????????????????????????????????????????????????????????????????????????????????????????????????????????????????????????????????????????????????????????????????????????????????????????????????????????????????????????????????????????????????????????????????????????????????????????????????????????????????????????????????????????????????????????????????????????????????????????????????????????????????????????????????????????????????????????????????????????????????????????????????????????????????????????????????????????????????????????????????????????

			>Tillandsia_juerg-rutschmannii
			??????????????????????????????????????????????????????????????????????????????????????????????????????????????????????????????????????????????????????????????????????????????????????????????????????????????????????????????????????????????????????????????????????????????????????????????????????????????????????????????????????????????????????????????????????????????????????????????????????????????????????????????????????????????????????????????????????????????????????????????????????????????????????????????????????????????????????????????????????????????????????????????????????????????????????????????????????????????????????????????????????????????????????????????????????????????????????????????????????????????????????????????????????????????????????????????

			>Tillandsia_juncea
			ATAATTATAATAAAGTGAAATATGTCGAAATCCTTTTTGGAATAATACCAAATCGAAAATAAATGTCCGATAGCAAGTTGATCAGTTAATTCAATAAGAGATAAATGGGAGATAGCACTCGATTTAGTTGGTACCGCCCAACCGAATCCGATTCAATCGTTTACTCATTCAATGAGTAAATTTTCAAGTTCAGCCAATCTTTTTTTC---AAAAAAAAAATTGCAAGTAAATGAAATCGTGAATAAATGTGTTTCATTTCTCTATCATTATAGAAAAAAAGATCCATATTATATTACTTATGGAATTCGAACCCGAACTCGATTTATGATTCATTATTTCGATCTTATTGGCCTTTGTTCTTTATTTTTTATTTTTCATATAGATTTCCGTCTTTATATTATACCCTTTCGTAGATGAATTATGCTTATTTTCACATCTAGGATTTACATATACAACATATATTACTGTCAAGAGGGAATTTTTCTCAGTATTTAGATATTTAGATTCAAAATAAGAAAGGGATCAATTCAATTATAAACCCGCAAAACAAAGATTAGGATTGGGTTGGGTTGCGCTATATATATCAAAGAGTATACAATAATGATGTATTTGGTGAATCAAATACATGGTCTAATAACAAACCATTTTAACATAACATTTTAATGAGTTGATAATATTAATTGAATATTGAATATTTTTTGAAAGATTTTTGTTAAAGGTTTCATTCATGCCTAATCCATATCGAGTAGACCTTGTTGTTGTGAG

			>Tillandsia_kammii
			??????????????????????????????????????????????????????????????????????????????????????????????????????????????????????????????????????????????????????????????????????????????????????????????????????????????????????????????????????????????????????????????????????????????????????????????????????????????????????????????????????????????????????????????????????????????????????????????????????????????????????????????????????????????????????????????????????????????????????????????????????????????????????????????????????????????????????????????????????????????????????????????????????????????????????????????????????????????????????????????????????????????????????????????????????????????????????????????????????????????????????????????????????????????????????????????

			>Tillandsia_karwinskyana
			??????????????????????????????????????????????????????????????????????????????????????????????????????????????????????????????????????????????????????????????????????????????????????????????????????????????????????????????????????????????????????????????????????????????????????????????????????????????????????????????????????????????????????????????????????????????????????????????????????????????????????????????????????????????????????????????????????????????????????????????????????????????????????????????????????????????????????????????????????????????????????????????????????????????????????????????????????????????????????????????????????????????????????????????????????????????????????????????????????????????????????????????????????????????????????????????

			>Tillandsia_kauffmannii
			ATAATTATAATAAAGTGAAATATGTCGAAATCCTTTTTGGAATAATACCAAATCGAAAATAAATGTCCGATAGCAAGTTGATCAGTTAATTCAATAAGAGATAAATGGGAGATAGCACTCGATTTAGTTGGTACCACCCAACCGAATCCGATTCAATCGTTTACTCATTCAATGAGTAAATTTTCAAGTTCAGCCAATCTTTATTTTT-------------------------------------------------------------------------------------------------------------------------------------------------------------------TTTTTCATATAGATTTCCGTCTTTATATTATACCCTTTCGTAGATGAATTATGCTTATTTTCACATCTAGGATTTACATATACAACATATATTACTGTCAAGAGGGAATTTTTCTCAGTATTTAGATATTTAGATTCAAAATAAGAAAGGGATCAATTCAATTATAAACCCGCAAAACAAAGATTAGGATTGGGTTGGGTTGCGCTATATATATCAAAGAGTATACAATAATGATGTATTTGGTGAATCAAATACATGGTCTAATAACAAACCATTTTAACATAACATTTTAATGAGTTGATAATATTAATTGAATATTGAATATTTTTTGAAAGATTTTTGTTAAAGGTTTCATTCATGCCTAATCCATATCGAGTAGACCTTGTTGTTGTGAG

			>Tillandsia_kautskyi
			??????????????????????????????????????????????????????????????????????????????????????????????????????????????????????????????????????????????????????????????????????????????????????????????????????????????????????????????????????????????????????????????????????????????????????????????????????????????????????????????????????????????????????????????????????????????????????????????????????????????????????????????????????????????????????????????????????????????????????????????????????????????????????????????????????????????????????????????????????????????????????????????????????????????????????????????????????????????????????????????????????????????????????????????????????????????????????????????????????????????????????????????????????????????????????????????

			>Tillandsia_kegeliana
			??????????????????????????????????????????????????????????????????????????????????????????????????????????????????????????????????????????????????????????????????????????????????????????????????????????????????????????????????????????????????????????????????????????????????????????????????????????????????????????????????????????????????????????????????????????????????????????????????????????????????????????????????????????????????????????????????????????????????????????????????????????????????????????????????????????????????????????????????????????????????????????????????????????????????????????????????????????????????????????????????????????????????????????????????????????????????????????????????????????????????????????????????????????????????????????????

			>Tillandsia_kirchhoffiana
			??????????????????????????????????????????????????????????????????????????????????????????????????????????????????????????????????????????????????????????????????????????????????????????????????????????????????????????????????????????????????????????????????????????????????????????????????????????????????????????????????????????????????????????????????????????????????????????????????????????????????????????????????????????????????????????????????????????????????????????????????????????????????????????????????????????????????????????????????????????????????????????????????????????????????????????????????????????????????????????????????????????????????????????????????????????????????????????????????????????????????????????????????????????????????????????????

			>Tillandsia_kirschnekii
			??????????????????????????????????????????????????????????????????????????????????????????????????????????????????????????????????????????????????????????????????????????????????????????????????????????????????????????????????????????????????????????????????????????????????????????????????????????????????????????????????????????????????????????????????????????????????????????????????????????????????????????????????????????????????????????????????????????????????????????????????????????????????????????????????????????????????????????????????????????????????????????????????????????????????????????????????????????????????????????????????????????????????????????????????????????????????????????????????????????????????????????????????????????????????????????????

			>Tillandsia_klausii
			ATAATTATAATAAAGTGAAATATGTCGAAATCCTTTTTGGAATAATACCAAATCGAAAATAAATGTCCGATAGCAAGTTGATCAGTTAATTCAATAAGAGATAAATGGGAGATAGCACTCGATTTAGTTGGTACCGCCCAACCGAATCCGATTCAATCGTTTACTCATTCAATGAGTAAATTTTCAAGTTCAGCCAATCTTTTTTTC---AAAAAAAAAATTGCAAGTAAATGAAATCGTGAATAAATGTGTTTCATTTCTCTATCATTATAGAAAAAAAGATCCATATTATATTACTTATGGAATTCGAACCCGAACTCGATTTATGATTCATTATTTCGATCTTATTGGCCTTTGTTCTTTATTTTTTATTTTTCATATAGATTTCCGTCTTTATATTATACCCTTTCGTAGATGAATTATGCTTATTTTCACATCTAGGATTTACATATACAACATATATTACTGTCAAGAGGGAATTTTTCTCAGTATTTAGATATTTAGATTCAAAATAAGAAAGGGATCAATTCAATTATAAACCCGCAAAACAAAGATTAGGATTGGGTTGGGTTGCGCTATATATATCAAAGAGTATACAATAATGATGTATTTGGTGAATCAAATACATGGTCTAATAACAAACCATTTTAACATAACATTTTAATGAGTTGATAATATTAATTGAATATTGAATATTTTTTGAAAGATTTTTGTTAAAGGTTTCATTCATGCCTAATCCATATCGAGTAGACCTTGTTGTTGTGAG

			>Tillandsia_koehresiana
			??????????????????????????????????????????????????????????????????????????????????????????????????????????????????????????????????????????????????????????????????????????????????????????????????????????????????????????????????????????????????????????????????????????????????????????????????????????????????????????????????????????????????????????????????????????????????????????????????????????????????????????????????????????????????????????????????????????????????????????????????????????????????????????????????????????????????????????????????????????????????????????????????????????????????????????????????????????????????????????????????????????????????????????????????????????????????????????????????????????????????????????????????????????????????????????????

			>Tillandsia_kuehhasii
			??????????????????????????????????????????????????????????????????????????????????????????????????????????????????????????????????????????????????????????????????????????????????????????????????????????????????????????????????????????????????????????????????????????????????????????????????????????????????????????????????????????????????????????????????????????????????????????????????????????????????????????????????????????????????????????????????????????????????????????????????????????????????????????????????????????????????????????????????????????????????????????????????????????????????????????????????????????????????????????????????????????????????????????????????????????????????????????????????????????????????????????????????????????????????????????????

			>Tillandsia_lajensis
			??????????????????????????????????????????????????????????????????????????????????????????????????????????????????????????????????????????????????????????????????????????????????????????????????????????????????????????????????????????????????????????????????????????????????????????????????????????????????????????????????????????????????????????????????????????????????????????????????????????????????????????????????????????????????????????????????????????????????????????????????????????????????????????????????????????????????????????????????????????????????????????????????????????????????????????????????????????????????????????????????????????????????????????????????????????????????????????????????????????????????????????????????????????????????????????????

			>Tillandsia_lampropoda
			??????????????????????????????????????????????????????????????????????????????????????????????????????????????????????????????????????????????????????????????????????????????????????????????????????????????????????????????????????????????????????????????????????????????????????????????????????????????????????????????????????????????????????????????????????????????????????????????????????????????????????????????????????????????????????????????????????????????????????????????????????????????????????????????????????????????????????????????????????????????????????????????????????????????????????????????????????????????????????????????????????????????????????????????????????????????????????????????????????????????????????????????????????????????????????????????

			>Tillandsia_landbeckii
			??????????????????????????????????????????????????????????????????????????????????????????????????????????????????????????????????????????????????????????????????????????????????????????????????????????????????????????????????????????????????????????????????????????????????????????????????????????????????????????????????????????????????????????????????????????????????????????????????????????????????????????????????????????????????????????????????????????????????????????????????????????????????????????????????????????????????????????????????????????????????????????????????????????????????????????????????????????????????????????????????????????????????????????????????????????????????????????????????????????????????????????????????????????????????????????????

			>Tillandsia_landbeckii_subsp._andina
			??????????????????????????????????????????????????????????????????????????????????????????????????????????????????????????????????????????????????????????????????????????????????????????????????????????????????????????????????????????????????????????????????????????????????????????????????????????????????????????????????????????????????????????????????????????????????????????????????????????????????????????????????????????????????????????????????????????????????????????????????????????????????????????????????????????????????????????????????????????????????????????????????????????????????????????????????????????????????????????????????????????????????????????????????????????????????????????????????????????????????????????????????????????????????????????????

			>Tillandsia_landbeckii_var._rigidor
			??????????????????????????????????????????????????????????????????????????????????????????????????????????????????????????????????????????????????????????????????????????????????????????????????????????????????????????????????????????????????????????????????????????????????????????????????????????????????????????????????????????????????????????????????????????????????????????????????????????????????????????????????????????????????????????????????????????????????????????????????????????????????????????????????????????????????????????????????????????????????????????????????????????????????????????????????????????????????????????????????????????????????????????????????????????????????????????????????????????????????????????????????????????????????????????????

			>Tillandsia_langlasseana
			??????????????????????????????????????????????????????????????????????????????????????????????????????????????????????????????????????????????????????????????????????????????????????????????????????????????????????????????????????????????????????????????????????????????????????????????????????????????????????????????????????????????????????????????????????????????????????????????????????????????????????????????????????????????????????????????????????????????????????????????????????????????????????????????????????????????????????????????????????????????????????????????????????????????????????????????????????????????????????????????????????????????????????????????????????????????????????????????????????????????????????????????????????????????????????????????

			>Tillandsia_latifolia_var._divaricata
			ATAATTATAATAAAGTGAAATATGTCGAAATCCTTTTTGGAATAATACCAAATCGAAAATAAATGTCCGATAGCAAGTTGATCAGTTAATTCAATAAGAGATAAATGGGAGATAGCACTCGATTTAGTTGGTACCACCCAACCGAATCCGATTCAATCGTTTACTCATTCAATGAGTAAATTTTCAAGTTCAGCCAATCTTTATTT---------------------------------------------------------------------------------------------------------------------------------------------------------------------TTTTTCATATAGATTTCCGTCTTTATATTATACCCTTTCGTAGATGAATTATGCTTATTTTCACATCTAGGATTTACATATACAACATATATTACTGTCAAGAGGGAATTTTTCTCAGTATTTAGATATTTAGATTCAAAATAAGAAAGGGATCAATTCAATTATAAACCCGCAAAACAAAAATTAGGATTGGGTTGGGTTGCGCTATATATATCAAAGAGTATACAATAATGATGTATTTGGTGAATCAAATACATGGTCTAATAACAAACCATTTTAACATAACATTTTAATGAGTTGATAATATTAATTGAATATTGAATATTTTTTGAAAGATTTTTGTTAAAGGTTTCATTCATGCCTAATCCATATCGAGTAGACCTTGTTGTTGTGAG

			>Tillandsia_laui
			??????????????????????????????????????????????????????????????????????????????????????????????????????????????????????????????????????????????????????????????????????????????????????????????????????????????????????????????????????????????????????????????????????????????????????????????????????????????????????????????????????????????????????????????????????????????????????????????????????????????????????????????????????????????????????????????????????????????????????????????????????????????????????????????????????????????????????????????????????????????????????????????????????????????????????????????????????????????????????????????????????????????????????????????????????????????????????????????????????????????????????????????????????????????????????????????

			>Tillandsia_lautneri
			??????????????????????????????????????????????????????????????????????????????????????????????????????????????????????????????????????????????????????????????????????????????????????????????????????????????????????????????????????????????????????????????????????????????????????????????????????????????????????????????????????????????????????????????????????????????????????????????????????????????????????????????????????????????????????????????????????????????????????????????????????????????????????????????????????????????????????????????????????????????????????????????????????????????????????????????????????????????????????????????????????????????????????????????????????????????????????????????????????????????????????????????????????????????????????????????

			>Tillandsia_lechneri
			??????????????????????????????????????????????????????????????????????????????????????????????????????????????????????????????????????????????????????????????????????????????????????????????????????????????????????????????????????????????????????????????????????????????????????????????????????????????????????????????????????????????????????????????????????????????????????????????????????????????????????????????????????????????????????????????????????????????????????????????????????????????????????????????????????????????????????????????????????????????????????????????????????????????????????????????????????????????????????????????????????????????????????????????????????????????????????????????????????????????????????????????????????????????????????????????

			>Tillandsia_leiboldiana
			??????????????????????????????????????????????????????????????????????????????????????????????????????????????????????????????????????????????????????????????????????????????????????????????????????????????????????????????????????????????????????????????????????????????????????????????????????????????????????????????????????????????????????????????????????????????????????????????????????????????????????????????????????????????????????????????????????????????????????????????????????????????????????????????????????????????????????????????????????????????????????????????????????????????????????????????????????????????????????????????????????????????????????????????????????????????????????????????????????????????????????????????????????????????????????????????

			>Tillandsia_lepidosepala
			ATAATTATAATAAAGTGAAATATGTCGAAATCCTTTTTGGAATAATACCAAATTGAAAATAAATGTCCGATAGCAAGTTGATCAGTTAATTCAATAAGAGATAAATGGGAGATAGCACTCGATTTAGTTGGTACCACCCAACCGAATCCGATTCAATCGTTTACTCATTCAATGAGTAAATTTTCAAGTTCAGCCAATCCTTTTTTTTTTCAAAAAAAAATTGCAAGTAAATGAAATCGTGAATAAATGTGTTTCATTTCTCTATCATTATAGAAAAAAAGATCCATATTATATTACTTATGGAATTCGAACCCGAACTCGATTTATGATTCATTATTTCGATCTTATTGGCCTTTGTTCTTTATTTTTTATTTTTCATATAGATTTCCGTCTTTATATTATACCCTTTCGTAGATGAATTATGCTTATTTTCACATCTAGGATTTACATATACAACATATATTACTGTCAAGAGGGAATTTTTCTCAGTATTTAGATATTTAGATTCAAAATAAGAAAGGGATCAATTCAATTATAAACCCGCAAAACAAAGATTAGGATTGGGTTGGGTTGCGCTATATATATCAAAGAGTATACAATAATGATGTATTTGGTGAATCAAATACATGGTCTAATAACAAACCATTTTAACATAACATTTTAATGAGTTGATAATATTAATTGAATATTGAATATTTTTTGAAAGATTTTTGTTAAAGGTTTCATTCATGCCTAATCCATATCGAGTAGACCTTGTTGTTGTGAG

			>Tillandsia_leucopetala
			??????????????????????????????????????????????????????????????????????????????????????????????????????????????????????????????????????????????????????????????????????????????????????????????????????????????????????????????????????????????????????????????????????????????????????????????????????????????????????????????????????????????????????????????????????????????????????????????????????????????????????????????????????????????????????????????????????????????????????????????????????????????????????????????????????????????????????????????????????????????????????????????????????????????????????????????????????????????????????????????????????????????????????????????????????????????????????????????????????????????????????????????????????????????????????????????

			>Tillandsia_limbata
			??????????????????????????????????????????????????????????????????????????????????????????????????????????????????????????????????????????????????????????????????????????????????????????????????????????????????????????????????????????????????????????????????????????????????????????????????????????????????????????????????????????????????????????????????????????????????????????????????????????????????????????????????????????????????????????????????????????????????????????????????????????????????????????????????????????????????????????????????????????????????????????????????????????????????????????????????????????????????????????????????????????????????????????????????????????????????????????????????????????????????????????????????????????????????????????????

			>Tillandsia_lorentziana
			??????????????????????????????????????????????????????????????????????????????????????????????????????????????????????????????????????????????????????????????????????????????????????????????????????????????????????????????????????????????????????????????????????????????????????????????????????????????????????????????????????????????????????????????????????????????????????????????????????????????????????????????????????????????????????????????????????????????????????????????????????????????????????????????????????????????????????????????????????????????????????????????????????????????????????????????????????????????????????????????????????????????????????????????????????????????????????????????????????????????????????????????????????????????????????????????

			>Tillandsia_lotteae
			??????????????????????????????????????????????????????????????????????????????????????????????????????????????????????????????????????????????????????????????????????????????????????????????????????????????????????????????????????????????????????????????????????????????????????????????????????????????????????????????????????????????????????????????????????????????????????????????????????????????????????????????????????????????????????????????????????????????????????????????????????????????????????????????????????????????????????????????????????????????????????????????????????????????????????????????????????????????????????????????????????????????????????????????????????????????????????????????????????????????????????????????????????????????????????????????

			>Tillandsia_lucida
			??????????????????????????????????????????????????????????????????????????????????????????????????????????????????????????????????????????????????????????????????????????????????????????????????????????????????????????????????????????????????????????????????????????????????????????????????????????????????????????????????????????????????????????????????????????????????????????????????????????????????????????????????????????????????????????????????????????????????????????????????????????????????????????????????????????????????????????????????????????????????????????????????????????????????????????????????????????????????????????????????????????????????????????????????????????????????????????????????????????????????????????????????????????????????????????????

			>Tillandsia_lymanii
			??????????????????????????????????????????????????????????????????????????????????????????????????????????????????????????????????????????????????????????????????????????????????????????????????????????????????????????????????????????????????????????????????????????????????????????????????????????????????????????????????????????????????????????????????????????????????????????????????????????????????????????????????????????????????????????????????????????????????????????????????????????????????????????????????????????????????????????????????????????????????????????????????????????????????????????????????????????????????????????????????????????????????????????????????????????????????????????????????????????????????????????????????????????????????????????????

			>Tillandsia_macbrideana
			ATAATTATAATAAAGTGAAATATGTCGAAATCCTTTTTGGAATAATACCAAATCGAAAATAAATGTCCGATAGCAAGTTGATCAGTTAATTCAATAAGAGATAAATGGGAGATAGCACTCGATTTAGTTGGTACCACCCAACCGAATCCGATTCAATCGTTTACTCATTCAATGAGTAAATTTTCAAGTTCAGCCAATCTTTATTTTT-------------------------------------------------------------------------------------------------------------------------------------------------------------------TTTTTCATATAGATTTCCGTCTTTATATTATACCCTTTCGTAGATGAATTATGCTTATTTTCACATCTAGGATTTACATATACAACATATATTACTGTCAAGAGGGAATTTTTCTCAGTATTTAGATATTTAGATTCAAAATAAGAAAGGGATCAATTCAATTATAAACCCGCAAAACAAAGATTAGGATTGGGTTGGGTTGCGCTATATATATCAAAGAGTATACAATAATGATGTATTTGGTGAATCAAATACATGGTCTAATAACAAACCATTTTAACATAACATTTTAATGAGTTGATAATATTAATTGAATATTGAATATTTTTTGAAAGATTTTTGTTAAAGGTTTCATTCATGCCTAATCCATATCGAGTAGACCTTGTTGTTGTGAG

			>Tillandsia_macdougallii
			??????????????????????????????????????????????????????????????????????????????????????????????????????????????????????????????????????????????????????????????????????????????????????????????????????????????????????????????????????????????????????????????????????????????????????????????????????????????????????????????????????????????????????????????????????????????????????????????????????????????????????????????????????????????????????????????????????????????????????????????????????????????????????????????????????????????????????????????????????????????????????????????????????????????????????????????????????????????????????????????????????????????????????????????????????????????????????????????????????????????????????????????????????????????????????????????

			>Tillandsia_maculata
			??????????????????????????????????????????????????????????????????????????????????????????????????????????????????????????????????????????????????????????????????????????????????????????????????????????????????????????????????????????????????????????????????????????????????????????????????????????????????????????????????????????????????????????????????????????????????????????????????????????????????????????????????????????????????????????????????????????????????????????????????????????????????????????????????????????????????????????????????????????????????????????????????????????????????????????????????????????????????????????????????????????????????????????????????????????????????????????????????????????????????????????????????????????????????????????????

			>Tillandsia_magnusiana
			??????????????????????????????????????????????????????????????????????????????????????????????????????????????????????????????????????????????????????????????????????????????????????????????????????????????????????????????????????????????????????????????????????????????????????????????????????????????????????????????????????????????????????????????????????????????????????????????????????????????????????????????????????????????????????????????????????????????????????????????????????????????????????????????????????????????????????????????????????????????????????????????????????????????????????????????????????????????????????????????????????????????????????????????????????????????????????????????????????????????????????????????????????????????????????????????

			>Tillandsia_makoyana
			??????????????????????????????????????????????????????????????????????????????????????????????????????????????????????????????????????????????????????????????????????????????????????????????????????????????????????????????????????????????????????????????????????????????????????????????????????????????????????????????????????????????????????????????????????????????????????????????????????????????????????????????????????????????????????????????????????????????????????????????????????????????????????????????????????????????????????????????????????????????????????????????????????????????????????????????????????????????????????????????????????????????????????????????????????????????????????????????????????????????????????????????????????????????????????????????

			>Tillandsia_malzinei
			ATAATTATAATAAAGTGAAATATGTCGAAATCCTTTTTGGAATAATACCAAATCAAAAATAAATGTCCAATAGCAAGTTGATCAGTTAATTCAATAAGAGATAAATGGGAGATAGCACTCGATTTAGTTGGTACCACCCAACCGAATCCGATTCAATCATTTACTCATTCAATGAGTAAATTTTCAAGTTCAGTCAATCCTTTTTTTTTT--------AATTGCAAGTAAATGAAATCGTGAATAAATGTGTTTCATTTCTCTATCATTATAGAAAAAAAGATCCATATTATATTACTTATGGAATTCGAACCCGAACTCGATTTATGATTCATTATTTCGATCTTATTGGCCTTTGTTCTTTATTTTTTATTTTTCATATAGATTTCCGTCTTTATATTATACCCTTTCGTAGATGAATTATGCTTATTTTCACATCTAGGATTTACATATACAACATATATTACTGTCAAGAGGGAATTTTTCTCAGTATTTAGATATTTAGATTCAAAATAAGAAAGGGATCAATTCAATTATAAACCCGCAAAACAAAGATTAGGATTGGGTTGGGTTGCGCTATATATATCAAAGAGTATACAATAATGATGTATTTGGTGAATCAAATACATGGTCTAATAACAAACCATTTTAACATAACATTTTAATGAGTTGATAATATTAATTGAATATTGAATATTTTTTGAAAGATTTTTGTTAAAGGTTTCATTCATGCCTAATCCATATCGAGTAGACCTTGTTGTTGTGAG

			>Tillandsia_marconae
			ATAATTATAATAAAGTGAAATATGTCGAAATCCTTTTTGGAATAATACCAAATCGAAAATAAATGTCCGATAGCAAGTTGATCAGTTAATTCAATAAGAGATAAATGGGAGATAGCACTCGATTTAGTTGGTACCACCCAACCGAATCCGATTCAATCGTTTACTCATTCAATGAGTAAATTTGCAAGTTCAGCCAATCTTTTTTTTG---AAAAAAAAATTGCAAGTAAATGAAATCGTGAATAAATGTGTTTCATTTCTCTATCATTATAGAAAAAAAGATCCATATTATATTACTTATGGAATTCGAACCCGAACTCGATTTATGATTCATTATTTCGATCTTATTGGCCTTTGTTCTTTATTTTTTATTTTTCATATAGATTTCCGTCTTTATATTATACCCTTTCGTAGATGAATTATGCTTATTTTCACATCTAGGATTTACATATACAACATATATTACTGTCAAGAGGGAATTTTTCTCAGTATTTAGATATTTAGATTCAAAATAAGAAAGGG-----TTCAATTATAAACCCGCAAAACAAAGATTAGGATTGGGTTGGGTTGCGCTATATATATCAAAGAGTATACAATAATGATGTATTTGGTGAATCAAATACATGGTCTAATAACAAACCATTTTAACATAACATTTTAATGAGTTGATAATATTAATTGAATATTGAATATTTTTTGAAAGATTTTTGTTAAAGGTTTCATTCATGCCTAATCCATATCGAGTAGACCTTGTTGTTGTGAG

			>Tillandsia_marnier-lapostollei
			??????????????????????????????????????????????????????????????????????????????????????????????????????????????????????????????????????????????????????????????????????????????????????????????????????????????????????????????????????????????????????????????????????????????????????????????????????????????????????????????????????????????????????????????????????????????????????????????????????????????????????????????????????????????????????????????????????????????????????????????????????????????????????????????????????????????????????????????????????????????????????????????????????????????????????????????????????????????????????????????????????????????????????????????????????????????????????????????????????????????????????????????????????????????????????????????

			>Tillandsia_matudae
			ATAATTATAATAAAGTGAAATATGTCGAAATCCTTTTTGGAATAATACCAAATCAAAAATAAATGTCCGATAGCAAGTTGATCAGTTAATTCAATAAGAGATAAATGGGAGATAGCACTCGATTTAGTTGGTACCGCCCAACCGAATCCGATTCAATCGTTTACTCATTCAATGAGTAAATTTTCAAGTTCAGCCAATCTTTTTTTC---AAAAAAAAAATTGCAAGTAAATGAAATCGTGAATAAATGTGTTTCATTTCTCTATCATTATAGAAAAAAAGATCCATATTATATTACTTATGGAATTCGAACCCGAACTCGATTTATGATTCATTATTTCGATCTTATTGGCCTTTGTTCTTTATTTTTTATTTTTCATATAGATTTCCGTCTTTATATTATACCCTTTCGTAGATGAATTATGCTTATTTTCACATCTAGGATTTACATATACAACATATATTACTGTCAAGAGGGAATTTTTCTCAGTATTTAGATATTTAGATTCAAAATAAGAAAGGGATCAATTCAATTATAAACCCGCAAAACAAAGATTAGGATTGGGTTGGGTTGCGCTATATATATCAAAGAGTATACAATAATGATGTATTTGGTGAATCAAATACATGGTCTAATAACAAACCATTTTAACATAACATTTTAATGAGTTGATAATATTAATTGAATATTGAATATTTTTTGAAAGATTTTTGTTAAAGGTTTCATTCATGCCTAATCCATATCGAGTAGACCTTGTTGTTGTGAG

			>Tillandsia_may-patii
			??????????????????????????????????????????????????????????????????????????????????????????????????????????????????????????????????????????????????????????????????????????????????????????????????????????????????????????????????????????????????????????????????????????????????????????????????????????????????????????????????????????????????????????????????????????????????????????????????????????????????????????????????????????????????????????????????????????????????????????????????????????????????????????????????????????????????????????????????????????????????????????????????????????????????????????????????????????????????????????????????????????????????????????????????????????????????????????????????????????????????????????????????????????????????????????????

			>Tillandsia_mazatlanensis
			??????????????????????????????????????????????????????????????????????????????????????????????????????????????????????????????????????????????????????????????????????????????????????????????????????????????????????????????????????????????????????????????????????????????????????????????????????????????????????????????????????????????????????????????????????????????????????????????????????????????????????????????????????????????????????????????????????????????????????????????????????????????????????????????????????????????????????????????????????????????????????????????????????????????????????????????????????????????????????????????????????????????????????????????????????????????????????????????????????????????????????????????????????????????????????????????

			>Tillandsia_milagrensis
			??????????????????????????????????????????????????????????????????????????????????????????????????????????????????????????????????????????????????????????????????????????????????????????????????????????????????????????????????????????????????????????????????????????????????????????????????????????????????????????????????????????????????????????????????????????????????????????????????????????????????????????????????????????????????????????????????????????????????????????????????????????????????????????????????????????????????????????????????????????????????????????????????????????????????????????????????????????????????????????????????????????????????????????????????????????????????????????????????????????????????????????????????????????????????????????????

			>Tillandsia_mima
			??????????????????????????????????????????????????????????????????????????????????????????????????????????????????????????????????????????????????????????????????????????????????????????????????????????????????????????????????????????????????????????????????????????????????????????????????????????????????????????????????????????????????????????????????????????????????????????????????????????????????????????????????????????????????????????????????????????????????????????????????????????????????????????????????????????????????????????????????????????????????????????????????????????????????????????????????????????????????????????????????????????????????????????????????????????????????????????????????????????????????????????????????????????????????????????????

			>Tillandsia_mirabilis
			??????????????????????????????????????????????????????????????????????????????????????????????????????????????????????????????????????????????????????????????????????????????????????????????????????????????????????????????????????????????????????????????????????????????????????????????????????????????????????????????????????????????????????????????????????????????????????????????????????????????????????????????????????????????????????????????????????????????????????????????????????????????????????????????????????????????????????????????????????????????????????????????????????????????????????????????????????????????????????????????????????????????????????????????????????????????????????????????????????????????????????????????????????????????????????????????

			>Tillandsia_mollis
			??????????????????????????????????????????????????????????????????????????????????????????????????????????????????????????????????????????????????????????????????????????????????????????????????????????????????????????????????????????????????????????????????????????????????????????????????????????????????????????????????????????????????????????????????????????????????????????????????????????????????????????????????????????????????????????????????????????????????????????????????????????????????????????????????????????????????????????????????????????????????????????????????????????????????????????????????????????????????????????????????????????????????????????????????????????????????????????????????????????????????????????????????????????????????????????????

			>Tillandsia_mooreana
			??????????????????????????????????????????????????????????????????????????????????????????????????????????????????????????????????????????????????????????????????????????????????????????????????????????????????????????????????????????????????????????????????????????????????????????????????????????????????????????????????????????????????????????????????????????????????????????????????????????????????????????????????????????????????????????????????????????????????????????????????????????????????????????????????????????????????????????????????????????????????????????????????????????????????????????????????????????????????????????????????????????????????????????????????????????????????????????????????????????????????????????????????????????????????????????????

			>Tillandsia_moronesensis
			??????????????????????????????????????????????????????????????????????????????????????????????????????????????????????????????????????????????????????????????????????????????????????????????????????????????????????????????????????????????????????????????????????????????????????????????????????????????????????????????????????????????????????????????????????????????????????????????????????????????????????????????????????????????????????????????????????????????????????????????????????????????????????????????????????????????????????????????????????????????????????????????????????????????????????????????????????????????????????????????????????????????????????????????????????????????????????????????????????????????????????????????????????????????????????????????

			>Tillandsia_multicaulis
			ATAATTATAATAAAGTGAAATATGTCGAAATCCTTTTTGGAATAATACCAAATCAAAAATAAATGTCCGATAGCAAGTTGATCAGTTAATTCAATAAGAGATAAATGGGAGATAGCACTCGATTTAGTTGGTACCACCCAACCGAATCCGATTCAATCGTTTACTCATTCAATGAGTAAATTTTCAAGTTCAGCCAATCTTTTTTTTT----------AATTGCAAGTAAATGAAATCGTGAATAAATGTGTTTCATTTCTCTATCATTATAGAAAAAAAGATCCATATTATATTACTTATGGAATTCGAACCCGAACTCGATTTATGATTCATTATTTCGATCTTATTGGCCTTTGTTCTTTATTTTTTATTTTTCATATAGATTTCCGTCTTTATATTATACCCTTTCGTAGATGAATTATGCTTATTTTCACATCTAGGATTTACATATACAACATATATTACTGTCAAGAGGGAATTTTTCTCAGTATTTAGATATTTAGATTCAAAATAAGAAAGGG-----TTCAATTATAAACCCGCAAAACAAAGATTAGGATTGGGTTGGGTTGCGCTATATATATCAAAGAGTATACAATAATGATGTATTTGGTGAATCAAATACATGGTCTAATAACAAACCATTTTAACATAACATTTTAATGAGTTGATAATATTAATTGAATATTGAATATTTTTTGAAAGATTTTTGTTAAAGGTTTCATTCATGCCTAATCCATATCGAGTAGACCTTGTTGTTGTGAG

			>Tillandsia_myriantha
			??????????????????????????????????????????????????????????????????????????????????????????????????????????????????????????????????????????????????????????????????????????????????????????????????????????????????????????????????????????????????????????????????????????????????????????????????????????????????????????????????????????????????????????????????????????????????????????????????????????????????????????????????????????????????????????????????????????????????????????????????????????????????????????????????????????????????????????????????????????????????????????????????????????????????????????????????????????????????????????????????????????????????????????????????????????????????????????????????????????????????????????????????????????????????????????????

			>Tillandsia_nana
			??????????????????????????????????????????????????????????????????????????????????????????????????????????????????????????????????????????????????????????????????????????????????????????????????????????????????????????????????????????????????????????????????????????????????????????????????????????????????????????????????????????????????????????????????????????????????????????????????????????????????????????????????????????????????????????????????????????????????????????????????????????????????????????????????????????????????????????????????????????????????????????????????????????????????????????????????????????????????????????????????????????????????????????????????????????????????????????????????????????????????????????????????????????????????????????????

			>Tillandsia_neglecta
			??????????????????????????????????????????????????????????????????????????????????????????????????????????????????????????????????????????????????????????????????????????????????????????????????????????????????????????????????????????????????????????????????????????????????????????????????????????????????????????????????????????????????????????????????????????????????????????????????????????????????????????????????????????????????????????????????????????????????????????????????????????????????????????????????????????????????????????????????????????????????????????????????????????????????????????????????????????????????????????????????????????????????????????????????????????????????????????????????????????????????????????????????????????????????????????????

			>Tillandsia_nicolasensis
			??????????????????????????????????????????????????????????????????????????????????????????????????????????????????????????????????????????????????????????????????????????????????????????????????????????????????????????????????????????????????????????????????????????????????????????????????????????????????????????????????????????????????????????????????????????????????????????????????????????????????????????????????????????????????????????????????????????????????????????????????????????????????????????????????????????????????????????????????????????????????????????????????????????????????????????????????????????????????????????????????????????????????????????????????????????????????????????????????????????????????????????????????????????????????????????????

			>Tillandsia_nizandaensis
			??????????????????????????????????????????????????????????????????????????????????????????????????????????????????????????????????????????????????????????????????????????????????????????????????????????????????????????????????????????????????????????????????????????????????????????????????????????????????????????????????????????????????????????????????????????????????????????????????????????????????????????????????????????????????????????????????????????????????????????????????????????????????????????????????????????????????????????????????????????????????????????????????????????????????????????????????????????????????????????????????????????????????????????????????????????????????????????????????????????????????????????????????????????????????????????????

			>Tillandsia_novakii
			??????????????????????????????????????????????????????????????????????????????????????????????????????????????????????????????????????????????????????????????????????????????????????????????????????????????????????????????????????????????????????????????????????????????????????????????????????????????????????????????????????????????????????????????????????????????????????????????????????????????????????????????????????????????????????????????????????????????????????????????????????????????????????????????????????????????????????????????????????????????????????????????????????????????????????????????????????????????????????????????????????????????????????????????????????????????????????????????????????????????????????????????????????????????????????????????

			>Tillandsia_nuptialis
			??????????????????????????????????????????????????????????????????????????????????????????????????????????????????????????????????????????????????????????????????????????????????????????????????????????????????????????????????????????????????????????????????????????????????????????????????????????????????????????????????????????????????????????????????????????????????????????????????????????????????????????????????????????????????????????????????????????????????????????????????????????????????????????????????????????????????????????????????????????????????????????????????????????????????????????????????????????????????????????????????????????????????????????????????????????????????????????????????????????????????????????????????????????????????????????????

			>Tillandsia_oaxacana
			??????????????????????????????????????????????????????????????????????????????????????????????????????????????????????????????????????????????????????????????????????????????????????????????????????????????????????????????????????????????????????????????????????????????????????????????????????????????????????????????????????????????????????????????????????????????????????????????????????????????????????????????????????????????????????????????????????????????????????????????????????????????????????????????????????????????????????????????????????????????????????????????????????????????????????????????????????????????????????????????????????????????????????????????????????????????????????????????????????????????????????????????????????????????????????????????

			>Tillandsia_oblivata
			??????????????????????????????????????????????????????????????????????????????????????????????????????????????????????????????????????????????????????????????????????????????????????????????????????????????????????????????????????????????????????????????????????????????????????????????????????????????????????????????????????????????????????????????????????????????????????????????????????????????????????????????????????????????????????????????????????????????????????????????????????????????????????????????????????????????????????????????????????????????????????????????????????????????????????????????????????????????????????????????????????????????????????????????????????????????????????????????????????????????????????????????????????????????????????????????

			>Tillandsia_oerstediana
			??????????????????????????????????????????????????????????????????????????????????????????????????????????????????????????????????????????????????????????????????????????????????????????????????????????????????????????????????????????????????????????????????????????????????????????????????????????????????????????????????????????????????????????????????????????????????????????????????????????????????????????????????????????????????????????????????????????????????????????????????????????????????????????????????????????????????????????????????????????????????????????????????????????????????????????????????????????????????????????????????????????????????????????????????????????????????????????????????????????????????????????????????????????????????????????????

			>Tillandsia_paleacea
			??????????????????????????????????????????????????????????????????????????????????????????????????????????????????????????????????????????????????????????????????????????????????????????????????????????????????????????????????????????????????????????????????????????????????????????????????????????????????????????????????????????????????????????????????????????????????????????????????????????????????????????????????????????????????????????????????????????????????????????????????????????????????????????????????????????????????????????????????????????????????????????????????????????????????????????????????????????????????????????????????????????????????????????????????????????????????????????????????????????????????????????????????????????????????????????????

			>Tillandsia_pamelae
			??????????????????????????????????????????????????????????????????????????????????????????????????????????????????????????????????????????????????????????????????????????????????????????????????????????????????????????????????????????????????????????????????????????????????????????????????????????????????????????????????????????????????????????????????????????????????????????????????????????????????????????????????????????????????????????????????????????????????????????????????????????????????????????????????????????????????????????????????????????????????????????????????????????????????????????????????????????????????????????????????????????????????????????????????????????????????????????????????????????????????????????????????????????????????????????????

			>Tillandsia_paniculata
			ATAATTATAATAAAGTGAAATATGTCGAAATCCTTTTTGGAATAATACCAAATCGAAAATAAATGTCCGATAGCAAGTTGATCAGTTAATTCAATAAGAGATAAATGGGAGATAGCACTCGATTTAGTTGGTACCACCCAACCGAATCCGATTCAATCGTTTACTCATTCAATGAGTAAATTTTCAAGTTCAGCCAATCTTTTTTTTTTT------TTAATTGCAAGTAAATGAAATCGTGAATAAATGTGTTTCATTTCTCTATCATTATAGAAAAAAAGATCCATATTATATTACTTATGGAATTCGAACCCGAACTCGATTTATGATTCATTATTTCGATCTTATTGGCCTTTGTTCTTTATTTTTTATTTTTCATATAGATTTCCGTCTTTATATTATACCCTTTCGTAGATGAATTATGCTTATTTTCACATCTAGGATTTACATATACAACATATATTACTGTCAAGAGAGAATTTTTCTCAGTATTTAGATATTTAGATTCAAAATAAGAAAGGGATCAATTCAATTATAAACCTGCAAAACAAAGATTAGGATTGGGTTGGGTTGCGCTATATATATCAAAGAGTATACAATAATGATGTATTTGGTGAATCAAATACATGGTCTAATAACAAACCATTTTAACATAACATTTTAATGAGTTGATAATATTAATTGAATATTGAATATTTTTTGAAAGATTTTTGTTAAAGGTTTCATTCATGCCTAATCCATATCGAGTAGACCTTGTTGTTGTGAG

			>Tillandsia_paraensis
			??????????????????????????????????????????????????????????????????????????????????????????????????????????????????????????????????????????????????????????????????????????????????????????????????????????????????????????????????????????????????????????????????????????????????????????????????????????????????????????????????????????????????????????????????????????????????????????????????????????????????????????????????????????????????????????????????????????????????????????????????????????????????????????????????????????????????????????????????????????????????????????????????????????????????????????????????????????????????????????????????????????????????????????????????????????????????????????????????????????????????????????????????????????????????????????????

			>Tillandsia_paraibensis
			??????????????????????????????????????????????????????????????????????????????????????????????????????????????????????????????????????????????????????????????????????????????????????????????????????????????????????????????????????????????????????????????????????????????????????????????????????????????????????????????????????????????????????????????????????????????????????????????????????????????????????????????????????????????????????????????????????????????????????????????????????????????????????????????????????????????????????????????????????????????????????????????????????????????????????????????????????????????????????????????????????????????????????????????????????????????????????????????????????????????????????????????????????????????????????????????

			>Tillandsia_paraisoensis
			??????????????????????????????????????????????????????????????????????????????????????????????????????????????????????????????????????????????????????????????????????????????????????????????????????????????????????????????????????????????????????????????????????????????????????????????????????????????????????????????????????????????????????????????????????????????????????????????????????????????????????????????????????????????????????????????????????????????????????????????????????????????????????????????????????????????????????????????????????????????????????????????????????????????????????????????????????????????????????????????????????????????????????????????????????????????????????????????????????????????????????????????????????????????????????????????

			>Tillandsia_parryi
			??????????????????????????????????????????????????????????????????????????????????????????????????????????????????????????????????????????????????????????????????????????????????????????????????????????????????????????????????????????????????????????????????????????????????????????????????????????????????????????????????????????????????????????????????????????????????????????????????????????????????????????????????????????????????????????????????????????????????????????????????????????????????????????????????????????????????????????????????????????????????????????????????????????????????????????????????????????????????????????????????????????????????????????????????????????????????????????????????????????????????????????????????????????????????????????????

			>Tillandsia_paucifolia
			??????????????????????????????TCCTTTTTGGAATAATACCAAATCGAAAATAAATGTCCGATAGCAAGTTGATCAGTTAATTCAATAAGAGATAAATGGGAGATAGCACTCGATTTAGTTGGTACCGCCCAACCGAATCCGATTCAATCGTTTACTCATTCAATGAGTAAATTTTCAAGTTCAGCCAATC--CTTTTTTTCAAAAAAAAAATTGCAAGTAAATGAAATCGTGAATAAATGTGTTTCATTTCTCTATCATTATAGAAAAAAAGATCCATATTATATTACTTATGGAATTCGAACCCGAACTCGATTTATGATTCATTATTTCGATCTTATTGGCCTTTGTTCTTTATTTTTTATTTTTCATATAGATTTCCGTCTTTATATTATACCCTTTCGTAGATGAATTATGCTTATTTTCACATCTAGGATTTACATATACAACATATATTACTGTCAAGAGGGAATTTTTCTCAGTATTTAGATATTTAGATTCAAAATAAGAAAGGGATCAATTCAATTATAAACCCGCAAAACAAAGATTAGGATTGGGTTGGGTTGCGCTATATATATCAAAGAGTATACAATAATGATGTATTTGGTGAATCAAATACATGGTCTAATAACAAACCATTTTAACATAACATTTTAATGAGTTGATAATATTAATTGAATATTGAATATTTTTTGAAAGATTTTTGTTAAAGGTTTCATTCATGCCTAATCCATATCGAGTAGACCTTGTTGTTGTGAG

			>Tillandsia_pinicola
			??????????????????????????????????????????????????????????????????????????????????????????????????????????????????????????????????????????????????????????????????????????????????????????????????????????????????????????????????????????????????????????????????????????????????????????????????????????????????????????????????????????????????????????????????????????????????????????????????????????????????????????????????????????????????????????????????????????????????????????????????????????????????????????????????????????????????????????????????????????????????????????????????????????????????????????????????????????????????????????????????????????????????????????????????????????????????????????????????????????????????????????????????????????????????????????????

			>Tillandsia_plagiotropica
			??????????????????????????????????????????????????????????????????????????????????????????????????????????????????????????????????????????????????????????????????????????????????????????????????????????????????????????????????????????????????????????????????????????????????????????????????????????????????????????????????????????????????????????????????????????????????????????????????????????????????????????????????????????????????????????????????????????????????????????????????????????????????????????????????????????????????????????????????????????????????????????????????????????????????????????????????????????????????????????????????????????????????????????????????????????????????????????????????????????????????????????????????????????????????????????????

			>Tillandsia_plumosa
			ATAATTATAATAAAGTGAAATATGTCGAAATCCTTTTTGGAATAATACCAAATTGAAAATAAATGTCCGATAGCAAGTTGATCAGTTAATTCAATAAGAGATAAATGGGAGATAGCACTCGATTTAGTTGGTACCACCCAACCGAATCCGATTCAATCGTTTACTCATTCAATGAGTAAATTTTCAAGTTCAGCCAATCTTTTTTTTTTT------AAAATTGCAAGTAAATGAAATCGTGAATAAATGTGTTTCATTTCTCTATCATTATAGAAAAAAAGATCCATATTATATTACTTATGGAATTCGAACCCGAACTCGATTTATGATTCATTATTTCGATCTTATTGGCCTTTGTTCTTTATTTTTTATTTTTCATATAGATTTCCGTCTTTATATTATACCCTTTCGTAGATGAATTATGCTTATTTTCACATCTAGGATTTACATATACAACATATATTACTGTCAAGAGGGAATTTTTCTCAGTATTTAGATATTTAGATTCAAAATAAGAAAGGGATCAATTCAATTATAAACCCGCAAAACAAAGATTAGGATTGGGTTGGGTTGCGCTATATATATCAAAGAGTATACAATAATGATGTATTTGGTGAATCAAATACATGGTCTAATAACAAACCATTTTAACATAACATTTTAATGAGTTGATAATATTAATTGAATATTGAATATTTTTTGAAAGATTTTTGTTAAAGGTTTCATTCATGCCTAATCCATATCGAGTAGACCTTGTTGTTGTGAG

			>Tillandsia_pohliana
			ATAATTATAATAAAGTGAAATATGTCGAAATCCTTTTTGGAATAATACCAAATCGAAAATAAATGTCCGATAACAAGTTGATCAGTTAATTCAATAAGAGATAAATGGGAGATAGCACTCGATTTAGTTGGTACCACCTAACCGAATCCGATTCAATCGTTTACTCATTCAATGAGTAAATTTTCAAGTTCAGCCAATCTTTATTTTT-------------------------------------------------------------------------------------------------------------------------------------------------------------------TTTTTCATATAGATTTCCGTCTTTATATTATACCCTTTCGTAGATGAATTATGCTTACTTTCACATCTAGGATTTACATATACAACATATATTACTGTCAAGAGGGAATTTTTCTCAGTATTTAGATATTTAGATTCCAAATAAGAAAGGGATCAATTCAATTATAAACCCGCAAAACAAAGATTAGGATTGGGTTGGGTTGCGCTATATATATCAAAGAGTATACAATAATGATGTATTTGGTGAATCAAATACATGGTCTAATAACAAACCATTTTAACATAACATTTTAATGAGTTGATAATATTAATTGACTATTGAATATTTTTTGAAAGATTTTTGTTAAAGGTTTCATTCATGCCTAATCCATATCGAGTAGACCTTGTTGTTGTGAG

			>Tillandsia_polita
			??????????????????????????????????????????????????????????????????????????????????????????????????????????????????????????????????????????????????????????????????????????????????????????????????????????????????????????????????????????????????????????????????????????????????????????????????????????????????????????????????????????????????????????????????????????????????????????????????????????????????????????????????????????????????????????????????????????????????????????????????????????????????????????????????????????????????????????????????????????????????????????????????????????????????????????????????????????????????????????????????????????????????????????????????????????????????????????????????????????????????????????????????????????????????????????????

			>Tillandsia_polystachia
			??????????????????????????????????????????????????????????????????????????????????????????????????????????????????????????????????????????????????????????????????????????????????????????????????????????????????????????????????????????????????????????????????????????????????????????????????????????????????????????????????????????????????????????????????????????????????????????????????????????????????????????????????????????????????????????????????????????????????????????????????????????????????????????????????????????????????????????????????????????????????????????????????????????????????????????????????????????????????????????????????????????????????????????????????????????????????????????????????????????????????????????????????????????????????????????????

			>Tillandsia_polzii
			??????????????????????????????????????????????????????????????????????????????????????????????????????????????????????????????????????????????????????????????????????????????????????????????????????????????????????????????????????????????????????????????????????????????????????????????????????????????????????????????????????????????????????????????????????????????????????????????????????????????????????????????????????????????????????????????????????????????????????????????????????????????????????????????????????????????????????????????????????????????????????????????????????????????????????????????????????????????????????????????????????????????????????????????????????????????????????????????????????????????????????????????????????????????????????????????

			>Tillandsia_ponderosa
			??????????????????????????????????????????????????????????????????????????????????????????????????????????????????????????????????????????????????????????????????????????????????????????????????????????????????????????????????????????????????????????????????????????????????????????????????????????????????????????????????????????????????????????????????????????????????????????????????????????????????????????????????????????????????????????????????????????????????????????????????????????????????????????????????????????????????????????????????????????????????????????????????????????????????????????????????????????????????????????????????????????????????????????????????????????????????????????????????????????????????????????????????????????????????????????????

			>Tillandsia_prodigiosa
			??????????????????????????????????????????????????????????????????????????????????????????????????????????????????????????????????????????????????????????????????????????????????????????????????????????????????????????????????????????????????????????????????????????????????????????????????????????????????????????????????????????????????????????????????????????????????????????????????????????????????????????????????????????????????????????????????????????????????????????????????????????????????????????????????????????????????????????????????????????????????????????????????????????????????????????????????????????????????????????????????????????????????????????????????????????????????????????????????????????????????????????????????????????????????????????????

			>Tillandsia_prolata
			??????????????????????????????????????????????????????????????????????????????????????????????????????????????????????????????????????????????????????????????????????????????????????????????????????????????????????????????????????????????????????????????????????????????????????????????????????????????????????????????????????????????????????????????????????????????????????????????????????????????????????????????????????????????????????????????????????????????????????????????????????????????????????????????????????????????????????????????????????????????????????????????????????????????????????????????????????????????????????????????????????????????????????????????????????????????????????????????????????????????????????????????????????????????????????????????

			>Tillandsia_propagulifera
			??????????????????????????????????????????????????????????????????????????????????????????????????????????????????????????????????????????????????????????????????????????????????????????????????????????????????????????????????????????????????????????????????????????????????????????????????????????????????????????????????????????????????????????????????????????????????????????????????????????????????????????????????????????????????????????????????????????????????????????????????????????????????????????????????????????????????????????????????????????????????????????????????????????????????????????????????????????????????????????????????????????????????????????????????????????????????????????????????????????????????????????????????????????????????????????????

			>Tillandsia_pseudocardenasii
			??????????????????????????????????????????????????????????????????????????????????????????????????????????????????????????????????????????????????????????????????????????????????????????????????????????????????????????????????????????????????????????????????????????????????????????????????????????????????????????????????????????????????????????????????????????????????????????????????????????????????????????????????????????????????????????????????????????????????????????????????????????????????????????????????????????????????????????????????????????????????????????????????????????????????????????????????????????????????????????????????????????????????????????????????????????????????????????????????????????????????????????????????????????????????????????????

			>Tillandsia_pseudomacbrideana
			ATAATTATAATAAAGTGAAATATGTCGAAATCCTTTTTGGAATAATACCAAATCGAAAATAAATGTCCGATAGCAAGTTGATCAGTTAATTCAATAAGAGATAAATGGGAGATAGCACTCGATTTAGTTGGTACCACCCAACCGAATCCGATTCAATCGTTTACTCATTCAATGAGTAAATTTTCAAGTTCAGCCAATCTTTATTTTTT------------------------------------------------------------------------------------------------------------------------------------------------------------------TTTTTCATATAGATTTCCGTCTTTATATTATACCCTTTCGTAGATGAATTATGCTTATTTTCACATCTAGGATTTACATATACAACATATATTACTGTCAAGAGGGAATTTTTCTCAGTATTTAGATATTTAGATTCAAAATAAGAAAGGGATCAATTCAATTATAAACCCGCAAAACAAAGATTAGGATTGGGTTGGGTTGCGCTATATATATCAAAGAGTATACAATAATGATGTATTTGGTGAATCAAATACATGGTCTAATAACAAACCATTTTAACATAACATTTTAATGAGTTGATAATATTAATTGAATATTGAATATTTTTTGAAAGATTTTTGTTAAAGGTTTCATTCATGCCTAATCCATATCGAGTAGACCTTGTTGTTGTGAG

			>Tillandsia_pseudomicans
			??????????????????????????????????????????????????????????????????????????????????????????????????????????????????????????????????????????????????????????????????????????????????????????????????????????????????????????????????????????????????????????????????????????????????????????????????????????????????????????????????????????????????????????????????????????????????????????????????????????????????????????????????????????????????????????????????????????????????????????????????????????????????????????????????????????????????????????????????????????????????????????????????????????????????????????????????????????????????????????????????????????????????????????????????????????????????????????????????????????????????????????????????????????????????????????????

			>Tillandsia_pueblensis
			??????????????????????????????????????????????????????????????????????????????????????????????????????????????????????????????????????????????????????????????????????????????????????????????????????????????????????????????????????????????????????????????????????????????????????????????????????????????????????????????????????????????????????????????????????????????????????????????????????????????????????????????????????????????????????????????????????????????????????????????????????????????????????????????????????????????????????????????????????????????????????????????????????????????????????????????????????????????????????????????????????????????????????????????????????????????????????????????????????????????????????????????????????????????????????????????

			>Tillandsia_punctulata
			ATAATTATAATAAAGTGAAATATGTCGAAATCCTTTTTGGAATAATACCAAATCGAAAATAAATGTCCGATAACAAGTTGATCAGTTAATTCAATAAGAGATAAATGGGAGATAGCACTCGATTTAGTTGGTACCGCCCAACCGAATCCGATTCAATCGTTTACTCATTCAATGAGTAAATTTTCAAGTTCAGCCAATCTTTTTTTC---AAAAAAAAAATTGCAAGTAAATGAAATCGTGAATAAATGTGTTTCATTTCTCTATCATTATAGAAAAAAAGATCCATATTATATTACTTATGGAATTCGAACCCGAACTCGATTTATGATTCATTATTTCGATCTTATTGGCCTTTGTTCTTTATTTTTTATTTTTCATATAGATTTCCGTCTTTATATTATACCCTTTCGTAGATGAATTATGCTTATTTTCACATCTAGGATTTACATATACAACATATATTACTGTCAAGAGGGAATTTTTCTCAGTATTTAGATATTTAGATTCAAAATAAGAAAGGGATCAATTCAATTATAAACCCGCAAAACAAAGATTAGGATTGGGTTGGGTTGCGCTATATATATCAAAGAGTATACAATAATGATGTATTTGGTGAATCAAATACATGGTCTAATAACAAACCATTTTAACATAACATTTTAATGAGTTGATAATATTAATTGAATATTGAATATTTTTTGAAAGATTTTTGTTAAAGGTTTCATTCATGCCTAATCCATATCGAGTAGACCTTGTTGTTGTGAG

			>Tillandsia_purpurea
			??????????????????????????????????????????????????????????????????????????????????????????????????????????????????????????????????????????????????????????????????????????????????????????????????????????????????????????????????????????????????????????????????????????????????????????????????????????????????????????????????????????????????????????????????????????????????????????????????????????????????????????????????????????????????????????????????????????????????????????????????????????????????????????????????????????????????????????????????????????????????????????????????????????????????????????????????????????????????????????????????????????????????????????????????????????????????????????????????????????????????????????????????????????????????????????????

			>Tillandsia_quaquaflorifera
			??????????????????????????????????????????????????????????????????????????????????????????????????????????????????????????????????????????????????????????????????????????????????????????????????????????????????????????????????????????????????????????????????????????????????????????????????????????????????????????????????????????????????????????????????????????????????????????????????????????????????????????????????????????????????????????????????????????????????????????????????????????????????????????????????????????????????????????????????????????????????????????????????????????????????????????????????????????????????????????????????????????????????????????????????????????????????????????????????????????????????????????????????????????????????????????????

			>Tillandsia_rauhii
			ATAATTATAATAAAGTGAAATATGTCGAAATCCTTTTTGGAATAATACCAAATCGAAAATAAATGTCCGATAGCAAGTTGATCAGTTAATTCAATAAGAGATAAATGGGAGATAGCACTCGATTTAGTTGGTACCACCCAACCGAATCCGATTCAATCGTTTACTCATTCAATGAGTAAATTTTCAAGTTCAGCCAATCTTTATTTTTTT----------TTGCAAGTAAATGAAATCGTGAATAAATGTGTTTCATTTCTCTATCATTATAGAAAAAAAGATCCATATTATATTACTTATGGAATTCAAACCCGAACTCAATTTATGATTCATTATTTCGATCTTATTGGCCTTTGTTCTTTATTTTTTATTTTTCATATAGATTTCCGTCTTTATATTATACCCTTTCGTAGATGAATTATGCTTATTTTCACATCTAGGATTTACATATACAACATATATTACTGTCAAGAGGGAATTTTTCTCAGTATTTAGATATTTAGATTCAAAATAAGAAAGGGATCAATTCAATTATAAACCCGCAAAACAAAGATTAGGATTGGGTTGGGTTGCGCTATATATATCAAAGAGTATACAATAATGATGTATTTGGTGAATCAAATACATGGTCTAATAACAAACCATTTTAACATAACATTTTAATGAGTTGATAATATTAATTGAATATTGAATATTTTTTGAAAGATTTTTGTTAAAGGTTTCATTCATGCCTAATCCATATCGAGTAGACCTTGTTGTTGTGAG

			>Tillandsia_rectangula
			??????????????????????????????????????????????????????????????????????????????????????????????????????????????????????????????????????????????????????????????????????????????????????????????????????????????????????????????????????????????????????????????????????????????????????????????????????????????????????????????????????????????????????????????????????????????????????????????????????????????????????????????????????????????????????????????????????????????????????????????????????????????????????????????????????????????????????????????????????????????????????????????????????????????????????????????????????????????????????????????????????????????????????????????????????????????????????????????????????????????????????????????????????????????????????????????

			>Tillandsia_recurvata
			??????????????????????????????????????????????????????????????????????????????????????????????????????????????????????????????????????????????????????????????????????????????????????????????????????????????????????????????????????????????????????????????????????????????????????????????????????????????????????????????????????????????????????????????????????????????????????????????????????????????????????????????????????????????????????????????????????????????????????????????????????????????????????????????????????????????????????????????????????????????????????????????????????????????????????????????????????????????????????????????????????????????????????????????????????????????????????????????????????????????????????????????????????????????????????????????

			>Tillandsia_recurvifolia
			??????????????????????????????????????????????????????????????????????????????????????????????????????????????????????????????????????????????????????????????????????????????????????????????????????????????????????????????????????????????????????????????????????????????????????????????????????????????????????????????????????????????????????????????????????????????????????????????????????????????????????????????????????????????????????????????????????????????????????????????????????????????????????????????????????????????????????????????????????????????????????????????????????????????????????????????????????????????????????????????????????????????????????????????????????????????????????????????????????????????????????????????????????????????????????????????

			>Tillandsia_recurvifolia_var._subsecundifolia
			??????????????????????????????????????????????????????????????????????????????????????????????????????????????????????????????????????????????????????????????????????????????????????????????????????????????????????????????????????????????????????????????????????????????????????????????????????????????????????????????????????????????????????????????????????????????????????????????????????????????????????????????????????????????????????????????????????????????????????????????????????????????????????????????????????????????????????????????????????????????????????????????????????????????????????????????????????????????????????????????????????????????????????????????????????????????????????????????????????????????????????????????????????????????????????????????

			>Tillandsia_remota
			ATAATTATAATAAAGTGAAATATGTCGAAATCCTTTTTGGAATAATACCAAATCGAAAATAAATGTCCGATAGCAAGTTGATCAGTTAATTCAATAAGAGATAAATGGGAGATAGCACTCGATTTAGTTGGTACCGCCCAACCGAATCCGATTCAATCGTTTACTCATTCAATGAGTAAATTTTCAAGTTCAGCCAATCTTTTTTTCA--AAAAAAAAAATTGCAAGTAAATGAAATCGTGAATAAATGTGTTTCATTTCTCTATCATTATAGAAAAAAAGATCCATATTATATTACTTATGGAATTCGAACCCGAACTCGATTTATGATTCATTATTTCGATCTTATTGGCCTTTGTTCTTTATTTTTTATTTTTCATATAGATTTCCGTCTTTATATTATACCCTTTCGTAGATGAATTATGCTTATTTTCACATCTAGGATTTACATATACAACATATATTACTGTCAAGAGGGAATTTTTCTCAGTATTTAGATATTTAGATTCAAAATAAGAAAGGGATCAATTCAATTATAAACCCGCAAAACAAAGATTAGGATTGGGTTGGGTTGCGCTATATATATCAAAGAGTATACAATAATGATGTATTTGGTGAATCAAATACATGGTCTAATAACAAACCATTTTAACATAACATTTTAATGAGTTGATAATATTAATTGAATATTGAATATTTTTTGAAAGATTTTTGTTAAAGGTTTCATTCATGCCTAATCCATATCGAGTAGACCTTGTTGTTGTGAG

			>Tillandsia_rosarioae
			??????????????????????????????????????????????????????????????????????????????????????????????????????????????????????????????????????????????????????????????????????????????????????????????????????????????????????????????????????????????????????????????????????????????????????????????????????????????????????????????????????????????????????????????????????????????????????????????????????????????????????????????????????????????????????????????????????????????????????????????????????????????????????????????????????????????????????????????????????????????????????????????????????????????????????????????????????????????????????????????????????????????????????????????????????????????????????????????????????????????????????????????????????????????????????????????

			>Tillandsia_roseoscapa
			??????????????????????????????????????????????????????????????????????????????????????????????????????????????????????????????????????????????????????????????????????????????????????????????????????????????????????????????????????????????????????????????????????????????????????????????????????????????????????????????????????????????????????????????????????????????????????????????????????????????????????????????????????????????????????????????????????????????????????????????????????????????????????????????????????????????????????????????????????????????????????????????????????????????????????????????????????????????????????????????????????????????????????????????????????????????????????????????????????????????????????????????????????????????????????????????

			>Tillandsia_rupicola
			??????????????????????????????????????????????????????????????????????????????????????????????????????????????????????????????????????????????????????????????????????????????????????????????????????????????????????????????????????????????????????????????????????????????????????????????????????????????????????????????????????????????????????????????????????????????????????????????????????????????????????????????????????????????????????????????????????????????????????????????????????????????????????????????????????????????????????????????????????????????????????????????????????????????????????????????????????????????????????????????????????????????????????????????????????????????????????????????????????????????????????????????????????????????????????????????

			>Tillandsia_sceptriformis
			??????????????????????????????????????????????????????????????????????????????????????????????????????????????????????????????????????????????????????????????????????????????????????????????????????????????????????????????????????????????????????????????????????????????????????????????????????????????????????????????????????????????????????????????????????????????????????????????????????????????????????????????????????????????????????????????????????????????????????????????????????????????????????????????????????????????????????????????????????????????????????????????????????????????????????????????????????????????????????????????????????????????????????????????????????????????????????????????????????????????????????????????????????????????????????????????

			>Tillandsia_schiedeana
			??????????????????????????????????????????????????????????????????????????????????????????????????????????????????????????????????????????????????????????????????????????????????????????????????????????????????????????????????????????????????????????????????????????????????????????????????????????????????????????????????????????????????????????????????????????????????????????????????????????????????????????????????????????????????????????????????????????????????????????????????????????????????????????????????????????????????????????????????????????????????????????????????????????????????????????????????????????????????????????????????????????????????????????????????????????????????????????????????????????????????????????????????????????????????????????????

			>Tillandsia_schimperiana
			??????????????????????????????????????????????????????????????????????????????????????????????????????????????????????????????????????????????????????????????????????????????????????????????????????????????????????????????????????????????????????????????????????????????????????????????????????????????????????????????????????????????????????????????????????????????????????????????????????????????????????????????????????????????????????????????????????????????????????????????????????????????????????????????????????????????????????????????????????????????????????????????????????????????????????????????????????????????????????????????????????????????????????????????????????????????????????????????????????????????????????????????????????????????????????????????

			>Tillandsia_secunda
			??????????????????????????????????????????????????????????????????????????????????????????????????????????????????????????????????????????????????????????????????????????????????????????????????????????????????????????????????????????????????????????????????????????????????????????????????????????????????????????????????????????????????????????????????????????????????????????????????????????????????????????????????????????????????????????????????????????????????????????????????????????????????????????????????????????????????????????????????????????????????????????????????????????????????????????????????????????????????????????????????????????????????????????????????????????????????????????????????????????????????????????????????????????????????????????????

			>Tillandsia_seleriana
			??????????????????????????????????????????????????????????????????????????????????????????????????????????????????????????????????????????????????????????????????????????????????????????????????????????????????????????????????????????????????????????????????????????????????????????????????????????????????????????????????????????????????????????????????????????????????????????????????????????????????????????????????????????????????????????????????????????????????????????????????????????????????????????????????????????????????????????????????????????????????????????????????????????????????????????????????????????????????????????????????????????????????????????????????????????????????????????????????????????????????????????????????????????????????????????????

			>Tillandsia_sessemocinoi
			??????????????????????????????????????????????????????????????????????????????????????????????????????????????????????????????????????????????????????????????????????????????????????????????????????????????????????????????????????????????????????????????????????????????????????????????????????????????????????????????????????????????????????????????????????????????????????????????????????????????????????????????????????????????????????????????????????????????????????????????????????????????????????????????????????????????????????????????????????????????????????????????????????????????????????????????????????????????????????????????????????????????????????????????????????????????????????????????????????????????????????????????????????????????????????????????

			>Tillandsia_setacea
			??????????????????????????????????????????????????????????????????????????????????????????????????????????????????????????????????????????????????????????????????????????????????????????????????????????????????????????????????????????????????????????????????????????????????????????????????????????????????????????????????????????????????????????????????????????????????????????????????????????????????????????????????????????????????????????????????????????????????????????????????????????????????????????????????????????????????????????????????????????????????????????????????????????????????????????????????????????????????????????????????????????????????????????????????????????????????????????????????????????????????????????????????????????????????????????????

			>Tillandsia_sierrajuarezensis
			??????????????????????????????????????????????????????????????????????????????????????????????????????????????????????????????????????????????????????????????????????????????????????????????????????????????????????????????????????????????????????????????????????????????????????????????????????????????????????????????????????????????????????????????????????????????????????????????????????????????????????????????????????????????????????????????????????????????????????????????????????????????????????????????????????????????????????????????????????????????????????????????????????????????????????????????????????????????????????????????????????????????????????????????????????????????????????????????????????????????????????????????????????????????????????????????

			>Tillandsia_socialis
			??????????????????????????????????????????????????????????????????????????????????????????????????????????????????????????????????????????????????????????????????????????????????????????????????????????????????????????????????????????????????????????????????????????????????????????????????????????????????????????????????????????????????????????????????????????????????????????????????????????????????????????????????????????????????????????????????????????????????????????????????????????????????????????????????????????????????????????????????????????????????????????????????????????????????????????????????????????????????????????????????????????????????????????????????????????????????????????????????????????????????????????????????????????????????????????????

			>Tillandsia_somnians
			??????????????????????????????????????????????????????????????????????????????????CAGTTAATTCAATAAGAGATAAATGGGAGATAGCACTCGATTTAGTTGGTACCACCCAACCGAATCCGATTCAATCGTTTACTCATTCAATGAGTAAATTTTAAAGTTCAGCCAATCCTTTATTTTTT-----------------------------------------------------------------------------------------------------------------------------------------------------------------TTTTTCATATAGATTTCCGTCTTTATATTATACCCTTTCGTAGATGAATTATGCTTATTTTCACATCTAGGATTTACATATACAACATATATTACTGTCAAGAGGGAATTTTTCTCAGTATTTAGATATTTAGATTCAAAATAAGAAAGGGATCAATTCAATTATAAACCCGCAAAACAAAGATTAGGATTGGGTTGGGTTGCGCTATATATATCAAAGAGTATACAATAATGATGTATTTGGTGAATCAAATACATGGTCTAATAACAAACCATTTTAACATAACATTTTAATGAGTTGATAATATTAATTGAATATTGAATATTTTTTGAAAGATTTTTGTTAAAGGTTTCATTCATGCCTAATCCATATCGAGTAGACCTTGTTGTTGTGAG

			>Tillandsia_spathacea
			??????????????????????????????????????????????????????????????????????????????????????????????????????????????????????????????????????????????????????????????????????????????????????????????????????????????????????????????????????????????????????????????????????????????????????????????????????????????????????????????????????????????????????????????????????????????????????????????????????????????????????????????????????????????????????????????????????????????????????????????????????????????????????????????????????????????????????????????????????????????????????????????????????????????????????????????????????????????????????????????????????????????????????????????????????????????????????????????????????????????????????????????????????????????????????????????

			>Tillandsia_sphaerocephala
			??????????????????????????????????????????????????????????????????????????????????????????????????????????????????????????????????????????????????????????????????????????????????????????????????????????????????????????????????????????????????????????????????????????????????????????????????????????????????????????????????????????????????????????????????????????????????????????????????????????????????????????????????????????????????????????????????????????????????????????????????????????????????????????????????????????????????????????????????????????????????????????????????????????????????????????????????????????????????????????????????????????????????????????????????????????????????????????????????????????????????????????????????????????????????????????????

			>Tillandsia_spiraliflora
			??????????????????????????????????????????????????????????????????????????????????????????????????????????????????????????????????????????????????????????????????????????????????????????????????????????????????????????????????????????????????????????????????????????????????????????????????????????????????????????????????????????????????????????????????????????????????????????????????????????????????????????????????????????????????????????????????????????????????????????????????????????????????????????????????????????????????????????????????????????????????????????????????????????????????????????????????????????????????????????????????????????????????????????????????????????????????????????????????????????????????????????????????????????????????????????????

			>Tillandsia_sprengeliana
			??????????????????????????????????????????????????????????????????????????????????????????????????????????????????????????????????????????????????????????????????????????????????????????????????????????????????????????????????????????????????????????????????????????????????????????????????????????????????????????????????????????????????????????????????????????????????????????????????????????????????????????????????????????????????????????????????????????????????????????????????????????????????????????????????????????????????????????????????????????????????????????????????????????????????????????????????????????????????????????????????????????????????????????????????????????????????????????????????????????????????????????????????????????????????????????????

			>Tillandsia_stenoura
			??????????????????????????????????????????????????????????????????????????????????????????????????????????????????????????????????????????????????????????????????????????????????????????????????????????????????????????????????????????????????????????????????????????????????????????????????????????????????????????????????????????????????????????????????????????????????????????????????????????????????????????????????????????????????????????????????????????????????????????????????????????????????????????????????????????????????????????????????????????????????????????????????????????????????????????????????????????????????????????????????????????????????????????????????????????????????????????????????????????????????????????????????????????????????????????????

			>Tillandsia_streptocarpa
			??????????????????????????????????????????????????????????????????????????????????????????????????????????????????????????????????????????????????????????????????????????????????????????????????????????????????????????????????????????????????????????????????????????????????????????????????????????????????????????????????????????????????????????????????????????????????????????????????????????????????????????????????????????????????????????????????????????????????????????????????????????????????????????????????????????????????????????????????????????????????????????????????????????????????????????????????????????????????????????????????????????????????????????????????????????????????????????????????????????????????????????????????????????????????????????????

			>Tillandsia_streptophylla
			??????????????????????????????????????????????????????????????????????????????????????????????????????????????????????????????????????????????????????????????????????????????????????????????????????????????????????????????????????????????????????????????????????????????????????????????????????????????????????????????????????????????????????????????????????????????????????????????????????????????????????????????????????????????????????????????????????????????????????????????????????????????????????????????????????????????????????????????????????????????????????????????????????????????????????????????????????????????????????????????????????????????????????????????????????????????????????????????????????????????????????????????????????????????????????????????

			>Tillandsia_stricta
			ATAATTATAATAAAGTGAAATATGTCGAAATCCTTTTTGGAATAATACCAAATCGAAAATAAATGTCCGATAGCAAGTTGATCAGTTAATTCAATAAGAGATAAATGGGAGATAGCACTCGATTTAGTTGGTACCACCCAACCGAATCCGATTCAATCGTTTACTCATTCAATGAGTAAATTTTCAAGTTCAGCCAATCTTTATTTTTTT----------TTGCAAGTAAATGAAATCGTGAATAAATGTGTTTCATTTCTCTATCATTATAGAAAAAAGGATCCATATTATATTACTTATGGAATTCGAACCCGAACTCGATTTATGATTCATTATTTCGATCTTATTGGCCTTTGTTCTTTATTTTTTATTTTTCATATGGATTTCCGTCTTTATATTATACCCTTTCGTAGATGAATTATGCTTACTTTCACATCTAGGATTTACATATACAACATATATTACTGTCAAGAGGGAATTTTTCTCAGTATTTAGATATTTAGATTCAAAATAAGAAAGGGATCAATTCAATTATAAACCCGCAAAACAAAGATTAGGATTGGGTTGGGTTGCGCTATATATATCAAAGAGTATACAATAATGATGTATTTGGTGAATCAAATACATGGTCTAATAACAAACCATTTTAACATAACATTTTAATGAGTTGATAATATTAATTGACTATTGAATATTTTTTGAAAGATTTTTGTTAAAGGTTTCATTCATGCCTAATCCATATCGAGTAGACCTTGTTGTTGTGAG

			>Tillandsia_stricta_var._albifolia
			??????????????????????????????????????????????????????????????????????????????????????????????????????????????????????????????????????????????????????????????????????????????????????????????????????????????????????????????????????????????????????????????????????????????????????????????????????????????????????????????????????????????????????????????????????????????????????????????????????????????????????????????????????????????????????????????????????????????????????????????????????????????????????????????????????????????????????????????????????????????????????????????????????????????????????????????????????????????????????????????????????????????????????????????????????????????????????????????????????????????????????????????????????????????????????????????

			>Tillandsia_subulifera
			??????????????????????????????????????????????????????????????????????????????????????????????????????????????????????????????????????????????????????????????????????????????????????????????????????????????????????????????????????????????????????????????????????????????????????????????????????????????????????????????????????????????????????????????????????????????????????????????????????????????????????????????????????????????????????????????????????????????????????????????????????????????????????????????????????????????????????????????????????????????????????????????????????????????????????????????????????????????????????????????????????????????????????????????????????????????????????????????????????????????????????????????????????????????????????????????

			>Tillandsia_suesilliae
			??????????????????????????????????????????????????????????????????????????????????????????????????????????????????????????????????????????????????????????????????????????????????????????????????????????????????????????????????????????????????????????????????????????????????????????????????????????????????????????????????????????????????????????????????????????????????????????????????????????????????????????????????????????????????????????????????????????????????????????????????????????????????????????????????????????????????????????????????????????????????????????????????????????????????????????????????????????????????????????????????????????????????????????????????????????????????????????????????????????????????????????????????????????????????????????????

			>Tillandsia_supermexicana
			??????????????????????????????????????????????????????????????????????????????????????????????????????????????????????????????????????????????????????????????????????????????????????????????????????????????????????????????????????????????????????????????????????????????????????????????????????????????????????????????????????????????????????????????????????????????????????????????????????????????????????????????????????????????????????????????????????????????????????????????????????????????????????????????????????????????????????????????????????????????????????????????????????????????????????????????????????????????????????????????????????????????????????????????????????????????????????????????????????????????????????????????????????????????????????????????

			>Tillandsia_tecolometl
			??????????????????????????????????????????????????????????????????????????????????????????????????????????????????????????????????????????????????????????????????????????????????????????????????????????????????????????????????????????????????????????????????????????????????????????????????????????????????????????????????????????????????????????????????????????????????????????????????????????????????????????????????????????????????????????????????????????????????????????????????????????????????????????????????????????????????????????????????????????????????????????????????????????????????????????????????????????????????????????????????????????????????????????????????????????????????????????????????????????????????????????????????????????????????????????????

			>Tillandsia_tectorum
			ATAATTATAATTAAGTGAAATATGTCGAAATCCTTTTTGGAATAATACCAAATTGAAAATAAATGTCCGATAGCAAGTTGATCAGTTAATTCAATAAGAGATAAATGGGAGATAGCACTCGATTTAGTTGGTACCACCCAACCGAATCCGATTCAATCGTTTACTCATTCAATGAGTAAATTTTCAAGTTCAGCCAATCTTATTTTTTTT--------AATTGCAAGTAAATGAAATCGTGAATAAATGTGTTTCATTTCTCTATCATTATAGAAAAAAAGATCCATATTATATTACTTATGGAATTCGAACCCGAACTCGATTTATGATTCATTATTTCGATCTTATTGGCCTTTGTTCTTTATTTTTTATTTTTCATATAGATTTCCGTCTTTATATTATACCCTTTCGTAGATGAATTATGCTTATTTTCACATCTAGGATTTACATATACAACATATATTACTGTCAAGAGGGAATTTTTCTCAGTATTTAGATATTTAGATTCAAAATAAGAAAGGGATCAATTCAATTATAAACCCGCAAAACAAAGATTAGGATTGGGTTGGGTTGCGCTATATATATCAAAGAGTATACAATAATGATGTATTTGGTGAATCAAATACATGGTCTAATAACAAACCATTTTAACATAACATTTTAATGAGTTGATAATATTAATTGAATATTGAATATTTTTTGAAAGATTTTTGTTAAAGGTTTCATTCATGCCTAATCCATATCGAGTAGACCTTGTTGTTGTGAG

			>Tillandsia_tehuacana
			??????????????????????????????????????????????????????????????????????????????????????????????????????????????????????????????????????????????????????????????????????????????????????????????????????????????????????????????????????????????????????????????????????????????????????????????????????????????????????????????????????????????????????????????????????????????????????????????????????????????????????????????????????????????????????????????????????????????????????????????????????????????????????????????????????????????????????????????????????????????????????????????????????????????????????????????????????????????????????????????????????????????????????????????????????????????????????????????????????????????????????????????????????????????????????????????

			>Tillandsia_tenuifolia
			ATAATTATAATAAAGTGAAATATGTCGAAATCCTTTTTGGAATAATACCAAATCGAAAATAAATGTCCGATAGCAAGTTGATCAGTTAATTCAATAAGAGATAAATGGGAGATAGCACTCGATTTAGTTGGTACCACCCAACCGAATCCGATTCAATCGTTTACTCATTCAATGAGTAAATTTTCAAGTTCAGCCAATCTTTTTTTTTTT------TTAATTTCAAGTAAATGAAATCGTGAATAAATGTGTTTCATTTCTCTATCATTATAGAAAAAAAGATTCATATTATATTACTTATGGAATTCGAACCCGAACTCGATTTATGATTCATTATTTCGATCTTATTGGCCTTTGTTCTTTATTTTTTATTTTTCATATAGATTTCCGTCTTTATATTATACCCTTTCGTAGATGAATTATGCTTACTTTCACATCTAGGATTTACATATACAACATATATTACTGTCAAGAGGGAATTTTTCTCAGTATTTAGATATTTAGATTCAAAATAAGAAAGGGATCAATTCAATTATAAACCCGCAAAACAAAGATTAGGATTGGGTTGGGTTGCGCTATATATATCAAAGAGTATACAATAATGATGTATTTGGTGAATCAAATACATGGTCTAATAACAAACCATTTTAACATAACATTTTAATGAGTTGATAATATTAATTGACTATTGAATATTTTTTGAAAGATTTTTGTTAAAGGTTTCATTCATGCCTAATCCATATCGAGTAGACCTTGTTGTTGTGAG

			>Tillandsia_tenuifolia_var._albifolia
			??????????????????????????????????????????????????????????????????????????????????????????????????????????????????????????????????????????????????????????????????????????????????????????????????????????????????????????????????????????????????????????????????????????????????????????????????????????????????????????????????????????????????????????????????????????????????????????????????????????????????????????????????????????????????????????????????????????????????????????????????????????????????????????????????????????????????????????????????????????????????????????????????????????????????????????????????????????????????????????????????????????????????????????????????????????????????????????????????????????????????????????????????????????????????????????????

			>Tillandsia_tequendamae
			??????????????????????????????????????????????????????????????????????????????????????????????????????????????????????????????????????????????????????????????????????????????????????????????????????????????????????????????????????????????????????????????????????????????????????????????????????????????????????????????????????????????????????????????????????????????????????????????????????????????????????????????????????????????????????????????????????????????????????????????????????????????????????????????????????????????????????????????????????????????????????????????????????????????????????????????????????????????????????????????????????????????????????????????????????????????????????????????????????????????????????????????????????????????????????????????

			>Tillandsia_teres
			??????????????????????????????????????????????????????????????????????????????????????????????????????????????????????????????????????????????????????????????????????????????????????????????????????????????????????????????????????????????????????????????????????????????????????????????????????????????????????????????????????????????????????????????????????????????????????????????????????????????????????????????????????????????????????????????????????????????????????????????????????????????????????????????????????????????????????????????????????????????????????????????????????????????????????????????????????????????????????????????????????????????????????????????????????????????????????????????????????????????????????????????????????????????????????????????

			>Tillandsia_toropiensis
			??????????????????????????????????????????????????????????????????????????????????????????????????????????????????????????????????????????????????????????????????????????????????????????????????????????????????????????????????????????????????????????????????????????????????????????????????????????????????????????????????????????????????????????????????????????????????????????????????????????????????????????????????????????????????????????????????????????????????????????????????????????????????????????????????????????????????????????????????????????????????????????????????????????????????????????????????????????????????????????????????????????????????????????????????????????????????????????????????????????????????????????????????????????????????????????????

			>Tillandsia_tortilis
			ATAATTATAATAAAGTGAAATATGTCGAAATCCTTTTTGGAATAATACCAAATTGAAAATAAATGTCCGATAGCAAGTTGATCAGTTAATTCAATAAGAGATAAATGGGAGATAGCACTCGATTTAGTTGGTACCACCCAACCGAATCCGATTCAATCGTTTACTCATTCAATGAGTAAATTTTCAAGTTCAGCCAATCTTTTTTTTT--------AAAATTGCAAGTAAATGAAATCGTGAATAAATGTGTTTCATTTCTCTATCATTATAGAAAAAAAGATCCATATTATATTACTTATGGAATTCGAACCCGAACTCGATTTATGATTCATTATTTCGATCTTATTGGCCTTTGTTCTTTATTTTTTATTTTTCATATAGATTTCCGTCTTTATATTATACCCTTTCGTAGATGAATTATGCTTATTTTCACATCTAGGATTTACATATACAACATATATTACTGTCAAGAGGGAATTTTTCTCAGTATTTAGATATTTAGATTCAAAATAAGAAAGGGATCAATTCAATTATAAACCCGCAAAACAAAGATTAGGATTGGGTTGGGTTGCGCTATATATATCAAAGAGTATACAATAATGATGTATTTGGTGAATCAAATACATGGTCTAATAACAAACCATTTTAACATAACATTTTAATGAGTTGATAATATTAATTGAATATTGAATATTTTTTGAAAGATTTTTGTTAAAGGTTTCATTCATGCCTAATCCATATCGAGTAGACCTTGTTGTTGTGAG

			>Tillandsia_tricholepis
			??????????????????????????????????????????????????????????????????????????????????????????????????????????????????????????????????????????????????????????????????????????????????????????????????????????????????????????????????????????????????????????????????????????????????????????????????????????????????????????????????????????????????????????????????????????????????????????????????????????????????????????????????????????????????????????????????????????????????????????????????????????????????????????????????????????????????????????????????????????????????????????????????????????????????????????????????????????????????????????????????????????????????????????????????????????????????????????????????????????????????????????????????????????????????????????????

			>Tillandsia_tricolor
			??????????????????????????????????????????????????????????????????????????????????????????????????????????????????????????????????????????????????????????????????????????????????????????????????????????????????????????????????????????????????????????????????????????????????????????????????????????????????????????????????????????????????????????????????????????????????????????????????????????????????????????????????????????????????????????????????????????????????????????????????????????????????????????????????????????????????????????????????????????????????????????????????????????????????????????????????????????????????????????????????????????????????????????????????????????????????????????????????????????????????????????????????????????????????????????????

			>Tillandsia_turneri
			??????????????????????????????????????????????????????????????????????????????????????????????????????????????????????????????????????????????????????????????????????????????????????????????????????????????????????????????????????????????????????????????????????????????????????????????????????????????????????????????????????????????????????????????????????????????????????????????????????????????????????????????????????????????????????????????????????????????????????????????????????????????????????????????????????????????????????????????????????????????????????????????????????????????????????????????????????????????????????????????????????????????????????????????????????????????????????????????????????????????????????????????????????????????????????????????

			>Tillandsia_turneri_var._orientalis
			??????????????????????????????????????????????????????????????????????????????????????????????????????????????????????????????????????????????????????????????????????????????????????????????????????????????????????????????????????????????????????????????????????????????????????????????????????????????????????????????????????????????????????????????????????????????????????????????????????????????????????????????????????????????????????????????????????????????????????????????????????????????????????????????????????????????????????????????????????????????????????????????????????????????????????????????????????????????????????????????????????????????????????????????????????????????????????????????????????????????????????????????????????????????????????????????

			>Tillandsia_usneoides
			ATAATTATAATAAAGTGAAATATGTCGAAATCCTTTTTGGAATAATACCAAATCGAAAATAAATGTCCGATAGCAAGTTGATCAGTTAATTCAATAAGAGATAAATGGGAGATAGCACTCGATTTAGTTGGTACCACCCAACCGAATCCGATTCAATCGTTTACTCATTCAATGAGTAAATTTTCAAGTTCAGCCAATCTTTATTTTT-------------------------------------------------------------------------------------------------------------------------------------------------------------------TTTTTCATATAGATTTCCGTCTTTATATTATACCCTTTCGTAGATGAATTATGCTTATTTTCACATCTAGGATTTACATATACAACATATATTACTGTCAAGAGGGAATTTTTCTCAGTATTTAGATATTTAGATTCAAAATAAGAAAGGGATCAATTCAATTATAAACCCGCAAAACAAAGATTAGGATTGGGTTGGGTTGCGCTATATATATCAAAGAGTATACAATAATGATGTATTTGGTGAATCAAATACATGGTCTAATAACAAACCATTTTAACATAACATTTTAATGAGTTGATAATATTAATTGACTATTGAATATTTTTTGAAAGATTTTTGTTAAAGGTTTCATTCATGCCTAATCCATATCGAGTAGACCTTGTTGTTGTGAG

			>Tillandsia_utriculata
			ATAATTATAATAAAGTGAAATATGTCGAAATCCTTTTTGGAATAATACCAAATCGAAAATAAATGTCCGATAGCAAGTTGATCAGTTAATTCAATAAGAGATAAATGGGAGATAGCACTCGATTTAGTTGGTACCGCCCAACCGAATCCGATTCAATCGTTTACTCATTCAATGAGTAAATTTTCAAGTTCAGCCAATCTTTTTTTTTTTAAAAAAAAAATTGCAAGTAAATGAAATCGTGAATAAATGTGTTTCATTTCTCTATCATTATAGAAAAAAAGATACATATTATATTACTTATGGAATTCGAACCCGAACTCGATTTATGATTCATTATTTCGATCTTATTGGCCTTTGTTCTTTATTTTTTATTTTTCATATAGATTTCCGTCTTTATATTATACCCTTTCGTAGATGAATTATGCTTATTTTCACATCTAGGATTTACATATACAACATATATTACTGTCAAGAGGGAATTTTTCTCAGTATTTAGATATTTAGATTCAAAATAAGAAAGGGATCAATTCAATTATAAACCCGCAAAACAAAGATTAGGATTGGGTTGGGTTGCGCTATATATATCAAAGAGTATACAATAATGATGTATTTGGTGAATCAAATACATGGTCTAATAACAAACCATTTTAACATAACATTTTAATGAGTTGATAATATTAATTGAATATTGAATATTTTTTGAAAGATTTTTGTTAAAGGTTTCATTCATGCCTAATCCATATCGAGTAGACCTTGTTGTTGTGAG

			>Tillandsia_utriculata_subsp._pringlei
			??????????????????????????????????????????????????????????????????????????????????????????????????????????????????????????????????????????????????????????????????????????????????????????????????????????????????????????????????????????????????????????????????????????????????????????????????????????????????????????????????????????????????????????????????????????????????????????????????????????????????????????????????????????????????????????????????????????????????????????????????????????????????????????????????????????????????????????????????????????????????????????????????????????????????????????????????????????????????????????????????????????????????????????????????????????????????????????????????????????????????????????????????????????????????????????????

			>Tillandsia_variabilis
			??????????????????????????????????????????????????????????????????????????????????????????????????????????????????????????????????????????????????????????????????????????????????????????????????????????????????????????????????????????????????????????????????????????????????????????????????????????????????????????????????????????????????????????????????????????????????????????????????????????????????????????????????????????????????????????????????????????????????????????????????????????????????????????????????????????????????????????????????????????????????????????????????????????????????????????????????????????????????????????????????????????????????????????????????????????????????????????????????????????????????????????????????????????????????????????????

			>Tillandsia_venusta
			??????????????????????????????????????????????????????????????????????????????????????????????????????????????????????????????????????????????????????????????????????????????????????????????????????????????????????????????????????????????????????????????????????????????????????????????????????????????????????????????????????????????????????????????????????????????????????????????????????????????????????????????????????????????????????????????????????????????????????????????????????????????????????????????????????????????????????????????????????????????????????????????????????????????????????????????????????????????????????????????????????????????????????????????????????????????????????????????????????????????????????????????????????????????????????????????

			>Tillandsia_violacea
			??????????????????????????????????????????????????????????????????????????????????????????????????????????????????????????????????????????????????????????????????????????????????????????????????????????????????????????????????????????????????????????????????????????????????????????????????????????????????????????????????????????????????????????????????????????????????????????????????????????????????????????????????????????????????????????????????????????????????????????????????????????????????????????????????????????????????????????????????????????????????????????????????????????????????????????????????????????????????????????????????????????????????????????????????????????????????????????????????????????????????????????????????????????????????????????????

			>Tillandsia_werdermannii
			??????????????????????????????????????????????????????????????????????????????????????????????????????????????????????????????????????????????????????????????????????????????????????????????????????????????????????????????????????????????????????????????????????????????????????????????????????????????????????????????????????????????????????????????????????????????????????????????????????????????????????????????????????????????????????????????????????????????????????????????????????????????????????????????????????????????????????????????????????????????????????????????????????????????????????????????????????????????????????????????????????????????????????????????????????????????????????????????????????????????????????????????????????????????????????????????

			>Tillandsia_werneriana
			ATAATTATAATAAAGTGAAATATGTCGAAATCCTTTTTGGAATAATACCAAATCGAAAATAAATGTCCGATAGCAAGTTGATCAGTTAATTCAATAAGAGATAAATGGGAGATAGCACTCGATTTAGTTGGTACCACCCAACCGAATCCGATTCAATCGTTTACTCATTCAATGAGTCAATTTTCAAGTTCAGCCAATC-TTTTTTTTTC-AAAAAAAAATTGCAAGTAAATGAAATCGTGAATAAATGTGTTTCATTTCTCTATCATTATAGAAAAAAAGATCCATATTATATTACTTATGGAATTCGAACCCGAACTCGATTTATGATTCATTATTTCGATCTTATTGGCCTTTGTTCTTTATTTTTTATTTTTCATATAGATTTCCGTCTTTATATTATACCCTTTCGTAGATGAATTATGCTTATTTTCACATCTAGGATTTACATATACAACATATATTACTGTCAAGAGGGAATTTTTCTCAGTATTTAGATATTTAGATTCAAAATAAGAAAGGGATCAATTCAATTATAAACCCGCAAAACAAAGATTAGGGTTGGGTTGGGTTGCGCTATATATATCAAAGAGTATACAATAATGATGTATTTGGTGAATCAAATACATGGTCTAATAACAAACCATTTTAACATAACATTTTAATGAGTTGATAATATTAATTGAATATTGAATATTTTTTGAAAGATTTTTGTTAAAGGTTTCATTCATGCCTAATCCATATCGAGTAGACCTTGTTGTTGTGAG

			>Tillandsia_winkleri
			??????????????????????????????????????????????????????????????????????????????????????????????????????????????????????????????????????????????????????????????????????????????????????????????????????????????????????????????????????????????????????????????????????????????????????????????????????????????????????????????????????????????????????????????????????????????????????????????????????????????????????????????????????????????????????????????????????????????????????????????????????????????????????????????????????????????????????????????????????????????????????????????????????????????????????????????????????????????????????????????????????????????????????????????????????????????????????????????????????????????????????????????????????????????????????????????

			>Tillandsia_witeckii
			??????????????????????????????????????????????????????????????????????????????????????????????????????????????????????????????????????????????????????????????????????????????????????????????????????????????????????????????????????????????????????????????????????????????????????????????????????????????????????????????????????????????????????????????????????????????????????????????????????????????????????????????????????????????????????????????????????????????????????????????????????????????????????????????????????????????????????????????????????????????????????????????????????????????????????????????????????????????????????????????????????????????????????????????????????????????????????????????????????????????????????????????????????????????????????????????

			>Tillandsia_xerographica
			ATAATTATAATAAAGTGAAATATGTCGAAATCCTTTTTGGAATAATACCAAATCGAAAATAAATGTCCGATAGCAAGTTGATCAGTTAATTCAATAAGAGATAAATGGGAGATAGCACTCGATTTAGTTGGTACCGCCCAACCGAATCCGATTCAATCGTTTACTCATTCAATGAGTAAATTTTCAAGTTCAGCCAATCTTTTTTTC---AAAAAAAAAATTGCAAGTAAATGAAATCGTGAATAAATGTGTTTCATTTCTCTATCATTATAGAAAAAAAGATCCATATTATATTACTTATGGAATTCGAACCCGAACTCGATTTATGATTCATTATTTCGATCTTATTGGCCTTTGTTCTTTATTTTTTATTTTTCATATAGATTTCCGTCTTTATATTATACCCTTTCGTAGATGAATTATGCTTATTTTCACATCTAGGATTTACATATACAACATATATTACTGTCAAGAGGGAATTTTTCTCAGTATTTAGATATTTAGATTCAAAATAAGAAAGGGATCAATTCAATTATAAACCCGCAAAACAAAGATTAGGATTGGGTTGGGTTGCGCTATATATATCAAAGAGTATACAATAATGATGTATTTGGTGAATCAAATACATGGTCTAATAACAAACCATTTTAACATAACATTTTAATGAGTTGATAATATTAATTGAATATTGAATATTTTTTGAAAGATTTTTGTTAAAGGTTTCATTCATGCCTAATCCATATCGAGTAGACCTTGTTGTTGTGAG

			>Tillandsia_xiphioides
			ATAATTATAATAAAGTGAAATATGTCGAAATCCTTTTTGGAATAATACCAAATCGAAAATAAATGTCCGATAGCAAGTTGATCAGTTAATTCAATAAGAGATAAATGGGAGATAGCACTCGATTTAGTTGGTACCACCCAACCGAATCCGATTCAATCGTTTACTCATTCAATGAGTAAATTTTCAAGTTCAGCCAATCTTTATTTTTT------------------------------------------------------------------------------------------------------------------------------------------------------------------TTTTTCATATAGATTTCCGTCTTTATATTATACCCTTTCATAGATGAATTATGCTTATTTTCACATCTAGGATTTACATATACAATATATATTACTGTCAAGAGGGAATTTTTCTCAGTATTTAGATATTTAGATTCAAAATAAGAAAGGGATCAATTCAATTATAAACCCGCAAAACAAAGATTAGGATTGGGTTGGGTTGCGCTATATATATCAAAGAGTATACAATAATGATGTATTTGGTGAATCAAATACATGGTCTAATAACAAACCATTTTAACATAACATTTTAATGAGTTGATAATATTAATTGACTATTGAATATTTTTTGAAAGATTTTTGTTAAAGGTTTCATTCATGCCTAATCCATATCGAGTAGACCTTGTTGTTGTGAG

			>Typha_latifolia
			ATAATTATAATAAAGTAAAATATGTCGAAATTTTATTTGGAATAGTGCCACATCGAAAATAAATGTCCGATAGCAAGTTGATCAGTTAATTCAATAAGAGATAAATGGGAGATAGAACTCTATTTAGTTGGTACCACCCAACCGAATCCGATTCAATCATTTACTCATTCAATGAGTCAATTTTCAAGTTCATCCAATC--TTTTTTTTTGAAAAAAAAATT---AGTAGATGAAATCAGGAGTAAATGTGTTTTATTTCTCTATCATTATA-GAAAAAAGATCCAAATTATATTACCTATGGAATTCGAACCCAAACTCGATTTATGATTTATTATTTCGATCTCATTGTCCATTGTTCCTTATTTTTTCTTTTC-ATATGGATTTCTATCTTTACAT-----CCTTTCATGGATGAATTATGTCTATTTTCACATCTAGGATTTACATATACAACATATATTACTGTCAAGAGGGAATTTTCCTCAGTATTTAGATATTGAGATCCAAAATAAGAGAGGGATCAATTCAATTCTCAACTTGCAAAACAAGGATTAGGATTGGATTGGGTTGCGCTATATATATCAAAGAGTATACAATAATGATGTATTTGGTGAATCAAATACATGGTCTAATAACGAACCATTTTAACATAACATTTTAATGAGTTGATAATATTAACTGAATA-------TTTTTTGAAAGATTTTTGTCAAAGATTTCATTCATGCCTAATCCATATCGAGTAGACCTTGTCGTTGTTAG

			>Ursulaea_macvaughii
			???????????????????????????????????TTTGGAATAATACCAAATCGAAAATAAATGTCCGATAGCAAGTTGATCAGTTAATTCAATAAGAGATAAATGGGAGATAGCACTCGATTTAGTTGGTACCACCCAACCGAATCCGATTCACTCGTTTACTCATTGAATGAGTAAATTTTCAAGTTCAGCCAATC--TTTTTTTTTTAAAAAAAAATTGCAAGTAGATGAAATCGTGAATAAATGTGTTTCATTTCTATATCATTATATAAAAAAAGATCCATATTATATTACTTATGGAATTCGAACCCGAACTCGATTTATGATTTATTATTTGGATCTTATTGGCCTTTGTTCTTTATTTTTGATTTTTCATATAGATTTCCGTCTTTATATTATACCCTTTCGTGGATGAATTATGCCTATTTTCACATCTAGGATTTACATATACAACATATATTACTGTCAAGAGGGAATTTTTCTCAGTATTTAGATATTTAGATTCAAAATAAGAAAGGGATCAATTCAATTATAAACCTGCAAAACAAAGATTAGGATTGGGTTGGGTTGCGCTATATATATCAAAGAGTATACAATAATGATGTATTTGGTGAATCAAATACATGGTCTAATAACAAACCATTTTAACATAACATTTTAATGAGTTGATAATATTAATTGAATATTGAATATTTTTTGAAAGATTTTTGTTAAAGGTTTCATTCATGCCTAATCCATATCGAGTAGACCTTGTTGTTGTG??

			>Ursulaea_tuitensis
			ATAATTATAATAAAGTGAAATATGTCGAAATCCTTTTTGGAATAATACCAAATCGAAAATAAATGTCCGATAGCAAGTTGATCAGTTAATTCAATAAGAGATAAATGGGAGATAGCACTCGATTTAGTTGGTACCACCCAACCGAATCCGATTCACTCGTTTACTCATTCAATGAGTAAATTTTCAAGTTCAGCCAATC-TTTTTTTTTTCAAAAAAAAATTGCAAGTAGATGAAATCGTGAATAAATGTGTTTCATTTCTATATCATTATATAAAAAAAGATCCATATTATATTACTTATGGAATTCGAACCCGAACTCGATTTATGATTTATTATTTGGATCTTATTGGCCTTTGTTCTTTATTTTTGATTTTTCATATAGATTTCCGTCTTTATATTATACCCTTTCGTGGATGAATTATGCCTATTTTCACATCTAGGATTTACATATACAACATATATTACTGTCAAGAGGGAATTTTTCTCAGTATTTAGATATTTAGATTCAAAATAAGAAAGGGATCAATTCAATTATAAACCTGCAAAACAAAGATTAGGATTGGGTTGGGTTGCGCTATATATATCAAAGAGTATACAATAATGATGTATTTGGTGAATCAAATACATGGTCTAATAACAAACCATTTTAACATAACATTTTAATGAGTTGATAATATTAATTGAATATTGAATATTTTTTGAAAGATTTTTGTTAAAGGTTTCATTCATGCCTAATCCATATCGAGTAGACCTTGTTGTTGTG??

			>Vriesea_agostiniana
			??????????????????????????????????????????????????????????????????????????????????????????????????????????????????????????????????????????????????????????????????????????????????????????????????????????????????????????????????????????????????????????????????????????????????????????????????????????????????????????????????????????????????????????????????????????????????????????????????????????????????????????????????????????????????????????????????????????????????????????????????????????????????????????????????????????????????????????????????????????????????????????????????????????????????????????????????????????????????????????????????????????????????????????????????????????????????????????????????????????????????????????????????????????????????????????????

			>Vriesea_amethystina
			??????????????????????????????????????????????????????????????????????????????????????????????????????????????????????????????????????????????????????????????????????????????????????????????????????????????????????????????????????????????????????????????????????????????????????????????????????????????????????????????????????????????????????????????????????????????????????????????????????????????????????????????????????????????????????????????????????????????????????????????????????????????????????????????????????????????????????????????????????????????????????????????????????????????????????????????????????????????????????????????????????????????????????????????????????????????????????????????????????????????????????????????????????????????????????????????

			>Vriesea_andaraiensis
			??????????????????????????????????????????????????????????????????????????????????????????????????????????????????????????????????????????????????????????????????????????????????????????????????????????????????????????????????????????????????????????????????????????????????????????????????????????????????????????????????????????????????????????????????????????????????????????????????????????????????????????????????????????????????????????????????????????????????????????????????????????????????????????????????????????????????????????????????????????????????????????????????????????????????????????????????????????????????????????????????????????????????????????????????????????????????????????????????????????????????????????????????????????????????????????????

			>Vriesea_arachnoidea
			??????????????????????????????????????????????????????????????????????????????????????????????????????????????????????????????????????????????????????????????????????????????????????????????????????????????????????????????????????????????????????????????????????????????????????????????????????????????????????????????????????????????????????????????????????????????????????????????????????????????????????????????????????????????????????????????????????????????????????????????????????????????????????????????????????????????????????????????????????????????????????????????????????????????????????????????????????????????????????????????????????????????????????????????????????????????????????????????????????????????????????????????????????????????????????????????

			>Vriesea_atra
			??????????????????????????????????????????????????????????????????????????????????????????????????????????????????????????????????????????????????????????????????????????????????????????????????????????????????????????????????????????????????????????????????????????????????????????????????????????????????????????????????????????????????????????????????????????????????????????????????????????????????????????????????????????????????????????????????????????????????????????????????????????????????????????????????????????????????????????????????????????????????????????????????????????????????????????????????????????????????????????????????????????????????????????????????????????????????????????????????????????????????????????????????????????????????????????????

			>Vriesea_bahiana
			??????????????????????????????????????????????????????????????????????????????????????????????????????????????????????????????????????????????????????????????????????????????????????????????????????????????????????????????????????????????????????????????????????????????????????????????????????????????????????????????????????????????????????????????????????????????????????????????????????????????????????????????????????????????????????????????????????????????????????????????????????????????????????????????????????????????????????????????????????????????????????????????????????????????????????????????????????????????????????????????????????????????????????????????????????????????????????????????????????????????????????????????????????????????????????????????

			>Vriesea_billbergioides
			??????????????????????????????????????????????????????????????????????????????????????????????????????????????????????????????????????????????????????????????????????????????????????????????????????????????????????????????????????????????????????????????????????????????????????????????????????????????????????????????????????????????????????????????????????????????????????????????????????????????????????????????????????????????????????????????????????????????????????????????????????????????????????????????????????????????????????????????????????????????????????????????????????????????????????????????????????????????????????????????????????????????????????????????????????????????????????????????????????????????????????????????????????????????????????????????

			>Vriesea_bituminosa
			ATAATTATAATAAAGTGAAATATGTCGAAATCCTTTTTGGAATAATACCAAATCGAAAATAAATGTCCGATAGCAAGTTGATCAGTTAATTCAATAAGAGATAAATGGGAGATAGCACTCGATTTAGTTGGTACCACCCAACCGAATCCGATTCAATCGTTTACTCATTCAATGAGTAAATTTTCAAGTTCAGCCAATCTTTTTTTTTTCAAAAAAAAAATTGCAAGTAAATGAAATCGTGAATAAATGTGTTTCATTTCTCTATCATTATAGAAAAAAGGATCCATATTATATTACTTATGGAATTCGAGCCCGAACTCGATTTATGATTCATTATTTCGATCTTATTGGCCTTTGTTCTTTATTTTTTATTTTTCATATAGATTTCCGTCTTTATATTATACCCTTTCGTAGATGAATTATGCTTATTTTCACATCTAGGATTTACATATACAACATATATTACTGTCAAGAGGGAATTTTTCTCAGTATTTAGATATTTAGATTCAAAATAAGAAAGGGATCAATTCAATTATAAACCTGCAAAACAAAGATTAGGATTGGGTTGGGTTGCGCTATATATATCAAAGAGTATACAATAATGATGTATTTGGTGAATCAAATACATGGTCTAATAACAAACCATTTTAACATAACATTTTAATGAGTTGATAATATTAATTGAATATTGAATATTTTTTGAAAGATTTTTGTTAAAGGTTTCATTCATGCCTAATCCATATCGAGTAGACCTTGTTGTTGTGAG

			>Vriesea_bleheri
			??????????????????????????????????????????????????????????????????????????????????????????????????????????????????????????????????????????????????????????????????????????????????????????????????????????????????????????????????????????????????????????????????????????????????????????????????????????????????????????????????????????????????????????????????????????????????????????????????????????????????????????????????????????????????????????????????????????????????????????????????????????????????????????????????????????????????????????????????????????????????????????????????????????????????????????????????????????????????????????????????????????????????????????????????????????????????????????????????????????????????????????????????????????????????????????????

			>Vriesea_botafogensis
			??????????????????????????????????????????????????????????????????????????????????????????????????????????????????????????????????????????????????????????????????????????????????????????????????????????????????????????????????????????????????????????????????????????????????????????????????????????????????????????????????????????????????????????????????????????????????????????????????????????????????????????????????????????????????????????????????????????????????????????????????????????????????????????????????????????????????????????????????????????????????????????????????????????????????????????????????????????????????????????????????????????????????????????????????????????????????????????????????????????????????????????????????????????????????????????????

			>Vriesea_cacuminis
			??????????????????????????????????????????????????????????????????????????????????????????????????????????????????????????????????????????????????????????????????????????????????????????????????????????????????????????????????????????????????????????????????????????????????????????????????????????????????????????????????????????????????????????????????????????????????????????????????????????????????????????????????????????????????????????????????????????????????????????????????????????????????????????????????????????????????????????????????????????????????????????????????????????????????????????????????????????????????????????????????????????????????????????????????????????????????????????????????????????????????????????????????????????????????????????????

			>Vriesea_calimaniana
			??????????????????????????????????????????????????????????????????????????????????????????????????????????????????????????????????????????????????????????????????????????????????????????????????????????????????????????????????????????????????????????????????????????????????????????????????????????????????????????????????????????????????????????????????????????????????????????????????????????????????????????????????????????????????????????????????????????????????????????????????????????????????????????????????????????????????????????????????????????????????????????????????????????????????????????????????????????????????????????????????????????????????????????????????????????????????????????????????????????????????????????????????????????????????????????????

			>Vriesea_capixabae
			??????????????????????????????????????????????????????????????????????????????????????????????????????????????????????????????????????????????????????????????????????????????????????????????????????????????????????????????????????????????????????????????????????????????????????????????????????????????????????????????????????????????????????????????????????????????????????????????????????????????????????????????????????????????????????????????????????????????????????????????????????????????????????????????????????????????????????????????????????????????????????????????????????????????????????????????????????????????????????????????????????????????????????????????????????????????????????????????????????????????????????????????????????????????????????????????

			>Vriesea_carinata
			ATAATTATAATAAAGTGAAATATGTCGAAATCCTTTTTGGAATAATACCAAATCGAAAATAAATGTCCGATAGCAAGTTGATCAGTTAATTCAATAAGAGATAAATGGGAGATAGCACTCGATTTAGTTGGTACCACCCAACCGAATCCGATTCAATCGTTTACTCATTCAATGAGTAAATTTTCAAGTTCAGCCAATCTTTTTTTTTCA-AAAAAAAAATTGCAAGTAAATGAAATCGTGAATAAATGTGTTTCATTTCTCTATCATTATAGAAAAAAGGATCCATATTATATTACTTATGGAATTCGAGCCCGAACTCGATTTATGATTCATTATTTCGATCTTATTGGCCTTTGTTCTTTATTTTTGATTTTTCATATAGATTTCCGTCTTTATATTATACCCTTTCGTAGATGAATTATGCTTATTTTCACATCTAGGATTTACATATACAACATATATTACTGTCAAGAGGGAATTTTTCTCAGTATTTAGATATTTAGATTCAAAATAAGAAAGGGATCAATTCAATTATAAACCTGCAAAACAAAGATTAGGATTGGGTTGGGTTGCGCTATATATATCAAAGAGTATACAATAATGATGTATTTGGTGAATCAAATACATGGTCTAATAACAAACCATTTTAACATAACATTTTAATGAGTTGATAATATTAATTGAATATTGAATATTTTTTGAAAGATTTTTGTTAAAGGTTTCATTCATGCCTAATCCATATCGAGTAGACCTTGTTGTTGTGAG

			>Vriesea_carinata_var._flavominiata
			??????????????????????????????????????????????????????????????????????????????????????????????????????????????????????????????????????????????????????????????????????????????????????????????????????????????????????????????????????????????????????????????????????????????????????????????????????????????????????????????????????????????????????????????????????????????????????????????????????????????????????????????????????????????????????????????????????????????????????????????????????????????????????????????????????????????????????????????????????????????????????????????????????????????????????????????????????????????????????????????????????????????????????????????????????????????????????????????????????????????????????????????????????????????????????????????

			>Vriesea_carinata_var._mangaratibensis
			??????????????????????????????????????????????????????????????????????????????????????????????????????????????????????????????????????????????????????????????????????????????????????????????????????????????????????????????????????????????????????????????????????????????????????????????????????????????????????????????????????????????????????????????????????????????????????????????????????????????????????????????????????????????????????????????????????????????????????????????????????????????????????????????????????????????????????????????????????????????????????????????????????????????????????????????????????????????????????????????????????????????????????????????????????????????????????????????????????????????????????????????????????????????????????????????

			>Vriesea_colnagoi
			??????????????????????????????????????????????????????????????????????????????????????????????????????????????????????????????????????????????????????????????????????????????????????????????????????????????????????????????????????????????????????????????????????????????????????????????????????????????????????????????????????????????????????????????????????????????????????????????????????????????????????????????????????????????????????????????????????????????????????????????????????????????????????????????????????????????????????????????????????????????????????????????????????????????????????????????????????????????????????????????????????????????????????????????????????????????????????????????????????????????????????????????????????????????????????????????

			>Vriesea_correia-araujoi
			ATAATTATAATAAAGTGAAATATGTCGAAATCCTTTTTGGAATAATACCAAATCGAAAATAAATGTCCGATAGCAAGTTGATCAGTTAATTCAATAAGAGATAAATGGGAGATAGCACTCGATTTAGTTGGTACCACCCAACCGAATCCGATTCAATCGTTTACTCATTCAATGAGTAAATTTTCAAGTTCAGCCAATCTTTTTTTTTCAAAAAAAAAAATTGCAAGTAAATGAAATCGTGAATAAATGTGTTTCATTTCTCTATCATTATAGAAAAAAGGATCCATATTATATTACTTATGGAATTCGAGCCCGAACTCGATTTATGATTCATTATTTCGATCTTATTGGCCTTTGTTCTTTATTTTTTATTTTTCATATAGATTTCCGTCTTTATATTATACCCTTTCGTAGATGAATTATGCTTATTTTCACATCTAGGATTTACATATACAACATATATTACTGTCAAGAGGGAATTTTTCTCAGTATTTAGATATTTAGATTCAAAATAAGAAAGGGATCAATTCAATTATAAACCTGCAAAACAAAGATTAGGATTGGGTTGGGTTGCGCTATATATATCAAAGAGTATACAATAATGATGTATTTGGTGAATCAAATACATGGTCTAATAACAAACCATTTTAACATAACATTTTAATGAGTTGATAATATTAATTGAATATTGAATATTTTTTGAAAGATTTTTGTTAAAGGTTTCATTCATGCCTAATCCATATCGAGTAGACCTTGTTGTTGTGAG

			>Vriesea_crassa
			??????????????????????????????????????????????????????????????????????????????????????????????????????????????????????????????????????????????????????????????????????????????????????????????????????????????????????????????????????????????????????????????????????????????????????????????????????????????????????????????????????????????????????????????????????????????????????????????????????????????????????????????????????????????????????????????????????????????????????????????????????????????????????????????????????????????????????????????????????????????????????????????????????????????????????????????????????????????????????????????????????????????????????????????????????????????????????????????????????????????????????????????????????????????????????????????

			>Vriesea_dubia
			??????????????????????????????????????????????????????????????????????????????????????????????????????????????????????????????????????????????????????????????????????????????????????????????????????????????????????????????????????????????????????????????????????????????????????????????????????????????????????????????????????????????????????????????????????????????????????????????????????????????????????????????????????????????????????????????????????????????????????????????????????????????????????????????????????????????????????????????????????????????????????????????????????????????????????????????????????????????????????????????????????????????????????????????????????????????????????????????????????????????????????????????????????????????????????????????

			>Vriesea_duvaliana
			??????????????????????????????????????????????????????????????????????????????????????????????????????????????????????????????????????????????????????????????????????????????????????????????????????????????????????????????????????????????????????????????????????????????????????????????????????????????????????????????????????????????????????????????????????????????????????????????????????????????????????????????????????????????????????????????????????????????????????????????????????????????????????????????????????????????????????????????????????????????????????????????????????????????????????????????????????????????????????????????????????????????????????????????????????????????????????????????????????????????????????????????????????????????????????????????

			>Vriesea_elata
			??????????????????????????????????????????????????????????????????????????????????????????????????????????????????????????????????????????????????????????????????????????????????????????????????????????????????????????????????????????????????????????????????????????????????????????????????????????????????????????????????????????????????????????????????????????????????????????????????????????????????????????????????????????????????????????????????????????????????????????????????????????????????????????????????????????????????????????????????????????????????????????????????????????????????????????????????????????????????????????????????????????????????????????????????????????????????????????????????????????????????????????????????????????????????????????????

			>Vriesea_eltoniana
			??????????????????????????????????????????????????????????????????????????????????????????????????????????????????????????????????????????????????????????????????????????????????????????????????????????????????????????????????????????????????????????????????????????????????????????????????????????????????????????????????????????????????????????????????????????????????????????????????????????????????????????????????????????????????????????????????????????????????????????????????????????????????????????????????????????????????????????????????????????????????????????????????????????????????????????????????????????????????????????????????????????????????????????????????????????????????????????????????????????????????????????????????????????????????????????????

			>Vriesea_ensiformis
			??????????????????????????????????????????????????????????????????????????????????????????????????????????????????????????????????????????????????????????????????????????????????????????????????????????????????????????????????????????????????????????????????????????????????????????????????????????????????????????????????????????????????????????????????????????????????????????????????????????????????????????????????????????????????????????????????????????????????????????????????????????????????????????????????????????????????????????????????????????????????????????????????????????????????????????????????????????????????????????????????????????????????????????????????????????????????????????????????????????????????????????????????????????????????????????????

			>Vriesea_ensiformis_var._bicolor
			??????????????????????????????????????????????????????????????????????????????????????????????????????????????????????????????????????????????????????????????????????????????????????????????????????????????????????????????????????????????????????????????????????????????????????????????????????????????????????????????????????????????????????????????????????????????????????????????????????????????????????????????????????????????????????????????????????????????????????????????????????????????????????????????????????????????????????????????????????????????????????????????????????????????????????????????????????????????????????????????????????????????????????????????????????????????????????????????????????????????????????????????????????????????????????????????

			>Vriesea_erythrodactylon
			??????????????????????????????????????????????????????????????????????????????????????????????????????????????????????????????????????????????????????????????????????????????????????????????????????????????????????????????????????????????????????????????????????????????????????????????????????????????????????????????????????????????????????????????????????????????????????????????????????????????????????????????????????????????????????????????????????????????????????????????????????????????????????????????????????????????????????????????????????????????????????????????????????????????????????????????????????????????????????????????????????????????????????????????????????????????????????????????????????????????????????????????????????????????????????????????

			>Vriesea_fenestralis
			??????????????????????????????????????????????????????????????????????????????????????????????????????????????????????????????????????????????????????????????????????????????????????????????????????????????????????????????????????????????????????????????????????????????????????????????????????????????????????????????????????????????????????????????????????????????????????????????????????????????????????????????????????????????????????????????????????????????????????????????????????????????????????????????????????????????????????????????????????????????????????????????????????????????????????????????????????????????????????????????????????????????????????????????????????????????????????????????????????????????????????????????????????????????????????????????

			>Vriesea_flammea
			??????????????????????????????????????????????????????????????????????????????????????????????????????????????????????????????????????????????????????????????????????????????????????????????????????????????????????????????????????????????????????????????????????????????????????????????????????????????????????????????????????????????????????????????????????????????????????????????????????????????????????????????????????????????????????????????????????????????????????????????????????????????????????????????????????????????????????????????????????????????????????????????????????????????????????????????????????????????????????????????????????????????????????????????????????????????????????????????????????????????????????????????????????????????????????????????

			>Vriesea_flava
			??????????????????????????????????????????????????????????????????????????????????????????????????????????????????????????????????????????????????????????????????????????????????????????????????????????????????????????????????????????????????????????????????????????????????????????????????????????????????????????????????????????????????????????????????????????????????????????????????????????????????????????????????????????????????????????????????????????????????????????????????????????????????????????????????????????????????????????????????????????????????????????????????????????????????????????????????????????????????????????????????????????????????????????????????????????????????????????????????????????????????????????????????????????????????????????????

			>Vriesea_fluviatilis
			??????????????????????????????????????????????????????????????????????????????????????????????????????????????????????????????????????????????????????????????????????????????????????????????????????????????????????????????????????????????????????????????????????????????????????????????????????????????????????????????????????????????????????????????????????????????????????????????????????????????????????????????????????????????????????????????????????????????????????????????????????????????????????????????????????????????????????????????????????????????????????????????????????????????????????????????????????????????????????????????????????????????????????????????????????????????????????????????????????????????????????????????????????????????????????????????

			>Vriesea_fosteriana
			??????????????????????????????????????????????????????????????????????????????????????????????????????????????????????????????????????????????????????????????????????????????????????????????????????????????????????????????????????????????????????????????????????????????????????????????????????????????????????????????????????????????????????????????????????????????????????????????????????????????????????????????????????????????????????????????????????????????????????????????????????????????????????????????????????????????????????????????????????????????????????????????????????????????????????????????????????????????????????????????????????????????????????????????????????????????????????????????????????????????????????????????????????????????????????????????

			>Vriesea_friburgensis
			??????????????????????????????????????????????????????????????????????????????????????????????????????????????????????????????????????????????????????????????????????????????????????????????????????????????????????????????????????????????????????????????????????????????????????????????????????????????????????????????????????????????????????????????????????????????????????????????????????????????????????????????????????????????????????????????????????????????????????????????????????????????????????????????????????????????????????????????????????????????????????????????????????????????????????????????????????????????????????????????????????????????????????????????????????????????????????????????????????????????????????????????????????????????????????????????

			>Vriesea_gigantea
			??????????????????????????????????????????????????????????????????????????????????????????????????????????????????????????????????????????????????????????????????????????????????????????????????????????????????????????????????????????????????????????????????????????????????????????????????????????????????????????????????????????????????????????????????????????????????????????????????????????????????????????????????????????????????????????????????????????????????????????????????????????????????????????????????????????????????????????????????????????????????????????????????????????????????????????????????????????????????????????????????????????????????????????????????????????????????????????????????????????????????????????????????????????????????????????????

			>Vriesea_gracilior
			??????????????????????????????????????????????????????????????????????????????????????????????????????????????????????????????????????????????????????????????????????????????????????????????????????????????????????????????????????????????????????????????????????????????????????????????????????????????????????????????????????????????????????????????????????????????????????????????????????????????????????????????????????????????????????????????????????????????????????????????????????????????????????????????????????????????????????????????????????????????????????????????????????????????????????????????????????????????????????????????????????????????????????????????????????????????????????????????????????????????????????????????????????????????????????????????

			>Vriesea_gradata
			??????????????????????????????????????????????????????????????????????????????????????????????????????????????????????????????????????????????????????????????????????????????????????????????????????????????????????????????????????????????????????????????????????????????????????????????????????????????????????????????????????????????????????????????????????????????????????????????????????????????????????????????????????????????????????????????????????????????????????????????????????????????????????????????????????????????????????????????????????????????????????????????????????????????????????????????????????????????????????????????????????????????????????????????????????????????????????????????????????????????????????????????????????????????????????????????

			>Vriesea_grandiflora
			??????????????????????????????????????????????????????????????????????????????????????????????????????????????????????????????????????????????????????????????????????????????????????????????????????????????????????????????????????????????????????????????????????????????????????????????????????????????????????????????????????????????????????????????????????????????????????????????????????????????????????????????????????????????????????????????????????????????????????????????????????????????????????????????????????????????????????????????????????????????????????????????????????????????????????????????????????????????????????????????????????????????????????????????????????????????????????????????????????????????????????????????????????????????????????????????

			>Vriesea_guttata
			??????????????????????????????????????????????????????????????????????????????????????????????????????????????????????????????????????????????????????????????????????????????????????????????????????????????????????????????????????????????????????????????????????????????????????????????????????????????????????????????????????????????????????????????????????????????????????????????????????????????????????????????????????????????????????????????????????????????????????????????????????????????????????????????????????????????????????????????????????????????????????????????????????????????????????????????????????????????????????????????????????????????????????????????????????????????????????????????????????????????????????????????????????????????????????????????

			>Vriesea_heterostachys
			??????????????????????????????????????????????????????????????????????????????????????????????????????????????????????????????????????????????????????????????????????????????????????????????????????????????????????????????????????????????????????????????????????????????????????????????????????????????????????????????????????????????????????????????????????????????????????????????????????????????????????????????????????????????????????????????????????????????????????????????????????????????????????????????????????????????????????????????????????????????????????????????????????????????????????????????????????????????????????????????????????????????????????????????????????????????????????????????????????????????????????????????????????????????????????????????

			>Vriesea_hydrophora
			??????????????????????????????????????????????????????????????????????????????????????????????????????????????????????????????????????????????????????????????????????????????????????????????????????????????????????????????????????????????????????????????????????????????????????????????????????????????????????????????????????????????????????????????????????????????????????????????????????????????????????????????????????????????????????????????????????????????????????????????????????????????????????????????????????????????????????????????????????????????????????????????????????????????????????????????????????????????????????????????????????????????????????????????????????????????????????????????????????????????????????????????????????????????????????????????

			>Vriesea_incurvata
			??????????????????????????????????????????????????????????????????????????????????????????????????????????????????????????????????????????????????????????????????????????????????????????????????????????????????????????????????????????????????????????????????????????????????????????????????????????????????????????????????????????????????????????????????????????????????????????????????????????????????????????????????????????????????????????????????????????????????????????????????????????????????????????????????????????????????????????????????????????????????????????????????????????????????????????????????????????????????????????????????????????????????????????????????????????????????????????????????????????????????????????????????????????????????????????????

			>Vriesea_inflata
			??????????????????????????????????????????????????????????????????????????????????????????????????????????????????????????????????????????????????????????????????????????????????????????????????????????????????????????????????????????????????????????????????????????????????????????????????????????????????????????????????????????????????????????????????????????????????????????????????????????????????????????????????????????????????????????????????????????????????????????????????????????????????????????????????????????????????????????????????????????????????????????????????????????????????????????????????????????????????????????????????????????????????????????????????????????????????????????????????????????????????????????????????????????????????????????????

			>Vriesea_interrogatoria
			??????????????????????????????????????????????????????????????????????????????????????????????????????????????????????????????????????????????????????????????????????????????????????????????????????????????????????????????????????????????????????????????????????????????????????????????????????????????????????????????????????????????????????????????????????????????????????????????????????????????????????????????????????????????????????????????????????????????????????????????????????????????????????????????????????????????????????????????????????????????????????????????????????????????????????????????????????????????????????????????????????????????????????????????????????????????????????????????????????????????????????????????????????????????????????????????

			>Vriesea_jonghei
			ATAATTATAATAAAGTGAAATATGTCGAAATCCTTTTTGGAATAATACCAAATCGAAAATAAATGTCCGATAGCAAGTTGATCAGTTAATTCAATAAGAGATAAATGGGAGATAGCACTCGATTTAGTTGGTACCACCCAACCGAATCCGATTCAATCGTTTACTCATTCAATGAGTAAATTTTCAAGTTCAGCCAATCTTTTTTTTTTCAAAAAAAAAATTGCAAGTAAATGAAATCGTGAATAAATGTGTTTCATTTCTCTATCATTATAGAAAAAAGGATCCATATTATATTACTTATGGAATTCGAGCCCGAACTCGATTTATGATTCATTATTTCGATCTTATTGGCCTTTGTTCTTTATTTTTTATTTTTCATATAGATTTCCGTCTTTATATTATACCCTTTCGTAGATGAATTATGCTTATTTTCACATCTAGGATTTACATATACAACATATATTACTGTCAAGAGGGAATTTTTCTCAGTATTTAGATATTTAGATTCAAAATAAGAAAGGGATCAATTCAATTATAAACCTGCAAAACAAAGATTAGGATTGGGTTGGGTTGCGCTATATATATCAAAGAGTATACAATAATGATGTATTTGGTGAATCAAATACATGGTCTAATAACAAACCATTTTAACATAACATTTTAATGAGTTGATAATATTAATTGAATATTGAATATTTTTTGAAAGATTTTTGTTAAAGGTTTCATTCATGCCTAATCCATATCGAGTAGACCTTGTTGTTGTGAG

			>Vriesea_limae
			??????????????????????????????????????????????????????????????????????????????????????????????????????????????????????????????????????????????????????????????????????????????????????????????????????????????????????????????????????????????????????????????????????????????????????????????????????????????????????????????????????????????????????????????????????????????????????????????????????????????????????????????????????????????????????????????????????????????????????????????????????????????????????????????????????????????????????????????????????????????????????????????????????????????????????????????????????????????????????????????????????????????????????????????????????????????????????????????????????????????????????????????????????????????????????????????

			>Vriesea_longicaulis
			??????????????????????????????????????????????????????????????????????????????????????????????????????????????????????????????????????????????????????????????????????????????????????????????????????????????????????????????????????????????????????????????????????????????????????????????????????????????????????????????????????????????????????????????????????????????????????????????????????????????????????????????????????????????????????????????????????????????????????????????????????????????????????????????????????????????????????????????????????????????????????????????????????????????????????????????????????????????????????????????????????????????????????????????????????????????????????????????????????????????????????????????????????????????????????????????

			>Vriesea_longistaminea
			??????????????????????????????????????????????????????????????????????????????????????????????????????????????????????????????????????????????????????????????????????????????????????????????????????????????????????????????????????????????????????????????????????????????????????????????????????????????????????????????????????????????????????????????????????????????????????????????????????????????????????????????????????????????????????????????????????????????????????????????????????????????????????????????????????????????????????????????????????????????????????????????????????????????????????????????????????????????????????????????????????????????????????????????????????????????????????????????????????????????????????????????????????????????????????????????

			>Vriesea_lubbersii
			??????????????????????????????????????????????????????????????????????????????????????????????????????????????????????????????????????????????????????????????????????????????????????????????????????????????????????????????????????????????????????????????????????????????????????????????????????????????????????????????????????????????????????????????????????????????????????????????????????????????????????????????????????????????????????????????????????????????????????????????????????????????????????????????????????????????????????????????????????????????????????????????????????????????????????????????????????????????????????????????????????????????????????????????????????????????????????????????????????????????????????????????????????????????????????????????

			>Vriesea_maxoniana
			??????????????????????????????????????????????????????????????????????????????????????????????????????????????????????????????????????????????????????????????????????????????????????????????????????????????????????????????????????????????????????????????????????????????????????????????????????????????????????????????????????????????????????????????????????????????????????????????????????????????????????????????????????????????????????????????????????????????????????????????????????????????????????????????????????????????????????????????????????????????????????????????????????????????????????????????????????????????????????????????????????????????????????????????????????????????????????????????????????????????????????????????????????????????????????????????

			>Vriesea_minuta
			??????????????????????????????????????????????????????????????????????????????????????????????????????????????????????????????????????????????????????????????????????????????????????????????????????????????????????????????????????????????????????????????????????????????????????????????????????????????????????????????????????????????????????????????????????????????????????????????????????????????????????????????????????????????????????????????????????????????????????????????????????????????????????????????????????????????????????????????????????????????????????????????????????????????????????????????????????????????????????????????????????????????????????????????????????????????????????????????????????????????????????????????????????????????????????????????

			>Vriesea_modesta
			??????????????????????????????????????????????????????????????????????????????????????????????????????????????????????????????????????????????????????????????????????????????????????????????????????????????????????????????????????????????????????????????????????????????????????????????????????????????????????????????????????????????????????????????????????????????????????????????????????????????????????????????????????????????????????????????????????????????????????????????????????????????????????????????????????????????????????????????????????????????????????????????????????????????????????????????????????????????????????????????????????????????????????????????????????????????????????????????????????????????????????????????????????????????????????????????

			>Vriesea_neoglutinosa
			??????????????????????????????????????????????????????????????????????????????????????????????????????????????????????????????????????????????????????????????????????????????????????????????????????????????????????????????????????????????????????????????????????????????????????????????????????????????????????????????????????????????????????????????????????????????????????????????????????????????????????????????????????????????????????????????????????????????????????????????????????????????????????????????????????????????????????????????????????????????????????????????????????????????????????????????????????????????????????????????????????????????????????????????????????????????????????????????????????????????????????????????????????????????????????????????

			>Vriesea_noblickii
			??????????????????????????????????????????????????????????????????????????????????????????????????????????????????????????????????????????????????????????????????????????????????????????????????????????????????????????????????????????????????????????????????????????????????????????????????????????????????????????????????????????????????????????????????????????????????????????????????????????????????????????????????????????????????????????????????????????????????????????????????????????????????????????????????????????????????????????????????????????????????????????????????????????????????????????????????????????????????????????????????????????????????????????????????????????????????????????????????????????????????????????????????????????????????????????????

			>Vriesea_ochracea
			??????????????????????????????????????????????????????????????????????????????????????????????????????????????????????????????????????????????????????????????????????????????????????????????????????????????????????????????????????????????????????????????????????????????????????????????????????????????????????????????????????????????????????????????????????????????????????????????????????????????????????????????????????????????????????????????????????????????????????????????????????????????????????????????????????????????????????????????????????????????????????????????????????????????????????????????????????????????????????????????????????????????????????????????????????????????????????????????????????????????????????????????????????????????????????????????

			>Vriesea_oligantha
			??????????????????????????????????????????????????????????????????????????????????????????????????????????????????????????????????????????????????????????????????????????????????????????????????????????????????????????????????????????????????????????????????????????????????????????????????????????????????????????????????????????????????????????????????????????????????????????????????????????????????????????????????????????????????????????????????????????????????????????????????????????????????????????????????????????????????????????????????????????????????????????????????????????????????????????????????????????????????????????????????????????????????????????????????????????????????????????????????????????????????????????????????????????????????????????????

			>Vriesea_pabstii
			??????????????????????????????????????????????????????????????????????????????????????????????????????????????????????????????????????????????????????????????????????????????????????????????????????????????????????????????????????????????????????????????????????????????????????????????????????????????????????????????????????????????????????????????????????????????????????????????????????????????????????????????????????????????????????????????????????????????????????????????????????????????????????????????????????????????????????????????????????????????????????????????????????????????????????????????????????????????????????????????????????????????????????????????????????????????????????????????????????????????????????????????????????????????????????????????

			>Vriesea_paraibica
			??????????????????????????????????????????????????????????????????????????????????????????????????????????????????????????????????????????????????????????????????????????????????????????????????????????????????????????????????????????????????????????????????????????????????????????????????????????????????????????????????????????????????????????????????????????????????????????????????????????????????????????????????????????????????????????????????????????????????????????????????????????????????????????????????????????????????????????????????????????????????????????????????????????????????????????????????????????????????????????????????????????????????????????????????????????????????????????????????????????????????????????????????????????????????????????????

			>Vriesea_pauperrima
			??????????????????????????????????????????????????????????????????????????????????????????????????????????????????????????????????????????????????????????????????????????????????????????????????????????????????????????????????????????????????????????????????????????????????????????????????????????????????????????????????????????????????????????????????????????????????????????????????????????????????????????????????????????????????????????????????????????????????????????????????????????????????????????????????????????????????????????????????????????????????????????????????????????????????????????????????????????????????????????????????????????????????????????????????????????????????????????????????????????????????????????????????????????????????????????????

			>Vriesea_platynema
			??????????????????????????????????????????????????????????????????????????????????????????????????????????????????????????????????????????????????????????????????????????????????????????????????????????????????????????????????????????????????????????????????????????????????????????????????????????????????????????????????????????????????????????????????????????????????????????????????????????????????????????????????????????????????????????????????????????????????????????????????????????????????????????????????????????????????????????????????????????????????????????????????????????????????????????????????????????????????????????????????????????????????????????????????????????????????????????????????????????????????????????????????????????????????????????????

			>Vriesea_platynema_var._rosea
			??????????????????????????????????????????????????????????????????????????????????????????????????????????????????????????????????????????????????????????????????????????????????????????????????????????????????????????????????????????????????????????????????????????????????????????????????????????????????????????????????????????????????????????????????????????????????????????????????????????????????????????????????????????????????????????????????????????????????????????????????????????????????????????????????????????????????????????????????????????????????????????????????????????????????????????????????????????????????????????????????????????????????????????????????????????????????????????????????????????????????????????????????????????????????????????????

			>Vriesea_platynema_var._variegata
			??????????????????????????????????????????????????????????????????????????????????????????????????????????????????????????????????????????????????????????????????????????????????????????????????????????????????????????????????????????????????????????????????????????????????????????????????????????????????????????????????????????????????????????????????????????????????????????????????????????????????????????????????????????????????????????????????????????????????????????????????????????????????????????????????????????????????????????????????????????????????????????????????????????????????????????????????????????????????????????????????????????????????????????????????????????????????????????????????????????????????????????????????????????????????????????????

			>Vriesea_procera
			??????????????????????????????????????????????????????????????????????????????????????????????????????????????????????????????????????????????????????????????????????????????????????????????????????????????????????????????????????????????????????????????????????????????????????????????????????????????????????????????????????????????????????????????????????????????????????????????????????????????????????????????????????????????????????????????????????????????????????????????????????????????????????????????????????????????????????????????????????????????????????????????????????????????????????????????????????????????????????????????????????????????????????????????????????????????????????????????????????????????????????????????????????????????????????????????

			>Vriesea_procera_var._tenuis
			??????????????????????????????????????????????????????????????????????????????????????????????????????????????????????????????????????????????????????????????????????????????????????????????????????????????????????????????????????????????????????????????????????????????????????????????????????????????????????????????????????????????????????????????????????????????????????????????????????????????????????????????????????????????????????????????????????????????????????????????????????????????????????????????????????????????????????????????????????????????????????????????????????????????????????????????????????????????????????????????????????????????????????????????????????????????????????????????????????????????????????????????????????????????????????????????

			>Vriesea_psittacina
			ATAATTATAATAAAGTGAAATATGTCGAAATCCTTTTTGGAATAATACCAAATCGAAAATAAATGTCCGATAGCAAGTTGATCAGTTAATTCAATAAGAGATAAATGGGAGATAGCACTCGATTTAGTTGGTACCACCCAACCGAATCCGATTCAATCGTTTACTCATTCAATGAGTAAATTTTCAAGTTCAGCCAATCTTTTTTTTTCA-AAAAAAAAATTGCAAGTAAATGAAATCGTGAATAAATGTGTTTCATTTCTCTATCATTATAGAAAAAAGGATCCATATTATATTACTTATGGAATTCGAGCCCGAACTCGATTTATGATTCATTATTTCGATCTTATTGGCCTTTGTTCTTTATTTTTGATTTTTCATATAGATTTCCGTCTTTATATTATACCCTTTCGTAGATGAATTATGCTTATTTTCACATCTAGGATTTACATATACAACATATATTACTGTCAAGAGGGAATTTTTCTCAGTATTTAGATATTTAGATTCAAAATAAGAAAGGGATCAATTCAATTATAAACCTGCAAAACAAAGATTAGGATTGGGTTGGGTTGCGCTATATATATCAAAGAGTATACAATAATGATGTATTTGGTGAATCAAATACATGGTCTAATAACAAACCATTTTAACATAACATTTTAATGAGTTGATAATATTAATTGAATATTGAATATTTTTTGAAAGATTTTTGTTAAAGGTTTCATTCATGCCTAATCCATATCGAGTAGACCTTGTTGTTGTGAG

			>Vriesea_recurvata
			??????????????????????????????????????????????????????????????????????????????????????????????????????????????????????????????????????????????????????????????????????????????????????????????????????????????????????????????????????????????????????????????????????????????????????????????????????????????????????????????????????????????????????????????????????????????????????????????????????????????????????????????????????????????????????????????????????????????????????????????????????????????????????????????????????????????????????????????????????????????????????????????????????????????????????????????????????????????????????????????????????????????????????????????????????????????????????????????????????????????????????????????????????????????????????????????

			>Vriesea_repandostachys
			??????????????????????????????????????????????????????????????????????????????????????????????????????????????????????????????????????????????????????????????????????????????????????????????????????????????????????????????????????????????????????????????????????????????????????????????????????????????????????????????????????????????????????????????????????????????????????????????????????????????????????????????????????????????????????????????????????????????????????????????????????????????????????????????????????????????????????????????????????????????????????????????????????????????????????????????????????????????????????????????????????????????????????????????????????????????????????????????????????????????????????????????????????????????????????????????

			>Vriesea_rhodostachys
			??????????????????????????????????????????????????????????????????????????????????????????????????????????????????????????????????????????????????????????????????????????????????????????????????????????????????????????????????????????????????????????????????????????????????????????????????????????????????????????????????????????????????????????????????????????????????????????????????????????????????????????????????????????????????????????????????????????????????????????????????????????????????????????????????????????????????????????????????????????????????????????????????????????????????????????????????????????????????????????????????????????????????????????????????????????????????????????????????????????????????????????????????????????????????????????????

			>Vriesea_rodigasiana
			??????????????????????????????????????????????????????????????????????????????????????????????????????????????????????????????????????????????????????????????????????????????????????????????????????????????????????????????????????????????????????????????????????????????????????????????????????????????????????????????????????????????????????????????????????????????????????????????????????????????????????????????????????????????????????????????????????????????????????????????????????????????????????????????????????????????????????????????????????????????????????????????????????????????????????????????????????????????????????????????????????????????????????????????????????????????????????????????????????????????????????????????????????????????????????????????

			>Vriesea_roethii
			??????????????????????????????????????????????????????????????????????????????????????????????????????????????????????????????????????????????????????????????????????????????????????????????????????????????????????????????????????????????????????????????????????????????????????????????????????????????????????????????????????????????????????????????????????????????????????????????????????????????????????????????????????????????????????????????????????????????????????????????????????????????????????????????????????????????????????????????????????????????????????????????????????????????????????????????????????????????????????????????????????????????????????????????????????????????????????????????????????????????????????????????????????????????????????????????

			>Vriesea_rubra
			??????????????????????????????????????????????????????????????????????????????????????????????????????????????????????????????????????????????????????????????????????????????????????????????????????????????????????????????????????????????????????????????????????????????????????????????????????????????????????????????????????????????????????????????????????????????????????????????????????????????????????????????????????????????????????????????????????????????????????????????????????????????????????????????????????????????????????????????????????????????????????????????????????????????????????????????????????????????????????????????????????????????????????????????????????????????????????????????????????????????????????????????????????????????????????????????

			>Vriesea_rubyae
			??????????????????????????????????????????????????????????????????????????????????????????????????????????????????????????????????????????????????????????????????????????????????????????????????????????????????????????????????????????????????????????????????????????????????????????????????????????????????????????????????????????????????????????????????????????????????????????????????????????????????????????????????????????????????????????????????????????????????????????????????????????????????????????????????????????????????????????????????????????????????????????????????????????????????????????????????????????????????????????????????????????????????????????????????????????????????????????????????????????????????????????????????????????????????????????????

			>Vriesea_ruschii_subsp._leonii
			??????????????????????????????????????????????????????????????????????????????????????????????????????????????????????????????????????????????????????????????????????????????????????????????????????????????????????????????????????????????????????????????????????????????????????????????????????????????????????????????????????????????????????????????????????????????????????????????????????????????????????????????????????????????????????????????????????????????????????????????????????????????????????????????????????????????????????????????????????????????????????????????????????????????????????????????????????????????????????????????????????????????????????????????????????????????????????????????????????????????????????????????????????????????????????????????

			>Vriesea_sanfranciscana
			??????????????????????????????????????????????????????????????????????????????????????????????????????????????????????????????????????????????????????????????????????????????????????????????????????????????????????????????????????????????????????????????????????????????????????????????????????????????????????????????????????????????????????????????????????????????????????????????????????????????????????????????????????????????????????????????????????????????????????????????????????????????????????????????????????????????????????????????????????????????????????????????????????????????????????????????????????????????????????????????????????????????????????????????????????????????????????????????????????????????????????????????????????????????????????????????

			>Vriesea_saundersii
			ATAATTATAATAAAGTGAAATATGTCGAAATCCTTTTTGGAATAATACCAAATCTAAAATAAATGTCCGATAGCAAGTTGATCAGTTAATTCAATAAGAGATAAATGGGAGATAGCACTCGATTTAGTTGGTACCACCCAACCGAATCCGATTCAATCGTTTACTCATTCAATGAGTAAATTTTCAAGTTCAGCCAATCTTTTTTTTTTCAAAAAAAAAATTGCAAGTAAATGAAATCGTGAATAAATGTGTTTCATTTCTCTATCATTATAGAAAAAAGGATCCATATTATATTACTTATGGAATTCGAGCCCGAACTCGATTTATGATTCATTATTTCGATCTTATTGGCCTTTGTTCTTTATTTTTTATTTTTCATATAGATTTCCGTCTTTATATTATACCCTTTCGTAGATGAATTATGCTTATTTTCACATCTAGGATTTACATATACAACATATATTACTGTCAAGAGGGAATTTTTCTCAGTATTTAGATATTTAGATTCAAAATAAGAAAGGGATCAATTCAATTATAAACCTGCAAAACAAAGATTAGGATTGGGTTGGGTTGCGCTATATATATCAAAGAGTATACAATAATGATGTATTTGGTGAATCAAATACATGGTCTAATAACAAACCATTTTAACATAACATTTTAATGAGTTGATAATATTAATTGAATATTGAATATTTTTTGAAAGATTTTTGTTAAAGGTTTCATTCATGCCTAATCCATATCGAGTAGACCTTGTTGTTGTGAG

			>Vriesea_sazimae
			??????????????????????????????????????????????????????????????????????????????????????????????????????????????????????????????????????????????????????????????????????????????????????????????????????????????????????????????????????????????????????????????????????????????????????????????????????????????????????????????????????????????????????????????????????????????????????????????????????????????????????????????????????????????????????????????????????????????????????????????????????????????????????????????????????????????????????????????????????????????????????????????????????????????????????????????????????????????????????????????????????????????????????????????????????????????????????????????????????????????????????????????????????????????????????????????

			>Vriesea_scalaris
			??????????????????????????????????????????????????????????????????????????????????????????????????????????????????????????????????????????????????????????????????????????????????????????????????????????????????????????????????????????????????????????????????????????????????????????????????????????????????????????????????????????????????????????????????????????????????????????????????????????????????????????????????????????????????????????????????????????????????????????????????????????????????????????????????????????????????????????????????????????????????????????????????????????????????????????????????????????????????????????????????????????????????????????????????????????????????????????????????????????????????????????????????????????????????????????????

			>Vriesea_scalaris_var._viridis
			??????????????????????????????????????????????????????????????????????????????????????????????????????????????????????????????????????????????????????????????????????????????????????????????????????????????????????????????????????????????????????????????????????????????????????????????????????????????????????????????????????????????????????????????????????????????????????????????????????????????????????????????????????????????????????????????????????????????????????????????????????????????????????????????????????????????????????????????????????????????????????????????????????????????????????????????????????????????????????????????????????????????????????????????????????????????????????????????????????????????????????????????????????????????????????????????

			>Vriesea_segadas-viannae
			??????????????????????????????????????????????????????????????????????????????????????????????????????????????????????????????????????????????????????????????????????????????????????????????????????????????????????????????????????????????????????????????????????????????????????????????????????????????????????????????????????????????????????????????????????????????????????????????????????????????????????????????????????????????????????????????????????????????????????????????????????????????????????????????????????????????????????????????????????????????????????????????????????????????????????????????????????????????????????????????????????????????????????????????????????????????????????????????????????????????????????????????????????????????????????????????

			>Vriesea_seideliana
			??????????????????????????????????????????????????????????????????????????????????????????????????????????????????????????????????????????????????????????????????????????????????????????????????????????????????????????????????????????????????????????????????????????????????????????????????????????????????????????????????????????????????????????????????????????????????????????????????????????????????????????????????????????????????????????????????????????????????????????????????????????????????????????????????????????????????????????????????????????????????????????????????????????????????????????????????????????????????????????????????????????????????????????????????????????????????????????????????????????????????????????????????????????????????????????????

			>Vriesea_simplex
			??????????????????????????????????????????????????????????????????????????????????????????????????????????????????????????????????????????????????????????????????????????????????????????????????????????????????????????????????????????????????????????????????????????????????????????????????????????????????????????????????????????????????????????????????????????????????????????????????????????????????????????????????????????????????????????????????????????????????????????????????????????????????????????????????????????????????????????????????????????????????????????????????????????????????????????????????????????????????????????????????????????????????????????????????????????????????????????????????????????????????????????????????????????????????????????????

			>Vriesea_sincorana
			??????????????????????????????????????????????????????????????????????????????????????????????????????????????????????????????????????????????????????????????????????????????????????????????????????????????????????????????????????????????????????????????????????????????????????????????????????????????????????????????????????????????????????????????????????????????????????????????????????????????????????????????????????????????????????????????????????????????????????????????????????????????????????????????????????????????????????????????????????????????????????????????????????????????????????????????????????????????????????????????????????????????????????????????????????????????????????????????????????????????????????????????????????????????????????????????

			>Vriesea_sparsiflora
			??????????????????????????????????????????????????????????????????????????????????????????????????????????????????????????????????????????????????????????????????????????????????????????????????????????????????????????????????????????????????????????????????????????????????????????????????????????????????????????????????????????????????????????????????????????????????????????????????????????????????????????????????????????????????????????????????????????????????????????????????????????????????????????????????????????????????????????????????????????????????????????????????????????????????????????????????????????????????????????????????????????????????????????????????????????????????????????????????????????????????????????????????????????????????????????????

			>Vriesea_sucrei
			??????????????????????????????????????????????????????????????????????????????????????????????????????????????????????????????????????????????????????????????????????????????????????????????????????????????????????????????????????????????????????????????????????????????????????????????????????????????????????????????????????????????????????????????????????????????????????????????????????????????????????????????????????????????????????????????????????????????????????????????????????????????????????????????????????????????????????????????????????????????????????????????????????????????????????????????????????????????????????????????????????????????????????????????????????????????????????????????????????????????????????????????????????????????????????????????

			>Vriesea_taritubensis
			??????????????????????????????????????????????????????????????????????????????????????????????????????????????????????????????????????????????????????????????????????????????????????????????????????????????????????????????????????????????????????????????????????????????????????????????????????????????????????????????????????????????????????????????????????????????????????????????????????????????????????????????????????????????????????????????????????????????????????????????????????????????????????????????????????????????????????????????????????????????????????????????????????????????????????????????????????????????????????????????????????????????????????????????????????????????????????????????????????????????????????????????????????????????????????????????

			>Vriesea_taritubensis_var._brevisepala
			??????????????????????????????????????????????????????????????????????????????????????????????????????????????????????????????????????????????????????????????????????????????????????????????????????????????????????????????????????????????????????????????????????????????????????????????????????????????????????????????????????????????????????????????????????????????????????????????????????????????????????????????????????????????????????????????????????????????????????????????????????????????????????????????????????????????????????????????????????????????????????????????????????????????????????????????????????????????????????????????????????????????????????????????????????????????????????????????????????????????????????????????????????????????????????????????

			>Vriesea_taritubensis_var._patens
			??????????????????????????????????????????????????????????????????????????????????????????????????????????????????????????????????????????????????????????????????????????????????????????????????????????????????????????????????????????????????????????????????????????????????????????????????????????????????????????????????????????????????????????????????????????????????????????????????????????????????????????????????????????????????????????????????????????????????????????????????????????????????????????????????????????????????????????????????????????????????????????????????????????????????????????????????????????????????????????????????????????????????????????????????????????????????????????????????????????????????????????????????????????????????????????????

			>Vriesea_teresopolitana
			??????????????????????????????????????????????????????????????????????????????????????????????????????????????????????????????????????????????????????????????????????????????????????????????????????????????????????????????????????????????????????????????????????????????????????????????????????????????????????????????????????????????????????????????????????????????????????????????????????????????????????????????????????????????????????????????????????????????????????????????????????????????????????????????????????????????????????????????????????????????????????????????????????????????????????????????????????????????????????????????????????????????????????????????????????????????????????????????????????????????????????????????????????????????????????????????

			>Vriesea_tijucana
			??????????????????????????????????????????????????????????????????????????????????????????????????????????????????????????????????????????????????????????????????????????????????????????????????????????????????????????????????????????????????????????????????????????????????????????????????????????????????????????????????????????????????????????????????????????????????????????????????????????????????????????????????????????????????????????????????????????????????????????????????????????????????????????????????????????????????????????????????????????????????????????????????????????????????????????????????????????????????????????????????????????????????????????????????????????????????????????????????????????????????????????????????????????????????????????????

			>Vriesea_unilateralis
			??????????????????????????????????????????????????????????????????????????????????????????????????????????????????????????????????????????????????????????????????????????????????????????????????????????????????????????????????????????????????????????????????????????????????????????????????????????????????????????????????????????????????????????????????????????????????????????????????????????????????????????????????????????????????????????????????????????????????????????????????????????????????????????????????????????????????????????????????????????????????????????????????????????????????????????????????????????????????????????????????????????????????????????????????????????????????????????????????????????????????????????????????????????????????????????????

			>Vriesea_vagans
			??????????????????????????????????????????????????????????????????????????????????????????????????????????????????????????????????????????????????????????????????????????????????????????????????????????????????????????????????????????????????????????????????????????????????????????????????????????????????????????????????????????????????????????????????????????????????????????????????????????????????????????????????????????????????????????????????????????????????????????????????????????????????????????????????????????????????????????????????????????????????????????????????????????????????????????????????????????????????????????????????????????????????????????????????????????????????????????????????????????????????????????????????????????????????????????????

			>Vriesea_zamorensis
			ATAATTATAATAAAGTGAAATATGTCGAAATCCTTTTTGGAATAATACCAAATCGAAAATAAATGTCCGATAGCAAGTTGATCAGTTAATTCAATAAGAGATAAATGGGAGATAGCACTCGATTTAGTTGGTACCACCCAACCGAATCCGATTCAATCGTTTACTCATTCAATGAGTCAATTTTCAAGTTCAGCCAATCTTTTTTTTTTT------TTTATTGCAAATAAATGAAATCGTGAATAAATGTGTTTCATTTCTCTATCATTATAGAAAAAAAGATCCATATTATATTACTTATGGAATTCGAACCCGAACTCGATTTATGATTCATTATTTCGATCTTATTGGCCTTTGTTCTTTATTTTTGATTTTTCATATAGATTTCCGTCTTTATATTATACCCTTTCGTAGATGAATTATGCTTATTTTCACATCTAGGATTTACATATACAACATATATTACTGTCAAGAGGGAATTTTTCTCAGTATTTAGATATTTAGATTCAAAATAAGAAAGGGATCAATTCAATTATAAACCTGCAAAACAAAGATTAGGATTGGGTTGGGTTGCGCTATATATATCAAAGAGTATACAATAATGATGTATTTGGTGAATCAAATACATGGTCTAATAACAAACCATTTTAACATAACATTTTAATGAGTTGATAATATTAATTGAATATTGAATATTTTTTGAAAGATTTTTGTTAAAGGTTTCATTCATGCCTAATCCATATCGAGTAGACCTTGTTGTTGTGAG

			>Vriesea_zonata
			??????????????????????????????????????????????????????????????????????????????????????????????????????????????????????????????????????????????????????????????????????????????????????????????????????????????????????????????????????????????????????????????????????????????????????????????????????????????????????????????????????????????????????????????????????????????????????????????????????????????????????????????????????????????????????????????????????????????????????????????????????????????????????????????????????????????????????????????????????????????????????????????????????????????????????????????????????????????????????????????????????????????????????????????????????????????????????????????????????????????????????????????????????????????????????????????

			>Wallisia_anceps
			??????????????????????????????????????????????????????????????????????????????????????????????????????????????????????????????????????????????????????????????????????????????????????????????????????????????????????????????????????????????????????????????????????????????????????????????????????????????????????????????????????????????????????????????????????????????????????????????????????????????????????????????????????????????????????????????????????????????????????????????????????????????????????????????????????????????????????????????????????????????????????????????????????????????????????????????????????????????????????????????????????????????????????????????????????????????????????????????????????????????????????????????????????????????????????????????

			>Wallisia_cyanea
			ATAATTATAATAAAGTGAAATATGTCGAAATCCTTTTTGGAATAATACCAAATCGAAAATAAATGTCCGATAGCAAGTTGATCAGTTAATTCAATAAGAGATAAATGGGAGATAGCACTCGATTTAGTTGGTACCACCCAACCGAATCCGATTCAATCGTTTACTCA------GAGTAAATTTTCAAGTTCAGCCAATC--------------------------------------------------------------------------------------------------------------------------------------------------------C--------TTTATTTTTTATTTTTCATATGGATTTCCGTCTTTATATTATACCCTTTCGTAGATGAATTATGCTTATTTTCACATCTAGGATTTACATATACAACATATATTACTGTCAAGAGGGAATTTTTCTCAGTATTTAGATATTTAGATTCAAAATAAGAAAGGGATCAATTCAATTATAAACCCGCAAAACAAAGATTAGGATTGGGTTGGGTTGCGCTATATATATCAAAGAGTATACAATAATGATGTATTTGGTGAATCAAATACATGGTCTAATAACAAACCATTTTAACATAACATTTTAATGAGTTGATAATATTAATTGAATA-------TTTTTTGAAAGATTTTTGTTAAAGGTTTCATTCATGCCTAATCCATATCGAGTAGACCTTGTTGTTGTGAG

			>Wallisia_lindeniana
			??????????????????????????????????????????????????????????????????????????????????????????????????????????????????????????????????????????????????????????????????????????????????????????????????????????????????????????????????????????????????????????????????????????????????????????????????????????????????????????????????????????????????????????????????????????????????????????????????????????????????????????????????????????????????????????????????????????????????????????????????????????????????????????????????????????????????????????????????????????????????????????????????????????????????????????????????????????????????????????????????????????????????????????????????????????????????????????????????????????????????????????????????????????????????????????????

			>Waltillia_hatschbachii
			??????????????????????????????????????????????????????????????????????????????????????????????????????????????????????????????????????????????????????????????????????????????????????????????????????????????????????????????????????????????????????????????????????????????????????????????????????????????????????????????????????????????????????????????????????????????????????????????????????????????????????????????????????????????????????????????????????????????????????????????????????????????????????????????????????????????????????????????????????????????????????????????????????????????????????????????????????????????????????????????????????????????????????????????????????????????????????????????????????????????????????????????????????????????????????????????

			>Werauhia_barii
			??????????????????????????????????????????????????????????????????????????????????????????????????????????????????????????????????????????????????????????????????????????????????????????????????????????????????????????????????????????????????????????????????????????????????????????????????????????????????????????????????????????????????????????????????????????????????????????????????????????????????????????????????????????????????????????????????????????????????????????????????????????????????????????????????????????????????????????????????????????????????????????????????????????????????????????????????????????????????????????????????????????????????????????????????????????????????????????????????????????????????????????????????????????????????????????????

			>Werauhia_bicolor
			??????????????????????????????????????????????????????????????????????????????????????????????????????????????????????????????????????????????????????????????????????????????????????????????????????????????????????????????????????????????????????????????????????????????????????????????????????????????????????????????????????????????????????????????????????????????????????????????????????????????????????????????????????????????????????????????????????????????????????????????????????????????????????????????????????????????????????????????????????????????????????????????????????????????????????????????????????????????????????????????????????????????????????????????????????????????????????????????????????????????????????????????????????????????????????????????

			>Werauhia_burgeri
			??????????????????????????????????????????????????????????????????????????????????????????????????????????????????????????????????????????????????????????????????????????????????????????????????????????????????????????????????????????????????????????????????????????????????????????????????????????????????????????????????????????????????????????????????????????????????????????????????????????????????????????????????????????????????????????????????????????????????????????????????????????????????????????????????????????????????????????????????????????????????????????????????????????????????????????????????????????????????????????????????????????????????????????????????????????????????????????????????????????????????????????????????????????????????????????????

			>Werauhia_gladioliflora
			??????????????????????????????????????????????????????????????????????????????????????????????????????????????????????????????????????????????????????????????????????????????????????????????????????????????????????????????????????????????????????????????????????????????????????????????????????????????????????????????????????????????????????????????????????????????????????????????????????????????????????????????????????????????????????????????????????????????????????????????????????????????????????????????????????????????????????????????????????????????????????????????????????????????????????????????????????????????????????????????????????????????????????????????????????????????????????????????????????????????????????????????????????????????????????????????

			>Werauhia_insignis
			ATAATTATAATAAAGTGAAATATGTCGAAATCCTTTTTGGAATAATACCAAATCGAAAATAAATGTCCGATAGCAAGTTGATCAGTTAATTCAATAAGAGATAAATGGGAGATAGCACTCGATTTAGTTGGTACCACCCAACCGAATCCGATTCAATCGTTTACTCATTCAATGAGTCAATTTTCAAGTTCAGCCAATCTTTTTTTTGAAAAAAAAAAAATTGCAAGTAAATGAAATCGTGAATAAATGTGTTTCATTTCTCTATCATTATAGAAAAAAGGATCCATATTATATTACTTATGGAATTCGAACCCGAACTCGATTTATGATTCATTATTTCGATCTTATTGGCCTTTGTTCTTTATTTTTGATTTTTCATATAGATTTCCGTCTTTATATTATACCCTTTCGTAGATGAATTATGCTTATTTTCACATCTAGGATTTACATATACAACATATATTACTGTCAAGAGGGAATTTTTCTCAGTATTTAGATATTTAGATTCAAAATAAGAAAGGGATCAATTCAATTATAAACCTGCAAAACAAAGATTAGGATTGGGTTGGGTTGCGCTATATATATCAAAGAGTATACAATAATGATGTATTTGGTGAATCAAATACATGGTCTAATAACAAACCATTTTAACATAACATTTTAATGAGTTGATAATATTAATTGAATATTGAATATTTTTTGAAAGATTTTTGTTAAAGGTTTCATTCATGCCTAATCCATATCGAGTAGACCTTGTTGTTGTGAG

			>Werauhia_ororiensis
			ATAATTATAATAAAGTGAAATATGTCGAAATCCTTTTTGGAATAATACCAAATCGAAAATAAATGTCCGATAGCAAGTTGATCAGTTAATTCAATAAGAGATAAATGGGAGATAGCACTCGATTTAGTTGGTACCACCCAACCGAATCCGATTCAATCGTTTACTCATTCAATGAGTCAATTTTCAAGTTCAGCCAATCTTTTTTTTTCAAAAAAAAAAATTGCAAGTAAATGAAATCGTGAATAAATGTGTTTCATTTCTCTATCATTATAGAAAAAAGGATCCATATTATATTACTTATGGAATTCGAACCCGAACTCGATTTATGATTCATTATTTCGATCTTATTGGCCTTTGTTCTCTATTTTTGATTTTTCATATAGATTTCCGTCTTTATATTATACCCTTTCGTAGATGAATTATGCTTATTTTCACATCTAGGATTTACATATACAACATATATTACTGTCAAGAGGGAATTTTTCTCAGTATTTAGATATTTAGATTCAAAATAAGAAAGGGATCAATTCAATTATAAACCTGCAAAACAAAGATTAGGATTGGGTTGGGTTGCGCTATATATATCAAAGAGTATACAATAATGATGTATTTGGTGAATCAAATACATGGTCTAATAACAAACCATTTTAACATAACATTTTAATGAGTTGATAATATTAATTGAATATTGAATATTTTTTGAAAGATTTTTGTTAAAGGTTTCATTCATGCCTAATCCATATCGAGTAGACCTTGTTGTTGTGAG

			>Werauhia_patzeltii
			??????????????????????????????????????????????????????????????????????????????????????????????????????????????????????????????????????????????????????????????????????????????????????????????????????????????????????????????????????????????????????????????????????????????????????????????????????????????????????????????????????????????????????????????????????????????????????????????????????????????????????????????????????????????????????????????????????????????????????????????????????????????????????????????????????????????????????????????????????????????????????????????????????????????????????????????????????????????????????????????????????????????????????????????????????????????????????????????????????????????????????????????????????????????????????????????

			>Werauhia_pectinata
			??????????????????????????????????????????????????????????????????????????????????????????????????????????????????????????????????????????????????????????????????????????????????????????????????????????????????????????????????????????????????????????????????????????????????????????????????????????????????????????????????????????????????????????????????????????????????????????????????????????????????????????????????????????????????????????????????????????????????????????????????????????????????????????????????????????????????????????????????????????????????????????????????????????????????????????????????????????????????????????????????????????????????????????????????????????????????????????????????????????????????????????????????????????????????????????????

			>Werauhia_pedicellata
			??????????????????????????????????????????????????????????????????????????????????????????????????????????????????????????????????????????????????????????????????????????????????????????????????????????????????????????????????????????????????????????????????????????????????????????????????????????????????????????????????????????????????????????????????????????????????????????????????????????????????????????????????????????????????????????????????????????????????????????????????????????????????????????????????????????????????????????????????????????????????????????????????????????????????????????????????????????????????????????????????????????????????????????????????????????????????????????????????????????????????????????????????????????????????????????????

			>Werauhia_ringens
			ATAATTATAATAAAGTGAAATATGTCGAAATCCTTTTTGGAATAATACCAAATCGAAAATAAATGTCCGATAGCAAGTTGATCAGTTAATTCAATAAGAGATAAATGGGAGATAGCACTCGATTTGGTTGGTACCACCCAACCGAATCCGATTCAATCGTTTACTCATTCAATGAGTCAATTTTCAAGTTCAGCCAATCTTTTTTTTTGAAAAAAAAAAATTGCAAGTAAATGAAATCGTGAATAAATGTGTTTCATTTCTCTATCATTATAGAAAAAAAGATCCATATTATATTACTTATGGAATTCGAACCCGAACTCGATTTATGATTCATTATTTCGATCTTATTGGCCTTTGTTCTTTATTTTTGATTTTTCATATAGATTTCCGTCTTTATATTATACCCTTTCGTAGATGAATTATGCTTATTTTCACATCTAGGATTTACATATACAACATATATTACTGTCAAGAGGGAATTTTTCTCAGTATTTAGATATTTAGATTCAAAATAAGAAAGGGATCAATTCAATTATAAACCTGCAAAACAAAGATTAGGATTGGGTTGGGTTGCGCTATATATATCAAAGAGTATACAATAATGATGTATTTGGTGAATCAAATACATGGTCTAATAACAAACCATTTTAACATAACATTTTAATGAGTTGATAATATTAATTGAATATTGAATATTTTTTGAAAGATTTTTGTTAAAGGTTTCATTCATGCCTAATCCATATCGAGTAGACCTTGTTGTTGTGAG

			>Werauhia_sanguinolenta
			??????????????????????????????????????????????????????????????????????????????????????????????????????????????????????????????????????????????????????????????????????????????????????????????????????????????????????????????????????????????????????????????????????????????????????????????????????????????????????????????????????????????????????????????????????????????????????????????????????????????????????????????????????????????????????????????????????????????????????????????????????????????????????????????????????????????????????????????????????????????????????????????????????????????????????????????????????????????????????????????????????????????????????????????????????????????????????????????????????????????????????????????????????????????????????????????

			>Werauhia_tarmaensis
			ATAATTATAATAAAGTGAAATATGTCGAAATCCTTTTTGGAATAATACCAAATCGAAAATAAATGTCCGATAGCAAGTTGATCAGTTAATTCAATAAGAGATAAATGGGAGATAGCACTCGATTTAGTTGGTACCACCCAACCGAATCCGATTCAATCGTTTACTCATTCAATGAGTCAATTTTCAAGTTCAGCCAATCTTTTTTTTTTCAAAAAAAAAATTGCAAGTAAATGAAATCGTGAATAAATGTGTTTCATTTCTCTATCATTATAGAAAAAAAGATCCATATTATATTACTTATGGAATTCGAACCCGAACTCGATTTATGATTCATTATTTCGATCTTATTGGCCTTTGTTCTTTATTTTTGATTTTTCATATAGATTTCCGTCTTTATATTATACCCTTTCGTAGATGAATTATGCTTATTTTCACATCTAGGATTTACATATACAACATATATTACTGTCAAGAGGGAATTTTTCTCAGTATTTAGATATTTAGATTCAAAATAAGAAAGGGA-----TCAATTATAAACCTGCAAAACAAAGATTAGGATTGGGTTGGGTTGCGCTATATATATCAAAGAGTATACAATAATGATGTATTTGGTGAATCAAATACATGGTCTAATAACAAACCATTTTAACATAACATTTTAATGAGTTGATAATATTAATTGAATATTGAATATTTTTTGAAAGATTTTTGTTAAAGGTTTCATTCATGCCTAATCCATATCGAGTAGACCTTGTTGTTGTGAG

			>Werauhia_viridiflora
			ATAATTATAATAAAGTGAAATATGTCGAAATCCTTTTTGGAATAATACCAAATCGAAAATAAATGTCCGATAGCAAGTTGATCAGTTAATTCAATAAGAGATAAATGGGAGATAGCACTCGATTTAGTTGGTACCACCCAACCGAATCCGATTCAATCGTTTACTCATTCAATGAGTCAATTTTCAAGTTCAGCCAATCTTTTTTTTTG--AAAAAAAAATTGCAAGTAAATGAAATCGTGAATAAATGTGTTTCATTTCTCTATCATTATAGAAAAAAAGATCCATATTATATTACTTATGGAATTCGAACCCGAACTCGATTTATGATTCATTATTTCGATCTTATTGGCCTTTGTTCTTTATTTTTGATTTTTCATATAGATTTCCGTCTTTATATTATACCCTTTCGTAGATGAATTATGCTTATTTTCACATCTAGGATTTACATATACAACATATATTACTGTCAAGAGGGAATTTTTCTCAGTATTTAGATATTTAGATTCAAAATAAGAAAGGGATCAATTCAATTATAAACCTGCAAAACAAAGATTAGGATTGGGTTGGGTTGCGCTATATATATCAAAGAGTATACAATAATGATGTATTTGGTGAATCAAATACATGGTCTAATAACAAACCATTTTAACATAACATTTTAATGAGTTGATAATATTAATTGAATATTGAATATTTTTTGAAAGATTTTTGTTAAAGGTTTCATTCATGCCTAATCCATATCGAGTAGACCTTGTTGTTGTGAG

			>Werauhia_vittata
			??????????????????????????????????????????????????????????????????????????????????????????????????????????????????????????????????????????????????????????????????????????????????????????????????????????????????????????????????????????????????????????????????????????????????????????????????????????????????????????????????????????????????????????????????????????????????????????????????????????????????????????????????????????????????????????????????????????????????????????????????????????????????????????????????????????????????????????????????????????????????????????????????????????????????????????????????????????????????????????????????????????????????????????????????????????????????????????????????????????????????????????????????????????????????????????????

			>Wittmackia_abbreviata
			??????????????????????????????????????????????????????????????????????????????????????????????????????????????????????????????????????????????????????????????????????????????????????????????????????????????????????????????????????????????????????????????????????????????????????????????????????????????????????????????????????????????????????????????????????????????????????????????????????????????????????????????????????????????????????????????????????????????????????????????????????????????????????????????????????????????????????????????????????????????????????????????????????????????????????????????????????????????????????????????????????????????????????????????????????????????????????????????????????????????????????????????????????????????????????????????

			>Wittmackia_altocaririensis
			??????????????????????????????????????????????????????????????????????????????????????????????????????????????????????????????????????????????????????????????????????????????????????????????????????????????????????????????????????????????????????????????????????????????????????????????????????????????????????????????????????????????????????????????????????????????????????????????????????????????????????????????????????????????????????????????????????????????????????????????????????????????????????????????????????????????????????????????????????????????????????????????????????????????????????????????????????????????????????????????????????????????????????????????????????????????????????????????????????????????????????????????????????????????????????????????

			>Wittmackia_amorimii
			??????????????????????????????????????????????????????????????????????????????????????????????????????????????????????????????????????????????????????????????????????????????????????????????????????????????????????????????????????????????????????????????????????????????????????????????????????????????????????????????????????????????????????????????????????????????????????????????????????????????????????????????????????????????????????????????????????????????????????????????????????????????????????????????????????????????????????????????????????????????????????????????????????????????????????????????????????????????????????????????????????????????????????????????????????????????????????????????????????????????????????????????????????????????????????????????

			>Wittmackia_andersoniana
			??????????????????????????????????????????????????????????????????????????????????????????????????????????????????????????????????????????????????????????????????????????????????????????????????????????????????????????????????????????????????????????????????????????????????????????????????????????????????????????????????????????????????????????????????????????????????????????????????????????????????????????????????????????????????????????????????????????????????????????????????????????????????????????????????????????????????????????????????????????????????????????????????????????????????????????????????????????????????????????????????????????????????????????????????????????????????????????????????????????????????????????????????????????????????????????????

			>Wittmackia_antillana
			??????????????????????????????????????????????????????????????????????????????????????????????????????????????????????????????????????????????????????????????????????????????????????????????????????????????????????????????????????????????????????????????????????????????????????????????????????????????????????????????????????????????????????????????????????????????????????????????????????????????????????????????????????????????????????????????????????????????????????????????????????????????????????????????????????????????????????????????????????????????????????????????????????????????????????????????????????????????????????????????????????????????????????????????????????????????????????????????????????????????????????????????????????????????????????????????

			>Wittmackia_bicolor
			??????????????????????????????????????????????????????????????????????????????????????????????????????????????????????????????????????????????????????????????????????????????????????????????????????????????????????????????????????????????????????????????????????????????????????????????????????????????????????????????????????????????????????????????????????????????????????????????????????????????????????????????????????????????????????????????????????????????????????????????????????????????????????????????????????????????????????????????????????????????????????????????????????????????????????????????????????????????????????????????????????????????????????????????????????????????????????????????????????????????????????????????????????????????????????????????

			>Wittmackia_brasiliensis
			??????????????????????????????????????????????????????????????????????????????????????????????????????????????????????????????????????????????????????????????????????????????????????????????????????????????????????????????????????????????????????????????????????????????????????????????????????????????????????????????????????????????????????????????????????????????????????????????????????????????????????????????????????????????????????????????????????????????????????????????????????????????????????????????????????????????????????????????????????????????????????????????????????????????????????????????????????????????????????????????????????????????????????????????????????????????????????????????????????????????????????????????????????????????????????????????

			>Wittmackia_burle-marxii
			??????????????????????????????????????????????????????????????????????????????????????????????????????????????????????????????????????????????????????????????????????????????????????????????????????????????????????????????????????????????????????????????????????????????????????????????????????????????????????????????????????????????????????????????????????????????????????????????????????????????????????????????????????????????????????????????????????????????????????????????????????????????????????????????????????????????????????????????????????????????????????????????????????????????????????????????????????????????????????????????????????????????????????????????????????????????????????????????????????????????????????????????????????????????????????????????

			>Wittmackia_canaliculata
			??????????????????????????????????????????????????????????????????????????????????????????????????????????????????????????????????????????????????????????????????????????????????????????????????????????????????????????????????????????????????????????????????????????????????????????????????????????????????????????????????????????????????????????????????????????????????????????????????????????????????????????????????????????????????????????????????????????????????????????????????????????????????????????????????????????????????????????????????????????????????????????????????????????????????????????????????????????????????????????????????????????????????????????????????????????????????????????????????????????????????????????????????????????????????????????????

			>Wittmackia_caymanensis
			??????????????????????????????????????????????????????????????????????????????????????????????????????????????????????????????????????????????????????????????????????????????????????????????????????????????????????????????????????????????????????????????????????????????????????????????????????????????????????????????????????????????????????????????????????????????????????????????????????????????????????????????????????????????????????????????????????????????????????????????????????????????????????????????????????????????????????????????????????????????????????????????????????????????????????????????????????????????????????????????????????????????????????????????????????????????????????????????????????????????????????????????????????????????????????????????

			>Wittmackia_distans
			??????????????????????????????????????????????????????????????????????????????????????????????????????????????????????????????????????????????????????????????????????????????????????????????????????????????????????????????????????????????????????????????????????????????????????????????????????????????????????????????????????????????????????????????????????????????????????????????????????????????????????????????????????????????????????????????????????????????????????????????????????????????????????????????????????????????????????????????????????????????????????????????????????????????????????????????????????????????????????????????????????????????????????????????????????????????????????????????????????????????????????????????????????????????????????????????

			>Wittmackia_eriostachya
			?????????ATAAAGTGAAATATGTCGAAATCCTTTTTGGAATAATACCAAATCGAAAATAAATGTCCGATAGCAAGTTGATCAGTTAATTCAATAAGAGATAAATGGGAGATAGCACTCGATTTAGTTGGTACCACCCAACCGAATCCGATTCACTCGTTTACTCATTCAATGAGTAAATTTTCAAGTTCAGCCAATC--TTTTTTTTGAAAAAAAAAATTGCAAGTAGATGAAATCGTGAATAAATGTGTTTCATTTCTATATCATTATATAAAAAAAGATCCATATTATATTACTTATGGAATTCGAACCCGAACTCGATTTATGATTTATTATTTGGATCTTATTGGCCTTTGTTCTTTATTTTTGATTTTTCATATGGATTTCCGTCTTTATATTATACCCTTTCGTGGATGAATTATGCCTATTTTCACATCTAGGATTTACATATACAACATATATTACTGTCAAGAGGGAATTTTTCTCAGTATTTAGATATTTAGATTCAAAATAAGAAAGGGATCAATTCAATTATAAACCTGCAAAACAAAGATTAGGATTGGGTTGGGTTGCGCTATATATATCAAAGAGTATACAATAATGATGTATTTGGTGAATCAAATACATGGTCTAATAACAAACCATTTTAACATAACATTTTAATGAGTTGATAATATTAATTGAATATTGAATATTTTTTGAAAGATTTTTGTTAAAGGTTTCATTCATGCCTAATCCATATCGAGTAGRCCTTGTTGTTGTG??

			>Wittmackia_fawcettii
			??????????????????????????????????????????????????????????????????????????????????????????????????????????????????????????????????????????????????????????????????????????????????????????????????????????????????????????????????????????????????????????????????????????????????????????????????????????????????????????????????????????????????????????????????????????????????????????????????????????????????????????????????????????????????????????????????????????????????????????????????????????????????????????????????????????????????????????????????????????????????????????????????????????????????????????????????????????????????????????????????????????????????????????????????????????????????????????????????????????????????????????????????????????????????????????????

			>Wittmackia_froesii
			??????????????????????????????????????????????????????????????????????????????????????????????????????????????????????????????????????????????????????????????????????????????????????????????????????????????????????????????????????????????????????????????????????????????????????????????????????????????????????????????????????????????????????????????????????????????????????????????????????????????????????????????????????????????????????????????????????????????????????????????????????????????????????????????????????????????????????????????????????????????????????????????????????????????????????????????????????????????????????????????????????????????????????????????????????????????????????????????????????????????????????????????????????????????????????????????

			>Wittmackia_incompta
			??????????????????????????????????????????????????????????????????????????????????????????????????????????????????????????????????????????????????????????????????????????????????????????????????????????????????????????????????????????????????????????????????????????????????????????????????????????????????????????????????????????????????????????????????????????????????????????????????????????????????????????????????????????????????????????????????????????????????????????????????????????????????????????????????????????????????????????????????????????????????????????????????????????????????????????????????????????????????????????????????????????????????????????????????????????????????????????????????????????????????????????????????????????????????????????????

			>Wittmackia_inermis
			??????????????????????????????????????????????????????????????????????????????????????????????????????????????????????????????????????????????????????????????????????????????????????????????????????????????????????????????????????????????????????????????????????????????????????????????????????????????????????????????????????????????????????????????????????????????????????????????????????????????????????????????????????????????????????????????????????????????????????????????????????????????????????????????????????????????????????????????????????????????????????????????????????????????????????????????????????????????????????????????????????????????????????????????????????????????????????????????????????????????????????????????????????????????????????????????

			>Wittmackia_ituberaensis
			??????????????????????????????????????????????????????????????????????????????????????????????????????????????????????????????????????????????????????????????????????????????????????????????????????????????????????????????????????????????????????????????????????????????????????????????????????????????????????????????????????????????????????????????????????????????????????????????????????????????????????????????????????????????????????????????????????????????????????????????????????????????????????????????????????????????????????????????????????????????????????????????????????????????????????????????????????????????????????????????????????????????????????????????????????????????????????????????????????????????????????????????????????????????????????????????

			>Wittmackia_jamaicana
			??????????????????????????????????????????????????????????????????????????????????????????????????????????????????????????????????????????????????????????????????????????????????????????????????????????????????????????????????????????????????????????????????????????????????????????????????????????????????????????????????????????????????????????????????????????????????????????????????????????????????????????????????????????????????????????????????????????????????????????????????????????????????????????????????????????????????????????????????????????????????????????????????????????????????????????????????????????????????????????????????????????????????????????????????????????????????????????????????????????????????????????????????????????????????????????????

			>Wittmackia_laesslei
			??????????????????????????????????????????????????????????????????????????????????????????????????????????????????????????????????????????????????????????????????????????????????????????????????????????????????????????????????????????????????????????????????????????????????????????????????????????????????????????????????????????????????????????????????????????????????????????????????????????????????????????????????????????????????????????????????????????????????????????????????????????????????????????????????????????????????????????????????????????????????????????????????????????????????????????????????????????????????????????????????????????????????????????????????????????????????????????????????????????????????????????????????????????????????????????????

			>Wittmackia_laevigata
			??????????????????????????????????????????????????????????????????????????????????????????????????????????????????????????????????????????????????????????????????????????????????????????????????????????????????????????????????????????????????????????????????????????????????????????????????????????????????????????????????????????????????????????????????????????????????????????????????????????????????????????????????????????????????????????????????????????????????????????????????????????????????????????????????????????????????????????????????????????????????????????????????????????????????????????????????????????????????????????????????????????????????????????????????????????????????????????????????????????????????????????????????????????????????????????????

			>Wittmackia_limae
			??????????????????????????????????????????????????????????????????????????????????????????????????????????????????????????????????????????????????????????????????????????????????????????????????????????????????????????????????????????????????????????????????????????????????????????????????????????????????????????????????????????????????????????????????????????????????????????????????????????????????????????????????????????????????????????????????????????????????????????????????????????????????????????????????????????????????????????????????????????????????????????????????????????????????????????????????????????????????????????????????????????????????????????????????????????????????????????????????????????????????????????????????????????????????????????????

			>Wittmackia_lingulata
			ATAATTATAATAAAGTGAAATATGTCGAAATCCTTTTTGGAATAATACCAAATCGAAAATAAATGTCCGATAGCAAGTTGATCAGTTAATTCAATAAGAGATAAATGGGAGATAGCACTCGATTTAGTTGGTACCACCCAACCGAATCCGATTCACTCGTTTACTCATTCAATGAGTAAATTTTCAAGTTCAGCCAATC--TTTTTTTTGAAAAAAAAAATTGCAAGTAGATGAAATCGTGAATAAATGTGTTTCATTTCTATATCATTATATAAAAAAAGATCCATATTATATTACTTATGGAATTCGAACCCGAACTCGATTTATGATTTATTATTTGGATCTTATTGGCCTTTGTTCTTTATTTTTGATTTTTCATATGGATTTCCGTCTTTATATTATACCCTTTCGTGGATGAATTATGCCTATTTTCACATCTAGGATTTACATATACAACATATATTACTGTCAAGAGGGAATTTTTCTCAGTATTTAGATATTTAGATTCAAAATAAGAAAGGGATCAATTCAATTATAAACCTGCAAAACAAAGATTAGGATTGGGTTGGGTTGCGCTATATATATCAAAGAGTATACAATAATGATGTATTTGGTGAATCAAATACATGGTCTAATAACAAACCATTTTAACATAACATTTTAATGAGTTGATAATATTAATTGAATATTGAATATTTTTTGAAAGATTTTTGTTAAAGGTTTCATTCATGCCTAATCCATATCGAGTAGACCTTGTTGTTGTGAG

			>Wittmackia_lingulatoides
			??????????????????????????????????????????????????????????????????????????????????????????????????????????????????????????????????????????????????????????????????????????????????????????????????????????????????????????????????????????????????????????????????????????????????????????????????????????????????????????????????????????????????????????????????????????????????????????????????????????????????????????????????????????????????????????????????????????????????????????????????????????????????????????????????????????????????????????????????????????????????????????????????????????????????????????????????????????????????????????????????????????????????????????????????????????????????????????????????????????????????????????????????????????????????????????????

			>Wittmackia_linharesiorum
			??????????????????????????????????????????????????????????????????????????????????????????????????????????????????????????????????????????????????????????????????????????????????????????????????????????????????????????????????????????????????????????????????????????????????????????????????????????????????????????????????????????????????????????????????????????????????????????????????????????????????????????????????????????????????????????????????????????????????????????????????????????????????????????????????????????????????????????????????????????????????????????????????????????????????????????????????????????????????????????????????????????????????????????????????????????????????????????????????????????????????????????????????????????????????????????????

			>Wittmackia_maranguapensis
			??????????????????????????????????????????????????????????????????????????????????????????????????????????????????????????????????????????????????????????????????????????????????????????????????????????????????????????????????????????????????????????????????????????????????????????????????????????????????????????????????????????????????????????????????????????????????????????????????????????????????????????????????????????????????????????????????????????????????????????????????????????????????????????????????????????????????????????????????????????????????????????????????????????????????????????????????????????????????????????????????????????????????????????????????????????????????????????????????????????????????????????????????????????????????????????????

			>Wittmackia_mesoamericana
			??????????????????????????????????????????????????????????????????????????????????????????????????????????????????????????????????????????????????????????????????????????????????????????????????????????????????????????????????????????????????????????????????????????????????????????????????????????????????????????????????????????????????????????????????????????????????????????????????????????????????????????????????????????????????????????????????????????????????????????????????????????????????????????????????????????????????????????????????????????????????????????????????????????????????????????????????????????????????????????????????????????????????????????????????????????????????????????????????????????????????????????????????????????????????????????????

			>Wittmackia_negrilensis
			??????????????????????????????????????????????????????????????????????????????????????????????????????????????????????????????????????????????????????????????????????????????????????????????????????????????????????????????????????????????????????????????????????????????????????????????????????????????????????????????????????????????????????????????????????????????????????????????????????????????????????????????????????????????????????????????????????????????????????????????????????????????????????????????????????????????????????????????????????????????????????????????????????????????????????????????????????????????????????????????????????????????????????????????????????????????????????????????????????????????????????????????????????????????????????????????

			>Wittmackia_neoregelioides
			??????????????????????????????????????????????????????????????????????????????????????????????????????????????????????????????????????????????????????????????????????????????????????????????????????????????????????????????????????????????????????????????????????????????????????????????????????????????????????????????????????????????????????????????????????????????????????????????????????????????????????????????????????????????????????????????????????????????????????????????????????????????????????????????????????????????????????????????????????????????????????????????????????????????????????????????????????????????????????????????????????????????????????????????????????????????????????????????????????????????????????????????????????????????????????????????

			>Wittmackia_patentissima
			??????????????????????????????????????????????????????????????????????????????????????????????????????????????????????????????????????????????????????????????????????????????????????????????????????????????????????????????????????????????????????????????????????????????????????????????????????????????????????????????????????????????????????????????????????????????????????????????????????????????????????????????????????????????????????????????????????????????????????????????????????????????????????????????????????????????????????????????????????????????????????????????????????????????????????????????????????????????????????????????????????????????????????????????????????????????????????????????????????????????????????????????????????????????????????????????

			>Wittmackia_penduliflora
			???????????????????????????????????TTTGGAATAATACCAAATCGAAAATAAATGTCCGATAGCAAGTTGATCAGTTAATTCAATAAGAGATAAATGGGAGATAGCACTCGATTTAGTTGGTACCACCCAACCGAATCCGATTCACTCGTTTACTCATTCAATGAGTAAATTTTCAAGTTCAGCCAATC--TTTTTTTTTAAAAAAAAAATTGCAAGTAGATGAAATCGTGAATAAATGTGTTTCATTTCTATATCATTATATAAAAAAAGATCCATATTATATTACTTATGGAATTCGAACCCGAACTCGATTTATGATTTATTATTTGGATCTTATTGGCYTTTGTTCTTTATTTTTGATTTTTCATATAGATTTCCGTCTTTATATTATACCCTTTCGTGGATGAATTATGCCTATTTTCACATCTAGGATTTACATATACAACATATATTACTGTCAAGAGGGAATTTTTCTCAGTATTTAGATATTTAGATTCAAAATAAGAAAGGGATCAATTCAATTATAAACCTGCAAAACAAAGATTAGGATTGGGTTGGGTTGCGCTATATATATCAAAGAGTATACAATAATGATGTATTTGGTGAATCAAATACATGGTCTAATAACAAACCATTTTAACATAACATTTTAATGAGTTGATAATATTAATTGAATATTGAATATTTTTTGAAAGATTTTTGTTAAAGGTTTCATTCATGCCTAATCCATATCGAGTAGACCTTGTTGTTGTG??

			>Wittmackia_pernambucentris
			??????????????????????????????????????????????????????????????????????????????????????????????????????????????????????????????????????????????????????????????????????????????????????????????????????????????????????????????????????????????????????????????????????????????????????????????????????????????????????????????????????????????????????????????????????????????????????????????????????????????????????????????????????????????????????????????????????????????????????????????????????????????????????????????????????????????????????????????????????????????????????????????????????????????????????????????????????????????????????????????????????????????????????????????????????????????????????????????????????????????????????????????????????????????????????????????

			>Wittmackia_polycephala
			???????????????????????????????????TTTTGGATAATACCAAATCGAAAATAAATGTCCAATAGCAAGTTGATCAGTTAATTCAATAAGAGATAAATGGGAGATAGCACTCGATTTAGTTGGTACCACCCAACCGAATCCGATTCACTCGTTTACTCATTCAATGAGTAAATTTTCAAGTTCAGCCAATCTTTTTTTTTT-AAAAAAAAAATTGCAAGTAGATGAAATCGTGAATAAATGTGTTTCATTTCTATATCATTATATAAAAAAAGATCCATATTATATTACTTATGGAATTCGAACCCGAACTCGATTTATGATTTATTATTTGGATCTTATTGGCCTTTGTTCTTTATTTTTGATTTTTAATATAGATTTCCGTCTTTATATTATACCCTTTCGTGGATGAATTATGCCTATTTTCACATCTAGGATTTACATATACAACATATATTACTGTCAAGAGGGAATTTTTCTCAGTATTTAGATATTTAGATTCAAAATAAGAAAGGGATCAATTCAATTATAAACCTGCAAAACAAAGATTAGGATTGGGTTGGGTTGCGCTATATATATCAAAGAGTATACAATAATGATGTATTTGGTGAATCAAATACATGGTCTAATAACAAACCATTTTAACATAACATTTTAATGAGTTGATAATATTAATTGAATATTGAATATTTTTTGAAAGATTTTTGTTAAAGGTTTCATTCATGCCTAATCCATATCGAGTAGACCTTGTTGTTGTG??

			>Wittmackia_portoricensis
			??????????????????????????????????????????????????????????????????????????????????????????????????????????????????????????????????????????????????????????????????????????????????????????????????????????????????????????????????????????????????????????????????????????????????????????????????????????????????????????????????????????????????????????????????????????????????????????????????????????????????????????????????????????????????????????????????????????????????????????????????????????????????????????????????????????????????????????????????????????????????????????????????????????????????????????????????????????????????????????????????????????????????????????????????????????????????????????????????????????????????????????????????????????????????????????????

			>Wittmackia_silvana
			??????????????????????????????????????????????????????????????????????????????????????????????????????????????????????????????????????????????????????????????????????????????????????????????????????????????????????????????????????????????????????????????????????????????????????????????????????????????????????????????????????????????????????????????????????????????????????????????????????????????????????????????????????????????????????????????????????????????????????????????????????????????????????????????????????????????????????????????????????????????????????????????????????????????????????????????????????????????????????????????????????????????????????????????????????????????????????????????????????????????????????????????????????????????????????????????

			>Wittmackia_spinulosa
			??????????????????????????????????????????????????????????????????????????????????????????????????????????????????????????????????????????????????????????????????????????????????????????????????????????????????????????????????????????????????????????????????????????????????????????????????????????????????????????????????????????????????????????????????????????????????????????????????????????????????????????????????????????????????????????????????????????????????????????????????????????????????????????????????????????????????????????????????????????????????????????????????????????????????????????????????????????????????????????????????????????????????????????????????????????????????????????????????????????????????????????????????????????????????????????????

			>Wittmackia_sulbahianensis
			??????????????????????????????????????????????????????????????????????????????????????????????????????????????????????????????????????????????????????????????????????????????????????????????????????????????????????????????????????????????????????????????????????????????????????????????????????????????????????????????????????????????????????????????????????????????????????????????????????????????????????????????????????????????????????????????????????????????????????????????????????????????????????????????????????????????????????????????????????????????????????????????????????????????????????????????????????????????????????????????????????????????????????????????????????????????????????????????????????????????????????????????????????????????????????????????

			>Wittmackia_tentaculifera
			??????????????????????????????????????????????????????????????????????????????????????????????????????????????????????????????????????????????????????????????????????????????????????????????????????????????????????????????????????????????????????????????????????????????????????????????????????????????????????????????????????????????????????????????????????????????????????????????????????????????????????????????????????????????????????????????????????????????????????????????????????????????????????????????????????????????????????????????????????????????????????????????????????????????????????????????????????????????????????????????????????????????????????????????????????????????????????????????????????????????????????????????????????????????????????????????

			>Wittmackia_turbinocalyx
			???????????????????????????????????TTTTGGATAATACC?AATCGAAAATAAATGTCCAATAGCAAGTTGATCAGTTAATTCAATAAGAGATAAATGGGAGATAGCACTCGATTTAGTTGGTACCACCCAACCGAATCCGATTCACTCGTTTACTCATTCAATGAGTAAATTTTCAAGTTCAGCCAATCTTTTTTTTTT-AAAAAAAAAATTGCAAGTAGATGAAATCGTGAATAAATGTGTTTCATTTCTATATCATTATATAAAAAAAGATCCATATTATATTACTTATGGAATTCGAACCCGAACTCGATTTATGATTTATTATTTGGATCTTATTGGCCTTTGTTCTTTATTTTTGATTTTTAATATAGATTTCCGTCTTTATATTATACCCTTTCGTGGATGAATTATGCCTATTTTCACATCTAGGATTTACATATACAACATATATTACTGTCAAGAGGGAATTTTTCTCAGTATTTAGATATTTAGATTCAAAATAAGAAAGGGATCAATTCAATTATAAACCTGCAAAACAAAGATTAGGATTGGGTTGGGTTGCGCTATATATATCAAAGAGTATACAATAATGATGTATTTGGTGAATCAAATACATGGTCTAATAACAAACCATTTTAACATAACATTTTAATGAGTTGATAATATTAATTGAATATTGAATATTTTTTGAAAGATTTTTGTTAAAGGTTTCATTCATGCCTAATCCATATCGAGTAGACCTTGTTGTTGTG??

			>Wittmackia_urbaniana
			????????????????????????????????????????GATAATACCAAACGGAAAATAAATGTCCAATAGCAAGTTGATCAGTTAATTCAATAAGAGATAAATGGGAGATAGCACTCGATTTAGTTGGTACCACCCAACCGAATCCGATTCACTCGTTTACTCATTCAATGAGTAAATTTTCAAGTTCAGCCAATCTTTTTTTTTT-AAAAAAAAAATTGCAAGTAGATGAAATCGTGAATAAATGTGTTTCATTTCTATATCATTATATAAAAAAAGATCCATATTATATTACTTATGGAATTCGAACCCGAACTCGATTTATGATTTATTATTTGGATCTTATTGGCCTTTGTTCTTTATTTTTGATTTTTAATATAGATTTCCGTCTTTATATTATACCCTTTCGTGGATGAATTATGCCTATTTTCACATCTAGGATTTACATATACAACATATATTACTGTCAAGAGGGAATTTTTCTCAGTATTTAGATATTTAGATTCAAAATAAGAAAGGGATCAATTCAATTATAAACCTGCAAAACAAAGATTAGGATTGGGTTGGGTTGCGCTATATATATCAAAGAGTATACAATAATGATGTATTTGGTGAATCAAATACATGGTCTAATAACAAACCATTTTAACATAACATTTTAATGAGTTGATAATATTAATTGAATATTGAATATTTTTTGAAAGATTTTTGTTAAAGGTTTCATTCATGCCTAATCCATATCGAGTAGACCTTGTTGTTGTG??

			>Wittmackia_viridostigma
			??????????????????????????????????????????????????????????????????????????????????????????????????????????????????????????????????????????????????????????????????????????????????????????????????????????????????????????????????????????????????????????????????????????????????????????????????????????????????????????????????????????????????????????????????????????????????????????????????????????????????????????????????????????????????????????????????????????????????????????????????????????????????????????????????????????????????????????????????????????????????????????????????????????????????????????????????????????????????????????????????????????????????????????????????????????????????????????????????????????????????????????????????????????????????????????????

			>Wittrockia_abbreviata
			??????????????????????????????????????????????????????????????????????????????????????????????????????????????????????????????????????????????????????????????????????????????????????????????????????????????????????????????????????????????????????????????????????????????????????????????????????????????????????????????????????????????????????????????????????????????????????????????????????????????????????????????????????????????????????????????????????????????????????????????????????????????????????????????????????????????????????????????????????????????????????????????????????????????????????????????????????????????????????????????????????????????????????????????????????????????????????????????????????????????????????????????????????????????????????????????

			>Wittrockia_antillana
			??????????????????????????????????????????????????????????????????????????????????????????????????????????????????????????????????????????????????????????????????????????????????????????????????????????????????????????????????????????????????????????????????????????????????????????????????????????????????????????????????????????????????????????????????????????????????????????????????????????????????????????????????????????????????????????????????????????????????????????????????????????????????????????????????????????????????????????????????????????????????????????????????????????????????????????????????????????????????????????????????????????????????????????????????????????????????????????????????????????????????????????????????????????????????????????????

			>Wittrockia_cyathiformis
			???????????????????????GTCGA---TCCTTTTTGGATAATACCAAATCGAAAATAAATGTCCGATAGCAAGTTGATCAGTTAATTCAATAAGAGATAAATGGGAGATAGCACTCGATTTAGTTGGTACCACCCAACCGAATCCGATTCACTCGTTTACTCATTGAATGAGTAAATTTTCAAGTTCAGCCAWTYTTTTTTTTTTTCAAAAAAAAATTGCAAGTAGATGAAATCGCGAATAAATGTGTTTCATTTCTATATCATTATATAAAAAAAGATCCATATTATATTACTTATGGAATTCGAACCCGAACTCGATTTATGATTTATTATTTGGATCTTATTGGCCTTTGTTCTTTATTTTTGATTTTTCATATAGATTTCCGTCTTTATATTATACCCTTTCGTGGATGAATTATGCCTATTTTCACATCTAGGATTTACATATACAACATATATTACTGTCAAGAGGGAATTTTTCTCAGTATTTAGATATTTAGATTCAAAATAAGAAAGGGATCAATTCAATTATAAACCTGCAAAACAAAGATTAGGATTGGGTTGGGTTGCGCTATATATATCAAAGAGTATACAATAATGATGTATTTGGTGAATCAAATACATGGTCTAATAACAAACCATTTTAACATAACATTTTAATGAGTTGATAATATTAATTGAATATTGAATATTTTTTGAAAGATTTTTGTTAAAGGTTTCATTCATGCCTAATCCATATCGAGTAGACCTTGTTGTTGTG??

			>Wittrockia_gigantea
			??????????????????????????????????????????????????????????????????????????????????????????????????????????????????????????????????????????????????????????????????????????????????????????????????????????????????????????????????????????????????????????????????????????????????????????????????????????????????????????????????????????????????????????????????????????????????????????????????????????????????????????????????????????????????????????????????????????????????????????????????????????????????????????????????????????????????????????????????????????????????????????????????????????????????????????????????????????????????????????????????????????????????????????????????????????????????????????????????????????????????????????????????????????????????????????????

			>Wittrockia_paulistana
			??????????????????????????????????????????????????????????????????????????????????????????????????????????????????????????????????????????????????????????????????????????????????????????????????????????????????????????????????????????????????????????????????????????????????????????????????????????????????????????????????????????????????????????????????????????????????????????????????????????????????????????????????????????????????????????????????????????????????????????????????????????????????????????????????????????????????????????????????????????????????????????????????????????????????????????????????????????????????????????????????????????????????????????????????????????????????????????????????????????????????????????????????????????????????????????????

			>Wittrockia_spiralipetala
			??????????????????????????????????????????????????????????????????????????????????????????????????????????????????????????????????????????????????????????????????????????????????????????????????????????????????????????????????????????????????????????????????????????????????????????????????????????????????????????????????????????????????????????????????????????????????????????????????????????????????????????????????????????????????????????????????????????????????????????????????????????????????????????????????????????????????????????????????????????????????????????????????????????????????????????????????????????????????????????????????????????????????????????????????????????????????????????????????????????????????????????????????????????????????????????????

			>Wittrockia_superba
			ATAATTATAATAAAGTGAAATATGTCGAAATCCTTTTTGGAATAATACCAAATCGAAAATAAATGTCCGATAGCAAGTTGATCAGTTAATTCAATAAGAGATAAATGGGAGATAGCACTCGATTTAGTTGGTACCACCCAACCGAATCCGATTCACTCGTTTACTCATTGAATGAGTAAATTTTCAAGTTCAGCCAATCTTTTTTTTTTTAAAAAAAAAATTGCAAGTAGATGAAATCGTGAATAAATGTGTTTCATTTCTATATCATTATATAAAAAAAGATCCATATTATATTACTTATGGAATTCGAACCCGAACTCGATTTATGATTTATTATTTGGATCTTATTGGCCTTTGTTCTTTATTTTTGATTTTTCATATGGATTTACGTCTTTATATTATACCCTTTCGTGGATGAATTATGCCTATTTTCACATCTAGGATTTACATATACAACATATATTACCCTCAAGAGGGAATTTTTCTCAGTATTTAGATATTTAGATTCAAAATAAGAAAGGGATCAATTCAATTATAAACCTGCAAAACAAAGATTAGGA-----TTGGGTTGCGCTATATATATCAAAGAGTATACAATAATGATGTATTTGGTGAATCAAATACATGGTCTAATAACAAACCATTTTAACATAACATTTTAATGAGTTGATAATATTAATTGAATATTGAATATTTTTTGAAAGATTTTTGTTAAAGGTTTCATTCATGCCTAATCCATATCGAGTAGACCTTGTTGTTGTG??

			>Zizkaea_tuerckheimii
			??????????????????????????????????????????????????????????????????????????????????????????????????????????????????????????????????????????????????????????????????????????????????????????????????????????????????????????????????????????????????????????????????????????????????????????????????????????????????????????????????????????????????????????????????????????????????????????????????????????????????????????????????????????????????????????????????????????????????????????????????????????????????????????????????????????????????????????????????????????????????????????????????????????????????????????????????????????????????????????????????????????????????????????????????????????????????????????????????????????????????????????????????????????????????????????????

;leafy

			>Acanthostachys_pitcairnioides
			?????????????????????????????????????????????????????????????????????????????????????????????????????????????????????????????????????????????????????????????????????????????????????????????????????????????????????????????????????????????????????????????????????????????????????????????????????????????????????????????????????????????????????????????????????????????????????????????????????????????????????????????????????????????????????????????????????????????????????????????????????????????????????????????????????????????????????????????????????????????????????????????????????

			>Acanthostachys_strobilacea
			?????????????????????????????????????????????????????????????????????????????????????????????????????????????????????????????????????????????????????????????????????????????????????????????????????????????????????????????????????????????????????????????????????????????????????????????????????????????????????????????????????????????????????????????????????????????????????????????????????????????????????????????????????????????????????????????????????????????????????????????????????????????????????????????????????????????????????????????????????????????????????????????????????

			>Aechmea_abbreviata
			?????????????????????????????????????????????????????????????????????????????????????????????????????????????????????????????????????????????????????????????????????????????????????????????????????????????????????????????????????????????????????????????????????????????????????????????????????????????????????????????????????????????????????????????????????????????????????????????????????????????????????????????????????????????????????????????????????????????????????????????????????????????????????????????????????????????????????????????????????????????????????????????????????

			>Aechmea_aculeatosepala
			?????????????????????????????????????????????????????????????????????????????????????????????????????????????????????????????????????????????????????????????????????????????????????????????????????????????????????????????????????????????????????????????????????????????????????????????????????????????????????????????????????????????????????????????????????????????????????????????????????????????????????????????????????????????????????????????????????????????????????????????????????????????????????????????????????????????????????????????????????????????????????????????????????

			>Aechmea_aiuruocensis
			?????????????????????????????????????????????????????????????????????????????????????????????????????????????????????????????????????????????????????????????????????????????????????????????????????????????????????????????????????????????????????????????????????????????????????????????????????????????????????????????????????????????????????????????????????????????????????????????????????????????????????????????????????????????????????????????????????????????????????????????????????????????????????????????????????????????????????????????????????????????????????????????????????

			>Aechmea_alba
			?????????????????????????????????????????????????????????????????????????????????????????????????????????????????????????????????????????????????????????????????????????????????????????????????????????????????????????????????????????????????????????????????????????????????????????????????????????????????????????????????????????????????????????????????????????????????????????????????????????????????????????????????????????????????????????????????????????????????????????????????????????????????????????????????????????????????????????????????????????????????????????????????????

			>Aechmea_alopecurus
			?????????????????????????????????????????????????????????????????????????????????????????????????????????????????????????????????????????????????????????????????????????????????????????????????????????????????????????????????????????????????????????????????????????????????????????????????????????????????????????????????????????????????????????????????????????????????????????????????????????????????????????????????????????????????????????????????????????????????????????????????????????????????????????????????????????????????????????????????????????????????????????????????????

			>Aechmea_angustifolia
			?????????????????????????????????????????????????????????????????????????????????????????????????????????????????????????????????????????????????????????????????????????????????????????????????????????????????????????????????????????????????????????????????????????????????????????????????????????????????????????????????????????????????????????????????????????????????????????????????????????????????????????????????????????????????????????????????????????????????????????????????????????????????????????????????????????????????????????????????????????????????????????????????????

			>Aechmea_aquilega
			?????????????????????????????????????????????????????????????????????????????????????????????????????????????????????????????????????????????????????????????????????????????????????????????????????????????????????????????????????????????????????????????????????????????????????????????????????????????????????????????????????????????????????????????????????????????????????????????????????????????????????????????????????????????????????????????????????????????????????????????????????????????????????????????????????????????????????????????????????????????????????????????????????

			>Aechmea_arenaria
			?????????????????????????????????????????????????????????????????????????????????????????????????????????????????????????????????????????????????????????????????????????????????????????????????????????????????????????????????????????????????????????????????????????????????????????????????????????????????????????????????????????????????????????????????????????????????????????????????????????????????????????????????????????????????????????????????????????????????????????????????????????????????????????????????????????????????????????????????????????????????????????????????????

			>Aechmea_bambusoides
			?????????????????????????????????????????????????????????????????????????????????????????????????????????????????????????????????????????????????????????????????????????????????????????????????????????????????????????????????????????????????????????????????????????????????????????????????????????????????????????????????????????????????????????????????????????????????????????????????????????????????????????????????????????????????????????????????????????????????????????????????????????????????????????????????????????????????????????????????????????????????????????????????????

			>Aechmea_blanchetiana
			?????????????????????????????????????????????????????????????????????????????????????????????????????????????????????????????????????????????????????????????????????????????????????????????????????????????????????????????????????????????????????????????????????????????????????????????????????????????????????????????????????????????????????????????????????????????????????????????????????????????????????????????????????????????????????????????????????????????????????????????????????????????????????????????????????????????????????????????????????????????????????????????????????

			>Aechmea_blumenavii
			?????????????????????????????????????????????????????????????????????????????????????????????????????????????????????????????????????????????????????????????????????????????????????????????????????????????????????????????????????????????????????????????????????????????????????????????????????????????????????????????????????????????????????????????????????????????????????????????????????????????????????????????????????????????????????????????????????????????????????????????????????????????????????????????????????????????????????????????????????????????????????????????????????

			>Aechmea_bocainensis
			?????????????????????????????????????????????????????????????????????????????????????????????????????????????????????????????????????????????????????????????????????????????????????????????????????????????????????????????????????????????????????????????????????????????????????????????????????????????????????????????????????????????????????????????????????????????????????????????????????????????????????????????????????????????????????????????????????????????????????????????????????????????????????????????????????????????????????????????????????????????????????????????????????

			>Aechmea_bracteata
			?????????????????????????????????????????????????????????????????????????????????????????????????????????????????????????????????????????????????????????????????????????????????????????????????????????????????????????????????????????????????????????????????????????????????????????????????????????????????????????????????????????????????????????????????????????????????????????????????????????????????????????????????????????????????????????????????????????????????????????????????????????????????????????????????????????????????????????????????????????????????????????????????????

			>Aechmea_bracteata_var._pacifica
			?????????????????????????????????????????????????????????????????????????????????????????????????????????????????????????????????????????????????????????????????????????????????????????????????????????????????????????????????????????????????????????????????????????????????????????????????????????????????????????????????????????????????????????????????????????????????????????????????????????????????????????????????????????????????????????????????????????????????????????????????????????????????????????????????????????????????????????????????????????????????????????????????????

			>Aechmea_brassicoides
			?????????????????????????????????????????????????????????????????????????????????????????????????????????????????????????????????????????????????????????????????????????????????????????????????????????????????????????????????????????????????????????????????????????????????????????????????????????????????????????????????????????????????????????????????????????????????????????????????????????????????????????????????????????????????????????????????????????????????????????????????????????????????????????????????????????????????????????????????????????????????????????????????????

			>Aechmea_brevicollis
			?????????????????????????????????????????????????????????????????????????????????????????????????????????????????????????????????????????????????????????????????????????????????????????????????????????????????????????????????????????????????????????????????????????????????????????????????????????????????????????????????????????????????????????????????????????????????????????????????????????????????????????????????????????????????????????????????????????????????????????????????????????????????????????????????????????????????????????????????????????????????????????????????????

			>Aechmea_bromeliifolia
			?????????????????????????????????????????????????????????????????????????????????????????????????????????????????????????????????????????????????????????????????????????????????????????????????????????????????????????????????????????????????????????????????????????????????????????????????????????????????????????????????????????????????????????????????????????????????????????????????????????????????????????????????????????????????????????????????????????????????????????????????????????????????????????????????????????????????????????????????????????????????????????????????????

			>Aechmea_bromeliifolia_var._albobracteata
			?????????????????????????????????????????????????????????????????????????????????????????????????????????????????????????????????????????????????????????????????????????????????????????????????????????????????????????????????????????????????????????????????????????????????????????????????????????????????????????????????????????????????????????????????????????????????????????????????????????????????????????????????????????????????????????????????????????????????????????????????????????????????????????????????????????????????????????????????????????????????????????????????????

			>Aechmea_bruggeri
			?????????????????????????????????????????????????????????????????????????????????????????????????????????????????????????????????????????????????????????????????????????????????????????????????????????????????????????????????????????????????????????????????????????????????????????????????????????????????????????????????????????????????????????????????????????????????????????????????????????????????????????????????????????????????????????????????????????????????????????????????????????????????????????????????????????????????????????????????????????????????????????????????????

			>Aechmea_caesia
			?????????????????????????????????????????????????????????????????????????????????????????????????????????????????????????????????????????????????????????????????????????????????????????????????????????????????????????????????????????????????????????????????????????????????????????????????????????????????????????????????????????????????????????????????????????????????????????????????????????????????????????????????????????????????????????????????????????????????????????????????????????????????????????????????????????????????????????????????????????????????????????????????????

			>Aechmea_calyculata
			?????????????????????????????????????????????????????????????????????????????????????????????????????????????????????????????????????????????????????????????????????????????????????????????????????????????????????????????????????????????????????????????????????????????????????????????????????????????????????????????????????????????????????????????????????????????????????????????????????????????????????????????????????????????????????????????????????????????????????????????????????????????????????????????????????????????????????????????????????????????????????????????????????

			>Aechmea_candida
			?????????????????????????????????????????????????????????????????????????????????????????????????????????????????????????????????????????????????????????????????????????????????????????????????????????????????????????????????????????????????????????????????????????????????????????????????????????????????????????????????????????????????????????????????????????????????????????????????????????????????????????????????????????????????????????????????????????????????????????????????????????????????????????????????????????????????????????????????????????????????????????????????????

			>Aechmea_capixabae
			?????????????????????????????????????????????????????????????????????????????????????????????????????????????????????????????????????????????????????????????????????????????????????????????????????????????????????????????????????????????????????????????????????????????????????????????????????????????????????????????????????????????????????????????????????????????????????????????????????????????????????????????????????????????????????????????????????????????????????????????????????????????????????????????????????????????????????????????????????????????????????????????????????

			>Aechmea_carvalhoi
			?????????????????????????????????????????????????????????????????????????????????????????????????????????????????????????????????????????????????????????????????????????????????????????????????????????????????????????????????????????????????????????????????????????????????????????????????????????????????????????????????????????????????????????????????????????????????????????????????????????????????????????????????????????????????????????????????????????????????????????????????????????????????????????????????????????????????????????????????????????????????????????????????????

			>Aechmea_castanea
			?????????????????????????????????????????????????????????????????????????????????????????????????????????????????????????????????????????????????????????????????????????????????????????????????????????????????????????????????????????????????????????????????????????????????????????????????????????????????????????????????????????????????????????????????????????????????????????????????????????????????????????????????????????????????????????????????????????????????????????????????????????????????????????????????????????????????????????????????????????????????????????????????????

			>Aechmea_castelnavi
			?????????????????????????????????????????????????????????????????????????????????????????????????????????????????????????????????????????????????????????????????????????????????????????????????????????????????????????????????????????????????????????????????????????????????????????????????????????????????????????????????????????????????????????????????????????????????????????????????????????????????????????????????????????????????????????????????????????????????????????????????????????????????????????????????????????????????????????????????????????????????????????????????????

			>Aechmea_castelnavii
			?????????????????????????????????????????????????????????????????????????????????????????????????????????????????????????????????????????????????????????????????????????????????????????????????????????????????????????????????????????????????????????????????????????????????????????????????????????????????????????????????????????????????????????????????????????????????????????????????????????????????????????????????????????????????????????????????????????????????????????????????????????????????????????????????????????????????????????????????????????????????????????????????????

			>Aechmea_caudata
			?????????????????????????????????????????????????????????????????????????????????????????????????????????????????????????????????????????????????????????????????????????????????????????????????????????????????????????????????????????????????????????????????????????????????????????????????????????????????????????????????????????????????????????????????????????????????????????????????????????????????????????????????????????????????????????????????????????????????????????????????????????????????????????????????????????????????????????????????????????????????????????????????????

			>Aechmea_cephaloides
			?????????????????????????????????????????????????????????????????????????????????????????????????????????????????????????????????????????????????????????????????????????????????????????????????????????????????????????????????????????????????????????????????????????????????????????????????????????????????????????????????????????????????????????????????????????????????????????????????????????????????????????????????????????????????????????????????????????????????????????????????????????????????????????????????????????????????????????????????????????????????????????????????????

			>Aechmea_chantinii
			?????????????????????????????????????????????????????????????????????????????????????????????????????????????????????????????????????????????????????????????????????????????????????????????????????????????????????????????????????????????????????????????????????????????????????????????????????????????????????????????????????????????????????????????????????????????????????????????????????????????????????????????????????????????????????????????????????????????????????????????????????????????????????????????????????????????????????????????????????????????????????????????????????

			>Aechmea_chrysocoma
			?????????????????????????????????????????????????????????????????????????????????????????????????????????????????????????????????????????????????????????????????????????????????????????????????????????????????????????????????????????????????????????????????????????????????????????????????????????????????????????????????????????????????????????????????????????????????????????????????????????????????????????????????????????????????????????????????????????????????????????????????????????????????????????????????????????????????????????????????????????????????????????????????????

			>Aechmea_coelestis
			?????????????????????????????????????????????????????????????????????????????????????????????????????????????????????????????????????????????????????????????????????????????????????????????????????????????????????????????????????????????????????????????????????????????????????????????????????????????????????????????????????????????????????????????????????????????????????????????????????????????????????????????????????????????????????????????????????????????????????????????????????????????????????????????????????????????????????????????????????????????????????????????????????

			>Aechmea_colombiana
			?????????????????????????????????????????????????????????????????????????????????????????????????????????????????????????????????????????????????????????????????????????????????????????????????????????????????????????????????????????????????????????????????????????????????????????????????????????????????????????????????????????????????????????????????????????????????????????????????????????????????????????????????????????????????????????????????????????????????????????????????????????????????????????????????????????????????????????????????????????????????????????????????????

			>Aechmea_comata
			?????????????????????????????????????????????????????????????????????????????????????????????????????????????????????????????????????????????????????????????????????????????????????????????????????????????????????????????????????????????????????????????????????????????????????????????????????????????????????????????????????????????????????????????????????????????????????????????????????????????????????????????????????????????????????????????????????????????????????????????????????????????????????????????????????????????????????????????????????????????????????????????????????

			>Aechmea_contracta
			?????????????????????????????????????????????????????????????????????????????????????????????????????????????????????????????????????????????????????????????????????????????????????????????????????????????????????????????????????????????????????????????????????????????????????????????????????????????????????????????????????????????????????????????????????????????????????????????????????????????????????????????????????????????????????????????????????????????????????????????????????????????????????????????????????????????????????????????????????????????????????????????????????

			>Aechmea_correia-araujoi
			?????????????????????????????????????????????????????????????????????????????????????????????????????????????????????????????????????????????????????????????????????????????????????????????????????????????????????????????????????????????????????????????????????????????????????????????????????????????????????????????????????????????????????????????????????????????????????????????????????????????????????????????????????????????????????????????????????????????????????????????????????????????????????????????????????????????????????????????????????????????????????????????????????

			>Aechmea_costantinii
			?????????????????????????????????????????????????????????????????????????????????????????????????????????????????????????????????????????????????????????????????????????????????????????????????????????????????????????????????????????????????????????????????????????????????????????????????????????????????????????????????????????????????????????????????????????????????????????????????????????????????????????????????????????????????????????????????????????????????????????????????????????????????????????????????????????????????????????????????????????????????????????????????????

			>Aechmea_cucullata
			?????????????????????????????????????????????????????????????????????????????????????????????????????????????????????????????????????????????????????????????????????????????????????????????????????????????????????????????????????????????????????????????????????????????????????????????????????????????????????????????????????????????????????????????????????????????????????????????????????????????????????????????????????????????????????????????????????????????????????????????????????????????????????????????????????????????????????????????????????????????????????????????????????

			>Aechmea_cylindrata
			?????????????????????????????????????????????????????????????????????????????????????????????????????????????????????????????????????????????????????????????????????????????????????????????????????????????????????????????????????????????????????????????????????????????????????????????????????????????????????????????????????????????????????????????????????????????????????????????????????????????????????????????????????????????????????????????????????????????????????????????????????????????????????????????????????????????????????????????????????????????????????????????????????

			>Aechmea_dactylina
			?????????????????????????????????????????????????????????????????????????????????????????????????????????????????????????????????????????????????????????????????????????????????????????????????????????????????????????????????????????????????????????????????????????????????????????????????????????????????????????????????????????????????????????????????????????????????????????????????????????????????????????????????????????????????????????????????????????????????????????????????????????????????????????????????????????????????????????????????????????????????????????????????????

			>Aechmea_dealbata
			?????????????????????????????????????????????????????????????????????????????????????????????????????????????????????????????????????????????????????????????????????????????????????????????????????????????????????????????????????????????????????????????????????????????????????????????????????????????????????????????????????????????????????????????????????????????????????????????????????????????????????????????????????????????????????????????????????????????????????????????????????????????????????????????????????????????????????????????????????????????????????????????????????

			>Aechmea_digitata
			?????????????????????????????????????????????????????????????????????????????????????????????????????????????????????????????????????????????????????????????????????????????????????????????????????????????????????????????????????????????????????????????????????????????????????????????????????????????????????????????????????????????????????????????????????????????????????????????????????????????????????????????????????????????????????????????????????????????????????????????????????????????????????????????????????????????????????????????????????????????????????????????????????

			>Aechmea_discordiae
			?????????????????????????????????????????????????????????????????????????????????????????????????????????????????????????????????????????????????????????????????????????????????????????????????????????????????????????????????????????????????????????????????????????????????????????????????????????????????????????????????????????????????????????????????????????????????????????????????????????????????????????????????????????????????????????????????????????????????????????????????????????????????????????????????????????????????????????????????????????????????????????????????????

			>Aechmea_disjuncta
			?????????????????????????????????????????????????????????????????????????????????????????????????????????????????????????????????????????????????????????????????????????????????????????????????????????????????????????????????????????????????????????????????????????????????????????????????????????????????????????????????????????????????????????????????????????????????????????????????????????????????????????????????????????????????????????????????????????????????????????????????????????????????????????????????????????????????????????????????????????????????????????????????????

			>Aechmea_distichantha
			?????????????????????????????????????????????????????????????????????????????????????????????????????????????????????????????????????????????????????????????????????????????????????????????????????????????????????????????????????????????????????????????????????????????????????????????????????????????????????????????????????????????????????????????????????????????????????????????????????????????????????????????????????????????????????????????????????????????????????????????????????????????????????????????????????????????????????????????????????????????????????????????????????

			>Aechmea_distichantha_var._glaziovii
			?????????????????????????????????????????????????????????????????????????????????????????????????????????????????????????????????????????????????????????????????????????????????????????????????????????????????????????????????????????????????????????????????????????????????????????????????????????????????????????????????????????????????????????????????????????????????????????????????????????????????????????????????????????????????????????????????????????????????????????????????????????????????????????????????????????????????????????????????????????????????????????????????????

			>Aechmea_egleriana
			?????????????????????????????????????????????????????????????????????????????????????????????????????????????????????????????????????????????????????????????????????????????????????????????????????????????????????????????????????????????????????????????????????????????????????????????????????????????????????????????????????????????????????????????????????????????????????????????????????????????????????????????????????????????????????????????????????????????????????????????????????????????????????????????????????????????????????????????????????????????????????????????????????

			>Aechmea_eurycorymbus
			?????????????????????????????????????????????????????????????????????????????????????????????????????????????????????????????????????????????????????????????????????????????????????????????????????????????????????????????????????????????????????????????????????????????????????????????????????????????????????????????????????????????????????????????????????????????????????????????????????????????????????????????????????????????????????????????????????????????????????????????????????????????????????????????????????????????????????????????????????????????????????????????????????

			>Aechmea_farinosa
			?????????????????????????????????????????????????????????????????????????????????????????????????????????????????????????????????????????????????????????????????????????????????????????????????????????????????????????????????????????????????????????????????????????????????????????????????????????????????????????????????????????????????????????????????????????????????????????????????????????????????????????????????????????????????????????????????????????????????????????????????????????????????????????????????????????????????????????????????????????????????????????????????????

			>Aechmea_fasciata
			?????????????????????????????????????????????????????????????????????????????????????????????????????????????????????????????????????????????????????????????????????????????????????????????????????????????????????????????????????????????????????????????????????????????????????????????????????????????????????????????????????????????????????????????????????????????????????????????????????????????????????????????????????????????????????????????????????????????????????????????????????????????????????????????????????????????????????????????????????????????????????????????????????

			>Aechmea_filicaulis
			?????????????????????????????????????????????????????????????????????????????????????????????????????????????????????????????????????????????????????????????????????????????????????????????????????????????????????????????????????????????????????????????????????????????????????????????????????????????????????????????????????????????????????????????????????????????????????????????????????????????????????????????????????????????????????????????????????????????????????????????????????????????????????????????????????????????????????????????????????????????????????????????????????

			>Aechmea_flavorosea
			?????????????????????????????????????????????????????????????????????????????????????????????????????????????????????????????????????????????????????????????????????????????????????????????????????????????????????????????????????????????????????????????????????????????????????????????????????????????????????????????????????????????????????????????????????????????????????????????????????????????????????????????????????????????????????????????????????????????????????????????????????????????????????????????????????????????????????????????????????????????????????????????????????

			>Aechmea_floribunda
			?????????????????????????????????????????????????????????????????????????????????????????????????????????????????????????????????????????????????????????????????????????????????????????????????????????????????????????????????????????????????????????????????????????????????????????????????????????????????????????????????????????????????????????????????????????????????????????????????????????????????????????????????????????????????????????????????????????????????????????????????????????????????????????????????????????????????????????????????????????????????????????????????????

			>Aechmea_fosteriana
			?????????????????????????????????????????????????????????????????????????????????????????????????????????????????????????????????????????????????????????????????????????????????????????????????????????????????????????????????????????????????????????????????????????????????????????????????????????????????????????????????????????????????????????????????????????????????????????????????????????????????????????????????????????????????????????????????????????????????????????????????????????????????????????????????????????????????????????????????????????????????????????????????????

			>Aechmea_fulgens
			?????????????????????????????????????????????????????????????????????????????????????????????????????????????????????????????????????????????????????????????????????????????????????????????????????????????????????????????????????????????????????????????????????????????????????????????????????????????????????????????????????????????????????????????????????????????????????????????????????????????????????????????????????????????????????????????????????????????????????????????????????????????????????????????????????????????????????????????????????????????????????????????????????

			>Aechmea_fulgens_var._discolor
			?????????????????????????????????????????????????????????????????????????????????????????????????????????????????????????????????????????????????????????????????????????????????????????????????????????????????????????????????????????????????????????????????????????????????????????????????????????????????????????????????????????????????????????????????????????????????????????????????????????????????????????????????????????????????????????????????????????????????????????????????????????????????????????????????????????????????????????????????????????????????????????????????????

			>Aechmea_gamosepala
			?????????????????????????????????????????????????????????????????????????????????????????????????????????????????????????????????????????????????????????????????????????????????????????????????????????????????????????????????????????????????????????????????????????????????????????????????????????????????????????????????????????????????????????????????????????????????????????????????????????????????????????????????????????????????????????????????????????????????????????????????????????????????????????????????????????????????????????????????????????????????????????????????????

			>Aechmea_gracilis
			?????????????????????????????????????????????????????????????????????????????????????????????????????????????????????????????????????????????????????????????????????????????????????????????????????????????????????????????????????????????????????????????????????????????????????????????????????????????????????????????????????????????????????????????????????????????????????????????????????????????????????????????????????????????????????????????????????????????????????????????????????????????????????????????????????????????????????????????????????????????????????????????????????

			>Aechmea_guainumbiorum
			?????????????????????????????????????????????????????????????????????????????????????????????????????????????????????????????????????????????????????????????????????????????????????????????????????????????????????????????????????????????????????????????????????????????????????????????????????????????????????????????????????????????????????????????????????????????????????????????????????????????????????????????????????????????????????????????????????????????????????????????????????????????????????????????????????????????????????????????????????????????????????????????????????

			>Aechmea_haltonii
			?????????????????????????????????????????????????????????????????????????????????????????????????????????????????????????????????????????????????????????????????????????????????????????????????????????????????????????????????????????????????????????????????????????????????????????????????????????????????????????????????????????????????????????????????????????????????????????????????????????????????????????????????????????????????????????????????????????????????????????????????????????????????????????????????????????????????????????????????????????????????????????????????????

			>Aechmea_heterosepala
			?????????????????????????????????????????????????????????????????????????????????????????????????????????????????????????????????????????????????????????????????????????????????????????????????????????????????????????????????????????????????????????????????????????????????????????????????????????????????????????????????????????????????????????????????????????????????????????????????????????????????????????????????????????????????????????????????????????????????????????????????????????????????????????????????????????????????????????????????????????????????????????????????????

			>Aechmea_hoppii
			?????????????????????????????????????????????????????????????????????????????????????????????????????????????????????????????????????????????????????????????????????????????????????????????????????????????????????????????????????????????????????????????????????????????????????????????????????????????????????????????????????????????????????????????????????????????????????????????????????????????????????????????????????????????????????????????????????????????????????????????????????????????????????????????????????????????????????????????????????????????????????????????????????

			>Aechmea_jungurudoensis
			?????????????????????????????????????????????????????????????????????????????????????????????????????????????????????????????????????????????????????????????????????????????????????????????????????????????????????????????????????????????????????????????????????????????????????????????????????????????????????????????????????????????????????????????????????????????????????????????????????????????????????????????????????????????????????????????????????????????????????????????????????????????????????????????????????????????????????????????????????????????????????????????????????

			>Aechmea_kertesziae
			?????????????????????????????????????????????????????????????????????????????????????????????????????????????????????????????????????????????????????????????????????????????????????????????????????????????????????????????????????????????????????????????????????????????????????????????????????????????????????????????????????????????????????????????????????????????????????????????????????????????????????????????????????????????????????????????????????????????????????????????????????????????????????????????????????????????????????????????????????????????????????????????????????

			>Aechmea_kleinii
			?????????????????????????????????????????????????????????????????????????????????????????????????????????????????????????????????????????????????????????????????????????????????????????????????????????????????????????????????????????????????????????????????????????????????????????????????????????????????????????????????????????????????????????????????????????????????????????????????????????????????????????????????????????????????????????????????????????????????????????????????????????????????????????????????????????????????????????????????????????????????????????????????????

			>Aechmea_lamarchei
			?????????????????????????????????????????????????????????????????????????????????????????????????????????????????????????????????????????????????????????????????????????????????????????????????????????????????????????????????????????????????????????????????????????????????????????????????????????????????????????????????????????????????????????????????????????????????????????????????????????????????????????????????????????????????????????????????????????????????????????????????????????????????????????????????????????????????????????????????????????????????????????????????????

			>Aechmea_leptantha
			?????????????????????????????????????????????????????????????????????????????????????????????????????????????????????????????????????????????????????????????????????????????????????????????????????????????????????????????????????????????????????????????????????????????????????????????????????????????????????????????????????????????????????????????????????????????????????????????????????????????????????????????????????????????????????????????????????????????????????????????????????????????????????????????????????????????????????????????????????????????????????????????????????

			>Aechmea_leucolepis
			?????????????????????????????????????????????????????????????????????????????????????????????????????????????????????????????????????????????????????????????????????????????????????????????????????????????????????????????????????????????????????????????????????????????????????????????????????????????????????????????????????????????????????????????????????????????????????????????????????????????????????????????????????????????????????????????????????????????????????????????????????????????????????????????????????????????????????????????????????????????????????????????????????

			>Aechmea_lilacinantha
			?????????????????????????????????????????????????????????????????????????????????????????????????????????????????????????????????????????????????????????????????????????????????????????????????????????????????????????????????????????????????????????????????????????????????????????????????????????????????????????????????????????????????????????????????????????????????????????????????????????????????????????????????????????????????????????????????????????????????????????????????????????????????????????????????????????????????????????????????????????????????????????????????????

			>Aechmea_longicuspis_var._kuntzeana
			?????????????????????????????????????????????????????????????????????????????????????????????????????????????????????????????????????????????????????????????????????????????????????????????????????????????????????????????????????????????????????????????????????????????????????????????????????????????????????????????????????????????????????????????????????????????????????????????????????????????????????????????????????????????????????????????????????????????????????????????????????????????????????????????????????????????????????????????????????????????????????????????????????

			>Aechmea_lueddemanniana
			?????????????????????????????????????????????????????????????????????????????????????????????????????????????????????????????????????????????????????????????????????????????????????????????????????????????????????????????????????????????????????????????????????????????????????????????????????????????????????????????????????????????????????????????????????????????????????????????????????????????????????????????????????????????????????????????????????????????????????????????????????????????????????????????????????????????????????????????????????????????????????????????????????

			>Aechmea_maasii
			?????????????????????????????????????????????????????????????????????????????????????????????????????????????????????????????????????????????????????????????????????????????????????????????????????????????????????????????????????????????????????????????????????????????????????????????????????????????????????????????????????????????????????????????????????????????????????????????????????????????????????????????????????????????????????????????????????????????????????????????????????????????????????????????????????????????????????????????????????????????????????????????????????

			>Aechmea_magdalenae
			?????????????????????????????????????????????????????????????????????????????????????????????????????????????????????????????????????????????????????????????????????????????????????????????????????????????????????????????????????????????????????????????????????????????????????????????????????????????????????????????????????????????????????????????????????????????????????????????????????????????????????????????????????????????????????????????????????????????????????????????????????????????????????????????????????????????????????????????????????????????????????????????????????

			>Aechmea_manzanaresiana
			?????????????????????????????????????????????????????????????????????????????????????????????????????????????????????????????????????????????????????????????????????????????????????????????????????????????????????????????????????????????????????????????????????????????????????????????????????????????????????????????????????????????????????????????????????????????????????????????????????????????????????????????????????????????????????????????????????????????????????????????????????????????????????????????????????????????????????????????????????????????????????????????????????

			>Aechmea_marauensis
			?????????????????????????????????????????????????????????????????????????????????????????????????????????????????????????????????????????????????????????????????????????????????????????????????????????????????????????????????????????????????????????????????????????????????????????????????????????????????????????????????????????????????????????????????????????????????????????????????????????????????????????????????????????????????????????????????????????????????????????????????????????????????????????????????????????????????????????????????????????????????????????????????????

			>Aechmea_marginalis
			?????????????????????????????????????????????????????????????????????????????????????????????????????????????????????????????????????????????????????????????????????????????????????????????????????????????????????????????????????????????????????????????????????????????????????????????????????????????????????????????????????????????????????????????????????????????????????????????????????????????????????????????????????????????????????????????????????????????????????????????????????????????????????????????????????????????????????????????????????????????????????????????????????

			>Aechmea_mariae-reginae
			?????????????????????????????????????????????????????????????????????????????????????????????????????????????????????????????????????????????????????????????????????????????????????????????????????????????????????????????????????????????????????????????????????????????????????????????????????????????????????????????????????????????????????????????????????????????????????????????????????????????????????????????????????????????????????????????????????????????????????????????????????????????????????????????????????????????????????????????????????????????????????????????????????

			>Aechmea_melinonii
			?????????????????????????????????????????????????????????????????????????????????????????????????????????????????????????????????????????????????????????????????????????????????????????????????????????????????????????????????????????????????????????????????????????????????????????????????????????????????????????????????????????????????????????????????????????????????????????????????????????????????????????????????????????????????????????????????????????????????????????????????????????????????????????????????????????????????????????????????????????????????????????????????????

			>Aechmea_mertensii
			?????????????????????????????????????????????????????????????????????????????????????????????????????????????????????????????????????????????????????????????????????????????????????????????????????????????????????????????????????????????????????????????????????????????????????????????????????????????????????????????????????????????????????????????????????????????????????????????????????????????????????????????????????????????????????????????????????????????????????????????????????????????????????????????????????????????????????????????????????????????????????????????????????

			>Aechmea_mexicana
			?????????????????????????????????????????????????????????????????????????????????????????????????????????????????????????????????????????????????????????????????????????????????????????????????????????????????????????????????????????????????????????????????????????????????????????????????????????????????????????????????????????????????????????????????????????????????????????????????????????????????????????????????????????????????????????????????????????????????????????????????????????????????????????????????????????????????????????????????????????????????????????????????????

			>Aechmea_microcephala
			?????????????????????????????????????????????????????????????????????????????????????????????????????????????????????????????????????????????????????????????????????????????????????????????????????????????????????????????????????????????????????????????????????????????????????????????????????????????????????????????????????????????????????????????????????????????????????????????????????????????????????????????????????????????????????????????????????????????????????????????????????????????????????????????????????????????????????????????????????????????????????????????????????

			>Aechmea_miniata
			?????????????????????????????????????????????????????????????????????????????????????????????????????????????????????????????????????????????????????????????????????????????????????????????????????????????????????????????????????????????????????????????????????????????????????????????????????????????????????????????????????????????????????????????????????????????????????????????????????????????????????????????????????????????????????????????????????????????????????????????????????????????????????????????????????????????????????????????????????????????????????????????????????

			>Aechmea_miniata_var._discolor
			?????????????????????????????????????????????????????????????????????????????????????????????????????????????????????????????????????????????????????????????????????????????????????????????????????????????????????????????????????????????????????????????????????????????????????????????????????????????????????????????????????????????????????????????????????????????????????????????????????????????????????????????????????????????????????????????????????????????????????????????????????????????????????????????????????????????????????????????????????????????????????????????????????

			>Aechmea_mira
			?????????????????????????????????????????????????????????????????????????????????????????????????????????????????????????????????????????????????????????????????????????????????????????????????????????????????????????????????????????????????????????????????????????????????????????????????????????????????????????????????????????????????????????????????????????????????????????????????????????????????????????????????????????????????????????????????????????????????????????????????????????????????????????????????????????????????????????????????????????????????????????????????????

			>Aechmea_moorei
			?????????????????????????????????????????????????????????????????????????????????????????????????????????????????????????????????????????????????????????????????????????????????????????????????????????????????????????????????????????????????????????????????????????????????????????????????????????????????????????????????????????????????????????????????????????????????????????????????????????????????????????????????????????????????????????????????????????????????????????????????????????????????????????????????????????????????????????????????????????????????????????????????????

			>Aechmea_murcae
			?????????????????????????????????????????????????????????????????????????????????????????????????????????????????????????????????????????????????????????????????????????????????????????????????????????????????????????????????????????????????????????????????????????????????????????????????????????????????????????????????????????????????????????????????????????????????????????????????????????????????????????????????????????????????????????????????????????????????????????????????????????????????????????????????????????????????????????????????????????????????????????????????????

			>Aechmea_muricata
			?????????????????????????????????????????????????????????????????????????????????????????????????????????????????????????????????????????????????????????????????????????????????????????????????????????????????????????????????????????????????????????????????????????????????????????????????????????????????????????????????????????????????????????????????????????????????????????????????????????????????????????????????????????????????????????????????????????????????????????????????????????????????????????????????????????????????????????????????????????????????????????????????????

			>Aechmea_mutica
			?????????????????????????????????????????????????????????????????????????????????????????????????????????????????????????????????????????????????????????????????????????????????????????????????????????????????????????????????????????????????????????????????????????????????????????????????????????????????????????????????????????????????????????????????????????????????????????????????????????????????????????????????????????????????????????????????????????????????????????????????????????????????????????????????????????????????????????????????????????????????????????????????????

			>Aechmea_nallyi
			?????????????????????????????????????????????????????????????????????????????????????????????????????????????????????????????????????????????????????????????????????????????????????????????????????????????????????????????????????????????????????????????????????????????????????????????????????????????????????????????????????????????????????????????????????????????????????????????????????????????????????????????????????????????????????????????????????????????????????????????????????????????????????????????????????????????????????????????????????????????????????????????????????

			>Aechmea_napoensis
			?????????????????????????????????????????????????????????????????????????????????????????????????????????????????????????????????????????????????????????????????????????????????????????????????????????????????????????????????????????????????????????????????????????????????????????????????????????????????????????????????????????????????????????????????????????????????????????????????????????????????????????????????????????????????????????????????????????????????????????????????????????????????????????????????????????????????????????????????????????????????????????????????????

			>Aechmea_nidularioides
			?????????????????????????????????????????????????????????????????????????????????????????????????????????????????????????????????????????????????????????????????????????????????????????????????????????????????????????????????????????????????????????????????????????????????????????????????????????????????????????????????????????????????????????????????????????????????????????????????????????????????????????????????????????????????????????????????????????????????????????????????????????????????????????????????????????????????????????????????????????????????????????????????????

			>Aechmea_nudicaulis
			?????????????????????????????????????????????????????????????????????????????????????????????????????????????????????????????????????????????????????????????????????????????????????????????????????????????????????????????????????????????????????????????????????????????????????????????????????????????????????????????????????????????????????????????????????????????????????????????????????????????????????????????????????????????????????????????????????????????????????????????????????????????????????????????????????????????????????????????????????????????????????????????????????

			>Aechmea_organensis
			?????????????????????????????????????????????????????????????????????????????????????????????????????????????????????????????????????????????????????????????????????????????????????????????????????????????????????????????????????????????????????????????????????????????????????????????????????????????????????????????????????????????????????????????????????????????????????????????????????????????????????????????????????????????????????????????????????????????????????????????????????????????????????????????????????????????????????????????????????????????????????????????????????

			>Aechmea_orlandiana
			?????????????????????????????????????????????????????????????????????????????????????????????????????????????????????????????????????????????????????????????????????????????????????????????????????????????????????????????????????????????????????????????????????????????????????????????????????????????????????????????????????????????????????????????????????????????????????????????????????????????????????????????????????????????????????????????????????????????????????????????????????????????????????????????????????????????????????????????????????????????????????????????????????

			>Aechmea_ornata
			?????????????????????????????????????????????????????????????????????????????????????????????????????????????????????????????????????????????????????????????????????????????????????????????????????????????????????????????????????????????????????????????????????????????????????????????????????????????????????????????????????????????????????????????????????????????????????????????????????????????????????????????????????????????????????????????????????????????????????????????????????????????????????????????????????????????????????????????????????????????????????????????????????

			>Aechmea_paniculata
			?????????????????????????????????????????????????????????????????????????????????????????????????????????????????????????????????????????????????????????????????????????????????????????????????????????????????????????????????????????????????????????????????????????????????????????????????????????????????????????????????????????????????????????????????????????????????????????????????????????????????????????????????????????????????????????????????????????????????????????????????????????????????????????????????????????????????????????????????????????????????????????????????????

			>Aechmea_paratiensis
			?????????????????????????????????????????????????????????????????????????????????????????????????????????????????????????????????????????????????????????????????????????????????????????????????????????????????????????????????????????????????????????????????????????????????????????????????????????????????????????????????????????????????????????????????????????????????????????????????????????????????????????????????????????????????????????????????????????????????????????????????????????????????????????????????????????????????????????????????????????????????????????????????????

			>Aechmea_patriciae
			?????????????????????????????????????????????????????????????????????????????????????????????????????????????????????????????????????????????????????????????????????????????????????????????????????????????????????????????????????????????????????????????????????????????????????????????????????????????????????????????????????????????????????????????????????????????????????????????????????????????????????????????????????????????????????????????????????????????????????????????????????????????????????????????????????????????????????????????????????????????????????????????????????

			>Aechmea_pectinata
			?????????????????????????????????????????????????????????????????????????????????????????????????????????????????????????????????????????????????????????????????????????????????????????????????????????????????????????????????????????????????????????????????????????????????????????????????????????????????????????????????????????????????????????????????????????????????????????????????????????????????????????????????????????????????????????????????????????????????????????????????????????????????????????????????????????????????????????????????????????????????????????????????????

			>Aechmea_pedicellata
			?????????????????????????????????????????????????????????????????????????????????????????????????????????????????????????????????????????????????????????????????????????????????????????????????????????????????????????????????????????????????????????????????????????????????????????????????????????????????????????????????????????????????????????????????????????????????????????????????????????????????????????????????????????????????????????????????????????????????????????????????????????????????????????????????????????????????????????????????????????????????????????????????????

			>Aechmea_penduliflora
			?????????????????????????????????????????????????????????????????????????????????????????????????????????????????????????????????????????????????????????????????????????????????????????????????????????????????????????????????????????????????????????????????????????????????????????????????????????????????????????????????????????????????????????????????????????????????????????????????????????????????????????????????????????????????????????????????????????????????????????????????????????????????????????????????????????????????????????????????????????????????????????????????????

			>Aechmea_perforata
			?????????????????????????????????????????????????????????????????????????????????????????????????????????????????????????????????????????????????????????????????????????????????????????????????????????????????????????????????????????????????????????????????????????????????????????????????????????????????????????????????????????????????????????????????????????????????????????????????????????????????????????????????????????????????????????????????????????????????????????????????????????????????????????????????????????????????????????????????????????????????????????????????????

			>Aechmea_phanerophlebia
			?????????????????????????????????????????????????????????????????????????????????????????????????????????????????????????????????????????????????????????????????????????????????????????????????????????????????????????????????????????????????????????????????????????????????????????????????????????????????????????????????????????????????????????????????????????????????????????????????????????????????????????????????????????????????????????????????????????????????????????????????????????????????????????????????????????????????????????????????????????????????????????????????????

			>Aechmea_pimenti-velosoi
			?????????????????????????????????????????????????????????????????????????????????????????????????????????????????????????????????????????????????????????????????????????????????????????????????????????????????????????????????????????????????????????????????????????????????????????????????????????????????????????????????????????????????????????????????????????????????????????????????????????????????????????????????????????????????????????????????????????????????????????????????????????????????????????????????????????????????????????????????????????????????????????????????????

			>Aechmea_pineliana
			?????????????????????????????????????????????????????????????????????????????????????????????????????????????????????????????????????????????????????????????????????????????????????????????????????????????????????????????????????????????????????????????????????????????????????????????????????????????????????????????????????????????????????????????????????????????????????????????????????????????????????????????????????????????????????????????????????????????????????????????????????????????????????????????????????????????????????????????????????????????????????????????????????

			>Aechmea_polyantha
			?????????????????????????????????????????????????????????????????????????????????????????????????????????????????????????????????????????????????????????????????????????????????????????????????????????????????????????????????????????????????????????????????????????????????????????????????????????????????????????????????????????????????????????????????????????????????????????????????????????????????????????????????????????????????????????????????????????????????????????????????????????????????????????????????????????????????????????????????????????????????????????????????????

			>Aechmea_pubescens
			?????????????????????????????????????????????????????????????????????????????????????????????????????????????????????????????????????????????????????????????????????????????????????????????????????????????????????????????????????????????????????????????????????????????????????????????????????????????????????????????????????????????????????????????????????????????????????????????????????????????????????????????????????????????????????????????????????????????????????????????????????????????????????????????????????????????????????????????????????????????????????????????????????

			>Aechmea_racinae
			?????????????????????????????????????????????????????????????????????????????????????????????????????????????????????????????????????????????????????????????????????????????????????????????????????????????????????????????????????????????????????????????????????????????????????????????????????????????????????????????????????????????????????????????????????????????????????????????????????????????????????????????????????????????????????????????????????????????????????????????????????????????????????????????????????????????????????????????????????????????????????????????????????

			>Aechmea_ramosa
			?????????????????????????????????????????????????????????????????????????????????????????????????????????????????????????????????????????????????????????????????????????????????????????????????????????????????????????????????????????????????????????????????????????????????????????????????????????????????????????????????????????????????????????????????????????????????????????????????????????????????????????????????????????????????????????????????????????????????????????????????????????????????????????????????????????????????????????????????????????????????????????????????????

			>Aechmea_recurvata
			?????????????????????????????????????????????????????????????????????????????????????????????????????????????????????????????????????????????????????????????????????????????????????????????????????????????????????????????????????????????????????????????????????????????????????????????????????????????????????????????????????????????????????????????????????????????????????????????????????????????????????????????????????????????????????????????????????????????????????????????????????????????????????????????????????????????????????????????????????????????????????????????????????

			>Aechmea_roberto-seidelii
			?????????????????????????????????????????????????????????????????????????????????????????????????????????????????????????????????????????????????????????????????????????????????????????????????????????????????????????????????????????????????????????????????????????????????????????????????????????????????????????????????????????????????????????????????????????????????????????????????????????????????????????????????????????????????????????????????????????????????????????????????????????????????????????????????????????????????????????????????????????????????????????????????????

			>Aechmea_rodriguesiana
			?????????????????????????????????????????????????????????????????????????????????????????????????????????????????????????????????????????????????????????????????????????????????????????????????????????????????????????????????????????????????????????????????????????????????????????????????????????????????????????????????????????????????????????????????????????????????????????????????????????????????????????????????????????????????????????????????????????????????????????????????????????????????????????????????????????????????????????????????????????????????????????????????????

			>Aechmea_roeseliae
			?????????????????????????????????????????????????????????????????????????????????????????????????????????????????????????????????????????????????????????????????????????????????????????????????????????????????????????????????????????????????????????????????????????????????????????????????????????????????????????????????????????????????????????????????????????????????????????????????????????????????????????????????????????????????????????????????????????????????????????????????????????????????????????????????????????????????????????????????????????????????????????????????????

			>Aechmea_romeroi
			?????????????????????????????????????????????????????????????????????????????????????????????????????????????????????????????????????????????????????????????????????????????????????????????????????????????????????????????????????????????????????????????????????????????????????????????????????????????????????????????????????????????????????????????????????????????????????????????????????????????????????????????????????????????????????????????????????????????????????????????????????????????????????????????????????????????????????????????????????????????????????????????????????

			>Aechmea_rubiginosa
			?????????????????????????????????????????????????????????????????????????????????????????????????????????????????????????????????????????????????????????????????????????????????????????????????????????????????????????????????????????????????????????????????????????????????????????????????????????????????????????????????????????????????????????????????????????????????????????????????????????????????????????????????????????????????????????????????????????????????????????????????????????????????????????????????????????????????????????????????????????????????????????????????????

			>Aechmea_serragrandensis
			?????????????????????????????????????????????????????????????????????????????????????????????????????????????????????????????????????????????????????????????????????????????????????????????????????????????????????????????????????????????????????????????????????????????????????????????????????????????????????????????????????????????????????????????????????????????????????????????????????????????????????????????????????????????????????????????????????????????????????????????????????????????????????????????????????????????????????????????????????????????????????????????????????

			>Aechmea_servitensis
			?????????????????????????????????????????????????????????????????????????????????????????????????????????????????????????????????????????????????????????????????????????????????????????????????????????????????????????????????????????????????????????????????????????????????????????????????????????????????????????????????????????????????????????????????????????????????????????????????????????????????????????????????????????????????????????????????????????????????????????????????????????????????????????????????????????????????????????????????????????????????????????????????????

			>Aechmea_servitensis_var._exigua
			?????????????????????????????????????????????????????????????????????????????????????????????????????????????????????????????????????????????????????????????????????????????????????????????????????????????????????????????????????????????????????????????????????????????????????????????????????????????????????????????????????????????????????????????????????????????????????????????????????????????????????????????????????????????????????????????????????????????????????????????????????????????????????????????????????????????????????????????????????????????????????????????????????

			>Aechmea_sphaerocephala
			?????????????????????????????????????????????????????????????????????????????????????????????????????????????????????????????????????????????????????????????????????????????????????????????????????????????????????????????????????????????????????????????????????????????????????????????????????????????????????????????????????????????????????????????????????????????????????????????????????????????????????????????????????????????????????????????????????????????????????????????????????????????????????????????????????????????????????????????????????????????????????????????????????

			>Aechmea_squarrosa
			?????????????????????????????????????????????????????????????????????????????????????????????????????????????????????????????????????????????????????????????????????????????????????????????????????????????????????????????????????????????????????????????????????????????????????????????????????????????????????????????????????????????????????????????????????????????????????????????????????????????????????????????????????????????????????????????????????????????????????????????????????????????????????????????????????????????????????????????????????????????????????????????????????

			>Aechmea_strobilacea
			?????????????????????????????????????????????????????????????????????????????????????????????????????????????????????????????????????????????????????????????????????????????????????????????????????????????????????????????????????????????????????????????????????????????????????????????????????????????????????????????????????????????????????????????????????????????????????????????????????????????????????????????????????????????????????????????????????????????????????????????????????????????????????????????????????????????????????????????????????????????????????????????????????

			>Aechmea_tayoensis
			?????????????????????????????????????????????????????????????????????????????????????????????????????????????????????????????????????????????????????????????????????????????????????????????????????????????????????????????????????????????????????????????????????????????????????????????????????????????????????????????????????????????????????????????????????????????????????????????????????????????????????????????????????????????????????????????????????????????????????????????????????????????????????????????????????????????????????????????????????????????????????????????????????

			>Aechmea_tessmannii
			?????????????????????????????????????????????????????????????????????????????????????????????????????????????????????????????????????????????????????????????????????????????????????????????????????????????????????????????????????????????????????????????????????????????????????????????????????????????????????????????????????????????????????????????????????????????????????????????????????????????????????????????????????????????????????????????????????????????????????????????????????????????????????????????????????????????????????????????????????????????????????????????????????

			>Aechmea_tillandsioides
			?????????????????????????????????????????????????????????????????????????????????????????????????????????????????????????????????????????????????????????????????????????????????????????????????????????????????????????????????????????????????????????????????????????????????????????????????????????????????????????????????????????????????????????????????????????????????????????????????????????????????????????????????????????????????????????????????????????????????????????????????????????????????????????????????????????????????????????????????????????????????????????????????????

			>Aechmea_triangularis
			?????????????????????????????????????????????????????????????????????????????????????????????????????????????????????????????????????????????????????????????????????????????????????????????????????????????????????????????????????????????????????????????????????????????????????????????????????????????????????????????????????????????????????????????????????????????????????????????????????????????????????????????????????????????????????????????????????????????????????????????????????????????????????????????????????????????????????????????????????????????????????????????????????

			>Aechmea_triticina
			?????????????????????????????????????????????????????????????????????????????????????????????????????????????????????????????????????????????????????????????????????????????????????????????????????????????????????????????????????????????????????????????????????????????????????????????????????????????????????????????????????????????????????????????????????????????????????????????????????????????????????????????????????????????????????????????????????????????????????????????????????????????????????????????????????????????????????????????????????????????????????????????????????

			>Aechmea_vallerandii
			?????????????????????????????????????????????????????????????????????????????????????????????????????????????????????????????????????????????????????????????????????????????????????????????????????????????????????????????????????????????????????????????????????????????????????????????????????????????????????????????????????????????????????????????????????????????????????????????????????????????????????????????????????????????????????????????????????????????????????????????????????????????????????????????????????????????????????????????????????????????????????????????????????

			>Aechmea_vanhoutteana
			?????????????????????????????????????????????????????????????????????????????????????????????????????????????????????????????????????????????????????????????????????????????????????????????????????????????????????????????????????????????????????????????????????????????????????????????????????????????????????????????????????????????????????????????????????????????????????????????????????????????????????????????????????????????????????????????????????????????????????????????????????????????????????????????????????????????????????????????????????????????????????????????????????

			>Aechmea_victoriana
			?????????????????????????????????????????????????????????????????????????????????????????????????????????????????????????????????????????????????????????????????????????????????????????????????????????????????????????????????????????????????????????????????????????????????????????????????????????????????????????????????????????????????????????????????????????????????????????????????????????????????????????????????????????????????????????????????????????????????????????????????????????????????????????????????????????????????????????????????????????????????????????????????????

			>Aechmea_weilbachii
			?????????????????????????????????????????????????????????????????????????????????????????????????????????????????????????????????????????????????????????????????????????????????????????????????????????????????????????????????????????????????????????????????????????????????????????????????????????????????????????????????????????????????????????????????????????????????????????????????????????????????????????????????????????????????????????????????????????????????????????????????????????????????????????????????????????????????????????????????????????????????????????????????????

			>Aechmea_werdermannii
			?????????????????????????????????????????????????????????????????????????????????????????????????????????????????????????????????????????????????????????????????????????????????????????????????????????????????????????????????????????????????????????????????????????????????????????????????????????????????????????????????????????????????????????????????????????????????????????????????????????????????????????????????????????????????????????????????????????????????????????????????????????????????????????????????????????????????????????????????????????????????????????????????????

			>Aechmea_winkleri
			?????????????????????????????????????????????????????????????????????????????????????????????????????????????????????????????????????????????????????????????????????????????????????????????????????????????????????????????????????????????????????????????????????????????????????????????????????????????????????????????????????????????????????????????????????????????????????????????????????????????????????????????????????????????????????????????????????????????????????????????????????????????????????????????????????????????????????????????????????????????????????????????????????

			>Aechmea_woronowii
			?????????????????????????????????????????????????????????????????????????????????????????????????????????????????????????????????????????????????????????????????????????????????????????????????????????????????????????????????????????????????????????????????????????????????????????????????????????????????????????????????????????????????????????????????????????????????????????????????????????????????????????????????????????????????????????????????????????????????????????????????????????????????????????????????????????????????????????????????????????????????????????????????????

			>Aechmea_zebrina
			?????????????????????????????????????????????????????????????????????????????????????????????????????????????????????????????????????????????????????????????????????????????????????????????????????????????????????????????????????????????????????????????????????????????????????????????????????????????????????????????????????????????????????????????????????????????????????????????????????????????????????????????????????????????????????????????????????????????????????????????????????????????????????????????????????????????????????????????????????????????????????????????????????

			>Alcantarea_aurantiaca
			?????????????????????????????????????????????????????????????????????????????????????????????????????????????????????????????????????????????????????????????????????????????????????????????????????????????????????????????????????????????????????????????????????????????????????????????????????????????????????????????????????????????????????????????????????????????????????????????????????????????????????????????????????????????????????????????????????????????????????????????????????????????????????????????????????????????????????????????????????????????????????????????????????

			>Alcantarea_benzingii
			?????????????????????????????????????????????????????????????????????????????????????????????????????????????????????????????????????????????????????????????????????????????????????????????????????????????????????????????????????????????????????????????????????????????????????????????????????????????????????????????????????????????????????????????????????????????????????????????????????????????????????????????????????????????????????????????????????????????????????????????????????????????????????????????????????????????????????????????????????????????????????????????????????

			>Alcantarea_burle-marxii
			??????????????????????GCGTACGGAGTGGGGGGAAGGAGCGGCTGGAGAGCGATTGGACGGAAAGCTGTGGATAGCTGGGGAGGGAGAGGGAGCCGTGAGTTGTCTTGTCGGGACCTGACATTGCATTTTCTCGGAGACACCCCGCTGTTGGGCGGCCGCCACGTGTCCGCGGAAGTGACCCGGGCCCCACGCTCGCGCGCGGTGGGGGCGTAATGAGTGCACAGCGGAGCTCCGAGAGGGAGCACTGTTGCTGCATCAAGATTCGCTCTCTCGCAGCTGCGTCGGGAGGAGTGTACGTTGAGTTGCAGTGTAGTGCGCTACCTGTGAATCCACAGCGTTGACTGCGTGGCCGACATTTGTCACAATCCTCAGTTTCAATTAATTTAGTTGGTGCGCCATCCCAGGCACTGATATATTTTTGCATTTTAGGCAGTGGTGAGTAGGAGGCTATTCCAAGCTTAATAAAGTACCCTGAAATGAAATAGAAAACTTTTTGTTCGATATGATTAGAGCAGCTTAAGTTAGAGGATACCCATAACAATTAAGATGTGCATGCAAAT??????????????

			>Alcantarea_compacta
			?????????????????????????????????????????????????????????????????????????????????????????????????????????????????????????????????????????????????????????????????????????????????????????????????????????????????????????????????????????????????????????????????????????????????????????????????????????????????????????????????????????????????????????????????????????????????????????????????????????????????????????????????????????????????????????????????????????????????????????????????????????????????????????????????????????????????????????????????????????????????????????????????????

			>Alcantarea_duarteana
			AGAAAGAAAGAAAGAAAGAAAGGCGTACGGAGGGGAGGGAAGGAGCGGCTGGAGAGCGATTGGACGGAAAGCTGTGGATAGCTGGGGAGGGAGAGGCAGCCGTGAGTTGTCTTGTCGGGACCTGACATTGCATTTTCTCGGAGACACCCCGCTGTTGGGCGGCCGCCACGTGTCCGCGGAAGTGACCCGGGCCCCACGCTCGCGCGCGGTGGGGGCGTAATGAGTGCACAGCGGAGCTCCGAGAGGGAGCACTGTTGCTGCATCAAGATTCGCTCTCTCGCAGCTGCGTCGGGAGGAGTGTACGTTGAGTTGCAGTGTAGTGCGCTACCTGTGAATCCACAGCGTTGACTGCGTGGCCGACATTTGTCACAATCCTCAGTTTCAATTAATTTAGTTGGTGCGCCATCCCAGGCACTGATATATTTTTGCGTTTTAGGCAGTGGTGAGTAGGAGGCTATTCCAAGCTTAATAAAGTACCCTGAAATGAAATAGAAAACTTTTTGTTCGATATGATTAGAGCAGCTTAAGTTAGAGGATACCCATAACAATTAAGATGTGCATGCAAATCTAGTTGCCAAAA?

			>Alcantarea_farneyi
			?????????????????????????????????????????????????????????????????????????????????????????????????????????????????????????????????????????????????????????????????????????????????????????????????????????????????????????????????????????????????????????????????????????????????????????????????????????????????????????????????????????????????????????????????????????????????????????????????????????????????????????????????????????????????????????????????????????????????????????????????????????????????????????????????????????????????????????????????????????????????????????????????????

			>Alcantarea_geniculata
			?????????????????????????????????GGGGGGAAGGAGCGGCTGGAGAGCGATTGGACGGAAAGCTGTGGATAGCTGGGGAGGGAGAGGCAGCCGTGAGTTGTCTTGTCGGGACCTGACATTGCATTTTCTCGGAGACACCCCGCTGTTGGGCGGCCGCCACGTGTCCGCGGAAGTGACCCGGGCCCCACGCTCGCGCGCGGTGGGGGCGTAATGAGTGCACAGCGGAGCTCCGAGAGGGAGCACTGTTGCTGCATCAAGATTCGCTCTCTCGCAGCTGCGTCGGGAGGAGTGTACGTTGAGTTGCAGTGTAGTGCGCTACCTGTGAATCCACAGCGTTGACTGCGTGGCCGACATTTGTCACAATCCTCAGCTTCAATTAATTTAGTAGGTGCGCCATCCCAGGCACTGATATATTTTTGCATTTTAGGCAGTGGTGAGTAGGAGGCTATTCCAAGCTTAATAAAGTACCCTGAAATGAAATAGAAAACTTTTTGTTCGATATGATTAGAGCAGCTTAAGTTAGAGGATACCCATAACAATTAAGATGTGCATGCAAATCTAGTTGCCAAAA?

			>Alcantarea_glaziouana
			AGAAAGAAGGAATGAAAGAAAGGCGTACGGAAGGGGGGAAAGAAGCGGCTGGAGAGCGATTGGACGGAAAGCTGGGATTAGCTGGGGAGGGAGAGGAAGCCGTGAGTTGTCTGGTCGGGACTTGACATTGCATTTTCTCGGAGACACCCCG-TGTTGGCCGGCCGCCACGTGTCCGCGGAAGTGACCCGGGCCCCACGCTCGCGCGCGGT-GGGGCGTAATGAGTGCACAGCGGAGCTCCGAGAGGGAGCACTGTTGCTGCATCAAGATTCGCTCTCTCGCAGCTGCGTCGGGAGGAGTGTACGTTGAGTTGCAGTGTAGTGCGCTACCTGTGAATCCACAGCGTTGACTGCGTGGCCGACATTTGTCACAATCCTCAGTTTCAATTAATTTAGTTGGTGCGCCATCCCAGGCACTGATATATTTTTGCATTTTAGGCAGTGGTGAGTAGGAGGCTATTCCAAGCTTAATAAAGTACCCTGAAATGAAATAGAAAACTTTTTGTTCGATATGATTAGAGCAGCTTAAGTTAGAGGATACCCA???????????????????????????????????????

			>Alcantarea_heloisae
			?????????????????????????????????????????????????????????????????????????????????????????????????????????????????????????????????????????????????????????????????????????????????????????????????????????????????????????????????????????????????????????????????????????????????????????????????????????????????????????????????????????????????????????????????????????????????????????????????????????????????????????????????????????????????????????????????????????????????????????????????????????????????????????????????????????????????????????????????????????????????????????????????????

			>Alcantarea_imperialis
			AGAGAGAAAGAAAGAAAGAAAGGCGTACGGAGGGGGGGGAAGGATTGGCTGGAGAGCGATTGGACGGAAAGCTGTGGATAGCTGGGGACGGAGAGGCAGCCGTGAGTTGTCTTGTCGGGACCTGACATTGCATTTTCTCGGAGACACCCCGCTGTTGGGCGGCCGCCACGTGTCCGCGGAAGTGACCCGGGCCCCACGCTCGCGCGCGGTGGGGGCGTAATGAGTGCACAGCGGAGCTCCGAGAGGGAGCACTGTTGCTGCATCAAGATTCGCTCTCTCGCAGCTGCGTCGGGAGGAGTGTACGTTGAGTTGCAGTGTAGTGCGCTACCTGTGAATCCACAGCGTTGACTGCGTGGCCGACATTTGTCACAATCCTCAGCTTCAATTAATTTAGTTGGTGCGCCATCCCATGCACTGATACATTTTTGCATTTTAGGCAGTGGTGAGTAGGAGGCTATTCCAAGCTTAATAAAGTACCCTGAAATGAAATAGAAAACTTTTTGTTCGATATGATTAGAGCAGCTTAAGTTAGAGGATACCCATAACAATTAAGATGTGCATGCAAATCTAGTTGCCAAAAC

			>Alcantarea_martinellii
			AGAAAGAAAGAAAGAAAGAAAGGCGTACGGAGGGGGGGGAAGGAGCGGCTGGAGAGCGATTGGACGGAAAGCTGTGGATAGCTGGGGAGGGAGAGGCAGCCGTGAGTTGTCTTGTCGGGACCTGACATTGCATTTTCTCGGAGAGACCCCGCTGTTGGGCGGCCGCCACGTGTCCGCGGAAGTGACCCGGGCCCCACGCTCGCGCGCGGTGGGGGCGTAATGAGTGCACAGCGGAGCTCCGAGAGGGAGCACTGTTGCTGCATCAAGATTCGCTCTCTCGCAGCTGCGTCGGGAGGAGTGTACGTTGAGTTGCAGTGTAGTGCGCTACCTGTGAATCCACAGCGTTGACTGCGTGGCCGACATTTGTCACAATCCTCAGTTTCAATTAATTTAGTAGGTGCGCCATCCCAGGCACTGATATATTTTTGCATTTTAGGCAGTGGTGAGTAGGAGGCTATTCCAAGCTTAATAAAGTACCCTGAAATGAAATAGAAAAATTTTTGTTCGATATGATTAGAGCAGCTTAAGTTAGAGGATACCCATAACAATTAAGATGTGCATGCAAATCTAGTTGCCAAAAC

			>Alcantarea_nahoumii
			??????????????????AAAGGCGTACGGAGGGGGGGGAAGGAGCGGCTGGAGAGCGATTGGACGGAAAGCTGTGGATAGCTGGGGAGGGAGAGGGAGCCGTGAGTTGTCTTGTCGGGACCTGACATTGCATTTTCTCGGAGACACCCCGCTGTTGGGCGGCCGCCACGTGTCCGCGGAAGTGACCCGGGCCCCACGCTCGCGCGCGGTGGGGGCGTAATGAGTGCACAGCGGAGCTCCGAGAGGGAGCACTGTTGCTGCATCAAGATTCGCTCTCTCGCAGCTGCGTCGGGAGGAGTGTACGTTGAGTTGCAGTGTAGTGCGCTACCTGTGAATCCACAGCGTTGACTGCGTGGCCGACATTTGTCACAATCCTCAGTTTCAATTAATTTAGTTGGTGCGCCATCCCAGGCACTGATATATTTTTGCATTTTAGGCAGTGGTGAATAGGAGGCTATTCCAAGCTTAATAAAGTACCCTGAAATGAAATAGAAAACTTTTTGTTCGATATGATTAGAGCAGCTTAAGTTAGAGGATACCCATAACAATTAAGATGTGCATGCAAATCTAGTTGCCAAAAC

			>Alcantarea_nevaresii
			AGAAAGAAAGAAAGAAAGAAAGGCGTACGGAGGGGGGGGAAGGAGCGGCTGGAGAGCGATTGGACGGAAAGCTGTGGATAGCTGGGGAGGGAGAGGCAGCCGTGAGTTGTCTTGTCGGGACCTGACATTGCATTTTCTCGGAGACACCCCGCTGTTGGGCGGCCGCCACGTGTCCGCGGAAGTGACCCGGGCCCCACGCTCGCGCGCGGTGGGGGCGTAATGAGTGCACAGCGGAGCTCCGAGAGGGAGCACTGTTGCTGCATCAAGATTCGCTCTCTCGCAGCTGCGTCGGGAGGAGTGTACGTTGAGTTGCAGTGTAGTGCGCTACCTGTGAATCCACAGCGTTGACTGCGTGGCCGACATTTGTCACAATCCTCAGCTTCAATTAATTTAGTAGGTGCGCCATCCCAGGCACTGATATATTTTTGCATTTTAGGCAGTGGTGAGTAGGAGGCTATTCCAAGCTTAATAAAGTACCCTGAAATGAAATAGAAAACTTTTTGTTCGATATGATTAGAGCAGCTTAAGTTAGAGGATACCCATAACAA?????????????????????????????????

			>Alcantarea_odorata
			AGAAAGAGAGAAAGAAAGAAAGGCGTACGGAGGGGGGGGAAGGAGCGGCTGGAGAGCGATTGGACGGAAAGCTGTGGATAGCTGGGGAGGGAGAGGGAGCCGTGAGTTGTCTTGTCGGGACCTGACATTGCATTTTCTCGGAGACACCCCGCTGTTGGGCGGCCGCCACGTGTCCGCGGAAGTGACCCGGGCCCCACGCTCGCGCGCGGTGGGGGCGTAATGAGTGCACAGCGGAGCTCCGAGAGGGAGCACTGTTGCTGCATCAAGATTCGCTCTCTCGCAGCTGCGTCGGGAGGAGTGTACGTTGAGTTGCAGTGTAGTGCGCTACCTGTGAATCCACAGCGTTGACTGCGTGGCCGACATTTGTCACAATCCTCAGTTTCAATTAATTTAGTTGGTGCGCCATCCCAGGCACTGATATATTTTTGCATTTTAGGCAGTGGTGAGTAGGAGGCTATTCCAAGCTTAATAAAGTACCCTGAAATGAAATAGAAAACTTTTTGTTCGATATGATTAGAGCAGCTTAAGTTAAGGGATACCCATAACAATTAAGATGTGCATGCAAATCTAGTTGCCAAAAC

			>Alcantarea_recurvifolia
			?????????????????????????????????????????????????????????????????????????????????????????????????????????????????????????????????????????????????????????????????????????????????????????????????????????????????????????????????????????????????????????????????????????????????????????????????????????????????????????????????????????????????????????????????????????????????????????????????????????????????????????????????????????????????????????????????????????????????????????????????????????????????????????????????????????????????????????????????????????????????????????????????????

			>Alcantarea_regina
			AGAGAGAAAGAAAGAAAGAAAGGCGTACGGAGGGGGGGGAAGGATTGGCTGGAGAGCGATTGGACGGAAAGCTGTGGATAGCTGGGGAGGGAGAGGCAGCCGTGAGTTGTCTTGTCGGGACCTGACATTGCATTTTCTCGGAGACACCCCGCTGTTGGGCGGCCGCCACGTGTCCGCGGAAGTGACCCGGGCCCCACGCTCGCGCGCGGTGGGGGCGTAATGAGTGCACAGCGGAGCTCCGAGAGGGAGCACTGTTGCTGCATCAAGATTCGCTCTCTCGCAGCTGCGTCGGGAGGAGTGTACGTTGAGTTGCAGTGTAGTGCGCTACCTGTGAATCCACAGCGTTGACTGCGTGGCCGACATTTGTCACAATCCTCGGCTTCAATTAATTTAGTTGGTGCGCCATCCCAGGCACTGATACATTTTTGCATTTTAGGCAGTGGTGAGTAGGAGGCTATTCCAAGCTTAATAAAGTACCCTGAAATGAAATAGAAAACTTTTTGTTCGATATGATTAGAGCAGCTTAAGTTAGAGGATACCCATAACAATTAAGATGTGCATGCAAATCTAGTTGCCAAAAC

			>Alcantarea_roberto-kautskyi
			AGAAAGAAAGAAAGAAAGAAAGGCGTACGGAGGGGGGGGGAGGAGCGGCTGGAGAGCGATTGGACGGAAAGCTGTGGATAGCTGGGGAGGGAGAGGCAGCCGTGAGTTGTCTTGTCGGGACCTGACATTGCATTTTCTCGGAAACACCCCGCTGTTGGGCGGCCGCCACGTGTCCGCGGAAGTGACCCGGGCCCCACGCTCGCGCGCGGTGGGGGCGTAATGAGTGCACAGCGGGGCTCCGAGAGGGAGCACTGTTGCTGCATCAAGATTCG--CTCTCGCA-TTGCATCGGGAGGAGTGTACGTTGAGTTACAGTGTAGTGCGCTACCTGTGAATCCACAGCGTTGACTGCGTGGCCGACATTTGTCACAATCCTCAGCTTCAATTAATTTAGTTGGTGCGCCATCCCAGGCACTGATATATTTTTGCATTTTAGGCAGTGGTGAGTAGGAGGCTATTCCAAGCTTAATAAAGTACCCTGAAATGAAATAGAAAACTTTTTGTTCGATATGATTAGAGCAGCTTAAGTTAGAGGATACCCATAACAATTAAGATGCGCATGCAAATCTAGTTGCCAAAAC

			>Alcantarea_vasconcelosiana
			?????????????????????????????????????????????????????????????????????????????????????????????????????????????????????????????????????????????????????????????????????????????????????????????????????????????????????????????????????????????????????????????????????????????????????????????????????????????????????????????????????????????????????????????????????????????????????????????????????????????????????????????????????????????????????????????????????????????????????????????????????????????????????????????????????????????????????????????????????????????????????????????????????

			>Ananas_ananassoides
			?????????????????????????????????????????????????????????????????????????????????????????????????????????????????????????????????????????????????????????????????????????????????????????????????????????????????????????????????????????????????????????????????????????????????????????????????????????????????????????????????????????????????????????????????????????????????????????????????????????????????????????????????????????????????????????????????????????????????????????????????????????????????????????????????????????????????????????????????????????????????????????????????????

			>Ananas_ananassoides_var._nanus
			?????????????????????????????????????????????????????????????????????????????????????????????????????????????????????????????????????????????????????????????????????????????????????????????????????????????????????????????????????????????????????????????????????????????????????????????????????????????????????????????????????????????????????????????????????????????????????????????????????????????????????????????????????????????????????????????????????????????????????????????????????????????????????????????????????????????????????????????????????????????????????????????????????

			>Ananas_bracteatus
			?????????????????????????????????????????????????????????????????????????????????????????????????????????????????????????????????????????????????????????????????????????????????????????????????????????????????????????????????????????????????????????????????????????????????????????????????????????????????????????????????????????????????????????????????????????????????????????????????????????????????????????????????????????????????????????????????????????????????????????????????????????????????????????????????????????????????????????????????????????????????????????????????????

			>Ananas_comosus
			?????????????????????????????????????????????????????????????????????????????????????????????????????????????????????????????????????????????????????????????????????????????????????????????????????????????????????????????????????????????????????????????????????????????????????????????????????????????????????????????????????????????????????????????????????????????????????????????????????????????????????????????????????????????????????????????????????????????????????????????????????????????????????????????????????????????????????????????????????????????????????????????????????

			>Ananas_comosus_var._microstachys
			?????????????????????????????????????????????????????????????????????????????????????????????????????????????????????????????????????????????????????????????????????????????????????????????????????????????????????????????????????????????????????????????????????????????????????????????????????????????????????????????????????????????????????????????????????????????????????????????????????????????????????????????????????????????????????????????????????????????????????????????????????????????????????????????????????????????????????????????????????????????????????????????????????

			>Ananas_comosus_var._parguazensis
			?????????????????????????????????????????????????????????????????????????????????????????????????????????????????????????????????????????????????????????????????????????????????????????????????????????????????????????????????????????????????????????????????????????????????????????????????????????????????????????????????????????????????????????????????????????????????????????????????????????????????????????????????????????????????????????????????????????????????????????????????????????????????????????????????????????????????????????????????????????????????????????????????????

			>Ananas_fritzmuelleri
			?????????????????????????????????????????????????????????????????????????????????????????????????????????????????????????????????????????????????????????????????????????????????????????????????????????????????????????????????????????????????????????????????????????????????????????????????????????????????????????????????????????????????????????????????????????????????????????????????????????????????????????????????????????????????????????????????????????????????????????????????????????????????????????????????????????????????????????????????????????????????????????????????????

			>Ananas_sagenaria
			?????????????????????????????????????????????????????????????????????????????????????????????????????????????????????????????????????????????????????????????????????????????????????????????????????????????????????????????????????????????????????????????????????????????????????????????????????????????????????????????????????????????????????????????????????????????????????????????????????????????????????????????????????????????????????????????????????????????????????????????????????????????????????????????????????????????????????????????????????????????????????????????????????

			>Androlepis_fragrans
			?????????????????????????????????????????????????????????????????????????????????????????????????????????????????????????????????????????????????????????????????????????????????????????????????????????????????????????????????????????????????????????????????????????????????????????????????????????????????????????????????????????????????????????????????????????????????????????????????????????????????????????????????????????????????????????????????????????????????????????????????????????????????????????????????????????????????????????????????????????????????????????????????????

			>Androlepis_skinneri
			?????????????????????????????????????????????????????????????????????????????????????????????????????????????????????????????????????????????????????????????????????????????????????????????????????????????????????????????????????????????????????????????????????????????????????????????????????????????????????????????????????????????????????????????????????????????????????????????????????????????????????????????????????????????????????????????????????????????????????????????????????????????????????????????????????????????????????????????????????????????????????????????????????

			>Araeococcus_chlorocarpus
			?????????????????????????????????????????????????????????????????????????????????????????????????????????????????????????????????????????????????????????????????????????????????????????????????????????????????????????????????????????????????????????????????????????????????????????????????????????????????????????????????????????????????????????????????????????????????????????????????????????????????????????????????????????????????????????????????????????????????????????????????????????????????????????????????????????????????????????????????????????????????????????????????????

			>Araeococcus_flagellifolius
			?????????????????????????????????????????????????????????????????????????????????????????????????????????????????????????????????????????????????????????????????????????????????????????????????????????????????????????????????????????????????????????????????????????????????????????????????????????????????????????????????????????????????????????????????????????????????????????????????????????????????????????????????????????????????????????????????????????????????????????????????????????????????????????????????????????????????????????????????????????????????????????????????????

			>Araeococcus_goeldianus
			?????????????????????????????????????????????????????????????????????????????????????????????????????????????????????????????????????????????????????????????????????????????????????????????????????????????????????????????????????????????????????????????????????????????????????????????????????????????????????????????????????????????????????????????????????????????????????????????????????????????????????????????????????????????????????????????????????????????????????????????????????????????????????????????????????????????????????????????????????????????????????????????????????

			>Araeococcus_micranthus
			?????????????????????????????????????????????????????????????????????????????????????????????????????????????????????????????????????????????????????????????????????????????????????????????????????????????????????????????????????????????????????????????????????????????????????????????????????????????????????????????????????????????????????????????????????????????????????????????????????????????????????????????????????????????????????????????????????????????????????????????????????????????????????????????????????????????????????????????????????????????????????????????????????

			>Araeococcus_pectinatus
			?????????????????????????????????????????????????????????????????????????????????????????????????????????????????????????????????????????????????????????????????????????????????????????????????????????????????????????????????????????????????????????????????????????????????????????????????????????????????????????????????????????????????????????????????????????????????????????????????????????????????????????????????????????????????????????????????????????????????????????????????????????????????????????????????????????????????????????????????????????????????????????????????????

			>Barfussia_laxissima
			?????????????????????????????????????????????????????????????????????????????????????????????????????????????????????????????????????????????????????????????????????????????????????????????????????????????????????????????????????????????????????????????????????????????????????????????????????????????????????????????????????????????????????????????????????????????????????????????????????????????????????????????????????????????????????????????????????????????????????????????????????????????????????????????????????????????????????????????????????????????????????????????????????

			>Barfussia_platyrhachis
			?????????????????????????????????????????????????????????????????????????????????????????????????????????????????????????????????????????????????????????????????????????????????????????????????????????????????????????????????????????????????????????????????????????????????????????????????????????????????????????????????????????????????????????????????????????????????????????????????????????????????????????????????????????????????????????????????????????????????????????????????????????????????????????????????????????????????????????????????????????????????????????????????????

			>Barfussia_wagneriana
			?????????????????????????????????????????????????????????????????????????????????????????????????????????????????????????????????????????????????????????????????????????????????????????????????????????????????????????????????????????????????????????????????????????????????????????????????????????????????????????????????????????????????????????????????????????????????????????????????????????????????????????????????????????????????????????????????????????????????????????????????????????????????????????????????????????????????????????????????????????????????????????????????????

			>Billbergia_amoena
			?????????????????????????????????????????????????????????????????????????????????????????????????????????????????????????????????????????????????????????????????????????????????????????????????????????????????????????????????????????????????????????????????????????????????????????????????????????????????????????????????????????????????????????????????????????????????????????????????????????????????????????????????????????????????????????????????????????????????????????????????????????????????????????????????????????????????????????????????????????????????????????????????????

			>Billbergia_brachysiphon
			?????????????????????????????????????????????????????????????????????????????????????????????????????????????????????????????????????????????????????????????????????????????????????????????????????????????????????????????????????????????????????????????????????????????????????????????????????????????????????????????????????????????????????????????????????????????????????????????????????????????????????????????????????????????????????????????????????????????????????????????????????????????????????????????????????????????????????????????????????????????????????????????????????

			>Billbergia_brasiliensis
			?????????????????????????????????????????????????????????????????????????????????????????????????????????????????????????????????????????????????????????????????????????????????????????????????????????????????????????????????????????????????????????????????????????????????????????????????????????????????????????????????????????????????????????????????????????????????????????????????????????????????????????????????????????????????????????????????????????????????????????????????????????????????????????????????????????????????????????????????????????????????????????????????????

			>Billbergia_chlorantha
			?????????????????????????????????????????????????????????????????????????????????????????????????????????????????????????????????????????????????????????????????????????????????????????????????????????????????????????????????????????????????????????????????????????????????????????????????????????????????????????????????????????????????????????????????????????????????????????????????????????????????????????????????????????????????????????????????????????????????????????????????????????????????????????????????????????????????????????????????????????????????????????????????????

			>Billbergia_decora
			?????????????????????????????????????????????????????????????????????????????????????????????????????????????????????????????????????????????????????????????????????????????????????????????????????????????????????????????????????????????????????????????????????????????????????????????????????????????????????????????????????????????????????????????????????????????????????????????????????????????????????????????????????????????????????????????????????????????????????????????????????????????????????????????????????????????????????????????????????????????????????????????????????

			>Billbergia_distachia
			?????????????????????????????????????????????????????????????????????????????????????????????????????????????????????????????????????????????????????????????????????????????????????????????????????????????????????????????????????????????????????????????????????????????????????????????????????????????????????????????????????????????????????????????????????????????????????????????????????????????????????????????????????????????????????????????????????????????????????????????????????????????????????????????????????????????????????????????????????????????????????????????????????

			>Billbergia_eloiseae
			?????????????????????????????????????????????????????????????????????????????????????????????????????????????????????????????????????????????????????????????????????????????????????????????????????????????????????????????????????????????????????????????????????????????????????????????????????????????????????????????????????????????????????????????????????????????????????????????????????????????????????????????????????????????????????????????????????????????????????????????????????????????????????????????????????????????????????????????????????????????????????????????????????

			>Billbergia_euphemiae
			?????????????????????????????????????????????????????????????????????????????????????????????????????????????????????????????????????????????????????????????????????????????????????????????????????????????????????????????????????????????????????????????????????????????????????????????????????????????????????????????????????????????????????????????????????????????????????????????????????????????????????????????????????????????????????????????????????????????????????????????????????????????????????????????????????????????????????????????????????????????????????????????????????

			>Billbergia_horrida
			?????????????????????????????????????????????????????????????????????????????????????????????????????????????????????????????????????????????????????????????????????????????????????????????????????????????????????????????????????????????????????????????????????????????????????????????????????????????????????????????????????????????????????????????????????????????????????????????????????????????????????????????????????????????????????????????????????????????????????????????????????????????????????????????????????????????????????????????????????????????????????????????????????

			>Billbergia_kautskyana
			?????????????????????????????????????????????????????????????????????????????????????????????????????????????????????????????????????????????????????????????????????????????????????????????????????????????????????????????????????????????????????????????????????????????????????????????????????????????????????????????????????????????????????????????????????????????????????????????????????????????????????????????????????????????????????????????????????????????????????????????????????????????????????????????????????????????????????????????????????????????????????????????????????

			>Billbergia_leptopoda
			?????????????????????????????????????????????????????????????????????????????????????????????????????????????????????????????????????????????????????????????????????????????????????????????????????????????????????????????????????????????????????????????????????????????????????????????????????????????????????????????????????????????????????????????????????????????????????????????????????????????????????????????????????????????????????????????????????????????????????????????????????????????????????????????????????????????????????????????????????????????????????????????????????

			>Billbergia_magnifica
			?????????????????????????????????????????????????????????????????????????????????????????????????????????????????????????????????????????????????????????????????????????????????????????????????????????????????????????????????????????????????????????????????????????????????????????????????????????????????????????????????????????????????????????????????????????????????????????????????????????????????????????????????????????????????????????????????????????????????????????????????????????????????????????????????????????????????????????????????????????????????????????????????????

			>Billbergia_manarae
			?????????????????????????????????????????????????????????????????????????????????????????????????????????????????????????????????????????????????????????????????????????????????????????????????????????????????????????????????????????????????????????????????????????????????????????????????????????????????????????????????????????????????????????????????????????????????????????????????????????????????????????????????????????????????????????????????????????????????????????????????????????????????????????????????????????????????????????????????????????????????????????????????????

			>Billbergia_meyeri
			?????????????????????????????????????????????????????????????????????????????????????????????????????????????????????????????????????????????????????????????????????????????????????????????????????????????????????????????????????????????????????????????????????????????????????????????????????????????????????????????????????????????????????????????????????????????????????????????????????????????????????????????????????????????????????????????????????????????????????????????????????????????????????????????????????????????????????????????????????????????????????????????????????

			>Billbergia_morelii
			?????????????????????????????????????????????????????????????????????????????????????????????????????????????????????????????????????????????????????????????????????????????????????????????????????????????????????????????????????????????????????????????????????????????????????????????????????????????????????????????????????????????????????????????????????????????????????????????????????????????????????????????????????????????????????????????????????????????????????????????????????????????????????????????????????????????????????????????????????????????????????????????????????

			>Billbergia_nana
			?????????????????????????????????????????????????????????????????????????????????????????????????????????????????????????????????????????????????????????????????????????????????????????????????????????????????????????????????????????????????????????????????????????????????????????????????????????????????????????????????????????????????????????????????????????????????????????????????????????????????????????????????????????????????????????????????????????????????????????????????????????????????????????????????????????????????????????????????????????????????????????????????????

			>Billbergia_nutans
			?????????????????????????????????????????????????????????????????????????????????????????????????????????????????????????????????????????????????????????????????????????????????????????????????????????????????????????????????????????????????????????????????????????????????????????????????????????????????????????????????????????????????????????????????????????????????????????????????????????????????????????????????????????????????????????????????????????????????????????????????????????????????????????????????????????????????????????????????????????????????????????????????????

			>Billbergia_pallidiflora
			?????????????????????????????????????????????????????????????????????????????????????????????????????????????????????????????????????????????????????????????????????????????????????????????????????????????????????????????????????????????????????????????????????????????????????????????????????????????????????????????????????????????????????????????????????????????????????????????????????????????????????????????????????????????????????????????????????????????????????????????????????????????????????????????????????????????????????????????????????????????????????????????????????

			>Billbergia_porteana
			?????????????????????????????????????????????????????????????????????????????????????????????????????????????????????????????????????????????????????????????????????????????????????????????????????????????????????????????????????????????????????????????????????????????????????????????????????????????????????????????????????????????????????????????????????????????????????????????????????????????????????????????????????????????????????????????????????????????????????????????????????????????????????????????????????????????????????????????????????????????????????????????????????

			>Billbergia_pyramidalis
			?????????????????????????????????????????????????????????????????????????????????????????????????????????????????????????????????????????????????????????????????????????????????????????????????????????????????????????????????????????????????????????????????????????????????????????????????????????????????????????????????????????????????????????????????????????????????????????????????????????????????????????????????????????????????????????????????????????????????????????????????????????????????????????????????????????????????????????????????????????????????????????????????????

			>Billbergia_robert-readii
			?????????????????????????????????????????????????????????????????????????????????????????????????????????????????????????????????????????????????????????????????????????????????????????????????????????????????????????????????????????????????????????????????????????????????????????????????????????????????????????????????????????????????????????????????????????????????????????????????????????????????????????????????????????????????????????????????????????????????????????????????????????????????????????????????????????????????????????????????????????????????????????????????????

			>Billbergia_rosea
			?????????????????????????????????????????????????????????????????????????????????????????????????????????????????????????????????????????????????????????????????????????????????????????????????????????????????????????????????????????????????????????????????????????????????????????????????????????????????????????????????????????????????????????????????????????????????????????????????????????????????????????????????????????????????????????????????????????????????????????????????????????????????????????????????????????????????????????????????????????????????????????????????????

			>Billbergia_speciosa
			?????????????????????????????????????????????????????????????????????????????????????????????????????????????????????????????????????????????????????????????????????????????????????????????????????????????????????????????????????????????????????????????????????????????????????????????????????????????????????????????????????????????????????????????????????????????????????????????????????????????????????????????????????????????????????????????????????????????????????????????????????????????????????????????????????????????????????????????????????????????????????????????????????

			>Billbergia_stenopetala
			?????????????????????????????????????????????????????????????????????????????????????????????????????????????????????????????????????????????????????????????????????????????????????????????????????????????????????????????????????????????????????????????????????????????????????????????????????????????????????????????????????????????????????????????????????????????????????????????????????????????????????????????????????????????????????????????????????????????????????????????????????????????????????????????????????????????????????????????????????????????????????????????????????

			>Billbergia_tweedieana
			?????????????????????????????????????????????????????????????????????????????????????????????????????????????????????????????????????????????????????????????????????????????????????????????????????????????????????????????????????????????????????????????????????????????????????????????????????????????????????????????????????????????????????????????????????????????????????????????????????????????????????????????????????????????????????????????????????????????????????????????????????????????????????????????????????????????????????????????????????????????????????????????????????

			>Billbergia_velascana
			?????????????????????????????????????????????????????????????????????????????????????????????????????????????????????????????????????????????????????????????????????????????????????????????????????????????????????????????????????????????????????????????????????????????????????????????????????????????????????????????????????????????????????????????????????????????????????????????????????????????????????????????????????????????????????????????????????????????????????????????????????????????????????????????????????????????????????????????????????????????????????????????????????

			>Billbergia_violacea
			?????????????????????????????????????????????????????????????????????????????????????????????????????????????????????????????????????????????????????????????????????????????????????????????????????????????????????????????????????????????????????????????????????????????????????????????????????????????????????????????????????????????????????????????????????????????????????????????????????????????????????????????????????????????????????????????????????????????????????????????????????????????????????????????????????????????????????????????????????????????????????????????????????

			>Billbergia_viridiflora
			?????????????????????????????????????????????????????????????????????????????????????????????????????????????????????????????????????????????????????????????????????????????????????????????????????????????????????????????????????????????????????????????????????????????????????????????????????????????????????????????????????????????????????????????????????????????????????????????????????????????????????????????????????????????????????????????????????????????????????????????????????????????????????????????????????????????????????????????????????????????????????????????????????

			>Billbergia_vittata
			?????????????????????????????????????????????????????????????????????????????????????????????????????????????????????????????????????????????????????????????????????????????????????????????????????????????????????????????????????????????????????????????????????????????????????????????????????????????????????????????????????????????????????????????????????????????????????????????????????????????????????????????????????????????????????????????????????????????????????????????????????????????????????????????????????????????????????????????????????????????????????????????????????

			>Billbergia_zebrina
			?????????????????????????????????????????????????????????????????????????????????????????????????????????????????????????????????????????????????????????????????????????????????????????????????????????????????????????????????????????????????????????????????????????????????????????????????????????????????????????????????????????????????????????????????????????????????????????????????????????????????????????????????????????????????????????????????????????????????????????????????????????????????????????????????????????????????????????????????????????????????????????????????????

			>Brewcaria_reflexa
			?????????????????????????????????????????????????????????????????????????????????????????????????????????????????????????????????????????????????????????????????????????????????????????????????????????????????????????????????????????????????????????????????????????????????????????????????????????????????????????????????????????????????????????????????????????????????????????????????????????????????????????????????????????????????????????????????????????????????????????????????????????????????????????????????????????????????????????????????????????????????????????????????????

			>Brocchinia_acuminata
			?????????????????????????????????????????????????????????????????????????????????????????????????????????????????????????????????????????????????????????????????????????????????????????????????????????????????????????????????????????????????????????????????????????????????????????????????????????????????????????????????????????????????????????????????????????????????????????????????????????????????????????????????????????????????????????????????????????????????????????????????????????????????????????????????????????????????????????????????????????????????????????????????????

			>Brocchinia_micrantha
			?????????????????????????????????????????????????????????????????????????????????????????????????????????????????????????????????????????????????????????????????????????????????????????????????????????????????????????????????????????????????????????????????????????????????????????????????????????????????????????????????????????????????????????????????????????????????????????????????????????????????????????????????????????????????????????????????????????????????????????????????????????????????????????????????????????????????????????????????????????????????????????????????????

			>Brocchinia_paniculata
			?????????????????????????????????????????????????????????????????????????????????????????????????????????????????????????????????????????????????????????????????????????????????????????????????????????????????????????????????????????????????????????????????????????????????????????????????????????????????????????????????????????????????????????????????????????????????????????????????????????????????????????????????????????????????????????????????????????????????????????????????????????????????????????????????????????????????????????????????????????????????????????????????????

			>Brocchinia_prismatica
			?????????????????????????????????????????????????????????????????????????????????????????????????????????????????????????????????????????????????????????????????????????????????????????????????????????????????????????????????????????????????????????????????????????????????????????????????????????????????????????????????????????????????????????????????????????????????????????????????????????????????????????????????????????????????????????????????????????????????????????????????????????????????????????????????????????????????????????????????????????????????????????????????????

			>Brocchinia_reducta
			?????????????????????????????????????????????????????????????????????????????????????????????????????????????????????????????????????????????????????????????????????????????????????????????????????????????????????????????????????????????????????????????????????????????????????????????????????????????????????????????????????????????????????????????????????????????????????????????????????????????????????????????????????????????????????????????????????????????????????????????????????????????????????????????????????????????????????????????????????????????????????????????????????

			>Brocchinia_steyermarkii
			?????????????????????????????????????????????????????????????????????????????????????????????????????????????????????????????????????????????????????????????????????????????????????????????????????????????????????????????????????????????????????????????????????????????????????????????????????????????????????????????????????????????????????????????????????????????????????????????????????????????????????????????????????????????????????????????????????????????????????????????????????????????????????????????????????????????????????????????????????????????????????????????????????

			>Brocchinia_tatei
			?????????????????????????????????????????????????????????????????????????????????????????????????????????????????????????????????????????????????????????????????????????????????????????????????????????????????????????????????????????????????????????????????????????????????????????????????????????????????????????????????????????????????????????????????????????????????????????????????????????????????????????????????????????????????????????????????????????????????????????????????????????????????????????????????????????????????????????????????????????????????????????????????????

			>Brocchinia_uaipanensis
			?????????????????????????????????????????????????????????????????????????????????????????????????????????????????????????????????????????????????????????????????????????????????????????????????????????????????????????????????????????????????????????????????????????????????????????????????????????????????????????????????????????????????????????????????????????????????????????????????????????????????????????????????????????????????????????????????????????????????????????????????????????????????????????????????????????????????????????????????????????????????????????????????????

			>Bromelia_agavifolia
			?????????????????????????????????????????????????????????????????????????????????????????????????????????????????????????????????????????????????????????????????????????????????????????????????????????????????????????????????????????????????????????????????????????????????????????????????????????????????????????????????????????????????????????????????????????????????????????????????????????????????????????????????????????????????????????????????????????????????????????????????????????????????????????????????????????????????????????????????????????????????????????????????????

			>Bromelia_antiacantha
			?????????????????????????????????????????????????????????????????????????????????????????????????????????????????????????????????????????????????????????????????????????????????????????????????????????????????????????????????????????????????????????????????????????????????????????????????????????????????????????????????????????????????????????????????????????????????????????????????????????????????????????????????????????????????????????????????????????????????????????????????????????????????????????????????????????????????????????????????????????????????????????????????????

			>Bromelia_balansae
			?????????????????????????????????????????????????????????????????????????????????????????????????????????????????????????????????????????????????????????????????????????????????????????????????????????????????????????????????????????????????????????????????????????????????????????????????????????????????????????????????????????????????????????????????????????????????????????????????????????????????????????????????????????????????????????????????????????????????????????????????????????????????????????????????????????????????????????????????????????????????????????????????????

			>Bromelia_chrysantha
			?????????????????????????????????????????????????????????????????????????????????????????????????????????????????????????????????????????????????????????????????????????????????????????????????????????????????????????????????????????????????????????????????????????????????????????????????????????????????????????????????????????????????????????????????????????????????????????????????????????????????????????????????????????????????????????????????????????????????????????????????????????????????????????????????????????????????????????????????????????????????????????????????????

			>Bromelia_flemingii
			?????????????????????????????????????????????????????????????????????????????????????????????????????????????????????????????????????????????????????????????????????????????????????????????????????????????????????????????????????????????????????????????????????????????????????????????????????????????????????????????????????????????????????????????????????????????????????????????????????????????????????????????????????????????????????????????????????????????????????????????????????????????????????????????????????????????????????????????????????????????????????????????????????

			>Bromelia_hemisphaerica
			?????????????????????????????????????????????????????????????????????????????????????????????????????????????????????????????????????????????????????????????????????????????????????????????????????????????????????????????????????????????????????????????????????????????????????????????????????????????????????????????????????????????????????????????????????????????????????????????????????????????????????????????????????????????????????????????????????????????????????????????????????????????????????????????????????????????????????????????????????????????????????????????????????

			>Bromelia_irwinii
			?????????????????????????????????????????????????????????????????????????????????????????????????????????????????????????????????????????????????????????????????????????????????????????????????????????????????????????????????????????????????????????????????????????????????????????????????????????????????????????????????????????????????????????????????????????????????????????????????????????????????????????????????????????????????????????????????????????????????????????????????????????????????????????????????????????????????????????????????????????????????????????????????????

			>Bromelia_karatas
			?????????????????????????????????????????????????????????????????????????????????????????????????????????????????????????????????????????????????????????????????????????????????????????????????????????????????????????????????????????????????????????????????????????????????????????????????????????????????????????????????????????????????????????????????????????????????????????????????????????????????????????????????????????????????????????????????????????????????????????????????????????????????????????????????????????????????????????????????????????????????????????????????????

			>Bromelia_pinguin
			?????????????????????????????????????????????????????????????????????????????????????????????????????????????????????????????????????????????????????????????????????????????????????????????????????????????????????????????????????????????????????????????????????????????????????????????????????????????????????????????????????????????????????????????????????????????????????????????????????????????????????????????????????????????????????????????????????????????????????????????????????????????????????????????????????????????????????????????????????????????????????????????????????

			>Bromelia_serra
			?????????????????????????????????????????????????????????????????????????????????????????????????????????????????????????????????????????????????????????????????????????????????????????????????????????????????????????????????????????????????????????????????????????????????????????????????????????????????????????????????????????????????????????????????????????????????????????????????????????????????????????????????????????????????????????????????????????????????????????????????????????????????????????????????????????????????????????????????????????????????????????????????????

			>Bromelia_trianae
			?????????????????????????????????????????????????????????????????????????????????????????????????????????????????????????????????????????????????????????????????????????????????????????????????????????????????????????????????????????????????????????????????????????????????????????????????????????????????????????????????????????????????????????????????????????????????????????????????????????????????????????????????????????????????????????????????????????????????????????????????????????????????????????????????????????????????????????????????????????????????????????????????????

			>Canistropsis_albiflora
			?????????????????????????????????????????????????????????????????????????????????????????????????????????????????????????????????????????????????????????????????????????????????????????????????????????????????????????????????????????????????????????????????????????????????????????????????????????????????????????????????????????????????????????????????????????????????????????????????????????????????????????????????????????????????????????????????????????????????????????????????????????????????????????????????????????????????????????????????????????????????????????????????????

			>Canistropsis_billbergioides
			?????????????????????????????????????????????????????????????????????????????????????????????????????????????????????????????????????????????????????????????????????????????????????????????????????????????????????????????????????????????????????????????????????????????????????????????????????????????????????????????????????????????????????????????????????????????????????????????????????????????????????????????????????????????????????????????????????????????????????????????????????????????????????????????????????????????????????????????????????????????????????????????????????

			>Canistropsis_burchellii
			?????????????????????????????????????????????????????????????????????????????????????????????????????????????????????????????????????????????????????????????????????????????????????????????????????????????????????????????????????????????????????????????????????????????????????????????????????????????????????????????????????????????????????????????????????????????????????????????????????????????????????????????????????????????????????????????????????????????????????????????????????????????????????????????????????????????????????????????????????????????????????????????????????

			>Canistropsis_correia-araujoi
			?????????????????????????????????????????????????????????????????????????????????????????????????????????????????????????????????????????????????????????????????????????????????????????????????????????????????????????????????????????????????????????????????????????????????????????????????????????????????????????????????????????????????????????????????????????????????????????????????????????????????????????????????????????????????????????????????????????????????????????????????????????????????????????????????????????????????????????????????????????????????????????????????????

			>Canistropsis_elata
			?????????????????????????????????????????????????????????????????????????????????????????????????????????????????????????????????????????????????????????????????????????????????????????????????????????????????????????????????????????????????????????????????????????????????????????????????????????????????????????????????????????????????????????????????????????????????????????????????????????????????????????????????????????????????????????????????????????????????????????????????????????????????????????????????????????????????????????????????????????????????????????????????????

			>Canistropsis_exigua
			?????????????????????????????????????????????????????????????????????????????????????????????????????????????????????????????????????????????????????????????????????????????????????????????????????????????????????????????????????????????????????????????????????????????????????????????????????????????????????????????????????????????????????????????????????????????????????????????????????????????????????????????????????????????????????????????????????????????????????????????????????????????????????????????????????????????????????????????????????????????????????????????????????

			>Canistropsis_marceloi
			?????????????????????????????????????????????????????????????????????????????????????????????????????????????????????????????????????????????????????????????????????????????????????????????????????????????????????????????????????????????????????????????????????????????????????????????????????????????????????????????????????????????????????????????????????????????????????????????????????????????????????????????????????????????????????????????????????????????????????????????????????????????????????????????????????????????????????????????????????????????????????????????????????

			>Canistropsis_microps
			?????????????????????????????????????????????????????????????????????????????????????????????????????????????????????????????????????????????????????????????????????????????????????????????????????????????????????????????????????????????????????????????????????????????????????????????????????????????????????????????????????????????????????????????????????????????????????????????????????????????????????????????????????????????????????????????????????????????????????????????????????????????????????????????????????????????????????????????????????????????????????????????????????

			>Canistropsis_pulcherrima
			?????????????????????????????????????????????????????????????????????????????????????????????????????????????????????????????????????????????????????????????????????????????????????????????????????????????????????????????????????????????????????????????????????????????????????????????????????????????????????????????????????????????????????????????????????????????????????????????????????????????????????????????????????????????????????????????????????????????????????????????????????????????????????????????????????????????????????????????????????????????????????????????????????

			>Canistropsis_simulans
			?????????????????????????????????????????????????????????????????????????????????????????????????????????????????????????????????????????????????????????????????????????????????????????????????????????????????????????????????????????????????????????????????????????????????????????????????????????????????????????????????????????????????????????????????????????????????????????????????????????????????????????????????????????????????????????????????????????????????????????????????????????????????????????????????????????????????????????????????????????????????????????????????????

			>Canistrum_alagoanum
			?????????????????????????????????????????????????????????????????????????????????????????????????????????????????????????????????????????????????????????????????????????????????????????????????????????????????????????????????????????????????????????????????????????????????????????????????????????????????????????????????????????????????????????????????????????????????????????????????????????????????????????????????????????????????????????????????????????????????????????????????????????????????????????????????????????????????????????????????????????????????????????????????????

			>Canistrum_aurantiacum
			?????????????????????????????????????????????????????????????????????????????????????????????????????????????????????????????????????????????????????????????????????????????????????????????????????????????????????????????????????????????????????????????????????????????????????????????????????????????????????????????????????????????????????????????????????????????????????????????????????????????????????????????????????????????????????????????????????????????????????????????????????????????????????????????????????????????????????????????????????????????????????????????????????

			>Canistrum_camacaense
			?????????????????????????????????????????????????????????????????????????????????????????????????????????????????????????????????????????????????????????????????????????????????????????????????????????????????????????????????????????????????????????????????????????????????????????????????????????????????????????????????????????????????????????????????????????????????????????????????????????????????????????????????????????????????????????????????????????????????????????????????????????????????????????????????????????????????????????????????????????????????????????????????????

			>Canistrum_fosterianum
			?????????????????????????????????????????????????????????????????????????????????????????????????????????????????????????????????????????????????????????????????????????????????????????????????????????????????????????????????????????????????????????????????????????????????????????????????????????????????????????????????????????????????????????????????????????????????????????????????????????????????????????????????????????????????????????????????????????????????????????????????????????????????????????????????????????????????????????????????????????????????????????????????????

			>Canistrum_pickelii
			?????????????????????????????????????????????????????????????????????????????????????????????????????????????????????????????????????????????????????????????????????????????????????????????????????????????????????????????????????????????????????????????????????????????????????????????????????????????????????????????????????????????????????????????????????????????????????????????????????????????????????????????????????????????????????????????????????????????????????????????????????????????????????????????????????????????????????????????????????????????????????????????????????

			>Canistrum_triangulare
			?????????????????????????????????????????????????????????????????????????????????????????????????????????????????????????????????????????????????????????????????????????????????????????????????????????????????????????????????????????????????????????????????????????????????????????????????????????????????????????????????????????????????????????????????????????????????????????????????????????????????????????????????????????????????????????????????????????????????????????????????????????????????????????????????????????????????????????????????????????????????????????????????????

			>Catopsis_berteroniana
			?????????????????????????????????????????????????????????????????????????????????????????????????????????????????????????????????????????????????????????????????????????????????????????????????????????????????????????????????????????????????????????????????????????????????????????????????????????????????????????????????????????????????????????????????????????????????????????????????????????????????????????????????????????????????????????????????????????????????????????????????????????????????????????????????????????????????????????????????????????????????????????????????????

			>Catopsis_floribunda
			?????????????????????????????????????????????????????????????????????????????????????????????????????????????????????????????????????????????????????????????????????????????????????????????????????????????????????????????????????????????????????????????????????????????????????????????????????????????????????????????????????????????????????????????????????????????????????????????????????????????????????????????????????????????????????????????????????????????????????????????????????????????????????????????????????????????????????????????????????????????????????????????????????

			>Catopsis_juncifolia
			?????????????????????????????????????????????????????????????????????????????????????????????????????????????????????????????????????????????????????????????????????????????????????????????????????????????????????????????????????????????????????????????????????????????????????????????????????????????????????????????????????????????????????????????????????????????????????????????????????????????????????????????????????????????????????????????????????????????????????????????????????????????????????????????????????????????????????????????????????????????????????????????????????

			>Catopsis_morreniana
			?????????????????????????????????????????????????????????????????????????????????????????????????????????????????????????????????????????????????????????????????????????????????????????????????????????????????????????????????????????????????????????????????????????????????????????????????????????????????????????????????????????????????????????????????????????????????????????????????????????????????????????????????????????????????????????????????????????????????????????????????????????????????????????????????????????????????????????????????????????????????????????????????????

			>Catopsis_nitida
			?????????????????????????????????????????????????????????????????????????????????????????????????????????????????????????????????????????????????????????????????????????????????????????????????????????????????????????????????????????????????????????????????????????????????????????????????????????????????????????????????????????????????????????????????????????????????????????????????????????????????????????????????????????????????????????????????????????????????????????????????????????????????????????????????????????????????????????????????????????????????????????????????????

			>Catopsis_nutans
			?????????????????????????????????????????????????????????????????????????????????????????????????????????????????????????????????????????????????????????????????????????????????????????????????????????????????????????????????????????????????????????????????????????????????????????????????????????????????????????????????????????????????????????????????????????????????????????????????????????????????????????????????????????????????????????????????????????????????????????????????????????????????????????????????????????????????????????????????????????????????????????????????????

			>Catopsis_nutans_var._nutans
			?????????????????????????????????????????????????????????????????????????????????????????????????????????????????????????????????????????????????????????????????????????????????????????????????????????????????????????????????????????????????????????????????????????????????????????????????????????????????????????????????????????????????????????????????????????????????????????????????????????????????????????????????????????????????????????????????????????????????????????????????????????????????????????????????????????????????????????????????????????????????????????????????????

			>Catopsis_paniculata
			?????????????????????????????????????????????????????????????????????????????????????????????????????????????????????????????????????????????????????????????????????????????????????????????????????????????????????????????????????????????????????????????????????????????????????????????????????????????????????????????????????????????????????????????????????????????????????????????????????????????????????????????????????????????????????????????????????????????????????????????????????????????????????????????????????????????????????????????????????????????????????????????????????

			>Catopsis_sessiliflora
			?????????????????????????????????????????????????????????????????????????????????????????????????????????????????????????????????????????????????????????????????????????????????????????????????????????????????????????????????????????????????????????????????????????????????????????????????????????????????????????????????????????????????????????????????????????????????????????????????????????????????????????????????????????????????????????????????????????????????????????????????????????????????????????????????????????????????????????????????????????????????????????????????????

			>Catopsis_subulata
			?????????????????????????????????????????????????????????????????????????????????????????????????????????????????????????????????????????????????????????????????????????????????????????????????????????????????????????????????????????????????????????????????????????????????????????????????????????????????????????????????????????????????????????????????????????????????????????????????????????????????????????????????????????????????????????????????????????????????????????????????????????????????????????????????????????????????????????????????????????????????????????????????????

			>Catopsis_wangerinii
			?????????????????????????????????????????????????????????????????????????????????????????????????????????????????????????????????????????????????????????????????????????????????????????????????????????????????????????????????????????????????????????????????????????????????????????????????????????????????????????????????????????????????????????????????????????????????????????????????????????????????????????????????????????????????????????????????????????????????????????????????????????????????????????????????????????????????????????????????????????????????????????????????????

			>Cipuropsis_amicorum
			?????????????????????????????????????????????????????????????????????????????????????????????????????????????????????????????????????????????????????????????????????????????????????????????????????????????????????????????????????????????????????????????????????????????????????????????????????????????????????????????????????????????????????????????????????????????????????????????????????????????????????????????????????????????????????????????????????????????????????????????????????????????????????????????????????????????????????????????????????????????????????????????????????

			>Connellia_nutans
			?????????????????????????????????????????????????????????????????????????????????????????????????????????????????????????????????????????????????????????????????????????????????????????????????????????????????????????????????????????????????????????????????????????????????????????????????????????????????????????????????????????????????????????????????????????????????????????????????????????????????????????????????????????????????????????????????????????????????????????????????????????????????????????????????????????????????????????????????????????????????????????????????????

			>Cottendorfia_florida
			?????????????????????????????????????????????????????????????????????????????????????????????????????????????????????????????????????????????????????????????????????????????????????????????????????????????????????????????????????????????????????????????????????????????????????????????????????????????????????????????????????????????????????????????????????????????????????????????????????????????????????????????????????????????????????????????????????????????????????????????????????????????????????????????????????????????????????????????????????????????????????????????????????

			>Cryptanthus_acaulis
			?????????????????????????????????????????????????????????????????????????????????????????????????????????????????????????????????????????????????????????????????????????????????????????????????????????????????????????????????????????????????????????????????????????????????????????????????????????????????????????????????????????????????????????????????????????????????????????????????????????????????????????????????????????????????????????????????????????????????????????????????????????????????????????????????????????????????????????????????????????????????????????????????????

			>Cryptanthus_bahianus
			?????????????????????????????????????????????????????????????????????????????????????????????????????????????????????????????????????????????????????????????????????????????????????????????????????????????????????????????????????????????????????????????????????????????????????????????????????????????????????????????????????????????????????????????????????????????????????????????????????????????????????????????????????????????????????????????????????????????????????????????????????????????????????????????????????????????????????????????????????????????????????????????????????

			>Cryptanthus_beuckeri
			?????????????????????????????????????????????????????????????????????????????????????????????????????????????????????????????????????????????????????????????????????????????????????????????????????????????????????????????????????????????????????????????????????????????????????????????????????????????????????????????????????????????????????????????????????????????????????????????????????????????????????????????????????????????????????????????????????????????????????????????????????????????????????????????????????????????????????????????????????????????????????????????????????

			>Cryptanthus_bromelioides
			?????????????????????????????????????????????????????????????????????????????????????????????????????????????????????????????????????????????????????????????????????????????????????????????????????????????????????????????????????????????????????????????????????????????????????????????????????????????????????????????????????????????????????????????????????????????????????????????????????????????????????????????????????????????????????????????????????????????????????????????????????????????????????????????????????????????????????????????????????????????????????????????????????

			>Cryptanthus_burle-marxii
			?????????????????????????????????????????????????????????????????????????????????????????????????????????????????????????????????????????????????????????????????????????????????????????????????????????????????????????????????????????????????????????????????????????????????????????????????????????????????????????????????????????????????????????????????????????????????????????????????????????????????????????????????????????????????????????????????????????????????????????????????????????????????????????????????????????????????????????????????????????????????????????????????????

			>Cryptanthus_colnagoi
			?????????????????????????????????????????????????????????????????????????????????????????????????????????????????????????????????????????????????????????????????????????????????????????????????????????????????????????????????????????????????????????????????????????????????????????????????????????????????????????????????????????????????????????????????????????????????????????????????????????????????????????????????????????????????????????????????????????????????????????????????????????????????????????????????????????????????????????????????????????????????????????????????????

			>Cryptanthus_diamantinensis
			?????????????????????????????????????????????????????????????????????????????????????????????????????????????????????????????????????????????????????????????????????????????????????????????????????????????????????????????????????????????????????????????????????????????????????????????????????????????????????????????????????????????????????????????????????????????????????????????????????????????????????????????????????????????????????????????????????????????????????????????????????????????????????????????????????????????????????????????????????????????????????????????????????

			>Cryptanthus_dianae
			?????????????????????????????????????????????????????????????????????????????????????????????????????????????????????????????????????????????????????????????????????????????????????????????????????????????????????????????????????????????????????????????????????????????????????????????????????????????????????????????????????????????????????????????????????????????????????????????????????????????????????????????????????????????????????????????????????????????????????????????????????????????????????????????????????????????????????????????????????????????????????????????????????

			>Cryptanthus_maritimus
			?????????????????????????????????????????????????????????????????????????????????????????????????????????????????????????????????????????????????????????????????????????????????????????????????????????????????????????????????????????????????????????????????????????????????????????????????????????????????????????????????????????????????????????????????????????????????????????????????????????????????????????????????????????????????????????????????????????????????????????????????????????????????????????????????????????????????????????????????????????????????????????????????????

			>Cryptanthus_ruthae
			?????????????????????????????????????????????????????????????????????????????????????????????????????????????????????????????????????????????????????????????????????????????????????????????????????????????????????????????????????????????????????????????????????????????????????????????????????????????????????????????????????????????????????????????????????????????????????????????????????????????????????????????????????????????????????????????????????????????????????????????????????????????????????????????????????????????????????????????????????????????????????????????????????

			>Cryptanthus_teretifolius
			?????????????????????????????????????????????????????????????????????????????????????????????????????????????????????????????????????????????????????????????????????????????????????????????????????????????????????????????????????????????????????????????????????????????????????????????????????????????????????????????????????????????????????????????????????????????????????????????????????????????????????????????????????????????????????????????????????????????????????????????????????????????????????????????????????????????????????????????????????????????????????????????????????

			>Cryptanthus_warren-loosei
			?????????????????????????????????????????????????????????????????????????????????????????????????????????????????????????????????????????????????????????????????????????????????????????????????????????????????????????????????????????????????????????????????????????????????????????????????????????????????????????????????????????????????????????????????????????????????????????????????????????????????????????????????????????????????????????????????????????????????????????????????????????????????????????????????????????????????????????????????????????????????????????????????????

			>Cryptanthus_zonatus
			?????????????????????????????????????????????????????????????????????????????????????????????????????????????????????????????????????????????????????????????????????????????????????????????????????????????????????????????????????????????????????????????????????????????????????????????????????????????????????????????????????????????????????????????????????????????????????????????????????????????????????????????????????????????????????????????????????????????????????????????????????????????????????????????????????????????????????????????????????????????????????????????????????

			>Deinacanthon_urbanianum
			?????????????????????????????????????????????????????????????????????????????????????????????????????????????????????????????????????????????????????????????????????????????????????????????????????????????????????????????????????????????????????????????????????????????????????????????????????????????????????????????????????????????????????????????????????????????????????????????????????????????????????????????????????????????????????????????????????????????????????????????????????????????????????????????????????????????????????????????????????????????????????????????????????

			>Deuterocohnia_abstrusa
			?????????????????????????????????????????????????????????????????????????????????????????????????????????????????????????????????????????????????????????????????????????????????????????????????????????????????????????????????????????????????????????????????????????????????????????????????????????????????????????????????????????????????????????????????????????????????????????????????????????????????????????????????????????????????????????????????????????????????????????????????????????????????????????????????????????????????????????????????????????????????????????????????????

			>Deuterocohnia_brevifolia
			?????????????????????????????????????????????????????????????????????????????????????????????????????????????????????????????????????????????????????????????????????????????????????????????????????????????????????????????????????????????????????????????????????????????????????????????????????????????????????????????????????????????????????????????????????????????????????????????????????????????????????????????????????????????????????????????????????????????????????????????????????????????????????????????????????????????????????????????????????????????????????????????????????

			>Deuterocohnia_brevispicata
			?????????????????????????????????????????????????????????????????????????????????????????????????????????????????????????????????????????????????????????????????????????????????????????????????????????????????????????????????????????????????????????????????????????????????????????????????????????????????????????????????????????????????????????????????????????????????????????????????????????????????????????????????????????????????????????????????????????????????????????????????????????????????????????????????????????????????????????????????????????????????????????????????????

			>Deuterocohnia_chrysantha
			?????????????????????????????????????????????????????????????????????????????????????????????????????????????????????????????????????????????????????????????????????????????????????????????????????????????????????????????????????????????????????????????????????????????????????????????????????????????????????????????????????????????????????????????????????????????????????????????????????????????????????????????????????????????????????????????????????????????????????????????????????????????????????????????????????????????????????????????????????????????????????????????????????

			>Deuterocohnia_digitata
			?????????????????????????????????????????????????????????????????????????????????????????????????????????????????????????????????????????????????????????????????????????????????????????????????????????????????????????????????????????????????????????????????????????????????????????????????????????????????????????????????????????????????????????????????????????????????????????????????????????????????????????????????????????????????????????????????????????????????????????????????????????????????????????????????????????????????????????????????????????????????????????????????????

			>Deuterocohnia_gableana
			?????????????????????????????????????????????????????????????????????????????????????????????????????????????????????????????????????????????????????????????????????????????????????????????????????????????????????????????????????????????????????????????????????????????????????????????????????????????????????????????????????????????????????????????????????????????????????????????????????????????????????????????????????????????????????????????????????????????????????????????????????????????????????????????????????????????????????????????????????????????????????????????????????

			>Deuterocohnia_glandulosa
			?????????????????????????????????????????????????????????????????????????????????????????????????????????????????????????????????????????????????????????????????????????????????????????????????????????????????????????????????????????????????????????????????????????????????????????????????????????????????????????????????????????????????????????????????????????????????????????????????????????????????????????????????????????????????????????????????????????????????????????????????????????????????????????????????????????????????????????????????????????????????????????????????????

			>Deuterocohnia_haumanii
			?????????????????????????????????????????????????????????????????????????????????????????????????????????????????????????????????????????????????????????????????????????????????????????????????????????????????????????????????????????????????????????????????????????????????????????????????????????????????????????????????????????????????????????????????????????????????????????????????????????????????????????????????????????????????????????????????????????????????????????????????????????????????????????????????????????????????????????????????????????????????????????????????????

			>Deuterocohnia_longipetala
			?????????????????????????????????????????????????????????????????????????????????????????????????????????????????????????????????????????????????????????????????????????????????????????????????????????????????????????????????????????????????????????????????????????????????????????????????????????????????????????????????????????????????????????????????????????????????????????????????????????????????????????????????????????????????????????????????????????????????????????????????????????????????????????????????????????????????????????????????????????????????????????????????????

			>Deuterocohnia_lotteae
			?????????????????????????????????????????????????????????????????????????????????????????????????????????????????????????????????????????????????????????????????????????????????????????????????????????????????????????????????????????????????????????????????????????????????????????????????????????????????????????????????????????????????????????????????????????????????????????????????????????????????????????????????????????????????????????????????????????????????????????????????????????????????????????????????????????????????????????????????????????????????????????????????????

			>Deuterocohnia_meziana
			?????????????????????????????????????????????????????????????????????????????????????????????????????????????????????????????????????????????????????????????????????????????????????????????????????????????????????????????????????????????????????????????????????????????????????????????????????????????????????????????????????????????????????????????????????????????????????????????????????????????????????????????????????????????????????????????????????????????????????????????????????????????????????????????????????????????????????????????????????????????????????????????????????

			>Deuterocohnia_meziana_subsp._carmineoviridiflora
			?????????????????????????????????????????????????????????????????????????????????????????????????????????????????????????????????????????????????????????????????????????????????????????????????????????????????????????????????????????????????????????????????????????????????????????????????????????????????????????????????????????????????????????????????????????????????????????????????????????????????????????????????????????????????????????????????????????????????????????????????????????????????????????????????????????????????????????????????????????????????????????????????????

			>Deuterocohnia_meziana_subsp._pedicellata
			?????????????????????????????????????????????????????????????????????????????????????????????????????????????????????????????????????????????????????????????????????????????????????????????????????????????????????????????????????????????????????????????????????????????????????????????????????????????????????????????????????????????????????????????????????????????????????????????????????????????????????????????????????????????????????????????????????????????????????????????????????????????????????????????????????????????????????????????????????????????????????????????????????

			>Deuterocohnia_recurvipetala
			?????????????????????????????????????????????????????????????????????????????????????????????????????????????????????????????????????????????????????????????????????????????????????????????????????????????????????????????????????????????????????????????????????????????????????????????????????????????????????????????????????????????????????????????????????????????????????????????????????????????????????????????????????????????????????????????????????????????????????????????????????????????????????????????????????????????????????????????????????????????????????????????????????

			>Deuterocohnia_scapigera
			?????????????????????????????????????????????????????????????????????????????????????????????????????????????????????????????????????????????????????????????????????????????????????????????????????????????????????????????????????????????????????????????????????????????????????????????????????????????????????????????????????????????????????????????????????????????????????????????????????????????????????????????????????????????????????????????????????????????????????????????????????????????????????????????????????????????????????????????????????????????????????????????????????

			>Deuterocohnia_schreiteri
			?????????????????????????????????????????????????????????????????????????????????????????????????????????????????????????????????????????????????????????????????????????????????????????????????????????????????????????????????????????????????????????????????????????????????????????????????????????????????????????????????????????????????????????????????????????????????????????????????????????????????????????????????????????????????????????????????????????????????????????????????????????????????????????????????????????????????????????????????????????????????????????????????????

			>Deuterocohnia_seramisiana
			?????????????????????????????????????????????????????????????????????????????????????????????????????????????????????????????????????????????????????????????????????????????????????????????????????????????????????????????????????????????????????????????????????????????????????????????????????????????????????????????????????????????????????????????????????????????????????????????????????????????????????????????????????????????????????????????????????????????????????????????????????????????????????????????????????????????????????????????????????????????????????????????????????

			>Deuterocohnia_strobilifera_var._inermis
			?????????????????????????????????????????????????????????????????????????????????????????????????????????????????????????????????????????????????????????????????????????????????????????????????????????????????????????????????????????????????????????????????????????????????????????????????????????????????????????????????????????????????????????????????????????????????????????????????????????????????????????????????????????????????????????????????????????????????????????????????????????????????????????????????????????????????????????????????????????????????????????????????????

			>Deuterocohnia_strobilifera_var._strobilifera
			?????????????????????????????????????????????????????????????????????????????????????????????????????????????????????????????????????????????????????????????????????????????????????????????????????????????????????????????????????????????????????????????????????????????????????????????????????????????????????????????????????????????????????????????????????????????????????????????????????????????????????????????????????????????????????????????????????????????????????????????????????????????????????????????????????????????????????????????????????????????????????????????????????

			>Disteganthus_basilateralis
			?????????????????????????????????????????????????????????????????????????????????????????????????????????????????????????????????????????????????????????????????????????????????????????????????????????????????????????????????????????????????????????????????????????????????????????????????????????????????????????????????????????????????????????????????????????????????????????????????????????????????????????????????????????????????????????????????????????????????????????????????????????????????????????????????????????????????????????????????????????????????????????????????????

			>Disteganthus_lateralis
			?????????????????????????????????????????????????????????????????????????????????????????????????????????????????????????????????????????????????????????????????????????????????????????????????????????????????????????????????????????????????????????????????????????????????????????????????????????????????????????????????????????????????????????????????????????????????????????????????????????????????????????????????????????????????????????????????????????????????????????????????????????????????????????????????????????????????????????????????????????????????????????????????????

			>Dyckia_acutiflora
			?????????????????????????????????????????????????????????????????????????????????????????????????????????????????????????????????????????????????????????????????????????????????????????????????????????????????????????????????????????????????????????????????????????????????????????????????????????????????????????????????????????????????????????????????????????????????????????????????????????????????????????????????????????????????????????????????????????????????????????????????????????????????????????????????????????????????????????????????????????????????????????????????????

			>Dyckia_areniticola
			?????????????????????????????????????????????????????????????????????????????????????????????????????????????????????????????????????????????????????????????????????????????????????????????????????????????????????????????????????????????????????????????????????????????????????????????????????????????????????????????????????????????????????????????????????????????????????????????????????????????????????????????????????????????????????????????????????????????????????????????????????????????????????????????????????????????????????????????????????????????????????????????????????

			>Dyckia_aurea
			?????????????????????????????????????????????????????????????????????????????????????????????????????????????????????????????????????????????????????????????????????????????????????????????????????????????????????????????????????????????????????????????????????????????????????????????????????????????????????????????????????????????????????????????????????????????????????????????????????????????????????????????????????????????????????????????????????????????????????????????????????????????????????????????????????????????????????????????????????????????????????????????????????

			>Dyckia_beateae
			?????????????????????????????????????????????????????????????????????????????????????????????????????????????????????????????????????????????????????????????????????????????????????????????????????????????????????????????????????????????????????????????????????????????????????????????????????????????????????????????????????????????????????????????????????????????????????????????????????????????????????????????????????????????????????????????????????????????????????????????????????????????????????????????????????????????????????????????????????????????????????????????????????

			>Dyckia_brachyphylla
			?????????????????????????????????????????????????????????????????????????????????????????????????????????????????????????????????????????????????????????????????????????????????????????????????????????????????????????????????????????????????????????????????????????????????????????????????????????????????????????????????????????????????????????????????????????????????????????????????????????????????????????????????????????????????????????????????????????????????????????????????????????????????????????????????????????????????????????????????????????????????????????????????????

			>Dyckia_bracteata
			?????????????????????????????????????????????????????????????????????????????????????????????????????????????????????????????????????????????????????????????????????????????????????????????????????????????????????????????????????????????????????????????????????????????????????????????????????????????????????????????????????????????????????????????????????????????????????????????????????????????????????????????????????????????????????????????????????????????????????????????????????????????????????????????????????????????????????????????????????????????????????????????????????

			>Dyckia_braunii
			?????????????????????????????????????????????????????????????????????????????????????????????????????????????????????????????????????????????????????????????????????????????????????????????????????????????????????????????????????????????????????????????????????????????????????????????????????????????????????????????????????????????????????????????????????????????????????????????????????????????????????????????????????????????????????????????????????????????????????????????????????????????????????????????????????????????????????????????????????????????????????????????????????

			>Dyckia_brevifolia
			?????????????????????????????????????????????????????????????????????????????????????????????????????????????????????????????????????????????????????????????????????????????????????????????????????????????????????????????????????????????????????????????????????????????????????????????????????????????????????????????????????????????????????????????????????????????????????????????????????????????????????????????????????????????????????????????????????????????????????????????????????????????????????????????????????????????????????????????????????????????????????????????????????

			>Dyckia_choristaminea
			?????????????????????????????????????????????????????????????????????????????????????????????????????????????????????????????????????????????????????????????????????????????????????????????????????????????????????????????????????????????????????????????????????????????????????????????????????????????????????????????????????????????????????????????????????????????????????????????????????????????????????????????????????????????????????????????????????????????????????????????????????????????????????????????????????????????????????????????????????????????????????????????????????

			>Dyckia_cinerea
			?????????????????????????????????????????????????????????????????????????????????????????????????????????????????????????????????????????????????????????????????????????????????????????????????????????????????????????????????????????????????????????????????????????????????????????????????????????????????????????????????????????????????????????????????????????????????????????????????????????????????????????????????????????????????????????????????????????????????????????????????????????????????????????????????????????????????????????????????????????????????????????????????????

			>Dyckia_dawsonii
			?????????????????????????????????????????????????????????????????????????????????????????????????????????????????????????????????????????????????????????????????????????????????????????????????????????????????????????????????????????????????????????????????????????????????????????????????????????????????????????????????????????????????????????????????????????????????????????????????????????????????????????????????????????????????????????????????????????????????????????????????????????????????????????????????????????????????????????????????????????????????????????????????????

			>Dyckia_delicata
			?????????????????????????????????????????????????????????????????????????????????????????????????????????????????????????????????????????????????????????????????????????????????????????????????????????????????????????????????????????????????????????????????????????????????????????????????????????????????????????????????????????????????????????????????????????????????????????????????????????????????????????????????????????????????????????????????????????????????????????????????????????????????????????????????????????????????????????????????????????????????????????????????????

			>Dyckia_densiflora
			?????????????????????????????????????????????????????????????????????????????????????????????????????????????????????????????????????????????????????????????????????????????????????????????????????????????????????????????????????????????????????????????????????????????????????????????????????????????????????????????????????????????????????????????????????????????????????????????????????????????????????????????????????????????????????????????????????????????????????????????????????????????????????????????????????????????????????????????????????????????????????????????????????

			>Dyckia_dissitiflora
			?????????????????????????????????????????????????????????????????????????????????????????????????????????????????????????????????????????????????????????????????????????????????????????????????????????????????????????????????????????????????????????????????????????????????????????????????????????????????????????????????????????????????????????????????????????????????????????????????????????????????????????????????????????????????????????????????????????????????????????????????????????????????????????????????????????????????????????????????????????????????????????????????????

			>Dyckia_distachya
			?????????????????????????????????????????????????????????????????????????????????????????????????????????????????????????????????????????????????????????????????????????????????????????????????????????????????????????????????????????????????????????????????????????????????????????????????????????????????????????????????????????????????????????????????????????????????????????????????????????????????????????????????????????????????????????????????????????????????????????????????????????????????????????????????????????????????????????????????????????????????????????????????????

			>Dyckia_duckei
			?????????????????????????????????????????????????????????????????????????????????????????????????????????????????????????????????????????????????????????????????????????????????????????????????????????????????????????????????????????????????????????????????????????????????????????????????????????????????????????????????????????????????????????????????????????????????????????????????????????????????????????????????????????????????????????????????????????????????????????????????????????????????????????????????????????????????????????????????????????????????????????????????????

			>Dyckia_dusenii
			?????????????????????????????????????????????????????????????????????????????????????????????????????????????????????????????????????????????????????????????????????????????????????????????????????????????????????????????????????????????????????????????????????????????????????????????????????????????????????????????????????????????????????????????????????????????????????????????????????????????????????????????????????????????????????????????????????????????????????????????????????????????????????????????????????????????????????????????????????????????????????????????????????

			>Dyckia_elisabethae
			?????????????????????????????????????????????????????????????????????????????????????????????????????????????????????????????????????????????????????????????????????????????????????????????????????????????????????????????????????????????????????????????????????????????????????????????????????????????????????????????????????????????????????????????????????????????????????????????????????????????????????????????????????????????????????????????????????????????????????????????????????????????????????????????????????????????????????????????????????????????????????????????????????

			>Dyckia_encholirioides
			?????????????????????????????????????????????????????????????????????????????????????????????????????????????????????????????????????????????????????????????????????????????????????????????????????????????????????????????????????????????????????????????????????????????????????????????????????????????????????????????????????????????????????????????????????????????????????????????????????????????????????????????????????????????????????????????????????????????????????????????????????????????????????????????????????????????????????????????????????????????????????????????????????

			>Dyckia_espiritosantensis
			?????????????????????????????????????????????????????????????????????????????????????????????????????????????????????????????????????????????????????????????????????????????????????????????????????????????????????????????????????????????????????????????????????????????????????????????????????????????????????????????????????????????????????????????????????????????????????????????????????????????????????????????????????????????????????????????????????????????????????????????????????????????????????????????????????????????????????????????????????????????????????????????????????

			>Dyckia_estevesii
			?????????????????????????????????????????????????????????????????????????????????????????????????????????????????????????????????????????????????????????????????????????????????????????????????????????????????????????????????????????????????????????????????????????????????????????????????????????????????????????????????????????????????????????????????????????????????????????????????????????????????????????????????????????????????????????????????????????????????????????????????????????????????????????????????????????????????????????????????????????????????????????????????????

			>Dyckia_estevesii_var._braunii
			?????????????????????????????????????????????????????????????????????????????????????????????????????????????????????????????????????????????????????????????????????????????????????????????????????????????????????????????????????????????????????????????????????????????????????????????????????????????????????????????????????????????????????????????????????????????????????????????????????????????????????????????????????????????????????????????????????????????????????????????????????????????????????????????????????????????????????????????????????????????????????????????????????

			>Dyckia_excelsa
			?????????????????????????????????????????????????????????????????????????????????????????????????????????????????????????????????????????????????????????????????????????????????????????????????????????????????????????????????????????????????????????????????????????????????????????????????????????????????????????????????????????????????????????????????????????????????????????????????????????????????????????????????????????????????????????????????????????????????????????????????????????????????????????????????????????????????????????????????????????????????????????????????????

			>Dyckia_ferox
			?????????????????????????????????????????????????????????????????????????????????????????????????????????????????????????????????????????????????????????????????????????????????????????????????????????????????????????????????????????????????????????????????????????????????????????????????????????????????????????????????????????????????????????????????????????????????????????????????????????????????????????????????????????????????????????????????????????????????????????????????????????????????????????????????????????????????????????????????????????????????????????????????????

			>Dyckia_ferruginea
			?????????????????????????????????????????????????????????????????????????????????????????????????????????????????????????????????????????????????????????????????????????????????????????????????????????????????????????????????????????????????????????????????????????????????????????????????????????????????????????????????????????????????????????????????????????????????????????????????????????????????????????????????????????????????????????????????????????????????????????????????????????????????????????????????????????????????????????????????????????????????????????????????????

			>Dyckia_floribunda
			?????????????????????????????????????????????????????????????????????????????????????????????????????????????????????????????????????????????????????????????????????????????????????????????????????????????????????????????????????????????????????????????????????????????????????????????????????????????????????????????????????????????????????????????????????????????????????????????????????????????????????????????????????????????????????????????????????????????????????????????????????????????????????????????????????????????????????????????????????????????????????????????????????

			>Dyckia_fosteriana
			?????????????????????????????????????????????????????????????????????????????????????????????????????????????????????????????????????????????????????????????????????????????????????????????????????????????????????????????????????????????????????????????????????????????????????????????????????????????????????????????????????????????????????????????????????????????????????????????????????????????????????????????????????????????????????????????????????????????????????????????????????????????????????????????????????????????????????????????????????????????????????????????????????

			>Dyckia_goehringii
			?????????????????????????????????????????????????????????????????????????????????????????????????????????????????????????????????????????????????????????????????????????????????????????????????????????????????????????????????????????????????????????????????????????????????????????????????????????????????????????????????????????????????????????????????????????????????????????????????????????????????????????????????????????????????????????????????????????????????????????????????????????????????????????????????????????????????????????????????????????????????????????????????????

			>Dyckia_gouveiana
			?????????????????????????????????????????????????????????????????????????????????????????????????????????????????????????????????????????????????????????????????????????????????????????????????????????????????????????????????????????????????????????????????????????????????????????????????????????????????????????????????????????????????????????????????????????????????????????????????????????????????????????????????????????????????????????????????????????????????????????????????????????????????????????????????????????????????????????????????????????????????????????????????????

			>Dyckia_grandidentata
			?????????????????????????????????????????????????????????????????????????????????????????????????????????????????????????????????????????????????????????????????????????????????????????????????????????????????????????????????????????????????????????????????????????????????????????????????????????????????????????????????????????????????????????????????????????????????????????????????????????????????????????????????????????????????????????????????????????????????????????????????????????????????????????????????????????????????????????????????????????????????????????????????????

			>Dyckia_granmogulensis
			?????????????????????????????????????????????????????????????????????????????????????????????????????????????????????????????????????????????????????????????????????????????????????????????????????????????????????????????????????????????????????????????????????????????????????????????????????????????????????????????????????????????????????????????????????????????????????????????????????????????????????????????????????????????????????????????????????????????????????????????????????????????????????????????????????????????????????????????????????????????????????????????????????

			>Dyckia_hebdingii
			?????????????????????????????????????????????????????????????????????????????????????????????????????????????????????????????????????????????????????????????????????????????????????????????????????????????????????????????????????????????????????????????????????????????????????????????????????????????????????????????????????????????????????????????????????????????????????????????????????????????????????????????????????????????????????????????????????????????????????????????????????????????????????????????????????????????????????????????????????????????????????????????????????

			>Dyckia_hohenbergioides
			?????????????????????????????????????????????????????????????????????????????????????????????????????????????????????????????????????????????????????????????????????????????????????????????????????????????????????????????????????????????????????????????????????????????????????????????????????????????????????????????????????????????????????????????????????????????????????????????????????????????????????????????????????????????????????????????????????????????????????????????????????????????????????????????????????????????????????????????????????????????????????????????????????

			>Dyckia_ibiramensis
			?????????????????????????????????????????????????????????????????????????????????????????????????????????????????????????????????????????????????????????????????????????????????????????????????????????????????????????????????????????????????????????????????????????????????????????????????????????????????????????????????????????????????????????????????????????????????????????????????????????????????????????????????????????????????????????????????????????????????????????????????????????????????????????????????????????????????????????????????????????????????????????????????????

			>Dyckia_jonesiana
			?????????????????????????????????????????????????????????????????????????????????????????????????????????????????????????????????????????????????????????????????????????????????????????????????????????????????????????????????????????????????????????????????????????????????????????????????????????????????????????????????????????????????????????????????????????????????????????????????????????????????????????????????????????????????????????????????????????????????????????????????????????????????????????????????????????????????????????????????????????????????????????????????????

			>Dyckia_kranziana
			?????????????????????????????????????????????????????????????????????????????????????????????????????????????????????????????????????????????????????????????????????????????????????????????????????????????????????????????????????????????????????????????????????????????????????????????????????????????????????????????????????????????????????????????????????????????????????????????????????????????????????????????????????????????????????????????????????????????????????????????????????????????????????????????????????????????????????????????????????????????????????????????????????

			>Dyckia_leptostachya
			?????????????????????????????????????????????????????????????????????????????????????????????????????????????????????????????????????????????????????????????????????????????????????????????????????????????????????????????????????????????????????????????????????????????????????????????????????????????????????????????????????????????????????????????????????????????????????????????????????????????????????????????????????????????????????????????????????????????????????????????????????????????????????????????????????????????????????????????????????????????????????????????????????

			>Dyckia_lindevaldae
			?????????????????????????????????????????????????????????????????????????????????????????????????????????????????????????????????????????????????????????????????????????????????????????????????????????????????????????????????????????????????????????????????????????????????????????????????????????????????????????????????????????????????????????????????????????????????????????????????????????????????????????????????????????????????????????????????????????????????????????????????????????????????????????????????????????????????????????????????????????????????????????????????????

			>Dyckia_lunaris
			?????????????????????????????????????????????????????????????????????????????????????????????????????????????????????????????????????????????????????????????????????????????????????????????????????????????????????????????????????????????????????????????????????????????????????????????????????????????????????????????????????????????????????????????????????????????????????????????????????????????????????????????????????????????????????????????????????????????????????????????????????????????????????????????????????????????????????????????????????????????????????????????????????

			>Dyckia_macedoi
			?????????????????????????????????????????????????????????????????????????????????????????????????????????????????????????????????????????????????????????????????????????????????????????????????????????????????????????????????????????????????????????????????????????????????????????????????????????????????????????????????????????????????????????????????????????????????????????????????????????????????????????????????????????????????????????????????????????????????????????????????????????????????????????????????????????????????????????????????????????????????????????????????????

			>Dyckia_machrisiana
			?????????????????????????????????????????????????????????????????????????????????????????????????????????????????????????????????????????????????????????????????????????????????????????????????????????????????????????????????????????????????????????????????????????????????????????????????????????????????????????????????????????????????????????????????????????????????????????????????????????????????????????????????????????????????????????????????????????????????????????????????????????????????????????????????????????????????????????????????????????????????????????????????????

			>Dyckia_maracasensis
			?????????????????????????????????????????????????????????????????????????????????????????????????????????????????????????????????????????????????????????????????????????????????????????????????????????????????????????????????????????????????????????????????????????????????????????????????????????????????????????????????????????????????????????????????????????????????????????????????????????????????????????????????????????????????????????????????????????????????????????????????????????????????????????????????????????????????????????????????????????????????????????????????????

			>Dyckia_maritima
			?????????????????????????????????????????????????????????????????????????????????????????????????????????????????????????????????????????????????????????????????????????????????????????????????????????????????????????????????????????????????????????????????????????????????????????????????????????????????????????????????????????????????????????????????????????????????????????????????????????????????????????????????????????????????????????????????????????????????????????????????????????????????????????????????????????????????????????????????????????????????????????????????????

			>Dyckia_marnier-lapostollei
			?????????????????????????????????????????????????????????????????????????????????????????????????????????????????????????????????????????????????????????????????????????????????????????????????????????????????????????????????????????????????????????????????????????????????????????????????????????????????????????????????????????????????????????????????????????????????????????????????????????????????????????????????????????????????????????????????????????????????????????????????????????????????????????????????????????????????????????????????????????????????????????????????????

			>Dyckia_marnier-lapostollei_var._estevesii
			?????????????????????????????????????????????????????????????????????????????????????????????????????????????????????????????????????????????????????????????????????????????????????????????????????????????????????????????????????????????????????????????????????????????????????????????????????????????????????????????????????????????????????????????????????????????????????????????????????????????????????????????????????????????????????????????????????????????????????????????????????????????????????????????????????????????????????????????????????????????????????????????????????

			>Dyckia_martinellii
			?????????????????????????????????????????????????????????????????????????????????????????????????????????????????????????????????????????????????????????????????????????????????????????????????????????????????????????????????????????????????????????????????????????????????????????????????????????????????????????????????????????????????????????????????????????????????????????????????????????????????????????????????????????????????????????????????????????????????????????????????????????????????????????????????????????????????????????????????????????????????????????????????????

			>Dyckia_mellobarretoi
			?????????????????????????????????????????????????????????????????????????????????????????????????????????????????????????????????????????????????????????????????????????????????????????????????????????????????????????????????????????????????????????????????????????????????????????????????????????????????????????????????????????????????????????????????????????????????????????????????????????????????????????????????????????????????????????????????????????????????????????????????????????????????????????????????????????????????????????????????????????????????????????????????????

			>Dyckia_microcalyx
			?????????????????????????????????????????????????????????????????????????????????????????????????????????????????????????????????????????????????????????????????????????????????????????????????????????????????????????????????????????????????????????????????????????????????????????????????????????????????????????????????????????????????????????????????????????????????????????????????????????????????????????????????????????????????????????????????????????????????????????????????????????????????????????????????????????????????????????????????????????????????????????????????????

			>Dyckia_microcalyx_var._ostenii
			?????????????????????????????????????????????????????????????????????????????????????????????????????????????????????????????????????????????????????????????????????????????????????????????????????????????????????????????????????????????????????????????????????????????????????????????????????????????????????????????????????????????????????????????????????????????????????????????????????????????????????????????????????????????????????????????????????????????????????????????????????????????????????????????????????????????????????????????????????????????????????????????????????

			>Dyckia_milagrensis
			?????????????????????????????????????????????????????????????????????????????????????????????????????????????????????????????????????????????????????????????????????????????????????????????????????????????????????????????????????????????????????????????????????????????????????????????????????????????????????????????????????????????????????????????????????????????????????????????????????????????????????????????????????????????????????????????????????????????????????????????????????????????????????????????????????????????????????????????????????????????????????????????????????

			>Dyckia_mirandiana
			?????????????????????????????????????????????????????????????????????????????????????????????????????????????????????????????????????????????????????????????????????????????????????????????????????????????????????????????????????????????????????????????????????????????????????????????????????????????????????????????????????????????????????????????????????????????????????????????????????????????????????????????????????????????????????????????????????????????????????????????????????????????????????????????????????????????????????????????????????????????????????????????????????

			>Dyckia_montezumensis
			?????????????????????????????????????????????????????????????????????????????????????????????????????????????????????????????????????????????????????????????????????????????????????????????????????????????????????????????????????????????????????????????????????????????????????????????????????????????????????????????????????????????????????????????????????????????????????????????????????????????????????????????????????????????????????????????????????????????????????????????????????????????????????????????????????????????????????????????????????????????????????????????????????

			>Dyckia_monticola
			?????????????????????????????????????????????????????????????????????????????????????????????????????????????????????????????????????????????????????????????????????????????????????????????????????????????????????????????????????????????????????????????????????????????????????????????????????????????????????????????????????????????????????????????????????????????????????????????????????????????????????????????????????????????????????????????????????????????????????????????????????????????????????????????????????????????????????????????????????????????????????????????????????

			>Dyckia_nana
			?????????????????????????????????????????????????????????????????????????????????????????????????????????????????????????????????????????????????????????????????????????????????????????????????????????????????????????????????????????????????????????????????????????????????????????????????????????????????????????????????????????????????????????????????????????????????????????????????????????????????????????????????????????????????????????????????????????????????????????????????????????????????????????????????????????????????????????????????????????????????????????????????????

			>Dyckia_niederleinii
			?????????????????????????????????????????????????????????????????????????????????????????????????????????????????????????????????????????????????????????????????????????????????????????????????????????????????????????????????????????????????????????????????????????????????????????????????????????????????????????????????????????????????????????????????????????????????????????????????????????????????????????????????????????????????????????????????????????????????????????????????????????????????????????????????????????????????????????????????????????????????????????????????????

			>Dyckia_oligantha
			?????????????????????????????????????????????????????????????????????????????????????????????????????????????????????????????????????????????????????????????????????????????????????????????????????????????????????????????????????????????????????????????????????????????????????????????????????????????????????????????????????????????????????????????????????????????????????????????????????????????????????????????????????????????????????????????????????????????????????????????????????????????????????????????????????????????????????????????????????????????????????????????????????

			>Dyckia_paraensis
			?????????????????????????????????????????????????????????????????????????????????????????????????????????????????????????????????????????????????????????????????????????????????????????????????????????????????????????????????????????????????????????????????????????????????????????????????????????????????????????????????????????????????????????????????????????????????????????????????????????????????????????????????????????????????????????????????????????????????????????????????????????????????????????????????????????????????????????????????????????????????????????????????????

			>Dyckia_pectinata
			?????????????????????????????????????????????????????????????????????????????????????????????????????????????????????????????????????????????????????????????????????????????????????????????????????????????????????????????????????????????????????????????????????????????????????????????????????????????????????????????????????????????????????????????????????????????????????????????????????????????????????????????????????????????????????????????????????????????????????????????????????????????????????????????????????????????????????????????????????????????????????????????????????

			>Dyckia_pernambucana
			?????????????????????????????????????????????????????????????????????????????????????????????????????????????????????????????????????????????????????????????????????????????????????????????????????????????????????????????????????????????????????????????????????????????????????????????????????????????????????????????????????????????????????????????????????????????????????????????????????????????????????????????????????????????????????????????????????????????????????????????????????????????????????????????????????????????????????????????????????????????????????????????????????

			>Dyckia_platyphylla
			?????????????????????????????????????????????????????????????????????????????????????????????????????????????????????????????????????????????????????????????????????????????????????????????????????????????????????????????????????????????????????????????????????????????????????????????????????????????????????????????????????????????????????????????????????????????????????????????????????????????????????????????????????????????????????????????????????????????????????????????????????????????????????????????????????????????????????????????????????????????????????????????????????

			>Dyckia_pottiorum
			?????????????????????????????????????????????????????????????????????????????????????????????????????????????????????????????????????????????????????????????????????????????????????????????????????????????????????????????????????????????????????????????????????????????????????????????????????????????????????????????????????????????????????????????????????????????????????????????????????????????????????????????????????????????????????????????????????????????????????????????????????????????????????????????????????????????????????????????????????????????????????????????????????

			>Dyckia_pseudococcinea
			?????????????????????????????????????????????????????????????????????????????????????????????????????????????????????????????????????????????????????????????????????????????????????????????????????????????????????????????????????????????????????????????????????????????????????????????????????????????????????????????????????????????????????????????????????????????????????????????????????????????????????????????????????????????????????????????????????????????????????????????????????????????????????????????????????????????????????????????????????????????????????????????????????

			>Dyckia_pulquinensis
			?????????????????????????????????????????????????????????????????????????????????????????????????????????????????????????????????????????????????????????????????????????????????????????????????????????????????????????????????????????????????????????????????????????????????????????????????????????????????????????????????????????????????????????????????????????????????????????????????????????????????????????????????????????????????????????????????????????????????????????????????????????????????????????????????????????????????????????????????????????????????????????????????????

			>Dyckia_pumila
			?????????????????????????????????????????????????????????????????????????????????????????????????????????????????????????????????????????????????????????????????????????????????????????????????????????????????????????????????????????????????????????????????????????????????????????????????????????????????????????????????????????????????????????????????????????????????????????????????????????????????????????????????????????????????????????????????????????????????????????????????????????????????????????????????????????????????????????????????????????????????????????????????????

			>Dyckia_rariflora
			?????????????????????????????????????????????????????????????????????????????????????????????????????????????????????????????????????????????????????????????????????????????????????????????????????????????????????????????????????????????????????????????????????????????????????????????????????????????????????????????????????????????????????????????????????????????????????????????????????????????????????????????????????????????????????????????????????????????????????????????????????????????????????????????????????????????????????????????????????????????????????????????????????

			>Dyckia_remotiflora
			?????????????????????????????????????????????????????????????????????????????????????????????????????????????????????????????????????????????????????????????????????????????????????????????????????????????????????????????????????????????????????????????????????????????????????????????????????????????????????????????????????????????????????????????????????????????????????????????????????????????????????????????????????????????????????????????????????????????????????????????????????????????????????????????????????????????????????????????????????????????????????????????????????

			>Dyckia_rondonopolitana
			?????????????????????????????????????????????????????????????????????????????????????????????????????????????????????????????????????????????????????????????????????????????????????????????????????????????????????????????????????????????????????????????????????????????????????????????????????????????????????????????????????????????????????????????????????????????????????????????????????????????????????????????????????????????????????????????????????????????????????????????????????????????????????????????????????????????????????????????????????????????????????????????????????

			>Dyckia_saxatilis
			?????????????????????????????????????????????????????????????????????????????????????????????????????????????????????????????????????????????????????????????????????????????????????????????????????????????????????????????????????????????????????????????????????????????????????????????????????????????????????????????????????????????????????????????????????????????????????????????????????????????????????????????????????????????????????????????????????????????????????????????????????????????????????????????????????????????????????????????????????????????????????????????????????

			>Dyckia_secunda
			?????????????????????????????????????????????????????????????????????????????????????????????????????????????????????????????????????????????????????????????????????????????????????????????????????????????????????????????????????????????????????????????????????????????????????????????????????????????????????????????????????????????????????????????????????????????????????????????????????????????????????????????????????????????????????????????????????????????????????????????????????????????????????????????????????????????????????????????????????????????????????????????????????

			>Dyckia_selloa
			?????????????????????????????????????????????????????????????????????????????????????????????????????????????????????????????????????????????????????????????????????????????????????????????????????????????????????????????????????????????????????????????????????????????????????????????????????????????????????????????????????????????????????????????????????????????????????????????????????????????????????????????????????????????????????????????????????????????????????????????????????????????????????????????????????????????????????????????????????????????????????????????????????

			>Dyckia_tenebrosa
			?????????????????????????????????????????????????????????????????????????????????????????????????????????????????????????????????????????????????????????????????????????????????????????????????????????????????????????????????????????????????????????????????????????????????????????????????????????????????????????????????????????????????????????????????????????????????????????????????????????????????????????????????????????????????????????????????????????????????????????????????????????????????????????????????????????????????????????????????????????????????????????????????????

			>Dyckia_tobatiensis
			?????????????????????????????????????????????????????????????????????????????????????????????????????????????????????????????????????????????????????????????????????????????????????????????????????????????????????????????????????????????????????????????????????????????????????????????????????????????????????????????????????????????????????????????????????????????????????????????????????????????????????????????????????????????????????????????????????????????????????????????????????????????????????????????????????????????????????????????????????????????????????????????????????

			>Dyckia_tomentella
			?????????????????????????????????????????????????????????????????????????????????????????????????????????????????????????????????????????????????????????????????????????????????????????????????????????????????????????????????????????????????????????????????????????????????????????????????????????????????????????????????????????????????????????????????????????????????????????????????????????????????????????????????????????????????????????????????????????????????????????????????????????????????????????????????????????????????????????????????????????????????????????????????????

			>Dyckia_ursina
			?????????????????????????????????????????????????????????????????????????????????????????????????????????????????????????????????????????????????????????????????????????????????????????????????????????????????????????????????????????????????????????????????????????????????????????????????????????????????????????????????????????????????????????????????????????????????????????????????????????????????????????????????????????????????????????????????????????????????????????????????????????????????????????????????????????????????????????????????????????????????????????????????????

			>Dyckia_velascana
			?????????????????????????????????????????????????????????????????????????????????????????????????????????????????????????????????????????????????????????????????????????????????????????????????????????????????????????????????????????????????????????????????????????????????????????????????????????????????????????????????????????????????????????????????????????????????????????????????????????????????????????????????????????????????????????????????????????????????????????????????????????????????????????????????????????????????????????????????????????????????????????????????????

			>Dyckia_vestita
			?????????????????????????????????????????????????????????????????????????????????????????????????????????????????????????????????????????????????????????????????????????????????????????????????????????????????????????????????????????????????????????????????????????????????????????????????????????????????????????????????????????????????????????????????????????????????????????????????????????????????????????????????????????????????????????????????????????????????????????????????????????????????????????????????????????????????????????????????????????????????????????????????????

			>Dyckia_vicentensis
			?????????????????????????????????????????????????????????????????????????????????????????????????????????????????????????????????????????????????????????????????????????????????????????????????????????????????????????????????????????????????????????????????????????????????????????????????????????????????????????????????????????????????????????????????????????????????????????????????????????????????????????????????????????????????????????????????????????????????????????????????????????????????????????????????????????????????????????????????????????????????????????????????????

			>Dyckia_walteriana
			?????????????????????????????????????????????????????????????????????????????????????????????????????????????????????????????????????????????????????????????????????????????????????????????????????????????????????????????????????????????????????????????????????????????????????????????????????????????????????????????????????????????????????????????????????????????????????????????????????????????????????????????????????????????????????????????????????????????????????????????????????????????????????????????????????????????????????????????????????????????????????????????????????

			>Edmundoa_ambigua
			?????????????????????????????????????????????????????????????????????????????????????????????????????????????????????????????????????????????????????????????????????????????????????????????????????????????????????????????????????????????????????????????????????????????????????????????????????????????????????????????????????????????????????????????????????????????????????????????????????????????????????????????????????????????????????????????????????????????????????????????????????????????????????????????????????????????????????????????????????????????????????????????????????

			>Edmundoa_lindenii
			?????????????????????????????????????????????????????????????????????????????????????????????????????????????????????????????????????????????????????????????????????????????????????????????????????????????????????????????????????????????????????????????????????????????????????????????????????????????????????????????????????????????????????????????????????????????????????????????????????????????????????????????????????????????????????????????????????????????????????????????????????????????????????????????????????????????????????????????????????????????????????????????????????

			>Edmundoa_lindenii_var._rosaea
			?????????????????????????????????????????????????????????????????????????????????????????????????????????????????????????????????????????????????????????????????????????????????????????????????????????????????????????????????????????????????????????????????????????????????????????????????????????????????????????????????????????????????????????????????????????????????????????????????????????????????????????????????????????????????????????????????????????????????????????????????????????????????????????????????????????????????????????????????????????????????????????????????????

			>Edmundoa_perplexa
			?????????????????????????????????????????????????????????????????????????????????????????????????????????????????????????????????????????????????????????????????????????????????????????????????????????????????????????????????????????????????????????????????????????????????????????????????????????????????????????????????????????????????????????????????????????????????????????????????????????????????????????????????????????????????????????????????????????????????????????????????????????????????????????????????????????????????????????????????????????????????????????????????????

			>Eduandrea_selloana
			?????????????????????????????????????????????????????????????????????????????????????????????????????????????????????????????????????????????????????????????????????????????????????????????????????????????????????????????????????????????????????????????????????????????????????????????????????????????????????????????????????????????????????????????????????????????????????????????????????????????????????????????????????????????????????????????????????????????????????????????????????????????????????????????????????????????????????????????????????????????????????????????????????

			>Encholirium_agavoides
			?????????????????????????????????????????????????????????????????????????????????????????????????????????????????????????????????????????????????????????????????????????????????????????????????????????????????????????????????????????????????????????????????????????????????????????????????????????????????????????????????????????????????????????????????????????????????????????????????????????????????????????????????????????????????????????????????????????????????????????????????????????????????????????????????????????????????????????????????????????????????????????????????????

			>Encholirium_biflorum
			?????????????????????????????????????????????????????????????????????????????????????????????????????????????????????????????????????????????????????????????????????????????????????????????????????????????????????????????????????????????????????????????????????????????????????????????????????????????????????????????????????????????????????????????????????????????????????????????????????????????????????????????????????????????????????????????????????????????????????????????????????????????????????????????????????????????????????????????????????????????????????????????????????

			>Encholirium_brachypodum
			?????????????????????????????????????????????????????????????????????????????????????????????????????????????????????????????????????????????????????????????????????????????????????????????????????????????????????????????????????????????????????????????????????????????????????????????????????????????????????????????????????????????????????????????????????????????????????????????????????????????????????????????????????????????????????????????????????????????????????????????????????????????????????????????????????????????????????????????????????????????????????????????????????

			>Encholirium_ctenophyllum
			?????????????????????????????????????????????????????????????????????????????????????????????????????????????????????????????????????????????????????????????????????????????????????????????????????????????????????????????????????????????????????????????????????????????????????????????????????????????????????????????????????????????????????????????????????????????????????????????????????????????????????????????????????????????????????????????????????????????????????????????????????????????????????????????????????????????????????????????????????????????????????????????????????

			>Encholirium_diamantinum
			?????????????????????????????????????????????????????????????????????????????????????????????????????????????????????????????????????????????????????????????????????????????????????????????????????????????????????????????????????????????????????????????????????????????????????????????????????????????????????????????????????????????????????????????????????????????????????????????????????????????????????????????????????????????????????????????????????????????????????????????????????????????????????????????????????????????????????????????????????????????????????????????????????

			>Encholirium_eddie-estevesii
			?????????????????????????????????????????????????????????????????????????????????????????????????????????????????????????????????????????????????????????????????????????????????????????????????????????????????????????????????????????????????????????????????????????????????????????????????????????????????????????????????????????????????????????????????????????????????????????????????????????????????????????????????????????????????????????????????????????????????????????????????????????????????????????????????????????????????????????????????????????????????????????????????????

			>Encholirium_erectiflorum
			?????????????????????????????????????????????????????????????????????????????????????????????????????????????????????????????????????????????????????????????????????????????????????????????????????????????????????????????????????????????????????????????????????????????????????????????????????????????????????????????????????????????????????????????????????????????????????????????????????????????????????????????????????????????????????????????????????????????????????????????????????????????????????????????????????????????????????????????????????????????????????????????????????

			>Encholirium_fragae
			?????????????????????????????????????????????????????????????????????????????????????????????????????????????????????????????????????????????????????????????????????????????????????????????????????????????????????????????????????????????????????????????????????????????????????????????????????????????????????????????????????????????????????????????????????????????????????????????????????????????????????????????????????????????????????????????????????????????????????????????????????????????????????????????????????????????????????????????????????????????????????????????????????

			>Encholirium_gracile
			?????????????????????????????????????????????????????????????????????????????????????????????????????????????????????????????????????????????????????????????????????????????????????????????????????????????????????????????????????????????????????????????????????????????????????????????????????????????????????????????????????????????????????????????????????????????????????????????????????????????????????????????????????????????????????????????????????????????????????????????????????????????????????????????????????????????????????????????????????????????????????????????????????

			>Encholirium_heloisae
			?????????????????????????????????????????????????????????????????????????????????????????????????????????????????????????????????????????????????????????????????????????????????????????????????????????????????????????????????????????????????????????????????????????????????????????????????????????????????????????????????????????????????????????????????????????????????????????????????????????????????????????????????????????????????????????????????????????????????????????????????????????????????????????????????????????????????????????????????????????????????????????????????????

			>Encholirium_horridum
			?????????????????????????????????????????????????????????????????????????????????????????????????????????????????????????????????????????????????????????????????????????????????????????????????????????????????????????????????????????????????????????????????????????????????????????????????????????????????????????????????????????????????????????????????????????????????????????????????????????????????????????????????????????????????????????????????????????????????????????????????????????????????????????????????????????????????????????????????????????????????????????????????????

			>Encholirium_irwinii
			?????????????????????????????????????????????????????????????????????????????????????????????????????????????????????????????????????????????????????????????????????????????????????????????????????????????????????????????????????????????????????????????????????????????????????????????????????????????????????????????????????????????????????????????????????????????????????????????????????????????????????????????????????????????????????????????????????????????????????????????????????????????????????????????????????????????????????????????????????????????????????????????????????

			>Encholirium_kranzianum
			?????????????????????????????????????????????????????????????????????????????????????????????????????????????????????????????????????????????????????????????????????????????????????????????????????????????????????????????????????????????????????????????????????????????????????????????????????????????????????????????????????????????????????????????????????????????????????????????????????????????????????????????????????????????????????????????????????????????????????????????????????????????????????????????????????????????????????????????????????????????????????????????????????

			>Encholirium_longiflorum
			?????????????????????????????????????????????????????????????????????????????????????????????????????????????????????????????????????????????????????????????????????????????????????????????????????????????????????????????????????????????????????????????????????????????????????????????????????????????????????????????????????????????????????????????????????????????????????????????????????????????????????????????????????????????????????????????????????????????????????????????????????????????????????????????????????????????????????????????????????????????????????????????????????

			>Encholirium_luxor
			?????????????????????????????????????????????????????????????????????????????????????????????????????????????????????????????????????????????????????????????????????????????????????????????????????????????????????????????????????????????????????????????????????????????????????????????????????????????????????????????????????????????????????????????????????????????????????????????????????????????????????????????????????????????????????????????????????????????????????????????????????????????????????????????????????????????????????????????????????????????????????????????????????

			>Encholirium_lymanianum
			?????????????????????????????????????????????????????????????????????????????????????????????????????????????????????????????????????????????????????????????????????????????????????????????????????????????????????????????????????????????????????????????????????????????????????????????????????????????????????????????????????????????????????????????????????????????????????????????????????????????????????????????????????????????????????????????????????????????????????????????????????????????????????????????????????????????????????????????????????????????????????????????????????

			>Encholirium_magalhaesii
			?????????????????????????????????????????????????????????????????????????????????????????????????????????????????????????????????????????????????????????????????????????????????????????????????????????????????????????????????????????????????????????????????????????????????????????????????????????????????????????????????????????????????????????????????????????????????????????????????????????????????????????????????????????????????????????????????????????????????????????????????????????????????????????????????????????????????????????????????????????????????????????????????????

			>Encholirium_maximum
			?????????????????????????????????????????????????????????????????????????????????????????????????????????????????????????????????????????????????????????????????????????????????????????????????????????????????????????????????????????????????????????????????????????????????????????????????????????????????????????????????????????????????????????????????????????????????????????????????????????????????????????????????????????????????????????????????????????????????????????????????????????????????????????????????????????????????????????????????????????????????????????????????????

			>Encholirium_pedicellatum
			?????????????????????????????????????????????????????????????????????????????????????????????????????????????????????????????????????????????????????????????????????????????????????????????????????????????????????????????????????????????????????????????????????????????????????????????????????????????????????????????????????????????????????????????????????????????????????????????????????????????????????????????????????????????????????????????????????????????????????????????????????????????????????????????????????????????????????????????????????????????????????????????????????

			>Encholirium_pulchrum
			?????????????????????????????????????????????????????????????????????????????????????????????????????????????????????????????????????????????????????????????????????????????????????????????????????????????????????????????????????????????????????????????????????????????????????????????????????????????????????????????????????????????????????????????????????????????????????????????????????????????????????????????????????????????????????????????????????????????????????????????????????????????????????????????????????????????????????????????????????????????????????????????????????

			>Encholirium_scrutor
			?????????????????????????????????????????????????????????????????????????????????????????????????????????????????????????????????????????????????????????????????????????????????????????????????????????????????????????????????????????????????????????????????????????????????????????????????????????????????????????????????????????????????????????????????????????????????????????????????????????????????????????????????????????????????????????????????????????????????????????????????????????????????????????????????????????????????????????????????????????????????????????????????????

			>Encholirium_spectabile
			?????????????????????????????????????????????????????????????????????????????????????????????????????????????????????????????????????????????????????????????????????????????????????????????????????????????????????????????????????????????????????????????????????????????????????????????????????????????????????????????????????????????????????????????????????????????????????????????????????????????????????????????????????????????????????????????????????????????????????????????????????????????????????????????????????????????????????????????????????????????????????????????????????

			>Encholirium_splendidum
			?????????????????????????????????????????????????????????????????????????????????????????????????????????????????????????????????????????????????????????????????????????????????????????????????????????????????????????????????????????????????????????????????????????????????????????????????????????????????????????????????????????????????????????????????????????????????????????????????????????????????????????????????????????????????????????????????????????????????????????????????????????????????????????????????????????????????????????????????????????????????????????????????????

			>Encholirium_structor
			?????????????????????????????????????????????????????????????????????????????????????????????????????????????????????????????????????????????????????????????????????????????????????????????????????????????????????????????????????????????????????????????????????????????????????????????????????????????????????????????????????????????????????????????????????????????????????????????????????????????????????????????????????????????????????????????????????????????????????????????????????????????????????????????????????????????????????????????????????????????????????????????????????

			>Encholirium_subsecundum
			?????????????????????????????????????????????????????????????????????????????????????????????????????????????????????????????????????????????????????????????????????????????????????????????????????????????????????????????????????????????????????????????????????????????????????????????????????????????????????????????????????????????????????????????????????????????????????????????????????????????????????????????????????????????????????????????????????????????????????????????????????????????????????????????????????????????????????????????????????????????????????????????????????

			>Encholirium_vogelii
			?????????????????????????????????????????????????????????????????????????????????????????????????????????????????????????????????????????????????????????????????????????????????????????????????????????????????????????????????????????????????????????????????????????????????????????????????????????????????????????????????????????????????????????????????????????????????????????????????????????????????????????????????????????????????????????????????????????????????????????????????????????????????????????????????????????????????????????????????????????????????????????????????????

			>Epidryos_allenii
			?????????????????????????????????????????????????????????????????????????????????????????????????????????????????????????????????????????????????????????????????????????????????????????????????????????????????????????????????????????????????????????????????????????????????????????????????????????????????????????????????????????????????????????????????????????????????????????????????????????????????????????????????????????????????????????????????????????????????????????????????????????????????????????????????????????????????????????????????????????????????????????????????????

			>Fascicularia_bicolor
			?????????????????????????????????????????????????????????????????????????????????????????????????????????????????????????????????????????????????????????????????????????????????????????????????????????????????????????????????????????????????????????????????????????????????????????????????????????????????????????????????????????????????????????????????????????????????????????????????????????????????????????????????????????????????????????????????????????????????????????????????????????????????????????????????????????????????????????????????????????????????????????????????????

			>Fernseea_itatiaiae
			?????????????????????????????????????????????????????????????????????????????????????????????????????????????????????????????????????????????????????????????????????????????????????????????????????????????????????????????????????????????????????????????????????????????????????????????????????????????????????????????????????????????????????????????????????????????????????????????????????????????????????????????????????????????????????????????????????????????????????????????????????????????????????????????????????????????????????????????????????????????????????????????????????

			>Forzzaea_leopoldo-horstii
			?????????????????????????????????????????????????????????????????????????????????????????????????????????????????????????????????????????????????????????????????????????????????????????????????????????????????????????????????????????????????????????????????????????????????????????????????????????????????????????????????????????????????????????????????????????????????????????????????????????????????????????????????????????????????????????????????????????????????????????????????????????????????????????????????????????????????????????????????????????????????????????????????????

			>Forzzaea_warasii
			?????????????????????????????????????????????????????????????????????????????????????????????????????????????????????????????????????????????????????????????????????????????????????????????????????????????????????????????????????????????????????????????????????????????????????????????????????????????????????????????????????????????????????????????????????????????????????????????????????????????????????????????????????????????????????????????????????????????????????????????????????????????????????????????????????????????????????????????????????????????????????????????????????

			>Fosterella_albicans
			?????????????????????????????????????????????????????????????????????????????????????????????????????????????????????????????????????????????????????????????????????????????????????????????????????????????????????????????????????????????????????????????????????????????????????????????????????????????????????????????????????????????????????????????????????????????????????????????????????????????????????????????????????????????????????????????????????????????????????????????????????????????????????????????????????????????????????????????????????????????????????????????????????

			>Fosterella_batistana
			?????????????????????????????????????????????????????????????????????????????????????????????????????????????????????????????????????????????????????????????????????????????????????????????????????????????????????????????????????????????????????????????????????????????????????????????????????????????????????????????????????????????????????????????????????????????????????????????????????????????????????????????????????????????????????????????????????????????????????????????????????????????????????????????????????????????????????????????????????????????????????????????????????

			>Fosterella_caulescens
			?????????????????????????????????????????????????????????????????????????????????????????????????????????????????????????????????????????????????????????????????????????????????????????????????????????????????????????????????????????????????????????????????????????????????????????????????????????????????????????????????????????????????????????????????????????????????????????????????????????????????????????????????????????????????????????????????????????????????????????????????????????????????????????????????????????????????????????????????????????????????????????????????????

			>Fosterella_christophii
			?????????????????????????????????????????????????????????????????????????????????????????????????????????????????????????????????????????????????????????????????????????????????????????????????????????????????????????????????????????????????????????????????????????????????????????????????????????????????????????????????????????????????????????????????????????????????????????????????????????????????????????????????????????????????????????????????????????????????????????????????????????????????????????????????????????????????????????????????????????????????????????????????????

			>Fosterella_cotacajensis
			?????????????????????????????????????????????????????????????????????????????????????????????????????????????????????????????????????????????????????????????????????????????????????????????????????????????????????????????????????????????????????????????????????????????????????????????????????????????????????????????????????????????????????????????????????????????????????????????????????????????????????????????????????????????????????????????????????????????????????????????????????????????????????????????????????????????????????????????????????????????????????????????????????

			>Fosterella_floridensis
			?????????????????????????????????????????????????????????????????????????????????????????????????????????????????????????????????????????????????????????????????????????????????????????????????????????????????????????????????????????????????????????????????????????????????????????????????????????????????????????????????????????????????????????????????????????????????????????????????????????????????????????????????????????????????????????????????????????????????????????????????????????????????????????????????????????????????????????????????????????????????????????????????????

			>Fosterella_gracilis
			?????????????????????????????????????????????????????????????????????????????????????????????????????????????????????????????????????????????????????????????????????????????????????????????????????????????????????????????????????????????????????????????????????????????????????????????????????????????????????????????????????????????????????????????????????????????????????????????????????????????????????????????????????????????????????????????????????????????????????????????????????????????????????????????????????????????????????????????????????????????????????????????????????

			>Fosterella_graminea
			?????????????????????????????????????????????????????????????????????????????????????????????????????????????????????????????????????????????????????????????????????????????????????????????????????????????????????????????????????????????????????????????????????????????????????????????????????????????????????????????????????????????????????????????????????????????????????????????????????????????????????????????????????????????????????????????????????????????????????????????????????????????????????????????????????????????????????????????????????????????????????????????????????

			>Fosterella_hatschbachii
			?????????????????????????????????????????????????????????????????????????????????????????????????????????????????????????????????????????????????????????????????????????????????????????????????????????????????????????????????????????????????????????????????????????????????????????????????????????????????????????????????????????????????????????????????????????????????????????????????????????????????????????????????????????????????????????????????????????????????????????????????????????????????????????????????????????????????????????????????????????????????????????????????????

			>Fosterella_heterophylla
			?????????????????????????????????????????????????????????????????????????????????????????????????????????????????????????????????????????????????????????????????????????????????????????????????????????????????????????????????????????????????????????????????????????????????????????????????????????????????????????????????????????????????????????????????????????????????????????????????????????????????????????????????????????????????????????????????????????????????????????????????????????????????????????????????????????????????????????????????????????????????????????????????????

			>Fosterella_kroemeri
			?????????????????????????????????????????????????????????????????????????????????????????????????????????????????????????????????????????????????????????????????????????????????????????????????????????????????????????????????????????????????????????????????????????????????????????????????????????????????????????????????????????????????????????????????????????????????????????????????????????????????????????????????????????????????????????????????????????????????????????????????????????????????????????????????????????????????????????????????????????????????????????????????????

			>Fosterella_micrantha
			?????????????????????????????????????????????????????????????????????????????????????????????????????????????????????????????????????????????????????????????????????????????????????????????????????????????????????????????????????????????????????????????????????????????????????????????????????????????????????????????????????????????????????????????????????????????????????????????????????????????????????????????????????????????????????????????????????????????????????????????????????????????????????????????????????????????????????????????????????????????????????????????????????

			>Fosterella_penduliflora
			?????????????????????????????????????????????????????????????????????????????????????????????????????????????????????????????????????????????????????????????????????????????????????????????????????????????????????????????????????????????????????????????????????????????????????????????????????????????????????????????????????????????????????????????????????????????????????????????????????????????????????????????????????????????????????????????????????????????????????????????????????????????????????????????????????????????????????????????????????????????????????????????????????

			>Fosterella_petiolata
			?????????????????????????????????????????????????????????????????????????????????????????????????????????????????????????????????????????????????????????????????????????????????????????????????????????????????????????????????????????????????????????????????????????????????????????????????????????????????????????????????????????????????????????????????????????????????????????????????????????????????????????????????????????????????????????????????????????????????????????????????????????????????????????????????????????????????????????????????????????????????????????????????????

			>Fosterella_rexiae
			?????????????????????????????????????????????????????????????????????????????????????????????????????????????????????????????????????????????????????????????????????????????????????????????????????????????????????????????????????????????????????????????????????????????????????????????????????????????????????????????????????????????????????????????????????????????????????????????????????????????????????????????????????????????????????????????????????????????????????????????????????????????????????????????????????????????????????????????????????????????????????????????????????

			>Fosterella_robertreadii
			?????????????????????????????????????????????????????????????????????????????????????????????????????????????????????????????????????????????????????????????????????????????????????????????????????????????????????????????????????????????????????????????????????????????????????????????????????????????????????????????????????????????????????????????????????????????????????????????????????????????????????????????????????????????????????????????????????????????????????????????????????????????????????????????????????????????????????????????????????????????????????????????????????

			>Fosterella_rusbyi
			?????????????????????????????????????????????????????????????????????????????????????????????????????????????????????????????????????????????????????????????????????????????????????????????????????????????????????????????????????????????????????????????????????????????????????????????????????????????????????????????????????????????????????????????????????????????????????????????????????????????????????????????????????????????????????????????????????????????????????????????????????????????????????????????????????????????????????????????????????????????????????????????????????

			>Fosterella_schidosperma
			?????????????????????????????????????????????????????????????????????????????????????????????????????????????????????????????????????????????????????????????????????????????????????????????????????????????????????????????????????????????????????????????????????????????????????????????????????????????????????????????????????????????????????????????????????????????????????????????????????????????????????????????????????????????????????????????????????????????????????????????????????????????????????????????????????????????????????????????????????????????????????????????????????

			>Fosterella_spectabilis
			?????????????????????????????????????????????????????????????????????????????????????????????????????????????????????????????????????????????????????????????????????????????????????????????????????????????????????????????????????????????????????????????????????????????????????????????????????????????????????????????????????????????????????????????????????????????????????????????????????????????????????????????????????????????????????????????????????????????????????????????????????????????????????????????????????????????????????????????????????????????????????????????????????

			>Fosterella_vasquezii
			?????????????????????????????????????????????????????????????????????????????????????????????????????????????????????????????????????????????????????????????????????????????????????????????????????????????????????????????????????????????????????????????????????????????????????????????????????????????????????????????????????????????????????????????????????????????????????????????????????????????????????????????????????????????????????????????????????????????????????????????????????????????????????????????????????????????????????????????????????????????????????????????????????

			>Fosterella_villosula
			?????????????????????????????????????????????????????????????????????????????????????????????????????????????????????????????????????????????????????????????????????????????????????????????????????????????????????????????????????????????????????????????????????????????????????????????????????????????????????????????????????????????????????????????????????????????????????????????????????????????????????????????????????????????????????????????????????????????????????????????????????????????????????????????????????????????????????????????????????????????????????????????????????

			>Fosterella_weberbaueri
			?????????????????????????????????????????????????????????????????????????????????????????????????????????????????????????????????????????????????????????????????????????????????????????????????????????????????????????????????????????????????????????????????????????????????????????????????????????????????????????????????????????????????????????????????????????????????????????????????????????????????????????????????????????????????????????????????????????????????????????????????????????????????????????????????????????????????????????????????????????????????????????????????????

			>Fosterella_weddelliana
			?????????????????????????????????????????????????????????????????????????????????????????????????????????????????????????????????????????????????????????????????????????????????????????????????????????????????????????????????????????????????????????????????????????????????????????????????????????????????????????????????????????????????????????????????????????????????????????????????????????????????????????????????????????????????????????????????????????????????????????????????????????????????????????????????????????????????????????????????????????????????????????????????????

			>Fosterella_windischii
			?????????????????????????????????????????????????????????????????????????????????????????????????????????????????????????????????????????????????????????????????????????????????????????????????????????????????????????????????????????????????????????????????????????????????????????????????????????????????????????????????????????????????????????????????????????????????????????????????????????????????????????????????????????????????????????????????????????????????????????????????????????????????????????????????????????????????????????????????????????????????????????????????????

			>Fosterella_yuvinkae
			?????????????????????????????????????????????????????????????????????????????????????????????????????????????????????????????????????????????????????????????????????????????????????????????????????????????????????????????????????????????????????????????????????????????????????????????????????????????????????????????????????????????????????????????????????????????????????????????????????????????????????????????????????????????????????????????????????????????????????????????????????????????????????????????????????????????????????????????????????????????????????????????????????

			>Glomeropitcairnia_erectiflora
			?????????????????????????????????????????????????????????????????????????????????????????????????????????????????????????????????????????????????????????????????????????????????????????????????????????????????????????????????????????????????????????????????????????????????????????????????????????????????????????????????????????????????????????????????????????????????????????????????????????????????????????????????????????????????????????????????????????????????????????????????????????????????????????????????????????????????????????????????????????????????????????????????????

			>Glomeropitcairnia_penduliflora
			?????????????????????????????????????????????????????????????????????????????????????????????????????????????????????????????????????????????????????????????????????????????????????????????????????????????????????????????????????????????????????????????????????????????????????????????????????????????????????????????????????????????????????????????????????????????????????????????????????????????????????????????????????????????????????????????????????????????????????????????????????????????????????????????????????????????????????????????????????????????????????????????????????

			>Goudaea_chrysostachys
			?????????????????????????????????????????????????????????????????????????????????????????????????????????????????????????????????????????????????????????????????????????????????????????????????????????????????????????????????????????????????????????????????????????????????????????????????????????????????????????????????????????????????????????????????????????????????????????????????????????????????????????????????????????????????????????????????????????????????????????????????????????????????????????????????????????????????????????????????????????????????????????????????????

			>Goudaea_ospinae
			?????????????????????????????????????????????????????????????????????????????????????????????????????????????????????????????????????????????????????????????????????????????????????????????????????????????????????????????????????????????????????????????????????????????????????????????????????????????????????????????????????????????????????????????????????????????????????????????????????????????????????????????????????????????????????????????????????????????????????????????????????????????????????????????????????????????????????????????????????????????????????????????????????

			>Goudaea_ospinae_var._gruberi
			?????????????????????????????????????????????????????????????????????????????????????????????????????????????????????????????????????????????????????????????????????????????????????????????????????????????????????????????????????????????????????????????????????????????????????????????????????????????????????????????????????????????????????????????????????????????????????????????????????????????????????????????????????????????????????????????????????????????????????????????????????????????????????????????????????????????????????????????????????????????????????????????????????

			>Gregbrownia_fulgens
			?????????????????????????????????????????????????????????????????????????????????????????????????????????????????????????????????????????????????????????????????????????????????????????????????????????????????????????????????????????????????????????????????????????????????????????????????????????????????????????????????????????????????????????????????????????????????????????????????????????????????????????????????????????????????????????????????????????????????????????????????????????????????????????????????????????????????????????????????????????????????????????????????????

			>Gregbrownia_hutchisonii
			?????????????????????????????????????????????????????????????????????????????????????????????????????????????????????????????????????????????????????????????????????????????????????????????????????????????????????????????????????????????????????????????????????????????????????????????????????????????????????????????????????????????????????????????????????????????????????????????????????????????????????????????????????????????????????????????????????????????????????????????????????????????????????????????????????????????????????????????????????????????????????????????????????

			>Greigia_mulfordii
			?????????????????????????????????????????????????????????????????????????????????????????????????????????????????????????????????????????????????????????????????????????????????????????????????????????????????????????????????????????????????????????????????????????????????????????????????????????????????????????????????????????????????????????????????????????????????????????????????????????????????????????????????????????????????????????????????????????????????????????????????????????????????????????????????????????????????????????????????????????????????????????????????????

			>Greigia_sphacelata
			?????????????????????????????????????????????????????????????????????????????????????????????????????????????????????????????????????????????????????????????????????????????????????????????????????????????????????????????????????????????????????????????????????????????????????????????????????????????????????????????????????????????????????????????????????????????????????????????????????????????????????????????????????????????????????????????????????????????????????????????????????????????????????????????????????????????????????????????????????????????????????????????????????

			>Guzmania_acorifolia
			?????????????????????????????????????????????????????????????????????????????????????????????????????????????????????????????????????????????????????????????????????????????????????????????????????????????????????????????????????????????????????????????????????????????????????????????????????????????????????????????????????????????????????????????????????????????????????????????????????????????????????????????????????????????????????????????????????????????????????????????????????????????????????????????????????????????????????????????????????????????????????????????????????

			>Guzmania_angustifolia
			?????????????????????????????????????????????????????????????????????????????????????????????????????????????????????????????????????????????????????????????????????????????????????????????????????????????????????????????????????????????????????????????????????????????????????????????????????????????????????????????????????????????????????????????????????????????????????????????????????????????????????????????????????????????????????????????????????????????????????????????????????????????????????????????????????????????????????????????????????????????????????????????????????

			>Guzmania_butcheri
			?????????????????????????????????????????????????????????????????????????????????????????????????????????????????????????????????????????????????????????????????????????????????????????????????????????????????????????????????????????????????????????????????????????????????????????????????????????????????????????????????????????????????????????????????????????????????????????????????????????????????????????????????????????????????????????????????????????????????????????????????????????????????????????????????????????????????????????????????????????????????????????????????????

			>Guzmania_condensata
			?????????????????????????????????????????????????????????????????????????????????????????????????????????????????????????????????????????????????????????????????????????????????????????????????????????????????????????????????????????????????????????????????????????????????????????????????????????????????????????????????????????????????????????????????????????????????????????????????????????????????????????????????????????????????????????????????????????????????????????????????????????????????????????????????????????????????????????????????????????????????????????????????????

			>Guzmania_coriostachya
			?????????????????????????????????????????????????????????????????????????????????????????????????????????????????????????????????????????????????????????????????????????????????????????????????????????????????????????????????????????????????????????????????????????????????????????????????????????????????????????????????????????????????????????????????????????????????????????????????????????????????????????????????????????????????????????????????????????????????????????????????????????????????????????????????????????????????????????????????????????????????????????????????????

			>Guzmania_donnellsmithii
			?????????????????????????????????????????????????????????????????????????????????????????????????????????????????????????????????????????????????????????????????????????????????????????????????????????????????????????????????????????????????????????????????????????????????????????????????????????????????????????????????????????????????????????????????????????????????????????????????????????????????????????????????????????????????????????????????????????????????????????????????????????????????????????????????????????????????????????????????????????????????????????????????????

			>Guzmania_glaucophylla
			?????????????????????????????????????????????????????????????????????????????????????????????????????????????????????????????????????????????????????????????????????????????????????????????????????????????????????????????????????????????????????????????????????????????????????????????????????????????????????????????????????????????????????????????????????????????????????????????????????????????????????????????????????????????????????????????????????????????????????????????????????????????????????????????????????????????????????????????????????????????????????????????????????

			>Guzmania_graminifolia
			?????????????????????????????????????????????????????????????????????????????????????????????????????????????????????????????????????????????????????????????????????????????????????????????????????????????????????????????????????????????????????????????????????????????????????????????????????????????????????????????????????????????????????????????????????????????????????????????????????????????????????????????????????????????????????????????????????????????????????????????????????????????????????????????????????????????????????????????????????????????????????????????????????

			>Guzmania_herrerae
			?????????????????????????????????????????????????????????????????????????????????????????????????????????????????????????????????????????????????????????????????????????????????????????????????????????????????????????????????????????????????????????????????????????????????????????????????????????????????????????????????????????????????????????????????????????????????????????????????????????????????????????????????????????????????????????????????????????????????????????????????????????????????????????????????????????????????????????????????????????????????????????????????????

			>Guzmania_lingulata
			?????????????????????????????????????????????????????????????????????????????????????????????????????????????????????????????????????????????????????????????????????????????????????????????????????????????????????????????????????????????????????????????????????????????????????????????????????????????????????????????????????????????????????????????????????????????????????????????????????????????????????????????????????????????????????????????????????????????????????????????????????????????????????????????????????????????????????????????????????????????????????????????????????

			>Guzmania_melinonis
			?????????????????????????????????????????????????????????????????????????????????????????????????????????????????????????????????????????????????????????????????????????????????????????????????????????????????????????????????????????????????????????????????????????????????????????????????????????????????????????????????????????????????????????????????????????????????????????????????????????????????????????????????????????????????????????????????????????????????????????????????????????????????????????????????????????????????????????????????????????????????????????????????????

			>Guzmania_monostachia
			?????????????????????????????????????????????????????????????????????????????????????????????????????????????????????????????????????????????????????????????????????????????????????????????????????????????????????????????????????????????????????????????????????????????????????????????????????????????????????????????????????????????????????????????????????????????????????????????????????????????????????????????????????????????????????????????????????????????????????????????????????????????????????????????????????????????????????????????????????????????????????????????????????

			>Guzmania_mucronata
			?????????????????????????????????????????????????????????????????????????????????????????????????????????????????????????????????????????????????????????????????????????????????????????????????????????????????????????????????????????????????????????????????????????????????????????????????????????????????????????????????????????????????????????????????????????????????????????????????????????????????????????????????????????????????????????????????????????????????????????????????????????????????????????????????????????????????????????????????????????????????????????????????????

			>Guzmania_multiflora
			?????????????????????????????????????????????????????????????????????????????????????????????????????????????????????????????????????????????????????????????????????????????????????????????????????????????????????????????????????????????????????????????????????????????????????????????????????????????????????????????????????????????????????????????????????????????????????????????????????????????????????????????????????????????????????????????????????????????????????????????????????????????????????????????????????????????????????????????????????????????????????????????????????

			>Guzmania_musaica
			?????????????????????????????????????????????????????????????????????????????????????????????????????????????????????????????????????????????????????????????????????????????????????????????????????????????????????????????????????????????????????????????????????????????????????????????????????????????????????????????????????????????????????????????????????????????????????????????????????????????????????????????????????????????????????????????????????????????????????????????????????????????????????????????????????????????????????????????????????????????????????????????????????

			>Guzmania_nicaraguensis
			?????????????????????????????????????????????????????????????????????????????????????????????????????????????????????????????????????????????????????????????????????????????????????????????????????????????????????????????????????????????????????????????????????????????????????????????????????????????????????????????????????????????????????????????????????????????????????????????????????????????????????????????????????????????????????????????????????????????????????????????????????????????????????????????????????????????????????????????????????????????????????????????????????

			>Guzmania_osyana
			?????????????????????????????????????????????????????????????????????????????????????????????????????????????????????????????????????????????????????????????????????????????????????????????????????????????????????????????????????????????????????????????????????????????????????????????????????????????????????????????????????????????????????????????????????????????????????????????????????????????????????????????????????????????????????????????????????????????????????????????????????????????????????????????????????????????????????????????????????????????????????????????????????

			>Guzmania_patula
			?????????????????????????????????????????????????????????????????????????????????????????????????????????????????????????????????????????????????????????????????????????????????????????????????????????????????????????????????????????????????????????????????????????????????????????????????????????????????????????????????????????????????????????????????????????????????????????????????????????????????????????????????????????????????????????????????????????????????????????????????????????????????????????????????????????????????????????????????????????????????????????????????????

			>Guzmania_plicatifolia
			?????????????????????????????????????????????????????????????????????????????????????????????????????????????????????????????????????????????????????????????????????????????????????????????????????????????????????????????????????????????????????????????????????????????????????????????????????????????????????????????????????????????????????????????????????????????????????????????????????????????????????????????????????????????????????????????????????????????????????????????????????????????????????????????????????????????????????????????????????????????????????????????????????

			>Guzmania_polycephala
			?????????????????????????????????????????????????????????????????????????????????????????????????????????????????????????????????????????????????????????????????????????????????????????????????????????????????????????????????????????????????????????????????????????????????????????????????????????????????????????????????????????????????????????????????????????????????????????????????????????????????????????????????????????????????????????????????????????????????????????????????????????????????????????????????????????????????????????????????????????????????????????????????????

			>Guzmania_rhonhofiana
			?????????????????????????????????????????????????????????????????????????????????????????????????????????????????????????????????????????????????????????????????????????????????????????????????????????????????????????????????????????????????????????????????????????????????????????????????????????????????????????????????????????????????????????????????????????????????????????????????????????????????????????????????????????????????????????????????????????????????????????????????????????????????????????????????????????????????????????????????????????????????????????????????????

			>Guzmania_roezlii
			?????????????????????????????????????????????????????????????????????????????????????????????????????????????????????????????????????????????????????????????????????????????????????????????????????????????????????????????????????????????????????????????????????????????????????????????????????????????????????????????????????????????????????????????????????????????????????????????????????????????????????????????????????????????????????????????????????????????????????????????????????????????????????????????????????????????????????????????????????????????????????????????????????

			>Guzmania_sanguinea
			?????????????????????????????????????????????????????????????????????????????????????????????????????????????????????????????????????????????????????????????????????????????????????????????????????????????????????????????????????????????????????????????????????????????????????????????????????????????????????????????????????????????????????????????????????????????????????????????????????????????????????????????????????????????????????????????????????????????????????????????????????????????????????????????????????????????????????????????????????????????????????????????????????

			>Guzmania_scherzeriana
			?????????????????????????????????????????????????????????????????????????????????????????????????????????????????????????????????????????????????????????????????????????????????????????????????????????????????????????????????????????????????????????????????????????????????????????????????????????????????????????????????????????????????????????????????????????????????????????????????????????????????????????????????????????????????????????????????????????????????????????????????????????????????????????????????????????????????????????????????????????????????????????????????????

			>Guzmania_spectabilis
			?????????????????????????????????????????????????????????????????????????????????????????????????????????????????????????????????????????????????????????????????????????????????????????????????????????????????????????????????????????????????????????????????????????????????????????????????????????????????????????????????????????????????????????????????????????????????????????????????????????????????????????????????????????????????????????????????????????????????????????????????????????????????????????????????????????????????????????????????????????????????????????????????????

			>Guzmania_subcorymbosa
			?????????????????????????????????????????????????????????????????????????????????????????????????????????????????????????????????????????????????????????????????????????????????????????????????????????????????????????????????????????????????????????????????????????????????????????????????????????????????????????????????????????????????????????????????????????????????????????????????????????????????????????????????????????????????????????????????????????????????????????????????????????????????????????????????????????????????????????????????????????????????????????????????????

			>Guzmania_variegata
			?????????????????????????????????????????????????????????????????????????????????????????????????????????????????????????????????????????????????????????????????????????????????????????????????????????????????????????????????????????????????????????????????????????????????????????????????????????????????????????????????????????????????????????????????????????????????????????????????????????????????????????????????????????????????????????????????????????????????????????????????????????????????????????????????????????????????????????????????????????????????????????????????????

			>Guzmania_wittmackii
			?????????????????????????????????????????????????????????????????????????????????????????????????????????????????????????????????????????????????????????????????????????????????????????????????????????????????????????????????????????????????????????????????????????????????????????????????????????????????????????????????????????????????????????????????????????????????????????????????????????????????????????????????????????????????????????????????????????????????????????????????????????????????????????????????????????????????????????????????????????????????????????????????????

			>Guzmania_zahnii
			?????????????????????????????????????????????????????????????????????????????????????????????????????????????????????????????????????????????????????????????????????????????????????????????????????????????????????????????????????????????????????????????????????????????????????????????????????????????????????????????????????????????????????????????????????????????????????????????????????????????????????????????????????????????????????????????????????????????????????????????????????????????????????????????????????????????????????????????????????????????????????????????????????

			>Hechtia_argentea
			?????????????????????????????????????????????????????????????????????????????????????????????????????????????????????????????????????????????????????????????????????????????????????????????????????????????????????????????????????????????????????????????????????????????????????????????????????????????????????????????????????????????????????????????????????????????????????????????????????????????????????????????????????????????????????????????????????????????????????????????????????????????????????????????????????????????????????????????????????????????????????????????????????

			>Hechtia_caerulea
			?????????????????????????????????????????????????????????????????????????????????????????????????????????????????????????????????????????????????????????????????????????????????????????????????????????????????????????????????????????????????????????????????????????????????????????????????????????????????????????????????????????????????????????????????????????????????????????????????????????????????????????????????????????????????????????????????????????????????????????????????????????????????????????????????????????????????????????????????????????????????????????????????????

			>Hechtia_carlsoniae
			?????????????????????????????????????????????????????????????????????????????????????????????????????????????????????????????????????????????????????????????????????????????????????????????????????????????????????????????????????????????????????????????????????????????????????????????????????????????????????????????????????????????????????????????????????????????????????????????????????????????????????????????????????????????????????????????????????????????????????????????????????????????????????????????????????????????????????????????????????????????????????????????????????

			>Hechtia_deceptrix
			?????????????????????????????????????????????????????????????????????????????????????????????????????????????????????????????????????????????????????????????????????????????????????????????????????????????????????????????????????????????????????????????????????????????????????????????????????????????????????????????????????????????????????????????????????????????????????????????????????????????????????????????????????????????????????????????????????????????????????????????????????????????????????????????????????????????????????????????????????????????????????????????????????

			>Hechtia_dichroantha
			?????????????????????????????????????????????????????????????????????????????????????????????????????????????????????????????????????????????????????????????????????????????????????????????????????????????????????????????????????????????????????????????????????????????????????????????????????????????????????????????????????????????????????????????????????????????????????????????????????????????????????????????????????????????????????????????????????????????????????????????????????????????????????????????????????????????????????????????????????????????????????????????????????

			>Hechtia_epigyna
			?????????????????????????????????????????????????????????????????????????????????????????????????????????????????????????????????????????????????????????????????????????????????????????????????????????????????????????????????????????????????????????????????????????????????????????????????????????????????????????????????????????????????????????????????????????????????????????????????????????????????????????????????????????????????????????????????????????????????????????????????????????????????????????????????????????????????????????????????????????????????????????????????????

			>Hechtia_fosteriana
			?????????????????????????????????????????????????????????????????????????????????????????????????????????????????????????????????????????????????????????????????????????????????????????????????????????????????????????????????????????????????????????????????????????????????????????????????????????????????????????????????????????????????????????????????????????????????????????????????????????????????????????????????????????????????????????????????????????????????????????????????????????????????????????????????????????????????????????????????????????????????????????????????????

			>Hechtia_ghiesbreghtii
			?????????????????????????????????????????????????????????????????????????????????????????????????????????????????????????????????????????????????????????????????????????????????????????????????????????????????????????????????????????????????????????????????????????????????????????????????????????????????????????????????????????????????????????????????????????????????????????????????????????????????????????????????????????????????????????????????????????????????????????????????????????????????????????????????????????????????????????????????????????????????????????????????????

			>Hechtia_glomerata
			?????????????????????????????????????????????????????????????????????????????????????????????????????????????????????????????????????????????????????????????????????????????????????????????????????????????????????????????????????????????????????????????????????????????????????????????????????????????????????????????????????????????????????????????????????????????????????????????????????????????????????????????????????????????????????????????????????????????????????????????????????????????????????????????????????????????????????????????????????????????????????????????????????

			>Hechtia_guatemalensis
			?????????????????????????????????????????????????????????????????????????????????????????????????????????????????????????????????????????????????????????????????????????????????????????????????????????????????????????????????????????????????????????????????????????????????????????????????????????????????????????????????????????????????????????????????????????????????????????????????????????????????????????????????????????????????????????????????????????????????????????????????????????????????????????????????????????????????????????????????????????????????????????????????????

			>Hechtia_hernandez-sandovalii
			?????????????????????????????????????????????????????????????????????????????????????????????????????????????????????????????????????????????????????????????????????????????????????????????????????????????????????????????????????????????????????????????????????????????????????????????????????????????????????????????????????????????????????????????????????????????????????????????????????????????????????????????????????????????????????????????????????????????????????????????????????????????????????????????????????????????????????????????????????????????????????????????????????

			>Hechtia_isthmusiana
			?????????????????????????????????????????????????????????????????????????????????????????????????????????????????????????????????????????????????????????????????????????????????????????????????????????????????????????????????????????????????????????????????????????????????????????????????????????????????????????????????????????????????????????????????????????????????????????????????????????????????????????????????????????????????????????????????????????????????????????????????????????????????????????????????????????????????????????????????????????????????????????????????????

			>Hechtia_lepidophylla
			?????????????????????????????????????????????????????????????????????????????????????????????????????????????????????????????????????????????????????????????????????????????????????????????????????????????????????????????????????????????????????????????????????????????????????????????????????????????????????????????????????????????????????????????????????????????????????????????????????????????????????????????????????????????????????????????????????????????????????????????????????????????????????????????????????????????????????????????????????????????????????????????????????

			>Hechtia_lundelliorum
			?????????????????????????????????????????????????????????????????????????????????????????????????????????????????????????????????????????????????????????????????????????????????????????????????????????????????????????????????????????????????????????????????????????????????????????????????????????????????????????????????????????????????????????????????????????????????????????????????????????????????????????????????????????????????????????????????????????????????????????????????????????????????????????????????????????????????????????????????????????????????????????????????????

			>Hechtia_malvernii
			?????????????????????????????????????????????????????????????????????????????????????????????????????????????????????????????????????????????????????????????????????????????????????????????????????????????????????????????????????????????????????????????????????????????????????????????????????????????????????????????????????????????????????????????????????????????????????????????????????????????????????????????????????????????????????????????????????????????????????????????????????????????????????????????????????????????????????????????????????????????????????????????????????

			>Hechtia_matudae
			?????????????????????????????????????????????????????????????????????????????????????????????????????????????????????????????????????????????????????????????????????????????????????????????????????????????????????????????????????????????????????????????????????????????????????????????????????????????????????????????????????????????????????????????????????????????????????????????????????????????????????????????????????????????????????????????????????????????????????????????????????????????????????????????????????????????????????????????????????????????????????????????????????

			>Hechtia_montana
			?????????????????????????????????????????????????????????????????????????????????????????????????????????????????????????????????????????????????????????????????????????????????????????????????????????????????????????????????????????????????????????????????????????????????????????????????????????????????????????????????????????????????????????????????????????????????????????????????????????????????????????????????????????????????????????????????????????????????????????????????????????????????????????????????????????????????????????????????????????????????????????????????????

			>Hechtia_myriantha
			?????????????????????????????????????????????????????????????????????????????????????????????????????????????????????????????????????????????????????????????????????????????????????????????????????????????????????????????????????????????????????????????????????????????????????????????????????????????????????????????????????????????????????????????????????????????????????????????????????????????????????????????????????????????????????????????????????????????????????????????????????????????????????????????????????????????????????????????????????????????????????????????????????

			>Hechtia_pretiosa
			?????????????????????????????????????????????????????????????????????????????????????????????????????????????????????????????????????????????????????????????????????????????????????????????????????????????????????????????????????????????????????????????????????????????????????????????????????????????????????????????????????????????????????????????????????????????????????????????????????????????????????????????????????????????????????????????????????????????????????????????????????????????????????????????????????????????????????????????????????????????????????????????????????

			>Hechtia_purpusii
			?????????????????????????????????????????????????????????????????????????????????????????????????????????????????????????????????????????????????????????????????????????????????????????????????????????????????????????????????????????????????????????????????????????????????????????????????????????????????????????????????????????????????????????????????????????????????????????????????????????????????????????????????????????????????????????????????????????????????????????????????????????????????????????????????????????????????????????????????????????????????????????????????????

			>Hechtia_schottii
			?????????????????????????????????????????????????????????????????????????????????????????????????????????????????????????????????????????????????????????????????????????????????????????????????????????????????????????????????????????????????????????????????????????????????????????????????????????????????????????????????????????????????????????????????????????????????????????????????????????????????????????????????????????????????????????????????????????????????????????????????????????????????????????????????????????????????????????????????????????????????????????????????????

			>Hechtia_stenopetala
			?????????????????????????????????????????????????????????????????????????????????????????????????????????????????????????????????????????????????????????????????????????????????????????????????????????????????????????????????????????????????????????????????????????????????????????????????????????????????????????????????????????????????????????????????????????????????????????????????????????????????????????????????????????????????????????????????????????????????????????????????????????????????????????????????????????????????????????????????????????????????????????????????????

			>Hechtia_subalata
			?????????????????????????????????????????????????????????????????????????????????????????????????????????????????????????????????????????????????????????????????????????????????????????????????????????????????????????????????????????????????????????????????????????????????????????????????????????????????????????????????????????????????????????????????????????????????????????????????????????????????????????????????????????????????????????????????????????????????????????????????????????????????????????????????????????????????????????????????????????????????????????????????????

			>Hechtia_texensis
			?????????????????????????????????????????????????????????????????????????????????????????????????????????????????????????????????????????????????????????????????????????????????????????????????????????????????????????????????????????????????????????????????????????????????????????????????????????????????????????????????????????????????????????????????????????????????????????????????????????????????????????????????????????????????????????????????????????????????????????????????????????????????????????????????????????????????????????????????????????????????????????????????????

			>Hechtia_tillandsioides
			?????????????????????????????????????????????????????????????????????????????????????????????????????????????????????????????????????????????????????????????????????????????????????????????????????????????????????????????????????????????????????????????????????????????????????????????????????????????????????????????????????????????????????????????????????????????????????????????????????????????????????????????????????????????????????????????????????????????????????????????????????????????????????????????????????????????????????????????????????????????????????????????????????

			>Hohenbergia_andina
			?????????????????????????????????????????????????????????????????????????????????????????????????????????????????????????????????????????????????????????????????????????????????????????????????????????????????????????????????????????????????????????????????????????????????????????????????????????????????????????????????????????????????????????????????????????????????????????????????????????????????????????????????????????????????????????????????????????????????????????????????????????????????????????????????????????????????????????????????????????????????????????????????????

			>Hohenbergia_belemii
			?????????????????????????????????????????????????????????????????????????????????????????????????????????????????????????????????????????????????????????????????????????????????????????????????????????????????????????????????????????????????????????????????????????????????????????????????????????????????????????????????????????????????????????????????????????????????????????????????????????????????????????????????????????????????????????????????????????????????????????????????????????????????????????????????????????????????????????????????????????????????????????????????????

			>Hohenbergia_brachycephala
			?????????????????????????????????????????????????????????????????????????????????????????????????????????????????????????????????????????????????????????????????????????????????????????????????????????????????????????????????????????????????????????????????????????????????????????????????????????????????????????????????????????????????????????????????????????????????????????????????????????????????????????????????????????????????????????????????????????????????????????????????????????????????????????????????????????????????????????????????????????????????????????????????????

			>Hohenbergia_capitata
			?????????????????????????????????????????????????????????????????????????????????????????????????????????????????????????????????????????????????????????????????????????????????????????????????????????????????????????????????????????????????????????????????????????????????????????????????????????????????????????????????????????????????????????????????????????????????????????????????????????????????????????????????????????????????????????????????????????????????????????????????????????????????????????????????????????????????????????????????????????????????????????????????????

			>Hohenbergia_catingae
			?????????????????????????????????????????????????????????????????????????????????????????????????????????????????????????????????????????????????????????????????????????????????????????????????????????????????????????????????????????????????????????????????????????????????????????????????????????????????????????????????????????????????????????????????????????????????????????????????????????????????????????????????????????????????????????????????????????????????????????????????????????????????????????????????????????????????????????????????????????????????????????????????????

			>Hohenbergia_correia-araujoi
			?????????????????????????????????????????????????????????????????????????????????????????????????????????????????????????????????????????????????????????????????????????????????????????????????????????????????????????????????????????????????????????????????????????????????????????????????????????????????????????????????????????????????????????????????????????????????????????????????????????????????????????????????????????????????????????????????????????????????????????????????????????????????????????????????????????????????????????????????????????????????????????????????????

			>Hohenbergia_edmundoi
			?????????????????????????????????????????????????????????????????????????????????????????????????????????????????????????????????????????????????????????????????????????????????????????????????????????????????????????????????????????????????????????????????????????????????????????????????????????????????????????????????????????????????????????????????????????????????????????????????????????????????????????????????????????????????????????????????????????????????????????????????????????????????????????????????????????????????????????????????????????????????????????????????????

			>Hohenbergia_humilis
			?????????????????????????????????????????????????????????????????????????????????????????????????????????????????????????????????????????????????????????????????????????????????????????????????????????????????????????????????????????????????????????????????????????????????????????????????????????????????????????????????????????????????????????????????????????????????????????????????????????????????????????????????????????????????????????????????????????????????????????????????????????????????????????????????????????????????????????????????????????????????????????????????????

			>Hohenbergia_loredanoana
			?????????????????????????????????????????????????????????????????????????????????????????????????????????????????????????????????????????????????????????????????????????????????????????????????????????????????????????????????????????????????????????????????????????????????????????????????????????????????????????????????????????????????????????????????????????????????????????????????????????????????????????????????????????????????????????????????????????????????????????????????????????????????????????????????????????????????????????????????????????????????????????????????????

			>Hohenbergia_pennae
			?????????????????????????????????????????????????????????????????????????????????????????????????????????????????????????????????????????????????????????????????????????????????????????????????????????????????????????????????????????????????????????????????????????????????????????????????????????????????????????????????????????????????????????????????????????????????????????????????????????????????????????????????????????????????????????????????????????????????????????????????????????????????????????????????????????????????????????????????????????????????????????????????????

			>Hohenbergia_ramageana
			?????????????????????????????????????????????????????????????????????????????????????????????????????????????????????????????????????????????????????????????????????????????????????????????????????????????????????????????????????????????????????????????????????????????????????????????????????????????????????????????????????????????????????????????????????????????????????????????????????????????????????????????????????????????????????????????????????????????????????????????????????????????????????????????????????????????????????????????????????????????????????????????????????

			>Hohenbergia_ridleyi
			?????????????????????????????????????????????????????????????????????????????????????????????????????????????????????????????????????????????????????????????????????????????????????????????????????????????????????????????????????????????????????????????????????????????????????????????????????????????????????????????????????????????????????????????????????????????????????????????????????????????????????????????????????????????????????????????????????????????????????????????????????????????????????????????????????????????????????????????????????????????????????????????????????

			>Hohenbergia_rosea
			?????????????????????????????????????????????????????????????????????????????????????????????????????????????????????????????????????????????????????????????????????????????????????????????????????????????????????????????????????????????????????????????????????????????????????????????????????????????????????????????????????????????????????????????????????????????????????????????????????????????????????????????????????????????????????????????????????????????????????????????????????????????????????????????????????????????????????????????????????????????????????????????????????

			>Hohenbergia_salzmannii
			?????????????????????????????????????????????????????????????????????????????????????????????????????????????????????????????????????????????????????????????????????????????????????????????????????????????????????????????????????????????????????????????????????????????????????????????????????????????????????????????????????????????????????????????????????????????????????????????????????????????????????????????????????????????????????????????????????????????????????????????????????????????????????????????????????????????????????????????????????????????????????????????????????

			>Hohenbergia_stellata
			?????????????????????????????????????????????????????????????????????????????????????????????????????????????????????????????????????????????????????????????????????????????????????????????????????????????????????????????????????????????????????????????????????????????????????????????????????????????????????????????????????????????????????????????????????????????????????????????????????????????????????????????????????????????????????????????????????????????????????????????????????????????????????????????????????????????????????????????????????????????????????????????????????

			>Hohenbergia_utriculosa
			?????????????????????????????????????????????????????????????????????????????????????????????????????????????????????????????????????????????????????????????????????????????????????????????????????????????????????????????????????????????????????????????????????????????????????????????????????????????????????????????????????????????????????????????????????????????????????????????????????????????????????????????????????????????????????????????????????????????????????????????????????????????????????????????????????????????????????????????????????????????????????????????????????

			>Hohenbergiopsis_guatemalensis
			?????????????????????????????????????????????????????????????????????????????????????????????????????????????????????????????????????????????????????????????????????????????????????????????????????????????????????????????????????????????????????????????????????????????????????????????????????????????????????????????????????????????????????????????????????????????????????????????????????????????????????????????????????????????????????????????????????????????????????????????????????????????????????????????????????????????????????????????????????????????????????????????????????

			>Hoplocryptanthus_caracensis
			?????????????????????????????????????????????????????????????????????????????????????????????????????????????????????????????????????????????????????????????????????????????????????????????????????????????????????????????????????????????????????????????????????????????????????????????????????????????????????????????????????????????????????????????????????????????????????????????????????????????????????????????????????????????????????????????????????????????????????????????????????????????????????????????????????????????????????????????????????????????????????????????????????

			>Hoplocryptanthus_ferrarius
			?????????????????????????????????????????????????????????????????????????????????????????????????????????????????????????????????????????????????????????????????????????????????????????????????????????????????????????????????????????????????????????????????????????????????????????????????????????????????????????????????????????????????????????????????????????????????????????????????????????????????????????????????????????????????????????????????????????????????????????????????????????????????????????????????????????????????????????????????????????????????????????????????????

			>Hoplocryptanthus_glaziovii
			?????????????????????????????????????????????????????????????????????????????????????????????????????????????????????????????????????????????????????????????????????????????????????????????????????????????????????????????????????????????????????????????????????????????????????????????????????????????????????????????????????????????????????????????????????????????????????????????????????????????????????????????????????????????????????????????????????????????????????????????????????????????????????????????????????????????????????????????????????????????????????????????????????

			>Hoplocryptanthus_lavrasensis
			?????????????????????????????????????????????????????????????????????????????????????????????????????????????????????????????????????????????????????????????????????????????????????????????????????????????????????????????????????????????????????????????????????????????????????????????????????????????????????????????????????????????????????????????????????????????????????????????????????????????????????????????????????????????????????????????????????????????????????????????????????????????????????????????????????????????????????????????????????????????????????????????????????

			>Hoplocryptanthus_regius
			?????????????????????????????????????????????????????????????????????????????????????????????????????????????????????????????????????????????????????????????????????????????????????????????????????????????????????????????????????????????????????????????????????????????????????????????????????????????????????????????????????????????????????????????????????????????????????????????????????????????????????????????????????????????????????????????????????????????????????????????????????????????????????????????????????????????????????????????????????????????????????????????????????

			>Hoplocryptanthus_schwackeanus
			?????????????????????????????????????????????????????????????????????????????????????????????????????????????????????????????????????????????????????????????????????????????????????????????????????????????????????????????????????????????????????????????????????????????????????????????????????????????????????????????????????????????????????????????????????????????????????????????????????????????????????????????????????????????????????????????????????????????????????????????????????????????????????????????????????????????????????????????????????????????????????????????????????

			>Hoplocryptanthus_tiradentesensis
			?????????????????????????????????????????????????????????????????????????????????????????????????????????????????????????????????????????????????????????????????????????????????????????????????????????????????????????????????????????????????????????????????????????????????????????????????????????????????????????????????????????????????????????????????????????????????????????????????????????????????????????????????????????????????????????????????????????????????????????????????????????????????????????????????????????????????????????????????????????????????????????????????????

			>Hoplocryptanthus_vidaliorum
			?????????????????????????????????????????????????????????????????????????????????????????????????????????????????????????????????????????????????????????????????????????????????????????????????????????????????????????????????????????????????????????????????????????????????????????????????????????????????????????????????????????????????????????????????????????????????????????????????????????????????????????????????????????????????????????????????????????????????????????????????????????????????????????????????????????????????????????????????????????????????????????????????????

			>Jagrantia_monstrum
			?????????????????????????????????????????????????????????????????????????????????????????????????????????????????????????????????????????????????????????????????????????????????????????????????????????????????????????????????????????????????????????????????????????????????????????????????????????????????????????????????????????????????????????????????????????????????????????????????????????????????????????????????????????????????????????????????????????????????????????????????????????????????????????????????????????????????????????????????????????????????????????????????????

			>Josemania_asplundii
			?????????????????????????????????????????????????????????????????????????????????????????????????????????????????????????????????????????????????????????????????????????????????????????????????????????????????????????????????????????????????????????????????????????????????????????????????????????????????????????????????????????????????????????????????????????????????????????????????????????????????????????????????????????????????????????????????????????????????????????????????????????????????????????????????????????????????????????????????????????????????????????????????????

			>Josemania_singularis
			?????????????????????????????????????????????????????????????????????????????????????????????????????????????????????????????????????????????????????????????????????????????????????????????????????????????????????????????????????????????????????????????????????????????????????????????????????????????????????????????????????????????????????????????????????????????????????????????????????????????????????????????????????????????????????????????????????????????????????????????????????????????????????????????????????????????????????????????????????????????????????????????????????

			>Karawata_depressa
			?????????????????????????????????????????????????????????????????????????????????????????????????????????????????????????????????????????????????????????????????????????????????????????????????????????????????????????????????????????????????????????????????????????????????????????????????????????????????????????????????????????????????????????????????????????????????????????????????????????????????????????????????????????????????????????????????????????????????????????????????????????????????????????????????????????????????????????????????????????????????????????????????????

			>Karawata_gustavoi
			?????????????????????????????????????????????????????????????????????????????????????????????????????????????????????????????????????????????????????????????????????????????????????????????????????????????????????????????????????????????????????????????????????????????????????????????????????????????????????????????????????????????????????????????????????????????????????????????????????????????????????????????????????????????????????????????????????????????????????????????????????????????????????????????????????????????????????????????????????????????????????????????????????

			>Karawata_hostilis
			?????????????????????????????????????????????????????????????????????????????????????????????????????????????????????????????????????????????????????????????????????????????????????????????????????????????????????????????????????????????????????????????????????????????????????????????????????????????????????????????????????????????????????????????????????????????????????????????????????????????????????????????????????????????????????????????????????????????????????????????????????????????????????????????????????????????????????????????????????????????????????????????????????

			>Karawata_multiflora
			?????????????????????????????????????????????????????????????????????????????????????????????????????????????????????????????????????????????????????????????????????????????????????????????????????????????????????????????????????????????????????????????????????????????????????????????????????????????????????????????????????????????????????????????????????????????????????????????????????????????????????????????????????????????????????????????????????????????????????????????????????????????????????????????????????????????????????????????????????????????????????????????????????

			>Karawata_nigribracteata
			?????????????????????????????????????????????????????????????????????????????????????????????????????????????????????????????????????????????????????????????????????????????????????????????????????????????????????????????????????????????????????????????????????????????????????????????????????????????????????????????????????????????????????????????????????????????????????????????????????????????????????????????????????????????????????????????????????????????????????????????????????????????????????????????????????????????????????????????????????????????????????????????????????

			>Karawata_saxicola
			?????????????????????????????????????????????????????????????????????????????????????????????????????????????????????????????????????????????????????????????????????????????????????????????????????????????????????????????????????????????????????????????????????????????????????????????????????????????????????????????????????????????????????????????????????????????????????????????????????????????????????????????????????????????????????????????????????????????????????????????????????????????????????????????????????????????????????????????????????????????????????????????????????

			>Lapanthus_duartei
			?????????????????????????????????????????????????????????????????????????????????????????????????????????????????????????????????????????????????????????????????????????????????????????????????????????????????????????????????????????????????????????????????????????????????????????????????????????????????????????????????????????????????????????????????????????????????????????????????????????????????????????????????????????????????????????????????????????????????????????????????????????????????????????????????????????????????????????????????????????????????????????????????????

			>Lapanthus_itambensis
			?????????????????????????????????????????????????????????????????????????????????????????????????????????????????????????????????????????????????????????????????????????????????????????????????????????????????????????????????????????????????????????????????????????????????????????????????????????????????????????????????????????????????????????????????????????????????????????????????????????????????????????????????????????????????????????????????????????????????????????????????????????????????????????????????????????????????????????????????????????????????????????????????????

			>Lemeltonia_dodsonii
			?????????????????????????????????????????????????????????????????????????????????????????????????????????????????????????????????????????????????????????????????????????????????????????????????????????????????????????????????????????????????????????????????????????????????????????????????????????????????????????????????????????????????????????????????????????????????????????????????????????????????????????????????????????????????????????????????????????????????????????????????????????????????????????????????????????????????????????????????????????????????????????????????????

			>Lemeltonia_monadelpha
			?????????????????????????????????????????????????????????????????????????????????????????????????????????????????????????????????????????????????????????????????????????????????????????????????????????????????????????????????????????????????????????????????????????????????????????????????????????????????????????????????????????????????????????????????????????????????????????????????????????????????????????????????????????????????????????????????????????????????????????????????????????????????????????????????????????????????????????????????????????????????????????????????????

			>Lemeltonia_narthecioides
			?????????????????????????????????????????????????????????????????????????????????????????????????????????????????????????????????????????????????????????????????????????????????????????????????????????????????????????????????????????????????????????????????????????????????????????????????????????????????????????????????????????????????????????????????????????????????????????????????????????????????????????????????????????????????????????????????????????????????????????????????????????????????????????????????????????????????????????????????????????????????????????????????????

			>Lemeltonia_triglochinoides
			?????????????????????????????????????????????????????????????????????????????????????????????????????????????????????????????????????????????????????????????????????????????????????????????????????????????????????????????????????????????????????????????????????????????????????????????????????????????????????????????????????????????????????????????????????????????????????????????????????????????????????????????????????????????????????????????????????????????????????????????????????????????????????????????????????????????????????????????????????????????????????????????????????

			>Lindmania_guianensis
			?????????????????????????????????????????????????????????????????????????????????????????????????????????????????????????????????????????????????????????????????????????????????????????????????????????????????????????????????????????????????????????????????????????????????????????????????????????????????????????????????????????????????????????????????????????????????????????????????????????????????????????????????????????????????????????????????????????????????????????????????????????????????????????????????????????????????????????????????????????????????????????????????????

			>Lindmania_longipes
			?????????????????????????????????????????????????????????????????????????????????????????????????????????????????????????????????????????????????????????????????????????????????????????????????????????????????????????????????????????????????????????????????????????????????????????????????????????????????????????????????????????????????????????????????????????????????????????????????????????????????????????????????????????????????????????????????????????????????????????????????????????????????????????????????????????????????????????????????????????????????????????????????????

			>Lutheria_glutinosa
			?????????????????????????????????????????????????????????????????????????????????????????????????????????????????????????????????????????????????????????????????????????????????????????????????????????????????????????????????????????????????????????????????????????????????????????????????????????????????????????????????????????????????????????????????????????????????????????????????????????????????????????????????????????????????????????????????????????????????????????????????????????????????????????????????????????????????????????????????????????????????????????????????????

			>Lutheria_splendens
			?????????????????????????????????????????????????????????????????????????????????????????????????????????????????????????????????????????????????????????????????????????????????????????????????????????????????????????????????????????????????????????????????????????????????????????????????????????????????????????????????????????????????????????????????????????????????????????????????????????????????????????????????????????????????????????????????????????????????????????????????????????????????????????????????????????????????????????????????????????????????????????????????????

			>Lymania_alvimii
			?????????????????????????????????????????????????????????????????????????????????????????????????????????????????????????????????????????????????????????????????????????????????????????????????????????????????????????????????????????????????????????????????????????????????????????????????????????????????????????????????????????????????????????????????????????????????????????????????????????????????????????????????????????????????????????????????????????????????????????????????????????????????????????????????????????????????????????????????????????????????????????????????????

			>Lymania_azurea
			?????????????????????????????????????????????????????????????????????????????????????????????????????????????????????????????????????????????????????????????????????????????????????????????????????????????????????????????????????????????????????????????????????????????????????????????????????????????????????????????????????????????????????????????????????????????????????????????????????????????????????????????????????????????????????????????????????????????????????????????????????????????????????????????????????????????????????????????????????????????????????????????????????

			>Lymania_brachycaulis
			?????????????????????????????????????????????????????????????????????????????????????????????????????????????????????????????????????????????????????????????????????????????????????????????????????????????????????????????????????????????????????????????????????????????????????????????????????????????????????????????????????????????????????????????????????????????????????????????????????????????????????????????????????????????????????????????????????????????????????????????????????????????????????????????????????????????????????????????????????????????????????????????????????

			>Lymania_corallina
			?????????????????????????????????????????????????????????????????????????????????????????????????????????????????????????????????????????????????????????????????????????????????????????????????????????????????????????????????????????????????????????????????????????????????????????????????????????????????????????????????????????????????????????????????????????????????????????????????????????????????????????????????????????????????????????????????????????????????????????????????????????????????????????????????????????????????????????????????????????????????????????????????????

			>Lymania_globosa
			?????????????????????????????????????????????????????????????????????????????????????????????????????????????????????????????????????????????????????????????????????????????????????????????????????????????????????????????????????????????????????????????????????????????????????????????????????????????????????????????????????????????????????????????????????????????????????????????????????????????????????????????????????????????????????????????????????????????????????????????????????????????????????????????????????????????????????????????????????????????????????????????????????

			>Lymania_languida
			?????????????????????????????????????????????????????????????????????????????????????????????????????????????????????????????????????????????????????????????????????????????????????????????????????????????????????????????????????????????????????????????????????????????????????????????????????????????????????????????????????????????????????????????????????????????????????????????????????????????????????????????????????????????????????????????????????????????????????????????????????????????????????????????????????????????????????????????????????????????????????????????????????

			>Lymania_smithii
			?????????????????????????????????????????????????????????????????????????????????????????????????????????????????????????????????????????????????????????????????????????????????????????????????????????????????????????????????????????????????????????????????????????????????????????????????????????????????????????????????????????????????????????????????????????????????????????????????????????????????????????????????????????????????????????????????????????????????????????????????????????????????????????????????????????????????????????????????????????????????????????????????????

			>Lymania_spiculata
			?????????????????????????????????????????????????????????????????????????????????????????????????????????????????????????????????????????????????????????????????????????????????????????????????????????????????????????????????????????????????????????????????????????????????????????????????????????????????????????????????????????????????????????????????????????????????????????????????????????????????????????????????????????????????????????????????????????????????????????????????????????????????????????????????????????????????????????????????????????????????????????????????????

			>Mezobromelia_bicolor
			?????????????????????????????????????????????????????????????????????????????????????????????????????????????????????????????????????????????????????????????????????????????????????????????????????????????????????????????????????????????????????????????????????????????????????????????????????????????????????????????????????????????????????????????????????????????????????????????????????????????????????????????????????????????????????????????????????????????????????????????????????????????????????????????????????????????????????????????????????????????????????????????????????

			>Mezobromelia_capituligera
			?????????????????????????????????????????????????????????????????????????????????????????????????????????????????????????????????????????????????????????????????????????????????????????????????????????????????????????????????????????????????????????????????????????????????????????????????????????????????????????????????????????????????????????????????????????????????????????????????????????????????????????????????????????????????????????????????????????????????????????????????????????????????????????????????????????????????????????????????????????????????????????????????????

			>Mezobromelia_pleiosticha
			?????????????????????????????????????????????????????????????????????????????????????????????????????????????????????????????????????????????????????????????????????????????????????????????????????????????????????????????????????????????????????????????????????????????????????????????????????????????????????????????????????????????????????????????????????????????????????????????????????????????????????????????????????????????????????????????????????????????????????????????????????????????????????????????????????????????????????????????????????????????????????????????????????

			>Navia_barbellata
			?????????????????????????????????????????????????????????????????????????????????????????????????????????????????????????????????????????????????????????????????????????????????????????????????????????????????????????????????????????????????????????????????????????????????????????????????????????????????????????????????????????????????????????????????????????????????????????????????????????????????????????????????????????????????????????????????????????????????????????????????????????????????????????????????????????????????????????????????????????????????????????????????????

			>Navia_igneosicola
			?????????????????????????????????????????????????????????????????????????????????????????????????????????????????????????????????????????????????????????????????????????????????????????????????????????????????????????????????????????????????????????????????????????????????????????????????????????????????????????????????????????????????????????????????????????????????????????????????????????????????????????????????????????????????????????????????????????????????????????????????????????????????????????????????????????????????????????????????????????????????????????????????????

			>Navia_phelpsiae
			?????????????????????????????????????????????????????????????????????????????????????????????????????????????????????????????????????????????????????????????????????????????????????????????????????????????????????????????????????????????????????????????????????????????????????????????????????????????????????????????????????????????????????????????????????????????????????????????????????????????????????????????????????????????????????????????????????????????????????????????????????????????????????????????????????????????????????????????????????????????????????????????????????

			>Navia_saxicola
			?????????????????????????????????????????????????????????????????????????????????????????????????????????????????????????????????????????????????????????????????????????????????????????????????????????????????????????????????????????????????????????????????????????????????????????????????????????????????????????????????????????????????????????????????????????????????????????????????????????????????????????????????????????????????????????????????????????????????????????????????????????????????????????????????????????????????????????????????????????????????????????????????????

			>Neoglaziovia_burle-marxii
			?????????????????????????????????????????????????????????????????????????????????????????????????????????????????????????????????????????????????????????????????????????????????????????????????????????????????????????????????????????????????????????????????????????????????????????????????????????????????????????????????????????????????????????????????????????????????????????????????????????????????????????????????????????????????????????????????????????????????????????????????????????????????????????????????????????????????????????????????????????????????????????????????????

			>Neoglaziovia_variegata
			?????????????????????????????????????????????????????????????????????????????????????????????????????????????????????????????????????????????????????????????????????????????????????????????????????????????????????????????????????????????????????????????????????????????????????????????????????????????????????????????????????????????????????????????????????????????????????????????????????????????????????????????????????????????????????????????????????????????????????????????????????????????????????????????????????????????????????????????????????????????????????????????????????

			>Neoregelia_binotii
			?????????????????????????????????????????????????????????????????????????????????????????????????????????????????????????????????????????????????????????????????????????????????????????????????????????????????????????????????????????????????????????????????????????????????????????????????????????????????????????????????????????????????????????????????????????????????????????????????????????????????????????????????????????????????????????????????????????????????????????????????????????????????????????????????????????????????????????????????????????????????????????????????????

			>Neoregelia_bragarum
			?????????????????????????????????????????????????????????????????????????????????????????????????????????????????????????????????????????????????????????????????????????????????????????????????????????????????????????????????????????????????????????????????????????????????????????????????????????????????????????????????????????????????????????????????????????????????????????????????????????????????????????????????????????????????????????????????????????????????????????????????????????????????????????????????????????????????????????????????????????????????????????????????????

			>Neoregelia_capixaba
			?????????????????????????????????????????????????????????????????????????????????????????????????????????????????????????????????????????????????????????????????????????????????????????????????????????????????????????????????????????????????????????????????????????????????????????????????????????????????????????????????????????????????????????????????????????????????????????????????????????????????????????????????????????????????????????????????????????????????????????????????????????????????????????????????????????????????????????????????????????????????????????????????????

			>Neoregelia_carolinae
			?????????????????????????????????????????????????????????????????????????????????????????????????????????????????????????????????????????????????????????????????????????????????????????????????????????????????????????????????????????????????????????????????????????????????????????????????????????????????????????????????????????????????????????????????????????????????????????????????????????????????????????????????????????????????????????????????????????????????????????????????????????????????????????????????????????????????????????????????????????????????????????????????????

			>Neoregelia_compacta
			?????????????????????????????????????????????????????????????????????????????????????????????????????????????????????????????????????????????????????????????????????????????????????????????????????????????????????????????????????????????????????????????????????????????????????????????????????????????????????????????????????????????????????????????????????????????????????????????????????????????????????????????????????????????????????????????????????????????????????????????????????????????????????????????????????????????????????????????????????????????????????????????????????

			>Neoregelia_concentrica
			?????????????????????????????????????????????????????????????????????????????????????????????????????????????????????????????????????????????????????????????????????????????????????????????????????????????????????????????????????????????????????????????????????????????????????????????????????????????????????????????????????????????????????????????????????????????????????????????????????????????????????????????????????????????????????????????????????????????????????????????????????????????????????????????????????????????????????????????????????????????????????????????????????

			>Neoregelia_cruenta
			?????????????????????????????????????????????????????????????????????????????????????????????????????????????????????????????????????????????????????????????????????????????????????????????????????????????????????????????????????????????????????????????????????????????????????????????????????????????????????????????????????????????????????????????????????????????????????????????????????????????????????????????????????????????????????????????????????????????????????????????????????????????????????????????????????????????????????????????????????????????????????????????????????

			>Neoregelia_dungsiana
			?????????????????????????????????????????????????????????????????????????????????????????????????????????????????????????????????????????????????????????????????????????????????????????????????????????????????????????????????????????????????????????????????????????????????????????????????????????????????????????????????????????????????????????????????????????????????????????????????????????????????????????????????????????????????????????????????????????????????????????????????????????????????????????????????????????????????????????????????????????????????????????????????????

			>Neoregelia_eleutheropetala
			?????????????????????????????????????????????????????????????????????????????????????????????????????????????????????????????????????????????????????????????????????????????????????????????????????????????????????????????????????????????????????????????????????????????????????????????????????????????????????????????????????????????????????????????????????????????????????????????????????????????????????????????????????????????????????????????????????????????????????????????????????????????????????????????????????????????????????????????????????????????????????????????????????

			>Neoregelia_lactea
			?????????????????????????????????????????????????????????????????????????????????????????????????????????????????????????????????????????????????????????????????????????????????????????????????????????????????????????????????????????????????????????????????????????????????????????????????????????????????????????????????????????????????????????????????????????????????????????????????????????????????????????????????????????????????????????????????????????????????????????????????????????????????????????????????????????????????????????????????????????????????????????????????????

			>Neoregelia_laevis
			?????????????????????????????????????????????????????????????????????????????????????????????????????????????????????????????????????????????????????????????????????????????????????????????????????????????????????????????????????????????????????????????????????????????????????????????????????????????????????????????????????????????????????????????????????????????????????????????????????????????????????????????????????????????????????????????????????????????????????????????????????????????????????????????????????????????????????????????????????????????????????????????????????

			>Neoregelia_leucophoea
			?????????????????????????????????????????????????????????????????????????????????????????????????????????????????????????????????????????????????????????????????????????????????????????????????????????????????????????????????????????????????????????????????????????????????????????????????????????????????????????????????????????????????????????????????????????????????????????????????????????????????????????????????????????????????????????????????????????????????????????????????????????????????????????????????????????????????????????????????????????????????????????????????????

			>Neoregelia_longisepala
			?????????????????????????????????????????????????????????????????????????????????????????????????????????????????????????????????????????????????????????????????????????????????????????????????????????????????????????????????????????????????????????????????????????????????????????????????????????????????????????????????????????????????????????????????????????????????????????????????????????????????????????????????????????????????????????????????????????????????????????????????????????????????????????????????????????????????????????????????????????????????????????????????????

			>Neoregelia_lymaniana
			?????????????????????????????????????????????????????????????????????????????????????????????????????????????????????????????????????????????????????????????????????????????????????????????????????????????????????????????????????????????????????????????????????????????????????????????????????????????????????????????????????????????????????????????????????????????????????????????????????????????????????????????????????????????????????????????????????????????????????????????????????????????????????????????????????????????????????????????????????????????????????????????????????

			>Neoregelia_mucugensis
			?????????????????????????????????????????????????????????????????????????????????????????????????????????????????????????????????????????????????????????????????????????????????????????????????????????????????????????????????????????????????????????????????????????????????????????????????????????????????????????????????????????????????????????????????????????????????????????????????????????????????????????????????????????????????????????????????????????????????????????????????????????????????????????????????????????????????????????????????????????????????????????????????????

			>Neoregelia_myrmecophila
			?????????????????????????????????????????????????????????????????????????????????????????????????????????????????????????????????????????????????????????????????????????????????????????????????????????????????????????????????????????????????????????????????????????????????????????????????????????????????????????????????????????????????????????????????????????????????????????????????????????????????????????????????????????????????????????????????????????????????????????????????????????????????????????????????????????????????????????????????????????????????????????????????????

			>Neoregelia_nevaresii
			?????????????????????????????????????????????????????????????????????????????????????????????????????????????????????????????????????????????????????????????????????????????????????????????????????????????????????????????????????????????????????????????????????????????????????????????????????????????????????????????????????????????????????????????????????????????????????????????????????????????????????????????????????????????????????????????????????????????????????????????????????????????????????????????????????????????????????????????????????????????????????????????????????

			>Neoregelia_nivea
			?????????????????????????????????????????????????????????????????????????????????????????????????????????????????????????????????????????????????????????????????????????????????????????????????????????????????????????????????????????????????????????????????????????????????????????????????????????????????????????????????????????????????????????????????????????????????????????????????????????????????????????????????????????????????????????????????????????????????????????????????????????????????????????????????????????????????????????????????????????????????????????????????????

			>Neoregelia_pascoaliana
			?????????????????????????????????????????????????????????????????????????????????????????????????????????????????????????????????????????????????????????????????????????????????????????????????????????????????????????????????????????????????????????????????????????????????????????????????????????????????????????????????????????????????????????????????????????????????????????????????????????????????????????????????????????????????????????????????????????????????????????????????????????????????????????????????????????????????????????????????????????????????????????????????????

			>Neoregelia_pauciflora
			?????????????????????????????????????????????????????????????????????????????????????????????????????????????????????????????????????????????????????????????????????????????????????????????????????????????????????????????????????????????????????????????????????????????????????????????????????????????????????????????????????????????????????????????????????????????????????????????????????????????????????????????????????????????????????????????????????????????????????????????????????????????????????????????????????????????????????????????????????????????????????????????????????

			>Neoregelia_pineliana
			?????????????????????????????????????????????????????????????????????????????????????????????????????????????????????????????????????????????????????????????????????????????????????????????????????????????????????????????????????????????????????????????????????????????????????????????????????????????????????????????????????????????????????????????????????????????????????????????????????????????????????????????????????????????????????????????????????????????????????????????????????????????????????????????????????????????????????????????????????????????????????????????????????

			>Neoregelia_retrorsa
			?????????????????????????????????????????????????????????????????????????????????????????????????????????????????????????????????????????????????????????????????????????????????????????????????????????????????????????????????????????????????????????????????????????????????????????????????????????????????????????????????????????????????????????????????????????????????????????????????????????????????????????????????????????????????????????????????????????????????????????????????????????????????????????????????????????????????????????????????????????????????????????????????????

			>Neoregelia_richteri
			?????????????????????????????????????????????????????????????????????????????????????????????????????????????????????????????????????????????????????????????????????????????????????????????????????????????????????????????????????????????????????????????????????????????????????????????????????????????????????????????????????????????????????????????????????????????????????????????????????????????????????????????????????????????????????????????????????????????????????????????????????????????????????????????????????????????????????????????????????????????????????????????????????

			>Neoregelia_rothinessa
			?????????????????????????????????????????????????????????????????????????????????????????????????????????????????????????????????????????????????????????????????????????????????????????????????????????????????????????????????????????????????????????????????????????????????????????????????????????????????????????????????????????????????????????????????????????????????????????????????????????????????????????????????????????????????????????????????????????????????????????????????????????????????????????????????????????????????????????????????????????????????????????????????????

			>Neoregelia_rubrovittata
			?????????????????????????????????????????????????????????????????????????????????????????????????????????????????????????????????????????????????????????????????????????????????????????????????????????????????????????????????????????????????????????????????????????????????????????????????????????????????????????????????????????????????????????????????????????????????????????????????????????????????????????????????????????????????????????????????????????????????????????????????????????????????????????????????????????????????????????????????????????????????????????????????????

			>Neoregelia_tenebrosa
			?????????????????????????????????????????????????????????????????????????????????????????????????????????????????????????????????????????????????????????????????????????????????????????????????????????????????????????????????????????????????????????????????????????????????????????????????????????????????????????????????????????????????????????????????????????????????????????????????????????????????????????????????????????????????????????????????????????????????????????????????????????????????????????????????????????????????????????????????????????????????????????????????????

			>Neoregelia_viridovinosa
			?????????????????????????????????????????????????????????????????????????????????????????????????????????????????????????????????????????????????????????????????????????????????????????????????????????????????????????????????????????????????????????????????????????????????????????????????????????????????????????????????????????????????????????????????????????????????????????????????????????????????????????????????????????????????????????????????????????????????????????????????????????????????????????????????????????????????????????????????????????????????????????????????????

			>Neoregelia_wilsoniana
			?????????????????????????????????????????????????????????????????????????????????????????????????????????????????????????????????????????????????????????????????????????????????????????????????????????????????????????????????????????????????????????????????????????????????????????????????????????????????????????????????????????????????????????????????????????????????????????????????????????????????????????????????????????????????????????????????????????????????????????????????????????????????????????????????????????????????????????????????????????????????????????????????????

			>Neoregelia_zonata
			?????????????????????????????????????????????????????????????????????????????????????????????????????????????????????????????????????????????????????????????????????????????????????????????????????????????????????????????????????????????????????????????????????????????????????????????????????????????????????????????????????????????????????????????????????????????????????????????????????????????????????????????????????????????????????????????????????????????????????????????????????????????????????????????????????????????????????????????????????????????????????????????????????

			>Nidularium_altimontanum
			?????????????????????????????????????????????????????????????????????????????????????????????????????????????????????????????????????????????????????????????????????????????????????????????????????????????????????????????????????????????????????????????????????????????????????????????????????????????????????????????????????????????????????????????????????????????????????????????????????????????????????????????????????????????????????????????????????????????????????????????????????????????????????????????????????????????????????????????????????????????????????????????????????

			>Nidularium_amazonicum
			?????????????????????????????????????????????????????????????????????????????????????????????????????????????????????????????????????????????????????????????????????????????????????????????????????????????????????????????????????????????????????????????????????????????????????????????????????????????????????????????????????????????????????????????????????????????????????????????????????????????????????????????????????????????????????????????????????????????????????????????????????????????????????????????????????????????????????????????????????????????????????????????????????

			>Nidularium_angustibracteatum
			?????????????????????????????????????????????????????????????????????????????????????????????????????????????????????????????????????????????????????????????????????????????????????????????????????????????????????????????????????????????????????????????????????????????????????????????????????????????????????????????????????????????????????????????????????????????????????????????????????????????????????????????????????????????????????????????????????????????????????????????????????????????????????????????????????????????????????????????????????????????????????????????????????

			>Nidularium_campoalegrense
			?????????????????????????????????????????????????????????????????????????????????????????????????????????????????????????????????????????????????????????????????????????????????????????????????????????????????????????????????????????????????????????????????????????????????????????????????????????????????????????????????????????????????????????????????????????????????????????????????????????????????????????????????????????????????????????????????????????????????????????????????????????????????????????????????????????????????????????????????????????????????????????????????????

			>Nidularium_camposportoi
			?????????????????????????????????????????????????????????????????????????????????????????????????????????????????????????????????????????????????????????????????????????????????????????????????????????????????????????????????????????????????????????????????????????????????????????????????????????????????????????????????????????????????????????????????????????????????????????????????????????????????????????????????????????????????????????????????????????????????????????????????????????????????????????????????????????????????????????????????????????????????????????????????????

			>Nidularium_fulgens
			?????????????????????????????????????????????????????????????????????????????????????????????????????????????????????????????????????????????????????????????????????????????????????????????????????????????????????????????????????????????????????????????????????????????????????????????????????????????????????????????????????????????????????????????????????????????????????????????????????????????????????????????????????????????????????????????????????????????????????????????????????????????????????????????????????????????????????????????????????????????????????????????????????

			>Nidularium_innocentii
			?????????????????????????????????????????????????????????????????????????????????????????????????????????????????????????????????????????????????????????????????????????????????????????????????????????????????????????????????????????????????????????????????????????????????????????????????????????????????????????????????????????????????????????????????????????????????????????????????????????????????????????????????????????????????????????????????????????????????????????????????????????????????????????????????????????????????????????????????????????????????????????????????????

			>Nidularium_krisgreeniae
			?????????????????????????????????????????????????????????????????????????????????????????????????????????????????????????????????????????????????????????????????????????????????????????????????????????????????????????????????????????????????????????????????????????????????????????????????????????????????????????????????????????????????????????????????????????????????????????????????????????????????????????????????????????????????????????????????????????????????????????????????????????????????????????????????????????????????????????????????????????????????????????????????????

			>Nidularium_marigoi
			?????????????????????????????????????????????????????????????????????????????????????????????????????????????????????????????????????????????????????????????????????????????????????????????????????????????????????????????????????????????????????????????????????????????????????????????????????????????????????????????????????????????????????????????????????????????????????????????????????????????????????????????????????????????????????????????????????????????????????????????????????????????????????????????????????????????????????????????????????????????????????????????????????

			>Nidularium_picinguabense
			?????????????????????????????????????????????????????????????????????????????????????????????????????????????????????????????????????????????????????????????????????????????????????????????????????????????????????????????????????????????????????????????????????????????????????????????????????????????????????????????????????????????????????????????????????????????????????????????????????????????????????????????????????????????????????????????????????????????????????????????????????????????????????????????????????????????????????????????????????????????????????????????????????

			>Nidularium_procerum
			?????????????????????????????????????????????????????????????????????????????????????????????????????????????????????????????????????????????????????????????????????????????????????????????????????????????????????????????????????????????????????????????????????????????????????????????????????????????????????????????????????????????????????????????????????????????????????????????????????????????????????????????????????????????????????????????????????????????????????????????????????????????????????????????????????????????????????????????????????????????????????????????????????

			>Nidularium_purpureum
			?????????????????????????????????????????????????????????????????????????????????????????????????????????????????????????????????????????????????????????????????????????????????????????????????????????????????????????????????????????????????????????????????????????????????????????????????????????????????????????????????????????????????????????????????????????????????????????????????????????????????????????????????????????????????????????????????????????????????????????????????????????????????????????????????????????????????????????????????????????????????????????????????????

			>Ochagavia_andina
			?????????????????????????????????????????????????????????????????????????????????????????????????????????????????????????????????????????????????????????????????????????????????????????????????????????????????????????????????????????????????????????????????????????????????????????????????????????????????????????????????????????????????????????????????????????????????????????????????????????????????????????????????????????????????????????????????????????????????????????????????????????????????????????????????????????????????????????????????????????????????????????????????????

			>Ochagavia_carnea
			?????????????????????????????????????????????????????????????????????????????????????????????????????????????????????????????????????????????????????????????????????????????????????????????????????????????????????????????????????????????????????????????????????????????????????????????????????????????????????????????????????????????????????????????????????????????????????????????????????????????????????????????????????????????????????????????????????????????????????????????????????????????????????????????????????????????????????????????????????????????????????????????????????

			>Ochagavia_elegans
			?????????????????????????????????????????????????????????????????????????????????????????????????????????????????????????????????????????????????????????????????????????????????????????????????????????????????????????????????????????????????????????????????????????????????????????????????????????????????????????????????????????????????????????????????????????????????????????????????????????????????????????????????????????????????????????????????????????????????????????????????????????????????????????????????????????????????????????????????????????????????????????????????????

			>Ochagavia_litoralis
			?????????????????????????????????????????????????????????????????????????????????????????????????????????????????????????????????????????????????????????????????????????????????????????????????????????????????????????????????????????????????????????????????????????????????????????????????????????????????????????????????????????????????????????????????????????????????????????????????????????????????????????????????????????????????????????????????????????????????????????????????????????????????????????????????????????????????????????????????????????????????????????????????????

			>Orthophytum_alvimii
			?????????????????????????????????????????????????????????????????????????????????????????????????????????????????????????????????????????????????????????????????????????????????????????????????????????????????????????????????????????????????????????????????????????????????????????????????????????????????????????????????????????????????????????????????????????????????????????????????????????????????????????????????????????????????????????????????????????????????????????????????????????????????????????????????????????????????????????????????????????????????????????????????????

			>Orthophytum_argentum
			?????????????????????????????????????????????????????????????????????????????????????????????????????????????????????????????????????????????????????????????????????????????????????????????????????????????????????????????????????????????????????????????????????????????????????????????????????????????????????????????????????????????????????????????????????????????????????????????????????????????????????????????????????????????????????????????????????????????????????????????????????????????????????????????????????????????????????????????????????????????????????????????????????

			>Orthophytum_benzingii
			?????????????????????????????????????????????????????????????????????????????????????????????????????????????????????????????????????????????????????????????????????????????????????????????????????????????????????????????????????????????????????????????????????????????????????????????????????????????????????????????????????????????????????????????????????????????????????????????????????????????????????????????????????????????????????????????????????????????????????????????????????????????????????????????????????????????????????????????????????????????????????????????????????

			>Orthophytum_boudetianum
			?????????????????????????????????????????????????????????????????????????????????????????????????????????????????????????????????????????????????????????????????????????????????????????????????????????????????????????????????????????????????????????????????????????????????????????????????????????????????????????????????????????????????????????????????????????????????????????????????????????????????????????????????????????????????????????????????????????????????????????????????????????????????????????????????????????????????????????????????????????????????????????????????????

			>Orthophytum_braunii
			?????????????????????????????????????????????????????????????????????????????????????????????????????????????????????????????????????????????????????????????????????????????????????????????????????????????????????????????????????????????????????????????????????????????????????????????????????????????????????????????????????????????????????????????????????????????????????????????????????????????????????????????????????????????????????????????????????????????????????????????????????????????????????????????????????????????????????????????????????????????????????????????????????

			>Orthophytum_compactum
			?????????????????????????????????????????????????????????????????????????????????????????????????????????????????????????????????????????????????????????????????????????????????????????????????????????????????????????????????????????????????????????????????????????????????????????????????????????????????????????????????????????????????????????????????????????????????????????????????????????????????????????????????????????????????????????????????????????????????????????????????????????????????????????????????????????????????????????????????????????????????????????????????????

			>Orthophytum_conquistense
			?????????????????????????????????????????????????????????????????????????????????????????????????????????????????????????????????????????????????????????????????????????????????????????????????????????????????????????????????????????????????????????????????????????????????????????????????????????????????????????????????????????????????????????????????????????????????????????????????????????????????????????????????????????????????????????????????????????????????????????????????????????????????????????????????????????????????????????????????????????????????????????????????????

			>Orthophytum_diamantinense
			?????????????????????????????????????????????????????????????????????????????????????????????????????????????????????????????????????????????????????????????????????????????????????????????????????????????????????????????????????????????????????????????????????????????????????????????????????????????????????????????????????????????????????????????????????????????????????????????????????????????????????????????????????????????????????????????????????????????????????????????????????????????????????????????????????????????????????????????????????????????????????????????????????

			>Orthophytum_disjunctum
			?????????????????????????????????????????????????????????????????????????????????????????????????????????????????????????????????????????????????????????????????????????????????????????????????????????????????????????????????????????????????????????????????????????????????????????????????????????????????????????????????????????????????????????????????????????????????????????????????????????????????????????????????????????????????????????????????????????????????????????????????????????????????????????????????????????????????????????????????????????????????????????????????????

			>Orthophytum_duartei
			?????????????????????????????????????????????????????????????????????????????????????????????????????????????????????????????????????????????????????????????????????????????????????????????????????????????????????????????????????????????????????????????????????????????????????????????????????????????????????????????????????????????????????????????????????????????????????????????????????????????????????????????????????????????????????????????????????????????????????????????????????????????????????????????????????????????????????????????????????????????????????????????????????

			>Orthophytum_estevesii
			?????????????????????????????????????????????????????????????????????????????????????????????????????????????????????????????????????????????????????????????????????????????????????????????????????????????????????????????????????????????????????????????????????????????????????????????????????????????????????????????????????????????????????????????????????????????????????????????????????????????????????????????????????????????????????????????????????????????????????????????????????????????????????????????????????????????????????????????????????????????????????????????????????

			>Orthophytum_falconii
			?????????????????????????????????????????????????????????????????????????????????????????????????????????????????????????????????????????????????????????????????????????????????????????????????????????????????????????????????????????????????????????????????????????????????????????????????????????????????????????????????????????????????????????????????????????????????????????????????????????????????????????????????????????????????????????????????????????????????????????????????????????????????????????????????????????????????????????????????????????????????????????????????????

			>Orthophytum_foliosum
			?????????????????????????????????????????????????????????????????????????????????????????????????????????????????????????????????????????????????????????????????????????????????????????????????????????????????????????????????????????????????????????????????????????????????????????????????????????????????????????????????????????????????????????????????????????????????????????????????????????????????????????????????????????????????????????????????????????????????????????????????????????????????????????????????????????????????????????????????????????????????????????????????????

			>Orthophytum_fosterianum
			?????????????????????????????????????????????????????????????????????????????????????????????????????????????????????????????????????????????????????????????????????????????????????????????????????????????????????????????????????????????????????????????????????????????????????????????????????????????????????????????????????????????????????????????????????????????????????????????????????????????????????????????????????????????????????????????????????????????????????????????????????????????????????????????????????????????????????????????????????????????????????????????????????

			>Orthophytum_glabrum
			?????????????????????????????????????????????????????????????????????????????????????????????????????????????????????????????????????????????????????????????????????????????????????????????????????????????????????????????????????????????????????????????????????????????????????????????????????????????????????????????????????????????????????????????????????????????????????????????????????????????????????????????????????????????????????????????????????????????????????????????????????????????????????????????????????????????????????????????????????????????????????????????????????

			>Orthophytum_graomogolense
			?????????????????????????????????????????????????????????????????????????????????????????????????????????????????????????????????????????????????????????????????????????????????????????????????????????????????????????????????????????????????????????????????????????????????????????????????????????????????????????????????????????????????????????????????????????????????????????????????????????????????????????????????????????????????????????????????????????????????????????????????????????????????????????????????????????????????????????????????????????????????????????????????????

			>Orthophytum_grossiorum
			?????????????????????????????????????????????????????????????????????????????????????????????????????????????????????????????????????????????????????????????????????????????????????????????????????????????????????????????????????????????????????????????????????????????????????????????????????????????????????????????????????????????????????????????????????????????????????????????????????????????????????????????????????????????????????????????????????????????????????????????????????????????????????????????????????????????????????????????????????????????????????????????????????

			>Orthophytum_gurkenii
			?????????????????????????????????????????????????????????????????????????????????????????????????????????????????????????????????????????????????????????????????????????????????????????????????????????????????????????????????????????????????????????????????????????????????????????????????????????????????????????????????????????????????????????????????????????????????????????????????????????????????????????????????????????????????????????????????????????????????????????????????????????????????????????????????????????????????????????????????????????????????????????????????????

			>Orthophytum_harleyi
			?????????????????????????????????????????????????????????????????????????????????????????????????????????????????????????????????????????????????????????????????????????????????????????????????????????????????????????????????????????????????????????????????????????????????????????????????????????????????????????????????????????????????????????????????????????????????????????????????????????????????????????????????????????????????????????????????????????????????????????????????????????????????????????????????????????????????????????????????????????????????????????????????????

			>Orthophytum_horridum
			?????????????????????????????????????????????????????????????????????????????????????????????????????????????????????????????????????????????????????????????????????????????????????????????????????????????????????????????????????????????????????????????????????????????????????????????????????????????????????????????????????????????????????????????????????????????????????????????????????????????????????????????????????????????????????????????????????????????????????????????????????????????????????????????????????????????????????????????????????????????????????????????????????

			>Orthophytum_lanuginosum
			?????????????????????????????????????????????????????????????????????????????????????????????????????????????????????????????????????????????????????????????????????????????????????????????????????????????????????????????????????????????????????????????????????????????????????????????????????????????????????????????????????????????????????????????????????????????????????????????????????????????????????????????????????????????????????????????????????????????????????????????????????????????????????????????????????????????????????????????????????????????????????????????????????

			>Orthophytum_lemei
			?????????????????????????????????????????????????????????????????????????????????????????????????????????????????????????????????????????????????????????????????????????????????????????????????????????????????????????????????????????????????????????????????????????????????????????????????????????????????????????????????????????????????????????????????????????????????????????????????????????????????????????????????????????????????????????????????????????????????????????????????????????????????????????????????????????????????????????????????????????????????????????????????????

			>Orthophytum_leprosum
			?????????????????????????????????????????????????????????????????????????????????????????????????????????????????????????????????????????????????????????????????????????????????????????????????????????????????????????????????????????????????????????????????????????????????????????????????????????????????????????????????????????????????????????????????????????????????????????????????????????????????????????????????????????????????????????????????????????????????????????????????????????????????????????????????????????????????????????????????????????????????????????????????????

			>Orthophytum_lucidum
			?????????????????????????????????????????????????????????????????????????????????????????????????????????????????????????????????????????????????????????????????????????????????????????????????????????????????????????????????????????????????????????????????????????????????????????????????????????????????????????????????????????????????????????????????????????????????????????????????????????????????????????????????????????????????????????????????????????????????????????????????????????????????????????????????????????????????????????????????????????????????????????????????????

			>Orthophytum_macroflorum
			?????????????????????????????????????????????????????????????????????????????????????????????????????????????????????????????????????????????????????????????????????????????????????????????????????????????????????????????????????????????????????????????????????????????????????????????????????????????????????????????????????????????????????????????????????????????????????????????????????????????????????????????????????????????????????????????????????????????????????????????????????????????????????????????????????????????????????????????????????????????????????????????????????

			>Orthophytum_magalhaesii
			?????????????????????????????????????????????????????????????????????????????????????????????????????????????????????????????????????????????????????????????????????????????????????????????????????????????????????????????????????????????????????????????????????????????????????????????????????????????????????????????????????????????????????????????????????????????????????????????????????????????????????????????????????????????????????????????????????????????????????????????????????????????????????????????????????????????????????????????????????????????????????????????????????

			>Orthophytum_maracasense
			?????????????????????????????????????????????????????????????????????????????????????????????????????????????????????????????????????????????????????????????????????????????????????????????????????????????????????????????????????????????????????????????????????????????????????????????????????????????????????????????????????????????????????????????????????????????????????????????????????????????????????????????????????????????????????????????????????????????????????????????????????????????????????????????????????????????????????????????????????????????????????????????????????

			>Orthophytum_mello-barretoi
			?????????????????????????????????????????????????????????????????????????????????????????????????????????????????????????????????????????????????????????????????????????????????????????????????????????????????????????????????????????????????????????????????????????????????????????????????????????????????????????????????????????????????????????????????????????????????????????????????????????????????????????????????????????????????????????????????????????????????????????????????????????????????????????????????????????????????????????????????????????????????????????????????????

			>Orthophytum_pseudostoloniferum
			?????????????????????????????????????????????????????????????????????????????????????????????????????????????????????????????????????????????????????????????????????????????????????????????????????????????????????????????????????????????????????????????????????????????????????????????????????????????????????????????????????????????????????????????????????????????????????????????????????????????????????????????????????????????????????????????????????????????????????????????????????????????????????????????????????????????????????????????????????????????????????????????????????

			>Orthophytum_pseudovagans
			?????????????????????????????????????????????????????????????????????????????????????????????????????????????????????????????????????????????????????????????????????????????????????????????????????????????????????????????????????????????????????????????????????????????????????????????????????????????????????????????????????????????????????????????????????????????????????????????????????????????????????????????????????????????????????????????????????????????????????????????????????????????????????????????????????????????????????????????????????????????????????????????????????

			>Orthophytum_riocontense
			?????????????????????????????????????????????????????????????????????????????????????????????????????????????????????????????????????????????????????????????????????????????????????????????????????????????????????????????????????????????????????????????????????????????????????????????????????????????????????????????????????????????????????????????????????????????????????????????????????????????????????????????????????????????????????????????????????????????????????????????????????????????????????????????????????????????????????????????????????????????????????????????????????

			>Orthophytum_roseolilacinum
			?????????????????????????????????????????????????????????????????????????????????????????????????????????????????????????????????????????????????????????????????????????????????????????????????????????????????????????????????????????????????????????????????????????????????????????????????????????????????????????????????????????????????????????????????????????????????????????????????????????????????????????????????????????????????????????????????????????????????????????????????????????????????????????????????????????????????????????????????????????????????????????????????????

			>Orthophytum_rubiginosum
			?????????????????????????????????????????????????????????????????????????????????????????????????????????????????????????????????????????????????????????????????????????????????????????????????????????????????????????????????????????????????????????????????????????????????????????????????????????????????????????????????????????????????????????????????????????????????????????????????????????????????????????????????????????????????????????????????????????????????????????????????????????????????????????????????????????????????????????????????????????????????????????????????????

			>Orthophytum_santaritense
			?????????????????????????????????????????????????????????????????????????????????????????????????????????????????????????????????????????????????????????????????????????????????????????????????????????????????????????????????????????????????????????????????????????????????????????????????????????????????????????????????????????????????????????????????????????????????????????????????????????????????????????????????????????????????????????????????????????????????????????????????????????????????????????????????????????????????????????????????????????????????????????????????????

			>Orthophytum_saxicola
			?????????????????????????????????????????????????????????????????????????????????????????????????????????????????????????????????????????????????????????????????????????????????????????????????????????????????????????????????????????????????????????????????????????????????????????????????????????????????????????????????????????????????????????????????????????????????????????????????????????????????????????????????????????????????????????????????????????????????????????????????????????????????????????????????????????????????????????????????????????????????????????????????????

			>Orthophytum_schulzianum
			?????????????????????????????????????????????????????????????????????????????????????????????????????????????????????????????????????????????????????????????????????????????????????????????????????????????????????????????????????????????????????????????????????????????????????????????????????????????????????????????????????????????????????????????????????????????????????????????????????????????????????????????????????????????????????????????????????????????????????????????????????????????????????????????????????????????????????????????????????????????????????????????????????

			>Orthophytum_sucrei
			?????????????????????????????????????????????????????????????????????????????????????????????????????????????????????????????????????????????????????????????????????????????????????????????????????????????????????????????????????????????????????????????????????????????????????????????????????????????????????????????????????????????????????????????????????????????????????????????????????????????????????????????????????????????????????????????????????????????????????????????????????????????????????????????????????????????????????????????????????????????????????????????????????

			>Orthophytum_toscanoi
			?????????????????????????????????????????????????????????????????????????????????????????????????????????????????????????????????????????????????????????????????????????????????????????????????????????????????????????????????????????????????????????????????????????????????????????????????????????????????????????????????????????????????????????????????????????????????????????????????????????????????????????????????????????????????????????????????????????????????????????????????????????????????????????????????????????????????????????????????????????????????????????????????????

			>Orthophytum_vagans
			?????????????????????????????????????????????????????????????????????????????????????????????????????????????????????????????????????????????????????????????????????????????????????????????????????????????????????????????????????????????????????????????????????????????????????????????????????????????????????????????????????????????????????????????????????????????????????????????????????????????????????????????????????????????????????????????????????????????????????????????????????????????????????????????????????????????????????????????????????????????????????????????????????

			>Orthophytum_vasconcelosianum
			?????????????????????????????????????????????????????????????????????????????????????????????????????????????????????????????????????????????????????????????????????????????????????????????????????????????????????????????????????????????????????????????????????????????????????????????????????????????????????????????????????????????????????????????????????????????????????????????????????????????????????????????????????????????????????????????????????????????????????????????????????????????????????????????????????????????????????????????????????????????????????????????????????

			>Orthophytum_zanonii
			?????????????????????????????????????????????????????????????????????????????????????????????????????????????????????????????????????????????????????????????????????????????????????????????????????????????????????????????????????????????????????????????????????????????????????????????????????????????????????????????????????????????????????????????????????????????????????????????????????????????????????????????????????????????????????????????????????????????????????????????????????????????????????????????????????????????????????????????????????????????????????????????????????

			>Pitcairnia_abundans
			?????????????????????????????????????????????????????????????????????????????????????????????????????????????????????????????????????????????????????????????????????????????????????????????????????????????????????????????????????????????????????????????????????????????????????????????????????????????????????????????????????????????????????????????????????????????????????????????????????????????????????????????????????????????????????????????????????????????????????????????????????????????????????????????????????????????????????????????????????????????????????????????????????

			>Pitcairnia_albiflos
			?????????????????????????????????????????????????????????????????????????????????????????????????????????????????????????????????????????????????????????????????????????????????????????????????????????????????????????????????????????????????????????????????????????????????????????????????????????????????????????????????????????????????????????????????????????????????????????????????????????????????????????????????????????????????????????????????????????????????????????????????????????????????????????????????????????????????????????????????????????????????????????????????????

			>Pitcairnia_atrorubens
			?????????????????????????????????????????????????????????????????????????????????????????????????????????????????????????????????????????????????????????????????????????????????????????????????????????????????????????????????????????????????????????????????????????????????????????????????????????????????????????????????????????????????????????????????????????????????????????????????????????????????????????????????????????????????????????????????????????????????????????????????????????????????????????????????????????????????????????????????????????????????????????????????????

			>Pitcairnia_beachiae
			?????????????????????????????????????????????????????????????????????????????????????????????????????????????????????????????????????????????????????????????????????????????????????????????????????????????????????????????????????????????????????????????????????????????????????????????????????????????????????????????????????????????????????????????????????????????????????????????????????????????????????????????????????????????????????????????????????????????????????????????????????????????????????????????????????????????????????????????????????????????????????????????????????

			>Pitcairnia_breedlovei
			?????????????????????????????????????????????????????????????????????????????????????????????????????????????????????????????????????????????????????????????????????????????????????????????????????????????????????????????????????????????????????????????????????????????????????????????????????????????????????????????????????????????????????????????????????????????????????????????????????????????????????????????????????????????????????????????????????????????????????????????????????????????????????????????????????????????????????????????????????????????????????????????????????

			>Pitcairnia_burle-marxii
			?????????????????????????????????????????????????????????????????????????????????????????????????????????????????????????????????????????????????????????????????????????????????????????????????????????????????????????????????????????????????????????????????????????????????????????????????????????????????????????????????????????????????????????????????????????????????????????????????????????????????????????????????????????????????????????????????????????????????????????????????????????????????????????????????????????????????????????????????????????????????????????????????????

			>Pitcairnia_carinata
			?????????????????????????????????????????????????????????????????????????????????????????????????????????????????????????????????????????????????????????????????????????????????????????????????????????????????????????????????????????????????????????????????????????????????????????????????????????????????????????????????????????????????????????????????????????????????????????????????????????????????????????????????????????????????????????????????????????????????????????????????????????????????????????????????????????????????????????????????????????????????????????????????????

			>Pitcairnia_chiapensis
			?????????????????????????????????????????????????????????????????????????????????????????????????????????????????????????????????????????????????????????????????????????????????????????????????????????????????????????????????????????????????????????????????????????????????????????????????????????????????????????????????????????????????????????????????????????????????????????????????????????????????????????????????????????????????????????????????????????????????????????????????????????????????????????????????????????????????????????????????????????????????????????????????????

			>Pitcairnia_corallina
			?????????????????????????????????????????????????????????????????????????????????????????????????????????????????????????????????????????????????????????????????????????????????????????????????????????????????????????????????????????????????????????????????????????????????????????????????????????????????????????????????????????????????????????????????????????????????????????????????????????????????????????????????????????????????????????????????????????????????????????????????????????????????????????????????????????????????????????????????????????????????????????????????????

			>Pitcairnia_elvirae
			?????????????????????????????????????????????????????????????????????????????????????????????????????????????????????????????????????????????????????????????????????????????????????????????????????????????????????????????????????????????????????????????????????????????????????????????????????????????????????????????????????????????????????????????????????????????????????????????????????????????????????????????????????????????????????????????????????????????????????????????????????????????????????????????????????????????????????????????????????????????????????????????????????

			>Pitcairnia_feliciana
			?????????????????????????????????????????????????????????????????????????????????????????????????????????????????????????????????????????????????????????????????????????????????????????????????????????????????????????????????????????????????????????????????????????????????????????????????????????????????????????????????????????????????????????????????????????????????????????????????????????????????????????????????????????????????????????????????????????????????????????????????????????????????????????????????????????????????????????????????????????????????????????????????????

			>Pitcairnia_flammea
			?????????????????????????????????????????????????????????????????????????????????????????????????????????????????????????????????????????????????????????????????????????????????????????????????????????????????????????????????????????????????????????????????????????????????????????????????????????????????????????????????????????????????????????????????????????????????????????????????????????????????????????????????????????????????????????????????????????????????????????????????????????????????????????????????????????????????????????????????????????????????????????????????????

			>Pitcairnia_flammea_var._glabrior
			?????????????????????????????????????????????????????????????????????????????????????????????????????????????????????????????????????????????????????????????????????????????????????????????????????????????????????????????????????????????????????????????????????????????????????????????????????????????????????????????????????????????????????????????????????????????????????????????????????????????????????????????????????????????????????????????????????????????????????????????????????????????????????????????????????????????????????????????????????????????????????????????????????

			>Pitcairnia_fuertesii
			?????????????????????????????????????????????????????????????????????????????????????????????????????????????????????????????????????????????????????????????????????????????????????????????????????????????????????????????????????????????????????????????????????????????????????????????????????????????????????????????????????????????????????????????????????????????????????????????????????????????????????????????????????????????????????????????????????????????????????????????????????????????????????????????????????????????????????????????????????????????????????????????????????

			>Pitcairnia_heterophylla
			?????????????????????????????????????????????????????????????????????????????????????????????????????????????????????????????????????????????????????????????????????????????????????????????????????????????????????????????????????????????????????????????????????????????????????????????????????????????????????????????????????????????????????????????????????????????????????????????????????????????????????????????????????????????????????????????????????????????????????????????????????????????????????????????????????????????????????????????????????????????????????????????????????

			>Pitcairnia_hirtzii
			?????????????????????????????????????????????????????????????????????????????????????????????????????????????????????????????????????????????????????????????????????????????????????????????????????????????????????????????????????????????????????????????????????????????????????????????????????????????????????????????????????????????????????????????????????????????????????????????????????????????????????????????????????????????????????????????????????????????????????????????????????????????????????????????????????????????????????????????????????????????????????????????????????

			>Pitcairnia_integrifolia
			?????????????????????????????????????????????????????????????????????????????????????????????????????????????????????????????????????????????????????????????????????????????????????????????????????????????????????????????????????????????????????????????????????????????????????????????????????????????????????????????????????????????????????????????????????????????????????????????????????????????????????????????????????????????????????????????????????????????????????????????????????????????????????????????????????????????????????????????????????????????????????????????????????

			>Pitcairnia_jimenezii
			?????????????????????????????????????????????????????????????????????????????????????????????????????????????????????????????????????????????????????????????????????????????????????????????????????????????????????????????????????????????????????????????????????????????????????????????????????????????????????????????????????????????????????????????????????????????????????????????????????????????????????????????????????????????????????????????????????????????????????????????????????????????????????????????????????????????????????????????????????????????????????????????????????

			>Pitcairnia_lignosa
			?????????????????????????????????????????????????????????????????????????????????????????????????????????????????????????????????????????????????????????????????????????????????????????????????????????????????????????????????????????????????????????????????????????????????????????????????????????????????????????????????????????????????????????????????????????????????????????????????????????????????????????????????????????????????????????????????????????????????????????????????????????????????????????????????????????????????????????????????????????????????????????????????????

			>Pitcairnia_loki-schmidtiae
			?????????????????????????????????????????????????????????????????????????????????????????????????????????????????????????????????????????????????????????????????????????????????????????????????????????????????????????????????????????????????????????????????????????????????????????????????????????????????????????????????????????????????????????????????????????????????????????????????????????????????????????????????????????????????????????????????????????????????????????????????????????????????????????????????????????????????????????????????????????????????????????????????????

			>Pitcairnia_macrochlamys
			?????????????????????????????????????????????????????????????????????????????????????????????????????????????????????????????????????????????????????????????????????????????????????????????????????????????????????????????????????????????????????????????????????????????????????????????????????????????????????????????????????????????????????????????????????????????????????????????????????????????????????????????????????????????????????????????????????????????????????????????????????????????????????????????????????????????????????????????????????????????????????????????????????

			>Pitcairnia_maidifolia
			?????????????????????????????????????????????????????????????????????????????????????????????????????????????????????????????????????????????????????????????????????????????????????????????????????????????????????????????????????????????????????????????????????????????????????????????????????????????????????????????????????????????????????????????????????????????????????????????????????????????????????????????????????????????????????????????????????????????????????????????????????????????????????????????????????????????????????????????????????????????????????????????????????

			>Pitcairnia_nigra
			?????????????????????????????????????????????????????????????????????????????????????????????????????????????????????????????????????????????????????????????????????????????????????????????????????????????????????????????????????????????????????????????????????????????????????????????????????????????????????????????????????????????????????????????????????????????????????????????????????????????????????????????????????????????????????????????????????????????????????????????????????????????????????????????????????????????????????????????????????????????????????????????????????

			>Pitcairnia_orchidifolia
			?????????????????????????????????????????????????????????????????????????????????????????????????????????????????????????????????????????????????????????????????????????????????????????????????????????????????????????????????????????????????????????????????????????????????????????????????????????????????????????????????????????????????????????????????????????????????????????????????????????????????????????????????????????????????????????????????????????????????????????????????????????????????????????????????????????????????????????????????????????????????????????????????????

			>Pitcairnia_piepenbringii
			?????????????????????????????????????????????????????????????????????????????????????????????????????????????????????????????????????????????????????????????????????????????????????????????????????????????????????????????????????????????????????????????????????????????????????????????????????????????????????????????????????????????????????????????????????????????????????????????????????????????????????????????????????????????????????????????????????????????????????????????????????????????????????????????????????????????????????????????????????????????????????????????????????

			>Pitcairnia_poortmanii
			?????????????????????????????????????????????????????????????????????????????????????????????????????????????????????????????????????????????????????????????????????????????????????????????????????????????????????????????????????????????????????????????????????????????????????????????????????????????????????????????????????????????????????????????????????????????????????????????????????????????????????????????????????????????????????????????????????????????????????????????????????????????????????????????????????????????????????????????????????????????????????????????????????

			>Pitcairnia_prolifera
			?????????????????????????????????????????????????????????????????????????????????????????????????????????????????????????????????????????????????????????????????????????????????????????????????????????????????????????????????????????????????????????????????????????????????????????????????????????????????????????????????????????????????????????????????????????????????????????????????????????????????????????????????????????????????????????????????????????????????????????????????????????????????????????????????????????????????????????????????????????????????????????????????????

			>Pitcairnia_pseudoundulata
			?????????????????????????????????????????????????????????????????????????????????????????????????????????????????????????????????????????????????????????????????????????????????????????????????????????????????????????????????????????????????????????????????????????????????????????????????????????????????????????????????????????????????????????????????????????????????????????????????????????????????????????????????????????????????????????????????????????????????????????????????????????????????????????????????????????????????????????????????????????????????????????????????????

			>Pitcairnia_pungens
			?????????????????????????????????????????????????????????????????????????????????????????????????????????????????????????????????????????????????????????????????????????????????????????????????????????????????????????????????????????????????????????????????????????????????????????????????????????????????????????????????????????????????????????????????????????????????????????????????????????????????????????????????????????????????????????????????????????????????????????????????????????????????????????????????????????????????????????????????????????????????????????????????????

			>Pitcairnia_punicea
			?????????????????????????????????????????????????????????????????????????????????????????????????????????????????????????????????????????????????????????????????????????????????????????????????????????????????????????????????????????????????????????????????????????????????????????????????????????????????????????????????????????????????????????????????????????????????????????????????????????????????????????????????????????????????????????????????????????????????????????????????????????????????????????????????????????????????????????????????????????????????????????????????????

			>Pitcairnia_rectiflora
			?????????????????????????????????????????????????????????????????????????????????????????????????????????????????????????????????????????????????????????????????????????????????????????????????????????????????????????????????????????????????????????????????????????????????????????????????????????????????????????????????????????????????????????????????????????????????????????????????????????????????????????????????????????????????????????????????????????????????????????????????????????????????????????????????????????????????????????????????????????????????????????????????????

			>Pitcairnia_recurvata
			?????????????????????????????????????????????????????????????????????????????????????????????????????????????????????????????????????????????????????????????????????????????????????????????????????????????????????????????????????????????????????????????????????????????????????????????????????????????????????????????????????????????????????????????????????????????????????????????????????????????????????????????????????????????????????????????????????????????????????????????????????????????????????????????????????????????????????????????????????????????????????????????????????

			>Pitcairnia_reflexiflora
			?????????????????????????????????????????????????????????????????????????????????????????????????????????????????????????????????????????????????????????????????????????????????????????????????????????????????????????????????????????????????????????????????????????????????????????????????????????????????????????????????????????????????????????????????????????????????????????????????????????????????????????????????????????????????????????????????????????????????????????????????????????????????????????????????????????????????????????????????????????????????????????????????????

			>Pitcairnia_riparia
			?????????????????????????????????????????????????????????????????????????????????????????????????????????????????????????????????????????????????????????????????????????????????????????????????????????????????????????????????????????????????????????????????????????????????????????????????????????????????????????????????????????????????????????????????????????????????????????????????????????????????????????????????????????????????????????????????????????????????????????????????????????????????????????????????????????????????????????????????????????????????????????????????????

			>Pitcairnia_roseoalba
			?????????????????????????????????????????????????????????????????????????????????????????????????????????????????????????????????????????????????????????????????????????????????????????????????????????????????????????????????????????????????????????????????????????????????????????????????????????????????????????????????????????????????????????????????????????????????????????????????????????????????????????????????????????????????????????????????????????????????????????????????????????????????????????????????????????????????????????????????????????????????????????????????????

			>Pitcairnia_rubronigriflora
			?????????????????????????????????????????????????????????????????????????????????????????????????????????????????????????????????????????????????????????????????????????????????????????????????????????????????????????????????????????????????????????????????????????????????????????????????????????????????????????????????????????????????????????????????????????????????????????????????????????????????????????????????????????????????????????????????????????????????????????????????????????????????????????????????????????????????????????????????????????????????????????????????????

			>Pitcairnia_sanguinea
			?????????????????????????????????????????????????????????????????????????????????????????????????????????????????????????????????????????????????????????????????????????????????????????????????????????????????????????????????????????????????????????????????????????????????????????????????????????????????????????????????????????????????????????????????????????????????????????????????????????????????????????????????????????????????????????????????????????????????????????????????????????????????????????????????????????????????????????????????????????????????????????????????????

			>Pitcairnia_smithiorum
			?????????????????????????????????????????????????????????????????????????????????????????????????????????????????????????????????????????????????????????????????????????????????????????????????????????????????????????????????????????????????????????????????????????????????????????????????????????????????????????????????????????????????????????????????????????????????????????????????????????????????????????????????????????????????????????????????????????????????????????????????????????????????????????????????????????????????????????????????????????????????????????????????????

			>Pitcairnia_spicata
			?????????????????????????????????????????????????????????????????????????????????????????????????????????????????????????????????????????????????????????????????????????????????????????????????????????????????????????????????????????????????????????????????????????????????????????????????????????????????????????????????????????????????????????????????????????????????????????????????????????????????????????????????????????????????????????????????????????????????????????????????????????????????????????????????????????????????????????????????????????????????????????????????????

			>Pitcairnia_sprucei
			?????????????????????????????????????????????????????????????????????????????????????????????????????????????????????????????????????????????????????????????????????????????????????????????????????????????????????????????????????????????????????????????????????????????????????????????????????????????????????????????????????????????????????????????????????????????????????????????????????????????????????????????????????????????????????????????????????????????????????????????????????????????????????????????????????????????????????????????????????????????????????????????????????

			>Pitcairnia_squarrosa
			?????????????????????????????????????????????????????????????????????????????????????????????????????????????????????????????????????????????????????????????????????????????????????????????????????????????????????????????????????????????????????????????????????????????????????????????????????????????????????????????????????????????????????????????????????????????????????????????????????????????????????????????????????????????????????????????????????????????????????????????????????????????????????????????????????????????????????????????????????????????????????????????????????

			>Pitcairnia_staminea
			?????????????????????????????????????????????????????????????????????????????????????????????????????????????????????????????????????????????????????????????????????????????????????????????????????????????????????????????????????????????????????????????????????????????????????????????????????????????????????????????????????????????????????????????????????????????????????????????????????????????????????????????????????????????????????????????????????????????????????????????????????????????????????????????????????????????????????????????????????????????????????????????????????

			>Pitcairnia_sulphurea
			?????????????????????????????????????????????????????????????????????????????????????????????????????????????????????????????????????????????????????????????????????????????????????????????????????????????????????????????????????????????????????????????????????????????????????????????????????????????????????????????????????????????????????????????????????????????????????????????????????????????????????????????????????????????????????????????????????????????????????????????????????????????????????????????????????????????????????????????????????????????????????????????????????

			>Pitcairnia_utcubambensis
			?????????????????????????????????????????????????????????????????????????????????????????????????????????????????????????????????????????????????????????????????????????????????????????????????????????????????????????????????????????????????????????????????????????????????????????????????????????????????????????????????????????????????????????????????????????????????????????????????????????????????????????????????????????????????????????????????????????????????????????????????????????????????????????????????????????????????????????????????????????????????????????????????????

			>Pitcairnia_villetaensis
			?????????????????????????????????????????????????????????????????????????????????????????????????????????????????????????????????????????????????????????????????????????????????????????????????????????????????????????????????????????????????????????????????????????????????????????????????????????????????????????????????????????????????????????????????????????????????????????????????????????????????????????????????????????????????????????????????????????????????????????????????????????????????????????????????????????????????????????????????????????????????????????????????????

			>Pitcairnia_wendlandii
			?????????????????????????????????????????????????????????????????????????????????????????????????????????????????????????????????????????????????????????????????????????????????????????????????????????????????????????????????????????????????????????????????????????????????????????????????????????????????????????????????????????????????????????????????????????????????????????????????????????????????????????????????????????????????????????????????????????????????????????????????????????????????????????????????????????????????????????????????????????????????????????????????????

			>Pitcairnia_xanthocalyx
			?????????????????????????????????????????????????????????????????????????????????????????????????????????????????????????????????????????????????????????????????????????????????????????????????????????????????????????????????????????????????????????????????????????????????????????????????????????????????????????????????????????????????????????????????????????????????????????????????????????????????????????????????????????????????????????????????????????????????????????????????????????????????????????????????????????????????????????????????????????????????????????????????????

			>Pitcairnia_yaupibajaensis
			?????????????????????????????????????????????????????????????????????????????????????????????????????????????????????????????????????????????????????????????????????????????????????????????????????????????????????????????????????????????????????????????????????????????????????????????????????????????????????????????????????????????????????????????????????????????????????????????????????????????????????????????????????????????????????????????????????????????????????????????????????????????????????????????????????????????????????????????????????????????????????????????????????

			>Portea_alatisepala
			?????????????????????????????????????????????????????????????????????????????????????????????????????????????????????????????????????????????????????????????????????????????????????????????????????????????????????????????????????????????????????????????????????????????????????????????????????????????????????????????????????????????????????????????????????????????????????????????????????????????????????????????????????????????????????????????????????????????????????????????????????????????????????????????????????????????????????????????????????????????????????????????????????

			>Portea_filifera
			?????????????????????????????????????????????????????????????????????????????????????????????????????????????????????????????????????????????????????????????????????????????????????????????????????????????????????????????????????????????????????????????????????????????????????????????????????????????????????????????????????????????????????????????????????????????????????????????????????????????????????????????????????????????????????????????????????????????????????????????????????????????????????????????????????????????????????????????????????????????????????????????????????

			>Portea_grandiflora
			?????????????????????????????????????????????????????????????????????????????????????????????????????????????????????????????????????????????????????????????????????????????????????????????????????????????????????????????????????????????????????????????????????????????????????????????????????????????????????????????????????????????????????????????????????????????????????????????????????????????????????????????????????????????????????????????????????????????????????????????????????????????????????????????????????????????????????????????????????????????????????????????????????

			>Portea_kermesina
			?????????????????????????????????????????????????????????????????????????????????????????????????????????????????????????????????????????????????????????????????????????????????????????????????????????????????????????????????????????????????????????????????????????????????????????????????????????????????????????????????????????????????????????????????????????????????????????????????????????????????????????????????????????????????????????????????????????????????????????????????????????????????????????????????????????????????????????????????????????????????????????????????????

			>Portea_nana
			?????????????????????????????????????????????????????????????????????????????????????????????????????????????????????????????????????????????????????????????????????????????????????????????????????????????????????????????????????????????????????????????????????????????????????????????????????????????????????????????????????????????????????????????????????????????????????????????????????????????????????????????????????????????????????????????????????????????????????????????????????????????????????????????????????????????????????????????????????????????????????????????????????

			>Portea_petropolitana
			?????????????????????????????????????????????????????????????????????????????????????????????????????????????????????????????????????????????????????????????????????????????????????????????????????????????????????????????????????????????????????????????????????????????????????????????????????????????????????????????????????????????????????????????????????????????????????????????????????????????????????????????????????????????????????????????????????????????????????????????????????????????????????????????????????????????????????????????????????????????????????????????????????

			>Portea_petropolitana_var._extensa
			?????????????????????????????????????????????????????????????????????????????????????????????????????????????????????????????????????????????????????????????????????????????????????????????????????????????????????????????????????????????????????????????????????????????????????????????????????????????????????????????????????????????????????????????????????????????????????????????????????????????????????????????????????????????????????????????????????????????????????????????????????????????????????????????????????????????????????????????????????????????????????????????????????

			>Portea_petropolitana_var._noettigii
			?????????????????????????????????????????????????????????????????????????????????????????????????????????????????????????????????????????????????????????????????????????????????????????????????????????????????????????????????????????????????????????????????????????????????????????????????????????????????????????????????????????????????????????????????????????????????????????????????????????????????????????????????????????????????????????????????????????????????????????????????????????????????????????????????????????????????????????????????????????????????????????????????????

			>Pseudalcantarea_grandis
			?????????????????????????????????????????????????????????????????????????????????????????????????????????????????????????????????????????????????????????????????????????????????????????????????????????????????????????????????????????????????????????????????????????????????????????????????????????????????????????????????????????????????????????????????????????????????????????????????????????????????????????????????????????????????????????????????????????????????????????????????????????????????????????????????????????????????????????????????????????????????????????????????????

			>Pseudalcantarea_macropetala
			?????????????????????????????????????????????????????????????????????????????????????????????????????????????????????????????????????????????????????????????????????????????????????????????????????????????????????????????????????????????????????????????????????????????????????????????????????????????????????????????????????????????????????????????????????????????????????????????????????????????????????????????????????????????????????????????????????????????????????????????????????????????????????????????????????????????????????????????????????????????????????????????????????

			>Pseudalcantarea_viridiflora
			?????????????????????????????????????????????????????????????????????????????????????????????????????????????????????????????????????????????????????????????????????????????????????????????????????????????????????????????????????????????????????????????????????????????????????????????????????????????????????????????????????????????????????????????????????????????????????????????????????????????????????????????????????????????????????????????????????????????????????????????????????????????????????????????????????????????????????????????????????????????????????????????????????

			>Pseudaraeococcus_parviflorus
			?????????????????????????????????????????????????????????????????????????????????????????????????????????????????????????????????????????????????????????????????????????????????????????????????????????????????????????????????????????????????????????????????????????????????????????????????????????????????????????????????????????????????????????????????????????????????????????????????????????????????????????????????????????????????????????????????????????????????????????????????????????????????????????????????????????????????????????????????????????????????????????????????????

			>Puya_aequatorialis
			?????????????????????????????????????????????????????????????????????????????????????????????????????????????????????????????????????????????????????????????????????????????????????????????????????????????????????????????????????????????????????????????????????????????????????????????????????????????????????????????????????????????????????????????????????????????????????????????????????????????????????????????????????????????????????????????????????????????????????????????????????????????????????????????????????????????????????????????????????????????????????????????????????

			>Puya_alpestris
			?????????????????????????????????????????????????????????????????????????????????????????????????????????????????????????????????????????????????????????????????????????????????????????????????????????????????????????????????????????????????????????????????????????????????????????????????????????????????????????????????????????????????????????????????????????????????????????????????????????????????????????????????????????????????????????????????????????????????????????????????????????????????????????????????????????????????????????????????????????????????????????????????????

			>Puya_angusta
			?????????????????????????????????????????????????????????????????????????????????????????????????????????????????????????????????????????????????????????????????????????????????????????????????????????????????????????????????????????????????????????????????????????????????????????????????????????????????????????????????????????????????????????????????????????????????????????????????????????????????????????????????????????????????????????????????????????????????????????????????????????????????????????????????????????????????????????????????????????????????????????????????????

			>Puya_bicolor
			?????????????????????????????????????????????????????????????????????????????????????????????????????????????????????????????????????????????????????????????????????????????????????????????????????????????????????????????????????????????????????????????????????????????????????????????????????????????????????????????????????????????????????????????????????????????????????????????????????????????????????????????????????????????????????????????????????????????????????????????????????????????????????????????????????????????????????????????????????????????????????????????????????

			>Puya_boliviensis
			?????????????????????????????????????????????????????????????????????????????????????????????????????????????????????????????????????????????????????????????????????????????????????????????????????????????????????????????????????????????????????????????????????????????????????????????????????????????????????????????????????????????????????????????????????????????????????????????????????????????????????????????????????????????????????????????????????????????????????????????????????????????????????????????????????????????????????????????????????????????????????????????????????

			>Puya_castellanosii
			?????????????????????????????????????????????????????????????????????????????????????????????????????????????????????????????????????????????????????????????????????????????????????????????????????????????????????????????????????????????????????????????????????????????????????????????????????????????????????????????????????????????????????????????????????????????????????????????????????????????????????????????????????????????????????????????????????????????????????????????????????????????????????????????????????????????????????????????????????????????????????????????????????

			>Puya_chilensis
			?????????????????????????????????????????????????????????????????????????????????????????????????????????????????????????????????????????????????????????????????????????????????????????????????????????????????????????????????????????????????????????????????????????????????????????????????????????????????????????????????????????????????????????????????????????????????????????????????????????????????????????????????????????????????????????????????????????????????????????????????????????????????????????????????????????????????????????????????????????????????????????????????????

			>Puya_coerulea
			?????????????????????????????????????????????????????????????????????????????????????????????????????????????????????????????????????????????????????????????????????????????????????????????????????????????????????????????????????????????????????????????????????????????????????????????????????????????????????????????????????????????????????????????????????????????????????????????????????????????????????????????????????????????????????????????????????????????????????????????????????????????????????????????????????????????????????????????????????????????????????????????????????

			>Puya_coerulea_var._violacea
			?????????????????????????????????????????????????????????????????????????????????????????????????????????????????????????????????????????????????????????????????????????????????????????????????????????????????????????????????????????????????????????????????????????????????????????????????????????????????????????????????????????????????????????????????????????????????????????????????????????????????????????????????????????????????????????????????????????????????????????????????????????????????????????????????????????????????????????????????????????????????????????????????????

			>Puya_compacta
			?????????????????????????????????????????????????????????????????????????????????????????????????????????????????????????????????????????????????????????????????????????????????????????????????????????????????????????????????????????????????????????????????????????????????????????????????????????????????????????????????????????????????????????????????????????????????????????????????????????????????????????????????????????????????????????????????????????????????????????????????????????????????????????????????????????????????????????????????????????????????????????????????????

			>Puya_dasylirioides
			?????????????????????????????????????????????????????????????????????????????????????????????????????????????????????????????????????????????????????????????????????????????????????????????????????????????????????????????????????????????????????????????????????????????????????????????????????????????????????????????????????????????????????????????????????????????????????????????????????????????????????????????????????????????????????????????????????????????????????????????????????????????????????????????????????????????????????????????????????????????????????????????????????

			>Puya_densiflora
			?????????????????????????????????????????????????????????????????????????????????????????????????????????????????????????????????????????????????????????????????????????????????????????????????????????????????????????????????????????????????????????????????????????????????????????????????????????????????????????????????????????????????????????????????????????????????????????????????????????????????????????????????????????????????????????????????????????????????????????????????????????????????????????????????????????????????????????????????????????????????????????????????????

			>Puya_dyckioides
			?????????????????????????????????????????????????????????????????????????????????????????????????????????????????????????????????????????????????????????????????????????????????????????????????????????????????????????????????????????????????????????????????????????????????????????????????????????????????????????????????????????????????????????????????????????????????????????????????????????????????????????????????????????????????????????????????????????????????????????????????????????????????????????????????????????????????????????????????????????????????????????????????????

			>Puya_ferreyrae
			?????????????????????????????????????????????????????????????????????????????????????????????????????????????????????????????????????????????????????????????????????????????????????????????????????????????????????????????????????????????????????????????????????????????????????????????????????????????????????????????????????????????????????????????????????????????????????????????????????????????????????????????????????????????????????????????????????????????????????????????????????????????????????????????????????????????????????????????????????????????????????????????????????

			>Puya_ferruginea
			?????????????????????????????????????????????????????????????????????????????????????????????????????????????????????????????????????????????????????????????????????????????????????????????????????????????????????????????????????????????????????????????????????????????????????????????????????????????????????????????????????????????????????????????????????????????????????????????????????????????????????????????????????????????????????????????????????????????????????????????????????????????????????????????????????????????????????????????????????????????????????????????????????

			>Puya_floccosa
			?????????????????????????????????????????????????????????????????????????????????????????????????????????????????????????????????????????????????????????????????????????????????????????????????????????????????????????????????????????????????????????????????????????????????????????????????????????????????????????????????????????????????????????????????????????????????????????????????????????????????????????????????????????????????????????????????????????????????????????????????????????????????????????????????????????????????????????????????????????????????????????????????????

			>Puya_gilmartiniae
			?????????????????????????????????????????????????????????????????????????????????????????????????????????????????????????????????????????????????????????????????????????????????????????????????????????????????????????????????????????????????????????????????????????????????????????????????????????????????????????????????????????????????????????????????????????????????????????????????????????????????????????????????????????????????????????????????????????????????????????????????????????????????????????????????????????????????????????????????????????????????????????????????????

			>Puya_goudotiana
			?????????????????????????????????????????????????????????????????????????????????????????????????????????????????????????????????????????????????????????????????????????????????????????????????????????????????????????????????????????????????????????????????????????????????????????????????????????????????????????????????????????????????????????????????????????????????????????????????????????????????????????????????????????????????????????????????????????????????????????????????????????????????????????????????????????????????????????????????????????????????????????????????????

			>Puya_hamata
			?????????????????????????????????????????????????????????????????????????????????????????????????????????????????????????????????????????????????????????????????????????????????????????????????????????????????????????????????????????????????????????????????????????????????????????????????????????????????????????????????????????????????????????????????????????????????????????????????????????????????????????????????????????????????????????????????????????????????????????????????????????????????????????????????????????????????????????????????????????????????????????????????????

			>Puya_harmsii
			?????????????????????????????????????????????????????????????????????????????????????????????????????????????????????????????????????????????????????????????????????????????????????????????????????????????????????????????????????????????????????????????????????????????????????????????????????????????????????????????????????????????????????????????????????????????????????????????????????????????????????????????????????????????????????????????????????????????????????????????????????????????????????????????????????????????????????????????????????????????????????????????????????

			>Puya_herrerae
			?????????????????????????????????????????????????????????????????????????????????????????????????????????????????????????????????????????????????????????????????????????????????????????????????????????????????????????????????????????????????????????????????????????????????????????????????????????????????????????????????????????????????????????????????????????????????????????????????????????????????????????????????????????????????????????????????????????????????????????????????????????????????????????????????????????????????????????????????????????????????????????????????????

			>Puya_herzogii
			?????????????????????????????????????????????????????????????????????????????????????????????????????????????????????????????????????????????????????????????????????????????????????????????????????????????????????????????????????????????????????????????????????????????????????????????????????????????????????????????????????????????????????????????????????????????????????????????????????????????????????????????????????????????????????????????????????????????????????????????????????????????????????????????????????????????????????????????????????????????????????????????????????

			>Puya_humilis
			?????????????????????????????????????????????????????????????????????????????????????????????????????????????????????????????????????????????????????????????????????????????????????????????????????????????????????????????????????????????????????????????????????????????????????????????????????????????????????????????????????????????????????????????????????????????????????????????????????????????????????????????????????????????????????????????????????????????????????????????????????????????????????????????????????????????????????????????????????????????????????????????????????

			>Puya_lanata
			?????????????????????????????????????????????????????????????????????????????????????????????????????????????????????????????????????????????????????????????????????????????????????????????????????????????????????????????????????????????????????????????????????????????????????????????????????????????????????????????????????????????????????????????????????????????????????????????????????????????????????????????????????????????????????????????????????????????????????????????????????????????????????????????????????????????????????????????????????????????????????????????????????

			>Puya_laxa
			?????????????????????????????????????????????????????????????????????????????????????????????????????????????????????????????????????????????????????????????????????????????????????????????????????????????????????????????????????????????????????????????????????????????????????????????????????????????????????????????????????????????????????????????????????????????????????????????????????????????????????????????????????????????????????????????????????????????????????????????????????????????????????????????????????????????????????????????????????????????????????????????????????

			>Puya_lineata
			?????????????????????????????????????????????????????????????????????????????????????????????????????????????????????????????????????????????????????????????????????????????????????????????????????????????????????????????????????????????????????????????????????????????????????????????????????????????????????????????????????????????????????????????????????????????????????????????????????????????????????????????????????????????????????????????????????????????????????????????????????????????????????????????????????????????????????????????????????????????????????????????????????

			>Puya_macrura
			?????????????????????????????????????????????????????????????????????????????????????????????????????????????????????????????????????????????????????????????????????????????????????????????????????????????????????????????????????????????????????????????????????????????????????????????????????????????????????????????????????????????????????????????????????????????????????????????????????????????????????????????????????????????????????????????????????????????????????????????????????????????????????????????????????????????????????????????????????????????????????????????????????

			>Puya_mima
			?????????????????????????????????????????????????????????????????????????????????????????????????????????????????????????????????????????????????????????????????????????????????????????????????????????????????????????????????????????????????????????????????????????????????????????????????????????????????????????????????????????????????????????????????????????????????????????????????????????????????????????????????????????????????????????????????????????????????????????????????????????????????????????????????????????????????????????????????????????????????????????????????????

			>Puya_mirabilis
			?????????????????????????????????????????????????????????????????????????????????????????????????????????????????????????????????????????????????????????????????????????????????????????????????????????????????????????????????????????????????????????????????????????????????????????????????????????????????????????????????????????????????????????????????????????????????????????????????????????????????????????????????????????????????????????????????????????????????????????????????????????????????????????????????????????????????????????????????????????????????????????????????????

			>Puya_nana
			?????????????????????????????????????????????????????????????????????????????????????????????????????????????????????????????????????????????????????????????????????????????????????????????????????????????????????????????????????????????????????????????????????????????????????????????????????????????????????????????????????????????????????????????????????????????????????????????????????????????????????????????????????????????????????????????????????????????????????????????????????????????????????????????????????????????????????????????????????????????????????????????????????

			>Puya_nitida
			?????????????????????????????????????????????????????????????????????????????????????????????????????????????????????????????????????????????????????????????????????????????????????????????????????????????????????????????????????????????????????????????????????????????????????????????????????????????????????????????????????????????????????????????????????????????????????????????????????????????????????????????????????????????????????????????????????????????????????????????????????????????????????????????????????????????????????????????????????????????????????????????????????

			>Puya_nutans
			?????????????????????????????????????????????????????????????????????????????????????????????????????????????????????????????????????????????????????????????????????????????????????????????????????????????????????????????????????????????????????????????????????????????????????????????????????????????????????????????????????????????????????????????????????????????????????????????????????????????????????????????????????????????????????????????????????????????????????????????????????????????????????????????????????????????????????????????????????????????????????????????????????

			>Puya_obconica
			?????????????????????????????????????????????????????????????????????????????????????????????????????????????????????????????????????????????????????????????????????????????????????????????????????????????????????????????????????????????????????????????????????????????????????????????????????????????????????????????????????????????????????????????????????????????????????????????????????????????????????????????????????????????????????????????????????????????????????????????????????????????????????????????????????????????????????????????????????????????????????????????????????

			>Puya_pygmaea
			?????????????????????????????????????????????????????????????????????????????????????????????????????????????????????????????????????????????????????????????????????????????????????????????????????????????????????????????????????????????????????????????????????????????????????????????????????????????????????????????????????????????????????????????????????????????????????????????????????????????????????????????????????????????????????????????????????????????????????????????????????????????????????????????????????????????????????????????????????????????????????????????????????

			>Puya_raimondii
			?????????????????????????????????????????????????????????????????????????????????????????????????????????????????????????????????????????????????????????????????????????????????????????????????????????????????????????????????????????????????????????????????????????????????????????????????????????????????????????????????????????????????????????????????????????????????????????????????????????????????????????????????????????????????????????????????????????????????????????????????????????????????????????????????????????????????????????????????????????????????????????????????????

			>Puya_santosii
			?????????????????????????????????????????????????????????????????????????????????????????????????????????????????????????????????????????????????????????????????????????????????????????????????????????????????????????????????????????????????????????????????????????????????????????????????????????????????????????????????????????????????????????????????????????????????????????????????????????????????????????????????????????????????????????????????????????????????????????????????????????????????????????????????????????????????????????????????????????????????????????????????????

			>Puya_spathacea
			?????????????????????????????????????????????????????????????????????????????????????????????????????????????????????????????????????????????????????????????????????????????????????????????????????????????????????????????????????????????????????????????????????????????????????????????????????????????????????????????????????????????????????????????????????????????????????????????????????????????????????????????????????????????????????????????????????????????????????????????????????????????????????????????????????????????????????????????????????????????????????????????????????

			>Puya_stenothyrsa
			?????????????????????????????????????????????????????????????????????????????????????????????????????????????????????????????????????????????????????????????????????????????????????????????????????????????????????????????????????????????????????????????????????????????????????????????????????????????????????????????????????????????????????????????????????????????????????????????????????????????????????????????????????????????????????????????????????????????????????????????????????????????????????????????????????????????????????????????????????????????????????????????????????

			>Puya_trianae
			?????????????????????????????????????????????????????????????????????????????????????????????????????????????????????????????????????????????????????????????????????????????????????????????????????????????????????????????????????????????????????????????????????????????????????????????????????????????????????????????????????????????????????????????????????????????????????????????????????????????????????????????????????????????????????????????????????????????????????????????????????????????????????????????????????????????????????????????????????????????????????????????????????

			>Puya_tuberosa
			?????????????????????????????????????????????????????????????????????????????????????????????????????????????????????????????????????????????????????????????????????????????????????????????????????????????????????????????????????????????????????????????????????????????????????????????????????????????????????????????????????????????????????????????????????????????????????????????????????????????????????????????????????????????????????????????????????????????????????????????????????????????????????????????????????????????????????????????????????????????????????????????????????

			>Puya_ultima
			?????????????????????????????????????????????????????????????????????????????????????????????????????????????????????????????????????????????????????????????????????????????????????????????????????????????????????????????????????????????????????????????????????????????????????????????????????????????????????????????????????????????????????????????????????????????????????????????????????????????????????????????????????????????????????????????????????????????????????????????????????????????????????????????????????????????????????????????????????????????????????????????????????

			>Puya_venusta
			?????????????????????????????????????????????????????????????????????????????????????????????????????????????????????????????????????????????????????????????????????????????????????????????????????????????????????????????????????????????????????????????????????????????????????????????????????????????????????????????????????????????????????????????????????????????????????????????????????????????????????????????????????????????????????????????????????????????????????????????????????????????????????????????????????????????????????????????????????????????????????????????????????

			>Puya_wrightii
			?????????????????????????????????????????????????????????????????????????????????????????????????????????????????????????????????????????????????????????????????????????????????????????????????????????????????????????????????????????????????????????????????????????????????????????????????????????????????????????????????????????????????????????????????????????????????????????????????????????????????????????????????????????????????????????????????????????????????????????????????????????????????????????????????????????????????????????????????????????????????????????????????????

			>Puya_yakespala
			?????????????????????????????????????????????????????????????????????????????????????????????????????????????????????????????????????????????????????????????????????????????????????????????????????????????????????????????????????????????????????????????????????????????????????????????????????????????????????????????????????????????????????????????????????????????????????????????????????????????????????????????????????????????????????????????????????????????????????????????????????????????????????????????????????????????????????????????????????????????????????????????????????

			>Quesnelia_arvensis
			?????????????????????????????????????????????????????????????????????????????????????????????????????????????????????????????????????????????????????????????????????????????????????????????????????????????????????????????????????????????????????????????????????????????????????????????????????????????????????????????????????????????????????????????????????????????????????????????????????????????????????????????????????????????????????????????????????????????????????????????????????????????????????????????????????????????????????????????????????????????????????????????????????

			>Quesnelia_conquistensis
			?????????????????????????????????????????????????????????????????????????????????????????????????????????????????????????????????????????????????????????????????????????????????????????????????????????????????????????????????????????????????????????????????????????????????????????????????????????????????????????????????????????????????????????????????????????????????????????????????????????????????????????????????????????????????????????????????????????????????????????????????????????????????????????????????????????????????????????????????????????????????????????????????????

			>Quesnelia_edmundoi
			?????????????????????????????????????????????????????????????????????????????????????????????????????????????????????????????????????????????????????????????????????????????????????????????????????????????????????????????????????????????????????????????????????????????????????????????????????????????????????????????????????????????????????????????????????????????????????????????????????????????????????????????????????????????????????????????????????????????????????????????????????????????????????????????????????????????????????????????????????????????????????????????????????

			>Quesnelia_edmundoi_var._rubrobracteata
			?????????????????????????????????????????????????????????????????????????????????????????????????????????????????????????????????????????????????????????????????????????????????????????????????????????????????????????????????????????????????????????????????????????????????????????????????????????????????????????????????????????????????????????????????????????????????????????????????????????????????????????????????????????????????????????????????????????????????????????????????????????????????????????????????????????????????????????????????????????????????????????????????????

			>Quesnelia_humilis
			?????????????????????????????????????????????????????????????????????????????????????????????????????????????????????????????????????????????????????????????????????????????????????????????????????????????????????????????????????????????????????????????????????????????????????????????????????????????????????????????????????????????????????????????????????????????????????????????????????????????????????????????????????????????????????????????????????????????????????????????????????????????????????????????????????????????????????????????????????????????????????????????????????

			>Quesnelia_kautskyi
			?????????????????????????????????????????????????????????????????????????????????????????????????????????????????????????????????????????????????????????????????????????????????????????????????????????????????????????????????????????????????????????????????????????????????????????????????????????????????????????????????????????????????????????????????????????????????????????????????????????????????????????????????????????????????????????????????????????????????????????????????????????????????????????????????????????????????????????????????????????????????????????????????????

			>Quesnelia_koltesii
			?????????????????????????????????????????????????????????????????????????????????????????????????????????????????????????????????????????????????????????????????????????????????????????????????????????????????????????????????????????????????????????????????????????????????????????????????????????????????????????????????????????????????????????????????????????????????????????????????????????????????????????????????????????????????????????????????????????????????????????????????????????????????????????????????????????????????????????????????????????????????????????????????????

			>Quesnelia_lateralis
			?????????????????????????????????????????????????????????????????????????????????????????????????????????????????????????????????????????????????????????????????????????????????????????????????????????????????????????????????????????????????????????????????????????????????????????????????????????????????????????????????????????????????????????????????????????????????????????????????????????????????????????????????????????????????????????????????????????????????????????????????????????????????????????????????????????????????????????????????????????????????????????????????????

			>Quesnelia_liboniana
			?????????????????????????????????????????????????????????????????????????????????????????????????????????????????????????????????????????????????????????????????????????????????????????????????????????????????????????????????????????????????????????????????????????????????????????????????????????????????????????????????????????????????????????????????????????????????????????????????????????????????????????????????????????????????????????????????????????????????????????????????????????????????????????????????????????????????????????????????????????????????????????????????????

			>Quesnelia_marmorata
			?????????????????????????????????????????????????????????????????????????????????????????????????????????????????????????????????????????????????????????????????????????????????????????????????????????????????????????????????????????????????????????????????????????????????????????????????????????????????????????????????????????????????????????????????????????????????????????????????????????????????????????????????????????????????????????????????????????????????????????????????????????????????????????????????????????????????????????????????????????????????????????????????????

			>Quesnelia_quesneliana
			?????????????????????????????????????????????????????????????????????????????????????????????????????????????????????????????????????????????????????????????????????????????????????????????????????????????????????????????????????????????????????????????????????????????????????????????????????????????????????????????????????????????????????????????????????????????????????????????????????????????????????????????????????????????????????????????????????????????????????????????????????????????????????????????????????????????????????????????????????????????????????????????????????

			>Quesnelia_strobilispica
			?????????????????????????????????????????????????????????????????????????????????????????????????????????????????????????????????????????????????????????????????????????????????????????????????????????????????????????????????????????????????????????????????????????????????????????????????????????????????????????????????????????????????????????????????????????????????????????????????????????????????????????????????????????????????????????????????????????????????????????????????????????????????????????????????????????????????????????????????????????????????????????????????????

			>Quesnelia_testudo
			?????????????????????????????????????????????????????????????????????????????????????????????????????????????????????????????????????????????????????????????????????????????????????????????????????????????????????????????????????????????????????????????????????????????????????????????????????????????????????????????????????????????????????????????????????????????????????????????????????????????????????????????????????????????????????????????????????????????????????????????????????????????????????????????????????????????????????????????????????????????????????????????????????

			>Quesnelia_tubifolia
			?????????????????????????????????????????????????????????????????????????????????????????????????????????????????????????????????????????????????????????????????????????????????????????????????????????????????????????????????????????????????????????????????????????????????????????????????????????????????????????????????????????????????????????????????????????????????????????????????????????????????????????????????????????????????????????????????????????????????????????????????????????????????????????????????????????????????????????????????????????????????????????????????????

			>Racinaea_cuspidata
			?????????????????????????????????????????????????????????????????????????????????????????????????????????????????????????????????????????????????????????????????????????????????????????????????????????????????????????????????????????????????????????????????????????????????????????????????????????????????????????????????????????????????????????????????????????????????????????????????????????????????????????????????????????????????????????????????????????????????????????????????????????????????????????????????????????????????????????????????????????????????????????????????????

			>Racinaea_dyeriana
			?????????????????????????????????????????????????????????????????????????????????????????????????????????????????????????????????????????????????????????????????????????????????????????????????????????????????????????????????????????????????????????????????????????????????????????????????????????????????????????????????????????????????????????????????????????????????????????????????????????????????????????????????????????????????????????????????????????????????????????????????????????????????????????????????????????????????????????????????????????????????????????????????????

			>Racinaea_elegans
			?????????????????????????????????????????????????????????????????????????????????????????????????????????????????????????????????????????????????????????????????????????????????????????????????????????????????????????????????????????????????????????????????????????????????????????????????????????????????????????????????????????????????????????????????????????????????????????????????????????????????????????????????????????????????????????????????????????????????????????????????????????????????????????????????????????????????????????????????????????????????????????????????????

			>Racinaea_fraseri
			?????????????????????????????????????????????????????????????????????????????????????????????????????????????????????????????????????????????????????????????????????????????????????????????????????????????????????????????????????????????????????????????????????????????????????????????????????????????????????????????????????????????????????????????????????????????????????????????????????????????????????????????????????????????????????????????????????????????????????????????????????????????????????????????????????????????????????????????????????????????????????????????????????

			>Racinaea_hamaleana
			?????????????????????????????????????????????????????????????????????????????????????????????????????????????????????????????????????????????????????????????????????????????????????????????????????????????????????????????????????????????????????????????????????????????????????????????????????????????????????????????????????????????????????????????????????????????????????????????????????????????????????????????????????????????????????????????????????????????????????????????????????????????????????????????????????????????????????????????????????????????????????????????????????

			>Racinaea_multiflora
			?????????????????????????????????????????????????????????????????????????????????????????????????????????????????????????????????????????????????????????????????????????????????????????????????????????????????????????????????????????????????????????????????????????????????????????????????????????????????????????????????????????????????????????????????????????????????????????????????????????????????????????????????????????????????????????????????????????????????????????????????????????????????????????????????????????????????????????????????????????????????????????????????????

			>Racinaea_parviflora
			?????????????????????????????????????????????????????????????????????????????????????????????????????????????????????????????????????????????????????????????????????????????????????????????????????????????????????????????????????????????????????????????????????????????????????????????????????????????????????????????????????????????????????????????????????????????????????????????????????????????????????????????????????????????????????????????????????????????????????????????????????????????????????????????????????????????????????????????????????????????????????????????????????

			>Racinaea_pugiformis
			?????????????????????????????????????????????????????????????????????????????????????????????????????????????????????????????????????????????????????????????????????????????????????????????????????????????????????????????????????????????????????????????????????????????????????????????????????????????????????????????????????????????????????????????????????????????????????????????????????????????????????????????????????????????????????????????????????????????????????????????????????????????????????????????????????????????????????????????????????????????????????????????????????

			>Racinaea_ropalocarpa
			?????????????????????????????????????????????????????????????????????????????????????????????????????????????????????????????????????????????????????????????????????????????????????????????????????????????????????????????????????????????????????????????????????????????????????????????????????????????????????????????????????????????????????????????????????????????????????????????????????????????????????????????????????????????????????????????????????????????????????????????????????????????????????????????????????????????????????????????????????????????????????????????????????

			>Racinaea_rothschuhiana
			?????????????????????????????????????????????????????????????????????????????????????????????????????????????????????????????????????????????????????????????????????????????????????????????????????????????????????????????????????????????????????????????????????????????????????????????????????????????????????????????????????????????????????????????????????????????????????????????????????????????????????????????????????????????????????????????????????????????????????????????????????????????????????????????????????????????????????????????????????????????????????????????????????

			>Racinaea_schumanniana
			?????????????????????????????????????????????????????????????????????????????????????????????????????????????????????????????????????????????????????????????????????????????????????????????????????????????????????????????????????????????????????????????????????????????????????????????????????????????????????????????????????????????????????????????????????????????????????????????????????????????????????????????????????????????????????????????????????????????????????????????????????????????????????????????????????????????????????????????????????????????????????????????????????

			>Racinaea_seemannii
			?????????????????????????????????????????????????????????????????????????????????????????????????????????????????????????????????????????????????????????????????????????????????????????????????????????????????????????????????????????????????????????????????????????????????????????????????????????????????????????????????????????????????????????????????????????????????????????????????????????????????????????????????????????????????????????????????????????????????????????????????????????????????????????????????????????????????????????????????????????????????????????????????????

			>Racinaea_sinuosa
			?????????????????????????????????????????????????????????????????????????????????????????????????????????????????????????????????????????????????????????????????????????????????????????????????????????????????????????????????????????????????????????????????????????????????????????????????????????????????????????????????????????????????????????????????????????????????????????????????????????????????????????????????????????????????????????????????????????????????????????????????????????????????????????????????????????????????????????????????????????????????????????????????????

			>Racinaea_spiculosa
			?????????????????????????????????????????????????????????????????????????????????????????????????????????????????????????????????????????????????????????????????????????????????????????????????????????????????????????????????????????????????????????????????????????????????????????????????????????????????????????????????????????????????????????????????????????????????????????????????????????????????????????????????????????????????????????????????????????????????????????????????????????????????????????????????????????????????????????????????????????????????????????????????????

			>Racinaea_spiculosa_var._micrantha
			?????????????????????????????????????????????????????????????????????????????????????????????????????????????????????????????????????????????????????????????????????????????????????????????????????????????????????????????????????????????????????????????????????????????????????????????????????????????????????????????????????????????????????????????????????????????????????????????????????????????????????????????????????????????????????????????????????????????????????????????????????????????????????????????????????????????????????????????????????????????????????????????????????

			>Racinaea_venusta
			?????????????????????????????????????????????????????????????????????????????????????????????????????????????????????????????????????????????????????????????????????????????????????????????????????????????????????????????????????????????????????????????????????????????????????????????????????????????????????????????????????????????????????????????????????????????????????????????????????????????????????????????????????????????????????????????????????????????????????????????????????????????????????????????????????????????????????????????????????????????????????????????????????

			>Rapatea_paludosa
			?????????????????????????????????????????????????????????????????????????????????????????????????????????????????????????????????????????????????????????????????????????????????????????????????????????????????????????????????????????????????????????????????????????????????????????????????????????????????????????????????????????????????????????????????????????????????????????????????????????????????????????????????????????????????????????????????????????????????????????????????????????????????????????????????????????????????????????????????????????????????????????????????????

			>Rokautskyia_exaltata
			?????????????????????????????????????????????????????????????????????????????????????????????????????????????????????????????????????????????????????????????????????????????????????????????????????????????????????????????????????????????????????????????????????????????????????????????????????????????????????????????????????????????????????????????????????????????????????????????????????????????????????????????????????????????????????????????????????????????????????????????????????????????????????????????????????????????????????????????????????????????????????????????????????

			>Rokautskyia_fernseeoides
			?????????????????????????????????????????????????????????????????????????????????????????????????????????????????????????????????????????????????????????????????????????????????????????????????????????????????????????????????????????????????????????????????????????????????????????????????????????????????????????????????????????????????????????????????????????????????????????????????????????????????????????????????????????????????????????????????????????????????????????????????????????????????????????????????????????????????????????????????????????????????????????????????????

			>Rokautskyia_latifolia
			?????????????????????????????????????????????????????????????????????????????????????????????????????????????????????????????????????????????????????????????????????????????????????????????????????????????????????????????????????????????????????????????????????????????????????????????????????????????????????????????????????????????????????????????????????????????????????????????????????????????????????????????????????????????????????????????????????????????????????????????????????????????????????????????????????????????????????????????????????????????????????????????????????

			>Rokautskyia_microglazioui
			?????????????????????????????????????????????????????????????????????????????????????????????????????????????????????????????????????????????????????????????????????????????????????????????????????????????????????????????????????????????????????????????????????????????????????????????????????????????????????????????????????????????????????????????????????????????????????????????????????????????????????????????????????????????????????????????????????????????????????????????????????????????????????????????????????????????????????????????????????????????????????????????????????

			>Rokautskyia_odoratissima
			?????????????????????????????????????????????????????????????????????????????????????????????????????????????????????????????????????????????????????????????????????????????????????????????????????????????????????????????????????????????????????????????????????????????????????????????????????????????????????????????????????????????????????????????????????????????????????????????????????????????????????????????????????????????????????????????????????????????????????????????????????????????????????????????????????????????????????????????????????????????????????????????????????

			>Rokautskyia_roberto-kautskyi
			?????????????????????????????????????????????????????????????????????????????????????????????????????????????????????????????????????????????????????????????????????????????????????????????????????????????????????????????????????????????????????????????????????????????????????????????????????????????????????????????????????????????????????????????????????????????????????????????????????????????????????????????????????????????????????????????????????????????????????????????????????????????????????????????????????????????????????????????????????????????????????????????????????

			>Rokautskyia_sanctaluciae
			?????????????????????????????????????????????????????????????????????????????????????????????????????????????????????????????????????????????????????????????????????????????????????????????????????????????????????????????????????????????????????????????????????????????????????????????????????????????????????????????????????????????????????????????????????????????????????????????????????????????????????????????????????????????????????????????????????????????????????????????????????????????????????????????????????????????????????????????????????????????????????????????????????

			>Rokautskyia_scaposa
			?????????????????????????????????????????????????????????????????????????????????????????????????????????????????????????????????????????????????????????????????????????????????????????????????????????????????????????????????????????????????????????????????????????????????????????????????????????????????????????????????????????????????????????????????????????????????????????????????????????????????????????????????????????????????????????????????????????????????????????????????????????????????????????????????????????????????????????????????????????????????????????????????????

			>Ronnbergia_aciculosa
			?????????????????????????????????????????????????????????????????????????????????????????????????????????????????????????????????????????????????????????????????????????????????????????????????????????????????????????????????????????????????????????????????????????????????????????????????????????????????????????????????????????????????????????????????????????????????????????????????????????????????????????????????????????????????????????????????????????????????????????????????????????????????????????????????????????????????????????????????????????????????????????????????????

			>Ronnbergia_allenii
			?????????????????????????????????????????????????????????????????????????????????????????????????????????????????????????????????????????????????????????????????????????????????????????????????????????????????????????????????????????????????????????????????????????????????????????????????????????????????????????????????????????????????????????????????????????????????????????????????????????????????????????????????????????????????????????????????????????????????????????????????????????????????????????????????????????????????????????????????????????????????????????????????????

			>Ronnbergia_columbiana
			?????????????????????????????????????????????????????????????????????????????????????????????????????????????????????????????????????????????????????????????????????????????????????????????????????????????????????????????????????????????????????????????????????????????????????????????????????????????????????????????????????????????????????????????????????????????????????????????????????????????????????????????????????????????????????????????????????????????????????????????????????????????????????????????????????????????????????????????????????????????????????????????????????

			>Ronnbergia_deleonii
			?????????????????????????????????????????????????????????????????????????????????????????????????????????????????????????????????????????????????????????????????????????????????????????????????????????????????????????????????????????????????????????????????????????????????????????????????????????????????????????????????????????????????????????????????????????????????????????????????????????????????????????????????????????????????????????????????????????????????????????????????????????????????????????????????????????????????????????????????????????????????????????????????????

			>Ronnbergia_drakeana
			?????????????????????????????????????????????????????????????????????????????????????????????????????????????????????????????????????????????????????????????????????????????????????????????????????????????????????????????????????????????????????????????????????????????????????????????????????????????????????????????????????????????????????????????????????????????????????????????????????????????????????????????????????????????????????????????????????????????????????????????????????????????????????????????????????????????????????????????????????????????????????????????????????

			>Ronnbergia_explodens
			?????????????????????????????????????????????????????????????????????????????????????????????????????????????????????????????????????????????????????????????????????????????????????????????????????????????????????????????????????????????????????????????????????????????????????????????????????????????????????????????????????????????????????????????????????????????????????????????????????????????????????????????????????????????????????????????????????????????????????????????????????????????????????????????????????????????????????????????????????????????????????????????????????

			>Ronnbergia_fraseri
