## Supplemental Folder for "Drivers of dispersal and diversification in bromeliads": Tables S1, S2 and S3.docx

Table S1 Main clade support and timing of crown and stem ages.

| Clade | PP | Mean (Ma)  crown | 95% CI (Ma)  crown | Mean (Ma)  stem | 95% CI (Ma)  stem |
| --- | --- | --- | --- | --- | --- |
| Brocchinioideae | 0.99 | 10.43 | 8.33–11.60 | 22.77 | 20.77–23.09 |
| Bromelioideae (except *Bromelia*) | 0.86 | 9.03 | 6.01–10.96 | 9.84 | 6.70–11.86 |
| *Bromelia* | 0.99 | 5.06 | 3.5–6.06 | 11.17 | 7.81–13.26 |
| Hechtioideae | 1 | 11.95 | 9.14–14.59 | 15.19 | 12.20–17.60 |
| Lindmanioideae | 1 | 4.6 | 3.54–5.27 | 18.70 | 15.67–20.32 |
| Navioideae | 1 | 10.45 | 7.77–12.29 | 15.14 | 12.21–17.49 |
| Pitcairnioideae | 0.95 | 11.97 | 8.6–14.03 | 13.17 | 9.79–15.13 |
| Puyoideae | 0.91 | 7.13 | 4.72–8.78 | 9.84 | 6.70–11.86 |
| Tillandsioideae | 0.99 | 13.77 | 12.20–17.60 | 15.19 | 12.20–17.60 |
| Bromelioideae+Puyoideae | 0.95 | 11.17 | 7.81–13.26 | 11.17 | 7.81–13.26 |
| Bromeliaceae | 1 | 22.77 | 20.77–23.09 | 96.33 | 96–97.63 |

*Notes:* Node calibration based on Givnish et al. (2018); Mean and CI= 95% resulted from the treePL analysis. PP = Posterior probability resulted from the BEAST MCC tree; CI= Confidence interval; Ma= Million years ago.

Table S2 Comparison of trait-based best-fit models of dispersal represented by mean distance (MD).

| Formula: MD ~ | λ | df | AIC |
| --- | --- | --- | --- |
| All traits | 0.736  (0.620, 0.815) | 7 | 5941.572 |
| All numeric traits | 0.753  (0.652, 0.824) | 4 | 5939.643 |
| AT+CH | 0.748  (0.645, 0.821) | 2 | 5937.534 |

*Notes*: The model comprising mean annual temperature and canopy height was the best-fitted. λ= Pagel’s lambda; AT= mean annual temperature; CH= canopy height; MD= mean distance; df= degrees of freedom.

Table S3 ANOVA of the PGLS complete model. Estimated value of lambda of 0.736 (0.620, 0.815), kappa and delta fixed in 1, considering the influence of all variables under the mean distances. The levels of berry (FT) and winged (ST) were considered points of comparison, thus not estimated in the analysis. All variables were scale 0 modified.

| Trait | df | Sum Sq | Mean Sq | F value | p-value |
| --- | --- | --- | --- | --- | --- |
| *Capsule (FT)* | 1 | 56 | 55.93 | 1.4347 | 0.23113 |
| *Naked (ST)* | 1 | 12 | 12.24 | 0.3140 | 0.57531 |
| *Plumose (ST)* | 1 | 93 | 93.46 | 2.3977 | 0.12166 |
| ***AP*** | **1** | **535** | **535.27** | **13.7314** | **0.00021** |
| ***AT*** | **1** | **1479** | **1478.51** | **37.9286** | **8.712e-10** |
| *EL* | 1 | 66 | 65.99 | 1.6929 | 0.1933 |
| ***CH*** | **1** | **544** | **543.93** | **13.9535** | **0.00019** |
| *Residuals* | 2180 | 84979 | 38.98 | - | - |
