## Supplementary figures and images for "Drivers of dispersal and diversification in bromeliads"

### Fig S1

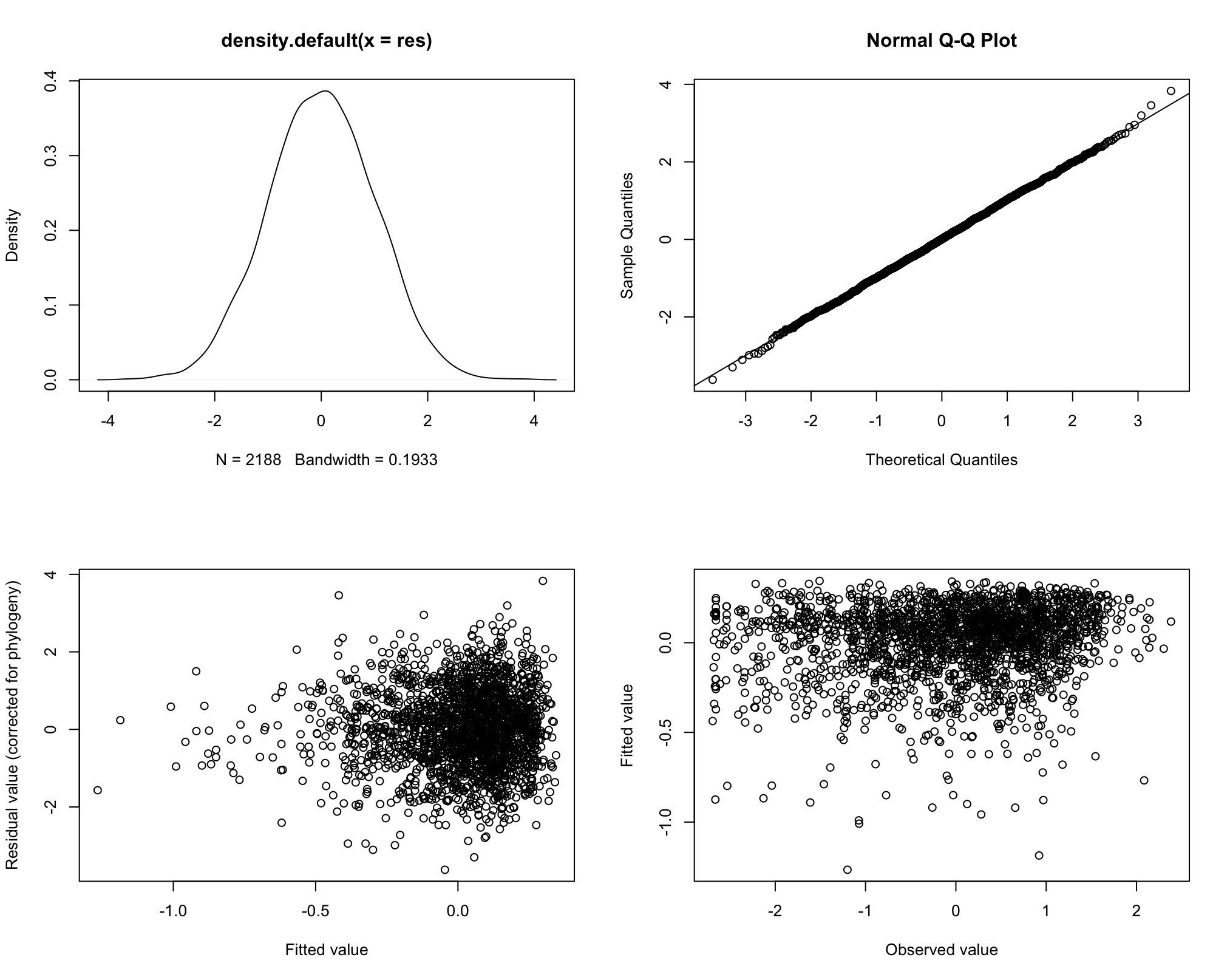

### Fig S2.pdf

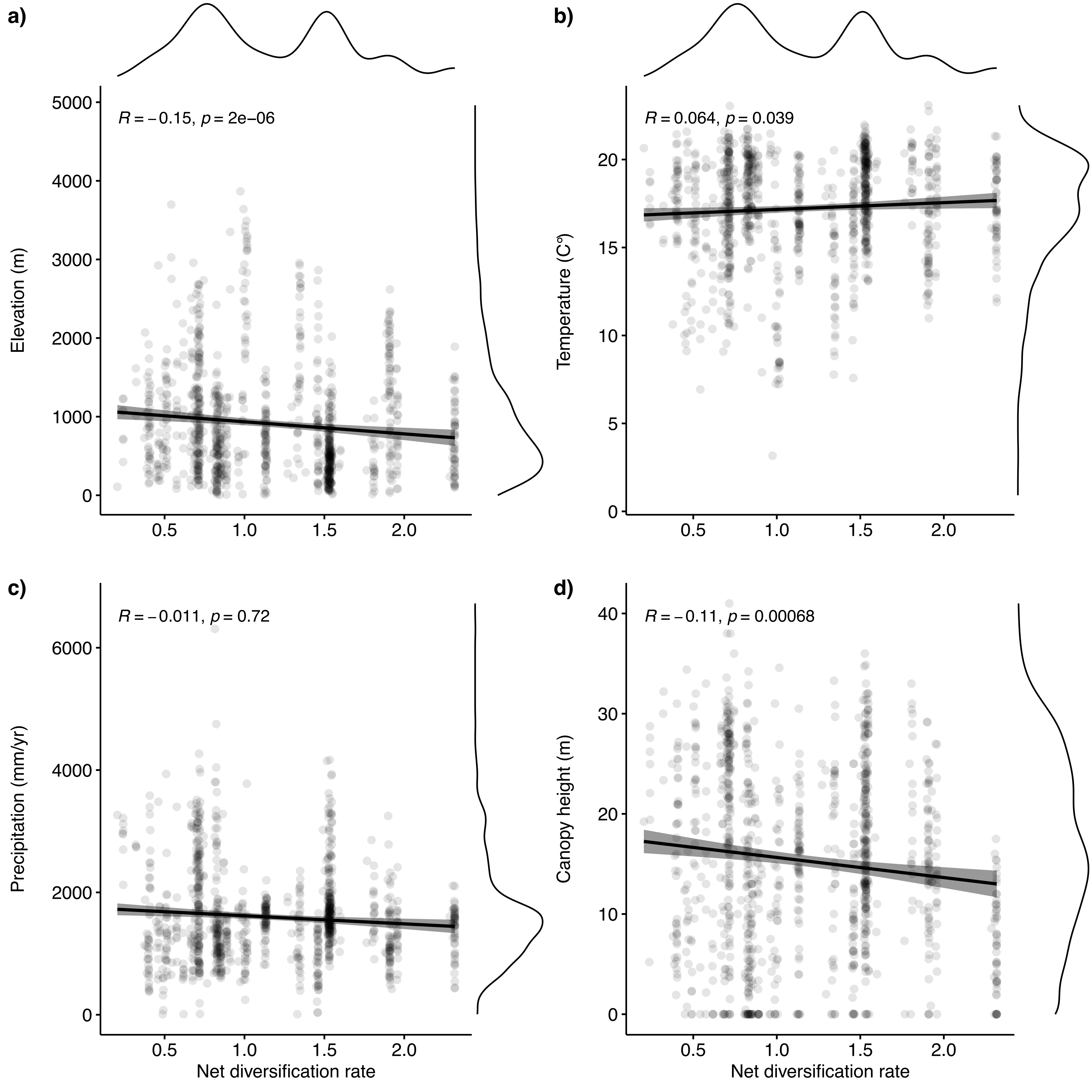

### Fig S3.pdf

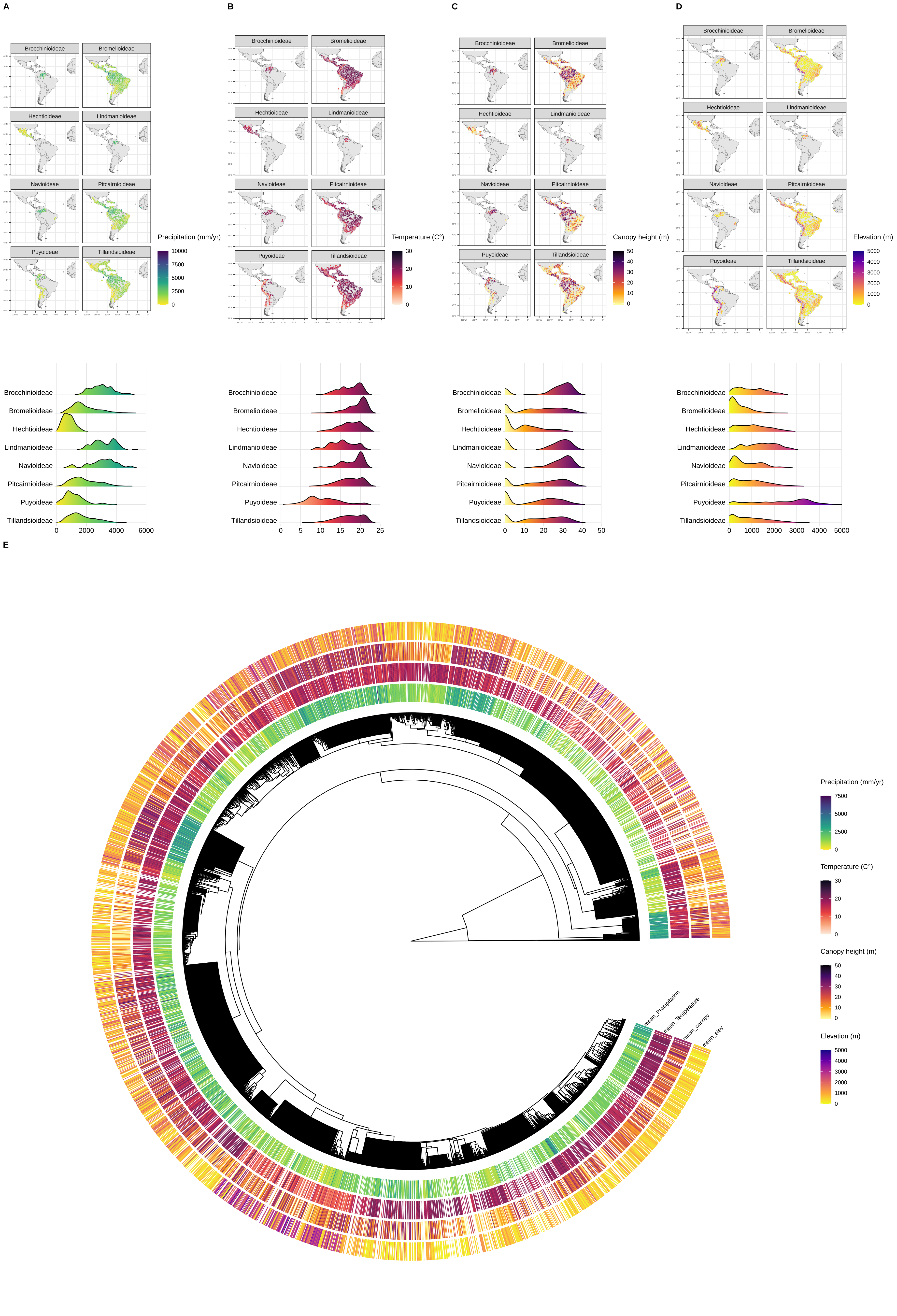

### Fig S4.pdf

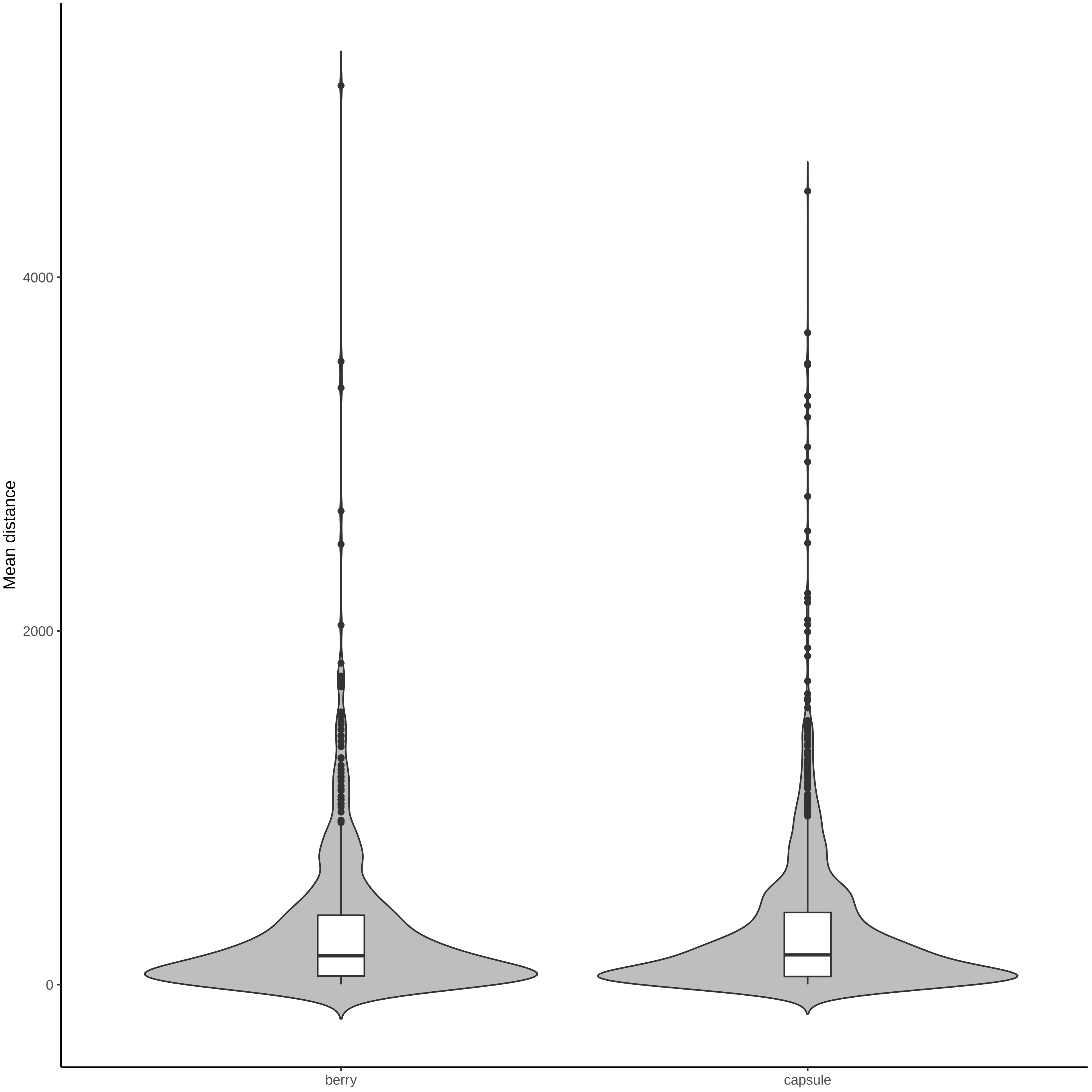
